# TALE-independent transcriptional activation of the rice executor gene *Xa23* is regulated via histone acetylation during zygote development

**DOI:** 10.64898/2026.09.02.748781

**Authors:** Kyrylo Schenstnyi, Kasidit Rattanawong, Aya Satoh, Annett Strauß, Natalie Faiß, Danalyn Rose Holmes, Laura Redzich, Erika Toda, Hanifah Aini, Atsuko Kinoshita, Thomas Lahaye, Takashi Okamoto

## Abstract

Transcription activator-like effectors (TALEs) from *Xanthomonas* activate transcription of “executor” (*E*) genes in host plants, leading to cell death and thereby restricting proliferation of biotrophic pathogens. Because *E* gene transcripts had only been detected upon activation by cognate *Xanthomonas* TALEs, *E* genes were thought to function exclusively in plant immunity. Here, we detect TALE-independent transcription of the rice *E* gene *Xa23* in zygotes 4-6 hours after gamete fusion. Histone deacetylase inhibition induces *Xa23* transcription in unfertilized egg cells, implicating histone acetylation in *Xa23* regulation. We identified potential *cis*-regulatory elements and transcription start sites associated with native *Xa23* transcription during zygote development. Together, our findings suggest that *Xa23* is a developmentally regulated gene with a native role during early zygote development. This supports a previously proposed model in which *E* genes have native functions in development, while fortuitous upstream polymorphisms can create TALE-binding sites that convert them into immune executors.

## Introduction

*Xanthomonas* bacteria inject transcription activator-like effectors (TALEs) into host cells via the type III secretion system (T3SS).^1^ Once in the nucleus, TALEs bind specific effector binding elements (*EBEs*) in host gene promoters and activate transcription.^2^ Unlike eukaryotic transcription factors (TFs), which recruit the RNA polymerase II pre-initiation complex (PIC) via the mediator complex, TALEs interact directly with the PIC through TFIIAγ, a general TF.^3,4^ This dual binding to the *EBE* and PIC enforces a fixed spacing (≈50 nucleotides; nt) between the *EBE* and the transcription start site (TSS).^5^

One of the best-studied examples of a TALE-*EBE* interaction involves AvrBs3, a TALE from pepper-pathogenic *Xanthomonas* strains, and the pepper gene *upa20*, which encodes a basic helix-loop-helix (bHLH) TF.^5^ Native and AvrBs3-induced *upa20* transcripts initiate at distinct TSSs, illustrating how TALE-dependent activation can generate transcripts that differ from their endogenously initiated counterparts.

TALEs activate ectopic expression of host genes, increasing host susceptibility to *Xanthomonas* strains. Such TALE targets are referred to as susceptibility (*S*) genes.^6–8^ However, some plant genotypes harbor a functionally and structurally unique class of resistance (*R*) genes that are preceded by *EBEs* that act as TALE traps. These *EBEs* might have evolved under pathogen selection to match specific TALEs.^9–15^ TALE-dependent transcriptional activation of these *R* genes triggers host cell death and inhibits *Xanthomonas* proliferation. Because the encoded proteins execute cell death rather than recognize TALEs, they are commonly referred to as “executor” (E) proteins.^16^

Two well-studied rice *E* genes, *Xa10* and *Xa23*, contain upstream *EBEs* that trap the *Xanthomonas oryzae* pv. *oryzae* (*Xoo*) TALE proteins AvrXa10 and AvrXa23, respectively. ^10,11,17,18^ Accordingly, *Xa10* and *Xa23* are activated by and confer resistance to *Xoo* strains containing AvrXa10 or AvrXa23. *Xa23* alleles lacking an AvrXa23-compatible *EBE* in their 5’ upstream region (*xa23*) do not confer resistance to AvrXa23-expressing *Xoo* strains due to the absence of TALE-mediated transcriptional activation. However, such *xa23* alleles can still trigger cell death and confer resistance when *Xanthomonas* strains express designer TALEs (dTALEs) tailored to user-defined upstream *xa23* sequences, demonstrating that *EBE*-lacking *xa23* alleles retain executor function.^19^ Similarly, the recently identified *Xa10*-like gene *xa10-Ni*, which lacks *EBEs* for known TALEs, can trigger cell death when *Xoo* strains deliver dTALEs that bind upstream of *xa10-Ni*.^19^ Together, these observations indicate that E proteins can retain cell death-inducing capacity despite their promoters lacking *EBEs* for naturally occurring TALEs. The widespread occurrence of functional *E* genes without recognizable *EBEs* argues against TALE recognition as the primary evolutionary function of this gene family.^20^ Instead, these findings suggest that TALE-mediated activation represents a secondary, opportunistic exploitation of a pre-existing executor function. However, the native functions of *E* genes remain unknown because TALE-independent transcriptional activation of *E* genes has not been reported.

We previously noted that tight transcriptional control of *E* genes mirrors the regulation of developmental programmed cell death (dPCD).^20^ We speculated that native *E*gene expression is confined to a few cells in specific organs at defined developmental stages, with E proteins functioning as transcriptionally regulated dPCD executors that confer *Xoo* resistance when ectopically activated by TALEs. This hypothesis challenges the conventional view of *E* genes as dedicated components of immunity.^19,21^

In this study, we investigated whether *E* genes might have developmental functions beyond immunity. We first assessed the abundance of *E* gene-like sequences across cultivated and wild rice and cutgrass species and tested whether newly discovered *E* gene homologs trigger cell death when overexpressed in *N. benthamiana*. *Xa23* homologs showed the highest sequence conservation, suggesting possible functions beyond immunity.

We next asked whether *Xa23* is transcribed outside the context of immunity. We searched public RNA-seq datasets for cell types, organs, developmental stages, or treatments in which non-TALE-induced *Xa23* transcripts are detectable. Notably, among the surveyed conditions, such transcripts were detected exclusively in rice zygotes produced via an *in vitro* fertilization (IVF) system, 4-6 hours after gamete fusion (after fusion, AF).^22,23^ Furthermore, we confirmed native, TALE-independent *Xa23* expression in IVF-produced zygotes at 6 hours AF.

Treatment of unfertilized egg cells, which do not express *Xa23*, with the histone deacetylase (HDAC) inhibitor (HDACi) trichostatin A (TSA) activated transcription of *Xa23*, suggesting that its regulation involves an epigenetic mechanism.

Additionally, we characterized TSA-induced *Xa23* transcripts via 5’ and 3’ rapid amplification of cDNA ends (RACE) and compared them to previously reported AvrXa23- and dTALE-induced *Xa23* transcripts.^11,19^ We discovered that TSA-induced *Xa23* transcripts are functionally equivalent to the cell death-inducing AvrXa23-induced transcripts. We also identified the transcription start sites (TSSs) of TSA-induced *Xa23* transcripts and potential *cis*-regulatory elements (CREs) in *Xa23* 5’ upstream genomic sequences that can be useful for further elucidation of pathways regulating *Xa23* transcription during zygote development.

Overall, our study established two experimental systems, IVF-produced zygotes (fertilization- dependent) and TSA-treated unfertilized egg cells (fertilization-independent), to investigate the putative role of *Xa23* beyond immunity and advance our understanding of the regulatory mechanisms governing cell death.

## Results

### Rice *E* genes display an evolutionary pattern atypical of canonical resistance genes

To gain insights into the native role of rice *E* genes, we asked whether their evolutionary conservation more closely resembles that of canonical race-specific immune receptor genes or genes with essential functions in plant development. Whereas many immune receptor genes display extensive presence/absence polymorphism and allelic diversity within a species,^24^ genes required for plant development are typically retained and highly conserved across accessions.^25^ Thus, gene conservation patterns within and across species can provide clues to their underlying biological functions.

We analyzed the conservation of *Xa10-Ni* and *Xa23* across 3,002 cultivated rice accessions. *Xa10* was excluded from the analysis because *Xa10* is not present in the rice reference genome (Nipponbare) to which the genomes of 3,002 cultivated accessions are aligned.^26,27^ *Xa10-Ni* was present as a full-length coding sequence (CDS) in 2,913 of the 3,002 cultivated rice accessions (97%), whereas only 89 accessions (3%) carried predicted premature stop codons (Data S1). XA10-NI proteins from 2,913 accessions shared 94-100% amino acid identity with XA10-Ni. *Xa23* was even more highly conserved: its full-length CDS was present in all 3,002 rice accessions (100%; Data S2). XA23 proteins from 3,002 accessions shared 96-100% amino acid identity with XA23. The near-universal retention and exceptionally high sequence conservation of *Xa10-Ni*, and especially *Xa23*, are therefore inconsistent with the evolutionary patterns typically observed for canonical race-specific resistance genes. Instead, these observations suggest that these genes have been subject to strong evolutionary constraint, consistent with an ancestral function in plant development.

### The *E* gene family has an ancient evolutionary origin

To determine whether *E* genes are restricted to cultivated rice or instead represent an evolutionarily ancient gene family, we examined their conservation across the tribe Oryzeae. If *E* genes predate the origin of cultivated rice, homologs would be expected to occur in wild *Oryza* species and, for a more ancient origin, in related genera that diverged before *Oryza* diversification. We therefore mined publicly available genomes of wild rice (*Oryza*) and cutgrass (*Leersia*) species, closely related genera within the tribe Oryzeae, for *E* gene-like sequences.^28,29^ Overall, we identified 53 *E* gene-like genomic sequences in 17 genomes from 13 wild rice species and one cutgrass species (Table S1).

Specifically, *Xa10*-like homologs were detected in seven of 13 wild rice species (54%) and in the single cutgrass species analyzed, while *Xa10-Ni*-like and *Xa23*-like homologs were present in eight (62%) and nine (69%) wild rice species, respectively. Together, these findings indicate that the *E* gene family members represent ancient evolutionary lineages maintained since before the divergence of the *Oryza* and *Leersia* lineages approximately 14 million years ago (MYA),^30^ consistent with an origin that predates their proposed recruitment into plant immunity.

### Sequence diversification of *E* gene family members

Although the *E* gene lineages are ancient, the number of genes within each lineage varied markedly among Oryzeae genomes. This is exemplified by two *O. longistaminata* accessions: whereas accession W11 contains only a single *Xa10*-like gene and one *Xa10-Ni*-like gene, accession RD23 contains three *Xa10*-like, one *Xa10-Ni*-like, and six *Xa23*-like genes (Figure 1; Table S1). Substantial copy-number variation was also evident across the broader set of Oryzeae genomes, ranging from single *E* gene-like loci in several genomes to pronounced expansions, including 12 loci in *O. officinalis* accession W0002 (Table S1). Moreover, these expansions involved different combinations of *Xa10*-, *Xa10-Ni*-, and *Xa23*-like sequences. Together, these observations indicate that recurrent, lineage-specific gene duplication and loss have dynamically shaped the *E* gene family during Oryzeae evolution.

**Figure 1.**
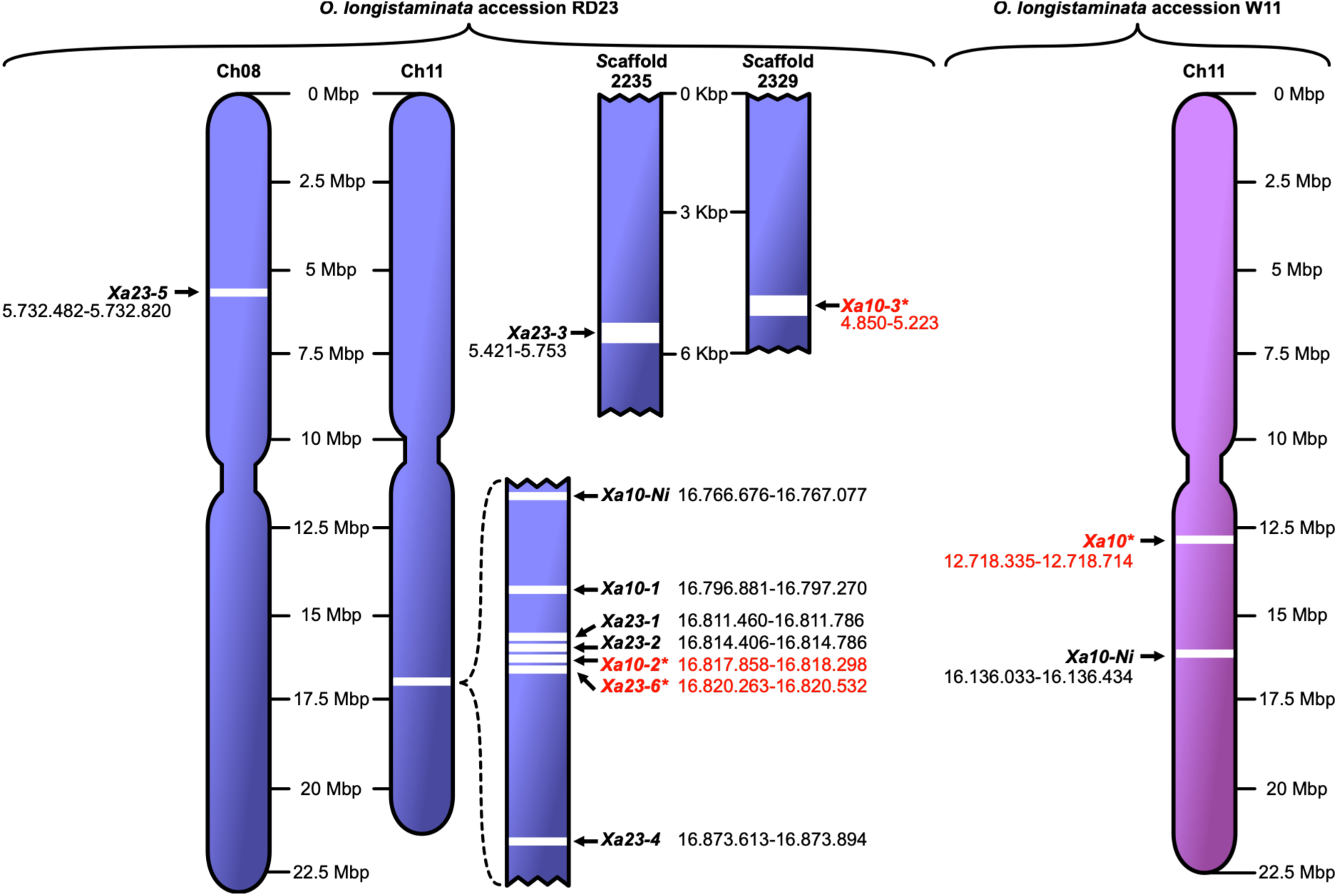
The rice *E* gene family has undergone duplication and diversification. The exact genomic positions of *E* gene-like loci in the genomes of *O. longistaminata* accessions RD23 and W11 are specified. Pseudogenes with predicted premature stop codons within their CDSs or encoding proteins that share <40% identity to *O. sativa* E proteins are marked with **red** asterisks (*).

Having established extensive copy-number variation across Oryzeae genomes, we next examined all *E* gene-like sequences to determine how the duplicated genes diversified. To do so, we aligned all newly identified *E* -like genomic sequences from wild rice and cutgrass to the corresponding *Xa10*, *Xa10-Ni*, and *Xa23* genomic sequences from cultivated rice, identified putative start and stop codons, and designated each protein according to the rice E protein with the highest sequence identity (Table S1 and Data S3-S5). For example, a predicted protein from cutgrass shared 45% identity with rice XA10, compared with 26% and 40% identity to rice XA10-Ni and XA23, respectively, and was therefore designated *L. perrieri* XA10-1. Pairwise sequence identity comparisons revealed amino acid identities of 45-99% for XA10-like proteins, 47-100% identity for XA10-Ni-like proteins and 48-100% for XA23-like proteins relative to their respective rice counterparts (Table S1). Thus, individual members within each of the three E protein lineages have accumulated substantial sequence variation following their diversification.

Consistent with this classification, phylogenetic analysis showed that E-like proteins form three paralogous clades derived from a shared ancestral gene, with XA10 and XA10-Ni representing more closely related sister lineages (Figure 2A). Together, these analyses indicate that the *E* gene family underwent recurrent lineage-specific expansion and substantial sequence diversification across Oryzeae while retaining three distinct paralogous lineages (Figures 1 and 2A; Table S1).

**Figure 2.**
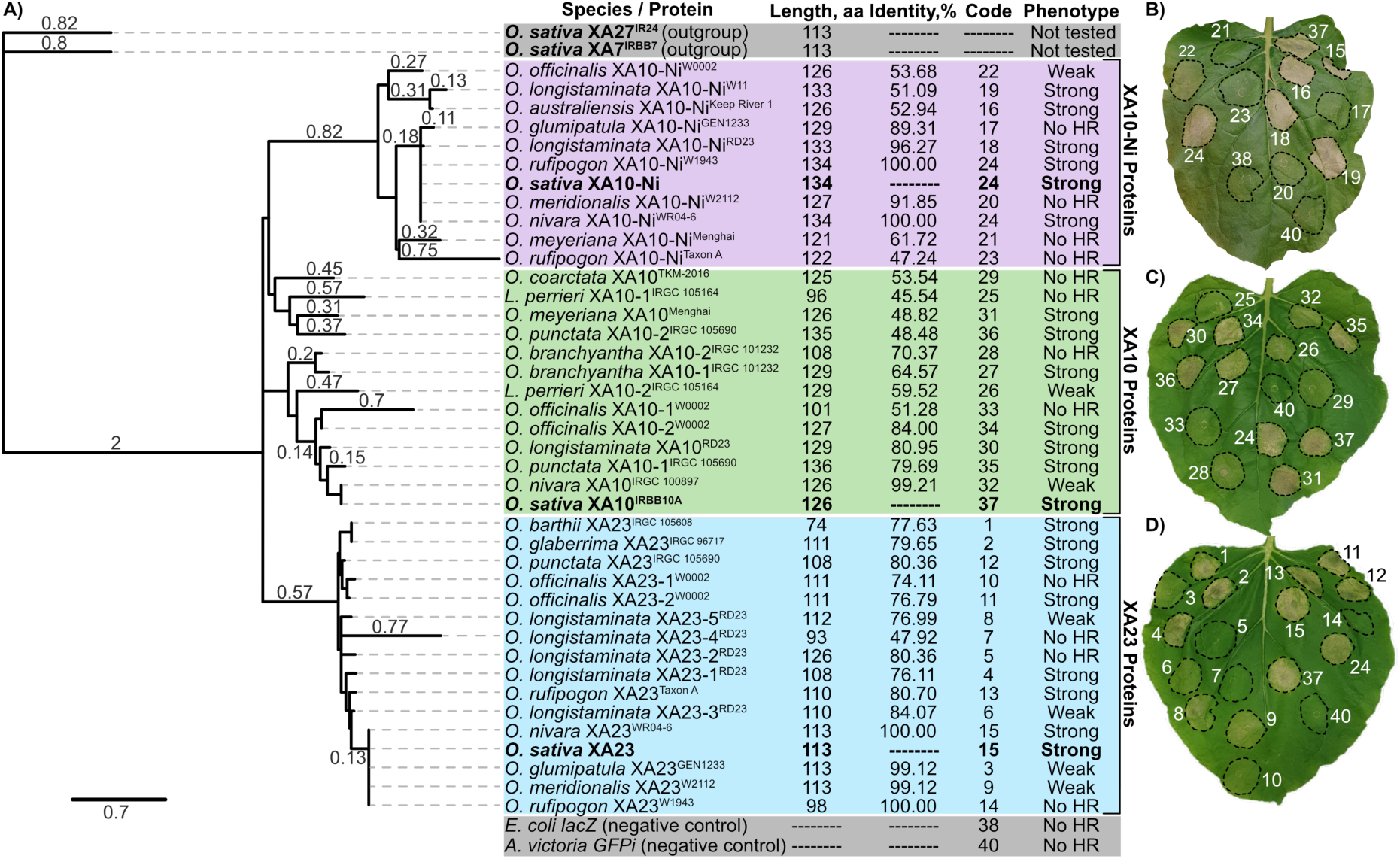
XA10, XA10-Ni, and XA23 homologous proteins from wild rice and cutgrass species trigger HR in *N. benthamiana*. **A)** Maximum-likelihood phylogenetic analysis of E protein sequences from cultivated and wild rice and cutgrass species. Values on tree branches represent substitutions per site based on 1000 bootstrap replications (*B* = 1000). Rice E proteins XA7 and XA27 were used as outgroups.^9,12^ Groups of XA10, XA10-Ni, and XA23 proteins are highlighted in green, purple, and blue, respectively. Values near the protein names represent the percent identity of homologous proteins to XA10, XA10-Ni, or XA23 from cultivated rice. Codes correspond to the assembled constructs used for the assay in B-D (Data S9). **B-D)** HR phenotypes induced by *Xa23* (B), *Xa10-Ni* (C), and *Xa10* (D) homologs upon transient overexpression in *N. benthamiana*. HR phenotypes were scored as “Strong”, “Weak”, or “No HR” (no cell death phenotype). *Aequorea victoria GFPi* and *Escherichia coli LacZ* served as negative controls. Photos were taken three days post-infiltration (3 dpi).

One evolutionary consequence of *E* gene family expansion is the degeneration of duplicated gene copies. Consistent with this, we found that 16 of 53 (30%) newly identified *E* gene-like loci show hallmarks of pseudogenization (Table S1). Specifically, two out of ten *E* gene-like loci in the genome of *O. longistaminata* accession RD23 contained predicted premature stop codons within the expected CDSs (Figure 1). In addition, one of six putative *Xa23* paralogs in the genome of *O. longistaminata* accession RD23 was predicted to encode a protein sharing only 22% amino acid identity with the rice XA23 executor. Together, the presence of premature stop codons and reduced protein sequence similarity (<40%) indicate that some *E* gene-like loci represent non-functional pseudogenes that likely arose through degenerative evolution following gene duplication.

After excluding 16 pseudogenes from further analysis, we examined whether the remaining 37 *E* gene-like loci contain *EBEs* compatible with the AvrXa10 or AvrXa23 TALEs, which would indicate that they can be transcriptionally activated by these effectors and thereby confer resistance to xanthomonads carrying these TALEs. None of the newly identified *E* gene-like loci contained *EBEs* compatible with AvrXa10 and AvrXa23 (Data S6-S8). This absence is consistent with the notion that *E* gene-like loci have not been maintained as TALE-responsive immune receptors in these species, further supporting the idea that immunity is not the primary or ancestral function shaping the evolution of this gene family.

### Functional diversification of *E* gene family members

To assess whether newly identified genes from wild rice and cutgrass species encode functional E proteins, we overexpressed predicted CDSs of these genes under the control of the constitutive *Cauliflower Mosaic Virus* (*CaMV*) *35S* promoter in *N. benthamiana* leaves via *Agrobacterium-*mediated transient delivery, and scored infection phenotypes. *E* gene overexpression induced hypersensitive response (HR) phenotypes of variable strength across all three gene classes, revealing substantial functional diversity. Among eight *Xa10-Ni*-like genes, four triggered strong HR, two weak HR, and two no visible response (Figure 2B). Similarly, six of 12 *Xa10*-like genes triggered strong HR, two weak, and four none (Figure 2C), while six of 14 *Xa23*-like genes triggered strong HR, three weak, and five none (Figure 2D).

Despite this variability, at least one HR-inducing *E* gene homolog was detected across nearly all tested wild rice and cutgrass species (Figure 2 and Table S1), suggesting that the presence of at least one functional *E* gene is selectively maintained. Thus, expansion of the *E* gene family has been accompanied by pronounced functional diversification, ranging from paralogs that retain strong cell death-inducing activity to paralogs with weak or undetectable activity. At the same time, the frequent absence of activity among paralogs is consistent with the idea that many *E* gene copies have accumulated inactivating mutations following duplication, leading to functional decay or pseudogenization.

### Domain-level analysis identifies critical regions for XA23-mediated cell death induction

Given the pronounced functional variation among XA23-like proteins, including both fully active and inactive homologs, we next exploited naturally occurring sequence variation to infer which protein regions are required for XA23-mediated cell death execution. Previous studies predicted that the 113-amino acid XA23 protein contains three transmembrane motifs (TMMs) of approximately 20 amino acids (TMM1-TMM3) whose length and hydrophobic properties are consistent with retention in the endoplasmic reticulum (ER) rather than localization to the plasma membrane.^11^ We therefore analyzed naturally occurring XA23-like proteins that differ in predicted domain composition or in the sequences within these motifs, and correlated these differences with their ability to induce HR in *N. benthamiana*.

We first tested variant XA23^W1943^ from *O. rufipogon* accession W1943, which lacks the first 15 N-terminal amino acids, including six residues of the predicted TMM1 (Data S5). This truncation abolished HR induction in *N. benthamiana*, indicating that the N-terminal region encompassing TMM1 is essential for XA23-mediated HR (Figure 2D). Next, we examined XA23-2 from *O. longistaminata* accession RD23 (XA23-2^RD23^), which carries a Tyr-to-Cys substitution at a conserved position 60 (Y60C) within TMM2 and failed to induce HR. This loss of activity implicates TMM2, and specifically this conserved residue, as critical for XA23-mediated cell death. Finally, we analyzed two near-identical XA23 variants from *O. glaberrima* accession IRGC 96717 (XA23^IRGC 96717^), which encodes a full-length, wild-type-like protein containing all three predicted TMMs, and *O. barthii* accession IRGC 105608 (XA23^IRGC 105608^), which encodes a C-terminally-truncated protein lacking the predicted TMM3. Notably, both variants triggered strong cell death when expressed in *N. benthamiana*, indicating that TMM3 is not required for XA23-mediated cell death execution. Taken together, these results identify TMM1 and TMM2 as essential determinants of XA23 function, while TMM3 is dispensable, thereby providing a domain-level framework for predicting the functional potential of XA23-like proteins based on sequence variation.

### Stage-specific, TALE-independent *Xa23* activation in rice zygotes

While TALE-induced *Xa23* expression in the immunity context is well studied, its native, TALE-independent expression has remained unexplored. If XA23 has a native function in development, *Xa23* would be expected to be expressed in specific tissues and/or at particular developmental stages. We therefore investigated when and where *Xa23* is transcribed independently of TALE-mediated activation, as a prerequisite for elucidating any native function distinct from its role as a TALE-inducible *R* gene.^20^ To detect native, TALE-independent expression of *E* genes, we mined published RNA-seq datasets from the RAP-DB database, comprising 46 datasets representing 840 rice samples collected from various rice tissues and conditions (Data S10).^31^ Because the rice Nipponbare reference genome lacks *Xa10*, precluding *Xa10*-specific read analysis, we focused on *Xa10-Ni* and *Xa23,* both of which are present in the reference genome.^19^

Notably, *Xa10-Ni*-specific reads were detected in none of the 46 datasets, whereas *Xa23*-specific reads were detected in only one dataset. This dataset included three related but distinct sample types: 1) unfertilized egg cells; 2) sperm cells; and 3) IVF-produced zygotes 4 hours after fusion (AF) of sperm and egg cells (Figures 3A and S1).^22^ Notably, *Xa23*-specific reads were detected in IVF-produced zygotes 4 hours AF, but not in sperm or egg cells. In rice, fusion of the two haploid nuclei (karyogamy) occurs approximately 2-4 hours AF and coincides with transcriptional changes known as zygotic genome activation (ZGA).^32–34^ The timing of *Xa23* expression therefore suggests that its transcriptional activation may be part of the ZGA program.

**Figure 3.**
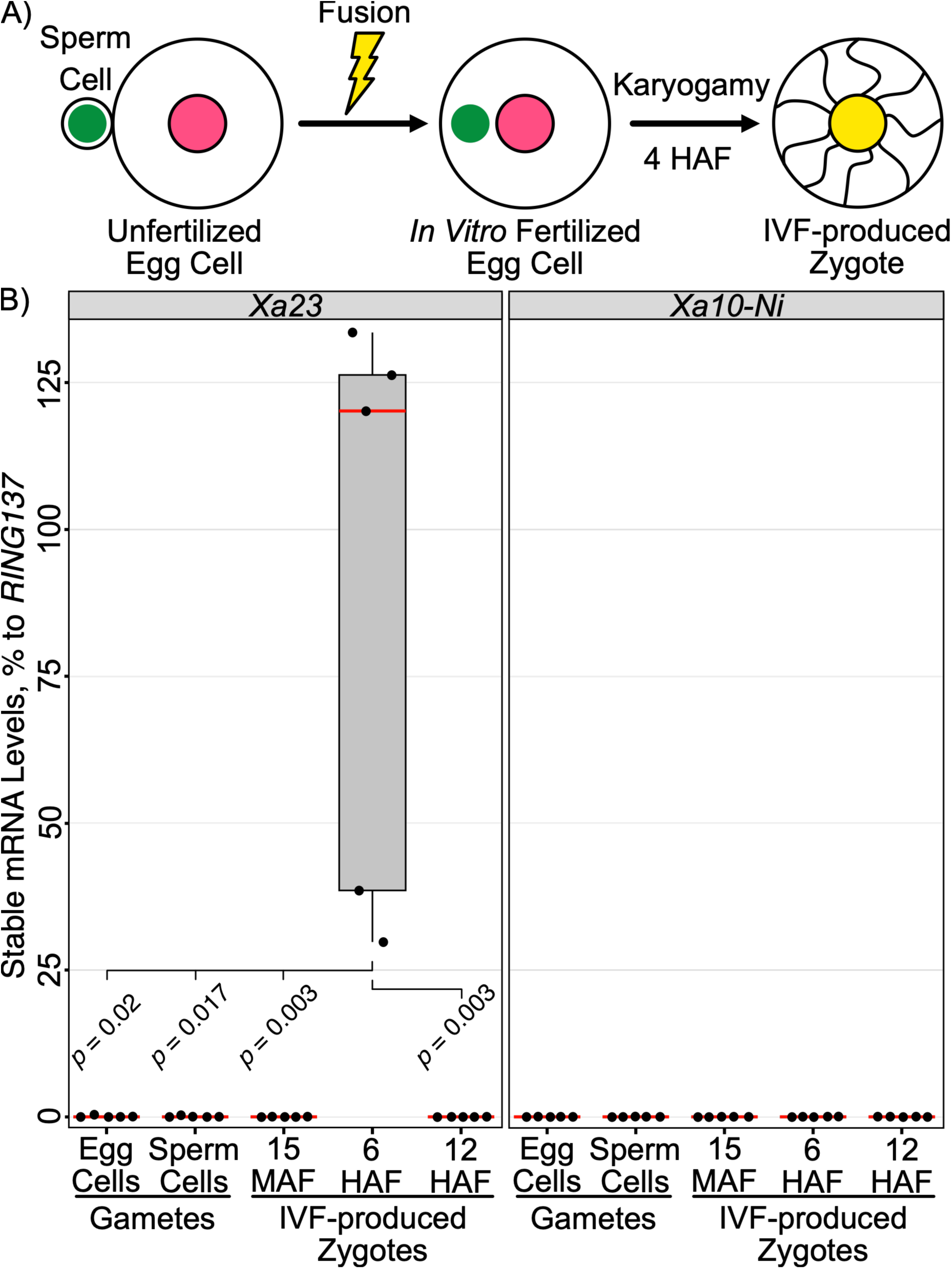
Stage-specific, native, TALE-independent transcriptional activation of *Xa23* is detected in IVF-produced zygotes. **A)** Schematic of the zygote production via IVF. **B)** Quantification of *Xa23* expression in gametes and IVF-produced zygotes via qRT-PCR. The housekeeping gene *RING137* served as a normalization control; *Xa10-Ni* served as a negative control. MAF / HAF, minutes/hours after fusion. Each sample contained 10 sperm cells, three unfertilized egg cells, or three IVF-produced zygotes (*n*=5). Black dots represent data points from individual biological replicates. Box plots show the median (center red line), lower and upper quartiles (box limits), and minimum and maximum values (whiskers). Statistical significance was assessed using the Independent-Samples Kruskal-Wallis Test and the Bonferroni *p*-value adjustment method for multiple comparisons. Only *p*-values ≤ 0.05 are shown.

Furthermore, we analyzed a follow-up time-series RNA-seq dataset covering the first 6 hours of zygote development.^23^ *Xa23*-specific reads first appeared 2 hours AF, remained detectable at 4 hours AF, and reached their highest abundance at 6 hours (Figure S2). No *Xa23*-specific reads were detected in either gamete or during the earlier stages of zygote development.

Taken together, these results demonstrate stage-specific, TALE-independent *Xa23* activation during early zygote development, consistent with a native role outside the immunity context. To exclude the possibility that *Xa23* activation was caused by the experimental conditions of the IVF system, including electrical stimulation and prolonged incubation in mannitol, we analyzed *Xa23* expression in IVF-treated gametes and zygotes collected at different stages after fertilization by quantitative reverse transcription polymerase chain reaction (qRT-PCR). Despite exposure to the IVF conditions, *Xa23* transcripts were not detected in unfertilized egg cells or in zygotes at 15 minutes AF (Figure 3B). Instead, *Xa23* expression was detected specifically at 6 hours AF (*p*-value = 0.003) and was no longer detectable at 12 hours AF. These results confirm the stage-specific, native, TALE-independent *de novo Xa23* transcription in IVF-produced zygotes at 6 hours AF and indicate that *Xa23* expression is tightly restricted to this developmental stage. Furthermore, our results suggest that *Xa23*, an *E* gene capable of triggering cell death upon TALE-mediated activation, might have a role during early zygote development.

### Histone deacetylase inhibition induces *Xa23* expression in unfertilized rice egg cells

Although we detected and confirmed stage-specific, native, TALE-independent *de novo Xa23* transcription in IVF-produced zygotes (Figures 3, S1, and S2), the mechanisms underlying *Xa23* transcriptional activation remained unknown. Because *Xa23* transcription in IVF-produced zygotes coincides with ZGA, we hypothesized that *Xa23* transcription is a consequence of epigenetic reprogramming of paternal and/or maternal chromatin during karyogamy, a process known to play a critical role in many developmental programs, including ZGA.^32–34^

To test whether *Xa23* activation can be achieved by epigenetic reprogramming of paternal and/or maternal chromatin, we searched for a chemical agent that can activate epigenetic reprogramming of chromatin in a fertilization-independent manner. We selected TSA, a non-selective HDACi that prevents HDAC-dependent removal of histone acetylation marks at regulatory chromatin regions, thereby upregulating proapoptotic genes in mammalian cancer cells.^35,36^ To examine the fertilization-independent *de novo* expression of *Xa23* in unfertilized egg cells that do not normally express *Xa23* (Figures 3, S1, and S2), we treated unfertilized egg cells with 10 µM TSA dissolved in dimethyl sulfoxide (DMSO) or with DMSO (solvent) only and quantified *Xa23* transcripts 24 hours after treatment via qRT-PCR (Figure 4A). Following this experiment, *Xa23* transcripts were detected exclusively in TSA-treated samples, but not in DMSO controls (*p*-value = 0.015; Figure 4B). Notably, TSA treatment did not induce *Xa10-Ni* expression (*p*-value = 1.0), indicating that transcriptional regulation of *Xa23* and *Xa10-Ni* differs, at least in some respects. Taken together, these findings support our initial hypothesis that chemical treatment can activate *Xa23* transcription in a fertilization-independent manner, likely through epigenetic mechanisms involving histone acetylation.

**Figure 4.**
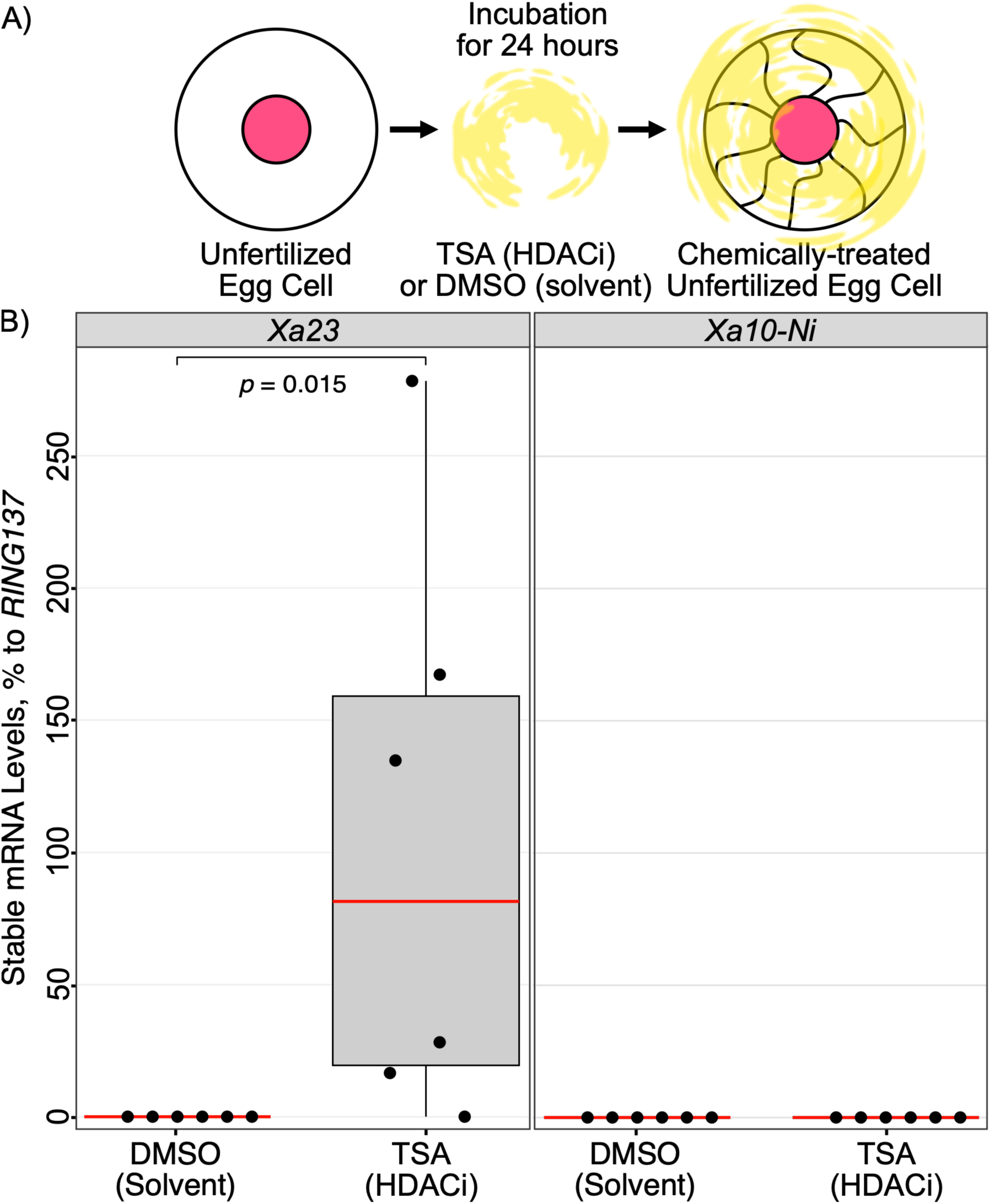
TSA treatment activates *Xa23* transcription in unfertilized rice egg cells. **A)** Schematic of the chemical treatment of unfertilized egg cells. **B)** Quantification of *Xa23* expression in DMSO- and TSA-treated unfertilized egg cells via qRT-PCR. *RING137* served as a normalization control; *Xa10-Ni* served as a negative control. Each sample contained three cells (*n*=6). Black dots represent data points from individual biological replicates. Box plots show the median (center red line), lower and upper quartiles (box limits), and minimum and maximum values (whiskers). Statistical significance was assessed using the Independent-Samples Mann-Whitney U Test. Only *p*-values ≤ 0.05 are shown.

### Native *Xa23* transcripts in zygotes structurally resemble cell death-inducing variants

While the qRT-PCR-based assay enabled detection and quantification of native *Xa23* mRNA transcripts in IVF-produced zygotes and TSA-treated unfertilized egg cells (Figures 3 and 4), it remained unclear whether native *Xa23* transcript variants induced during developmental processes differ from the (d)TALE-induced *Xa23* transcript variants within the immunity context. In this regard, previous studies have demonstrated that native and (d)TALE-induced mRNA transcripts of host *S* genes often have different TSSs and may therefore encode distinct protein variants.^5,37^

To investigate whether native *Xa23* mRNAs encode potentially cell death-inducing XA23 proteins, we reconstructed putative full-length native *Xa23* transcripts *in silico* using short RNA-seq reads from previously analyzed datasets.^22,23^ Following our analysis, we identified six splice variants of native *Xa23* transcripts that differ in their start sites, lengths, and *Xa23* CDS coverage (Figures 5 and S3 and Data S11). However, short-read RNA-seq is not well suited to reliably reconstruct transcripts, identify TSSs, and characterize UTRs,^38^ especially when transcript abundance is low, as is the case for native *Xa23* transcripts.

**Figure 5.**
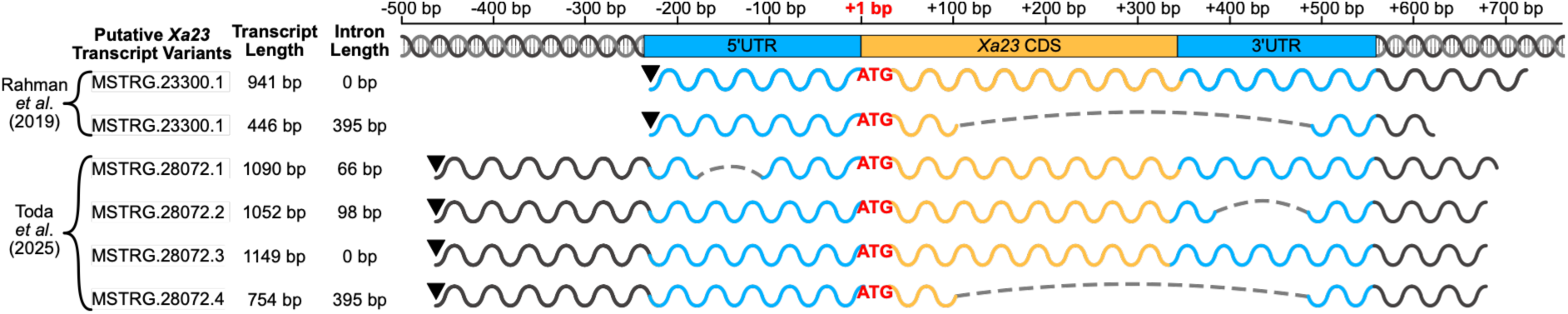
Native, TALE-independent transcription of *Xa23* might generate multiple transcript variants. Putative native *Xa23* transcripts were reconstructed *in silico* using short RNA-seq reads from previously analyzed datasets (Figure S3).^22,23^ *Xa23* UTRs are shown in **blue** and the *Xa23* CDS in **orange**. The first nucleotide of the *Xa23* start codon (**ATG**) is designated as “**+1**”. Thick wavy lines represent reconstructed putative *Xa23* transcript variants with TSSs (▾) and introns (grey dashed arc lines). Full-length sequences of the reconstructed putative *Xa23* transcripts are available in Data S11.

To accurately characterize native *Xa23* transcripts, we performed 5’RACE and 3’RACE on mRNA extracted from TSA-treated and DMSO-treated unfertilized egg cells. 5’RACE and 3’RACE PCR amplicons were obtained only from TSA-treated unfertilized egg cells (Figure 6A), again confirming that transcriptional activation of *Xa23* can be achieved via TSA treatment. Sequencing of the 5’RACE products revealed two distinct types of 5’UTRs: a long 5’UTR (L 5’UTR), represented by a single cloned amplicon, and a short 5’UTR (S 5’UTR), represented by four cloned amplicons of non-identical but similar size (Figure 6B and Data S12). The L 5’UTR is 120 nucleotides (nt) long and contains a 91-nt intron with a TSS located 211 nt upstream of the start codon. In contrast, the four amplicons with an S 5’UTR are intronless and range in length from 35 to 56 nt.

**Figure 6.**
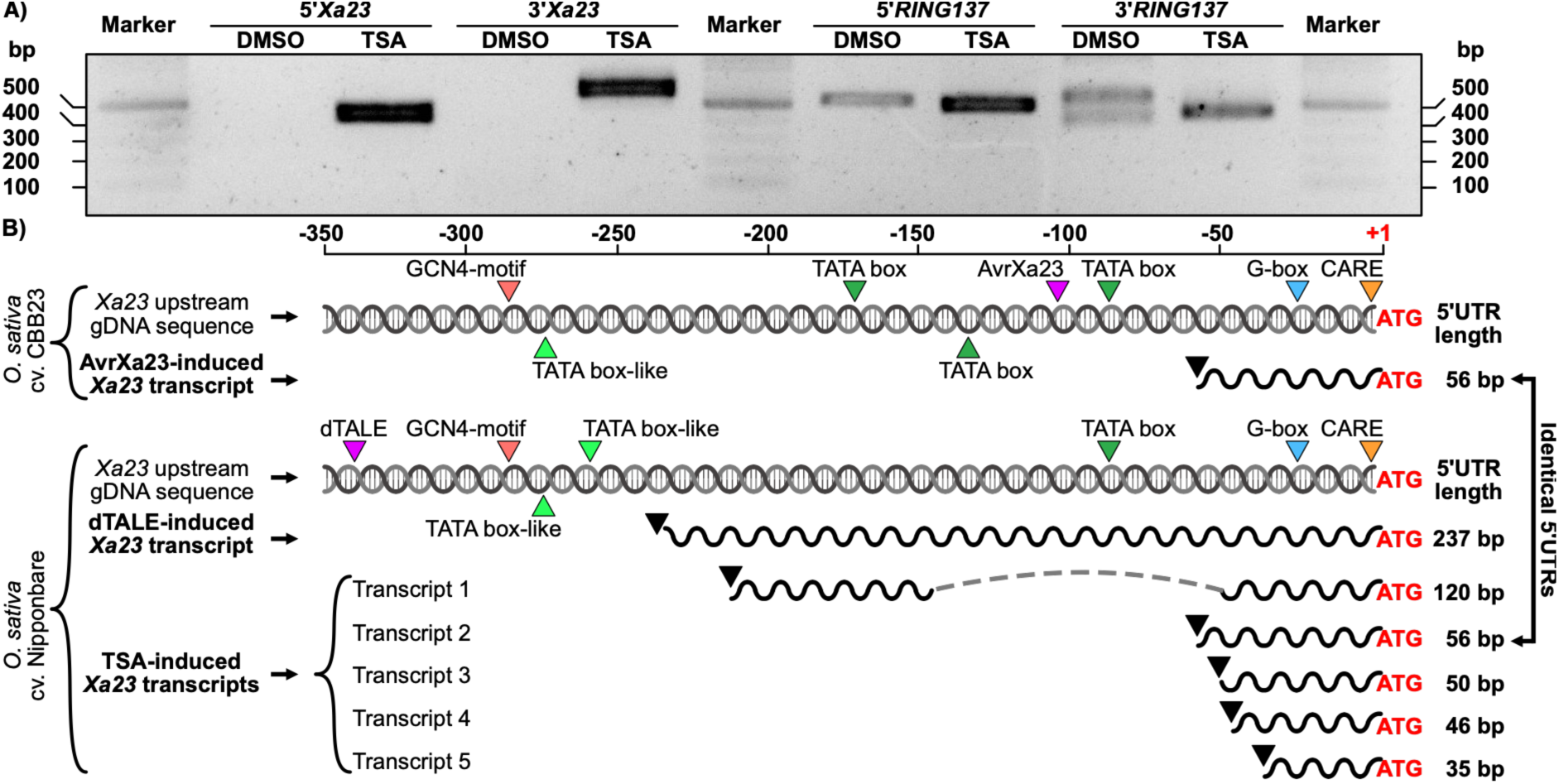
TSA-induced *Xa23* transcripts contain short and long 5’UTRs. **A)** 5ʹand 3’RACE analysis of *Xa23* and *RING137* using mRNA from unfertilized egg cells treated with DMSO and 10 μM TSA. **B)** Schematic comparison of the 5’UTRs of TSA-induced, dTALE-induced, and AvrXa23-induced *Xa23* transcripts.^11,19^ The first nucleotide of the *Xa23* start codon (**ATG**) is designated as “**+1**”. Thick wavy lines represent 5’UTRs of *Xa23* transcripts with TSSs (▾) and introns (grey dashed arc lines). Other annotations include AvrXa23 and the dTALE binding elements (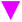),^11,19^ TATA boxes (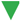), TATA box-like sequences (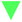), GCN4 motifs (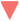), G boxes (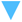), and CARE (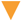). The genomic sequences upstream of the TATA boxes (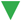; position −87) are polymorphic. Full-length sequences of the characterized TSA-induced *Xa23* transcripts are available in Data S9 and Data S10.

We also compared newly identified 5’UTRs from TSA-induced *Xa23* transcripts with those induced by AvrXa23 TALE (56-nt 5’UTR) and dTALE (237-nt 5’UTR).^11,19^ We discovered that the TSA- and AvrXa23-induced transcripts share an identical 56-nt 5’UTR (Figure 6B and Data S12). This structural similarity strongly suggests that TSA-induced *Xa23* transcripts are functionally equivalent to the cell death-inducing AvrXa23-induced transcripts.

Following analysis of 5’RACE products, we subcloned and characterized the 3’RACE amplicons of TSA-induced *Xa23* transcripts and found no splicing of the *Xa23* CDS and no major differences in 3’UTR length compared with dTALE-induced and AvrXa23-induced *Xa23* transcripts (Data S13).

In summary, the structural comparison of TSA-induced and AvrXa23-induced *Xa23* transcripts suggests they are functionally equivalent, which may imply that native *Xa23* transcripts encode the functional XA23 protein capable of cell death execution in IVF-produced zygotes and unfertilized egg cells.

### Conserved *cis*-elements may mediate *Xa23* transcription in zygotes

Following the detection of native *Xa23* transcripts in IVF-produced zygotes (Figure 3), we asked whether CREs upstream of *Xa23* might contribute to its stage-specific activation. To identify candidate CREs, we compared the 350 nt regions upstream of the *Xa23* start codon in two rice genotypes, Nipponbare and CBB23, both previously used for dTALE- and AvrXa23-mediated transcriptional activation of *Xa23*.^11,19^ Despite only 48% sequence identity, both genotypes shared several candidate CREs, including GCN4 motifs, canonical TATA boxes, TATA-like sequences, G boxes, and “CAACTC” regulatory elements (CAREs; Figure 6B and Data S12). Among these CREs, the TATA box is a well-characterized eukaryotic core promoter element.^39^ A canonical TATA box (“TATAAAA”) was conserved 79 nt upstream of the *Xa23* start codon in both genotypes (Figure 6B and Data S12). It is located 23-44 nt upstream of the TSSs of *Xa23* transcripts with S 5’UTRs, consistent with a potential role in TALE-independent transcription. Additionally, Nipponbare contained two TATA-like sequences (“TATATTA” and “TATTATA”) 63 and 48 nt upstream of the TSS of *Xa23* transcripts with L 5’UTRs, suggesting that these sequences may support the generation of *Xa23* transcripts with L 5’UTRs.

Furthermore, we identified GCN4 motifs upstream of the TATA boxes in both genotypes (Figure 6B and Data S12). Previous work links GCN4 motifs to five basic leucine zipper TFs (RISBZ1-5).^40,41^ Intriguingly, *RISBZ1*/*bZIP58* and *RISBZ5*/*bZIP52* are expressed in IVF-produced zygotes, but not in gametes (Figure S4).^22^ However, whether RISBZ1/bZIP58 or RISBZ5/bZIP52 can activate *Xa23* transcription remains to be tested.

In summary, the conservation of TATA boxes and other elements across divergent promoters supports their role in regulating native, TALE-independent *Xa23* transcription in diverse genetic backgrounds.

## Discussion

### Evolutionary conservation supports functions of *E* genes beyond immunity

Originally, *E* genes were discovered in the immunity context.^10,11,19^ Transcriptional activation of *E*genes by cognate TALEs from *Xoo* or dTALEs triggers host cell death and confers resistance to xanthomonads. However, whether *E* genes have native functions beyond immunity has remained unknown. Prior to this study, TALE-independent *E* gene transcription in its endogenous biological context had not been reported.

We started this study by asking whether the evolutionary patterns of rice *E* genes resemble those of canonical race-specific immune receptor genes or instead point to functions beyond immunity. Analysis of 3,002 cultivated rice accessions revealed remarkably strong conservation of *Xa10-Ni* and, particularly, *Xa23* (Data S1 and S2). Full-length *Xa10-Ni* coding sequences were retained in 97% of accessions, whereas a full-length *Xa23* coding sequence was present in all 3,002 accessions, with XA23 proteins sharing 96–100% amino acid identity. Such widespread retention and high sequence conservation contrast with the extensive presence/absence variation and allelic diversification characteristic of many race-specific immune receptor genes and suggest that these *E* genes are subject to evolutionary constraints unrelated to their established role in TALE-triggered immunity.

Extending the analysis across the Oryzeae revealed that the *E* gene family is evolutionarily ancient. *Xa10*-, *Xa10-Ni*-, and *Xa23*-like genes were detected in multiple wild *Oryza* species, and *Xa10*-like sequences were also present in the related genus *Leersia* (Figure 2 and Table S1). Thus, the major *E* gene lineages originated before the divergence of *Oryza* and *Leersia* approximately 14 MYA.^42^ At the same time, pronounced differences in *E* gene copy number among species and even between accessions of the same species indicate recurrent lineage-specific duplication and gene loss. Consistent with such dynamic family evolution, a subset of the identified loci showed hallmarks of pseudogenization, while the remaining proteins displayed substantial variation in their ability to induce cell death upon overexpression in *N. benthamiana*. Similar functional variation has been reported for members of the *Xa27* family, in which only a subset of paralogs and alleles retain cell death-inducing activity.^43,44^ Together, these observations suggest that *E* gene duplication has been followed by divergent evolutionary trajectories, including retention of cell death-inducing activity, functional diversification, and degenerative loss.

Importantly, none of the newly identified *E* gene-like loci contained *EBEs* compatible with AvrXa10 or AvrXa23 (Data S6-S8). Thus, although the *E* gene family is ancient, there is no indication that maintenance of TALE-responsive promoters has driven its long-term evolution. Instead, these findings support a model in which *E* genes originated and were maintained for functions unrelated to TALE recognition, while individual alleles were secondarily recruited into immunity through the acquisition of promoter sequences recognized by pathogen TALEs.^20^ In this scenario, TALE-responsive *E* genes represent evolutionary traps that exploit the intrinsic cell death-inducing potential of an older gene family rather than immune receptors that evolved primarily for pathogen recognition.

Among the analyzed E proteins, XA23 showed particularly strong sequence conservation (Table S1 and Data S5), making it an attractive model for investigating the molecular basis of executor-mediated cell death and the native function of this protein family. Natural variation among XA23-like proteins indicates that TMM1 and TMM2 are required for cell death induction, whereas TMM3 appears dispensable (Figure 2 and Data S5). Together with the detection of native, TALE-independent *Xa23* transcription in zygotes (Figure 3), these evolutionary and functional observations support the hypothesis that *E* genes have native roles beyond immunity. The nature of these native functions, however, remains to be determined.

### *Xa23* native function can be discovered in IVF-produced zygotes and TSA-treated egg cells

To identify potential native functions of rice *E* genes, we searched for their native, TALE-independent expression across cell types, tissues, organs, and developmental stages. RNA-seq mining revealed *Xa23* transcripts exclusively in rice zygotes (Figures S1 and S2).^22,23^ Transcripts first appeared 2 hours AF, remained detectable at 4 hours, and peaked at 6 hours AF; qRT-PCR independently confirmed expression at 6 hours AF, whereas transcripts were no longer detectable at 12 hours AF (Figure 3). This narrow temporal expression window represents evidence of TALE-independent transcriptional activation of an *E* gene in its endogenous context, supporting the hypothesis that native expression of *E* genes is tightly controlled and restricted to specific cell types and developmental stages.^20^

Given that constitutive expression of *E* genes can cause severe developmental defects,^10,45^ the XA23 activity may be restricted to the earliest stages of zygote development. Although the native function of *Xa23* remains unknown, the IVF system provides an experimental platform to investigate it.

Notably, TSA treatment of unfertilized rice egg cells activated *Xa23* transcription (Figure 4). This proof-of-principle experiment demonstrates that *Xa23* transcription can be induced independently of both TALEs and fertilization and provides a complementary system for dissecting the mechanisms controlling *Xa23* transcription and determining the consequences of its activation.

### Transcription of *Xa23* is likely epigenetically regulated during zygote development

Transcriptome profiling of IVF-produced zygotes revealed tightly controlled *de novo Xa23* transcription post-karyogamy (Figures 3, S1, and S2). Treatment of unfertilized egg cells with TSA activated *Xa23* transcription (Figure 4), implicating histone acetylation and chromatin accessibility in its regulation. Consistent with this possibility, 3D genomes from rice gametes undergo dynamic reorganization during karyogamy,^46^ including sperm-derived chromatin decondensation and remodeling of paternal DNA methylation.^33,47^ Therefore, *Xa23* transcription in zygotes may be enabled by chromatin decondensation caused by histone acetylation.

However, chromatin accessibility alone is unlikely to explain *Xa23* transcription, suggesting that *Xa23* transcription additionally requires sequence-specific TFs. Using *in silico* analysis, we identified numerous known CREs within the putative *Xa23* promoter, including the canonical TATA box adjacent to the TSS of *Xa23* transcripts with S 5’UTRs, TATA box-like sequences adjacent to the TSS of *Xa23* transcripts with L 5’UTRs, and GCN4 motifs upstream of TATA box-like sequences (Figure 6). The known GCN4 motif-binding activity of RISBZ1/bZIP58 and RISBZ5/bZIP52,^40,41^ and their spatiotemporal expression patterns (Figure S4), identify these TFs as candidate upstream regulators of *Xa23* during zygote development. Future studies using targeted candidate CRE mutagenesis,^48^ TurboCas proximity labeling,^49^ yeast one-hybrid screening,^50^ and dual-visible reporter assays^51^ could experimentally define the CREs controlling native *Xa23* expression and identify TFs interacting with the *Xa23* promoter during zygote development.

### XA23 may contribute to Ca^2+^ signaling during early zygote development

The transient *Xa23* activation during early zygote development raises questions about its potential function in this developmental context (Figure 3). Known properties of XA23 provide a possible link to Ca²⁺ signaling. XA23 forms oligomers through self-interaction, localizes to the ER, and initiates free Ca^2+^ depletion.^19^ Because fertilization triggers elevation of intracellular Ca^2+^ levels,^52–54^ which facilitate karyogamy,^34^ ER reorganization,^55^ and exocytosis of cell wall materials presumably to prevent polyspermy,^56,57^ we broadly hypothesize that XA23 may act as a Ca^2+^ channel that contributes to fertilization-induced Ca^2+^ signaling. This hypothesis is consistent with the accumulation of Ca^2+^ precipitates in intracellular compartments of IVF-produced zygotes 6 hours AF.^58^

One potential downstream process affected by XA23 is cell-wall remodeling. Previous reports have linked other rice E proteins to cell wall-associated processes: ectopic *Xa27* expression leads to secondary cell wall thickening and reduced pit diameter, ^12,59^ while ectopic *Xa7* expression upregulates lignin biosynthesis pathways.^60^ Because native, TALE-independent *Xa23* expression peaks at 6 hours AF (Figure 3), much after primary cell wall biosynthesis (20 minutes to 2 hours AF),^34,53,56^ XA23 is unlikely to contribute to early cell wall formation. Instead, XA23 may contribute to secondary cell wall formation via Ca^2+^ release during early zygote development.

Alternatively, *Xa23* activation may be part of a fertilization-dependent stress response. The DNA damage checkpoint kinase WEE1 from rice is transiently expressed in IVF-produced zygotes following the parental genome mixing and restoration events at the same developmental stage as *Xa23* (6 hours AF; Figure 3).^33,61^ Similarly, *Arabidopsis thaliana WEE1* is upregulated in response to DNA stress and controls cell-cycle arrest upon activation of the DNA integrity checkpoint.^62^ Therefore, the expression of rice *Xa23* and *WEE1* in IVF-produced zygotes at 6 hours AF may represent a molecular mechanism protecting against fertilization-dependent cellular stress.

Reactive oxygen species (ROS) may provide another link between XA23 and zygote physiology. ROS levels are high in unfertilized egg cells, progressively decline during early zygote development (1-8 hours AF), and then gradually increase again before the first cell division (16 hours AF).^63^ Inhibition of intracellular ROS production by malonate and diphenyleneiodonium in zygotes causes cell division arrest.^63^ Because ectopic, continuous, or transient expression of *E* genes coincides with increased ROS levels,^43,60^ *Xa23* activation might serve as a cellular tool in shifting ROS dynamics in developing rice zygotes.

### Does E protein-mediated cell death depend on the level of *E* gene expression?

*E* genes are primarily characterized by their ability to trigger cell death in plant cells upon TALE-mediated ectopic expression.^10,11,19^ However, it is possible that such cell death does not occur under native, host-driven expression conditions, and the observed cell death may instead result from abnormally high expression levels not reached in the developmental context. In our system, native *Xa23* expression does not correlate with an obvious cell death response (Figure 3), supporting the hypothesis that executor-mediated cell death may occur only upon ectopic, not native, expression.

This raises the question of whether the structure and predicted molecular function of known E proteins can serve as a basis for a working hypothesis that reconciles both their role in development, where native expression does not lead to cell death, and their ability to induce cell death when overexpressed.^9–12^

Most E proteins encode ER-localized transmembrane proteins with similarity to calcium transporters.^10,19,64^ The ER serves as a major intracellular calcium reservoir, and controlled calcium release from the ER through specific channels is a well-established mechanism for activating signaling cascades during development.^34,53–55,57,65^ However, if the expression levels or activities of proteins that mediate Ca^2+^ export from the ER are abnormally high, this may lead to excessive Ca^2+^ release and ultimately trigger a cell death response.^66^

In summary, the proposed function of E proteins as ER-localized transmembrane proteins mediating Ca^2+^ release is consistent with both their ability to induce cell death upon TALE-mediated overexpression and their potential native role as transcriptionally activated signaling components that modulate developmental processes via ER-derived calcium signaling.

### Epigenetics-targeting drugs as tools for the putative discovery of genes with pro-apoptotic properties

*E* genes may be widespread across plant species but remain unannotated or unidentified in model species such as Arabidopsis due to the tightly controlled spatiotemporal expression.^20^ Therefore, if some *E* genes are yet-unannotated in plant genomes, their corresponding reads would be discarded as unmapped reads during RNA-seq data analysis.^67^ Given the roles of HDACis, DNA methyltransferase inhibitors, and other epigenetics-targeting drugs in altering gene transcription,^68–71^ treating plant tissues with these chemical agents, followed by RNA-seq analysis, could uncover novel genes with pro-apoptotic properties.^67^ This approach would lay the foundation for expanding our understanding of cell death control mechanisms and help to elucidate developmental cell death programs and pathways.

As a proof-of-principle experiment, we used TSA to activate *Xa23* transcription ectopically (Figures 4 and 6). Although *Xa10-Ni* is located only 22,177 nucleotides upstream of *Xa23,*^10^ TSA treatment did not activate *Xa10-Ni* transcription. The mechanism behind the differential expression of *Xa10-Ni* and *Xa23* upon TSA treatment remains unclear. Therefore, future studies should examine diverse cell types, organs, and tissues across different developmental stages to determine whether, when, and where *Xa10-Ni* is natively expressed.

Additionally, one could test whether treatments with epigenetics-targeting drugs can activate transcription of other known rice *E* genes, namely *Xa7*,^9^ *Xa10*,^10^ *Xa27*,^12^ and *Xa27B*,^43^ in different tissues and at different developmental stages. Recently, Xu *et al*.^72^ tested four HDACis in their ability to enhance rice resistance to *Magnaporthe oryzae* and showed that sodium butyrate treatment significantly enhanced disease resistance by upregulating numerous rice defense-related genes. Further mining of RNA-seq datasets might help detect TALE-independent activation of other known *E* genes and potentially yet-unannotated genes with pro-apoptotic properties.

### Limitations of the study

Although we identified native, TALE-independent transcriptional activation of *Xa23* during early zygote development, it remains unknown which upstream pathway components activate *Xa23* transcription and whether *Xa23* transcripts are translated in the developmental context. Additionally, future studies should determine the role of *Xa23* in zygote development. Lastly, it would be advantageous to mine publicly available RNA-seq datasets for evidence of transcription of other *E* genes in contexts unrelated to immunity and to test whether treatments with epigenetics-targeting drugs activate transcription of these *E* genes in various plant tissues.

## Resource availability Lead contact

Further information and requests for resources and reagents should be directed to and will be fulfilled by the lead contact, Kirill Schenstniy.

## Materials availability

Generated plasmids will be available upon request from the lead contact with a completed materials transfer agreement.

## Data and code availability

- RNA-seq data are deposited in the Data Bank of Japan (ID: DRA007969) and the DDBJ Sequence Read Archive (ID: DRA014663).^22,23^
- This paper does not report original code.
- Any additional information required to reanalyze the data reported in this paper is available from the lead contact upon request.

## Acknowledgments

We thank Dr. Tom Schreiber and Dr. Sylvestre Marillonnet (Leibniz Institute of Plant Biochemistry, Germany) for the pAGM53151 plasmid, Dr. Niels Gallas (Eberhard Karls University Tübingen, Germany) for technical suggestions on RACE, Ms. Tomoko Mochizuki (Tokyo Metropolitan University, Japan) for isolating egg cells, and RIKEN Bio Resource Center (Tsukuba, Japan) for cultured rice cells (Oc line). We gratefully acknowledge funding from 1) the Deutsche Forschungsgemeinschaft (German Research Foundation) EXC-2048/1 (project ID: 390686111), LA1338/20-1, LA1338/19-1, LA1338/18-1, and TRR356 (project ID: 491090170, subproject B03); 2) the BMFTR (INNO-TOM; 031B1538B); and 3) JSPS KAKENHI (Grant-in-Aid for Scientific Research(B), grant no. 22H02315 and 25K01990 to T.O.).

## Author contributions

KS conceived and designed experiments. KS and LR performed genome mining and phylogenetic analysis of *E* gene alleles and homologs. KS, AS, and NF performed transient overexpression of *E* gene homologs in *N. benthamiana*. KS, KR, and AH performed IVF experiments. KS and KR performed TSA treatment of unfertilized egg cells. KS performed qRT-PCRs and data analysis. KS, ASH, ET, and AK characterized *Xa23* transcripts. KS prepared figures. KS, DRH, and TL wrote the draft. All authors reviewed and edited the final version of the manuscript. TL and TO provided funding and supervision.

## Declaration of generative AI and AI-assisted technologies in the writing process

The authors used ChatGPT-5.6 Sol and Grammarly to improve the readability and language of the draft manuscript, subsequently reviewed and edited the content, and take full responsibility for the published article.

## STAR ★ Methods

Detailed methods are provided in the online version of this paper and include the following:

- KEY RESOURCES TABLE
- EXPERIMENTAL MODEL AND STUDY PARTICIPANT DETAILS
- `o` Plant materials and growth conditions
- METHOD DETAILS
- `o` Analysis of *Xa10-Ni* and *Xa23* alleles from 3,002 cultivated rice accessions
- `o` Mining for *E* gene-like sequences in genomes of wild rice and cutgrass species
- `o` Assembly of constructs
- `o` Overexpression of *E* genes in *N. benthamiana*
- `o` *In vitro* gamete fusion and zygote production
- `o` HDACi treatment of unfertilized egg cells
- `o` Total RNA isolation and cDNA synthesis
- `o` Quantitative real-time PCR
- `o` Characterization of native *Xa23* transcripts
- `o` Characterization of the TSA-induced *Xa23* transcripts via 5’ and 3’ RACE
- QUANTIFICATION AND STATISTICAL ANALYSIS

### Key resources table

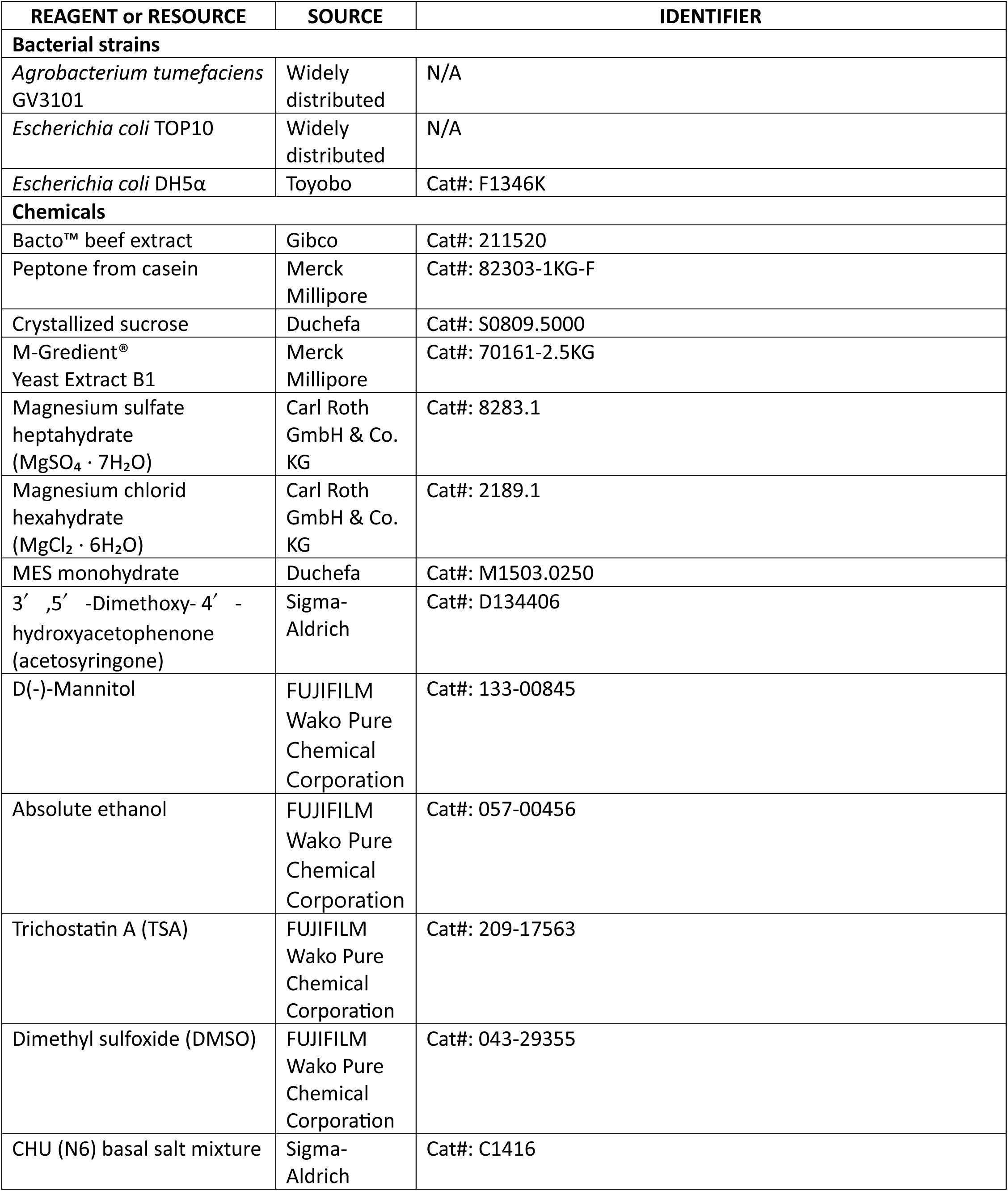
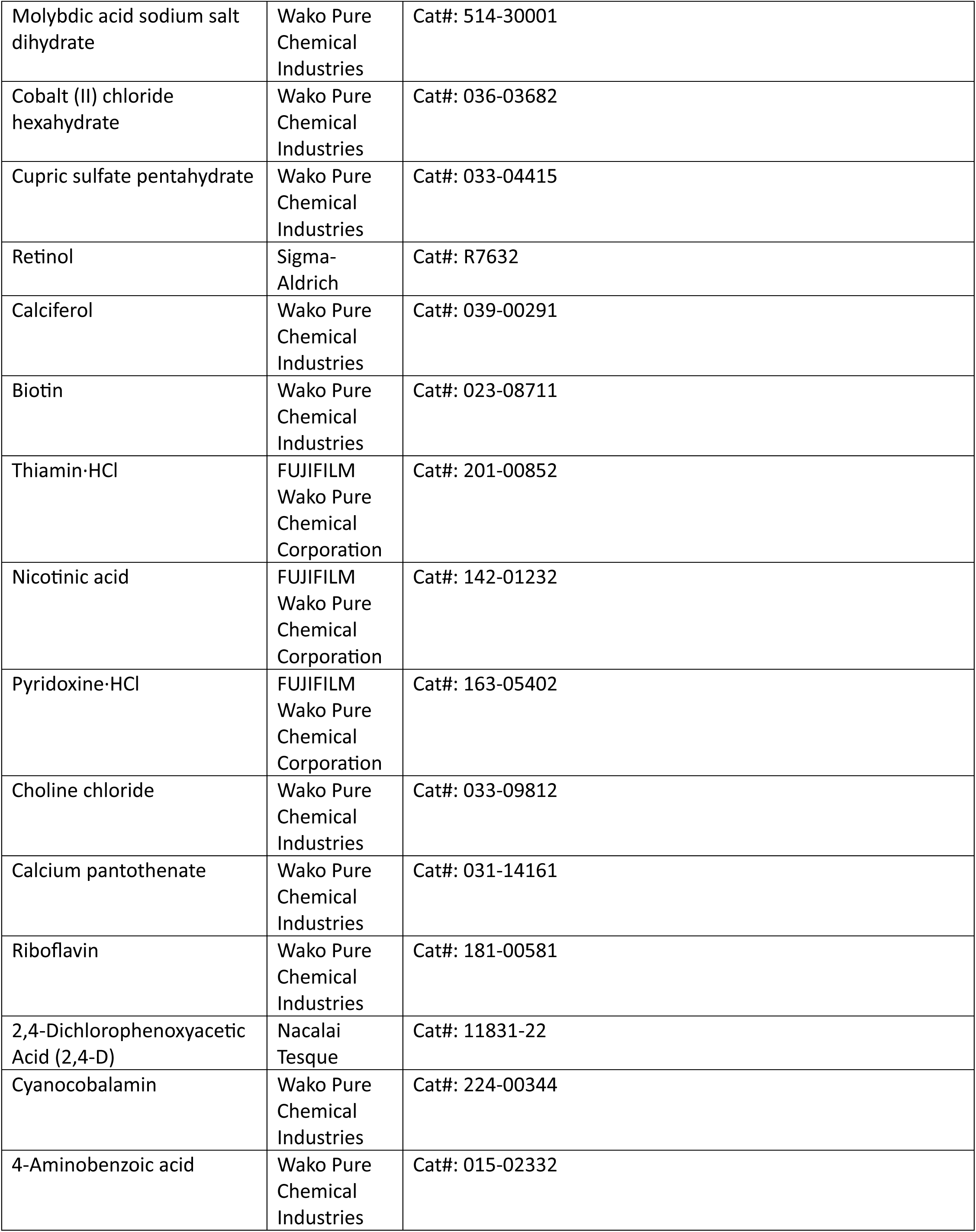
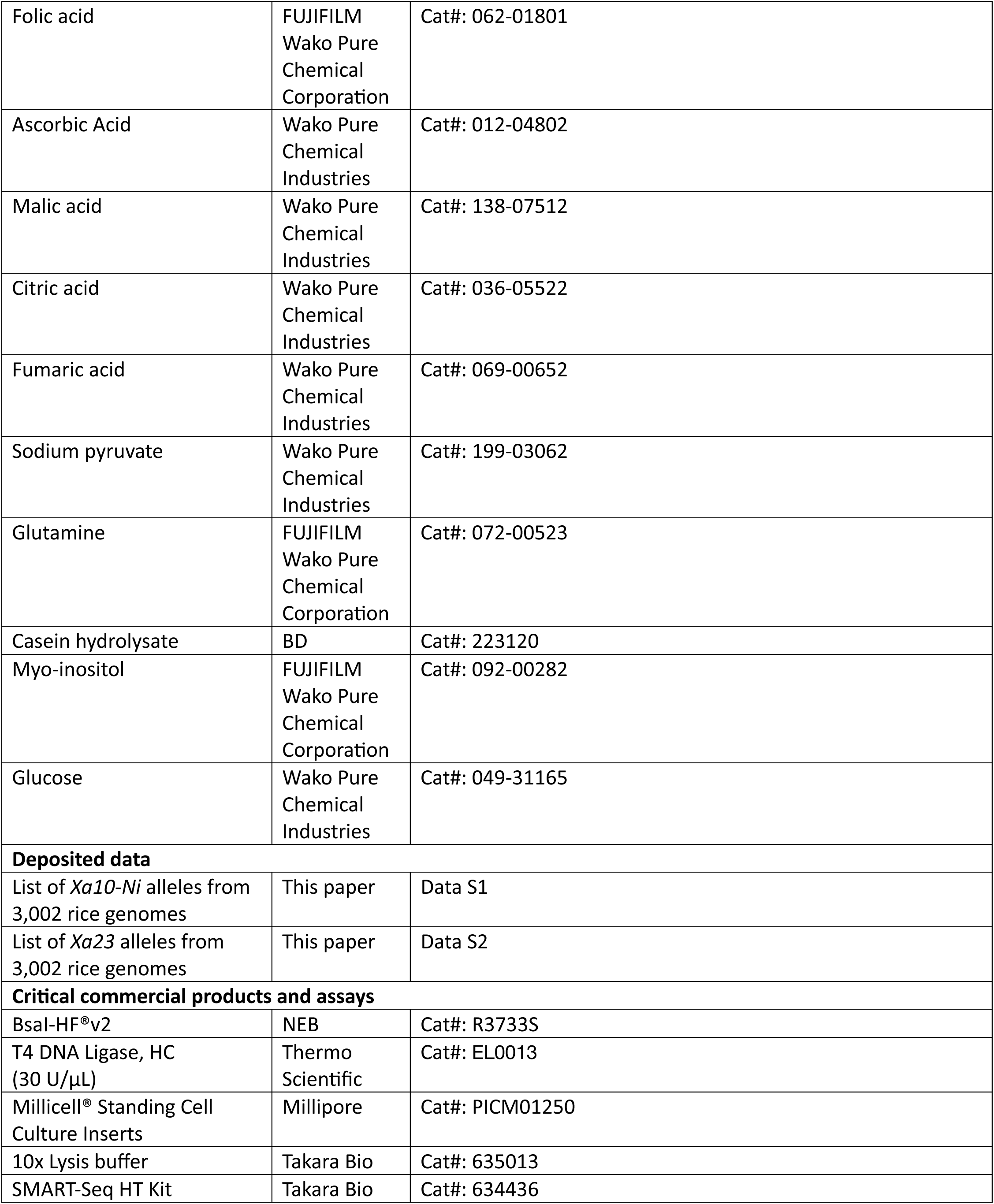
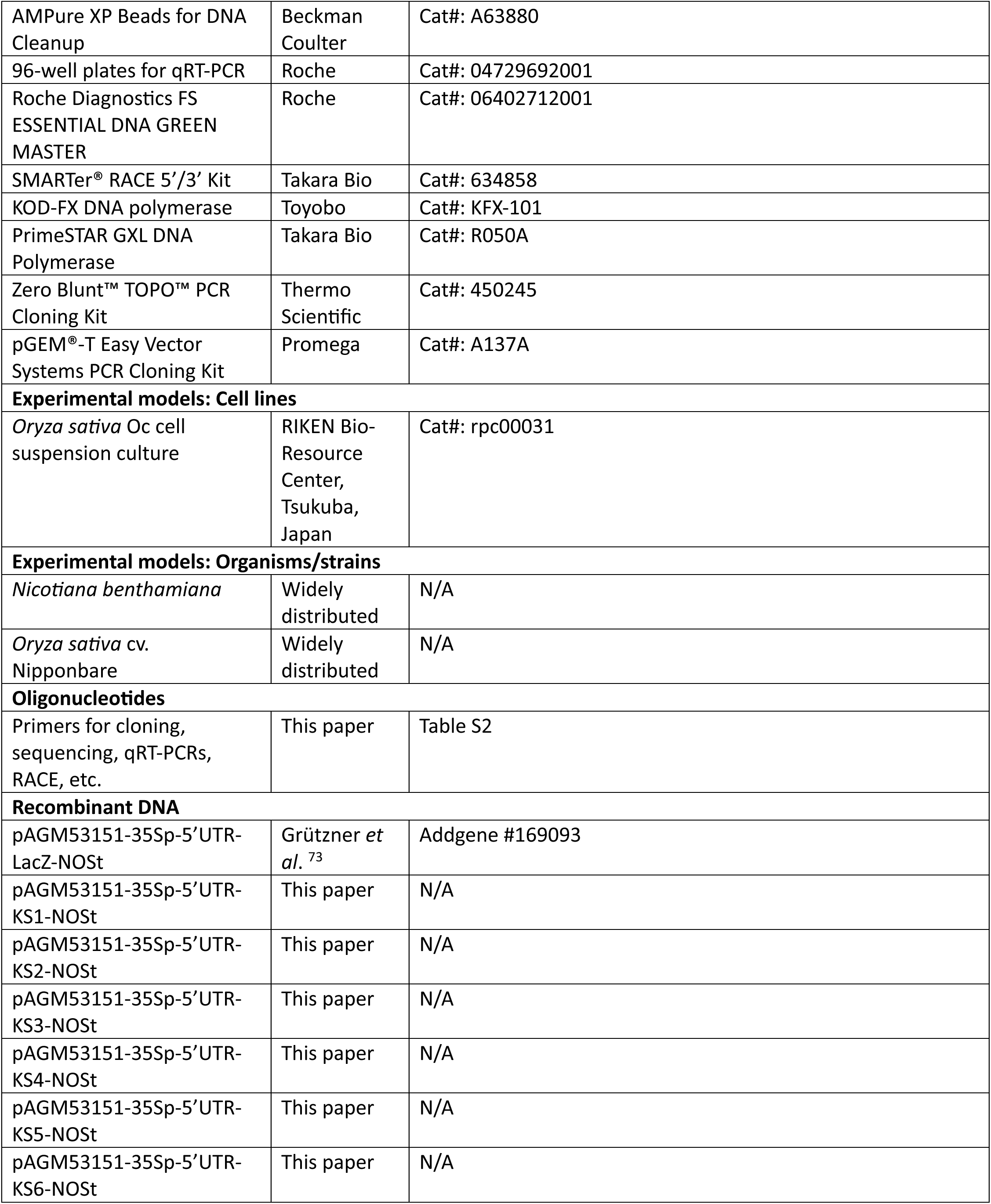
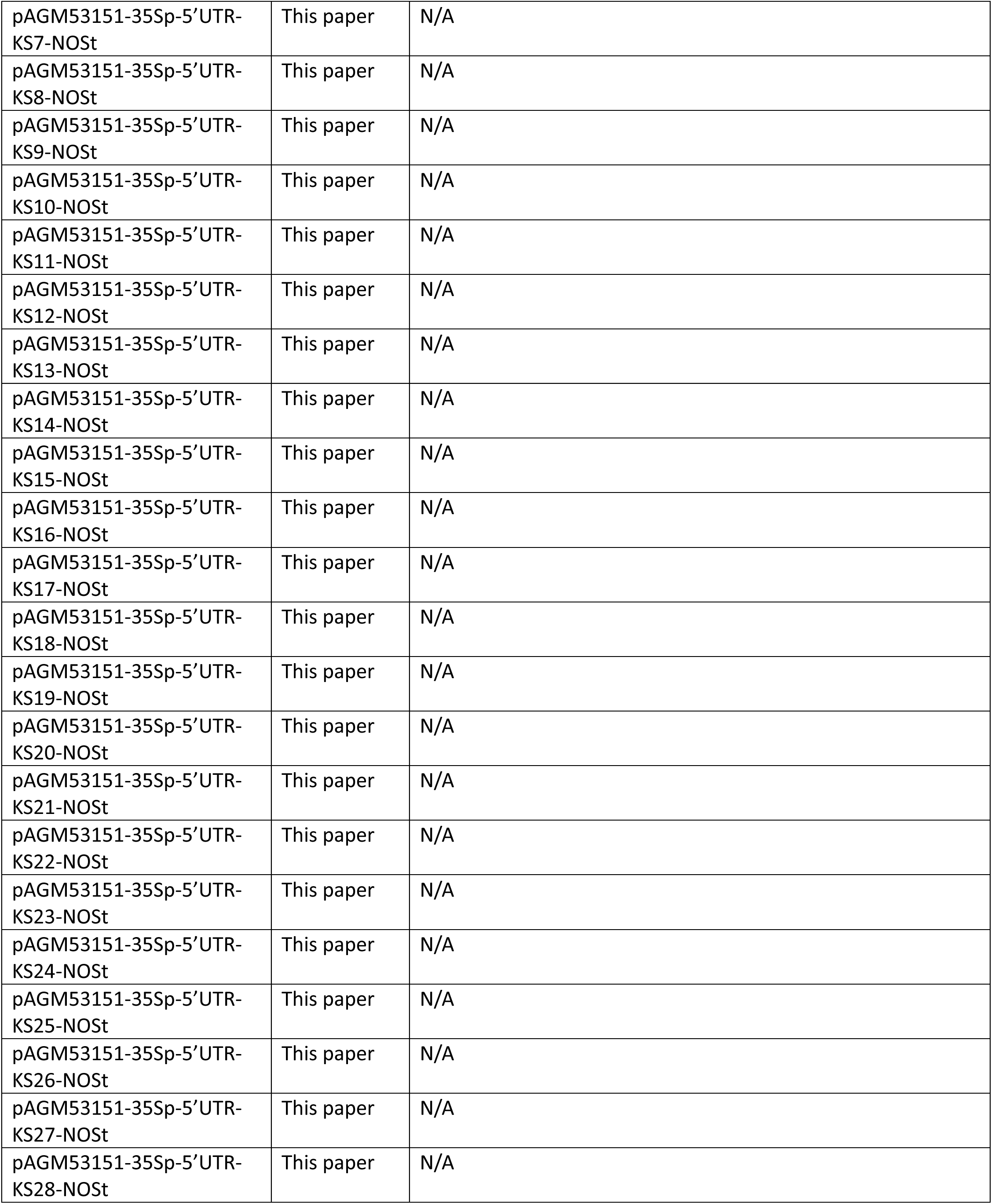
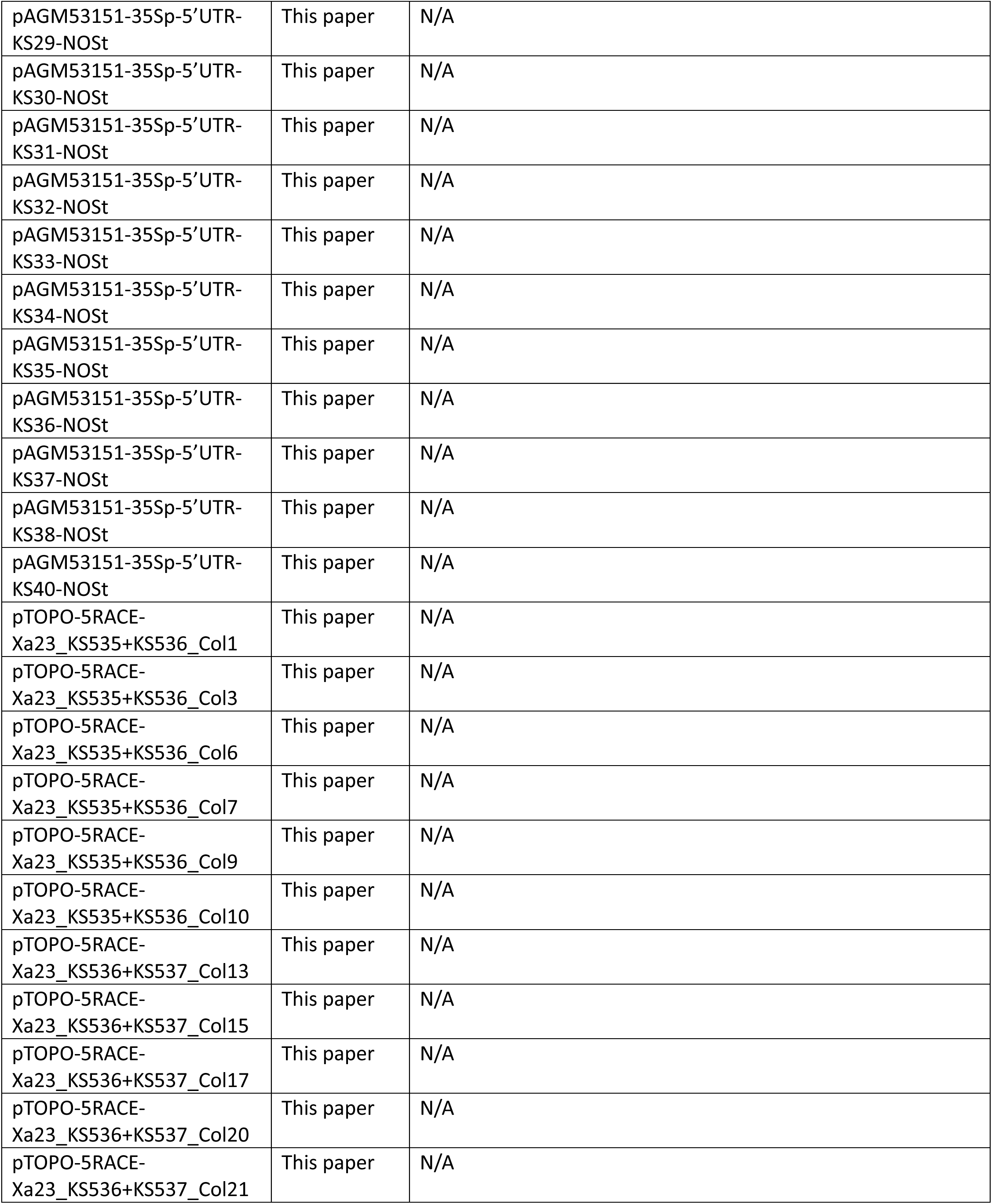
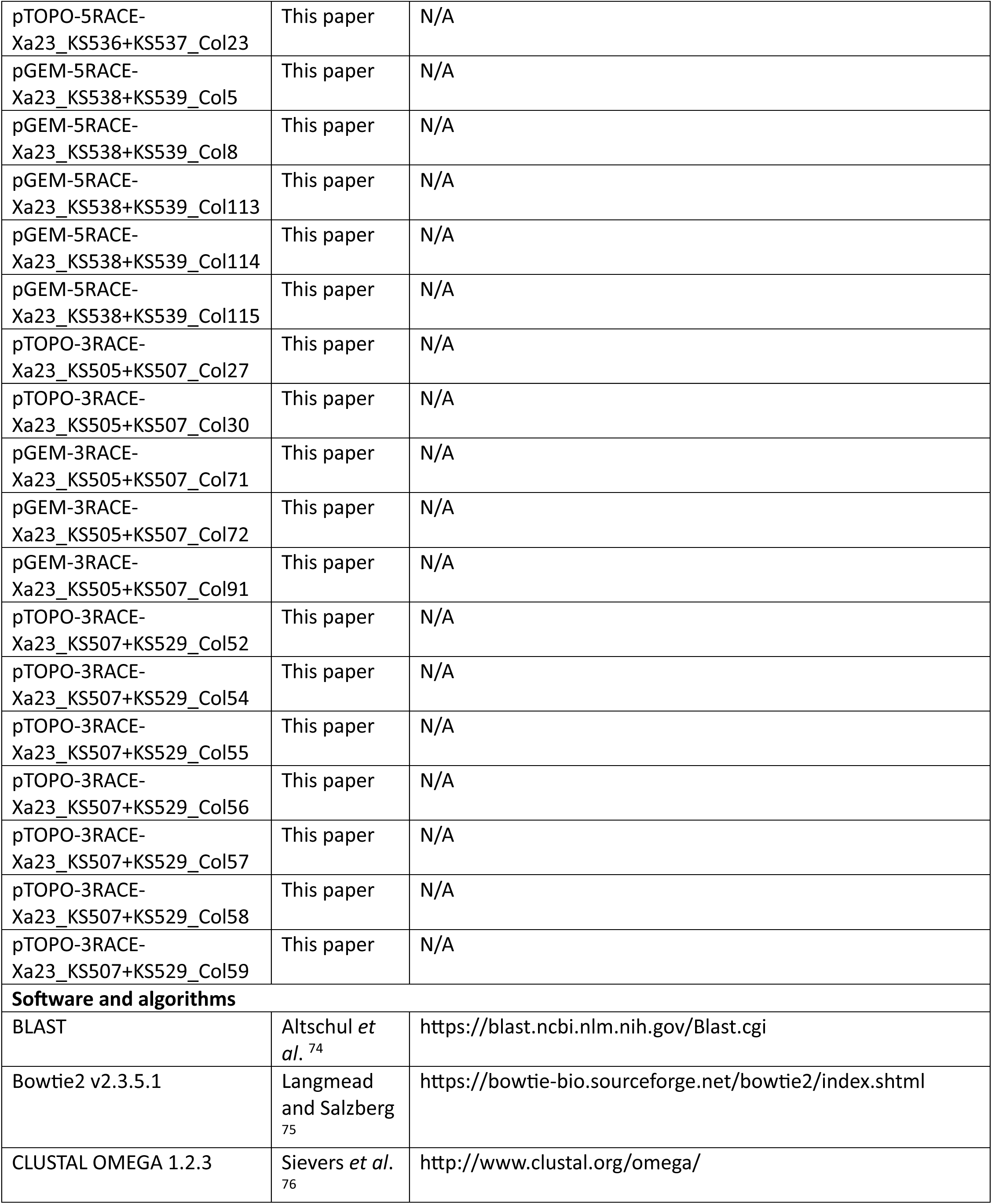
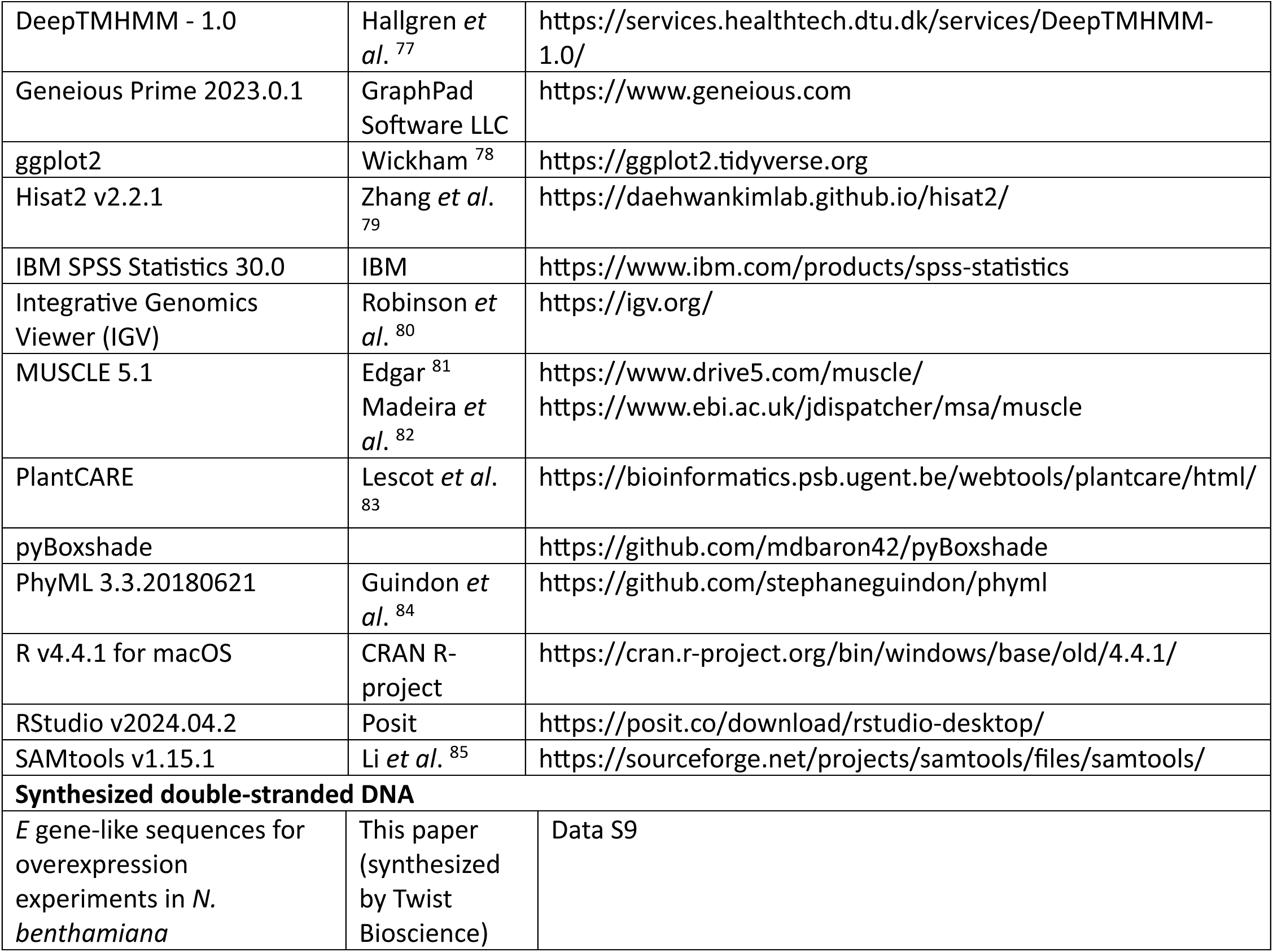

| REAGENT or RESOURCE | SOURCE | IDENTIFIER |
| --- | --- | --- |
| **Bacterial strains** |  |  |
| *Agrobacterium tumefaciens* GV3101 | Widely distributed | N/A |
| *Escherichia coli* TOP10 | Widely distributed | N/A |
| *Escherichia coli* DH5α | Toyobo | Cat#: F1346K |
| **Chemicals** |  |  |
| Bacto™ beef extract | Gibco | Cat#: 211520 |
| Peptone from casein | Merck Millipore | Cat#: 82303-1KG-F |
| Crystallized sucrose | Duchefa | Cat#: S0809.5000 |
| M-Gredient® Yeast Extract B1 | Merck Millipore | Cat#: 70161-2.5KG |
| Magnesium sulfate heptahydrate (MgSO<sub>4</sub> · 7H<sub>2</sub>O) | Carl Roth GmbH & Co. KG | Cat#: 8283.1 |
| Magnesium chlorid hexahydrate (MgCl<sub>2</sub> · 6H<sub>2</sub>O) | Carl Roth GmbH & Co. KG | Cat#: 2189.1 |
| MES monohydrate | Duchefa | Cat#: M1503.0250 |
| 3',5'-Dimethoxy-4'-hydroxyacetophenone (acetosyringone) | Sigma-Aldrich | Cat#: D134406 |
| D(-)-Mannitol | FUJIFILM Wako Pure Chemical Corporation | Cat#: 133-00845 |
| Absolute ethanol | FUJIFILM Wako Pure Chemical Corporation | Cat#: 057-00456 |
| Trichostatin A (TSA) | FUJIFILM Wako Pure Chemical Corporation | Cat#: 209-17563 |
| Dimethyl sulfoxide (DMSO) | FUJIFILM Wako Pure Chemical Corporation | Cat#: 043-29355 |
| CHU (N6) basal salt mixture | Sigma-Aldrich | Cat#: C1416 |
| Molybdic acid sodium salt dihydrate | Wako Pure Chemical Industries | Cat#: 514-30001 |
| Cobalt (II) chloride hexahydrate | Wako Pure Chemical Industries | Cat#: 036-03682 |
| Cupric sulfate pentahydrate | Wako Pure Chemical Industries | Cat#: 033-04415 |
| Retinol | Sigma-Aldrich | Cat#: R7632 |
| Calciferol | Wako Pure Chemical Industries | Cat#: 039-00291 |
| Biotin | Wako Pure Chemical Industries | Cat#: 023-08711 |
| Thiamin·HCl | FUJIFILM Wako Pure Chemical Corporation | Cat#: 201-00852 |
| Nicotinic acid | FUJIFILM Wako Pure Chemical Corporation | Cat#: 142-01232 |
| Pyridoxine·HCl | FUJIFILM Wako Pure Chemical Corporation | Cat#: 163-05402 |
| Choline chloride | Wako Pure Chemical Industries | Cat#: 033-09812 |
| Calcium pantothenate | Wako Pure Chemical Industries | Cat#: 031-14161 |
| Riboflavin | Wako Pure Chemical Industries | Cat#: 181-00581 |
| 2,4-Dichlorophenoxyacetic Acid (2,4-D) | Nacalai Tesque | Cat#: 11831-22 |
| Cyanocobalamin | Wako Pure Chemical Industries | Cat#: 224-00344 |
| 4-Aminobenzoic acid | Wako Pure Chemical Industries | Cat#: 015-02332 |
| Folic acid | FUJIFILM<br>Wako Pure<br>Chemical<br>Corporation | Cat#: 062-01801 |
| Ascorbic Acid | Wako Pure<br>Chemical<br>Industries | Cat#: 012-04802 |
| Malic acid | Wako Pure<br>Chemical<br>Industries | Cat#: 138-07512 |
| Citric acid | Wako Pure<br>Chemical<br>Industries | Cat#: 036-05522 |
| Fumaric acid | Wako Pure<br>Chemical<br>Industries | Cat#: 069-00652 |
| Sodium pyruvate | Wako Pure<br>Chemical<br>Industries | Cat#: 199-03062 |
| Glutamine | FUJIFILM<br>Wako Pure<br>Chemical<br>Corporation | Cat#: 072-00523 |
| Casein hydrolysate | BD | Cat#: 223120 |
| Myo-inositol | FUJIFILM<br>Wako Pure<br>Chemical<br>Corporation | Cat#: 092-00282 |
| Glucose | Wako Pure<br>Chemical<br>Industries | Cat#: 049-31165 |
| **Deposited data** |  |  |
| List of *Xa10-Ni* alleles from 3,002 rice genomes | This paper | Data S1 |
| List of *Xa23* alleles from 3,002 rice genomes | This paper | Data S2 |
| **Critical commercial products and assays** |  |  |
| BsaI-HF<sup>®</sup>v2 | NEB | Cat#: R3733S |
| T4 DNA Ligase, HC (30 U/μL) | Thermo Scientific | Cat#: EL0013 |
| Millicell<sup>®</sup> Standing Cell Culture Inserts | Millipore | Cat#: PICM01250 |
| 10x Lysis buffer | Takara Bio | Cat#: 635013 |
| SMART-Seq HT Kit | Takara Bio | Cat#: 634436 |
| AMPure XP Beads for DNA Cleanup | Beckman Coulter | Cat#: A63880 |
| 96-well plates for qRT-PCR | Roche | Cat#: 04729692001 |
| Roche Diagnostics FS ESSENTIAL DNA GREEN MASTER | Roche | Cat#: 06402712001 |
| SMARTer® RACE 5'/3' Kit | Takara Bio | Cat#: 634858 |
| KOD-FX DNA polymerase | Toyobo | Cat#: KFX-101 |
| PrimeSTAR GXL DNA Polymerase | Takara Bio | Cat#: R050A |
| Zero Blunt™ TOPO™ PCR Cloning Kit | Thermo Scientific | Cat#: 450245 |
| pGEM®-T Easy Vector Systems PCR Cloning Kit | Promega | Cat#: A137A |
| **Experimental models: Cell lines** |  |  |
| *Oryza sativa* Oc cell suspension culture | RIKEN Bio-Resource Center, Tsukuba, Japan | Cat#: rpc00031 |
| **Experimental models: Organisms/strains** |  |  |
| *Nicotiana benthamiana* | Widely distributed | N/A |
| *Oryza sativa* cv. Nipponbare | Widely distributed | N/A |
| **Oligonucleotides** |  |  |
| Primers for cloning, sequencing, qRT-PCRs, RACE, etc. | This paper | Table S2 |
| **Recombinant DNA** |  |  |
| pAGM53151-35Sp-5'UTR-LacZ-NOST | Grützner *et al.* <sup>73</sup> | Addgene #169093 |
| pAGM53151-35Sp-5'UTR-KS1-NOST | This paper | N/A |
| pAGM53151-35Sp-5'UTR-KS2-NOST | This paper | N/A |
| pAGM53151-35Sp-5'UTR-KS3-NOST | This paper | N/A |
| pAGM53151-35Sp-5'UTR-KS4-NOST | This paper | N/A |
| pAGM53151-35Sp-5'UTR-KS5-NOST | This paper | N/A |
| pAGM53151-35Sp-5'UTR-KS6-NOST | This paper | N/A |
| pAGM53151-35Sp-5'UTR-KS7-NOST | This paper | N/A |
| pAGM53151-35Sp-5'UTR-KS8-NOST | This paper | N/A |
| pAGM53151-35Sp-5'UTR-KS9-NOST | This paper | N/A |
| pAGM53151-35Sp-5'UTR-KS10-NOST | This paper | N/A |
| pAGM53151-35Sp-5'UTR-KS11-NOST | This paper | N/A |
| pAGM53151-35Sp-5'UTR-KS12-NOST | This paper | N/A |
| pAGM53151-35Sp-5'UTR-KS13-NOST | This paper | N/A |
| pAGM53151-35Sp-5'UTR-KS14-NOST | This paper | N/A |
| pAGM53151-35Sp-5'UTR-KS15-NOST | This paper | N/A |
| pAGM53151-35Sp-5'UTR-KS16-NOST | This paper | N/A |
| pAGM53151-35Sp-5'UTR-KS17-NOST | This paper | N/A |
| pAGM53151-35Sp-5'UTR-KS18-NOST | This paper | N/A |
| pAGM53151-35Sp-5'UTR-KS19-NOST | This paper | N/A |
| pAGM53151-35Sp-5'UTR-KS20-NOST | This paper | N/A |
| pAGM53151-35Sp-5'UTR-KS21-NOST | This paper | N/A |
| pAGM53151-35Sp-5'UTR-KS22-NOST | This paper | N/A |
| pAGM53151-35Sp-5'UTR-KS23-NOST | This paper | N/A |
| pAGM53151-35Sp-5'UTR-KS24-NOST | This paper | N/A |
| pAGM53151-35Sp-5'UTR-KS25-NOST | This paper | N/A |
| pAGM53151-35Sp-5'UTR-KS26-NOST | This paper | N/A |
| pAGM53151-35Sp-5'UTR-KS27-NOST | This paper | N/A |
| pAGM53151-35Sp-5'UTR-KS28-NOST | This paper | N/A |
| pAGM53151-35Sp-5'UTR-KS29-NOST | This paper | N/A |
| pAGM53151-35Sp-5'UTR-KS30-NOST | This paper | N/A |
| pAGM53151-35Sp-5'UTR-KS31-NOST | This paper | N/A |
| pAGM53151-35Sp-5'UTR-KS32-NOST | This paper | N/A |
| pAGM53151-35Sp-5'UTR-KS33-NOST | This paper | N/A |
| pAGM53151-35Sp-5'UTR-KS34-NOST | This paper | N/A |
| pAGM53151-35Sp-5'UTR-KS35-NOST | This paper | N/A |
| pAGM53151-35Sp-5'UTR-KS36-NOST | This paper | N/A |
| pAGM53151-35Sp-5'UTR-KS37-NOST | This paper | N/A |
| pAGM53151-35Sp-5'UTR-KS38-NOST | This paper | N/A |
| pAGM53151-35Sp-5'UTR-KS40-NOST | This paper | N/A |
| pTOPO-5RACE-Xa23_KS535+KS536_Col1 | This paper | N/A |
| pTOPO-5RACE-Xa23_KS535+KS536_Col3 | This paper | N/A |
| pTOPO-5RACE-Xa23_KS535+KS536_Col6 | This paper | N/A |
| pTOPO-5RACE-Xa23_KS535+KS536_Col7 | This paper | N/A |
| pTOPO-5RACE-Xa23_KS535+KS536_Col9 | This paper | N/A |
| pTOPO-5RACE-Xa23_KS535+KS536_Col10 | This paper | N/A |
| pTOPO-5RACE-Xa23_KS536+KS537_Col13 | This paper | N/A |
| pTOPO-5RACE-Xa23_KS536+KS537_Col15 | This paper | N/A |
| pTOPO-5RACE-Xa23_KS536+KS537_Col17 | This paper | N/A |
| pTOPO-5RACE-Xa23_KS536+KS537_Col20 | This paper | N/A |
| pTOPO-5RACE-Xa23_KS536+KS537_Col21 | This paper | N/A |
| pTOPO-5RACE-Xa23_KS536+KS537_Col23 | This paper | N/A |
| pGEM-5RACE-Xa23_KS538+KS539_Col5 | This paper | N/A |
| pGEM-5RACE-Xa23_KS538+KS539_Col8 | This paper | N/A |
| pGEM-5RACE-Xa23_KS538+KS539_Col113 | This paper | N/A |
| pGEM-5RACE-Xa23_KS538+KS539_Col114 | This paper | N/A |
| pGEM-5RACE-Xa23_KS538+KS539_Col115 | This paper | N/A |
| pTOPO-3RACE-Xa23_KS505+KS507_Col27 | This paper | N/A |
| pTOPO-3RACE-Xa23_KS505+KS507_Col30 | This paper | N/A |
| pGEM-3RACE-Xa23_KS505+KS507_Col71 | This paper | N/A |
| pGEM-3RACE-Xa23_KS505+KS507_Col72 | This paper | N/A |
| pGEM-3RACE-Xa23_KS505+KS507_Col91 | This paper | N/A |
| pTOPO-3RACE-Xa23_KS507+KS529_Col52 | This paper | N/A |
| pTOPO-3RACE-Xa23_KS507+KS529_Col54 | This paper | N/A |
| pTOPO-3RACE-Xa23_KS507+KS529_Col55 | This paper | N/A |
| pTOPO-3RACE-Xa23_KS507+KS529_Col56 | This paper | N/A |
| pTOPO-3RACE-Xa23_KS507+KS529_Col57 | This paper | N/A |
| pTOPO-3RACE-Xa23_KS507+KS529_Col58 | This paper | N/A |
| pTOPO-3RACE-Xa23_KS507+KS529_Col59 | This paper | N/A |
| **Software and algorithms** |  |  |
| BLAST | Altschul *et al.*<sup>74</sup> | https://blast.ncbi.nlm.nih.gov/Blast.cgi |
| Bowtie2 v2.3.5.1 | Langmead and Salzberg<sup>75</sup> | https://bowtie-bio.sourceforge.net/bowtie2/index.shtml |
| CLUSTAL OMEGA 1.2.3 | Sievers *et al.*<sup>76</sup> | http://www.clustal.org/omega/ |
| DeepTMHMM - 1.0 | Hallgren *et al.* <sup>77</sup> | https://services.healthtech.dtu.dk/services/DeepTMHMM-1.0/ |
| Geneious Prime 2023.0.1 | GraphPad Software LLC | https://www.geneious.com |
| ggplot2 | Wickham <sup>78</sup> | https://ggplot2.tidyverse.org |
| Hisat2 v2.2.1 | Zhang *et al.* <sup>79</sup> | https://daehwankimlab.github.io/hisat2/ |
| IBM SPSS Statistics 30.0 | IBM | https://www.ibm.com/products/spss-statistics |
| Integrative Genomics Viewer (IGV) | Robinson *et al.* <sup>80</sup> | https://igv.org/ |
| MUSCLE 5.1 | Edgar <sup>81</sup><br>Madeira *et al.* <sup>82</sup> | https://www.drive5.com/muscle/<br>https://www.ebi.ac.uk/jdispatcher/msa/muscle |
| PlantCARE | Lescot *et al.* <sup>83</sup> | https://bioinformatics.psb.ugent.be/webtools/plantcare/html/ |
| pyBoxshade |  | https://github.com/mdbaron42/pyBoxshade |
| PhyML 3.3.20180621 | Guindon *et al.* <sup>84</sup> | https://github.com/stephaneguindon/phyml |
| R v4.4.1 for macOS | CRAN R-project | https://cran.r-project.org/bin/windows/base/old/4.4.1/ |
| RStudio v2024.04.2 | Posit | https://posit.co/download/rstudio-desktop/ |
| SAMtools v1.15.1 | Li *et al.* <sup>85</sup> | https://sourceforge.net/projects/samtools/files/samtools/ |
| **Synthesized double-stranded DNA** |  |  |
| *E* gene-like sequences for overexpression experiments in *N. benthamiana* | This paper (synthesized by Twist Bioscience) | Data S9 |

### Experimental model and study participant details Plant materials and growth conditions

*Oryza sativa* cv. Nipponbare plants were grown in an environmental chamber (K30-7248; Koito Industries, Yokohama, Japan) at 26°C under a 13-hour light/11-hour dark photoperiod. The isolation of egg cells and sperm cells from rice flowers, and the electrofusion of gametes were conducted as previously described.^86–88^

*Nicotiana benthamiana* plants were grown at 20-24°C, 35-60% humidity, and a 16-hour light/8-hour dark photoperiod. Five-to six-week-old plants were used for transient overexpression of *E* genes.

### Method details

#### Analysis of *Xa10-Ni* and *Xa23* alleles from 3,002 cultivated rice accessions

*Xa10* was excluded from the analysis because *Xa10* is not present in the reference genome (*O. sativa* cv. Nipponbare) to which the genomes of 3,002 cultivated rice accessions are aligned.^26,27^ For *Xa10-Ni* and *Xa23* alleles, genomic sequences corresponding to the chr11_22179001:22182000 (*Xa10-Ni*) and chr11:22,202,401-22,205,400 (*Xa23*) of the reference genome were extracted from the IRRI SNP-Seek Database (http://snp-seek.irri.org).^27^ Start and stop codons of *Xa10-Ni* and *Xa23* alleles from 3,002 cultivated rice accessions were identified based on the genomic structures of *Xa10-Ni* and *Xa23* from Nipponbare. The percentage identity matrix was generated using the CLUSTAL OMEGA algorithm^76^ in Geneious Prime 2023.0.1. The lists of XA10 and XA23 protein sequences from 3,002 cultivated rice accessions are included as Data S1 and S2.

### Mining for *E* gene-like sequences in genomes of wild rice and cutgrass species

Genomic sequences of *Xa10* from *O. sativa* cv. IRBB10A (NCBI GenBank ID: JX025645.1), *Xa10-Ni* from *O. sativa* cv. Nipponbare (RAP-DB: Os11g0586400; MSU: LOC_Os11g37570), and *Xa23* from *O. sativa* cv. Nipponbare (RAP-DB: Os11g0586701; MSU: LOC_Os11g37620) were used as queries to align NCBI database sequences (“Whole-Genome Shotgun Contigs” and “RefSeq Genome Database”) using the Basic Local Alignment Search Tool (BLAST).^74^ The search was limited to genomes of *Oryza* and *Leersia* species (“Organisms”) available in the aforementioned databases. The search was optimized for “More dissimilar sequences (discontiguous megablast)”. All sequences with the query cover above 30% and identity above 30% were extracted and aligned to the *Xa10*, *Xa10-Ni*, and *Xa23* reference sequences in Geneious Prime 2023.0.1 using the MUSCLE algorithm (Table S1).^81^ Putative CDSs within *E* gene-like sequences were determined by mimicking gene structures of *Xa10*, *Xa10-Ni*, and *Xa23*: a single exon from the nearest start codon to the nearest stop codon. Putative CDSs containing premature stop codons and encoding proteins with < 40% identity to XA10, XA10-Ni, and XA23 were considered pseudogenes and excluded from further analysis. Only the putative open reading frames (ORFs) potentially encoding full-length proteins were selected for further alignments (Data S3-S5). Putative TMMs within the protein sequences were predicted using DeepTMHMM v1.0.^77^ A percentage identity matrix for putative full-length protein sequences was generated in Geneious Prime 2023.0.1 using the CLUSTAL OMEGA algorithm (Table S1).^76^ Maximum-likelihood phylogenetic analysis of E protein sequences from the genomes of sequenced *Oryza* and *Leersia* species was performed in Geneious Prime 2023.0.1 using the PhyML 3.3.20180621 method (Figure 2).^84^ The maximum-likelihood phylogenetic analysis is based on the Jones-Taylor-Thornton substitution model and 1000 bootstrap replications (*B* = 1000). XA7 and XA27, two rice E proteins, were used as outgroups for the phylogenetic analysis.^9,12^

### Assembly of constructs

Due to their short lengths, CDSs of *E* gene homologs from *Oryza* and *Leersia* species were synthesized with *Bsa*I overhangs for GoldenGate cloning as gene fragments using commercial services from Twist Bioscience (Data S9). To ensure the presence of *Bsa*I overhangs in the synthesized gene fragments, 5’-TTTGGTCTCAA-3’ was added in front of the start codon and 5’-GCTTTGAGACCAAA-3’ was added after the expected stop codon of each CDS. The *GFP* containing an artificial intron (*GFPi*) was synthesized the same way as CDSs of putative *E* genes. Synthesized fragments were subcloned into the *Bsa*I-compatible pAGM53151-35Sp-5’UTR-NOSt vector via GoldenGate cloning reaction (Addgene #169093).^73^ To verify correct cloning assembly, plasmids were analyzed via Sanger sequencing. Plasmids with the correct sequence fragments were individually transformed into *Agrobacterium tumefaciens* GV3101 strain.

### Overexpression of *E* genes in *N. benthamiana*

*A. tumefaciens* strains were grown in YEB medium containing 0.5% (w/v) Bacto™ beef extract (Gibco, Catalog #211520), 0.5% (w/v) peptone from casein (Merck Millipore, Catalog #82303-1KG-F), 0.5% (w/v) crystallized sucrose (Duchefa, Catalog #S0809.5000), 0.1% (w/v) M-Gredient® Yeast Extract B1 (Merck Millipore, Catalog #70161-2.5KG), 2 mM MgSO₄ ⋅ 7H₂O (Carl Roth GmbH & Co. KG, Catalog #8283.1) with the respective antibiotics. Cells were harvested and resuspended in the *Agrobacterium* infiltration medium (AIM; pH 5.3) containing 10 mM MgCl₂ ⋅ 6H₂O (Carl Roth GmbH & Co. KG, Catalog #2189.1), 5 mM MES monohydrate (Duchefa, Catalog #M1503.0250), and 150 μM acetosyringone (Sigma-Aldrich, Catalog #D134406). The inocula (OD_600nm_ = 0.8) were infiltrated into the leaves of 5-6-week-old *N. benthamiana* plants using a needleless syringe. Leaves were harvested, and photos were taken three days post-infiltration (3 dpi).

### *In vitro* gamete fusion and zygote production

Extraction, isolation, fusion of rice gametes, and culturing of IVF-produced zygotes were performed as previously described.^88^ Individual unfertilized egg cells were treated with an electric impulse AC 5V / DC 15V to imitate the fusion process and kept in 370 mosmol/kg H_2_O mannitol solution for six hours. Rice sperm cells were incubated in 370 mosmol/kg H_2_O mannitol solution for a maximum of 15 minutes, since longer incubation leads to rice sperm burst.^88^ Production of zygotes was performed via electrofusion of one rice egg cell with one rice sperm cell using standard settings (AC 5V / DC 15V). IVF-produced zygotes were cultured in 450 mosmol/kg H_2_O mannitol solution without feeder cells for harvesting at 15 minutes AF and 6 hours AF. IVF-produced zygotes were cultured in N6Z liquid medium with feeder cells (RIKEN Bio-Resource Center, Tsukuba, Japan; Catalog #: rpc00031) for later harvest (12 hours AF).

### HDACi treatment of unfertilized egg cells

Extracted unfertilized egg cells were cultured in N6Z liquid medium supplemented with either 10 μM TSA (FUJIFILM Wako Pure Chemical Corporation; Catalog #: 209-17563) or DMSO (FUJIFILM Wako Pure Chemical Corporation; Catalog #: 043-29355) for 24 hours. N6Z medium contained feeder cells (RIKEN Bio-Resource Center, Tsukuba, Japan; Catalog #: rpc00031).^88^ **Total RNA isolation and cDNA synthesis**

Three egg cells/zygotes per sample were harvested for total RNA extraction. For sperm cells, 10 cells were harvested per sample. Harvested cells were lysed in 10x Lysis buffer (TAKARA; Catalog #635013). Total RNA was converted to complementary DNA (cDNA) using the SMART-Seq HT Kit (TAKARA; Catalog #634436) and the producer-recommended standard protocol. 18 PCR cycles were used to amplify cDNA. The resulting cDNA was purified using 1.8x concentration AMPure XP Beads (Beckman Coulter; Catalog #A63880) and the standard protocol recommended by the manufacturer.

### Quantitative real-time PCR

qRT-PCR was performed in 10 μl reactions using 96-well qRT-PCR Plates (Roche; Catalog #04729692001) and LightCycler® 96 (Roche; Instrument Serial Number: 000000000014640). Each qRT-PCR reaction contained 1 pmol of forward and 1 pmol of reverse primers (Table S2), 7 ng of synthesized and purified cDNA, 5 μl of Roche Diagnostics FastStart Essential DNA Green Master (Roche; Catalog #06402712001), and Milli-Q water. The same program was used for all qRT-PCRs: preincubation at 95°C for 10 min; 3 step amplification for 40 cycles of 95°C for 10 sec, 60°C for 10 sec, and 72°C for 10 sec; and high-resolution melting with continuous acquisition at 95°C for 60 sec, 40°C for 60 sec, 65°C for 1 sec, and 97°C for 1 sec. Each sample had two technical replicates. The mean Ct values for five biological replicates and a minimum of five different cDNA and gDNA samples of various dilutions (1:4, 1:16, 1:64, and 1:256) were used to determine primer efficiency. *Xa10-Ni* (RAP-DB: Os11g0586400; MSU: LOC_Os11g37570) was used as a negative control. *RING137* (RAP-DB: Os08g0384900; MSU: LOC_Os08g29590), a housekeeping gene (HKG) encoding an E3 ligase, was used for normalization.

### Quantification and characterization of transcripts via bioinformatic approaches

The pre-processed RNA-seq reads from the datasets published by Rahman *et al.* ^22^ (Runs #DRR166403 – DRR166403) and Toda *et al.* ^23^ (Runs #DRR398287 – DRR398291) were mapped to the *Xa23* locus (RAP-DB: Os11g0586701; MSU: LOC_Os11g37620) from the Nipponbare reference genome using Bowtie2 v2.3.5.1.^75^ The mapping data were converted and sorted using Samtools v1.15.1^93^ for visualization in IGV (Figure S3).^80^ Full-length sequences of the reconstructed putative native *Xa23* transcripts are available in Data S11. Transcripts per million (TPM) values were used to quantify abundance of *Xa23*, *Xa10-Ni*, *RING137*, *RISBZ1*/*bZIP58*, and *RISBZ5*/*bZIP52* in RNA-seq samples.

### Characterization of the TSA-induced *Xa23* transcripts via 5’ and 3’ RACE

Total RNA was harvested from 7 unfertilized egg cells treated with DMSO (solvent) for 24 hours and from 7 unfertilized egg cells treated with 10 μM TSA for 24 hours via cell lysis in 2 μl of 10x Lysis buffer (Takara Bio; Catalog #635013). SMARTer® RACE 5’/3’ Kit (Takara Bio; Catalog #634858) was used for synthesis of first-strand cDNA for 5’ and 3’ RACE according to the standard protocol. To improve specificity, nested PCRs were performed. The first PCR was performed using KOD-FX DNA polymerase (Toyobo; Catalog #KFX-101), 1.5 µL of 1:10 diluted RACE-ready cDNA as a template, and target gene-specific primers for *Xa23* (RAP-DB: Os11g0586701; MSU: LOC_Os11g37620) and *RING137* (RAP-DB: Os08g0384900; MSU: LOC_Os08g29590), used as a positive control. The following PCR program was used for the first PCR: 98°C for 5 min; 13 cycles of 98°C for 30 sec, 68°C (-1°C per cycle) for 30 sec, and 68°C for 30 sec; 17 cycles of 98°C for 30 sec, 55°C for 30 sec, and 68°C for 30 sec. The second PCR was performed using PrimeSTAR GXL DNA Polymerase (Takara Bio; Catalog # R050A), 1.5 µL of 1:5000 diluted product of the first PCR as a template, and target gene-specific primers for *Xa23* and *RING137* used as a positive control. The following PCR program was used for the second PCR: 98°C for 5 min; 35 cycles of 98°C for 20 sec, 60°C for 20 sec, and 68°C for 60 sec; 68°C for 2 min. In the case of 5’ *RING137* PCR, the annealing temperature was reduced to 55°C. *RING137-*specific amplicons from the 5’ and 3’ RACE PCRs were analyzed via Sanger sequencing. *Xa23-*specific amplicons from the 5’ and 3’ RACE PCRs were cloned using Zero Blunt TOPO PCR Cloning Kit (Thermo Scientific Invitrogen; Catalog #450245) and pGEM-T Easy Vector Systems PCR Cloning Kit (Promega; Catalog # A137A), and several colonies from each cloning were analyzed via Sanger sequencing. The list of all oligonucleotides used in this study is available in Table S2.

### *In silico* analysis of putative CREs within the *Xa23* upstream sequences

Putative CREs within the *Xa23* upstream sequences were predicted using PlantCARE.^83^ 350 nt regions upstream of the *Xa23* start codon from the rice genotypes Nipponbare and CBB23 were used for *in silico* analysis. Only putative CREs that are common to both genotypes are annotated (Figure 6B and Data S12).

### Statistical analysis

Boxplots were constructed in RStudio v2024.04.2 using the ggplot2 package^78^ for R v4.4.1. Statistical analyses were performed in IBM SPSS Statistics 30.0. Independent-Samples Mann-Whitney U Test and Independent-Samples Kruskal-Wallis Test, followed by the Bonferroni *p*-value adjustment for multiple comparisons, were used to assess significance levels.

## Supplemental information

**Document S1. Figures S1–S4, Tables S1and S2, and Data S1-S13**

## Supplemental Figures

**Figure S1**. Native *Xa23* transcripts are detectable in IVF-produced zygotes.

**Figure S2**. *Xa23* expression in IVF-produced zygotes peaks at six hours after fertilization.

**Figure S3**. RNA-seq reads from the published datasets map to the *Xa23* locus.

**Figure S4**. *RISBZ1* and *RISBZ5* transcripts are detectable in IVF-produced zygotes.

## Supplemental Tables

**Table S1.** Percentage of residues that are identical between XA10, XA10-NI, and XA23 proteins and the homologs from wild rice and cutgrass species.

**Table S2.** List of primers used in this study.

## Supplemental Data

**Data S1.** XA10-Ni protein variants from 3,002 cultivated rice accessions.

**Data S2**. XA23 protein variants from 3,002 cultivated rice accessions.

**Data S3**. Alignment of the XA10 protein from the cultivated rice accession IRBB10A and XA10 homologs from wild rice and cutgrass species.

**Data S4**. Alignment of the XA10-Ni protein from the cultivated rice accession Nipponbare and XA10-Ni homologs from wild rice species.

**Data S5**. Alignment of the XA23 protein from the cultivated rice accession Nipponbare and XA23 homologs from wild rice species.

**Data S6**. Alignment of genomic sequences upstream of the *Xa10*-like loci from cultivated rice, wild rice, and cutgrass species.

**Data S7**. Alignment of genomic sequences upstream of the *Xa10-Ni*-like loci from cultivated and wild rice species.

**Data S8**. Alignment of genomic sequences upstream of the *Xa23*-like loci from cultivated and wild rice species.

**Data S9**. Synthesized *E* gene homologs from wild rice and cutgrass species for transient overexpression in *N. benthamiana*.

**Data S10**. Publicly available RNA-seq datasets from the RAP-DB database screened for RNA-seq reads mapping to *O. sativa Xa10-Ni* and *Xa23* loci.

**Data S11**. Alignment of the genomic *Xa23* sequence and putative full-length native *Xa23* transcripts reconstructed *in silico* using the published RNA-Seq datasets.

**Data S12**. Alignment of the genomic *Xa23* sequences from two cultivated rice accessions (Nipponbare and CBB23) and 5’RACE-derived parts of the TSA-induced *Xa23* transcripts.

**Data S13**. Alignment of the genomic *Xa23* sequences from two cultivated rice accessions (Nipponbare and CBB23) and 3’RACE-derived parts of the TSA-induced *Xa23* transcripts.

## References

1. Yang, B., and White, F.F. (2004). Diverse members of the AvrBs3/PthA family of type III effectors are major virulence determinants in bacterial blight disease of rice. Mol. Plant-Microbe Interact. 17, 1192–1200. 10.1094/MPMI.2004.17.11.1192.

2. Boch, J., Scholze, H., Schornack, S., Landgraf, A., Hahn, S., Kay, S., Lahaye, T., Nickstadt, A., and Bonas, U. (2009). Breaking the code of DNA binding specificity of TAL-type III effectors. Science (New York, N.Y.) 326, 1509–1512. 10.1126/science.1178811.

3. Yuan, M., Ke, Y., Huang, R., Ma, L., Yang, Z., Chu, Z., Xiao, J., Li, X., and Wang, S. (2016). A host basal transcription factor is a key component for infection of rice by TALE-carrying bacteria. eLife 5, e19605. 10.7554/eLife.19605.

4. Robinson, P.J., Trnka, M.J., Bushnell, D.A., Davis, R.E., Mattei, P.-J., Burlingame, A.L., and Kornberg, R.D. (2016). Structure of a complete mediator-RNA polymerase II pre-initiation complex. Cell 166, 1411–1422.e1416. 10.1016/j.cell.2016.08.050.

5. Kay, S., Hahn, S., Marois, E., Hause, G., and Bonas, U. (2007). A bacterial effector acts as a plant transcription factor and induces a cell size regulator. Science (New York, N.Y.) 318, 648–651. 10.1126/science.1144956.

6. Yang, B., Sugio, A., and White, F.F. (2006). *Os8N3* is a host disease-susceptibility gene for bacterial blight of rice. Proceedings of the National Academy of Sciences 103, 10503–10508. 10.1073/pnas.0604088103.

7. Zhou, J., Peng, Z., Long, J., Sosso, D., Liu, B., Eom, J.-S., Huang, S., Liu, S., Vera Cruz, C., Frommer, W.B., et al. (2015). Gene targeting by the TAL effector PthXo2 reveals cryptic resistance gene for bacterial blight of rice. TPJ 82, 632–643. 10.1111/tpj.12838.

8. Streubel, J., Pesce, C., Hutin, M., Koebnik, R., Boch, J., and Szurek, B. (2013). Five phylogenetically close rice *SWEET* genes confer TAL effector-mediated susceptibility to *Xanthomonas oryzae* pv. *oryzae*. New Phytol. 200, 808–819. 10.1111/nph.12411.

9. Chen, X., Liu, P., Mei, L., He, X., Chen, L., Liu, H., Shen, S., Ji, Z., Zheng, X., Zhang, Y., et al. (2021). *Xa7*, a new executor *R* gene that confers durable and broad-spectrum resistance to bacterial blight disease in rice. Plant Commun. 2, 100143. 10.1016/j.xplc.2021.100143.

10. Tian, D., Wang, J., Zeng, X., Gu, K., Qiu, C., Yang, X., Zhou, Z., Goh, M., Luo, Y., Murata-Hori, M., et al. (2014). The rice TAL effector–dependent resistance protein XA10 triggers cell death and calcium depletion in the endoplasmic reticulum. The Plant Cell 26, 497–515. 10.1105/tpc.113.119255.

11. Wang, C., Zhang, X., Fan, Y., Gao, Y., Zhu, Q., Zheng, C., Qin, T., Li, Y., Che, J., Zhang, M., et al. (2015). XA23 is an executor R protein and confers broad-spectrum disease resistance in rice. Molecular Plant 8, 290–302. 10.1016/j.molp.2014.10.010.

12. Gu, K., Yang, B., Tian, D., Wu, L., Wang, D., Sreekala, C., Yang, F., Chu, Z., Wang, G.-L., White, F.F., and Yin, Z. (2005). *R* gene expression induced by a type-III effector triggers disease resistance in rice. Nature 435, 1122–1125. 10.1038/nature03630.

13. Römer, P., Hahn, S., Jordan, T., Strauß, T., Bonas, U., and Lahaye, T. (2007). Plant pathogen recognition mediated by promoter activation of the pepper *Bs3* resistance gene. Science (New York, N.Y.) 318, 645–648. 10.1126/science.1144958.

14. Strauß, T., van Poecke, R.M.P., Strauß, A., Römer, P., Minsavage, G.V., Singh, S., Wolf, C., Strauß, A., Kim, S., Lee, H.-A., et al. (2012). RNA-seq pinpoints a *Xanthomonas* TAL-effector activated resistance gene in a large-crop genome. Proceedings of the National Academy of Sciences 109, 19480–19485. 10.1073/pnas.121241510.

15. Roeschlin, R.A., Uviedo, F., García, L., Molina, M.C., Favaro, M.A., Chiesa, M.A., Tasselli, S., Franco-Zorrilla, J.M., Forment, J., Gadea, J., and Marano, M.R. (2019). PthA4AT, a 7.5-repeats transcription activator-like (TAL) effector from *Xanthomonas citri* ssp. *citri*, triggers citrus canker resistance. Mol. Plant Pathol. 20, 1394–1407. 10.1111/mpp.12844.

16. Bogdanove, A.J., Schornack, S., and Lahaye, T. (2010). TAL effectors: finding plant genes for disease and defense. Curr. Opin. Plant Biol. 13, 394–401. 10.1016/j.pbi.2010.04.010.

17. Yoshimura, A., Mew, T., Khush, G., and Omura, T. (1983). Inheritance of resistance to bacterial blight in rice cultivar Cas 209. Genetics 73, 1409–1412. 10.1094/Phyto-73-1409.

18. Wang, C.-L., Qin, T.-F., Yu, H.-M., Zhang, X.-P., Che, J.-Y., Gao, Y., Zheng, C.-K., Yang, B., and Zhao, K.-J. (2014). The broad bacterial blight resistance of rice line CBB23 is triggered by a novel transcription activator-like (TAL) effector of *Xanthomonas oryzae* pv. *oryzae*. Mol. Plant Pathol. 15, 333–341. 10.1111/mpp.12092.

19. Wang, J., Tian, D., Gu, K., Yang, X., Wang, L., Zeng, X., and Yin, Z. (2017). Induction of *Xa10*-like genes in rice cultivar Nipponbare confers disease resistance to rice bacterial blight. Mol. Plant-Microbe Interact. 30, 466–477. 10.1094/MPMI-11-16-0229-R.

20. Nowack, M.K., Holmes, D.R., and Lahaye, T. (2022). TALE-induced cell death executors: an origin outside immunity? Trends in Plant Science. 10.1016/j.tplants.2021.11.003.

21. Zhang, J., Yin, Z., and White, F. (2015). TAL effectors and the executor *R* genes. Front. Plant. Sci. 6. 10.3389/fpls.2015.00641.

22. Rahman, M.H., Toda, E., Kobayashi, M., Kudo, T., Koshimizu, S., Takahara, M., Iwami, M., Watanabe, Y., Sekimoto, H., Yano, K., and Okamoto, T. (2019). Expression of genes from paternal alleles in rice zygotes and involvement of *OsASGR-BBML1* in initiation of zygotic development. Plant and Cell Physiology 60, 725–737. 10.1093/pcp/pcz030.

23. Toda, E., Koshimizu, S., Kinoshita, A., Higashiyama, T., Izawa, T., Yano, K., and Okamoto, T. (2025). Transcriptional dynamics during karyogamy in rice zygotes. Development. 10.1242/dev.204497.

24. Gladieux, P., van Oosterhout, C., Fairhead, S., Jouet, A., Ortiz, D., Ravel, S., Shrestha, R.-K., Frouin, J., He, X., Zhu, Y., et al. (2024). Extensive immune receptor repertoire diversity in disease-resistant rice landraces. Curr. Biol. 34, 3983–3995.e3986. 10.1016/j.cub.2024.07.061.

25. Fan, C., Walling, J.G., Zhang, J., Hirsch, C.D., Jiang, J., and Wing, R.A. (2011). Conservation and purifying selection of transcribed genes located in a rice centromere. The Plant Cell 23, 2821–2830. 10.1105/tpc.111.085605.

26. Wang, W., Mauleon, R., Hu, Z., Chebotarov, D., Tai, S., Wu, Z., Li, M., Zheng, T., Fuentes, R.R., Zhang, F., et al. (2018). Genomic variation in 3,010 diverse accessions of Asian cultivated rice. Nature 557, 43–49. 10.1038/s41586-018-0063-9.

27. Mansueto, L., Fuentes, R.R., Borja, F.N., Detras, J., Abriol-Santos, J.M., Chebotarov, D., Sanciangco, M., Palis, K., Copetti, D., Poliakov, A., et al. (2017). Rice SNP-seek database update: new SNPs, indels, and queries. Nucleic Acids Res. 45, D1075–D1081. 10.1093/nar/gkw1135.

28. Kellogg, E.A. (2009). The evolutionary history of Ehrhartoideae, Oryzeae, and Oryza. Rice 2, 1–14. 10.1007/s12284-009-9022-2.

29. Stein, J.C., Yu, Y., Copetti, D., Zwickl, D.J., Zhang, L., Zhang, C., Chougule, K., Gao, D., Iwata, A., Goicoechea, J.L., et al. (2018). Genomes of 13 domesticated and wild rice relatives highlight genetic conservation, turnover and innovation across the genus *Oryza*. Nat. Genet. 50, 285–296. 10.1038/s41588-018-0040-0.

30. Tang, L., Zou, X.-h., Achoundong, G., Potgieter, C., Second, G., Zhang, D.-y., and Ge, S. (2010). Phylogeny and biogeography of the rice tribe (Oryzeae): evidence from combined analysis of 20 chloroplast fragments. Mol. Phylogen. Evol. 54, 266–277. 10.1016/j.ympev.2009.08.007.

31. Sakai, H., Lee, S.S., Tanaka, T., Numa, H., Kim, J., Kawahara, Y., Wakimoto, H., Yang, C.-c., Iwamoto, M., Abe, T., et al. (2013). Rice annotation project database (RAP-DB): an integrative and interactive database for rice genomics. Plant and Cell Physiology 54, e6–e6. 10.1093/pcp/pcs183.

32. Chen, J., Strieder, N., Krohn, N.G., Cyprys, P., Sprunck, S., Engelmann, J.C., and Dresselhaus, T. (2017). Zygotic genome activation occurs shortly after fertilization in maize. The Plant Cell 29, 2106–2125. 10.1105/tpc.17.00099.

33. Ohnishi, Y., Hoshino, R., and Okamoto, T. (2014). Dynamics of male and female chromatin during karyogamy in rice zygotes. Plant Physiol. 165, 1533–1543. 10.1104/pp.114.236059.

34. Ohnishi, Y., Kokubu, I., Kinoshita, T., and Okamoto, T. (2019). Sperm entry into the egg cell induces the progression of karyogamy in rice zygotes. Plant and Cell Physiology 60, 1656–1665. 10.1093/pcp/pcz077.

35. Kang, Y., Marischuk, K., Castelvecchi, G.D., and Bashirullah, A. (2017). HDAC inhibitors disrupt programmed resistance to apoptosis during *Drosophila* development. G3 Genes|Genomes|Genetics 7, 1985–1993. 10.1534/g3.117.041541.

36. Kim, H.-R., Kim, E.-J., Yang, S.-H., Jeong, E.-T., Park, C., Lee, J.-H., Youn, M.-J., So, H.-S., and Park, R. (2006). Trichostatin A induces apoptosis in lung cancer cells via simultaneous activation of the death receptor-mediated and mitochondrial pathway. Experimental & Molecular Medicine 38, 616–624. 10.1038/emm.2006.73.

37. Wu, D., von Roepenack-Lahaye, E., Buntru, M., de Lange, O., Schandry, N., Pérez-Quintero, A.L., Weinberg, Z., Lowe-Power, T.M., Szurek, B., Michael, A.J., et al. (2019). A plant pathogen type III effector protein subverts translational regulation to boost host polyamine levels. Cell Host & Microbe 26, 638–649.e635. 10.1016/j.chom.2019.09.014.

38. Steijger, T., Abril, J.F., Engström, P.G., Kokocinski, F., Abril, J.F., Akerman, M., Alioto, T., Ambrosini, G., Antonarakis, S.E., Behr, J., et al. (2013). Assessment of transcript reconstruction methods for RNA-seq. Nat. Methods 10, 1177–1184. 10.1038/nmeth.2714.

39. Juven-Gershon, T., Hsu, J.-Y., Theisen, J.W.M., and Kadonaga, J.T. (2008). The RNA polymerase II core promoter — the gateway to transcription. Curr. Opin. Cell Biol. 20, 253–259. 10.1016/j.ceb.2008.03.003.

40. Onodera, Y., Suzuki, A., Wu, C.-Y., Washida, H., and Takaiwa, F. (2001). A rice functional transcriptional activator, RISBZ1, responsible for endosperm-specific expression of storage protein genes through GCN4 motif. Journal of Biological Chemistry 276, 14139–14152. 10.1074/jbc.M007405200.

41. Yamamoto, M.P., Onodera, Y., Touno, S.M., and Takaiwa, F. (2006). Synergism between RPBF Dof and RISBZ1 bZIP activators in the regulation of rice seed expression genes. Plant Physiol. 141, 1694–1707. 10.1104/pp.106.082826.

42. Guo, Y.-L., and Ge, S. (2005). Molecular phylogeny of Oryzeae (Poaceae) based on DNA sequences from chloroplast, mitochondrial, and nuclear genomes. Am. J. Bot. 92, 1548–1558. 10.3732/ajb.92.9.1548.

43. Tian, D., Teo, J., and Yin, Z. (2024). Ectopic expression of the executor-type *R* gene paralog *Xa27B* in rice leads to spontaneous lesions and enhanced disease resistance. Mol. Plant-Microbe Interact. 37, 143–154. 10.1094/MPMI-10-23-0153-R.

44. Li, T., Huang, S., Zhou, J., and Yang, B. (2013). Designer TAL effectors induce disease susceptibility and resistance to *Xanthomonas oryzae* pv. *oryzae* in rice. Molecular Plant 6, 781–789. 10.1093/mp/sst034.

45. Ji, Z., Sun, H., Wei, Y., Li, M., Wang, H., Xu, J., Lei, C., Wang, C., and Zhao, K. (2022). Ectopic expression of executor gene *Xa23* enhances resistance to both bacterial and fungal diseases in rice. International Journal of Molecular Sciences 23, 6545. 10.3390/ijms23126545.

46. Zhou, S., Jiang, W., Zhao, Y., and Zhou, D.-X. (2019). Single-cell three-dimensional genome structures of rice gametes and unicellular zygotes. Nature Plants 5, 795–800. 10.1038/s41477-019-0471-3.

47. Liu, Q., Ma, X., Li, X., Zhang, X., Zhou, S., Xiong, L., Zhao, Y., and Zhou, D.-X. (2023). Paternal DNA methylation is remodeled to maternal levels in rice zygote. Nat. Commun. 14, 6571. 10.1038/s41467-023-42394-0.

48. Wang, L., O’Conner, S., Tanvir, R., Zheng, W., Cothron, S., Towery, K., Bi, H., Ellison, E.E., Yang, B., Voytas, D.F., and Li, L. (2025). CRISPR/Cas9-based editing of *NF-YC4* promoters yields high-protein rice and soybean. New Phytol. 245, 2103–2116. 10.1111/nph.20141.

49. Cenik, B.K., Aoi, Y., Iwanaszko, M., Howard, B.C., Morgan, M.A., Andersen, G.D., Bartom, E.T., and Shilatifard, A. (2024). TurboCas: A method for locus-specific labeling of genomic regions and isolating their associated protein interactome. Mol. Cell 84, 4929–4944.e4928. 10.1016/j.molcel.2024.11.007.

50. Hua, L., Wang, N., Stanley, S., Donald, R.M., Eeda, S.K., Billakurthi, K., Borba, A.R., and Hibberd, J.M. (2025). A transcription factor ensemble orchestrates bundle sheath expression in rice. Nat. Commun. 16, 7040. 10.1038/s41467-025-62087-0.

51. Sun, H., Wang, S., Yang, K., Zhu, C., Liu, Y., and Gao, Z. (2023). Development of dual-visible reporter assays to determine the DNA–protein interaction. TPJ 113, 1095–1101. 10.1111/tpj.16094.

52. Digonnet, C., Aldon, D., Leduc, N., Dumas, C., and Rougier, M. (1997). First evidence of a calcium transient in flowering plants at fertilization. Development 124, 2867–2874. 10.1242/dev.124.15.2867.

53. Antoine, A.-F., Faure, J.-E., Dumas, C., and Feijó, J.A. (2001). Differential contribution of cytoplasmic Ca^2+^ and Ca^2+^ influx to gamete fusion and egg activation in maize. Nat. Cell Biol. 3, 1120–1123. 10.1038/ncb1201-1120.

54. Ohnishi, Y., and Okamoto, T. (2017). Nuclear migration during karyogamy in rice zygotes is mediated by continuous convergence of actin meshwork toward the egg nucleus. J. Plant Res. 130, 339–348. 10.1007/s10265-016-0892-2.

55. Pónya, Z., Kristòf, Z., Ciampolini, F., Faleri, C., and Cresti, M. (2004). Structural change in the endoplasmic reticulum during the *in situ* development and *in vitro* fertilisation of wheat egg cells. Sexual Plant Reproduction 17, 177–188. 10.1007/s00497-004-0226-8.

56. Kranz, E., von Wiegen, P., and Lörz, H. (1995). Early cytological events after induction of cell division in egg cells and zygote development following *in vitro* fertilization with angiosperm gametes. TPJ 8, 9–23. 10.1046/j.1365-313X.1995.08010009.x.

57. Denninger, P., Bleckmann, A., Lausser, A., Vogler, F., Ott, T., Ehrhardt, D.W., Frommer, W.B., Sprunck, S., Dresselhaus, T., and Grossmann, G. (2014). Male–female communication triggers calcium signatures during fertilization in *Arabidopsis*. Nat. Commun. 5, 4645. 10.1038/ncomms5645.

58. Zhao, J., Yu, F., Liang, S., Zhou, C., and Yang, H. (2002). Changes of calcium distribution in egg cells, zygotes and two-celled proembryos of rice (*Oryza sativa* L.). Sexual Plant Reproduction 14, 331–337. 10.1007/s00497-002-0127-7.

59. Tian, D., and Yin, Z. (2009). Constitutive heterologous expression of *avrXa27* in rice containing the *R* gene *Xa27* confers enhanced resistance to compatible *Xanthomonas oryzae* strains. Mol. Plant Pathol. 10, 29–39. 10.1111/j.1364-3703.2008.00509.x.

60. He, L., Liu, P., Mei, L., Luo, H., Ban, T., Chen, X., and Ma, B. (2024). Disease resistance features of the executor *R* gene *Xa7* reveal novel insights into the interaction between rice and *Xanthomonas oryzae* pv. *oryzae*. Front. Plant. Sci. Volume 15–2024. 10.3389/fpls.2024.1365989.

61. Sukawa, Y., and Okamoto, T. (2018). Cell cycle in egg cell and its progression during zygotic development in rice. Plant Reproduction 31, 107–116. 10.1007/s00497-017-0318-x.

62. De Schutter, K., Joubès, J.r.m., Cools, T., Verkest, A., Corellou, F., Babiychuk, E., Van Der Schueren, E., Beeckman, T., Kushnir, S., Inzé, D., and De Veylder, L. (2007). *Arabidopsis* WEE1 kinase controls cell cycle arrest in response to activation of the DNA integrity checkpoint. The Plant Cell 19, 211–225. 10.1105/tpc.106.045047.

63. Rattanawong, K., Koiso, N., Toda, E., Kinoshita, A., Tanaka, M., Tsuji, H., and Okamoto, T. (2021). Regulatory functions of ROS dynamics via glutathione metabolism and glutathione peroxidase activity in developing rice zygote. TPJ 108, 1097–1115. 10.1111/tpj.15497.

64. Wang, J., Zeng, X., Tian, D., Yang, X., Wang, L., and Yin, Z. (2018). The pepper Bs4C proteins are localized to the endoplasmic reticulum (ER) membrane and confer disease resistance to bacterial blight in transgenic rice. Mol. Plant Pathol. 19, 2025–2035. 10.1111/mpp.12684.

65. Hamamura, Y., Nishimaki, M., Takeuchi, H., Geitmann, A., Kurihara, D., and Higashiyama, T. (2014). Live imaging of calcium spikes during double fertilization in *Arabidopsis*. Nat. Commun. 5, 4722. 10.1038/ncomms5722.

66. Wang, Q., Cang, X., Yan, H., Zhang, Z., Li, W., He, J., Zhang, M., Lou, L., Wang, R., and Chang, M. (2024). Activating plant immunity: the hidden dance of intracellular Ca^2+^ stores. New Phytol. 242, 2430–2439. 10.1111/nph.19717.

67. Boivin, V., Reulet, G., Boisvert, O., Couture, S., Elela, S.A., and Scott, M.S. (2020). Reducing the structure bias of RNA-Seq reveals a large number of non-annotated non-coding RNA. Nucleic Acids Res. 48, 2271–2286. 10.1093/nar/gkaa028.

68. Bolden, J.E., Shi, W., Jankowski, K., Kan, C.Y., Cluse, L., Martin, B.P., MacKenzie, K.L., Smyth, G.K., and Johnstone, R.W. (2013). HDAC inhibitors induce tumor-cell-selective pro-apoptotic transcriptional responses. Cell Death & Disease 4, e519–e519. 10.1038/cddis.2013.9.

69. Pu, Y., Wang, Z., Tao, S., Yang, E.J., Zhang, J., Han, Y., Wu, S., Ren, G., Chen, L.-J., Zhang, X., et al. (2026). DNA methyltransferase inhibition is a therapeutic vulnerability in VHL-deficient renal cell carcinoma cells. Experimental & Molecular Medicine 58, 798–812. 10.1038/s12276-026-01663-w.

70. Hartmanis, L., Ramsköld, D., Hendriks, G.-J., Johnsson, P., Hallén, G., Ma, R., Larsson, A.J.M., Hahne, S., Ziegenhain, C., Hartman, J., and Sandberg, R. (2025). Deciphering direct transcriptional effects of epigenetic compounds through large-scale new RNA profiling. Nat. Commun. 16, 6629. 10.1038/s41467-025-61769-z.

71. Brocks, D., Schmidt, C.R., Daskalakis, M., Jang, H.S., Shah, N.M., Li, D., Li, J., Zhang, B., Hou, Y., Laudato, S., et al. (2017). DNMT and HDAC inhibitors induce cryptic transcription start sites encoded in long terminal repeats. Nat. Genet. 49, 1052–1060. 10.1038/ng.3889.

72. Xu, Y., Miao, Y., Cai, B., Yi, Q., Tian, X., Wang, Q., Ma, D., Luo, Q., Tan, F., and Hu, Y. (2022). A histone deacetylase inhibitor enhances rice immunity by derepressing the expression of defense-related genes. Front. Plant. Sci. Volume 13–2022. 10.3389/fpls.2022.1041095.

73. Grützner, R., Schubert, R., Horn, C., Yang, C., Vogt, T., and Marillonnet, S. (2021). Engineering betalain biosynthesis in tomato for high level betanin production in fruits. Front. Plant. Sci. 12. 10.3389/fpls.2021.682443.

74. Altschul, S.F., Gish, W., Miller, W., Myers, E.W., and Lipman, D.J. (1990). Basic local alignment search tool. J. Mol. Biol. 215, 403–410. 10.1016/S0022-2836(05)80360-2.

75. Langmead, B., and Salzberg, S.L. (2012). Fast gapped-read alignment with Bowtie 2. Nat. Methods 9, 357–359. 10.1038/nmeth.1923.

76. Sievers, F., Wilm, A., Dineen, D., Gibson, T.J., Karplus, K., Li, W., Lopez, R., McWilliam, H., Remmert, M., Söding, J., et al. (2011). Fast, scalable generation of high-quality protein multiple sequence alignments using Clustal Omega. Mol. Syst. Biol. 7, 539. 10.1038/msb.2011.75.

77. Hallgren, J., Tsirigos, K.D., Pedersen, M.D., Almagro Armenteros, J.J., Marcatili, P., Nielsen, H., Krogh, A., and Winther, O. (2022). DeepTMHMM predicts alpha and beta transmembrane proteins using deep neural networks. bioRxiv, 2022.2004.2008.487609. 10.1101/2022.04.08.487609.

78. Wickham, H. (2016). Toolbox. In ggplot2: Elegant Graphics for Data Analysis, ed. (Springer International Publishing), pp. 33–74. 10.1007/978-3-319-24277-4_3.

79. Zhang, Y., Park, C., Bennett, C., Thornton, M., and Kim, D. (2021). Rapid and accurate alignment of nucleotide conversion sequencing reads with HISAT-3N. Genome Res. 31, 1290–1295. 10.1101/gr.275193.120.

80. Robinson, J.T., Thorvaldsdóttir, H., Winckler, W., Guttman, M., Lander, E.S., Getz, G., and Mesirov, J.P. (2011). Integrative genomics viewer. Nat. Biotechnol. 29, 24–26. 10.1038/nbt.1754.

81. Edgar, R.C. (2022). Muscle5: high-accuracy alignment ensembles enable unbiased assessments of sequence homology and phylogeny. Nat. Commun. 13, 6968. 10.1038/s41467-022-34630-w.

82. Madeira, F., Madhusoodanan, N., Lee, J., Eusebi, A., Niewielska, A., Tivey, A.R.N., Lopez, R., and Butcher, S. (2024). The EMBL-EBI job dispatcher sequence analysis tools framework in 2024. Nucleic Acids Res. 52, W521–W525. 10.1093/nar/gkae241.

83. Lescot, M., Déhais, P., Thijs, G., Marchal, K., Moreau, Y., Van de Peer, Y., Rouzé, P., and Rombauts, S. (2002). PlantCARE, a database of plant *cis*-acting regulatory elements and a portal to tools for *in silico* analysis of promoter sequences. Nucleic Acids Res. 30, 325–327. 10.1093/nar/30.1.325.

84. Guindon, S., Dufayard, J.-F., Lefort, V., Anisimova, M., Hordijk, W., and Gascuel, O. (2010). New algorithms and methods to estimate maximum-likelihood phylogenies: assessing the performance of PhyML 3.0. Syst. Biol. 59, 307–321. 10.1093/sysbio/syq010.

85. Li, H., Handsaker, B., Wysoker, A., Fennell, T., Ruan, J., Homer, N., Marth, G., Abecasis, G., Durbin, R., and Subgroup, G.P.D.P. (2009). The sequence alignment / map format and SAMtools. Bioinformatics 25, 2078–2079. 10.1093/bioinformatics/btp352.

86. Uchiumi, T., Komatsu, S., Koshiba, T., and Okamoto, T. (2006). Isolation of gametes and central cells from *Oryza sativa* L. Sexual Plant Reproduction 19, 37–45. 10.1007/s00497-006-0020-x.

87. Uchiumi, T., Uemura, I., and Okamoto, T. (2007). Establishment of an *in vitro* fertilization system in rice (*Oryza sativa* L.). Planta 226, 581–589. 10.1007/s00425-007-0506-2.

88. Toda, E., Ohnishi, Y., and Okamoto, T. (2016). Electro-fusion of gametes and subsequent culture of zygotes in rice. Bio-protocol 6, e2074. 10.21769/BioProtoc.2074.

