## Supplemental Data 1 for "TALE-independent transcriptional activation of the rice executor gene *Xa23* is regulated via histone acetylation during zygote development"

**Supplemental Figures**

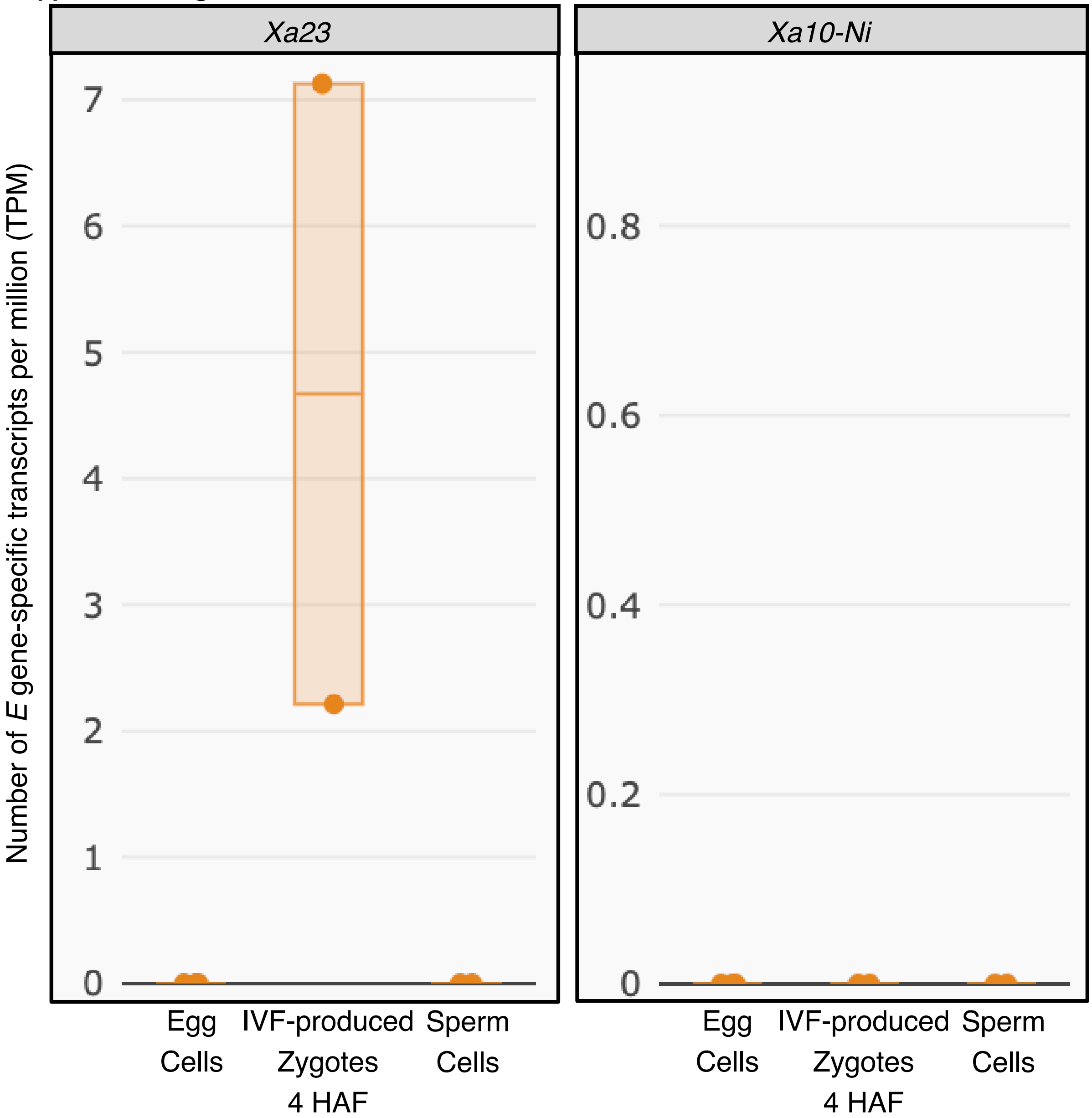


**Figure S1**. **Native *Xa23* transcripts are detectable in IVF-produced zygotes.** Data representing the number of *Xa23*- and *Xa10-Ni*-specific transcripts per million (TPM) of total transcripts in RNA-seq samples prepared from rice unfertilized egg cells, sperm cells, and IVF-produced zygotes 4 hours after fusion (4 HAF) of gametes were extracted from the Rahman *et al.* [1] database using RAP-DB.[2] *Xa10-Ni* serves as a negative control.

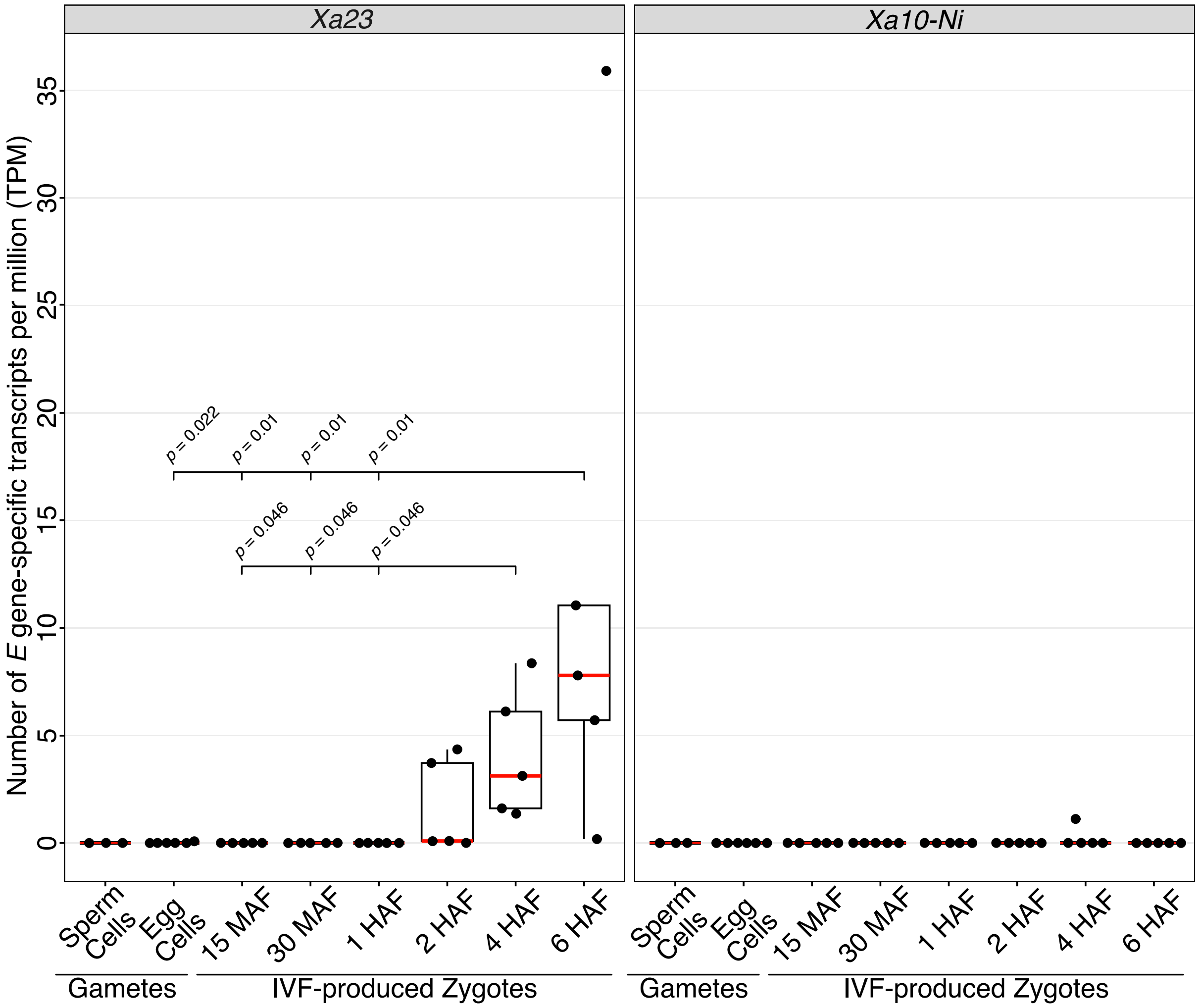


**Figure S2**. ***Xa23* expression in IVF-produced zygotes peaks at six hours after fertilization.** Number of *Xa23*- and *Xa10-Ni*-specific transcripts per million (TPM) of total transcripts in RNA-seq samples prepared from rice sperm cells, unfertilized egg cells, and IVF-produced zygotes 15 and 30 minutes after fusion (MAF) of gametes and 1 hour, 2 hours, 4 hours, and 6 hours after fusion (HAF) of gametes. *Xa10-Ni* serves as a negative control. Data representing the number of *Xa23*- and *Xa10-Ni*-specific transcripts per million were extracted from the database published by Toda *et al.* [3] . Black dots represent data points from individual biological replicates (*n*=5). Box plots show the median (center red line), lower and upper quartiles (box limits) and minimum and maximum values (whiskers). The Independent-Samples Kruskal-Wallis Test and the Bonferroni *p*-value adjustment method for multiple comparisons were used to assess statistical significance. Only adjusted *p*-values ≤ 0.05 are shown.

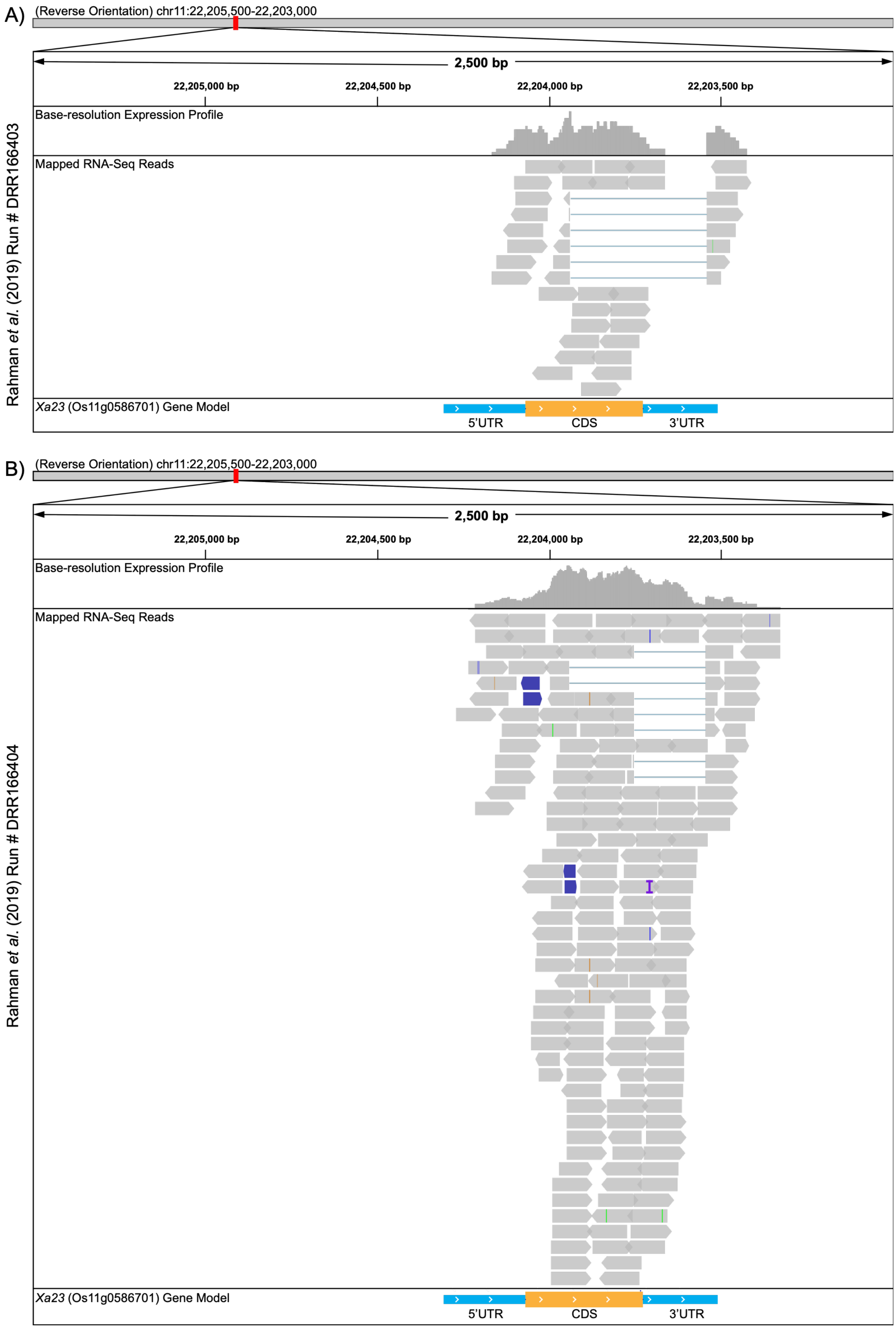
A)
(Reverse Orientation) chr11:22,205,500-22,203,000
2,500 bp
22,205,000 bp
22,204,500 bp
22,204,000 bp
22,203,500 bp
Base-resolution Expression Profile
Mapped RNA-Seq Reads
Rahman et al. (2019) Run # DRR166403
Xa23 (Os11g0586701) Gene Model
5'UTR
CDS
3'UTR
B)
(Reverse Orientation) chr11:22,205,500-22,203,000
2,500 bp
22,205,000 bp
22,204,500 bp
22,204,000 bp
22,203,500 bp
Base-resolution Expression Profile
Mapped RNA-Seq Reads
Rahman et al. (2019) Run # DRR166404
Xa23 (Os11g0586701) Gene Model
5'UTR
CDS
3'UTR

50

C)

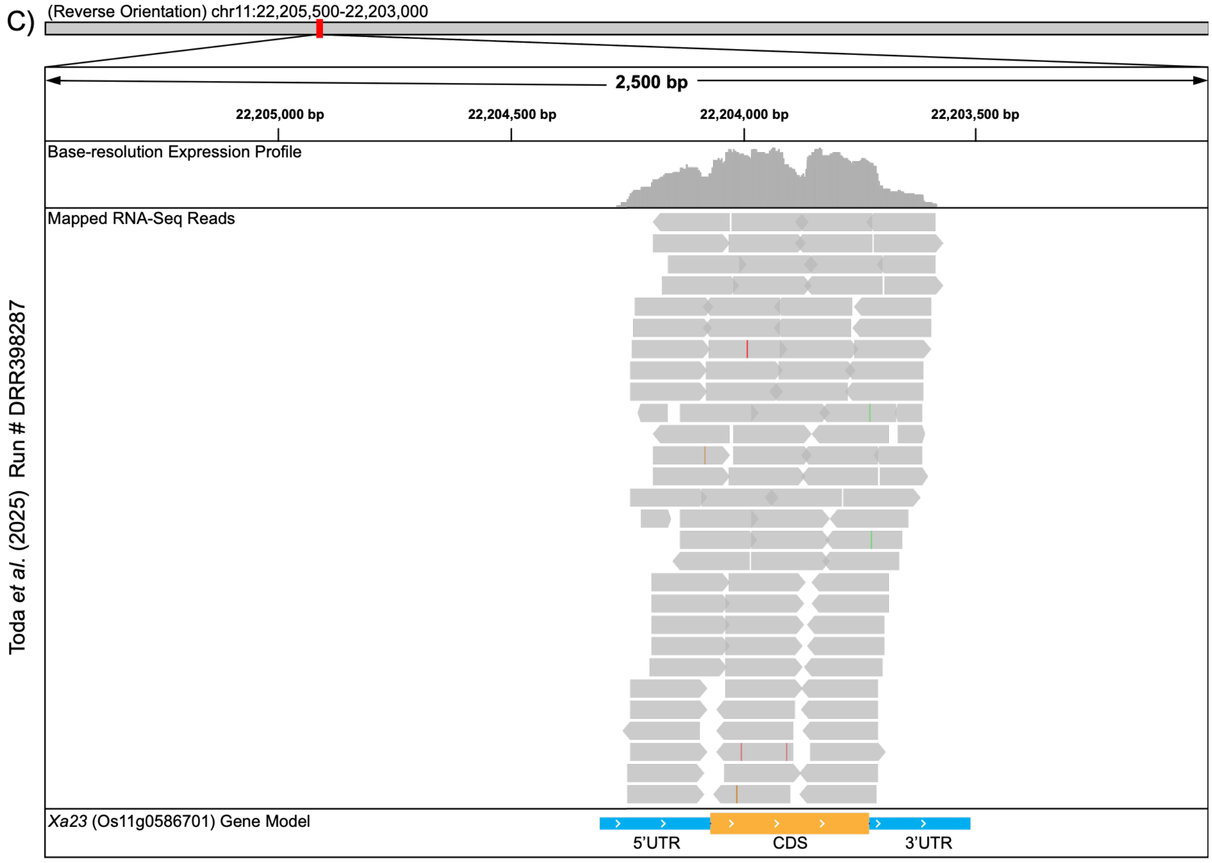
(Reverse Orientation) chr11:22,205,500-22,203,000
2,500 bp
22,205,000 bp
22,204,500 bp
22,204,000 bp
22,203,500 bp
Base-resolution Expression Profile
Mapped RNA-Seq Reads
*Xa23* (Os11g0586701) Gene Model
5'UTR
CDS
3'UTR
Toda *et al.* (2025) Run # DRR398287

D)

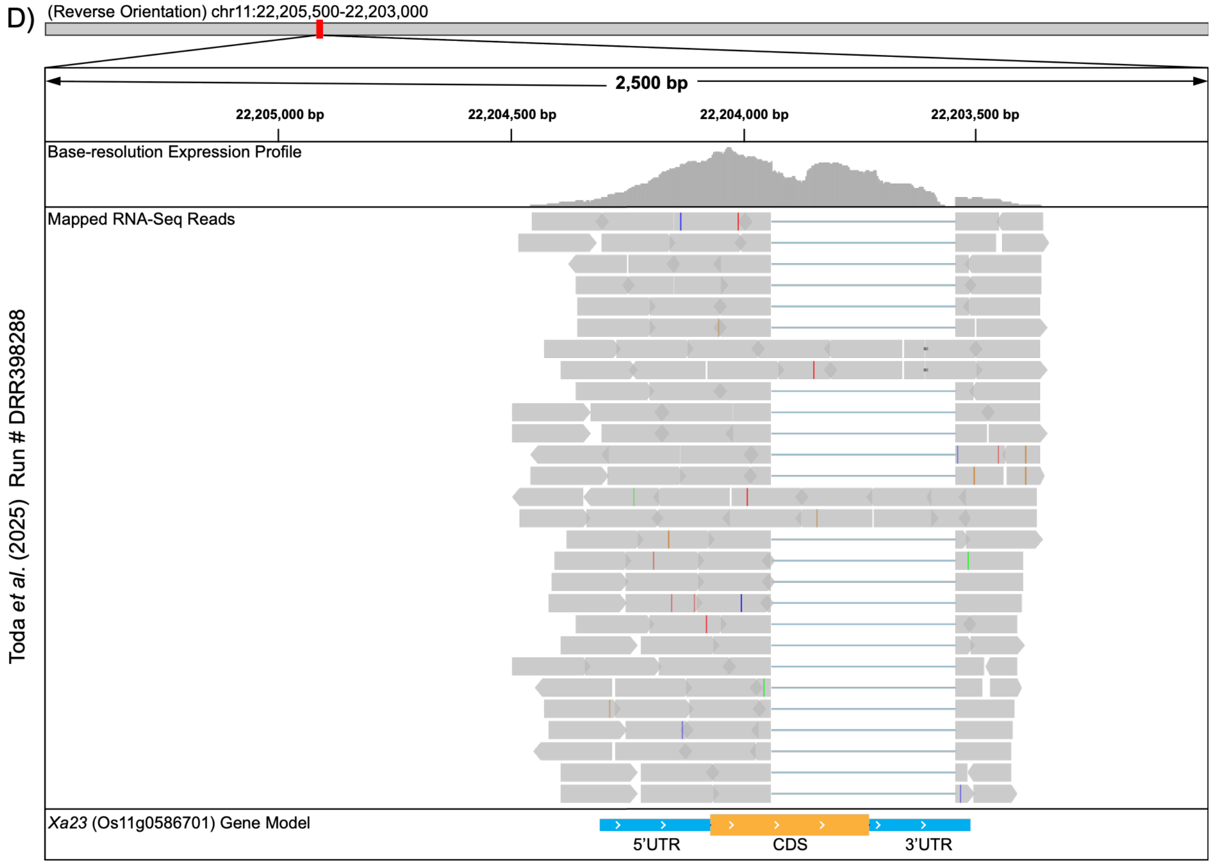
(Reverse Orientation) chr11:22,205,500-22,203,000
2,500 bp
22,205,000 bp
22,204,500 bp
22,204,000 bp
22,203,500 bp
Base-resolution Expression Profile
Mapped RNA-Seq Reads
*Xa23* (Os11g0586701) Gene Model
5'UTR
CDS
3'UTR
Toda *et al.* (2025) Run # DRR398288

51

E)

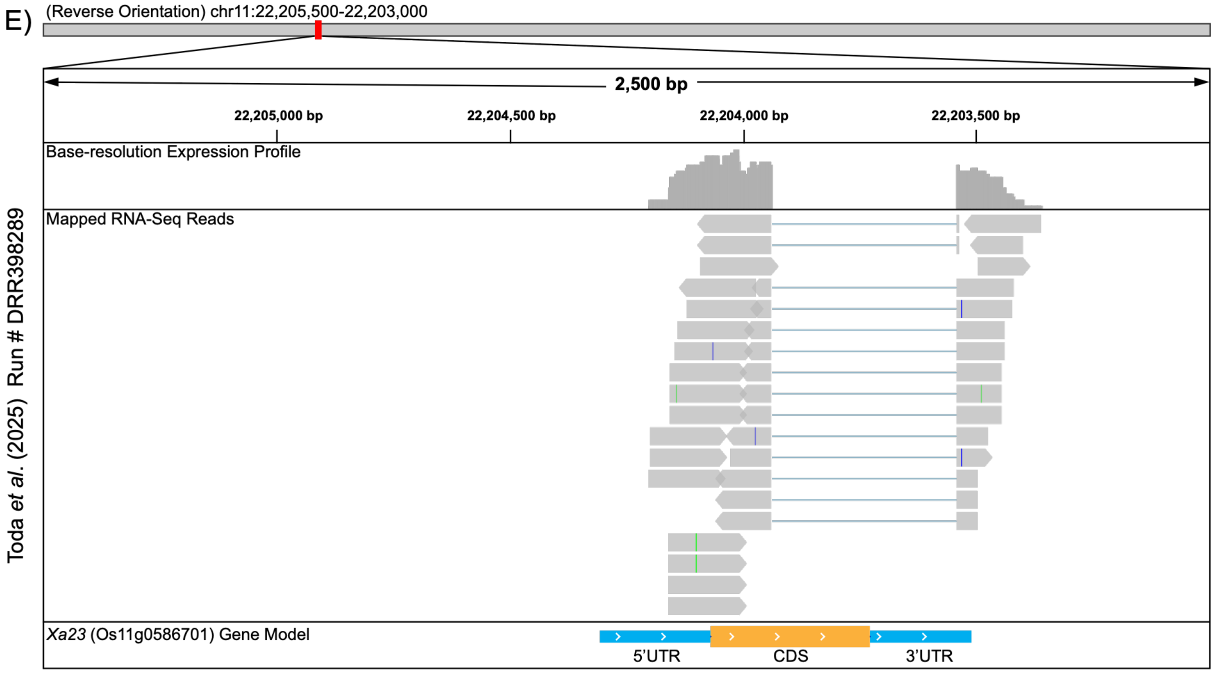
(Reverse Orientation) chr11:22,205,500-22,203,000
2,500 bp
22,205,000 bp
22,204,500 bp
22,204,000 bp
22,203,500 bp
Base-resolution Expression Profile
Mapped RNA-Seq Reads
Toda et al. (2025) Run # DRR398289
Xa23 (Os11g0586701) Gene Model
5'UTR
CDS
3'UTR

F)

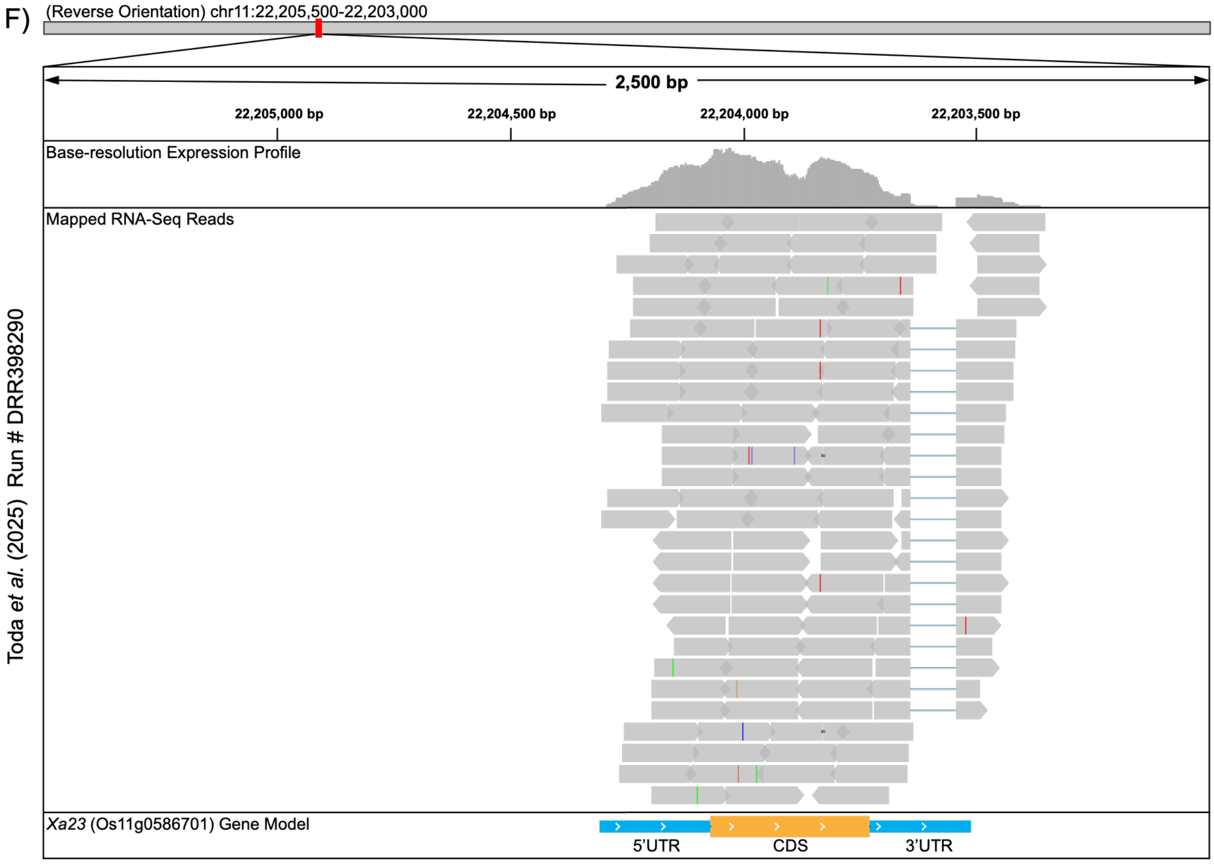
(Reverse Orientation) chr11:22,205,500-22,203,000
2,500 bp
22,205,000 bp
22,204,500 bp
22,204,000 bp
22,203,500 bp
Base-resolution Expression Profile
Mapped RNA-Seq Reads
Toda et al. (2025) Run # DRR398290
Xa23 (Os11g0586701) Gene Model
5'UTR
CDS
3'UTR

52

G)

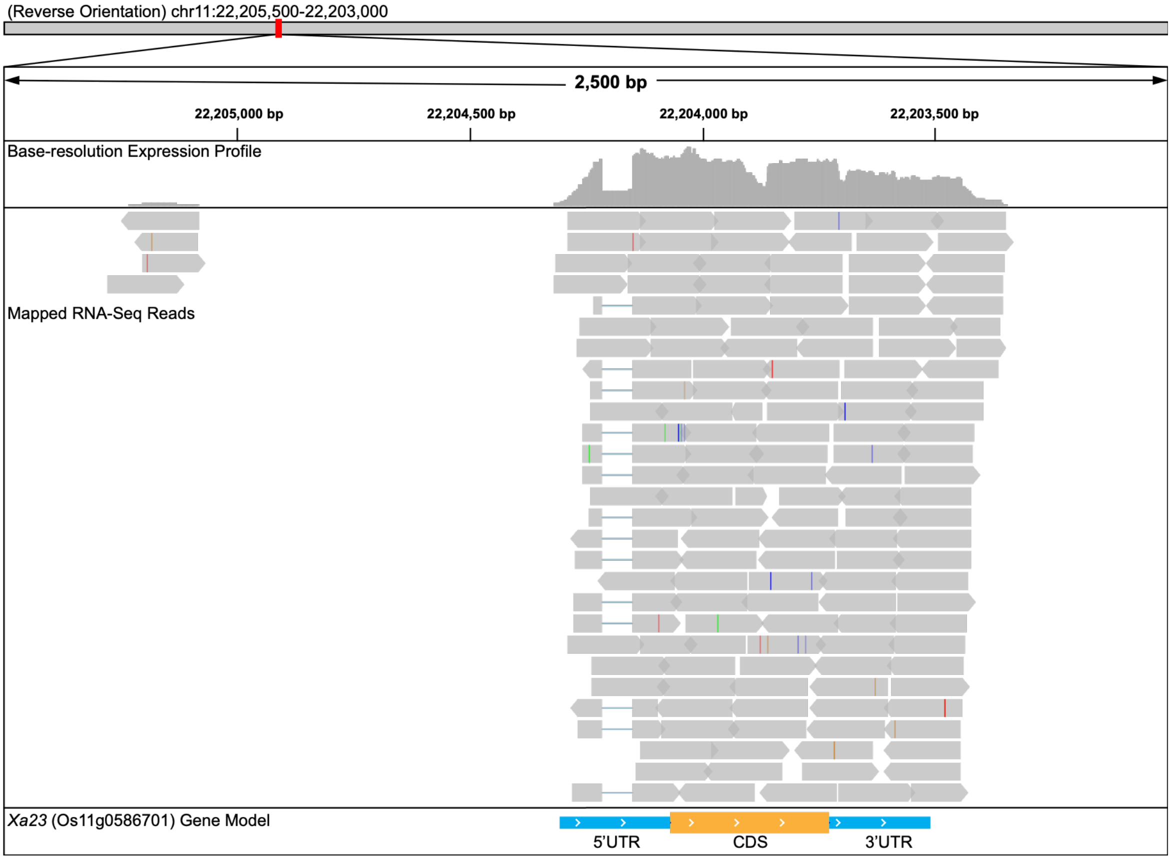


**Figure S3**. **RNA-seq reads from the published datasets map to the *Xa23* locus.** Pre-processed RNA-seq reads from datasets published by Rahman *et al.* [1] (A-B) and Toda *et al.* [3] (C-G) were mapped to the Nipponbare reference genome sequence using Bowtie2 v2.3.5.1.[4] The mapping data were converted and sorted using Samtools v1.15.1 and visualized in IGV.[5,6] The grey bar at the top of each figure represents the full length of rice chromosome 11. The vertical red line on chromosome 11 marks the location of *Xa23* (chr11:22,203,00-22,205,500 in reverse orientation). The coverage track represents the number of RNA-seq reads aligned to each nucleotide (base-resolution expression profile). Grey bars represent individual reads mapped to the *Xa23* locus. Thin lines within the grey bars represent splice junctions of individual reads. Vertical, colorful, thin lines within grey bars represent mismatches between a particular nucleotide in a read and the reference nucleotide at the same position. The *Xa23* gene model is provided. White arrows indicate the direction of *Xa23* transcription. 5' and 3' UTRs are colored in **blue**. *Xa23* CDS is colored in **orange**. Full-length sequences of the reconstructed putative native *Xa23* transcripts are available in Data S11.

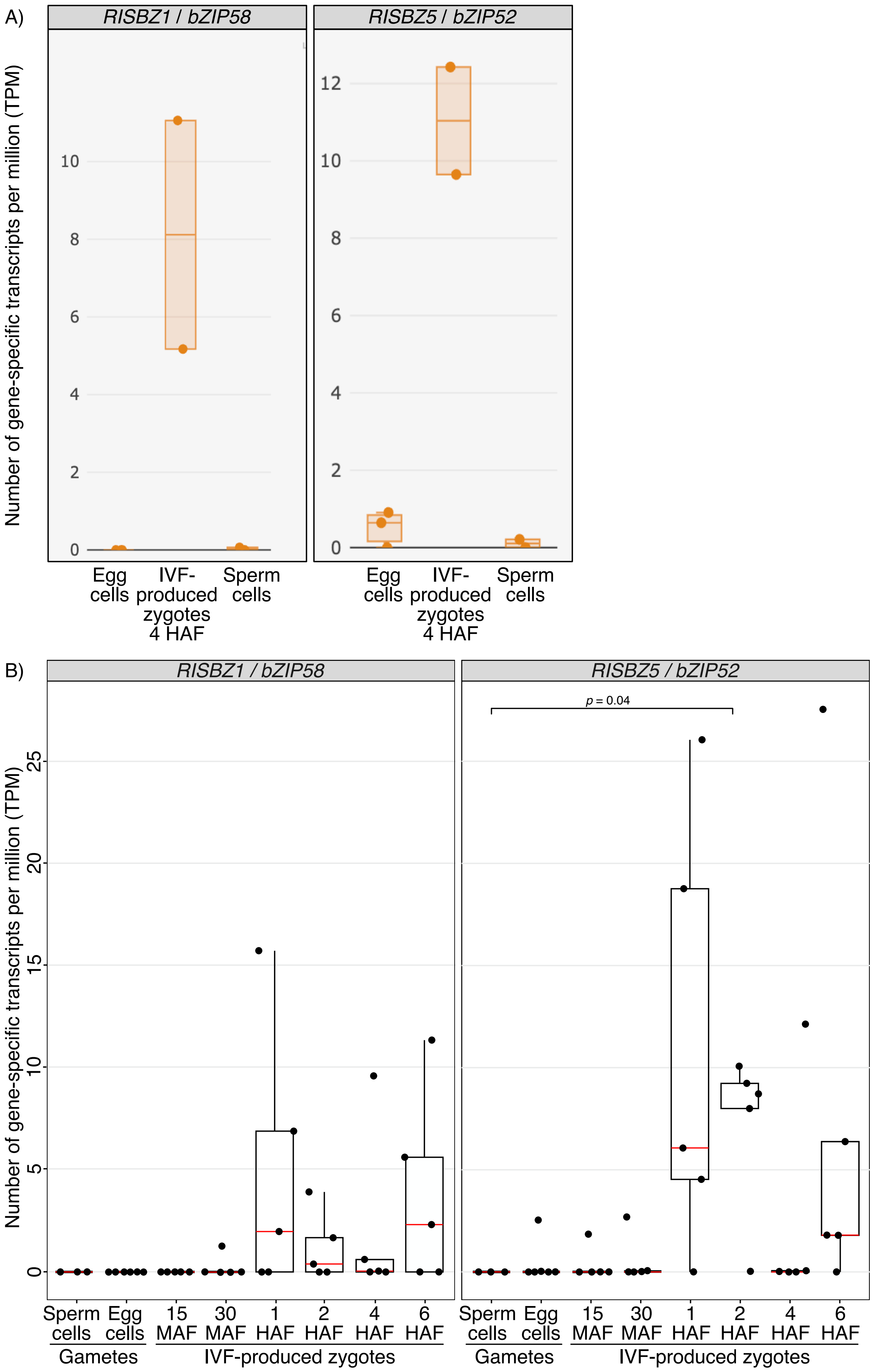
A)
RISBZ1 / bZIP58
RISBZ5 / bZIP52
Number of gene-specific transcripts per million (TPM)
Egg cells
IVF-produced zygotes 4 HAF
Sperm cells
B)
RISBZ1 / bZIP58
RISBZ5 / bZIP52
p = 0.04
Sperm cells
Egg cells
15 MAF
30 MAF
1 HAF
2 HAF
4 HAF
6 HAF
Gametes
IVF-produced zygotes

68

**Figure S4. *RISBZ1* and *RISBZ5* transcripts are detectable in IVF-produced zygotes.**
**A)** Number of *RISBZ1- and RISBZ5*-specific transcripts per million (TPM) of total transcripts in RNA-seq samples prepared from rice unfertilized egg cells, sperm cells, and IVF-produced zygotes four hours after fusion (4 HAF) of gametes. Data representing the number of *RISBZ1- and RISBZ5*-specific transcripts per million was extracted from the database published by Rahman *et al.* [1] using RAP-DB.[2]
**B)** Number of *RISBZ1- and RISBZ5*-specific transcripts per million of total transcripts in RNA-seq samples prepared from rice sperm cells, unfertilized egg cells, and IVF-produced zygotes 15 and 30 minutes after fusion (MAF) of gametes and one, two, four, and six hours after fusion (HAF) of gametes. Data representing the number of *RISBZ1- and RISBZ5*-specific transcripts per million was extracted from the database published by Toda *et al.* [3]. Black dots represent data points from individual biological replicates ($n$=5). Box plots show the median (center red line), lower and upper quartiles (box limits), and minimum and maximum values (whiskers). The Independent-Samples Kruskal-Wallis Test and the Bonferroni $p$-value adjustment method for multiple comparisons were used to assess statistical significance. Only adjusted $p$-values ≤ 0.05 are shown.

### Supplemental Tables

**Table S1. Percentage of residues that are identical between XA10, XA10-NI, and XA23 proteins and the homologs from wild rice and cutgrass species.** *E* gene-like sequences were aligned in Geneious Prime 2023.0.1 using the MUSCLE algorithm.[7] Putative CDSs were determined based on the gene structures of *Xa10* from *O. sativa* cv. IRBB10A, *Xa10-Ni* from *O. sativa* cv. Nipponbare, and *Xa23* from *O. sativa* cv. Nipponbare. Putative CDSs containing premature stop codons and encoding proteins with < 40% identity to XA10, XA10-Ni, and XA23 were considered pseudogenes and excluded from further analysis. Putative full-length proteins were selected for subsequent alignments (Data S3-S5). The percentage identity matrix was generated in Geneious Prime 2023.0.1 using the CLUSTAL OMEGA algorithm.[8] Color coding highlights putative proteins with the highest identity to XA10 (green), XA10-NI (purple), and XA23 (blue). Codes of assembled constructs and HR phenotypes induced by *Xa10*, *Xa10-Ni,* and *Xa23* homologs in *N. benthamiana* are specified (Figure 2 and Data S9).

| # | Species | Accession | # *E* gene-like sequences within the same species | Located on a chromosome or a scaffold (GeneBank Accession) | Percentage of residues identical to | | | Denomination based on amino acid identity | Transient overexpression in *N. benthamiana* | |
|---|---|---|---|---|---|---|---|---|---|---|
| | | | | | XA10 | XA10-Ni | XA23 | | Code | Phenotype |
| 1 | *L. perrieri* | IRGC 105164 | 2 | ALNV02008700.1 | 45.54 | 26.17 | 40.48 | XA10-1 | 25 | No HR |
| 2 | | | | ALNV02008701.1 | 59.52 | 27.54 | 40.54 | XA10-2 | 26 | Weak HR |
| 3 | *O. australiensis* | KR-Keep [illegible] | 1 | JAIFGZ010001913.1 | 34.38 | 52.94 | 30.43 | XA10-Ni | 16 | Strong HR |
| 4 | *O. barthii* | LB120 | 1 | PQXQ01045564.1 | 50.65 | 30.12 | 77.63 | XA23 | 1 | Strong HR |
| 5 | *O. brachyantha* | IRGC 101232 | 3 | NC_023173.2 | Premature Stop Codon |  |  | --------- | Excluded | --------- |
| 6 | | | | | 64.57 | 29.29 | 48.67 | XA10-1 | 27 | Strong HR |
| 7 | | | | | 70.37 | 29.17 | 47.32 | XA10-2 | 28 | No HR |
| 8 | *O. coarctata* | TKM-2016 | 3 | WLYV01018886.1 | < 40% amino acid identity |  |  | --------- | Excluded | --------- |
| 9 |  |  |  |  | < 40% amino acid identity |  |  | --------- | Excluded | --------- |
| 10 | | | | WLYV01049991.1 | 53.54 | 27.54 | 38.74 | XA10 | 29 | No HR |
| 11 | *O. glaberrima* | IRGC 96717 | 1 | ADWL02000023.1 | 50.91 | 28.93 | 79.65 | XA23 | 2 | Strong HR |
| 12 | *O. glumipatula* | GEN 1233 | 2 | CM002514.2 | 28.89 | 89.31 | 27.35 | XA10-Ni | 17 | No HR |
| 13 | | | | | 45.95 | 26.23 | 99.12 | XA23 | 3 | Weak HR |
| 14 | *O. longistaminata* | RD23 | 10 | CM003667.1 | 51.82 | 29.75 | 76.99 | XA23-5 | 8 | Weak HR |
| 15 | | | | CM003670.1 | 28.47 | 96.27 | 26.45 | XA10-Ni | 18 | Strong HR |
| 16 | | | | | 80.95 | 27.86 | 44.74 | XA10 | 30 | Strong HR |
| 17 | | | | | 50.45 | 27.87 | 76.11 | XA23-1 | 4 | Strong HR |
| 18 | | | | | 48.18 | 30.58 | 80.36 | XA23-2 | 5 | No HR |
| 19 |  |  |  |  | Premature Stop Codon |  |  | --------- | Excluded | --------- |
| 20 |  |  |  |  | < 40% amino acid identity |  |  | --------- | Excluded | --------- |
| 21 | | | | | 34.02 | 19.81 | 47.92 | XA23-4 | 7 | No HR |
| 22 | | | | LQBC01002215 | 47.75 | 29.51 | 84.07 | XA23-3 | 6 | Weak HR |
| 23 |  |  |  | LQBC01002312 | Premature Stop Codon |  |  | --------- | Excluded | --------- |
| 24 | *O. longistaminata* | W11 | 2 | WNHE01001006.1 | 30.3 | 51.09 | 28.57 | XA10-Ni | 19 | Strong HR |
| 25 |  |  |  |  | Premature Stop Codon |  |  | --------- | Excluded | --------- |

| # | Species | Accession | # *E-like* sequences within the same species | Located on a chromosome or a scaffold (GeneBank Accession) | Percentage of residues identical to | | | Denomination based on amino acid identity | Overexpression in *N. benthamiana* | |
|---|---|---|---|---|---|---|---|---|---|---|
| | | | | | XA10 | XA10-Ni | Code | | Code | Phenotype |
| 26 | *O. meridionalis* | OR44 (W2112) | 2 | CM002534.2 | 27.21 | 91.85 | 24.17 | XA10-Ni | 20 | No HR |
| 27 | | | | | 45.95 | 27.05 | 99.12 | XA23 | 9 | Weak HR |
| 28 | *O. meyeriana* | Menghai | 2 | SPHZ02000010.1 | 48.82 | 33.58 | 40 | XA10 | 31 | Strong HR |
| 29 | | | | | 30.65 | 61.72 | 27.97 | XA10-Ni | 21 | No HR |
| 30 | *O. nivara* | WR04-6 | 2 | QKSA01000344.1 | 28.26 | 100.00 | 27.05 | XA10-Ni | 24 | Strong HR |
| 31 | | | | | 46.85 | 27.05 | 100.00 | XA23 | 15 | Strong HR |
| 32 | *O. nivara* | IRGC 100897 | 3 | CM002502.2 | Premature Stop Codon |  |  | --------- | Excluded | --------- |
| 33 | | | | | 99.21 | 28.26 | 45.95 | XA10 | 32 | Weak HR |
| 34 |  |  |  |  | No Start Codon |  |  | --------- | Excluded | --------- |
| 35 | *O. officinalis* | W0002 | 12 | BDMV01000002.1 | < 40% amino acid identity |  |  | --------- | Excluded | --------- |
| 36 |  |  |  |  | Premature Stop Codon |  |  | --------- | Excluded | --------- |
| 37 | | | | | 35.16 | 53.68 | 34.51 | XA10-Ni | 22 | Weak HR |
| 38 |  |  |  |  | Premature Stop Codon |  |  | --------- | Excluded | --------- |
| 39 | | | | | 51.28 | 18.6 | 21.36 | XA10-1 | 33 | No HR |
| 40 |  |  |  |  | Premature Stop Codon |  |  | --------- | Excluded | --------- |
| 41 | | | | | 50 | 27.27 | 74.11 | XA23-1 | 10 | No HR |
| 42 | | | | | 52.73 | 27.27 | 76.79 | XA23-2 | 11 | Strong HR |
| 43 |  |  |  |  | Premature Stop Codon |  |  | --------- | Excluded | --------- |
| 44 |  |  |  |  | Premature Stop Codon |  |  | --------- | Excluded | --------- |
| 45 |  |  |  |  | Premature Stop Codon |  |  | --------- | Excluded | --------- |
| 46 | | | | BDMV01000011.1 | 84.00 | 27.86 | 43.86 | XA10-2 | 34 | Strong HR |
| 47 | *O. punctata* | IRGC 105690 | 3 | CM002498.2 | 79.69 | 24.29 | 45.22 | XA10-1 | 35 | Strong HR |
| 48 | | | | | 51.82 | 27.27 | 80.36 | XA23 | 12 | Strong HR |
| 49 | | | | | 48.48 | 30.43 | 38.98 | XA10-2 | 36 | Strong HR |
| 50 | *O. rufipogon* | Taxon A | 2 | LONB01001864.1 | 49.55 | 30.33 | 80.70 | XA23 | 13 | Strong HR |
| 51 | | | | | 27.56 | 47.24 | 19.63 | XA10-Ni | 23 | No HR |
| 52 | *O. rufipogon* | W1943 | 2 | CBQP010058267.1 | 49.47 | 27.36 | 100.00 | XA23 | 14 | No HR |
| 53 | | | | CBQP010058253.1 | 28.26 | 100.00 | 27.05 | XA10-Ni | 24 | Strong HR |
| 54 | *O. sativa* | Nipponbare | 2 | NC_089045.1 | 28.26 | -------- | 27.05 | XA10-Ni | 24 | Strong HR |
| 55 |  |  |  |  | 46.85 | 27.05 | -------- | XA23 | 15 | Strong HR |
| 56 | *O. sativa* | IRBB10A | 1 | JX025645.1 | -------- | 28.26 | 46.85 | XA10 | 37 | Strong HR |

**Table S2. List of primers used in this study.**

| # | Oligo | Sequence 5’>3’ | Purpose |
|---|---|---|---|
| 1 | KS481 | GGCTTGCTTCATAAACCAAG | Colony PCR for insertion of *E* gene sequences in pAGM53151 |
| 2 | KS482 | ATCCTAGTTTGCGCGCTATA | Colony PCR for insertion of *E* gene sequences in pAGM53151 & sequencing of constructs |
| 3 | KS483 | TTTGGTCTCAAATGTCTAAGGGCGAA GAACTCTTCAC | Amplification of *GFPi* for cloning into pAGM53151 |
| 4 | KS484 | TTTGGTCTCAAAGCTTATTTGTACAAC TCGTCCATGCCG | Amplification of *GFPi* for cloning into pAGM53151 |
| 5 | KS499 | CAGTGATGGTTGGGATCGCT | Forward primer for qRT-PCR of HKG *RING137* (Os08g0384900) |
| 6 | KS500 | GCTAGGCATGGGTCTTGGTT | Reverse primer for qRT-PCR of HKG *RING137* (Os08g0384900) |
| 7 | KS503 | CGTGGTGGTGAGATTATGCCT | Forward primer for qRT-PCR of *Xa23* (Os11g0586701) |
| 8 | KS504 | AGTGCTGCCAGAGAGGAAGT | Reverse primer for qRT-PCR of *Xa23* (Os11g0586701) |
| 9 | $Xa10^{Ni}$-FW | CATTGCTCCTCTCACGCCAT | Forward primer for qRT-PCR of *Xa10-Ni* (Os11g0586400) |
| 10 | $Xa10^{Ni}$-RV | TGGACATCGCCTTCGGTTTC | Reverse primer for qRT-PCR of *Xa10-Ni* (Os11g0586400) |
| 11 | KS535 | CAGATACAACTCACAACACAAGCTAG CC | Reverse primer for *Xa23* 5’RACE PCR (first PCR) |
| 12 | KS536 | AGGCCCTGAAGGCTTAAACAGGGAG | Reverse primer for *Xa23* 5’RACE PCR (first and second PCRs) |
| 13 | KS537 | CGTCGTTGTCAGCGATGTAGCGAAA GC | Reverse primer for *Xa23* 5’RACE PCR (second PCRs) |
| 14 | KS538 | GGAAAGATGAAGGCGTAGGCGAGCT CG | Reverse primer for *Xa23* 5’RACE PCR (first PCR) |
| 15 | KS539 | GATGAGGATCATGTGTATACCGGCTA CG | Reverse primer for *Xa23* 5’RACE PCR (second PCR) |
| 16 | KS505 | GATTACGCCAAGCTTCTCAAGGAGCT GGCAGCCGTAGCCG | Forward primer for *Xa23* 3’RACE PCR (first PCR) |
| 17 | KS507 | GATTACGCCAAGCTTCACATGATCCT CATCTACCTCTGCCGC | Forward primer for *Xa23* 3’RACE PCR (first and second PCRs) |
| 18 | KS529 | TCACCGTTTCCAACAGCCTCCG | Forward primer for *Xa23* 3’RACE PCR (second PCR) |
| 19 | KS540 | CGCCATGAATTGTCCTCCCAAGTTTC CA | Reverse primer for *RING137* 5’RACE PCR (first PCR) |
| 20 | KS541 | CAGATCTCCAGTGGTCAAGGCATGA GCG | Reverse primer for *RING137* 5’RACE PCR (second PCR) |
| 21 | KS543 | GTGCCGGAACTGCTGCTATGGTTGG GG | Forward primer for *RING137* 3’RACE PCR (first PCR) |
| 22 | KS544 | TTGTCGTGTTCGCCATCCTCGGTATC GC | Forward primer for *RING137* 3’RACE PCR (second PCR) |
| 23 | M13F | GTAAAACGACGGCCAG | Sequencing of the cloned RACE PCR amplicons in pCR-Blunt II-TOPO and pGEM-T Easy vectors |
| 24 | M13R | CAGGAAACAGCTATGAC | Sequencing of the cloned RACE PCR amplicons in pCR-Blunt II-TOPO and pGEM-T Easy vectors |

**Supplemental Data**

**Data S1. XA10-Ni protein variants from 3,002 cultivated rice accessions.** Genomic sequences corresponding to the chr11:22,179,001-22,182,000 of the reference genome (*O. sativa* cv. Nipponbare) were extracted from the IRRI SNP-Seek Database (http://snp-seek.irri.org).[9] Start and stop codons of *Xa10-Ni* genes from 3,002 cultivated rice accessions were identified based on the genomic structure of *Xa10-Ni* from *O. sativa* cv. Nipponbare. 89 accessions containing premature stop codons (*) within the *Xa10-Ni* CDS are highlighted in red.

```
B001    MELITACALLLIAVHIAIVISCPEPLAAVALVDVPLLYLCRYLLIIRRGVIIVSGAGAGGGRLRRFRLGAAMAIIYMAMSTLFFLRIAPLTPWWGALAAWVMILLIVEAIFAFFFPYRCCFNETEGDVQNNPHV
B002    MELITACALLLIAVHIAIVISCPEPLAAVALVDVPLLYLCRYLLIIRRGVIIVSGAGAGGGRLRRFRLGAAMAIIYMAMSTLFFLRIAPLTPWWGALAAWVMILLIVEAIFAFFFPYRCCFNETEGDVQNNPHV
B003    MELITACALLLIAVHIAIVISCPEPLAAVALVDVPLLYLYRYLLIIRRGVIIVSGAGGGGGRLRRFRLGAAMAIIYMAMSTLFFLRIAPLTPWWGALAAWVMILLIVEAIFAFFFPYRCCFNETEGDVQNNPHV
B004    MELITACALLLIAVHIAIVISCPEPLAAVALVDVPLLYLCRYLLIIRRGVIIVSGAGAGGGRLRRFRLGAAMAIIYMAMSTLFFLRIAPLTPWWGALAAWVMILLIVEAIFAFFFPYRCCFNETEGDVQNNPHV
B005    MELITACALLLIAVHIAIVISCPEPLAAVALVDVPLLYLCRYLLIIRRGVIIVSGAGAGGGRLRRFRLGAAMAIIYMAMSTLFFLRIAPLTPWWGALAAWVMILLIVEAIFAFFFPYRCCFNETEGDVQNNPHV
B006    MELITACALLLIAVHIAIVISCPEPLAAVALVDVPLLYLCRYLLIIRRGVIIVSGGGGGGRLRRFRLGAAMAIIYMAMSTLFFLRIAPLTPWWGALAAWVMILLIVEAIFAFFFPYRCCFNETEGDVQNNPHV
B007    MELITACALLLIAVHIAIVISCPEPLAAVALVDVPLLYLYRYLLIIRRGVIIVSGAGGGGGRLRRFRLGAAMAIIYMAMSTLFFLRIAPLTPWWGALAAWVMILLIVEAIFAFFFPYRCCFNETEGDVQNNPHV
B008    MELITACALLLIAVHIAIVISCPEPLAAVALVDVPLLYLCRYLLIIRRGVIIVSGAGAGGGRLRRFRLGAAMAIIYMAMSTLFFLRIAPLTPWWGALAAWVMILLIVEAIFAFFFPYRCCFNETEGDVQNNPHV
B009    MELITACALLLIAVHIAIVISCPEPLAAVALVDVPLLYLCRYLLIIRRGVIIVSGAGAGGGRLRRFRLGAAMAIIYMAMSTLFFLRIAPLTPWWGALAAWVMILLIVEAIFAFFFPYRCCFNETEGDVQNNPHV
B010    MELITACALLLIAVHIAIVISCPEPLAAVALVDVPLLYLCRYLLIIRRGVIIVSGAGGGGGRLRRFRLGAAMAIIYMAMSTLFFLRIAPLTPWWGALAAWVMILLIVEAIFAFFFPYRCCFNETEGDVQNNPHV
B011    MELITACALLLIAVHIAIVISCPEPLAAVALVDVPLLYLCRYLLIIRRGVIIVSGGGGGGRLRRFRLGAAMAIIYMAMSTLFFLRIAPLTPWWGALAAWVMILLIVEAIFAFFFPYRCCFNETEGDVQNNPHV
B012    MELITACALLLIAVHIAIVISCPEPLAAVALVDVPLLYLCRYLLIIRRGVIIVSGGGGGGRLRRFRLGAAMAIIYMAMSTLFFLRIAPLTPWWGALAAWVMILLIVEAIFAFFFPYRCCFNETEGDVQNNPHV
B013    MELITACALLLIAVHIAIVISCPEPLAAVALVDVPLLYLCRYLLIIRRGVIIVSGAGGAAAAGCAASASGQPWRSSTWPCRPSSSSALLLSRHGGAPSPPGS*SCSSSRPSSPSSSPTAAASTKPKAMSRIIPM
B015    MELITACALLLIAVHIAIVISCPEPLAAVALVDVPLLYLCRYLLIIRRGVIIVSGAGAGGGRLRRFRLGAAMAIIYMAMSTLFFLRIAPLTPWWGALAAWVMILLIVEAIFAFFFPYRCCFNETEGDVQNNPHV
B016    MELITACALLLIAVHIAIVISCPEPLAAVALVDVPLLYLCRYLLIIRRGVIIVSGAGAGGGRLRRFRLGAAMAIIYMAMSTLFFLRIAPLTPWWGALAAWVMILLIVEAIFAFFFPYRCCFNETEGDVQNNPHV
B017    MELITACALLLIAVHIAIVISCPEPLAAVALVDVPLLYLCRYLLIIRRGVIIVSGAGAGGGRLRRFRLGAAMAIIYMAMSTLFFLRIAPLTPWWGALAAWVMILLIVEAIFAFFFPYRCCFNETEGDVQNNPHV
B018    MELITACALLLIAVHIAIVISCPEPLAAVALVDVPLLYLYRYLLIIRRGVIIVSGAGGGGGRLRRFRLGAAMAIIYMAMSTLFFLRIAPLTPWWGALAAWVMILLIVEAIFAFFFPYRCCFNETEGDVQNNPHV
B019    MELITACALLLIAVHIAIVISCPEPLAAVALVDVPLLYLCRYLLIIRRGVIIVSGGGGGGRLRRFRLGAAMAIIYMAMSTLFFLRIAPLTPWWGALAAWVMILLIVEAIFAFFFPYRCCFNETEGDVQNNPHV
B020    MELITACALLLIAVHIAIVISCPEPLAAVALVDVPLLYLCRYLLIIRRGVIIVSGGGGGGRLRRFRLGAAMAIIYMAMSTLFFLRIAPLTPWWGALAAWVMILLIVEAIFAFFFPYRCCFNETEGDVQNNPHV
B021    MELITACALLLIAVHIAIVISCPEPLAAVALVDVPLLYLYRYLLIIRRGVIIVSGAGGGGGRLRRFRLGAAMAIIYMAMSTLFFLRIAPLTPWWGALAAWVMILLIVEAIFAFFFPYRCCFNETEGDVQNNPHV
B023    MELITACALLLIAVHIAIVISCPEPLAAVALVDVPLLYLCRYLLIIRRGVIIVSGAGAGGGRLRRFRLGAAMAIIYMAMSTLFFLRIAPLTPWWGALAAWVMILLIVEAIFAFFFPYRCCFNETEGDVQNNPHV
B024    MELITACALLLIAVHIAIVISCPEPLAAVALVDVPLLYLCRYLLIIRRGVIIVSGAGAGGGRLRRFRLGAAMAIIYMAMSTLFFLRIAPLTPWWGALAAWVMILLIVEAIFAFFFPYRCCFNETEGDVQNNPHV
B025    MELITACALLLIAVHIAIVISCPEPLAAVALVDVPLLYLYRYLLIIRRGVIIVSGAGGGGGRLRRFRLGAAMAIIYMAMSTLFFLRIAPLTPWWGALAAWVMILLIVEAIFAFFFPYRCCFNETEGDVQNNPHV
B026    MELITACALLLIAVHIAIVISCPEPLAAVALVDVPLLYLCRYLLIIRRGVIIVSGGGGGGRLRRFRLGAAMAIIYMAMSTLFFLRIAPLTPWWGALAAWVMILLIVEAIFAFFFPYRCCFNETEGDVQNNPHV
B027    MELITACALLLIAVHIAIVISCPEPLAAVALVDVPLLYLCRYLLIIRRGVIIVSGGGGGGRLRRFRLGAAMAIIYMAMSTLFFLRIAPLTPWWGALAAWVMILLIVEAIFAFFFPYRCCFNETEGDVQNNPHV
B028    MELITACALLLIAVHIAIVISCPEPLAAVALVDVPLLYLCRYLLIIRRGVIIVSGGGGGGRLRRFRLGAAMAIIYMAMSTLFFLRIAPLTPWWGALAAWVMILLIVEAIFAFFFPYRCCFNETEGDVQNNPHV
B029    MELITACALLLIAVHIAIVISCPEPLAAVALVDVPLLYLCRYLLIIRRGVIIVSGGGGGGRLRRFRLGAAMAIIYMAMSTLFFLRIAPLTPWWGALAAWVMILLIVEAIFAFFFPYRCCFNETEGDVQNNPHV
B030    MELITACALLLIAVHIAIVISCPEPLAAVALVDVPLLYLCRYLLIIRRGVIIVSGAGAGGGRLRRFRLGAAMAIIYMAMSTLFFLRIAPLTPWWGALAAWVMILLIVEAIFAFFFPYRCCFNETEGDVQNNPHV
B031    MELITACALLLIAVHIAIVISCPEPLAAVALVDVPLLYLCRYLLIIRRGVIIVSGGGGGGGGRLRRFRLGAAMAIIYMAMSTLFFLRIAPLTPWWGALAAWVMILLIVEAIFAFFFPYRCCFNETEGDVQNNPHV
B032    MELITACALLLIAVHIAIVISCPEPLAAVALVDVPLLYLCRYLLIIRRGVIIVSGGGGGGRLRRFRLGAAMAIIYMAMSTLFFLRIAPLTPWWGALAAWVMILLIVEAIFAFFFPYRCCFNETEGDVQNNPHV
B033    MELITACALLLIAVHIAIVISCPEPLAAVALVDVPLLYLCRYLLIIRRGVIIVSGGGGGGRLRRFRLGAAMAIIYMAMSTLFFLRIAPLTPWWGALAAWVMILLIVEAIFAFFFPYRCCFNETEGDVQNNPHV
B034    MELITACALLLIAVHIAIVISCPEPLAAVALVDVPLLYLCRYLLIIRRGVIIVSGAGAGGGRLRRFRLGAAMAIIYMAMSTLFFLRIAPLTPWWGALAAWVMILLIVEAIFAFFFPYRCCFNETEGDVQNNPHV
B035    MELITACALLLIAVHIAIVISCPEPLAAVALVDVPLLYLCRYLLIIRRGVIIVSGAGAGGGRLRRFRLGAAMAIIYMAMSTLFFLRIAPLTPWWGALAAWVMILLIVEAIFAFFFPYRCCFNETEGDVQNNPHV
B036    MELITACALLLIAVHIAIVISCPEPLAAVALVDVPLLYLYRYLLIIRRGVIIVSGAGGGGGRLRRFRLGAAMAIIYMAMSTLFFLRIAPLTPWWGALAAWVMILLIVEAIFAFFFPYRCCFNETEGDVQNNPHV
B037    MELITACALLLIAVHIAIVISCPEPLAAVALVDVPLLYLYRYLLIIRRGVIIVSGAGGGGGRLRRFRLGAAMAIIYMAMSTLFFLRIAPLTPWWGALAAWVMILLIVEAIFAFFFPYRCCFNETEGDVQNNPHV
B038    MELITACALLLIAVHIAIVISCPEPLAAVALVDVPLLYLCRYLLIIRRGVIIVSGAGAGGGRLRRFRLGAAMAIIYMAMSTLFFLRIAPLTPWWGALAAWVMILLIVEAIFAFFFPYRCCFNETEGDVQNNPHV
B039    MELITACALLLIAVHIAIVISCPEPLAAVALVDVPLLYLCRYLLIIRRGVIIVSGAGAGGGRLRRFRLGAAMAIIYMAMSTLFFLRIAPLTPWWGALAAWVMILLIVEAIFAFFFPYRCCFNETEGDVQNNPHV
B040    MELITACALLLIAVHIAIVISCPEPLAAVALVDVPLLYLCRYLLIIRRGVIIVSGGGGGGRLRRFRLGAAMAIIYMAMSTLFFLRIAPLTPWWGALAAWVMILLIVEAIFAFFFPYRCCFNETEGDVQNNPHV
B043    MELITACALLLIAVHIAIVISCPEPLAAVALVDVPLLYLCRYLLIIRRGVIIVSGAGAGGGRLRRFRLGAAMAIIYMAMSTLFFLRIAPLTPWWGALAAWVMILLIVEAIFAFFFPYRCCFNETEGDVQNNPHV
B044    MELITACALLLIAVHIAIVISCPEPLAAVALVDVPLLYLCRYLLIIRRGVIIVSGGGGGGRLRRFRLGAAMAIIYMAMSTLFFLRIAPLTPWWGALAAWVMILLIVEAIFAFFFPYRCCFNETEGDVQNNPHV
B045    MELITACALLLIAVHIAIVISCPEPLAAVALVDVPLLYLCRYLLIIRRGVIIVSGAGAGGGRLRRFRLGAAMAIIYMAMSTLFFLRIAPLTPWWGALAAWVMILLIVEAIFAFFFPYRCCFNETEGDVQNNPHV
B046    MELITACALLLIAVHIAIVISCPEPLAAVALVDVPLLYLCRYLLIIRRGVIIVSGAGAGGGRLRRFRLGAAMAIIYMAMSTLFFLRIAPLTPWWGALAAWVMILLIVEAIFAFFFPYRCCFNETEGDVQNNPHV
B047    MELITACALLLIAVHIAIVISCPEPLAAVALVDVPLLYLCRYLLIIRRGVIIVSGAGAGGGRLRRFRLGAAMAIIYMAMSTLFFLRIAPLTPWWGALAAWVMILLIVEAIFAFFFPYRCCFNETEGDVQNNPHV
B048    MELITACALLLIAVHIAIVISCPEPLAAVALVDVPLLYLCRYLLIIRRGVIIVSGGGGGGRLRRFRLGAAMAIIYMAMSTLFFLRIAPLTPWWGALAAWVMILLIVEAIFAFFFPYRCCFNETEGDVQNNPHV
B049    MELITACALLLIAVHIAIVISCPEPLAAVAFVNVPLLYLCRYLLIIRRGVIIVSGGGGGGGRLRRFRLGAAMAVIYMAMSTLFFLRIAPLTPWWGALAAWVMILLIVEAIFAFFFPYRCCFNETEGDVQNNPHV
B051    MELITACALLLIAVHIAIVISCPEPLAAVALVDVPLLYLCRYLLIIRRGVIIVSGAGAGGGRLRRFRLGAAMAIIYMAMSTLFFLRIAPLTPWWGALAAWVMILLIVEAIFAFFFPYRCCFNETEGDVQNNPHV
B052    MELITACALLLIAVHIAIVISCPEPLAAVALVDVPLLYLCRYLLIIRRGVIIVSGAGAGGGRLRRFRLGAAMAIIYMAMSTLFFLRIAPLTPWWGALAAWVMILLIVEAIFAFFFPYRCCFNETEGDVQNNPHV
B053    MELITACALLLIAVHIAIVISCPEPLAAVALVDVPLLYLYRYLLIIRRGVIIVSGAGAGGGRLRRFRLGAAMAIIYMAMSTLFFLRIAPLTPWWGALAAWVMILLIVEAIFAFFFPYRCCFNETEGDVQNNPHV
B054    MELITACALLLIAVHIAIVISCPEPLAAVALVDVPLLYLYRYLLIIRRGVIIVSGAGGGGGRLRRFRLGAAMAIIYMAMSTLFFLRIAPLTPWWGALAAWVMILLIVEAIFAFFFPYRCCFNETEGDVQNNPHV
B055    MELITACALLLIAVHIAIVISCPEPLAAVALVDVPLLYLCRYLLIIRRGVIIVSGAGAGGGRLRRFRLGAAMAIIYMAMSTLFFLRIAPLTPWWGALAAWVMILLIVEAIFAFFFPYRCCFNETEGDVQNNPHV
B056    MELITACALLLIAVHIAIVISCPEPLAAVALVDVPLLYLCRYLLIIRRGVIIVSGAGAGGGRLRRFRLGAAMAIIYMAMSTLFFLRIAPLTPWWGALAAWVMILLIVEAIFAFFFPYRCCFNETEGDVQNNPHV
B057    MELITACALLLIAVHIAIVISCPEPLAAVALVDVPLLYLCRYLLIIRRGVIIVSGAGAGGGRLRRFRLGAAMAIIYMAMSTLFFLRIAPLTPWWGALAAWVMILLIVEAIFAFFFPYRCCFNETEGDVQNNPHV
B058    MELITACALLLIAVHIAIVISCPEPLAAVALVDVPLLYLCRYLLIIRRGVIIVSGGGGGGRLRRFRLGAAMAIIYMAMSTLFFLRIAPLTPWWGALAAWVMILLIVEAIFAFFFPYRCCFNETEGDVQNNPHV
B059    MELITACALLLIAVHIAIVISCPEPLAAVALVDVPLLYLCRYLLIIRRGVIIVSGAGAGGGRLRRFRLGAAMAIIYMAMSTLFFLRIAPLTPWWGALAAWVMILLIVEAIFAFFFPYRCCFNETEGDVQNNPHV
B060    MELITACALLLIAVHIAIVISCPEPLAAVALVDVPLLYLYRYLLIIRRGVIIVSGAGGGGGRLRRFRLGAAMAIIYMAMSTLFFLRIAPLTPWWGALAAWVMILLIVEAIFAFFFPYRCCFNETEGDVQNNPHV
B061    MELITACALLLIAVHIAIVISCPEPLAAVALVDVPLLYLCRYLLIIRRGVIIVSGAGAGGGRLRRFRLGAAMAIIYMAMSTLFFLRIAPLTPWWGALAAWVMILLIVEAIFAFFFPYRCCFNETEGDVQNNPHV
B062    MELITACALLLIAVHIAIVISCPEPLAAVALVDVPLLYLCRYLLIIRRGVIIVSGGGGGGRLRRFRLGAAMAIIYMAMSTLFFLRIAPLTPWWGALAAWVMILLIVEAIFAFFFPYRCCFNETEGDVQNNPHV
B063    MELITACALLLIAVHIAIVISCPEPLAAVALVDVPLLYLCRYLLIIRRGVIIVSGAGAGGGRLRRFRLGAAMAIIYMAMSTLFFLRIAPLTPWWGALAAWVMILLIVEAIFAFFFPYRCCFNETEGDVQNNPHV
B064    MELITACALLLIAVHIAIVISCPEPLAAVALVDVPLLYLCRYLLIIRRGVIIVSGGGGGGRLRRFRLGAAMAIIYMAMSTLFFLRIAPLTPWWGALAAWVMILLIVEAIFAFFFPYRCCFNETEGDVQNNPHV
B065    MELITACALLLIAVHIAIVISCPEPLAAVALVDVPLLYLCRYLLIIRRGVIIVSGGGGGGRLRRFRLGAAMAIIYMAMSTLFFLRIAPLTPWWGALAAWVMILLIVEAIFAFFFPYRCCFNETEGDVQNNPHV
B066    MELITACALLLIAVHIAIVISCPEPLAAVALVDVPLLYLCRYLLIIRRGVIIVSGAGGGGGRLRRFRLGAAMAIIYMAMSTLFFLRIAPLTPWWGALAAWVMILLIVEAIFAFFFPYRCCFNETEGDVQNNPHV
B067    MELITACALLLIAVHIAIVISCPEPLAAVALVDVPLLYLCRYLLIIRRGVIIVSGAGAGGGRLRRFRLGAAMAIIYMAMSTLFFLRIAPLTPWWGALAAWVMILLIVEAIFAFFFPYRCCFNETEGDVQNNPHV
B068    MELITACALLLIAVHIAIVISCPEPLAAVALVDVPLLYLCRYLLIIRRGVIIVSGAGAGGGRLRRFRLGAAMAIIYMAMSTLFFLRIAPLTPWWGALAAWVMILLIVEAIFAFFFPYRCCFNETEGDVQNNPHV
B069    MELITACALLLIAVHIAIVISCPEPLAAVALVDVPLLYLYRYLLIIRRGVIIVSGAGGGGGRLRRFRLGAAMAIIYMAMSTLFFLRIAPLTPWWGALAAWVMILLIVEAIFAFFFPYRCCFNETEGDVQNNPHV
B070    MELITACALLLIAVHIAIVISCPEPLAAVALVDVPLLYLYRYLLIIRRGVIIVSGAGGGGGRLRRFRLGAAMAIIYMAMSTLFFLRIAPLTPWWGALAAWVMILLIVEAIFAFFFPYRCCFNETEGDVQNNPHV
B071    MELITACALLLIAVHIAIVISCPEPLAAVALVDVPLLYLCRYLLIIRRGVIIVSGAGAGGGRLRRFRLGAAMAIIYMAMSTLFFLRIAPLTPWWGALAAWVMILLIVEAIFAFFFPYRCCFNETEGDVQNNPHV
B072    MELITACALLLIAVHIAIVISCPEPLAAVALVDVPLLYLCRYLLIIRRGVIIVSGAGAGGGRLRRFRLGAAMAIIYMAMSTLFFLRIAPLTPWWGALAAWVMILLIVEAIFAFFFPYRCCFNETEGDVQNNPHV
B073    MELITACALLLIAVHIAIVISCPEPLAAVALVDVPLLYLCRYLLIIRRGVIIVSGGGGGGRLRRFRLGAAMAIIYMAMSTLFFLRIAPLTPWWGALAAWVMILLIVEAIFAFFFPYRCCFNETEGDVQNNPHV
B074    MELITACALLLIAVHIAIVISCPEPLAAVALVDVPLLYLCRYLLIIRRGVIIVSGAGAGGGRLRRFRLGAAMAIIYMAMSTLFFLRIAPLTPWWGALAAWVMILLIVEAIFAFFFPYRCCFNETEGDVQNNPHV
B075    MELITACALLLIAVHIAIVISCPEPLAAVALVDVPLLYLCRYLLIIRRGVIIVSGGGGGGRLRRFRLGAAMAIIYMAMSTLFFLRIAPLTPWWGALAAWVMILLIVEAIFAFFFPYRCCFNETEGDVQNNPHV
B076    MELITACALLLIAVHIAIVISCPEPLAAVALVDVPLLYLCRYLLIIRRGVIIVSGGGGGGRLRRFRLGAAMAIIYMAMSTLFFLRIAPLTPWWGALAAWVMILLIVEAIFAFFFPYRCCFNETEGDVQNNPHV
B077    MELITACALLLIAVHIAIVISCPEPLAAVALVDVPLLYLCRYLLIIRRGVIIVSGAGAGGGRLRRFRLGAAMAIIYMAMSTLFFLRIAPLTPWWGALAAWVMILLIVEAIFAFFFPYRCCFNETEGDVQNNPHV
B079    MELITACALLLIAVHIAIVISCPEPLAAVALVDVPLLYLCRYLLIIRRGVIIVSGGGGGGRLRRFRLGAAMAIIYMAMSTLFFLRIAPLTPWWGALAAWVMILLIVEAIFAFFFPYRCCFNETEGDVQNNPHV
B081    MELITACALLLIAVHIAIVISCPEPLAAVALVDVPLLYLCRYLLIIRRGVIIVSGGGGGGRLRRFRLGAAMAIIYMAMSTLFFLRIAPLTPWWGALAAWVMILLIVEAIFAFFFPYRCCFNETEGDVQNNPHV
B082    MELITACALLLIAVHIAIVISCPEPLAAVALVDVPLLYLCRYLLIIRRGVIIVSGGGGGGRLRRFRLGAAMAIIYMAMSTLFFLRIAPLTPWWGALAAWVMILLIVEAIFAFFFPYRCCFNETEGDVQNNPHV
B083    MELITACALLLIAVHIAIVISCPEPLAAVALVDVPLLYLCRYLLIIRRGVIIVSGAGAGGGRLRRFRLGAAMAIIYMAMSTLFFLRIAPLTPWWGALAAWVMILLIVEAIFAFFFPYRCCFNETEGDVQNNPHV
B084    MELITACALLLIAVHIAIVISCPEPLAAVALVDVPLLYLCRYLLIIRRGVIIVSGAGAGGGRLRRFRLGAAMAIIYMAMSTLFFLRIAPLTPWWGALAAWVMILLIVEAIFAFFFPYRCCFNETEGDVQNNPHV
B085    MELITACALLLIAVHIAIVISCPEPLAAVALVDVPLLYLCRYLLIIRRGVIIVSGAGAGGGRLRRFRLGAAMAIIYMAMSTLFFLRIAPLTPWWGALAAWVMILLIVEAIFAFFFPYRCCFNETEGDVQNNPHV
B086    MELITACALLLIAVHIAIVISCPEPLAAVALVDVPLLYLCRYLLIIRRGVIIVSGGGGGGRLRRFRLGAAMAIIYMAMSTLFFLRIAPLTPWWGALAAWVMILLIVEAIFAFFFPYRCCFNETEGDVQNNPHV
B087    MELITACALLLIAVHIAIVISCPEPLAAVALVDVPLLYLCRYLLIIRRGVIIVSGAGAGGGRLRRFRLGAAMAIIYMAMSTLFFLRIAPLTPWWGALAAWVMILLIVEAIFAFFFPYRCCFNETEGDVQNNPHV
B088    MELITACALLLIAVHIAIVISCPEPLAAVALVDVPLLYLCRYLLIIRRGVIIVSGAGAGGGRLRRFRLGAAMAIIYMAMSTLFFLRIAPLTPWWGALAAWVMILLIVEAIFAFFFPYRCCFNETEGDVQNNPHV
B089    MELITACALLLIAVHIAIVISCPEPLAAVALVDVPLLYLYRYLLIIRRGVIIVSGAGGGGGRLRRFRLGAAMAIIYMAMSTLFFLRIAPLTPWWGALAAWVMILLIVEAIFAFFFPYRCCFNETEGDVQNNPHV
B090    MELITACALLLIAVHIAIVISCPEPLAAVALVDVPLLYLYRYLLIIRRGVIIVSGAGGGGGRLRRFRLGAAMAIIYMAMSTLFFLRIAPLTPWWGALAAWVMILLIVEAIFAFFFPYRCCFNETEGDVQNNPHV
B091    MELITACALLLIAVHIAIVISCPEPLAAVALVDVPLLYLYRYLLIIRRGVIIVSGAGAGGGRLRRFRLGAAMAIIYMAMSTLFFLRIAPLTPWWGALAAWVMILLIVEAIFAFFFPYRCCFNETEGDVQNNPHV
B092    MELITACALLLIAVHIAIVISCPEPLAAVALVDVPLLYLCRYLLIIRRGVIIVSGAGAGGGRLRRFRLGAAMAIIYMAMSTLFFLRIAPLTPWWGALAAWVMILLIVEAIFAFFFPYRCCFNETEGDVQNNPHV
B093    MELITACALLLIAVHIAIVISCPEPLAAVALVDVPLLYLYRYLLIIRRGVIIVSGAGGGGGRLRRFRLGAAMAIIYMAMSTLFFLRIAPLTPWWGALAAWVMILLIVEAIFAFFFPYRCCFNETEGDVQNNPHV
B094    MELITACALLLIAVHIAIVISCPEPLAAVALVDVPLLYLYRYLLIIRRGVIIVSGAGAGGGRLRRFRLGAAMAIIYMAMSTLFFLRIAPLTPWWGALAAWVMILLIVEAIFAFFFPYRCCFNETEGDVQNNPHV
B095    MELITACALLLIAVHIAIVISCPEPLAAVALVDVPLLYLCRYLLIIRRGVIIVSGAGGAAAAGCAASASGQPWRSSTWPCRPSSSSALLLSRHGGAPSPPGS*SCSSSRPSSPSSSPTAAASTKPKAMSRIIPM
B096    MELITACALLLIAVHIAIVISCPEPLAAVALVDVPLLYLCRYLLIIRRGVIIVSGAGGAAAAGCAASASGQPWRSSTWPCRPSSSSALLLSRHGGAPSPPGS*SCSSSRPSSPSSSPTAAASTKPKAMSRIIPM
B097    MELITACALLLIAVHIAIVISCPEPLAAVALVDVPLLYLCRYLLIIRRGVIIVSGGGGGGRLRRFRLGAAMAIIYMAMSTLFFLRIAPLTPWWGALAAWVMILLIVEAIFAFFFPYRCCFNETEGDVQNNPHV
B100    MELITACALLLIAVHIAIVISCPEPLAAVALVDVPLLYLCRYLLIIRRGVIIVSGAGGGGGRLRRFRLGAAMAIIYMAMSTLFFLRIAPLTPWWGALAAWVMILLIVEAIFAFFFPYRCCFNETEGDVQNNPHV
B102    MELITACALLLIAVHIAIVISCPEPLAAVALVDVPLLYLCRYLLIIRRGVIIVSGAGAGGGRLRRFRLGAAMAIIYMAMSTLFFLRIAPLTPWWGALAAWVMILLIVEAIFAFFFPYRCCFNETEGDVQNNPHV
B103    MELITACALLLIAVHIAIVISCPEPLAAVALVDVPLLYLCRYLLIIRRGVIIVSGAGGGGGRLRRFRLGAAMAIIYMAMSTLFFLRIAPLTPWWGALAAWVMILLIVEAIFAFFFPYRCCFNETEGDVQNNPHV
B104    MELITACALLLIAVHIAIVISCPEPLAAVALVDVPLLYLYRYLLIIRRGVIIVSGAGGGGGRLRRFRLGAAMAIIYMAMSTLFFLRIAPLTPWWGALAAWVMILLIVEAIFAFFFPYRCCFNETEGDVQNNPHV
B105    MELITACALLLIAVHIAIVISCPEPLAAVALVDVPLLYLCRYLLIIRRGVIIVSGGGGGGRLRRFRLGAAMAIIYMAMSTLFFLRIAPLTPWWGALAAWVMILLIVEAIFAFFFPYRCCFNETEGDVQNNPHV
B106    MELITACALLLIAVHIAIVISCPEPLAAVALVDVPLLYLCRYLLIIRRGVIIVSGGAGGGRLRRFRLGAAMAIIYMAMSTLFFLRIAPLTPWWGALAAWVMILLIVEAIFAFFFPYRCCFNETEGDVQNNPHV
B107    MELITACALLLIAVHIAIVISCPEPLAAVALVDVPLLYLCRYLLIIRRGVIIVSGAGGAAAAGCAASASGQPWRSSTWPCRPSSSSALLLSRHGGAPSPPGS*SCSSSRPSSPSSSPTAAASTKPKAMSRIIPM
B108    MELITACALLLIAVHIAIVISCPEPLAAVALVDVPLLYLYRYLLIIRRGVIIVSGAGAGGGRLRRFRLGAAMAIIYMAMSTLFFLRIAPLTPWWGALAAWVMILLIVEAIFAFFFPYRCCFNETEGDVQNNPHV
B109    MELITACALLLIAVHIAIVISCPEPLAAVALVDVPLLYLYRYLLIIRRGVIIVSGAGGGGGRLRRFRLGAAMAIIYMAMSTLFFLRIAPLTPWWGALAAWVMILLIVEAIFAFFFPYRCCFNETEGDVQNNPHV
B110    MELITACALLLIAVHIAIVISCPEPLAAVALVDVPLLYLYRYLLIIRRGVIIVSGAGGGGGRLRRFRLGAAMAIIYMAMSTLFFLRIAPLTPWWGALAAWVMILLIVEAIFAFFFPYRCCFNETEGDVQNNPHV
B111    MELITACALLLIAVHIAIVISCPEPLAAVALVDVPLLYLYRYLLIIRRGVIIVSGAGGGGGRLRRFRLGAAMAIIYMAMSTLFFLRIAPLTPWWGALAAWVMILLIVEAIFAFFFPYRCCFNETEGDVQNNPHV
B112    MELITACALLLIAVHIAIVISCPEPLAAVALVDVPLLYLCRYLLIIRRGVIIVSGAGAGGGRLRRFRLGAAMAIIYMAMSTLFFLRIAPLTPWWGALAAWVMILLIVEAIFAFFFPYRCCFNETEGDVQNNPHV
B113    MELITACALLLIAVHIAIVISCPEPLAAVALVDVPLLYLYRYLLIIRRGVIIVSGAGGGGGRLRRFRLGAAMAIIYMAMSTLFFLRIAPLTPWWGALAAWVMILLIVEAIFAFFFPYRCCFNETEGDVQNNPHV
B114    MELITACALLLIAVHIAIVISCPEPLAAVALVDVPLLYLCRYLLIIRRGVIIVSGGGGGGRLRRFRLGAAMAIIYMAMSTLFFLRIAPLTPWWGALAAWVMILLIVEAIFAFFFPYRCCFNETEGDVQNNPHV
B115    MELITACALLLIAVHIAIVISCPEPLAAVALVDVPLLYLCRYLLIIRRGVIIVSGAGAGGGRLRRFRLGAAMAIIYMAMSTLFFLRIAPLTPWWGALAAWVMILLIVEAIFAFFFPYRCCFNETEGDVQNNPHV
```

B116 MELITACALLLIAVHIAIVISCPEPLAAVALVDVPLLYLYRYLLIIRRGVIIVSGAGGGGGRLRRFRLGAAMAIIYMAMSTLFFLRIAPLTPWWGALAAWVMILLIVEAIFAFFFPYRCCFNETEGDVQNNPHV
B117 MELITACALLLIAVHIAIVISCPEPLAAVALVDVPLLYLCRYLLIIRRGVIIVSGAGAGGGRLRRFRLGAAMAIIYMAMSTLFFLRIAPLTPWWGALAAWVMILLIVEAIFAFFFPYRCCFNETEGDVQNNPHV
B118 MELITACALLLIAVHIAIVISCPEPLAAVALVDVPLLYLCRYLLIIRRGVIIVSGGGGGGRLRRFRLGAAMAIIYMAMSTLFFLRIAPLTPWWGALAAWVMILLIVEAIFAFFFPYRCCFNETEGDVQNNPHV
B119 MELITACALLLIAVHIAIVISCPEPLAAVALVDVPLLYLCRYLLIIRRGVIIVSGAGAGGGRLRRFRLGAAMAIIYMAMSTLFFLRIAPLTPWWGALAAWVMILLIVEAIFAFFFPYRCCFNETEGDVQNNPHV
B120 MELITACALLLIAVHIAIVISCPEPLAAVALVDVPLLYLCRYLLIIRRGVIIVSGGGGGGRLRRFRLGAAMAIIYMAMSTLFFLRIAPLTPWWGALAAWVMILLIVEAIFAFFFPYRCCFNETEGDVQNNPHV
B121 MELITACALLLIAVHIAIVISCPEPLAAVALVDVPLLYLCRYLLIIRRGVIIVSGAGAGGGRLRRFRLGAAMAIIYMAMSTLFFLRIAPLTPWWGALAAWVMILLIVEAIFAFFFPYRCCFNETEGDVQNNPHV
B122 MELITACALLLIAVHIAIVISCPEPLAAVALVDVPLLYLCRYLLIIRRGVIIVSGAGAGGGRLRRFRLGAAMAIIYMAMSTLFFLRIAPLTPWWGALAAWVMILLIVEAIFAFFFPYRCCFNETEGDVQNNPHV
B123 MELITACALLLIAVHIAIVISCPEPLAAVALVDVPLLYLYRYLLIIRRGVIIVSGAGGGGGRLRRFRLGAAMAIIYMAMSTLFFLRIAPLTPWWGALAAWVMILLIVEAIFAFFFPYRCCFNETEGDVQNNPHV
B124 MELITACALLLIAVHIAIVISCPEPLAAVALVDVPLLYLCRYLLIIRRGVIIVSGGGGGGRLRRFRLGAAMAIIYMAMSTLFFLRIAPLTPWWGALAAWVMILLIVEAIFAFFFPYRCCFNETEGDVQNNPHV
B125 MELITACALLLIAVHIAIVISCPEPLAAVALVDVPLLYLCRYLLIIRRGVIIVSGGGGGGRLRRFRLGAAMAIIYMAMSTLFFLRIAPLTPWWGALAAWVMILLIVEAIFAFFFPYRCCFNETEGDVQNNPHV
B126 MELITACALLLIAVHIAIVISCPEPLAAVALVDVPLLYLCRYLLIIRRGVIIVSGAGAGGGRLRRFRLGAAMAIIYMAMSTLFFLRIAPLTPWWGALAAWVMILLIVEAIFAFFFPYRCCFNETEGDVQNNPHV
B127 MELITACALLLIAVHIAIVISCPEPLAAVALVDVPLLYLCRYLLIIRRGVIIVSGAGAGGGRLRRFRLGAAMAIIYMAMSTLFFLRIAPLTPWWGALAAWVMILLIVEAIFAFFFPYRCCFNETEGDVQNNPHV
B128 MELITACALLLIAVHIAIVISCPEPLAAVALVDVPLLYLCRYLLIIRRGVIIVSGGGGGGRLRRFRLGAAMAIIYMAMSTLFFLRIAPLTPWWGALAAWVMILLIVEAIFAFFFPYRCCFNETEGDVQNNPHV
B129 MELITACALLLIAVHIAIVISCPEPLAAVALVDVPLLYLYRYLLIIRRGVIIVSGGGGGGRLRRFRLGAAMAIIYMAMSTLFFLRIAPLTPWWGALAAWVMILLIVEAIFAFFFPYRCCFNETEGDVQNNPHV
B130 MELITACALLLIAVHIAIVISCPEPLAAVALVDVPLLYLYRYLLIIRRGVIIVSGAGAGGGRLRRFRLGAAMAIIYMAMSTLFFLRIAPLTPWWGALAAWVMILLIVEAIFAFFFPYRCCFNETEGDVQNNPHV
B131 MELITACALLLIAVHIAIVISCPEPLAAVALVDVPLLYLCRYLLIIRRGVIIVSGGGGGGRLRRFRLGAAMAIIYMAMSTLFFLRIAPLTPWWGALAAWVMILLIVEAIFAFFFPYRCCFNETEGDVQNNPHV
B132 MELITACALLLIAVHIAIVISCPEPLAAVALVDVPLLYLCRYLLIIRRGVIIVSGGGGGGRLRRFRLGAAMAIIYMAMSTLFFLRIAPLTPWWGALAAWVMILLIVEAIFAFFFPYRCCFNETEGDVQNNPHV
B133 MELITACALLLIAVHIAIVISCPEPLAAVALVDVPLLYLYRYLLIIRRGVIIVSGAGGGGGRLRRFRLGAAMAIIYMAMSTLFFLRIAPLTPWWGALAAWVMILLIVEAIFAFFFPYRCCFNETEGDVQNNPHV
B134 MELITACALLLIAVHIAIVISCPEPLAAVALVDVPLLYLCRYLLIIRRGVIIVSGAGGGGGRLRRFRLGAAMAIIYMAMSTLFFLRIAPLTPWWGALAAWVMILLIVEAIFAFFFPYRCCFNETEGDVQNNPHV
B135 MELITACALLLIAVHIAIVISCPEPLAAVALVDVPLLYLYRYLLIIRRGVIIVSGAGGGGGRLRRFRLGAAMAIIYMAMSTLFFLRIAPLTPWWGALAAWVMILLIVEAIFAFFFPYRCCFNETEGDVQNNPHV
B136 MELITACALLLIAVHIAIVISCPEPLAAVALVDVPLLYLYRYLLIIRRGVIIVSGAGGGGGRLRRFRLGAAMAIIYMAMSTLFFLRIAPLTPWWGALAAWVMILLIVEAIFAFFFPYRCCFNETEGDVQNNPHV
B137 MELITACALLLIAVHIAIVISCPEPLAAVALVDVPLLYLCRYLLIIRRGVIIVSGAGAGGGRLRRFRLGAAMAIIYMAMSTLFFLRIAPLTPWWGALAAWVMILLIVEAIFAFFFPYRCCFNETEGDVQNNPHV
B138 MELITACALLLIAVHIAIVISCPEPLAAVALVDVPLLYLCRYLLIIRRGVIIVSGGGAGGGRLRRFRLGAAMAIIYMAMSTLFFLRIAPLTPWWGALAAWVMILLIVEAIFAFFFPYRCCFNETEGDVQNNPHV
B139 MELITACALLLIAVHIAIVISCPEPLAAVALVDVPLLYLCRYLLIIRRGVIIVSGAGAGGGRLRRFRLGAAMAIIYMAMSTLFFLRIAPLTPWWGALAAWVMILLIVEAIFAFFFPYRCCFNETEGDVQNNPHV
B141 MELITACALLLIAVHIAIVISCPEPLAAVALVDVPLLYLCRYLLIIRRGVIIVSGAGAGGGRLRRFRLGAAMAIIYMAMSTLFFLRIAPLTPWWGALAAWVMILLIVEAIFAFFFPYRCCFNETEGDVQNNPHV
B142 MELITACALLLIAVHIAIVISCPEPLAAVALVDVPLLYLCRYLLIIRRGVIIVSGAGAGGGRLRRFRLGAAMAIIYMAMSTLFFLRIAPLTPWWGALAAWVMILLIVEAIFAFFFPYRCCFNETEGDVQNNPHV
B143 MELITACALLLIAVHIAIVISCPEPLAAVALVDVPLLYLCRYLLIIRRGVIIVSGAGAGGGRLRRFRLGAAMAIIYMAMSTLFFLRIAPLTPWWGALAAWVMILLIVEAIFAFFFPYRCCFNETEGDVQNNPHV
B144 MELITACALLLIAVHIAIVISCPEPLAAVALVDVPLLYLCRYLLIIRRGVIIVSGAGGGGGRLRRFRLGAAMAIIYMAMSTLFFLRIAPLTPWWGALAAWVMILLIVEAIFAFFFPYRCCFNETEGDVQNNPHV
B145 MELITACALLLIAVHIAIVISCPEPLAAVALVDVPLLYLCRYLLIIRRGVIIVSGGGGGGRLRRFRLGAAMAIIYMAMSTLFFLRIAPLTPWWGALAAWVMILLIVEAIFAFFFPYRCCFNETEGDVQNNPHV
B146 MELITACALLLIAVHIAIVISCPEPLAAVALVDVPLLYLCRYLLIIRRGVIIVSGGGGGGRLRRFRLGAAMAIIYMAMSTLFFLRIAPLTPWWGALAAWVMILLIVEAIFAFFFPYRCCFNETEGDVQNNPHV
B147 MELITACALLLIAVHIAIVISCPEPLAAVALVDVPLLYLCRYLLIIRRGVIIVSGGGGGGRLRRFRLGAAMAIIYMAMSTLFFLRIAPLTPWWGALAAWVMILLIVEAIFAFFFPYRCCFNETEGDVQNNPHV
B148 MELITACALLLIAVHIAIVISCPEPLAAVALVDVPLLYLCRYLLIIRRGVIIVSGAGAGGGRLRRFRLGAAMAIIYMAMSTLFFLRIAPLTPWWGALAAWVMILLIVEAIFAFFFPYRCCFNETEGDVQNNPHV
B149 MELITACALLLIAVHIAIVISCPEPLAAVALVDVPLLYLCRYLLIIRRGVIIVSGAGAGGGRLRRFRLGAAMAIIYMAMSTLFFLRIAPLTPWWGALAAWVMILLIVEAIFAFFFPYRCCFNETEGDVQNNPHV
B150 MELITACALLLIAVHIAIVISCPEPLAAVALVDVPLLYLCRYLLIIRRGVIIVSGAGAGGGRLRRFRLGAAMAIIYMAMSTLFFLRIAPLTPWWGALAAWVMILLIVEAIFAFFFPYRCCFNETEGDVQNNPHV
B151 MELITACALLLIAVHIAIVISCPEPLAAVALVDVPLLYLYRYLLIIRRGVIIVSGAGAGGGRLRRFRLGAAMAIIYMAMSTLFFLRIAPLTPWWGALAAWVMILLIVEAIFAFFFPYRCCFNETEGDVQNNPHV
B152 MELITACALLLIAVHIAIVISCPEPLAAVALVDVPLLYLCRYLLIIRRGVIIVSGAGAGGGRLRRFRLGAAMAIIYMAMSTLFFLRIAPLTPWWGALAAWVMILLIVEAIFAFFFPYRCCFNETEGDVQNNPHV
B153 MELITACALLLIAVHIAIVISCPEPLAAVALVDVPLLYLYRYLLIIRRGVIIVSGAGGGGGRLRRFRLGAAMAIIYMAMSTLFFLRIAPLTPWWGALAAWVMILLIVEAIFAFFFPYRCCFNETEGDVQNNPHV
B154 MELITACALLLIAVHIAIVISCPEPLAAVALVDVPLLYLCRYLLIIRRGVIIVSGAGAGGGRLRRFRLGAAMAIIYMAMSTLFFLRIAPLTPWWGALAAWVMILLIVEAIFAFFFPYRCCFNETEGDVQNNPHV
B155 MELITACALLLIAVHIAIVISCPEPLAAVALVDVPLLYLCRYLLIIRRGVIIVSGAGAGGGRLRRFRLGAAMAIIYMAMSTLFFLRIAPLTPWWGALAAWVMILLIVEAIFAFFFPYRCCFNETEGDVQNNPHV
B156 MELITACALLLIAVHIAIVISCPEPLAAVALVDVPLLYLYRYLLIIRRGVIIVSGAGAGGGRLRRFRLGAAMAIIYMAMSTLFFLRIAPLTPWWGALAAWVMILLIVEAIFAFFFPYRCCFNETEGDVQNNPHV
B157 MELITACALLLIAVHIAIVISCPEPLAAVALVDVPLLYLCRYLLIIRRGVIIVSGGGGGGRLRRFRLGAAMAIIYMAMSTLFFLRIAPLTPWWGALAAWVMILLIVEAIFAFFFPYRCCFNETEGDVQNNPHV
B158 MELITACALLLIAVHIAIVISCPEPLAAVALVDVPLLYLCRYLLIIRRGVIIVSGAGAGGGRLRRFRLGAAMAIIYMAMSTLFFLRIAPLTPWWGALAAWVMILLIVEAIFAFFFPYRCCFNETEGDVQNNPHV
B159 MELITACALLLIAVHIAIVISCPEPLAAVALVDVPLLYLCRYLLIIRRGVIIVSGAGAGGGRLRRFRLGAAMAIIYMAMSTLFFLRIAPLTPWWGALAAWVMILLIVEAIFAFFFPYRCCFNETEGDVQNNPHV
B160 MELITACALLLIAVHIAIVISCPEPLAAVALVDVPLLYLCRYLLIIRRGVIIVSGAGAGGGRLRRFRLGAAMAIIYMAMSTLFFLRIAPLTPWWGALAAWVMILLIVEAIFAFFFPYRCCFNETEGDVQNNPHV
B161 MELITACALLLIAVHIAIVISCPEPLAAVALVDVPLLYLCRYLLIIRRGVIIVSGAGAGGGRLRRFRLGAAMAIIYMAMSTLFFLRIAPLTPWWGALAAWVMILLIVEAIFAFFFPYRCCFNETEGDVQNNPHV
B162 MELITACALLLIAVHIAIVISCPEPLAAVALVDVPLLYLCRYLLIIRRGVIIVSGAGAGGGRLRRFRLGAAMAIIYMAMSTLFFLRIAPLTPWWGALAAWVMILLIVEAIFAFFFPYRCCFNETEGDVQNNPHV
B163 MELITACALLLIAVHIAIVISCPEPLAAVALVDVPLLYLCRYLLIIRRGVIIVSGGGGGGRLRRFRLGAAMAIIYMAMSTLFFLRIAPLTPWWGALAAWVMILLIVEAIFAFFFPYRCCFNETEGDVQNNPHV
B164 MELITACALLLIAVHIAIVISCPEPLAAVAFVNVPLLYLCRYLLIIRRGVIIVSGGGGGGGRLRRFRLGAAMAVIYMAMSTLFFLRIAPLTPWWGALAAWVMILLIVEAIFAFFFPYRCCFNETEGDVQNNPHV
B165 MELITACALLLIAVHIAIVISCPEPLAAVALVDVPLLYLCRYLLIIRRGVIIVSGGGGGGRLRRFRLGAAMAIIYMAMSTLFFLRIAPLTPWWGALAAWVMILLIVEAIFAFFFPYRCCFNETEGDVQNNPHV
B167 MELITACALLLIAVHIAIVISCPEPLAAVALVDVPLLYLCRYLLIIRRGVIIVSGAGAGGGRLRRFRLGAAMAIIYMAMSTLFFLRIAPLTPWWGALAAWVMILLIVEAIFAFFFPYRCCFNETEGDVQNNPHV
B168 MELITACALLLIAVHIAIVISCPEPLAAVALVDVPLLYLYRYLLIIRRGVIIVSGAGGGGGRLRRFRLGAAMAIIYMAMSTLFFLRIAPLTPWWGALAAWVMILLIVEAIFAFFFPYRCCFNETEGDVQNNPHV
B169 MELITACALLLIAVHIAIVISCPEPLAAVALVDVPLLYLCRYLLIIRRGVIIVSGAGAGGGRLRRFRLGAAMAIIYMAMSTLFFLRIAPLTPWWGALAAWVMILLIVEAIFAFFFPYRCCFNETEGDVQNNPHV
B170 MELITACALLLIAVHIAIVISCPEPLAAVALVDVPLLYLCRYLLIIRRGVIIVSGAGAGGGRLRRFRLGAAMAIIYMAMSTLFFLRIAPLTPWWGALAAWVMILLIVEAIFAFFFPYRCCFNETEGDVQNNPHV
B171 MELITACALLLIAVHIAIVISCPEPLAAVALVDVPLLYLCRYLLIIRRGVIIVSGAGAGGGRLRRFRLGAAMAIIYMAMSTLFFLRIAPLTPWWGALAAWVMILLIVEAIFAFFFPYRCCFNETEGDVQNNPHV
B173 MELITACALLLIAVHIAIVISCPEPLAAVALVDVPLLYLYRYLLIIRRGVIIVSGAGGGGGRLRRFRLGAAMAIIYMAMSTLFFLRIAPLTPWWGALAAWVMILLIVEAIFAFFFPYRCCFNETEGDVQNNPHV
B176 MELITACALLLIAVHIAIVISCPEPLAAVALVDVPLLYLCRYLLIIRRGVIIVSGGGGGGRLRRFRLGAAMAIIYMAMSTLFFLRIAPLTPWWGALAAWVMILLIVEAIFAFFFPYRCCFNETEGDVQNNPHV
B179 MELITACALLLIAVHIAIVISCPEPLAAVALVDVPLLYLCRYLLIIRRGVIIVSGAGAGGGRLRRFRLGAAMAIIYMAMSTLFFLRIAPLTPWWGALAAWVMILLIVEAIFAFFFPYRCCFNETEGDVQNNPHV
B180 MELITACALLLIAVHIAIVISCPEPLAAVALVDVPLLYLCRYLLIIRRGVIIVSGAGAGGGRLRRFRLGAAMAIIYMAMSTLFFLRIAPLTPWWGALAAWVMILLIVEAIFAFFFPYRCCFNETEGDVQNNPHV
B182 MELITACALLLIAVHIAIVISCPEPLAAVALVDVPLLYLCRYLLIIRRGVIIVSGAGAGGGRLRRFRLGAAMAIIYMAMSTLFFLRIAPLTPWWGALAAWVMILLIVEAIFAFFFPYRCCFNETEGDVQNNPHV
B183 MELITACALLLIAVHIAIVISCPEPLAAVALVDVPLLYLCRYLLIIRRGVIIVSGAGAGGGRLRRFRLGAAMAIIYMAMSTLFFLRIAPLTPWWGALAAWVMILLIVEAIFAFFFPYRCCFNETEGDVQNNPHV
B184 MELITACALLLIAVHIAIVISCPEPLAAVALVDVPLLYLCRYLLIIRRGVIIVSGAGAGGGRLRRFRLGAAMAIIYMAMSTLFFLRIAPLTPWWGALAAWVMILLIVEAIFAFFFPYRCCFNETEGDVQNNPHV
B185 MELITACALLLIAVHIAIVISCPEPLAAVALVDVPLLYLCRYLLIIRRGVIIVSGAGAGGGRLRRFRLGAAMAIIYMAMSTLFFLRIAPLTPWWGALAAWVMILLIVEAIFAFFFPYRCCFNETEGDVQNNPHV
B187 MELITACALLLIAVHIAIVISCPEPLAAVALVDVPLLYLCRYLLIIRRGVIIVSGAGAGGGRLRRFRLGAAMAIIYMAMSTLFFLRIAPLTPWWGALAAWVMILLIVEAIFAFFFPYRCCFNETEGDVQNNPHV
B188 MELITACALLLIAVHIAIVISCPEPLAAVALVDVPLLYLCRYLLIIRRGVIIVSGAGGGGGRLRRFRLGAAMAIIYMAMSTLFFLRIAPLTPWWGALAAWVMILLIVEAIFAFFFPYRCCFNETEGDVQNNPHV
B189 MELITACALLLIAVHIAIVISCPEPLAAVALVDVPLLYLCRYLLIIRRGVIIVSGAGAGGGRLRRFRLGAAMAIIYMAMSTLFFLRIAPLTPWWGALAAWVMILLIVEAIFAFFFPYRCCFNETEGDVQNNPHV
B190 MELITACALLLIAVHIAIVISCPEPLAAVALVDVPLLYLCRYLLIIRRGVIIVSGAGAGGGRLRRFRLGAAMAIIYMAMSTLFFLRIAPLTPWWGALAAWVMILLIVEAIFAFFFPYRCCFNETEGDVQNNPHV
B191 MELITACALLLIAVHIAIVISCPEPLAAVALVDVPLLYLYRYLLIIRRGVIIVSGAGGGGGRLRRFRLGAAMAIIYMAMSTLFFLRIAPLTPWWGALAAWVMILLIVEAIFAFFFPYRCCFNETEGDVQNNPHV
B192 MELITACALLLIAVHIAIVISCPEPLAAVALVDVPLLYLCRYLLIIRRGVIIVSGAGAGGGRLRRFRLGAAMAIIYMAMSTLFFLRIAPLTPWWGALAAWVMILLIVEAIFAFFFPYRCCFNETEGDVQNNPHV
B193 MELITACALLLIAVHIAIVISCPEPLAAVALVDVPLLYLCRYLLIIRRGVIIVSGGGGGGRLRRFRLGAAMAIIYMAMSTLFFLRIAPLTPWWGALAAWVMILLIVEAIFAFFFPYRCCFNETEGDVQNNPHV
B194 MELITACALLLIAVHIAIVISCPEPLAAVALVDVPLLYLCRYLLIIRRGVIIVSGGGGGGRLRRFRLGAAMAIIYMVMSTLFFLRIAPLTPWWGALAAWVMILLIVEAIFAFFFPYRCCFNETEGDVQNNPHV
B195 MELITACALLLIAVHIAIVISCPEPLAAVALVDVPLLYLCRYLLIIRRGVIIVSGGGGGGRLRRFRLGAAMAIIYMAMSTLFFLRIAPLTPWWGALAAWVMILLIVEAIFAFFFPYRCCFNETEGDVQNNPHV
B196 MELITACALLLIAVHIAIVISCPEPLAAVALVDVPLLYLYRYLLIIRRGVIIVSGAGGGGGRLRRFRLGAAMAIIYMAMSTLFFLRIAPLTPWWGALAAWVMILLIVEAIFAFFFPYRCCFNETEGDVQNNPHV
B197 MELITACALLLIAVHIAIVISCPEPLAAVALVDVPLLYLYRYLLIIRRGVIIVSGAGGGGGRLRRFRLGAAMAIIYMAMSTLFFLRIAPLTPWWGALAAWVMILLIVEAIFAFFFPYRCCFNETEGDVQNNPHV
B198 MELITACALLLIAVHIAIVISCPEPLAAVALVDVPLLYLCRYLLIIRRGVIIVSGAGGGGGRLRRFRLGAAMAIIYMAMSTLFFLRIAPLTPWWGALAAWVMILLIVEAIFAFFFPYRCCFNETEGDVQNNPHV
B199 MELITACALLLIAVHIAIVISCPEPLAAVALVDVPLLYLCRYLLIIRRGVIIVSGAGAGGGRLRRFRLGAAMAIIYMAMSTLFFLRIAPLTPWWGALAAWVMILLIVEAIFAFFFPYRCCFNETEGDVQNNPHV
B200 MELITACALLLIAVHIAIVISCPEPLAAVALVDVPLLYLCRYLLIIRRGVIIVSGAGAGGGRLRRFRLGAAMAIIYMAMSTLFFLRIAPLTPWWGALAAWVMILLIVEAIFAFFFPYRCCFNETEGDVQNNPHV
B201 MELITACALLLIAVHIAIVISCPEPLAAVALVDVPLLYLCRYLLIIRRGVIIVSGAGAGGGRLRRFRLGAAMAIIYMAMSTLFFLRIAPLTPWWGALAAWVMILLIVEAIFAFFFPYRCCFNETEGDVQNNPHV
B202 MELITACALLLIAVHIAIVISCPEPLAAVALVDVPLLYLCRYLLIIRRGVIIVSGAGAGGGRLRRFRLGAAMAIIYMAMSTLFFLRIAPLTPWWGALAAWVMILLIVEAIFAFFFPYRCCFNETEGDVQNNPHV
B203 MELITACALLLIAVHIAIVISCPEPLAAVALVDVPLLYLCRYLLIIRRGVIIVSGGGGGGRLRRFRLGAAMAIIYMAMSTLFFLRIAPLTPWWGALAAWVMILLIVEAIFAFFFPYRCCFNETEGDVQNNPHV
B204 MELITACALLLIAVHIAIVISCPEPLAAVALVDVPLLYLCRYLLIIRRGVIIVSGAGAGGGRLRRFRLGAAMAIIYMAMSTLFFLRIAPLTPWWGALAAWVMILLIVEAIFAFFFPYRCCFNETEGDVQNNPHV
B205 MELITACALLLIAVHIAIVISCPEPLAAVALVDVPLLYLCRYLLIIRRGVIIVSGAGGGGGRLRRFRLGAAMAIIYMAMSTLFFLRIAPLTPWWGALAAWVMILLIVEAIFAFFFPYRCCFNETEGDVQNNPHV
B207 MELITACALLLIAVHIAIVISCPEPLAAVALVDVPLLYLCRYLLIIRRGVIIVSGGGGGGRLRRFRLGAAMAIIYMAMSTLFFLRIAPLTPWWGALAAWVMILLIVEAIFAFFFPYRCCFNETEGDVQNNPHV
B208 MELITACALLLIAVHIAIVISCPEPLAAVALVDVPLLYLYRYLLIIRRGVIIVSGAGGGGGRLRRFRLGAAMAIIYMAMSTLFFLRIAPLTPWWGALAAWVMILLIVEAIFAFFFPYRCCFNETEGDVQNNPHV
B210 MELITACALLLIAVHIAIVISCPEPLAAVALVDVPLLYLYRYLLIIRRGVIIVSGAGGGGGRLRRFRLGAAMAIIYMAMSTLFFLRIAPLTPWWGALAAWVMILLIVEAIFAFFFPYRCCFNETEGDVQNNPHV
B212 MELITACALLLIAVHIAIVISCPEPLAAVALVDVPLLYLCRYLLIIRRGVIIVSGAGAGGGRLRRFRLGAAMAIIYMAMSTLFFLRIAPLTPWWGALAAWVMILLIVEAIFAFFFPYRCCFNETEGDVQNNPHV
B213 MELITACALLLIAVHIAIVISCPEPLAAVALVDVPLLYLYRYLLIIRRGVIIVSGAGGGGGRLRRFRLGAAMAIIYMAMSTLFFLRIAPLTPWWGALAAWVMILLIVEAIFAFFFPYRCCFNETEGDVQNNPHV
B214 MELITACALLLIAVHIAIVISCPEPLAAVALVDVPLLYLCRYLLIIRRGVIIVSGGGGGGRLRRFRLGAAMAIIYMAMSTLFFLRIAPLTPWWGALAAWVMILLIVEAIFAFFFPYRCCFNETEGDVQNNPHV
B215 MELITACALLLIAVHIAIVISCPEPLAAVALVDVPLLYLCRYLLIIRRGVIIVSGAGAGGGRLRRFRLGAAMAIIYMAMSTLFFLRIAPLTPWWGALAAWVMILLIVEAIFAFFFPYRCCFNETEGDVQNNPHV
B216 MELITACALLLIAVHIAIVISCPEPLAAVALVDVPLLYLYRYLLIIRRGVIIVSGAGGGGGRLRRFRLGAAMAIIYMAMSTLFFLRIAPLTPWWGALAAWVMILLIVEAIFAFFFPYRCCFNETEGDVQNNPHV
B217 MELITACALLLIAVHIAIVISCPEPLAAVALVDVPLLYLCRYLLIIRRGVIIVSGAGAGGGRLRRFRLGAAMAIIYMAMSTLFFLRIAPLTPWWGALAAWVMILLIVEAIFAFFFPYRCCFNETEGDVQNNPHV
B218 MELITACALLLIAVHIAIVISCPEPLAAVALVDVPLLYLCRYLLIIRRGVIIVSGAGGGGGRLRRFRLGAAMAIIYMAMSTLFFLRIAPLTPWWGALAAWVMILLIVEAIFAFFFPYRCCFNETEGDVQNNPHV
B219 MELITACALLLIAVHIAIVISCPEPLAAVALVDVPLLYLCRYLLIIRRGVIIVSGGGGGGRLRRFRLGAAMAIIYMAMSTLFFLRIAPLTPWWGALAAWVMILLIVEAIFAFFFPYRCCFNETEGDVQNNPHV
B221 MELITACALLLIAVHIAIVISCPEPLAAVAFVNVPLLYLCRYLLIIRRGVIIVSGGGGGGGRLRRFRLGAAMAVIYMAMSTLFFLRIAPLTPWWGALAAWVMILLIVEAIFAFFFPYRCCFNETEGDVQNNPHV
B222 MELITACALLLIAVHIAIVISCPEPLAAVALVDVPLLYLYRYLLIIRRGVIIVSGGGGGGRLRRFRLGAAMAIIYMAMSTLFFLRIAPLTPWWGALAAWVMILLIVEAIFAFFFPYRCCFNETEGDVQNNPHV
B223 MELITACALLLIAVHIAIVISCPEPLAAVALVDVPLLYLCRYLLIIRRGVIIVSGAGAGGGRLRRFRLGAAMAIIYMAMSTLFFLRIAPLTPWWGALAAWVMILLIVEAIFAFFFPYRCCFNETEGDVQNNPHV
B224 MELITACALLLIAVHIAIVISCPEPLAAVALVDVPLLYLCRYLLIIRRGVIIVSGGGGGGRLRRFRLGAAMAIIYMAMSTLFFLRIAPLTPWWGALAAWVMILLIVEAIFAFFFPYRCCFNETEGDVQNNPHV
B225 MELITACALLLIAVHIAIVISCPEPLAAVALVDVPLLYLCRYLLIIRRGVIIVSGAGAGGGRLRRFRLGAAMAIIYMAMSTLFFLRIAPLTPWWGALAAWVMILLIVEAIFAFFFPYRCCFNETEGDVQNNPHV
B226 MELITACALLLIAVHIAIVISCPEPLAAVALVDVPLLYLCRYLLIIRRGVIIVSGAGAGGGRLRRFRLGAAMAIIYMAMSTLFFLRIAPLTPWWGALAAWVMILLIVEAIFAFFFPYRCCFNETEGDVQNNPHV
B227 MELITACALLLIAVHIAIVISCPEPLAAVALVDVPLLYLCRYLLIIRRGVIIVSGAGAGGGRLRRFRLGAAMAIIYMAMSTLFFLRIAPLTPWWGALAAWVMILLIVEAIFAFFFPYRCCFNETEGDVQNNPHV
B228 MELITACALLLIAVHIAIVISCPEPLAAVALVDVPLLYLCRYLLIIRRGVIIVSGAGAGGGRLRRFRLGAAMAIIYMAMSTLFFLRIAPLTPWWGALAAWVMILLIVEAIFAFFFPYRCCFNETEGDVQNNPHV
B229 MELITACALLLIAVHIAIVISCPEPLAAVALVDVPLLYLYRYLLIIRRGVIIVSGAGGGGGRLRRFRLGAAMAIIYMAMSTLFFLRIAPLTPWWGALAAWVMILLIVEAIFAFFFPYRCCFNETEGDVQNNPHV
B230 MELITACALLLIAVHIAIVISCPEPLAAVALVDVPLLYLYRYLLIIRRGVIIVSGAGGGGGRLRRFRLGAAMAIIYMAMSTLFFLRIAPLTPWWGALAAWVMILLIVEAIFAFFFPYRCCFNETEGDVQNNPHV
B231 MELITACALLLIAVHIAIVISCPEPLAAVALVDVPLLYLCRYLLIIRRGVIIVSGGGGGGRLRRFRLGAAMAIIYMAMSTLFFLRIAPLTPWWGALAAWVMILLIVEAIFAFFFPYRCCFNETEGDVQNNPHV
B232 MELITACALLLIAVHIAIVISCPEPLAAVALVDVPLLYLCRYLLIIRRGVIIVSGAGAGGGRLRRFRLGAAMAIIYMAMSTLFFLRIAPLTPWWGALAAWVMILLIVEAIFAFFFPYRCCFNETEGDVQNNPHV
B233 MELITACALLLIAVHIAIVISCPEPLAAVALVDVPLLYLCRYLLIIRRGVIIVSGGGGGGRLRRFRLGAAMAIIYMAMSTLFFLRIAPLTPWWGALAAWVMILLIVEAIFAFFFPYRCCFNETEGDVQNNPHV
B234 MELITACALLLIAVHIAIVISCPEPLAAVALVDVPLLYLYRYLLIIRRGVIIVSGAGGGGGRLRRFRLGAAMAIIYMAMSTLFFLRIAPLTPWWGALAAWVMILLIVEAIFAFFFPYRCCFNETEGDVQNNPHV
B235 MELITACALLLIAVHIAIVISCPEPLAAVALVDVPLLYLCRYLLIIRRGVIIVSGAGAGGGRLRRFRLGAAMAIIYMAMSTLFFLRIAPLTPWWGALAAWVMILLIVEAIFAFFFPYRCCFNETEGDVQNNPHV
B236 MELITACALLLIAVHIAIVISCPEPLAAVALVDVPLLYLCRYLLIIRRGVIIVSGAGAGGGRLRRFRLGAAMAIIYMAMSTLFFLRIAPLTPWWGALAAWVMILLIVEAIFAFFFPYRCCFNETEGDVQNNPHV
B238 MELITACALLLIAVHIAIVISCPEPLAAVALVDVPLLYLYRYLLIIRRGVIIVSGAGGGGGRLRRFRLGAAMAIIYMAMSTLFFLRIAPLTPWWGALAAWVMILLIVEAIFAFFFPYRCCFNETEGDVQNNPHV
B239 MELITACALLLIAVHIAIVISCPEPLAAVALVDVPLLYLYRYLLIIRRGVIIVSGAGGGGGRLRRFRLGAAMAIIYMAMSTLFFLRIAPLTPWWGALAAWVMILLIVEAIFAFFFPYRCCFNETEGDVQNNPHV
B240 MELITACALLLIAVHIAIVISCPEPLAAVALVDVPLLYLCRYLLIIRRGVIIVSGAGAGGGRLRRFRLGAAMAIIYMAMSTLFFLRIAPLTPWWGALAAWVMILLIVEAIFAFFFPYRCCFNETEGDVQNNPHV
B241 MELITACALLLIAVHIAIVISCPEPLAAVALVDVPLLYLCRYLLIIRRGVIIVSGAGAGGGRLRRFRLGAAMAIIYMAMSTLFFLRIAPLTPWWGALAAWVMILLIVEAIFAFFFPYRCCFNETEGDVQNNPHV
B242 MELITACALLLIAVHIAIVISCPEPLAAVALVDVPLLYLCRYLLIIRRGVIIVSGGGGGGRLRRFRLGAAMAIIYMAMSTLFFLRIAPLTPWWGALAAWVMILLIVEAIFAFFFPYRCCFNETEGDVQNNPHV
B243 MELITACALLLIAVHIAIVISCPEPLAAVAFVNVPLLYLCRYLLIIRRGVIIVSGGGGGGGRLRRFRLGAAMAVIYMAMSTLFFLRIAPLTPWWGALAAWVMILLIVEAIFAFFFPYRCCFNETEGDVQNNPHV
B244 MELITACALLLIAVHIAIVISCPEPLAAVALVDVPLLYLCRYLLIIRRGVIIVSGAGAGGGRLRRFRLGAAMAIIYMAMSTLFFLRIAPLTPWWGALAAWVMILLIVEAIFAFFFPYRCCFNETEGDVQNNPHV
B245 MELITACALLLIAVHIAIVISCPEPLAAVALVDVPLLYLCRYLLIIRRGVIIVSGAGGGGGRLRRFRLGAAMAIIYMAMSTLFFLRIAPLTPWWGALAAWVMILLIVEAIFAFFFPYRCCFNETEGDVQNNPHV
B246 MELITACALLLIAVHIAIVISCPEPLAAVALVDVPLLYLCRYLLIIRRGVIIVSGAGGAAAAGCAASASGQPWRSSTWPCRPSSSSALLLSRHGGAPSPPGS*SCSSSRPSSPSSSPTAAASTKPKAMSRIIPM
B247 MELITACALLLIAVHIAIVISCPEPLAAVALVDVPLLYLYRYLLIIRRGVIIVSGAGGGGGRLRRFRLGAAMAIIYMAMSTLFFLRIAPLTPWWGALAAWVMILLIVEAIFAFFFPYRCCFNETEGDVQNNPHV
B248 MELITACALLLIAVHIAIVISCPEPLAAVALVDVPLLYLYRYLLIIRRGVIIVSGAGGGGGRLRRFRLGAAMAIIYMAMSTLFFLRIAPLTPWWGALAAWVMILLIVEAIFAFFFPYRCCFNETEGDVQNNPHV
B249 MELITACALLLIAVHIAIVISCPEPLAAVALVDVPLLYLCRYLLIIRRGVIIVSGAGAGGGRLRRFRLGAAMAIIYMAMSTLFFLRIAPLTPWWGALAAWVMILLIVEAIFAFFFPYRCCFNETEGDVQNNPHV
B250 MELITACALLLIAVHIAIVISCPEPLAAVALVDVPLLYLCRYLLIIRRGVIIVSGAGAGGGRLRRFRLGAAMAIIYMAMSTLFFLRIAPLTPWWGALAAWVMILLIVEAIFAFFFPYRCCFNETEGDVQNNPHV
B252 MELITACALLLIAVHIAIVISCPEPLAAVALVDVPLLYLCRYLLIIRRGVIIVSGAGAGGGRLRRFRLGAAMAIIYMAMSTLFFLRIAPLTPWWGALAAWVMILLIVEAIFAFFFPYRCCFNETEGDVQNNPHV
B253 MELITACALLLIAVHIAIVISCPEPLAAVALVDVPLLYLCRYLLIIRRGVIIVSGAGAGGGRLRRFRLGAAMAIIYMAMSTLFFLRIAPLTPWWGALAAWVMILLIVEAIFAFFFPYRCCFNETEGDVQNNPHV

| | |
|---|---|
| B254 | MELITACALLLIAVHIAIVISCPEPLAAVALVDVPLLYLCRYLLIIRRGVIIVSGAGAGGGRLRRFRLGAAMAIIYMAMSTLFFLRIAPLTPWWGALAAWVMILLIVEAIFAFFFPYRCCFNETEGDVQNNPHV |
| B255 | MELITACALLLIAVHIAIVISCPEPLAAVALVDVPLLYLCRYLLIIRRGVIIVSGGGAGGGRLRRFRLGAAMAIIYMAMSTLFFLRIAPLTPWWGALAAWVMILLIVEAIFAFFFPYRCCFNETEGDVQNNPHV |
| B258 | MELITACALLLIAVHIAIVISCPEPLAAVALVDVPLLYLCRYLLIIRRGVIIVSGAGAGGGRLRRFRLGAAMAIIYMAMSTLFFLRIAPLTPWWGALAAWVMILLIVEAIFAFFFPYRCCFNETEGDVQNNPHV |
| B259 | MELITACALLLIAVHIAIVISCPEPLAAVALVDVPLLYLYRYLLIIRRGVIIVSGAGGGGGRLRRFRLGAAMAIIYMAMSTLFFLRIAPLTPWWGALAAWVMILLIVEAIFAFFFPYRCCFNETEGDVQNNPHV |
| B260 | MELITACALLLIAVHIAIVISCPEPLAAVALVDVPLLYLYRYLLIIRRGVIIVSGAGGGGGRLRRFRLGAAMAIIYMAMSTLFFLRIAPLTPWWGALAAWVMILLIVEAIFAFFFPYRCCFNETEGDVQNNPHV |
| B261 | MELITACALLLIAVHIAIVISCPEPLAAVALVDVPLLYLCRYLLIIRRGVIIVSGGGGGGRLRRFRLGAAMAIIYMAMSTLFFLRIAPLTPWWGALAAWVMILLIVEAIFAFFFPYRCCFNETEGDVQNNPHV |
| B263 | MELITACALLLIAVHIAIVISCPEPLAAVALVDVPLLYLCRYLLIIRRGVIIVSGAGAGGGRLRRFRLGAAMAIIYMAMSTLFFLRIAPLTPWWGALAAWVMILLIVEAIFAFFFPYRCCFNETEGDVQNNPHV |
| B264 | MELITACALLLIAVHIAIVISCPEPLAAVALVDVPLLYLYRYLLIIRRGVIIVSGAGGGGGRLRRFRLGAAMAIIYMAMSTLFFLRIAPLTPWWGALAAWVMILLIVEAIFAFFFPYRCCFNETEGDVQNNPHV |
| B265 | MELITACALLLIAVHIAIVISCPEPLAAVALVDVPLLYLCRYLLIIRRGVIIVSGAGGAAAAGCAASASGQPWRSSTWPCRPSSSSALLLSRHGGAPSPPGS*SCSSSRPSSPSSSPTAAASTKPKAMSRIIPM |
| B266 | MELITACALLLIAVHIAIVISCPEPLAAVALVDVPLLYLYRYLLIIRRGVIIVSGAGGGGGRLRRFRLGAAMAIIYMAMSTLFFLRIAPLTPWWGALAAWVMILLIVEAIFAFFFPYRCCFNETEGDVQNNPHV |
| B267 | MELITACALLLIAVHIAIVISCPEPLAAVALVDVPLLYLCRYLLIIRRGVIIVSGGGGGGRLRRFRLGAAMAIIYMAMSTLFFLRIAPLTPWWGALAAWVMILLIVEAIFAFFFPYRCCFNETEGDVQNNPHV |
| B268 | MELITACALLLIAVHIAIVISCPEPLAAVALVDVPLLYLYRYLLIIRRGVIIVSGAGGGGGRLRRFRLGAAMAIIYMAMSTLFFLRIAPLTPWWGALAAWVMILLIVEAIFAFFFPYRCCFNETEGDVQNNPHV |
| B269 | MELITACALLLIAVHIAIVISCPEPLAAVALVDVPLLYLCRYLLIIRRGVIIVSGAGAGGGRLRRFRLGAAMAIIYMAMSTLFFLRIAPLTPWWGALAAWVMILLIVEAIFAFFFPYRCCFNETEGDVQNNPHV |
| CX2 | MELITACALLLIAVHIAIVISCPEPLAAVALVDVPLLYLCRYLLIIRRGVIIVSGAGGAAAAGCAASASGQPWRSSTWPCRPSSSSALLLSRHGGAPSPPGS*SCSSSRPSSPSSSPTAAASTKPKAMSRIIPM |
| CX3 | MELITACALLLIAVHIAIVISCPEPLAAVALVDVPLLYLCRYLLIIRRGVIIVSGGGGGGRLRRFRLGAAMAIIYMAMSTLFFLRIAPLTPWWGALAAWVMILLIVEAIFAFFFPYRCCFNETEGDVQNNPHV |
| CX4 | MELITACALLLIAVHIAIVISCPEPLAAVALVDVPLLYLCRYLLIIRRGVIIVSGAGAGGGRLRRFRLGAAMAIIYMAMSTLFFLRIAPLTPWWGALAAWVMILLIVEAIFAFFFPYRCCFNETEGDVQNNPHV |
| CX5 | MELITACALLLIAVHIAIVISCPEPLAAVALVDVPLLYLCRYLLIIRRGVIIVSGGGGGGRLRRFRLGAAMAIIYMAMSTLFFLRIAPLTPWWGALAAWVMILLIVEAIFAFFFPYRCCFNETEGDVQNNPHV |
| CX6 | MELITACALLLIAVHIAIVISCPEPLAAVALVDVPLLYLCRYLLIIRRGVIIVSGGGGGGRLRRFRLGAAMAIIYMAMSTLFFLRIAPLTPWWGALAAWVMILLIVEAIFAFFFPYRCCFNETEGDVQNNPHV |
| CX8 | MELITACALLLIAVHIAIVISCPEPLAAVALVDVPLLYLCRYLLIIRRGVIIVSGGGGGGRLRRFRLGAAMAIIYMAMSTLFFLRIAPLTPWWGALAAWVMILLIVEAIFAFFFPYRCCFNETEGDVQNNPHV |
| CX9 | MELITACALLLIAVHIAIVISCPEPLAAVALVDVPLLYLCRYLLIIRRGVIIVSGAGAGGGRLRRFRLGAAMAIIYMAMSTLFFLRIAPLTPWWGALAAWVMILLIVEAIFAFFFPYRCCFNETEGDVQNNPHV |
| CX10 | MELITACALLLIAVHIAIVISCPEPLAAVALVDVPLLYLYRYLLIIRRGVIIVSGAGGGGGRLRRFRLGAAMAIIYMAMSTLFFLRIAPLTPWWGALAAWVMILLIVEAIFAFFFPYRCCFNETEGDVQNNPHV |
| CX11 | MELITACALLLIAVHIAIVISCPEPLAAVALVDVPLLYLCRYLLIIRRGVIIVSGAGAGGGRLRRFRLGAAMAIIYMAMSTLFFLRIAPLTPWWGALAAWVMILLIVEAIFAFFFPYRCCFNETEGDVQNNPHV |
| CX12 | MELITACALLLIAVHIAIVISCPEPLAAVALVDVPLLYLCRYLLIIRRGVIIVSGAGAGGGRLRRFRLGAAMAIIYMAMSTLFFLRIAPLTPWWGALAAWVMILLIVEAIFAFFFPYRCCFNETEGDVQNNPHV |
| CX13 | MELITACALLLIAVHIAIVISCPEPLAAVALVDVPLLYLCRYLLIIRRGVIIVSGGGGGGRLRRFRLGAAMAIIYMAMSTLFFLRIAPLTPWWGALAAWVMILLIVEAIFAFFFPYRCCFNETEGDVQNNPHV |
| CX14 | MELITACALLLIAVHIAIVISCPEPLAAVALVDVPLLYLCRYLLIIRRGVIIVSGAGAGGGRLRRFRLGAAMAIIYMAMSTLFFLRIAPLTPWWGALAAWVMILLIVEAIFAFFFPYRCCFNETEGDVQNNPHV |
| CX15 | MELITACALLLIAVHIAIVISCPEPLAAVALVDVPLLYLCRYLLIIRRGVIIVSGGGGGGRLRRFRLGAAMAIIYMAMSTLFFLRIAPLTPWWGALAAWVMILLIVEAIFAFFFPYRCCFNETEGDVQNNPHV |
| CX16 | MELITACALLLIAVHIAIVISCPEPLAAVALVDVPLLYLCRYLLIIRRGVIIVSGAGAGGGRLRRFRLGAAMAIIYMAMSTLFFLRIAPLTPWWGALAAWVMILLIVEAIFAFFFPYRCCFNETEGDVQNNPHV |
| CX17 | MELITACALLLIAVHIAIVISCPEPLAAVALVDVPLLYLCRYLLIIRRGVIIVSGGGGGGRLRRFRLGAAMAIIYMAMSTLFFLRIAPLTPWWGALAAWVMILLIVEAIFAFFFPYRCCFNETEGDVQNNPHV |
| CX18 | MELITACALLLIAVHIAIVISCPEPLAAVALVDVPLLYLCRYLLIIRRGVIIVSGAGAGGGRLRRFRLGAAMAIIYMAMSTLFFLRIAPLTPWWGALAAWVMILLIVEAIFAFFFPYRCCFNETEGDVQNNPHV |
| CX19 | MELITACALLLIAVHIAIVISCPEPLAAVALVDVPLLYLCRYLLIIRRGVIIVSGAGAGGGRLRRFRLGAAMAIIYMAMSTLFFLRIAPLTPWWGALAAWVMILLIVEAIFAFFFPYRCCFNETEGDVQNNPHV |
| CX20 | MELITACALLLIAVHIAIVISCPEPLAAVALVDVPLLYLCRYLLIIRRGVIIVSGGGGGGRLRRFRLGAAMAIIYMAMSTLFFLRIAPLTPWWGALAAWVMILLIVEAIFAFFFPYRCCFNETEGDVQNNPHV |
| CX21 | MELITACALLLIAVHIAIVISCPEPLAAVALVDVPLLYLCRYLLIIRRGVIIVSGGGGGGRLRRFRLGAAMAIIYMAMSTLFFLRIAPLTPWWGALAAWVMILLIVEAIFAFFFPYRCCFNETEGDVQNNPHV |
| CX22 | MELITACALLLIAVHIAIVISCPEPLAAVALVDVPLLYLCRYLLIIRRGVIIVSGAGAGGGRLRRFRLGAAMAIIYMAMSTLFFLRIAPLTPWWGALAAWVMILLIVEAIFAFFFPYRCCFNETEGDVQNNPHV |
| CX23 | MELITACALLLIAVHIAIVISCPEPLAAVALVDVPLLYLCRYLLIIRRGVIIVSGAGGGGRLRRFRLGAAMAIIYMAMSTLFFLRIAPLTPWWGALAAWVMILLIVEAIFAFFFPYRCCFNETEGDVQNNPHV |
| CX24 | MELITACALLLIAVHIAIVISCPEPLAAVALVDVPLLYLCRYLLIIRRGVIIVSGGGGGGRLRRFRLGAAMAIIYMAMSTLFFLRIAPLTPWWGALAAWVMILLIVEAIFAFFFPYRCCFNETEGDVQNNPHV |
| CX25 | MELITACALLLIAVHIAIVISCPEPLAAVALVDVPLLYLCRYLLIIRRGVIIVSGGGGGGRLRRFRLGAAMAIIYMAMSTLFFLRIAPLTPWWGALAAWVMILLIVEAIFAFFFPYRCCFNETEGDVQNNPHV |
| CX26 | MELITACALLLIAVHIAIVISCPEPLAAVALVDVPLLYLCRYLLIIRRGVIIVSGGGGGGRLRRFRLGAAMAIIYMAMSTLFFLRIAPLTPWWGALAAWVMILLIVEAIFAFFFPYRCCFNETEGDVQNNPHV |
| CX27 | MELITACALLLIAVHIAIVISCPEPLAAVALVDVPLLYLCRYLLIIRRGVIIVSGAGAGGGRLRRFRLGAAMAIIYMAMSTLFFLRIAPLTPWWGALAAWVMILLIVEAIFAFFFPYRCCFNETEGDVQNNPHV |
| CX28 | MELITACALLLIAVHIAIVISCPEPLAAVALVDVPLLYLCRYLLIIRRGVIIVSGAGAGGGRLRRFRLGAAMAIIYMAMSTLFFLRIAPLTPWWGALAAWVMILLIVEAIFAFFFPYRCCFNETEGDVQNNPHV |
| CX29 | MELITACALLLIAVHIAIVISCPEPLAAVAFVNVPLLYLCRYLLIIRRGVIIVSGGGGGGGRLRRFRLGAAMAVIYMAMSTLFFLRIAPLTPWWGALAAWVMILLIVEAIFAFFFPYRCCFNETEGDVQNNPHV |
| CX30 | MELITACALLLIAVHIAIVISCPEPLAAVALVDVPLLYLCRYLLIIRRGVIIVSGAGAGGGRLRRFRLGAAMAIIYMAMSTLFFLRIAPLTPWWGALAAWVMILLIVEAIFAFFFPYRCCFNETEGDVQNNPHV |
| CX31 | MELITACALLLIAVHIAIVISCPEPLAAVALVDVPLLYLCRYLLIIRRGVIIVSGGGGGGRLRRFRLGAAMAIIYMAMSTLFFLRIAPLTPWWGALAAWVMILLIVEAIFAFFFPYRCCFNETEGDVQNNPHV |
| CX32 | MELITACALLLIAVHIAIVISCPEPLAAVALVDVPLLYLYRYLLIIRRGVIIVSGAGAGGGRLRRFRLGAAMAIIYMAMSTLFFLRIAPLTPWWGALAAWVMILLIVEAIFAFFFPYRCCFNETEGDVQNNPHV |
| CX33 | MELITACALLLIAVHIAIVISCPEPLAAVALVDVPLLYLCRYLLIIRRGVIIVSGAGAGGGRLRRFRLGAAMAIIYMAMSTLFFLRIAPLTPWWGALAAWVMILLIVEAIFAFFFPYRCCFNETEGDVQNNPHV |
| CX34 | MELITACALLLIAVHIAIVISCPEPLAAVALVDVPLLYLCRYLLIIRRGVIIVSGGGAGGGRLRRFRLGAAMAIIYMAMSTLFFLRIAPLTPWWGALAAWVMILLIVEAIFAFFFPYRCCFNETEGDVQNNPHV |
| CX35 | MELITACALLLIAVHIAIVISCPEPLAAVALVDVPLLYLCRYLLIIRRGVIIVSGGGGGGRLRRFRLGAAMAIIYMAMSTLFFLRIAPLTPWWGALAAWVMILLIVEAIFAFFFPYRCCFNETEGDVQNNPHV |
| CX37 | MELITACALLLIAVHIAIVISCPEPLAAVALVDVPLLYLCRYLLIIRRGVIIVSGAGGGGRLRRFRLGAAMAIIYMAMSTLFFLRIAPLTPWWGALAAWVMILLIVEAIFAFFFPYRCCFNETEGDVQNNPHV |
| CX42 | MELITACALLLIAVHIAIVISCPEPLAAVALVDVPLLYLCRYLLIIRRGVIIVSGAGAGGGRLRRFRLGAAMAIIYMAMSTLFFLRIAPLTPWWGALAAWVMILLIVEAIFAFFFPYRCCFNETEGDVQNNPHV |
| CX43 | MELITACALLLIAVHIAIVISCPEPLAAVALVDVPLLYLCRYLLIIRRGVIIVSGAGAGGGRLRRFRLGAAMAIIYMAMSTLFFLRIAPLTPWWGALAAWVMILLIVEAIFAFFFPYRCCFNETEGDVQNNPHV |
| CX44 | MELITACALLLIAVHIAIVISCPEPLAAVALVDVPLLYLCRYLLIIRRGVIIVSGAGGGGRLRRFRLGAAMAIIYMAMSTLFFLRIAPLTPWWGALAAWVMILLIVEAIFAFFFPYRCCFNETEGDVQNNPHV |
| CX45 | MELITACALLLIAVHIAIVISCPEPLAAVALVDVPLLYLCRYLLIIRRGVIIVSGGGGGGRLRRFRLGAAMAIIYMAMSTLFFLRIAPLTPWWGALAAWVMILLIVEAIFAFFFPYRCCFNETEGDVQNNPHV |
| CX46 | MELITACALLLIAVHIAIVISCPEPLAAVALVDVPLLYLCRYLLIIRRGVIIVSGAGAGGGRLRRFRLGAAMAIIYMAMSTLFFLRIAPLTPWWGALAAWVMILLIVEAIFAFFFPYRCCFNETEGDVQNNPHV |
| CX47 | MELITACALLLIAVHIAIVISCPEPLAAVALVDVPLLYLCRYLLIIRRGVIIVSGAGAGGGRLRRFRLGAAMAIIYMAMSTLFFLRIAPLTPWWGALAAWVMILLIVEAIFAFFFPYRCCFNETEGDVQNNPHV |
| CX48 | MELITACALLLIAVHIAIVISCPEPLAAVALVDVPLLYLCRYLLIIRRGVIIVSGGGGGGRLRRFRLGAAMAIIYMAMSTLFFLRIAPLTPWWGALAAWVMILLIVEAIFAFFFPYRCCFNETEGDVQNNPHV |
| CX49 | MELITACALLLIAVHIAIVISCPEPLAAVALVDVPLLYLCRYLLIIRRGVIIVSGAGAGGGRLRRFRLGAAMAIIYMAMSTLFFLRIAPLTPWWGALAAWVMILLIVEAIFAFFFPYRCCFNETEGDVQNNPHV |
| CX50 | MELITACALLLIAVHIAIVISCPEPLAAVALVDVPLLYLCRYLLIIRRGVIIVSGAGAGGGRLRRFRLGAAMAIIYMAMSTLFFLRIAPLTPWWGALAAWVMILLIVEAIFAFFFPYRCCFNETEGDVQNNPHV |
| CX51 | MELITACALLLIAVHIAIVISCPEPLAAVALVDVPLLYLCRYLLIIRRGVIIVSGGGAGGGRLRRFRLGAAMAIIYMAMSTLFFLRIAPLTPWWGALAAWVMILLIVEAIFAFFFPYRCCFNETEGDVQNNPHV |
| CX52 | MELITACALLLIAVHIAIVISCPEPLAAVALVDVPLLYLCRYLLIIRRGVIIVSGGGGGGRLRRFRLGAAMAIIYMAMSTLFFLRIAPLTPWWGALAAWVMILLIVEAIFAFFFPYRCCFNETEGDVQNNPHV |
| CX53 | MELITACALLLIAVHIAIVISCPEPLAAVALVDVPLLYLCRYLLIIRRGVIIVSGGGGGGRLRRFRLGAAMAIIYMAMSTLFFLRIAPLTPWWGALAAWVMILLIVEAIFAFFFPYRCCFNETEGDVQNNPHV |
| CX54 | MELITACALLLIAVHIAIVISCPEPLAAVALVDVPLLYLYRYLLIIRRGVIIVSGAGGGGGRLRRFRLGAAMAIIYMAMSTLFFLRIAPLTPWWGALAAWVMILLIVEAIFAFFFPYRCCFNETEGDVQNNPHV |
| CX55 | MELITACALLLIAVHIAIVISCPEPLAAVALVDVPLLYLCRYLLIIRRGVIIVSGAGGGGGRLRRFRLGAAMAIIYMAMSTLFFLRIAPLTPWWGALAAWVMILLIVEAIFAFFFPYRCCFNETEGDVQNNPHV |
| CX56 | MELITACALLLIAVHIAIVISCPEPLAAVALVDVPLLYLCRYLLIIRRGVIIVSGAGGGGGRLRRFRLGAAMAIIYMAMSTLFFLRIAPLTPWWGALAAWVMILLIVEAIFAFFFPYRCCFNETEGDVQNNPHV |
| CX57 | MELITACALLLIAVHIAIVISCPEPLAAVALVDVPLLYLCRYLLIIRRGVIIVSGGGGGGRLRRFRLGAAMAIIYMAMSTLFFLRIAPLTPWWGALAAWVMILLIVEAIFAFFFPYRCCFNETEGDVQNNPHV |
| CX58 | MELITACALLLIAVHIAIVISCPEPLAAVALVDVPLLYLCRYLLIIRRGVIIVSGAGAGGGRLRRFRLGAAMAIIYMAMSTLFFLRIAPLTPWWGALAAWVMILLIVEAIFAFFFPYRCCFNETEGDVQNNPHV |
| CX59 | MELITACALLLIAVHIAIVISCPEPLAAVALVDVPLLYLCRYLLIIRRGVIIVSGAGAGGGRLRRFRLGAAMAIIYMAMSTLFFLRIAPLTPWWGALAAWVMILLIVEAIFAFFFPYRCCFNETEGDVQNNPHV |
| CX60 | MELITACALLLIAVHIAIVISCPEPLAAVALVDVPLLYLCRYLLIIRRGVIIVSGGGGGGRLRRFRLGAAMAIIYMAMSTLFFLRIAPLTPWWGALAAWVMILLIVEAIFAFFFPYRCCFNETEGDVQNNPHV |
| CX61 | MELITACALLLIAVHIAIVISCPEPLAAVALVDVPLLYLCRYLLIIRRGVIIVSGGGGGGRLRRFRLGAAMAIIYMAMSTLFFLRIAPLTPWWGALAAWVMILLIVEAIFAFFFPYRCCFNETEGDVQNNPHV |
| CX63 | MELITACALLLIAVHIAIVISCPEPLAAVALVDVPLLYLCRYLLIIRRGVIIVSGAGAGGGRLRRFRLGAAMAIIYMAMSTLFFLRIAPLTPWWGALAAWVMILLIVEAIFAFFFPYRCCFNETEGDVQNNPHV |
| CX64 | MELITACALLLIAVHIAIVISCPEPLAAVALVDVPLLYLCRYLLIIRRGVIIVSGGGGGGRLRRFRLGAAMAIIYMAMSTLFFLRIAPLTPWWGALAAWVMILLIVEAIFAFFFPYRCCFNETEGDVQNNPHV |
| CX65 | MELITACALLLIAVHIAIVISCPEPLAAVALVDVPLLYLCRYLLIIRRGVIIVSGAGAGGGRLRRFRLGAAMAIIYMAMSTLFFLRIAPLTPWWGALAAWVMILLIVEAIFAFFFPYRCCFNETEGDVQNNPHV |
| CX66 | MELITACALLLIAVHIAIVISCPEPLAAVALVDVPLLYLCRYLLIIRRGVIIVSGAGAGGGRLRRFRLGAAMAIIYMAMSTLFFLRIAPLTPWWGALAAWVMILLIVEAIFAFFFPYRCCFNETEGDVQNNPHV |
| CX67 | MELITACALLLIAVHIAIVISCPEPLAAVALVDVPLLYLCRYLLIIRRGVIIVSGAGAGGGRLRRFRLGAAMAIIYMAMSTLFFLRIAPLTPWWGALAAWVMILLIVEAIFAFFFPYRCCFNETEGDVQNNPHV |
| CX68 | MELITACALLLIAVHIAIVISCPEPLAAVALVDVPLLYLCRYLLIIRRGVIIVSGGGGGGRLRRFRLGAAMAIIYMAMSTLFFLRIAPLTPWWGALAAWVMILLIVEAIFAFFFPYRCCFNETEGDVQNNPHV |
| CX69 | MELITACALLLIAVHIAIVISCPEPLAAVALVDVPLLYLCRYLLIIRRGVIIVSGAGAGGGRLRRFRLGAAMAIIYMAMSTLFFLRIAPLTPWWGALAAWVMILLIVEAIFAFFFPYRCCFNETEGDVQNNPHV |
| CX70 | MELITACALLLIAVHIAIVISCPEPLAAVALVDVPLLYLCRYLLIIRRGVIIVSGAGGGGGGRLRRFRLGAAMAIIYMAMSTLFFLRIAPLTPWWGALAAWVMILLIVEAIFAFFFPYRCCFNETEGDVQNNPHV |
| CX71 | MELITACALLLIAVHIAIVISCPEPLAAVALVDVPLLYLCRYLLIIRRGVIIVSGGGGGGRLRRFRLGAAMAIIYMAMSTLFFLRIAPLTPWWGALAAWVMILLIVEAIFAFFFPYRCCFNETEGDVQNNPHV |
| CX72 | MELITACALLLIAVHIAIVISCPEPLAAVALVDVPLLYLCRYLLIIRRGVIIVSGAGAGGGRLRRFRLGAAMAIIYMAMSTLFFLRIAPLTPWWGALAAWVMILLIVEAIFAFFFPYRCCFNETEGDVQNNPHV |
| CX73 | MELITACALLLIAVHIAIVISCPEPLAAVALVDVPLLYLCRYLLIIRRGVIIVSGGGGGGRLRRFRLGAAMAIIYMAMSTLFFLRIAPLTPWWGALAAWVMILLIVEAIFAFFFPYRCCFNETEGDVQNNPHV |
| CX74 | MELITACALLLIAVHIAIVISCPEPLAAVALVDVPLLYLCRYLLIIRRGVIIVSGAGAGGGRLRRFRLGAAMAIIYMAMSTLFFLRIAPLTPWWGALAAWVMILLIVEAIFAFFFPYRCCFNETEGDVQNNPHV |
| CX75 | MELITACALLLIAVHIAIVISCPEPLAAVALVDVPLLYLCRYLLIIRRGVIIVSGAGGGGGRLRRFRLGAAMAIIYMAMSTLFFLRIAPLTPWWGALAAWVMILLIVEAIFAFFFPYRCCFNETEGDVQNNPHV |
| CX76 | MELITACALLLIAVHIAIVISCPEPLAAVALVDVPLLYLCRYLLIIRRGVIIVSGGGGGGRLRRFRLGAAMAIIYMAMSTLFFLRIAPLTPWWGALAAWVMILLIVEAIFAFFFPYRCCFNETEGDVQNNPHV |
| CX77 | MELITACALLLIAVHIAIVISCPEPLAAVALVDVPLLYLYRYLLIIRRGVIIVSGAGGGGGRLRRFRLGAAMAIIYMAMSTLFFLRIAPLTPWWGALAAWVMILLIVEAIFAFFFPYRCCFNETEGDVQNNPHV |
| CX78 | MELITACALLLIAVHIAIVISCPEPLAAVALVDVPLLYLCRYLLIIRRGVIIVSGAGAGGGRLRRFRLGAAMAIIYMAMSTLFFLRIAPLTPWWGALAAWVMILLIVEAIFAFFFPYRCCFNETEGDVQNNPHV |
| CX79 | MELITACALLLIAVHIAIVISCPEPLAAVALVDVPLLYLCRYLLIIRRGVIIVSGAGAGGGRLRRFRLGAAMAIIYMAMSTLFFLRIAPLTPWWGALAAWVMILLIVEAIFAFFFPYRCCFNETEGDVQNNPHV |
| CX80 | MELITACALLLIAVHIAIVISCPEPLAAVALVDVPLLYLCRYLLIIRRGVIIVSGAGAGGGRLRRFRLGAAMAIIYMAMSTLFFLRIAPLTPWWGALAAWVMILLIVEAIFAFFFPYRCCFNETEGDVQNNPHV |
| CX82 | MELITACALLLIAVHIAIVISCPEPLAAVALVDVPLLYLCRYLLIIRRGVIIVSGGGGGGRLRRFRLGAAMAIIYMAMSTLFFLRIAPLTPWWGALAAWVMILLIVEAIFAFFFPYRCCFNETEGDVQNNPHV |
| CX83 | MELITACALLLIAVHIAIVISCPEPLAAVALVDVPLLYLCRYLLIIRRGVIIVSGGGGGGRLRRFRLGAAMAIIYMAMSTLFFLRIAPLTPWWGALAAWVMILLIVEAIFAFFFPYRCCFNETEGDVQNNPHV |
| CX84 | MELITACALLLIAVHIAIVISCPEPLAAVALVDVPLLYLCRYLLIIRRGVIIVSGGGGGGRLRRFRLGAAMAIIYMAMSTLFFLRIAPLTPWWGALAAWVMILLIVEAIFAFFFPYRCCFNETEGDVQNNPHV |
| CX85 | MELITACALLLIAVHIAIVISCPEPLAAVALVDVPLLYLCRYLLIIRRGVIIVSGGGGGGRLRRFRLGAAMAIIYMAMSTLFFLRIAPLTPWWGALAAWVMILLIVEAIFAFFFPYRCCFNETEGDVQNNPHV |
| CX86 | MELITACALLLIAVHIAIVISCPEPLAAVALVDVPLLYLCRYLLIIRRGVIIVSGAGAGGGRLRRFRLGAAMAIIYMAMSTLFFLRIAPLTPWWGALAAWVMILLIVEAIFAFFFPYRCCFNETEGDVQNNPHV |
| CX87 | MELITACALLLIAVHIAIVISCPEPLAAVALVDVPLLYLCRYLLIIRRGVIIVSGAGAGGGRLRRFRLGAAMAIIYMAMSTLFFLRIAPLTPWWGALAAWVMILLIVEAIFAFFFPYRCCFNETEGDVQNNPHV |
| CX88 | MELITACALLLIAVHIAIVISCPEPLAAVALVDVPLLYLCRYLLIIRRGVIIVSGGGGGGRLRRFRLGAAMAIIYMAMSTLFFLRIAPLTPWWGALAAWVMILLIVEAIFAFFFPYRCCFNETEGDVQNNPHV |
| CX89 | MELITACALLLIAVHIAIVISCPEPLAAVALVDVPLLYLCRYLLIIRRGVIIVSGGGGGGRLRRFRLGAAMAIIYMAMSTLFFLRIAPLTPWWGALAAWVMILLIVEAIFAFFFPYRCCFNETEGDVQNNPHV |
| CX90 | MELITACALLLIAVHIAIVISCPEPLAAVALVDVPLLYLCRYLLIIRRGVIIVSGGGGGGRLRRFRLGAAMAIIYMAMSTLFFLRIAPLTPWWGALAAWVMILLIVEAIFAFFFPYRCCFNETEGDVQNNPHV |
| CX91 | MELITACALLLIAVHIAIVISCPEPLAAVALVDVPLLYLCRYLLIIRRGVIIVSGAGAGGGRLRRFRLGAAMAIIYMAMSTLFFLRIAPLTPWWGALAAWVMILLIVEAIFAFFFPYRCCFNETEGDVQNNPHV |
| CX92 | MELITACALLLIAVHIAIVISCPEPLAAVALVDVPLLYLCRYLLIIRRGVIIVSGGGGGGRLRRFRLGAAMAIIYMAMSTLFFLRIAPLTPWWGALAAWVMILLIVEAIFAFFFPYRCCFNETEGDVQNNPHV |
| CX93 | MELITACALLLIAVHIAIVISCPEPLAAVALVDVPLLYLCRYLLIIRRGVIIVSGGGGGGRLRRFRLGAAMAIIYMAMSTLFFLRIAPLTPWWGALAAWVMILLIVEAIFAFFFPYRCCFNETEGDVQNNPHV |
| CX94 | MELITACALLLIAVHIAIVISCPEPLAAVALVDVPLLYLCRYLLIIRRGVIIVSGAGGAAAAGCAASASGQPWRSSTWPCRPSSSSALLLSRHGGAPSPPGS*SCSSSRPSSPSSSPTAAASTKPKAMSRIIPM |
| CX96 | MELITACALLLIAVHIAIVISCPEPLAAVALVDVPLLYLCRYLLIIRRGVIIVSGAGAGGGRLRRFRLGAAMAIIYMAMSTLFFLRIAPLTPWWGALAAWVMILLIVEAIFAFFFPYRCCFNETEGDVQNNPHV |
| CX97 | MELITACALLLIAVHIAIVISCPEPLAAVALVDVPLLYLCRYLLIIRRGVIIVSGAGAGGGRLRRFRLGAAMAIIYMAMSTLFFLRIAPLTPWWGALAAWVMILLIVEAIFAFFFPYRCCFNETEGDVQNNPHV |
| CX98 | MELITACALLLIAVHIAIVISCPEPLAAVALVDVPLLYLCRYLLIIRRGVIIVSGAGGAAAAGCAASASGQPWRSSTWPCRPSSSSALLLSRHGGAPSPPGS*SCSSSRPSSPSSSPTAAASTKPKAMSRIIPM |
| CX99 | MELITACALLLIAVHIAIVISCPEPLAAVALVDVPLLYLCRYLLIIRRGVIIVSGGGGGGRLRRFRLGAAMAIIYMAMSTLFFLRIAPLTPWWGALAAWVMILLIVEAIFAFFFPYRCCFNETEGDVQNNPHV |
| CX100 | MELITACALLLIAVHIAIVISCPEPLAAVALVDVPLLYLCRYLLIIRRGVIIVSGGGGGGRLRRFRLGAAMAIIYMAMSTLFFLRIAPLTPWWGALAAWVMILLIVEAIFAFFFPYRCCFNETEGDVQNNPHV |
| CX101 | MELITACALLLIAVHIAIVISCPEPLAAVALVDVPLLYLCRYLLIIRRGVIIVSGGGGGGRLRRFRLGAAMAIIYMAMSTLFFLRIAPLTPWWGALAAWVMILLIVEAIFAFFFPYRCCFNETEGDVQNNPHV |
| CX102 | MELITACALLLIAVHIAIVISCPEPLAAVALVDVPLLYLCRYLLIIRRGVIIVSGGGGGGRLRRFRLGAAMAIIYMAMSTLFFLRIAPLTPWWGALAAWVMILLIVEAIFAFFFPYRCCFNETEGDVQNNPHV |
| CX103 | MELITACALLLIAVHIAIVISCPEPLAAVALVDVPLLYLCRYLLIIRRGVIIVSGAGAGGGRLRRFRLGAAMAIIYMAMSTLFFLRIAPLTPWWGALAAWVMILLIVEAIFAFFFPYRCCFNETEGDVQNNPHV |
| CX104 | MELITACALLLIAVHIAIVISCPEPLAAVALVDVPLLYLCRYLLIIRRGVIIVSGAGAGGGRLRRFRLGAAMAIIYMAMSTLFFLRIAPLTPWWGALAAWVMILLIVEAIFAFFFPYRCCFNETEGDVQNNPHV |
| CX106 | MELITACALLLIAVHIAIVISCPEPLAAVALVDVPLLYLYRYLLIIRRGVIIVSGAGGGGGRLRRFRLGAAMAIIYMAMSTLFFLRIAPLTPWWGALAAWVMILLIVEAIFAFFFPYRCCFNETEGDVQNNPHV |
| CX107 | MELITACALLLIAVHIAIVISCPEPLAAVALVDVPLLYLCRYLLIIRRGVIIVSGGGGGGRLRRFRLGAAMAIIYMAMSTLFFLRIAPLTPWWGALAAWVMILLIVEAIFAFFFPYRCCFNETEGDVQNNPHV |
| CX108 | MELITACALLLIAVHIAIVISCPEPLAAVAFVNVPLLYLCRYLLIIRRGVIIVSGGGGGGGRLRRFRLGAAMAVIYMAMSTLFFLRIAPLTPWWGALAAWVMILLIVEAIFAFFFPYRCCFNETEGDVQNNPHV |
| CX109 | MELITACALLLIAVHIAIVISCPEPLAAVALVDVPLLYLCRYLLIIRRGVIIVSGAGAGGGRLRRFRLGAAMAIIYMAMSTLFFLRIAPLTPWWGALAAWVMILLIVEAIFAFFFPYRCCFNETEGDVQNNPHV |
| CX110 | MELITACALLLIAVHIAIVISCPEPLAAVALVDVPLLYLCRYLLIIRRGVIIVSGAGAGGGRLRRFRLGAAMAIIYMAMSTLFFLRIAPLTPWWGALAAWVMILLIVEAIFAFFFPYRCCFNETEGDVQNNPHV |
| CX111 | MELITACALLLIAVHIAIVISCPEPLAAVALVDVPLLYLCRYLLIIRRGVIIVSGAGAGGGRLRRFRLGAAMAIIYMAMSTLFFLRIAPLTPWWGALAAWVMILLIVEAIFAFFFPYRCCFNETEGDVQNNPHV |
| CX112 | MELITACALLLIAVHIAIVISCPEPLAAVAFVNVPLLYLCRYLLIIRRGVIIVSGGGGGGGRLRRFRLGAAMAVIYMAMSTLFFLRIAPLTPWWGALAAWVMILLIVEAIFAFFFPYRCCFNETEGDVQNNPHV |
| CX113 | MELITACALLLIAVHIAIVISCPEPLAAVALVDVPLLYLCRYLLIIRRGVIIVSGAGAGGGRLRRFRLGAAMAIIYMAMSTLFFLRIAPLTPWWGALAAWVMILLIVEAIFAFFFPYRCCFNETEGDVQNNPHV |
| CX114 | MELITACALLLIAVHIAIVISCPEPLAAVALVDVPLLYLCRYLLIIRRGVIIVSGAGAGGGRLRRFRLGAAMAIIYMAMSTLFFLRIAPLTPWWGALAAWVMILLIVEAIFAFFFPYRCCFNETEGDVQNNPHV |
| CX115 | MELITACALLLIAVHIAIVISCPEPLAAVALVDVPLLYLCRYLLIIRRGVIIVSGGGGGGRLRRFRLGAAMAIIYMAMSTLFFLRIAPLTPWWGALAAWVMILLIVEAIFAFFFPYRCCFNETEGDVQNNPHV |
| CX116 | MELITACALLLIAVHIAIVISCPEPLAAVALVDVPLLYLCRYLLIIRRGVIIVSGAGAGGGRLRRFRLGAAMAIIYMAMSTLFFLRIAPLTPWWGALAAWVMILLIVEAIFAFFFPYRCCFNETEGDVQNNPHV |
| CX117 | MELITACALLLIAVHIAIVISCPEPLAAVALVDVPLLYLCRYLLIIRRGVIIVSGGGGGGRLRRFRLGAAMAIIYMAMSTLFFLRIAPLTPWWGALAAWVMILLIVEAIFAFFFPYRCCFNETEGDVQNNPHV |
| CX118 | MELITACALLLIAVHIAIVISCPEPLAAVALVDVPLLYLCRYLLIIRRGVIIVSGGGGGGRLRRFRLGAAMAIIYMAMSTLFFLRIAPLTPWWGALAAWVMILLIVEAIFAFFFPYRCCFNETEGDVQNNPHV |
| CX119 | MELITACALLLIAVHIAIVISCPEPLAAVALVDVPLLYLCRYLLIIRRGVIIVSGGGGGGRLRRFRLGAAMAIIYMAMSTLFFLRIAPLTPWWGALAAWVMILLIVEAIFAFFFPYRCCFNETEGDVQNNPHV |
| CX120 | MELITACALLLIAVHIAIVISCPEPLAAVALVDVPLLYLCRYLLIIRRGVIIVSGGGGGGRLRRFRLGAAMAIIYMAMSTLFFLRIAPLTPWWGALAAWVMILLIVEAIFAFFFPYRCCFNETEGDVQNNPHV |
| CX121 | MELITACALLLIAVHIAIVISCPEPLAAVALVDVPLLYLCRYLLIIRRGVIIVSGGGGGGRLRRFRLGAAMAIIYMAMSTLFFLRIAPLTPWWGALAAWVMILLIVEAIFAFFFPYRCCFNETEGDVQNNPHV |

| | |
|---|---|
| CX122 | MELITACALLLIAVHIAIVISCPEPLAAVALVDVPLLYLCRYLLIIRRGVIIVSGAGAGGGRLRRFRLGAAMAIIYMAMSTLFFLRIAPLTPWWGALAAWVMILLIVEAIFAFFFPYRCCFNETEGDVQNNPHV |
| CX123 | MELITACALLLIAVHIAIVISCPEPLAAVALVDVPLLYLYRYLLIIRRGVIIVSGAGGGGGRLRRFRLGAAMAIIYMAMSTLFFLRIAPLTPWWGALAAWVMILLIVEAIFAFFFPYRCCFNETEGDVQNNPHV |
| CX124 | MELITACALLLIAVHIAIVISCPEPLAAVALVDVPLLYLCRYLLIIRRGVIIVSGGGGGGRLRRFRLGAAMAIIYMAMSTLFFLRIAPLTPWWGALAAWVMILLIVEAIFAFFFPYRCCFNETEGDVQNNPHV |
| CX125 | MELITACALLLIAVHIAIVISCPEPLAAVALVDVPLLYLCRYLLIIRRGVIIVSGGGGGGRLRRFRLGAAMAIIYMAMSTLFFLRIAPLTPWWGALAAWVMILLIVEAIFAFFFPYRCCFNETEGDVQNNPHV |
| CX126 | MELITACALLLIAVHIAIVISCPEPLAAVALVDVPLLYLCRYLLIIRRGVIIVSGAGAGGGRLRRFRLGAAMAIIYMAMSTLFFLRIAPLTPWWGALAAWVMILLIVEAIFAFFFPYRCCFNETEGDVQNNPHV |
| CX128 | MELITACALLLIAVHIAIVISCPEPLAAVALVDVPLLYLCRYLLIIRRGVIIVSGGGGGGRLRRFRLGAAMAIIYMAMSTLFFLRIAPLTPWWGALAAWVMILLIVEAIFAFFFPYRCCFNETEGDVQNNPHV |
| CX129 | MELITACALLLIAVHIAIVISCPEPLAAVALVDVPLLYLYRYLLIIRRGVIIVSGAGGGGGRLRRFRLGAAMAIIYMAMSTLFFLRIAPLTPWWGALAAWVMILLIVEAIFAFFFPYRCCFNETEGDVQNNPHV |
| CX130 | MELITACALLLIAVHIAIVISCPEPLAAVALVDVPLLYLCRYLLIIRRGVIIVSGGGGGGRLRRFRLGAAMAIIYMAMSTLFFLRIAPLTPWWGALAAWVMILLIVEAIFAFFFPYRCCFNETEGDVQNNPHV |
| CX131 | MELITACALLLIAVHIAIVISCPEPLAAVALVDVPLLYLCRYLLIIRRGVIIVSGGGGGGRLRRFRLGAAMAIIYMAMSTLFFLRIAPLTPWWGALAAWVMILLIVEAIFAFFFPYRCCFNETEGDVQNNPHV |
| CX132 | MELITACALLLIAVHIAIVISCPEPLAAVALVDVPLLYLCRYLLIIRRGVIIVSGAGAGGGRLRRFRLGAAMAIIYMAMSTLFFLRIAPLTPWWGALAAWVMILLIVEAIFAFFFPYRCCFNETEGDVQNNPHV |
| CX133 | MELITACALLLIAVHIAIVISCPEPLAAVALVDVPLLYLCRYLLIIRRGVIIVSGAGAGGGRLRRFRLGAAMAIIYMAMSTLFFLRIAPLTPWWGALAAWVMILLIVEAIFAFFFPYRCCFNETEGDVQNNPHV |
| CX134 | MELITACALLLIAVHIAIVISCPEPLAAVALVDVPLLYLCRYLLIIRRGVIIVSGAGAGGGRLRRFRLGAAMAIIYMAMSTLFFLRIAPLTPWWGALAAWVMILLIVEAIFAFFFPYRCCFNETEGDVQNNPHV |
| CX138 | MELITACALLLIAVHIAIVISCPEPLAAVALVDVPLLYLCRYLLIIRRGVIIVSGAGGGGGRLRRFRLGAAMAIIYMAMSTLFFLRIAPLTPWWGALAAWVMILLIVEAIFAFFFPYRCCFNETEGDVQNNPHV |
| CX139 | MELITACALLLIAVHIAIVISCPEPLAAVALVDVPLLYLCRYLLIIRRGVIIVSGAGAGGGRLRRFRLGAAMAIIYMAMSTLFFLRIAPLTPWWGALAAWVMILLIVEAIFAFFFPYRCCFNETEGDVQNNPHV |
| CX140 | MELITACALLLIAVHIAIVISCPEPLAAVALVDVPLLYLCRYLLIIRRGVIIVSGAGAGGGRLRRFRLGAAMAIIYMAMSTLFFLRIAPLTPWWGALAAWVMILLIVEAIFAFFFPYRCCFNETEGDVQNNPHV |
| CX141 | MELITACALLLIAVHIAIVISCPEPLAAVALVDVPLLYLCRYLLIIRRGVIIVSGGGGGGRLRRFRLGAAMAIIYMAMSTLFFLRIAPLTPWWGALAAWVMILLIVEAIFAFFFPYRCCFNETEGDVQNNPHV |
| CX142 | MELITACALLLIAVHIAIVISCPEPLAAVALVDVPLLYLYRYLLIIRRGVIIVSGAGGGGGRLRRFRLGAAMAIIYMAMSTLFFLRIAPLTPWWGALAAWVMILLIVEAIFAFFFPYRCCFNETEGDVQNNPHV |
| CX143 | MELITACALLLIAVHIAIVISCPEPLAAVALVDVPLLYLCRYLLIIRRGVIIVSGAGAGGGRLRRFRLGAAMAIIYMAMSTLFFLRIAPLTPWWGALAAWVMILLIVEAIFAFFFPYRCCFNETEGDVQNNPHV |
| CX144 | MELITACALLLIAVHIAIVISCPEPLAAVALVDVPLLYLCRYLLIIRRGVIIVSGAGAGGGRLRRFRLGAAMAIIYMAMSTLFFLRIAPLTPWWGALAAWVMILLIVEAIFAFFFPYRCCFNETEGDVQNNPHV |
| CX145 | MELITACALLLIAVHIAIVISCPEPLAAVALVDVPLLYLCRYLLIIRRGVIIVSGGGGGGRLRRFRLGAAMAIIYMAMSTLFFLRIAPLTPWWGALAAWVMILLIVEAIFAFFFPYRCCFNETEGDVQNNPHV |
| CX146 | MELITACALLLIAVHIAIVISCPEPLAAVALVDVPLLYLCRYLLIIRRGVIIVSGGGGGGRLRRFRLGAAMAIIYMAMSTLFFLRIAPLTPWWGALAAWVMILLIVEAIFAFFFPYRCCFNETEGDVQNNPHV |
| CX147 | MELITACALLLIAVHIAIVISCPEPLAAVALVDVPLLYLCRYLLIIRRGVIIVSGAGGAAAAGCAASASGQPWRSSTWPCRPSSSSALLLSRHGGAPSPPGS*SCSSSRPSSPSSSPTAAASTKPKAMSRIIPM |
| CX148 | MELITACALLLIAVHIAIVISCPEPLAAVALVDVPLLYLCRYLLIIRRGVIIVSGAGGGGGGRLRRFRLGAAMAIIYMAMSTLFFLRIAPLTPWWGALAAWVMILLIVEAIFAFFFPYRCCFNETEGDVQNNPHV |
| CX149 | MELITACALLLIAVHIAIVISCPEPLAAVALVDVPLLYLCRYLLIIRRGVIIVSGAGAGGGRLRRFRLGAAMAIIYMAMSTLFFLRIAPLTPWWGALAAWVMILLIVEAIFAFFFPYRCCFNETEGDVQNNPHV |
| CX150 | MELITACALLLIAVHIAIVISCPEPLAAVALVDVPLLYLCRYLLIIRRGVIIVSGGGGGGRLRRFRLGAAMAIIYMAMSTLFFLRIAPLTPWWGALAAWVMILLIVEAIFAFFFPYRCCFNETEGDVQNNPHV |
| CX151 | MELITACALLLIAVHIAIVISCPEPLAAVALVDVPLLYLYRYLLIIRRGVIIVSGAGGGGGRLRRFRLGAAMAIIYMAMSTLFFLRIAPLTPWWGALAAWVMILLIVEAIFAFFFPYRCCFNETEGDVQNNPHV |
| CX152 | MELITACALLLIAVHIAIVISCPEPLAAVALVDVPLLYLCRYLLIIRRGVIIVSGGGGGGRLRRFRLGAAMAIIYMAMSTLFFLRIAPLTPWWGALAAWVMILLIVEAIFAFFFPYRCCFNETEGDVQNNPHV |
| CX153 | MELITACALLLIAVHIAIVISCPEPLAAVALVDVPLLYLCRYLLIIRRGVIIVSGAGGAAAAGCAASASGQPWRSSTWPCRPSSSSALLLSRHGGAPSPPGS*SCSSSRPSSPSSSPTAAASTKPKAMSRIIPM |
| CX154 | MELITACALLLIAVHIAIVISCPEPLAAVALVDVPLLYLCRYLLIIRRGVIIVSGGGGGGRLRRFRLGAAMAIIYMAMSTLFFLRIAPLTPWWGALAAWVMILLIVEAIFAFFFPYRCCFNETEGDVQNNPHV |
| CX155 | MELITACALLLIAVHIAIVISCPEPLAAVALVDVPLLYLCRYLLIIRRGVIIVSGGGGGGRLRRFRLGAAMAIIYMAMSTLFFLRIAPLTPWWGALAAWVMILLIVEAIFAFFFPYRCCFNETEGDVQNNPHV |
| CX156 | MELITACALLLIAVHIAIVISCPEPLAAVALVDVPLLYLCRYLLIIRRGVIIVSGAGAGGGRLRRFRLGAAMAIIYMAMSTLFFLRIAPLTPWWGALAAWVMILLIVEAIFAFFFPYRCCFNETEGDVQNNPHV |
| CX157 | MELITACALLLIAVHIAIVISCPEPLAAVALVDVPLLYLYRYLLIIRRGVIIVSGAGGGGGRLRRFRLGAAMAIIYMAMSTLFFLRIAPLTPWWGALAAWVMILLIVEAIFAFFFPYRCCFNETEGDVQNNPHV |
| CX158 | MELITACALLLIAVHIAIVISCPEPLAAVALVDVPLLYLCRYLLIIRRGVIIVSGAGGAAAAGCAASASGQPWRSSTWPCRPSSSSALLLSRHGGAPSPPGS*SCSSSRPSSPSSSPTAAASTKPKAMSRIIPM |
| CX160 | MELITACALLLIAVHIAIVISCPEPLAAVALVDVPLLYLCRYLLIIRRGVIIVSGGGGGGRLRRFRLGAAMAIIYMAMSTLFFLRIAPLTPWWGALAAWVMILLIVEAIFAFFFPYRCCFNETEGDVQNNPHV |
| CX161 | MELITACALLLIAVHIAIVISCPEPLAAVALVDVPLLYLCRYLLIIRRGVIIVSGGGGGGRLRRFRLGAAMAIIYMAMSTLFFLRIAPLTPWWGALAAWVMILLIVEAIFAFFFPYRCCFNETEGDVQNNPHV |
| CX162 | MELITACALLLIAVHIAIVISCPEPLAAVALVDVPLLYLCRYLLIIRRGVIIVSGAGAGGGRLRRFRLGAAMAIIYMAMSTLFFLRIAPLTPWWGALAAWVMILLIVEAIFAFFFPYRCCFNETEGDVQNNPHV |
| CX165 | MELITACALLLIAVHIAIVISCPEPLAAVALVDVPLLYLCRYLLIIRRGVIIVSGAGAGGGRLRRFRLGAAMAIIYMAMSTLFFLRIAPLTPWWGALAAWVMILLIVEAIFAFFFPYRCCFNETEGDVQNNPHV |
| CX182 | MELITACALLLIAVHIAIVISCPEPLAAVALVDVPLLYLCRYLLIIRRGVIIVSGGGGGGRLRRFRLGAAMAIIYMAMSTLFFLRIAPLTPWWGALAAWVMILLIVEAIFAFFFPYRCCFNETEGDVQNNPHV |
| CX205 | MELITACALLLIAVHIAIVISCPEPLAAVALVDVPLLYLYRYLLIIRRGVIIVSGAGGGGGRLRRFRLGAAMAIIYMAMSTLFFLRIAPLTPWWGALAAWVMILLIVEAIFAFFFPYRCCFNETEGDVQNNPHV |
| CX206 | MELITACALLLIAVHIAIVISCPEPLAAVALVDVPLLYLCRYLLIIRRGVIIVSGGGGGGRLRRFRLGAAMAIIYMAMSTLFFLRIAPLTPWWGALAAWVMILLIVEAIFAFFFPYRCCFNETEGDVQNNPHV |
| CX207 | MELITACALLLIAVHIAIVISCPEPLAAVALVDVPLLYLCRYLLIIRRGVIIVSGGGGGGRLRRFRLGAAMAIIYMAMSTLFFLRIAPLTPWWGALAAWVMILLIVEAIFAFFFPYRCCFNETEGDVQNNPHV |
| CX210 | MELITACALLLIAVHIAIVISCPEPLAAVALVDVPLLYLCRYLLIIRRGVIIVSGGGGGGRLRRFRLGAAMAIIYMAMSTLFFLRIAPLTPWWGALAAWVMILLIVEAIFAFFFPYRCCFNETEGDVQNNPHV |
| CX211 | MELITACALLLIAVHIAIVISCPEPLAAVALVDVPLLYLCRYLLIIRRGVIIVSGAGAGGGRLRRFRLGAAMAIIYMAMSTLFFLRIAPLTPWWGALAAWVMILLIVEAIFAFFFPYRCCFNETEGDVQNNPHV |
| CX212 | MELITACALLLIAVHIAIVISCPEPLAAVALVDVPLLYLCRYLLIIRRGVIIVSGAGAGGGRLRRFRLGAAMAIIYMAMSTLFFLRIAPLTPWWGALAAWVMILLIVEAIFAFFFPYRCCFNETEGDVQNNPHV |
| CX213 | MELITACALLLIAVHIAIVISCPEPLAAVALVDVPLLYLCRYLLIIRRGVIIVSGAGAGGGRLRRFRLGAAMAIIYMAMSTLFFLRIAPLTPWWGALAAWVMILLIVEAIFAFFFPYRCCFNETEGDVQNNPHV |
| CX214 | MELITACALLLIAVHIAIVISCPEPLAAVALVDVPLLYLCRYLLIIRRGVIIVSGAGAGGGRLRRFRLGAAMAIIYMAMSTLFFLRIAPLTPWWGALAAWVMILLIVEAIFAFFFPYRCCFNETEGDVQNNPHV |
| CX218 | MELITACALLLIAVHIAIVISCPEPLAAVALVDVPLLYLCRYLLIIRRGVIIVSGAGAGGGRLRRFRLGAAMAIIYMAMSTLFFLRIAPLTPWWGALAAWVMILLIVEAIFAFFFPYRCCFNETEGDVQNNPHV |
| CX219 | MELITACALLLIAVHIAIVISCPEPLAAVALVDVPLLYLCRYLLIIRRGVIIVSGGGGGGRLRRFRLGAAMAIIYMAMSTLFFLRIAPLTPWWGALAAWVMILLIVEAIFAFFFPYRCCFNETEGDVQNNPHV |
| CX220 | MELITACALLLIAVHIAIVISCPEPLAAVALVDVPLLYLCRYLLIIRRGVIIVSGAGAGGGRLRRFRLGAAMAIIYMAMSTLFFLRIAPLTPWWGALAAWVMILLIVEAIFAFFFPYRCCFNETEGDVQNNPHV |
| CX221 | MELITACALLLIAVHIAIVISCPEPLAAVALVDVPLLYLCRYLLIIRRGVIIVSGGGGGGRLRRFRLGAAMAIIYMAMSTLFFLRIAPLTPWWGALAAWVMILLIVEAIFAFFFPYRCCFNETEGDVQNNPHV |
| CX225 | MELITACALLLIAVHIAIVISCPEPLAAVALVDVPLLYLCRYLLIIRRGVIIVSGAGAGGGRLRRFRLGAAMAIIYMAMSTLFFLRIAPLTPWWGALAAWVMILLIVEAIFAFFFPYRCCFNETEGDVQNNPHV |
| CX226 | MELITACALLLIAVHIAIVISCPEPLAAVALVDVPLLYLCRYLLIIRRGVIIVSGAGAGGGRLRRFRLGAAMAIIYMAMSTLFFLRIAPLTPWWGALAAWVMILLIVEAIFAFFFPYRCCFNETEGDVQNNPHV |
| CX227 | MELITACALLLIAVHIAIVISCPEPLAAVALVDVPLLYLCRYLLIIRRGVIIVSGAGAGGGRLRRFRLGAAMAIIYMAMSTLFFLRIAPLTPWWGALAAWVMILLIVEAIFAFFFPYRCCFNETEGDVQNNPHV |
| CX228 | MELITACALLLIAVHIAIVISCPEPLAAVALVDVPLLYLCRYLLIIRRGVIIVSGAGAGGGRLRRFRLGAAMAIIYMAMSTLFFLRIAPLTPWWGALAAWVMILLIVEAIFAFFFPYRCCFNETEGDVQNNPHV |
| CX230 | MELITACALLLIAVHIAIVISCPEPLAAVALVDVPLLYLCRYLLIIRRGVIIVSGGGGGGRLRRFRLGAAMAIIYMAMSTLFFLRIAPLTPWWGALAAWVMILLIVEAIFAFFFPYRCCFNETEGDVQNNPHV |
| CX231 | MELITACALLLIAVHIAIVISCPEPLAAVALVDVPLLYLCRYLLIIRRGVIIVSGAGAGGGRLRRFRLGAAMAIIYMAMSTLFFLRIAPLTPWWGALAAWVMILLIVEAIFAFFFPYRCCFNETEGDVQNNPHV |
| CX232 | MELITACALLLIAVHIAIVISCPEPLAAVALVDVPLLYLCRYLLIIRRGVIIVSGGGGGGRLRRFRLGAAMAIIYMAMSTLFFLRIAPLTPWWGALAAWVMILLIVEAIFAFFFPYRCCFNETEGDVQNNPHV |
| CX233 | MELITACALLLIAVHIAIVISCPEPLAAVALVDVPLLYLCRYLLIIRRGVIIVSGGGGGGRLRRFRLGAAMAIIYMAMSTLFFLRIAPLTPWWGALAAWVMILLIVEAIFAFFFPYRCCFNETEGDVQNNPHV |
| CX234 | MELITACALLLIAVHIAIVISCPEPLAAVALVDVPLLYLCRYLLIIRRGVIIVSGGGAGGGRLRRFRLGAAMAIIYMAMSTLFFLRIAPLTPWWGALAAWVMILLIVEAIFAFFFPYRCCFNETEGDVQNNPHV |
| CX235 | MELITACALLLIAVHIAIVISCPEPLAAVALVDVPLLYLCRYLLIIRRGVIIVSGGGGGGRLRRFRLGAAMAIIYMAMSTLFFLRIAPLTPWWGALAAWVMILLIVEAIFAFFFPYRCCFNETEGDVQNNPHV |
| CX236 | MELITACALLLIAVHIAIVISCPEPLAAVALVDVPLLYLCRYLLIIRRGVIIVSGAGGGGRLRRFRLGAAMAIIYMAMSTLFFLRIAPLTPWWGALAAWVMILLIVEAIFAFFFPYRCCFNETEGDVQNNPHV |
| CX237 | MELITACALLLIAVHIAIVISCPEPLAAVALVDVPLLYLCRYLLIIRRGVIIVSGGGGGGRLRRFRLGAAMAIIYMAMSTLFFLRIAPLTPWWGALAAWVMILLIVEAIFAFFFPYRCCFNETEGDVQNNPHV |
| CX238 | MELITACALLLIAVHIAIVISCPEPLAAVALVDVPLLYLCRYLLIIRRGVIIVSGGGGGGRLRRFRLGAAMAIIYMAMSTLFFLRIAPLTPWWGALAAWVMILLIVEAIFAFFFPYRCCFNETEGDVQNNPHV |
| CX240 | MELITACALLLIAVHIAIVISCPEPLAAVALVDVPLLYLCRYLLIIRRGVIIVSGGGGGGRLRRFRLGAAMAIIYMAMSTLFFLRIAPLTPWWGALAAWVMILLIVEAIFAFFFPYRCCFNETEGDVQNNPHV |
| CX241 | MELITACALLLIAVHIAIVISCPEPLAAVALVDVPLLYLCRYLLIIRRGVIIVSGAGAGGGRLRRFRLGAAMAIIYMAMSTLFFLRIAPLTPWWGALAAWVMILLIVEAIFAFFFPYRCCFNETEGDVQNNPHV |
| CX242 | MELITACALLLIAVHIAIVISCPEPLAAVALVDVPLLYLCRYLLIIRRGVIIVSGGGGGGRLRRFRLGAAMAIIYMAMSTLFFLRIAPLTPWWGALAAWVMILLIVEAIFAFFFPYRCCFNETEGDVQNNPHV |
| CX243 | MELITACALLLIAVHIAIVISCPEPLAAVALVDVPLLYLCRYLLIIRRGVIIVSGAGAGGGRLRRFRLGAAMAIIYMAMSTLFFLRIAPLTPWWGALAAWVMILLIVEAIFAFFFPYRCCFNETEGDVQNNPHV |
| CX247 | MELITACALLLIAVHIAIVISCPEPLAAVALVDVPLLYLCRYLLIIRRGVIIVSGAGAGGGRLRRFRLGAAMAIIYMAMSTLFFLRIAPLTPWWGALAAWVMILLIVEAIFAFFFPYRCCFNETEGDVQNNPHV |
| CX248 | MELITACALLLIAVHIAIVISCPEPLAAVALVDVPLLYLCRYLLIIRRGVIIVSGAGGGGGRLRRFRLGAAMAIIYMAMSTLFFLRIAPLTPWWGALAAWVMILLIVEAIFAFFFPYRCCFNETEGDVQNNPHV |
| CX249 | MELITACALLLIAVHIAIVISCPEPLAAVALVDVPLLYLCRYLLIIRRGVIIVSGGGGGGRLRRFRLGAAMAIIYMAMSTLFFLRIAPLTPWWGALAAWVMILLIVEAIFAFFFPYRCCFNETEGDVQNNPHV |
| CX250 | MELITACALLLIAVHIAIVISCPEPLAAVALVDVPLLYLCRYLLIIRRGVIIVSGAGAGGGRLRRFRLGAAMAIIYMAMSTLFFLRIAPLTPWWGALAAWVMILLIVEAIFAFFFPYRCCFNETEGDVQNNPHV |
| CX251 | MELITACALLLIAVHIAIVISCPEPLAAVAFVNVPLLYLCRYLLIIRRGVIIVSGGGGGGGRLRRFRLGAAMAVIYMAMSTLFFLRIAPLTPWWGALAAWVMILLIVEAIFAFFFPYRCCFNETEGDVQNNPHV |
| CX262 | MELITACALLLIAVHIAIVISCPEPLAAVALVDVPLLYLYRYLLIIRRGVIIVSGAGGGGGRLRRFRLGAAMAIIYMAMSTLFFLRIAPLTPWWGALAAWVMILLIVEAIFAFFFPYRCCFNETEGDVQNNPHV |
| CX263 | MELITACALLLIAVHIAIVISCPEPLAAVALVDVPLLYLCRYLLIIRRGVIIVSGAGAGGGRLRRFRLGAAMAIIYMAMSTLFFLRIAPLTPWWGALAAWVMILLIVEAIFAFFFPYRCCFNETEGDVQNNPHV |
| CX265 | MELITACALLLIAVHIAIVISCPEPLAAVALVDVPLLYLCRYLLIIRRGVIIVSGGGGGGRLRRFRLGAAMAIIYMAMSTLFFLRIAPLTPWWGALAAWVMILLIVEAIFAFFFPYRCCFNETEGDVQNNPHV |
| CX266 | MELITACALLLIAVHIAIVISCPEPLAAVALVDVPLLYLYRYLLIIRRGVIIVSGAGGGGGRLRRFRLGAAMAIIYMAMSTLFFLRIAPLTPWWGALAAWVMILLIVEAIFAFFFPYRCCFNETEGDVQNNPHV |
| CX267 | MELITACALLLIAVHIAIVISCPEPLAAVALVDVPLLYLCRYLLIIRRGVIIVSGAGAGGGRLRRFRLGAAMAIIYMAMSTLFFLRIAPLTPWWGALAAWVMILLIVEAIFAFFFPYRCCFNETEGDVQNNPHV |
| CX268 | MELITACALLLIAVHIAIVISCPEPLAAVALVDVPLLYLCRYLLIIRRGVIIVSGGGAGGGRLRRFRLGAAMAIIYMAMSTLFFLRIAPLTPWWGALAAWVMILLIVEAIFAFFFPYRCCFNETEGDVQNNPHV |
| CX269 | MELITACALLLIAVHIAIVISCPEPLAAVALVDVPLLYLCRYLLIIRRGVIIVSGAGAGGGRLRRFRLGAAMAIIYMAMSTLFFLRIAPLTPWWGALAAWVMILLIVEAIFAFFFPYRCCFNETEGDVQNNPHV |
| CX270 | MELITACALLLIAVHIAIVISCPEPLAAVALVDVPLLYLCRYLLIIRRGVIIVSGAGAGGGRLRRFRLGAAMAIIYMAMSTLFFLRIAPLTPWWGALAAWVMILLIVEAIFAFFFPYRCCFNETEGDVQNNPHV |
| CX273 | MELITACALLLIAVHIAIVISCPEPLAAVALVDVPLLYLCRYLLIIRRGVIIVSGAGAGGGRLRRFRLGAAMAIIYMAMSTLFFLRIAPLTPWWGALAAWVMILLIVEAIFAFFFPYRCCFNETEGDVQNNPHV |
| CX274 | MELITACALLLIAVHIAIVISCPEPLAAVALVDVPLLYLCRYLLIIRRGVIIVSGAGAGGGRLRRFRLGAAMAIIYMAMSTLFFLRIAPLTPWWGALAAWVMILLIVEAIFAFFFPYRCCFNETEGDVQNNPHV |
| CX275 | MELITACALLLIAVHIAIVISCPEPLAAVALVDVPLLYLCRYLLIIRRGVIIVSGAGAGGGRLRRFRLGAAMAIIYMAMSTLFFLRIAPLTPWWGALAAWVMILLIVEAIFAFFFPYRCCFNETEGDVQNNPHV |
| CX276 | MELITACALLLIAVHIAIVISCPEPLAAVALVDVPLLYLCRYLLIIRRGVIIVSGGGGGGRLRRFRLGAAMAIIYMAMSTLFFLRIAPLTPWWGALAAWVMILLIVEAIFAFFFPYRCCFNETEGDVQNNPHV |
| CX277 | MELITACALLLIAVHIAIVISCPEPLAAVALVDVPLLYLCRYLLIIRRGVIIVSGAGGGGGRLRRFRLGAAMAIIYMAMSTLFFLRIAPLTPWWGALAAWVMILLIVEAIFAFFFPYRCCFNETEGDVQNNPHV |
| CX278 | MELITACALLLIAVHIAIVISCPEPLAAVALVDVPLLYLCRYLLIIRRGVIIVSGGGGGGRLRRFRLGAAMAIIYMAMSTLFFLRIAPLTPWWGALAAWVMILLIVEAIFAFFFPYRCCFNETEGDVQNNPHV |
| CX280 | MELITACALLLIAVHIAIVISCPEPLAAVALVDVPLLYLCRYLLIIRRGVIIVSGAGAGGGRLRRFRLGAAMAIIYMAMSTLFFLRIAPLTPWWGALAAWVMILLIVEAIFAFFFPYRCCFNETEGDVQNNPHV |
| CX281 | MELITACALLLIAVHIAIVISCPEPLAAVALVDVPLLYLCRYLLIIRRGVIIVSGAGAGGGRLRRFRLGAAMAIIYMAMSTLFFLRIAPLTPWWGALAAWVMILLIVEAIFAFFFPYRCCFNETEGDVQNNPHV |
| CX282 | MELITACALLLIAVHIAIVISCPEPLAAVALVDVPLLYLCRYLLIIRRGVIIVSGAGGGGGRLRRFRLGAAMAIIYMAMSTLFFLRIAPLTPWWGALAAWVMILLIVEAIFAFFFPYRCCFNETEGDVQNNPHV |
| CX284 | MELITACALLLIAVHIAIVISCPEPLAAVALVDVPLLYLCRYLLIIRRGVIIVSGAGGGGGRLRRFRLGAAMAIIYMAMSTLFFLRIAPLTPWWGALAAWVMILLIVEAIFAFFFPYRCCFNETEGDVQNNPHV |
| CX285 | MELITACALLLIAVHIAIVISCPEPLAAVALVDVPLLYLYRYLLIIRRGVIIVSGAGGGGGRLRRFRLGAAMAIIYMAMSTLFFLRIAPLTPWWGALAAWVMILLIVEAIFAFFFPYRCCFNETEGDVQNNPHV |
| CX286 | MELITACALLLIAVHIAIVISCPEPLAAVALVDVPLLYLCRYLLIIRRGVIIVSGGGGGGRLRRFRLGAAMAIIYMAMSTLFFLRIAPLTPWWGALAAWVMILLIVEAIFAFFFPYRCCFNETEGDVQNNPHV |
| CX287 | MELITACALLLIAVHIAIVISCPEPLAAVALVDVPLLYLCRYLLIIRRGVIIVSGAGAGGGRLRRFRLGAAMAIIYMAMSTLFFLRIAPLTPWWGALAAWVMILLIVEAIFAFFFPYRCCFNETEGDVQNNPHV |
| CX288 | MELITACALLLIAVHIAIVISCPEPLAAVALVDVPLLYLCRYLLIIRRGVIIVSGGGGGGRLRRFRLGAAMAIIYMAMSTLFFLRIAPLTPWWGALAAWVMILLIVEAIFAFFFPYRCCFNETEGDVQNNPHV |
| CX290 | MELITACALLLIAVHIAIVISCPEPLAAVALVDVPLLYLCRYLLIIRRGVIIVSGGGGGGRLRRFRLGAAMAIIYMAMSTLFFLRIAPLTPWWGALAAWVMILLIVEAIFAFFFPYRCCFNETEGDVQNNPHV |
| CX291 | MELITACALLLIAVHIAIVISCPEPLAAVALVDVPLLYLCRYLLIIRRGVIIVSGGGGGGRLRRFRLGAAMAIIYMAMSTLFFLRIAPLTPWWGALAAWVMILLIVEAIFAFFFPYRCCFNETEGDVQNNPHV |
| CX296 | MELITACALLLIAVHIAIVISCPEPLAAVALVDVPLLYLCRYLLIIRRGVIIVSGAGAGGGRLRRFRLGAAMAIIYMAMSTLFFLRIAPLTPWWGALAAWVMILLIVEAIFAFFFPYRCCFNETEGDVQNNPHV |
| CX303 | MELITACALLLIAVHIAIVISCPEPLAAVALVDVPLLYLCRYLLIIRRGVIIVSGGGGGGRLRRFRLGAAMAIIYMAMSTLFFLRIAPLTPWWGALAAWVMILLIVEAIFAFFFPYRCCFNETEGDVQNNPHV |
| CX304 | MELITACALLLIAVHIAIVISCPEPLAAVALVDVPLLYLCRYLLIIRRGVIIVSGAGAGGGRLRRFRLGAAMAIIYMAMSTLFFLRIAPLTPWWGALAAWVMILLIVEAIFAFFFPYRCCFNETEGDVQNNPHV |
| CX305 | MELITACALLLIAVHIAIVISCPEPLAAVALVDVPLLYLCRYLLIIRRGVIIVSGAGAGGGRLRRFRLGAAMAIIYMAMSTLFFLRIAPLTPWWGALAAWVMILLIVEAIFAFFFPYRCCFNETEGDVQNNPHV |
| CX306 | MELITACALLLIAVHIAIVISCPEPLAAVALVDVPLLYLCRYLLIIRRGVIIVSGAGAGGGRLRRFRLGAAMAIIYMAMSTLFFLRIAPLTPWWGALAAWVMILLIVEAIFAFFFPYRCCFNETEGDVQNNPHV |
| CX307 | MELITACALLLIAVHIAIVISCPEPLAAVALVDVPLLYLYRYLLIIRRGVIIVSGAGGGGGRLRRFRLGAAMAIIYMAMSTLFFLRIAPLTPWWGALAAWVMILLIVEAIFAFFFPYRCCFNETEGDVQNNPHV |
| CX313 | MELITACALLLIAVHIAIVISCPEPLAAVALVDVPLLYLCRYLLIIRRGVIIVSGAGAGGGRLRRFRLGAAMAIIYMAMSTLFFLRIAPLTPWWGALAAWVMILLIVEAIFAFFFPYRCCFNETEGDVQNNPHV |
| CX314 | MELITACALLLIAVHIAIVISCPEPLAAVALVDVPLLYLCRYLLIIRRGVIIVSGGGGGGRLRRFRLGAAMAIIYMAMSTLFFLRIAPLTPWWGALAAWVMILLIVEAIFAFFFPYRCCFNETEGDVQNNPHV |
| CX315 | MELITACALLLIAVHIAIVISCPEPLAAVALVDVPLLYLCRYLLIIRRGVIIVSGAGGGGGRLRRFRLGAAMAIIYMAMSTLFFLRIAPLTPWWGALAAWVMILLIVEAIFAFFFPYRCCFNETEGDVQNNPHV |
| CX316 | MELITACALLLIAVHIAIVISCPEPLAAVALVDVPLLYLCRYLLIIRRGVIIVSGGGGGGRLRRFRLGAAMAIIYMAMSTLFFLRIAPLTPWWGALAAWVMILLIVEAIFAFFFPYRCCFNETEGDVQNNPHV |
| CX317 | MELITACALLLIAVHIAIVISCPEPLAAVALVDVPLLYLCRYLLIIRRGVIIVSGGGGGGRLRRFRLGAAMAIIYMAMSTLFFLRIAPLTPWWGALAAWVMILLIVEAIFAFFFPYRCCFNETEGDVQNNPHV |
| CX318 | MELITACALLLIAVHIAIVISCPEPLAAVALVDVPLLYLCRYLLIIRRGVIIVSGGGGGGRLRRFRLGAAMAIIYMAMSTLFFLRIAPLTPWWGALAAWVMILLIVEAIFAFFFPYRCCFNETEGDVQNNPHV |
| CX319 | MELITACALLLIAVHIAIVISCPEPLAAVALVDVPLLYLCRYLLIIRRGVIIVSGGGGGGRLRRFRLGAAMAIIYMAMSTLFFLRIAPLTPWWGALAAWVMILLIVEAIFAFFFPYRCCFNETEGDVQNNPHV |
| CX328 | MELITACALLLIAVHIAIVISCPEPLAAVALVDVPLLYLYRYLLIIRRGVIIVSGAGGGGGRLRRFRLGAAMAIIYMAMSTLFFLRIAPLTPWWGALAAWVMILLIVEAIFAFFFPYRCCFNETEGDVQNNPHV |
| CX329 | MELITACALLLIAVHIAIVISCPEPLAAVALVDVPLLYLCRYLLIIRRGVIIVSGAGAGGGRLRRFRLGAAMAIIYMAMSTLFFLRIAPLTPWWGALAAWVMILLIVEAIFAFFFPYRCCFNETEGDVQNNPHV |
| CX330 | MELITACALLLIAVHIAIVISCPEPLAAVALVDVPLLYLCRYLLIIRRGVIIVSGAGAGGGRLRRFRLGAAMAIIYMAMSTLFFLRIAPLTPWWGALAAWVMILLIVEAIFAFFFPYRCCFNETEGDVQNNPHV |
| CX340 | MELITACALLLIAVHIAIVISCPEPLAAVALVDVPLLYLCRYLLIIRRGVIIVSGGGGGGRLRRFRLGAAMAIIYMAMSTLFFLRIAPLTPWWGALAAWVMILLIVEAIFAFFFPYRCCFNETEGDVQNNPHV |
| CX341 | MELITACALLLIAVHIAIVISCPEPLAAVALVDVPLLYLCRYLLIIRRGVIIVSGGGGGGRLRRFRLGAAMAIIYMAMSTLFFLRIAPLTPWWGALAAWVMILLIVEAIFAFFFPYRCCFNETEGDVQNNPHV |
| CX342 | MELITACALLLIAVHIAIVISCPEPLAAVALVDVPLLYLCRYLLIIRRGVIIVSGGGGGGRLRRFRLGAAMAIIYMAMSTLFFLRIAPLTPWWGALAAWVMILLIVEAIFAFFFPYRCCFNETEGDVQNNPHV |
| CX343 | MELITACALLLIAVHIAIVISCPEPLAAVALVDVPLLYLCRYLLIIRRGVIIVSGGGGGGRLRRFRLGAAMAIIYMAMSTLFFLRIAPLTPWWGALAAWVMILLIVEAIFAFFFPYRCCFNETEGDVQNNPHV |
| CX344 | MELITACALLLIAVHIAIVISCPEPLAAVALVDVPLLYLYRYLLIIRRGVIIVSGAGGGGGRLRRFRLGAAMAIIYMAMSTLFFLRIAPLTPWWGALAAWVMILLIVEAIFAFFFPYRCCFNETEGDVQNNPHV |
| CX345 | MELITACALLLIAVHIAIVISCPEPLAAVAFVNVPLLYLCRYLLIIRRGVIIVSGGGAGGGRLRRFRLGAAMAVIYMAMSTLFFLRIAPLTPWWGALAAWVMILLIVEAIFAFFFPYRCCFNETEGDVQNNPHV |
| CX346 | MELITACALLLIAVHIAIVISCPEPLAAVALVDVPLLYLCRYLLIIRRGVIIVSGAGAGGGRLRRFRLGAAMAIIYMAMSTLFFLRIAPLTPWWGALAAWVMILLIVEAIFAFFFPYRCCFNETEGDVQNNPHV |
| CX347 | MELITACALLLIAVHIAIVISCPEPLAAVALVDVPLLYLCRYLLIIRRGVIIVSGAGAGGGRLRRFRLGAAMAIIYMAMSTLFFLRIAPLTPWWGALAAWVMILLIVEAIFAFFFPYRCCFNETEGDVQNNPHV |
| CX348 | MELITACALLLIAVHIAIVISCPEPLAAVALVDVPLLYLCRYLLIIRRGVIIVSGAGAGGGRLRRFRLGAAMAIIYMAMSTLFFLRIAPLTPWWGALAAWVMILLIVEAIFAFFFPYRCCFNETEGDVQNNPHV |
| CX349 | MELITACALLLIAVHIAIVISCPEPLAAVALVDVPLLYLCRYLLIIRRGVIIVSGAGAGGGRLRRFRLGAAMAIIYMAMSTLFFLRIAPLTPWWGALAAWVMILLIVEAIFAFFFPYRCCFNETEGDVQNNPHV |
| CX350 | MELITACALLLIAVHIAIVISCPEPLAAVALVDVPLLYLCRYLLIIRRGVIIVSGAGAGGGRLRRFRLGAAMAIIYMAMSTLFFLRIAPLTPWWGALAAWVMILLIVEAIFAFFFPYRCCFNETEGDVQNNPHV |

| | |
|---|---|
| CX351 | MELITACALLLIAVHIAIVISCPEPLAAVALVDVPLLYLCRYLLIIRRGVIIVSGAGGGGGRLRRFRLGAAMAIIYMAMSTLFFLRIAPLTPWWGALAAWVMILLIVEAIFAFFFPYRCCFNETEGDVQNNPHV |
| CX352 | MELITACALLLIAVHIAIVISCPEPLAAVALVDVPLLYLCRYLLIIRRGVIIVSGAGAGGGRLRRFRLGAAMAIIYMAMSTLFFLRIAPLTPWWGALAAWVMILLIVEAIFAFFFPYRCCFNETEGDVQNNPHV |
| CX353 | MELITACALLLIAVHIAIVISCPEPLAAVALVDVPLLYLCRYLLIIRRGVIIVSGAGAGGGRLRRFRLGAAMAIIYMAMSTLFFLRIAPLTPWWGALAAWVMILLIVEAIFAFFFPYRCCFNETEGDVQNNPHV |
| CX354 | MELITACALLLIAVHIAIVISCPEPLAAVALVDVPLLYLYRYLLIIRRGVIIVSGAGAGGGRLRRFRLGAAMAIIYMAMSTLFFLRIAPLTPWWGALAAWVMILLIVEAIFAFFFPYRCCFNETEGDVQNNPHV |
| CX355 | MELITACALLLIAVHIAIVISCPEPLAAVALVDVPLLYLCRYLLIIRRGVIIVSGAGAGGGRLRRFRLGAAMAIIYMAMSTLFFLRIAPLTPWWGALAAWVMILLIVEAIFAFFFPYRCCFNETEGDVQNNPHV |
| CX356 | MELITACALLLIAVHIAIVISCPEPLAAVAFVNVPLLYLCRYLLIIRRGVIIVSGGGGGGGRLRRFRLGAAMAVIYMAMSTLFFLRIAPLTPWWGALAAWVMILLIVEAIFAFFFPYRCCFNETEGDVQNNPHV |
| CX357 | MELITACALLLIAVHIAIVISCPEPLAAVALVDVPLLYLCRYLLIIRRGVIIVSGAGAGGGRLRRFRLGAAMAIIYMAMSTLFFLRIAPLTPWWGALAAWVMILLIVEAIFAFFFPYRCCFNETEGDVQNNPHV |
| CX358 | MELITACALLLIAVHIAIVISCPEPLAAVALVDVPLLYLCRYLLIIRRGVIIVSGGGGGGRLRRFRLGAAMAIIYMAMSTLFFLRIAPLTPWWGALAAWVMILLIVEAIFAFFFPYRCCFNETEGDVQNNPHV |
| CX359 | MELITACALLLIAVHIAIVISCPEPLAAVALVDVPLLYLCRYLLIIRRGVIIVSGAGAGGGRLRRFRLGAAMAIIYMAMSTLFFLRIAPLTPWWGALAAWVMILLIVEAIFAFFFPYRCCFNETEGDVQNNPHV |
| CX360 | MELITACALLLIAVHIAIVISCPEPLAAVALVDVPLLYLCRYLLIIRRGVIIVSGGGGGGRLRRFRLGAAMAIIYMAMSTLFFLRIAPLTPWWGALAAWVMILLIVEAIFAFFFPYRCCFNETEGDVQNNPHV |
| CX361 | MELITACALLLIAVHIAIVISCPEPLAAVALVDVPLLYLCRYLLIIRRGVIIVSGGGGGGRLRRFRLGAAMAIIYMAMSTLFFLRIAPLTPWWGALAAWVMILLIVEAIFAFFFPYRCCFNETEGDVQNNPHV |
| CX362 | MELITACALLLIAVHIAIVISCPEPLAAVALVDVPLLYLCRYLLIIRRGVIIVSGGGGGGRLRRFRLGAAMAIIYMAMSTLFFLRIAPLTPWWGALAAWVMILLIVEAIFAFFFPYRCCFNETEGDVQNNPHV |
| CX363 | MELITACALLLIAVHIAIVISCPEPLAAVALVDVPLLYLCRYLLIIRRGVIIVSGGGGGGRLRRFRLGAAMAIIYMAMSTLFFLRIAPLTPWWGALAAWVMILLIVEAIFAFFFPYRCCFNETEGDVQNNPHV |
| CX364 | MELITACALLLIAVHIAIVISCPEPLAAVALVDVPLLYLCRYLLIIRRGVIIVSGGGGGGRLRRFRLGAAMAIIYMAMSTLFFLRIAPLTPWWGALAAWVMILLIVEAIFAFFFPYRCCFNETEGDVQNNPHV |
| CX365 | MELITACALLLIAVHIAIVISCPEPLAAVALVDVPLLYLCRYLLIIRRGVIIVSGGGGGGRLRRFRLGAAMAIIYMAMSTLFFLRIAPLTPWWGALAAWVMILLIVEAIFAFFFPYRCCFNETEGDVQNNPHV |
| CX366 | MELITACALLLIAVHIAIVISCPEPLAAVALVDVPLLYLCRYLLIIRRGVIIVSGGGGGGRLRRFRLGAAMAIIYMAMSTLFFLRIAPLTPWWGALAAWVMILLIVEAIFAFFFPYRCCFNETEGDVQNNPHV |
| CX367 | MELITACALLLIAVHIAIVISCPEPLAAVAFVNVPLLYLCRYLLIIRRGVIIVSGAGAGGGRLRRFRLGAAMAIIYMAMSTLFFLRIAPLTPWWGALAAWVMILLIVEAIFAFFFPYRCCFNETEGDVQNNPHV |
| CX368 | MELITACALLLIAVHIAIVISCPEPLAAVAFVNVPLLYLCRYLLIIRRGVIIVSGGGGGGGRLRRFRLGAAMAVIYMAMSTLFFLRIAPLTPWWGALAAWVMILLIVEAIFAFFFPYRCCFNETEGDVQNNPHV |
| CX369 | MELITACALLLIAVHIAIVISCPEPLAAVALVDVPLLYLCRYLLIIRRGVIIVSGAGAGGGRLRRFRLGAAMAIIYMAMSTLFFLRIAPLTPWWGALAAWVMILLIVEAIFAFFFPYRCCFNETEGDVQNNPHV |
| CX370 | MELITACALLLIAVHIAIVISCPEPLAAVALVDVPLLYLCRYLLIIRRGVIIVSGGGGGGRLRRFRLGAAMAIIYMAMSTLFFLRIAPLTPWWGALAAWVMILLIVEAIFAFFFPYRCCFNETEGDVQNNPHV |
| CX371 | MELITACALLLIAVHIAIVISCPEPLAAVALVDVPLLYLCRYLLIIRRGVIIVSGAGAGGGRLRRFRLGAAMAIIYMAMSTLFFLRIAPLTPWWGALAAWVMILLIVEAIFAFFFPYRCCFNETEGDVQNNPHV |
| CX372 | MELITACALLLIAVHIAIVISCPEPLAAVALVDVPLLYLCRYLLIIRRGVIIVSGAGAGGGRLRRFRLGAAMAIIYMAMSTLFFLRIAPLTPWWGALAAWVMILLIVEAIFAFFFPYRCCFNETEGDVQNNPHV |
| CX373 | MELITACALLLIAVHIAIVISCPEPLAAVALVDVPLLYLCRYLLIIRRGVIIVSGAGAGGGRLRRFRLGAAMAIIYMAMSTLFFLRIAPLTPWWGALAAWVMILLIVEAIFAFFFPYRCCFNETEGDVQNNPHV |
| CX374 | MELITACALLLIAVHIAIVISCPEPLAAVALVDVPLLYLCRYLLIIRRGVIIVSGAGAGGGRLRRFRLGAAMAIIYMAMSTLFFLRIAPLTPWWGALAAWVMILLIVEAIFAFFFPYRCCFNETEGDVQNNPHV |
| CX375 | MELITACALLLIAVHIAIVISCPEPLAAVALVDVPLLYLCRYLLIIRRGVIIVSGGGGGGRLRRFRLGAAMAIIYMAMSTLFFLRIAPLTPWWGALAAWVMILLIVEAIFAFFFPYRCCFNETEGDVQNNPHV |
| CX376 | MELITACALLLIAVHIAIVISCPEPLAAVALVDVPLLYLCRYLLIIRRGVIIVSGGGGGGRLRRFRLGAAMAIIYMAMSTLFFLRIAPLTPWWGALAAWVMILLIVEAIFAFFFPYRCCFNETEGDVQNNPHV |
| CX377 | MELITACALLLIAVHIAIVISCPEPLAAVALVDVPLLYLCRYLLIIRRGVIIVSGAGGGGRLRRFRLGAAMAIIYMAMSTLFFLRIAPLTPWWGALAAWVMILLIVEAIFAFFFPYRCCFNETEGDVQNNPHV |
| CX378 | MELITACALLLIAVHIAIVISCPEPLAAVALVDVPLLYLCRYLLIIRRGVIIVSGGGGGGRLRRFRLGAAMAIIYMAMSTLFFLRIAPLTPWWGALAAWVMILLIVEAIFAFFFPYRCCFNETEGDVQNNPHV |
| CX379 | MELITACALLLIAVHIAIVISCPEPLAAVALVDVPLLYLCRYLLIIRRGVIIVSGGGGGGRLRRFRLGAAMAIIYMAMSTLFFLRIAPLTPWWGALAAWVMILLIVEAIFAFFFPYRCCFNETEGDVQNNPHV |
| CX380 | MELITACALLLIAVHIAIVISCPEPLAAVALVDVPLLYLCRYLLIIRRGVIIVSGAGAGGGRLRRFRLGAAMAIIYMAMSTLFFLRIAPLTPWWGALAAWVMILLIVEAIFAFFFPYRCCFNETEGDVQNNPHV |
| CX381 | MELITACALLLIAVHIAIVISCPEPLAAVALVDVPLLYLCRYLLIIRRGVIIVSGGGGGGRLRRFRLGAAMAIIYMAMSTLFFLRIAPLTPWWGALAAWVMILLIVEAIFAFFFPYRCCFNETEGDVQNNPHV |
| CX382 | MELITACALLLISVHIAIVISCPEPLAAVALVDVPLLYLCRYLLIIRRGVIIVSGGGGGGRLRRFRLGAAMAIIYMAMSTLFFLRIAPLTPWWGALAAWVMILLIVEAIFAFFFPYRCCFNETEGDVQNNPHV |
| CX383 | MELITACALLLIAVHIAIVISCPEPLAAVALVDVPLLYLCRYLLIIRRGVIIVSGAGGGGGRLRRFRLGAAMAIIYMAMSTLFFLRIAPLTPWWGALAAWVMILLIVEAIFAFFFPYRCCFNETEGDVQNNPHV |
| CX384 | MELITACALLLIAVHIAIVISCPEPLAAVALVDVPLLYLCRYLLIIRRGVIIVSGAGGGGGRLRRFRLGAAMAIIYMAMSTLFFLRIAPLTPWWGALAAWVMILLIVEAIFAFFFPYRCCFNETEGDVQNNPHV |
| CX385 | MELITACALLLIAVHIAIVISCPEPLAAVALVDVPLLYLCRYLLIIRRGVIIVSGGGGGGRLRRFRLGAAMAIIYMAMSTLFFLRIAPLTPWWGALAAWVMILLIVEAIFAFFFPYRCCFNETEGDVQNNPHV |
| CX386 | MELITACALLLIAVHIAIVISCPEPLAAVALVDVPLLYLCRYLLIIRRGVIIVSGGGGGGRLRRFRLGAAMAIIYMAMSTLFFLRIAPLTPWWGALAAWVMILLIVEAIFAFFFPYRCCFNETEGDVQNNPHV |
| CX387 | MELITACALLLIAVHIAIVISCPEPLAAVALVDVPLLYLCRYLLIIRRGVIIVSGAGAGGGRLRRFRLGAAMAIIYMAMSTLFFLRIAPLTPWWGALAAWVMILLIVEAIFAFFFPYRCCFNETEGDVQNNPHV |
| CX388 | MELITACALLLIAVHIAIVISCPEPLAAVALVDVPLLYLCRYLLIIRRGVIIVSGGGGGGRLRRFRLGAAMAIIYMAMSTLFFLRIAPLTPWWGALAAWVMILLIVEAIFAFFFPYRCCFNETEGDVQNNPHV |
| CX389 | MELITACALLLIAVHIAIVISCPEPLAAVALVDVPLLYLCRYLLIIRRGVIIVSGGGGGGRLRRFRLGAAMAIIYMAMSTLFFLRIAPLTPWWGALAAWVMILLIVEAIFAFFFPYRCCFNETEGDVQNNPHV |
| CX390 | MELITACALLLIAVHIAIVISCPEPLAAVALVDVPLLYLYRYLLIIRRGVIIVSGAGAGGGRLRRFRLGAAMAIIYMAMSTLFFLRIAPLTPWWGALAAWVMILLIVEAIFAFFFPYRCCFNETEGDVQNNPHV |
| CX391 | MELITACALLLIAVHIAIVISCPEPLAAVAFVNVPLLYLCRYLLIIRRGVIIVSGGGGGGGRLRRFRLGAAMAVIYMAMSTLFFLRIAPLTPWWGALAAWVMILLIVEAIFAFFFPYRCCFNETEGDVQNNPHV |
| CX392 | MELITACALLLIAVHIAIVISCPEPLAAVALVDVPLLYLCRYLLIIRRGVIIVSGGGGGGRLRRFRLGAAMAIIYMAMSTLFFLRIAPLTPWWGALAAWVMILLIVEAIFAFFFPYRCCFNETEGDVQNNPHV |
| CX393 | MELITACALLLIAVHIAIVISCPEPLAAVALVDVPLLYLCRYLLIIRRGVIIVSGGGGGGRLRRFRLGAAMAIIYMAMSTLFFLRIAPLTPWWGALAAWVMILLIVEAIFAFFFPYRCCFNETEGDVQNNPHV |
| CX394 | MELITACALLLIAVHIAIVISCPEPLAAVALVDVPLLYLCRYLLIIRRGVIIVSGGGGGGRLRRFRLGAAMAIIYMAMSTLFFLRIAPLTPWWGALAAWVMILLIVEAIFAFFFPYRCCFNETEGDVQNNPHV |
| CX395 | MELITACALLLIAVHIAIVISCPEPLAAVALVDVPLLYLYRYLLIIRRGVIIVSGGGAGGGRLRRFRLGAAMAIIYMAMSTLFFLRIAPLTPWWGALAAWVMILLIVEAIFAFFFPYRCCFNETEGDVQNNPHV |
| CX396 | MELITACALLLIAVHIAIVISCPEPLAAVAFVNVPLLYLCRYLLIIRRGVIIVSGGGGGGGRLRRFRLGAAMAVIYMAMSTLFFLRIAPLTPWWGALAAWVMILLIVEAIFAFFFPYRCCFNETEGDVQNNPHV |
| CX397 | MELITACALLLIAVHIAIVISCPEPLAAVALVDVPLLYLCRYLLIIRRGVIIVSGAGAGGGRLRRFRLGAAMAIIYMAMSTLFFLRIAPLTPWWGALAAWVMILLIVEAIFAFFFPYRCCFNETEGDVQNNPHV |
| CX399 | MELITACALLLIAVHIAIVISCPEPLAAVALVDVPLLYLCRYLLIIRRGVIIVSGGGGGGRLRRFRLGAAMAIIYMAMSTLFFLRIAPLTPWWGALAAWVMILLIVEAIFAFFFPYRCCFNETEGDVQNNPHV |
| CX403 | MELITACALLLIAVHIAIVISCPEPLAAVALVDVPLLYLCRYLLIIRRGVIIVSGGGGGGRLRRFRLGAAMAIIYMAMSTLFFLRIAPLTPWWGALAAWVMILLIVEAIFAFFFPYRCCFNETEGDVQNNPHV |
| CX408 | MELITACALLLIAVHIAIVISCPEPLAAVALVDVPLLYLCRYLLIIRRGVIIVSGAGAGGGRLRRFRLGAAMAIIYMAMSTLFFLRIAPLTPWWGALAAWVMILLIVEAIFAFFFPYRCCFNETEGDVQNNPHV |
| CX409 | MELITACALLLIAVHIAIVISCPEPLAAVALVDVPLLYLCRYLLIIRRGVIIVSGAGAGGGRLRRFRLGAAMAIIYMAMSTLFFLRIAPLTPWWGALAAWVMILLIVEAIFAFFFPYRCCFNETEGDVQNNPHV |
| CX410 | MELITACALLLIAVHIAIVISCPEPLAAVALVDVPLLYLCRYLLIIRRGVIIVSGAGAGGGRLRRFRLGAAMAIIYMAMSTLFFLRIAPLTPWWGALAAWVMILLIVEAIFAFFFPYRCCFNETEGDVQNNPHV |
| CX411 | MELITACALLLIAVHIAIVISCPEPLAAVALVDVPLLYLCRYLLIIRRGVIIVSGAGAGGGRLRRFRLGAAMAIIYMAMSTLFFLRIAPLTPWWGALAAWVMILLIVEAIFAFFFPYRCCFNETEGDVQNNPHV |
| CX413 | MELITACALLLIAVHIAIVISCPEPLAAVALVDVPLLYLCRYLLIIRRGVIIVSGAGAGGGRLRRFRLGAAMAIIYMAMSTLFFLRIAPLTPWWGALAAWVMILLIVEAIFAFFFPYRCCFNETEGDVQNNPHV |
| CX414 | MELITACALLLIAVHIAIVISCPEPLAAVALVDVPLLYLCRYLLIIRRGVIIVSGAGAGGGRLRRFRLGAAMAIIYMAMSTLFFLRIAPLTPWWGALAAWVMILLIVEAIFAFFFPYRCCFNETEGDVQNNPHV |
| CX416 | MELITACALLLIAVHIAIVISCPEPLAAVALVDVPLLYLCRYLLIIRRGVIIVSGAGAGGGRLRRFRLGAAMAIIYMAMSTLFFLRIAPLTPWWGALAAWVMILLIVEAIFAFFFPYRCCFNETEGDVQNNPHV |
| CX417 | MELITACALLLIAVHIAIVISCPEPLAAVALVDVPLLYLCRYLLIIRRGVIIVSGAGAGGGRLRRFRLGAAMAIIYMAMSTLFFLRIAPLTPWWGALAAWVMILLIVEAIFAFFFPYRCCFNETEGDVQNNPHV |
| CX418 | MELITACALLLIAVHIAIVISCPEPLAAVALVDVPLLYLCRYLLIIRRGVIIVSGAGAGGGRLRRFRLGAAMAIIYMAMSTLFFLRIAPLTPWWGALAAWVMILLIVEAIFAFFFPYRCCFNETEGDVQNNPHV |
| CX419 | MELITACALLLIAVHIAIVISCPEPLAAVALVDVPLLYLCRYLLIIRRGVIIVSGAGAGGGRLRRFRLGAAMAIIYMAMSTLFFLRIAPLTPWWGALAAWVMILLIVEAIFAFFFPYRCCFNETEGDVQNNPHV |
| CX420 | MELITACALLLIAVHIAIVISCPEPLAAVALVDVPLLYLCRYLLIIRRGVIIVSGAGAGGGRLRRFRLGAAMAIIYMAMSTLFFLRIAPLTPWWGALAAWVMILLIVEAIFAFFFPYRCCFNETEGDVQNNPHV |
| CX421 | MELITACALLLIAVHIAIVISCPEPLAAVALVDVPLLYLCRYLLIIRRGVIIVSGAGAGGGRLRRFRLGAAMAIIYMAMSTLFFLRIAPLTPWWGALAAWVMILLIVEAIFAFFFPYRCCFNETEGDVQNNPHV |
| CX422 | MELITACALLLIAVHIAIVISCPEPLAAVALVDVPLLYLCRYLLIIRRGVIIVSGAGAGGGRLRRFRLGAAMAIIYMAMSTLFFLRIAPLTPWWGALAAWVMILLIVEAIFAFFFPYRCCFNETEGDVQNNPHV |
| CX423 | MELITACALLLIAVHIAIVISCPEPLAAVALVDVPLLYLCRYLLIIRRGVIIVSGAGAGGGRLRRFRLGAAMAIIYMAMSTLFFLRIAPLTPWWGALAAWVMILLIVEAIFAFFFPYRCCFNETEGDVQNNPHV |
| CX424 | MELITACALLLIAVHIAIVISCPEPLAAVALVDVPLLYLCRYLLIIRRGVIIVSGAGAGGGRLRRFRLGAAMAIIYMAMSTLFFLRIAPLTPWWGALAAWVMILLIVEAIFAFFFPYRCCFNETEGDVQNNPHV |
| CX425 | MELITACALLLIAVHIAIVISCPEPLAAVALVDVPLLYLCRYLLIIRRGVIIVSGAGAGGGRLRRFRLGAAMAIIYMAMSTLFFLRIAPLTPWWGALAAWVMILLIVEAIFAFFFPYRCCFNETEGDVQNNPHV |
| CX426 | MELITACALLLIAVHIAIVISCPEPLAAVALVDVPLLYLCRYLLIIRRGVIIVSGAGAGGGRLRRFRLGAAMAIIYMAMSTLFFLRIAPLTPWWGALAAWVMILLIVEAIFAFFFPYRCCFNETEGDVQNNPHV |
| CX427 | MELITACALLLIAVHIAIVISCPEPLAAVALVDVPLLYLYRYLLIIRRGVIIVSGAGGGGGRLRRFRLGAAMAIIYMAMSTLFFLRIAPLTPWWGALAAWVMILLIVEAIFAFFFPYRCCFNETEGDVQNNPHV |
| CX428 | MELITACALLLIAVHIAIVISCPEPLAAVALVDVPLLYLYRYLLIIRRGVIIVSGAGGGGGRLRRFRLGAAMAIIYMAMSTLFFLRIAPLTPWWGALAAWVMILLIVEAIFAFFFPYRCCFNETEGDVQNNPHV |
| CX429 | MELITACALLLIAVHIAIVISCPEPLAAVALVDVPLLYLYRYLLIIRRGVIIVSGAGGGGGRLRRFRLGAAMAIIYMAMSTLFFLRIAPLTPWWGALAAWVMILLIVEAIFAFFFPYRCCFNETEGDVQNNPHV |
| CX431 | MELITACALLLIAVHIAIVISCPEPLAAVALVDVPLLYLCRYLLIIRRGVIIVSGGGGGGRLRRFRLGAAMAIIYMAMSTLFFLRIAPLTPWWGALAAWVMILLIVEAIFAFFFPYRCCFNETEGDVQNNPHV |
| CX534 | MELITACALLLIAVHIAIVISCPEPLAAVALVDVPLLYLCRYLLIIRRGVIIVSGAGAGGGRLRRFRLGAAMAIIYMAMSTLFFLRIAPLTPWWGALAAWVMILLIVEAIFAFFFPYRCCFNETEGDVQNNPHV |
| CX542 | MELITACALLLIAVHIAIVISCPEPLAAVALVDVPLLYLCRYLLIIRRGVIIVSGGGGGGRLRRFRLGAAMAVIYMAMSTLFFLRIAPLTPWWGALAAWVMILLIVEAIFAFFFPYRCCFNETEGDVQNNPHV |
| CX548 | MELITACALLLIAVHIAIVISCPEPLAAVALVDVPLLYLCRYLLIIRRGVIIVSGGGAGGGRLRRFRLGAAMAIIYMAMSTLFFLRIAPLTPWWGALAAWVMILLIVEAIFAFFFPYRCCFNETEGDVQNNPHV |
| CX561 | MELITACALLLIAVHIAIVISCPEPLAAVALVDVPLLYLCRYLLIIRRGVIIVSGAGGGGRLRRFRLGAAMAIIYMAMSTLFFLRIAPLTPWWGALAAWVMILLIVEAIFAFFFPYRCCFNETEGDVQNNPHV |
| CX578 | MELITACALLLIAVHIAIVISCPEPLAAVALVDVPLLYLCRYLLIIRRGVIIVSGAGAGGGRLRRFRLGAAMAIIYMAMSTLFFLRIAPLTPWWGALAAWVMILLIVEAIFAFFFPYRCCFNETEGDVQNNPHV |
| CX579 | MELITACALLLIAVHIAIVISCPEPLAAVALVDVPLLYLCRYLLIIRRGVIIVSGGGGGGRLRRFRLGAAMAIIYMAMSTLFFLRIAPLTPWWGALAAWVMILLIVEAIFAFFFPYRCCFNETEGDVQNNPHV |
| IRIS_313-7620 | MELITACALLLIAVHIAIVISCPEPLAAVALVDVPLLYLYRYLLIIRRGVIIVSGAGGGGGRLRRFRLGAAMAIIYMAMSTLFFLRIAPLTPWWGALAAWVMILLIVEAIFAFFFPYRCCFNETEGDVQNNPHV |
| IRIS_313-7635 | MELITACALLLIAVHIAIVISCPEPLAAVALVDVPLLYLCRYLLIIRRGVIIVSGGGGGGRLRRFRLGAAMAIIYMAMSTLFFLRIAPLTPWWGALAAWVMILLIVEAIFAFFFPYRCCFNETEGDVQNNPHV |
| IRIS_313-7636 | MELITACALLLIAVHIAIVISCPEPLAAVALVDVPLLYLCRYLLIIRRGVIIVSGGGGGGRLRRFRLGAAMAIIYMAMSTLFFLRIAPLTPWWGALAAWVMILLIVEAIFAFFFPYRCCFNETEGDVQNNPHV |
| IRIS_313-7638 | MELITACALLLIAVHIAIVISCPEPLAAVALVDVPLLYLCRYLLIIRRGVIIVSGAGAGGGRLRRFRLGAAMAIIYMAMSTLFFLRIAPLTPWWGALAAWVMILLIVEAIFAFFFPYRCCFNETEGDVQNNPHV |
| IRIS_313-7641 | MELITACALLLIAVHIAIVISCPEPLAAVALVDVPLLYLCRYLLIIRRGVIIVSGAGAGGGRLRRFRLGAAMAIIYMAMSTLFFLRIAPLTPWWGALAAWVMILLIVEAIFAFFFPYRCCFNETEGDVQNNPHV |
| IRIS_313-7646 | MELITACALLLIAVHIAIVISCPEPLAAVALVDVPLLYLCRYLLIIRRGVIIVSGAGAGGGRLRRFRLGAAMAIIYMAMSTLFFLRIAPLTPWWGALAAWVMILLIVEAIFAFFFPYRCCFNETEGDVQNNPHV |
| IRIS_313-7650 | MELITACALLLIAVHIAIVISCPEPLAAVALVDVPLLYLCRYLLIIRRGVIIVSGAGAGGGRLRRFRLGAAMAIIYMAMSTLFFLRIAPLTPWWGALAAWVMILLIVEAIFAFFFPYRCCFNETEGDVQNNPHV |
| IRIS_313-7651 | MELITACALLLIAVHIAIVISCPEPLAAVALVDVPLLYLCRYLLIIRRGVIIVSGAGAGGGRLRRFRLGAAMAIIYMAMSTLFFLRIAPLTPWWGALAAWVMILLIVEAIFAFFFPYRCCFNETEGDVQNNPHV |
| IRIS_313-7654 | MELITACALLLIAVHIAIVISCPEPLAAVALVDVPLLYLCRYLLIIRRGVIIVSGGGGGGRLRRFRLGAAMAIIYMAMSTLFFLRIAPLTPWWGALAAWVMILLIVEAIFAFFFPYRCCFNETEGDVQNNPHV |
| IRIS_313-7664 | MELITACALLLIAVHIAIVISCPEPLAAVALVDVPLLYLCRYLLIIRRGVIIVSGAGAGGGRLRRFRLGAAMAIIYMAMSTLFFLRIAPLTPWWGALAAWVMILLIVEAIFAFFFPYRCCFNETEGDVQNNPHV |
| IRIS_313-7665 | MELITACALLLIAVHIAIVISCPEPLAAVALVDVPLLYLCRYLLIIRRGVIIVSGGGGGGRLRRFRLGAAMAIIYMAMSTLFFLRIAPLTPWWGALAAWVMILLIVEAIFAFFFPYRCCFNETEGDVQNNPHV |
| IRIS_313-7668 | MELITACALLLIAVHIAIVISCPEPLAAVALVDVPLLYLCRYLLIIRRGVIIVSGGGGGGRLRRFRLGAAMAIIYMAMSTLFFLRIAPLTPWWGALAAWVMILLIVEAIFAFFFPYRCCFNETEGDVQNNPHV |
| IRIS_313-7681 | MELITACALLLIAVHIAIVISCPEPLAAVALVDVPLLYLCRYLLIIRRGVIIVSGGGGGGRLRRFRLGAAMAIIYMAMSTLFFLRIAPLTPWWGALAAWVMILLIVEAIFAFFFPYRCCFNETEGDVQNNPHV |
| IRIS_313-7684 | MELITACALLLIAVHIAIVISCPEPLAAVALVDVPLLYLCRYLLIIRRGVIIVSGGGGGGRLRRFRLGAAMAIIYMAMSTLFFLRIAPLTPWWGALAAWVMILLIVEAIFAFFFPYRCCFNETEGDVQNNPHV |
| IRIS_313-7685 | MELITACALLLIAVHIAIVISCPEPLAAVALVDVPLLYLCRYLLIIRRGVIIVSGGGGGGRLRRFRLGAAMAIIYMAMSTLFFLRIAPLTPWWGALAAWVMILLIVEAIFAFFFPYRCCFNETEGDVQNNPHV |
| IRIS_313-7688 | MELITACALLLIAVHIAIVISCPEPLAAVALVDVPLLYLCRYLLIIRRGVIIVSGGGGGGRLRRFRLGAAMAIIYMAMSTLFFLRIAPLTPWWGALAAWVMILLIVEAIFAFFFPYRCCFNETEGDVQNNPHV |
| IRIS_313-7689 | MELITACALLLIAVHIAIVISCPEPLAAVALVDVPLLYLCRYLLIIRRGVIIVSGGGGGGRLRRFRLGAAMAIIYMAMSTLFFLRIAPLTPWWGALAAWVMILLIVEAIFAFFFPYRCCFNETEGDVQNNPHV |
| IRIS_313-7690 | MELITACALLLIAVHIAIVISCPEPLAAVALVDVPLLYLCRYLLIIRRGVIIVSGGGGGGRLRRFRLGAAMAIIYMAMSTLFFLRIAPLTPWWGALAAWVMILLIVEAIFAFFFPYRCCFNETEGDVQNNPHV |
| IRIS_313-7691 | MELITACALLLIAVHIAIVISCPEPLAAVALVDVPLLYLCRYLLIIRRGVIIVSGGGGGGRLRRFRLGAAMAIIYMAMSTLFFLRIAPLTPWWGALAAWVMILLIVEAIFAFFFPYRCCFNETEGDVQNNPHV |
| IRIS_313-7696 | MELITACALLLIAVHIAIVISCPEPLAAVALVDVPLLYLCRYLLIIRRGVIIVSGGGAGGGRLRRFRLGAAMAIIYMAMSTLFFLRIAPLTPWWGALAAWVMILLIVEAIFAFFFPYRCCFNETEGDVQNNPHV |
| IRIS_313-7698 | MELITACALLLIAVHIAIVISCPEPLAAVALVDVPLLYLCRYLLIIRRGVIIVSGGGGGGRLRRFRLGAAMAIIYMAMSTLFFLRIAPLTPWWGALAAWVMILLIVEAIFAFFFPYRCCFNETEGDVQNNPHV |
| IRIS_313-7699 | MELITACALLLIAVHIAIVISCPEPLAAVALVDVPLLYLCRYLLIIRRGVIIVSGGGGGGRLRRFRLGAAMAIIYMAMSTLFFLRIAPLTPWWGALAAWVMILLIVEAIFAFFFPYRCCFNETEGDVQNNPHV |
| IRIS_313-7719 | MELITACALLLIAVHIAIVISCPEPLAAVALVDVPLLYLYRYLLIIRRGVIIVSGAGGGGGRLRRFRLGAAMAIIYMAMSTLFFLRIAPLTPWWGALAAWVMILLIVEAIFAFFFPYRCCFNETEGDVQNNPHV |
| IRIS_313-7720 | MELITACALLLIAVHIAIVISCPEPLAAVALVDVPLLYLCRYLLIIRRGVIIVSGAGGAAAAGCAASASGQPWRSSTWPCRPSSSSALLLSRHGGAPSPPGS*SCSSSRPSSPSSSPTAAASTKPKAMSRIIPM |
| IRIS_313-7722 | MELITACALLLIAVHIAIVISCPEPLAAVALVDVPLLYLCRYLLIIRRGVIIVSGAGAGGGRLRRFRLGAAMAIIYMAMSTLFFLRIAPLTPWWGALAAWVMILLIVEAIFAFFFPYRCCFNETEGDVQNNPHV |
| IRIS_313-7725 | MELITACALLLIAVHIAIVISCPEPLAAVALVDVPLLYLCRYLLIIRRGVIIVSGAGAGGGRLRRFRLGAAMAIIYMAMSTLFFLRIAPLTPWWGALAAWVMILLIVEAIFAFFFPYRCCFNETEGDVQNNPHV |
| IRIS_313-7728 | MELITACALLLIAVHIAIVISCPEPLAAVALVDVPLLYLCRYLLIIRRGVIIVSGAGGGGGRLRRFRLGAAMAIIYMAMSTLFFLRIAPLTPWWGALAAWVMILLIVEAIFAFFFPYRCCFNETEGDVQNNPHV |
| IRIS_313-7736 | MELITACALLLIAVHIAIVISCPEPLAAVALVDVPLLYLCRYLLIIRRGVIIVSGGGAGGGRLRRFRLGAAMAIIYMAMSTLFFLRIAPLTPWWGALAAWVMILLIVEAIFAFFFPYRCCFNETEGDVQNNPHV |
| IRIS_313-7758 | MELITACALLLIAVHIAIVISCPEPLAAVALVDVPLLYLCRYLLIIRRGVIIVSGAGAGGGRLRRFRLGAAMAIIYMAMSTLFFLRIAPLTPWWGALAAWVMILLIVEAIFAFFFPYRCCFNETEGDVQNNPHV |
| IRIS_313-7766 | MELITACALLLIAVHIAIVISCPEPLAAVALVDVPLLYLCRYLLIIRRGVIIVSGGGGGGRLRRFRLGAAMAIIYMAMSTLFFLRIAPLTPWWGALAAWVMILLIVEAIFAFFFPYRCCFNETEGDVQNNPHV |
| IRIS_313-7769 | MELITACALLLIAVHIAIVISCPEPLAAVALVDVPLLYLCRYLLIIRRGVIIVSAGGGGGGRLRRFRLGAAMAIIYMAMSTLFFLRIAPLTPWWGALAAWVMILLIVEAIFAFFFPYRCCFNETEGDVQNNPHV |
| IRIS_313-7770 | MELITACALLLIAVHIAIVISCPEPLAAVALVDVPLLYLCRYLLIIRRGVIIVSGGGGGGRLRRFRLGAAMAIIYMAMSTLFFLRIAPLTPWWGALAAWVMILLIVEAIFAFFFPYRCCFNETEGDVQNNPHV |
| IRIS_313-7773 | MELITACALLLIAVHIAIVISCPEPLAAVALVDVPLLYLCRYLLIIRRGVIIVSGAGAGGGRLRRFRLGAAMAIIYMAMSTLFFLRIAPLTPWWGALAAWVMILLIVEAIFAFFFPYRCCFNETEGDVQNNPHV |
| IRIS_313-7778 | MELITACALLLIAVHIAIVISCPEPLAAVALVDVPLLYLYRYLLIIRRGVIIVSGAGGGGGRLRRFRLGAAMAIIYMAMSTLFFLRIAPLTPWWGALAAWVMILLIVEAIFAFFFPYRCCFNETEGDVQNNPHV |
| IRIS_313-7780 | MELITACALLLIAVHIAIVISCPEPLAAVALVDVPLLYLCRYLLIIRRGVIIVSGGGGGGRLRRFRLGAAMAIIYMAMSTLFFLRIAPLTPWWGALAAWVMILLIVEAIFAFFFPYRCCFNETEGDVQNNPHV |
| IRIS_313-7792 | MELITACALLLIAVHIAIVISCPEPLAAVALVDVPLLYLCRYLLIIRRGVIIVSGGGGGGRLRRFRLGAAMAIIYMAMSTLFFLRIAPLTPWWGALAAWVMILLIVEAIFAFFFPYRCCFNETEGDVQNNPHV |
| IRIS_313-7793 | MELITACALLLIAVHIAIVISCPEPLAAVALVDVPLLYLCRYLLIIRRGVIIVSGAGAGGGRLRRFRLGAAMAIIYMAMSTLFFLRIAPLTPWWGALAAWVMILLIVEAIFAFFFPYRCCFNETEGDVQNNPHV |
| IRIS_313-7795 | MELITACALLLIAVHIAIVISCPEPLAAVALVDVPLLYLCRYLLIIRRGVIIVSGAGAGGGRLRRFRLGAAMAIIYMAMSTLFFLRIAPLTPWWGALAAWVMILLIVEAIFAFFFPYRCCFNETEGDVQNNPHV |
| IRIS_313-7797 | MELITACALLLIAVHIAIVISCPEPLAAVALVDVPLLYLCRYLLIIRRGVIIVSGGGGGGRLRRFRLGAAMAIIYMAMSTLFFLRIAPLTPWWGALAAWVMILLIVEAIFAFFFPYRCCFNETEGDVQNNPHV |
| IRIS_313-7799 | MELITACALLLIAVHIAIVISCPEPLAAVALVDVPLLYLCRYLLIIRRGVIIVSGAGGAAAAGCAASASGQPWRSSTWPCRPSSSSALLLSRHGGAPSPPGS*SCSSSRPSSPSSSPTAAASTKPKAMSRIIPM |
| IRIS_313-7807 | MELITACALLLIAVHIAIVISCPEPLAAVALVDVPLLYLCRYLLIIRRGVIIVSGGGGGGRLRRFRLGAAMAIIYMAMSTLFFLRIAPLTPWWGALAAWVMILLIVEAIFAFFFPYRCCFNETEGDVQNNPHV |
| IRIS_313-7808 | MELITACALLLIAVHIAIVISCPEPLAAVALVDVPLLYLCRYLLIIRRGVIIVSGGGGGGRLRRFRLGAAMAIIYMAMSTLFFLRIAPLTPWWGALAAWVMILLIVEAIFAFFFPYRCCFNETEGDVQNNPHV |
| IRIS_313-7809 | MELITACALLLIAVHIAIVISCPEPLAAVALVDVPLLYLCRYLLIIRRGVIIVSGGGGGGRLRRFRLGAAMAIIYMAMSTLFFLRIAPLTPWWGALAAWVMILLIVEAIFAFFFPYRCCFNETEGDVQNNPHV |
| IRIS_313-7815 | MELITACALLLIAVHIAIVISCPEPLAAVALVDVPLLYLYRYLLIIRRGVIIVSGAGGGGGRLRRFRLGAAMAIIYMAMSTLFFLRIAPLTPWWGALAAWVMILLIVEAIFAFFFPYRCCFNETEGDVQNNPHV |
| IRIS_313-7816 | MELITACALLLIAVHIAIVISCPEPLAAVALVDVPLLYLCRYLLIIRRGVIIVSGGGGGGRLRRFRLGAAMAIIYMAMSTLFFLRIAPLTPWWGALAAWVMILLIVEAIFAFFFPYRCCFNETEGDVQNNPHV |
| IRIS_313-7819 | MELITACALLLIAVHIAIVISCPEPLAAVALVDVPLLYLCRYLLIIRRGVIIVSGGGGGGRLRRFRLGAAMAIIYMAMSTLFFLRIAPLTPWWGALAAWVMILLIVEAIFAFFFPYRCCFNETEGDVQNNPHV |
| IRIS_313-7820 | MELITACALLLIAVHIAIVISCPEPLAAVALVDVPLLYLCRYLLIIRRGVIIVSGGGGGGRLRRFRLGAAMAIIYMAMSTLFFLRIAPLTPWWGALAAWVMILLIVEAIFAFFFPYRCCFNETEGDVQNNPHV |

| | |
|---|---|
| IRIS_313-7824 | MELITACALLLIAVHIAIVISCPEPLAAVALVDVPLLYLCRYLLIIRRGVIIVSGGGGGGRLRRFRLGAAMAIIYMAMSTLFFLRIAPLTPWWGALAAWVMILLIVEAIFAFFFPYRCCFNETEGDVQNNPHV |
| IRIS_313-7826 | MELITACALLLIAVHIAIVISCPEPLAAVALVDVPLLYLCRYLLIIRRGVIIVSGGGGGGRLRRFRLGAAMAIIYMAMSTLFFLRIAPLTPWWGALAAWVMILLIVEAIFAFFFPYRCCFNETEGDVQNNPHV |
| IRIS_313-7832 | MELITACALLLIAVHIAIVISCPEPLAAVALVDVPLLYLCRYLLIIRRGVIIVSGGGGGGRLRRFRLGAAMAIIYMAMSTLFFLRIAPLTPWWGALAAWVMILLIVEAIFAFFFPYRCCFNETEGDVQNNPHV |
| IRIS_313-7838 | MELITACALLLIAVHIAIVISCPEPLAAVALVDVPLLYLCRYLLIIRRGVIIVSGAGAGGGRLRRFRLGAAMAIIYMAMSTLFFLRIAPLTPWWGALAAWVMILLIVEAIFAFFFPYRCCFNETEGDVQNNPHV |
| IRIS_313-7850 | MELITACALLLIAVHIAIVISCPEPLAAVALVDVPLLYLCRYLLIIRRGVIIVSGAGAGGGRLRRFRLGAAMAIIYMAMSTLFFLRIAPLTPWWGALAAWVMILLIVEAIFAFFFPYRCCFNETEGDVQNNPHV |
| IRIS_313-7856 | MELITACALLLIAVHIAIVISCPEPLAAVALVDVPLLYLYRYLLIIRRGVIIVSGAGGGGGRLRRFRLGAAMAIIYMAMSTLFFLRIAPLTPWWGALAAWVMILLIVEAIFAFFFPYRCCFNETEGDVQNNPHV |
| IRIS_313-7859 | MELITACALLLIAVHIAIVISCPEPLAAVALVDVPLLYLCRYLLIIRRGVIIVSGAGGGGGRLRRFRLGAAMAIIYMAMSTLFFLRIAPLTPWWGALAAWVMILLIVEAIFAFFFPYRCCFNETEGDVQNNPHV |
| IRIS_313-7863 | MELITACALLLIAVHIAIVISCPEPLAAVALVDVPLLYLCRYLLIIRRGVIIVSGAGAGGGRLRRFRLGAAMAIIYMAMSTLFFLRIAPLTPWWGALAAWVMILLIVEAIFAFFFPYRCCFNETEGDVQNNPHV |
| IRIS_313-7866 | MELITACALLLIAVHIAIVISCPEPLAAVALVDVPLLYLCRYLLIIRRGVIIVSGGGGGGRLRRFRLGAAMAIIYMAMSTLFFLRIAPLTPWWGALAAWVMILLIVEAIFAFFFPYRCCFNETEGDVQNNPHV |
| IRIS_313-7868 | MELITACALLLIAVHIAIVISCPEPLAAVALVDVPLLYLCRYLLIIRRGVIIVSGAGAGGGRLRRFRLGAAMAIIYMAMSTLFFLRIAPLTPWWGALAAWVMILLIVEAIFAFFFPYRCCFNETEGDVQNNPHV |
| IRIS_313-7870 | MELITACALLLIAVHIAIVISCPEPLAAVALVDVPLLYLCRYLLIIRRGVIIVSGAGGGGGRLRRFRLGAAMAIIYMAMSTLFFLRIAPLTPWWGALAAWVMILLIVEAIFAFFFPYRCCFNETEGDVQNNPHV |
| IRIS_313-7876 | MELITACALLLIAVHIAIVISCPEPLAAVALVDVPLLYLYRYLLIIRRGVIIVSGAGGGGGRLRRFRLGAAMAIIYMAMSTLFFLRIAPLTPWWGALAAWVMILLIVEAIFAFFFPYRCCFNETEGDVQNNPHV |
| IRIS_313-7883 | MELITACALLLIAVHIAIVISCPEPLAAVALVDVPLLYLYRYLLIIRRGVIIVSGAGGGGGRLRRFRLGAAMAIIYMAMSTLFFLRIAPLTPWWGALAAWVMILLIVEAIFAFFFPYRCCFNETEGDVQNNPHV |
| IRIS_313-7885 | MELITACALLLIAVHIAIVISCPEPLAAVALVDVPLLYLYRYLLIIRRGVIIVSGAGGGGGRLRRFRLGAAMAIIYMAMSTLFFLRIAPLTPWWGALAAWVMILLIVEAIFAFFFPYRCCFNETEGDVQNNPHV |
| IRIS_313-7902 | MELITACALLLIAVHIAIVISCPEPLAAVALVDVPLLYLCRYLLIIRRGVIIVSGAGAGGGRLRRFRLGAAMAIIYMAMSTLFFLRIAPLTPWWGALAAWVMILLIVEAIFAFFFPYRCCFNETEGDVQNNPHV |
| IRIS_313-7907 | MELITACALLLIAVHIAIVISCPEPLAAVALVDVPLLYLCRYLLIIRRGVIIVSGAGAGGGRLRRFRLGAAMAIIYMAMSTLFFLRIAPLTPWWGALAAWVMILLIVEAIFAFFFPYRCCFNETEGDVQNNPHV |
| IRIS_313-7909 | MELITACALLLIAVHIAIVISCPEPLAAVALVDVPLLYLCRYLLIIRRGVIIVSGAGAGGGRLRRFRLGAAMAIIYMAMSTLFFLRIAPLTPWWGALAAWVMILLIVEAIFAFFFPYRCCFNETEGDVQNNPHV |
| IRIS_313-7911 | MELITACALLLIAVHIAIVISCPEPLAAVALVDVPLLYLCRYLLIIRRGVIIVSGGGGGGRLRRFRLGAAMAIIYMAMSTLFFLRIAPLTPWWGALAAWVMILLIVEAIFAFFFPYRCCFNETEGDVQNNPHV |
| IRIS_313-7912 | MELITACALLLIAVHIAIVISCPEPLAAVALVDVPLLYLCRYLLIIRRGVIIVSGAGAGGGRLRRFRLGAAMAIIYMAMSTLFFLRIAPLTPWWGALAAWVMILLIVEAIFAFFFPYRCCFNETEGDVQNNPHV |
| IRIS_313-7914 | MELITACALLLIAVHIAIVISCPEPLAAVALVDVPLLYLCRYLLIIRRGVIIVSGAGAGGGRLRRFRLGAAMAIIYMAMSTLFFLRIAPLTPWWGALAAWVMILLIVEAIFAFFFPYRCCFNETEGDVQNNPHV |
| IRIS_313-7922 | MELITACALLLIAVHIAIVISCPEPLAAVALVDVPLLYLCRYLLIIRRGVIIVSGAGAGGGRLRRFRLGAAMAIIYMAMSTLFFLRIAPLTPWWGALAAWVMILLIVEAIFAFFFPYRCCFNETEGDVQNNPHV |
| IRIS_313-7924 | MELITACALLLIAVHIAIVISCPEPLAAVALVDVPLLYLCRYLLIIRRGVIIVSGAGAGGGRLRRFRLGAAMAIIYMAMSTLFFLRIAPLTPWWGALAAWVMILLIVEAIFAFFFPYRCCFNETEGDVQNNPHV |
| IRIS_313-7933 | MELITACALLLIAVHIAIVISCPEPLAAVALVDVPLLYLCRYLLIIRRGVIIVSGAGAGGGRLRRFRLGAAMAIIYMAMSTLFFLRIAPLTPWWGALAAWVMILLIVEAIFAFFFPYRCCFNETEGDVQNNPHV |
| IRIS_313-7959 | MELITACALLLIAVHIAIVISCPEPLAAVALVDVPLLYLCRYLLIIRRGVIIVSGAGAGGGRLRRFRLGAAMAIIYMAMSTLFFLRIAPLTPWWGALAAWVMILLIVEAIFAFFFPYRCCFNETEGDVQNNPHV |
| IRIS_313-7992 | MELITACALLLIAVHIAIVISCPEPLAAVALVDVPLLYLCRYLLIIRRGVIIVSGAGAGGGRLRRFRLGAAMAIIYMAMSTLFFLRIAPLTPWWGALAAWVMILLIVEAIFAFFFPYRCCFNETEGDVQNNPHV |
| IRIS_313-7993 | MELITACALLLIAVHIAIVISCPEPLAAVALVDVPLLYLCRYLLIIRRGVIIVSGAGAGGGRLRRFRLGAAMAIIYMAMSTLFFLRIAPLTPWWGALAAWVMILLIVEAIFAFFFPYRCCFNETEGDVQNNPHV |
| IRIS_313-7994 | MELITACALLLIAVHIAIVISCPEPLAAVALVDVPLLYLYRYLLIIRRGVIIVSGAGGGGGRLRRFRLGAAMAIIYMAMSTLFFLRIAPLTPWWGALAAWVMILLIVEAIFAFFFPYRCCFNETEGDVQNNPHV |
| IRIS_313-8003 | MELITACALLLIAVHIAIVISCPEPLAAVALVDVPLLYLCRYLLIIRRGVIIVSGAGAGGGRLRRFRLGAAMAIIYMAMSTLFFLRIAPLTPWWGALAAWVMILLIVEAIFAFFFPYRCCFNETEGDVQNNPHV |
| IRIS_313-8010 | MELITACALLLIAVHIAIVISCPEPLAAVALVDVPLLYLCRYLLIIRRGVIIVSGAGAGGGRLRRFRLGAAMAIIYMAMSTLFFLRIAPLTPWWGALAAWVMILLIVEAIFAFFFPYRCCFNETEGDVQNNPHV |
| IRIS_313-8011 | MELITACALLLIAVHIAIVISCPEPLAAVALVDVPLLYLCRYLLIIRRGVIIVSGAGGGGGRLRRFRLGAAMAIIYMAMSTLFFLRIAPLTPWWGALAAWVMILLIVEAIFAFFFPYRCCFNETEGDVQNNPHV |
| IRIS_313-8023 | MELITACALLLIAVHIAIVISCPEPLAAVALVDVPLLYLCRYLLIIRRGVIIVSGAGAGGGRLRRFRLGAAMAIIYMAMSTLFFLRIAPLTPWWGALAAWVMILLIVEAIFAFFFPYRCCFNETEGDVQNNPHV |
| IRIS_313-8024 | MELITACALLLIAVHIAIVISCPEPLAAVALVDVPLLYLYRYLLIIRRGVIIVSGAGGGGGRLRRFRLGAAMAIIYMAMSTLFFLRIAPLTPWWGALAAWVMILLIVEAIFAFFFPYRCCFNETEGDVQNNPHV |
| IRIS_313-8025 | MELITACALLLIAVHIAIVISCPEPLAAVALVDVPLLYLYRYLLIIRRGVIIVSGAGGGGGRLRRFRLGAAMAIIYMAMSTLFFLRIAPLTPWWGALAAWVMILLIVEAIFAFFFPYRCCFNETEGDVQNNPHV |
| IRIS_313-8026 | MELITACALLLIAVHIAIVISCPEPLAAVALVDVPLLYLCRYLLIIRRGVIIVSGAGAGGGRLRRFRLGAAMAIIYMAMSTLFFLRIAPLTPWWGALAAWVMILLIVEAIFAFFFPYRCCFNETEGDVQNNPHV |
| IRIS_313-8027 | MELITACALLLIAVHIAIVISCPEPLAAVALVDVPLLYLYRYLLIIRRGVIIVSGAGGGGGRLRRFRLGAAMAIIYMAMSTLFFLRIAPLTPWWGALAAWVMILLIVEAIFAFFFPYRCCFNETEGDVQNNPHV |
| IRIS_313-8029 | MELITACALLLIAVHIAIVISCPEPLAAVALVDVPLLYLYRYLLIIRRGVIIVSGAGGGGGRLRRFRLGAAMAIIYMAMSTLFFLRIAPLTPWWGALAAWVMILLIVEAIFAFFFPYRCCFNETEGDVQNNPHV |
| IRIS_313-8031 | MELITACALLLIAVHIAIVISCPEPLAAVALVDVPLLYLYRYLLIIRRGVIIVSGAGGGGGRLRRFRLGAAMAIIYMAMSTLFFLRIAPLTPWWGALAAWVMILLIVEAIFAFFFPYRCCFNETEGDVQNNPHV |
| IRIS_313-8032 | MELITACALLLIAVHIAIVISCPEPLAAVALVDVPLLYLYRYLLIIRRGVIIVSGAGGGGGRLRRFRLGAAMAIIYMAMSTLFFLRIAPLTPWWGALAAWVMILLIVEAIFAFFFPYRCCFNETEGDVQNNPHV |
| IRIS_313-8033 | MELITACALLLIAVHIAIVISCPEPLAAVALVDVPLLYLCRYLLIIRRGVIIVSGAGAGGGRLRRFRLGAAMAIIYMAMSTLFFLRIAPLTPWWGALAAWVMILLIVEAIFAFFFPYRCCFNETEGDVQNNPHV |
| IRIS_313-8037 | MELITACALLLIAVHIAIVISCPEPLAAVALVDVPLLYLCRYLLIIRRGVIIVSGAGAGGGRLRRFRLGAAMAIIYMAMSTLFFLRIAPLTPWWGALAAWVMILLIVEAIFAFFFPYRCCFNETEGDVQNNPHV |
| IRIS_313-8039 | MELITACALLLIAVHIAIVISCPEPLAAVALVDVPLLYLCRYLLIIRRGVIIVSGAGAGGGRLRRFRLGAAMAIIYMAMSTLFFLRIAPLTPWWGALAAWVMILLIVEAIFAFFFPYRCCFNETEGDVQNNPHV |
| IRIS_313-8041 | MELITACALLLIAVHIAIVISCPEPLAAVALVDVPLLYLYRYLLIIRRGVIIVSGAGGGGGRLRRFRLGAAMAIIYMAMSTLFFLRIAPLTPWWGALAAWVMILLIVEAIFAFFFPYRCCFNETEGDVQNNPHV |
| IRIS_313-8044 | MELITACALLLIAVHIAIVISCPEPLAAVALVDVPLLYLYRYLLIIRRGVIIVSGAGGGGGRLRRFRLGAAMAIIYMAMSTLFFLRIAPLTPWWGALAAWVMILLIVEAIFAFFFPYRCCFNETEGDVQNNPHV |
| IRIS_313-8046 | MELITACALLLIAVHIAIVISCPEPLAAVALVDVPLLYLYRYLLIIRRGVIIVSGAGGGGGRLRRFRLGAAMAIIYMAMSTLFFLRIAPLTPWWGALAAWVMILLIVEAIFAFFFPYRCCFNETEGDVQNNPHV |
| IRIS_313-8048 | MELITACALLLIAVHIAIVISCPEPLAAVALVDVPLLYLYRYLLIIRRGVIIVSGAGGGGGRLRRFRLGAAMAIIYMAMSTLFFLRIAPLTPWWGALAAWVMILLIVEAIFAFFFPYRCCFNETEGDVQNNPHV |
| IRIS_313-8049 | MELITACALLLIAVHIAIVISCPEPLAAVALVDVPLLYLYRYLLIIRRGVIIVSGAGGGGGRLRRFRLGAAMAIIYMAMSTLFFLRIAPLTPWWGALAAWVMILLIVEAIFAFFFPYRCCFNETEGDVQNNPHV |
| IRIS_313-8050 | MELITACALLLIAVHIAIVISCPEPLAAVALVDVPLLYLYRYLLIIRRGVIIVSGAGGGGGRLRRFRLGAAMAIIYMAMSTLFFLRIAPLTPWWGALAAWVMILLIVEAIFAFFFPYRCCFNETEGDVQNNPHV |
| IRIS_313-8052 | MELITACALLLIAVHIAIVISCPEPLAAVALVDVPLLYLYRYLLIIRRGVIIVSGAGGGGGRLRRFRLGAAMAIIYMAMSTLFFLRIAPLTPWWGALAAWVMILLIVEAIFAFFFPYRCCFNETEGDVQNNPHV |
| IRIS_313-8053 | MELITACALLLIAVHIAIVISCPEPLAAVALVDVPLLYLYRYLLIIRRGVIIVSGAGGGGGRLRRFRLGAAMAIIYMAMSTLFFLRIAPLTPWWGALAAWVMILLIVEAIFAFFFPYRCCFNETEGDVQNNPHV |
| IRIS_313-8057 | MELITACALLLIAVHIAIVISCPEPLAAVALVDVPLLYLYRYLLIIRRGVIIVSGAGGGGGRLRRFRLGAAMAIIYMAMSTLFFLRIAPLTPWWGALAAWVMILLIVEAIFAFFFPYRCCFNETEGDVQNNPHV |
| IRIS_313-8058 | MELITACALLLIAVHIAIVISCPEPLAAVALVDVPLLYLYRYLLIIRRGVIIVSGAGGGGGRLRRFRLGAAMAIIYMAMSTLFFLRIAPLTPWWGALAAWVMILLIVEAIFAFFFPYRCCFNETEGDVQNNPHV |
| IRIS_313-8060 | MELITACALLLIAVHIAIVISCPEPLAAVALVDVPLLYLCRYLLIIRRGVIIVSGAGGGGGRLRRFRLGAAMAIIYMAMSTLFFLRIAPLTPWWGALAAWVMILLIVEAIFAFFFPYRCCFNETEGDVQNNPHV |
| IRIS_313-8061 | MELITACALLLIAVHIAIVISCPEPLAAVALVDVPLLYLYRYLLIIRRGVIIVSGAGGGGGRLRRFRLGAAMAIIYMAMSTLFFLRIAPLTPWWGALAAWVMILLIVEAIFAFFFPYRCCFNETEGDVQNNPHV |
| IRIS_313-8062 | MELITACALLLIAVHIAIVISCPEPLAAVALVDVPLLYLCRYLLIIRRGVIIVSGAGAGGGRLRRFRLGAAMAIIYMAMSTLFFLRIAPLTPWWGALAAWVMILLIVEAIFAFFFPYRCCFNETEGDVQNNPHV |
| IRIS_313-8063 | MELITACALLLIAVHIAIVISCPEPLAAVALVDVPLLYLCRYLLIIRRGVIIVSGAGAGGGRLRRFRLGAAMAIIYMAMSTLFFLRIAPLTPWWGALAAWVMILLIVEAIFAFFFPYRCCFNETEGDVQNNPHV |
| IRIS_313-8064 | MELITACALLLIAVHIAIVISCPEPLAAVALVDVPLLYLCRYLLIIRRGVIIVSGAGAGGGRLRRFRLGAAMAIIYMAMSTLFFLRIAPLTPWWGALAAWVMILLIVEAIFAFFFPYRCCFNETEGDVQNNPHV |
| IRIS_313-8065 | MELITACALLLIAVHIAIVISCPEPLAAVALVDVPLLYLYRYLLIIRRGVIIVSGAGGGGGRLRRFRLGAAMAIIYMAMSTLFFLRIAPLTPWWGALAAWVMILLIVEAIFAFFFPYRCCFNETEGDVQNNPHV |
| IRIS_313-8066 | MELITACALLLIAVHIAIVISCPEPLAAVALVDVPLLYLCRYLLIIRRGVIIVSGAGAGGGRLRRFRLGAAMAIIYMAMSTLFFLRIAPLTPWWGALAAWVMILLIVEAIFAFFFPYRCCFNETEGDVQNNPHV |
| IRIS_313-8068 | MELITACALLLIAVHIAIVISCPEPLAAVALVDVPLLYLCRYLLIIRRGVIIVSGGGGGGRLRRFRLGAAMAIIYMAMSTLFFLRIAPLTPWWGALAAWVMILLIVEAIFAFFFPYRCCFNETEGDVQNNPHV |
| IRIS_313-8069 | MELITACALLLIAVHIAIVISCPEPLAAVALVDVPLLYLCRYLLIIRRGVIIVSGGGGGGRLRRFRLGAAMAIIYMAMSTLFFLRIAPLTPWWGALAAWVMILLIVEAIFAFFFPYRCCFNETEGDVQNNPHV |
| IRIS_313-8072 | MELITACALLLIAVHIAIVISCPEPLAAVALVDVPLLYLCRYLLIIRRGVIIVSGAGAGGGRLRRFRLGAAMAIIYMAMSTLFFLRIAPLTPWWGALAAWVMILLIVEAIFAFFFPYRCCFNETEGDVQNNPHV |
| IRIS_313-8073 | MELITACALLLIAVHIAIVISCPEPLAAVAFVNVPLLYLCRYLLIIRRGVIIVSGGGGGGGRLRRFRLGAAMAVIYMAMSTLFFLRIAPLTPWWGALAAWVMILLIVEAIFAFFFPYRCCFNETEGDVQNNPHV |
| IRIS_313-8074 | MELITACALLLIAVHIAIVISCPEPLAAVALVDVPLLYLYRYLLIIRRGVIIVSGAGGGGGRLRRFRLGAAMAIIYMAMSTLFFLRIAPLTPWWGALAAWVMILLIVEAIFAFFFPYRCCFNETEGDVQNNPHV |
| IRIS_313-8075 | MELITACALLLIAVHIAIVISCPEPLAAVALVDVPLLYLCRYLLIIRRGVIIVSGAGAGGGRLRRFRLGAAMAIIYMAMSTLFFLRIAPLTPWWGALAAWVMILLIVEAIFAFFFPYRCCFNETEGDVQNNPHV |
| IRIS_313-8076 | MELITACALLLIAVHIAIVISCPEPLAAVALVDVPLLYLYRYLLIIRRGVIIVSGAGGGGGRLRRFRLGAAMAIIYMAMSTLFFLRIAPLTPWWGALAAWVMILLIVEAIFAFFFPYRCCFNETEGDVQNNPHV |
| IRIS_313-8084 | MELITACALLLIAVHIAIVISCPEPLAAVALVDVPLLYLCRYLLIIRRGVIIVSGAGAGGGRLRRFRLGAAMAIIYMAMSTLFFLRIAPLTPWWGALAAWVMILLIVEAIFAFFFPYRCCFNETEGDVQNNPHV |
| IRIS_313-8085 | MELITACALLLIAVHIAIVISCPEPLAAVALVDVPLLYLCRYLLIIRRGVIIVSGAGAGGGRLRRFRLGAAMAIIYMAMSTLFFLRIAPLTPWWGALAAWVMILLIVEAIFAFFFPYRCCFNETEGDVQNNPHV |
| IRIS_313-8087 | MELITACALLLIAVHIAIVISCPEPLAAVALVDVPLLYLCRYLLIIRRGVIIVSGAGAGGGRLRRFRLGAAMAIIYMAMSTLFFLRIAPLTPWWGALAAWVMILLIVEAIFAFFFPYRCCFNETEGDVQNNPHV |
| IRIS_313-8090 | MELITACALLLIAVHIAIVISCPEPLAAVALVDVPLLYLCRYLLIIRRGVIIVSGAGAGGGRLRRFRLGAAMAIIYMAMSTLFFLRIAPLTPWWGALAAWVMILLIVEAIFAFFFPYRCCFNETEGDVQNNPHV |
| IRIS_313-8095 | MELITACALLLIAVHIAIVISCPEPLAAVALVDVPLLYLCRYLLIIRRGVIIVSGAGAGGGRLRRFRLGAAMAIIYMAMSTLFFLRIAPLTPWWGALAAWVMILLIVEAIFAFFFPYRCCFNETEGDVQNNPHV |
| IRIS_313-8096 | MELITACALLLIAVHIAIVISCPEPLAAVALVDVPLLYLCRYLLIIRRGVIIVSGAGAGGGRLRRFRLGAAMAIIYMAMSTLFFLRIAPLTPWWGALAAWVMILLIVEAIFAFFFPYRCCFNETEGDVQNNPHV |
| IRIS_313-8097 | MELITACALLLIAVHIAIVISCPEPLAAVALVDVPLLYLCRYLLIIRRGVIIVSGAGAGGGRLRRFRLGAAMAIIYMAMSTLFFLRIAPLTPWWGALAAWVMILLIVEAIFAFFFPYRCCFNETEGDVQNNPHV |
| IRIS_313-8099 | MELITACALLLIAVHIAIVISCPEPLAAVALVDVPLLYLCRYLLIIRRGVIIVSGAGAGGGRLRRFRLGAAMAIIYMAMSTLFFLRIAPLTPWWGALAAWVMILLIVEAIFAFFFPYRCCFNETEGDVQNNPHV |
| IRIS_313-8102 | MELITACALLLIAVHIAIVISCPEPLAAVALVDVPLLYLYRYLLIIRRGVIIVSGAGGGGGRLRRFRLGAAMAIIYMAMSTLFFLRIAPLTPWWGALAAWVMILLIVEAIFAFFFPYRCCFNETEGDVQNNPHV |
| IRIS_313-8105 | MELITACALLLIAVHIAIVISCPEPLAAVALVDVPLLYLCRYLLIIRRGVIIVSGAGAGGGRLRRFRLGAAMAIIYMAMSTLFFLRIAPLTPWWGALAAWVMILLIVEAIFAFFFPYRCCFNETEGDVQNNPHV |
| IRIS_313-8109 | MELITACALLLIAVHIAIVISCPEPLAAVALVDVPLLYLYRYLLIIRRGVIIVSGAGGGGGRLRRFRLGAAMAIIYMAMSTLFFLRIAPLTPWWGALAAWVMILLIVEAIFAFFFPYRCCFNETEGDVQNNPHV |
| IRIS_313-8111 | MELITACALLLIAVHIAIVISCPEPLAAVALVDVPLLYLCRYLLIIRRGVIIVSGAGAGGGRLRRFRLGAAMAIIYMAMSTLFFLRIAPLTPWWGALAAWVMILLIVEAIFAFFFPYRCCFNETEGDVQNNPHV |
| IRIS_313-8112 | MELITACALLLIAVHIAIVISCPEPLAAVALVDVPLLYLCRYLLIIRRGVIIVSGAGAGGGRLRRFRLGAAMAIIYMAMSTLFFLRIAPLTPWWGALAAWVMILLIVEAIFAFFFPYRCCFNETEGDVQNNPHV |
| IRIS_313-8113 | MELITACALLLIAVHIAIVISCPEPLAAVALVDVPLLYLCRYLLIIRRGVIIVSGAGAGGGRLRRFRLGAAMAIIYMAMSTLFFLRIAPLTPWWGALAAWVMILLIVEAIFAFFFPYRCCFNETEGDVQNNPHV |
| IRIS_313-8114 | MELITACALLLIAVHIAIVISCPEPLAAVALVDVPLLYLCRYLLIIRRGVIIVSGAGAGGGRLRRFRLGAAMAIIYMAMSTLFFLRIAPLTPWWGALAAWVMILLIVEAIFAFFFPYRCCFNETEGDVQNNPHV |
| IRIS_313-8115 | MELITACALLLIAVHIAIVISCPEPLAAVALVDVPLLYLCRYLLIIRRGVIIVSGAGAGGGRLRRFRLGAAMAIIYMAMSTLFFLRIAPLTPWWGALAAWVMILLIVEAIFAFFFPYRCCFNETEGDVQNNPHV |
| IRIS_313-8116 | MELITACALLLIAVHIAIVISCPEPLAAVALVDVPLLYLYRYLLIIRRGVIIVSGAGGGGGRLRRFRLGAAMAIIYMAMSTLFFLRIAPLTPWWGALAAWVMILLIVEAIFAFFFPYRCCFNETEGDVQNNPHV |
| IRIS_313-8118 | MELITACALLLIAVHIAIVISCPEPLAAVALVDVPLLYLCRYLLIIRRGVIIVSGAGAGGGRLRRFRLGAAMAIIYMAMSTLFFLRIAPLTPWWGALAAWVMILLIVEAIFAFFFPYRCCFNETEGDVQNNPHV |
| IRIS_313-8119 | MELITACALLLIAVHIAIVISCPEPLAAVALVDVPLLYLCRYLLIIRRGVIIVSGAGAGGGRLRRFRLGAAMAIIYMAMSTLFFLRIAPLTPWWGALAAWVMILLIVEAIFAFFFPYRCCFNETEGDVQNNPHV |
| IRIS_313-8121 | MELITACALLLIAVHIAIVISCPEPLAAVALVDVPLLYLCRYLLIIRRGVIIVSGAGAGGGRLRRFRLGAAMAIIYMAMSTLFFLRIAPLTPWWGALAAWVMILLIVEAIFAFFFPYRCCFNETEGDVQNNPHV |
| IRIS_313-8123 | MELITACALLLIAVHIAIVISCPEPLAAVALVDVPLLYLCRYLLIIRRGVIIVSGAGAGGGRLRRFRLGAAMAIIYMAMSTLFFLRIAPLTPWWGALAAWVMILLIVEAIFAFFFPYRCCFNETEGDVQNNPHV |
| IRIS_313-8124 | MELITACALLLIAVHIAIVISCPEPLAAVALVDVPLLYLYRYLLIIRRGVIIVSGAGGGGGRLRRFRLGAAMAIIYMAMSTLFFLRIAPLTPWWGALAAWVMILLIVEAIFAFFFPYRCCFNETEGDVQNNPHV |
| IRIS_313-8125 | MELITACALLLIAVHIAIVISCPEPLAAVALVDVPLLYLCRYLLIIRRGVIIVSGAGAGGGRLRRFRLGAAMAIIYMAMSTLFFLRIAPLTPWWGALAAWVMILLIVEAIFAFFFPYRCCFNETEGDVQNNPHV |
| IRIS_313-8126 | MELITACALLLIAVHIAIVISCPEPLAAVALVDVPLLYLCRYLLIIRRGVIIVSGAGAGGGRLRRFRLGAAMAIIYMAMSTLFFLRIAPLTPWWGALAAWVMILLIVEAIFAFFFPYRCCFNETEGDVQNNPHV |
| IRIS_313-8127 | MELITACALLLIAVHIAIVISCPEPLAAVALVDVPLLYLCRYLLIIRRGVIIVSGAGAGGGRLRRFRLGAAMAIIYMAMSTLFFLRIAPLTPWWGALAAWVMILLIVEAIFAFFFPYRCCFNETEGDVQNNPHV |
| IRIS_313-8128 | MELITACALLLIAVHIAIVISCPEPLAAVALVDVPLLYLCRYLLIIRRGVIIVSGAGAGGGRLRRFRLGAAMAIIYMAMSTLFFLRIAPLTPWWGALAAWVMILLIVEAIFAFFFPYRCCFNETEGDVQNNPHV |
| IRIS_313-8129 | MELITACALLLIAVHIAIVISCPEPLAAVALVDVPLLYLCRYLLIIRRGVIIVSGAGAGGGRLRRFRLGAAMAIIYMAMSTLFFLRIAPLTPWWGALAAWVMILLIVEAIFAFFFPYRCCFNETEGDVQNNPHV |
| IRIS_313-8132 | MELITACALLLIAVHIAIVISCPEPLAAVALVDVPLLYLCRYLLIIRRGVIIVSGAGAGGGRLRRFRLGAAMAIIYMAMSTLFFLRIAPLTPWWGALAAWVMILLIVEAIFAFFFPYRCCFNETEGDVQNNPHV |
| IRIS_313-8134 | MELITACALLLIAVHIAIVISCPEPLAAVALVDVPLLYLCRYLLIIRRGVIIVSGAGAGGGRLRRFRLGAAMAIIYMAMSTLFFLRIAPLTPWWGALAAWVMILLIVEAIFAFFFPYRCCFNETEGDVQNNPHV |
| IRIS_313-8135 | MELITACALLLIAVHIAIVISCPEPLAAVALVDVPLLYLCRYLLIIRRGVIIVSGAGAGGGRLRRFRLGAAMAIIYMAMSTLFFLRIAPLTPWWGALAAWVMILLIVEAIFAFFFPYRCCFNETEGDVQNNPHV |
| IRIS_313-8136 | MELITACALLLIAVHIAIVISCPEPLAAVALVDVPLLYLYRYLLIIRRGVIIVSGAGGGGGRLRRFRLGAAMAIIYMAMSTLFFLRIAPLTPWWGALAAWVMILLIVEAIFAFFFPYRCCFNETEGDVQNNPHV |
| IRIS_313-8137 | MELITACALLLIAVHIAIVISCPEPLAAVALVDVPLLYLYRYLLIIRRGVIIVSGAGGGGGRLRRFRLGAAMAIIYMAMSTLFFLRIAPLTPWWGALAAWVMILLIVEAIFAFFFPYRCCFNETEGDVQNNPHV |
| IRIS_313-8138 | MELITACALLLIAVHIAIVISCPEPLAAVALVDVPLLYLCRYLLIIRRGVIIVSGAGAGGGRLRRFRLGAAMAIIYMAMSTLFFLRIAPLTPWWGALAAWVMILLIVEAIFAFFFPYRCCFNETEGDVQNNPHV |
| IRIS_313-8139 | MELITACALLLIAVHIAIVISCPEPLAAVALVDVPLLYLYRYLLIIRRGVIIVSGAGGGGGRLRRFRLGAAMAIIYMAMSTLFFLRIAPLTPWWGALAAWVMILLIVEAIFAFFFPYRCCFNETEGDVQNNPHV |
| IRIS_313-8140 | MELITACALLLIAVHIAIVISCPEPLAAVALVDVPLLYLYRYLLIIRRGVIIVSGAGGGGGRLRRFRLGAAMAIIYMAMSTLFFLRIAPLTPWWGALAAWVMILLIVEAIFAFFFPYRCCFNETEGDVQNNPHV |
| IRIS_313-8141 | MELITACALLLIAVHIAIVISCPEPLAAVALVDVPLLYLCRYLLIIRRGVIIVSGAGAGGGRLRRFRLGAAMAIIYMAMSTLFFLRIAPLTPWWGALAAWVMILLIVEAIFAFFFPYRCCFNETEGDVQNNPHV |
| IRIS_313-8142 | MELITACALLLIAVHIAIVISCPEPLAAVALVDVPLLYLYRYLLIIRRGVIIVSGAGGGGGRLRRFRLGAAMAIIYMAMSTLFFLRIAPLTPWWGALAAWVMILLIVEAIFAFFFPYRCCFNETEGDVQNNPHV |
| IRIS_313-8143 | MELITACALLLIAVHIAIVISCPEPLAAVALVDVPLLYLCRYLLIIRRGVIIVSGAGAGGGRLRRFRLGAAMAIIYMAMSTLFFLRIAPLTPWWGALAAWVMILLIVEAIFAFFFPYRCCFNETEGDVQNNPHV |
| IRIS_313-8145 | MELITACALLLIAVHIAIVISCPEPLAAVALVDVPLLYLYRYLLIIRRGVIIVSGAGGGGGRLRRFRLGAAMAIIYMAMSTLFFLRIAPLTPWWGALAAWVMILLIVEAIFAFFFPYRCCFNETEGDVQNNPHV |
| IRIS_313-8147 | MELITACALLLIAVHIAIVISCPEPLAAVALVDVPLLYLCRYLLIIRRGVIIVSGAGAGGGRLRRFRLGAAMAIIYMAMSTLFFLRIAPLTPWWGALAAWVMILLIVEAIFAFFFPYRCCFNETEGDVQNNPHV |
| IRIS_313-8148 | MELITACALLLIAVHIAIVISCPEPLAAVALVDVPLLYLCRYLLIIRRGVIIVSGAGAGGGRLRRFRLGAAMAIIYMAMSTLFFLRIAPLTPWWGALAAWVMILLIVEAIFAFFFPYRCCFNETEGDVQNNPHV |
| IRIS_313-8149 | MELITACALLLIAVHIAIVISCPEPLAAVALVDVPLLYLCRYLLIIRRGVIIVSGAGAGGGRLRRFRLGAAMAIIYMAMSTLFFLRIAPLTPWWGALAAWVMILLIVEAIFAFFFPYRCCFNETEGDVQNNPHV |
| IRIS_313-8151 | MELITACALLLIAVHIAIVISCPEPLAAVALVDVPLLYLCRYLLIIRRGVIIVSGAGAGGGRLRRFRLGAAMAIIYMAMSTLFFLRIAPLTPWWGALAAWVMILLIVEAIFAFFFPYRCCFNETEGDVQNNPHV |
| IRIS_313-8154 | MELITACALLLIAVHIAIVISCPEPLAAVALVDVPLLYLCRYLLIIRRGVIIVSGAGAGGGRLRRFRLGAAMAIIYMAMSTLFFLRIAPLTPWWGALAAWVMILLIVEAIFAFFFPYRCCFNETEGDVQNNPHV |
| IRIS_313-8155 | MELITACALLLIAVHIAIVISCPEPLAAVALVDVPLLYLYRYLLIIRRGVIIVSGAGGGGGRLRRFRLGAAMAIIYMAMSTLFFLRIAPLTPWWGALAAWVMILLIVEAIFAFFFPYRCCFNETEGDVQNNPHV |
| IRIS_313-8158 | MELITACALLLIAVHIAIVISCPEPLAAVALVDVPLLYLCRYLLIIRRGVIIVSGAGAGGGRLRRFRLGAAMAIIYMAMSTLFFLRIAPLTPWWGALAAWVMILLIVEAIFAFFFPYRCCFNETEGDVQNNPHV |
| IRIS_313-8159 | MELITACALLLIAVHIAIVISCPEPLAAVALVDVPLLYLCRYLLIIRRGVIIVSGAGAGGGRLRRFRLGAAMAIIYMAMSTLFFLRIAPLTPWWGALAAWVMILLIVEAIFAFFFPYRCCFNETEGDVQNNPHV |
| IRIS_313-8160 | MELITACALLLIAVHIAIVISCPEPLAAVALVDVPLLYLCRYLLIIRRGVIIVSGAGAGGGRLRRFRLGAAMAIIYMAMSTLFFLRIAPLTPWWGALAAWVMILLIVEAIFAFFFPYRCCFNETEGDVQNNPHV |
| IRIS_313-8161 | MELITACALLLIAVHIAIVISCPEPLAAVALVDVPLLYLCRYLLIIRRGVIIVSGAGAGGGRLRRFRLGAAMAIIYMAMSTLFFLRIAPLTPWWGALAAWVMILLIVEAIFAFFFPYRCCFNETEGDVQNNPHV |
| IRIS_313-8162 | MELITACALLLIAVHIAIVISCPEPLAAVALVDVPLLYLCRYLLIIRRGVIIVSGAGAGGGRLRRFRLGAAMAIIYMAMSTLFFLRIAPLTPWWGALAAWVMILLIVEAIFAFFFPYRCCFNETEGDVQNNPHV |
| IRIS_313-8164 | MELITACALLLIAVHIAIVISCPEPLAAVALVDVPLLYLCRYLLIIRRGVIIVSGAGAGGGRLRRFRLGAAMAIIYMAMSTLFFLRIAPLTPWWGALAAWVMILLIVEAIFAFFFPYRCCFNETEGDVQNNPHV |
| IRIS_313-8165 | MELITACALLLIAVHIAIVISCPEPLAAVALVDVPLLYLCRYLLIIRRGVIIVSGAGAGGGRLRRFRLGAAMAIIYMAMSTLFFLRIAPLTPWWGALAAWVMILLIVEAIFAFFFPYRCCFNETEGDVQNNPHV |
| IRIS_313-8166 | MELITACALLLIAVHIAIVISCPEPLAAVALVDVPLLYLYRYLLIIRRGVIIVSGAGGGGGRLRRFRLGAAMAIIYMAMSTLFFLRIAPLTPWWGALAAWVMILLIVEAIFAFFFPYRCCFNETEGDVQNNPHV |
| IRIS_313-8167 | MELITACALLLIAVHIAIVISCPEPLAAVALVDVPLLYLYRYLLIIRRGVIIVSGAGGGGGRLRRFRLGAAMAIIYMAMSTLFFLRIAPLTPWWGALAAWVMILLIVEAIFAFFFPYRCCFNETEGDVQNNPHV |
| IRIS_313-8168 | MELITACALLLIAVHIAIVISCPEPLAAVALVDVPLLYLCRYLLIIRRGVIIVSGAGAGGGRLRRFRLGAAMAIIYMAMSTLFFLRIAPLTPWWGALAAWVMILLIVEAIFAFFFPYRCCFNETEGDVQNNPHV |
| IRIS_313-8170 | MELITACALLLIAVHIAIVISCPEPLAAVALVDVPLLYLYRYLLIIRRGVIIVSGAGGGGGRLRRFRLGAAMAIIYMAMSTLFFLRIAPLTPWWGALAAWVMILLIVEAIFAFFFPYRCCFNETEGDVQNNPHV |
| IRIS_313-8171 | MELITACALLLIAVHIAIVISCPEPLAAVALVDVPLLYLCRYLLIIRRGVIIVSGAGAGGGRLRRFRLGAAMAIIYMAMSTLFFLRIAPLTPWWGALAAWVMILLIVEAIFAFFFPYRCCFNETEGDVQNNPHV |
| IRIS_313-8172 | MELITACALLLIAVHIAIVISCPEPLAAVALVDVPLLYLCRYLLIIRRGVIIVSGAGAGGGRLRRFRLGAAMAIIYMAMSTLFFLRIAPLTPWWGALAAWVMILLIVEAIFAFFFPYRCCFNETEGDVQNNPHV |

IRIS_313-8173  MELITACALLLIAVHIAIVISCPEPLAAVALVDVPLLYLCRYLLIIRRGVIIVSGAGAGGGRLRRFRLGAAMAIIYMAMSTLFFLRIAPLTPWWGALAAWVMILLIVEAIFAFFFPYRCCFNETEGDVQNNPHV
IRIS_313-8177  MELITACALLLIAVHIAIVISCPEPLAAVALVDVPLLYLCRYLLIIRRGVIIVSGAGAGGGRLRRFRLGAAMAIIYMAMSTLFFLRIAPLTPWWGALAAWVMILLIVEAIFAFFFPYRCCFNETEGDVQNNPHV
IRIS_313-8180  MELITACALLLIAVHIAIVISCPEPLAAVALVDVPLLYLYRYLLIIRRGVIIVSGAGGGGGRLRRFRLGAAMAIIYMAMSTLFFLRIAPLTPWWGALAAWVMILLIVEAIFAFFFPYRCCFNETEGDVQNNPHV
IRIS_313-8183  MELITACALLLIAVHIAIVISCPEPLAAVALVDVPLLYLYRYLLIIRRGVIIVSGAGGGGGRLRRFRLGAAMAIIYMAMSTLFFLRIAPLTPWWGALAAWVMILLIVEAIFAFFFPYRCCFNETEGDVQNNPHV
IRIS_313-8184  MELITACALLLIAVHIAIVISCPEPLAAVALVDVPLLYLCRYLLIIRRGVIIVSGAGAGGGRLRRFRLGAAMAIIYMAMSTLFFLRIAPLTPWWGALAAWVMILLIVEAIFAFFFPYRCCFNETEGDVQNNPHV
IRIS_313-8185  MELITACALLLIAVHIAIVISCPEPLAAVALVDVPLLYLCRYLLIIRRGVIIVSGAGAGGGRLRRFRLGAAMAIIYMAMSTLFFLRIAPLTPWWGALAAWVMILLIVEAIFAFFFPYRCCFNETEGDVQNNPHV
IRIS_313-8186  MELITACALLLIAVHIAIVISCPEPLAAVALVDVPLLYLYRYLLIIRRGVIIVSGAGGGGGRLRRFRLGAAMAIIYMAMSTLFFLRIAPLTPWWGALAAWVMILLIVEAIFAFFFPYRCCFNETEGDVQNNPHV
IRIS_313-8192  MELITACALLLIAVHIAIVISCPEPLAAVALVDVPLLYLCRYLLIIRRGVIIVSGAGAGGGRLRRFRLGAAMAIIYMAMSTLFFLRIAPLTPWWGALAAWVMILLIVEAIFAFFFPYRCCFNETEGDVQNNPHV
IRIS_313-8193  MELITACALLLIAVHIAIVISCPEPLAAVALVDVPLLYLYRYLLIIRRGVIIVSGAGGGGGRLRRFRLGAAMAIIYMAMSTLFFLRIAPLTPWWGALAAWVMILLIVEAIFAFFFPYRCCFNETEGDVQNNPHV
IRIS_313-8195  MELITACALLLIAVHIAIVISCPEPLAAVALVDVPLLYLCRYLLIIRRGVIIVSGAGAGGGRLRRFRLGAAMAIIYMAMSTLFFLRIAPLTPWWGALAAWVMILLIVEAIFAFFFPYRCCFNETEGDVQNNPHV
IRIS_313-8200  MELITACALLLIAVHIAIVISCPEPLAAVALVDVPLLYLCRYLLIIRRGVIIVSGAGAGGGRLRRFRLGAAMAIIYMAMSTLFFLRIAPLTPWWGALAAWVMILLIVEAIFAFFFPYRCCFNETEGDVQNNPHV
IRIS_313-8202  MELITACALLLIAVHIAIVISCPEPLAAVALVDVPLLYLCRYLLIIRRGVIIVSGAGAGGGRLRRFRLGAAMAIIYMAMSTLFFLRIAPLTPWWGALAAWVMILLIVEAIFAFFFPYRCCFNETEGDVQNNPHV
IRIS_313-8204  MELITACALLLIAVHIAIVISCPEPLAAVALVDVPLLYLCRYLLIIRRGVIIVSGAGAGGGRLRRFRLGAAMAIIYMAMSTLFFLRIAPLTPWWGALAAWVMILLIVEAIFAFFFPYRCCFNETEGDVQNNPHV
IRIS_313-8205  MELITACALLLIAVHIAIVISCPEPLAAVALVDVPLLYLCRYLLIIRRGVIIVSGAGAGGGRLRRFRLGAAMAIIYMAMSTLFFLRIAPLTPWWGALAAWVMILLIVEAIFAFFFPYRCCFNETEGDVQNNPHV
IRIS_313-8208  MELITACALLLIAVHIAIVISCPEPLAAVALVDVPLLYLCRYLLIIRRGVIIVSGAGAGGGRLRRFRLGAAMAIIYMAMSTLFFLRIAPLTPWWGALAAWVMILLIVEAIFAFFFPYRCCFNETEGDVQNNPHV
IRIS_313-8209  MELITACALLLIAVHIAIVISCPEPLAAVALVDVPLLYLCRYLLIIRRGVIIVSGAGAGGGRLRRFRLGAAMAIIYMAMSTLFFLRIAPLTPWWGALAAWVMILLIVEAIFAFFFPYRCCFNETEGDVQNNPHV
IRIS_313-8212  MELITACALLLIAVHIAIVISCPEPLAAVALVDVPLLYLCRYLLIIRRGVIIVSGGGGGGRLRRFRLGAAMAIIYMAMSTLFFLRIAPLTPWWGALAAWVMILLIVEAIFAFFFPYRCCFNETEGDVQNNPHV
IRIS_313-8213  MELITACALLLIAVHIAIVISCPEPLAAVALVDVPLLYLYRYLLIIRRGVIIVSGAGGGGGRLRRFRLGAAMAIIYMAMSTLFFLRIAPLTPWWGALAAWVMILLIVEAIFAFFFPYRCCFNETEGDVQNNPHV
IRIS_313-8214  MELITACALLLIAVHIAIVISCPEPLAAVALVDVPLLYLYRYLLIIRRGVIIVSGAGGGGGRLRRFRLGAAMAIIYMAMSTLFFLRIAPLTPWWGALAAWVMILLIVEAIFAFFFPYRCCFNETEGDVQNNPHV
IRIS_313-8215  MELITACALLLIAVHIAIVISCPEPLAAVALVDVPLLYLCRYLLIIRRGVIIVSGGGGGGRLRRFRLGAAMAIIYMAMSTLFFLRIAPLTPWWGALAAWVMILLIVEAIFAFFFPYRCCFNETEGDVQNNPHV
IRIS_313-8216  MELITACALLLIAVHIAIVISCPEPLAAVALVDVPLLYLCRYLLIIRRGVIIVSGAGAGGGRLRRFRLGAAMAIIYMAMSTLFFLRIAPLTPWWGALAAWVMILLIVEAIFAFFFPYRCCFNETEGDVQNNPHV
IRIS_313-8217  MELITACALLLIAVHIAIVISCPEPLAAVALVDVPLLYLCRYLLIIRRGVIIVSGAGAGGGRLRRFRLGAAMAIIYMAMSTLFFLRIAPLTPWWGALAAWVMILLIVEAIFAFFFPYRCCFNETEGDVQNNPHV
IRIS_313-8218  MELITACALLLIAVHIAIVISCPEPLAAVALVDVPLLYLCRYLLIIRRGVIIVSGAGAGGGRLRRFRLGAAMAIIYMAMSTLFFLRIAPLTPWWGALAAWVMILLIVEAIFAFFFPYRCCFNETEGDVQNNPHV
IRIS_313-8232  MELITACALLLIAVHIAIVISCPEPLAAVALVDVPLLYLCRYLLIIRRGVIIVSGAGAGGGRLRRFRLGAAMAIIYMAMSTLFFLRIAPLTPWWGALAAWVMILLIVEAIFAFFFPYRCCFNETEGDVQNNPHV
IRIS_313-8244  MELITACALLLIAVHIAIVISCPEPLAAVALVDVPLLYLCRYLLIIRRGVIIVSGGGGGGRLRRFRLGAAMAIIYMAMSTLFFLRIAPLTPWWGALAAWVMILLIVEAIFAFFFPYRCCFNETEGDVQNNPHV
IRIS_313-8252  MELITACALLLIAVHIAIVISCPEPLAAVAFVNVPLLYLCRYLLIIRRGVIIVSGGGGGGGRLRRFRLGAAMAVIYMAMSTLFFLRIAPLTPWWGALAAWVMILLIVEAIFAFFFPYRCCFNETEGDVQNNPHV
IRIS_313-8253  MELITACALLLIAVHIAIVISCPEPLAAVALVDVPLLYLCRYLLIIRRGVIIVSGGGGGGRLRRFRLGAAMAIIYMAMSTLFFLRIAPLTPWWGALAAWVMILLIVEAIFAFFFPYRCCFNETEGDVQNNPHV
IRIS_313-8256  MELITACALLLIAVHIAIVISCPEPLAAVALVDVPLLYLCRYLLIIRRGVIIVSGAGAGGGRLRRFRLGAAMAIIYMAMSTLFFLRIAPLTPWWGALAAWVMILLIVEAIFAFFFPYRCCFNETEGDVQNNPHV
IRIS_313-8265  MELITACALLLIAVHIAIVISCPEPLAAVALVDVPLLYLCRYLLIIRRGVIIVSGGGGGGRLRRFRLGAAMAIIYMAMSTLFFLRIAPLTPWWGALAAWVMILLIVEAIFAFFFPYRCCFNETEGDVQNNPHV
IRIS_313-8268  MELITACALLLIAVHIAIVISCPEPLAAVALVDVPLLYLCRYLLIIRRGVIIVSGAGAGGGRLRRFRLGAAMAIIYMAMSTLFFLRIAPLTPWWGALAAWVMILLIVEAIFAFFFPYRCCFNETEGDVQNNPHV
IRIS_313-8277  MELITACALLLIAVHIAIVISCPEPLAAVALVDVPLLYLCRYLLIIRRGVIIVSGAGAGGGRLRRFRLGAAMAIIYMAMSTLFFLRIAPLTPWWGALAAWVMILLIVEAIFAFFFPYRCCFNETEGDVQNNPHV
IRIS_313-8279  MELITACALLLIAVHIAIVISCPEPLAAVALVDVPLLYLYRYLLIIRRGVIIVSGAGGGGGRLRRFRLGAAMAIIYMAMSTLFFLRIAPLTPWWGALAAWVMILLIVEAIFAFFFPYRCCFNETEGDVQNNPHV
IRIS_313-8283  MELITACALLLIAVHIAIVISCPEPLAAVALVDVPLLYLCRYLLIIRRGVIIVSGAGAGGGRLRRFRLGAAMAIIYMAMSTLFFLRIAPLTPWWGALAAWVMILLIVEAIFAFFFPYRCCFNETEGDVQNNPHV
IRIS_313-8285  MELITACALLLIAVHIAIVISCPEPLAAVALVDVPLLYLYRYLLIIRRGVIIVSGAGGGGGRLRRFRLGAAMAIIYMAMSTLFFLRIAPLTPWWGALAAWVMILLIVEAIFAFFFPYRCCFNETEGDVQNNPHV
IRIS_313-8288  MELITACALLLIAVHIAIVISCPEPLAAVALVDVPLLYLCRYLLIIRRGVIIVSGGGAGGGRLRRFRLGAAMAIIYMAMSTLFFLRIAPLTPWWGALAAWVMILLIVEAIFAFFFPYRCCFNETEGDVQNNPHV
IRIS_313-8291  MELITACALLLIAVHIAIVISCPEPLAAVALVDVPLLYLCRYLLIIRRGVIIVSGGGGGGRLRRFRLGAAMAIIYMAMSTLFFLRIAPLTPWWGALAAWVMILLIVEAIFAFFFPYRCCFNETEGDVQNNPHV
IRIS_313-8292  MELITACALLLIAVHIAIVISCPEPLAAVALVDVPLLYLCRYLLIIRRGVIIVSGAGGGGGGRLRRFRLGAAMAIIYMAMSTLFFLRIAPLTPWWGALAAWVMILLIVEAIFAFFFPYRCCFNETEGDVQNNPHV
IRIS_313-8293  MELITACALLLIAVHIAIVISCPEPLAAVALVDVPLLYLCRYLLIIRRGVIIVSGAGGGGGGRLRRFRLGAAMAIIYMAMSTLFFLRIAPLTPWWGALAAWVMILLIVEAIFAFFFPYRCCFNETEGDVQNNPHV
IRIS_313-8302  MELITACALLLIAVHIAIVISCPEPLAAVALVDVPLLYLYRYLLIIRRGVIIVSGAGGGGGRLRRFRLGAAMAIIYMAMSTLFFLRIAPLTPWWGALAAWVMILLIVEAIFAFFFPYRCCFNETEGDVQNNPHV
IRIS_313-8303  MELITACALLLIAVHIAIVISCPEPLAAVALVDVPLLYLCRYLLIIRRGVIIVSGGGGGGRLRRFRLGAAMAIIYMAMSTLFFLRIAPLTPWWGALAAWVMILLIVEAIFAFFFPYRCCFNETEGDVQNNPHV
IRIS_313-8305  MELITACALLLIAVHIAIVISCPEPLAAVALVDVPLLYLCRYLLIIRRGVIIVSGGGGGGRLRRFRLGAAMAIIYMAMSTLFFLRIAPLTPWWGALAAWVMILLIVEAIFAFFFPYRCCFNETEGDVQNNPHV
IRIS_313-8306  MELITACALLLIAVHIAIVISCPEPLAAVALVDVPLLYLCRYLLIIRRGVIIVSGGGGGGRLRRFRLGAAMAIIYMAMSTLFFLRIAPLTPWWGALAAWVMILLIVEAIFAFFFPYRCCFNETEGDVQNNPHV
IRIS_313-8312  MELITACALLLIAVHIAIVISCPEPLAAVALVDVPLLYLCRYLLIIRRGVIIVSGAGAGGGRLRRFRLGAAMAIIYMAMSTLFFLRIAPLTPWWGALAAWVMILLIVEAIFAFFFPYRCCFNETEGDVQNNPHV
IRIS_313-8314  MELITACALLLIAVHIAIVISCPEPLAAVALVDVPLLYLCRYLLIIRRGVIIVSGAGAGGGRLRRFRLGAAMAIIYMAMSTLFFLRIAPLTPWWGALAAWVMILLIVEAIFAFFFPYRCCFNETEGDVQNNPHV
IRIS_313-8315  MELITACALLLIAVHIAIVISCPEPLAAVALVDVPLLYLCRYLLIIRRGVIIVSGAGAGGGRLRRFRLGAAMAIIYMAMSTLFFLRIAPLTPWWGALAAWVMILLIVEAIFAFFFPYRCCFNETEGDVQNNPHV
IRIS_313-8316  MELITACALLLIAVHIAIVISCPEPLAAVALVDVPLLYLCRYLLIIRRGVIIVSGGGGGGRLRRFRLGAAMAIIYMAMSTLFFLRIAPLTPWWGALAAWVMILLIVEAIFAFFFPYRCCFNETEGDVQNNPHV
IRIS_313-8321  MELITACALLLIAVHIAIVISCPEPLAAVALVDVPLLYLCRYLLIIRRGVIIVSGAGGGGGRLRRFRLGAAMAIIYMAMSTLFFLRIAPLTPWWGALAAWVMILLIVEAIFAFFFPYRCCFNETEGDVQNNPHV
IRIS_313-8323  MELITACALLLIAVHIAIVISCPEPLAAVALVDVPLLYLYRYLLIIRRGVIIVSGAGGGGGRLRRFRLGAAMAIIYMAMSTLFFLRIAPLTPWWGALAAWVMILLIVEAIFAFFFPYRCCFNETEGDVQNNPHV
IRIS_313-8324  MELITACALLLIAVHIAIVISCPEPLAAVALVDVPLLYLYRYLLIIRRGVIIVSGAGGGGGRLRRFRLGAAMAIIYMAMSTLFFLRIAPLTPWWGALAAWVMILLIVEAIFAFFFPYRCCFNETEGDVQNNPHV
IRIS_313-8326  MELITACALLLIAVHIAIVISCPEPLAAVALVDVPLLYLCRYLLIIRRGVIIVSGAGAGGGRLRRFRLGAAMAIIYMAMSTLFFLRIAPLTPWWGALAAWVMILLIVEAIFAFFFPYRCCFNETEGDVQNNPHV
IRIS_313-8332  MELITACALLLIAVHIAIVISCPEPLAAVALVDVPLLYLCRYLLIIRRGVIIVSGGGGGGRLRRFRLGAAMAIIYMAMSTLFFLRIAPLTPWWGALAAWVMILLIVEAIFAFFFPYRCCFNETEGDVQNNPHV
IRIS_313-8339  MELITACALLLIAVHIAIVISCPEPLAAVALVDVPLLYLYRYLLIIRRGVIIVSGAGGGGGRLRRFRLGAAMAIIYMAMSTLFFLRIAPLTPWWGALAAWVMILLIVEAIFAFFFPYRCCFNETEGDVQNNPHV
IRIS_313-8341  MELITACALLLIAVHIAIVISCPEPLAAVALVDVPLLYLYRYLLIIRRGVIIVSGAGGGGGRLRRFRLGAAMAIIYMAMSTLFFLRIAPLTPWWGALAAWVMILLIVEAIFAFFFPYRCCFNETEGDVQNNPHV
IRIS_313-8342  MELITACALLLIAVHIAIVISCPEPLAAVAFVNVPLLYLCRYLLIIRRGVIIVSGGGGGGGRLRRFRLGAAMAVIYMAMSTLFFLRIAPLTPWWGALAAWVMILLIVEAIFAFFFPYRCCFNETEGDVQNNPHV
IRIS_313-8349  MELITACALLLIAVHIAIVISCPEPLAAVALVDVPLLYLCRYLLIIRRGVIIVSGAGAGGGRLRRFRLGAAMAIIYMAMSTLFFLRIAPLTPWWGALAAWVMILLIVEAIFAFFFPYRCCFNETEGDVQNNPHV
IRIS_313-8356  MELITACALLLIAVHIAIVISCPEPLAAVALVDVPLLYLCRYLLIIRRGVIIVSGAGAGGGRLRRFRLGAAMAIIYMAMSTLFFLRIAPLTPWWGALAAWVMILLIVEAIFAFFFPYRCCFNETEGDVQNNPHV
IRIS_313-8368  MELITACALLLIAVHIAIVISCPEPLAAVALVDVPLLYLCRYLLIIRRGVIIVSGGGGGGRLRRFRLGAAMAIIYMAMSTLFFLRIAPLTPWWGALAAWVMILLIVEAIFAFFFPYRCCFNETEGDVQNNPHV
IRIS_313-8380  MELITACALLLIAVHIAIVISCPEPLAAVALVDVPLLYLCRYLLIIRRGVIIVSGGGAGGGRLRRFRLGAAMAIIYMAMSTLFFLRIAPLTPWWGALAAWVMILLIVEAIFAFFFPYRCCFNETEGDVQNNPHV
IRIS_313-8381  MELITACALLLIAVHIAIVISCPEPLAAVALVDVPLLYLYRYLLIIRRGVIIVSGAGGGGGRLRRFRLGAAMAIIYMAMSTLFFLRIAPLTPWWGALAAWVMILLIVEAIFAFFFPYRCCFNETEGDVQNNPHV
IRIS_313-8382  MELITACALLLIAVHIAIVISCPEPLAAVALVDVPLLYLCRYLLIIRRGVIIVSGGGGGGRLRRFRLGAAMAIIYMAMSTLFFLRIAPLTPWWGALAAWVMILLIVEAIFAFFFPYRCCFNETEGDVQNNPHV
IRIS_313-8383  MELITACALLLIAVHIAIVISCPEPLAAVALVDVPLLYLCRYLLIIRRGVIIVSGAGAGGGRLRRFRLGAAMAIIYMAMSTLFFLRIAPLTPWWGALAAWVMILLIVEAIFAFFFPYRCCFNETEGDVQNNPHV
IRIS_313-8385  MELITACALLLIAVHIAIVISCPEPLAAVALVDVPLLYLCRYLLIIRRGVIIVSGAGAGGGRLRRFRLGAAMAIIYMAMSTLFFLRIAPLTPWWGALAAWVMILLIVEAIFAFFFPYRCCFNETEGDVQNNPHV
IRIS_313-8386  MELITACALLLIAVHIAIVISCPEPLAAVALVDVPLLYLCRYLLIIRRGVIIVSGGGAGGGRLRRFRLGAAMAIIYMAMSTLFFLRIAPLTPWWGALAAWVMILLIVEAIFAFFFPYRCCFNETEGDVQNNPHV
IRIS_313-8387  MELITACALLLIAVHIAIVISCPEPLAAVALVDVPLLYLCRYLLIIRRGVIIVSGAGAGGGRLRRFRLGAAMAIIYMAMSTLFFLRIAPLTPWWGALAAWVMILLIVEAIFAFFFPYRCCFNETEGDVQNNPHV
IRIS_313-8390  MELITACALLLIAVHIAIVISCPEPLAAVAFVNVPLLYLCRYLLIIRRGVIIVSGGGGGGGRLRRFRLGAAMAVIYMAMSTLFFLRIAPLTPWWGALAAWVMILLIVEAIFAFFFPYRCCFNETEGDVQNNPHV
IRIS_313-8391  MELITACALLLIAVHIAIVISCPEPLAAVALVDVPLLYLCRYLLIIRRGVIIVSGAGAGGGRLRRFRLGAAMAIIYMAMSTLFFLRIAPLTPWWGALAAWVMILLIVEAIFAFFFPYRCCFNETEGDVQNNPHV
IRIS_313-8392  MELITACALLLIAVHIAIVISCPEPLAAVALVDVPLLYLCRYLLIIRRGVIIVSGGGGGGRLRRFRLGAAMAIIYMAMSTLFFLRIAPLTPWWGALAAWVMILLIVEAIFAFFFPYRCCFNETEGDVQNNPHV
IRIS_313-8398  MELITACALLLIAVHIAIVISCPEPLAAVALVDVPLLYLCRYLLIIRRGVIIVSGAGAGGGRLRRFRLGAAMAIIYMAMSTLFFLRIAPLTPWWGALAAWVMILLIVEAIFAFFFPYRCCFNETEGDVQNNPHV
IRIS_313-8399  MELITACALLLIAVHIAIVISCPEPLAAVALVDVPLLYLCRYLLIIRRGVIIVSGAGGGGGRLRRFRLGAAMAIIYMAMSTLFFLRIAPLTPWWGALAAWVMILLIVEAIFAFFFPYRCCFNETEGDVQNNPHV
IRIS_313-8400  MELITACALLLIAVHIAIVISCPEPLAAVALVDVPLLYLYRYLLIIRRGVIIVSGAGGGGGRLRRFRLGAAMAIIYMAMSTLFFLRIAPLTPWWGALAAWVMILLIVEAIFAFFFPYRCCFNETEGDVQNNPHV
IRIS_313-8401  MELITACALLLIAVHIAIVISCPEPLAAVALVDVPLLYLCRYLLIIRRGVIIVSGAGAGGGRLRRFRLGAAMAIIYMAMSTLFFLRIAPLTPWWGALAAWVMILLIVEAIFAFFFPYRCCFNETEGDVQNNPHV
IRIS_313-8405  MELITACALLLIAVHIAIVISCPEPLAAVALVDVPLLYLCRYLLIIRRGVIIVSGGGGGGRLRRFRLGAAMAIIYMAMSTLFFLRIAPLTPWWGALAAWVMILLIVEAIFAFFFPYRCCFNETEGDVQNNPHV
IRIS_313-8407  MELITACALLLIAVHIAIVISCPEPLAAVALVDVPLLYLCRYLLIIRRGVIIVSGAGAGGGRLRRFRLGAAMAIIYMAMSTLFFLRIAPLTPWWGALAAWVMILLIVEAIFAFFFPYRCCFNETEGDVQNNPHV
IRIS_313-8409  MELITACALLLIAVHIAIVISCPEPLAAVALVDVPLLYLYRYLLIIRRGVIIVSGAGGGGGRLRRFRLGAAMAIIYMAMSTLFFLRIAPLTPWWGALAAWVMILLIVEAIFAFFFPYRCCFNETEGDVQNNPHV
IRIS_313-8410  MELITACALLLIAVHIAIVISCPEPLAAVAFVNVPLLYLCRYLLIIRRGVIIVSGGGGGGGRLRRFRLGAAMAVIYMAMSTLFFLRIAPLTPWWGALAAWVMILLIVEAIFAFFFPYRCCFNETEGDVQNNPHV
IRIS_313-8412  MELITACALLLIAVHIAIVISCPEPLAAVALVDVPLLYLCRYLLIIRRGVIIVSGGGGGGRLRRFRLGAAMAIIYMAMSTLFFLRIAPLTPWWGALAAWVMILLIVEAIFAFFFPYRCCFNETEGDVQNNPHV
IRIS_313-8414  MELITACALLLIAVHIAIVISCPEPLAAVALVDVPLLYLYRYLLIIRRGVIIVSGAGGGGGRLRRFRLGAAMAIIYMAMSTLFFLRIAPLTPWWGALAAWVMILLIVEAIFAFFFPYRCCFNETEGDVQNNPHV
IRIS_313-8431  MELITACALLLIAVHIAIVISCPEPLAAVALVDVPLLYLCRYLLIIRRGVIIVSGAGAGGGRLRRFRLGAAMAIIYMAMSTLFFLRIAPLTPWWGALAAWVMILLIVEAIFAFFFPYRCCFNETEGDVQNNPHV
IRIS_313-8433  MELITACALLLIAVHIAIVISCPEPLAAVALVDVPLLYLCRYLLIIRRGVIIVSGAGAGGGRLRRFRLGAAMAIIYMAMSTLFFLRIAPLTPWWGALAAWVMILLIVEAIFAFFFPYRCCFNETEGDVQNNPHV
IRIS_313-8434  MELITACALLLIAVHIAIVISCPEPLAAVALVDVPLLYLYRYLLIIRRGVIIVSGAGGGGGRLRRFRLGAAMAIIYMAMSTLFFLRIAPLTPWWGALAAWVMILLIVEAIFAFFFPYRCCFNETEGDVQNNPHV
IRIS_313-8435  MELITACALLLIAVHIAIVISCPEPLAAVALVDVPLLYLCRYLLIIRRGVIIVSGGGGGGRLRRFRLGAAMAIIYMAMSTLFFLRIAPLTPWWGALAAWVMILLIVEAIFAFFFPYRCCFNETEGDVQNNPHV
IRIS_313-8436  MELITACALLLIAVHIAIVISCPEPLAAVALVDVPLLYLCRYLLIIRRGVIIVSGAGGGGGRLRRFRLGAAMAIIYMAMSTLFFLRIAPLTPWWGALAAWVMILLIVEAIFAFFFPYRCCFNETEGDVQNNPHV
IRIS_313-8437  MELITACALLLIAVHIAIVISCPEPLAAVALVDVPLLYLCRYLLIIRRGVIIVSGAGAGGGRLRRFRLGAAMAIIYMAMSTLFFLRIAPLTPWWGALAAWVMILLIVEAIFAFFFPYRCCFNETEGDVQNNPHV
IRIS_313-8439  MELITACALLLIAVHIAIVISCPEPLAAVALVDVPLLYLCRYLLIIRRGVIIVSGAGGGGGGRLRRFRLGAAMAIIYMAMSTLFFLRIAPLTPWWGALAAWVMILLIVEAIFAFFFPYRCCFNETEGDVQNNPHV
IRIS_313-8450  MELITACALLLIAVHIAIVISCPEPLAAVALVDVPLLYLCRYLLIIRRGVIIVSGAGAGGGRLRRFRLGAAMAIIYMAMSTLFFLRIAPLTPWWGALAAWVMILLIVEAIFAFFFPYRCCFNETEGDVQNNPHV
IRIS_313-8453  MELITACALLLIAVHIAIVISCPEPLAAVALVDVPLLYLCRYLLIIRRGVIIVSGGGGGGRLRRFRLGAAMAIIYMAMSTLFFLRIAPLTPWWGALAAWVMILLIVEAIFAFFFPYRCCFNETEGDVQNNPHV
IRIS_313-8454  MELITACALLLIAVHIAIVISCPEPLAAVALVDVPLLYLCRYLLIIRRGVIIVSGGGGGGRLRRFRLGAAMAIIYMAMSTLFFLRIAPLTPWWGALAAWVMILLIVEAIFAFFFPYRCCFNETEGDVQNNPHV
IRIS_313-8457  MELITACALLLIAVHIAIVISCPEPLAAVALVDVPLLYLCRYLLIIRRGVIIVSGGGAGGGRLRRFRLGAAMAIIYMAMSTLFFLRIAPLTPWWGALAAWVMILLIVEAIFAFFFPYRCCFNETEGDVQNNPHV
IRIS_313-8458  MELITACALLLIAVHIAIVISCPEPLAAVALVDVPLLYLCRYLLIIRRGVIIVSGGGAGGGRLRRFRLGAAMAIIYMAMSTLFFLRIAPLTPWWGALAAWVMILLIVEAIFAFFFPYRCCFNETEGDVQNNPHV
IRIS_313-8466  MELITACALLLIAVHIAIVISCPEPLAAVALVDVPLLYLCRYLLIIRRGVIIVSGGGGGGRLRRFRLGAAMAIIYMAMSTLFFLRIAPLTPWWGALAAWVMILLIVEAIFAFFFPYRCCFNETEGDVQNNPHV
IRIS_313-8468  MELITACALLLIAVHIAIVISCPEPLAAVALVDVPLLYLCRYLLIIRRGVIIVSGAGGAAAAGCAASASGQPWRSSTWPCRPSSSSALLLSRHGGAPSPPGS*SCSSSRPSSPSSSPTAAASTKPAKAMSRIIPM
IRIS_313-8474  MELITACALLLIAVHIAIVISCPEPLAAVALVDVPLLYLYRYLLIIRRGVIIVSGAGGGGGRLRRFRLGAAMAIIYMAMSTLFFLRIAPLTPWWGALAAWVMILLIVEAIFAFFFPYRCCFNETEGDVQNNPHV
IRIS_313-8481  MELITACALLLIAVHIAIVISCPEPLAAVALVDVPLLYLCRYLLIIRRGVIIVSGAGAGGGRLRRFRLGAAMAIIYMAMSTLFFLRIAPLTPWWGALAAWVMILLIVEAIFAFFFPYRCCFNETEGDVQNNPHV
IRIS_313-8485  MELITACALLLIAVHIAIVISCPEPLAAVALVDVPLLYLCRYLLIIRRGVIIVSGGGGGGRLRRFRLGAAMAIIYMAMSTLFFLRIAPLTPWWGALAAWVMILLIVEAIFAFFFPYRCCFNETEGDVQNNPHV
IRIS_313-8486  MELITACALLLIAVHIAIVISCPEPLAAVALVDVPLLYLYRYLLIIRRGVIIVSGAGGGGGRLRRFRLGAAMAIIYMAMSTLFFLRIAPLTPWWGALAAWVMILLIVEAIFAFFFPYRCCFNETEGDVQNNPHV
IRIS_313-8492  MELITACALLLIAVHIAIVISCPEPLAAVALVDVPLLYLCRYLLIIRRGVIIVSGAGAGGGRLRRFRLGAAMAIIYMAMSTLFFLRIAPLTPWWGALAAWVMILLIVEAIFAFFFPYRCCFNETEGDVQNNPHV
IRIS_313-8493  MELITACALLLIAVHIAIVISCPEPLAAVALVDVPLLYLCRYLLIIRRGVIIVSGGGGGGRLRRFRLGAAMAIIYMAMSTLFFLRIAPLTPWWGALAAWVMILLIVEAIFAFFFPYRCCFNETEGDVQNNPHV
IRIS_313-8498  MELITACALLLIAVHIAIVISCPEPLAAVALVDVPLLYLCRYLLIIRRGVIIVSGAGAGGGRLRRFRLGAAMAIIYMAMSTLFFLRIAPLTPWWGALAAWVMILLIVEAIFAFFFPYRCCFNETEGDVQNNPHV
IRIS_313-8509  MELITACALLLIAVHIAIVISCPEPLAAVALVDVPLLYLCRYLLIIRRGVIIVSGAGAGGGRLRRFRLGAAMAIIYMAMSTLFFLRIAPLTPWWGALAAWVMILLIVEAIFAFFFPYRCCFNETEGDVQNNPHV
IRIS_313-8514  MELITACALLLIAVHIAIVISCPEPLAAVALVDVPLLYLCRYLLIIRRGVIIVSGGGGGGRLRRFRLGAAMAIIYMAMSTLFFLRIAPLTPWWGALAAWVMILLIVEAIFAFFFPYRCCFNETEGDVQNNPHV
IRIS_313-8519  MELITACALLLIAVHIAIVISCPEPLAAVALVDVPLLYLYRYLLIIRRGVIIVSGAGGGGGRLRRFRLGAAMAIIYMAMSTLFFLRIAPLTPWWGALAAWVMILLIVEAIFAFFFPYRCCFNETEGDVQNNPHV
IRIS_313-8523  MELITACALLLIAVHIAIVISCPEPLAAVALVDVPLLYLCRYLLIIRRGVIIVSGAGGGGGRLRRFRLGAAMAIIYMAMSTLFFLRIAPLTPWWGALAAWVMILLIVEAIFAFFFPYRCCFNETEGDVQNNPHV
IRIS_313-8530  MELITACALLLIAVHIAIVISCPEPLAAVALVDVPLLYLCRYLLIIRRGVIIVSGGGGGGRLRRFRLGAAMAIIYMAMSTLFFLRIAPLTPWWGALAAWVMILLIVEAIFAFFFPYRCCFNETEGDVQNNPHV
IRIS_313-8536  MELITACALLLIAVHIAIVISCPEPLAAVALVDVPLLYLCRYLLIIRRGVIIVSGGGGGGRLRRFRLGAAMAIIYMAMSTLFFLRIAPLTPWWGALAAWVMILLIVEAIFAFFFPYRCCFNETEGDVQNNPHV
IRIS_313-8554  MELITACALLLIAVHIAIVISCPEPLAAVALVDVPLLYLCRYLLIIRRGVIIVSGAGAGGGRLRRFRLGAAMAIIYMAMSTLFFLRIAPLTPWWGALAAWVMILLIVEAIFAFFFPYRCCFNETEGDVQNNPHV
IRIS_313-8557  MELITACALLLIAVHIAIVISCPEPLAAVALVDVPLLYLCRYLLIIRRGVIIVSGAGGAAAAGCAASASGQPWRSSTWPCRPSSSSALLLSRHGGAPSPPGS*SCSSSRPSSPSSSPTAAASTKPAKAMSRIIPM
IRIS_313-8559  MELITACALLLIAVHIAIVISCPEPLAAVALVDVPLLYLCRYLLIIRRGVIIVSGAGGAAAAGCAASASGQPWRSSTWPCRPSSSSALLLSRHGGAPSPPGS*SCSSSRPSSPSSSPTAAASTKPAKAMSRIIPM
IRIS_313-8565  MELITACALLLIAVHIAIVISCPEPLAAVALVDVPLLYLCRYLLIIRRGVIIVSGAGGGGGRLRRFRLGAAMAIIYMAMSTLFFLRIAPLTPWWGALAAWVMILLIVEAIFAFFFPYRCCFNETEGDVQNNPHV
IRIS_313-8567  MELITACALLLIAVHIAIVISCPEPLAAVALVDVPLLYLYRYLLIIRRGVIIVSGAGGGGGRLRRFRLGAAMAIIYMAMSTLFFLRIAPLTPWWGALAAWVMILLIVEAIFAFFFPYRCCFNETEGDVQNNPHV
IRIS_313-8568  MELITACALLLIAVHIAIVISCPEPLAAVALVDVPLLYLCRYLLIIRRGVIIVSGAGGAAAAGCAASASGQPWRSSTWPCRPSSSSALLLSRHGGAPSPPGS*SCSSSRPSSPSSSPTAAASTKPAKAMSRIIPM
IRIS_313-8571  MELITACALLLIAVHIAIVISCPEPLAAVALVDVPLLYLCRYLLIIRRGVIIVSGAGGGGRLRRFRLGAAMAIIYMAMSTLFFLRIAPLTPWWGALAAWVMILLIVEAIFAFFFPYRCCFNETEGDVQNNPHV
IRIS_313-8572  MELITACALLLIAVHIAIVISCPEPLAAVALVDVPLLYLCRYLLIIRRGVIIVSGAGAGGGRLRRFRLGAAMAIIYMAMSTLFFLRIAPLTPWWGALAAWVMILLIVEAIFAFFFPYRCCFNETEGDVQNNPHV
IRIS_313-8578  MELITACALLLIAVHIAIVISCPEPLAAVALVDVPLLYLYRYLLIIRRGVIIVSGAGGGGGRLRRFRLGAAMAIIYMAMSTLFFLRIAPLTPWWGALAAWVMILLIVEAIFAFFFPYRCCFNETEGDVQNNPHV
IRIS_313-8580  MELITACALLLIAVHIAIVISCPEPLAAVALVDVPLLYLCRYLLIIRRGVIIVSGAGAGGGRLRRFRLGAAMAIIYMAMSTLFFLRIAPLTPWWGALAAWVMILLIVEAIFAFFFPYRCCFNETEGDVQNNPHV
IRIS_313-8585  MELITACALLLIAVHIAIVISCPEPLAAVALVDVPLLYLCRYLLIIRRGVIIVSGGGGGGRLRRFRLGAAMAIIYMAMSTLFFLRIAPLTPWWGALAAWVMILLIVEAIFAFFFPYRCCFNETEGDVQNNPHV
IRIS_313-8586  MELITACALLLIAVHIAIVISCPEPLAAVALVDVPLLYLYRYLLIIRRGVIIVSGAGGGGGRLRRFRLGAAMAIIYMAMSTLFFLRIAPLTPWWGALAAWVMILLIVEAIFAFFFPYRCCFNETEGDVQNNPHV
IRIS_313-8591  MELITACALLLIAVHIAIVISCPEPLAAVALVDVPLLYLCRYLLIIRRGVIIVSGGGGGGRLRRFRLGAAMAIIYMAMSTLFFLRIAPLTPWWGALAAWVMILLIVEAIFAFFFPYRCCFNETEGDVQNNPHV
IRIS_313-8594  MELITACALLLIAVHIAIVISCPEPLAAVALVDVPLLYLYRYLLIIRRGVIIVSGAGGGGGRLRRFRLGAAMAIIYMAMSTLFFLRIAPLTPWWGALAAWVMILLIVEAIFAFFFPYRCCFNETEGDVQNNPHV
IRIS_313-8595  MELITACALLLIAVHIAIVISCPEPLAAVALVDVPLLYLCRYLLIIRRGVIIVSGAGGAAAAGCAASASGQPWRSSTWPCRPSSSSALLLSRHGGAPSPPGS*SCSSSRPSSPSSSPTAAASTKPAKAMSRIIPM
IRIS_313-8599  MELITACALLLIAVHIAIVISCPEPLAAVALVDVPLLYLYRYLLIIRRGVIIVSGAGGGGGRLRRFRLGAAMAIIYMAMSTLFFLRIAPLTPWWGALAAWVMILLIVEAIFAFFFPYRCCFNETEGDVQNNPHV
IRIS_313-8603  MELITACALLLIAVHIAIVISCPEPLAAVALVDVPLLYLCRYLLIIRRGVIIVSGAGGGGGGRLRRFRLGAAMAIIYMAMSTLFFLRIAPLTPWWGALAAWVMILLIVEAIFAFFFPYRCCFNETEGDVQNNPHV
IRIS_313-8606  MELITACALLLIAVHIAIVISCPEPLAAVALVDVPLLYLCRYLLIIRRGVIIVSGAGAGGGRLRRFRLGAAMAIIYMAMSTLFFLRIAPLTPWWGALAAWVMILLIVEAIFAFFFPYRCCFNETEGDVQNNPHV
IRIS_313-8608  MELITACALLLIAVHIAIVISCPEPLAAVALVDVPLLYLCRYLLIIRRGVIIVSGGGGGGRLRRFRLGAAMAIIYMAMSTLFFLRIAPLTPWWGALAAWVMILLIVEAIFAFFFPYRCCFNETEGDVQNNPHV

947 IRIS_313-8614 MELITACALLLIAVHIAIVISCPEPLAAVALVDVPLLYLCRYLLIIRRGVIIVSGGGGGGRLRRFRLGAAMAIIYMAMSTLFFLRIAPLTPWWGALAAWVMILLIVEAIFAFFFPYRCCFNETEGDVQNNPHV
948 IRIS_313-8616 MELITACALLLIAVHIAIVISCPEPLAAVALVDVPLLYLCRYLLIIRRGVIIVSGGGGGGRLRRFRLGAAMAIIYMAMSTLFFLRIAPLTPWWGALAAWVMILLIVEAIFAFFFPYRCCFNETEGDVQNNPHV
949 IRIS_313-8621 MELITACALLLIAVHIAIVISCPEPLAAVALVDVPLLYLCRYLLIIRRGVIIVSGAGAGGGRLRRFRLGAAMAIIYMAMSTLFFLRIAPLTPWWGALAAWVMILLIVEAIFAFFFPYRCCFNETEGDVQNNPHV
950 IRIS_313-8622 MELITACALLLIAVHIAIVISCPEPLAAVALVDVPLLYLCRYLLIIRRGVIIVSGAGGGGGRLRRFRLGAAMAIIYMAMSTLFFLRIAPLTPWWGALAAWVMILLIVEAIFAFFFPYRCCFNETEGDVQNNPHV
951 IRIS_313-8626 MELITACALLLIAVHIAIVISCPEPLAAVALVDVPLLYLCRYLLIIRRGVIIVSGAGGGGGRLRRFRLGAAMAIIYMAMSTLFFLRIAPLTPWWGALAAWVMILLIVEAIFAFFFPYRCCFNETEGDVQNNPHV
952 IRIS_313-8627 MELITACALLLIAVHIAIVISCPEPLAAVALVDVPLLYLYRYLLIIRRGVIIVSGAGGGGGRLRRFRLGAAMAIIYMAMSTLFFLRIAPLTPWWGALAAWVMILLIVEAIFAFFFPYRCCFNETEGDVQNNPHV
953 IRIS_313-8631 MELITACALLLIAVHIAIVISCPEPLAAVALVDVPLLYLCRYLLIIRRGVIIVSGAGGAAAAGCAASASGQPWRSSTWPCRPSSSSALLLSRHGGAPSPPGS*SCSSSRPSSPSSSPTAAASTKPKAMSRIIPM
954 IRIS_313-8632 MELITACALLLIAVHIAIVISCPEPLAAVALVDVPLLYLCRYLLIIRRGVIIVSGAGGGGGRLRRFRLGAAMAIIYMAMSTLFFLRIAPLTPWWGALAAWVMILLIVEAIFAFFFPYRCCFNETEGDVQNNPHV
955 IRIS_313-8637 MELITACALLLIAVHIAIVISCPEPLAAVALVDVPLLYLCRYLLIIRRGVIIVSGAGAGGGRLRRFRLGAAMAIIYMAMSTLFFLRIAPLTPWWGALAAWVMILLIVEAIFAFFFPYRCCFNETEGDVQNNPHV
956 IRIS_313-8638 MELITACALLLIAVHIAIVISCPEPLAAVALVDVPLLYLCRYLLIIRRGVIIVSGGGGGGRLRRFRLGAAMAIIYMAMSTLFFLRIAPLTPWWGALAAWVMILLIVEAIFAFFFPYRCCFNETEGDVQNNPHV
957 IRIS_313-8641 MELITACALLLIAVHIAIVISCPEPLAAVAFVNVPLLYLCRYLLIIRRGVIIVSGGGGGGGRLRRFRLGAAMAVIYMAMSTLFFLRIAPLTPWWGALAAWVMILLIVEAIFAFFFPYRCCFNETEGDVQNNPHV
958 IRIS_313-8643 MELITACALLLIAVHIAIVISCPEPLAAVALVDVPLLYLCRYLLIIRRGVIIVSGGGGGGRLRRFRLGAAMAIIYMAMSTLFFLRIAPLTPWWGALAAWVMILLIVEAIFAFFFPYRCCFNETEGDVQNNPHV
959 IRIS_313-8645 MELITACALLLIAVHIAIVISCPEPLAAVALVDVPLLYLCRYLLIIRRGVIIVSGGGAGGGRLRRFRLGAAMAIIYMAMSTLFFLRIAPLTPWWGALAAWVMILLIVEAIFAFFFPYRCCFNETEGDVQNNPHV
960 IRIS_313-8647 MELITACALLLIAVHIAIVISCPEPLAAVALVDVPLLYLCRYLLIIRRGVIIVSGAGGGGGGRLRRFRLGAAMAIIYMAMSTLFFLRIAPLTPWWGALAAWVMILLIVEAIFAFFFPYRCCFNETEGDVQNNPHV
961 IRIS_313-8655 MELITACALLLIAVHIAIVISCPEPLAAVAFVNVPLLYLCRYLLIIRRGVIIVSGGGGGGGRLRRFRLGAAMAVIYMAMSTLFFLRIAPLTPWWGALAAWVMILLIVEAIFAFFFPYRCCFNETEGDVQNNPHV
962 IRIS_313-8656 MELITACALLLIAVHIAIVISCPEPLAAVALVDVPLLYLCRYLLIIRRGVIIVSGAGAGGGRLRRFRLGAAMAIIYMAMSTLFFLRIAPLTPWWGALAAWVMILLIVEAIFAFFFPYRCCFNETEGDVQNNPHV
963 IRIS_313-8657 MELITACALLLIAVHIAIVISCPEPLAAVALVDVPLLYLCRYLLIIRRGVIIVSGGGGGGRLRRFRLGAAMAIIYMAMSTLFFLRIAPLTPWWGALAAWVMILLIVEAIFAFFFPYRCCFNETEGDVQNNPHV
964 IRIS_313-8658 MELITACALLLIAVHIAIVISCPEPLAAVALVDVPLLYLCRYLLIIRRGVIIVSGAGAGGGRLRRFRLGAAMAIIYMAMSTLFFLRIAPLTPWWGALAAWVMILLIVEAIFAFFFPYRCCFNETEGDVQNNPHV
965 IRIS_313-8659 MELITACALLLIAVHIAIVISCPEPLAAVALVDVPLLYLYRYLLIIRRGVIIVSGAGGGGGRLRRFRLGAAMAIIYMAMSTLFFLRIAPLTPWWGALAAWVMILLIVEAIFAFFFPYRCCFNETEGDVQNNPHV
966 IRIS_313-8660 MELITACALLLIAVHIAIVISCPEPLAAVALVDVPLLYLCRYLLIIRRGVIIVSGGGGGGRLRRFRLGAAMAIIYMAMSTLFFLRIAPLTPWWGALAAWVMILLIVEAIFAFFFPYRCCFNETEGDVQNNPHV
967 IRIS_313-8664 MELITACALLLIAVHIAIVISCPEPLAAVALVDVPLLYLYRYLLIIRRGVIIVSGAGGGGGRLRRFRLGAAMAIIYMAMSTLFFLRIAPLTPWWGALAAWVMILLIVEAIFAFFFPYRCCFNETEGDVQNNPHV
968 IRIS_313-8665 MELITACALLLIAVHIAIVISCPEPLAAVALVDVPLLYLCRYLLIIRRGVIIVSGAGAGGGRLRRFRLGAAMAIIYMAMSTLFFLRIAPLTPWWGALAAWVMILLIVEAIFAFFFPYRCCFNETEGDVQNNPHV
969 IRIS_313-8669 MELITACALLLIAVHIAIVISCPEPLAAVALVDVPLLYLCRYLLIIRRGVIIVSGAGGGGGRLRRFRLGAAMAIIYMAMSTLFFLRIAPLTPWWGALAAWVMILLIVEAIFAFFFPYRCCFNETEGDVQNNPHV
970 IRIS_313-8674 MELITACALLLIAVHIAIVISCPEPLAAVALVDVPLLYLCRYLLIIRRGVIIVSGGGAGGGRLRRFRLGAAMAIIYMAMSTLFFLRIAPLTPWWGALAAWVMILLIVEAIFAFFFPYRCCFNETEGDVQNNPHV
971 IRIS_313-8679 MELITACALLLIAVHIAIVISCPEPLAAVALVDVPLLYLCRYLLIIRRGVIIVSGAGAGGGRLRRFRLGAAMAIIYMVMSTLFFLRIAPLTPWWGALAAWVMILLIVEAIFAFFFPYRCCFNETEGDVQNNPHV
972 IRIS_313-8681 MELITACALLLIAVHIAIVISCPEPLAAVALVDVPLLYLCRYLLIIRRGVIIVSGAGAGGGRLRRFRLGAAMAIIYMAMSTLFFLRIAPLTPWWGALAAWVMILLIVEAIFAFFFPYRCCFNETEGDVQNNPHV
973 IRIS_313-8683 MELITACALLLIAVHIAIVISCPEPLAAVALVDVPLLYLCRYLLIIRRGVIIVSGGGGGGRLRRFRLGAAMAIIYMAMSTLFFLRIAPLTPWWGALAAWVMILLIVEAIFAFFFPYRCCFNETEGDVQNNPHV
974 IRIS_313-8687 MELITACALLLIAVHIAIVISCPEPLAAVALVDVPLLYLCRYLLIIRRGVIIVSGAGAGGGRLRRFRLGAAMAIIYMAMSTLFFLRIAPLTPWWGALAAWVMILLIVEAIFAFFFPYRCCFNETEGDVQNNPHV
975 IRIS_313-8690 MELITACALLLIAVHIAIVISCPEPLAAVALVDVPLLYLCRYLLIIRRGVIIVSGAGAGGGRLRRFRLGAAMAIIYMAMSTLFFLRIAPLTPWWGALAAWVMILLIVEAIFAFFFPYRCCFNETEGDVQNNPHV
976 IRIS_313-8694 MELITACALLLIAVHIAIVISCPEPLAAVALVDVPLLYLCRYLLIIRRGVIIVSGAGAGGGRLRRFRLGAAMAIIYMAMSTLFFLRIAPLTPWWGALAAWVMILLIVEAIFAFFFPYRCCFNETEGDVQNNPHV
977 IRIS_313-8697 MELITACALLLIAVHIAIVISCPEPLAAVALVDVPLLYLCRYLLIIRRGVIIVSGGGGGGRLRRFRLGAAMAIIYMAMSTLFFLRIAPLTPWWGALAAWVMILLIVEAIFAFFFPYRCCFNETEGDVQNNPHV
978 IRIS_313-8699 MELITACALLLIAVHIAIVISCPEPLAAVALVDVPLLYLCRYLLIIRRGVIIVSGAGAGGGRLRRFRLGAAMAIIYMAMSTLFFLRIAPLTPWWGALAAWVMILLIVEAIFAFFFPYRCCFNETEGDVQNNPHV
979 IRIS_313-8702 MELITACALLLIAVHIAIVISCPEPLAAVALVDVPLLYLCRYLLIIRRGVIIVSGGGAGGGRLRRFRLGAAMAIIYMAMSTLFFLRIAPLTPWWGALAAWVMILLIVEAIFAFFFPYRCCFNETEGDVQNNPHV
980 IRIS_313-8703 MELITACALLLIAVHIAIVISCPEPLAAVALVDVPLLYLCRYLLIIRRGVIIVSGAGAGGGRLRRFRLGAAMAIIYMAMSTLFFLRIAPLTPWWGALAAWVMILLIVEAIFAFFFPYRCCFNETEGDVQNNPHV
981 IRIS_313-8704 MELITACALLLIAVHIAIVISCPEPLAAVALVDVPLLYLCRYLLIIRRGVIIVSGAGAGGGRLRRFRLGAAMAIIYMAMSTLFFLRIAPLTPWWGALAAWVMILLIVEAIFAFFFPYRCCFNETEGDVQNNPHV
982 IRIS_313-8712 MELITACALLLIAVHIAIVISCPEPLAAVALVDVPLLYLCRYLLIIRRGVIIVSGAGAGGGRLRRFRLGAAMAIIYMAMSTLFFLRIAPLTPWWGALAAWVMILLIVEAIFAFFFPYRCCFNETEGDVQNNPHV
983 IRIS_313-8713 MELITACALLLIAVHIAIVISCPEPLAAVALVDVPLLYLCRYLLIIRRGVIIVSGGGGGGRLRRFRLGAAMAIIYMAMSTLFFLRIAPLTPWWGALAAWVMILLIVEAIFAFFFPYRCCFNETEGDVQNNPHV
984 IRIS_313-8717 MELITACALLLIAVHIAIVISCPEPLAAVALVDVPLLYLCRYLLIIRRGVIIVSGGGGGGRLRRFRLGAAMAIIYMAMSTLFFLRIAPLTPWWGALAAWVMILLIVEAIFAFFFPYRCCFNETEGDVQNNPHV
985 IRIS_313-8721 MELITACALLLIAVHIAIVISCPEPLAAVAFVNVPLLYLCRYLLIIRRGVIIVSGGGGGGGRLRRFRLGAAMAVIYMAMSTLFFLRIAPLTPWWGALAAWVMILLIVEAIFAFFFPYRCCFNETEGDVQNNPHV
986 IRIS_313-8722 MELITACALLLIAVHIAIVISCPEPLAAVALVDVPLLYLYRYLLIIRRGVIIVSGAGGGGGRLRRFRLGAAMAIIYMAMSTLFFLRIAPLTPWWGALAAWVMILLIVEAIFAFFFPYRCCFNETEGDVQNNPHV
987 IRIS_313-8723 MELITACALLLIAVHIAIVISCPEPLAAVALVDVPLLYLCRYLLIIRRGVIIVSGAGAGGGRLRRFRLGAAMAIIYMAMSTLFFLRIAPLTPWWGALAAWVMILLIVEAIFAFFFPYRCCFNETEGDVQNNPHV
988 IRIS_313-8725 MELITACALLLIAVHIAIVISCPEPLAAVALVDVPLLYLCRYLLIIRRGVIIVSGGGGGGRLRRFRLGAAMAIIYMAMSTLFFLRIAPLTPWWGALAAWVMILLIVEAIFAFFFPYRCCFNETEGDVQNNPHV
989 IRIS_313-8727 MELITACALLLIAVHIAIVISCPEPLAAVALVDVPLLYLCRYLLIIRRGVIIVSGAGGAAAAGCAASASGQPWRSSTWPCRPSSSSALLLSRHGGAPSPPGS*SCSSSRPSSPSSSPTAAASTKPKAMSRIIPM
990 IRIS_313-8731 MELITACALLLIAVHIAIVISCPEPLAAVALVDVPLLYLCRYLLIIRRGVIIVSGGGGGGRLRRFRLGAAMAIIYMAMSTLFFLRIAPLTPWWGALAAWVMILLIVEAIFAFFFPYRCCFNETEGDVQNNPHV
991 IRIS_313-8732 MELITACALLLIAVHIAIVISCPEPLAAVALVDVPLLYLCRYLLIIRRGVIIVSGAGAGGGRLRRFRLGAAMAIIYMAMSTLFFLRIAPLTPWWGALAAWVMILLIVEAIFAFFFPYRCCFNETEGDVQNNPHV
992 IRIS_313-8733 MELITACALLLIAVHIAIVISCPEPLAAVALVDVPLLYLCRYLLIIRRGVIIVSGAGAGGGRLRRFRLGAAMAIIYMAMSTLFFLRIAPLTPWWGALAAWVMILLIVEAIFAFFFPYRCCFNETEGDVQNNPHV
993 IRIS_313-8735 MELITACALLLIAVHIAIVISCPEPLAAVAFVNVPLLYLCRYLLIIRRGVIIVSGGGGGGGRLRRFRLGAAMAVIYMAMSTLFFLRIAPLTPWWGALAAWVMILLIVEAIFAFFFPYRCCFNETEGDVQNNPHV
994 IRIS_313-8737 MELITACALLLIAVHIAIVISCPEPLAAVALVDVPLLYLCRYLLIIRRGVIIVSGAGAGGGRLRRFRLGAAMAIIYMAMSTLFFLRIAPLTPWWGALAAWVMILLIVEAIFAFFFPYRCCFNETEGDVQNNPHV
995 IRIS_313-8739 MELITACALLLIAVHIAIVISCPEPLAAVALVDVPLLYLCRYLLIIRRGVIIVSGAGAGGGRLRRFRLGAAMAIIYMAMSTLFFLRIAPLTPWWGALAAWVMILLIVEAIFAFFFPYRCCFNETEGDVQNNPHV
996 IRIS_313-8743 MELITACALLLIAVHIAIVISCPEPLAAVALVDVPLLYLCRYLLIIRRGVIIVSGGGGGGRLRRFRLGAAMAIIYMAMSTLFFLRIAPLTPWWGALAAWVMILLIVEAIFAFFFPYRCCFNETEGDVQNNPHV
997 IRIS_313-8744 MELITACALLLIAVHIAIVISCPEPLAAVALVDVPLLYLCRYLLIIRRGVIIVSGGGAGGGRLRRFRLGAAMAIIYMAMSTLFFLRIAPLTPWWGALAAWVMILLIVEAIFAFFFPYRCCFNETEGDVQNNPHV
998 IRIS_313-8745 MELITACALLLIAVHIAIVISCPEPLAAVALVDVPLLYLCRYLLIIRRGVIIVSGAGAGGGRLRRFRLGAAMAIIYMAMSTLFFLRIAPLTPWWGALAAWVMILLIVEAIFAFFFPYRCCFNETEGDVQNNPHV
999 IRIS_313-8747 MELITACALLLIAVHIAIVISCPEPLAAVALVDVPLLYLCRYLLIIRRGVIIVSGAGAGGGRLRRFRLGAAMAIIYMAMSTLFFLRIAPLTPWWGALAAWVMILLIVEAIFAFFFPYRCCFNETEGDVQNNPHV
1000 IRIS_313-8751 MELITACALLLIAVHIAIVISCPEPLAAVALVDVPLLYLCRYLLIIRRGVIIVSGGGGGGRLRRFRLGAAMAIIYMAMSTLFFLRIAPLTPWWGALAAWVMILLIVEAIFAFFFPYRCCFNETEGDVQNNPHV
1001 IRIS_313-8754 MELITACALLLIAVHIAIVISCPEPLAAVALVDVPLLYLCRYLLIIRRGVIIVSGGGGGGRLRRFRLGAAMAIIYMAMSTLFFLRIAPLTPWWGALAAWVMILLIVEAIFAFFFPYRCCFNETEGDVQNNPHV
1002 IRIS_313-8755 MELITACALLLIAVHIAIVISCPEPLAAVALVDVPLLYLCRYLLIIRRGVIIVSGAGAGGGRLRRFRLGAAMAIIYMAMSTLFFLRIAPLTPWWGALAAWVMILLIVEAIFAFFFPYRCCFNETEGDVQNNPHV
1003 IRIS_313-8757 MELITACALLLIAVHIAIVISCPEPLAAVALVDVPLLYLCRYLLIIRRGVIIVSGGGGGGRLRRFRLGAAMAIIYMAMSTLFFLRIAPLTPWWGALAAWVMILLIVEAIFAFFFPYRCCFNETEGDVQNNPHV
1004 IRIS_313-8765 MELITACALLLIAVHIAIVISCPEPLAAVALVDVPLLYLCRYLLIIRRGVIIVSGAGAGGGRLRRFRLGAAMAIIYMAMSTLFFLRIAPLTPWWGALAAWVMILLIVEAIFAFFFPYRCCFNETEGDVQNNPHV
1005 IRIS_313-8767 MELITACALLLIAVHIAIVISCPEPLAAVALVDVPLLYLCRYLLIIRRGVIIVSGGGGGGRLRRFRLGAAMAIIYMAMSTLFFLRIAPLTPWWGALAAWVMILLIVEAIFAFFFPYRCCFNETEGDVQNNPHV
1006 IRIS_313-8768 MELITACALLLIAVHIAIVISCPEPLAAVALVDVPLLYLCRYLLIIRRGVIIVSGAGAGGGRLRRFRLGAAMAIIYMAMSTLFFLRIAPLTPWWGALAAWVMILLIVEAIFAFFFPYRCCFNETEGDVQNNPHV
1007 IRIS_313-8769 MELITACALLLIAVHIAIVISCPEPLAAVALVDVPLLYLYRYLLIIRRGVIIVSGAGGGGGRLRRFRLGAAMAIIYMAMSTLFFLRIAPLTPWWGALAAWVMILLIVEAIFAFFFPYRCCFNETEGDVQNNPHV
1008 IRIS_313-8771 MELITACALLLIAVHIAIVISCPEPLAAVALVDVPLLYLCRYLLIIRRGVIIVSGAGAGGGRLRRFRLGAAMAIIYMAMSTLFFLRIAPLTPWWGALAAWVMILLIVEAIFAFFFPYRCCFNETEGDVQNNPHV
1009 IRIS_313-8778 MELITACALLLIAVHIAIVISCPEPLAAVALVDVPLLYLCRYLLIIRRGVIIVSGAGAGGGRLRRFRLGAAMAIIYMAMSTLFFLRIAPLTPWWGALAAWVMILLIVEAIFAFFFPYRCCFNETEGDVQNNPHV
1010 IRIS_313-8781 MELITACALLLIAVHIAIVISCPEPLAAVALVDVPLLYLCRYLLIIRRGVIIVSGGGGGGRLRRFRLGAAMAIIYMAMSTLFFLRIAPLTPWWGALAAWVMILLIVEAIFAFFFPYRCCFNETEGDVQNNPHV
1011 IRIS_313-8783 MELITACALLLIAVHIAIVISCPEPLAAVALVDVPLLYLCRYLLIIRRGVIIVSGGGGGGGRLRRFRLGAAMAVIYMAMSTLFFLRIAPLTPWWGALAAWVMILLIVEAIFAFFFPYRCCFNETEGDVQNNPHV
1012 IRIS_313-8785 MELITACALLLIAVHIAIVISCPEPLAAVALVDVPLLYLCRYLLIIRRGVIIVSGAGAGGGRLRRFRLGAAMAIIYMVMSTLFFLRIAPLTPWWGALAAWVMILLIVEAIFAFFFPYRCCFNETEGDVQNNPHV
1013 IRIS_313-8789 MELITACALLLIAVHIAIVISCPEPLAAVAFVNVPLLYLCRYLLIIRRGVIIVSGGGGGGGRLRRFRLGAAMAVIYMAMSTLFFLRIAPLTPWWGALAAWVMILLIVEAIFAFFFPYRCCFNETEGDVQNNPHV
1014 IRIS_313-8791 MELITACALLLIAVHIAIVISCPEPLAAVALVDVPLLYLCRYLLIIRRGVIIVSGGGAGGGRLRRFRLGAAMAIIYMAMSTLFFLRIAPLTPWWGALAAWVMILLIVEAIFAFFFPYRCCFNETEGDVQNNPHV
1015 IRIS_313-8793 MELITACALLLIAVHIAIVISCPEPLAAVALVDVPLLYLCRYLLIIRRGVIIVSGGGAGGGRLRRFRLGAAMAIIYMAMSTLFFLRIAPLTPWWGALAAWVMILLIVEAIFAFFFPYRCCFNETEGDVQNNPHV
1016 IRIS_313-8796 MELITACALLLIAVHIAIVISCPEPLAAVALVDVPLLYLCRYLLIIRRGVIIVSGGGGGGRLRRFRLGAAMAIIYMAMSTLFFLRIAPLTPWWGALAAWVMILLIVEAIFAFFFPYRCCFNETEGDVQNNPHV
1017 IRIS_313-8803 MELITACALLLIAVHIAIVISCPEPLAAVALVDVPLLYLYRYLLIIRRGVIIVSGAGGGGGRLRRFRLGAAMAIIYMAMSTLFFLRIAPLTPWWGALAAWVMILLIVEAIFAFFFPYRCCFNETEGDVQNNPHV
1018 IRIS_313-8811 MELITACALLLIAVHIAIVISCPEPLAAVALVDVPLLYLCRYLLIIRRGVIIVSGAGGGGGGRLRRFRLGAAMAIIYMAMSTLFFLRIAPLTPWWGALAAWVMILLIVEAIFAFFFPYRCCFNETEGDVQNNPHV
1019 IRIS_313-8812 MELITACALLLIAVHIAIVISCPEPLAAVALVDVPLLYLCRYLLIIRRGVIIVSGGGGGGRLRRFRLGAAMAIIYMAMSTLFFLRIAPLTPWWGALAAWVMILLIVEAIFAFFFPYRCCFNETEGDVQNNPHV
1020 IRIS_313-8813 MELITACALLLIAVHIAIVISCPEPLAAVALVDVPLLYLCRYLLIIRRGVIIVSGAGAGGGRLRRFRLGAAMAIIYMAMSTLFFLRIAPLTPWWGALAAWVMILLIVEAIFAFFFPYRCCFNETEGDVQNNPHV
1021 IRIS_313-8814 MELITACALLLIAVHIAIVISCPEPLAAVALVDVPLLYLCRYLLIIRRGVIIVSGAGAGGGRLRRFRLGAAMAIIYMAMSTLFFLRIAPLTPWWGALAAWVMILLIVEAIFAFFFPYRCCFNETEGDVQNNPHV
1022 IRIS_313-8815 MELITACALLLIAVHIAIVISCPEPLAAVALVDVPLLYLCRYLLIIRRGVIIVSGAGAGGGRLRRFRLGAAMAIIYMAMSTLFFLRIAPLTPWWGALAAWVMILLIVEAIFAFFFPYRCCFNETEGDVQNNPHV
1023 IRIS_313-8822 MELITACALLLIAVHIAIVISCPEPLAAVAFVNVPLLYLCRYLLIIRRGVIIVSGGGGGGGRLRRFRLGAAMAVIYMAMSTLFFLRIAPLTPWWGALAAWVMILLIVEAIFAFFFPYRCCFNETEGDVQNNPHV
1024 IRIS_313-8831 MELITACALLLIAVHIAIVISCPEPLAAVALVDVPLLYLCRYLLIIRRGVIIVSGAGGAAAAGCAASASGQPWRSSTWPCRPSSSSALLLSRHGGAPSPPGS*SCSSSRPSSPSSSPTAAASTKPKAMSRIIPM
1025 IRIS_313-8833 MELITACALLLIAVHIAIVISCPEPLAAVALVDVPLLYLCRYLLIIRRGVIIVSGGGGGGRLRRFRLGAAMAIIYMAMSTLFFLRIAPLTPWWGALAAWVMILLIVEAIFAFFFPYRCCFNETEGDVQNNPHV
1026 IRIS_313-8844 MELITACALLLIAVHIAIVISCPEPLAAVALVDVPLLYLCRYLLIIRRGVIIVSGGGGGGRLRRFRLGAAMAIIYMAMSTLFFLRIAPLTPWWGALAAWVMILLIVEAIFAFFFPYRCCFNETEGDVQNNPHV
1027 IRIS_313-8846 MELITACALLLIAVHIAIVISCPEPLAAVALVDVPLLYLCRYLLIIRRGVIIVSGAGAGGGRLRRFRLGAAMAIIYMAMSTLFFLRIAPLTPWWGALAAWVMILLIVEAIFAFFFPYRCCFNETEGDVQNNPHV
1028 IRIS_313-8850 MELITACALLLIAVHIAIVISCPEPLAAVALVDVPLLYLCRYLLIIRRGVIIVSGAGAGGGRLRRFHLGAAMAIIYMAMSTLFFLRIAPLTPWWGALAAWVMILLIVEAIFAFFFPYRCCFNETEGDVQNNPHV
1029 IRIS_313-8854 MELITACALLLIAVHIAIVISCPEPLAAVALVDVPLLYLCRYLLIIRRGVIIVSGAGAGGGRLRRFRLGAAMAIIYMAMSTLFFLRIAPLTPWWGALAAWVMILLIVEAIFAFFFPYRCCFNETEGDVQNNPHV
1030 IRIS_313-8856 MELITACALLLIAVHIAIVISCPEPLAAVALVDVPLLYLCRYLLIIRRGVIIVSGAGAGGGRLRRFRLGAAMAIIYMAMSTLFFLRIAPLTPWWGALAAWVMILLIVEAIFAFFFPYRCCFNETEGDVQNNPHV
1031 IRIS_313-8857 MELITACALLLIAVHIAIVISCPEPLAAVALVDVPLLYLCRYLLIIRRGVIIVSGAGAGGGRLRRFRLGAAMAIIYMAMSTLFFLRIAPLTPWWGALAAWVMILLIVEAIFAFFFPYRCCFNETEGDVQNNPHV
1032 IRIS_313-8859 MELITACALLLIAVHIAIVISCPEPLAAVALVDVPLLYLYRYLLIIRRGVIIVSGAGGGGGRLRRFRLGAAMAIIYMAMSTLFFLRIAPLTPWWGALAAWVMILLIVEAIFAFFFPYRCCFNETEGDVQNNPHV
1033 IRIS_313-8864 MELITACALLLIAVHIAIVISCPEPLAAVAFVNVPLLYLCRYLLIIRRGVIIVSGGGGGGGRLRRFRLGAAMAVIYMAMSTLFFLRIAPLTPWWGALAAWVMILLIVEAIFAFFFPYRCCFNETEGDVQNNPHV
1034 IRIS_313-8865 MELITACALLLIAVHIAIVISCPEPLAAVALVDVPLLYLCRYLLIIRRGVIIVSGAGGGGGRLRRFRLGAAMAIIYMAMSTLFFLRIAPLTPWWGALAAWVMILLIVEAIFAFFFPYRCCFNETEGDVQNNPHV
1035 IRIS_313-8870 MELITACALLLIAVHIAIVISCPEPLAAVALVDVPLLYLYRYLLIIRRGVIIVSGAGGGGGRLRRFRLGAAMAIIYMAMSTLFFLRIAPLTPWWGALAAWVMILLIVEAIFAFFFPYRCCFNETEGDVQNNPHV
1036 IRIS_313-8872 MELITACALLLIAVHIAIVISCPEPLAAVALVDVPLLYLCRYLLIIRRGVIIVSGAGGGGGRLRRFRLGAAMAIIYMAMSTLFFLRIAPLTPWWGALAAWVMILLIVEAIFAFFFPYRCCFNETEGDVQNNPHV
1037 IRIS_313-8873 MELITACALLLIAVHIAIVISCPEPLAAVALVDVPLLYLYRYLLIIRRGVIIVSGAGGGGGRLRRFRLGAAMAIIYMAMSTLFFLRIAPLTPWWGALAAWVMILLIVEAIFAFFFPYRCCFNETEGDVQNNPHV
1038 IRIS_313-8876 MELITACALLLIAVHIAIVISCPEPLAAVALVDVPLLYLCRYLLIIRRGVIIVSGAGAGGGRLRRFRLGAAMAIIYMAMSTLFFLRIAPLTPWWGALAAWVMILLIVEAIFAFFFPYRCCFNETEGDVQNNPHV
1039 IRIS_313-8880 MELITACALLLIAVHIAIVISCPEPLAAVALVDVPLLYLCRYLLIIRRGVIIVSGAGGGGGRLRRFRLGAAMAIIYMAMSTLFFLRIAPLTPWWGALAAWVMILLIVEAIFAFFFPYRCCFNETEGDVQNNPHV
1040 IRIS_313-8883 MELITACALLLIAVHIAIVISCPEPLAAVALVDVPLLYLCRYLLIIRRGVIIVSGAGGGGGRLRRFRLGAAMAIIYMAMSTLFFLRIAPLTPWWGALAAWVMILLIVEAIFAFFFPYRCCFNETEGDVQNNPHV
1041 IRIS_313-8884 MELITACALLLIAVHIAIVISCPEPLAAVALVDVPLLYLYRYLLIIRRGVIIVSGAGGGGGRLRRFRLGAAMAIIYMAMSTLFFLRIAPLTPWWGALAAWVMILLIVEAIFAFFFPYRCCFNETEGDVQNNPHV
1042 IRIS_313-8889 MELITACALLLIAVHIAIVISCPEPLAAVALVDVPLLYLCRYLLIIRRGVIIVSGGGAGGGRLRRFRLGAAMAIIYMAMSTLFFLRIAPLTPWWGALAAWVMILLIVEAIFAFFFPYRCCFNETEGDVQNNPHV
1043 IRIS_313-8890 MELITACALLLIAVHIAIVISCPEPLAAVALVDVPLLYLCRYLLIIRRGVIIVSGAGAGGGRLRRFRLGAAMAIIYMAMSTLFFLRIAPLTPWWGALAAWVMILLIVEAIFAFFFPYRCCFNETEGDVQNNPHV
1044 IRIS_313-8894 MELITACALLLIAVHIAIVISCPEPLAAVALVDVPLLYLCRYLLIIRRGVIIVSGAGAGGGRLRRFRLGAAMAIIYMAMSTLFFLRIAPLTPWWGALAAWVMILLIVEAIFAFFFPYRCCFNETEGDVQNNPHV
1045 IRIS_313-8895 MELITACALLLIAVHIAIVISCPEPLAAVALVDVPLLYLCRYLLIIRRGVIIVSGAGAGGGRLRRFRLGAAMAIIYMAMSTLFFLRIAPLTPWWGALAAWVMILLIVEAIFAFFFPYRCCFNETEGDVQNNPHV
1046 IRIS_313-8900 MELITACALLLIAVHIAIVISCPEPLAAVALVDVPLLYLCRYLLIIRRGVIIVSGAGAGGGRLRRFRLGAAMAIIYMAMSTLFFLRIAPLTPWWGALAAWVMILLIVEAIFAFFFPYRCCFNETEGDVQNNPHV
1047 IRIS_313-8903 MELITACALLLIAVHIAIVISCPEPLAAVALVDVPLLYLCRYLLIIRRGVIIVSGAGAGGGRLRRFRLGAAMAIIYMAMSTLFFLRIAPLTPWWGALAAWVMILLIVEAIFAFFFPYRCCFNETEGDVQNNPHV
1048 IRIS_313-8909 MELITACALLLIAVHIAIVISCPEPLAAVALVDVPLLYLCRYLLIIRRGVIIVSGGGGGGRLRRFRLGAAMAIIYMAMSTLFFLRIAPLTPWWGALAAWVMILLIVEAIFAFFFPYRCCFNETEGDVQNNPHV
1049 IRIS_313-8911 MELITACALLLIAVHIAIVISCPEPLAAVALVDVPLLYLCRYLLIIRRGVIIVSGAGAGGGRLRRFRLGAAMAIIYMAMSTLFFLRIAPLTPWWGALAAWVMILLIVEAIFAFFFPYRCCFNETEGDVQNNPHV
1050 IRIS_313-8914 MELITACALLLIAVHIAIVISCPEPLAAVALVDVPLLYLCRYLLIIRRGVIIVSGGGGGGRLRRFRLGAAMAIIYMAMSTLFFLRIAPLTPWWGALAAWVMILLIVEAIFAFFFPYRCCFNETEGDVQNNPHV
1051 IRIS_313-8920 MELITACALLLIAVHIAIVISCPEPLAAVALVDVPLLYLCRYLLIIRRGVIIVSGGGGGGRLRRFRLGAAMAIIYMAMSTLFFLRIAPLTPWWGALAAWVMILLIVEAIFAFFFPYRCCFNETEGDVQNNPHV
1052 IRIS_313-8922 MELITACALLLIAVHIAIVISCPEPLAAVALVDVPLLYLCRYLLIIRRGVIIVSGGGAGGGRLRRFRLGAAMAIIYMAMSTLFFLRIAPLTPWWGALAAWVMILLIVEAIFAFFFPYRCCFNETEGDVQNNPHV
1053 IRIS_313-8923 MELITACALLLIAVHIAIVISCPEPLAAVALVDVPLLYLYRYLLIIRRGVIIVSGAGGGGGRLRRFRLGAAMAIIYMAMSTLFFLRIAPLTPWWGALAAWVMILLIVEAIFAFFFPYRCCFNETEGDVQNNPHV
1054 IRIS_313-8924 MELITACALLLIAVHIAIVISCPEPLAAVALVDVPLLYLCRYLLIIRRGVIIVSGGGGGGRLRRFRLGAAMAIIYMAMSTLFFLRIAPLTPWWGALAAWVMILLIVEAIFAFFFPYRCCFNETEGDVQNNPHV
1055 IRIS_313-8925 MELITACALLLIAVHIAIVISCPEPLAAVALVDVPLLYLCRYLLIIRRGVIIVSGAGAGGGRLRRFRLGAAMAIIYMAMSTLFFLRIAPLTPWWGALAAWVMILLIVEAIFAFFFPYRCCFNETEGDVQNNPHV
1056 IRIS_313-8927 MELITACALLLIAVHIAIVISCPEPLAAVALVDVPLLYLCRYLLIIRRGVIIVSGAGGGGGRLRRFRLGAAMAIIYMAMSTLFFLRIAPLTPWWGALAAWVMILLIVEAIFAFFFPYRCCFNETEGDVQNNPHV
1057 IRIS_313-8930 MELITACALLLIAVHIAIVISCPEPLAAVALVDVPLLYLCRYLLIIRRGVIIVSGAGAGGGRLRRFRLGAAMAIIYMAMSTLFFLRIAPLTPWWGALAAWVMILLIVEAIFAFFFPYRCCFNETEGDVQNNPHV
1058 IRIS_313-8932 MELITACALLLIAVHIAIVISCPEPLAAVALVDVPLLYLCRYLLIIRRGVIIVSGAGGAAAAGCAASASGQPWRSSTWPCRPSSSSALLLSRHGGAPSPPGS*SCSSSRPSSPSSSPTAAASTKPKAMSRIIPM
1059 IRIS_313-8935 MELITACALLLIAVHIAIVISCPEPLAAVALVDVPLLYLCRYLLIIRRGVIIVSGGGGGGRLRRFRLGAAMAIIYMAMSTLFFLRIAPLTPWWGALAAWVMILLIVEAIFAFFFPYRCCFNETEGDVQNNPHV
1060 IRIS_313-8940 MELITACALLLIAVHIAIVISCPEPLAAVALVDVPLLYLCRYLLIIRRGVIIVSGAGAGGGRLRRFRLGAAMAIIYMAMSTLFFLRIAPLTPWWGALAAWVMILLIVEAIFAFFFPYRCCFNETEGDVQNNPHV
1061 IRIS_313-8946 MELITACALLLIAVHIAIVISCPEPLAAVALVDVPLLYLCRYLLIIRRGVIIVSGGGGGGRLRRFRLGAAMAIIYMAMSTLFFLRIAPLTPWWGALAAWVMILLIVEAIFAFFFPYRCCFNETEGDVQNNPHV
1062 IRIS_313-8948 MELITACALLLIAVHIAIVISCPEPLAAVALVDVPLLYLCRYLLIIRRGVIIVSGGGGGGRLRRFRLGAAMAIIYMAMSTLFFLRIAPLTPWWGALAAWVMILLIVEAIFAFFFPYRCCFNETEGDVQNNPHV
1063 IRIS_313-8955 MELITACALLLIAVHIAIVISCPEPLAAVALVDVPLLYLCRYLLIIRRGVIIVSGAGGGGGRLRRFRLGAAMAIIYMAMSTLFFLRIAPLTPWWGALAAWVMILLIVEAIFAFFFPYRCCFNETEGDVQNNPHV
1064 IRIS_313-8956 MELITACALLLIAVHIAIVISCPEPLAAVALVDVPLLYLCRYLLIIRRGVIIVSGGGGGGRLRRFRLGAAMAIIYMAMSTLFFLRIAPLTPWWGALAAWVMILLIVEAIFAFFFPYRCCFNETEGDVQNNPHV
1065 IRIS_313-8957 MELITACALLLIAVHIAIVISCPEPLAAVALVDVPLLYLCRYLLIIRRGVIIVSGGGGGGRLRRFRLGAAMAIIYMAMSTLFFLRIAPLTPWWGALAAWVMILLIVEAIFAFFFPYRCCFNETEGDVQNNPHV
1066 IRIS_313-8960 MELITACALLLIAVHIAIVISCPEPLAAVALVDVPLLYLYRYLLIIRRGVIIVSGAGGGGGRLRRFRLGAAMAIIYMAMSTLFFLRIAPLTPWWGALAAWVMILLIVEAIFAFFFPYRCCFNETEGDVQNNPHV
1067 IRIS_313-8963 MELITACALLLIAVHIAIVISCPEPLAAVALVDVPLLYLCRYLLIIRRGVIIVSGAGAGGGRLRRFRLGAAMAIIYMAMSTLFFLRIAPLTPWWGALAAWVMILLIVEAIFAFFFPYRCCFNETEGDVQNNPHV
1068 IRIS_313-8967 MELITACALLLIAVHIAIVISCPEPLAAVALVDVPLLYLCRYLLIIRRGVIIVSGGGGGGRLRRFRLGAAMAIIYMAMSTLFFLRIAPLTPWWGALAAWVMILLIVEAIFAFFFPYRCCFNETEGDVQNNPHV
1069 IRIS_313-8968 MELITACALLLIAVHIAIVISCPEPLAAVALVDVPLLYLCRYLLIIRRGVIIVSGGGGGGRLRRFRLGAAMAIIYMAMSTLFFLRIAPLTPWWGALAAWVMILLIVEAIFAFFFPYRCCFNETEGDVQNNPHV

| | |
|---|---|
| IRIS_313-8976 | MELITACALLLIAVHIAIVISCPEPLAAVALVDVPLLYLYRYLLIIRRGVIIVSGAGGGGGRLRRFRLGAAMAIIYMAMSTLFFLRIAPLTPWWGALAAWVMILLIVEAIFAFFFPYRCCFNETEGDVQNNPHV |
| IRIS_313-8978 | MELITACALLLIAVHIAIVISCPEPLAAVALVDVPLLYLYRYLLIIRRGVIIVSGAGGGGGRLRRFRLGAAMAIIYMAMSTLFFLRIAPLTPWWGALAAWVMILLIVEAIFAFFFPYRCCFNETEGDVQNNPHV |
| IRIS_313-8980 | MELITACALLLIAVHIAIVISCPEPLAAVALVDVPLLYLYRYLLIIRRGVIIVSGAGGGGGRLRRFRLGAAMAIIYMAMSTLFFLRIAPLTPWWGALAAWVMILLIVEAIFAFFFPYRCCFNETEGDVQNNPHV |
| IRIS_313-8982 | MELITACALLLIAVHIAIVISCPEPLAAVALVDVPLLYLCRYLLIIRRGVIIVSGAGAGGGRLRRFRLGAAMAIIYMAMSTLFFLRIAPLTPWWGALAAWVMILLIVEAIFAFFFPYRCCFNETEGDVQNNPHV |
| IRIS_313-8985 | MELITACALLLIAVHIAIVISCPEPLAAVALVDVPLLYLCRYLLIIRRGVIIVSGGGGGGRLRRFRLGAAMAIIYMAMSTLFFLRIAPLTPWWGALAAWVMILLIVEAIFAFFFPYRCCFNETEGDVQNNPHV |
| IRIS_313-8986 | MELITACALLLIAVHIAIVISCPEPLAAVALVDVPLLYLCRYLLIIRRGVIIVSGAGAGGGRLRRFRLGAAMAIIYMAMSTLFFLRIAPLTPWWGALAAWVMILLIVEAIFAFFFPYRCCFNETEGDVQNNPHV |
| IRIS_313-8987 | MELITACALLLIAVHIAIVISCPEPLAAVALVDVPLLYLCRYLLIIRRGVIIVSGGGGGGRLRRFRLGAAMAIIYMAMSTLFFLRIAPLTPWWGALAAWVMILLIVEAIFAFFFPYRCCFNETEGDVQNNPHV |
| IRIS_313-8988 | MELITACALLLIAVHIAIVISCPEPLAAVALVDVPLLYLCRYLLIIRRGVIIVSGGGGGGRLRRFRLGAAMAIIYMAMSTLFFLRIAPLTPWWGALAAWVMILLIVEAIFAFFFPYRCCFNETEGDVQNNPHV |
| IRIS_313-8994 | MELITACALLLIAVHIAIVISCPEPLAAVALVDVPLLYLCRYLLIIRRGVIIVSGAGGGGRLRRFRLGAAMAIIYMAMSTLFFLRIAPLTPWWGALAAWVMILLIVEAIFAFFFPYRCCFNETEGDVQNNPHV |
| IRIS_313-8996 | MELITACALLLIAVHIAIVISCPEPLAAVALVDVPLLYLYRYLLIIRRGVIIVSGAGGGGGRLRRFRLGAAMAIIYMAMSTLFFLRIAPLTPWWGALAAWVMILLIVEAIFAFFFPYRCCFNETEGDVQNNPHV |
| IRIS_313-8998 | MELITACALLLIAVHIAIVISCPEPLAAVALVDVPLLYLCRYLLIIRRGVIIVSGAGAGGGRLRRFRLGAAMAIIYMAMSTLFFLRIAPLTPWWGALAAWVMILLIVEAIFAFFFPYRCCFNETEGDVQNNPHV |
| IRIS_313-8999 | MELITACALLLIAVHIAIVISCPEPLAAVALVDVPLLYLYRYLLIIRRGVIIVSGAGGGGGRLRRFRLGAAMAIIYMAMSTLFFLRIAPLTPWWGALAAWVMILLIVEAIFAFFFPYRCCFNETEGDVQNNPHV |
| IRIS_313-9002 | MELITACALLLIAVHIAIVISCPEPLAAVALVDVPLLYLCRYLLIIRRGVIIVSGAGAGGGRLRRFRLGAAMAIIYMAMSTLFFLRIAPLTPWWGALAAWVMILLIVEAIFAFFFPYRCCFNETEGDVQNNPHV |
| IRIS_313-9005 | MELITACALLLIAVHIAIVISCPEPLAAVALVDVPLLYLCRYLLIIRRGVIIVSGGGGGGRLRRFRLGAAMAIIYMAMSTLFFLRIAPLTPWWGALAAWVMILLIVEAIFAFFFPYRCCFNETEGDVQNNPHV |
| IRIS_313-9006 | MELITACALLLIAVHIAIVISCPEPLAAVALVDVPLLYLCRYLLIIRRGVIIVSGGGGGGRLRRFRLGAAMAIIYMAMSTLFFLRIAPLTPWWGALAAWVMILLIVEAIFAFFFPYRCCFNETEGDVQNNPHV |
| IRIS_313-9017 | MELITACALLLIAVHIAIVISCPEPLAAVALVDVPLLYLCRYLLIIRRGVIIVSGGGGGGRLRRFRLGAAMAIIYMAMSTLFFLRIAPLTPWWGALAAWVMILLIVEAIFAFFFPYRCCFNETEGDVQNNPHV |
| IRIS_313-9019 | MELITACALLLIAVHIAIVISCPEPLAAVALVDVPLLYLYRYLLIIRRGVIIVSGAGGGGGRLRRFRLGAAMAIIYMAMSTLFFLRIAPLTPWWGALAAWVMILLIVEAIFAFFFPYRCCFNETEGDVQNNPHV |
| IRIS_313-9020 | MELITACALLLIAVHIAIVISCPEPLAAVALVDVPLLYLYRYLLIIRRGVIIVSGAGGGGGRLRRFRLGAAMAIIYMAMSTLFFLRIAPLTPWWGALAAWVMILLIVEAIFAFFFPYRCCFNETEGDVQNNPHV |
| IRIS_313-9023 | MELITACALLLIAVHIAIVISCPEPLAAVALVDVPLLYLCRYLLIIRRGVIIVSGAGAGGGRLRRFRLGAAMAIIYMAMSTLFFLRIAPLTPWWGALAAWVMILLIVEAIFAFFFPYRCCFNETEGDVQNNPHV |
| IRIS_313-9032 | MELITACALLLIAVHIAIVISCPEPLAAVALVDVPLLYLCRYLLIIRRGVIIVSGGGGGGRLRRFRLGAAMAIIYMAMSTLFFLRIAPLTPWWGALAAWVMILLIVEAIFAFFFPYRCCFNETEGDVQNNPHV |
| IRIS_313-9039 | MELITACALLLIAVHIAIVISCPEPLAAVALVDVPLLYLCRYLLIIRRGVIIVSGGGAGGGRLRRFRLGAAMAIIYMAMSTLFFLRIAPLTPWWGALAAWVMILLIVEAIFAFFFPYRCCFNETEGDVQNNPHV |
| IRIS_313-9044 | MELITACALLLIAVHIAIVISCPEPLAAVALVDVPLLYLCRYLLIIRRGVIIVSGGGGGGRLRRFRLGAAMAIIYMAMSTLFFLRIAPLTPWWGALAAWVMILLIVEAIFAFFFPYRCCFNETEGDVQNNPHV |
| IRIS_313-9048 | MELITACALLLIAVHIAIVISCPEPLAAVALVDVPLLYLYRYLLIIRRGVIIVSGAGGGGGRLRRFRLGAAMAIIYMAMSTLFFLRIAPLTPWWGALAAWVMILLIVEAIFAFFFPYRCCFNETEGDVQNNPHV |
| IRIS_313-9049 | MELITACALLLIAVHIAIVISCPEPLAAVALVDVPLLYLCRYLLIIRRGVIIVSGGGGGGRLRRFRLGAAMAIIYMAMSTLFFLRIAPLTPWWGALAAWVMILLIVEAIFAFFFPYRCCFNETEGDVQNNPHV |
| IRIS_313-9050 | MELITACALLLIAVHIAIVISCPEPLAAVALVDVPLLYLCRYLLIIRRGVIIVSGAGGGGGGRLRRFRLGAAMAIIYMAMSTLFFLRIAPLTPWWGALAAWVMILLIVEAIFAFFFPYRCCFNETEGDVQNNPHV |
| IRIS_313-9053 | MELITACALLLIAVHIAIVISCPEPLAAVALVDVPLLYLCRYLLIIRRGVIIVSGAGAGGGRLRRFRLGAAMAIIYMAMSTLFFLRIAPLTPWWGALAAWVMILLIVEAIFAFFFPYRCCFNETEGDVQNNPHV |
| IRIS_313-9054 | MELITACALLLIAVHIAIVISCPEPLAAVALVDVPLLYLCRYLLIIRRGVIIVSGGGGGGRLRRFRLGAAMAIIYMAMSTLFFLRIAPLTPWWGALAAWVMILLIVEAIFAFFFPYRCCFNETEGDVQNNPHV |
| IRIS_313-9062 | MELITACALLLIAVHIAIVISCPEPLAAVALVDVPLLYLYRYLLIIRRGVIIVSGAGGGGGRLRRFRLGAAMAIIYMAMSTLFFLRIAPLTPWWGALAAWVMILLIVEAIFAFFFPYRCCFNETEGDVQNNPHV |
| IRIS_313-9065 | MELITACALLLIAVHIAIVISCPEPLAAVALVDVPLLYLCRYLLIIRRGVIIVSGAGAGGGRLRRFRLGAAMAIIYMAMSTLFFLRIAPLTPWWGALAAWVMILLIVEAIFAFFFPYRCCFNETEGDVQNNPHV |
| IRIS_313-9066 | MELITACALLLIAVHIAIVISCPEPLAAVALVDVPLLYLCRYLLIIRRGVIIVSGGGGGGRLRRFRLGAAMAIIYMAMSTLFFLRIAPLTPWWGALAAWVMILLIVEAIFAFFFPYRCCFNETEGDVQNNPHV |
| IRIS_313-9067 | MELITACALLLIAVHIAIVISCPEPLAAVALVDVPLLYLCRYLLIIRRGVIIVSGAGAGGGRLRRFRLGAAMAIIYMAMSTLFFLRIAPLTPWWGALAAWVMILLIVEAIFAFFFPYRCCFNETEGDVQNNPHV |
| IRIS_313-9070 | MELITACALLLIAVHIAIVISCPEPLAAVALVDVPLLYLCRYLLIIRRGVIIVSGAGAGGGRLRRFRLGAAMAIIYMAMSTLFFLRIAPLTPWWGALAAWVMILLIVEAIFAFFFPYRCCFNETEGDVQNNPHV |
| IRIS_313-9072 | MELITACALLLIAVHIAIVISCPEPLAAVALVDVPLLYLCRYLLIIRRGVIIVSGAGAGGGRLRRFRLGAAMAIIYMAMSTLFFLRIAPLTPWWGALAAWVMILLIVEAIFAFFFPYRCCFNETEGDVQNNPHV |
| IRIS_313-9081 | MELITACALLLIAVHIAIVISCPEPLAAVALVDVPLLYLCRYLLIIRRGVIIVSGAGAGGGRLRRFRLGAAMAIIYMAMSTLFFLRIAPLTPWWGALAAWVMILLIVEAIFAFFFPYRCCFNETEGDVQNNPHV |
| IRIS_313-9083 | MELITACALLLIAVHIAIVISCPEPLAAVALVDVPLLYLCRYLLIIRRGVIIVSGAGAGGGRLRRFRLGAAMAIIYMAMSTLFFLRIAPLTPWWGALAAWVMILLIVEAIFAFFFPYRCCFNETEGDVQNNPHV |
| IRIS_313-9091 | MELITACALLLIAVHIAIVISCPEPLAAVALVDVPLLYLCRYLLIIRRGVIIVSGGGGGGRLRRFRLGAAMAIIYMAMSTLFFLRIAPLTPWWGALAAWVMILLIVEAIFAFFFPYRCCFNETEGDVQNNPHV |
| IRIS_313-9097 | MELITACALLLIAVHIAIVISCPEPLAAVALVDVPLLYLCRYLLIIRRGVIIVSGGGGGGRLRRFRLGAAMAIIYMAMSTLFFLRIAPLTPWWGALAAWVMILLIVEAIFAFFFPYRCCFNETEGDVQNNPHV |
| IRIS_313-9098 | MELITACALLLIAVHIAIVISCPEPLAAVALVDVPLLYLCRYLLIIRRGVIIVSGGGGGGRLRRFRLGAAMAIIYMAMSTLFFLRIAPLTPWWGALAAWVMILLIVEAIFAFFFPYRCCFNETEGDVQNNPHV |
| IRIS_313-9101 | MELITACALLLIAVHIAIVISCPEPLAAVALVDVPLLYLCRYLLIIRRGVIIVSGAGGGGGRLRRFRLGAAMAIIYMAMSTLFFLRIAPLTPWWGALAAWVMILLIVEAIFAFFFPYRCCFNETEGDVQNNPHV |
| IRIS_313-9102 | MELITACALLLIAVHIAIVISCPEPLAAVALVDVPLLYLCRYLLIIRRGVIIVSGAGGAAAAGCAASASGQPWRSSTWPCRPSSSSALLLSRHGGAPSPPGS*SCSSSRPSSPSSSPTAAASTKPKAMSRIIPM |
| IRIS_313-9108 | MELITACALLLIAVHIAIVISCPEPLAAVALVDVPLLYLCRYLLIIRRGVIIVSGAGAGGGRLRRFRLGAAMAIIYMAMSTLFFLRIAPLTPWWGALAAWVMILLIVEAIFAFFFPYRCCFNETEGDVQNNPHV |
| IRIS_313-9111 | MELITACALLLIAVHIAIVISCPEPLAAVALVDVPLLYLCRYLLIIRRGVIIVSGAGAGGGRLRRFRLGAAMAIIYMAMSTLFFLRIAPLTPWWGALAAWVMILLIVEAIFAFFFPYRCCFNETEGDVQNNPHV |
| IRIS_313-9112 | MELITACALLLIAVHIAIVISCPEPLAAVALVDVPLLYLCRYLLIIRRGVIIVSGGGGGGRLRRFRLGAAMAIIYMAMSTLFFLRIAPLTPWWGALAAWVMILLIVEAIFAFFFPYRCCFNETEGDVQNNPHV |
| IRIS_313-9114 | MELITACALLLIAVHIAIVISCPEPLAAVALVDVPLLYLCRYLLIIRRGVIIVSGGGGGGRLRRFRLGAAMAIIYMAMSTLFFLRIAPLTPWWGALAAWVMILLIVEAIFAFFFPYRCCFNETEGDVQNNPHV |
| IRIS_313-9115 | MELITACALLLIAVHIAIVISCPEPLAAVALVDVPLLYLYRYLLIIRRGVIIVSGAGGGGGRLRRFRLGAAMAIIYMAMSTLFFLRIAPLTPWWGALAAWVMILLIVEAIFAFFFPYRCCFNETEGDVQNNPHV |
| IRIS_313-9116 | MELITACALLLIAVHIAIVISCPEPLAAVALVDVPLLYLCRYLLIIRRGVIIVSGAGGGGGRLRRFRLGAAMAIIYMAMSTLFFLRIAPLTPWWGALAAWVMILLIVEAIFAFFFPYRCCFNETEGDVQNNPHV |
| IRIS_313-9117 | MELITACALLLIAVHIAIVISCPEPLAAVALVDVPLLYLCRYLLIIRRGVIIVSGGGGGGRLRRFRLGAAMAIIYMAMSTLFFLRIAPLTPWWGALAAWVMILLIVEAIFAFFFPYRCCFNETEGDVQNNPHV |
| IRIS_313-9119 | MELITACALLLIAVHIAIVISCPEPLAAVALVDVPLLYLYRYLLIIRRGVIIVSGAGGGGGRLRRFRLGAAMAIIYMAMSTLFFLRIAPLTPWWGALAAWVMILLIVEAIFAFFFPYRCCFNETEGDVQNNPHV |
| IRIS_313-9120 | MELITACALLLIAVHIAIVISCPEPLAAVALVDVPLLYLCRYLLIIRRGVIIVSGGGGGGRLRRFRLGAAMAIIYMAMSTLFFLRIAPLTPWWGALAAWVMILLIVEAIFAFFFPYRCCFNETEGDVQNNPHV |
| IRIS_313-9121 | MELITACALLLIAVHIAIVISCPEPLAAVALVDVPLLYLCRYLLIIRRGVIIVSGAGAGGGRLRRFRLGAAMAIIYMAMSTLFFLRIAPLTPWWGALAAWVMILLIVEAIFAFFFPYRCCFNETEGDVQNNPHV |
| IRIS_313-9123 | MELITACALLLIAVHIAIVISCPEPLAAVALVDVPLLYLCRYLLIIRRGVIIVSGAGAGGGRLRRFRLGAAMAIIYMAMSTLFFLRIAPLTPWWGALAAWVMILLIVEAIFAFFFPYRCCFNETEGDVQNNPHV |
| IRIS_313-9129 | MELITACALLLIAVHIAIVISCPEPLAAVALVDVPLLYLYRYLLIIRRGVIIVSGAGGGGGRLRRFRLGAAMAIIYMAMSTLFFLRIAPLTPWWGALAAWVMILLIVEAIFAFFFPYRCCFNETEGDVQNNPHV |
| IRIS_313-9131 | MELITACALLLIAVHIAIVISCPEPLAAVALVDVPLLYLYRYLLIIRRGVIIVSGAGGGGGRLRRFRLGAAMAIIYMAMSTLFFLRIAPLTPWWGALAAWVMILLIVEAIFAFFFPYRCCFNETEGDVQNNPHV |
| IRIS_313-9137 | MELITACALLLIAVHIAIVISCPEPLAAVAFVNVPLLYLCRYLLIIRRGVIIVSGGGGGGGRLRRFRLGAAMAVIYMAMSTLFFLRIAPLTPWWGALAAWVMILLIVEAIFAFFFPYRCCFNETEGDVQNNPHV |
| IRIS_313-9139 | MELITACALLLIAVHIAIVISCPEPLAAVALVDVPLLYLCRYLLIIRRGVIIVSGAGAGGGRLRRFRLGAAMAIIYMAMSTLFFLRIAPLTPWWGALAAWVMILLIVEAIFAFFFPYRCCFNETEGDVQNNPHV |
| IRIS_313-9140 | MELITACALLLIAVHIAIVISCPEPLAAVALVDVPLLYLCRYLLIIRRGVIIVSGAGAGGGRLRRFRLGAAMAIIYMAMSTLFFLRIAPLTPWWGALAAWVMILLIVEAIFAFFFPYRCCFNETEGDVQNNPHV |
| IRIS_313-9148 | MELITACALLLIAVHIAIVISCPEPLAAVALVDVPLLYLCRYLLIIRRGVIIVSGAGAGGGRLRRFRLGAAMAIIYMAMSTLFFLRIAPLTPWWGALAAWVMILLIVEAIFAFFFPYRCCFNETEGDVQNNPHV |
| IRIS_313-9156 | MELITACALLLIAVHIAIVISCPEPLAAVALVDVPLLYLCRYLLIIRRGVIIVSGGGGGGRLRRFRLGAAMAIIYMAMSTLFFLRIAPLTPWWGALAAWVMILLIVEAIFAFFFPYRCCFNETEGDVQNNPHV |
| IRIS_313-9160 | MELITACALLLIAVHIAIVISCPEPLAAVALVDVPLLYLCRYLLIIRRGVIIVSGAGGAAAAGCAASASGQPWRSSTWPCRPSSSSALLLSRHGGAPSPPGS*SCSSSRPSSPSSSPTAAASTKPKAMSRIIPM |
| IRIS_313-9170 | MELITACALLLIAVHIAIVISCPEPLAAVALVDVPLLYLCRYLLIIRRGVIIVSGAGAGGGRLRRFRLGAAMAIIYMAMSTLFFLRIAPLTPWWGALAAWVMILLIVEAIFAFFFPYRCCFNETEGDVQNNPHV |
| IRIS_313-9172 | MELITACALLLIAVHIAIVISCPEPLAAVALVDVPLLYLCRYLLIIRRGVIIVSGGGGGGRLRRFRLGAAMAIIYMAMSTLFFLRIAPLTPWWGALAAWVMILLIVEAIFAFFFPYRCCFNETEGDVQNNPHV |
| IRIS_313-9174 | MELITACALLLIAVHIAIVISCPEPLAAVALVDVPLLYLCRYLLIIRRGVIIVSGAGAGGGRLRRFRLGAAMAIIYMAMSTLFFLRIAPLTPWWGALAAWVMILLIVEAIFAFFFPYRCCFNETEGDVQNNPHV |
| IRIS_313-9176 | MELITACALLLIAVHIAIVISCPEPLAAVALVDVPLLYLYRYLLIIRRGVIIVSGAGGGGGRLRRFRLGAAMAIIYMAMSTLFFLRIAPLTPWWGALAAWVMILLIVEAIFAFFFPYRCCFNETEGDVQNNPHV |
| IRIS_313-9182 | MELITACALLLIAVHIAIVISCPEPLAAVALVDVPLLYLYRYLLIIRRGVIIVSGAGGGGGRLRRFRLGAAMAIIYMAMSTLFFLRIAPLTPWWGALAAWVMILLIVEAIFAFFFPYRCCFNETEGDVQNNPHV |
| IRIS_313-9187 | MELITACALLLIAVHIAIVISCPEPLAAVALVDVPLLYLCRYLLIIRRGVIIVSGAGGGGGGRLRRFRLGAAMAIIYMAMSTLFFLRIAPLTPWWGALAAWVMILLIVEAIFAFFFPYRCCFNETEGDVQNNPHV |
| IRIS_313-9188 | MELITACALLLIAVHIAIVISCPEPLAAVALVDVPLLYLCRYLLIIRRGVIIVSGGGGGGRLRRFRLGAAMAIIYMAMSTLFFLRIAPLTPWWGALAAWVMILLIVEAIFAFFFPYRCCFNETEGDVQNNPHV |
| IRIS_313-9190 | MELITACALLLIAVHIAIVISCPEPLAAVALVDVPLLYLCRYLLIIRRGVIIVSGGGGGGRLRRFRLGAAMAIIYMAMSTLFFLRIAPLTPWWGALAAWVMILLIVEAIFAFFFPYRCCFNETEGDVQNNPHV |
| IRIS_313-9193 | MELITACALLLIAVHIAIVISCPEPLAAVALVDVPLLYLYRYLLIIRRGVIIVSGAGGGGGRLRRFRLGAAMAIIYMAMSTLFFLRIAPLTPWWGALAAWVMILLIVEAIFAFFFPYRCCFNETEGDVQNNPHV |
| IRIS_313-9197 | MELITACALLLIAVHIAIVISCPEPLAAVALVDVPLLYLCRYLLIIRRGVIIVSGGGGGGRLRRFRLGAAMAIIYMAMSTLFFLRIAPLTPWWGALAAWVMILLIVEAIFAFFFPYRCCFNETEGDVQNNPHV |
| IRIS_313-9198 | MELITACALLLIAVHIAIVISCPEPLAAVALVDVPLLYLCRYLLIIRRGVIIVSGGGGGGRLRRFRLGAAMAIIYMAMSTLFFLRIAPLTPWWGALAAWVMILLIVEAIFAFFFPYRCCFNETEGDVQNNPHV |
| IRIS_313-9201 | MELITACALLLIAVHIAIVISCPEPLAAVALVDVPLLYLYRYLLIIRRGVIIVSGAGGGGGRLRRFRLGAAMAIIYMAMSTLFFLRIAPLTPWWGALAAWVMILLIVEAIFAFFFPYRCCFNETEGDVQNNPHV |
| IRIS_313-9204 | MELITACALLLIAVHIAIVISCPEPLAAVALVDVPLLYLCRYLLIIRRGVIIVSGAGGGGGRLRRFRLGAAMAIIYMAMSTLFFLRIAPLTPWWGALAAWVMILLIVEAIFAFFFPYRCCFNETEGDVQNNPHV |
| IRIS_313-9208 | MELITACALLLIAVHIAIVISCPEPLAAVALVDVPLLYLCRYLLIIRRGVIIVSGGGGGGRLRRFRLGAAMAIIYMAMSTLFFLRIAPLTPWWGALAAWVMILLIVEAIFAFFFPYRCCFNETEGDVQNNPHV |
| IRIS_313-9209 | MELITACALLLIAVHIAIVISCPEPLAAVALVDVPLLYLCRYLLIIRRGVIIVSGAGGGGRLRRFRLGAAMAIIYMAMSTLFFLRIAPLTPWWGALAAWVMILLIVEAIFAFFFPYRCCFNETEGDVQNNPHV |
| IRIS_313-9210 | MELITACALLLIAVHIAIVISCPEPLAAVALVDVPLLYLCRYLLIIRRGVIIVSGGGGGGRLRRFRLGAAMAIIYMAMSTLFFLRIAPLTPWWGALAAWVMILLIVEAIFAFFFPYRCCFNETEGDVQNNPHV |
| IRIS_313-9218 | MELITACALLLIAVHIAIVISCPEPLAAVALVDVPLLYLCRYLLIIRRGVIIVSGGGGGGRLRRFRLGAAMAIIYMAMSTLFFLRIAPLTPWWGALAAWVMILLIVEAIFAFFFPYRCCFNETEGDVQNNPHV |
| IRIS_313-9227 | MELITACALLLIAVHIAIVISCPEPLAAVALVDVPLLYLCRYLLIIRRGVIIVSGAGAGGGRLRRFRLGAAMAIIYMAMSTLFFLRIAPLTPWWGALAAWVMILLIVEAIFAFFFPYRCCFNETEGDVQNNPHV |
| IRIS_313-9228 | MELITACALLLIAVHIAIVISCPEPLAAVALVDVPLLYLCRYLLIIRRGVIIVSGAGAGGGRLRRFRLGAAMAIIYMAMSTLFFLRIAPLTPWWGALAAWVMILLIVEAIFAFFFPYRCCFNETEGDVQNNPHV |
| IRIS_313-9239 | MELITACALLLIAVHIAIVISCPEPLAAVALVDVPLLYLYRYLLIIRRGVIIVSGAGGGGGRLRRFRLGAAMAIIYMAMSTLFFLRIAPLTPWWGALAAWVMILLIVEAIFAFFFPYRCCFNETEGDVQNNPHV |
| IRIS_313-9242 | MELITACALLLIAVHIAIVISCPEPLAAVALVDVPLLYLYRYLLIIRRGVIIVSGAGGGGGRLRRFRLGAAMAIIYMAMSTLFFLRIAPLTPWWGALAAWVMILLIVEAIFAFFFPYRCCFNETEGDVQNNPHV |
| IRIS_313-9243 | MELITACALLLIAVHIAIVISCPEPLAAVALVDVPLLYLCRYLLIIRRGVIIVSGAGGGGGRLRRFRLGAAMAIIYMAMSTLFFLRIAPLTPWWGALAAWVMILLIVEAIFAFFFPYRCCFNETEGDVQNNPHV |
| IRIS_313-9249 | MELITACALLLIAVHIAIVISCPEPLAAVALVDVPLLYLCRYLLIIRRGVIIVSGAGAGGGRLRRFRLGAAMAIIYMAMSTLFFLRIAPLTPWWGALAAWVMILLIVEAIFAFFFPYRCCFNETEGDVQNNPHV |
| IRIS_313-9251 | MELITACALLLIAVHIAIVISCPEPLAAVALVDVPLLYLCRYLLIIRRGVIIVSGGGGGGRLRRFRLGAAMAIIYMAMSTLFFLRIAPLTPWWGALAAWVMILLIVEAIFAFFFPYRCCFNETEGDVQNNPHV |
| IRIS_313-9253 | MELITACALLLIAVHIAIVISCPEPLAAVALVDVPLLYLCRYLLIIRRGVIIVSGAGAGGGRLRRFRLGAAMAIIYMAMSTLFFLRIAPLTPWWGALAAWVMILLIVEAIFAFFFPYRCCFNETEGDVQNNPHV |
| IRIS_313-9256 | MELITACALLLIAVHIAIVISCPEPLAAVALVDVPLLYLCRYLLIIRRGVIIVSGAGAGGGRLRRFRLGAAMAIIYMAMSTLFFLRIAPLTPWWGALAAWVMILLIVEAIFAFFFPYRCCFNETEGDVQNNPHV |
| IRIS_313-9258 | MELITACALLLIAVHIAIVISCPEPLAAVALVDVPLLYLCRYLLIIRRGVIIVSGAGAGGGRLRRFRLGAAMAIIYMAMSTLFFLRIAPLTPWWGALAAWVMILLIVEAIFAFFFPYRCCFNETEGDVQNNPHV |
| IRIS_313-9259 | MELITACALLLIAVHIAIVISCPEPLAAVALVDVPLLYLCRYLLIIRRGVIIVSGGGGGGRLRRFRLGAAMAIIYMAMSTLFFLRIAPLTPWWGALAAWVMILLIVEAIFAFFFPYRCCFNETEGDVQNNPHV |
| IRIS_313-9262 | MELITACALLLIAVHIAIVISCPEPLAAVALVDVPLLYLCRYLLIIRRGVIIVSGGGGGGRLRRFRLGAAMAIIYMAMSTLFFLRIAPLTPWWGALAAWVMILLIVEAIFAFFFPYRCCFNETEGDVQNNPHV |
| IRIS_313-9267 | MELITACALLLIAVHIAIVISCPEPLAAVALVDVPLLYLCRYLLIIRRGVIIVSGAGAGGGRLRRFRLGAAMAIIYMAMSTLFFLRIAPLTPWWGALAAWVMILLIVEAIFAFFFPYRCCFNETEGDVQNNPHV |
| IRIS_313-9271 | MELITACALLLIAVHIAIVISCPEPLAAVALVDVPLLYLCRYLLIIRRGVIIVSGAGAGGGRLRRFRLGAAMAIIYMAMSTLFFLRIAPLTPWWGALAAWVMILLIVEAIFAFFFPYRCCFNETEGDVQNNPHV |
| IRIS_313-9273 | MELITACALLLIAVHIAIVISCPEPLAAVALVDVPLLYLCRYLLIIRRGVIIVSGGGGGGRLRRFRLGAAMAIIYMAMSTLFFLRIAPLTPWWGALAAWVMILLIVEAIFAFFFPYRCCFNETEGDVQNNPHV |
| IRIS_313-9281 | MELITACALLLIAVHIAIVISCPEPLAAVALVDVPLLYLYRYLLIIRRGVIIVSGAGGGGGRLRRFRLGAAMAIIYMAMSTLFFLRIAPLTPWWGALAAWVMILLIVEAIFAFFFPYRCCFNETEGDVQNNPHV |
| IRIS_313-9283 | MELITACALLLIAVHIAIVISCPEPLAAVALVDVPLLYLCRYLLIIRRGVIIVSGAGAGGGRLRRFRLGAAMAIIYMAMSTLFFLRIAPLTPWWGALAAWVMILLIVEAIFAFFFPYRCCFNETEGDVQNNPHV |
| IRIS_313-9285 | MELITACALLLIAVHIAIVISCPEPLAAVALVDVPLLYLCRYLLIIRRGVIIVSGGGGGGRLRRFRLGAAMAIIYMAMSTLFFLRIAPLTPWWGALAAWVMILLIVEAIFAFFFPYRCCFNETEGDVQNNPHV |
| IRIS_313-9286 | MELITACALLLIAVHIAIVISCPEPLAAVALVDVPLLYLCRYLLIIRRGVIIVSGAGAGGGRLRRFRLGAAMAIIYMAMSTLFFLRIAPLTPWWGALAAWVMILLIVEAIFAFFFPYRCCFNETEGDVQNNPHV |
| IRIS_313-9287 | MELITACALLLIAVHIAIVISCPEPLAAVALVDVPLLYLCRYLLIIRRGVIIVSGGGGGGRLRRFRLGAAMAIIYMAMSTLFFLRIAPLTPWWGALAAWVMILLIVEAIFAFFFPYRCCFNETEGDVQNNPHV |
| IRIS_313-9288 | MELITACALLLIAVHIAIVISCPEPLAAVALVDVPLLYLCRYLLIIRRGVIIVSGGGGGGRLRRFRLGAAMAIIYMAMSTLFFLRIAPLTPWWGALAAWVMILLIVEAIFAFFFPYRCCFNETEGDVQNNPHV |
| IRIS_313-9294 | MELITACALLLIAVHIAIVISCPEPLAAVALVDVPLLYLCRYLLIIRRGVIIVSGAGAGGGRLRRFRLGAAMAIIYMAMSTLFFLRIAPLTPWWGALAAWVMILLIVEAIFAFFFPYRCCFNETEGDVQNNPHV |
| IRIS_313-9297 | MELITACALLLIAVHIAIVISCPEPLAAVALVDVPLLYLYRYLLIIRRGVIIVSGAGGGGGRLRRFRLGAAMAIIYMAMSTLFFLRIAPLTPWWGALAAWVMILLIVEAIFAFFFPYRCCFNETEGDVQNNPHV |
| IRIS_313-9301 | MELITACALLLIAVHIAIVISCPEPLAAVALVDVPLLYLYRYLLIIRRGVIIVSGAGGGGGRLRRFRLGAAMAIIYMAMSTLFFLRIAPLTPWWGALAAWVMILLIVEAIFAFFFPYRCCFNETEGDVQNNPHV |
| IRIS_313-9302 | MELITACALLLIAVHIAIVISCPEPLAAVALVDVPLLYLCRYLLIIRRGVIIVSGGGGGGRLRRFRLGAAMAIIYMAMSTLFFLRIAPLTPWWGALAAWVMILLIVEAIFAFFFPYRCCFNETEGDVQNNPHV |
| IRIS_313-9310 | MELITACALLLIAVHIAIVISCPEPLAAVALVDVPLLYLCRYLLIIRRGVIIVSGGGGGGRLRRFRLGAAMAIIYMAMSTLFFLRIAPLTPWWGALAAWVMILLIVEAIFAFFFPYRCCFNETEGDVQNNPHV |
| IRIS_313-9313 | MELITACALLLIAVHIAIVISCPEPLAAVALVDVPLLYLCRYLLIIRRGVIIVSGAGAGGGRLRRFRLGAAMAIIYMAMSTLFFLRIAPLTPWWGALAAWVMILLIVEAIFAFFFPYRCCFNETEGDVQNNPHV |
| IRIS_313-9314 | MELITACALLLIAVHIAIVISCPEPLAAVALVDVPLLYLCRYLLIIRRGVIIVSGGGGGGRLRRFRLGAAMAIIYMAMSTLFFLRIAPLTPWWGALAAWVMILLIVEAIFAFFFPYRCCFNETEGDVQNNPHV |
| IRIS_313-9317 | MELITACALLLIAVHIAIVISCPEPLAAVALVDVPLLYLYRYLLIIRRGVIIVSGAGGGGGRLRRFRLGAAMAIIYMAMSTLFFLRIAPLTPWWGALAAWVMILLIVEAIFAFFFPYRCCFNETEGDVQNNPHV |
| IRIS_313-9320 | MELITACALLLIAVHIAIVISCPEPLAAVALVDVPLLYLCRYLLIIRRGVIIVSGAGAGGGRLRRFRLGAAMAIIYMAMSTLFFLRIAPLTPWWGALAAWVMILLIVEAIFAFFFPYRCCFNETEGDVQNNPHV |
| IRIS_313-9324 | MELITACALLLIAVHIAIVISCPEPLAAVALVDVPLLYLCRYLLIIRRGVIIVSGAGAGGGRLRRFRLGAAMAIIYMAMSTLFFLRIAPLTPWWGALAAWVMILLIVEAIFAFFFPYRCCFNETEGDVQNNPHV |
| IRIS_313-9325 | MELITACALLLIAVHIAIVISCPEPLAAVALVDVPLLYLCRYLLIIRRGVIIVSGGGGGGRLRRFRLGAAMAIIYMAMSTLFFLRIAPLTPWWGALAAWVMILLIVEAIFAFFFPYRCCFNETEGDVQNNPHV |
| IRIS_313-9329 | MELITACALLLIAVHIAIVISCPEPLAAVALVDVPLLYLCRYLLIIRRGVIIVSGGGGGGRLRRFRLGAAMAIIYMAMSTLFFLRIAPLTPWWGALAAWVMILLIVEAIFAFFFPYRCCFNETEGDVQNNPHV |
| IRIS_313-9342 | MELITACALLLIAVHIAIVISCPEPLAAVALVDVPLLYLYRYLLIIRRGVIIVSGAGGGGGRLRRFRLGAAMAIIYMAMSTLFFLRIAPLTPWWGALAAWVMILLIVEAIFAFFFPYRCCFNETEGDVQNNPHV |
| IRIS_313-9346 | MELITACALLLIAVHIAIVISCPEPLAAVALVDVPLLYLYRYLLIIRRGVIIVSGAGGGGGRLRRFRLGAAMAIIYMAMSTLFFLRIAPLTPWWGALAAWVMILLIVEAIFAFFFPYRCCFNETEGDVQNNPHV |
| IRIS_313-9347 | MELITACALLLIAVHIAIVISCPEPLAAVALVDVPLLYLCRYLLIIRRGVIIVSGGGGGGRLRRFRLGAAMAIIYMAMSTLFFLRIAPLTPWWGALAAWVMILLIVEAIFAFFFPYRCCFNETEGDVQNNPHV |
| IRIS_313-9348 | MELITACALLLIAVHIAIVISCPEPLAAVALVDVPLLYLCRYLLIIRRGVIIVSGAGGAAAAGCAASASGQPWRSSTWPCRPSSSSALLLSRHGGAPSPPGS*SCSSSRPSSPSSSPTAAASTKPKAMSRIIPM |
| IRIS_313-9351 | MELITACALLLIAVHIAIVISCPEPLAAVALVDVPLLYLCRYLLIIRRGVIIVSGGGGGGRLRRFRLGAAMAIIYMAMSTLFFLRIAPLTPWWGALAAWVMILLIVEAIFAFFFPYRCCFNETEGDVQNNPHV |
| IRIS_313-9357 | MELITACALLLIAVHIAIVISCPEPLAAVALVDVPLLYLCRYLLIIRRGVIIVSGAGAGGGRLRRFRLGAAMAIIYMVMSTLFFLRIAPLTPWWGALAAWVMILLIVEAIFAFFFPYRCCFNETEGDVQNNPHV |
| IRIS_313-9360 | MELITACALLLIAVHIAIVISCPEPLAAVALVDVPLLYLCRYLLIIRRGVIIVSGAGAGGGRLRRFRLGAAMAIIYMAMSTLFFLRIAPLTPWWGALAAWVMILLIVEAIFAFFFPYRCCFNETEGDVQNNPHV |
| IRIS_313-9363 | MELITACALLLIAVHIAIVISCPEPLAAVALVDVPLLYLCRYLLIIRRGVIIVSGAGAGGGRLRRFRLGAAMAIIYMAMSTLFFLRIAPLTPWWGALAAWVMILLIVEAIFAFFFPYRCCFNETEGDVQNNPHV |
| IRIS_313-9366 | MELITACALLLIAVHIAIVISCPEPLAAVALVDVPLLYLCRYLLIIRRGVIIVSGAGAGGGRLRRFRLGAAMAIIYMAMSTLFFLRIAPLTPWWGALAAWVMILLIVEAIFAFFFPYRCCFNETEGDVQNNPHV |
| IRIS_313-9368 | MELITACALLLIAVHIAIVISCPEPLAAVAFVNVPLLYLCRYLLIIRRGVIIVSGGGGGGGRLRRFRLGAAMAVIYMAMSTLFFLRIAPLTPWWGALAAWVMILLIVEAIFAFFFPYRCCFNETEGDVQNNPHV |
| IRIS_313-9372 | MELITACALLLIAVHIAIVISCPEPLAAVALVDVPLLYLCRYLLIIRRGVIIVSGGGGGGRLRRFRLGAAMAIIYMAMSTLFFLRIAPLTPWWGALAAWVMILLIVEAIFAFFFPYRCCFNETEGDVQNNPHV |
| IRIS_313-9375 | MELITACALLLIAVHIAIVISCPEPLAAVALVDVPLLYLCRYLLIIRRGVIIVSGAGAGGGRLRRFRLGAAMAIIYMAMSTLFFLRIAPLTPWWGALAAWVMILLIVEAIFAFFFPYRCCFNETEGDVQNNPHV |
| IRIS_313-9379 | MELITACALLLIAVHIAIVISCPEPLAAVALVDVPLLYLCRYLLIIRRGVIIVSGAGAGGGRLRRFRLGAAMAIIYMAMSTLFFLRIAPLTPWWGALAAWVMILLIVEAIFAFFFPYRCCFNETEGDVQNNPHV |
| IRIS_313-9384 | MELITACALLLIAVHIAIVISCPEPLAAVALVDVPLLYLCRYLLIIRRGVIIVSGGGGGGRLRRFRLGAAMAIIYMAMSTLFFLRIAPLTPWWGALAAWVMILLIVEAIFAFFFPYRCCFNETEGDVQNNPHV |

| | |
|---|---|
| IRIS_313-9388 | MELITACALLLIAVHIAIVISCPEPLAAVALVDVPLLYLCRYLLIIRRGVIIVSGAGGAAAAGCAASASGQPWRSSTWPCRPSSSSALLLSRHGGAPSPPGS*SCSSSRPSSPSSSPTAAASTKPKAMSRIIPM |
| IRIS_313-9389 | MELITACALLLIAVHIAIVISCPEPLAAVALVDVPLLYLYRYLLIIRRGVIIVSGAGGGGGRLRRFRLGAAMAIIYMAMSTLFFLRIAPLTPWWGALAAWVMILLIVEAIFAFFFPYRCCFNETEGDVQNNPHV |
| IRIS_313-9391 | MELITACALLLIAVHIAIVISCPEPLAAVALVDVPLLYLCRYLLIIRRGVIIVSGGGGGGRLRRFRLGAAMAIIYMAMSTLFFLRIAPLTPWWGALAAWVMILLIVEAIFAFFFPYRCCFNETEGDVQNNPHV |
| IRIS_313-9392 | MELITACALLLIAVHIAIVISCPEPLAAVALVDVPLLYLYRYLLIIRRGVIIVSGGGGGGRLRRFRLGAAMAIIYMAMSTLFFLRIAPLTPWWGALAAWVMILLIVEAIFAFFFPYRCCFNETEGDVQNNPHV |
| IRIS_313-9397 | MELITACALLLIAVHIAIVISCPEPLAAVALVDVPLLYLCRYLLIIRRGVIIVSGGGGGGRLRRFRLGAAMAIIYMAMSTLFFLRIAPLTPWWGALAAWVMILLIVEAIFAFFFPYRCCFNETEGDVQNNPHV |
| IRIS_313-9400 | MELITACALLLIAVHIAIVISCPEPLAAVALVDVPLLYLCRYLLIIRRGVIIVSGAGAGGGRLRRFRLGAAMAIIYMAMSTLFFLRIAPLTPWWGALAAWVMILLIVEAIFAFFFPYRCCFNETEGDVQNNPHV |
| IRIS_313-9402 | MELITACALLLIAVHIAIVISCPEPLAAVALVDVPLLYLCRYLLIIRRGVIIVSGAGAGGGRLRRFRLGAAMAIIYMAMSTLFFLRIAPLTPWWGALAAWVMILLIVEAIFAFFFPYRCCFNETEGDVQNNPHV |
| IRIS_313-9403 | MELITACALLLIAVHIAIVISCPEPLAAVALVDVPLLYLCRYLLIIRRGVIIVSGAGAGGGRLRRFRLGAAMAIIYMAMSTLFFLRIAPLTPWWGALAAWVMILLIVEAIFAFFFPYRCCFNETEGDVQNNPHV |
| IRIS_313-9405 | MELITACALLLIAVHIAIVISCPEPLAAVALVDVPLLYLCRYLLIIRRGVIIVSGAGAGGGRLRRFRLGAAMAIIYMAMSTLFFLRIAPLTPWWGALAAWVMILLIVEAIFAFFFPYRCCFNETEGDVQNNPHV |
| IRIS_313-9406 | MELITACALLLIAVHIAIVISCPEPLAAVALVDVPLLYLCRYLLIIRRGVIIVSGGGGGGRLRRFRLGAAMAIIYMAMSTLFFLRIAPLTPWWGALAAWVMILLIVEAIFAFFFPYRCCFNETEGDVQNNPHV |
| IRIS_313-9409 | MELITACALLLIAVHIAIVISCPEPLAAVALVDVPLLYLCRYLLIIRRGVIIVSGGGGGGRLRRFRLGAAMAIIYMAMSTLFFLRIAPLTPWWGALAAWVMILLIVEAIFAFFFPYRCCFNETEGDVQNNPHV |
| IRIS_313-9410 | MELITACALLLIAVHIAIVISCPEPLAAVALVDVPLLYLCRYLLIIRRGVIIVSGAGAGGGRLRRFRLGAAMAIIYMAMSTLFFLRIAPLTPWWGALAAWVMILLIVEAIFAFFFPYRCCFNETEGDVQNNPHV |
| IRIS_313-9415 | MELITACALLLIAVHIAIVISCPEPLAAVALVDVPLLYLCRYLLIIRRGVIIVSGGGGGGRLRRFRLGAAMAIIYMAMSTLFFLRIAPLTPWWGALAAWVMILLIVEAIFAFFFPYRCCFNETEGDVQNNPHV |
| IRIS_313-9422 | MELITACALLLIAVHIAIVISCPEPLAAVALVDVPLLYLCRYLLIIRRGVIIVSGAGAGGGRLRRFRLGAAMAIIYMAMSTLFFLRIAPLTPWWGALAAWVMILLIVEAIFAFFFPYRCCFNETEGDVQNNPHV |
| IRIS_313-9423 | MELITACALLLIAVHIAIVISCPEPLAAVALVDVPLLYLCRYLLIIRRGVIIVSGAGGGGGRLRRFRLGAAMAIIYMAMSTLFFLRIAPLTPWWGALAAWVMILLIVEAIFAFFFPYRCCFNETEGDVQNNPHV |
| IRIS_313-9424 | MELITACALLLIAVHIAIVISCPEPLAAVALVDVPLLYLCRYLLIIRRGVIIVSGGGGGGRLRRFRLGAAMAIIYMAMSTLFFLRIAPLTPWWGALAAWVMILLIVEAIFAFFFPYRCCFNETEGDVQNNPHV |
| IRIS_313-9427 | MELITACALLLIAVHIAIVISCPEPLAAVALVDVPLLYLYRYLLIIRRGVIIVSGAGGGGGRLRRFRLGAAMAIIYMAMSTLFFLRIAPLTPWWGALAAWVMILLIVEAIFAFFFPYRCCFNETEGDVQNNPHV |
| IRIS_313-9429 | MELITACALLLIAVHIAIVISCPEPLAAVALVDVPLLYLYRYLLIIRRGVIIVSGAGAGGGRLRRFRLGAAMAIIYMAMSTLFFLRIAPLTPWWGALAAWVMILLIVEAIFAFFFPYRCCFNETEGDVQNNPHV |
| IRIS_313-9433 | MELITACALLLIAVHIAIVISCPEPLAAVALVDVPLLYLCRYLLIIRRGVIIVSGGGGGGRLRRFRLGAAMAIIYMAMSTLFFLRIAPLTPWWGALAAWVMILLIVEAIFAFFFPYRCCFNETEGDVQNNPHV |
| IRIS_313-9438 | MELITACALLLIAVHIAIVISCPEPLAAVALVDVPLLYLCRYLLIIRRGVIIVSGAGAGGGRLRRFRLGAAMAIIYMAMSTLFFLRIAPLTPWWGALAAWVMILLIVEAIFAFFFPYRCCFNETEGDVQNNPHV |
| IRIS_313-9445 | MELITACALLLIAVHIAIVISCPEPLAAVALVDVPLLYLCRYLLIIRRGVIIVSGAGAGGGRLRRFRLGAAMAIIYMAMSTLFFLRIAPLTPWWGALAAWVMILLIVEAIFAFFFPYRCCFNETEGDVQNNPHV |
| IRIS_313-9449 | MELITACALLLIAVHIAIVISCPEPLAAVAFVNVPLLYLCRYLLIIRRGVIIVSGGGAGGGRLRRFRLGAAMAVIYMAMSTLFFLRIAPLTPWWGALAAWVMILLIVEAIFAFFFPYRCCFNETEGDVQNNPHV |
| IRIS_313-9451 | MELITACALLLIAVHIAIVISCPEPLAAVALVDVPLLYLYRYLLIIRRGVIIVSGAGGGGGRLRRFRLGAAMAIIYMAMSTLFFLRIAPLTPWWGALAAWVMILLIVEAIFAFFFPYRCCFNETEGDVQNNPHV |
| IRIS_313-9452 | MELITACALLLIAVHIAIVISCPEPLAAVALVDVPLLYLYRYLLIIRRGVIIVSGAGGGGGRLRRFRLGAAMAIIYMAMSTLFFLRIAPLTPWWGALAAWVMILLIVEAIFAFFFPYRCCFNETEGDVQNNPHV |
| IRIS_313-9461 | MELITACALLLIAVHIAIVISCPEPLAAVALVDVPLLYLCRYLLIIRRGVIIVSGAGAGGGRLRRFRLGAAMAIIYMAMSTLFFLRIAPLTPWWGALAAWVMILLIVEAIFAFFFPYRCCFNETEGDVQNNPHV |
| IRIS_313-9463 | MELITACALLLIAVHIAIVISCPEPLAAVALVDVPLLYLCRYLLIIRRGVIIVSGAGAGGGRLRRFRLGAAMAIIYMAMSTLFFLRIAPLTPWWGALAAWVMILLIVEAIFAFFFPYRCCFNETEGDVQNNPHV |
| IRIS_313-9464 | MELITACALLLIAVHIAIVISCPEPLAAVALVDVPLLYLCRYLLIIRRGVIIVSGGGGGGRLRRFRLGAAMAIIYMAMSTLFFLRIAPLTPWWGALAAWVMILLIVEAIFAFFFPYRCCFNETEGDVQNNPHV |
| IRIS_313-9468 | MELITACALLLIAVHIAIVISCPEPLAAVALVDVPLLYLCRYLLIIRRGVIIVSGAGAGGGRLRRFRLGAAMAIIYMAMSTLFFLRIAPLTPWWGALAAWVMILLIVEAIFAFFFPYRCCFNETEGDVQNNPHV |
| IRIS_313-9469 | MELITACALLLIAVHIAIVISCPEPLAAVALVDVPLLYLYRYLLIIRRGVIIVSGAGGGGGRLRRFRLGAAMAIIYMAMSTLFFLRIAPLTPWWGALAAWVMILLIVEAIFAFFFPYRCCFNETEGDVQNNPHV |
| IRIS_313-9470 | MELITACALLLIAVHIAIVISCPEPLAAVALVDVPLLYLCRYLLIIRRGVIIVSGAGGGGGRLRRFRLGAAMAIIYMAMSTLFFLRIAPLTPWWGALAAWVMILLIVEAIFAFFFPYRCCFNETEGDVQNNPHV |
| IRIS_313-9472 | MELITACALLLIAVHIAIVISCPEPLAAVALVDVPLLYLCRYLLIIRRGVIIVSGGGGGGRLRRFRLGAAMAIIYMAMSTLFFLRIAPLTPWWGALAAWVMILLIVEAIFAFFFPYRCCFNETEGDVQNNPHV |
| IRIS_313-9482 | MELITACALLLIAVHIAIVISCPEPLAAVALVDVPLLYLCRYLLIIRRGVIIVSGGGGGGRLRRFRLGAAMAIIYMAMSTLFFLRIAPLTPWWGALAAWVMILLIVEAIFAFFFPYRCCFNETEGDVQNNPHV |
| IRIS_313-9484 | MELITACALLLIAVHIAIVISCPEPLAAVALVDVPLLYLCRYLLIIRRGVIIVSGAGGGGGGRLRRFRLGAAMAIIYMAMSTLFFLRIAPLTPWWGALAAWVMILLIVEAIFAFFFPYRCCFNETEGDVQNNPHV |
| IRIS_313-9491 | MELITACALLLIAVHIAIVISCPEPLAAVALVDVPLLYLYRYLLIIRRGVIIVSGAGGGGGRLRRFRLGAAMAIIYMAMSTLFFLRIAPLTPWWGALAAWVMILLIVEAIFAFFFPYRCCFNETEGDVQNNPHV |
| IRIS_313-9492 | MELITACALLLIAVHIAIVISCPEPLAAVALVDVPLLYLCRYLLIIRRGVIIVSGAGAGGGRLRRFRLGAAMAIIYMAMSTLFFLRIAPLTPWWGALAAWVMILLIVEAIFAFFFPYRCCFNETEGDVQNNPHV |
| IRIS_313-9503 | MELITACALLLIAVHIAIVISCPEPLAAVALVDVPLLYLCRYLLIIRRGVIIVSGGGAGGGRLRRFRLGAAMAIIYMAMSTLFFLRIAPLTPWWGALAAWVMILLIVEAIFAFFFPYRCCFNETEGDVQNNPHV |
| IRIS_313-9505 | MELITACALLLIAVHIAIVISCPEPLAAVALVDVPLLYLYRYLLIIRRGVIIVSGAGGGGGRLRRFRLGAAMAIIYMAMSTLFFLRIAPLTPWWGALAAWVMILLIVEAIFAFFFPYRCCFNETEGDVQNNPHV |
| IRIS_313-9506 | MELITACALLLIAVHIAIVISCPEPLAAVALVDVPLLYLCRYLLIIRRGVIIVSGGGGGGRLRRFRLGAAMAIIYMAMSTLFFLRIAPLTPWWGALAAWVMILLIVEAIFAFFFPYRCCFNETEGDVQNNPHV |
| IRIS_313-9516 | MELITACALLLIAVHIAIVISCPEPLAAVALVDVPLLYLCRYLLIIRRGVIIVSGAGAGGGRLRRFRLGAAMAIIYMAMSTLFFLRIAPLTPWWGALAAWVMILLIVEAIFAFFFPYRCCFNETEGDVQNNPHV |
| IRIS_313-9519 | MELITACALLLIAVHIAIVISCPEPLAAVALVDVPLLYLCRYLLIIRRGVIIVSGAGAGGGRLRRFRLGAAMAIIYMAMSTLFFLRIAPLTPWWGALAAWVMILLIVEAIFAFFFPYRCCFNETEGDVQNNPHV |
| IRIS_313-9522 | MELITACALLLIAVHIAIVISCPEPLAAVALVDVPLLYLCRYLLIIRRGVIIVSGGGGGGRLRRFRLGAAMAIIYMAMSTLFFLRIAPLTPWWGALAAWVMILLIVEAIFAFFFPYRCCFNETEGDVQNNPHV |
| IRIS_313-9523 | MELITACALLLIAVHIAIVISCPEPLAAVALVDVPLLYLYRYLLIIRRGVIIVSGAGGGGGRLRRFRLGAAMAIIYMAMSTLFFLRIAPLTPWWGALAAWVMILLIVEAIFAFFFPYRCCFNETEGDVQNNPHV |
| IRIS_313-9529 | MELITACALLLIAVHIAIVISCPEPLAAVALVDVPLLYLCRYLLIIRRGVIIVSGAGAGGGRLRRFRLGAAMAIIYMAMSTLFFLRIAPLTPWWGALAAWVMILLIVEAIFAFFFPYRCCFNETEGDVQNNPHV |
| IRIS_313-9533 | MELITACALLLIAVHIAIVISCPEPLAAVALVDVPLLYLCRYLLIIRRGVIIVSGGGGGGRLRRFRLGAAMAIIYMAMSTLFFLRIAPLTPWWGALAAWVMILLIVEAIFAFFFPYRCCFNETEGDVQNNPHV |
| IRIS_313-9539 | MELITACALLLIAVHIAIVISCPEPLAAVALVDVPLLYLCRYLLIIRRGVIIVSGAGAGGGRLRRFRLGAAMAIIYMAMSTLFFLRIAPLTPWWGALAAWVMILLIVEAIFAFFFPYRCCFNETEGDVQNNPHV |
| IRIS_313-9547 | MELITACALLLIAVHIAIVISCPEPLAAVALVDVPLLYLCRYLLIIRRGVIIVSGAGGGGGGRLRRFRLGAAMAIIYMAMSTLFFLRIAPLTPWWGALAAWVMILLIVEAIFAFFFPYRCCFNETEGDVQNNPHV |
| IRIS_313-9550 | MELITACALLLIAVHIAIVISCPEPLAAVALVDVPLLYLYRYLLIIRRGVIIVSGAGGGGGRLRRFRLGAAMAIIYMAMSTLFFLRIAPLTPWWGALAAWVMILLIVEAIFAFFFPYRCCFNETEGDVQNNPHV |
| IRIS_313-9551 | MELITACALLLIAVHIAIVISCPEPLAAVALVDVPLLYLCRYLLIIRRGVIIVSGAGAGGGRLRRFRLGAAMAIIYMAMSTLFFLRIAPLTPWWGALAAWVMILLIVEAIFAFFFPYRCCFNETEGDVQNNPHV |
| IRIS_313-9555 | MELITACALLLIAVHIAIVISCPEPLAAVALVDVPLLYLYRYLLIIRRGVIIVSGAGGGGGRLRRFRLGAAMAIIYMAMSTLFFLRIAPLTPWWGALAAWVMILLIVEAIFAFFFPYRCCFNETEGDVQNNPHV |
| IRIS_313-9557 | MELITACALLLIAVHIAIVISCPEPLAAVALVDVPLLYLCRYLLIIRRGVIIVSGAGGAAAAGCAASASGQPWRSSTWPCRPSSSSALLLSRHGGAPSPPGS*SCSSSRPSSPSSSPTAAASTKPKAMSRIIPM |
| IRIS_313-9558 | MELITACALLLIAVHIAIVISCPEPLAAVALVDVPLLYLCRYLLIIRRGVIIVSGAGGGGGGRLRRFRLGAAMAIIYMAMSTLFFLRIAPLTPWWGALAAWVMILLIVEAIFAFFFPYRCCFNETEGDVQNNPHV |
| IRIS_313-9560 | MELITACALLLIAVHIAIVISCPEPLAAVALVDVPLLYLCRYLLIIRRGVIIVSGGGGGGRLRRFRLGAAMAIIYMAMSTLFFLRIAPLTPWWGALAAWVMILLIVEAIFAFFFPYRCCFNETEGDVQNNPHV |
| IRIS_313-9566 | MELITACALLLIAVHIAIVISCPEPLAAVALVDVPLLYLCRYLLIIRRGVIIVSGGGGGGRLRRFRLGAAMAIIYMAMSTLFFLRIAPLTPWWGALAAWVMILLIVEAIFAFFFPYRCCFNETEGDVQNNPHV |
| IRIS_313-9568 | MELITACALLLIAVHIAIVISCPEPLAAVALVDVPLLYLYRYLLIIRRGVIIVSGAGGGGGRLRRFRLGAAMAIIYMAMSTLFFLRIAPLTPWWGALAAWVMILLIVEAIFAFFFPYRCCFNETEGDVQNNPHV |
| IRIS_313-9570 | MELITACALLLIAVHIAIVISCPEPLAAVALVDVPLLYLCRYLLIIRRGVIIVSGAGAGGGRLRRFRLGAAMAIIYMAMSTLFFLRIAPLTPWWGALAAWVMILLIVEAIFAFFFPYRCCFNETEGDVQNNPHV |
| IRIS_313-9572 | MELITACALLLIAVHIAIVISCPEPLAAVALVDVPLLYLCRYLLIIRRGVIIVSGAGAGGGRLRRFRLGAAMAIIYMAMSTLFFLRIAPLTPWWGALAAWVMILLIVEAIFAFFFPYRCCFNETEGDVQNNPHV |
| IRIS_313-9574 | MELITACALLLIAVHIAIVISCPEPLAAVALVDVPLLYLCRYLLIIRRGVIIVSGAGGGGGGRLRRFRLGAAMAIIYMAMSTLFFLRIAPLTPWWGALAAWVMILLIVEAIFAFFFPYRCCFNETEGDVQNNPHV |
| IRIS_313-9575 | MELITACALLLIAVHIAIVISCPEPLAAVALVDVPLLYLCRYLLIIRRGVIIVSGGGGGGRLRRFRLGAAMAIIYMAMSTLFFLRIAPLTPWWGALAAWVMILLIVEAIFAFFFPYRCCFNETEGDVQNNPHV |
| IRIS_313-9582 | MELITACALLLIAVHIAIVISCPEPLAAVALVDVPLLYLCRYLLIIRRGVIIVSGGGGGGRLRRFRLGAAMAIIYMAMSTLFFLRIAPLTPWWGALAAWVMILLIVEAIFAFFFPYRCCFNETEGDVQNNPHV |
| IRIS_313-9590 | MELITACALLLIAVHIAIVISCPEPLAAVALVDVPLLYLCRYLLIIRRGVIIVSGGGGGGRLRRFRLGAAMAIIYMAMSTLFFLRIAPLTPWWGALAAWVMILLIVEAIFAFFFPYRCCFNETEGDVQNNPHV |
| IRIS_313-9593 | MELITACALLLIAVHIAIVISCPEPLAAVALVDVPLLYLCRYLLIIRRGVIIVSGAGAGGGRLRRFRLGAAMAIIYMAMSTLFFLRIAPLTPWWGALAAWVMILLIVEAIFAFFFPYRCCFNETEGDVQNNPHV |
| IRIS_313-9594 | MELITACALLLIAVHIAIVISCPEPLAAVALVDVPLLYLCRYLLIIRRGVIIVSGAGAGGGRLRRFRLGAAMAIIYMAMSTLFFLRIAPLTPWWGALAAWVMILLIVEAIFAFFFPYRCCFNETEGDVQNNPHV |
| IRIS_313-9600 | MELITACALLLIAVHIAIVISCPEPLAAVALVDVPLLYLCRYLLIIRRGVIIVSGAGAGGGRLRRFRLGAAMAIIYMAMSTLFFLRIAPLTPWWGALAAWVMILLIVEAIFAFFFPYRCCFNETEGDVQNNPHV |
| IRIS_313-9601 | MELITACALLLIAVHIAIVISCPEPLAAVALVDVPLLYLCRYLLIIRRGVIIVSGAGAGGGRLRRFRLGAAMAIIYMAMSTLFFLRIAPLTPWWGALAAWVMILLIVEAIFAFFFPYRCCFNETEGDVQNNPHV |
| IRIS_313-9602 | MELITACALLLIAVHIAIVISCPEPLAAVALVDVPLLYLCRYLLIIRRGVIIVSGGGGGGRLRRFRLGAAMAIIYMAMSTLFFLRIAPLTPWWGALAAWVMILLIVEAIFAFFFPYRCCFNETEGDVQNNPHV |
| IRIS_313-9604 | MELITACALLLIAVHIAIVISCPEPLAAVALVDVPLLYLCRYLLIIRRGVIIVSGGGGGGRLRRFRLGAAMAIIYMAMSTLFFLRIAPLTPWWGALAAWVMILLIVEAIFAFFFPYRCCFNETEGDVQNNPHV |
| IRIS_313-9605 | MELITACALLLIAVHIAIVISCPEPLAAVALVDVPLLYLCRYLLIIRRGVIIVSGGGGGGRLRRFRLGAAMAIIYMAMSTLFFLRIAPLTPWWGALAAWVMILLIVEAIFAFFFPYRCCFNETEGDVQNNPHV |
| IRIS_313-9606 | MELITACALLLIAVHIAIVISCPEPLAAVALVDVPLLYLCRYLLIIRRGVIIVSGGGGGGRLRRFRLGAAMAIIYMAMSTLFFLRIAPLTPWWGALAAWVMILLIVEAIFAFFFPYRCCFNETEGDVQNNPHV |
| IRIS_313-9609 | MELITACALLLIAVHIAIVISCPEPLAAVALVDVPLLYLCRYLLIIRRGVIIVSGAGGGGGGRLRRFRLGAAMAIIYMAMSTLFFLRIAPLTPWWGALAAWVMILLIVEAIFAFFFPYRCCFNETEGDVQNNPHV |
| IRIS_313-9610 | MELITACALLLIAVHIAIVISCPEPLAAVAFVNVPLLYLCRYLLIIRRGVIIVSGGGGGGGRLRRFRLGAAMAVIYMAMSTLFFLRIAPLTPWWGALAAWVMILLIVEAIFAFFFPYRCCFNETEGDVQNNPHV |
| IRIS_313-9611 | MELITACALLLIAVHIAIVISCPEPLAAVALVDVPLLYLCRYLLIIRRGVIIVSGGGGGGRLRRFRLGAAMAIIYMAMSTLFFLRIAPLTPWWGALAAWVMILLIVEAIFAFFFPYRCCFNETEGDVQNNPHV |
| IRIS_313-9616 | MELITACALLLIAVHIAIVISCPEPLAAVALVDVPLLYLCRYLLIIRRGVIIVSGAGAGGGRLRRFRLGAAMAIIYMAMSTLFFLRIAPLTPWWGALAAWVMILLIVEAIFAFFFPYRCCFNETEGDVQNNPHV |
| IRIS_313-9617 | MELITACALLLIAVHIAIVISCPEPLAAVALVDVPLLYLCRYLLIIRRGVIIVSGAGAGGGRLRRFRLGAAMAIIYMAMSTLFFLRIAPLTPWWGALAAWVMILLIVEAIFAFFFPYRCCFNETEGDVQNNPHV |
| IRIS_313-9619 | MELITACALLLIAVHIAIVISCPEPLAAVALVDVPLLYLCRYLLIIRRGVIIVSGGGGGGRLRRFRLGAAMAIIYMAMSTLFFLRIAPLTPWWGALAAWVMILLIVEAIFAFFFPYRCCFNETEGDVQNNPHV |
| IRIS_313-9623 | MELITACALLLIAVHIAIVISCPEPLAAVALVDVPLLYLCRYLLIIRRGVIIVSGGGGGGRLRRFRLGAAMAIIYMAMSTLFFLRIAPLTPWWGALAAWVMILLIVEAIFAFFFPYRCCFNETEGDVQNNPHV |
| IRIS_313-9626 | MELITACALLLIAVHIAIVISCPEPLAAVALVDVPLLYLCRYLLIIRRGVIIVSGAGAGGGRLRRFRLGAAMAIIYMAMSTLFFLRIAPLTPWWGALAAWVMILLIVEAIFAFFFPYRCCFNETEGDVQNNPHV |
| IRIS_313-9629 | MELITACALLLIAVHIAIVISCPEPLAAVALVDVPLLYLCRYLLIIRRGVIIVSGAGAGGGRLRRFRLGAAMAIIYMAMSTLFFLRIAPLTPWWGALAAWVMILLIVEAIFAFFFPYRCCFNETEGDVQNNPHV |
| IRIS_313-9634 | MELITACALLLIAVHIAIVISCPEPLAAVALVDVPLLYLCRYLLIIRRGVIIVSGGGGGGRLRRFRLGAAMAIIYMAMSTLFFLRIAPLTPWWGALAAWVMILLIVEAIFAFFFPYRCCFNETEGDVQNNPHV |
| IRIS_313-9636 | MELITACALLLIAVHIAIVISCPEPLAAVALVDVPLLYLCRYLLIIRRGVIIVSGAGAGGGRLRRFRLGAAMAIIYMAMSTLFFLRIAPLTPWWGALAAWVMILLIVEAIFAFFFPYRCCFNETEGDVQNNPHV |
| IRIS_313-9637 | MELITACALLLIAVHIAIVISCPEPLAAVALVDVPLLYLCRYLLIIRRGVIIVSGAGAGGGRLRRFRLGAAMAIIYMAMSTLFFLRIAPLTPWWGALAAWVMILLIVEAIFAFFFPYRCCFNETEGDVQNNPHV |
| IRIS_313-9641 | MELITACALLLIAVHIAIVISCPEPLAAVALVDVPLLYLCRYLLIIRRGVIIVSGAGAGGGRLRRFRLGAAMAIIYMAMSTLFFLRIAPLTPWWGALAAWVMILLIVEAIFAFFFPYRCCFNETEGDVQNNPHV |
| IRIS_313-9648 | MELITACALLLIAVHIAIVISCPEPLAAVALVDVPLLYLYRYLLIIRRGVIIVSGAGGGGGRLRRFRLGAAMAIIYMAMSTLFFLRIAPLTPWWGALAAWVMILLIVEAIFAFFFPYRCCFNETEGDVQNNPHV |
| IRIS_313-9661 | MELITACALLLIAVHIAIVISCPEPLAAVAFVNVPLLYLCRYLLIIRRGVIIVSGGGGGGGRLRRFRLGAAMAVIYMAMSTLFFLRIAPLTPWWGALAAWVMILLIVEAIFAFFFPYRCCFNETEGDVQNNPHV |
| IRIS_313-9669 | MELITACALLLIAVHIAIVISCPEPLAAVALVDVPLLYLCRYLLIIRRGVIIVSGAGGAAAAGCAASASGQPWRSSTWPCRPSSSSALLLSRHGGAPSPPGS*SCSSSRPSSPSSSPTAAASTKPKAMSRIIPM |
| IRIS_313-9682 | MELITACALLLIAVHIAIVISCPEPLAAVALVDVPLLYLCRYLLIIRRGVIIVSGAGAGGGRLRRFRLGAAMAIIYMAMSTLFFLRIAPLTPWWGALAAWVMILLIVEAIFAFFFPYRCCFNETEGDVQNNPHV |
| IRIS_313-9685 | MELITACALLLIAVHIAIVISCPEPLAAVALVDVPLLYLCRYLLIIRRGVIIVSGGGGGGRLRRFRLGAAMAIIYMAMSTLFFLRIAPLTPWWGALAAWVMILLIVEAIFAFFFPYRCCFNETEGDVQNNPHV |
| IRIS_313-9687 | MELITACALLLIAVHIAIVISCPEPLAAVALVDVPLLYLCRYLLIIRRGVIIVSGGGGGGRLRRFRLGAAMAIIYMAMSTLFFLRIAPLTPWWGALAAWVMILLIVEAIFAFFFPYRCCFNETEGDVQNNPHV |
| IRIS_313-9691 | MELITACALLLIAVHIAIVISCPEPLAAVALVDVPLLYLCRYLLIIRRGVIIVSGAGAGGGRLRRFRLGAAMAIIYMAMSTLFFLRIAPLTPWWGALAAWVMILLIVEAIFAFFFPYRCCFNETEGDVQNNPHV |
| IRIS_313-9694 | MELITACALLLIAVHIAIVISCPEPLAAVALVDVPLLYLCRYLLIIRRGVIIVSGAGGAAAAGCAASASGQPWRSSTWPCRPSSSSALLLSRHGGAPSPPGS*SCSSSRPSSPSSSPTAAASTKPKAMSRIIPM |
| IRIS_313-9695 | MELITACALLLIAVHIAIVISCPEPLAAVALVDVPLLYLCRYLLIIRRGVIIVSGAGAGGGRLRRFRLGAAMAIIYMAMSTLFFLRIAPLTPWWGALAAWVMILLIVEAIFAFFFPYRCCFNETEGDVQNNPHV |
| IRIS_313-9696 | MELITACALLLIAVHIAIVISCPEPLAAVALVDVPLLYLCRYLLIIRRGVIIVSGGGGGGRLRRFRLGAAMAIIYMAMSTLFFLRIAPLTPWWGALAAWVMILLIVEAIFAFFFPYRCCFNETEGDVQNNPHV |
| IRIS_313-9697 | MELITACALLLIAVHIAIVISCPEPLAAVALVDVPLLYLCRYLLIIRRGVIIVSGGGGGGRLRRFRLGAAMAIIYMAMSTLFFLRIAPLTPWWGALAAWVMILLIVEAIFAFFFPYRCCFNETEGDVQNNPHV |
| IRIS_313-9698 | MELITACALLLIAVHIAIVISCPEPLAAVALVDVPLLYLCRYLLIIRRGVIIVSGAGAGGGRLRRFRLGAAMAIIYMAMSTLFFLRIAPLTPWWGALAAWVMILLIVEAIFAFFFPYRCCFNETEGDVQNNPHV |
| IRIS_313-9699 | MELITACALLLIAVHIAIVISCPEPLAAVALVDVPLLYLCRYLLIIRRGVIIVSGGGGGGRLRRFRLGAAMAIIYMAMSTLFFLRIAPLTPWWGALAAWVMILLIVEAIFAFFFPYRCCFNETEGDVQNNPHV |
| IRIS_313-9701 | MELITACALLLIAVHIAIVISCPEPLAAVALVDVPLLYLCRYLLIIRRGVIIVSGAGAGGGRLRRFRLGAAMAIIYMAMSTLFFLRIAPLTPWWGALAAWVMILLIVEAIFAFFFPYRCCFNETEGDVQNNPHV |
| IRIS_313-9702 | MELITACALLLIAVHIAIVISCPEPLAAVALVDVPLLYLCRYLLIIRRGVIIVSGAGAGGGRLRRFRLGAAMAIIYMAMSTLFFLRIAPLTPWWGALAAWVMILLIVEAIFAFFFPYRCCFNETEGDVQNNPHV |
| IRIS_313-9703 | MELITACALLLIAVHIAIVISCPEPLAAVALVDVPLLYLCRYLLIIRRGVIIVSGAGAGGGRLRRFRLGAAMAIIYMAMSTLFFLRIAPLTPWWGALAAWVMILLIVEAIFAFFFPYRCCFNETEGDVQNNPHV |
| IRIS_313-9705 | MELITACALLLIAVHIAIVISCPEPLAAVALVDVPLLYLCRYLLIIRRGVIIVSGGGGGGRLRRFRLGAAMAIIYMAMSTLFFLRIAPLTPWWGALAAWVMILLIVEAIFAFFFPYRCCFNETEGDVQNNPHV |
| IRIS_313-9706 | MELITACALLLIAVHIAIVISCPEPLAAVALVDVPLLYLCRYLLIIRRGVIIVSGAGAGGGRLRRFRLGAAMAIIYMAMSTLFFLRIAPLTPWWGALAAWVMILLIVEAIFAFFFPYRCCFNETEGDVQNNPHV |
| IRIS_313-9708 | MELITACALLLIAVHIAIVISCPEPLAAVALVDVPLLYLCRYLLIIRRGVIIVSGAGAGGGRLRRFRLGAAMAIIYMAMSTLFFLRIAPLTPWWGALAAWVMILLIVEAIFAFFFPYRCCFNETEGDVQNNPHV |
| IRIS_313-9709 | MELITACALLLIAVHIAIVISCPEPLAAVALVDVPLLYLCRYLLIIRRGVIIVSGAGAGGGRLRRFRLGAAMAIIYMAMSTLFFLRIAPLTPWWGALAAWVMILLIVEAIFAFFFPYRCCFNETEGDVQNNPHV |
| IRIS_313-9723 | MELITACALLLIAVHIAIVISCPEPLAAVALVDVPLLYLCRYLLIIRRGVIIVSGAGGGGGRLRRFRLGAAMAIIYMAMSTLFFLRIAPLTPWWGALAAWVMILLIVEAIFAFFFPYRCCFNETEGDVQNNPHV |
| IRIS_313-9724 | MELITACALLLIAVHIAIVISCPEPLAAVALVDVPLLYLCRYLLIIRRGVIIVSGAGAGGGRLRRFRLGAAMAIIYMAMSTLFFLRIAPLTPWWGALAAWVMILLIVEAIFAFFFPYRCCFNETEGDVQNNPHV |
| IRIS_313-9725 | MELITACALLLIAVHIAIVISCPEPLAAVALVDVPLLYLCRYLLIIRRGVIIVSGAGAGGGRLRRFRLGAAMAIIYMAMSTLFFLRIAPLTPWWGALAAWVMILLIVEAIFAFFFPYRCCFNETEGDVQNNPHV |
| IRIS_313-9727 | MELITACALLLIAVHIAIVISCPEPLAAVAFVNVPLLYLCRYLLIIRRGVIIVSGGGGGGGRLRRFRLGAAMAVIYMAMSTLFFLRIAPLTPWWGALAAWVMILLIVEAIFAFFFPYRCCFNETEGDVQNNPHV |
| IRIS_313-9730 | MELITACALLLIAVHIAIVISCPEPLAAVALVDVPLLYLYRYLLIIRRGVIIVSGAGGGGGRLRRFRLGAAMAIIYMAMSTLFFLRIAPLTPWWGALAAWVMILLIVEAIFAFFFPYRCCFNETEGDVQNNPHV |
| IRIS_313-9732 | MELITACALLLIAVHIAIVISCPEPLAAVALVDVPLLYLCRYLLIIRRGVIIVSGGGGGGRLRRFRLGAAMAIIYMAMSTLFFLRIAPLTPWWGALAAWVMILLIVEAIFAFFFPYRCCFNETEGDVQNNPHV |
| IRIS_313-9740 | MELITACALLLIAVHIAIVISCPEPLAAVALVDVPLLYLCRYLLIIRRGVIIVSGAGAGGGRLRRFRLGAAMAIIYMAMSTLFFLRIAPLTPWWGALAAWVMILLIVEAIFAFFFPYRCCFNETEGDVQNNPHV |
| IRIS_313-9742 | MELITACALLLIAVHIAIVISCPEPLAAVALVDVPLLYLCRYLLIIRRGVIIVSGAGAGGGRLRRFRLGAAMAIIYMAMSTLFFLRIAPLTPWWGALAAWVMILLIVEAIFAFFFPYRCCFNETEGDVQNNPHV |
| IRIS_313-9745 | MELITACALLLIAVHIAIVISCPEPLAAVALVDVPLLYLYRYLLIIRRGVIIVSGAGGGGGRLRRFRLGAAMAIIYMAMSTLFFLRIAPLTPWWGALAAWVMILLIVEAIFAFFFPYRCCFNETEGDVQNNPHV |
| IRIS_313-9758 | MELITACALLLIAVHIAIVISCPEPLAAVALVDVPLLYLCRYLLIIRRGVIIVSGGGGGGRLRRFRLGAAMAIIYMAMSTLFFLRIAPLTPWWGALAAWVMILLIVEAIFAFFFPYRCCFNETEGDVQNNPHV |
| IRIS_313-9759 | MELITACALLLIAVHIAIVISCPEPLAAVALVDVPLLYLCRYLLIIRRGVIIVSGAGAGGGRLRRFRLGAAMAIIYMAMSTLFFLRIAPLTPWWGALAAWVMILLIVEAIFAFFFPYRCCFNETEGDVQNNPHV |
| IRIS_313-9767 | MELITACALLLIAVHIAIVISCPEPLAAVALVDVPLLYLCRYLLIIRRGVIIVSGGGGGGRLRRFRLGAAMAIIYMAMSTLFFLRIAPLTPWWGALAAWVMILLIVEAIFAFFFPYRCCFNETEGDVQNNPHV |
| IRIS_313-9769 | MELITACALLLIAVHIAIVISCPEPLAAVALVDVPLLYLCRYLLIIRRGVIIVSGAGAGGGRLRRFRLGAAMAIIYMAMSTLFFLRIAPLTPWWGALAAWVMILLIVEAIFAFFFPYRCCFNETEGDVQNNPHV |
| IRIS_313-9770 | MELITACALLLIAVHIAIVISCPEPLAAVALVDVPLLYLCRYLLIIRRGVIIVSGAGAGGGRLRRFRLGAAMAIIYMAMSTLFFLRIAPLTPWWGALAAWVMILLIVEAIFAFFFPYRCCFNETEGDVQNNPHV |
| IRIS_313-9771 | MELITACALLLIAVHIAIVISCPEPLAAVALVDVPLLYLCRYLLIIRRGVIIVSGAGAGGGRLRRFRLGAAMAIIYMAMSTLFFLRIAPLTPWWGALAAWVMILLIVEAIFAFFFPYRCCFNETEGDVQNNPHV |
| IRIS_313-9774 | MELITACALLLIAVHIAIVISCPEPLAAVALVDVPLLYLCRYLLIIRRGVIIVSGAGGGGGRLRRFRLGAAMAIIYMAMSTLFFLRIAPLTPWWGALAAWVMILLIVEAIFAFFFPYRCCFNETEGDVQNNPHV |
| IRIS_313-9778 | MELITACALLLIAVHIAIVISCPEPLAAVALVDVPLLYLCRYLLIIRRGVIIVSGAGGAAAAGCAASASGQPWRSSTWPCRPSSSSALLLSRHGGAPSPPGS*SCSSSRPSSPSSSPTAAASTKPKAMSRIIPM |
| IRIS_313-9782 | MELITACALLLIAVHIAIVISCPEPLAAVALVDVPLLYLCRYLLIIRRGVIIVSGAGAGGGRLRRFRLGAAMAIIYMAMSTLFFLRIAPLTPWWGALAAWVMILLIVEAIFAFFFPYRCCFNETEGDVQNNPHV |
| IRIS_313-9783 | MELITACALLLIAVHIAIVISCPEPLAAVAFVNVPLLYLCRYLLIIRRGVIIVSGGGGGGGRLRRFRLGAAMAVIYMAMSTLFFLRIAPLTPWWGALAAWVMILLIVEAIFAFFFPYRCCFNETEGDVQNNPHV |
| IRIS_313-9789 | MELITACALLLIAVHIAIVISCPEPLAAVALVDVPLLYLYRYLLIIRRGVIIVSGAGGGGGRLRRFRLGAAMAIIYMAMSTLFFLRIAPLTPWWGALAAWVMILLIVEAIFAFFFPYRCCFNETEGDVQNNPHV |
| IRIS_313-9790 | MELITACALLLIAVHIAIVISCPEPLAAVALVDVPLLYLYRYLLIIRRGVIIVSGAGGGGGRLRRFRLGAAMAIIYMAMSTLFFLRIAPLTPWWGALAAWVMILLIVEAIFAFFFPYRCCFNETEGDVQNNPHV |
| IRIS_313-9791 | MELITACALLLIAVHIAIVISCPEPLAAVALVDVPLLYLCRYLLIIRRGVIIVSGAGAGGGRLRRFRLGAAMAIIYMAMSTLFFLRIAPLTPWWGALAAWVMILLIVEAIFAFFFPYRCCFNETEGDVQNNPHV |

```
IRIS_313-9795  MELITACALLLIAVHIAIVISCPEPLAAVALVDVPLLYLCRYLLIIRRGVIIVSGGGGGGRLRRFRLGAAMAIIYMAMSTLFFLRIAPLTPWWGALAAWVMILLIVEAIFAFFFPYRCCFNETEGDVQNNPHV
IRIS_313-9800  MELITACALLLIAVHIAIVISCPEPLAAVALVDVPLLYLYRYLLIIRRGVIIVSGAGGGGGRLRRFRLGAAMAIIYMAMSTLFFLRIAPLTPWWGALAAWVMILLIVEAIFAFFFPYRCCFNETEGDVQNNPHV
IRIS_313-9809  MELITACALLLIAVHIAIVISCPEPLAAVALVDVPLLYLCRYLLIIRRGVIIVSGGGGGGRLRRFRLGAAMAIIYMAMSTLFFLRIAPLTPWWGALAAWVMILLIVEAIFAFFFPYRCCFNETEGDVQNNPHV
IRIS_313-9811  MELITACALLLIAVHIAIVISCPEPLAAVALVDVPLLYLCRYLLIIRRGVIIVSGAGAGGGRLRRFRLGAAMAIIYMAMSTLFFLRIAPLTPWWGALAAWVMILLIVEAIFAFFFPYRCCFNETEGDVQNNPHV
IRIS_313-9813  MELITACALLLIAVHIAIVISCPEPLAAVALVDVPLLYLCRYLLIIRRGVIIVSGAGAGGGRLRRFRLGAAMAIIYMAMSTLFFLRIAPLTPWWGALAAWVMILLIVEAIFAFFFPYRCCFNETEGDVQNNPHV
IRIS_313-9814  MELITACALLLIAVHIAIVISCPEPLAAVALVDVPLLYLCRYLLIIRRGVIIVSGAGAGGGRLRRFRLGAAMAIIYMAMSTLFFLRIAPLTPWWGALAAWVMILLIVEAIFAFFFPYRCCFNETEGDVQNNPHV
IRIS_313-9817  MELITACALLLIAVHIAIVISCPEPLAAVALVDVPLLYLCRYLLIIRRGVIIVSGAGAGGGRLRRFRLGAAMAIIYMAMSTLFFLRIAPLTPWWGALAAWVMILLIVEAIFAFFFPYRCCFNETEGDVQNNPHV
IRIS_313-9818  MELITACALLLIAVHIAIVISCPEPLAAVALVDVPLLYLCRYLLIIRRGVIIVSGGGGGGRLRRFRLGAAMAIIYMAMSTLFFLRIAPLTPWWGALAAWVMILLIVEAIFAFFFPYRCCFNETEGDVQNNPHV
IRIS_313-9822  MELITACALLLIAVHIAIVISCPEPLAAVALVDVPLLYLCRYLLIIRRGVIIVSGGGGGGRLRRFRLGAAMAIIYMAMSTLFFLRIAPLTPWWGALAAWVMILLIVEAIFAFFFPYRCCFNETEGDVQNNPHV
IRIS_313-9825  MELITACALLLIAVHIAIVISCPEPLAAVALVDVPLLYLCRYLLIIRRGVIIVSGAGAGGGRLRRFRLGAAMAIIYMAMSTLFFLRIAPLTPWWGALAAWVMILLIVEAIFAFFFPYRCCFNETEGDVQNNPHV
IRIS_313-9831  MELITACALLLIAVHIAIVISCPEPLAAVALVDVPLLYLYRYLLIIRRGVIIVSGAGAGGGRLRRFRLGAAMAIIYMAMSTLFFLRIAPLTPWWGALAAWVMILLIVEAIFAFFFPYRCCFNETEGDVQNNPHV
IRIS_313-9832  MELITACALLLIAVHIAIVISCPEPLAAVALVDVPLLYLCRYLLIIRRGVIIVSGGGGGGRLRRFRLGAAMAIIYMAMSTLFFLRIAPLTPWWGALAAWVMILLIVEAIFAFFFPYRCCFNETEGDVQNNPHV
IRIS_313-9838  MELITACALLLIAVHIAIVISCPEPLAAVALVDVPLLYLCRYLLIIRRGVIIVSGAGAGGGRLRRFRLGAAMAIIYMAMSTLFFLRIAPLTPWWGALAAWVMILLIVEAIFAFFFPYRCCFNETEGDVQNNPHV
IRIS_313-9839  MELITACALLLIAVHIAIVISCPEPLAAVALVDVPLLYLCRYLLIIRRGVIIVSGAGAGGGRLRRFRLGAAMAIIYMAMSTLFFLRIAPLTPWWGALAAWVMILLIVEAIFAFFFPYRCCFNETEGDVQNNPHV
IRIS_313-9841  MELITACALLLIAVHIAIVISCPEPLAAVALVDVPLLYLCRYLLIIRRGVIIVSGAGAGGGRLRRFRLGAAMAIIYMAMSTLFFLRIAPLTPWWGALAAWVMILLIVEAIFAFFFPYRCCFNETEGDVQNNPHV
IRIS_313-9851  MELITACALLLIAVHIAIVISCPEPLAAVALVDVPLLYLCRYLLIIRRGVIIVSGAGAGGGRLRRFRLGAAMAIIYMAMSTLFFLRIAPLTPWWGALAAWVMILLIVEAIFAFFFPYRCCFNETEGDVQNNPHV
IRIS_313-9861  MELITACALLLIAVHIAIVISCPEPLAAVALVDVPLLYLCRYLLIIRRGVIIVSGAGAGGGRLRRFRLGAAMAIIYMAMSTLFFLRIAPLTPWWGALAAWVMILLIVEAIFAFFFPYRCCFNETEGDVQNNPHV
IRIS_313-9862  MELITACALLLIAVHIAIVISCPEPLAAVALVDVPLLYLCRYLLIIRRGVIIVSGGGGGGRLRRFRLGAAMAIIYMAMSTLFFLRIAPLTPWWGALAAWVMILLIVEAIFAFFFPYRCCFNETEGDVQNNPHV
IRIS_313-9867  MELITACALLLIAVHIAIVISCPEPLAAVALVDVPLLYLCRYLLIIRRGVIIVSGGGAGGGRLRRFRLGAAMAIIYMAMSTLFFLRIAPLTPWWGALAAWVMILLIVEAIFAFFFPYRCCFNETEGDVQNNPHV
IRIS_313-9880  MELITACALLLIAVHIAIVISCPEPLAAVALVDVPLLYLCRYLLIIRRGVIIVSGAGAGGGRLRRFRLGAAMAIIYMAMSTLFFLRIAPLTPWWGALAAWVMILLIVEAIFAFFFPYRCCFNETEGDVQNNPHV
IRIS_313-9882  MELITACALLLIAVHIAIVISCPEPLAAVALVDVPLLYLCRYLLIIRRGVIIVSGGGGGGRLRRFRLGAAMAIIYMAMSTLFFLRIAPLTPWWGALAAWVMILLIVEAIFAFFFPYRCCFNETEGDVQNNPHV
IRIS_313-9884  MELITACALLLIAVHIAIVISCPEPLAAVALVDVPLLYLCRYLLIIRRGVIIVSGAGAGGGRLRRFRLGAAMAIIYMAMSTLFFLRIAPLTPWWGALAAWVMILLIVEAIFAFFFPYRCCFNETEGDVQNNPHV
IRIS_313-9886  MELITACALLLIAVHIAIVISCPEPLAAVALVDVPLLYLCRYLLIIRRGVIIVSGAGAGGGRLRRFRLGAAMAIIYMAMSTLFFLRIAPLTPWWGALAAWVMILLIVEAIFAFFFPYRCCFNETEGDVQNNPHV
IRIS_313-9887  MELITACALLLIAVHIAIVISCPEPLAAVALVDVPLLYLCRYLLIIRRGVIIVSGAGAGGGRLRRFRLGAAMAIIYMAMSTLFFLRIAPLTPWWGALAAWVMILLIVEAIFAFFFPYRCCFNETEGDVQNNPHV
IRIS_313-9890  MELITACALLLIAVHIAIVISCPEPLAAVALVDVPLLYLCRYLLIIRRGVIIVSGAGAGGGRLRRFRLGAAMAIIYMAMSTLFFLRIAPLTPWWGALAAWVMILLIVEAIFAFFFPYRCCFNETEGDVQNNPHV
IRIS_313-9891  MELITACALLLIAVHIAIVISCPEPLAAVALVDVPLLYLCRYLLIIRRGVIIVSGAGAGGGRLRRFRLGAAMAIIYMAMSTLFFLRIAPLTPWWGALAAWVMILLIVEAIFAFFFPYRCCFNETEGDVQNNPHV
IRIS_313-9897  MELITACALLLIAVHIAIVISCPEPLAAVALVDVPLLYLCRYLLIIRRGVIIVSGAGAGGGRLRRFRLGAAMAIIYMAMSTLFFLRIAPLTPWWGALAAWVMILLIVEAIFAFFFPYRCCFNETEGDVQNNPHV
IRIS_313-9898  MELITACALLLIAVHIAIVISCPEPLAAVALVDVPLLYLYRYLLIIRRGVIIVSGAGGGGGRLRRFRLGAAMAIIYMAMSTLFFLRIAPLTPWWGALAAWVMILLIVEAIFAFFFPYRCCFNETEGDVQNNPHV
IRIS_313-9917  MELITACALLLIAVHIAIVISCPEPLAAVALVDVPLLYLCRYLLIIRRGVIIVSGGGGGGRLRRFRLGAAMAIIYMAMSTLFFLRIAPLTPWWGALAAWVMILLIVEAIFAFFFPYRCCFNETEGDVQNNPHV
IRIS_313-9922  MELITACALLLIAVHIAIVISCPEPLAAVALVDVPLLYLCRYLLIIRRGVIIVSGGGGGGRLRRFRLGAAMAIIYMAMSTLFFLRIAPLTPWWGALAAWVMILLIVEAIFAFFFPYRCCFNETEGDVQNNPHV
IRIS_313-9924  MELITACALLLIAVHIAIVISCPEPLAAVALVDVPLLYLCRYLLIIRRGVIIVSGGGGGGRLRRFRLGAAMAIIYMAMSTLFFLRIAPLTPWWGALAAWVMILLIVEAIFAFFFPYRCCFNETEGDVQNNPHV
IRIS_313-9925  MELITACALLLIAVHIAIVISCPEPLAAVALVDVPLLYLCRYLLIIRRGVIIVSGAGAGGGRLRRFRLGAAMAIIYMAMSTLFFLRIAPLTPWWGALAAWVMILLIVEAIFAFFFPYRCCFNETEGDVQNNPHV
IRIS_313-9926  MELITACALLLIAVHIAIVISCPEPLAAVALVDVPLLYLYRYLLIIRRGVIIVSGAGGGGGRLRRFRLGAAMAIIYMAMSTLFFLRIAPLTPWWGALAAWVMILLIVEAIFAFFFPYRCCFNETEGDVQNNPHV
IRIS_313-9928  MELITACALLLIAVHIAIVISCPEPLAAVALVDVPLLYLYRYLLIIRRGVIIVSGAGAGGGRLRRFRLGAAMAIIYMAMSTLFFLRIAPLTPWWGALAAWVMILLIVEAIFAFFFPYRCCFNETEGDVQNNPHV
IRIS_313-9929  MELITACALLLIAVHIAIVISCPEPLAAVALVDVPLLYLCRYLLIIRRGVIIVSGAGGGGGRLRRFRLGAAMAIIYMAMSTLFFLRIAPLTPWWGALAAWVMILLIVEAIFAFFFPYRCCFNETEGDVQNNPHV
IRIS_313-9935  MELITACALLLIAVHIAIVISCPEPLAAVALVDVPLLYLCRYLLIIRRGVIIVSGAGAGGGRLRRFRLGAAMAIIYMAMSTLFFLRIAPLTPWWGALAAWVMILLIVEAIFAFFFPYRCCFNETEGDVQNNPHV
IRIS_313-9936  MELITACALLLIAVHIAIVISCPEPLAAVALVDVPLLYLCRYLLIIRRGVIIVSGGGGGGRLRRFRLGAAMAIIYMAMSTLFFLRIAPLTPWWGALAAWVMILLIVEAIFAFFFPYRCCFNETEGDVQNNPHV
IRIS_313-9937  MELITACALLLIAVHIAIVISCPEPLAAVALVDVPLLYLYRYLLIIRRGVIIVSGAGGGGGRLRRFRLGAAMAIIYMAMSTLFFLRIAPLTPWWGALAAWVMILLIVEAIFAFFFPYRCCFNETEGDVQNNPHV
IRIS_313-9939  MELITACALLLIAVHIAIVISCPEPLAAVALVDVPLLYLCRYLLIIRRGVIIVSGGGAGGGRLRRFRLGAAMAIIYMAMSTLFFLRIAPLTPWWGALAAWVMILLIVEAIFAFFFPYRCCFNETEGDVQNNPHV
IRIS_313-9940  MELITACALLLIAVHIAIVISCPEPLAAVALVDVPLLYLCRYLLIIRRGVIIVSGGGGGGRLRRFRLGAAMAIIYMAMSTLFFLRIAPLTPWWGALAAWVMILLIVEAIFAFFFPYRCCFNETEGDVQNNPHV
IRIS_313-9944  MELITACALLLIAVHIAIVISCPEPLAAVALVDVPLLYLCRYLLIIRRGVIIVSGGGGGGRLRRFRLGAAMAIIYMAMSTLFFLRIAPLTPWWGALAAWVMILLIVEAIFAFFFPYRCCFNETEGDVQNNPHV
IRIS_313-9949  MELITACALLLIAVHIAIVISCPEPLAAVALVDVPLLYLYRYLLIIRRGVIIVSGAGGGGGRLRRFRLGAAMAIIYMAMSTLFFLRIAPLTPWWGALAAWVMILLIVEAIFAFFFPYRCCFNETEGDVQNNPHV
IRIS_313-9953  MELITACALLLIAVHIAIVISCPEPLAAVALVDVPLLYLYRYLLIIRRGVIIVSGAGGGGGRLRRFRLGAAMAIIYMAMSTLFFLRIAPLTPWWGALAAWVMILLIVEAIFAFFFPYRCCFNETEGDVQNNPHV
IRIS_313-9961  MELITACALLLIAVHIAIVISCPEPLAAVALVDVPLLYLCRYLLIIRRGVIIVSGAGAGGGRLRRFRLGAAMAIIYMAMSTLFFLRIAPLTPWWGALAAWVMILLIVEAIFAFFFPYRCCFNETEGDVQNNPHV
IRIS_313-9963  MELITACALLLIAVHIAIVISCPEPLAAVAFVNVPLLYLCRYLLIIRRGVIIVSGGGGGGGRLRRFRLGAAMAVIYMAMSTLFFLRIAPLTPWWGALAAWVMILLIVEAIFAFFFPYRCCFNETEGDVQNNPHV
IRIS_313-9964  MELITACALLLIAVHIAIVISCPEPLAAVALVDVPLLYLCRYLLIIRRGVIIVSGAGAGGGRLRRFRLGAAMAIIYMAMSTLFFLRIAPLTPWWGALAAWVMILLIVEAIFAFFFPYRCCFNETEGDVQNNPHV
IRIS_313-9966  MELITACALLLIAVHIAIVISCPEPLAAVALVDVPLLYLCRYLLIIRRGVIIVSGGGGGGRLRRFRLGAAMAIIYMAMSTLFFLRIAPLTPWWGALAAWVMILLIVEAIFAFFFPYRCCFNETEGDVQNNPHV
IRIS_313-9968  MELITACALLLIAVHIAIVISCPEPLAAVALVDVPLLYLYRYLLIIRRGVIIVSGAGGGGGRLRRFRLGAAMAIIYMAMSTLFFLRIAPLTPWWGALAAWVMILLIVEAIFAFFFPYRCCFNETEGDVQNNPHV
IRIS_313-9969  MELITACALLLIAVHIAIVISCPEPLAAVALVDVPLLYLCRYLLIIRRGVIIVSGAGGGGGRLRRFRLGAAMAIIYMAMSTLFFLRIAPLTPWWGALAAWVMILLIVEAIFAFFFPYRCCFNETEGDVQNNPHV
IRIS_313-9970  MELITACALLLIAVHIAIVISCPEPLAAVALVDVPLLYLCRYLLIIRRGVIIVSGGGGGGRLRRFRLGAAMAIIYMAMSTLFFLRIAPLTPWWGALAAWVMILLIVEAIFAFFFPYRCCFNETEGDVQNNPHV
IRIS_313-9974  MELITACALLLIAVHIAIVISCPEPLAAVALVDVPLLYLCRYLLIIRRGVIIVSGAGAGGGRLRRFRLGAAMAIIYMAMSTLFFLRIAPLTPWWGALAAWVMILLIVEAIFAFFFPYRCCFNETEGDVQNNPHV
IRIS_313-9976  MELITACALLLIAVHIAIVISCPEPLAAVALVDVPLLYLCRYLLIIRRGVIIVSGGGGGGRLRRFRLGAAMAIIYMAMSTLFFLRIAPLTPWWGALAAWVMILLIVEAIFAFFFPYRCCFNETEGDVQNNPHV
IRIS_313-9978  MELITACALLLIAVHIAIVISCPEPLAAVALVDVPLLYLCRYLLIIRRGVIIVSGAGAGGGRLRRFRLGAAMAIIYMAMSTLFFLRIAPLTPWWGALAAWVMILLIVEAIFAFFFPYRCCFNETEGDVQNNPHV
IRIS_313-9980  MELITACALLLIAVHIAIVISCPEPLAAVALVDVPLLYLCRYLLIIRRGVIIVSGAGAGGGRLRRFRLGAAMAIIYMAMSTLFFLRIAPLTPWWGALAAWVMILLIVEAIFAFFFPYRCCFNETEGDVQNNPHV
IRIS_313-9986  MELITACALLLIAVHIAIVISCPEPLAAVALVDVPLLYLCRYLLIIRRGVIIVSGAGAGGGRLRRFRLGAAMAIIYMAMSTLFFLRIAPLTPWWGALAAWVMILLIVEAIFAFFFPYRCCFNETEGDVQNNPHV
IRIS_313-9989  MELITACALLLIAVHIAIVISCPEPLAAVALVDVPLLYLCRYLLIIRRGVIIVSGGGGGGRLRRFRLGAAMAIIYMAMSTLFFLRIAPLTPWWGALAAWVMILLIVEAIFAFFFPYRCCFNETEGDVQNNPHV
IRIS_313-9995  MELITACALLLIAVHIAIVISCPEPLAAVALVDVPLLYLYRYLLIIRRGVIIVSGAGGGGGRLRRFRLGAAMAIIYMAMSTLFFLRIAPLTPWWGALAAWVMILLIVEAIFAFFFPYRCCFNETEGDVQNNPHV
IRIS_313-9996  MELITACALLLIAVHIAIVISCPEPLAAVALVDVPLLYLCRYLLIIRRGVIIVSGAGAGGGRLRRFRLGAAMAIIYMAMSTLFFLRIAPLTPWWGALAAWVMILLIVEAIFAFFFPYRCCFNETEGDVQNNPHV
IRIS_313-10000 MELITACALLLIAVHIAIVISCPEPLAAVALVDVPLLYLCRYLLIIRRGVIIVSGAGAGGGRLRRFRLGAAMAIIYMAMSTLFFLRIAPLTPWWGALAAWVMILLIVEAIFAFFFPYRCCFNETEGDVQNNPHV
IRIS_313-10001 MELITACALLLIAVHIAIVISCPEPLAAVALVDVPLLYLCRYLLIIRRGVIIVSGGGGGGRLRRFRLGAAMAIIYMAMSTLFFLRIAPLTPWWGALAAWVMILLIVEAIFAFFFPYRCCFNETEGDVQNNPHV
IRIS_313-10002 MELITACALLLIAVHIAIVISCPEPLAAVALVDVPLLYLCRYLLIIRRGVIIVSGGGAGGGRLRRFRLGAAMAIIYMAMSTLFFLRIAPLTPWWGALAAWVMILLIVEAIFAFFFPYRCCFNETEGDVQNNPHV
IRIS_313-10007 MELITACALLLIAVHIAIVISCPEPLAAVALVDVPLLYLCRYLLIIRRGVIIVSGAGAGGGRLRRFRLGAAMAIIYMAMSTLFFLRIAPLTPWWGALAAWVMILLIVEAIFAFFFPYRCCFNETEGDVQNNPHV
IRIS_313-10010 MELITACALLLIAVHIAIVISCPEPLAAVALVDVPLLYLCRYLLIIRRGVIIVSGAGAGGGRLRRFRLGAAMAIIYMAMSTLFFLRIAPLTPWWGALAAWVMILLIVEAIFAFFFPYRCCFNETEGDVQNNPHV
IRIS_313-10014 MELITACALLLIAVHIAIVISCPEPLAAVALVDVPLLYLCRYLLIIRRGVIIVSGAGAGGGRLRRFRLGAAMAIIYMAMSTLFFLRIAPLTPWWGALAAWVMILLIVEAIFAFFFPYRCCFNETEGDVQNNPHV
IRIS_313-10016 MELITACALLLIAVHIAIVISCPEPLAAVALVDVPLLYLCRYLLIIRRGVIIVSGAGAGGGRLRRFRLGAAMAIIYMAMSTLFFLRIAPLTPWWGALAAWVMILLIVEAIFAFFFPYRCCFNETEGDVQNNPHV
IRIS_313-10020 MELITACALLLIAVHIAIVISCPEPLAAVALVDVPLLYLCRYLLIIRRGVIIVSGAGAGGGRLRRFRLGAAMAIIYMAMSTLFFLRIAPLTPWWGALAAWVMILLIVEAIFAFFFPYRCCFNETEGDVQNNPHV
IRIS_313-10025 MELITACALLLIAVHIAIVISCPEPLAAVALVDVPLLYLCRYLLIIRRGVIIVSGAGAGGGRLRRFRLGAAMAIIYMAMSTLFFLRIAPLTPWWGALAAWVMILLIVEAIFAFFFPYRCCFNETEGDVQNNPHV
IRIS_313-10026 MELITACALLLIAVHIAIVISCPEPLAAVALVDVPLLYLCRYLLIIRRGVIIVSGAGAGGGRLRRFRLGAAMAIIYMAMSTLFFLRIAPLTPWWGALAAWVMILLIVEAIFAFFFPYRCCFNETEGDVQNNPHV
IRIS_313-10030 MELITACALLLIAVHIAIVISCPEPLAAVALVDVPLLYLCRYLLIIRRGVIIVSGAGGGGGGRLRRFRLGAAMAIIYMAMSTLFFLRIAPLTPWWGALAAWVMILLIVEAIFAFFFPYRCCFNETEGDVQNNPHV
IRIS_313-10032 MELITACALLLIAVHIAIVISCPEPLAAVALVDVPLLYLCRYLLIIRRGVIIVSGAGAGGGRLRRFRLGAAMAIIYMAMSTLFFLRIAPLTPWWGALAAWVMILLIVEAIFAFFFPYRCCFNETEGDVQNNPHV
IRIS_313-10034 MELITACALLLIAVHIAIVISCPEPLAAVALVDVPLLYLCRYLLIIRRGVIIVSGGGGGGRLRRFRLGAAMAIIYMAMSTLFFLRIAPLTPWWGALAAWVMILLIVEAIFAFFFPYRCCFNETEGDVQNNPHV
IRIS_313-10035 MELITACALLLIAVHIAIVISCPEPLAAVALVDVPLLYLCRYLLIIRRGVIIVSGAGAGGGRLRRFRLGAAMAIIYMAMSTLFFLRIAPLTPWWGALAAWVMILLIVEAIFAFFFPYRCCFNETEGDVQNNPHV
IRIS_313-10040 MELITACALLLIAVHIAIVISCPEPLAAVALVDVPLLYLCRYLLIIRRGVIIVSGGGGGGRLRRFRLGAAMAIIYMAMSTLFFLRIAPLTPWWGALAAWVMILLIVEAIFAFFFPYRCCFNETEGDVQNNPHV
IRIS_313-10041 MELITACALLLIAVHIAIVISCPEPLAAVALVDVPLLYLYRYLLIIRRGVIIVSGAGGGGGRLRRFRLGAAMAIIYMAMSTLFFLRIAPLTPWWGALAAWVMILLIVEAIFAFFFPYRCCFNETEGDVQNNPHV
IRIS_313-10045 MELITACALLLIAVHIAIVISCPEPLAAVALVDVPLLYLCRYLLIIRRGVIIVSGAGAGGGRLRRFRLGAAMAIIYMAMSTLFFLRIAPLTPWWGALAAWVMILLIVEAIFAFFFPYRCCFNETEGDVQNNPHV
IRIS_313-10046 MELITACALLLIAVHIAIVISCPEPLAAVALVDVPLLYLCRYLLIIRRGVIIVSGGGAGGGRLRRFRLGAAMAIIYMAMSTLFFLRIAPLTPWWGALAAWVMILLIVEAIFAFFFPYRCCFNETEGDVQNNPHV
IRIS_313-10047 MELITACALLLIAVHIAIVISCPEPLAAVALVDVPLLYLCRYLLIIRRGVIIVSGAGAGGGRLRRFRLGAAMAIIYMAMSTLFFLRIAPLTPWWGALAAWVMILLIVEAIFAFFFPYRCCFNETEGDVQNNPHV
IRIS_313-10048 MELITACALLLIAVHIAIVISCPEPLAAVALVDVPLLYLCRYLLIIRRGVIIVSGAGAGGGRLRRFRLGAAMAIIYMAMSTLFFLRIAPLTPWWGALAAWVMILLIVEAIFAFFFPYRCCFNETEGDVQNNPHV
IRIS_313-10050 MELITACALLLIAVHIAIVISCPEPLAAVALVDVPLLYLCRYLLIIRRGVIIVSGGGGGGRLRRFRLGAAMAIIYMAMSTLFFLRIAPLTPWWGALAAWVMILLIVEAIFAFFFPYRCCFNETEGDVQNNPHV
IRIS_313-10051 MELITACALLLIAVHIAIVISCPEPLAAVALVDVPLLYLYRYLLIIRRGVIIVSGAGGGGGRLRRFRLGAAMAIIYMAMSTLFFLRIAPLTPWWGALAAWVMILLIVEAIFAFFFPYRCCFNETEGDVQNNPHV
IRIS_313-10054 MELITACALLLIAVHIAIVISCPEPLAAVALVDVPLLYLCRYLLIIRRGVIIVSGAGAGGGRLRRFRLGAAMAIIYMAMSTLFFLRIAPLTPWWGALAAWVMILLIVEAIFAFFFPYRCCFNETEGDVQNNPHV
IRIS_313-10056 MELITACALLLIAVHIAIVISCPEPLAAVALVDVPLLYLCRYLLIIRRGVIIVSGAGGGGGRLRRFRLGAAMAIIYMAMSTLFFLRIAPLTPWWGALAAWVMILLIVEAIFAFFFPYRCCFNETEGDVQNNPHV
IRIS_313-10059 MELITACALLLIAVHIAIVISCPEPLAAVAFVNVPLLYLCRYLLIIRRGVIIVSGGGGGGGRLRRFRLGAAMAVIYMAMSTLFFLRIAPLTPWWGALAAWVMILLIVEAIFAFFFPYRCCFNETEGDVQNNPHV
IRIS_313-10061 MELITACALLLIAVHIAIVISCPEPLAAVALVDVPLLYLYRYLLIIRRGVIIVSGAGGGGGRLRRFRLGAAMAIIYMAMSTLFFLRIAPLTPWWGALAAWVMILLIVEAIFAFFFPYRCCFNETEGDVQNNPHV
IRIS_313-10062 MELITACALLLIAVHIAIVISCPEPLAAVALVDVPLLYLYRYLLIIRRGVIIVSGAGGGGGRLRRFRLGAAMAIIYMAMSTLFFLRIAPLTPWWGALAAWVMILLIVEAIFAFFFPYRCCFNETEGDVQNNPHV
IRIS_313-10065 MELITACALLLIAVHIAIVISCPEPLAAVALVDVPLLYLYRYLLIIRRGVIIVSGAGGGGGRLRRFRLGAAMAIIYMAMSTLFFLRIAPLTPWWGALAAWVMILLIVEAIFAFFFPYRCCFNETEGDVQNNPHV
IRIS_313-10067 MELITACALLLIAVHIAIVISCPEPLAAVALVDVPLLYLYRYLLIIRRGVIIVSGAGGGGGRLRRFRLGAAMAIIYMAMSTLFFLRIAPLTPWWGALAAWVMILLIVEAIFAFFFPYRCCFNETEGDVQNNPHV
IRIS_313-10071 MELITACALLLIAVHIAIVISCPEPLAAVALVDVPLLYLYRYLLIIRRGVIIVSGAGGGGGRLRRFRLGAAMAIIYMAMSTLFFLRIAPLTPWWGALAAWVMILLIVEAIFAFFFPYRCCFNETEGDVQNNPHV
IRIS_313-10073 MELITACALLLIAVHIAIVISCPEPLAAVALVDVPLLYLYRYLLIIRRGVIIVSGAGGGGGRLRRFRLGAAMAIIYMAMSTLFFLRIAPLTPWWGALAAWVMILLIVEAIFAFFFPYRCCFNETEGDVQNNPHV
IRIS_313-10074 MELITACALLLIAVHIAIVISCPEPLAAVALVDVPLLYLYRYLLIIRRGVIIVSGAGGGGGRLRRFRLGAAMAIIYMAMSTLFFLRIAPLTPWWGALAAWVMILLIVEAIFAFFFPYRCCFNETEGDVQNNPHV
IRIS_313-10075 MELITACALLLIAVHIAIVISCPEPLAAVALVDVPLLYLYRYLLIIRRGVIIVSGAGGGGGRLRRFRLGAAMAIIYMAMSTLFFLRIAPLTPWWGALAAWVMILLIVEAIFAFFFPYRCCFNETEGDVQNNPHV
IRIS_313-10076 MELITACALLLIAVHIAIVISCPEPLAAVALVDVPLLYLCRYLLIIRRGVIIVSGAGGGGGRLRRFRLGAAMAIIYMAMSTLFFLRIAPLTPWWGALAAWVMILLIVEAIFAFFFPYRCCFNETEGDVQNNPHV
IRIS_313-10077 MELITACALLLIAVHIAIVISCPEPLAAVALVDVPLLYLCRYLLIIRRGVIIVSGAGAGGGRLRRFRLGAAMAIIYMAMSTLFFLRIAPLTPWWGALAAWVMILLIVEAIFAFFFPYRCCFNETEGDVQNNPHV
IRIS_313-10078 MELITACALLLIAVHIAIVISCPEPLAAVALVDVPLLYLYRYLLIIRRGVIIVSGAGGGGGRLRRFRLGAAMAIIYMAMSTLFFLRIAPLTPWWGALAAWVMILLIVEAIFAFFFPYRCCFNETEGDVQNNPHV
IRIS_313-10079 MELITACALLLIAVHIAIVISCPEPLAAVALVDVPLLYLYRYLLIIRRGVIIVSGAGGGGGRLRRFRLGAAMAIIYMAMSTLFFLRIAPLTPWWGALAAWVMILLIVEAIFAFFFPYRCCFNETEGDVQNNPHV
IRIS_313-10080 MELITACALLLIAVHIAIVISCPEPLAAVALVDVPLLYLYRYLLIIRRGVIIVSGAGGGGGRLRRFRLGAAMAIIYMAMSTLFFLRIAPLTPWWGALAAWVMILLIVEAIFAFFFPYRCCFNETEGDVQNNPHV
IRIS_313-10082 MELITACALLLIAVHIAIVISCPEPLAAVALVDVPLLYLYRYLLIIRRGVIIVSGAGGGGGRLRRFRLGAAMAIIYMAMSTLFFLRIAPLTPWWGALAAWVMILLIVEAIFAFFFPYRCCFNETEGDVQNNPHV
IRIS_313-10083 MELITACALLLIAVHIAIVISCPEPLAAVALVDVPLLYLCRYLLIIRRGVIIVSGAGAGGGRLRRFRLGAAMAIIYMAMSTLFFLRIAPLTPWWGALAAWVMILLIVEAIFAFFFPYRCCFNETEGDVQNNPHV
IRIS_313-10084 MELITACALLLIAVHIAIVISCPEPLAAVALVDVPLLYLCRYLLIIRRGVIIVSGAGAGGGRLRRFRLGAAMAIIYMAMSTLFFLRIAPLTPWWGALAAWVMILLIVEAIFAFFFPYRCCFNETEGDVQNNPHV
IRIS_313-10089 MELITACALLLIAVHIAIVISCPEPLAAVALVDVPLLYLCRYLLIIRRGVIIVSGAGAGGGRLRRFRLGAAMAIIYMAMSTLFFLRIAPLTPWWGALAAWVMILLIVEAIFAFFFPYRCCFNETEGDVQNNPHV
IRIS_313-10092 MELITACALLLIAVHIAIVISCPEPLAAVALVDVPLLYLCRYLLIIRRGVIIVSGAGAGGGRLRRFRLGAAMAIIYMAMSTLFFLRIAPLTPWWGALAAWVMILLIVEAIFAFFFPYRCCFNETEGDVQNNPHV
IRIS_313-10093 MELITACALLLIAVHIAIVISCPEPLAAVALVDVPLLYLCRYLLIIRRGVIIVSGAGAGGGRLRRFRLGAAMAIIYMAMSTLFFLRIAPLTPWWGALAAWVMILLIVEAIFAFFFPYRCCFNETEGDVQNNPHV
IRIS_313-10094 MELITACALLLIAVHIAIVISCPEPLAAVALVDVPLLYLYRYLLIIRRGVIIVSGAGGGGGRLRRFRLGAAMAIIYMAMSTLFFLRIAPLTPWWGALAAWVMILLIVEAIFAFFFPYRCCFNETEGDVQNNPHV
IRIS_313-10096 MELITACALLLIAVHIAIVISCPEPLAAVALVDVPLLYLYRYLLIIRRGVIIVSGAGGGGGRLRRFRLGAAMAIIYMAMSTLFFLRIAPLTPWWGALAAWVMILLIVEAIFAFFFPYRCCFNETEGDVQNNPHV
IRIS_313-10097 MELITACALLLIAVHIAIVISCPEPLAAVALVDVPLLYLCRYLLIIRRGVIIVSGAGAGGGRLRRFRLGAAMAIIYMAMSTLFFLRIAPLTPWWGALAAWVMILLIVEAIFAFFFPYRCCFNETEGDVQNNPHV
IRIS_313-10099 MELITACALLLIAVHIAIVISCPEPLAAVALVDVPLLYLCRYLLIIRRGVIIVSGAGAGGGRLRRFRLGAAMAIIYMAMSTLFFLRIAPLTPWWGALAAWVMILLIVEAIFAFFFPYRCCFNETEGDVQNNPHV
IRIS_313-10102 MELITACALLLIAVHIAIVISCPEPLAAVALVDVPLLYLCRYLLIIRRGVIIVSGAGAGGGRLRRFRLGAAMAIIYMAMSTLFFLRIAPLTPWWGALAAWVMILLIVEAIFAFFFPYRCCFNETEGDVQNNPHV
IRIS_313-10103 MELITACALLLIAVHIAIVISCPEPLAAVALVDVPLLYLCRYLLIIRRGVIIVSGAGAGGGRLRRFRLGAAMAIIYMAMSTLFFLRIAPLTPWWGALAAWVMILLIVEAIFAFFFPYRCCFNETEGDVQNNPHV
IRIS_313-10109 MELITACALLLIAVHIAIVISCPEPLAAVALVDVPLLYLCRYLLIIRRGVIIVSGGGGAAAAGCAASASGQPWRSSTWPCRPSSSSALLLSRHGGAPSPPGS*SCSSSRPSSPSSSPTAAASTKPKAMSRIIPM
IRIS_313-10111 MELITACALLLIAVHIAIVISCPEPLAAVALVDVPLLYLCRYLLIIRRGVIIVSGAGAGGGRLRRFRLGAAMAIIYMAMSTLFFLRIAPLTPWWGALAAWVMILLIVEAIFAFFFPYRCCFNETEGDVQNNPHV
IRIS_313-10113 MELITACALLLIAVHIAIVISCPEPLAAVALVDVPLLYLCRYLLIIRRGVIIVSGGGGGGRLRRFRLGAAMAIIYMAMSTLFFLRIAPLTPWWGALAAWVMILLIVEAIFAFFFPYRCCFNETEGDVQNNPHV
IRIS_313-10114 MELITACALLLIAVHIAIVISCPEPLAAVALVDVPLLYLCRYLLIIRRGVIIVSGAGAGGGRLRRFRLGAAMAIIYMAMSTLFFLRIAPLTPWWGALAAWVMILLIVEAIFAFFFPYRCCFNETEGDVQNNPHV
IRIS_313-10119 MELITACALLLIAVHIAIVISCPEPLAAVALVDVPLLYLYRYLLIIRRGVIIVSGAGGGGGRLRRFRLGAAMAIIYMAMSTLFFLRIAPLTPWWGALAAWVMILLIVEAIFAFFFPYRCCFNETEGDVQNNPHV
IRIS_313-10124 MELITACALLLIAVHIAIVISCPEPLAAVALVDVPLLYLCRYLLIIRRGVIIVSGAGAGGGRLRRFRLGAAMAIIYMAMSTLFFLRIAPLTPWWGALAAWVMILLIVEAIFAFFFPYRCCFNETEGDVQNNPHV
IRIS_313-10129 MELITACALLLIAVHIAIVISCPEPLAAVALVDVPLLYLCRYLLIIRRGVIIVSGGGGGGRLRRFRLGAAMAIIYMAMSTLFFLRIAPLTPWWGALAAWVMILLIVEAIFAFFFPYRCCFNETEGDVQNNPHV
IRIS_313-10134 MELITACALLLIAVHIAIVISCPEPLAAVALVDVPLLYLCRYLLIIRRGVIIVSGGGGGGRLRRFRLGAAMAIIYMAMSTLFFLRIAPLTPWWGALAAWVMILLIVEAIFAFFFPYRCCFNETEGDVQNNPHV
IRIS_313-10147 MELITACALLLIAVHIAIVISCPEPLAAVALVDVPLLYLCRYLLIIRRGVIIVSGGGGGGRLRRFRLGAAMAIIYMAMSTLFFLRIAPLTPWWGALAAWVMILLIVEAIFAFFFPYRCCFNETEGDVQNNPHV
IRIS_313-10148 MELITACALLLIAVHIAIVISCPEPLAAVALVDVPLLYLCRYLLIIRRGVIIVSGAGAGGGRLRRFRLGAAMAIIYMAMSTLFFLRIAPLTPWWGALAAWVMILLIVEAIFAFFFPYRCCFNETEGDVQNNPHV
IRIS_313-10150 MELITACALLLIAVHIAIVISCPEPLAAVAFVNVPLLYLCRYLLIIRRRVIIVSGGGGGGGRLRRFRLGAAMAVIYMAMSTLFFLRIAPLTPWWGALAAWVMILLIVEAIFAFFFPYRCCFNETEGDVQNNPHV
IRIS_313-10151 MELITACALLLIAVHIAIVISCPEPLAAVALVDVPLLYLCRYLLIIRRGVIIVSGGGGGGRLRRFRLGAAMAIIYMAMSTLFFLRIAPLTPWWGALAAWVMILLIVEAIFAFFFPYRCCFNETEGDVQNNPHV
IRIS_313-10152 MELITACALLLIAVHIAIVISCPEPLAAVALVDVPLLYLCRYLLIIRRGVIIVSGAGAGGGRLRRFRLGAAMAIIYMAMSTLFFLRIAPLTPWWGALAAWVMILLIVEAIFAFFFPYRCCFNETEGDVQNNPHV
IRIS_313-10154 MELITACALLLIAVHIAIVISCPEPLAAVALVDVPLLYLYRYLLIIRRGVIIVSGAGGGGGRLRRFRLGAAMAIIYMAMSTLFFLRIAPLTPWWGALAAWVMILLIVEAIFAFFFPYRCCFNETEGDVQNNPHV
IRIS_313-10158 MELITACALLLIAVHIAIVISCPEPLAAVALVDVPLLYLCRYLLIIRRGVIIVSGAGAGGGRLRRFRLGAAMAIIYMAMSTLFFLRIAPLTPWWGALAAWVMILLIVEAIFAFFFPYRCCFNETEGDVQNNPHV
```

IRIS_313-10161 MELITACALLLIAVHIAIVISCPEPLAAVALVDVPLLYLCRYLLIIRRGVIIVSGGGGGGRLRRFRLGAAMAIIYMAMSTLFFLRIAPLTPWWGALAAWVMILLIVEAIFAFFFPYRCCFNETEGDVQNNPHV
IRIS_313-10164 MELITACALLLIAVHIAIVISCPEPLAAVALVDVPLLYLCRYLLIIRRGVIIVSGAGAGGGRLRRFRLGAAMAIIYMAMSTLFFLRIAPLTPWWGALAAWVMILLIVEAIFAFFFPYRCCFNETEGDVQNNPHV
IRIS_313-10167 MELITACALLLIAVHIAIVISCPEPLAAVALVDVPLLYLYRYLLIIRRGVIIVSGAGGGGGRLRRFRLGAAMAIIYMAMSTLFFLRIAPLTPWWGALAAWVMILLIVEAIFAFFFPYRCCFNETEGDVQNNPHV
IRIS_313-10168 MELITACALLLIAVHIAIVISCPEPLAAVALVDVPLLYLYRYLLIIRRGVIIVSGAGAGGGRLRRFRLGAAMAIIYMAMSTLFFLRIAPLTPWWGALAAWVMILLIVEAIFAFFFPYRCCFNETEGDVQNNPHV
IRIS_313-10170 MELITACALLLIAVHIAIVISCPEPLAAVALVDVPLLYLCRYLLIIRRGVIIVSGAGAGGGRLRRFRLGAAMAIIYMAMSTLFFLRIAPLTPWWGALAAWVMILLIVEAIFAFFFPYRCCFNETEGDVQNNPHV
IRIS_313-10171 MELITACALLLIAVHIAIVISCPEPLAAVALVDVPLLYLCRYLLIIRRGVIIVSGAGAGGGRLRRFRLGAAMAIIYMAMSTLFFLRIAPLTPWWGALAAWVMILLIVEAIFAFFFPYRCCFNETEGDVQNNPHV
IRIS_313-10176 MELITACALLLIAVHIAIVISCPEPLAAVALVDVPLLYLCRYLLIIRRGVIIVSGAGAGGGRLRRFRLGAAMAIIYMAMSTLFFLRIAPLTPWWGALAAWVMILLIVEAIFAFFFPYRCCFNETEGDVQNNPHV
IRIS_313-10177 MELITACALLLIAVHIAIVISCPEPLAAVALVDVPLLYLCRYLLIIRRGVIIVSGGGGGGRLRRFRLGAAMAIIYMAMSTLFFLRIAPLTPWWGALAAWVMILLIVEAIFAFFFPYRCCFNETEGDVQNNPHV
IRIS_313-10178 MELITACALLLIAVHIAIVISCPEPLAAVALVDVPLLYLCRYLLIIRRGVIIVSGAGAGGGRLRRFRLGAAMAIIYMAMSTLFFLRIAPLTPWWGALAAWVMILLIVEAIFAFFFPYRCCFNETEGDVQNNPHV
IRIS_313-10179 MELITACALLLIAVHIAIVISCPEPLAAVALVDVPLLYLCRYLLIIRRGVIIVSGGGGGGRLRRFRLGAAMAIIYMAMSTLFFLRIAPLTPWWGALAAWVMILLIVEAIFAFFFPYRCCFNETEGDVQNNPHV
IRIS_313-10189 MELITACALLLIAVHIAIVISCPEPLAAVALVDVPLLYLYRYLLIIRRGVIIVSGAGGGGGRLRRFRLGAAMAIIYMAMSTLFFLRIAPLTPWWGALAAWVMILLIVEAIFAFFFPYRCCFNETEGDVQNNPHV
IRIS_313-10190 MELITACALLLIAVHIAIVISCPEPLAAVALVDVPLLYLCRYLLIIRRGVIIVSGAGGGGGRLRRFRLGAAMAIIYMAMSTLFFLRIAPLTPWWGALAAWVMILLIVEAIFAFFFPYRCCFNETEGDVQNNPHV
IRIS_313-10191 MELITACALLLIAVHIAIVISCPEPLAAVALVDVPLLYLCRYLLIIRRGVIIVSGAGAGGGRLRRFRLGAAMAIIYMAMSTLFFLRIAPLTPWWGALAAWVMILLIVEAIFAFFFPYRCCFNETEGDVQNNPHV
IRIS_313-10196 MELITACALLLIAVHIAIVISCPEPLAAVALVDVPLLYLCRYLLIIRRGVIIVSGAGGGGGGRLRRFRLGAAMAIIYMAMSTLFFLRIAPLTPWWGALAAWVMILLIVEAIFAFFFPYRCCFNETEGDVQNNPHV
IRIS_313-10211 MELITACALLLIAVHIAIVISCPEPLAAVALVDVPLLYLYRYLLIIRRGVIIVSGAGGGGGRLRRFRLGAAMAIIYMAMSTLFFLRIAPLTPWWGALAAWVMILLIVEAIFAFFFPYRCCFNETEGDVQNNPHV
IRIS_313-10220 MELITACALLLIAVHIAIVISCPEPLAAVALVDVPLLYLCRYLLIIRRGVIIVSGAGAGGGRLRRFRLGAAMAIIYMAMSTLFFLRIAPLTPWWGALAAWVMILLIVEAIFAFFFPYRCCFNETEGDVQNNPHV
IRIS_313-10221 MELITACALLLIAVHIAIVISCPEPLAAVALVDVPLLYLCRYLLIIRRGVIIVSGGGGGGRLRRFRLGAAMAIIYMAMSTLFFLRIAPLTPWWGALAAWVMILLIVEAIFAFFFPYRCCFNETEGDVQNNPHV
IRIS_313-10224 MELITACALLLIAVHIAIVISCPEPLAAVALVDVPLLYLCRYLLIIRRGVIIVSGGGGGGRLRRFRLGAAMAIIYMAMSTLFFLRIAPLTPWWGALAAWVMILLIVEAIFAFFFPYRCCFNETEGDVQNNPHV
IRIS_313-10226 MELITACALLLIAVHIAIVISCPEPLAAVALVDVPLLYLCRYLLIIRRGVIIVSGGGGGGRLRRFRLGAAMAIIYMAMSTLFFLRIAPLTPWWGALAAWVMILLIVEAIFAFFFPYRCCFNETEGDVQNNPHV
IRIS_313-10228 MELITACALLLIAVHIAIVISCPEPLAAVALVDVPLLYLCRYLLIIRRGVIIVSGAGAGGGRLRRFRLGAAMAIIYMAMSTLFFLRIAPLTPWWGALAAWVMILLIVEAIFAFFFPYRCCFNETEGDVQNNPHV
IRIS_313-10234 MELITACALLLIAVHIAIVISCPEPLAAVALVDVPLLYLCRYLLIIRRGVIIVSGGGGGGRLRRFRLGAAMAIIYMAMSTLFFLRIAPLTPWWGALAAWVMILLIVEAIFAFFFPYRCCFNETEGDVQNNPHV
IRIS_313-10235 MELITACALLLIAVHIAIVISCPEPLAAVALVDVPLLYLCRYLLIIRRGVIIVSGGGGGGRLRRFRLGAAMAIIYMAMSTLFFLRIAPLTPWWGALAAWVMILLIVEAIFAFFFPYRCCFNETEGDVQNNPHV
IRIS_313-10237 MELITACALLLIAVHIAIVISCPEPLAAVALVDVPLLYLCRYLLIIRRGVIIVSGGGGGGRLRRFRLGAAMAIIYMAMSTLFFLRIAPLTPWWGALAAWVMILLIVEAIFAFFFPYRCCFNETEGDVQNNPHV
IRIS_313-10238 MELITACALLLIAVHIAIVISCPEPLAAVALVDVPLLYLCRYLLIIRRGVIIVSGGGGGGRLRRFRLGAAMAIIYMAMSTLFFLRIAPLTPWWGALAAWVMILLIVEAIFAFFFPYRCCFNETEGDVQNNPHV
IRIS_313-10239 MELITACALLLIAVHIAIVISCPEPLAAVALVDVPLLYLCRYLLIIRRGVIIVSGAGGAAAAGCAASASGQPWRSSTWPCRPSSSSALLLSRHGGAPSPPGS*SCSSSRPSSPSSSPTAAASTKPKAMSRIIPM
IRIS_313-10242 MELITACALLLIAVHIAIVISCPEPLAAVALVDVPLLYLCRYLLIIRRGVIIVSGAGAGGGRLRRFRLGAAMAIIYMAMSTLFFLRIAPLTPWWGALAAWVMILLIVEAIFAFFFPYRCCFNETEGDVQNNPHV
IRIS_313-10247 MELITACALLLIAVHIAIVISCPEPLAAVALVDVPLLYLCRYLLIIRRGVIIVSGAGAGGGRLRRFRLGAAMAIIYMAMSTLFFLRIAPLTPWWGALAAWVMILLIVEAIFAFFFPYRCCFNETEGDVQNNPHV
IRIS_313-10257 MELITACALLLIAVHIAIVISCPEPLAAVALVDVPLLYLCRYLLIIRRGVIIVSGAGAGGGRLRRFRLGAAMAIIYMAMSTLFFLRIAPLTPWWGALAAWVMILLIVEAIFAFFFPYRCCFNETEGDVQNNPHV
IRIS_313-10258 MELITACALLLIAVHIAIVISCPEPLAAVALVDVPLLYLCRYLLIIRRGVIIVSGAGAGGGRLRRFRLGAAMAIIYMAMSTLFFLRIAPLTPWWGALAAWVMILLIVEAIFAFFFPYRCCFNETEGDVQNNPHV
IRIS_313-10260 MELITACALLLIAVHIAIVISCPEPLAAVALVDVPLLYLCRYLLIIRRGVIIVSGAGAGGGRLRRFRLGAAMAIIYMAMSTLFFLRIAPLTPWWGALAAWVMILLIVEAIFAFFFPYRCCFNETEGDVQNNPHV
IRIS_313-10263 MELITACALLLIAVHIAIVISCPEPLAAVALVDVPLLYLCRYLLIIRRGVIIVSGAGGGGGGRLRRFRLGAAMAIIYMAMSTLFFLRIAPLTPWWGALAAWVMILLIVEAIFAFFFPYRCCFNETEGDVQNNPHV
IRIS_313-10268 MELITACALLLIAVHIAIVISCPEPLAAVALVDVPLLYLCRYLLIIRRGVIIVSGAGAGGGRLRRFRLGAAMAIIYMAMSTLFFLRIAPLTPWWGALAAWVMILLIVEAIFAFFFPYRCCFNETEGDVQNNPHV
IRIS_313-10271 MELITACALLLIAVHIAIVISCPEPLAAVALVDVPLLYLCRYLLIIRRGVIIVSGAGAGGGRLRRFRLGAAMAIIYMVMSTLFFLRIAPLTPWWGALAAWVMILLIVEAIFAFFFPYRCCFNETEGDVQNNPHV
IRIS_313-10272 MELITACALLLIAVHIAIVISCPEPLAAVALVDVPLLYLCRYLLIIRRGVIIVSGAGAGGGRLRRFRLGAAMAIIYMAMSTLFFLRIAPLTPWWGALAAWVMILLIVEAIFAFFFPYRCCFNETEGDVQNNPHV
IRIS_313-10274 MELITACALLLIAVHIAIVISCPEPLAAVALVDVPLLYLCRYLLIIRRGVIIVSGGGAGGGRLRRFRLGAAMAIIYMAMSTLFFLRIAPLTPWWGALAAWVMILLIVEAIFAFFFPYRCCFNETEGDVQNNPHV
IRIS_313-10275 MELITACALLLIAVHIAIVISCPEPLAAVALVDVPLLYLCRYLLIIRRGVIIVSGAGAGGGRLRRFRLGAAMAIIYMAMSTLFFLRIAPLTPWWGALAAWVMILLIVEAIFAFFFPYRCCFNETEGDVQNNPHV
IRIS_313-10279 MELITACALLLIAVHIAIVISCPEPLAAVALVDVPLLYLCRYLLIIRRGVIIVSGGGGGGRLRRFRLGAAMAIIYMAMSTLFFLRIAPLTPWWGALAAWVMILLIVEAIFAFFFPYRCCFNETEGDVQNNPHV
IRIS_313-10285 MELITACALLLIAVHIAIVISCPEPLAAVALVDVPLLYLCRYLLIIRRGVIIVSGGGGGGRLRRFRLGAAMAIIYMAMSTLFFLRIAPLTPWWGALAAWVMILLIVEAIFAFFFPYRCCFNETEGDVQNNPHV
IRIS_313-10287 MELITACALLLIAVHIAIVISCPEPLAAVALVDVPLLYLYRYLLIIRRGVIIVSGAGGGGGRLRRFRLGAAMAIIYMAMSTLFFLRIAPLTPWWGALAAWVMILLIVEAIFAFFFPYRCCFNETEGDVQNNPHV
IRIS_313-10289 MELITACALLLIAVHIAIVISCPEPLAAVALVDVPLLYLCRYLLIIRRGVIIVSGGGGGGRLRRFRLGAAMAIIYMAMSTLFFLRIAPLTPWWGALAAWVMILLIVEAIFAFFFPYRCCFNETEGDVQNNPHV
IRIS_313-10290 MELITACALLLIAVHIAIVISCPEPLAAVALVDVPLLYLCRYLLIIRRGVIIVSGGGGGGRLRRFRLGAAMAIIYMAMSTLFFLRIAPLTPWWGALAAWVMILLIVEAIFAFFFPYRCCFNETEGDVQNNPHV
IRIS_313-10293 MELITACALLLIAVHIAIVISCPEPLAAVALVDVPLLYLCRYLLIIRRGVIIVSGAGAGGGRLRRFRLGAAMAIIYMAMSTLFFLRIAPLTPWWGALAAWVMILLIVEAIFAFFFPYRCCFNETEGDVQNNPHV
IRIS_313-10294 MELITACALLLIAVHIAIVISCPEPLAAVALVDVPLLYLYRYLLIIRRGVIIVSGAGGGGGRLRRFRLGAAMAIIYMAMSTLFFLRIAPLTPWWGALAAWVMILLIVEAIFAFFFPYRCCFNETEGDVQNNPHV
IRIS_313-10295 MELITACALLLIAVHIAIVISCPEPLAAVALVDVPLLYLCRYLLIIRRGVIIVSGAGAGGGRLRRFRLGAAMAIIYMAMSTLFFLRIAPLTPWWGALAAWVMILLIVEAIFAFFFPYRCCFNETEGDVQNNPHV
IRIS_313-10298 MELITACALLLIAVHIAIVISCPEPLAAVALVDVPLLYLCRYLLIIRRGVIIVSGAGAGGGRLRRFRLGAAMAIIYMAMSTLFFLRIAPLTPWWGALAAWVMILLIVEAIFAFFFPYRCCFNETEGDVQNNPHV
IRIS_313-10300 MELITACALLLIAVHIAIVISCPEPLAAVALVDVPLLYLCRYLLIIRRGVIIVSGGGGGGRLRRFRLGAAMAIIYMAMSTLFFLRIAPLTPWWGALAAWVMILLIVEAIFAFFFPYRCCFNETEGDVQNNPHV
IRIS_313-10301 MELITACALLLIAVHIAIVISCPEPLAAVALVDVPLLYLCRYLLIIRRGVIIVSGAGAGGGRLRRFRLGAAMAIIYMAMSTLFFLRIAPLTPWWGALAAWVMILLIVEAIFAFFFPYRCCFNETEGDVQNNPHV
IRIS_313-10307 MELITACALLLIAVHIAIVISCPEPLAAVALVDVPLLYLCRYLLIIRRGVIIVSGGGGGGRLRRFRLGAAMAIIYMAMSTLFFLRIAPLTPWWGALAAWVMILLIVEAIFAFFFPYRCCFNETEGDVQNNPHV
IRIS_313-10314 MELITACALLLIAVHIAIVISCPEPLAAVALVDVPLLYLCRYLLIIRRGVIIVSGGGGGGRLRRFRLGAAMAIIYMAMSTLFFLRIAPLTPWWGALAAWVMILLIVEAIFAFFFPYRCCFNETEGDVQNNPHV
IRIS_313-10318 MELITACALLLIAVHIAIVISCPEPLAAVALVDVPLLYLCRYLLIIRRGVIIVSGAGAGGGRLRRFRLGAAMAIIYMAMSTLFFLRIAPLTPWWGALAAWVMILLIVEAIFAFFFPYRCCFNETEGDVQNNPHV
IRIS_313-10325 MELITACALLLIAVHIAIVISCPEPLAAVALVDVPLLYLCRYLLIIRRGVIIVSGAGAGGGRLRRFRLGAAMAIIYMAMSTLFFLRIAPLTPWWGALAAWVMILLIVEAIFAFFFPYRCCFNETEGDVQNNPHV
IRIS_313-10327 MELITACALLLIAVHIAIVISCPEPLAAVALVDVPLLYLCRYLLIIRRGVIIVSGAGAGGGRLRRFRLGAAMAIIYMAMSTLFFLRIAPLTPWWGALAAWVMILLIVEAIFAFFFPYRCCFNETEGDVQNNPHV
IRIS_313-10332 MELITACALLLIAVHIAIVISCPEPLAAVALVDVPLLYLCRYLLIIRRGVIIVSGGGGGGRLRRFRLGAAMAIIYMAMSTLFFLRIAPLTPWWGALAAWVMILLIVEAIFAFFFPYRCCFNETEGDVQNNPHV
IRIS_313-10333 MELITACALLLIAVHIAIVISCPEPLAAVALVDVPLLYLCRYLLIIRRGVIIVSGGGGGGRLRRFRLGAAMAIIYMAMSTLFFLRIAPLTPWWGALAAWVMILLIVEAIFAFFFPYRCCFNETEGDVQNNPHV
IRIS_313-10334 MELITACALLLIAVHIAIVISCPEPLAAVALVDVPLLYLCRYLLIIRRGVIIVSGGGGGGRLRRFRLGAAMAIIYMAMSTLFFLRIAPLTPWWGALAAWVMILLIVEAIFAFFFPYRCCFNETEGDVQNNPHV
IRIS_313-10336 MELITACALLLIAVHIAIVISCPEPLAAVALVDVPLLYLCRYLLIIRRGVIIVSGAGAGGGRLRRFRLGAAMAIIYMAMSTLFFLRIAPLTPWWGALAAWVMILLIVEAIFAFFFPYRCCFNETEGDVQNNPHV
IRIS_313-10337 MELITACALLLIAVHIAIVISCPEPLAAVALVDVPLLYLCRYLLIIRRGVIIVSGAGAGGGRLRRFRLGAAMAIIYMAMSTLFFLRIAPLTPWWGALAAWVMILLIVEAIFAFFFPYRCCFNETEGDVQNNPHV
IRIS_313-10340 MELITACALLLIAVHIAIVISCPEPLAAVALVDVPLLYLCRYLLIIRRGVIIVSGGGGGGRLRRFRLGAAMAIIYMAMSTLFFLRIAPLTPWWGALAAWVMILLIVEAIFAFFFPYRCCFNETEGDVQNNPHV
IRIS_313-10341 MELITACALLLIAVHIAIVISCPEPLAAVALVDVPLLYLCRYLLIIRRGVIIVSGGAGGGRLRRFRLGAAMAIIYMAMSTLFFLRIAPLTPWWGALAAWVMILLIVEAIFAFFFPYRCCFNETEGDVQNNPHV
IRIS_313-10348 MELITACALLLIAVHIAIVISCPEPLAAVALVDVPLLYLCRYLLIIRRGVIIVSGGGGGGRLRRFRLGAAMAIIYMAMSTLFFLRIAPLTPWWGALAAWVMILLIVEAIFAFFFPYRCCFNETEGDVQNNPHV
IRIS_313-10349 MELITACALLLIAVHIAIVISCPEPLAAVALVDVPLLYLCRYLLIIRRGVIIVSGAGAGGGRLRRFRLGAAMAIIYMAMSTLFFLRIAPLTPWWGALAAWVMILLIVEAIFAFFFPYRCCFNETEGDVQNNPHV
IRIS_313-10352 MELITACALLLIAVHIAIVISCPEPLAAVALVDVPLLYLCRYLLIIRRGVIIVSGAGAGGGRLRRFRLGAAMAIIYMAMSTLFFLRIAPLTPWWGALAAWVMILLIVEAIFAFFFPYRCCFNETEGDVQNNPHV
IRIS_313-10353 MELITACALLLIAVHIAIVISCPEPLAAVALVDVPLLYLCRYLLIIRRGVIIVSGGGGGGRLRRFRLGAAMAIIYMAMSTLFFLRIAPLTPWWGALAAWVMILLIVEAIFAFFFPYRCCFNETEGDVQNNPHV
IRIS_313-10355 MELITACALLLIAVHIAIVISCPEPLAAVALVDVPLLYLCRYLLIIRRGVIIVSGAGAGGGRLRRFRLGAAMAIIYMAMSTLFFLRIAPLTPWWGALAAWVMILLIVEAIFAFFFPYRCCFNETEGDVQNNPHV
IRIS_313-10357 MELITACALLLIAVHIAIVISCPEPLAAVALVDVPLLYLCRYLLIIRRGVIIVSGGGGGGRLRRFRLGAAMAIIYMAMSTLFFLRIAPLTPWWGALAAWVMILLIVEAIFAFFFPYRCCFNETEGDVQNNPHV
IRIS_313-10359 MELITACALLLIAVHIAIVISCPEPLAAVALVDVPLLYLCRYLLIIRRGVIIVSGAGAGGGRLRRFRLGAAMAIIYMAMSTLFFLRIAPLTPWWGALAAWVMILLIVEAIFAFFFPYRCCFNETEGDVQNNPHV
IRIS_313-10360 MELITACALLLIAVHIAIVISCPEPLAAVALVDVPLLYLCRYLLIIRRGVIIVSGGGGGGRLRRFRLGAAMAIIYMAMSTLFFLRIAPLTPWWGALAAWVMILLIVEAIFAFFFPYRCCFNETEGDVQNNPHV
IRIS_313-10361 MELITACALLLIAVHIAIVISCPEPLAAVALVDVPLLYLCRYLLIIRRGVIIVSGAGGAAAAGCAASASGQPWRSSTWPCRPSSSSALLLSRHGGAPSPPGS*SCSSSRPSSPSSSPTAAASTKPKAMSRIIPM
IRIS_313-10366 MELITACALLLIAVHIAIVISCPEPLAAVALVDVPLLYLYRYLLIIRRGVIIVSGAGGGGGRLRRFRLGAAMAIIYMAMSTLFFLRIAPLTPWWGALAAWVMILLIVEAIFAFFFPYRCCFNETEGDVQNNPHV
IRIS_313-10371 MELITACALLLIAVHIAIVISCPEPLAAVALVDVPLLYLCRYLLIIRRGVIIVSGAGAGGGRLRRFRLGAAMAIIYMAMSTLFFLRIAPLTPWWGALAAWVMILLIVEAIFAFFFPYRCCFNETEGDVQNNPHV
IRIS_313-10373 MELITACALLLIAVHIAIVISCPEPLAAVALVDVPLLYLCRYLLIIRRGVIIVSGAGAGGGRLRRFRLGAAMAIIYMAMSTLFFLRIAPLTPWWGALAAWVMILLIVEAIFAFFFPYRCCFNETEGDVQNNPHV
IRIS_313-10374 MELITACALLLIAVHIAIVISCPEPLAAVALVDVPLLYLCRYLLIIRRGVIIVSGGGAGGGRLRRFRLGAAMAIIYMAMSTLFFLRIAPLTPWWGALAAWVMILLIVEAIFAFFFPYRCCFNETEGDVQNNPHV
IRIS_313-10375 MELITACALLLIAVHIAIVISCPEPLAAVALVDVPLLYLCRYLLIIRRGVIIVSGGGAGGGRLRRFRLGAAMAIIYMAMSTLFFLRIAPLTPWWGALAAWVMILLIVEAIFAFFFPYRCCFNETEGDVQNNPHV
IRIS_313-10379 MELITACALLLIAVHIAIVISCPEPLAAVALVDVPLLYLCRYLLIIRRGVIIVSGAGAGGGRLRRFRLGAAMAIIYMAMSTLFFLRIAPLTPWWGALAAWVMILLIVEAIFAFFFPYRCCFNETEGDVQNNPHV
IRIS_313-10380 MELITACALLLIAVHIAIVISCPEPLAAVALVDVPLLYLCRYLLIIRRGVIIVSGAGAGGGRLRRFRLGAAMAIIYMAMSTLFFLRIAPLTPWWGALAAWVMILLIVEAIFAFFFPYRCCFNETEGDVQNNPHV
IRIS_313-10385 MELITACALLLIAVHIAIVISCPEPLAAVALVDVPLLYLCRYLLIIRRGVIIVSGAGAGGGRLRRFRLGAAMAIIYMAMSTLFFLRIAPLTPWWGALAAWVMILLIVEAIFAFFFPYRCCFNETEGDVQNNPHV
IRIS_313-10392 MELITACALLLIAVHIAIVISCPEPLAAVALVDVPLLYLCRYLLIIRRGVIIVSGGGGGGRLRRFRLGAAMAIIYMAMSTLFFLRIAPLTPWWGALAAWVMILLIVEAIFAFFFPYRCCFNETEGDVQNNPHV
IRIS_313-10394 MELITACALLLIAVHIAIVISCPEPLAAVALVDVPLLYLCRYLLIIRRGVIIVSGGGGGGRLRRFRLGAAMAIIYMAMSTLFFLRIAPLTPWWGALAAWVMILLIVEAIFAFFFPYRCCFNETEGDVQNNPHV
IRIS_313-10396 MELITACALLLIAVHIAIVISCPEPLAAVALVDVPLLYLCRYLLIIRRGVIIVSGGGGGGRLRRFRLGAAMAIIYMAMSTLFFLRIAPLTPWWGALAAWVMILLIVEAIFAFFFPYRCCFNETEGDVQNNPHV
IRIS_313-10397 MELITACALLLIAVHIAIVISCPEPLAAVALVDVPLLYLCRYLLIIRRGVIIVSGGGGGGRLRRFRLGAAMAIIYMAMSTLFFLRIAPLTPWWGALAAWVMILLIVEAIFAFFFPYRCCFNETEGDVQNNPHV
IRIS_313-10398 MELITACALLLIAVHIAIVISCPEPLAAVALVDVPLLYLCRYLLIIRRGVIIVSGGGGGGRLRRFRLGAAMAIIYMAMSTLFFLRIAPLTPWWGALAAWVMILLIVEAIFAFFFPYRCCFNETEGDVQNNPHV
IRIS_313-10399 MELITACALLLIAVHIAIVISCPEPLAAVALVDVPLLYLCRYLLIIRRGVIIVSGAGAGGGRLRRFRLGAAMAIIYMAMSTLFFLRIAPLTPWWGALAAWVMILLIVEAIFAFFFPYRCCFNETEGDVQNNPHV
IRIS_313-10400 MELITACALLLIAVHIAIVISCPEPLAAVALVDVPLLYLCRYLLIIRRGVIIVSGGGAGGGRLRRFRLGAAMAIIYMAMSTLFFLRIAPLTPWWGALAAWVMILLIVEAIFAFFFPYRCCFNETEGDVQNNPHV
IRIS_313-10401 MELITACALLLIAVHIAIVISCPEPLAAVALVDVPLLYLCRYLLIIRRGVIIVSGGGAGGGRLRRFRLGAAMAIIYMAMSTLFFLRIAPLTPWWGALAAWVMILLIVEAIFAFFFPYRCCFNETEGDVQNNPHV
IRIS_313-10402 MELITACALLLIAVHIAIVISCPEPLAAVALVDVPLLYLCRYLLIIRRGVIIVSGGGGGGRLRRFRLGAAMAIIYMAMSTLFFLRIAPLTPWWGALAAWVMILLIVEAIFAFFFPYRCCFNETEGDVQNNPHV
IRIS_313-10403 MELITACALLLIAVHIAIVISCPEPLAAVALVDVPLLYLCRYLLIIRRGVIIVSGGGAGGGRLRRFRLGAAMAIIYMAMSTLFFLRIAPLTPWWGALAAWVMILLIVEAIFAFFFPYRCCFNETEGDVQNNPHV
IRIS_313-10404 MELITACALLLIAVHIAIVISCPEPLAAVALVDVPLLYLCRYLLIIRRGVIIVSGAGGAAAAGCAASASGQPWRSSTWPCRPSSSSALLLSRHGGAPSPPGS*SCSSSRPSSPSSSPTAAASTKPKAMSRIIPM
IRIS_313-10412 MELITACALLLIAVHIAIVISCPEPLAAVALVDVPLLYLCRYLLIIRRGVIIVSGAGAGGGRLRRFRLGAAMAIIYMAMSTLFFLRIAPLTPWWGALAAWVMILLIVEAIFAFFFPYRCCFNETEGDVQNNPHV
IRIS_313-10417 MELITACALLLIAVHIAIVISCPEPLAAVALVDVPLLYLCRYLLIIRRGVIIVSGGGAGGGRLRRFRLGAAMAIIYMAMSTLFFLRIAPLTPWWGALAAWVMILLIVEAIFAFFFPYRCCFNETEGDVQNNPHV
IRIS_313-10421 MELITACALLLIAVHIAIVISCPEPLAAVALVDVPLLYLCRYLLIIRRGVIIVSGAGAGGGRLRRFRLGAAMAIIYMAMSTLFFLRIAPLTPWWGALAAWVMILLIVEAIFAFFFPYRCCFNETEGDVQNNPHV
IRIS_313-10422 MELITACALLLIAVHIAIVISCPEPLAAVALVDVPLLYLCRYLLIIRRGVIIVSGAGAGGGRLRRFRLGAAMAIIYMAMSTLFFLRIAPLTPWWGALAAWVMILLIVEAIFAFFFPYRCCFNETEGDVQNNPHV
IRIS_313-10423 MELITACALLLIAVHIAIVISCPEPLAAVALVDVPLLYLCRYLLIIRRGVIIVSGAGAGGGRLRRFRLGAAMAIIYMAMSTLFFLRIAPLTPWWGALAAWVMILLIVEAIFAFFFPYRCCFNETEGDVQNNPHV
IRIS_313-10428 MELITACALLLIAVHIAIVISCPEPLAAVALVDVPLLYLCRYLLIIRRGVIIVSGAGGAAAAGCAASASGQPWRSSTWPCRPSSSSALLLSRHGGAPSPPGS*SCSSSRPSSPSSSPTAAASTKPKAMSRIIPM
IRIS_313-10429 MELITACALLLIAVHIAIVISCPEPLAAVALVDVPLLYLYRYLLIIRRGVIIVSGAGGGGGRLRRFRLGAAMAIIYMAMSTLFFLRIAPLTPWWGALAAWVMILLIVEAIFAFFFPYRCCFNETEGDVQNNPHV
IRIS_313-10430 MELITACALLLIAVHIAIVISCPEPLAAVALVDVPLLYLCRYLLIIRRGVIIVSGAGAGGGRLRRFRLGAAMAIIYMAMSTLFFLRIAPLTPWWGALAAWVMILLIVEAIFAFFFPYRCCFNETEGDVQNNPHV
IRIS_313-10433 MELITACALLLIAVHIAIVISCPEPLAAVALVDVPLLYLCRYLLIIRRGVIIVSGAGAGGGRLRRFRLGAAMAIIYMAMSTLFFLRIAPLTPWWGALAAWVMILLIVEAIFAFFFPYRCCFNETEGDVQNNPHV
IRIS_313-10437 MELITACALLLIAVHIAIVISCPEPLAAVALVDVPLLYLCRYLLIIRRGVIIVSGAGAGGGRLRRFRLGAAMAIIYMAMSTLFFLRIAPLTPWWGALAAWVMILLIVEAIFAFFFPYRCCFNETEGDVQNNPHV
IRIS_313-10440 MELITACALLLIAVHIAIVISCPEPLAAVALVDVPLLYLCRYLLIIRRGVIIVSGAGAGGGRLRRFRLGAAMAIIYMAMSTLFFLRIAPLTPWWGALAAWVMILLIVEAIFAFFFPYRCCFNETEGDVQNNPHV
IRIS_313-10441 MELITACALLLIAVHIAIVISCPEPLAAVALVDVPLLYLCRYLLIIRRGVIIVSGGGGGGRLRRFRLGAAMAIIYMAMSTLFFLRIAPLTPWWGALAAWVMILLIVEAIFAFFFPYRCCFNETEGDVQNNPHV
IRIS_313-10444 MELITACALLLIAVHIAIVISCPEPLAAVALVDVPLLYLYRYLLIIRRGVIIVSGAGGGGGRLRRFRLGAAMAIIYMAMSTLFFLRIAPLTPWWGALAAWVMILLIVEAIFAFFFPYRCCFNETEGDVQNNPHV
IRIS_313-10448 MELITACALLLIAVHIAIVISCPEPLAAVALVDVPLLYLCRYLLIIRRGVIIVSGGGGAAAAGCAASASGQPWRSSTWPCRPSSSSALLLSRHGGAPSPPGS*SCSSSRPSSPSSSPTAAASTKPKAMSRIIPM
IRIS_313-10449 MELITACALLLIAVHIAIVISCPEPLAAVALVDVPLLYLCRYLLIIRRGVIIVSGGGGGGRLRRFRLGAAMAIIYMAMSTLFFLRIAPLTPWWGALAAWVMILLIVEAIFAFFFPYRCCFNETEGDVQNNPHV
IRIS_313-10450 MELITACALLLIAVHIAIVISCPEPLAAVALVDVPLLYLYRYLLIIRRGVIIVSGAGGGGGRLRRFRLGAAMAIIYMAMSTLFFLRIAPLTPWWGALAAWVMILLIVEAIFAFFFPYRCCFNETEGDVQNNPHV
IRIS_313-10452 MELITACALLLIAVHIAIVISCPEPLAAVALVDVPLLYLCRYLLIIRRGVIIVSGAGGGGGRLRRFRLGAAMAIIYMAMSTLFFLRIAPLTPWWGALAAWVMILLIVEAIFAFFFPYRCCFNETEGDVQNNPHV
IRIS_313-10453 MELITACALLLIAVHIAIVISCPEPLAAVALVDVPLLYLYRYLLIIRRGVIIVSGAGGGGGRLRRFRLGAAMAIIYMAMSTLFFLRIAPLTPWWGALAAWVMILLIVEAIFAFFFPYRCCFNETEGDVQNNPHV
IRIS_313-10458 MELITACALLLIAVHIAIVISCPEPLAAVALVDVPLLYLCRYLLIIRRGVIIVSGGGGGGRLRRFRLGAAMAIIYMAMSTLFFLRIAPLTPWWGALAAWVMILLIVEAIFAFFFPYRCCFNETEGDVQNNPHV
IRIS_313-10459 MELITACALLLIAVHIAIVISCPEPLAAVALVDVPLLYLYRYLLIIRRGVIIVSGGGGGGRLRRFRLGAAMAIIYMAMSTLFFLRIAPLTPWWGALAAWVMILLIVEAIFAFFFPYRCCFNETEGDVQNNPHV
IRIS_313-10469 MELITACALLLIAVHIAIVISCPEPLAAVALVDVPLLYLCRYLLIIRRGVIIVSGAGAGGGRLRRFRLGAAMAIIYMAMSTLFFLRIAPLTPWWGALAAWVMILLIVEAIFAFFFPYRCCFNETEGDVQNNPHV
IRIS_313-10476 MELITACALLLIAVHIAIVISCPEPLAAVALVDVPLLYLCRYLLIIRRGVIIVSGGGGGGRLRRFRLGAAMAIIYMAMSTLFFLRIAPLTPWWGALAAWVMILLIVEAIFAFFFPYRCCFNETEGDVQNNPHV
IRIS_313-10477 MELITACALLLIAVHIAIVISCPEPLAAVALVDVPLLYLCRYLLIIRRGVIIVSGGGGGGRLRRFRLGAAMAIIYMAMSTLFFLRIAPLTPWWGALAAWVMILLIVEAIFAFFFPYRCCFNETEGDVQNNPHV
IRIS_313-10480 MELITACALLLIAVHIAIVISCPEPLAAVALVDVPLLYLCRYLLIIRRGVIIVSGGGGGGRLRRFRLGAAMAIIYMAMSTLFFLRIAPLTPWWGALAAWVMILLIVEAIFAFFFPYRCCFNETEGDVQNNPHV
IRIS_313-10484 MELITACALLLIAVHIAIVISCPEPLAAVALVDVPLLYLCRYLLIIRRGVIIVSGGGGGGRLRRFRLGAAMAIIYMAMSTLFFLRIAPLTPWWGALAAWVMILLIVEAIFAFFFPYRCCFNETEGDVQNNPHV
IRIS_313-10485 MELITACALLLIAVHIAIVISCPEPLAAVALVDVPLLYLYRYLLIIRRGVIIVSGAGGGGGRLRRFRLGAAMAIIYMAMSTLFFLRIAPLTPWWGALAAWVMILLIVEAIFAFFFPYRCCFNETEGDVQNNPHV
IRIS_313-10489 MELITACALLLIAVHIAIVISCPEPLAAVALVDVPLLYLCRYLLIIRRGVIIVSGAGGGGGRLRRFRLGAAMAIIYMAMSTLFFLRIAPLTPWWGALAAWVMILLIVEAIFAFFFPYRCCFNETEGDVQNNPHV
IRIS_313-10497 MELITACALLLIAVHIAIVISCPEPLAAVALVDVPLLYLCRYLLIIRRGVIIVSGAGAGGGRLRRFRLGAAMAIIYMAMSTLFFLRIAPLTPWWGALAAWVMILLIVEAIFAFFFPYRCCFNETEGDVQNNPHV
IRIS_313-10502 MELITACALLLIAVHIAIVISCPEPLAAVALVDVPLLYLCRYLLIIRRGVIIVSGAGGGGRLRRFRLGAAMAIIYMAMSTLFFLRIAPLTPWWGALAAWVMILLIVEAIFAFFFPYRCCFNETEGDVQNNPHV
IRIS_313-10503 MELITACALLLIAVHIAIVISCPEPLAAVALVDVPLLYLYRYLLIIRRGVIIVSGAGGGGGRLRRFRLGAAMAIIYMAMSTLFFLRIAPLTPWWGALAAWVMILLIVEAIFAFFFPYRCCFNETEGDVQNNPHV
IRIS_313-10504 MELITACALLLIAVHIAIVISCPEPLAAVALVDVPLLYLCRYLLIIRRGVIIVSGGGGGGRLRRFRLGAAMAIIYMAMSTLFFLRIAPLTPWWGALAAWVMILLIVEAIFAFFFPYRCCFNETEGDVQNNPHV
IRIS_313-10506 MELITACALLLIAVHIAIVISCPEPLAAVALVDVPLLYLCRYLLIIRRGVIIVSGGGAGGGRLRRFRLGAAMAIIYMAMSTLFFLRIAPLTPWWGALAAWVMILLIVEAIFAFFFPYRCCFNETEGDVQNNPHV
IRIS_313-10507 MELITACALLLIAVHIAIVISCPEPLAAVALVDVPLLYLYRYLLIIRRGVIIVSGAGGGGGRLRRFRLGAAMAIIYMAMSTLFFLRIAPLTPWWGALAAWVMILLIVEAIFAFFFPYRCCFNETEGDVQNNPHV
IRIS_313-10509 MELITACALLLIAVHIAIVISCPEPLAAVALVDVPLLYLCRYLLIIRRGVIIVSGGGGGGRLRRFRLGAAMAIIYMAMSTLFFLRIAPLTPWWGALAAWVMILLIVEAIFAFFFPYRCCFNETEGDVQNNPHV
IRIS_313-10510 MELITACALLLIAVHIAIVISCPEPLAAVALVDVPLLYLCRYLLIIRRGVIIVSGGGGGGRLRRFRLGAAMAIIYMAMSTLFFLRIAPLTPWWGALAAWVMILLIVEAIFAFFFPYRCCFNETEGDVQNNPHV
IRIS_313-10511 MELITACALLLIAVHIAIVISCPEPLAAVALVDVPLLYLCRYLLIIRRGVIIVSGGGGGGRLRRFRLGAAMAIIYMAMSTLFFLRIAPLTPWWGALAAWVMILLIVEAIFAFFFPYRCCFNETEGDVQNNPHV

```
IRIS_313-10513 MELITACALLLIAVHIAIVISCPEPLAAVALVDVPLLYLCRYLLIIRRGVIIVSGAGAAAAAGCAASASGQPWRSSTWPCRPSSSSALLLSRHGGAPSPPGS*SCSSSRPSSPSSSPTAAASTKPKAMSRIIPM
IRIS_313-10514 MELITACALLLIAVHIAIVISCPEPLAAVALVDVPLLYLCRYLLIIRRGVIIVSGAGAGGGRLRRFRLGAAMAIIYMAMSTLFFLRIAPLTPWWGALAAWVMILLIVEAIFAFFFPYRCCFNETEGDVQNNPHV
IRIS_313-10515 MELITACALLLIAVHIAIVISCPEPLAAVALVDVPLLYLYRYLLIIRRGVIIVSGAGGGGGRLRRFRLGAAMAIIYMAMSTLFFLRIAPLTPWWGALAAWVMILLIVEAIFAFFFPYRCCFNETEGDVQNNPHV
IRIS_313-10516 MELITACALLLIAVHIAIVISCPEPLAAVALVDVPLLYLYRYLLIIRRGVIIVSGAGGGGGRLRRFRLGAAMAIIYMAMSTLFFLRIAPLTPWWGALAAWVMILLIVEAIFAFFFPYRCCFNETEGDVQNNPHV
IRIS_313-10517 MELITACALLLIAVHIAIVISCPEPLAAVALVDVPLLYLYRYLLIIRRGVIIVSGAGGGGGRLRRFRLGAAMAIIYMAMSTLFFLRIAPLTPWWGALAAWVMILLIVEAIFAFFFPYRCCFNETEGDVQNNPHV
IRIS_313-10518 MELITACALLLIAVHIAIVISCPEPLAAVALVDVPLLYLCRYLLIIRRGVIIVSGGGGGGRLRRFRLGAAMAIIYMAMSTLFFLRIAPLTPWWGALAAWVMILLIVEAIFAFFFPYRCCFNETEGDVQNNPHV
IRIS_313-10519 MELITACALLLIAVHIAIVISCPEPLAAVALVDVPLLYLCRYLLIIRRGVIIVSGAGAGGGRLRRFRLGAAMAIIYMAMSTLFFLRIAPLTPWWGALAAWVMILLIVEAIFAFFFPYRCCFNETEGDVQNNPHV
IRIS_313-10520 MELITACALLLIAVHIAIVISCPEPLAAVALVDVPLLYLCRYLLIIRRGVIIVSGGGGGGRLRRFRLGAAMAIIYMAMSTLFFLRIAPLTPWWGALAAWVMILLIVEAIFAFFFPYRCCFNETEGDVQNNPHV
IRIS_313-10521 MELITACALLLIAVHIAIVISCPEPLAAVALVDVPLLYLCRYLLIIRRGVIIVSGAGAGGGRLRRFRLGAAMAIIYMAMSTLFFLRIAPLTPWWGALAAWVMILLIVEAIFAFFFPYRCCFNETEGDVQNNPHV
IRIS_313-10522 MELITACALLLIAVHIAIVISCPEPLAAVALVDVPLLYLCRYLLIIRRGVIIVSGAGGGGGRLRRFRLGAAMAIIYMAMSTLFFLRIAPLTPWWGALAAWVMILLIVEAIFAFFFPYRCCFNETEGDVQNNPHV
IRIS_313-10523 MELITACALLLIAVHIAIVISCPEPLAAVALVDVPLLYLCRYLLIIRRGVIIVSGAGAGGGRLRRFRLGAAMAIIYMAMSTLFFLRIAPLTPWWGALAAWVMILLIVEAIFAFFFPYRCCFNETEGDVQNNPHV
IRIS_313-10524 MELITACALLLIAVHIAIVISCPEPLAAVALVDVPLLYLCRYLLIIRRGVIIVSGAGAGGGRLRRFRLGAAMAIIYMVMSTLFFLRIAPLTPWWGALAAWVMILLIVEAIFAFFFPYRCCFNETEGDVQNNPHV
IRIS_313-10525 MELITACALLLIAVHIAIVISCPEPLAAVALVDVPLLYLCRYLLIIRRGVIIVSGGGGGGRLRRFRLGAAMAIIYMAMSTLFFLRIAPLTPWWGALAAWVMILLIVEAIFAFFFPYRCCFNETEGDVQNNPHV
IRIS_313-10526 MELITACALLLIAVHIAIVISCPEPLAAVALVDVPLLYLCRYLLIIRRGVIIVSGAGGGGGGRLRRFRLGAAMAIIYMAMSTLFFLRIAPLTPWWGALAAWVMILLIVEAIFAFFFPYRCCFNETEGDVQNNPHV
IRIS_313-10527 MELITACALLLIAVHIAIVISCPEPLAAVALVDVPLLYLCRYLLIIRRGVIIVSGGGGGGRLRRFRLGAAMAIIYMAMSTLFFLRIAPLTPWWGALAAWVMILLIVEAIFAFFFPYRCCFNETEGDVQNNPHV
IRIS_313-10531 MELITACALLLIAVHIAIVISCPEPLAAVALVDVPLLYLCRYLLIIRRGVIIVSGAGAGGGRLRRFRLGAAMAIIYMAMSTLFFLRIAPLTPWWGALAAWVMILLIVEAIFAFFFPYRCCFNETEGDVQNNPHV
IRIS_313-10534 MELITACALLLIAVHIAIVISCPEPLAAVALVDVPLLYLCRYLLIIRRGVIIVSGAGAGGGRLRRFRLGAAMAIIYMAMSTLFFLRIAPLTPWWGALAAWVMILLIVEAIFAFFFPYRCCFNETEGDVQNNPHV
IRIS_313-10536 MELITACALLLIAVHIAIVISCPEPLAAVALVDVPLLYLCRYLLIIRRGVIIVSGAGAGGGRLRRFRLGAAMAIIYMAMSTLFFLRIAPLTPWWGALAAWVMILLIVEAIFAFFFPYRCCFNETEGDVQNNPHV
IRIS_313-10537 MELITACALLLIAVHIAIVISCPEPLAAVALVDVPLLYLCRYLLIIRRGVIIVSGAGAGGGRLRRFRLGAAMAIIYMAMSTLFFLRIAPLTPWWGALAAWVMILLIVEAIFAFFFPYRCCFNETEGDVQNNPHV
IRIS_313-10539 MELITACALLLIAVHIAIVISCPEPLAAVALVDVPLLYLCRYLLIIRRGVIIVSGAGAGGGRLRRFRLGAAMAIIYMAMSTLFFLRIAPLTPWWGALAAWVMILLIVEAIFAFFFPYRCCFNETEGDVQNNPHV
IRIS_313-10541 MELITACALLLIAVHIAIVISCPEPLAAVALVDVPLLYLCRYLLIIRRGVIIVSGAGAGGGRLRRFRLGAAMAIIYMAMSTLFFLRIAPLTPWWGALAAWVMILLIVEAIFAFFFPYRCCFNETEGDVQNNPHV
IRIS_313-10542 MELITACALLLIAVHIAIVISCPEPLAAVALVDVPLLYLCRYLLIIRRGVIIVSGAGGAAAAAGCAASASGQPWRSSTWPCRPSSSSALLLSRHGGAPSPPGS*SCSSSRPSSPSSSPTAAASTKPKAMSRIIPM
IRIS_313-10543 MELITACALLLIAVHIAIVISCPEPLAAVAFVNVPLLYLCRYLLIIRRGVIIVSGGGGGGGRLRRFRLGAAMAVIYMAMSTLFFLRIAPLTPWWGALAAWVMILLIVEAIFAFFFPYRCCFNETEGDVQNNPHV
IRIS_313-10544 MELITACALLLIAVHIAIVISCPEPLAAVALVDVPLLYLCRYLLIIRRGVIIVSGGGGGGRLRRFRLGAAMAIIYMAMSTLFFLRIAPLTPWWGALAAWVMILLIVEAIFAFFFPYRCCFNETEGDVQNNPHV
IRIS_313-10545 MELITACALLLIAVHIAIVISCPEPLAAVALVDVPLLYLCRYLLIIRRGVIIVSGAGAGGGRLRRFRLGAAMAIIYMAMSTLFFLRIAPLTPWWGALAAWVMILLIVEAIFAFFFPYRCCFNETEGDVQNNPHV
IRIS_313-10547 MELITACALLLIAVHIAIVISCPEPLAAVALVDVPLLYLCRYLLIIRRGVIIVSGAGAGGGRLRRFRLGAAMAIIYMAMSTLFFLRIAPLTPWWGALAAWVMILLIVEAIFAFFFPYRCCFNETEGDVQNNPHV
IRIS_313-10549 MELITACALLLIAVHIAIVISCPEPLAAVAFVNVPLLYLCRYLLIIRRGVIIVSGGGGGGGRLRRFRLGAAMAVIYMAMSTLFFLRIAPLTPWWGALAAWVMILLIVEAIFAFFFPYRCCFNETEGDVQNNPHV
IRIS_313-10550 MELITACALLLIAVHIAIVISCPEPLAAVALVDVPLLYLCRYLLIIRRGVIIVSGAGAGGGRLRRFRLGAAMAIIYMAMSTLFFLRIAPLTPWWGALAAWVMILLIVEAIFAFFFPYRCCFNETEGDVQNNPHV
IRIS_313-10552 MELITACALLLIAVHIAIVISCPEPLAAVALVDVPLLYLCRYLLIIRRGVIIVSGAGAGGGRLRRFRLGAAMAIIYMAMSTLFFLRIAPLTPWWGALAAWVMILLIVEAIFAFFFPYRCCFNETEGDVQNNPHV
IRIS_313-10554 MELITACALLLIAVHIAIVISCPEPLAAVALVDVPLLYLYRYLLIIRRGVIIVSGAGGGGGRLRRFRLGAAMAIIYMAMSTLFFLRIAPLTPWWGALAAWVMILLIVEAIFAFFFPYRCCFNETEGDVQNNPHV
IRIS_313-10555 MELITACALLLIAVHIAIVISCPEPLAAVALVDVPLLYLYRYLLIIRRGVIIVSGAGGGGGRLRRFRLGAAMAIIYMAMSTLFFLRIAPLTPWWGALAAWVMILLIVEAIFAFFFPYRCCFNETEGDVQNNPHV
IRIS_313-10556 MELITACALLLIAVHIAIVISCPEPLAAVALVDVPLLYLCRYLLIIRRGVIIVSGGGGGGRLRRFRLGAAMAIIYMAMSTLFFLRIAPLTPWWGALAAWVMILLIVEAIFAFFFPYRCCFNETEGDVQNNPHV
IRIS_313-10557 MELITACALLLIAVHIAIVISCPEPLAAVALVDVPLLYLYRYLLIIRRGVIIVSGAGGGGGRLRRFRLGAAMAIIYMAMSTLFFLRIAPLTPWWGALAAWVMILLIVEAIFAFFFPYRCCFNETEGDVQNNPHV
IRIS_313-10558 MELITACALLLIAVHIAIVISCPEPLAAVALVDVPLLYLCRYLLIIRRGVIIVSGAGAGGGRLRRFRLGAAMAIIYMAMSTLFFLRIAPLTPWWGALAAWVMILLIVEAIFAFFFPYRCCFNETEGDVQNNPHV
IRIS_313-10559 MELITACALLLIAVHIAIVISCPEPLAAVALVDVPLLYLCRYLLIIRRGVIIVSGAGAGGGRLRRFRLGAAMAIIYMAMSTLFFLRIAPLTPWWGALAAWVMILLIVEAIFAFFFPYRCCFNETEGDVQNNPHV
IRIS_313-10560 MELITACALLLIAVHIAIVISCPEPLAAVALVDVPLLYLCRYLLIIRRGVIIVSGGGGGGRLRRFRLGAAMAIIYMAMSTLFFLRIAPLTPWWGALAAWVMILLIVEAIFAFFFPYRCCFNETEGDVQNNPHV
IRIS_313-10561 MELITACALLLIAVHIAIVISCPEPLAAVALVDVPLLYLYRYLLIIRRGVIIVSGAGGGGGRLRRFRLGAAMAIIYMAMSTLFFLRIAPLTPWWGALAAWVMILLIVEAIFAFFFPYRCCFNETEGDVQNNPHV
IRIS_313-10562 MELITACALLLIAVHIAIVISCPEPLAAVALVDVPLLYLCRYLLIIRRGVIIVSGAGGGGGGRLRRFRLGAAMAIIYMAMSTLFFLRIAPLTPWWGALAAWVMILLIVEAIFAFFFPYRCCFNETEGDVQNNPHV
IRIS_313-10563 MELITACALLLIAVHIAIVISCPEPLAAVALVDVPLLYLCRYLLIIRRGVIIVSGAGAGGGRLRRFRLGAAMAIIYMAMSTLFFLRIAPLTPWWGALAAWVMILLIVEAIFAFFFPYRCCFNETEGDVQNNPHV
IRIS_313-10564 MELITACALLLIAVHIAIVISCPEPLAAVALVDVPLLYLCRYLLIIRRGVIIVSGAGAGGGRLRRFRLGAAMAIIYMAMSTLFFLRIAPLTPWWGALAAWVMILLIVEAIFAFFFPYRCCFNETEGDVQNNPHV
IRIS_313-10565 MELITACALLLIAVHIAIVISCPEPLAAVALVDVPLLYLCRYLLIIRRGVIIVSGGGGGGRLRRFRLGAAMAIIYMAMSTLFFLRIAPLTPWWGALAAWVMILLIVEAIFAFFFPYRCCFNETEGDVQNNPHV
IRIS_313-10566 MELITACALLLIAVHIAIVISCPEPLAAVALVDVPLLYLCRYLLIIRRGVIIVSGAGAGGGRLRRFRLGAAMAIIYMAMSTLFFLRIAPLTPWWGALAAWVMILLIVEAIFAFFFPYRCCFNETEGDVQNNPHV
IRIS_313-10567 MELITACALLLIAVHIAIVISCPEPLAAVALVDVPLLYLCRYLLIIRRGVIIVSGAGAGGGRLRRFRLGAAMAIIYMAMSTLFFLRIAPLTPWWGALAAWVMILLIVEAIFAFFFPYRCCFNETEGDVQNNPHV
IRIS_313-10568 MELITACALLLIAVHIAIVISCPEPLAAVALVDVPLLYLCRYLLIIRRGVIIVSGAGAGGGRLRRFRLGAAMAIIYMAMSTLFFLRIAPLTPWWGALAAWVMILLIVEAIFAFFFPYRCCFNETEGDVQNNPHV
IRIS_313-10569 MELITACALLLIAVHIAIVISCPEPLAAVALVDVPLLYLCRYLLIIRRGVIIVSGAGAGGGRLRRFRLGAAMAIIYMAMSTLFFLRIAPLTPWWGALAAWVMILLIVEAIFAFFFPYRCCFNETEGDVQNNPHV
IRIS_313-10570 MELITACALLLIAVHIAIVISCPEPLAAVALVDVPLLYLCRYLLIIRRGVIIVSGAGAGGGRLRRFRLGAAMAIIYMAMSTLFFLRIAPLTPWWGALAAWVMILLIVEAIFAFFFPYRCCFNETEGDVQNNPHV
IRIS_313-10572 MELITACALLLIAVHIAIVISCPEPLAAVALVDVPLLYLCRYLLIIRRGVIIVSGGGGGGRLRRFRLGAAMAIIYMAMSTLFFLRIAPLTPWWGALAAWVMILLIVEAIFAFFFPYRCCFNETEGDVQNNPHV
IRIS_313-10573 MELITACALLLIAVHIAIVISCPEPLAAVALVDVPLLYLCRYLLIIRRGVIIVSGAGAGGGRLRRFRLGAAMAIIYMAMSTLFFLRIAPLTPWWGALAAWVMILLIVEAIFAFFFPYRCCFNETEGDVQNNPHV
IRIS_313-10574 MELITACALLLIAVHIAIVISCPEPLAAVALVDVPLLYLCRYLLIIRRGVIIVSGAGAGGGRLRRFRLGAAMAIIYMAMSTLFFLRIAPLTPWWGALAAWVMILLIVEAIFAFFFPYRCCFNETEGDVQNNPHV
IRIS_313-10575 MELITACALLLIAVHIAIVISCPEPLAAVALVDVPLLYLCRYLLIIRRGVIIVSGGGGGGRLRRFRLGAAMAIIYMAMSTLFFLRIAPLTPWWGALAAWVMILLIVEAIFAFFFPYRCCFNETEGDVQNNPHV
IRIS_313-10576 MELITACALLLIAVHIAIVISCPEPLAAVALVDVPLLYLCRYLLIIRRGVIIVSGAGAGGGRLRRFRLGAAMAIIYMAMSTLFFLRIAPLTPWWGALAAWVMILLIVEAIFAFFFPYRCCFNETEGDVQNNPHV
IRIS_313-10577 MELITACALLLIAVHIAIVISCPEPLAAVALVDVPLLYLCRYLLIIRRGVIIVSGAGAGGGRLRRFRLGAAMAIIYMAMSTLFFLRIAPLTPWWGALAAWVMILLIVEAIFAFFFPYRCCFNETEGDVQNNPHV
IRIS_313-10578 MELITACALLLIAVHIAIVISCPEPLAAVALVDVPLLYLCRYLLIIRRGVIIVSGAGGGGGRLRRFRLGAAMAIIYMAMSTLFFLRIAPLTPWWGALAAWVMILLIVEAIFAFFFPYRCCFNETEGDVQNNPHV
IRIS_313-10580 MELITACALLLIAVHIAIVISCPEPLAAVALVDVPLLYLCRYLLIIRRGVIIVSGAGGGGGRLRRFRLGAAMAIIYMAMSTLFFLRIAPLTPWWGALAAWVMILLIVEAIFAFFFPYRCCFNETEGDVQNNPHV
IRIS_313-10581 MELITACALLLIAVHIAIVISCPEPLAAVALVDVPLLYLYRYLLIIRRGVIIVSGAGGGGGRLRRFRLGAAMAIIYMAMSTLFFLRIAPLTPWWGALAAWVMILLIVEAIFAFFFPYRCCFNETEGDVQNNPHV
IRIS_313-10582 MELITACALLLIAVHIAIVISCPEPLAAVALVDVPLLYLCRYLLIIRRGVIIVSGAGAGGGRLRRFRLGAAMAIIYMAMSTLFFLRIAPLTPWWGALAAWVMILLIVEAIFAFFFPYRCCFNETEGDVQNNPHV
IRIS_313-10583 MELITACALLLIAVHIAIVISCPEPLAAVALVDVPLLYLCRYLLIIRRGVIIVSGAGAGGGRLRRFRLGAAMAIIYMAMSTLFFLRIAPLTPWWGALAAWVMILLIVEAIFAFFFPYRCCFNETEGDVQNNPHV
IRIS_313-10585 MELITACALLLIAVHIAIVISCPEPLAAVALVDVPLLYLCRYLLIIRRGVIIVSGAGAGGGRLRRFRLGAAMAIIYMAMSTLFFLRIAPLTPWWGALAAWVMILLIVEAIFAFFFPYRCCFNETEGDVQNNPHV
IRIS_313-10587 MELITACALLLIAVHIAIVISCPEPLAAVAFVNVPLLYLCRYLLIIRRGVIIVSGGGGGGGRLRRFRLGAAMAVIYMAMSTLFFLRIAPLTPWWGALAAWVMILLIVEAIFAFFFPYRCCFNETEGDVQNNPHV
IRIS_313-10592 MELITACALLLIAVHIAIVISCPEPLAAVAFVNVPLLYLCRYLLIIRRGVIIVSGGGGGGGRLRRFRLGAAMAVIYMAMSTLFFLRIAPLTPWWGALAAWVMILLIVEAIFAFFFPYRCCFNETEGDVQNNPHV
IRIS_313-10593 MELITACALLLIAVHIAIVISCPEPLAAVAFVNVPLLYLCRYLLIIRRGVIIVSGGGGGGGRLRRFRLGAAMAVIYMAMSTLFFLRIAPLTPWWGALAAWVMILLIVEAIFAFFFPYRCCFNETEGDVQNNPHV
IRIS_313-10594 MELITACALLLIAVHIAIVISCPEPLAAVAFVNVPLLYLCRYLLIIRRGVIIVSGGGGGGGRLRRFRLGAAMAVIYMAMSTLFFLRIAPLTPWWGALAAWVMILLIVEAIFAFFFPYRCCFNETEGDVQNNPHV
IRIS_313-10595 MELITACALLLIAVHIAIVISCPEPLAAVALVDVPLLYLCRYLLIIRRGVIIVSGAGAGGGRLRRFRLGAAMAIIYMAMSTLFFLRIAPLTPWWGALAAWVMILLIVEAIFAFFFPYRCCFNETEGDVQNNPHV
IRIS_313-10598 MELITACALLLIAVHIAIVISCPEPLAAVAFVNVPLLYLCRYLLIIRRGVIIVSGGGGGGGRLRRFRLGAAMAVIYMAMSTLFFLRIAPLTPWWGALAAWVMILLIVEAIFAFFFPYRCCFNETEGDVQNNPHV
IRIS_313-10600 MELITACALLLIAVHIAIVISCPEPLAAVAFVNVPLLYLCRYLLIIRRGVIIVSGGGGGGGRLRRFRLGAAMAVIYMAMSTLFFLRIAPLTPWWGALAAWVMILLIVEAIFAFFFPYRCCFNETEGDVQNNPHV
IRIS_313-10602 MELITACALLLIAVHIAIVISCPEPLAAVALVDVPLLYLCRYLLIIRRGVIIVSGAGGGGGGRLRRFRLGAAMAIIYMAMSTLFFLRIAPLTPWWGALAAWVMILLIVEAIFAFFFPYRCCFNETEGDVQNNPHV
IRIS_313-10603 MELITACALLLIAVHIAIVISCPEPLAAVALVDVPLLYLCRYLLIIRRGVIIVSGAGAGGGRLRRFRLGAAMAIIYMAMSTLFFLRIAPLTPWWGALAAWVMILLIVEAIFAFFFPYRCCFNETEGDVQNNPHV
IRIS_313-10604 MELITACALLLIAVHIAIVISCPEPLAAVAFVNVPLLYLCRYLLIIRRGVIIVSGGGGGGGRLRRFRLGAAMAVIYMAMSTLFFLRIAPLTPWWGALAAWVMILLIVEAIFAFFFPYRCCFNETEGDVQNNPHV
IRIS_313-10605 MELITACALLLIAVHIAIVISCPEPLAAVAFVNVPLLYLCRYLLIIRRGVIIVSGGGGGGGRLRRFRLGAAMAVIYMAMSTLFFLRIAPLTPWWGALAAWVMILLIVEAIFAFFFPYRCCFNETEGDVQNNPHV
IRIS_313-10606 MELITACALLLIAVHIAIVISCPEPLAAVAFVNVPLLYLCRYLLIIRRGVIIVSGGGGGGGRLRRFRLGAAMAVIYMAMSTLFFLRIAPLTPWWGALAAWVMILLIVEAIFAFFFPYRCCFNETEGDVQNNPHV
IRIS_313-10608 MELITACALLLIAVHIAIVISCPEPLAAVALVDVPLLYLCRYLLIIRRGVIIVSGAGAGGGRLRRFRLGAAMAIIYMAMSTLFFLRIAPLTPWWGALAAWVMILLIVEAIFAFFFPYRCCFNETEGDVQNNPHV
IRIS_313-10609 MELITACALLLIAVHIAIVISCPEPLAAVALVDVPLLYLCRYLLIIRRGVIIVSGAGAGGGRLRRFRLGAAMAIIYMAMSTLFFLRIAPLTPWWGALAAWVMILLIVEAIFAFFFPYRCCFNETEGDVQNNPHV
IRIS_313-10610 MELITACALLLIAVHIAIVISCPEPLAAVALVDVPLLYLCRYLLIIRRGVIIVSGGGGGGRLRRFRLGAAMAIIYMAMSTLFFLRIAPLTPWWGALAAWVMILLIVEAIFAFFFPYRCCFNETEGDVQNNPHV
IRIS_313-10614 MELITACALLLIAVHIAIVISCPEPLAAVALVDVPLLYLYRYLLIIRRGVIIVSGAGGGGGRLRRFRLGAAMAIIYMAMSTLFFLRIAPLTPWWGALAAWVMILLIVEAIFAFFFPYRCCFNETEGDVQNNPHV
IRIS_313-10618 MELITACALLLIAVHIAIVISCPEPLAAVALVDVPLLYLCRYLLIIRRGVIIVSGAGAGGGRLRRFRLGAAMAIIYMAMSTLFFLRIAPLTPWWGALAAWVMILLIVEAIFAFFFPYRCCFNETEGDVQNNPHV
IRIS_313-10619 MELITACALLLIAVHIAIVISCPEPLAAVALVDVPLLYLCRYLLIIRRGVIIVSGAGAGGGRLRRFRLGAAMAIIYMAMSTLFFLRIAPLTPWWGALAAWVMILLIVEAIFAFFFPYRCCFNETEGDVQNNPHV
IRIS_313-10620 MELITACALLLIAVHIAIVISCPEPLAAVALVDVPLLYLCRYLLIIRRGVIIVSGAGAGGGRLRRFRLGAAMAIIYMAMSTLFFLRIAPLTPWWGALAAWVMILLIVEAIFAFFFPYRCCFNETEGDVQNNPHV
IRIS_313-10623 MELITACALLLIAVHIAIVISCPEPLAAVALVDVPLLYLCRYLLIIRRGVIIVSGAGAGGGRLRRFRLGAAMAIIYMAMSTLFFLRIAPLTPWWGALAAWVMILLIVEAIFAFFFPYRCCFNETEGDVQNNPHV
IRIS_313-10628 MELITACALLLIAVHIAIVISCPEPLAAVALVDVPLLYLCRYLLIIRRGVIIVSGAGGGGGGRLRRFRLGAAMAIIYMAMSTLFFLRIAPLTPWWGALAAWVMILLIVEAIFAFFFPYRCCFNETEGDVQNNPHV
IRIS_313-10631 MELITACALLLIAVHIAIVISCPEPLAAVALVDVPLLYLYRYLLIIRRGVIIVSGAGGGGGRLRRFRLGAAMAIIYMAMSTLFFLRIAPLTPWWGALAAWVMILLIVEAIFAFFFPYRCCFNETEGDVQNNPHV
IRIS_313-10640 MELITACALLLIAVHIAIVISCPEPLAAVALVDVPLLYLCRYLLIIRRGVIIVSGAGAGGGRLRRFRLGAAMAIIYMAMSTLFFLRIAPLTPWWGALAAWVMILLIVEAIFAFFFPYRCCFNETEGDVQNNPHV
IRIS_313-10642 MELITACALLLIAVHIAIVISCPEPLAAVALVDVPLLYLCRYLLIIRRGVIIVSGAGAGGGRLRRFRLGAAMAIIYMAMSTLFFLRIAPLTPWWGALAAWVMILLIVEAIFAFFFPYRCCFNETEGDVQNNPHV
IRIS_313-10644 MELITACALLLIAVHIAIVISCPEPLAAVALVDVPLLYLCRYLLIIRRGVIIVSGAGAGGGRLRRFRLGAAMAIIYMAMSTLFFLRIAPLTPWWGALAAWVMILLIVEAIFAFFFPYRCCFNETEGDVQNNPHV
IRIS_313-10645 MELITACALLLIAVHIAIVISCPEPLAAVALVDVPLLYLCRYLLIIRRGVIIVSGAGAGGGRLRRFRLGAAMAIIYMAMSTLFFLRIAPLTPWWGALAAWVMILLIVEAIFAFFFPYRCCFNETEGDVQNNPHV
IRIS_313-10648 MELITACALLLIAVHIAIVISCPEPLAAVALVDVPLLYLYRYLLIIRRGVIIVSGAGGGGGRLRRFRLGAAMAIIYMAMSTLFFLRIAPLTPWWGALAAWVMILLIVEAIFAFFFPYRCCFNETEGDVQNNPHV
IRIS_313-10649 MELITACALLLIAVHIAIVISCPEPLAAVALVDVPLLYLYRYLLIIRRGVIIVSGAGGGGGRLRRFRLGAAMAIIYMAMSTLFFLRIAPLTPWWGALAAWVMILLIVEAIFAFFFPYRCCFNETEGDVQNNPHV
IRIS_313-10650 MELITACALLLIAVHIAIVISCPEPLAAVALVDVPLLYLCRYLLIIRRGVIIVSGGGGGGRLRRFRLGAAMAIIYMAMSTLFFLRIAPLTPWWGALAAWVMILLIVEAIFAFFFPYRCCFNETEGDVQNNPHV
IRIS_313-10651 MELITACALLLIAVHIAIVISCPEPLAAVALVDVPLLYLCRYLLIIRRGVIIVSGGGAGGGRLRRFRLGAAMAIIYMAMSTLFFLRIAPLTPWWGALAAWVMILLIVEAIFAFFFPYRCCFNETEGDVQNNPHV
IRIS_313-10653 MELITACALLLIAVHIAIVISCPEPLAAVALVDVPLLYLCRYLLIIRRGVIIVSGAGGGGGRLRRFRLGAAMAIIYMAMSTLFFLRIAPLTPWWGALAAWVMILLIVEAIFAFFFPYRCCFNETEGDVQNNPHV
IRIS_313-10654 MELITACALLLIAVHIAIVISCPEPLAAVALVDVPLLYLYRYLLIIRRGVIIVSGAGGGGGRLRRFRLGAAMAIIYMAMSTLFFLRIAPLTPWWGALAAWVMILLIVEAIFAFFFPYRCCFNETEGDVQNNPHV
IRIS_313-10655 MELITACALLLIAVHIAIVISCPEPLAAVALVDVPLLYLCRYLLIIRRGVIIVSGAGAGGGRLRRFRLGAAMAIIYMAMSTLFFLRIAPLTPWWGALAAWVMILLIVEAIFAFFFPYRCCFNETEGDVQNNPHV
IRIS_313-10656 MELITACALLLIAVHIAIVISCPEPLAAVALVDVPLLYLCRYLLIIRRGVIIVSGAGAGGGRLRRFRLGAAMAIIYMAMSTLFFLRIAPLTPWWGALAAWVMILLIVEAIFAFFFPYRCCFNETEGDVQNNPHV
IRIS_313-10657 MELITACALLLIAVHIAIVISCPEPLAAVALVDVPLLYLYRYLLIIRRGVIIVSGAGGGGGRLRRFRLGAAMAIIYMAMSTLFFLRIAPLTPWWGALAAWVMILLIVEAIFAFFFPYRCCFNETEGDVQNNPHV
IRIS_313-10659 MELITACALLLIAVHIAIVISCPEPLAAVALVDVPLLYLCRYLLIIRRGVIIVSGGGGGGRLRRFRLGAAMAIIYMAMSTLFFLRIAPLTPWWGALAAWVMILLIVEAIFAFFFPYRCCFNETEGDVQNNPHV
IRIS_313-10660 MELITACALLLIAVHIAIVISCPEPLAAVALVDVPLLYLYRYLLIIRRGVIIVSGAGGGGGRLRRFRLGAAMAIIYMAMSTLFFLRIAPLTPWWGALAAWVMILLIVEAIFAFFFPYRCCFNETEGDVQNNPHV
IRIS_313-10661 MELITACALLLIAVHIAIVISCPEPLAAVALVDVPLLYLCRYLLIIRRGVIIVSGGGGGGRLRRFRLGAAMAIIYMAMSTLFFLRIAPLTPWWGALAAWVMILLIVEAIFAFFFPYRCCFNETEGDVQNNPHV
IRIS_313-10662 MELITACALLLIAVHIAIVISCPEPLAAVALVDVPLLYLCRYLLIIRRGVIIVSGGGGGGRLRRFRLGAAMAIIYMAMSTLFFLRIAPLTPWWGALAAWVMILLIVEAIFAFFFPYRCCFNETEGDVQNNPHV
IRIS_313-10664 MELITACALLLIAVHIAIVISCPEPLAAVALVDVPLLYLCRYLLIIRRGVIIVSGGGGGGRLRRFRLGAAMAIIYMAMSTLFFLRIAPLTPWWGALAAWVMILLIVEAIFAFFFPYRCCFNETEGDVQNNPHV
IRIS_313-10666 MELITACALLLIAVHIAIVISCPEPLAAVALVDVPLLYLCRYLLIIRRGVIIVSGGGGGGRLRRFRLGAAMAIIYMAMSTLFFLRIAPLTPWWGALAAWVMILLIVEAIFAFFFPYRCCFNETEGDVQNNPHV
IRIS_313-10667 MELITACALLLIAVHIAIVISCPEPLAAVALVDVPLLYLCRYLLIIRRGVIIVSGAGGGGGGRLRRFRLGAAMAIIYMAMSTLFFLRIAPLTPWWGALAAWVMILLIVEAIFAFFFPYRCCFNETEGDVQNNPHV
IRIS_313-10668 MELITACALLLIAVHIAIVISCPEPLAAVALVDVPLLYLCRYLLIIRRGVIIVSGAGAGGGRLRRFRLGAAMAIIYMAMSTLFFLRIAPLTPWWGALAAWVMILLIVEAIFAFFFPYRCCFNETEGDVQNNPHV
IRIS_313-10670 MELITACALLLIAVHIAIVISCPEPLAAVALVDVPLLYLCRYLLIIRRGVIIVSGAGAGGGRLRRFRLGAAMAIIYMAMSTLFFLRIAPLTPWWGALAAWVMILLIVEAIFAFFFPYRCCFNETEGDVQNNPHV
IRIS_313-10671 MELITACALLLIAVHIAIVISCPEPLAAVALVDVPLLYLCRYLLIIRRGVIIVSGAGAGGGRLRRFRLGAAMAIIYMAMSTLFFLRIAPLTPWWGALAAWVMILLIVEAIFAFFFPYRCCFNETEGDVQNNPHV
IRIS_313-10673 MELITACALLLIAVHIAIVISCPEPLAAVALVDVPLLYLCRYLLIIRRGVIIVSGAGAGGGRLRRFRLGAAMAIIYMAMSTLFFLRIAPLTPWWGALAAWVMILLIVEAIFAFFFPYRCCFNETEGDVQNNPHV
IRIS_313-10674 MELITACALLLIAVHIAIVISCPEPLAAVALVDVPLLYLCRYLLIIRRGVIIVSGAGAGGGRLRRFRLGAAMAIIYMAMSTLFFLRIAPLTPWWGALAAWVMILLIVEAIFAFFFPYRCCFNETEGDVQNNPHV
IRIS_313-10675 MELITACALLLIAVHIAIVISCPEPLAAVALVDVPLLYLCRYLLIIRRGVIIVSGAGAGGGRLRRFRLGAAMAIIYMAMSTLFFLRIAPLTPWWGALAAWVMILLIVEAIFAFFFPYRCCFNETEGDVQNNPHV
IRIS_313-10676 MELITACALLLIAVHIAIVISCPEPLAAVALVDVPLLYLCRYLLIIRRGVIIVSGGGGGGRLRRFRLGAAMAIIYMAMSTLFFLRIAPLTPWWGALAAWVMILLIVEAIFAFFFPYRCCFNETEGDVQNNPHV
IRIS_313-10677 MELITACALLLIAVHIAIVISCPEPLAAVALVDVPLLYLCRYLLIIRRGVIIVSGAGAGGGRLRRFRLGAAMAIIYMAMSTLFFLRIAPLTPWWGALAAWVMILLIVEAIFAFFFPYRCCFNETEGDVQNNPHV
IRIS_313-10678 MELITACALLLIAVHIAIVISCPEPLAAVALVDVPLLYLCRYLLIIRRGVIIVSGAGAGGGRLRRFRLGAAMAIIYMAMSTLFFLRIAPLTPWWGALAAWVMILLIVEAIFAFFFPYRCCFNETEGDVQNNPHV
IRIS_313-10679 MELITACALLLIAVHIAIVISCPEPLAAVALVDVPLLYLCRYLLIIRRGVIIVSGAGAGGGRLRRFRLGAAMAIIYMAMSTLFFLRIAPLTPWWGALAAWVMILLIVEAIFAFFFPYRCCFNETEGDVQNNPHV
IRIS_313-10680 MELITACALLLIAVHIAIVISCPEPLAAVALVDVPLLYLYRYLLIIRRGVIIVSGAGGGGGRLRRFRLGAAMAIIYMAMSTLFFLRIAPLTPWWGALAAWVMILLIVEAIFAFFFPYRCCFNETEGDVQNNPHV
IRIS_313-10681 MELITACALLLIAVHIAIVISCPEPLAAVALVDVPLLYLYRYLLIIRRGVIIVSGAGGGGGRLRRFRLGAAMAIIYMAMSTLFFLRIAPLTPWWGALAAWVMILLIVEAIFAFFFPYRCCFNETEGDVQNNPHV
IRIS_313-10682 MELITACALLLIAVHIAIVISCPEPLAAVALVDVPLLYLYRYLLIIRRGVIIVSGAGGGGGRLRRFRLGAAMAIIYMAMSTLFFLRIAPLTPWWGALAAWVMILLIVEAIFAFFFPYRCCFNETEGDVQNNPHV
IRIS_313-10683 MELITACALLLIAVHIAIVISCPEPLAAVALVDVPLLYLYRYLLIIRRGVIIVSGAGGGGGRLRRFRLGAAMAIIYMAMSTLFFLRIAPLTPWWGALAAWVMILLIVEAIFAFFFPYRCCFNETEGDVQNNPHV
IRIS_313-10684 MELITACALLLIAVHIAIVISCPEPLAAVALVDVPLLYLCRYLLIIRRGVIIVSGAGAGGGRLRRFRLGAAMAIIYMAMSTLFFLRIAPLTPWWGALAAWVMILLIVEAIFAFFFPYRCCFNETEGDVQNNPHV
IRIS_313-10687 MELITACALLLIAVHIAIVISCPEPLAAVALVDVPLLYLCRYLLIIRRGVIIVSGAGAGGGRLRRFRLGAAMAIIYMAMSTLFFLRIAPLTPWWGALAAWVMILLIVEAIFAFFFPYRCCFNETEGDVQNNPHV
IRIS_313-10688 MELITACALLLIAVHIAIVISCPEPLAAVALVDVPLLYLCRYLLIIRRGVIIVSGGGGGGRLRRFRLGAAMAIIYMAMSTLFFLRIAPLTPWWGALAAWVMILLIVEAIFAFFFPYRCCFNETEGDVQNNPHV
IRIS_313-10689 MELITACALLLIAVHIAIVISCPEPLAAVALVDVPLLYLCRYLLIIRRGVIIVSGAGAGGGRLRRFRLGAAMAIIYMAMSTLFFLRIAPLTPWWGALAAWVMILLIVEAIFAFFFPYRCCFNETEGDVQNNPHV
IRIS_313-10690 MELITACALLLIAVHIAIVISCPEPLAAVALVDVPLLYLCRYLLIIRRGVIIVSGGGAGGGRLRRFRLGAAMAIIYMAMSTLFFLRIAPLTPWWGALAAWVMILLIVEAIFAFFFPYRCCFNETEGDVQNNPHV
IRIS_313-10692 MELITACALLLIAVHIAIVISCPEPLAAVALVDVPLLYLCRYLLIIRRGVIIVSGGGGGGRLRRFRLGAAMAIIYMAMSTLFFLRIAPLTPWWGALAAWVMILLIVEAIFAFFFPYRCCFNETEGDVQNNPHV
IRIS_313-10693 MELITACALLLIAVHIAIVISCPEPLAAVALVDVPLLYLCRYLLIIRRGVIIVSGAGAGGGRLRRFRLGAAMAIIYMAMSTLFFLRIAPLTPWWGALAAWVMILLIVEAIFAFFFPYRCCFNETEGDVQNNPHV
IRIS_313-10694 MELITACALLLIAVHIAIVISCPEPLAAVALVDVPLLYLCRYLLIIRRGVIIVSGAGAGGGRLRRFRLGAAMAIIYMAMSTLFFLRIAPLTPWWGALAAWVMILLIVEAIFAFFFPYRCCFNETEGDVQNNPHV
IRIS_313-10695 MELITACALLLIAVHIAIVISCPEPLAAVALVDVPLLYLCRYLLIIRRGVIIVSGAGAGGGRLRRFRLGAAMAIIYMAMSTLFFLRIAPLTPWWGALAAWVMILLIVEAIFAFFFPYRCCFNETEGDVQNNPHV
```

IRIS_313-10697 MELITACALLLIAVHIAIVISCPEPLAAVALVDVPLLYLCRYLLIIRRGVIIVSGGGGGGRLRRFRLGAAMAIIYMAMSTLFFLRIAPLTPWWGALAAWVMILLIVEAIFAFFFPYRCCFNETEGDVQNNPHV
IRIS_313-10698 MELITACALLLIAVHIAIVISCPEPLAAVALVDVPLLYLCRYLLIIRRGVIIVSGAGAGGGRLRRFRLGAAMAIIYMAMSTLFFLRIAPLTPWWGALAAWVMILLIVEAIFAFFFPYRCCFNETEGDVQNNPHV
IRIS_313-10699 MELITACALLLIAVHIAIVISCPEPLAAVALVDVPLLYLCRYLLIIRRGVIIVSGGGGGGRLRRFRLGAAMAIIYMAMSTLFFLRIAPLTPWWGALAAWVMILLIVEAIFAFFFPYRCCFNETEGDVQNNPHV
IRIS_313-10700 MELITACALLLIAVHIAIVISCPEPLAAVALVDVPLLYLCRYLLIIRRGVIIVSGGGGGGRLRRFRLGAAMAIIYMAMSTLFFLRIAPLTPWWGALAAWVMILLIVEAIFAFFFPYRCCFNETEGDVQNNPHV
IRIS_313-10701 MELITACALLLIAVHIAIVISCPEPLAAVALVDVPLLYLCRYLLIIRRGVIIVSGGGGGGRLRRFRLGAAMAIIYMAMSTLFFLRIAPLTPWWGALAAWVMILLIVEAIFAFFFPYRCCFNETEGDVQNNPHV
IRIS_313-10702 MELITACALLLIAVHIAIVISCPEPLAAVALVDVPLLYLCRYLLIIRRGVIIVSGAGAGGGGRLRRFRLGAAMAIIYMAMSTLFFLRIAPLTPWWGALAAWVMILLIVEAIFAFFFPYRCCFNETEGDVQNNPHV
IRIS_313-10703 MELITACALLLIAVHIAIVISCPEPLAAVALVDVPLLYLYRYLLIIRRGVIIVSGAGGGGGRLRRFRLGAAMAIIYMAMSTLFFLRIAPLTPWWGALAAWVMILLIVEAIFAFFFPYRCCFNETEGDVQNNPHV
IRIS_313-10704 MELITACALLLIAVHIAIVISCPEPLAAVALVDVPLLYLCRYLLIIRRGVIIVSGAGAGGGRLRRFRLGAAMAIIYMAMSTLFFLRIAPLTPWWGALAAWVMILLIVEAIFAFFFPYRCCFNETEGDVQNNPHV
IRIS_313-10705 MELITACALLLIAVHIAIVISCPEPLAAVALVDVPLLYLCRYLLIIRRGVIIVSGGGAGGGRLRRFRLGAAMAIIYMAMSTLFFLRIAPLTPWWGALAAWVMILLIVEAIFAFFFPYRCCFNETEGDVQNNPHV
IRIS_313-10706 MELITACALLLIAVHIAIVISCPEPLAAVALVDVPLLYLCRYLLIIRRGVIIVSGAGAGGGRLRRFRLGAAMAIIYMAMSTLFFLRIAPLTPWWGALAAWVMILLIVEAIFAFFFPYRCCFNETEGDVQNNPHV
IRIS_313-10707 MELITACALLLIAVHIAIVISCPEPLAAVALVDVPLLYLCRYLLIIRRGVIIVSGAGAGGGRLRRFRLGAAMAIIYMAMSTLFFLRIAPLTPWWGALAAWVMILLIVEAIFAFFFPYRCCFNETEGDVQNNPHV
IRIS_313-10708 MELITACALLLIAVHIAIVISCPEPLAAVALVDVPLLYLYRYLLIIRRGVIIVSGAGGGGGRLRRFRLGAAMAIIYMAMSTLFFLRIAPLTPWWGALAAWVMILLIVEAIFAFFFPYRCCFNETEGDVQNNPHV
IRIS_313-10710 MELITACALLLIAVHIAIVISCPEPLAAVALVDVPLLYLCRYLLIIRRGVIIVSGAGAGGGRLRRFRLGAAMAIIYMAMSTLFFLRIAPLTPWWGALAAWVMILLIVEAIFAFFFPYRCCFNETEGDVQNNPHV
IRIS_313-10711 MELITACALLLIAVHIAIVISCPEPLAAVALVDVPLLYLCRYLLIIRRGVIIVSGAGAGGGRLRRFRLGAAMAIIYMAMSTLFFLRIAPLTPWWGALAAWVMILLIVEAIFAFFFPYRCCFNETEGDVQNNPHV
IRIS_313-10712 MELITACALLLIAVHIAIVISCPEPLAAVALVDVPLLYLCRYLLIIRRGVIIVSGAGAGGGRLRRFRLGAAMAIIYMAMSTLFFLRIAPLTPWWGALAAWVMILLIVEAIFAFFFPYRCCFNETEGDVQNNPHV
IRIS_313-10715 MELITACALLLIAVHIAIVISCPEPLAAVALVDVPLLYLCRYLLIIRRGVIIVSGAGAGGGRLRRFRLGAAMAIIYMAMSTLFFLRIAPLTPWWGALAAWVMILLIVEAIFAFFFPYRCCFNETEGDVQNNPHV
IRIS_313-10716 MELITACALLLIAVHIAIVISCPEPLAAVALVDVPLLYLCRYLLIIRRGVIIVSGGGGGGRLRRFRLGAAMAIIYMAMSTLFFLRIAPLTPWWGALAAWVMILLIVEAIFAFFFPYRCCFNETEGDVQNNPHV
IRIS_313-10717 MELITACALLLIAVHIAIVISCPEPLAAVALVDVPLLYLCRYLLIIRRGVIIVSGGGGGGRLRRFRLGAAMAIIYMAMSTLFFLRIAPLTPWWGALAAWVMILLIVEAIFAFFFPYRCCFNETEGDVQNNPHV
IRIS_313-10718 MELITACALLLIAVHIAIVISCPEPLAAVAFVNVPLLYLCRYLLIIRRGVIIVSGGGGGGGRLRRFRLGAAMAVIYMAMSTLFFLRIAPLTPWWGALAAWVMILLIVEAIFAFFFPYRCCFNETEGDVQNNPHV
IRIS_313-10719 MELITACALLLIAVHIAIVISCPEPLAAVALVDVPLLYLCRYLLIIRRGVIIVSGGGAGGGRLRRFRLGAAMAIIYMAMSTLFFLRIAPLTPWWGALAAWVMILLIVEAIFAFFFPYRCCFNETEGDVQNNPHV
IRIS_313-10720 MELITACALLLIAVHIAIVISCPEPLAAVALVDVPLLYLCRYLLIIRRGVIIVSGGGAGGGRLRRFRLGAAMAIIYMAMSTLFFLRIAPLTPWWGALAAWVMILLIVEAIFAFFFPYRCCFNETEGDVQNNPHV
IRIS_313-10721 MELITACALLLIAVHIAIVISCPEPLAAVALVDVPLLYLCRYLLIIRRGVIIVSGGGGGGRLRRFRLGAAMAIIYMAMSTLFFLRIAPLTPWWGALAAWVMILLIVEAIFAFFFPYRCCFNETEGDVQNNPHV
IRIS_313-10722 MELITACALLLIAVHIAIVISCPEPLAAVALVDVPLLYLCRYLLIIRRGVIIVSGAGAGGGRLRRFRLGAAMAIIYMAMSTLFFLRIAPLTPWWGALAAWVMILLIVEAIFAFFFPYRCCFNETEGDVQNNPHV
IRIS_313-10723 MELITACALLLIAVHIAIVISCPEPLAAVALVDVPLLYLCRYLLIIRRGVIIVSGAGAGGGRLRRFRLGAAMAIIYMAMSTLFFLRIAPLTPWWGALAAWVMILLIVEAIFAFFFPYRCCFNETEGDVQNNPHV
IRIS_313-10724 MELITACALLLIAVHIAIVISCPEPLAAVALVDVPLLYLYRYLLIIRRGVIIVSGAGGGGGRLRRFRLGAAMAIIYMAMSTLFFLRIAPLTPWWGALAAWVMILLIVEAIFAFFFPYRCCFNETEGDVQNNPHV
IRIS_313-10725 MELITACALLLIAVHIAIVISCPEPLAAVALVDVPLLYLCRYLLIIRRGVIIVSGAGAGGGRLRRFRLGAAMAIIYMAMSTLFFLRIAPLTPWWGALAAWVMILLIVEAIFAFFFPYRCCFNETEGDVQNNPHV
IRIS_313-10726 MELITACALLLIAVHIAIVISCPEPLAAVALVDVPLLYLCRYLLIIRRGVIIVSGAGGAAAAGCAASASGQPWRSSTWPCRPSSSSALLLSRHGGAPSPPGS*SCSSSRPSSPSSSPTAAASTKPKAMSRIIPM
IRIS_313-10727 MELITACALLLIAVHIAIVISCPEPLAAVALVDVPLLYLCRYLLIIRRGVIIVSGAGGAAAAGCAASASGQPWRSSTWPCRPSSSSALLLSRHGGAPSPPGS*SCSSSRPSSPSSSPTAAASTKPKAMSRIIPM
IRIS_313-10728 MELITACALLLIAVHIAIVISCPEPLAAVALVDVPLLYLCRYLLIIRRGVIIVSGAGAGGGRLRRFRLGAAMAIIYMAMSTLFFLRIAPLTPWWGALAAWVMILLIVEAIFAFFFPYRCCFNETEGDVQNNPHV
IRIS_313-10730 MELITACALLLIAVHIAIVISCPEPLAAVALVDVPLLYLYRYLLIIRRGVIIVSGAGGGGGRLRRFRLGAAMAIIYMAMSTLFFLRIAPLTPWWGALAAWVMILLIVEAIFAFFFPYRCCFNETEGDVQNNPHV
IRIS_313-10731 MELITACALLLIAVHIAIVISCPEPLAAVALVDVPLLYLCRYLLIIRRGVIIVSGGGGGGRLRRFRLGAAMAIIYMAMSTLFFLRIAPLTPWWGALAAWVMILLIVEAIFAFFFPYRCCFNETEGDVQNNPHV
IRIS_313-10732 MELITACALLLIAVHIAIVISCPEPLAAVALVDVPLLYLCRYLLIIRRGVIIVSGAGAGGGRLRRFRLGAAMAIIYMAMSTLFFLRIAPLTPWWGALAAWVMILLIVEAIFAFFFPYRCCFNETEGDVQNNPHV
IRIS_313-10733 MELITACALLLIAVHIAIVISCPEPLAAVALVDVPLLYLCRYLLIIRRGVIIVSGGGGGGRLRRFRLGAAMAIIYMAMSTLFFLRIAPLTPWWGALAAWVMILLIVEAIFAFFFPYRCCFNETEGDVQNNPHV
IRIS_313-10734 MELITACALLLIAVHIAIVISCPEPLAAVAFVNVPLLYLCRYLLIIRRGVIIVSGGGGGGGRLRRFRLGAAMAVIYMAMSTLFFLRIAPLTPWWGALAAWVMILLIVEAIFAFFFPYRCCFNETEGDVQNNPHV
IRIS_313-10735 MELITACALLLIAVHIAIVISCPEPLAAVALVDVPLLYLCRYLLIIRRGVIIVSGAGAGGGRLRRFRLGAAMAIIYMAMSTLFFLRIAPLTPWWGALAAWVMILLIVEAIFAFFFPYRCCFNETEGDVQNNPHV
IRIS_313-10736 MELITACALLLIAVHIAIVISCPEPLAAVAFVNVPLLYLCRYLLIIRRGVIIVSGGGGGGGRLRRFRLGAAMAVIYMAMSTLFFLRIAPLTPWWGALAAWVMILLIVEAIFAFFFPYRCCFNETEGDVQNNPHV
IRIS_313-10737 MELITACALLLIAVHIAIVISCPEPLAAVALVDVPLLYLCRYLLIIRRGVIIVSGAGAGGGRLRRFRLGAAMAIIYMAMSTLFFLRIAPLTPWWGALAAWVMILLIVEAIFAFFFPYRCCFNETEGDVQNNPHV
IRIS_313-10738 MELITACALLLIAVHIAIVISCPEPLAAVALVDVPLLYLCRYLLIIRRGVIIVSGAGAGGGRLRRFRLGAAMAIIYMAMSTLFFLRIAPLTPWWGALAAWVMILLIVEAIFAFFFPYRCCFNETEGDVQNNPHV
IRIS_313-10739 MELITACALLLIAVHIAIVISCPEPLAAVALVDVPLLYLCRYLLIIRRGVIIVSGGGGGGRLRRFRLGAAMAIIYMAMSTLFFLRIAPLTPWWGALAAWVMILLIVEAIFAFFFPYRCCFNETEGDVQNNPHV
IRIS_313-10740 MELITACALLLIAVHIAIVISCPEPLAAVALVDVPLLYLYRYLLIIRRGVIIVSGAGGGGGRLRRFRLGAAMAIIYMAMSTLFFLRIAPLTPWWGALAAWVMILLIVEAIFAFFFPYRCCFNETEGDVQNNPHV
IRIS_313-10741 MELITACALLLIAVHIAIVISCPEPLAAVALVDVPLLYLYRYLLIIRRGVIIVSGAGGGGGRLRRFRLGAAMAIIYMAMSTLFFLRIAPLTPWWGALAAWVMILLIVEAIFAFFFPYRCCFNETEGDVQNNPHV
IRIS_313-10742 MELITACALLLIAVHIAIVISCPEPLAAVALVDVPLLYLCRYLLIIRRGVIIVSGGGAGGGRLRRFRLGAAMAIIYMAMSTLFFLRIAPLTPWWGALAAWVMILLIVEAIFAFFFPYRCCFNETEGDVQNNPHV
IRIS_313-10743 MELITACALLLIAVHIAIVISCPEPLAAVALVDVPLLYLCRYLLIIRRGVIIVSGAGAGGGRLRRFRLGAAMAIIYMAMSTLFFLRIAPLTPWWGALAAWVMILLIVEAIFAFFFPYRCCFNETEGDVQNNPHV
IRIS_313-10744 MELITACALLLIAVHIAIVISCPEPLAAVALVDVPLLYLCRYLLIIRRGVIIVSGAGGGGGRLRRFRLGAAMAIIYMAMSTLFFLRIAPLTPWWGALAAWVMILLIVEAIFAFFFPYRCCFNETEGDVQNNPHV
IRIS_313-10745 MELITACALLLIAVHIAIVISCPEPLAAVALVDVPLLYLCRYLLIIRRGVIIVSGAGAGGGRLRRFRLGAAMAIIYMAMSTLFFLRIAPLTPWWGALAAWVMILLIVEAIFAFFFPYRCCFNETEGDVQNNPHV
IRIS_313-10746 MELITACALLLIAVHIAIVISCPEPLAAVALVDVPLLYLCRYLLIIRRGVIIVSGAGAGGGRLRRFRLGAAMAIIYMVMSTLFFLRIAPLTPWWGALAAWVMILLIVEAIFAFFFPYRCCFNETEGDVQNNPHV
IRIS_313-10747 MELITACALLLIAVHIAIVISCPEPLAAVALVDVPLLYLYRYLLIIRRGVIIVSGAGGGGGRLRRFRLGAAMAIIYMAMSTLFFLRIAPLTPWWGALAAWVMILLIVEAIFAFFFPYRCCFNETEGDVQNNPHV
IRIS_313-10748 MELITACALLLIAVHIAIVISCPEPLAAVALVDVPLLYLYRYLLIIRRGVIIVSGAGGGGGRLRRFRLGAAMAIIYMAMSTLFFLRIAPLTPWWGALAAWVMILLIVEAIFAFFFPYRCCFNETEGDVQNNPHV
IRIS_313-10749 MELITACALLLIAVHIAIVISCPEPLAAVALVDVPLLYLYRYLLIIRRGVIIVSGAGGGGGRLRRFRLGAAMAIIYMAMSTLFFLRIAPLTPWWGALAAWVMILLIVEAIFAFFFPYRCCFNETEGDVQNNPHV
IRIS_313-10750 MELITACALLLIAVHIAIVISCPEPLAAVALVDVPLLYLCRYLLIIRRGVIIVSGGGGGGRLRRFRLGAAMAIIYMAMSTLFFLRIAPLTPWWGALAAWVMILLIVEAIFAFFFPYRCCFNETEGDVQNNPHV
IRIS_313-10751 MELITACALLLIAVHIAIVISCPEPLAAVALVDVPLLYLCRYLLIIRRGVIIVSGAGAGGGRLRRFRLGAAMAIIYMAMSTLFFLRIAPLTPWWGALAAWVMILLIVEAIFAFFFPYRCCFNETEGDVQNNPHV
IRIS_313-10752 MELITACALLLIAVHIAIVISCPEPLAAVALVDVPLLYLCRYLLIIRRGVIIVSGAGAGGGRLRRFRLGAAMAIIYMAMSTLFFLRIAPLTPWWGALAAWVMILLIVEAIFAFFFPYRCCFNETEGDVQNNPHV
IRIS_313-10753 MELITACALLLIAVHIAIVISCPEPLAAVALVDVPLLYLCRYLLIIRRGVIIVSGAGAGGGRLRRFRLGAAMAIIYMAMSTLFFLRIAPLTPWWGALAAWVMILLIVEAIFAFFFPYRCCFNETEGDVQNNPHV
IRIS_313-10754 MELITACALLLIAVHIAIVISCPEPLAAVALVDVPLLYLCRYLLIIRRGVIIVSGGGGGGRLRRFRLGAAMAIIYMAMSTLFFLRIAPLTPWWGALAAWVMILLIVEAIFAFFFPYRCCFNETEGDVQNNPHV
IRIS_313-10755 MELITACALLLIAVHIAIVISCPEPLAAVALVDVPLLYLYRYLLIIRRGVIIVSGAGGGGGRLRRFRLGAAMAIIYMAMSTLFFLRIAPLTPWWGALAAWVMILLIVEAIFAFFFPYRCCFNETEGDVQNNPHV
IRIS_313-10756 MELITACALLLIAVHIAIVISCPEPLAAVALVDVPLLYLCRYLLIIRRGVIIVSGGGGGGRLRRFRLGAAMAIIYMAMSTLFFLRIAPLTPWWGALAAWVMILLIVEAIFAFFFPYRCCFNETEGDVQNNPHV
IRIS_313-10757 MELITACALLLIAVHIAIVISCPEPLAAVALVDVPLLYLCRYLLIIRRGVIIVSGAGGGGGGRLRRFRLGAAMAIIYMAMSTLFFLRIAPLTPWWGALAAWVMILLIVEAIFAFFFPYRCCFNETEGDVQNNPHV
IRIS_313-10758 MELITACALLLIAVHIAIVISCPEPLAAVALVDVPLLYLYRYLLIIRRGVIIVSGAGGGGGRLRRFRLGAAMAIIYMAMSTLFFLRIAPLTPWWGALAAWVMILLIVEAIFAFFFPYRCCFNETEGDVQNNPHV
IRIS_313-10759 MELITACALLLIAVHIAIVISCPEPLAAVALVDVPLLYLCRYLLIIRRGVIIVSGAGAGGGRLRRFRLGAAMAIIYMAMSTLFFLRIAPLTPWWGALAAWVMILLIVEAIFAFFFPYRCCFNETEGDVQNNPHV
IRIS_313-10760 MELITACALLLIAVHIAIVISCPEPLAAVALVDVPLLYLCRYLLIIRRGVIIVSGGGGGGRLRRFRLGAAMAIIYMAMSTLFFLRIAPLTPWWGALAAWVMILLIVEAIFAFFFPYRCCFNETEGDVQNNPHV
IRIS_313-10761 MELITACALLLIAVHIAIVISCPEPLAAVALVDVPLLYLCRYLLIIRRGVIIVSGAGAGGGRLRRFRLGAAMAIIYMAMSTLFFLRIAPLTPWWGALAAWVMILLIVEAIFAFFFPYRCCFNETEGDVQNNPHV
IRIS_313-10762 MELITACALLLIAVHIAIVISCPEPLAAVALVDVPLLYLCRYLLIIRRGVIIVSGAGAGGGRLRRFRLGAAMAIIYMAMSTLFFLRIAPLTPWWGALAAWVMILLIVEAIFAFFFPYRCCFNETEGDVQNNPHV
IRIS_313-10765 MELITACALLLIAVHIAIVISCPEPLAAVALVDVPLLYLCRYLLIIRRGVIIVSGAGAGGGRLRRFRLGAAMAIIYMVMSTLFFLRIAPLTPWWGALAAWVMILLIVEAIFAFFFPYRCCFNETEGDVQNNPHV
IRIS_313-10766 MELITACALLLIAVHIAIVISCPEPLAAVALVDVPLLYLCRYLLIIRRGVIIVSGAGAGGGRLRRFRLGAAMAIIYMAMSTLFFLRIAPLTPWWGALAAWVMILLIVEAIFAFFFPYRCCFNETEGDVQNNPHV
IRIS_313-10767 MELITACALLLIAVHIAIVISCPEPLAAVALVDVPLLYLYRYLLIIRRGVIIVSGAGGGGGRLRRFRLGAAMAIIYMAMSTLFFLRIAPLTPWWGALAAWVMILLIVEAIFAFFFPYRCCFNETEGDVQNNPHV
IRIS_313-10768 MELITACALLLIAVHIAIVISCPEPLAAVALVDVPLLYLCRYLLIIRRGVIIVSGAGGGGGGRLRRFRLGAAMAIIYMAMSTLFFLRIAPLTPWWGALAAWVMILLIVEAIFAFFFPYRCCFNETEGDVQNNPHV
IRIS_313-10769 MELITACALLLIAVHIAIVISCPEPLAAVALVDVPLLYLCRYLLIIRRGVIIVSGGGGGGRLRRFRLGAAMAIIYMAMSTLFFLRIAPLTPWWGALAAWVMILLIVEAIFAFFFPYRCCFNETEGDVQNNPHV
IRIS_313-10770 MELITACALLLIAVHIAIVISCPEPLAAVALVDVPLLYLYRYLLIIRRGVIIVSGAGGGGGRLRRFRLGAAMAIIYMAMSTLFFLRIAPLTPWWGALAAWVMILLIVEAIFAFFFPYRCCFNETEGDVQNNPHV
IRIS_313-10771 MELITACALLLIAVHIAIVISCPEPLAAVALVDVPLLYLCRYLLIIRRGVIIVSGAGAGGGRLRRFRLGAAMAIIYMAMSTLFFLRIAPLTPWWGALAAWVMILLIVEAIFAFFFPYRCCFNETEGDVQNNPHV
IRIS_313-10772 MELITACALLLIAVHIAIVISCPEPLAAVALVDVPLLYLCRYLLIIRRGVIIVSGGGGGGRLRRFRLGAAMAIIYMAMSTLFFLRIAPLTPWWGALAAWVMILLIVEAIFAFFFPYRCCFNETEGDVQNNPHV
IRIS_313-10773 MELITACALLLIAVHIAIVISCPEPLAAVALVDVPLLYLCRYLLIIRRGVIIVSGAGAGGGRLRRFRLGAAMAIIYMAMSTLFFLRIAPLTPWWGALAAWVMILLIVEAIFAFFFPYRCCFNETEGDVQNNPHV
IRIS_313-10774 MELITACALLLIAVHIAIVISCPEPLAAVALVDVPLLYLCRYLLIIRRGVIIVSGGGAGGGRLRRFRLGAAMAIIYMAMSTLFFLRIAPLTPWWGALAAWVMILLIVEAIFAFFFPYRCCFNETEGDVQNNPHV
IRIS_313-10775 MELITACALLLIAVHIAIVISCPEPLAAVALVDVPLLYLCRYLLIIRRGVIIVSGGGGGGRLRRFRLGAAMAIIYMAMSTLFFLRIAPLTPWWGALAAWVMILLIVEAIFAFFFPYRCCFNETEGDVQNNPHV
IRIS_313-10776 MELITACALLLIAVHIAIVISCPEPLAAVALVDVPLLYLCRYLLIIRRGVIIVSGAGAGGGRLRRFRLGAAMAIIYMAMSTLFFLRIAPLTPWWGALAAWVMILLIVEAIFAFFFPYRCCFNETEGDVQNNPHV
IRIS_313-10777 MELITACALLLIAVHIAIVISCPEPLAAVALVDVPLLYLCRYLLIIRRGVIIVSGGGGGGRLRRFRLGAAMAIIYMAMSTLFFLRIAPLTPWWGALAAWVMILLIVEAIFAFFFPYRCCFNETEGDVQNNPHV
IRIS_313-10778 MELITACALLLIAVHIAIVISCPEPLAAVALVDVPLLYLCRYLLIIRRGVIIVSGAGAGGGRLRRFRLGAAMAIIYMAMSTLFFLRIAPLTPWWGALAAWVMILLIVEAIFAFFFPYRCCFNETEGDVQNNPHV
IRIS_313-10779 MELITACALLLIAVHIAIVISCPEPLAAVALVDVPLLYLCRYLLIIRRGVIIVSGAGGAAAAGCAASASGQPWRSSTWPCRPSSSSALLLSRHGGAPSPPGS*SCSSSRPSSPSSSPTAAASTKPKAMSRIIPM
IRIS_313-10780 MELITACALLLIAVHIAIVISCPEPLAAVALVDVPLLYLCRYLLIIRRGVIIVSGAGAGGGRLRRFRLGAAMAIIYMAMSTLFFLRIAPLTPWWGALAAWVMILLIVEAIFAFFFPYRCCFNETEGDVQNNPHV
IRIS_313-10781 MELITACALLLIAVHIAIVISCPEPLAAVALVDVPLLYLYRYLLIIRRGVIIVSGAGGGGGRLRRFRLGAAMAIIYMAMSTLFFLRIAPLTPWWGALAAWVMILLIVEAIFAFFFPYRCCFNETEGDVQNNPHV
IRIS_313-10782 MELITACALLLIAVHIAIVISCPEPLAAVALVDVPLLYLCRYLLIIRRGVIIVSGAGAGGGRLRRFRLGAAMAIIYMAMSTLFFLRIAPLTPWWGALAAWVMILLIVEAIFAFFFPYRCCFNETEGDVQNNPHV
IRIS_313-10783 MELITACALLLIAVHIAIVISCPEPLAAVALVDVPLLYLYRYLLIIRRGVIIVSGAGGGGGRLRRFRLGAAMAIIYMAMSTLFFLRIAPLTPWWGALAAWVMILLIVEAIFAFFFPYRCCFNETEGDVQNNPHV
IRIS_313-10784 MELITACALLLIAVHIAIVISCPEPLAAVALVDVPLLYLYRYLLIIRRGVIIVSGAGGGGGRLRRFRLGAAMAIIYMAMSTLFFLRIAPLTPWWGALAAWVMILLIVEAIFAFFFPYRCCFNETEGDVQNNPHV
IRIS_313-10785 MELITACALLLIAVHIAIVISCPEPLAAVALVDVPLLYLYRYLLIIRRGVIIVSGAGGGGGRLRRFRLGAAMAIIYMAMSTLFFLRIAPLTPWWGALAAWVMILLIVEAIFAFFFPYRCCFNETEGDVQNNPHV
IRIS_313-10786 MELITACALLLIAVHIAIVISCPEPLAAVALVDVPLLYLCRYLLIIRRGVIIVSGAGGAAAAGCAASASGQPWRSSTWPCRPSSSSALLLSRHGGAPSPPGS*SCSSSRPSSPSSSPTAAASTKPKAMSRIIPM
IRIS_313-10787 MELITACALLLIAVHIAIVISCPEPLAAVALVDVPLLYLCRYLLIIRRGVIIVSGAGGGGGRLRRFRLGAAMAIIYMAMSTLFFLRIAPLTPWWGALAAWVMILLIVEAIFAFFFPYRCCFNETEGDVQNNPHV
IRIS_313-10788 MELITACALLLIAVHIAIVISCPEPLAAVALVDVPLLYLYRYLLIIRRGVIIVSGAGGGGGRLRRFRLGAAMAIIYMAMSTLFFLRIAPLTPWWGALAAWVMILLIVEAIFAFFFPYRCCFNETEGDVQNNPHV
IRIS_313-10789 MELITACALLLIAVHIAIVISCPEPLAAVALVDVPLLYLCRYLLIIRRGVIIVSGAGAGGGRLRRFRLGAAMAIIYMAMSTLFFLRIAPLTPWWGALAAWVMILLIVEAIFAFFFPYRCCFNETEGDVQNNPHV
IRIS_313-10790 MELITACALLLIAVHIAIVISCPEPLAAVALVDVPLLYLCRYLLIIRRGVIIVSGAGAGGGRLRRFRLGAAMAIIYMAMSTLFFLRIAPLTPWWGALAAWVMILLIVEAIFAFFFPYRCCFNETEGDVQNNPHV
IRIS_313-10791 MELITACALLLIAVHIAIVISCPEPLAAVALVDVPLLYLCRYLLIIRRGVIIVSGGGGGGRLRRFRLGAAMAIIYMAMSTLFFLRIAPLTPWWGALAAWVMILLIVEAIFAFFFPYRCCFNETEGDVQNNPHV
IRIS_313-10793 MELITACALLLIAVHIAIVISCPEPLAAVALVDVPLLYLCRYLLIIRRGVIIVSGAGAGGGRLRRFRLGAAMAIIYMAMSTLFFLRIAPLTPWWGALAAWVMILLIVEAIFAFFFPYRCCFNETEGDVQNNPHV
IRIS_313-10794 MELITACALLLIAVHIAIVISCPEPLAAVALVDVPLLYLYRYLLIIRRGVIIVSGAGGGGGRLRRFRLGAAMAIIYMAMSTLFFLRIAPLTPWWGALAAWVMILLIVEAIFAFFFPYRCCFNETEGDVQNNPHV
IRIS_313-10795 MELITACALLLIAVHIAIVISCPEPLAAVALVDVPLLYLCRYLLIIRRGVIIVSGAGAGGGRLRRFRLGAAMAIIYMAMSTLFFLRIAPLTPWWGALAAWVMILLIVEAIFAFFFPYRCCFNETEGDVQNNPHV
IRIS_313-10796 MELITACALLLIAVHIAIVISCPEPLAAVALVDVPLLYLCRYLLIIRRGVIIVSGAGAGGGRLRRFRLGAAMAIIYMAMSTLFFLRIAPLTPWWGALAAWVMILLIVEAIFAFFFPYRCCFNETEGDVQNNPHV
IRIS_313-10797 MELITACALLLIAVHIAIVISCPEPLAAVALVDVPLLYLCRYLLIIRRGVIIVSGGGGGGRLRRFRLGAAMAIIYMAMSTLFFLRIAPLTPWWGALAAWVMILLIVEAIFAFFFPYRCCFNETEGDVQNNPHV
IRIS_313-10798 MELITACALLLIAVHIAIVISCPEPLAAVALVDVPLLYLCRYLLIIRRGVIIVSGAGAGGGRLRRFRLGAAMAIIYMAMSTLFFLRIAPLTPWWGALAAWVMILLIVEAIFAFFFPYRCCFNETEGDVQNNPHV
IRIS_313-10799 MELITACALLLIAVHIAIVISCPEPLAAVALVDVPLLYLYRYLLIIRRGVIIVSGAGGGGGRLRRFRLGAAMAIIYMAMSTLFFLRIAPLTPWWGALAAWVMILLIVEAIFAFFFPYRCCFNETEGDVQNNPHV
IRIS_313-10800 MELITACALLLIAVHIAIVISCPEPLAAVALVDVPLLYLCRYLLIIRRGVIIVSGAGAGGGRLRRFRLGAAMAIIYMAMSTLFFLRIAPLTPWWGALAAWVMILLIVEAIFAFFFPYRCCFNETEGDVQNNPHV
IRIS_313-10801 MELITACALLLIAVHIAIVISCPEPLAAVALVDVPLLYLCRYLLIIRRGVIIVSGAGAGGGRLRRFRLGAAMAIIYMAMSTLFFLRIAPLTPWWGALAAWVMILLIVEAIFAFFFPYRCCFNETEGDVQNNPHV
IRIS_313-10802 MELITACALLLIAVHIAIVISCPEPLAAVALVDVPLLYLCRYLLIIRRGVIIVSGAGAGGGRLRRFRLGAAMAIIYMAMSTLFFLRIAPLTPWWGALAAWVMILLIVEAIFAFFFPYRCCFNETEGDVQNNPHV
IRIS_313-10803 MELITACALLLIAVHIAIVISCPEPLAAVALVDVPLLYLCRYLLIIRRGVIIVSGAGAGGGRLRRFRLGAAMAIIYMAMSTLFFLRIAPLTPWWGALAAWVMILLIVEAIFAFFFPYRCCFNETEGDVQNNPHV
IRIS_313-10804 MELITACALLLIAVHIAIVISCPEPLAAVALVDVPLLYLCRYLLIIRRGVIIVSGAGGAAAAGCAASASGQPWRSSTWPCRPSSSSALLLSRHGGAPSPPGS*SCSSSRPSSPSSSPTAAASTKPKAMSRIIPM
IRIS_313-10805 MELITACALLLIAVHIAIVISCPEPLAAVALVDVPLLYLYRYLLIIRRGVIIVSGAGGGGGRLRRFRLGAAMAIIYMAMSTLFFLRIAPLTPWWGALAAWVMILLIVEAIFAFFFPYRCCFNETEGDVQNNPHV
IRIS_313-10806 MELITACALLLIAVHIAIVISCPEPLAAVALVDVPLLYLCRYLLIIRRGVIIVSGAGGGGGRLRRFRLGAAMAIIYMAMSTLFFLRIAPLTPWWGALAAWVMILLIVEAIFAFFFPYRCCFNETEGDVQNNPHV
IRIS_313-10807 MELITACALLLIAVHIAIVISCPEPLAAVALVDVPLLYLCRYLLIIRRGVIIVSGGGGGGRLRRFRLGAAMAIIYMAMSTLFFLRIAPLTPWWGALAAWVMILLIVEAIFAFFFPYRCCFNETEGDVQNNPHV
IRIS_313-10808 MELITACALLLIAVHIAIVISCPEPLAAVALVDVPLLYLCRYLLIIRRGVIIVSGAGAGGGRLRRFRLGAAMAIIYMAMSTLFFLRIAPLTPWWGALAAWVMILLIVEAIFAFFFPYRCCFNETEGDVQNNPHV
IRIS_313-10809 MELITACALLLIAVHIAIVISCPEPLAAVALVDVPLLYLYRYLLIIRRGVIIVSGAGGGGGRLRRFRLGAAMAIIYMAMSTLFFLRIAPLTPWWGALAAWVMILLIVEAIFAFFFPYRCCFNETEGDVQNNPHV
IRIS_313-10810 MELITACALLLIAVHIAIVISCPEPLAAVALVDVPLLYLCRYLLIIRRGVIIVSGGGGGGRLRRFRLGAAMAIIYMAMSTLFFLRIAPLTPWWGALAAWVMILLIVEAIFAFFFPYRCCFNETEGDVQNNPHV
IRIS_313-10811 MELITACALLLIAVHIAIVISCPEPLAAVALVDVPLLYLCRYLLIIRRGVIIVSGGGGGGRLRRFRLGAAMAIIYMAMSTLFFLRIAPLTPWWGALAAWVMILLIVEAIFAFFFPYRCCFNETEGDVQNNPHV
IRIS_313-10812 MELITACALLLIAVHIAIVISCPEPLAAVALVDVPLLYLCRYLLIIRRGVIIVSGGGGGGRLRRFRLGAAMAIIYMAMSTLFFLRIAPLTPWWGALAAWVMILLIVEAIFAFFFPYRCCFNETEGDVQNNPHV
IRIS_313-10813 MELITACALLLIAVHIAIVISCPEPLAAVALVDVPLLYLCRYLLIIRRGVIIVSGAGGGGRLRRFRLGAAMAIIYMAMSTLFFLRIAPLTPWWGALAAWVMILLIVEAIFAFFFPYRCCFNETEGDVQNNPHV
IRIS_313-10814 MELITACALLLIAVHIAIVISCPEPLAAVALVDVPLLYLCRYLLIIRRGVIIVSGGGGGGRLRRFRLGAAMAIIYMAMSTLFFLRIAPLTPWWGALAAWVMILLIVEAIFAFFFPYRCCFNETEGDVQNNPHV
IRIS_313-10815 MELITACALLLIAVHIAIVISCPEPLAAVALVDVPLLYLYRYLLIIRRGVIIVSGAGGGGGRLRRFRLGAAMAIIYMAMSTLFFLRIAPLTPWWGALAAWVMILLIVEAIFAFFFPYRCCFNETEGDVQNNPHV
IRIS_313-10816 MELITACALLLIAVHIAIVISCPEPLAAVALVDVPLLYLYRYLLIIRRGVIIVSGAGAGGGRLRRFRLGAAMAIIYMAMSTLFFLRIAPLTPWWGALAAWVMILLIVEAIFAFFFPYRCCFNETEGDVQNNPHV
IRIS_313-10817 MELITACALLLIAVHIAIVISCPEPLAAVALVDVPLLYLCRYLLIIRRGVIIVSGAGGGGGRLRRFRLGAAMAIIYMAMSTLFFLRIAPLTPWWGALAAWVMILLIVEAIFAFFFPYRCCFNETEGDVQNNPHV
IRIS_313-10818 MELITACALLLIAVHIAIVISCPEPLAAVALVDVPLLYLCRYLLIIRRGVIIVSGGGGGGRLRRFRLGAAMAIIYMAMSTLFFLRIAPLTPWWGALAAWVMILLIVEAIFAFFFPYRCCFNETEGDVQNNPHV
IRIS_313-10819 MELITACALLLIAVHIAIVISCPEPLAAVALVDVPLLYLCRYLLIIRRGVIIVSGGGGGGRLRRFRLGAAMAIIYMAMSTLFFLRIAPLTPWWGALAAWVMILLIVEAIFAFFFPYRCCFNETEGDVQNNPHV
IRIS_313-10820 MELITACALLLIAVHIAIVISCPEPLAAVALVDVPLLYLCRYLLIIRRGVIIVSGGGGGGRLRRFRLGAAMAIIYMAMSTLFFLRIAPLTPWWGALAAWVMILLIVEAIFAFFFPYRCCFNETEGDVQNNPHV
IRIS_313-10821 MELITACALLLIAVHIAIVISCPEPLAAVALVDVPLLYLCRYLLIIRRGVIIVSGGGGGGRLRRFRLGAAMAIIYMAMSTLFFLRIAPLTPWWGALAAWVMILLIVEAIFAFFFPYRCCFNETEGDVQNNPHV
IRIS_313-10822 MELITACALLLIAVHIAIVISCPEPLAAVALVDVPLLYLCRYLLIIRRGVIIVSGGGGGGRLRRFRLGAAMAIIYMAMSTLFFLRIAPLTPWWGALAAWVMILLIVEAIFAFFFPYRCCFNETEGDVQNNPHV
IRIS_313-10823 MELITACALLLIAVHIAIVISCPEPLAAVALVDVPLLYLCRYLLIIRRGVIIVSGGGGGGRLRRFRLGAAMAIIYMAMSTLFFLRIAPLTPWWGALAAWVMILLIVEAIFAFFFPYRCCFNETEGDVQNNPHV
IRIS_313-10824 MELITACALLLIAVHIAIVISCPEPLAAVALVDVPLLYLCRYLLIIRRGVIIVSGAGGGGRLRRFRLGAAMAIIYMAMSTLFFLRIAPLTPWWGALAAWVMILLIVEAIFAFFFPYRCCFNETEGDVQNNPHV
IRIS_313-10825 MELITACALLLIAVHIAIVISCPEPLAAVALVDVPLLYLCRYLLIIRRGVIIVSGAGAGGGRLRRFRLGAAMAIIYMAMSTLFFLRIAPLTPWWGALAAWVMILLIVEAIFAFFFPYRCCFNETEGDVQNNPHV
IRIS_313-10827 MELITACALLLIAVHIAIVISCPEPLAAVALVDVPLLYLCRYLLIIRRGVIIVSGAGAGGGRLRRFRLGAAMAIIYMAMSTLFFLRIAPLTPWWGALAAWVMILLIVEAIFAFFFPYRCCFNETEGDVQNNPHV

IRIS_313-10828 MELITACALLLIAVHIAIVISCPEPLAAVALVDVPLLYLCRYLLIIRRGVIIVSGAGGGGGRLRRFRLGAAMAIIYMAMSTLFFLRIAPLTPWWGALAAWVMILLIVEAIFAFFFPYRCCFNETEGDVQNNPHV
IRIS_313-10829 MELITACALLLIAVHIAIVISCPEPLAAVALVDVPLLYLCRYLLIIRRGVIIVSGAGGGGGRLRRFRLGAAMAIIYMAMSTLFFLRIAPLTPWWGALAAWVMILLIVEAIFAFFFPYRCCFNETEGDVQNNPHV
IRIS_313-10830 MELITACALLLIAVHIAIVISCPEPLAAVALVDVPLLYLCRYLLIIRRGVIIVSGAGGGGGRLRRFRLGAAMAIIYMAMSTLFFLRIAPLTPWWGALAAWVMILLIVEAIFAFFFPYRCCFNETEGDVQNNPHV
IRIS_313-10831 MELITACALLLIAVHIAIVISCPEPLAAVALVDVPLLYLCRYLLIIRRGVIIVSGAGAGGGRLRRFRLGAAMAIIYMAMSTLFFLRIAPLTPWWGALAAWVMILLIVEAIFAFFFPYRCCFNETEGDVQNNPHV
IRIS_313-10832 MELITACALLLIAVHIAIVISCPEPLAAVALVDVPLLYLYRYLLIIRRGVIIVSGAGGGGGRLRRFRLGAAMAIIYMAMSTLFFLRIAPLTPWWGALAAWVMILLIVEAIFAFFFPYRCCFNETEGDVQNNPHV
IRIS_313-10833 MELITACALLLIAVHIAIVISCPEPLAAVALVDVPLLYLCRYLLIIRRGVIIVSGAGGGGGRLRRFRLGAAMAIIYMAMSTLFFLRIAPLTPWWGALAAWVMILLIVEAIFAFFFPYRCCFNETEGDVQNNPHV
IRIS_313-10834 MELITACALLLIAVHIAIVISCPEPLAAVALVDVPLLYLCRYLLIIRRGVIIVSGAGAGGGRLRRFRLGAAMAIIYMAMSTLFFLRIAPLTPWWGALAAWVMILLIVEAIFAFFFPYRCCFNETEGDVQNNPHV
IRIS_313-10835 MELITACALLLIAVHIAIVISCPEPLAAVALVDVPLLYLCRYLLIIRRGVIIVSGGGGGGRLRRFRLGAAMAIIYMAMSTLFFLRIAPLTPWWGALAAWVMILLIVEAIFAFFFPYRCCFNETEGDVQNNPHV
IRIS_313-10836 MELITACALLLIAVHIAIVISCPEPLAAVALVDVPLLYLCRYLLIIRRGVIIVSGGGGGGRLRRFRLGAAMAIIYMAMSTLFFLRIAPLTPWWGALAAWVMILLIVEAIFAFFFPYRCCFNETEGDVQNNPHV
IRIS_313-10837 MELITACALLLIAVHIAIVISCPEPLAAVALVDVPLLYLCRYLLIIRRGVIIVSGAGAGGGRLRRFRLGAAMAIIYMAMSTLFFLRIAPLTPWWGALAAWVMILLIVEAIFAFFFPYRCCFNETEGDVQNNPHV
IRIS_313-10838 MELITACALLLIAVHIAIVISCPEPLAAVALVDVPLLYLCRYLLIIRRGVIIVSGAGAGGGRLRRFRLGAAMAIIYMAMSTLFFLRIAPLTPWWGALAAWVMILLIVEAIFAFFFPYRCCFNETEGDVQNNPHV
IRIS_313-10839 MELITACALLLIAVHIAIVISCPEPLAAVALVDVPLLYLCRYLLIIRRGVIIVSGAGAGGGRLRRFRLGAAMAIIYMAMSTLFFLRIAPLTPWWGALAAWVMILLIVEAIFAFFFPYRCCFNETEGDVQNNPHV
IRIS_313-10840 MELITACALLLIAVHIAIVISCPEPLAAVALVDVPLLYLCRYLLIIRRGVIIVSGAGAGGGRLRRFRLGAAMAIIYMAMSTLFFLRIAPLTPWWGALAAWVMILLIVEAIFAFFFPYRCCFNETEGDVQNNPHV
IRIS_313-10841 MELITACALLLIAVHIAIVISCPEPLAAVALVDVPLLYLYRYLLIIRRGVIIVSGAGGGGGRLRRFRLGAAMAIIYMVMSTLFFLRIAPLTPWWGALAAWVMILLIVEAIFAFFFPYRCCFNETEGDVQNNPHV
IRIS_313-10842 MELITACALLLIAVHIAIVISCPEPLAAVALVDVPLLYLCRYLLIIRRGVIIVSGAGGGGGGRLRRFRLGAAMAIIYMAMSTLFFLRIAPLTPWWGALAAWVMILLIVEAIFAFFFPYRCCFNETEGDVQNNPHV
IRIS_313-10844 MELITACALLLIAVHIAIVISCPEPLAAVALVDVPLLYLCRYLLIIRRGVIIVSGAGAGGGRLRRFRLGAAMAIIYMAMSTLFFLRIAPLTPWWGALAAWVMILLIVEAIFAFFFPYRCCFNETEGDVQNNPHV
IRIS_313-10845 MELITACALLLIAVHIAIVISCPEPLAAVALVDVPLLYLCRYLLIIRRGVIIVSGAGAGGGRLRRFRLGAAMAIIYMAMSTLFFLRIAPLTPWWGALAAWVMILLIVEAIFAFFFPYRCCFNETEGDVQNNPHV
IRIS_313-10847 MELITACALLLIAVHIAIVISCPEPLAAVALVDVPLLYLCRYLLIIRRGVIIVSGGGGGGRLRRFRLGAAMAIIYMAMSTLFFLRIAPLTPWWGALAAWVMILLIVEAIFAFFFPYRCCFNETEGDVQNNPHV
IRIS_313-10848 MELITACALLLIAVHIAIVISCPEPLAAVALVDVPLLYLYRYLLIIRRGVIIVSGAGGGGGRLRRFRLGAAMAIIYMAMSTLFFLRIAPLTPWWGALAAWVMILLIVEAIFAFFFPYRCCFNETEGDVQNNPHV
IRIS_313-10849 MELITACALLLIAVHIAIVISCPEPLAAVAFVNVPLLYLCRYLLIIRRGVIIVSGGGGGGGRLRRFRLGAAMAVIYMAMSTLFFLRIAPLTPWWGALAAWVMILLIVEAIFAFFFPYRCCFNETEGDVQNNPHV
IRIS_313-10850 MELITACALLLIAVHIAIVISCPEPLAAVALVDVPLLYLCRYLLIIRRGVIIVSGAGAGGGRLRRFRLGAAMAIIYMAMSTLFFLRIAPLTPWWGALAAWVMILLIVEAIFAFFFPYRCCFNETEGDVQNNPHV
IRIS_313-10851 MELITACALLLIAVHIAIVISCPEPLAAVALVDVPLLYLCRYLLIIRRGVIIVSGAGAGGGRLRRFRLGAAMAIIYMAMSTLFFLRIAPLTPWWGALAAWVMILLIVEAIFAFFFPYRCCFNETEGDVQNNPHV
IRIS_313-10852 MELITACALLLIAVHIAIVISCPEPLAAVAFVNVPLLYLCRYLLIIRRGVIIVSGGGGGGGRLRRFRLGAAMAVIYMAMSTLFFLRIAPLTPWWGALAAWVMILLIVEAIFAFFFPYRCCFNETEGDVQNNPHV
IRIS_313-10853 MELITACALLLIAVHIAIVISCPEPLAAVALVDVPLLYLYRYLLIIRRGVIIVSGAGGGGGRLRRFRLGAAMAIIYMAMSTLFFLRIAPLTPWWGALAAWVMILLIVEAIFAFFFPYRCCFNETEGDVQNNPHV
IRIS_313-10854 MELITACALLLIAVHIAIVISCPEPLAAVAFVNVPLLYLCRYLLIIRRGVIIVSGGGGGGGRLRRFRLGAAMAVIYMAMSTLFFLRIAPLTPWWGALAAWVMILLIVEAIFAFFFPYRCCFNETEGDVQNNPHV
IRIS_313-10855 MELITACALLLIAVHIAIVISCPEPLAAVALVDVPLLYLCRYLLIIRRGVIIVSGAGGAAAAGCAASASGQPWRSSTWPCRPSSSSALLLSRHGGAPSPPGS*SCSSSRPSSPSSSPTAAASTKPKAMSRIIPM
IRIS_313-10856 MELITACALLLIAVHIAIVISCPEPLAAVAFVNVPLLYLCRYLLIIRRGVIIVSGGGAGGGRLRRFRLGAAMAVIYMAMSTLFFLRIAPLTPWWGALAAWVMILLIVEAIFAFFFPYRCCFNETEGDVQNNPHV
IRIS_313-10857 MELITACALLLIAVHIAIVISCPEPLAAVALVDVPLLYLCRYLLIIRRGVIIVSGGGGGGRLRRFRLGAAMAIIYMAMSTLFFLRIAPLTPWWGALAAWVMILLIVEAIFAFFFPYRCCFNETEGDVQNNPHV
IRIS_313-10858 MELITACALLLIAVHIAIVISCPEPLAAVALVDVPLLYLCRYLLIIRRGVIIVSGGGGGGRLRRFRLGAAMAIIYMAMSTLFFLRIAPLTPWWGALAAWVMILLIVEAIFAFFFPYRCCFNETEGDVQNNPHV
IRIS_313-10859 MELITACALLLIAVHIAIVISCPEPLAAVALVDVPLLYLCRYLLIIRRGVIIVSGAGAGGGRLRRFRLGAAMAIIYMAMSTLFFLRIAPLTPWWGALAAWVMILLIVEAIFAFFFPYRCCFNETEGDVQNNPHV
IRIS_313-10861 MELITACALLLIAVHIAIVISCPEPLAAVAFVNVPLLYLCRYLLIIRRGVIIVSGGGGGGGRLRRFRLGAAMAVIYMAMSTLFFLRIAPLTPWWGALAAWVMILLIVEAIFAFFFPYRCCFNETEGDVQNNPHV
IRIS_313-10862 MELITACALLLIAVHIAIVISCPEPLAAVALVDVPLLYLCRYLLIIRRGVIIVSGAGGGGGRLRRFRLGAAMAIIYMAMSTLFFLRIAPLTPWWGALAAWVMILLIVEAIFAFFFPYRCCFNETEGDVQNNPHV
IRIS_313-10863 MELITACALLLIAVHIAIVISCPEPLAAVALVDVPLLYLCRYLLIIRRGVIIVSGGGGGGRLRRFRLGAAMAIIYMAMSTLFFLRIAPLTPWWGALAAWVMILLIVEAIFAFFFPYRCCFNETEGDVQNNPHV
IRIS_313-10864 MELITACALLLIAVHIAIVISCPEPLAAVALVDVPLLYLCRYLLIIRRGVIIVSGAGAGGGRLRRFRLGAAMAIIYMAMSTLFFLRIAPLTPWWGALAAWVMILLIVEAIFAFFFPYRCCFNETEGDVQNNPHV
IRIS_313-10865 MELITACALLLIAVHIAIVISCPEPLAAVALVDVPLLYLYRYLLIIRRGVIIVSGAGGGGGRLRRFRLGAAMAIIYMAMSTLFFLRIAPLTPWWGALAAWVMILLIVEAIFAFFFPYRCCFNETEGDVQNNPHV
IRIS_313-10866 MELITACALLLIAVHIAIVISCPEPLAAVALVDVPLLYLCRYLLIIRRGVIIVSGAGAGGGRLRRFRLGAAMAIIYMAMSTLFFLRIAPLTPWWGALAAWVMILLIVEAIFAFFFPYRCCFNETEGDVQNNPHV
IRIS_313-10867 MELITACALLLIAVHIAIVISCPEPLAAVALVDVPLLYLYRYLLIIRRGVIIVSGAGGGGGRLRRFRLGAAMAIIYMAMSTLFFLRIAPLTPWWGALAAWVMILLIVEAIFAFFFPYRCCFNETEGDVQNNPHV
IRIS_313-10868 MELITACALLLIAVHIAIVISCPEPLAAVALVDVPLLYLCRYLLIIRRGVIIVSGAGAGGGRLRRFRLGAAMAIIYMAMSTLFFLRIAPLTPWWGALAAWVMILLIVEAIFAFFFPYRCCFNETEGDVQNNPHV
IRIS_313-10869 MELITACALLLIAVHIAIVISCPEPLAAVALVDVPLLYLCRYLLIIRRGVIIVSGAGAGGGRLRRFRLGAAMAIIYMAMSTLFFLRIAPLTPWWGALAAWVMILLIVEAIFAFFFPYRCCFNETEGDVQNNPHV
IRIS_313-10870 MELITACALLLIAVHIAIVISCPEPLAAVALVDVPLLYLCRYLLIIRRGVIIVSGAGGGGGRLRRFRLGAAMAIIYMAMSTLFFLRIAPLTPWWGALAAWVMILLIVEAIFAFFFPYRCCFNETEGDVQNNPHV
IRIS_313-10871 MELITACALLLIAVHIAIVISCPEPLAAVAFVNVPLLYLCRYLLIIRRGVIIVSGGGGGGGRLRRFRLGAAMAVIYMAMSTLFFLRIAPLTPWWGALAAWVMILLIVEAIFAFFFPYRCCFNETEGDVQNNPHV
IRIS_313-10872 MELITACALLLIAVHIAIVISCPEPLAAVALVDVPLLYLYRYLLIIRRGVIIVSGAGGGGGRLRRFRLGAAMAIIYMAMSTLFFLRIAPLTPWWGALAAWVMILLIVEAIFAFFFPYRCCFNETEGDVQNNPHV
IRIS_313-10873 MELITACALLLIAVHIAIVISCPEPLAAVALVDVPLLYLCRYLLIIRRGVIIVSGAGAGGGRLRRFRLGAAMAIIYMAMSTLFFLRIAPLTPWWGALAAWVMILLIVEAIFAFFFPYRCCFNETEGDVQNNPHV
IRIS_313-10874 MELITACALLLIAVHIAIVISCPEPLAAVALVDVPLLYLYRYLLIIRRGVIIVSGAGGGGGRLRRFRLGAAMAIIYMAMSTLFFLRIAPLTPWWGALAAWVMILLIVEAIFAFFFPYRCCFNETEGDVQNNPHV
IRIS_313-10875 MELITACALLLIAVHIAIVISCPEPLAAVALVDVPLLYLCRYLLIIRRGVIIVSGAGGGGGRLRRFRLGAAMAIIYMAMSTLFFLRIAPLTPWWGALAAWVMILLIVEAIFAFFFPYRCCFNETEGDVQNNPHV
IRIS_313-10876 MELITACALLLIAVHIAIVISCPEPLAAVAFVNVPLLYLCRYLLIIRRGVIIVSGGGGGGGRLRRFRLGAAMAVIYMAMSTLFFLRIAPLTPWWGALAAWVMILLIVEAIFAFFFPYRCCFNETEGDVQNNPHV
IRIS_313-10877 MELITACALLLIAVHIAIVISCPEPLAAVALVDVPLLYLCRYLLIIRRGVIIVSGAGAGGGRLRRFRLGAAMAIIYMAMSTLFFLRIAPLTPWWGALAAWVMILLIVEAIFAFFFPYRCCFNETEGDVQNNPHV
IRIS_313-10878 MELITACALLLIAVHIAIVISCPEPLAAVALVDVPLLYLCRYLLIIRRGVIIVSGAGAGGGRLRRFRLGAAMAIIYMAMSTLFFLRIAPLTPWWGALAAWVMILLIVEAIFAFFFPYRCCFNETEGDVQNNPHV
IRIS_313-10879 MELITACALLLIAVHIAIVISCPEPLAAVALVDVPLLYLCRYLLIIRRGVIIVSGAGAGGGRLRRFRLGAAMAIIYMAMSTLFFLRIAPLTPWWGALAAWVMILLIVEAIFAFFFPYRCCFNETEGDVQNNPHV
IRIS_313-10880 MELITACALLLIAVHIAIVISCPEPLAAVALVDVPLLYLCRYLLIIRRGVIIVSGGGGGGRLRRFRLGAAMAIIYMAMSTLFFLRIAPLTPWWGALAAWVMILLIVEAIFAFFFPYRCCFNETEGDVQNNPHV
IRIS_313-10881 MELITACALLLIAVHIAIVISCPEPLAAVALVDVPLLYLCRYLLIIRRGVIIVSGAGAGGGRLRRFRLGAAMAIIYMAMSTLFFLRIAPLTPWWGALAAWVMILLIVEAIFAFFFPYRCCFNETEGDVQNNPHV
IRIS_313-10882 MELITACALLLIAVHIAIVISCPEPLAAVALVDVPLLYLCRYLLIIRRGVIIVSGAGAGGGRLRRFRLGAAMAIIYMAMSTLFFLRIAPLTPWWGALAAWVMILLIVEAIFAFFFPYRCCFNETEGDVQNNPHV
IRIS_313-10883 MELITACALLLIAVHIAIVISCPEPLAAVAFVNVPLLYLCRYLLIIRRGVIIVSGGGGGGGRLRRFRLGAAMAVIYMAMSTLFFLRIAPLTPWWGALAAWVMILLIVEAIFAFFFPYRCCFNETEGDVQNNPHV
IRIS_313-10884 MELITACALLLIAVHIAIVISCPEPLAAVALVDVPLLYLYRYLLIIRRGVIIVSGAGGGGGRLRRFRLGAAMAIIYMAMSTLFFLRIAPLTPWWGALAAWVMILLIVEAIFAFFFPYRCCFNETEGDVQNNPHV
IRIS_313-10885 MELITACALLLIAVHIAIVISCPEPLAAVALVDVPLLYLCRYLLIIRRGVIIVSGGGGGGRLRRFRLGAAMAIIYMAMSTLFFLRIAPLTPWWGALAAWVMILLIVEAIFAFFFPYRCCFNETEGDVQNNPHV
IRIS_313-10886 MELITACALLLIAVHIAIVISCPEPLAAVALVDVPLLYLCRYLLIIRRGVIIVSGAGAGGGRLRRFRLGAAMAIIYMAMSTLFFLRIAPLTPWWGALAAWVMILLIVEAIFAFFFPYRCCFNETEGDVQNNPHV
IRIS_313-10887 MELITACALLLIAVHIAIVISCPEPLAAVALVDVPLLYLCRYLLIIRRGVIIVSGGGGGGRLRRFRLGAAMAIIYMAMSTLFFLRIAPLTPWWGALAAWVMILLIVEAIFAFFFPYRCCFNETEGDVQNNPHV
IRIS_313-10888 MELITACALLLIAVHIAIVISCPEPLAAVALVDVPLLYLYRYLLIIRRGVIIVSGAGGGGGRLRRFRLGAAMAIIYMAMSTLFFLRIAPLTPWWGALAAWVMILLIVEAIFAFFFPYRCCFNETEGDVQNNPHV
IRIS_313-10889 MELITACALLLIAVHIAIVISCPEPLAAVALVDVPLLYLYRYLLIIRRGVIIVSGAGGGGGRLRRFRLGAAMAIIYMAMSTLFFLRIAPLTPWWGALAAWVMILLIVEAIFAFFFPYRCCFNETEGDVQNNPHV
IRIS_313-10890 MELITACALLLIAVHIAIVISCPEPLAAVALVDVPLLYLCRYLLIIRRGVIIVSGAGGGGGRLRRFRLGAAMAIIYMAMSTLFFLRIAPLTPWWGALAAWVMILLIVEAIFAFFFPYRCCFNETEGDVQNNPHV
IRIS_313-10891 MELITACALLLIAVHIAIVISCPEPLAAVAFVNVPLLYLCRYLLIIRRGVIIVSGGGGGGGRLRRFRLGAAMAVIYMAMSTLFFLRIAPLTPWWGALAAWVMILLIVEAIFAFFFPYRCCFNETEGDVQNNPHV
IRIS_313-10892 MELITACALLLIAVHIAIVISCPEPLAAVALVDVPLLYLCRYLLIIRRGVIIVSGAGAGGGRLRRFRLGAAMAIIYMAMSTLFFLRIAPLTPWWGALAAWVMILLIVEAIFAFFFPYRCCFNETEGDVQNNPHV
IRIS_313-10893 MELITACALLLIAVHIAIVISCPEPLAAVALVDVPLLYLCRYLLIIRRGVIIVSGAGAGGGRLRRFRLGAAMAIIYMAMSTLFFLRIAPLTPWWGALAAWVMILLIVEAIFAFFFPYRCCFNETEGDVQNNPHV
IRIS_313-10894 MELITACALLLIAVHIAIVISCPEPLAAVALVDVPLLYLCRYLLIIRRGVIIVSGAGAGGGRLRRFRLGAAMAIIYMAMSTLFFLRIAPLTPWWGALAAWVMILLIVEAIFAFFFPYRCCFNETEGDVQNNPHV
IRIS_313-10895 MELITACALLLIAVHIAIVISCPEPLAAVALVDVPLLYLYRYLLIIRRGVIIVSGAGGGGGRLRRFRLGAAMAIIYMAMSTLFFLRIAPLTPWWGALAAWVMILLIVEAIFAFFFPYRCCFNETEGDVQNNPHV
IRIS_313-10896 MELITACALLLIAVHIAIVISCPEPLAAVALVDVPLLYLCRYLLIIRRGVIIVSGAGAGGGRLRRFRLGAAMAIIYMAMSTLFFLRIAPLTPWWGALAAWVMILLIVEAIFAFFFPYRCCFNETEGDVQNNPHV
IRIS_313-10897 MELITACALLLIAVHIAIVISCPEPLAAVALVDVPLLYLYRYLLIIRRGVIIVSGAGGGGGRLRRFRLGAAMAIIYMAMSTLFFLRIAPLTPWWGALAAWVMILLIVEAIFAFFFPYRCCFNETEGDVQNNPHV
IRIS_313-10898 MELITACALLLIAVHIAIVISCPEPLAAVALVDVPLLYLYRYLLIIRRGVIIVSGAGGGGGRLRRFRLGAAMAIIYMAMSTLFFLRIAPLTPWWGALAAWVMILLIVEAIFAFFFPYRCCFNETEGDVQNNPHV
IRIS_313-10899 MELITACALLLIAVHIAIVISCPEPLAAVALVDVPLLYLYRYLLIIRRGVIIVSGAGGGGGRLRRFRLGAAMAIIYMAMSTLFFLRIAPLTPWWGALAAWVMILLIVEAIFAFFFPYRCCFNETEGDVQNNPHV
IRIS_313-10900 MELITACALLLIAVHIAIVISCPEPLAAVALVDVPLLYLYRYLLIIRRGVIIVSGAGGGGGRLRRFRLGAAMAIIYMAMSTLFFLRIAPLTPWWGALAAWVMILLIVEAIFAFFFPYRCCFNETEGDVQNNPHV
IRIS_313-10901 MELITACALLLIAVHIAIVISCPEPLAAVALVDVPLLYLCRYLLIIRRGVIIVSGAGAGGGRLRRFRLGAAMAIIYMAMSTLFFLRIAPLTPWWGALAAWVMILLIVEAIFAFFFPYRCCFNETEGDVQNNPHV
IRIS_313-10902 MELITACALLLIAVHIAIVISCPEPLAAVALVDVPLLYLYRYLLIIRRGVIIVSGGGAGGGRLRRFRLGAAMAIIYMAMSTLFFLRIAPLTPWWGALAAWVMILLIVEAIFAFFFPYRCCFNETEGDVQNNPHV
IRIS_313-10903 MELITACALLLIAVHIAIVISCPEPLAAVALVDVPLLYLCRYLLIIRRGVIIVSGAGAGGGRLRRFRLGAAMAIIYMAMSTLFFLRIAPLTPWWGALAAWVMILLIVEAIFAFFFPYRCCFNETEGDVQNNPHV
IRIS_313-10904 MELITACALLLIAVHIAIVISCPEPLAAVALVDVPLLYLCRYLLIIRRGVIIVSGAGAGGGRLRRFRLGAAMAIIYMAMSTLFFLRIAPLTPWWGALAAWVMILLIVEAIFAFFFPYRCCFNETEGDVQNNPHV
IRIS_313-10905 MELITACALLLIAVHIAIVISCPEPLAAVALVDVPLLYLCRYLLIIRRGVIIVSGAGAGGGRLRRFRLGAAMAIIYMAMSTLFFLRIAPLTPWWGALAAWVMILLIVEAIFAFFFPYRCCFNETEGDVQNNPHV
IRIS_313-10906 MELITACALLLIAVHIAIVISCPEPLAAVALVDVPLLYLCRYLLIIRRGVIIVSGGGGGGRLRRFRLGAAMAIIYMAMSTLFFLRIAPLTPWWGALAAWVMILLIVEAIFAFFFPYRCCFNETEGDVQNNPHV
IRIS_313-10907 MELITACALLLIAVHIAIVISCPEPLAAVAFVNVPLLYLCRYLLIIRRGVIIVSGGSGGGGRLRRFRLGAAMAVIYMAMSTLFFLRIAPLTPWWGALAAWVMILLIVEAIFAFFFPYRCCFNETEGDVQNNPHV
IRIS_313-10908 MELITACALLLIAVHIAIVISCPEPLAAVALVDVPLLYLYRYLLIIRRGVIIVSGAGGGGGRLRRFRLGAAMAIIYMAMSTLFFLRIAPLTPWWGALAAWVMILLIVEAIFAFFFPYRCCFNETEGDVQNNPHV
IRIS_313-10909 MELITACALLLIAVHIAIVISCPEPLAAVALVDVPLLYLYRYLLIIRRGVIIVSGAGGGGGRLRRFRLGAAMAIIYMAMSTLFFLRIAPLTPWWGALAAWVMILLIVEAIFAFFFPYRCCFNETEGDVQNNPHV
IRIS_313-10910 MELITACALLLIAVHIAIVISCPEPLAAVALVDVPLLYLCRYLLIIRRGVIIVSGAGAGGGRLRRFRLGAAMAIIYMVMSTLFFLRIAPLTPWWGALAAWVMILLIVEAIFAFFFPYRCCFNETEGDVQNNPHV
IRIS_313-10911 MELITACALLLIAVHIAIVISCPEPLAAVAFVNVPLLYLCRYLLIIRRGVIIVSGGSAGGGRLRRFRLGAAMAVIYMAMSTLFFLRIAPLTPWWGALAAWVMILLIVEAIFAFFFPYRCCFNETEGDVQNNPHV
IRIS_313-10912 MELITACALLLIAVHIAIVISCPEPLAAVALVDVPLLYLYRYLLIIRRGVIIVSGAGGGGGRLRRFRLGAAMAIIYMAMSTLFFLRIAPLTPWWGALAAWVMILLIVEAIFAFFFPYRCCFNETEGDVQNNPHV
IRIS_313-10913 MELITACALLLIAVHIAIVISCPEPLAAVALVDVPLLYLYRYLLIIRRGVIIVSGAGGGGGRLRRFRLGAAMAIIYMAMSTLFFLRIAPLTPWWGALAAWVMILLIVEAIFAFFFPYRCCFNETEGDVQNNPHV
IRIS_313-10915 MELITACALLLIAVHIAIVISCPEPLAAVALVDVPLLYLCRYLLIIRRGVIIVSGGGGGGRLRRFRLGAAMAIIYMAMSTLFFLRIAPLTPWWGALAAWVMILLIVEAIFAFFFPYRCCFNETEGDVQNNPHV
IRIS_313-10916 MELITACALLLIAVHIAIVISCPEPLAAVALVDVPLLYLYRYLLIIRRGVIIVSGAGGGGGRLRRFRLGAAMAIIYMAMSTLFFLRIAPLTPWWGALAAWVMILLIVEAIFAFFFPYRCCFNETEGDVQNNPHV
IRIS_313-10917 MELITACALLLIAVHIAIVISCPEPLAAVALVDVPLLYLCRYLLIIRRGVIIVSGAGGGGGRLRRFRLGAAMAIIYMAMSTLFFLRIAPLTPWWGALAAWVMILLIVEAIFAFFFPYRCCFNETEGDVQNNPHV
IRIS_313-10918 MELITACALLLIAVHIAIVISCPEPLAAVALVDVPLLYLCRYLLIIRRGVIIVSGAGAGGGRLRRFRLGAAMAIIYMAMSTLFFLRIAPLTPWWGALAAWVMILLIVEAIFAFFFPYRCCFNETEGDVQNNPHV
IRIS_313-10919 MELITACALLLIAVHIAIVISCPEPLAAVALVDVPLLYLYRYLLIIRRGVIIVSGAGGGGGRLRRFRLGAAMAIIYMAMSTLFFLRIAPLTPWWGALAAWVMILLIVEAIFAFFFPYRCCFNETEGDVQNNPHV
IRIS_313-10920 MELITACALLLIAVHIAIVISCPEPLAAVALVDVPLLYLYRYLLIIRRGVIIVSGAGGGGGRLRRFRLGAAMAIIYMAMSTLFFLRIAPLTPWWGALAAWVMILLIVEAIFAFFFPYRCCFNETEGDVQNNPHV
IRIS_313-10921 MELITACALLLIAVHIAIVISCPEPLAAVALVDVPLLYLYRYLLIIRRGVIIVSGAGGGGGRLRRFRLGAAMAIIYMAMSTLFFLRIAPLTPWWGALAAWVMILLIVEAIFAFFFPYRCCFNETEGDVQNNPHV
IRIS_313-10922 MELITACALLLIAVHIAIVISCPEPLAAVALVDVPLLYLYRYLLIIRRGVIIVSGAGGGGGRLRRFRLGAAMAIIYMAMSTLFFLRIAPLTPWWGALAAWVMILLIVEAIFAFFFPYRCCFNETEGDVQNNPHV
IRIS_313-10923 MELITACALLLIAVHIAIVISCPEPLAAVALVDVPLLYLYRYLLIIRRGVIIVSGAGAGGGRLRRFRLGAAMAIIYMAMSTLFFLRIAPLTPWWGALAAWVMILLIVEAIFAFFFPYRCCFNETEGDVQNNPHV
IRIS_313-10924 MELITACALLLIAVHIAIVISCPEPLAAVALVDVPLLYLYRYLLIIRRGVIIVSGAGAGGGRLRRFRLGAAMAIIYMAMSTLFFLRIAPLTPWWGALAAWVMILLIVEAIFAFFFPYRCCFNETEGDVQNNPHV
IRIS_313-10925 MELITACALLLIAVHIAIVISCPEPLAAVAFVNVPLLYLCRYLLIIRRGVIIVSGGGGGGGRLRRFRLGAAMAVIYMAMSTLFFLRIAPLTPWWGALAAWVMILLIVEAIFAFFFPYRCCFNETEGDVQNNPHV
IRIS_313-10926 MELITACALLLIAVHIAIVISCPEPLAAVALVDVPLLYLCRYLLIIRRGVIIVSGAGAGGGRLRRFRLGAAMAIIYMAMSTLFFLRIAPLTPWWGALAAWVMILLIVEAIFAFFFPYRCCFNETEGDVQNNPHV
IRIS_313-10927 MELITACALLLIAVHIAIVISCPEPLAAVALVDVPLLYLCRYLLIIRRGVIIVSGAGAGGGRLRRFRLGAAMAIIYMAMSTLFFLRIAPLTPWWGALAAWVMILLIVEAIFAFFFPYRCCFNETEGDVQNNPHV
IRIS_313-10928 MELITACALLLIAVHIAIVISCPEPLAAVALVDVPLLYLYRYLLIIRRGVIIVSGAGGGGGRLRRFRLGAAMAIIYMAMSTLFFLRIAPLTPWWGALAAWVMILLIVEAIFAFFFPYRCCFNETEGDVQNNPHV
IRIS_313-10929 MELITACALLLIAVHIAIVISCPEPLAAVALVDVPLLYLCRYLLIIRRGVIIVSGAGGAAAAGCAASASGQPWRSSTWPCRPSSSSALLLSRHGGAPSPPGS*SCSSSRPSSPSSSPTAAASTKPKAMSRIIPM
IRIS_313-10930 MELITACALLLIAVHIAIVISCPEPLAAVAFVNVPLLYLCRYLLIIRRGVIIVSGGGGGGGRLRRFRLGAAMAVIYMAMSTLFFLRIAPLTPWWGALAAWVMILLIVEAIFAFFFPYRCCFNETEGDVQNNPHV
IRIS_313-10931 MELITACALLLIAVHIAIVISCPEPLAAVALVDVPLLYLCRYLLIIRRGVIIVSGAGAGGGRLRRFRLGAAMAIIYMAMSTLFFLRIAPLTPWWGALAAWVMILLIVEAIFAFFFPYRCCFNETEGDVQNNPHV
IRIS_313-10932 MELITACALLLIAVHIAIVISCPEPLAAVALVDVPLLYLYRYLLIIRRGVIIVSGAGGGGGRLRRFRLGAAMAIIYMAMSTLFFLRIAPLTPWWGALAAWVMILLIVEAIFAFFFPYRCCFNETEGDVQNNPHV
IRIS_313-10933 MELITACALLLIAVHIAIVISCPEPLAAVALVDVPLLYLCRYLLIIRRGVIIVSGAGGAAAAGCAASASGQPWRSSTWPCRPSSSSALLLSRHGGAPSPPGS*SCSSSRPSSPSSSPTAAASTKPKAMSRIIPM
IRIS_313-10934 MELITACALLLIAVHIAIVISCPEPLAAVALVDVPLLYLCRYLLIIRRGVIIVSGGGGGGRLRRFRLGAAMAIIYMAMSTLFFLRIAPLTPWWGALAAWVMILLIVEAIFAFFFPYRCCFNETEGDVQNNPHV
IRIS_313-10935 MELITACALLLIAVHIAIVISCPEPLAAVALVDVPLLYLCRYLLIIRRGVIIVSGGGGGGRLRRFRLGAAMAIIYMAMSTLFFLRIAPLTPWWGALAAWVMILLIVEAIFAFFFPYRCCFNETEGDVQNNPHV
IRIS_313-10936 MELITACALLLIAVHIAIVISCPEPLAAVALVDVPLLYLYRYLLIIRRGVIIVSGAGGGGGRLRRFRLGAAMAIIYMAMSTLFFLRIAPLTPWWGALAAWVMILLIVEAIFAFFFPYRCCFNETEGDVQNNPHV
IRIS_313-10937 MELITACALLLIAVHIAIVISCPEPLAAVALVDVPLLYLCRYLLIIRRGVIIVSGGGGGGRLRRFRLGAAMAIIYMAMSTLFFLRIAPLTPWWGALAAWVMILLIVEAIFAFFFPYRCCFNETEGDVQNNPHV
IRIS_313-10938 MELITACALLLIAVHIAIVISCPEPLAAVALVDVPLLYLCRYLLIIRRGVIIVSGGGGGGRLRRFRLGAAMAIIYMAMSTLFFLRIAPLTPWWGALAAWVMILLIVEAIFAFFFPYRCCFNETEGDVQNNPHV
IRIS_313-10940 MELITACALLLIAVHIAIVISCPEPLAAVALVDVPLLYLCRYLLIIRRGVIIVSGGGGGGRLRRFRLGAAMAIIYMAMSTLFFLRIAPLTPWWGALAAWVMILLIVEAIFAFFFPYRCCFNETEGDVQNNPHV
IRIS_313-10941 MELITACALLLIAVHIAIVISCPEPLAAVALVDVPLLYLCRYLLIIRRGVIIVSGGGGGGRLRRFRLGAAMAIIYMAMSTLFFLRIAPLTPWWGALAAWVMILLIVEAIFAFFFPYRCCFNETEGDVQNNPHV
IRIS_313-10942 MELITACALLLIAVHIAIVISCPEPLAAVALVDVPLLYLCRYLLIIRRGVIIVSGAGAGGGRLRRFRLGAAMAIIYMAMSTLFFLRIAPLTPWWGALAAWVMILLIVEAIFAFFFPYRCCFNETEGDVQNNPHV
IRIS_313-10943 MELITACALLLIAVHIAIVISCPEPLAAVALVDVPLLYLCRYLLIIRRGVIIVSGAGGAAAAGCAASASGQPWRSSTWPCRPSSSSALLLSRHGGAPSPPGS*SCSSSRPSSPSSSPTAAASTKPKAMSRIIPM
IRIS_313-10944 MELITACALLLIAVHIAIVISCPEPLAAVALVDVPLLYLCRYLLIIRRGVIIVSGGGGGGRLRRFRLGAAMAIIYMAMSTLFFLRIAPLTPWWGALAAWVMILLIVEAIFAFFFPYRCCFNETEGDVQNNPHV
IRIS_313-10945 MELITACALLLIAVHIAIVISCPEPLAAVALVDVPLLYLCRYLLIIRRGVIIVSGAGGGGGRLRRFRLGAAMAIIYMAMSTLFFLRIAPLTPWWGALAAWVMILLIVEAIFAFFFPYRCCFNETEGDVQNNPHV
IRIS_313-10946 MELITACALLLIAVHIAIVISCPEPLAAVALVDVPLLYLCRYLLIIRRGVIIVSGAGAGGGRLRRFRLGAAMAIIYMAMSTLFFLRIAPLTPWWGALAAWVMILLIVEAIFAFFFPYRCCFNETEGDVQNNPHV
IRIS_313-10947 MELITACALLLIAVHIAIVISCPEPLAAVALVDVPLLYLYRYLLIIRRGVIIVSGAGGGGGRLRRFRLGAAMAIIYMAMSTLFFLRIAPLTPWWGALAAWVMILLIVEAIFAFFFPYRCCFNETEGDVQNNPHV
IRIS_313-10948 MELITACALLLIAVHIAIVISCPEPLAAVALVDVPLLYLYRYLLIIRRGVIIVSGAGGGGGRLRRFRLGAAMAIIYMAMSTLFFLRIAPLTPWWGALAAWVMILLIVEAIFAFFFPYRCCFNETEGDVQNNPHV
IRIS_313-10949 MELITACALLLIAVHIAIVISCPEPLAAVALVDVPLLYLCRYLLIIRRGVIIVSGAGAGGGRLRRFRLGAAMAIIYMAMSTLFFLRIAPLTPWWGALAAWVMILLIVEAIFAFFFPYRCCFNETEGDVQNNPHV
IRIS_313-10950 MELITACALLLIAVHIAIVISCPEPLAAVALVDVPLLYLYRYLLIIRRGVIIVSGAGGGGGRLRRFRLGAAMAIIYMAMSTLFFLRIAPLTPWWGALAAWVMILLIVEAIFAFFFPYRCCFNETEGDVQNNPHV
IRIS_313-10951 MELITACALLLIAVHIAIVISCPEPLAAVALVDVPLLYLCRYLLIIRRGVIIVSGAGGGGGGRLRRFRLGAAMAIIYMAMSTLFFLRIAPLTPWWGALAAWVMILLIVEAIFAFFFPYRCCFNETEGDVQNNPHV
IRIS_313-10952 MELITACALLLIAVHIAIVISCPEPLAAVALVDVPLLYLYRYLLIIRRGVIIVSGAGGGGGRLRRFRLGAAMAIIYMAMSTLFFLRIAPLTPWWGALAAWVMILLIVEAIFAFFFPYRCCFNETEGDVQNNPHV
IRIS_313-10953 MELITACALLLIAVHIAIVISCPEPLAAVALVDVPLLYLCRYLLIIRRGVIIVSGAGAGGGRLRRFRLGAAMAIIYMAMSTLFFLRIAPLTPWWGALAAWVMILLIVEAIFAFFFPYRCCFNETEGDVQNNPHV
IRIS_313-10954 MELITACALLLIAVHIAIVISCPEPLAAVALVDVPLLYLCRYLLIIRRGVIIVSGGGGGGRLRRFRLGAAMAIIYMAMSTLFFLRIAPLTPWWGALAAWVMILLIVEAIFAFFFPYRCCFNETEGDVQNNPHV
IRIS_313-10955 MELITACALLLIAVHIAIVISCPEPLAAVALVDVPLLYLCRYLLIIRRGVIIVSGGGGGGRLRRFRLGAAMAIIYMAMSTLFFLRIAPLTPWWGALAAWVMILLIVEAIFAFFFPYRCCFNETEGDVQNNPHV

IRIS_313-10956 MELITACALLLIAVHIAIVISCPEPLAAVALVDVPLLYLYRYLLIIRRGVIIVSGAGGGGGRLRRFRLGAAMAIIYMAMSTLFFLRIAPLTPWWGALAAWVMILLIVEAIFAFFFPYRCCFNETEGDVQNNPHV
IRIS_313-10957 MELITACALLLIAVHIAIVISCPEPLAAVALVDVPLLYLCRYLLIIRRGVIIVSGAGGGGGRLRRFRLGAAMAIIYMAMSTLFFLRIAPLTPWWGALAAWVMILLIVEAIFAFFFPYRCCFNETEGDVQNNPHV
IRIS_313-10958 MELITACALLLIAVHIAIVISCPEPLAAVALVDVPLLYLCRYLLIIRRGVIIVSGAGAGGGRLRRFRLGAAMAIIYMAMSTLFFLRIAPLTPWWGALAAWVMILLIVEAIFAFFFPYRCCFNETEGDVQNNPHV
IRIS_313-10959 MELITACALLLIAVHIAIVISCPEPLAAVALVDVPLLYLYRYLLIIRRGVIIVSGAGGGGGRLRRFRLGAAMAIIYMAMSTLFFLRIAPLTPWWGALAAWVMILLIVEAIFAFFFPYRCCFNETEGDVQNNPHV
IRIS_313-10960 MELITACALLLIAVHIAIVISCPEPLAAVALVDVPLLYLYRYLLIIRRGVIIVSGAGGGGGRLRRFRLGAAMAIIYMAMSTLFFLRIAPLTPWWGALAAWVMILLIVEAIFAFFFPYRCCFNETEGDVQNNPHV
IRIS_313-10961 MELITACALLLIAVHIAIVISCPEPLAAVALVDVPLLYLCRYLLIIRRGVIIVSGGGGGGRLRRFRLGAAMAIIYMAMSTLFFLRIAPLTPWWGALAAWVMILLIVEAIFAFFFPYRCCFNETEGDVQNNPHV
IRIS_313-10962 MELITACALLLIAVHIAIVISCPEPLAAVALVDVPLLYLCRYLLIIRRGVIIVSGGGAGGGRLRRFRLGAAMAIIYMAMSTLFFLRIAPLTPWWGALAAWVMILLIVEAIFAFFFPYRCCFNETEGDVQNNPHV
IRIS_313-10963 MELITACALLLIAVHIAIVISCPEPLAAVAFVNVPLLYLCRYLLIIRRGVIIVSGGGGGGGRLRRFRLGAAMAVIYMAMSTLFFLRIAPLTPWWGALAAWVMILLIVEAIFAFFFPYRCCFNETEGDVQNNPHV
IRIS_313-10964 MELITACALLLIAVHIAIVISCPEPLAAVAFVNVPLLYLCRYLLIIRRGVIIVSGGGGGGGRLRRFRLGAAMAVIYMAMSTLFFLRIAPLTPWWGALAAWVMILLIVEAIFAFFFPYRCCFNETEGDVQNNPHV
IRIS_313-10965 MELITACALLLIAVHIAIVISCPEPLAAVAFVNVPLLYLCRYLLIIRRGVIIVSGGGGGGGRLRRFRLGAAMAVIYMAMSTLFFLRIAPLTPWWGALAAWVMILLIVEAIFAFFFPYRCCFNETEGDVQNNPHV
IRIS_313-10966 MELITACALLLIAVHIAIVISCPEPLAAVALVDVPLLYLCRYLLIIRRGVIIVSGAGGGGGGRLRRFRLGAAMAIIYMAMSTLFFLRIAPLTPWWGALAAWVMILLIVEAIFAFFFPYRCCFNETEGDVQNNPHV
IRIS_313-10967 MELITACALLLIAVHIAIVISCPEPLAAVALVDVPLLYLCRYLLIIRRGVIIVSGAGAGGGRLRRFRLGAAMAIIYMAMSTLFFLRIAPLTPWWGALAAWVMILLIVEAIFAFFFPYRCCFNETEGDVQNNPHV
IRIS_313-10968 MELITACALLLIAVHIAIVISCPEPLAAVALVDVPLLYLYRYLLIIRRGVIIVSGAGGGGGRLRRFRLGAAMAIIYMAMSTLFFLRIAPLTPWWGALAAWVMILLIVEAIFAFFFPYRCCFNETEGDVQNNPHV
IRIS_313-10969 MELITACALLLIAVHIAIVISCPEPLAAVAFVNVPLLYLCRYLLIIRRGVIIVSGGGGGGGRLRRFRLGAAMAVIYMAMSTLFFLRIAPLTPWWGALAAWVMILLIVEAIFAFFFPYRCCFNETEGDVQNNPHV
IRIS_313-10970 MELITACALLLIAVHIAIVISCPEPLAAVALVDVPLLYLCRYLLIIRRGVIIVSGAGGAAAAGCAASASGQPWRSSTWPCRPSSSSALLLSRHGGAPSPPGS*SCSSSRPSSPSSSPTAAASTKPKAMSRIIPM
IRIS_313-10971 MELITACALLLIAVHIAIVISCPEPLAAVALVDVPLLYLCRYLLIIRRGVIIVSGAGAGGGRLRRFRLGAAMAIIYMAMSTLFFLRIAPLTPWWGALAAWVMILLIVEAIFAFFFPYRCCFNETEGDVQNNPHV
IRIS_313-10972 MELITACALLLIAVHIAIVISCPEPLAAVALVDVPLLYLYRYLLIIRRGVIIVSGAGGGGGRLRRFRLGAAMAIIYMAMSTLFFLRIAPLTPWWGALAAWVMILLIVEAIFAFFFPYRCCFNETEGDVQNNPHV
IRIS_313-10973 MELITACALLLIAVHIAIVISCPEPLAAVALVDVPLLYLCRYLLIIRRGVIIVSGAGAGGGRLRRFRLGAAMAIIYMAMSTLFFLRIAPLTPWWGALAAWVMILLIVEAIFAFFFPYRCCFNETEGDVQNNPHV
IRIS_313-10974 MELITACALLLIAVHIAIVISCPEPLAAVALVDVPLLYLCRYLLIIRRGVIIVSGAGAGGGRLRRFRLGAAMAIIYMAMSTLFFLRIAPLTPWWGALAAWVMILLIVEAIFAFFFPYRCCFNETEGDVQNNPHV
IRIS_313-10975 MELITACALLLIAVHIAIVISCPEPLAAVALVDVPLLYLCRYLLIIRRGVIIVSGAGGGGGGRLRRFRLGAAMAIIYMAMSTLFFLRIAPLTPWWGALAAWVMILLIVEAIFAFFFPYRCCFNETEGDVQNNPHV
IRIS_313-10976 MELITACALLLIAVHIAIVISCPEPLAAVALVDVPLLYLCRYLLIIRRGVIIVSGAGAGGGRLRRFRLGAAMAIIYMAMSTLFFLRIAPLTPWWGALAAWVMILLIVEAIFAFFFPYRCCFNETEGDVQNNPHV
IRIS_313-10977 MELITACALLLIAVHIAIVISCPEPLAAVALVDVPLLYLCRYLLIIRRGVIIVSGAGAGGGRLRRFRLGAAMAIIYMAMSTLFFLRIAPLTPWWGALAAWVMILLIVEAIFAFFFPYRCCFNETEGDVQNNPHV
IRIS_313-10978 MELITACALLLIAVHIAIVISCPEPLAAVALVDVPLLYLCRYLLIIRRGVIIVSGAGAGGGRLRRFRLGAAMAIIYMAMSTLFFLRIAPLTPWWGALAAWVMILLIVEAIFAFFFPYRCCFNETEGDVQNNPHV
IRIS_313-10979 MELITACALLLIAVHIAIVISCPEPLAAVALVDVPLLYLCRYLLIIRRGVIIVSGAGGGGGRLRRFRLGAAMAIIYMAMSTLFFLRIAPLTPWWGALAAWVMILLIVEAIFAFFFPYRCCFNETEGDVQNNPHV
IRIS_313-10980 MELITACALLLIAVHIAIVISCPEPLAAVALVDVPLLYLCRYLLIIRRGVIIVSGAGAGGGRLRRFRLGAAMAIIYMAMSTLFFLRIAPLTPWWGALAAWVMILLIVEAIFAFFFPYRCCFNETEGDVQNNPHV
IRIS_313-10981 MELITACALLLIAVHIAIVISCPEPLAAVALVDVPLLYLCRYLLIIRRGVIIVSGAGGGGGRLRRFRLGAAMAIIYMAMSTLFFLRIAPLTPWWGALAAWVMILLIVEAIFAFFFPYRCCFNETEGDVQNNPHV
IRIS_313-10982 MELITACALLLIAVHIAIVISCPEPLAAVALVDVPLLYLCRYLLIIRRGVIIVSGAGAGGGRLRRFRLGAAMAIIYMAMSTLFFLRIAPLTPWWGALAAWVMILLIVEAIFAFFFPYRCCFNETEGDVQNNPHV
IRIS_313-10983 MELITACALLLIAVHIAIVISCPEPLAAVALVDVPLLYLCRYLLIIRRGVIIVSGGGGGGRLRRFRLGAAMAIIYMAMSTLFFLRIAPLTPWWGALAAWVMILLIVEAIFAFFFPYRCCFNETEGDVQNNPHV
IRIS_313-10984 MELITACALLLIAVHIAIVISCPEPLAAVALVDVPLLYLCRYLLIIRRGVIIVSGAGAGGGRLRRFRLGAAMAIIYMAMSTLFFLRIAPLTPWWGALAAWVMILLIVEAIFAFFFPYRCCFNETEGDVQNNPHV
IRIS_313-10985 MELITACALLLIAVHIAIVISCPEPLAAVALVDVPLLYLCRYLLIIRRGVIIVSGAGAGGGRLRRFRLGAAMAIIYMAMSTLFFLRIAPLTPWWGALAAWVMILLIVEAIFAFFFPYRCCFNETEGDVQNNPHV
IRIS_313-10986 MELITACALLLIAVHIAIVISCPEPLAAVALVDVPLLYLCRYLLIIRRGVIIVSGAGAGGGRLRRFRLGAAMAIIYMAMSTLFFLRIAPLTPWWGALAAWVMILLIVEAIFAFFFPYRCCFNETEGDVQNNPHV
IRIS_313-10987 MELITACALLLIAVHIAIVISCPEPLAAVALVDVPLLYLCRYLLIIRRGVIIVSGAGAGGGRLRRFRLGAAMAIIYMAMSTLFFLRIAPLTPWWGALAAWVMILLIVEAIFAFFFPYRCCFNETEGDVQNNPHV
IRIS_313-10988 MELITACALLLIAVHIAIVISCPEPLAAVALVDVPLLYLCRYLLIIRRGVIIVSGGGGGGRLRRFRLGAAMAIIYMAMSTLFFLRIAPLTPWWGALAAWVMILLIVEAIFAFFFPYRCCFNETEGDVQNNPHV
IRIS_313-10989 MELITACALLLIAVHIAIVISCPEPLAAVALVDVPLLYLCRYLLIIRRGVIIVSGAGAGGGRLRRFRLGAAMAIIYMAMSTLFFLRIAPLTPWWGALAAWVMILLIVEAIFAFFFPYRCCFNETEGDVQNNPHV
IRIS_313-10990 MELITACALLLIAVHIAIVISCPEPLAAVALVDVPLLYLCRYLLIIRRGVIIVSGAGGGGGGRLRRFRLGAAMAIIYMAMSTLFFLRIAPLTPWWGALAAWVMILLIVEAIFAFFFPYRCCFNETEGDVQNNPHV
IRIS_313-10991 MELITACALLLIAVHIAIVISCPEPLAAVALVDVPLLYLCRYLLIIRRGVIIVSGAGGGGGRLRRFRLGAAMAIIYMAMSTLFFLRIAPLTPWWGALAAWVMILLIVEAIFAFFFPYRCCFNETEGDVQNNPHV
IRIS_313-10992 MELITACALLLIAVHIAIVISCPEPLAAVALVDVPLLYLCRYLLIIRRGVIIVSGAGAGGGRLRRFRLGAAMAIIYMAMSTLFFLRIAPLTPWWGALAAWVMILLIVEAIFAFFFPYRCCFNETEGDVQNNPHV
IRIS_313-10993 MELITACALLLIAVHIAIVISCPEPLAAVALVDVPLLYLCRYLLIIRRGVIIVSGAGAGGGRLRRFRLGAAMAIIYMAMSTLFFLRIAPLTPWWGALAAWVMILLIVEAIFAFFFPYRCCFNETEGDVQNNPHV
IRIS_313-10994 MELITACALLLIAVHIAIVISCPEPLAAVALVDVPLLYLYRYLLIIRRGVIIVSGAGGGGGRLRRFRLGAAMAIIYMAMSTLFFLRIAPLTPWWGALAAWVMILLIVEAIFAFFFPYRCCFNETEGDVQNNPHV
IRIS_313-10995 MELITACALLLIAVHIAIVISCPEPLAAVALVDVPLLYLYRYLLIIRRGVIIVSGAGGGGGRLRRFRLGAAMAIIYMAMSTLFFLRIAPLTPWWGALAAWVMILLIVEAIFAFFFPYRCCFNETEGDVQNNPHV
IRIS_313-10996 MELITACALLLIAVHIAIVISCPEPLAAVALVDVPLLYLCRYLLIIRRGVIIVSGGGGGGRLRRFRLGAAMAIIYMAMSTLFFLRIAPLTPWWGALAAWVMILLIVEAIFAFFFPYRCCFNETEGDVQNNPHV
IRIS_313-10997 MELITACALLLIAVHIAIVISCPEPLAAVALVDVPLLYLCRYLLIIRRGVIIVSGGGGGGRLRRFRLGAAMAIIYMAMSTLFFLRIAPLTPWWGALAAWVMILLIVEAIFAFFFPYRCCFNETEGDVQNNPHV
IRIS_313-10998 MELITACALLLIAVHIAIVISCPEPLAAVALVDVPLLYLCRYLLIIRRGVIIVSGAGAGGGRLRRFRLGAAMAIIYMAMSTLFFLRIAPLTPWWGALAAWVMILLIVEAIFAFFFPYRCCFNETEGDVQNNPHV
IRIS_313-10999 MELITACALLLIAVHIAIVISCPEPLAAVALVDVPLLYLCRYLLIIRRGVIIVSGAGGGGGRLRRFRLGAAMAIIYMAMSTLFFLRIAPLTPWWGALAAWVMILLIVEAIFAFFFPYRCCFNETEGDVQNNPHV
IRIS_313-11000 MELITACALLLIAVHIAIVISCPEPLAAVALVDVPLLYLCRYLLIIRRGVIIVSGGGGGGRLRRFRLGAAMAIIYMAMSTLFFLRIAPLTPWWGALAAWVMILLIVEAIFAFFFPYRCCFNETEGDVQNNPHV
IRIS_313-11001 MELITACALLLIAVHIAIVISCPEPLAAVALVDVPLLYLYRYLLIIRRGVIIVSGAGGGGGRLRRFRLGAAMAIIYMAMSTLFFLRIAPLTPWWGALAAWVMILLIVEAIFAFFFPYRCCFNETEGDVQNNPHV
IRIS_313-11003 MELITACALLLIAVHIAIVISCPEPLAAVALVDVPLLYLYRYLLIIRRGVIIVSGAGGGGGRLRRFRLGAAMAIIYMAMSTLFFLRIAPLTPWWGALAAWVMILLIVEAIFAFFFPYRCCFNETEGDVQNNPHV
IRIS_313-11004 MELITACALLLIAVHIAIVISCPEPLAAVALVDVPLLYLYRYLLIIRRGVIIVSGAGGGGGRLRRFRLGAAMAIIYMAMSTLFFLRIAPLTPWWGALAAWVMILLIVEAIFAFFFPYRCCFNETEGDVQNNPHV
IRIS_313-11005 MELITACALLLIAVHIAIVISCPEPLAAVALVDVPLLYLYRYLLIIRRGVIIVSGAGGGGGRLRRFRLGAAMAIIYMAMSTLFFLRIAPLTPWWGALAAWVMILLIVEAIFAFFFPYRCCFNETEGDVQNNPHV
IRIS_313-11006 MELITACALLLIAVHIAIVISCPEPLAAVALVDVPLLYLCRYLLIIRRGVIIVSGGGGGGRLRRFRLGAAMAIIYMAMSTLFFLRIAPLTPWWGALAAWVMILLIVEAIFAFFFPYRCCFNETEGDVQNNPHV
IRIS_313-11007 MELITACALLLIAVHIAIVISCPEPLAAVALVDVPLLYLYRYLLIIRRGVIIVSGAGGGGGRLRRFRLGAAMAIIYMAMSTLFFLRIAPLTPWWGALAAWVMILLIVEAIFAFFFPYRCCFNETEGDVQNNPHV
IRIS_313-11008 MELITACALLLIAVHIAIVISCPEPLAAVALVDVPLLYLYRYLLIIRRGVIIVSGAGGGGGRLRRFRLGAAMAIIYMAMSTLFFLRIAPLTPWWGALAAWVMILLIVEAIFAFFFPYRCCFNETEGDVQNNPHV
IRIS_313-11009 MELITACALLLIAVHIAIVISCPEPLAAVALVDVPLLYLYRYLLIIRRGVIIVSGAGGGGGRLRRFRLGAAMAIIYMAMSTLFFLRIAPLTPWWGALAAWVMILLIVEAIFAFFFPYRCCFNETEGDVQNNPHV
IRIS_313-11010 MELITACALLLIAVHIAIVISCPEPLAAVALVDVPLLYLCRYLLIIRRGVIIVSGGGGGGRLRRFRLGAAMAIIYMAMSTLFFLRIAPLTPWWGALAAWVMILLIVEAIFAFFFPYRCCFNETEGDVQNNPHV
IRIS_313-11011 MELITACALLLIAVHIAIVISCPEPLAAVALVDVPLLYLCRYLLIIRRGVIIVSGAGAGGGRLRRFRLGAAMAIIYMAMSTLFFLRIAPLTPWWGALAAWVMILLIVEAIFAFFFPYRCCFNETEGDVQNNPHV
IRIS_313-11012 MELITACALLLIAVHIAIVISCPEPLAAVALVDVPLLYLCRYLLIIRRGVIIVSGGGGGGRLRRFRLGAAMAIIYMAMSTLFFLRIAPLTPWWGALAAWVMILLIVEAIFAFFFPYRCCFNETEGDVQNNPHV
IRIS_313-11013 MELITACALLLIAVHIAIVISCPEPLAAVAFVNVPLLYLCRYLLIIRRGVIIVSGGGGGGGRLRRFRLGAAMAVIYMAMSTLFFLRIAPLTPWWGALAAWVMILLIVEAIFAFFFPYRCCFNETEGDVQNNPHV
IRIS_313-11014 MELITACALLLIAVHIAIVISCPEPLAAVALVDVPLLYLCRYLLIIRRGVIIVSGAGGGGGRLRRFRLGAAMAIIYMAMSTLFFLRIAPLTPWWGALAAWVMILLIVEAIFAFFFPYRCCFNETEGDVQNNPHV
IRIS_313-11015 MELITACALLLIAVHIAIVISCPEPLAAVALVDVPLLYLCRYLLIIRRGVIIVSGAGGGGGRLRRFRLGAAMAIIYMAMSTLFFLRIAPLTPWWGALAAWVMILLIVEAIFAFFFPYRCCFNETEGDVQNNPHV
IRIS_313-11016 MELITACALLLIAVHIAIVISCPEPLAAVALVDVPLLYLCRYLLIIRRGVIIVSGAGAGGGRLRRFRLGAAMAIIYMAMSTLFFLRIAPLTPWWGALAAWVMILLIVEAIFAFFFPYRCCFNETEGDVQNNPHV
IRIS_313-11017 MELITACALLLIAVHIAIVISCPEPLAAVAFVNVPLLYLCRYLLIIRRGVIIVSGGGGGGGRLRRFRLGAAMAVIYMAMSTLFFLRIAPLTPWWGALAAWVMILLIVEAIFAFFFPYRCCFNETEGDVQNNPHV
IRIS_313-11018 MELITACALLLIAVHIAIVISCPEPLAAVALVDVPLLYLCRYLLIIRRGVIIVSGAGAGGGRLRRFRLGAAMAIIYMAMSTLFFLRIAPLTPWWGALAAWVMILLIVEAIFAFFFPYRCCFNETEGDVQNNPHV
IRIS_313-11019 MELITACALLLIAVHIAIVISCPEPLAAVALVDVPLLYLCRYLLIIRRGVIIVSGAGAGGGRLRRFRLGAAMAIIYMAMSTLFFLRIAPLTPWWGALAAWVMILLIVEAIFAFFFPYRCCFNETEGDVQNNPHV
IRIS_313-11020 MELITACALLLIAVHIAIVISCPEPLAAVAFVNVPLLYLCRYLLIIRRGVIIVSGGGGGGGRLRRFRLGAAMAVIYMAMSTLFFLRIAPLTPWWGALAAWVMILLIVEAIFAFFFPYRCCFNETEGDVQNNPHV
IRIS_313-11021 MELITACALLLIAVHIAIVISCPEPLAAVALVDVPLLYLCRYLLIIRRGVIIVSGAGAGGGRLRRFRLGAAMAIIYMAMSTLFFLRIAPLTPWWGALAAWVMILLIVEAIFAFFFPYRCCFNETEGDVQNNPHV
IRIS_313-11022 MELITACALLLIAVHIAIVISCPEPLAAVALVDVPLLYLCRYLLIIRRGVIIVSGAGAGGGRLRRFRLGAAMAIIYMAMSTLFFLRIAPLTPWWGALAAWVMILLIVEAIFAFFFPYRCCFNETEGDVQNNPHV
IRIS_313-11023 MELITACALLLIAVHIAIVISCPEPLAAVALVDVPLLYLCRYLLIIRRGVIIVSGAGAGGGRLRRFRLGAAMAIIYMAMSTLFFLRIAPLTPWWGALAAWVMILLIVEAIFAFFFPYRCCFNETEGDVQNNPHV
IRIS_313-11024 MELITACALLLIAVHIAIVISCPEPLAAVAFVNVPLLYLCRYLLIIRRGVIIVSGGGGGGGRLRRFRLGAAMAVIYMAMSTLFFLRIAPLTPWWGALAAWVMILLIVEAIFAFFFPYRCCFNETEGDVQNNPHV
IRIS_313-11025 MELITACALLLIAVHIAIVISCPEPLAAVALVDVPLLYLCRYLLIIRRGVIIVSGAGAGGGRLRRFRLGAAMAIIYMAMSTLFFLRIAPLTPWWGALAAWVMILLIVEAIFAFFFPYRCCFNETEGDVQNNPHV
IRIS_313-11026 MELITACALLLIAVHIAIVISCPEPLAAVALVDVPLLYLCRYLLIIRRGVIIVSGAGAGGGRLRRFRLGAAMAIIYMAMSTLFFLRIAPLTPWWGALAAWVMILLIVEAIFAFFFPYRCCFNETEGDVQNNPHV
IRIS_313-11027 MELITACALLLIAVHIAIVISCPEPLAAVAFVNVPLLYLCRYLLIIRRGVIIVSGGGGGGGRLRRFRLGAAMAVIYMAMSTLFFLRIAPLTPWWGALAAWVMILLIVEAIFAFFFPYRCCFNETEGDVQNNPHV
IRIS_313-11028 MELITACALLLIAVHIAIVISCPEPLAAVAFVNVPLLYLCRYLLIIRRGVIIVSGGGGGGGRLRRFRLGAAMAVIYMAMSTLFFLRIAPLTPWWGALAAWVMILLIVEAIFAFFFPYRCCFNETEGDVQNNPHV
IRIS_313-11029 MELITACALLLIAVHIAIVISCPEPLAAVALVDVPLLYLCRYLLIIRRGVIIVSGAGAGGGRLRRFRLGAAMAIIYMAMSTLFFLRIAPLTPWWGALAAWVMILLIVEAIFAFFFPYRCCFNETEGDVQNNPHV
IRIS_313-11030 MELITACALLLIAVHIAIVISCPEPLAAVALVDVPLLYLCRYLLIIRRGVIIVSGAGAGGGRLRRFRLGAAMAIIYMAMSTLFFLRIAPLTPWWGALAAWVMILLIVEAIFAFFFPYRCCFNETEGDVQNNPHV
IRIS_313-11031 MELITACALLLIAVHIAIVISCPEPLAAVALVDVPLLYLCRYLLIIRRGVIIVSGGGAGGGRLRRFRLGAAMAIIYMAMSTLFFLRIAPLTPWWGALAAWVMILLIVEAIFAFFFPYRCCFNETEGDVQNNPHV
IRIS_313-11032 MELITACALLLIAVHIAIVISCPEPLAAVALVDVPLLYLCRYLLIIRRGVIIVSGAGAGGGRLRRFRLGAAMAIIYMAMSTLFFLRIAPLTPWWGALAAWVMILLIVEAIFAFFFPYRCCFNETEGDVQNNPHV
IRIS_313-11033 MELITACALLLIAVHIAIVISCPEPLAAVAFVNVPLLYLCRYLLIIRRGVIIVSGAGAGGGRLRRFRLGAAMAVIYMAMSTLFFLRIAPLTPWWGALAAWVMILLIVEAIFAFFFPYRCCFNETEGDVQNNPHV
IRIS_313-11034 MELITACALLLIAVHIAIVISCPEPLAAVAFVNVPLLYLCRYLLIIRRGVIIVSGGGGGGGRLRRFRLGAAMAVIYMAMSTLFFLRIAPLTPWWGALAAWVMILLIVEAIFAFFFPYRCCFNETEGDVQNNPHV
IRIS_313-11035 MELITACALLLIAVHIAIVISCPEPLAAVALVDVPLLYLCRYLLIIRRGVIIVSGGGGGGRLRRFRLGAAMAIIYMAMSTLFFLRIAPLTPWWGALAAWVMILLIVEAIFAFFFPYRCCFNETEGDVQNNPHV
IRIS_313-11036 MELITACALLLIAVHIAIVISCPEPLAAVALVDVPLLYLCRYLLIIRRGVIIVSGAGAGGGRLRRFRLGAAMAIIYMAMSTLFFLRIAPLTPWWGALAAWVMILLIVEAIFAFFFPYRCCFNETEGDVQNNPHV
IRIS_313-11037 MELITACALLLIAVHIAIVISCPEPLAAVAFVNVPLLYLCRYLLIIRRGVIIVSGGGGGGGRLRRFRLGAAMAVIYMAMSTLFFLRIAPLTPWWGALAAWVMILLIVEAIFAFFFPYRCCFNETEGDVQNNPHV
IRIS_313-11038 MELITACALLLIAVHIAIVISCPEPLAAVALVDVPLLYLYRYLLIIRRGVIIVSGAGGGGGRLRRFRLGAAMAIIYMAMSTLFFLRIAPLTPWWGALAAWVMILLIVEAIFAFFFPYRCCFNETEGDVQNNPHV
IRIS_313-11039 MELITACALLLIAVHIAIVISCPEPLAAVALVDVPLLYLCRYLLIIRRGVIIVSGAGAGGGRLRRFRLGAAMAIIYMAMSTLFFLRIAPLTPWWGALAAWVMILLIVEAIFAFFFPYRCCFNETEGDVQNNPHV
IRIS_313-11040 MELITACALLLIAVHIAIVISCPEPLAAVALVDVPLLYLCRYLLIIRRGVIIVSGGGGGGRLRRFRLGAAMAIIYMAMSTLFFLRIAPLTPWWGALAAWVMILLIVEAIFAFFFPYRCCFNETEGDVQNNPHV
IRIS_313-11041 MELITACALLLIAVHIAIVISCPEPLAAVALVDVPLLYLCRYLLIIRRGVIIVSGAGGGGRLRRFRLGAAMAIIYMAMSTLFFLRIAPLTPWWGALAAWVMILLIVEAIFAFFFPYRCCFNETEGDVQNNPHV
IRIS_313-11042 MELITACALLLIAVHIAIVISCPEPLAAVALVDVPLLYLCRYLLIIRRGVIIVSGAGAGGGRLRRFRLGAAMAIIYMAMSTLFFLRIAPLTPWWGALAAWVMILLIVEAIFAFFFPYRCCFNETEGDVQNNPHV
IRIS_313-11043 MELITACALLLIAVHIAIVISCPEPLAAVALVDVPLLYLCRYLLIIRRGVIIVSGAGAGGGRLRRFRLGAAMAIIYMAMSTLFFLRIAPLTPWWGALAAWVMILLIVEAIFAFFFPYRCCFNETEGDVQNNPHV
IRIS_313-11044 MELITACALLLIAVHIAIVISCPEPLAAVALVDVPLLYLYRYLLIIRRGVIIVSGAGGGGGRLRRFRLGAAMAIIYMAMSTLFFLRIAPLTPWWGALAAWVMILLIVEAIFAFFFPYRCCFNETEGDVQNNPHV
IRIS_313-11045 MELITACALLLIAVHIAIVISCPEPLAAVALVDVPLLYLCRYLLIIRRGVIIVSGAGAGGGRLRRFRLGAAMAIIYMAMSTLFFLRIAPLTPWWGALAAWVMILLIVEAIFAFFFPYRCCFNETEGDVQNNPHV
IRIS_313-11046 MELITACALLLIAVHIAIVISCPEPLAAVALVDVPLLYLYRYLLIIRRGVIIVSGAGGGGGRLRRFRLGAAMAIIYMAMSTLFFLRIAPLTPWWGALAAWVMILLIVEAIFAFFFPYRCCFNETEGDVQNNPHV
IRIS_313-11047 MELITACALLLIAVHIAIVISCPEPLAAVALVDVPLLYLCRYLLIIRRGVIIVSGAGAGGGRLRRFRLGAAMAIIYMAMSTLFFLRIAPLTPWWGALAAWVMILLIVEAIFAFFFPYRCCFNETEGDVQNNPHV
IRIS_313-11048 MELITACALLLIAVHIAIVISCPEPLAAVAFVNVPLLYLCRYLLIIRRGVIIVSGGGGGGGRLRRFRLGAAMAVIYMAMSTLFFLRIAPLTPWWGALAAWVMILLIVEAIFAFFFPYRCCFNETEGDVQNNPHV
IRIS_313-11049 MELITACALLLIAVHIAIVISCPEPLAAVALVDVPLLYLCRYLLIIRRGVIIVSGAGAGGGRLRRFRLGAAMAIIYMAMSTLFFLRIAPLTPWWGALAAWVMILLIVEAIFAFFFPYRCCFNETEGDVQNNPHV
IRIS_313-11050 MELITACALLLIAVHIAIVISCPEPLAAVALVDVPLLYLCRYLLIIRRGVIIVSGAGAGGGRLRRFRLGAAMAIIYMAMSTLFFLRIAPLTPWWGALAAWVMILLIVEAIFAFFFPYRCCFNETEGDVQNNPHV
IRIS_313-11051 MELITACALLLIAVHIAIVISCPEPLAAVAFVNVPLLYLCRYLLIIRRGVIIVSGGGGGGGRLRRFRLGAAMAVIYMAMSTLFFLRIAPLTPWWGALAAWVMILLIVEAIFAFFFPYRCCFNETEGDVQNNPHV
IRIS_313-11052 MELITACALLLIAVHIAIVISCPEPLAAVAFVNVPLLYLCRYLLIIRRGVIIVSGGGGGGGRLRRFRLGAAMAVIYMAMSTLFFLRIAPLTPWWGALAAWVMILLIVEAIFAFFFPYRCCFNETEGDVQNNPHV
IRIS_313-11053 MELITACALLLIAVHIAIVISCPEPLAAVAFVNVPLLYLCRYLLIIRRGVIIVSGGGGGGGRLRRFRLGAAMAVIYMAMSTLFFLRIAPLTPWWGALAAWVMILLIVEAIFAFFFPYRCCFNETEGDVQNNPHV
IRIS_313-11054 MELITACALLLIAVHIAIVISCPEPLAAVAFVNVPLLYLCRYLLIIRRGVIIVSGGGGGGGRLRRFRLGAAMAVIYMAMSTLFFLRIAPLTPWWGALAAWVMILLIVEAIFAFFFPYRCCFNETEGDVQNNPHV
IRIS_313-11055 MELITACALLLIAVHIAIVISCPEPLAAVAFVNVPLLYLCRYLLIIRRGVIIVSGGGGGGGRLRRFRLGAAMAVIYMAMSTLFFLRIAPLTPWWGALAAWVMILLIVEAIFAFFFPYRCCFNETEGDVQNNPHV
IRIS_313-11056 MELITACALLLIAVHIAIVISCPEPLAAVALVDVPLLYLCRYLLIIRRGVIIVSGAGAGGGRLRRFRLGAAMAIIYMAMSTLFFLRIAPLTPWWGALAAWVMILLIVEAIFAFFFPYRCCFNETEGDVQNNPHV
IRIS_313-11057 MELITACALLLIAVHIAIVISCPEPLAAVAFVNVPLLYLCRYLLIIRRGVIIVSGGGGGGGRLRRFRLGAAMAVIYMAMSTLFFLRIAPLTPWWGALAAWVMILLIVEAIFAFFFPYRCCFNETEGDVQNNPHV
IRIS_313-11058 MELITACALLLIAVHIAIVISCPEPLAAVAFVNVPLLYLCRYLLIIRRGVIIVSGGGGGGGRLRRFRLGAAMAVIYMAMSTLFFLRIAPLTPWWGALAAWVMILLIVEAIFAFFFPYRCCFNETEGDVQNNPHV
IRIS_313-11059 MELITACALLLIAVHIAIVISCPEPLAAVALVDVPLLYLCRYLLIIRRGVIIVSGAGAGGGRLRRFRLGAAMAIIYMAMSTLFFLRIAPLTPWWGALAAWVMILLIVEAIFAFFFPYRCCFNETEGDVQNNPHV
IRIS_313-11060 MELITACALLLIAVHIAIVISCPEPLAAVAFVNVPLLYLCRYLLIIRRGVIIVSGGGGGGGRLRRFRLGAAMAVIYMAMSTLFFLRIAPLTPWWGALAAWVMILLIVEAIFAFFFPYRCCFNETEGDVQNNPHV
IRIS_313-11061 MELITACALLLIAVHIAIVISCPEPLAAVALVDVPLLYLCRYLLIIRRGVIIVSGAGGGGGGRLRRFRLGAAMAIIYMAMSTLFFLRIAPLTPWWGALAAWVMILLIVEAIFAFFFPYRCCFNETEGDVQNNPHV
IRIS_313-11062 MELITACALLLIAVHIAIVISCPEPLAAVAFVNVPLLYLCRYLLIIRRGVIIVSGGGGGGGRLRRFRLGAAMAVIYMAMSTLFFLRIAPLTPWWGALAAWVMILLIVEAIFAFFFPYRCCFNETEGDVQNNPHV
IRIS_313-11063 MELITACALLLIAVHIAIVISCPEPLAAVAFVNVPLLYLCRYLLIIRRGVIIVSGGGGGGGRLRRFRLGAAMAVIYMAMSTLFFLRIAPLTPWWGALAAWVMILLIVEAIFAFFFPYRCCFNETEGDVQNNPHV
IRIS_313-11064 MELITACALLLIAVHIAIVISCPEPLAAVAFVNVPLLYLCRYLLIIRRGVIIVSGGGGGGGRLRRFRLGAAMAVIYMAMSTLFFLRIAPLTPWWGALAAWVMILLIVEAIFAFFFPYRCCFNETEGDVQNNPHV
IRIS_313-11065 MELITACALLLIAVHIAIVISCPEPLAAVAFVNVPLLYLCRYLLIIRRGVIIVSGGGGGGGRLRRFRLGAAMAVIYMAMSTLFFLRIAPLTPWWGALAAWVMILLIVEAIFAFFFPYRCCFNETEGDVQNNPHV
IRIS_313-11066 MELITACALLLIAVHIAIVISCPEPLAAVAFVNVPLLYLCRYLLIIRRGVIIVSGGGGGGGRLRRFRLGAAMAVIYMAMSTLFFLRIAPLTPWWGALAAWVMILLIVEAIFAFFFPYRCCFNETEGDVQNNPHV
IRIS_313-11067 MELITACALLLIAVHIAIVISCPEPLAAVAFVNVPLLYLCRYLLIIRRGVIIVSGGGGGGGRLRRFRLGAAMAVIYMAMSTLFFLRIAPLTPWWGALAAWVMILLIVEAIFAFFFPYRCCFNETEGDVQNNPHV
IRIS_313-11068 MELITACALLLIAVHIAIVISCPEPLAAVAFVNVPLLYLCRYLLIIRRGVIIVSGGGGGGGRLRRFRLGAAMAVIYMAMSTLFFLRIAPLTPWWGALAAWVMILLIVEAIFAFFFPYRCCFNETEGDVQNNPHV
IRIS_313-11069 MELITACALLLIAVHIAIVISCPEPLAAVAFVNVPLLYLCRYLLIIRRGVIIVSGGGGGGGRLRRFRLGAAMAVIYMAMSTLFFLRIAPLTPWWGALAAWVMILLIVEAIFAFFFPYRCCFNETEGDVQNNPHV
IRIS_313-11070 MELITACALLLIAVHIAIVISCPEPLAAVAFVNVPLLYLCRYLLIIRRGVIIVSGGGGGGGRLRRFRLGAAMAVIYMAMSTLFFLRIAPLTPWWGALAAWVMILLIVEAIFAFFFPYRCCFNETEGDVQNNPHV
IRIS_313-11071 MELITACALLLIAVHIAIVISCPEPLAAVALVDVPLLYLCRYLLIIRRGVIIVSGGGGGGRLRRFRLGAAMAIIYMAMSTLFFLRIAPLTPWWGALAAWVMILLIVEAIFAFFFPYRCCFNETEGDVQNNPHV
IRIS_313-11072 MELITACALLLIAVHIAIVISCPEPLAAVALVDVPLLYLYRYLLIIRRGVIIVSGAGGGGGRLRRFRLGAAMAIIYMAMSTLFFLRIAPLTPWWGALAAWVMILLIVEAIFAFFFPYRCCFNETEGDVQNNPHV
IRIS_313-11073 MELITACALLLIAVHIAIVISCPEPLAAVALVDVPLLYLCRYLLIIRRGVIIVSGAGAGGGRLRRFRLGAAMAIIYMAMSTLFFLRIAPLTPWWGALAAWVMILLIVEAIFAFFFPYRCCFNETEGDVQNNPHV
IRIS_313-11074 MELITACALLLIAVHIAIVISCPEPLAAVALVDVPLLYLCRYLLIIRRGVIIVSGGGGGGRLRRFRLGAAMAIIYMAMSTLFFLRIAPLTPWWGALAAWVMILLIVEAIFAFFFPYRCCFNETEGDVQNNPHV
IRIS_313-11075 MELITACALLLIAVHIAIVISCPEPLAAVALVDVPLLYLCRYLLIIRRGVIIVSGAGAGGGRLRRFRLGAAMAIIYMAMSTLFFLRIAPLTPWWGALAAWVMILLIVEAIFAFFFPYRCCFNETEGDVQNNPHV
IRIS_313-11076 MELITACALLLIAVHIAIVISCPEPLAAVALVDVPLLYLCRYLLIIRRGVIIVSGGGGGGRLRRFRLGAAMAIIYMAMSTLFFLRIAPLTPWWGALAAWVMILLIVEAIFAFFFPYRCCFNETEGDVQNNPHV
IRIS_313-11077 MELITACALLLIAVHIAIVISCPEPLAAVALVDVPLLYLCRYLLIIRRGVIIVSGAGGGGGRLRRFRLGAAMAIIYMAMSTLFFLRIAPLTPWWGALAAWVMILLIVEAIFAFFFPYRCCFNETEGDVQNNPHV
IRIS_313-11078 MELITACALLLIAVHIAIVISCPEPLAAVALVDVPLLYLCRYLLIIRRGVIIVSGGGGGGRLRRFRLGAAMAIIYMAMSTLFFLRIAPLTPWWGALAAWVMILLIVEAIFAFFFPYRCCFNETEGDVQNNPHV
IRIS_313-11079 MELITACALLLIAVHIAIVISCPEPLAAVALVDVPLLYLYRYLLIIRRGVIIVSGGGGGGRLRRFRLGAAMAIIYMAMSTLFFLRIAPLTPWWGALAAWVMILLIVEAIFAFFFPYRCCFNETEGDVQNNPHV

IRIS_313-11080 MELITACALLLIAVHIAIVISCPEPLAAVALVDVPLLYLYRYLLIIRRGVIIVSGAGGGGGRLRRFRLGAAMAIIYMAMSTLFFLRIAPLTPWWGALAAWVMILLIVEAIFAFFFPYRCCFNETEGDVQNNPHV
IRIS_313-11081 MELITACALLLIAVHIAIVISCPEPLAAVALVDVPLLYLCRYLLIIRRGVIIVSGGGGGGRLRRFRLGAAMAIIYMAMSTLFFLRIAPLTPWWGALAAWVMILLIVEAIFAFFFPYRCCFNETEGDVQNNPHV
IRIS_313-11082 MELITACALLLIAVHIAIVISCPEPLAAVALVDVPLLYLCRYLLIIRRGVIIVSGGGGGGRLRRFRLGAAMAIIYMAMSTLFFLRIAPLTPWWGALAAWVMILLIVEAIFAFFFPYRCCFNETEGDVQNNPHV
IRIS_313-11083 MELITACALLLIAVHIAIVISCPEPLAAVALVDVPLLYLYRYLLIIRRGVIIVSGAGGGGGRLRRFRLGAAMAIIYMAMSTLFFLRIAPLTPWWGALAAWVMILLIVEAIFAFFFPYRCCFNETEGDVQNNPHV
IRIS_313-11084 MELITACALLLIAVHIAIVISCPEPLAAVALVDVPLLYLCRYLLIIRRGVIIVSGGGAGGGRLRRFRLGAAMAIIYMAMSTLFFLRIAPLTPWWGALAAWVMILLIVEAIFAFFFPYRCCFNETEGDVQNNPHV
IRIS_313-11085 MELITACALLLIAVHIAIVISCPEPLAAVALVDVPLLYLYRYLLIIRRGVIIVSGAGGGGGRLRRFRLGAAMAIIYMAMSTLFFLRIAPLTPWWGALAAWVMILLIVEAIFAFFFPYRCCFNETEGDVQNNPHV
IRIS_313-11086 MELITACALLLIAVHIAIVISCPEPLAAVALVDVPLLYLCRYLLIIRRGVIIVSGAGAGGGRLRRFRLGAAMAIIYMAMSTLFFLRIAPLTPWWGALAAWVMILLIVEAIFAFFFPYRCCFNETEGDVQNNPHV
IRIS_313-11087 MELITACALLLIAVHIAIVISCPEPLAAVALVDVPLLYLCRYLLIIRRGVIIVSGAGAGGGRLRRFRLGAAMAIIYMAMSTLFFLRIAPLTPWWGALAAWVMILLIVEAIFAFFFPYRCCFNETEGDVQNNPHV
IRIS_313-11088 MELITACALLLIAVHIAIVISCPEPLAAVALVDVPLLYLCRYLLIIRRGVIIVSGAGAGGGRLRRFRLGAAMAIIYMAMSTLFFLRIAPLTPWWGALAAWVMILLIVEAIFAFFFPYRCCFNETEGDVQNNPHV
IRIS_313-11089 MELITACALLLIAVHIAIVISCPEPLAAVALVDVPLLYLYRYLLIIRRGVIIVSGAGGGGGRLRRFRLGAAMAIIYMAMSTLFFLRIAPLTPWWGALAAWVMILLIVEAIFAFFFPYRCCFNETEGDVQNNPHV
IRIS_313-11090 MELITACALLLIAVHIAIVISCPEPLAAVALVDVPLLYLCRYLLIIRRGVIIVSGAGAGGGRLRRFRLGAAMAIIYMAMSTLFFLRIAPLTPWWGALAAWVMILLIVEAIFAFFFPYRCCFNETEGDVQNNPHV
IRIS_313-11091 MELITACALLLIAVHIAIVISCPEPLAAVALVDVPLLYLCRYLLIIRRGVIIVSGAGGGGGRLRRFRLGAAMAIIYMAMSTLFFLRIAPLTPWWGALAAWVMILLIVEAIFAFFFPYRCCFNETEGDVQNNPHV
IRIS_313-11092 MELITACALLLIAVHIAIVISCPEPLAAVALVDVPLLYLYRYLLIIRRGVIIVSGAGGGGGRLRRFRLGAAMAIIYMAMSTLFFLRIAPLTPWWGALAAWVMILLIVEAIFAFFFPYRCCFNETEGDVQNNPHV
IRIS_313-11093 MELITACALLLIAVHIAIVISCPEPLAAVALVDVPLLYLYRYLLIIRRGVIIVSGAGAGGGRLRRFRLGAAMAIIYMAMSTLFFLRIAPLTPWWGALAAWVMILLIVEAIFAFFFPYRCCFNETEGDVQNNPHV
IRIS_313-11094 MELITACALLLIAVHIAIVISCPEPLAAVALVDVPLLYLYRYLLIIRRGVIIVSGAGGGGGRLRRFRLGAAMAIIYMAMSTLFFLRIAPLTPWWGALAAWVMILLIVEAIFAFFFPYRCCFNETEGDVQNNPHV
IRIS_313-11095 MELITACALLLIAVHIAIVISCPEPLAAVALVDVPLLYLCRYLLIIRRGVIIVSGGGGGGRLRRFRLGAAMAIIYMAMSTLFFLRIAPLTPWWGALAAWVMILLIVEAIFAFFFPYRCCFNETEGDVQNNPHV
IRIS_313-11096 MELITACALLLIAVHIAIVISCPEPLAAVALVDVPLLYLCRYLLIIRRGVIIVSGAGAGGGRLRRFRLGAAMAIIYMAMSTLFFLRIAPLTPWWGALAAWVMILLIVEAIFAFFFPYRCCFNETEGDVQNNPHV
IRIS_313-11097 MELITACALLLIAVHIAIVISCPEPLAAVALVDVPLLYLYRYLLIIRRGVIIVSGAGGGGGRLRRFRLGAAMAIIYMAMSTLFFLRIAPLTPWWGALAAWVMILLIVEAIFAFFFPYRCCFNETEGDVQNNPHV
IRIS_313-11098 MELITACALLLIAVHIAIVISCPEPLAAVALVDVPLLYLCRYLLIIRRGVIIVSGGGGGGRLRRFRLGAAMAIIYMAMSTLFFLRIAPLTPWWGALAAWVMILLIVEAIFAFFFPYRCCFNETEGDVQNNPHV
IRIS_313-11099 MELITACALLLIAVHIAIVISCPEPLAAVALVDVPLLYLCRYLLIIRRGVIIVSGAGAGGGRLRRFRLGAAMAIIYMAMSTLFFLRIAPLTPWWGALAAWVMILLIVEAIFAFFFPYRCCFNETEGDVQNNPHV
IRIS_313-11100 MELITACALLLIAVHIAIVISCPEPLAAVALVDVPLLYLCRYLLIIRRGVIIVSGAGAGGGRLRRFRLGAAMAIIYMAMSTLFFLRIAPLTPWWGALAAWVMILLIVEAIFAFFFPYRCCFNETEGDVQNNPHV
IRIS_313-11101 MELITACALLLIAVHIAIVISCPEPLAAVALVDVPLLYLCRYLLIIRRGVIIVSGAGAGGGRLRRFRLGAAMAIIYMAMSTLFFLRIAPLTPWWGALAAWVMILLIVEAIFAFFFPYRCCFNETEGDVQNNPHV
IRIS_313-11102 MELITACALLLIAVHIAIVISCPEPLAAVALVDVPLLYLYRYLLIIRRGVIIVSGAGGGGGRLRRFRLGAAMAIIYMAMSTLFFLRIAPLTPWWGALAAWVMILLIVEAIFAFFFPYRCCFNETEGDVQNNPHV
IRIS_313-11103 MELITACALLLIAVHIAIVISCPEPLAAVALVDVPLLYLCRYLLIIRRGVIIVSGAGAGGGRLRRFRLGAAMAIIYMAMSTLFFLRIAPLTPWWGALAAWVMILLIVEAIFAFFFPYRCCFNETEGDVQNNPHV
IRIS_313-11104 MELITACALLLIAVHIAIVISCPEPLAAVALVDVPLLYLCRYLLIIRRGVIIVSGAGAGGGRLRRFRLGAAMAIIYMAMSTLFFLRIAPLTPWWGALAAWVMILLIVEAIFAFFFPYRCCFNETEGDVQNNPHV
IRIS_313-11105 MELITACALLLIAVHIAIVISCPEPLAAVALVDVPLLYLCRYLLIIRRGVIIVSGAGAGGGRLRRFRLGAAMAIIYMAMSTLFFLRIAPLTPWWGALAAWVMILLIVEAIFAFFFPYRCCFNETEGDVQNNPHV
IRIS_313-11106 MELITACALLLIAVHIAIVISCPEPLAAVALVDVPLLYLYRYLLIIRRGVIIVSGAGGGGGRLRRFRLGAAMAIIYMAMSTLFFLRIAPLTPWWGALAAWVMILLIVEAIFAFFFPYRCCFNETEGDVQNNPHV
IRIS_313-11107 MELITACALLLIAVHIAIVISCPEPLAAVALVDVPLLYLCRYLLIIRRGVIIVSGAGGGGGRLRRFRLGAAMAIIYMAMSTLFFLRIAPLTPWWGALAAWVMILLIVEAIFAFFFPYRCCFNETEGDVQNNPHV
IRIS_313-11108 MELITACALLLIAVHIAIVISCPEPLAAVALVDVPLLYLCRYLLIIRRGVIIVSGAGAGGGRLRRFRLGAAMAIIYMAMSTLFFLRIAPLTPWWGALAAWVMILLIVEAIFAFFFPYRCCFNETEGDVQNNPHV
IRIS_313-11109 MELITACALLLIAVHIAIVISCPEPLAAVALVDVPLLYLCRYLLIIRRGVIIVSGAGAGGGRLRRFRLGAAMAIIYMAMSTLFFLRIAPLTPWWGALAAWVMILLIVEAIFAFFFPYRCCFNETEGDVQNNPHV
IRIS_313-11110 MELITACALLLIAVHIAIVISCPEPLAAVALVDVPLLYLCRYLLIIRRGVIIVSGAGAGGGRLRRFRLGAAMAIIYMAMSTLFFLRIAPLTPWWGALAAWVMILLIVEAIFAFFFPYRCCFNETEGDVQNNPHV
IRIS_313-11111 MELITACALLLIAVHIAIVISCPEPLAAVALVDVPLLYLCRYLLIIRRGVIIVSGAGAGGGRLRRFRLGAAMAIIYMAMSTLFFLRIAPLTPWWGALAAWVMILLIVEAIFAFFFPYRCCFNETEGDVQNNPHV
IRIS_313-11112 MELITACALLLIAVHIAIVISCPEPLAAVALVDVPLLYLCRYLLIIRRGVIIVSGAGAGGGRLRRFRLGAAMAIIYMAMSTLFFLRIAPLTPWWGALAAWVMILLIVEAIFAFFFPYRCCFNETEGDVQNNPHV
IRIS_313-11113 MELITACALLLIAVHIAIVISCPEPLAAVALVDVPLLYLCRYLLIIRRGVIIVSGAGGGGGGRLRRFRLGAAMAIIYMAMSTLFFLRIAPLTPWWGALAAWVMILLIVEAIFAFFFPYRCCFNETEGDVQNNPHV
IRIS_313-11114 MELITACALLLIAVHIAIVISCPEPLAAVALVDVPLLYLCRYLLIIRRGVIIVSGGGAGGGRLRRFRLGAAMAIIYMAMSTLFFLRIAPLTPWWGALAAWVMILLIVEAIFAFFFPYRCCFNETEGDVQNNPHV
IRIS_313-11115 MELITACALLLIAVHIAIVISCPEPLAAVALVDVPLLYLCRYLLIIRRGVIIVSGAGAGGGRLRRFRLGAAMAIIYMAMSTLFFLRIAPLTPWWGALAAWVMILLIVEAIFAFFFPYRCCFNETEGDVQNNPHV
IRIS_313-11116 MELITACALLLIAVHIAIVISCPEPLAAVALVDVPLLYLCRYLLIIRRGVIIVSGAGAGGGRLRRFRLGAAMAIIYMAMSTLFFLRIAPLTPWWGALAAWVMILLIVEAIFAFFFPYRCCFNETEGDVQNNPHV
IRIS_313-11117 MELITACALLLIAVHIAIVISCPEPLAAVALVDVPLLYLCRYLLIIRRGVIIVSGAGAGGGRLRRFRLGAAMAIIYMAMSTLFFLRIAPLTPWWGALAAWVMILLIVEAIFAFFFPYRCCFNETEGDVQNNPHV
IRIS_313-11118 MELITACALLLIAVHIAIVISCPEPLAAVALVDVPLLYLYRYLLIIRRGVIIVSGAGGGGGRLRRFRLGAAMAIIYMAMSTLFFLRIAPLTPWWGALAAWVMILLIVEAIFAFFFPYRCCFNETEGDVQNNPHV
IRIS_313-11119 MELITACALLLIAVHIAIVISCPEPLAAVALVDVPLLYLCRYLLIIRRGVIIVSGAGAGGGRLRRFRLGAAMAIIYMAMSTLFFLRIAPLTPWWGALAAWVMILLIVEAIFAFFFPYRCCFNETEGDVQNNPHV
IRIS_313-11120 MELITACALLLIAVHIAIVISCPEPLAAVALVDVPLLYLCRYLLIIRRGVIIVSGAGAGGGRLRRFRLGAAMAIIYMAMSTLFFLRIAPLTPWWGALAAWVMILLIVEAIFAFFFPYRCCFNETEGDVQNNPHV
IRIS_313-11121 MELITACALLLIAVHIAIVISCPEPLAAVALVDVPLLYLCRYLLIIRRGVIIVSGAGAGGGRLRRFRLGAAMAIIYMAMSTLFFLRIAPLTPWWGALAAWVMILLIVEAIFAFFFPYRCCFNETEGDVQNNPHV
IRIS_313-11122 MELITACALLLIAVHIAIVISCPEPLAAVALVDVPLLYLCRYLLIIRRGVIIVSGGGGGGRLRRFRLGAAMAIIYMAMSTLFFLRIAPLTPWWGALAAWVMILLIVEAIFAFFFPYRCCFNETEGDVQNNPHV
IRIS_313-11123 MELITACALLLIAVHIAIVISCPEPLAAVALVDVPLLYLCRYLLIIRRGVIIVSGAGAGGGRLRRFRLGAAMAIIYMAMSTLFFLRIAPLTPWWGALAAWVMILLIVEAIFAFFFPYRCCFNETEGDVQNNPHV
IRIS_313-11124 MELITACALLLIAVHIAIVISCPEPLAAVALVDVPLLYLCRYLLIIRRGVIIVSGAGAGGGRLRRFRLGAAMAIIYMAMSTLFFLRIAPLTPWWGALAAWVMILLIVEAIFAFFFPYRCCFNETEGDVQNNPHV
IRIS_313-11125 MELITACALLLIAVHIAIVISCPEPLAAVALVDVPLLYLCRYLLIIRRGVIIVSGAGAGGGRLRRFRLGAAMAIIYMAMSTLFFLRIAPLTPWWGALAAWVMILLIVEAIFAFFFPYRCCFNETEGDVQNNPHV
IRIS_313-11126 MELITACALLLIAVHIAIVISCPEPLAAVALVDVPLLYLCRYLLIIRRGVIIVSGGGGGGRLRRFRLGAAMAIIYMAMSTLFFLRIAPLTPWWGALAAWVMILLIVEAIFAFFFPYRCCFNETEGDVQNNPHV
IRIS_313-11127 MELITACALLLIAVHIAIVISCPEPLAAVALVDVPLLYLYRYLLIIRRGVIIVSGAGGGGGRLRRFRLGAAMAIIYMAMSTLFFLRIAPLTPWWGALAAWVMILLIVEAIFAFFFPYRCCFNETEGDVQNNPHV
IRIS_313-11128 MELITACALLLIAVHIAIVISCPEPLAAVALVDVPLLYLCRYLLIIRRGVIIVSGGGAGGGRLRRFRLGAAMAIIYMAMSTLFFLRIAPLTPWWGALAAWVMILLIVEAIFAFFFPYRCCFNETEGDVQNNPHV
IRIS_313-11129 MELITACALLLIAVHIAIVISCPEPLAAVALVDVPLLYLCRYLLIIRRGVIIVSGGGGGGRLRRFRLGAAMAIIYMAMSTLFFLRIAPLTPWWGALAAWVMILLIVEAIFAFFFPYRCCFNETEGDVQNNPHV
IRIS_313-11130 MELITACALLLIAVHIAIVISCPEPLAAVALVDVPLLYLCRYLLIIRRGVIIVSGGGGGGRLRRFRLGAAMAIIYMAMSTLFFLRIAPLTPWWGALAAWVMILLIVEAIFAFFFPYRCCFNETEGDVQNNPHV
IRIS_313-11131 MELITACALLLIAVHIAIVISCPEPLAAVALVDVPLLYLCRYLLIIRRGVIIVSGAGAGGGRLRRFRLGAAMAIIYMAMSTLFFLRIAPLTPWWGALAAWVMILLIVEAIFAFFFPYRCCFNETEGDVQNNPHV
IRIS_313-11132 MELITACALLLIAVHIAIVISCPEPLAAVALVDVPLLYLYRYLLIIRRGVIIVSGAGGGGGRLRRFRLGAAMAIIYMAMSTLFFLRIAPLTPWWGALAAWVMILLIVEAIFAFFFPYRCCFNETEGDVQNNPHV
IRIS_313-11133 MELITACALLLIAVHIAIVISCPEPLAAVALVDVPLLYLYRYLLIIRRGVIIVSGAGAGGGRLRRFRLGAAMAIIYMAMSTLFFLRIAPLTPWWGALAAWVMILLIVEAIFAFFFPYRCCFNETEGDVQNNPHV
IRIS_313-11134 MELITACALLLIAVHIAIVISCPEPLAAVALVDVPLLYLCRYLLIIRRGVIIVSGGGGGGRLRRFRLGAAMAIIYMAMSTLFFLRIAPLTPWWGALAAWVMILLIVEAIFAFFFPYRCCFNETEGDVQNNPHV
IRIS_313-11135 MELITACALLLIAVHIAIVISCPEPLAAVALVDVPLLYLCRYLLIIRRGVIIVSGAGAGGGRLRRFRLGAAMAIIYMAMSTLFFLRIAPLTPWWGALAAWVMILLIVEAIFAFFFPYRCCFNETEGDVQNNPHV
IRIS_313-11136 MELITACALLLIAVHIAIVISCPEPLAAVALVDVPLLYLCRYLLIIRRGVIIVSGAGAGGGRLRRFRLGAAMAIIYMAMSTLFFLRIAPLTPWWGALAAWVMILLIVEAIFAFFFPYRCCFNETEGDVQNNPHV
IRIS_313-11137 MELITACALLLIAVHIAIVISCPEPLAAVALVDVPLLYLCRYLLIIRRGVIIVSGAGAGGGRLRRFRLGAAMAIIYMAMSTLFFLRIAPLTPWWGALAAWVMILLIVEAIFAFFFPYRCCFNETEGDVQNNPHV
IRIS_313-11138 MELITACALLLIAVHIAIVISCPEPLAAVALVDVPLLYLCRYLLIIRRGVIIVSGAGGGGGRLRRFRLGAAMAIIYMAMSTLFFLRIAPLTPWWGALAAWVMILLIVEAIFAFFFPYRCCFNETEGDVQNNPHV
IRIS_313-11139 MELITACALLLIAVHIAIVISCPEPLAAVALVDVPLLYLCRYLLIIRRGVIIVSGAGAGGGRLRRFRLGAAMAIIYMAMSTLFFLRIAPLTPWWGALAAWVMILLIVEAIFAFFFPYRCCFNETEGDVQNNPHV
IRIS_313-11140 MELITACALLLIAVHIAIVISCPEPLAAVALVDVPLLYLCRYLLIIRRGVIIVSGGGGGGRLRRFRLGAAMAIIYMAMSTLFFLRIAPLTPWWGALAAWVMILLIVEAIFAFFFPYRCCFNETEGDVQNNPHV
IRIS_313-11141 MELITACALLLIAVHIAIVISCPEPLAAVALVDVPLLYLCRYLLIIRRGVIIVSGAGAGGGRLRRFRLGAAMAIIYMAMSTLFFLRIAPLTPWWGALAAWVMILLIVEAIFAFFFPYRCCFNETEGDVQNNPHV
IRIS_313-11142 MELITACALLLIAVHIAIVISCPEPLAAVALVDVPLLYLCRYLLIIRRGVIIVSGGGAGGGRLRRFRLGAAMAIIYMAMSTLFFLRIAPLTPWWGALAAWVMILLIVEAIFAFFFPYRCCFNETEGDVQNNPHV
IRIS_313-11143 MELITACALLLIAVHIAIVISCPEPLAAVALVDVPLLYLYRYLLIIRRGVIIVSGAGGGGGRLRRFRLGAAMAIIYMAMSTLFFLRIAPLTPWWGALAAWVMILLIVEAIFAFFFPYRCCFNETEGDVQNNPHV
IRIS_313-11144 MELITACALLLIAVHIAIVISCPEPLAAVALVDVPLLYLCRYLLIIRRGVIIVSGAGGGGGRLRRFRLGAAMAIIYMAMSTLFFLRIAPLTPWWGALAAWVMILLIVEAIFAFFFPYRCCFNETEGDVQNNPHV
IRIS_313-11145 MELITACALLLIAVHIAIVISCPEPLAAVALVDVPLLYLCRYLLIIRRGVIIVSGGGGGGRLRRFRLGAAMAIIYMAMSTLFFLRIAPLTPWWGALAAWVMILLIVEAIFAFFFPYRCCFNETEGDVQNNPHV
IRIS_313-11146 MELITACALLLIAVHIAIVISCPEPLAAVALVDVPLLYLCRYLLIIRRGVIIVSGAGAGGGRLRRFRLGAAMAIIYMAMSTLFFLRIAPLTPWWGALAAWVMILLIVEAIFAFFFPYRCCFNETEGDVQNNPHV
IRIS_313-11147 MELITACALLLIAVHIAIVISCPEPLAAVALVDVPLLYLCRYLLIIRRGVIIVSGAGAGGGRLRRFRLGAAMAIIYMAMSTLFFLRIAPLTPWWGALAAWVMILLIVEAIFAFFFPYRCCFNETEGDVQNNPHV
IRIS_313-11148 MELITACALLLIAVHIAIVISCPEPLAAVALVDVPLLYLCRYLLIIRRGVIIVSGAGAGGGRLRRFRLGAAMAIIYMAMSTLFFLRIAPLTPWWGALAAWVMILLIVEAIFAFFFPYRCCFNETEGDVQNNPHV
IRIS_313-11149 MELITACALLLIAVHIAIVISCPEPLAAVALVDVPLLYLYRYLLIIRRGVIIVSGAGGGGGRLRRFRLGAAMAIIYMAMSTLFFLRIAPLTPWWGALAAWVMILLIVEAIFAFFFPYRCCFNETEGDVQNNPHV
IRIS_313-11150 MELITACALLLIAVHIAIVISCPEPLAAVALVDVPLLYLCRYLLIIRRGVIIVSGGGAGGGRLRRFRLGAAMAIIYMAMSTLFFLRIAPLTPWWGALAAWVMILLIVEAIFAFFFPYRCCFNETEGDVQNNPHV
IRIS_313-11151 MELITACALLLIAVHIAIVISCPEPLAAVALVDVPLLYLCRYLLIIRRGVIIVSGGGGGGRLRRFRLGAAMAIIYMAMSTLFFLRIAPLTPWWGALAAWVMILLIVEAIFAFFFPYRCCFNETEGDVQNNPHV
IRIS_313-11152 MELITACALLLIAVHIAIVISCPEPLAAVALVDVPLLYLCRYLLIIRRGVIIVSGAGAGGGRLRRFRLGAAMAIIYMAMSTLFFLRIAPLTPWWGALAAWVMILLIVEAIFAFFFPYRCCFNETEGDVQNNPHV
IRIS_313-11153 MELITACALLLIAVHIAIVISCPEPLAAVALVDVPLLYLCRYLLIIRRGVIIVSGAGAGGGRLRRFRLGAAMAIIYMAMSTLFFLRIAPLTPWWGALAAWVMILLIVEAIFAFFFPYRCCFNETEGDVQNNPHV
IRIS_313-11154 MELITACALLLIAVHIAIVISCPEPLAAVALVDVPLLYLCRYLLIIRRGVIIVSGAGGGGGRLRRFRLGAAMAIIYMAMSTLFFLRIAPLTPWWGALAAWVMILLIVEAIFAFFFPYRCCFNETEGDVQNNPHV
IRIS_313-11155 MELITACALLLIAVHIAIVISCPEPLAAVALVDVPLLYLYRYLLIIRRGVIIVSGAGGGGGRLRRFRLGAAMAIIYMAMSTLFFLRIAPLTPWWGALAAWVMILLIVEAIFAFFFPYRCCFNETEGDVQNNPHV
IRIS_313-11156 MELITACALLLIAVHIAIVISCPEPLAAVALVDVPLLYLYRYLLIIRRGVIIVSGAGGGGGRLRRFRLGAAMAIIYMAMSTLFFLRIAPLTPWWGALAAWVMILLIVEAIFAFFFPYRCCFNETEGDVQNNPHV
IRIS_313-11157 MELITACALLLIAVHIAIVISCPEPLAAVALVDVPLLYLCRYLLIIRRGVIIVSGGGGGGRLRRFRLGAAMAIIYMAMSTLFFLRIAPLTPWWGALAAWVMILLIVEAIFAFFFPYRCCFNETEGDVQNNPHV
IRIS_313-11158 MELITACALLLIAVHIAIVISCPEPLAAVALVDVPLLYLCRYLLIIRRGVIIVSGAGAGGGRLRRFRLGAAMAIIYMAMSTLFFLRIAPLTPWWGALAAWVMILLIVEAIFAFFFPYRCCFNETEGDVQNNPHV
IRIS_313-11159 MELITACALLLIAVHIAIVISCPEPLAAVALVDVPLLYLCRYLLIIRRGVIIVSGAGAGGGRLRRFRLGAAMAIIYMAMSTLFFLRIAPLTPWWGALAAWVMILLIVEAIFAFFFPYRCCFNETEGDVQNNPHV
IRIS_313-11160 MELITACALLLIAVHIAIVISCPEPLAAVALVDVPLLYLCRYLLIIRRGVIIVSGAGAGGGRLRRFRLGAAMAIIYMAMSTLFFLRIAPLTPWWGALAAWVMILLIVEAIFAFFFPYRCCFNETEGDVQNNPHV
IRIS_313-11161 MELITACALLLIAVHIAIVISCPEPLAAVALVDVPLLYLCRYLLIIRRGVIIVSGAGAGGGRLRRFRLGAAMAIIYMAMSTLFFLRIAPLTPWWGALAAWVMILLIVEAIFAFFFPYRCCFNETEGDVQNNPHV
IRIS_313-11162 MELITACALLLIAVHIAIVISCPEPLAAVALVDVPLLYLCRYLLIIRRGVIIVSGGGAGGGRLRRFRLGAAMAIIYMAMSTLFFLRIAPLTPWWGALAAWVMILLIVEAIFAFFFPYRCCFNETEGDVQNNPHV
IRIS_313-11163 MELITACALLLIAVHIAIVISCPEPLAAVAFVNVPLLYLCRYLLIIRRGVIIVSGGGGGGGRLRRFRLGAAMAVIYMAMSTLFFLRIAPLTPWWGALAAWVMILLIVEAIFAFFFPYRCCFNETEGDVQNNPHV
IRIS_313-11164 MELITACALLLIAVHIAIVISCPEPLAAVALVDVPLLYLCRYLLIIRRGVIIVSGAGAGGGRLRRFRLGAAMAIIYMAMSTLFFLRIAPLTPWWGALAAWVMILLIVEAIFAFFFPYRCCFNETEGDVQNNPHV
IRIS_313-11165 MELITACALLLIAVHIAIVISCPEPLAAVALVDVPLLYLCRYLLIIRRGVIIVSGGGAGGGRLRRFRLGAAMAIIYMAMSTLFFLRIAPLTPWWGALAAWVMILLIVEAIFAFFFPYRCCFNETEGDVQNNPHV
IRIS_313-11166 MELITACALLLIAVHIAIVISCPEPLAAVALVDVPLLYLCRYLLIIRRGVIIVSGAGAGGGRLRRFRLGAAMAIIYMAMSTLFFLRIAPLTPWWGALAAWVMILLIVEAIFAFFFPYRCCFNETEGDVQNNPHV
IRIS_313-11167 MELITACALLLIAVHIAIVISCPEPLAAVALVDVPLLYLCRYLLIIRRGVIIVSGGGGGGRLRRFRLGAAMAIIYMAMSTLFFLRIAPLTPWWGALAAWVMILLIVEAIFAFFFPYRCCFNETEGDVQNNPHV
IRIS_313-11168 MELITACALLLIAVHIAIVISCPEPLAAVALVDVPLLYLCRYLLIIRRGVIIVSGAGAGGGRLRRFRLGAAMAIIYMAMSTLFFLRIAPLTPWWGALAAWVMILLIVEAIFAFFFPYRCCFNETEGDVQNNPHV
IRIS_313-11169 MELITACALLLIAVHIAIVISCPEPLAAVALVDVPLLYLCRYLLIIRRGVIIVSGAGAGGGRLRRFRLGAAMAIIYMAMSTLFFLRIAPLTPWWGALAAWVMILLIVEAIFAFFFPYRCCFNETEGDVQNNPHV
IRIS_313-11170 MELITACALLLIAVHIAIVISCPEPLAAVALVDVPLLYLCRYLLIIRRGVIIVSGAGAGGGRLRRFRLGAAMAIIYMAMSTLFFLRIAPLTPWWGALAAWVMILLIVEAIFAFFFPYRCCFNETEGDVQNNPHV
IRIS_313-11171 MELITACALLLIAVHIAIVISCPEPLAAVALVDVPLLYLCRYLLIIRRGVIIVSGAGAGGGRLRRFRLGAAMAIIYMAMSTLFFLRIAPLTPWWGALAAWVMILLIVEAIFAFFFPYRCCFNETEGDVQNNPHV
IRIS_313-11172 MELITACALLLIAVHIAIVISCPEPLAAVALVDVPLLYLCRYLLIIRRGVIIVSGAGAGGGRLRRFRLGAAMAIIYMAMSTLFFLRIAPLTPWWGALAAWVMILLIVEAIFAFFFPYRCCFNETEGDVQNNPHV
IRIS_313-11173 MELITACALLLIAVHIAIVISCPEPLAAVAFVNVPLLYLCRYLLIIRRGVIIVSGGGAGGGRLRRFRLGAAMAVIYMAMSTLFFLRIAPLTPWWGALAAWVMILLIVEAIFAFFFPYRCCFNETEGDVQNNPHV
IRIS_313-11174 MELITACALLLIAVHIAIVISCPEPLAAVALVDVPLLYLCRYLLIIRRGVIIVSGGGGGGGRLRRFRLGAAMAVIYMAMSTLFFLRIAPLTPWWGALAAWVMILLIVEAIFAFFFPYRCCFNETEGDVQNNPHV
IRIS_313-11175 MELITACALLLIAVHIAIVISCPEPLAAVAFVNVPLLYLCRYLLIIRRGVIIVSGGGGGGGRLRRFRLGAAMAVIYMAMSTLFFLRIAPLTPWWGALAAWVMILLIVEAIFAFFFPYRCCFNETEGDVQNNPHV
IRIS_313-11176 MELITACALLLIAVHIAIVISCPEPLAAVALVDVPLLYLCRYLLIIRRGVIIVSGGGAGGGRLRRFRLGAAMAIIYMAMSTLFFLRIAPLTPWWGALAAWVMILLIVEAIFAFFFPYRCCFNETEGDVQNNPHV
IRIS_313-11177 MELITACALLLIAVHIAIVISCPEPLAAVALVDVPLLYLCRYLLIIRRGVIIVSGGGAGGGRLRRFRLGAAMAIIYMAMSTLFFLRIAPLTPWWGALAAWVMILLIVEAIFAFFFPYRCCFNETEGDVQNNPHV
IRIS_313-11178 MELITACALLLIAVHIAIVISCPEPLAAVALVDVPLLYLCRYLLIIRRGVIIVSGGGAGGGRLRRFRLGAAMAIIYMAMSTLFFLRIAPLTPWWGALAAWVMILLIVEAIFAFFFPYRCCFNETEGDVQNNPHV
IRIS_313-11179 MELITACALLLIAVHIAIVISCPEPLAAVALVDVPLLYLCRYLLIIRRGVIIVSGGGAGGGRLRRFRLGAAMAIIYMAMSTLFFLRIAPLTPWWGALAAWVMILLIVEAIFAFFFPYRCCFNETEGDVQNNPHV
IRIS_313-11180 MELITACALLLIAVHIAIVISCPEPLAAVALVDVPLLYLCRYLLIIRRGVIIVSGGGAGGGRLRRFRLGAAMAIIYMAMSTLFFLRIAPLTPWWGALAAWVMILLIVEAIFAFFFPYRCCFNETEGDVQNNPHV
IRIS_313-11181 MELITACALLLIAVHIAIVISCPEPLAAVALVDVPLLYLCRYLLIIRRGVIIVSGAGAGGGRLRRFRLGAAMAIIYMAMSTLFFLRIAPLTPWWGALAAWVMILLIVEAIFAFFFPYRCCFNETEGDVQNNPHV
IRIS_313-11182 MELITACALLLIAVHIAIVISCPEPLAAVALVDVPLLYLCRYLLIIRRGVIIVSGGGGGGRLRRFRLGAAMAIIYMAMSTLFFLRIAPLTPWWGALAAWVMILLIVEAIFAFFFPYRCCFNETEGDVQNNPHV
IRIS_313-11183 MELITACALLLIAVHIAIVISCPEPLAAVALVDVPLLYLCRYLLIIRRGVIIVSGAGGGGGRLRRFRLGAAMAIIYMAMSTLFFLRIAPLTPWWGALAAWVMILLIVEAIFAFFFPYRCCFNETEGDVQNNPHV
IRIS_313-11189 MELITACALLLIAVHIAIVISCPEPLAAVALVDVPLLYLCRYLLIIRRGVIIVSGAGAGGGRLRRFRLGAAMAIIYMAMSTLFFLRIAPLTPWWGALAAWVMILLIVEAIFAFFFPYRCCFNETEGDVQNNPHV
IRIS_313-11191 MELITACALLLIAVHIAIVISCPEPLAAVALVDVPLLYLCRYLLIIRRGVIIVSGAGAGGGRLRRFRLGAAMAIIYMAMSTLFFLRIAPLTPWWGALAAWVMILLIVEAIFAFFFPYRCCFNETEGDVQNNPHV
IRIS_313-11192 MELITACALLLIAVHIAIVISCPEPLAAVALVDVPLLYLCRYLLIIRRGVIIVSGGGGGGRLRRFRLGAAMAIIYMAMSTLFFLRIAPLTPWWGALAAWVMILLIVEAIFAFFFPYRCCFNETEGDVQNNPHV
IRIS_313-11193 MELITACALLLIAVHIAIVISCPEPLAAVALVDVPLLYLCRYLLIIRRGVIIVSGGGGGGRLRRFRLGAAMAIIYMAMSTLFFLRIAPLTPWWGALAAWVMILLIVEAIFAFFFPYRCCFNETEGDVQNNPHV
IRIS_313-11194 MELITACALLLIAVHIAIVISCPEPLAAVALVDVPLLYLCRYLLIIRRGVIIVSGGGGGGRLRRFRLGAAMAIIYMAMSTLFFLRIAPLTPWWGALAAWVMILLIVEAIFAFFFPYRCCFNETEGDVQNNPHV
IRIS_313-11195 MELITACALLLIAVHIAIVISCPEPLAAVALVDVPLLYLCRYLLIIRRGVIIVSGAGGGGGRLRRFRLGAAMAIIYMAMSTLFFLRIAPLTPWWGALAAWVMILLIVEAIFAFFFPYRCCFNETEGDVQNNPHV
IRIS_313-11196 MELITACALLLIAVHIAIVISCPEPLAAVALVDVPLLYLCRYLLIIRRGVIIVSGAGAGGGRLRRFRLGAAMAIIYMAMSTLFFLRIAPLTPWWGALAAWVMILLIVEAIFAFFFPYRCCFNETEGDVQNNPHV
IRIS_313-11197 MELITACALLLIAVHIAIVISCPEPLAAVALVDVPLLYLCRYLLIIRRGVIIVSGGGGGGRLRRFRLGAAMAIIYMAMSTLFFLRIAPLTPWWGALAAWVMILLIVEAIFAFFFPYRCCFNETEGDVQNNPHV
IRIS_313-11198 MELITACALLLIAVHIAIVISCPEPLAAVALVDVPLLYLCRYLLIIRRGVIIVSGAGAGGGRLRRFRLGAAMAIIYMAMSTLFFLRIAPLTPWWGALAAWVMILLIVEAIFAFFFPYRCCFNETEGDVQNNPHV
IRIS_313-11199 MELITACALLLIAVHIAIVISCPEPLAAVALVDVPLLYLCRYLLIIRRGVIIVSGGGAGGGRLRRFRLGAAMAIIYMAMSTLFFLRIAPLTPWWGALAAWVMILLIVEAIFAFFFPYRCCFNETEGDVQNNPHV
IRIS_313-11200 MELITACALLLIAVHIAIVISCPEPLAAVALVDVPLLYLCRYLLIIRRGVIIVSGGGAGGGRLRRFRLGAAMAIIYMAMSTLFFLRIAPLTPWWGALAAWVMILLIVEAIFAFFFPYRCCFNETEGDVQNNPHV
IRIS_313-11201 MELITACALLLIAVHIAIVISCPEPLAAVALVDVPLLYLCRYLLIIRRGVIIVSGAGAGGGRLRRFRLGAAMAIIYMAMSTLFFLRIAPLTPWWGALAAWVMILLIVEAIFAFFFPYRCCFNETEGDVQNNPHV
IRIS_313-11202 MELITACALLLIAVHIAIVISCPEPLAAVALVDVPLLYLCRYLLIIRRGVIIVSGAGAGGGRLRRFRLGAAMAIIYMAMSTLFFLRIAPLTPWWGALAAWVMILLIVEAIFAFFFPYRCCFNETEGDVQNNPHV
IRIS_313-11203 MELITACALLLIAVHIAIVISCPEPLAAVALVDVPLLYLCRYLLIIRRGVIIVSGAGAGGGRLRRFRLGAAMAIIYMAMSTLFFLRIAPLTPWWGALAAWVMILLIVEAIFAFFFPYRCCFNETEGDVQNNPHV
IRIS_313-11204 MELITACALLLIAVHIAIVISCPEPLAAVALVDVPLLYLCRYLLIIRRGVIIVSGAGAGGGRLRRFRLGAAMAIIYMAMSTLFFLRIAPLTPWWGALAAWVMILLIVEAIFAFFFPYRCCFNETEGDVQNNPHV
IRIS_313-11205 MELITACALLLIAVHIAIVISCPEPLAAVALVDVPLLYLCRYLLIIRRGVIIVSGGGGGGRLRRFRLGAAMAIIYMAMSTLFFLRIAPLTPWWGALAAWVMILLIVEAIFAFFFPYRCCFNETEGDVQNNPHV
IRIS_313-11206 MELITACALLLIAVHIAIVISCPEPLAAVALVDVPLLYLCRYLLIIRRGVIIVSGAGAGGGRLRRFRLGAAMAIIYMAMSTLFFLRIAPLTPWWGALAAWVMILLIVEAIFAFFFPYRCCFNETEGDVQNNPHV
IRIS_313-11207 MELITACALLLIAVHIAIVISCPEPLAAVALVDVPLLYLCRYLLIIRRGVIIVSGAGAGGGRLRRFRLGAAMAIIYMAMSTLFFLRIAPLTPWWGALAAWVMILLIVEAIFAFFFPYRCCFNETEGDVQNNPHV
IRIS_313-11208 MELITACALLLIAVHIAIVISCPEPLAAVALVDVPLLYLCRYLLIIRRGVIIVSGGGAGGGRLRRFRLGAAMAIIYMAMSTLFFLRIAPLTPWWGALAAWVMILLIVEAIFAFFFPYRCCFNETEGDVQNNPHV

IRIS_313-11209 MELITACALLLIAVHIAIVISCPEPLAAVALVDVPLLYLCRYLLIIRRGVIIVSGGGGGGRLRRFRLGAAMAIIYMAMSTLFFLRIAPLTPWWGALAAWVMILLIVEAIFAFFFPYRCCFNETEGDVQNNPHV
IRIS_313-11210 MELITACALLLIAVHIAIVISCPEPLAAVALVDVPLLYLCRYLLIIRRGVIIVSGAGAGGGRLRRFRLGAAMAIIYMAMSTLFFLRIAPLTPWWGALAAWVMILLIVEAIFAFFFPYRCCFNETEGDVQNNPHV
IRIS_313-11211 MELITACALLLIAVHIAIVISCPEPLAAVALVDVPLLYLCRYLLIIRRGVIIVSGAGAGGGRLRRFRLGAAMAIIYMAMSTLFFLRIAPLTPWWGALAAWVMILLIVEAIFAFFFPYRCCFNETEGDVQNNPHV
IRIS_313-11212 MELITACALLLIAVHIAIVISCPEPLAAVALVDVPLLYLCRYLLIIRRGVIIVSGAGAGGGRLRRFRLGAAMAIIYMAMSTLFFLRIAPLTPWWGALAAWVMILLIVEAIFAFFFPYRCCFNETEGDVQNNPHV
IRIS_313-11213 MELITACALLLIAVHIAIVISCPEPLAAVALVDVPLLYLCRYLLIIRRGVIIVSGGGGGGGRLRRFRLGAAMAVIYMAMSTLFFLRIAPLTPWWGALAAWVMILLIVEAIFAFFFPYRCCFNETEGDVQNNPHV
IRIS_313-11214 MELITACALLLIAVHIAIVISCPEPLAAVALVDVPLLYLCRYLLIIRRGVIIVSGAGAGGGRLRRFRLGAAMAIIYMAMSTLFFLRIAPLTPWWGALAAWVMILLIVEAIFAFFFPYRCCFNETEGDVQNNPHV
IRIS_313-11216 MELITACALLLIAVHIAIVISCPEPLAAVALVDVPLLYLCRYLLIIRRGVIIVSGAGGGGGRLRRFRLGAAMAIIYMAMSTLFFLRIAPLTPWWGALAAWVMILLIVEAIFAFFFPYRCCFNETEGDVQNNPHV
IRIS_313-11217 MELITACALLLIAVHIAIVISCPEPLAAVALVDVPLLYLCRYLLIIRRGVIIVSGAGAGGGRLRRFRLGAAMAIIYMAMSTLFFLRIAPLTPWWGALAAWVMILLIVEAIFAFFFPYRCCFNETEGDVQNNPHV
IRIS_313-11218 MELITACALLLIAVHIAIVISCPEPLAAVALVDVPLLYLCRYLLIIRRGVIIVSGAGAGGGRLRRFRLGAAMAIIYMAMSTLFFLRIAPLTPWWGALAAWVMILLIVEAIFAFFFPYRCCFNETEGDVQNNPHV
IRIS_313-11219 MELITACALLLIAVHIAIVISCPEPLAAVALVDVPLLYLCRYLLIIRRGVIIVSGGGGGGRLRRFRLGAAMAIIYMAMSTLFFLRIAPLTPWWGALAAWVMILLIVEAIFAFFFPYRCCFNETEGDVQNNPHV
IRIS_313-11220 MELITACALLLIAVHIAIVISCPEPLAAVALVDVPLLYLCRYLLIIRRGVIIVSGAGGGGRLRRFRLGAAMAIIYMAMSTLFFLRIAPLTPWWGALAAWVMILLIVEAIFAFFFPYRCCFNETEGDVQNNPHV
IRIS_313-11221 MELITACALLLIAVHIAIVISCPEPLAAVALVDVPLLYLCRYLLIIRRGVIIVSGAGAGGGRLRRFRLGAAMAIIYMAMSTLFFLRIAPLTPWWGALAAWVMILLIVEAIFAFFFPYRCCFNETEGDVQNNPHV
IRIS_313-11222 MELITACALLLIAVHIAIVISCPEPLAAVALVDVPLLYLCRYLLIIRRGVIIVSGGGGGGRLRRFRLGAAMAIIYMAMSTLFFLRIAPLTPWWGALAAWVMILLIVEAIFAFFFPYRCCFNETEGDVQNNPHV
IRIS_313-11223 MELITACALLLIAVHIAIVISCPEPLAAVALVDVPLLYLCRYLLIIRRGVIIVSGAGAGGGRLRRFRLGAAMAIIYMAMSTLFFLRIAPLTPWWGALAAWVMILLIVEAIFAFFFPYRCCFNETEGDVQNNPHV
IRIS_313-11224 MELITACALLLIAVHIAIVISCPEPLAAVALVDVPLLYLCRYLLIIRRGVIIVSGAGAGGGRLRRFRLGAAMAIIYMAMSTLFFLRIAPLTPWWGALAAWVMILLIVEAIFAFFFPYRCCFNETEGDVQNNPHV
IRIS_313-11225 MELITACALLLIAVHIAIVISCPEPLAAVALVDVPLLYLCRYLLIIRRGVIIVSGGGGGGRLRRFRLGAAMAIIYMAMSTLFFLRIAPLTPWWGALAAWVMILLIVEAIFAFFFPYRCCFNETEGDVQNNPHV
IRIS_313-11226 MELITACALLLIAVHIAIVISCPEPLAAVALVDVPLLYLCRYLLIIRRGVIIVSGAGAGGGRLRRFRLGAAMAIIYMAMSTLFFLRIAPLTPWWGALAAWVMILLIVEAIFAFFFPYRCCFNETEGDVQNNPHV
IRIS_313-11227 MELITACALLLIAVHIAIVISCPEPLAAVALVDVPLLYLCRYLLIIRRGVIIVSGAGAGGGRLRRFRLGAAMAIIYMAMSTLFFLRIAPLTPWWGALAAWVMILLIVEAIFAFFFPYRCCFNETEGDVQNNPHV
IRIS_313-11228 MELITACALLLIAVHIAIVISCPEPLAAVALVDVPLLYLCRYLLIIRRGVIIVSGAGAGGGRLRRFRLGAAMAIIYMAMSTLFFLRIAPLTPWWGALAAWVMILLIVEAIFAFFFPYRCCFNETEGDVQNNPHV
IRIS_313-11229 MELITACALLLIAVHIAIVISCPEPLAAVALVDVPLLYLCRYLLIIRRGVIIVSGGGGGGRLRRFRLGAAMAIIYMAMSTLFFLRIAPLTPWWGALAAWVMILLIVEAIFAFFFPYRCCFNETEGDVQNNPHV
IRIS_313-11230 MELITACALLLIAVHIAIVISCPEPLAAVALVDVPLLYLCRYLLIIRRGVIIVSGGGAGGGRLRRFRLGAAMAIIYMAMSTLFFLRIAPLTPWWGALAAWVMILLIVEAIFAFFFPYRCCFNETEGDVQNNPHV
IRIS_313-11231 MELITACALLLIAVHIAIVISCPEPLAAVALVDVPLLYLCRYLLIIRRGVIIVSGAGAGGGRLRRFRLGAAMAIIYMAMSTLFFLRIAPLTPWWGALAAWVMILLIVEAIFAFFFPYRCCFNETEGDVQNNPHV
IRIS_313-11232 MELITACALLLIAVHIAIVISCPEPLAAVALVDVPLLYLCRYLLIIRRGVIIVSGAGAGGGRLRRFRLGAAMAIIYMAMSTLFFLRIAPLTPWWGALAAWVMILLIVEAIFAFFFPYRCCFNETEGDVQNNPHV
IRIS_313-11233 MELITACALLLIAVHIAIVISCPEPLAAVALVDVPLLYLCRYLLIIRRGVIIVSGGGAGGGRLRRFRLGAAMAIIYMAMSTLFFLRIAPLTPWWGALAAWVMILLIVEAIFAFFFPYRCCFNETEGDVQNNPHV
IRIS_313-11234 MELITACALLLIAVHIAIVISCPEPLAAVALVDVPLLYLCRYLLIIRRGVIIVSGAGAGGGRLRRFRLGAAMAIIYMAMSTLFFLRIAPLTPWWGALAAWVMILLIVEAIFAFFFPYRCCFNETEGDVQNNPHV
IRIS_313-11235 MELITACALLLIAVHIAIVISCPEPLAAVALVDVPLLYLCRYLLIIRRGVIIVSGAGAGGGRLRRFRLGAAMAIIYMAMSTLFFLRIAPLTPWWGALAAWVMILLIVEAIFAFFFPYRCCFNETEGDVQNNPHV
IRIS_313-11236 MELITACALLLIAVHIAIVISCPEPLAAVALVDVPLLYLCRYLLIIRRGVIIVSGGGAGGGRLRRFRLGAAMAIIYMAMSTLFFLRIAPLTPWWGALAAWVMILLIVEAIFAFFFPYRCCFNETEGDVQNNPHV
IRIS_313-11237 MELITACALLLIAVHIAIVISCPEPLAAVALVDVPLLYLCRYLLIIRRGVIIVSGAGAGGGRLRRFRLGAAMAIIYMAMSTLFFLRIAPLTPWWGALAAWVMILLIVEAIFAFFFPYRCCFNETEGDVQNNPHV
IRIS_313-11238 MELITACALLLIAVHIAIVISCPEPLAAVALVDVPLLYLCRYLLIIRRGVIIVSGAGAGGGRLRRFRLGAAMAIIYMAMSTLFFLRIAPLTPWWGALAAWVMILLIVEAIFAFFFPYRCCFNETEGDVQNNPHV
IRIS_313-11239 MELITACALLLIAVHIAIVISCPEPLAAVALVDVPLLYLCRYLLIIRRGVIIVSGGGGGGRLRRFRLGAAMAIIYMAMSTLFFLRIAPLTPWWGALAAWVMILLIVEAIFAFFFPYRCCFNETEGDVQNNPHV
IRIS_313-11240 MELITACALLLIAVHIAIVISCPEPLAAVALVDVPLLYLCRYLLIIRRGVIIVSGAGAGGGRLRRFRLGAAMAIIYMAMSTLFFLRIAPLTPWWGALAAWVMILLIVEAIFAFFFPYRCCFNETEGDVQNNPHV
IRIS_313-11241 MELITACALLLIAVHIAIVISCPEPLAAVALVDVPLLYLCRYLLIIRRGVIIVSGGGGGGRLRRFRLGAAMAIIYMAMSTLFFLRIAPLTPWWGALAAWVMILLIVEAIFAFFFPYRCCFNETEGDVQNNPHV
IRIS_313-11242 MELITACALLLIAVHIAIVISCPEPLAAVALVDVPLLYLCRYLLIIRRGVIIVSGGGGGGRLRRFRLGAAMAIIYMAMSTLFFLRIAPLTPWWGALAAWVMILLIVEAIFAFFFPYRCCFNETEGDVQNNPHV
IRIS_313-11243 MELITACALLLIAVHIAIVISCPEPLAAVALVDVPLLYLCRYLLIIRRGVIIVSGAGAGGGRLRRFRLGAAMAIIYMAMSTLFFLRIAPLTPWWGALAAWVMILLIVEAIFAFFFPYRCCFNETEGDVQNNPHV
IRIS_313-11244 MELITACALLLIAVHIAIVISCPEPLAAVALVDVPLLYLCRYLLIIRRGVIIVSGGGGGGRLRRFRLGAAMAIIYMAMSTLFFLRIAPLTPWWGALAAWVMILLIVEAIFAFFFPYRCCFNETEGDVQNNPHV
IRIS_313-11245 MELITACALLLIAVHIAIVISCPEPLAAVALVDVPLLYLCRYLLIIRRGVIIVSGGGGGGRLRRFRLGAAMAIIYMAMSTLFFLRIAPLTPWWGALAAWVMILLIVEAIFAFFFPYRCCFNETEGDVQNNPHV
IRIS_313-11246 MELITACALLLIAVHIAIVISCPEPLAAVALVDVPLLYLCRYLLIIRRGVIIVSGAGAGGGRLRRFRLGAAMAIIYMAMSTLFFLRIAPLTPWWGALAAWVMILLIVEAIFAFFFPYRCCFNETEGDVQNNPHV
IRIS_313-11247 MELITACALLLIAVHIAIVISCPEPLAAVALVDVPLLYLCRYLLIIRRGVIIVSGAGAGGGRLRRFRLGAAMAIIYMAMSTLFFLRIAPLTPWWGALAAWVMILLIVEAIFAFFFPYRCCFNETEGDVQNNPHV
IRIS_313-11248 MELITACALLLIAVHIAIVISCPEPLAAVALVDVPLLYLYRYLLIIRRGVIIVSGAGGGGGRLRRFRLGAAMAIIYMAMSTLFFLRIAPLTPWWGALAAWVMILLIVEAIFAFFFPYRCCFNETEGDVQNNPHV
IRIS_313-11249 MELITACALLLIAVHIAIVISCPEPLAAVALVDVPLLYLCRYLLIIRRGVIIVSGGGAGGGRLRRFRLGAAMAIIYMAMSTLFFLRIAPLTPWWGALAAWVMILLIVEAIFAFFFPYRCCFNETEGDVQNNPHV
IRIS_313-11250 MELITACALLLIAVHIAIVISCPEPLAAVALVDVPLLYLCRYLLIIRRGVIIVSGAGAGGGRLRRFRLGAAMAIIYMAMSTLFFLRIAPLTPWWGALAAWVMILLIVEAIFAFFFPYRCCFNETEGDVQNNPHV
IRIS_313-11251 MELITACALLLIAVHIAIVISCPEPLAAVALVDVPLLYLCRYLLIIRRGVIIVSGGGGGGRLRRFRLGAAMAIIYMAMSTLFFLRIAPLTPWWGALAAWVMILLIVEAIFAFFFPYRCCFNETEGDVQNNPHV
IRIS_313-11252 MELITACALLLIAVHIAIVISCPEPLAAVALVDVPLLYLCRYLLIIRRGVIIVSGAGAGGGRLRRFRLGAAMAIIYMAMSTLFFLRIAPLTPWWGALAAWVMILLIVEAIFAFFFPYRCCFNETEGDVQNNPHV
IRIS_313-11253 MELITACALLLIAVHIAIVISCPEPLAAVALVDVPLLYLCRYLLIIRRGVIIVSGAGAGGGRLRRFRLGAAMAIIYMAMSTLFFLRIAPLTPWWGALAAWVMILLIVEAIFAFFFPYRCCFNETEGDVQNNPHV
IRIS_313-11254 MELITACALLLIAVHIAIVISCPEPLAAVALVDVPLLYLCRYLLIIRRGVIIVSGAGAGGGRLRRFRLGAAMAIIYMAMSTLFFLRIAPLTPWWGALAAWVMILLIVEAIFAFFFPYRCCFNETEGDVQNNPHV
IRIS_313-11255 MELITACALLLIAVHIAIVISCPEPLAAVALVDVPLLYLCRYLLIIRRGVIIVSGGGAGGGRLRRFRLGAAMAIIYMAMSTLFFLRIAPLTPWWGALAAWVMILLIVEAIFAFFFPYRCCFNETEGDVQNNPHV
IRIS_313-11256 MELITACALLLIAVHIAIVISCPEPLAAVALVDVPLLYLCRYLLIIRRGVIIVSGGGGGGRLRRFRLGAAMAIIYMAMSTLFFLRIAPLTPWWGALAAWVMILLIVEAIFAFFFPYRCCFNETEGDVQNNPHV
IRIS_313-11257 MELITACALLLIAVHIAIVISCPEPLAAVALVDVPLLYLCRYLLIIRRGVIIVSGGGGGGRLRRFRLGAAMAIIYMAMSTLFFLRIAPLTPWWGALAAWVMILLIVEAIFAFFFPYRCCFNETEGDVQNNPHV
IRIS_313-11258 MELITACALLLIAVHIAIVISCPEPLAAVALVDVPLLYLCRYLLIIRRGVIIVSGAGAGGGRLRRFRLGAAMAIIYMAMSTLFFLRIAPLTPWWGALAAWVMILLIVEAIFAFFFPYRCCFNETEGDVQNNPHV
IRIS_313-11259 MELITACALLLIAVHIAIVISCPEPLAAVALVDVPLLYLCRYLLIIRRGVIIVSGAGAGGGRLRRFRLGAAMAIIYMAMSTLFFLRIAPLTPWWGALAAWVMILLIVEAIFAFFFPYRCCFNETEGDVQNNPHV
IRIS_313-11260 MELITACALLLIAVHIAIVISCPEPLAAVALVDVPLLYLCRYLLIIRRGVIIVSGGGGGGRLRRFRLGAAMAIIYMAMSTLFFLRIAPLTPWWGALAAWVMILLIVEAIFAFFFPYRCCFNETEGDVQNNPHV
IRIS_313-11261 MELITACALLLIAVHIAIVISCPEPLAAVALVDVPLLYLCRYLLIIRRGVIIVSGGGAGGGRLRRFRLGAAMAIIYMAMSTLFFLRIAPLTPWWGALAAWVMILLIVEAIFAFFFPYRCCFNETEGDVQNNPHV
IRIS_313-11262 MELITACALLLIAVHIAIVISCPEPLAAVALVDVPLLYLCRYLLIIRRGVIIVSGAGAGGGRLRRFRLGAAMAIIYMAMSTLFFLRIAPLTPWWGALAAWVMILLIVEAIFAFFFPYRCCFNETEGDVQNNPHV
IRIS_313-11263 MELITACALLLIAVHIAIVISCPEPLAAVALVDVPLLYLCRYLLIIRRGVIIVSGAGGAAAAGCAASASGQPWRSSTWPCRPSSSSALLLSRHGGAPSPPGS*SCSSSRPSSPSSSPTAAASTKPKAMSRIIPM
IRIS_313-11264 MELITACALLLIAVHIAIVISCPEPLAAVALVDVPLLYLCRYLLIIRRGVIIVSGGGGGGRLRRFRLGAAMAIIYMAMSTLFFLRIAPLTPWWGALAAWVMILLIVEAIFAFFFPYRCCFNETEGDVQNNPHV
IRIS_313-11265 MELITACALLLIAVHIAIVISCPEPLAAVAFVNVPLLYLCRYLLIIRRGVIIVSGGGGGGGRLRRFRLGAAMAVIYMAMSTLFFLRIAPLTPWWGALAAWVMILLIVEAIFAFFFPYRCCFNETEGDVQNNPHV
IRIS_313-11266 MELITACALLLIAVHIAIVISCPEPLAAVALVDVPLLYLCRYLLIIRRGVIIVSGAGAGGGRLRRFRLGAAMAIIYMAMSTLFFLRIAPLTPWWGALAAWVMILLIVEAIFAFFFPYRCCFNETEGDVQNNPHV
IRIS_313-11267 MELITACALLLIAVHIAIVISCPEPLAAVALVDVPLLYLCRYLLIIRRGVIIVSGAGAGGGRLRRFRLGAAMAIIYMAMSTLFFLRIAPLTPWWGALAAWVMILLIVEAIFAFFFPYRCCFNETEGDVQNNPHV
IRIS_313-11268 MELITACALLLIAVHIAIVISCPEPLAAVALVDVPLLYLCRYLLIIRRGVIIVSGAGAGGGRLRRFRLGAAMAIIYMAMSTLFFLRIAPLTPWWGALAAWVMILLIVEAIFAFFFPYRCCFNETEGDVQNNPHV
IRIS_313-11269 MELITACALLLIAVHIAIVISCPEPLAAVALVDVPLLYLCRYLLIIRRGVIIVSGGGGGGRLRRFRLGAAMAIIYMAMSTLFFLRIAPLTPWWGALAAWVMILLIVEAIFAFFFPYRCCFNETEGDVQNNPHV
IRIS_313-11270 MELITACALLLIAVHIAIVISCPEPLAAVALVDVPLLYLCRYLLIIRRGVIIVSGGGGGGRLRRFRLGAAMAIIYMAMSTLFFLRIAPLTPWWGALAAWVMILLIVEAIFAFFFPYRCCFNETEGDVQNNPHV
IRIS_313-11271 MELITACALLLIAVHIAIVISCPEPLAAVALVDVPLLYLCRYLLIIRRGVIIVSGGGAGGGRLRRFRLGAAMAIIYMAMSTLFFLRIAPLTPWWGALAAWVMILLIVEAIFAFFFPYRCCFNETEGDVQNNPHV
IRIS_313-11272 MELITACALLLIAVHIAIVISCPEPLAAVALVDVPLLYLCRYLLIIRRGVIIVSGAGAGGGRLRRFRLGAAMAIIYMAMSTLFFLRIAPLTPWWGALAAWVMILLIVEAIFAFFFPYRCCFNETEGDVQNNPHV
IRIS_313-11273 MELITACALLLIAVHIAIVISCPEPLAAVALVDVPLLYLCRYLLIIRRGVIIVSGAGAGGGRLRRFRLGAAMAIIYMAMSTLFFLRIAPLTPWWGALAAWVMILLIVEAIFAFFFPYRCCFNETEGDVQNNPHV
IRIS_313-11274 MELITACALLLIAVHIAIVISCPEPLAAVALVDVPLLYLCRYLLIIRRGVIIVSGAGAGGGRLRRFRLGAAMAIIYMAMSTLFFLRIAPLTPWWGALAAWVMILLIVEAIFAFFFPYRCCFNETEGDVQNNPHV
IRIS_313-11275 MELITACALLLIAVHIAIVISCPEPLAAVALVDVPLLYLCRYLLIIRRGVIIVSGAGAGGGRLRRFRLGAAMAIIYMAMSTLFFLRIAPLTPWWGALAAWVMILLIVEAIFAFFFPYRCCFNETEGDVQNNPHV
IRIS_313-11276 MELITACALLLIAVHIAIVISCPEPLAAVALVDVPLLYLCRYLLIIRRGVIIVSGGGAGGGRLRRFRLGAAMAIIYMAMSTLFFLRIAPLTPWWGALAAWVMILLIVEAIFAFFFPYRCCFNETEGDVQNNPHV
IRIS_313-11277 MELITACALLLIAVHIAIVISCPEPLAAVALVDVPLLYLCRYLLIIRRGVIIVSGAGAGGGRLRRFRLGAAMAIIYMAMSTLFFLRIAPLTPWWGALAAWVMILLIVEAIFAFFFPYRCCFNETEGDVQNNPHV
IRIS_313-11278 MELITACALLLIAVHIAIVISCPEPLAAVALVDVPLLYLCRYLLIIRRGVIIVSGGGGGGRLRRFRLGAAMAIIYMAMSTLFFLRIAPLTPWWGALAAWVMILLIVEAIFAFFFPYRCCFNETEGDVQNNPHV
IRIS_313-11279 MELITACALLLIAVHIAIVISCPEPLAAVALVDVPLLYLYRYLLIIRRGVIIVSGAGGGGGRLRRFRLGAAMAIIYMAMSTLFFLRIAPLTPWWGALAAWVMILLIVEAIFAFFFPYRCCFNETEGDVQNNPHV
IRIS_313-11280 MELITACALLLIAVHIAIVISCPEPLAAVALVDVPLLYLCRYLLIIRRGVIIVSGGGGGGRLRRFRLGAAMAIIYMAMSTLFFLRIAPLTPWWGALAAWVMILLIVEAIFAFFFPYRCCFNETEGDVQNNPHV
IRIS_313-11281 MELITACALLLIAVHIAIVISCPEPLAAVALVDVPLLYLYRYLLIIRRGVIIVSGAGGGGGRLRRFRLGAAMAIIYMAMSTLFFLRIAPLTPWWGALAAWVMILLIVEAIFAFFFPYRCCFNETEGDVQNNPHV
IRIS_313-11282 MELITACALLLIAVHIAIVISCPEPLAAVALVDVPLLYLCRYLLIIRRGVIIVSGGGAGGGRLRRFRLGAAMAIIYMAMSTLFFLRIAPLTPWWGALAAWVMILLIVEAIFAFFFPYRCCFNETEGDVQNNPHV
IRIS_313-11283 MELITACALLLIAVHIAIVISCPEPLAAVALVDVPLLYLCRYLLIIRRGVIIVSGAGGGGGRLRRFRLGAAMAIIYMAMSTLFFLRIAPLTPWWGALAAWVMILLIVEAIFAFFFPYRCCFNETEGDVQNNPHV
IRIS_313-11284 MELITACALLLIAVHIAIVISCPEPLAAVALVDVPLLYLCRYLLIIRRGVIIVSGAGAGGGRLRRFRLGAAMAIIYMAMSTLFFLRIAPLTPWWGALAAWVMILLIVEAIFAFFFPYRCCFNETEGDVQNNPHV
IRIS_313-11285 MELITACALLLIAVHIAIVISCPEPLAAVALVDVPLLYLCRYLLIIRRGVIIVSGGGGGGRLRRFRLGAAMAIIYMAMSTLFFLRIAPLTPWWGALAAWVMILLIVEAIFAFFFPYRCCFNETEGDVQNNPHV
IRIS_313-11286 MELITACALLLIAVHIAIVISCPEPLAAVALVDVPLLYLCRYLLIIRRGVIIVSGGGGGGRLRRFRLGAAMAIIYMAMSTLFFLRIAPLTPWWGALAAWVMILLIVEAIFAFFFPYRCCFNETEGDVQNNPHV
IRIS_313-11287 MELITACALLLIAVHIAIVISCPEPLAAVALVDVPLLYLCRYLLIIRRGVIIVSGGGGGGRLRRFRLGAAMAIIYMAMSTLFFLRIAPLTPWWGALAAWVMILLIVEAIFAFFFPYRCCFNETEGDVQNNPHV
IRIS_313-11288 MELITACALLLIAVHIAIVISCPEPLAAVALVDVPLLYLCRYLLIIRRGVIIVSGGGGGGRLRRFRLGAAMAIIYMAMSTLFFLRIAPLTPWWGALAAWVMILLIVEAIFAFFFPYRCCFNETEGDVQNNPHV
IRIS_313-11289 MELITACALLLIAVHIAIVISCPEPLAAVALVDVPLLYLCRYLLIIRRGVIIVSGAGAGGGRLRRFRLGAAMAIIYMAMSTLFFLRIAPLTPWWGALAAWVMILLIVEAIFAFFFPYRCCFNETEGDVQNNPHV
IRIS_313-11290 MELITACALLLIAVHIAIVISCPEPLAAVALVDVPLLYLCRYLLIIRRGVIIVSGAGAGGGRLRRFRLGAAMAIIYMAMSTLFFLRIAPLTPWWGALAAWVMILLIVEAIFAFFFPYRCCFNETEGDVQNNPHV
IRIS_313-11291 MELITACALLLIAVHIAIVISCPEPLAAVALVDVPLLYLCRYLLIIRRGVIIVSGAGAGGGRLRRFRLGAAMAIIYMAMSTLFFLRIAPLTPWWGALAAWVMILLIVEAIFAFFFPYRCCFNETEGDVQNNPHV
IRIS_313-11292 MELITACALLLIAVHIAIVISCPEPLAAVALVDVPLLYLCRYLLIIRRGVIIVSGAGAGGGRLRRFRLGAAMAIIYMAMSTLFFLRIAPLTPWWGALAAWVMILLIVEAIFAFFFPYRCCFNETEGDVQNNPHV
IRIS_313-11293 MELITACALLLIAVHIAIVISCPEPLAAVALVDVPLLYLCRYLLIIRRGVIIVSGGGAGGGRLRRFRLGAAMAIIYMAMSTLFFLRIAPLTPWWGALAAWVMILLIVEAIFAFFFPYRCCFNETEGDVQNNPHV
IRIS_313-11294 MELITACALLLIAVHIAIVISCPEPLAAVALVDVPLLYLCRYLLIIRRGVIIVSGGGGGGRLRRFRLGAAMAIIYMAMSTLFFLRIAPLTPWWGALAAWVMILLIVEAIFAFFFPYRCCFNETEGDVQNNPHV
IRIS_313-11295 MELITACALLLIAVHIAIVISCPEPLAAVALVDVPLLYLYRYLLIIRRGVIIVSGAGGGGGRLRRFRLGAAMAIIYMAMSTLFFLRIAPLTPWWGALAAWVMILLIVEAIFAFFFPYRCCFNETEGDVQNNPHV
IRIS_313-11296 MELITACALLLIAVHIAIVISCPEPLAAVALVDVPLLYLCRYLLIIRRGVIIVSGAGAGGGRLRRFRLGAAMAIIYMAMSTLFFLRIAPLTPWWGALAAWVMILLIVEAIFAFFFPYRCCFNETEGDVQNNPHV
IRIS_313-11297 MELITACALLLIAVHIAIVISCPEPLAAVALVDVPLLYLYRYLLIIRRGVIIVSGAGGGGGRLRRFRLGAAMAIIYMAMSTLFFLRIAPLTPWWGALAAWVMILLIVEAIFAFFFPYRCCFNETEGDVQNNPHV
IRIS_313-11298 MELITACALLLIAVHIAIVISCPEPLAAVALVDVPLLYLCRYLLIIRRGVIIVSGAGGGGGRLRRFRLGAAMAIIYMAMSTLFFLRIAPLTPWWGALAAWVMILLIVEAIFAFFFPYRCCFNETEGDVQNNPHV
IRIS_313-11299 MELITACALLLIAVHIAIVISCPEPLAAVALVDVPLLYLCRYLLIIRRGVIIVSGAGGGGGRLRRFRLGAAMAIIYMAMSTLFFLRIAPLTPWWGALAAWVMILLIVEAIFAFFFPYRCCFNETEGDVQNNPHV
IRIS_313-11300 MELITACALLLIAVHIAIVISCPEPLAAVALVDVPLLYLCRYLLIIRRGVIIVSGAGAGGGRLRRFRLGAAMAIIYMAMSTLFFLRIAPLTPWWGALAAWVMILLIVEAIFAFFFPYRCCFNETEGDVQNNPHV
IRIS_313-11301 MELITACALLLIAVHIAIVISCPEPLAAVALVDVPLLYLCRYLLIIRRGVIIVSGGGGGGRLRRFRLGAAMAIIYMAMSTLFFLRIAPLTPWWGALAAWVMILLIVEAIFAFFFPYRCCFNETEGDVQNNPHV
IRIS_313-11302 MELITACALLLIAVHIAIVISCPEPLAAVALVDVPLLYLCRYLLIIRRGVIIVSGGGAGGGRLRRFRLGAAMAIIYMAMSTLFFLRIAPLTPWWGALAAWVMILLIVEAIFAFFFPYRCCFNETEGDVQNNPHV
IRIS_313-11303 MELITACALLLIAVHIAIVISCPEPLAAVALVDVPLLYLCRYLLIIRRGVIIVSGGGAGGGRLRRFRLGAAMAIIYMAMSTLFFLRIAPLTPWWGALAAWVMILLIVEAIFAFFFPYRCCFNETEGDVQNNPHV
IRIS_313-11304 MELITACALLLIAVHIAIVISCPEPLAAVALVDVPLLYLCRYLLIIRRGVIIVSGGGGGGRLRRFRLGAAMAIIYMAMSTLFFLRIAPLTPWWGALAAWVMILLIVEAIFAFFFPYRCCFNETEGDVQNNPHV
IRIS_313-11305 MELITACALLLIAVHIAIVISCPEPLAAVALVDVPLLYLCRYLLIIRRGVIIVSGAGAGGGRLRRFRLGAAMAIIYMAMSTLFFLRIAPLTPWWGALAAWVMILLIVEAIFAFFFPYRCCFNETEGDVQNNPHV
IRIS_313-11306 MELITACALLLIAVHIAIVISCPEPLAAVALVDVPLLYLCRYLLIIRRGVIIVSGAGAGGGRLRRFRLGAAMAIIYMAMSTLFFLRIAPLTPWWGALAAWVMILLIVEAIFAFFFPYRCCFNETEGDVQNNPHV
IRIS_313-11307 MELITACALLLIAVHIAIVISCPEPLAAVALVDVPLLYLYRYLLIIRRGVIIVSGAGGGGGRLRRFRLGAAMAIIYMAMSTLFFLRIAPLTPWWGALAAWVMILLIVEAIFAFFFPYRCCFNETEGDVQNNPHV
IRIS_313-11308 MELITACALLLIAVHIAIVISCPEPLAAVALVDVPLLYLCRYLLIIRRGVIIVSGAGAGGGRLRRFRLGAAMAIIYMAMSTLFFLRIAPLTPWWGALAAWVMILLIVEAIFAFFFPYRCCFNETEGDVQNNPHV
IRIS_313-11309 MELITACALLLIAVHIAIVISCPEPLAAVALVDVPLLYLYRYLLIIRRGVIIVSGAGGGGGRLRRFRLGAAMAIIYMAMSTLFFLRIAPLTPWWGALAAWVMILLIVEAIFAFFFPYRCCFNETEGDVQNNPHV
IRIS_313-11310 MELITACALLLIAVHIAIVISCPEPLAAVALVDVPLLYLCRYLLIIRRGVIIVSGGGAGGGRLRRFRLGAAMAIIYMAMSTLFFLRIAPLTPWWGALAAWVMILLIVEAIFAFFFPYRCCFNETEGDVQNNPHV
IRIS_313-11311 MELITACALLLIAVHIAIVISCPEPLAAVALVDVPLLYLCRYLLIIRRGVIIVSGAGAGGGRLRRFRLGAAMAIIYMAMSTLFFLRIAPLTPWWGALAAWVMILLIVEAIFAFFFPYRCCFNETEGDVQNNPHV
IRIS_313-11312 MELITACALLLIAVHIAIVISCPEPLAAVALVDVPLLYLCRYLLIIRRGVIIVSGAGAGGGRLRRFRLGAAMAIIYMAMSTLFFLRIAPLTPWWGALAAWVMILLIVEAIFAFFFPYRCCFNETEGDVQNNPHV
IRIS_313-11313 MELITACALLLIAVHIAIVISCPEPLAAVALVDVPLLYLCRYLLIIRRGVIIVSGAGAGGGRLRRFRLGAAMAIIYMAMSTLFFLRIAPLTPWWGALAAWVMILLIVEAIFAFFFPYRCCFNETEGDVQNNPHV
IRIS_313-11314 MELITACALLLIAVHIAIVISCPEPLAAVALVDVPLLYLCRYLLIIRRGVIIVSGAGAGGGRLRRFRLGAAMAIIYMAMSTLFFLRIAPLTPWWGALAAWVMILLIVEAIFAFFFPYRCCFNETEGDVQNNPHV
IRIS_313-11315 MELITACALLLIAVHIAIVISCPEPLAAVALVDVPLLYLCRYLLIIRRGVIIVSGGGAGGGRLRRFRLGAAMAIIYMAMSTLFFLRIAPLTPWWGALAAWVMILLIVEAIFAFFFPYRCCFNETEGDVQNNPHV
IRIS_313-11316 MELITACALLLIAVHIAIVISCPEPLAAVALVDVPLLYLCRYLLIIRRGVIIVSGAGAGGGRLRRFRLGAAMAIIYMAMSTLFFLRIAPLTPWWGALAAWVMILLIVEAIFAFFFPYRCCFNETEGDVQNNPHV
IRIS_313-11317 MELITACALLLIAVHIAIVISCPEPLAAVALVDVPLLYLCRYLLIIRRGVIIVSGAGAGGGRLRRFRLGAAMAIIYMAMSTLFFLRIAPLTPWWGALAAWVMILLIVEAIFAFFFPYRCCFNETEGDVQNNPHV
IRIS_313-11318 MELITACALLLIAVHIAIVISCPEPLAAVALVDVPLLYLCRYLLIIRRGVIIVSGAGAGGGRLRRFRLGAAMAIIYMAMSTLFFLRIAPLTPWWGALAAWVMILLIVEAIFAFFFPYRCCFNETEGDVQNNPHV
IRIS_313-11319 MELITACALLLIAVHIAIVISCPEPLAAVALVDVPLLYLCRYLLIIRRGVIIVSGAGAGGGRLRRFRLGAAMAIIYMAMSTLFFLRIAPLTPWWGALAAWVMILLIVEAIFAFFFPYRCCFNETEGDVQNNPHV
IRIS_313-11320 MELITACALLLIAVHIAIVISCPEPLAAVALVDVPLLYLCRYLLIIRRGVIIVSGAGGGGGGRLRRFRLGAAMAIIYMAMSTLFFLRIAPLTPWWGALAAWVMILLIVEAIFAFFFPYRCCFNETEGDVQNNPHV
IRIS_313-11321 MELITACALLLIAVHIAIVISCPEPLAAVALVDVPLLYLCRYLLIIRRGVIIVSGAGAGGGRLRRFRLGAAMAIIYMAMSTLFFLRIAPLTPWWGALAAWVMILLIVEAIFAFFFPYRCCFNETEGDVQNNPHV
IRIS_313-11322 MELITACALLLIAVHIAIVISCPEPLAAVAFVNVPLLYLCRYLLIIRRGVIIVSGGGGGGGRLRRFRLGAAMAVIYMAMSTLFFLRIAPLTPWWGALAAWVMILLIVEAIFAFFFPYRCCFNETEGDVQNNPHV
IRIS_313-11323 MELITACALLLIAVHIAIVISCPEPLAAVALVDVPLLYLCRYLLIIRRGVIIVSGAGAGGGRLRRFRLGAAMAIIYMAMSTLFFLRIAPLTPWWGALAAWVMILLIVEAIFAFFFPYRCCFNETEGDVQNNPHV
IRIS_313-11324 MELITACALLLIAVHIAIVISCPEPLAAVAFVNVPLLYLCRYLLIIRRGVIIVSGGGGGGGRLRRFRLGAAMAVIYMAMSTLFFLRIAPLTPWWGALAAWVMILLIVEAIFAFFFPYRCCFNETEGDVQNNPHV
IRIS_313-11325 MELITACALLLIAVHIAIVISCPEPLAAVALVDVPLLYLCRYLLIIRRGVIIVSGGGAGGGRLRRFRLGAAMAIIYMAMSTLFFLRIAPLTPWWGALAAWVMILLIVEAIFAFFFPYRCCFNETEGDVQNNPHV
IRIS_313-11326 MELITACALLLIAVHIAIVISCPEPLAAVALVDVPLLYLCRYLLIIRRGVIIVSGGGGGGRLRRFRLGAAMAIIYMAMSTLFFLRIAPLTPWWGALAAWVMILLIVEAIFAFFFPYRCCFNETEGDVQNNPHV
IRIS_313-11327 MELITACALLLIAVHIAIVISCPEPLAAVALVDVPLLYLCRYLLIIRRGVIIVSGAGAGGGRLRRFRLGAAMAIIYMVMSTLFFLRIAPLTPWWGALAAWVMILLIVEAIFAFFFPYRCCFNETEGDVQNNPHV
IRIS_313-11328 MELITACALLLIAVHIAIVISCPEPLAAVALVDVPLLYLCRYLLIIRRGVIIVSGAGGGGGRLRRFRLGAAMAIIYMAMSTLFFLRIAPLTPWWGALAAWVMILLIVEAIFAFFFPYRCCFNETEGDVQNNPHV
IRIS_313-11329 MELITACALLLIAVHIAIVISCPEPLAAVALVDVPLLYLCRYLLIIRRGVIIVSGAGAGGGRLRRFRLGAAMAIIYMAMSTLFFLRIAPLTPWWGALAAWVMILLIVEAIFAFFFPYRCCFNETEGDVQNNPHV
IRIS_313-11330 MELITACALLLIAVHIAIVISCPEPLAAVALVDVPLLYLCRYLLIIRRGVIIVSGGGAGGGRLRRFRLGAAMAIIYMAMSTLFFLRIAPLTPWWGALAAWVMILLIVEAIFAFFFPYRCCFNETEGDVQNNPHV
IRIS_313-11331 MELITACALLLIAVHIAIVISCPEPLAAVALVDVPLLYLCRYLLIIRRGVIIVSGGGAGGGRLRRFRLGAAMAIIYMAMSTLFFLRIAPLTPWWGALAAWVMILLIVEAIFAFFFPYRCCFNETEGDVQNNPHV
IRIS_313-11334 MELITACALLLIAVHIAIVISCPEPLAAVALVDVPLLYLCRYLLIIRRGVIIVSGGGGGGRLRRFRLGAAMAIIYMAMSTLFFLRIAPLTPWWGALAAWVMILLIVEAIFAFFFPYRCCFNETEGDVQNNPHV

```
IRIS_313-11335 MELITACALLLIAVHIAIVISCPEPLAAVALVDVPLLYLCRYLLIIRRGVIIVSGAGAGGGRLRRFRLGAAMAIIYMAMSTLFFLRIAPLTPWWGALAAWVMILLIVEAIFAFFFPYRCCFNETEGDVQNNPHV
IRIS_313-11336 MELITACALLLIAVHIAIVISCPEPLAAVALVDVPLLYLCRYLLIIRRGVIIVSGAGAGGGRLRRFRLGAAMAIIYMAMSTLFFLRIAPLTPWWGALAAWVMILLIVEAIFAFFFPYRCCFNETEGDVQNNPHV
IRIS_313-11337 MELITACALLLIAVHIAIVISCPEPLAAVALVDVPLLYLCRYLLIIRRGVIIVSGAGGGGGRLRRFRLGAAMAIIYMAMSTLFFLRIAPLTPWWGALAAWVMILLIVEAIFAFFFPYRCCFNETEGDVQNNPHV
IRIS_313-11338 MELITACALLLIAVHIAIVISCPEPLAAVALVDVPLLYLCRYLLIIRRGVIIVSGAGGAAAAGCAASASGQPWRSSTWPCRPSSSSALLLSRHGGAPSPPGS*SCSSSRPSSPSSSPTAAASTKPKAMSRIIPM
IRIS_313-11339 MELITACALLLIAVHIAIVISCPEPLAAVALVDVPLLYLCRYLLIIRRGVIIVSGAGAGGGRLRRFRLGAAMAIIYMAMSTLFFLRIAPLTPWWGALAAWVMILLIVEAIFAFFFPYRCCFNETEGDVQNNPHV
IRIS_313-11340 MELITACALLLIAVHIAIVISCPEPLAAVALVDVPLLYLCRYLLIIRRGVIIVSGAGGGGGRLRRFRLGAAMAIIYMAMSTLFFLRIAPLTPWWGALAAWVMILLIVEAIFAFFFPYRCCFNETEGDVQNNPHV
IRIS_313-11341 MELITACALLLIAVHIAIVISCPEPLAAVALVDVPLLYLCRYLLIIRRGVIIVSGAGGGGGGRLRRFRLGAAMAIIYMAMSTLFFLRIAPLTPWWGALAAWVMILLIVEAIFAFFFPYRCCFNETEGDVQNNPHV
IRIS_313-11342 MELITACALLLIAVHIAIVISCPEPLAAVALVDVPLLYLCRYLLIIRRGVIIVSGAGAGGGRLRRFRLGAAMAIIYMAMSTLFFLRIAPLTPWWGALAAWVMILLIVEAIFAFFFPYRCCFNETEGDVQNNPHV
IRIS_313-11343 MELITACALLLIAVHIAIVISCPEPLAAVALVDVPLLYLCRYLLIIRRGVIIVSGGGAGGGRLRRFRLGAAMAIIYMAMSTLFFLRIAPLTPWWGALAAWVMILLIVEAIFAFFFPYRCCFNETEGDVQNNPHV
IRIS_313-11344 MELITACALLLIAVHIAIVISCPEPLAAVALVDVPLLYLCRYLLIIRRGVIIVSGAGAGGGRLRRFRLGAAMAIIYMAMSTLFFLRIAPLTPWWGALAAWVMILLIVEAIFAFFFPYRCCFNETEGDVQNNPHV
IRIS_313-11345 MELITACALLLIAVHIAIVISCPEPLAAVALVDVPLLYLCRYLLIIRRGVIIVSGAGGGGGGRLRRFRLGAAMAIIYMAMSTLFFLRIAPLTPWWGALAAWVMILLIVEAIFAFFFPYRCCFNETEGDVQNNPHV
IRIS_313-11346 MELITACALLLIAVHIAIVISCPEPLAAVALVDVPLLYLCRYLLIIRRGVIIVSGGGGGGRLRRFRLGAAMAIIYMAMSTLFFLRIAPLTPWWGALAAWVMILLIVEAIFAFFFPYRCCFNETEGDVQNNPHV
IRIS_313-11347 MELITACALLLIAVHIAIVISCPEPLAAVALVDVPLLYLCRYLLIIRRGVIIVSGAGGGGGRLRRFRLGAAMAIIYMAMSTLFFLRIAPLTPWWGALAAWVMILLIVEAIFAFFFPYRCCFNETEGDVQNNPHV
IRIS_313-11348 MELITACALLLIAVHIAIVISCPEPLAAVALVDVPLLYLCRYLLIIRRGVIIVSGAGGGGGRLRRFRLGAAMAIIYMAMSTLFFLRIAPLTPWWGALAAWVMILLIVEAIFAFFFPYRCCFNETEGDVQNNPHV
IRIS_313-11349 MELITACALLLIAVHIAIVISCPEPLAAVALVDVPLLYLCRYLLIIRRGVIIVSGGGAGGGRLRRFRLGAAMAIIYMAMSTLFFLRIAPLTPWWGALAAWVMILLIVEAIFAFFFPYRCCFNETEGDVQNNPHV
IRIS_313-11350 MELITACALLLIAVHIAIVISCPEPLAAVALVDVPLLYLCRYLLIIRRGVIIVSGAGAGGGRLRRFRLGAAMAIIYMAMSTLFFLRIAPLTPWWGALAAWVMILLIVEAIFAFFFPYRCCFNETEGDVQNNPHV
IRIS_313-11351 MELITACALLLIAVHIAIVISCPEPLAAVALVDVPLLYLCRYLLIIRRGVIIVSGAGAGGGRLRRFRLGAAMAIIYMAMSTLFFLRIAPLTPWWGALAAWVMILLIVEAIFAFFFPYRCCFNETEGDVQNNPHV
IRIS_313-11352 MELITACALLLIAVHIAIVISCPEPLAAVALVDVPLLYLCRYLLIIRRGVIIVSGAGAGGGRLRRFRLGAAMAIIYMAMSTLFFLRIAPLTPWWGALAAWVMILLIVEAIFAFFFPYRCCFNETEGDVQNNPHV
IRIS_313-11353 MELITACALLLIAVHIAIVISCPEPLAAVALVDVPLLYLCRYLLIIRRGVIIVSGAGAGGGRLRRFRLGAAMAIIYMAMSTLFFLRIAPLTPWWGALAAWVMILLIVEAIFAFFFPYRCCFNETEGDVQNNPHV
IRIS_313-11354 MELITACALLLIAVHIAIVISCPEPLAAVALVDVPLLYLCRYLLIIRRGVIIVSGGGAGGGRLRRFRLGAAMAIIYMAMSTLFFLRIAPLTPWWGALAAWVMILLIVEAIFAFFFPYRCCFNETEGDVQNNPHV
IRIS_313-11355 MELITACALLLIAVHIAIVISCPEPLAAVALVDVPLLYLCRYLLIIRRGVIIVSGAGAGGGRLRRFRLGAAMAIIYMAMSTLFFLRIAPLTPWWGALAAWVMILLIVEAIFAFFFPYRCCFNETEGDVQNNPHV
IRIS_313-11356 MELITACALLLIAVHIAIVISCPEPLAAVALVDVPLLYLCRYLLIIRRGVIIVSGAGAGGGRLRRFRLGAAMAIIYMAMSTLFFLRIAPLTPWWGALAAWVMILLIVEAIFAFFFPYRCCFNETEGDVQNNPHV
IRIS_313-11357 MELITACALLLIAVHIAIVISCPEPLAAVALVDVPLLYLCRYLLIIRRGVIIVSGGGGGGRLRRFRLGAAMAIIYMAMSTLFFLRIAPLTPWWGALAAWVMILLIVEAIFAFFFPYRCCFNETEGDVQNNPHV
IRIS_313-11358 MELITACALLLIAVHIAIVISCPEPLAAVALVDVPLLYLCRYLLIIRRGVIIVSGGGGGGRLRRFRLGAAMAIIYMAMSTLFFLRIAPLTPWWGALAAWVMILLIVEAIFAFFFPYRCCFNETEGDVQNNPHV
IRIS_313-11359 MELITACALLLIAVHIAIVISCPEPLAAVALVDVPLLYLCRYLLIIRRGVIIVSGAGAGGGRLRRFRLGAAMAIIYMAMSTLFFLRIAPLTPWWGALAAWVMILLIVEAIFAFFFPYRCCFNETEGDVQNNPHV
IRIS_313-11360 MELITACALLLIAVHIAIVISCPEPLAAVALVDVPLLYLCRYLLIIRRGVIIVSGAGAGGGRLRRFRLGAAMAIIYMAMSTLFFLRIAPLTPWWGALAAWVMILLIVEAIFAFFFPYRCCFNETEGDVQNNPHV
IRIS_313-11361 MELITACALLLIAVHIAIVISCPEPLAAVALVDVPLLYLCRYLLIIRRGVIIVSGAGAGGGRLRRFRLGAAMAIIYMAMSTLFFLRIAPLTPWWGALAAWVMILLIVEAIFAFFFPYRCCFNETEGDVQNNPHV
IRIS_313-11362 MELITACALLLIAVHIAIVISCPEPLAAVALVDVPLLYLCRYLLIIRRGVIIVSGAGAGGGRLRRFRLGAAMAIIYMAMSTLFFLRIAPLTPWWGALAAWVMILLIVEAIFAFFFPYRCCFNETEGDVQNNPHV
IRIS_313-11363 MELITACALLLIAVHIAIVISCPEPLAAVALVDVPLLYLCRYLLIIRRGVIIVSGAGAGGGRLRRFRLGAAMAIIYMAMSTLFFLRIAPLTPWWGALAAWVMILLIVEAIFAFFFPYRCCFNETEGDVQNNPHV
IRIS_313-11364 MELITACALLLIAVHIAIVISCPEPLAAVALVDVPLLYLCRYLLIIRRGVIIVSGAGAGGGRLRRFRLGAAMAIIYMAMSTLFFLRIAPLTPWWGALAAWVMILLIVEAIFAFFFPYRCCFNETEGDVQNNPHV
IRIS_313-11365 MELITACALLLIAVHIAIVISCPEPLAAVALVDVPLLYLCRYLLIIRRGVIIVSGAGAGGGRLRRFRLGAAMAIIYMAMSTLFFLRIAPLTPWWGALAAWVMILLIVEAIFAFFFPYRCCFNETEGDVQNNPHV
IRIS_313-11367 MELITACALLLIAVHIAIVISCPEPLAAVALVDVPLLYLCRYLLIIRRGVIIVSGAGAGGGRLRRFRLGAAMAIIYMAMSTLFFLRIAPLTPWWGALAAWVMILLIVEAIFAFFFPYRCCFNETEGDVQNNPHV
IRIS_313-11368 MELITACALLLIAVHIAIVISCPEPLAAVALVDVPLLYLCRYLLIIRRGVIIVSGAGAGGGRLRRFRLGAAMAIIYMAMSTLFFLRIAPLTPWWGALAAWVMILLIVEAIFAFFFPYRCCFNETEGDVQNNPHV
IRIS_313-11369 MELITACALLLIAVHIAIVISCPEPLAAVALVDVPLLYLCRYLLIIRRGVIIVSGAGAGGGRLRRFRLGAAMAIIYMAMSTLFFLRIAPLTPWWGALAAWVMILLIVEAIFAFFFPYRCCFNETEGDVQNNPHV
IRIS_313-11370 MELITACALLLIAVHIAIVISCPEPLAAVALVDVPLLYLCRYLLIIRRGVIIVSGGGAGGGRLRRFRLGAAMAIIYMAMSTLFFLRIAPLTPWWGALAAWVMILLIVEAIFAFFFPYRCCFNETEGDVQNNPHV
IRIS_313-11371 MELITACALLLIAVHIAIVISCPEPLAAVALVDVPLLYLCRYLLIIRRGVIIVSGGGGGGGRLRRFRLGAAMAVIYMAMSTLFFLRIAPLTPWWGALAAWVMILLIVEAIFAFFFPYRCCFNETEGDVQNNPHV
IRIS_313-11372 MELITACALLLIAVHIAIVISCPEPLAAVALVDVPLLYLCRYLLIIRRGVIIVSGAGGAAAAGCAASASGQPWRSSTWPCRPSSSSALLLSRHGGAPSPPGS*SCSSSRPSSPSSSPTAAASTKPKAMSRIIPM
IRIS_313-11373 MELITACALLLIAVHIAIVISCPEPLAAVALVDVPLLYLCRYLLIIRRGVIIVSGAGAGGGRLRRFRLGAAMAIIYMAMSTLFFLRIAPLTPWWGALAAWVMILLIVEAIFAFFFPYRCCFNETEGDVQNNPHV
IRIS_313-11374 MELITACALLLIAVHIAIVISCPEPLAAVALVDVPLLYLCRYLLIIRRGVIIVSGAGAGGGRLRRFRLGAAMAIIYMAMSTLFFLRIAPLTPWWGALAAWVMILLIVEAIFAFFFPYRCCFNETEGDVQNNPHV
IRIS_313-11375 MELITACALLLIAVHIAIVISCPEPLAAVALVDVPLLYLCRYLLIIRRGVIIVSGAGAGGGRLRRFRLGAAMAIIYMAMSTLFFLRIAPLTPWWGALAAWVMILLIVEAIFAFFFPYRCCFNETEGDVQNNPHV
IRIS_313-11376 MELITACALLLIAVHIAIVISCPEPLAAVALVDVPLLYLCRYLLIIRRGVIIVSGAGAGGGRLRRFRLGAAMAIIYMAMSTLFFLRIAPLTPWWGALAAWVMILLIVEAIFAFFFPYRCCFNETEGDVQNNPHV
IRIS_313-11377 MELITACALLLIAVHIAIVISCPEPLAAVALVDVPLLYLCRYLLIIRRGVIIVSGGGAGGGRLRRFRLGAAMAIIYMAMSTLFFLRIAPLTPWWGALAAWVMILLIVEAIFAFFFPYRCCFNETEGDVQNNPHV
IRIS_313-11379 MELITACALLLIAVHIAIVISCPEPLAAVALVDVPLLYLCRYLLIIRRGVIIVSGAGAGGGRLRRFRLGAAMAIIYMAMSTLFFLRIAPLTPWWGALAAWVMILLIVEAIFAFFFPYRCCFNETEGDVQNNPHV
IRIS_313-11380 MELITACALLLIAVHIAIVISCPEPLAAVALVDVPLLYLCRYLLIIRRGVIIVSGAGAGGGRLRRFRLGAAMAIIYMAMSTLFFLRIAPLTPWWGALAAWVMILLIVEAIFAFFFPYRCCFNETEGDVQNNPHV
IRIS_313-11381 MELITACALLLIAVHIAIVISCPEPLAAVALVDVPLLYLCRYLLIIRRGVIIVSGAGGGGGGRLRRFRLGAAMAIIYMAMSTLFFLRIAPLTPWWGALAAWVMILLIVEAIFAFFFPYRCCFNETEGDVQNNPHV
IRIS_313-11382 MELITACALLLIAVHIAIVISCPEPLAAVALVDVPLLYLYRYLLIIRRGVIIVSGAGGGGGRLRRFRLGAAMAIIYMAMSTLFFLRIAPLTPWWGALAAWVMILLIVEAIFAFFFPYRCCFNETEGDVQNNPHV
IRIS_313-11383 MELITACALLLIAVHIAIVISCPEPLAAVALVDVPLLYLCRYLLIIRRGVIIVSGAGAGGGRLRRFRLGAAMAIIYMAMSTLFFLRIAPLTPWWGALAAWVMILLIVEAIFAFFFPYRCCFNETEGDVQNNPHV
IRIS_313-11384 MELITACALLLIAVHIAIVISCPEPLAAVALVDVPLLYLCRYLLIIRRGVIIVSGAGGGGGRLRRFRLGAAMAIIYMAMSTLFFLRIAPLTPWWGALAAWVMILLIVEAIFAFFFPYRCCFNETEGDVQNNPHV
IRIS_313-11385 MELITACALLLIAVHIAIVISCPEPLAAVALVDVPLLYLCRYLLIIRRGVIIVSGGGAGGGRLRRFRLGAAMAIIYMAMSTLFFLRIAPLTPWWGALAAWVMILLIVEAIFAFFFPYRCCFNETEGDVQNNPHV
IRIS_313-11386 MELITACALLLIAVHIAIVISCPEPLAAVALVDVPLLYLCRYLLIIRRGVIIVSGGGAGGGRLRRFRLGAAMAIIYMAMSTLFFLRIAPLTPWWGALAAWVMILLIVEAIFAFFFPYRCCFNETEGDVQNNPHV
IRIS_313-11387 MELITACALLLIAVHIAIVISCPEPLAAVALVDVPLLYLCRYLLIIRRGVIIVSGAGAGGGRLRRFRLGAAMAIIYMAMSTLFFLRIAPLTPWWGALAAWVMILLIVEAIFAFFFPYRCCFNETEGDVQNNPHV
IRIS_313-11388 MELITACALLLIAVHIAIVISCPEPLAAVALVDVPLLYLCRYLLIIRRGVIIVSGAGAGGGRLRRFRLGAAMAIIYMAMSTLFFLRIAPLTPWWGALAAWVMILLIVEAIFAFFFPYRCCFNETEGDVQNNPHV
IRIS_313-11389 MELITACALLLIAVHIAIVISCPEPLAAVALVDVPLLYLCRYLLIIRRGVIIVSGGGGGGRLRRFRLGAAMAIIYMAMSTLFFLRIAPLTPWWGALAAWVMILLIVEAIFAFFFPYRCCFNETEGDVQNNPHV
IRIS_313-11390 MELITACALLLIAVHIAIVISCPEPLAAVALVDVPLLYLCRYLLIIRRGVIIVSGAGAGGGRLRRFRLGAAMAIIYMAMSTLFFLRIAPLTPWWGALAAWVMILLIVEAIFAFFFPYRCCFNETEGDVQNNPHV
IRIS_313-11391 MELITACALLLIAVHIAIVISCPEPLAAVALVDVPLLYLCRYLLIIRRGVIIVSGGGAGGGRLRRFRLGAAMAIIYMAMSTLFFLRIAPLTPWWGALAAWVMILLIVEAIFAFFFPYRCCFNETEGDVQNNPHV
IRIS_313-11392 MELITACALLLIAVHIAIVISCPEPLAAVALVDVPLLYLCRYLLIIRRGVIIVSGGGAGGGRLRRFRLGAAMAIIYMAMSTLFFLRIAPLTPWWGALAAWVMILLIVEAIFAFFFPYRCCFNETEGDVQNNPHV
IRIS_313-11393 MELITACALLLIAVHIAIVISCPEPLAAVALVDVPLLYLCRYLLIIRRGVIIVSGAGAGGGRLRRFRLGAAMAIIYMAMSTLFFLRIAPLTPWWGALAAWVMILLIVEAIFAFFFPYRCCFNETEGDVQNNPHV
IRIS_313-11394 MELITACALLLIAVHIAIVISCPEPLAAVALVDVPLLYLCRYLLIIRRGVIIVSGAGAGGGRLRRFRLGAAMAIIYMAMSTLFFLRIAPLTPWWGALAAWVMILLIVEAIFAFFFPYRCCFNETEGDVQNNPHV
IRIS_313-11395 MELITACALLLIAVHIAIVISCPEPLAAVALVDVPLLYLCRYLLIIRRGVIIVSGAGGAAAAGCAASASGQPWRSSTWPCRPSSSSALLLSRHGGAPSPPGS*SCSSSRPSSPSSSPTAAASTKPKAMSRIIPM
IRIS_313-11396 MELITACALLLIAVHIAIVISCPEPLAAVALVDVPLLYLCRYLLIIRRGVIIVSGAGAGGGRLRRFRLGAAMAIIYMAMSTLFFLRIAPLTPWWGALAAWVMILLIVEAIFAFFFPYRCCFNETEGDVQNNPHV
IRIS_313-11397 MELITACALLLIAVHIAIVISCPEPLAAVALVDVPLLYLYRYLLIIRRGVIIVSGAGGGGGRLRRFRLGAAMAIIYMAMSTLFFLRIAPLTPWWGALAAWVMILLIVEAIFAFFFPYRCCFNETEGDVQNNPHV
IRIS_313-11398 MELITACALLLIAVHIAIVISCPEPLAAVALVDVPLLYLCRYLLIIRRGVIIVSGAGAGGGRLRRFRLGAAMAIIYMAMSTLFFLRIAPLTPWWGALAAWVMILLIVEAIFAFFFPYRCCFNETEGDVQNNPHV
IRIS_313-11399 MELITACALLLIAVHIAIVISCPEPLAAVALVDVPLLYLCRYLLIIRRGVIIVSGAGAGGGRLRRFRLGAAMAIIYMAMSTLFFLRIAPLTPWWGALAAWVMILLIVEAIFAFFFPYRCCFNETEGDVQNNPHV
IRIS_313-11400 MELITACALLLIAVHIAIVISCPEPLAAVALVDVPLLYLCRYLLIIRRGVIIVSGGGAGGGRLRRFRLGAAMAIIYMAMSTLFFLRIAPLTPWWGALAAWVMILLIVEAIFAFFFPYRCCFNETEGDVQNNPHV
IRIS_313-11401 MELITACALLLIAVHIAIVISCPEPLAAVALVDVPLLYLCRYLLIIRRGVIIVSGAGAGGGRLRRFRLGAAMAIIYMAMSTLFFLRIAPLTPWWGALAAWVMILLIVEAIFAFFFPYRCCFNETEGDVQNNPHV
IRIS_313-11402 MELITACALLLIAVHIAIVISCPEPLAAVALVDVPLLYLYRYLLIIRRGVIIVSGAGGGGGRLRRFRLGAAMAIIYMAMSTLFFLRIAPLTPWWGALAAWVMILLIVEAIFAFFFPYRCCFNETEGDVQNNPHV
IRIS_313-11403 MELITACALLLIAVHIAIVISCPEPLAAVALVDVPLLYLCRYLLIIRRGVIIVSGGGAGGGRLRRFRLGAAMAIIYMAMSTLFFLRIAPLTPWWGALAAWVMILLIVEAIFAFFFPYRCCFNETEGDVQNNPHV
IRIS_313-11404 MELITACALLLIAVHIAIVISCPEPLAAVALVDVPLLYLCRYLLIIRRGVIIVSGAGAGGGRLRRFRLGAAMAIIYMAMSTLFFLRIAPLTPWWGALAAWVMILLIVEAIFAFFFPYRCCFNETEGDVQNNPHV
IRIS_313-11405 MELITACALLLIAVHIAIVISCPEPLAAVALVDVPLLYLCRYLLIIRRGVIIVSGGGGGGRLRRFRLGAAMAIIYMAMSTLFFLRIAPLTPWWGALAAWVMILLIVEAIFAFFFPYRCCFNETEGDVQNNPHV
IRIS_313-11406 MELITACALLLIAVHIAIVISCPEPLAAVALVDVPLLYLYRYLLIIRRGVIIVSGAGGGGGRLRRFRLGAAMAIIYMAMSTLFFLRIAPLTPWWGALAAWVMILLIVEAIFAFFFPYRCCFNETEGDVQNNPHV
IRIS_313-11407 MELITACALLLIAVHIAIVISCPEPLAAVALVDVPLLYLCRYLLIIRRGVIIVSGGGGGGRLRRFRLGAAMAIIYMAMSTLFFLRIAPLTPWWGALAAWVMILLIVEAIFAFFFPYRCCFNETEGDVQNNPHV
IRIS_313-11408 MELITACALLLIAVHIAIVISCPEPLAAVALVDVPLLYLCRYLLIIRRGVIIVSGAGAGGGRLRRFRLGAAMAIIYMAMSTLFFLRIAPLTPWWGALAAWVMILLIVEAIFAFFFPYRCCFNETEGDVQNNPHV
IRIS_313-11409 MELITACALLLIAVHIAIVISCPEPLAAVALVDVPLLYLCRYLLIIRRGVIIVSGGGGGGGRLRRFRLGAAMAIIYMAMSTLFFLRIAPLTPWWGALAAWVMILLIVEAIFAFFFPYRCCFNETEGDVQNNPHV
IRIS_313-11410 MELITACALLLIAVHIAIVISCPEPLAAVALVDVPLLYLCRYLLIIRRGVIIVSGAGAGGGRLRRFRLGAAMAIIYMAMSTLFFLRIAPLTPWWGALAAWVMILLIVEAIFAFFFPYRCCFNETEGDVQNNPHV
IRIS_313-11411 MELITACALLLIAVHIAIVISCPEPLAAVALVDVPLLYLCRYLLIIRRGVIIVSGGGAGGGRLRRFRLGAAMAIIYMAMSTLFFLRIAPLTPWWGALAAWVMILLIVEAIFAFFFPYRCCFNETEGDVQNNPHV
IRIS_313-11413 MELITACALLLIAVHIAIVISCPEPLAAVAFVNVPLLYLCRYLLIIRRGVIIVSGGGAGGGRLRRFRLGAAMAVIYMAMSTLFFLRIAPLTPWWGALAAWVMILLIVEAIFAFFFPYRCCFNETEGDVQNNPHV
IRIS_313-11414 MELITACALLLIAVHIAIVISCPEPLAAVALVDVPLLYLCRYLLIIRRGVIIVSGGGAGGGRLRRFRLGAAMAIIYMAMSTLFFLRIAPLTPWWGALAAWVMILLIVEAIFAFFFPYRCCFNETEGDVQNNPHV
IRIS_313-11416 MELITACALLLIAVHIAIVISCPEPLAAVALVDVPLLYLCRYLLIIRRGVIIVSGGGGGGRLRRFRLGAAMAIIYMAMSTLFFLRIAPLTPWWGALAAWVMILLIVEAIFAFFFPYRCCFNETEGDVQNNPHV
IRIS_313-11417 MELITACALLLIAVHIAIVISCPEPLAAVALVDVPLLYLCRYLLIIRRGVIIVSGAGAGGGRLRRFRLGAAMAIIYMAMSTLFFLRIAPLTPWWGALAAWVMILLIVEAIFAFFFPYRCCFNETEGDVQNNPHV
IRIS_313-11418 MELITACALLLIAVHIAIVISCPEPLAAVALVDVPLLYLCRYLLIIRRGVIIVSGAGAGGGRLRRFRLGAAMAIIYMAMSTLFFLRIAPLTPWWGALAAWVMILLIVEAIFAFFFPYRCCFNETEGDVQNNPHV
IRIS_313-11419 MELITACALLLIAVHIAIVISCPEPLAAVALVDVPLLYLCRYLLIIRRGVIIVSGAGAGGGRLRRFRLGAAMAIIYMAMSTLFFLRIAPLTPWWGALAAWVMILLIVEAIFAFFFPYRCCFNETEGDVQNNPHV
IRIS_313-11420 MELITACALLLIAVHIAIVISCPEPLAAVALVDVPLLYLCRYLLIIRRGVIIVSGAGAGGGRLRRFRLGAAMAIIYMAMSTLFFLRIAPLTPWWGALAAWVMILLIVEAIFAFFFPYRCCFNETEGDVQNNPHV
IRIS_313-11421 MELITACALLLIAVHIAIVISCPEPLAAVALVDVPLLYLCRYLLIIRRGVIIVSGGGAGGGRLRRFRLGAAMAIIYMAMSTLFFLRIAPLTPWWGALAAWVMILLIVEAIFAFFFPYRCCFNETEGDVQNNPHV
IRIS_313-11422 MELITACALLLIAVHIAIVISCPEPLAAVALVDVPLLYLYRYLLIIRRGVIIVSGAGGGGGRLRRFRLGAAMAIIYMAMSTLFFLRIAPLTPWWGALAAWVMILLIVEAIFAFFFPYRCCFNETEGDVQNNPHV
IRIS_313-11423 MELITACALLLIAVHIAIVISCPEPLAAVALVDVPLLYLCRYLLIIRRGVIIVSGAGAGGGRLRRFRLGAAMAIIYMAMSTLFFLRIAPLTPWWGALAAWVMILLIVEAIFAFFFPYRCCFNETEGDVQNNPHV
IRIS_313-11424 MELITACALLLIAVHIAIVISCPEPLAAVALVDVPLLYLYRYLLIIRRGVIIVSGAGGGGGRLRRFRLGAAMAIIYMAMSTLFFLRIAPLTPWWGALAAWVMILLIVEAIFAFFFPYRCCFNETEGDVQNNPHV
IRIS_313-11425 MELITACALLLIAVHIAIVISCPEPLAAVALVDVPLLYLCRYLLIIRRGVIIVSGAGAGGGRLRRFRLGAAMAIIYMAMSTLFFLRIAPLTPWWGALAAWVMILLIVEAIFAFFFPYRCCFNETEGDVQNNPHV
IRIS_313-11426 MELITACALLLIAVHIAIVISCPEPLAAVALVDVPLLYLCRYLLIIRRGVIIVSGAGGGGGRLRRFRLGAAMAIIYMAMSTLFFLRIAPLTPWWGALAAWVMILLIVEAIFAFFFPYRCCFNETEGDVQNNPHV
IRIS_313-11427 MELITACALLLIAVHIAIVISCPEPLAAVALVDVPLLYLCRYLLIIRRGVIIVSGAGAGGGRLRRFRLGAAMAIIYMAMSTLFFLRIAPLTPWWGALAAWVMILLIVEAIFAFFFPYRCCFNETEGDVQNNPHV
IRIS_313-11428 MELITACALLLIAVHIAIVISCPEPLAAVALVDVPLLYLCRYLLIIRRGVIIVSGAGAGGGRLRRFRLGAAMAIIYMAMSTLFFLRIAPLTPWWGALAAWVMILLIVEAIFAFFFPYRCCFNETEGDVQNNPHV
IRIS_313-11429 MELITACALLLIAVHIAIVISCPEPLAAVALVDVPLLYLCRYLLIIRRGVIIVSGAGGGGGRLRRFRLGAAMAIIYMAMSTLFFLRIAPLTPWWGALAAWVMILLIVEAIFAFFFPYRCCFNETEGDVQNNPHV
IRIS_313-11431 MELITACALLLIAVHIAIVISCPEPLAAVALVDVPLLYLCRYLLIIRRGVIIVSGGGGGGRLRRFRLGAAMAIIYMAMSTLFFLRIAPLTPWWGALAAWVMILLIVEAIFAFFFPYRCCFNETEGDVQNNPHV
IRIS_313-11432 MELITACALLLIAVHIAIVISCPEPLAAVALVDVPLLYLCRYLLIIRRGVIIVSGGGGGGRLRRFRLGAAMAIIYMAMSTLFFLRIAPLTPWWGALAAWVMILLIVEAIFAFFFPYRCCFNETEGDVQNNPHV
IRIS_313-11433 MELITACALLLIAVHIAIVISCPEPLAAVALVDVPLLYLCRYLLIIRRGVIIVSGAGAGGGRLRRFRLGAAMAIIYMAMSTLFFLRIAPLTPWWGALAAWVMILLIVEAIFAFFFPYRCCFNETEGDVQNNPHV
IRIS_313-11434 MELITACALLLIAVHIAIVISCPEPLAAVALVDVPLLYLYRYLLIIRRGVIIVSGAGGGGGRLRRFRLGAAMAIIYMAMSTLFFLRIAPLTPWWGALAAWVMILLIVEAIFAFFFPYRCCFNETEGDVQNNPHV
IRIS_313-11435 MELITACALLLIAVHIAIVISCPEPLAAVALVDVPLLYLCRYLLIIRRGVIIVSGAGAGGGRLRRFRLGAAMAIIYMAMSTLFFLRIAPLTPWWGALAAWVMILLIVEAIFAFFFPYRCCFNETEGDVQNNPHV
IRIS_313-11436 MELITACALLLIAVHIAIVISCPEPLAAVALVDVPLLYLCRYLLIIRRGVIIVSGAGAGGGRLRRFRLGAAMAIIYMAMSTLFFLRIAPLTPWWGALAAWVMILLIVEAIFAFFFPYRCCFNETEGDVQNNPHV
IRIS_313-11437 MELITACALLLIAVHIAIVISCPEPLAAVALVDVPLLYLCRYLLIIRRGVIIVSGGGGGGRLRRFRLGAAMAIIYMAMSTLFFLRIAPLTPWWGALAAWVMILLIVEAIFAFFFPYRCCFNETEGDVQNNPHV
IRIS_313-11438 MELITACALLLIAVHIAIVISCPEPLAAVALVDVPLLYLCRYLLIIRRGVIIVSGGGGGGRLRRFRLGAAMAIIYMAMSTLFFLRIAPLTPWWGALAAWVMILLIVEAIFAFFFPYRCCFNETEGDVQNNPHV
IRIS_313-11439 MELITACALLLIAVHIAIVISCPEPLAAVALVDVPLLYLYRYLLIIRRGVIIVSGAGGGGGRLRRFRLGAAMAIIYMAMSTLFFLRIAPLTPWWGALAAWVMILLIVEAIFAFFFPYRCCFNETEGDVQNNPHV
IRIS_313-11441 MELITACALLLIAVHIAIVISCPEPLAAVALVDVPLLYLCRYLLIIRRGVIIVSGAGAGGGRLRRFRLGAAMAIIYMAMSTLFFLRIAPLTPWWGALAAWVMILLIVEAIFAFFFPYRCCFNETEGDVQNNPHV
IRIS_313-11442 MELITACALLLIAVHIAIVISCPEPLAAVALVDVPLLYLCRYLLIIRRGVIIVSGAGGAAAAGCAASASGQPWRSSTWPCRPSSSSALLLSRHGGAPSPPGS*SCSSSRPSSPSSSPTAAASTKPKAMSRIIPM
IRIS_313-11443 MELITACALLLIAVHIAIVISCPEPLAAVALVDVPLLYLCRYLLIIRRGVIIVSGGGGGGRLRRFRLGAAMAIIYMAMSTLFFLRIAPLTPWWGALAAWVMILLIVEAIFAFFFPYRCCFNETEGDVQNNPHV
IRIS_313-11445 MELITACALLLIAVHIAIVISCPEPLAAVALVDVPLLYLCRYLLIIRRGVIIVSGAGGAAAAGCAASASGQPWRSSTWPCRPSSSSALLLSRHGGAPSPPGS*SCSSSRPSSPSSSPTAAASTKPKAMSRIIPM
IRIS_313-11446 MELITACALLLIAVHIAIVISCPEPLAAVALVDVPLLYLCRYLLIIRRGVIIVSGGGGGGRLRRFRLGAAMAIIYMAMSTLFFLRIAPLTPWWGALAAWVMILLIVEAIFAFFFPYRCCFNETEGDVQNNPHV
IRIS_313-11447 MELITACALLLIAVHIAIVISCPEPLAAVALVDVPLLYLCRYLLIIRRGVIIVSGGGGGGRLRRFRLGAAMAIIYMAMSTLFFLRIAPLTPWWGALAAWVMILLIVEAIFAFFFPYRCCFNETEGDVQNNPHV
IRIS_313-11448 MELITACALLLIAVHIAIVISCPEPLAAVALVDVPLLYLCRYLLIIRRGVIIVSGAGGAAAAGCAASASGQPWRSSTWPCRPSSSSALLLSRHGGAPSPPGS*SCSSSRPSSPSSSPTAAASTKPKAMSRIIPM
IRIS_313-11449 MELITACALLLIAVHIAIVISCPEPLAAVALVDVPLLYLCRYLLIIRRGVIIVSGGGGGGRLRRFRLGAAMAIIYMAMSTLFFLRIAPLTPWWGALAAWVMILLIVEAIFAFFFPYRCCFNETEGDVQNNPHV
IRIS_313-11451 MELITACALLLIAVHIAIVISCPEPLAAVALVDVPLLYLCRYLLIIRRGVIIVSGAGGGGGGRLRRFRLGAAMAIIYMAMSTLFFLRIAPLTPWWGALAAWVMILLIVEAIFAFFFPYRCCFNETEGDVQNNPHV
IRIS_313-11452 MELITACALLLIAVHIAIVISCPEPLAAVALVDVPLLYLCRYLLIIRRGVIIVSGAGAGGGRLRRFRLGAAMAIIYMAMSTLFFLRIAPLTPWWGALAAWVMILLIVEAIFAFFFPYRCCFNETEGDVQNNPHV
IRIS_313-11453 MELITACALLLIAVHIAIVISCPEPLAAVALVDVPLLYLCRYLLIIRRGVIIVSGGGGGGRLRRFRLGAAMAIIYMAMSTLFFLRIAPLTPWWGALAAWVMILLIVEAIFAFFFPYRCCFNETEGDVQNNPHV
IRIS_313-11454 MELITACALLLIAVHIAIVISCPEPLAAVAFVNVPLLYLCRYLLIIRRGVIIVSGGGGGGGRLRRFRLGAAMAVIYMAMSTLFFLRIAPLTPWWGALAAWVMILLIVEAIFAFFFPYRCCFNETEGDVQNNPHV
IRIS_313-11455 MELITACALLLIAVHIAIVISCPEPLAAVALVDVPLLYLCRYLLIIRRGVIIVSGAGAGGGRLRRFRLGAAMAIIYMAMSTLFFLRIAPLTPWWGALAAWVMILLIVEAIFAFFFPYRCCFNETEGDVQNNPHV
IRIS_313-11456 MELITACALLLIAVHIAIVISCPEPLAAVAFVNVPLLYLCRYLLIIRRGVIIVSGGGGGGGRLRRFRLGAAMAVIYMAMSTLFFLRIAPLTPWWGALAAWVMILLIVEAIFAFFFPYRCCFNETEGDVQNNPHV
IRIS_313-11458 MELITACALLLIAVHIAIVISCPEPLAAVALVDVPLLYLCRYLLIIRRGVIIVSGAGAGGGRLRRFRLGAAMAIIYMAMSTLFFLRIAPLTPWWGALAAWVMILLIVEAIFAFFFPYRCCFNETEGDVQNNPHV
IRIS_313-11460 MELITACALLLIAVHIAIVISCPEPLAAVALVDVPLLYLCRYLLIIRRGVIIVSGGGGGGRLRRFRLGAAMAIIYMAMSTLFFLRIAPLTPWWGALAAWVMILLIVEAIFAFFFPYRCCFNETEGDVQNNPHV
IRIS_313-11461 MELITACALLLIAVHIAIVISCPEPLAAVALVDVPLLYLCRYLLIIRRGVIIVSGAGAGGGRLRRFRLGAAMAIIYMAMSTLFFLRIAPLTPWWGALAAWVMILLIVEAIFAFFFPYRCCFNETEGDVQNNPHV
IRIS_313-11462 MELITACALLLIAVHIAIVISCPEPLAAVAFVNVPLLYLCRYLLIIRRGVIIVSGGGGGGGRLRRFRLGAAMAVIYMAMSTLFFLRIAPLTPWWGALAAWVMILLIVEAIFAFFFPYRCCFNETEGDVQNNPHV
IRIS_313-11463 MELITACALLLIAVHIAIVISCPEPLAAVALVDVPLLYLYRYLLIIRRGVIIVSGAGGGGGRLRRFRLGAAMAIIYMAMSTLFFLRIAPLTPWWGALAAWVMILLIVEAIFAFFFPYRCCFNETEGDVQNNPHV
IRIS_313-11464 MELITACALLLIAVHIAIVISCPEPLAAVALVDVPLLYLYRYLLIIRRGVIIVSGAGGGGGRLRRFRLGAAMAIIYMAMSTLFFLRIAPLTPWWGALAAWVMILLIVEAIFAFFFPYRCCFNETEGDVQNNPHV
IRIS_313-11465 MELITACALLLIAVHIAIVISCPEPLAAVALVDVPLLYLCRYLLIIRRGVIIVSGAGGGGGRLRRFRLGAAMAIIYMAMSTLFFLRIAPLTPWWGALAAWVMILLIVEAIFAFFFPYRCCFNETEGDVQNNPHV
IRIS_313-11467 MELITACALLLIAVHIAIVISCPEPLAAVALVDVPLLYLCRYLLIIRRGVIIVSGAGAGGGRLRRFRLGAAMAIIYMAMSTLFFLRIAPLTPWWGALAAWVMILLIVEAIFAFFFPYRCCFNETEGDVQNNPHV
IRIS_313-11471 MELITACALLLIAVHIAIVISCPEPLAAVALVDVPLLYLCRYLLIIRRGVIIVSGAGAGGGRLRRFRLGAAMAIIYMAMSTLFFLRIAPLTPWWGALAAWVMILLIVEAIFAFFFPYRCCFNETEGDVQNNPHV
```

IRIS_313-11472 MELITACALLLIAVHIAIVISCPEPLAAVALVDVPLLYLCRYLLIIRRGVIIVSGGGGGGRLRRFRLGAAMAIIYMAMSTLFFLRIAPLTPWWGALAAWVMILLIVEAIFAFFFPYRCCFNETEGDVQNNPHV
IRIS_313-11473 MELITACALLLIAVHIAIVISCPEPLAAVALVDVPLLYLCRYLLIIRRGVIIVSGAGAGGGRLRRFRLGAAMAIIYMAMSTLFFLRIAPLTPWWGALAAWVMILLIVEAIFAFFFPYRCCFNETEGDVQNNPHV
IRIS_313-11476 MELITACALLLIAVHIAIVISCPEPLAAVALVDVPLLYLCRYLLIIRRGVIIVSGGGGGGRLRRFRLGAAMAIIYMAMSTLFFLRIAPLTPWWGALAAWVMILLIVEAIFAFFFPYRCCFNETEGDVQNNPHV
IRIS_313-11477 MELITACALLLIAVHIAIVISCPEPLAAVAFVNVPLLYLCRYLLIIRRGVIIVSGGGGGGGRLRRFRLGAAMAVIYMAMSTLFFLRIAPLTPWWGALAAWVMILLIVEAIFAFFFPYRCCFNETEGDVQNNPHV
IRIS_313-11478 MELITACALLLIAVHIAIVISCPEPLAAVALVDVPLLYLCRYLLIIRRGVIIVSGGGGGGRLRRFRLGAAMAIIYMAMSTLFFLRIAPLTPWWGALAAWVMILLIVEAIFAFFFPYRCCFNETEGDVQNNPHV
IRIS_313-11479 MELITACALLLIAVHIAIVISCPEPLAAVALVDVPLLYLYRYLLIIRRGVIIVSGAGGGGGRLRRFRLGAAMAIIYMAMSTLFFLRIAPLTPWWGALAAWVMILLIVEAIFAFFFPYRCCFNETEGDVQNNPHV
IRIS_313-11480 MELITACALLLIAVHIAIVISCPEPLAAVALVDVPLLYLCRYLLIIRRGVIIVSGAGAGGGRLRRFRLGAAMAIIYMAMSTLFFLRIAPLTPWWGALAAWVMILLIVEAIFAFFFPYRCCFNETEGDVQNNPHV
IRIS_313-11481 MELITACALLLIAVHIAIVISCPEPLAAVALVDVPLLYLCRYLLIIRRGVIIVSGAGAGGGRLRRFRLGAAMAIIYMAMSTLFFLRIAPLTPWWGALAAWVMILLIVEAIFAFFFPYRCCFNETEGDVQNNPHV
IRIS_313-11482 MELITACALLLIAVHIAIVISCPEPLAAVALVDVPLLYLCRYLLIIRRGVIIVSGAGAGGGRLRRFRLGAAMAIIYMAMSTLFFLRIAPLTPWWGALAAWVMILLIVEAIFAFFFPYRCCFNETEGDVQNNPHV
IRIS_313-11483 MELITACALLLIAVHIAIVISCPEPLAAVAFVNVPLLYLCRYLLIIRRGVIIVSGGGGGGGRLRRFRLGAAMAVIYMAMSTLFFLRIAPLTPWWGALAAWVMILLIVEAIFAFFFPYRCCFNETEGDVQNNPHV
IRIS_313-11484 MELITACALLLIAVHIAIVISCPEPLAAVALVDVPLLYLCRYLLIIRRGVIIVSGAGGGGGRLRRFRLGAAMAIIYMAMSTLFFLRIAPLTPWWGALAAWVMILLIVEAIFAFFFPYRCCFNETEGDVQNNPHV
IRIS_313-11485 MELITACALLLIAVHIAIVISCPEPLAAVALVDVPLLYLCRYLLIIRRGVIIVSGAGAGGGRLRRFRLGAAMAIIYMAMSTLFFLRIAPLTPWWGALAAWVMILLIVEAIFAFFFPYRCCFNETEGDVQNNPHV
IRIS_313-11486 MELITACALLLIAVHIAIVISCPEPLAAVALVDVPLLYLCRYLLIIRRGVIIVSGAGAGGGRLRRFRLGAAMAIIYMAMSTLFFLRIAPLTPWWGALAAWVMILLIVEAIFAFFFPYRCCFNETEGDVQNNPHV
IRIS_313-11487 MELITACALLLIAVHIAIVISCPEPLAAVALVDVPLLYLCRYLLIIRRGVIIVSGGGGGGRLRRFRLGAAMAIIYMAMSTLFFLRIAPLTPWWGALAAWVMILLIVEAIFAFFFPYRCCFNETEGDVQNNPHV
IRIS_313-11489 MELITACALLLIAVHIAIVISCPEPLAAVAFVNVPLLYLCRYLLIIRRGVIIVSGGGGGGGRLRRFRLGAAMAVIYMAMSTLFFLRIAPLTPWWGALAAWVMILLIVEAIFAFFFPYRCCFNETEGDVQNNPHV
IRIS_313-11490 MELITACALLLIAVHIAIVISCPEPLAAVALVDVPLLYLCRYLLIIRRGVIIVSGAGGGGRLRRFRLGAAMAIIYMAMSTLFFLRIAPLTPWWGALAAWVMILLIVEAIFAFFFPYRCCFNETEGDVQNNPHV
IRIS_313-11491 MELITACALLLIAVHIAIVISCPEPLAAVALVDVPLLYLCRYLLIIRRGVIIVSGGGGGGRLRRFRLGAAMAIIYMAMSTLFFLRIAPLTPWWGALAAWVMILLIVEAIFAFFFPYRCCFNETEGDVQNNPHV
IRIS_313-11492 MELITACALLLIAVHIAIVISCPEPLAAVALVDVPLLYLCRYLLIIRRGVIIVSGAGAGGGRLRRFRLGAAMAIIYMAMSTLFFLRIAPLTPWWGALAAWVMILLIVEAIFAFFFPYRCCFNETEGDVQNNPHV
IRIS_313-11493 MELITACALLLIAVHIAIVISCPEPLAAVALVDVPLLYLCRYLLIIRRGVIIVSGGGGGGRLRRFRLGAAMAIIYMAMSTLFFLRIAPLTPWWGALAAWVMILLIVEAIFAFFFPYRCCFNETEGDVQNNPHV
IRIS_313-11494 MELITACALLLIAVHIAIVISCPEPLAAVALVDVPLLYLCRYLLIIRRGVIIVSGAGAGGGRLRRFRLGAAMAIIYMAMSTLFFLRIAPLTPWWGALAAWVMILLIVEAIFAFFFPYRCCFNETEGDVQNNPHV
IRIS_313-11495 MELITACALLLIAVHIAIVISCPEPLAAVALVDVPLLYLCRYLLIIRRGVIIVSGAGAGGGRLRRFRLGAAMAIIYMAMSTLFFLRIAPLTPWWGALAAWVMILLIVEAIFAFFFPYRCCFNETEGDVQNNPHV
IRIS_313-11496 MELITACALLLIAVHIAIVISCPEPLAAVALVDVPLLYLCRYLLIIRRGVIIVSGAGAGGGRLRRFRLGAAMAIIYMAMSTLFFLRIAPLTPWWGALAAWVMILLIVEAIFAFFFPYRCCFNETEGDVQNNPHV
IRIS_313-11497 MELITACALLLIAVHIAIVISCPEPLAAVALVDVPLLYLCRYLLIIRRGVIIVSGGGGGGRLRRFRLGAAMAIIYMAMSTLFFLRIAPLTPWWGALAAWVMILLIVEAIFAFFFPYRCCFNETEGDVQNNPHV
IRIS_313-11498 MELITACALLLIAVHIAIVISCPEPLAAVALVDVPLLYLYRYLLIIRRGVIIVSGAGGGGGRLRRFRLGAAMAIIYMAMSTLFFLRIAPLTPWWGALAAWVMILLIVEAIFAFFFPYRCCFNETEGDVQNNPHV
IRIS_313-11499 MELITACALLLIAVHIAIVISCPEPLAAVALVDVPLLYLCRYLLIIRRGVIIVSGAGAGGGRLRRFRLGAAMAIIYMAMSTLFFLRIAPLTPWWGALAAWVMILLIVEAIFAFFFPYRCCFNETEGDVQNNPHV
IRIS_313-11500 MELITACALLLIAVHIAIVISCPEPLAAVALVDVPLLYLCRYLLIIRRGVIIVSGAGAGGGRLRRFRLGAAMAIIYMAMSTLFFLRIAPLTPWWGALAAWVMILLIVEAIFAFFFPYRCCFNETEGDVQNNPHV
IRIS_313-11505 MELITACALLLIAVHIAIVISCPEPLAAVALVDVPLLYLCRYLLIIRRGVIIVSGAGAGGGRLRRFRLGAAMAIIYMAMSTLFFLRIAPLTPWWGALAAWVMILLIVEAIFAFFFPYRCCFNETEGDVQNNPHV
IRIS_313-11506 MELITACALLLIAVHIAIVISCPEPLAAVALVDVPLLYLCRYLLIIRRGVIIVSGGGGGGRLRRFRLGAAMAIIYMAMSTLFFLRIAPLTPWWGALAAWVMILLIVEAIFAFFFPYRCCFNETEGDVQNNPHV
IRIS_313-11507 MELITACALLLIAVHIAIVISCPEPLAAVALVDVPLLYLCRYLLIIRRGVIIVSGGGGGGRLRRFRLGAAMAIIYMAMSTLFFLRIAPLTPWWGALAAWVMILLIVEAIFAFFFPYRCCFNETEGDVQNNPHV
IRIS_313-11508 MELITACALLLIAVHIAIVISCPEPLAAVALVDVPLLYLCRYLLIIRRGVIIVSGAGGGGGGRLRRFRLGAAMAIIYMAMSTLFFLRIAPLTPWWGALAAWVMILLIVEAIFAFFFPYRCCFNETEGDVQNNPHV
IRIS_313-11509 MELITACALLLIAVHIAIVISCPEPLAAVALVDVPLLYLCRYLLIIRRGVIIVSGGGGGGRLRRFRLGAAMAIIYMAMSTLFFLRIAPLTPWWGALAAWVMILLIVEAIFAFFFPYRCCFNETEGDVQNNPHV
IRIS_313-11510 MELITACALLLIAVHIAIVISCPEPLAAVALVDVPLLYLCRYLLIIRRGVIIVSGAGGAAAAGCAASASGQPWRSSTWPCRPSSSSALLLSRHGGAPSPPGS*SCSSSRPSSPSSSPTAAASTKPKAMSRIIPM
IRIS_313-11511 MELITACALLLIAVHIAIVISCPEPLAAVALVDVPLLYLCRYLLIIRRGVIIVSGAGAGGGRLRRFRLGAAMAIIYMAMSTLFFLRIAPLTPWWGALAAWVMILLIVEAIFAFFFPYRCCFNETEGDVQNNPHV
IRIS_313-11512 MELITACALLLIAVHIAIVISCPEPLAAVALVDVPLLYLCRYLLIIRRGVIIVSGAGGAAAAGCAASASGQPWRSSTWPCRPSSSSALLLSRHGGAPSPPGS*SCSSSRPSSPSSSPTAAASTKPKAMSRIIPM
IRIS_313-11513 MELITACALLLIAVHIAIVISCPEPLAAVALVDVPLLYLCRYLLIIRRGVIIVSGAGAGGGRLRRFRLGAAMAIIYMAMSTLFFLRIAPLTPWWGALAAWVMILLIVEAIFAFFFPYRCCFNETEGDVQNNPHV
IRIS_313-11515 MELITACALLLIAVHIAIVISCPEPLAAVALVDVPLLYLCRYLLIIRRGVIIVSGGGGGGRLRRFRLGAAMAIIYMAMSTLFFLRIAPLTPWWGALAAWVMILLIVEAIFAFFFPYRCCFNETEGDVQNNPHV
IRIS_313-11516 MELITACALLLIAVHIAIVISCPEPLAAVALVDVPLLYLCRYLLIIRRGVIIVSGGGGGGRLRRFRLGAAMAIIYMAMSTLFFLRIAPLTPWWGALAAWVMILLIVEAIFAFFFPYRCCFNETEGDVQNNPHV
IRIS_313-11517 MELITACALLLIAVHIAIVISCPEPLAAVALVDVPLLYLCRYLLIIRRGVIIVSGAGAGGGRLRRFRLGAAMAIIYMAMSTLFFLRIAPLTPWWGALAAWVMILLIVEAIFAFFFPYRCCFNETEGDVQNNPHV
IRIS_313-11521 MELITACALLLIAVHIAIVISCPEPLAAVALVDVPLLYLCRYLLIIRRGVIIVSGGGGGGRLRRFRLGAAMAIIYMAMSTLFFLRIAPLTPWWGALAAWVMILLIVEAIFAFFFPYRCCFNETEGDVQNNPHV
IRIS_313-11522 MELITACALLLIAVHIAIVISCPEPLAAVALVDVPLLYLCRYLLIIRRGVIIVSGAGGGGGRLRRFRLGAAMAIIYMAMSTLFFLRIAPLTPWWGALAAWVMILLIVEAIFAFFFPYRCCFNETEGDVQNNPHV
IRIS_313-11523 MELITACALLLIAVHIAIVISCPEPLAAVALVDVPLLYLCRYLLIIRRGVIIVSGAGAGGGRLRRFRLGAAMAIIYMAMSTLFFLRIAPLTPWWGALAAWVMILLIVEAIFAFFFPYRCCFNETEGDVQNNPHV
IRIS_313-11524 MELITACALLLIAVHIAIVISCPEPLAAVALVDVPLLYLYRYLLIIRRGVIIVSGAGGGGGRLRRFRLGAAMAIIYMAMSTLFFLRIAPLTPWWGALAAWVMILLIVEAIFAFFFPYRCCFNETEGDVQNNPHV
IRIS_313-11525 MELITACALLLIAVHIAIVISCPEPLAAVALVDVPLLYLYRYLLIIRRGVIIVSGAGAGGGRLRRFRLGAAMAIIYMAMSTLFFLRIAPLTPWWGALAAWVMILLIVEAIFAFFFPYRCCFNETEGDVQNNPHV
IRIS_313-11526 MELITACALLLIAVHIAIVISCPEPLAAVALVDVPLLYLYRYLLIIRRGVIIVSGAGGGGGRLRRFRLGAAMAIIYMAMSTLFFLRIAPLTPWWGALAAWVMILLIVEAIFAFFFPYRCCFNETEGDVQNNPHV
IRIS_313-11527 MELITACALLLIAVHIAIVISCPEPLAAVALVDVPLLYLCRYLLIIRRGVIIVSGAGAGGGRLRRFRLGAAMAIIYMAMSTLFFLRIAPLTPWWGALAAWVMILLIVEAIFAFFFPYRCCFNETEGDVQNNPHV
IRIS_313-11528 MELITACALLLIAVHIAIVISCPEPLAAVALVDVPLLYLCRYLLIIRRGVIIVSGAGGAAAAGCAASASGQPWRSSTWPCRPSSSSALLLSRHGGAPSPPGS*SCSSSRPSSPSSSPTAAASTKPKAMSRIIPM
IRIS_313-11530 MELITACALLLIAVHIAIVISCPEPLAAVALVDVPLLYLCRYLLIIRRGVIIVSGGGGGGRLRRFRLGAAMAIIYMAMSTLFFLRIAPLTPWWGALAAWVMILLIVEAIFAFFFPYRCCFNETEGDVQNNPHV
IRIS_313-11532 MELITACALLLIAVHIAIVISCPEPLAAVALVDVPLLYLCRYLLIIRRGVIIVSGAGAGGGRLRRFRLGAAMAIIYMAMSTLFFLRIAPLTPWWGALAAWVMILLIVEAIFAFFFPYRCCFNETEGDVQNNPHV
IRIS_313-11534 MELITACALLLIAVHIAIVISCPEPLAAVALVDVPLLYLCRYLLIIRRGVIIVSGGGGGGRLRRFRLGAAMAIIYMAMSTLFFLRIAPLTPWWGALAAWVMILLIVEAIFAFFFPYRCCFNETEGDVQNNPHV
IRIS_313-11535 MELITACALLLIAVHIAIVISCPEPLAAVALVDVPLLYLCRYLLIIRRGVIIVSGGGGGGRLRRFRLGAAMAIIYMAMSTLFFLRIAPLTPWWGALAAWVMILLIVEAIFAFFFPYRCCFNETEGDVQNNPHV
IRIS_313-11536 MELITACALLLIAVHIAIVISCPEPLAAVALVDVPLLYLCRYLLIIRRGVIIVSGAGAGGGRLRRFRLGAAMAIIYMAMSTLFFLRIAPLTPWWGALAAWVMILLIVEAIFAFFFPYRCCFNETEGDVQNNPHV
IRIS_313-11537 MELITACALLLIAVHIAIVISCPEPLAAVALVDVPLLYLCRYLLIIRRGVIIVSGAGAGGGRLRRFRLGAAMAIIYMAMSTLFFLRIAPLTPWWGALAAWVMILLIVEAIFAFFFPYRCCFNETEGDVQNNPHV
IRIS_313-11538 MELITACALLLIAVHIAIVISCPEPLAAVALVDVPLLYLCRYLLIIRRGVIIVSGAGAGGGRLRRFRLGAAMAIIYMAMSTLFFLRIAPLTPWWGALAAWVMILLIVEAIFAFFFPYRCCFNETEGDVQNNPHV
IRIS_313-11539 MELITACALLLIAVHIAIVISCPEPLAAVALVDVPLLYLYRYLLIIRRGVIIVSGAGGGGGRLRRFRLGAAMAIIYMAMSTLFFLRIAPLTPWWGALAAWVMILLIVEAIFAFFFPYRCCFNETEGDVQNNPHV
IRIS_313-11540 MELITACALLLIAVHIAIVISCPEPLAAVALVDVPLLYLCRYLLIIRRGVIIVSGAGAGGGRLRRFRLGAAMAIIYMAMSTLFFLRIAPLTPWWGALAAWVMILLIVEAIFAFFFPYRCCFNETEGDVQNNPHV
IRIS_313-11541 MELITACALLLIAVHIAIVISCPEPLAAVALVDVPLLYLCRYLLIIRRGVIIVSGAGGAAAAGCAASASGQPWRSSTWPCRPSSSSALLLSRHGGAPSPPGS*SCSSSRPSSPSSSPTAAASTKPKAMSRIIPM
IRIS_313-11542 MELITACALLLIAVHIAIVISCPEPLAAVALVDVPLLYLCRYLLIIRRGVIIVSGAGAGGGRLRRFRLGAAMAIIYMAMSTLFFLRIAPLTPWWGALAAWVMILLIVEAIFAFFFPYRCCFNETEGDVQNNPHV
IRIS_313-11543 MELITACALLLIAVHIAIVISCPEPLAAVALVDVPLLYLYRYLLIIRRGVIIVSGAGGGGGRLRRFRLGAAMAIIYMAMSTLFFLRIAPLTPWWGALAAWVMILLIVEAIFAFFFPYRCCFNETEGDVQNNPHV
IRIS_313-11544 MELITACALLLIAVHIAIVISCPEPLAAVALVDVPLLYLCRYLLIIRRGVIIVSGGGGGGRLRRFRLGAAMAIIYMAMSTLFFLRIAPLTPWWGALAAWVMILLIVEAIFAFFFPYRCCFNETEGDVQNNPHV
IRIS_313-11545 MELITACALLLIAVHIAIVISCPEPLAAVALVDVPLLYLCRYLLIIRRGVIIVSGGGGGGRLRRFRLGAAMAIIYMAMSTLFFLRIAPLTPWWGALAAWVMILLIVEAIFAFFFPYRCCFNETEGDVQNNPHV
IRIS_313-11546 MELITACALLLIAVHIAIVISCPEPLAAVALVDVPLLYLYRYLLIIRRGVIIVSGAGGGGGRLRRFRLGAAMAIIYMAMSTLFFLRIAPLTPWWGALAAWVMILLIVEAIFAFFFPYRCCFNETEGDVQNNPHV
IRIS_313-11547 MELITACALLLIAVHIAIVISCPEPLAAVALVDVPLLYLCRYLLIIRRGVIIVSGGGGGGRLRRFRLGAAMAIIYMAMSTLFFLRIAPLTPWWGALAAWVMILLIVEAIFAFFFPYRCCFNETEGDVQNNPHV
IRIS_313-11548 MELITACALLLIAVHIAIVISCPEPLAAVALVDVPLLYLCRYLLIIRRGVIIVSGAGAGGGRLRRFRLGAAMAIIYMAMSTLFFLRIAPLTPWWGALAAWVMILLIVEAIFAFFFPYRCCFNETEGDVQNNPHV
IRIS_313-11549 MELITACALLLIAVHIAIVISCPEPLAAVALVDVPLLYLCRYLLIIRRGVIIVSGGGGGGRLRRFRLGAAMAIIYMAMSTLFFLRIAPLTPWWGALAAWVMILLIVEAIFAFFFPYRCCFNETEGDVQNNPHV
IRIS_313-11551 MELITACALLLIAVHIAIVISCPEPLAAVALVDVPLLYLCRYLLIIRRGVIIVSGAGGGGGRLRRFRLGAAMAIIYMAMSTLFFLRIAPLTPWWGALAAWVMILLIVEAIFAFFFPYRCCFNETEGDVQNNPHV
IRIS_313-11554 MELITACALLLIAVHIAIVISCPEPLAAVALVDVPLLYLCRYLLIIRRGVIIVSGAGAGGGRLRRFRLGAAMAIIYMAMSTLFFLRIAPLTPWWGALAAWVMILLIVEAIFAFFFPYRCCFNETEGDVQNNPHV
IRIS_313-11555 MELITACALLLIAVHIAIVISCPEPLAAVALVDVPLLYLCRYLLIIRRGVIIVSGAGGAAAAGCAASASGQPWRSSTWPCRPSSSSALLLSRHGGAPSPPGS*SCSSSRPSSPSSSPTAAASTKPKAMSRIIPM
IRIS_313-11556 MELITACALLLIAVHIAIVISCPEPLAAVALVDVPLLYLCRYLLIIRRGVIIVSGGGGGGRLRRFRLGAAMAIIYMAMSTLFFLRIAPLTPWWGALAAWVMILLIVEAIFAFFFPYRCCFNETEGDVQNNPHV
IRIS_313-11557 MELITACALLLIAVHIAIVISCPEPLAAVALVDVPLLYLCRYLLIIRRGVIIVSGAGGGGGRLRRFRLGAAMAIIYMAMSTLFFLRIAPLTPWWGALAAWVMILLIVEAIFAFFFPYRCCFNETEGDVQNNPHV
IRIS_313-11558 MELITACALLLIAVHIAIVISCPEPLAAVALVDVPLLYLCRYLLIIRRGVIIVSGAGAGGGRLRRFRLGAAMAIIYMAMSTLFFLRIAPLTPWWGALAAWVMILLIVEAIFAFFFPYRCCFNETEGDVQNNPHV
IRIS_313-11561 MELITACALLLIAVHIAIVISCPEPLAAVALVDVPLLYLCRYLLIIRRGVIIVSGAGGGGGRLRRFRLGAAMAIIYMAMSTLFFLRIAPLTPWWGALAAWVMILLIVEAIFAFFFPYRCCFNETEGDVQNNPHV
IRIS_313-11563 MELITACALLLIAVHIAIVISCPEPLAAVALVDVPLLYLCRYLLIIRRGVIIVSGAGAGGGRLRRFRLGAAMAIIYMAMSTLFFLRIAPLTPWWGALAAWVMILLIVEAIFAFFFPYRCCFNETEGDVQNNPHV
IRIS_313-11564 MELITACALLLIAVHIAIVISCPEPLAAVALVDVPLLYLCRYLLIIRRGVIIVSGAGAGGGRLRRFRLGAAMAIIYMAMSTLFFLRIAPLTPWWGALAAWVMILLIVEAIFAFFFPYRCCFNETEGDVQNNPHV
IRIS_313-11565 MELITACALLLIAVHIAIVISCPEPLAAVALVDVPLLYLCRYLLIIRRGVIIVSGGGGGGRLRRFRLGAAMAIIYMAMSTLFFLRIAPLTPWWGALAAWVMILLIVEAIFAFFFPYRCCFNETEGDVQNNPHV
IRIS_313-11566 MELITACALLLIAVHIAIVISCPEPLAAVALVDVPLLYLCRYLLIIRRGVIIVSGAGGGGGRLRRFRLGAAMAIIYMAMSTLFFLRIAPLTPWWGALAAWVMILLIVEAIFAFFFPYRCCFNETEGDVQNNPHV
IRIS_313-11567 MELITACALLLIAVHIAIVISCPEPLAAVALVDVPLLYLCRYLLIIRRGVIIVSGAGAGGGRLRRFRLGAAMAIIYMAMSTLFFLRIAPLTPWWGALAAWVMILLIVEAIFAFFFPYRCCFNETEGDVQNNPHV
IRIS_313-11568 MELITACALLLIAVHIAIVISCPEPLAAVALVDVPLLYLCRYLLIIRRGVIIVSGGGGGGRLRRFRLGAAMAIIYMAMSTLFFLRIAPLTPWWGALAAWVMILLIVEAIFAFFFPYRCCFNETEGDVQNNPHV
IRIS_313-11570 MELITACALLLIAVHIAIVISCPEPLAAVALVDVPLLYLCRYLLIIRRGVIIVSGAGGGGGRLRRFRLGAAMAIIYMAMSTLFFLRIAPLTPWWGALAAWVMILLIVEAIFAFFFPYRCCFNETEGDVQNNPHV
IRIS_313-11571 MELITACALLLIAVHIAIVISCPEPLAAVALVDVPLLYLCRYLLIIRRGVIIVSGAGAGGGRLRRFRLGAAMAIIYMAMSTLFFLRIAPLTPWWGALAAWVMILLIVEAIFAFFFPYRCCFNETEGDVQNNPHV
IRIS_313-11572 MELITACALLLIAVHIAIVISCPEPLAAVALVDVPLLYLCRYLLIIRRGVIIVSGAGGAAAAGCAASASGQPWRSSTWPCRPSSSSALLLSRHGGAPSPPGS*SCSSSRPSSPSSSPTAAASTKPKAMSRIIPM
IRIS_313-11573 MELITACALLLIAVHIAIVISCPEPLAAVALVDVPLLYLCRYLLIIRRGVIIVSGAGGGGGGRLRRFRLGAAMAIIYMAMSTLFFLRIAPLTPWWGALAAWVMILLIVEAIFAFFFPYRCCFNETEGDVQNNPHV
IRIS_313-11574 MELITACALLLIAVHIAIVISCPEPLAAVALVDVPLLYLCRYLLIIRRGVIIVSGAGAGGGRLRRFRLGAAMAIIYMAMSTLFFLRIAPLTPWWGALAAWVMILLIVEAIFAFFFPYRCCFNETEGDVQNNPHV
IRIS_313-11575 MELITACALLLIAVHIAIVISCPEPLAAVALVDVPLLYLCRYLLIIRRGVIIVSGAGAGGGRLRRFRLGAAMAIIYMAMSTLFFLRIAPLTPWWGALAAWVMILLIVEAIFAFFFPYRCCFNETEGDVQNNPHV
IRIS_313-11576 MELITACALLLIAVHIAIVISCPEPLAAVALVDVPLLYLCRYLLIIRRGVIIVSGGGGGGRLRRFRLGAAMAIIYMAMSTLFFLRIAPLTPWWGALAAWVMILLIVEAIFAFFFPYRCCFNETEGDVQNNPHV
IRIS_313-11577 MELITACALLLIAVHIAIVISCPEPLAAVALVDVPLLYLCRYLLIIRRGVIIVSGAGGAAAAGCAASASGQPWRSSTWPCRPSSSSALLLSRHGGAPSPPGS*SCSSSRPSSPSSSPTAAASTKPKAMSRIIPM
IRIS_313-11578 MELITACALLLIAVHIAIVISCPEPLAAVALVDVPLLYLCRYLLIIRRGVIIVSGAGAGGGRLRRFRLGAAMAIIYMAMSTLFFLRIAPLTPWWGALAAWVMILLIVEAIFAFFFPYRCCFNETEGDVQNNPHV
IRIS_313-11579 MELITACALLLIAVHIAIVISCPEPLAAVALVDVPLLYLCRYLLIIRRGVIIVSGGGGGGRLRRFRLGAAMAIIYMAMSTLFFLRIAPLTPWWGALAAWVMILLIVEAIFAFFFPYRCCFNETEGDVQNNPHV
IRIS_313-11580 MELITACALLLIAVHIAIVISCPEPLAAVALVDVPLLYLCRYLLIIRRGVIIVSGGGGGGRLRRFRLGAAMAIIYMAMSTLFFLRIAPLTPWWGALAAWVMILLIVEAIFAFFFPYRCCFNETEGDVQNNPHV
IRIS_313-11582 MELITACALLLIAVHIAIVISCPEPLAAVALVDVPLLYLCRYLLIIRRGVIIVSGAGAGGGRLRRFRLGAAMAIIYMAMSTLFFLRIAPLTPWWGALAAWVMILLIVEAIFAFFFPYRCCFNETEGDVQNNPHV
IRIS_313-11584 MELITACALLLIAVHIAIVISCPEPLAAVALVDVPLLYLCRYLLIIRRGVIIVSGAGAGGGRLRRFRLGAAMAIIYMAMSTLFFLRIAPLTPWWGALAAWVMILLIVEAIFAFFFPYRCCFNETEGDVQNNPHV
IRIS_313-11585 MELITACALLLIAVHIAIVISCPEPLAAVALVDVPLLYLCRYLLIIRRGVIIVSGAGAGGGRLRRFRLGAAMAIIYMAMSTLFFLRIAPLTPWWGALAAWVMILLIVEAIFAFFFPYRCCFNETEGDVQNNPHV
IRIS_313-11586 MELITACALLLIAVHIAIVISCPEPLAAVALVDVPLLYLCRYLLIIRRGVIIVSGAGAGGGRLRRFRLGAAMAIIYMAMSTLFFLRIAPLTPWWGALAAWVMILLIVEAIFAFFFPYRCCFNETEGDVQNNPHV
IRIS_313-11588 MELITACALLLIAVHIAIVISCPEPLAAVALVDVPLLYLCRYLLIIRRGVIIVSGAGGAAAAGCAASASGQPWRSSTWPCRPSSSSALLLSRHGGAPSPPGS*SCSSSRPSSPSSSPTAAASTKPKAMSRIIPM
IRIS_313-11591 MELITACALLLIAVHIAIVISCPEPLAAVALVDVPLLYLCRYLLIIRRGVIIVSGAGAGGGRLRRFRLGAAMAIIYMAMSTLFFLRIAPLTPWWGALAAWVMILLIVEAIFAFFFPYRCCFNETEGDVQNNPHV
IRIS_313-11594 MELITACALLLIAVHIAIVISCPEPLAAVALVDVPLLYLYRYLLIIRRGVIIVSGAGAGGGRLRRFRLGAAMAIIYMAMSTLFFLRIAPLTPWWGALAAWVMILLIVEAIFAFFFPYRCCFNETEGDVQNNPHV
IRIS_313-11595 MELITACALLLIAVHIAIVISCPEPLAAVALVDVPLLYLCRYLLIIRRGVIIVSGAGAGGGRLRRFRLGAAMAIIYMAMSTLFFLRIAPLTPWWGALAAWVMILLIVEAIFAFFFPYRCCFNETEGDVQNNPHV
IRIS_313-11596 MELITACALLLIAVHIAIVISCPEPLAAVALVDVPLLYLCRYLLIIRRGVIIVSGAGAGGGRLRRFRLGAAMAIIYMAMSTLFFLRIAPLTPWWGALAAWVMILLIVEAIFAFFFPYRCCFNETEGDVQNNPHV
IRIS_313-11597 MELITACALLLIAVHIAIVISCPEPLAAVALVDVPLLYLCRYLLIIRRGVIIVSGGGGGGRLRRFRLGAAMAIIYMAMSTLFFLRIAPLTPWWGALAAWVMILLIVEAIFAFFFPYRCCFNETEGDVQNNPHV
IRIS_313-11598 MELITACALLLIAVHIAIVISCPEPLAAVALVDVPLLYLCRYLLIIRRGVIIVSGAGAGGGRLRRFRLGAAMAIIYMAMSTLFFLRIAPLTPWWGALAAWVMILLIVEAIFAFFFPYRCCFNETEGDVQNNPHV
IRIS_313-11599 MELITACALLLIAVHIAIVISCPEPLAAVALVDVPLLYLCRYLLIIRRGVIIVSGGGGGGRLRRFRLGAAMAIIYMAMSTLFFLRIAPLTPWWGALAAWVMILLIVEAIFAFFFPYRCCFNETEGDVQNNPHV
IRIS_313-11600 MELITACALLLIAVHIAIVISCPEPLAAVALVDVPLLYLCRYLLIIRRGVIIVSGGGGGGRLRRFRLGAAMAIIYMAMSTLFFLRIAPLTPWWGALAAWVMILLIVEAIFAFFFPYRCCFNETEGDVQNNPHV
IRIS_313-11602 MELITACALLLIAVHIAIVISCPEPLAAVAFVNVPLLYLCRYLLIIRRGVIIVSGGGAGGGRLRRFRLGAAMAVIYMAMSTLFFLRIAPLTPWWGALAAWVMILLIVEAIFAFFFPYRCCFNETEGDVQNNPHV
IRIS_313-11603 MELITACALLLIAVHIAIVISCPEPLAAVALVDVPLLYLCRYLLIIRRGVIIVSGAGAGGGRLRRFRLGAAMAIIYMAMSTLFFLRIAPLTPWWGALAAWVMILLIVEAIFAFFFPYRCCFNETEGDVQNNPHV
IRIS_313-11604 MELITACALLLIAVHIAIVISCPEPLAAVALVDVPLLYLCRYLLIIRRGVIIVSGAGAGGGRLRRFRLGAAMAIIYMAMSTLFFLRIAPLTPWWGALAAWVMILLIVEAIFAFFFPYRCCFNETEGDVQNNPHV
IRIS_313-11606 MELITACALLLIAVHIAIVISCPEPLAAVALVDVPLLYLCRYLLIIRRGVIIVSGGGGGGRLRRFRLGAAMAIIYMAMSTLFFLRIAPLTPWWGALAAWVMILLIVEAIFAFFFPYRCCFNETEGDVQNNPHV
IRIS_313-11607 MELITACALLLIAVHIAIVISCPEPLAAVALVDVPLLYLCRYLLIIRRGVIIVSGAGAGGGRLRRFRLGAAMAIIYMAMSTLFFLRIAPLTPWWGALAAWVMILLIVEAIFAFFFPYRCCFNETEGDVQNNPHV
IRIS_313-11608 MELITACALLLIAVHIAIVISCPEPLAAVALVDVPLLYLCRYLLIIRRGVIIVSGAGAGGGRLRRFRLGAAMAIIYMAMSTLFFLRIAPLTPWWGALAAWVMILLIVEAIFAFFFPYRCCFNETEGDVQNNPHV
IRIS_313-11609 MELITACALLLIAVHIAIVISCPEPLAAVALVDVPLLYLCRYLLIIRRGVIIVSGGGGGGRLRRFRLGAAMAIIYMAMSTLFFLRIAPLTPWWGALAAWVMILLIVEAIFAFFFPYRCCFNETEGDVQNNPHV
IRIS_313-11610 MELITACALLLIAVHIAIVISCPEPLAAVALVDVPLLYLCRYLLIIRRGVIIVSGGGGGGRLRRFRLGAAMAIIYMAMSTLFFLRIAPLTPWWGALAAWVMILLIVEAIFAFFFPYRCCFNETEGDVQNNPHV
IRIS_313-11611 MELITACALLLIAVHIAIVISCPEPLAAVALVDVPLLYLCRYLLIIRRGVIIVSGAGAGGGRLRRFRLGAAMAIIYMAMSTLFFLRIAPLTPWWGALAAWVMILLIVEAIFAFFFPYRCCFNETEGDVQNNPHV
IRIS_313-11615 MELITACALLLIAVHIAIVISCPEPLAAVALVDVPLLYLCRYLLIIRRGVIIVSGAGAGGGRLRRFRLGAAMAIIYMAMSTLFFLRIAPLTPWWGALAAWVMILLIVEAIFAFFFPYRCCFNETEGDVQNNPHV
IRIS_313-11616 MELITACALLLIAVHIAIVISCPEPLAAVALVDVPLLYLCRYLLIIRRGVIIVSGAGAGGGRLRRFRLGAAMAIIYMAMSTLFFLRIAPLTPWWGALAAWVMILLIVEAIFAFFFPYRCCFNETEGDVQNNPHV
IRIS_313-11617 MELITACALLLIAVHIAIVISCPEPLAAVALVDVPLLYLCRYLLIIRRGVIIVSGAGAGGGRLRRFRLGAAMAIIYMAMSTLFFLRIAPLTPWWGALAAWVMILLIVEAIFAFFFPYRCCFNETEGDVQNNPHV
IRIS_313-11618 MELITACALLLIAVHIAIVISCPEPLAAVAFVNVPLLYLCRYLLIIRRGVIIVSGGGGGGGRLRRFRLGAAMAVIYMAMSTLFFLRIAPLTPWWGALAAWVMILLIVEAIFAFFFPYRCCFNETEGDVQNNPHV
IRIS_313-11619 MELITACALLLIAVHIAIVISCPEPLAAVAFVNVPLLYLCRYLLIIRRGVIIVSGGGGGGGRLRRFRLGAAMAVIYMAMSTLFFLRIAPLTPWWGALAAWVMILLIVEAIFAFFFPYRCCFNETEGDVQNNPHV
IRIS_313-11620 MELITACALLLIAVHIAIVISCPEPLAAVALVDVPLLYLCRYLLIIRRGVIIVSGAGAGGGRLRRFRLGAAMAIIYMAMSTLFFLRIAPLTPWWGALAAWVMILLIVEAIFAFFFPYRCCFNETEGDVQNNPHV
IRIS_313-11621 MELITACALLLIAVHIAIVISCPEPLAAVALVDVPLLYLCRYLLIIRRGVIIVSGGGGGGRLRRFRLGAAMAIIYMAMSTLFFLRIAPLTPWWGALAAWVMILLIVEAIFAFFFPYRCCFNETEGDVQNNPHV
IRIS_313-11622 MELITACALLLIAVHIAIVISCPEPLAAVALVDVPLLYLYRYLLIIRRGVIIVSGAGGGGGRLRRFRLGAAMAIIYMAMSTLFFLRIAPLTPWWGALAAWVMILLIVEAIFAFFFPYRCCFNETEGDVQNNPHV
IRIS_313-11623 MELITACALLLIAVHIAIVISCPEPLAAVALVDVPLLYLCRYLLIIRRGVIIVSGAGGGGGRLRRFRLGAAMAIIYMAMSTLFFLRIAPLTPWWGALAAWVMILLIVEAIFAFFFPYRCCFNETEGDVQNNPHV
IRIS_313-11624 MELITACALLLIAVHIAIVISCPEPLAAVALVDVPLLYLCRYLLIIRRGVIIVSGGGGGGRLRRFRLGAAMAIIYMAMSTLFFLRIAPLTPWWGALAAWVMILLIVEAIFAFFFPYRCCFNETEGDVQNNPHV
IRIS_313-11625 MELITACALLLIAVHIAIVISCPEPLAAVALVDVPLLYLCRYLLIIRRGVIIVSGAGAGGGRLRRFRLGAAMAIIYMAMSTLFFLRIAPLTPWWGALAAWVMILLIVEAIFAFFFPYRCCFNETEGDVQNNPHV
IRIS_313-11626 MELITACALLLIAVHIAIVISCPEPLAAVALVDVPLLYLCRYLLIIRRGVIIVSGAGAGGGRLRRFRLGAAMAIIYMAMSTLFFLRIAPLTPWWGALAAWVMILLIVEAIFAFFFPYRCCFNETEGDVQNNPHV
IRIS_313-11627 MELITACALLLIAVHIAIVISCPEPLAAVALVDVPLLYLCRYLLIIRRGVIIVSGAGAGGGRLRRFRLGAAMAIIYMAMSTLFFLRIAPLTPWWGALAAWVMILLIVEAIFAFFFPYRCCFNETEGDVQNNPHV

IRIS_313-11628 MELITACALLLIAVHIAIVISCPEPLAAVALVDVPLLYLCRYLLIIRRGVIIVSGAGAGGGRLRRFRLGAAMAIIYMAMSTLFFLRIAPLTPWWGALAAWVMILLIVEAIFAFFFPYRCCFNETEGDVQNNPHV
IRIS_313-11629 MELITACALLLIAVHIAIVISCPEPLAAVALVDVPLLYLCRYLLIIRRGVIIVSGAGAGGGRLRRFRLGAAMAIIYMAMSTLFFLRIAPLTPWWGALAAWVMILLIVEAIFAFFFPYRCCFNETEGDVQNNPHV
IRIS_313-11630 MELITACALLLIAVHIAIVISCPEPLAAVALVDVPLLYLCRYLLIIRRGVIIVSGAGAGGGRLRRFRLGAAMAIIYMAMSTLFFLRIAPLTPWWGALAAWVMILLIVEAIFAFFFPYRCCFNETEGDVQNNPHV
IRIS_313-11632 MELITACALLLIAVHIAIVISCPEPLAAVALVDVPLLYLCRYLLIIRRGVIIVSGAGAGGGRLRRFRLGAAMAIIYMAMSTLFFLRIAPLTPWWGALAAWVMILLIVEAIFAFFFPYRCCFNETEGDVQNNPHV
IRIS_313-11635 MELITACALLLIAVHIAIVISCPEPLAAVALVDVPLLYLCRYLLIIRRGVIIVSGAGGAAAAGCAASASGQPWRSSTWPCRPSSSSALLLSRHGGAPSPPGS*SCSSSRPSSPSSSPTAAASTKPKAMSRIIPM
IRIS_313-11636 MELITACALLLIAVHIAIVISCPEPLAAVAFVNVPLLYLCRYLLIIRRGVIIVSGGGGGGGRLRRFRLGAAMAVIYMAMSTLFFLRIAPLTPWWGALAAWVMILLIVEAIFAFFFPYRCCFNETEGDVQNNPHV
IRIS_313-11638 MELITACALLLIAVHIAIVISCPEPLAAVALVDVPLLYLCRYLLIIRRGVIIVSGAGAGGGRLRRFRLGAAMAIIYMAMSTLFFLRIAPLTPWWGALAAWVMILLIVEAIFAFFFPYRCCFNETEGDVQNNPHV
IRIS_313-11639 MELITACALLLIAVHIAIVISCPEPLAAVALVDVPLLYLCRYLLIIRRGVIIVSGGGGGGRLRRFRLGAAMAIIYMAMSTLFFLRIAPLTPWWGALAAWVMILLIVEAIFAFFFPYRCCFNETEGDVQNNPHV
IRIS_313-11640 MELITACALLLIAVHIAIVISCPEPLAAVALVDVPLLYLCRYLLIIRRGVIIVSGGGGGGRLRRFRLGAAMAIIYMAMSTLFFLRIAPLTPWWGALAAWVMILLIVEAIFAFFFPYRCCFNETEGDVQNNPHV
IRIS_313-11641 MELITACALLLIAVHIAIVISCPEPLAAVALVDVPLLYLCRYLLIIRRGVIIVSGGGGGGRLRRFRLGAAMAIIYMAMSTLFFLRIAPLTPWWGALAAWVMILLIVEAIFAFFFPYRCCFNETEGDVQNNPHV
IRIS_313-11642 MELITACALLLIAVHIAIVISCPEPLAAVALVDVPLLYLCRYLLIIRRGVIIVSGAGAGGGRLRRFRLGAAMAIIYMAMSTLFFLRIAPLTPWWGALAAWVMILLIVEAIFAFFFPYRCCFNETEGDVQNNPHV
IRIS_313-11643 MELITACALLLIAVHIAIVISCPEPLAAVALVDVPLLYLCRYLLIIRRGVIIVSGAGAGGGRLRRFRLGAAMAIIYMAMSTLFFLRIAPLTPWWGALAAWVMILLIVEAIFAFFFPYRCCFNETEGDVQNNPHV
IRIS_313-11644 MELITACALLLIAVHIAIVISCPEPLAAVALVDVPLLYLCRYLLIIRRGVIIVSGAGAGGGRLRRFRLGAAMAIIYMAMSTLFFLRIAPLTPWWGALAAWVMILLIVEAIFAFFFPYRCCFNETEGDVQNNPHV
IRIS_313-11645 MELITACALLLIAVHIAIVISCPEPLAAVALVDVPLLYLCRYLLIIRRGVIIVSGGGGGGRLRRFRLGAAMAIIYMAMSTLFFLRIAPLTPWWGALAAWVMILLIVEAIFAFFFPYRCCFNETEGDVQNNPHV
IRIS_313-11646 MELITACALLLIAVHIAIVISCPEPLAAVALVDVPLLYLCRYLLIIRRGVIIVSGAGAGGGRLRRFRLGAAMAIIYMAMSTLFFLRIAPLTPWWGALAAWVMILLIVEAIFAFFFPYRCCFNETEGDVQNNPHV
IRIS_313-11647 MELITACALLLIAVHIAIVISCPEPLAAVALVDVPLLYLCRYLLIIRRGVIIVSGGGGGGRLRRFRLGAAMAIIYMAMSTLFFLRIAPLTPWWGALAAWVMILLIVEAIFAFFFPYRCCFNETEGDVQNNPHV
IRIS_313-11648 MELITACALLLIAVHIAIVISCPEPLAAVALVDVPLLYLCRYLLIIRRGVIIVSGAGAGGGRLRRFRLGAAMAIIYMAMSTLFFLRIAPLTPWWGALAAWVMILLIVEAIFAFFFPYRCCFNETEGDVQNNPHV
IRIS_313-11650 MELITACALLLIAVHIAIVISCPEPLAAVALVDVPLLYLCRYLLIIRRGVIIVSGAGAGGGRLRRFRLGAAMAIIYMAMSTLFFLRIAPLTPWWGALAAWVMILLIVEAIFAFFFPYRCCFNETEGDVQNNPHV
IRIS_313-11651 MELITACALLLIAVHIAIVISCPEPLAAVALVDVPLLYLCRYLLIIRRGVIIVSGAGGGGGRLRRFRLGAAMAIIYMAMSTLFFLRIAPLTPWWGALAAWVMILLIVEAIFAFFFPYRCCFNETEGDVQNNPHV
IRIS_313-11652 MELITACALLLIAVHIAIVISCPEPLAAVALVDVPLLYLCRYLLIIRRGVIIVSGAGAGGGRLRRFRLGAAMAIIYMAMSTLFFLRIAPLTPWWGALAAWVMILLIVEAIFAFFFPYRCCFNETEGDVQNNPHV
IRIS_313-11653 MELITACALLLIAVHIAIVISCPEPLAAVALVDVPLLYLCRYLLIIRRGVIIVSGAGGGGGRLRRFRLGAAMAIIYMAMSTLFFLRIAPLTPWWGALAAWVMILLIVEAIFAFFFPYRCCFNETEGDVQNNPHV
IRIS_313-11654 MELITACALLLIAVHIAIVISCPEPLAAVALVDVPLLYLCRYLLIIRRGVIIVSGAGAGGGRLRRFRLGAAMAIIYMAMSTLFFLRIAPLTPWWGALAAWVMILLIVEAIFAFFFPYRCCFNETEGDVQNNPHV
IRIS_313-11655 MELITACALLLIAVHIAIVISCPEPLAAVALVDVPLLYLCRYLLIIRRGVIIVSGAGAGGGRLRRFRLGAAMAIIYMAMSTLFFLRIAPLTPWWGALAAWVMILLIVEAIFAFFFPYRCCFNETEGDVQNNPHV
IRIS_313-11656 MELITACALLLIAVHIAIVISCPEPLAAVALVDVPLLYLCRYLLIIRRGVIIVSGGGGGGRLRRFRLGAAMAIIYMAMSTLFFLRIAPLTPWWGALAAWVMILLIVEAIFAFFFPYRCCFNETEGDVQNNPHV
IRIS_313-11657 MELITACALLLIAVHIAIVISCPEPLAAVALVDVPLLYLCRYLLIIRRGVIIVSGAGAGGGRLRRFRLGAAMAIIYMAMSTLFFLRIAPLTPWWGALAAWVMILLIVEAIFAFFFPYRCCFNETEGDVQNNPHV
IRIS_313-11658 MELITACALLLIAVHIAIVISCPEPLAAVALVDVPLLYLCRYLLIIRRGVIIVSGAGAGGGRLRRFRLGAAMAIIYMAMSTLFFLRIAPLTPWWGALAAWVMILLIVEAIFAFFFPYRCCFNETEGDVQNNPHV
IRIS_313-11659 MELITACALLLIAVHIAIVISCPEPLAAVALVDVPLLYLYRYLLIIRRGVIIVSGAGGGGGRLRRFRLGAAMAIIYMAMSTLFFLRIAPLTPWWGALAAWVMILLIVEAIFAFFFPYRCCFNETEGDVQNNPHV
IRIS_313-11660 MELITACALLLIAVHIAIVISCPEPLAAVALVDVPLLYLYRYLLIIRRGVIIVSGAGGGGGRLRRFRLGAAMAIIYMAMSTLFFLRIAPLTPWWGALAAWVMILLIVEAIFAFFFPYRCCFNETEGDVQNNPHV
IRIS_313-11661 MELITACALLLIAVHIAIVISCPEPLAAVALVDVPLLYLCRYLLIIRRGVIIVSGAGAGGGRLRRFRLGAAMAIIYMAMSTLFFLRIAPLTPWWGALAAWVMILLIVEAIFAFFFPYRCCFNETEGDVQNNPHV
IRIS_313-11663 MELITACALLLIAVHIAIVISCPEPLAAVALVDVPLLYLCRYLLIIRRGVIIVSGAGGAAAAGCAASASGQPWRSSTWPCRPSSSSALLLSRHGGAPSPPGS*SCSSSRPSSPSSSPTAAASTKPKAMSRIIPM
IRIS_313-11664 MELITACALLLIAVHIAIVISCPEPLAAVALVDVPLLYLYRYLLIIRRGVIIVSGAGGGGGRLRRFRLGAAMAIIYMAMSTLFFLRIAPLTPWWGALAAWVMILLIVEAIFAFFFPYRCCFNETEGDVQNNPHV
IRIS_313-11665 MELITACALLLIAVHIAIVISCPEPLAAVALVDVPLLYLCRYLLIIRRGVIIVSGAGAGGGRLRRFRLGAAMAIIYMAMSTLFFLRIAPLTPWWGALAAWVMILLIVEAIFAFFFPYRCCFNETEGDVQNNPHV
IRIS_313-11666 MELITACALLLIAVHIAIVISCPEPLAAVALVDVPLLYLCRYLLIIRRGVIIVSGAGAGGGRLRRFRLGAAMAIIYMAMSTLFFLRIAPLTPWWGALAAWVMILLIVEAIFAFFFPYRCCFNETEGDVQNNPHV
IRIS_313-11667 MELITACALLLIAVHIAIVISCPEPLAAVALVDVPLLYLYRYLLIIRRGVIIVSGAGGGGGRLRRFRLGAAMAIIYMAMSTLFFLRIAPLTPWWGALAAWVMILLIVEAIFAFFFPYRCCFNETEGDVQNNPHV
IRIS_313-11668 MELITACALLLIAVHIAIVISCPEPLAAVALVDVPLLYLCRYLLIIRRGVIIVSGAGAGGGRLRRFRLGAAMAIIYMAMSTLFFLRIAPLTPWWGALAAWVMILLIVEAIFAFFFPYRCCFNETEGDVQNNPHV
IRIS_313-11669 MELITACALLLIAVHIAIVISCPEPLAAVALVDVPLLYLYRYLLIIRRGVIIVSGAGGGGGRLRRFRLGAAMAIIYMAMSTLFFLRIAPLTPWWGALAAWVMILLIVEAIFAFFFPYRCCFNETEGDVQNNPHV
IRIS_313-11671 MELITACALLLIAVHIAIVISCPEPLAAVALVDVPLLYLCRYLLIIRRGVIIVSGAGAGGGRLRRFRLGAAMAIIYMAMSTLFFLRIAPLTPWWGALAAWVMILLIVEAIFAFFFPYRCCFNETEGDVQNNPHV
IRIS_313-11672 MELITACALLLIAVHIAIVISCPEPLAAVALVDVPLLYLCRYLLIIRRGVIIVSGAGAGGGRLRRFRLGAAMAIIYMAMSTLFFLRIAPLTPWWGALAAWVMILLIVEAIFAFFFPYRCCFNETEGDVQNNPHV
IRIS_313-11673 MELITACALLLIAVHIAIVISCPEPLAAVALVDVPLLYLCRYLLIIRRGVIIVSGAGGGGGRLRRFRLGAAMAIIYMAMSTLFFLRIAPLTPWWGALAAWVMILLIVEAIFAFFFPYRCCFNETEGDVQNNPHV
IRIS_313-11674 MELITACALLLIAVHIAIVISCPEPLAAVALVDVPLLYLYRYLLIIRRGVIIVSGAGGGGGRLRRFRLGAAMAIIYMAMSTLFFLRIAPLTPWWGALAAWVMILLIVEAIFAFFFPYRCCFNETEGDVQNNPHV
IRIS_313-11676 MELITACALLLIAVHIAIVISCPEPLAAVALVDVPLLYLCRYLLIIRRGVIIVSGGGGGGRLRRFRLGAAMAIIYMAMSTLFFLRIAPLTPWWGALAAWVMILLIVEAIFAFFFPYRCCFNETEGDVQNNPHV
IRIS_313-11677 MELITACALLLIAVHIAIVISCPEPLAAVALVDVPLLYLYRYLLIIRRGVIIVSGAGGGGGRLRRFRLGAAMAIIYMAMSTLFFLRIAPLTPWWGALAAWVMILLIVEAIFAFFFPYRCCFNETEGDVQNNPHV
IRIS_313-11678 MELITACALLLIAVHIAIVISCPEPLAAVALVDVPLLYLCRYLLIIRRGVIIVSGAGGGGRLRRFRLGAAMAIIYMAMSTLFFLRIAPLTPWWGALAAWVMILLIVEAIFAFFFPYRCCFNETEGDVQNNPHV
IRIS_313-11679 MELITACALLLIAVHIAIVISCPEPLAAVALVDVPLLYLCRYLLIIRRGVIIVSGGGGGGRLRRFRLGAAMAIIYMAMSTLFFLRIAPLTPWWGALAAWVMILLIVEAIFAFFFPYRCCFNETEGDVQNNPHV
IRIS_313-11680 MELITACALLLIAVHIAIVISCPEPLAAVALVDVPLLYLYRYLLIIRRGVIIVSGAGGGGGRLRRFRLGAAMAIIYMAMSTLFFLRIAPLTPWWGALAAWVMILLIVEAIFAFFFPYRCCFNETEGDVQNNPHV
IRIS_313-11681 MELITACALLLIAVHIAIVISCPEPLAAVALVDVPLLYLCRYLLIIRRGVIIVSGAGAGGGRLRRFRLGAAMAIIYMAMSTLFFLRIAPLTPWWGALAAWVMILLIVEAIFAFFFPYRCCFNETEGDVQNNPHV
IRIS_313-11682 MELITACALLLIAVHIAIVISCPEPLAAVALVDVPLLYLCRYLLIIRRGVIIVSGGGGGGRLRRFRLGAAMAIIYMAMSTLFFLRIAPLTPWWGALAAWVMILLIVEAIFAFFFPYRCCFNETEGDVQNNPHV
IRIS_313-11683 MELITACALLLIAVHIAIVISCPEPLAAVALVDVPLLYLCRYLLIIRRGVIIVSGGGGGGRLRRFRLGAAMAIIYMAMSTLFFLRIAPLTPWWGALAAWVMILLIVEAIFAFFFPYRCCFNETEGDVQNNPHV
IRIS_313-11684 MELITACALLLIAVHIAIVISCPEPLAAVALVDVPLLYLCRYLLIIRRGVIIVSGGGGGGRLRRFRLGAAMAIIYMAMSTLFFLRIAPLTPWWGALAAWVMILLIVEAIFAFFFPYRCCFNETEGDVQNNPHV
IRIS_313-11685 MELITACALLLIAVHIAIVISCPEPLAAVALVDVPLLYLYRYLLIIRRGVIIVSGAGGGGGRLRRFRLGAAMAIIYMAMSTLFFLRIAPLTPWWGALAAWVMILLIVEAIFAFFFPYRCCFNETEGDVQNNPHV
IRIS_313-11686 MELITACALLLIAVHIAIVISCPEPLAAVALVDVPLLYLYRYLLIIRRGVIIVSGAGGGGGRLRRFRLGAAMAIIYMAMSTLFFLRIAPLTPWWGALAAWVMILLIVEAIFAFFFPYRCCFNETEGDVQNNPHV
IRIS_313-11687 MELITACALLLIAVHIAIVISCPEPLAAVALVDVPLLYLCRYLLIIRRGVIIVSGAGAGGGRLRRFRLGAAMAIIYMVMSTLFFLRIAPLTPWWGALAAWVMILLIVEAIFAFFFPYRCCFNETEGDVQNNPHV
IRIS_313-11688 MELITACALLLIAVHIAIVISCPEPLAAVALVDVPLLYLYRYLLIIRRGVIIVSGAGGGGGRLRRFRLGAAMAIIYMAMSTLFFLRIAPLTPWWGALAAWVMILLIVEAIFAFFFPYRCCFNETEGDVQNNPHV
IRIS_313-11689 MELITACALLLIAVHIAIVISCPEPLAAVALVDVPLLYLCRYLLIIRRGVIIVSGAGAGGGRLRRFRLGAAMAIIYMAMSTLFFLRIAPLTPWWGALAAWVMILLIVEAIFAFFFPYRCCFNETEGDVQNNPHV
IRIS_313-11690 MELITACALLLIAVHIAIVISCPEPLAAVALVDVPLLYLCRYLLIIRRGVIIVSGAGGGGGRLRRFRLGAAMAIIYMAMSTLFFLRIAPLTPWWGALAAWVMILLIVEAIFAFFFPYRCCFNETEGDVQNNPHV
IRIS_313-11691 MELITACALLLIAVHIAIVISCPEPLAAVALVDVPLLYLCRYLLIIRRGVIIVSGAGAGGGRLRRFRLGAAMAIIYMAMSTLFFLRIAPLTPWWGALAAWVMILLIVEAIFAFFFPYRCCFNETEGDVQNNPHV
IRIS_313-11692 MELITACALLLIAVHIAIVISCPEPLAAVALVDVPLLYLCRYLLIIRRGVIIVSGAGAGGGRLRRFRLGAAMAIIYMAMSTLFFLRIAPLTPWWGALAAWVMILLIVEAIFAFFFPYRCCFNETEGDVQNNPHV
IRIS_313-11693 MELITACALLLIAVHIAIVISCPEPLAAVALVDVPLLYLCRYLLIIRRGVIIVSGGGGGGRLRRFRLGAAMAIIYMAMSTLFFLRIAPLTPWWGALAAWVMILLIVEAIFAFFFPYRCCFNETEGDVQNNPHV
IRIS_313-11694 MELITACALLLIAVHIAIVISCPEPLAAVALVDVPLLYLCRYLLIIRRGVIIVSGGGGGGRLRRFRLGAAMAIIYMAMSTLFFLRIAPLTPWWGALAAWVMILLIVEAIFAFFFPYRCCFNETEGDVQNNPHV
IRIS_313-11695 MELITACALLLIAVHIAIVISCPEPLAAVALVDVPLLYLCRYLLIIRRGVIIVSGGGGGGRLRRFRLGAAMAIIYMAMSTLFFLRIAPLTPWWGALAAWVMILLIVEAIFAFFFPYRCCFNETEGDVQNNPHV
IRIS_313-11698 MELITACALLLIAVHIAIVISCPEPLAAVALVDVPLLYLCRYLLIIRRGVIIVSGAGAGGGRLRRFRLGAAMAIIYMAMSTLFFLRIAPLTPWWGALAAWVMILLIVEAIFAFFFPYRCCFNETEGDVQNNPHV
IRIS_313-11700 MELITACALLLIAVHIAIVISCPEPLAAVALVDVPLLYLYRYLLIIRRGVIIVSGAGGGGGRLRRFRLGAAMAIIYMAMSTLFFLRIAPLTPWWGALAAWVMILLIVEAIFAFFFPYRCCFNETEGDVQNNPHV
IRIS_313-11702 MELITACALLLIAVHIAIVISCPEPLAAVALVDVPLLYLCRYLLIIRRGVIIVSGAGAGGGRLRRFRLGAAMAIIYMAMSTLFFLRIAPLTPWWGALAAWVMILLIVEAIFAFFFPYRCCFNETEGDVQNNPHV
IRIS_313-11704 MELITACALLLIAVHIAIVISCPEPLAAVALVDVPLLYLCRYLLIIRRGVIIVSGAGAGGGRLRRFRLGAAMAIIYMAMSTLFFLRIAPLTPWWGALAAWVMILLIVEAIFAFFFPYRCCFNETEGDVQNNPHV
IRIS_313-11705 MELITACALLLIAVHIAIVISCPEPLAAVALVDVPLLYLCRYLLIIRRGVIIVSGGGGGGRLRRFRLGAAMAIIYMAMSTLFFLRIAPLTPWWGALAAWVMILLIVEAIFAFFFPYRCCFNETEGDVQNNPHV
IRIS_313-11706 MELITACALLLIAVHIAIVISCPEPLAAVALVDVPLLYLCRYLLIIRRGVIIVSGAGGAAAAGCAASASGQPWRSSTWPCRPSSSSALLLSRHGGAPSPPGS*SCSSSRPSSPSSSPTAAASTKPKAMSRIIPM
IRIS_313-11707 MELITACALLLIAVHIAIVISCPEPLAAVALVDVPLLYLCRYLLIIRRGVIIVSGGGGGGRLRRFRLGAAMAIIYMAMSTLFFLRIAPLTPWWGALAAWVMILLIVEAIFAFFFPYRCCFNETEGDVQNNPHV
IRIS_313-11708 MELITACALLLIAVHIAIVISCPEPLAAVALVDVPLLYLCRYLLIIRRGVIIVSGGGGGGRLRRFRLGAAMAIIYMAMSTLFFLRIAPLTPWWGALAAWVMILLIVEAIFAFFFPYRCCFNETEGDVQNNPHV
IRIS_313-11709 MELITACALLLIAVHIAIVISCPEPLAAVALVDVPLLYLCRYLLIIRRGVIIVSGGGGGGRLRRFRLGAAMAIIYMAMSTLFFLRIAPLTPWWGALAAWVMILLIVEAIFAFFFPYRCCFNETEGDVQNNPHV
IRIS_313-11710 MELITACALLLIAVHIAIVISCPEPLAAVALVDVPLLYLCRYLLIIRRGVIIVSGGGGGGRLRRFRLGAAMAIIYMAMSTLFFLRIAPLTPWWGALAAWVMILLIVEAIFAFFFPYRCCFNETEGDVQNNPHV
IRIS_313-11711 MELITACALLLIAVHIAIVISCPEPLAAVALVDVPLLYLYRYLLIIRRGVIIVSGAGGGGGRLRRFRLGAAMAIIYMAMSTLFFLRIAPLTPWWGALAAWVMILLIVEAIFAFFFPYRCCFNETEGDVQNNPHV
IRIS_313-11712 MELITACALLLIAVHIAIVISCPEPLAAVALVDVPLLYLCRYLLIIRRGVIIVSGAGAGGGRLRRFRLGAAMAIIYMAMSTLFFLRIAPLTPWWGALAAWVMILLIVEAIFAFFFPYRCCFNETEGDVQNNPHV
IRIS_313-11713 MELITACALLLIAVHIAIVISCPEPLAAVALVDVPLLYLCRYLLIIRRGVIIVSGAGAGGGRLRRFRLGAAMAIIYMAMSTLFFLRIAPLTPWWGALAAWVMILLIVEAIFAFFFPYRCCFNETEGDVQNNPHV
IRIS_313-11714 MELITACALLLIAVHIAIVISCPEPLAAVALVDVPLLYLYRYLLIIRRGVIIVSGAGGGGGRLRRFRLGAAMAIIYMAMSTLFFLRIAPLTPWWGALAAWVMILLIVEAIFAFFFPYRCCFNETEGDVQNNPHV
IRIS_313-11716 MELITACALLLIAVHIAIVISCPEPLAAVALVDVPLLYLCRYLLIIRRGVIIVSGAGGAAAAGCAASASGQPWRSSTWPCRPSSSSALLLSRHGGAPSPPGS*SCSSSRPSSPSSSPTAAASTKPKAMSRIIPM
IRIS_313-11717 MELITACALLLIAVHIAIVISCPEPLAAVALVDVPLLYLCRYLLIIRRGVIIVSGGGGGGRLRRFRLGAAMAIIYMAMSTLFFLRIAPLTPWWGALAAWVMILLIVEAIFAFFFPYRCCFNETEGDVQNNPHV
IRIS_313-11719 MELITACALLLIAVHIAIVISCPEPLAAVALVDVPLLYLCRYLLIIRRGVIIVSGGGGGGRLRRFRLGAAMAIIYMAMSTLFFLRIAPLTPWWGALAAWVMILLIVEAIFAFFFPYRCCFNETEGDVQNNPHV
IRIS_313-11720 MELITACALLLIAVHIAIVISCPEPLAAVALVDVPLLYLYRYLLIIRRGVIIVSGAGGGGGRLRRFRLGAAMAIIYMAMSTLFFLRIAPLTPWWGALAAWVMILLIVEAIFAFFFPYRCCFNETEGDVQNNPHV
IRIS_313-11721 MELITACALLLIAVHIAIVISCPEPLAAVALVDVPLLYLCRYLLIIRRGVIIVSGGGGGGRLRRFRLGAAMAIIYMAMSTLFFLRIAPLTPWWGALAAWVMILLIVEAIFAFFFPYRCCFNETEGDVQNNPHV
IRIS_313-11722 MELITACALLLIAVHIAIVISCPEPLAAVALVDVPLLYLCRYLLIIRRGVIIVSGGGGGGRLRRFRLGAAMAIIYMAMSTLFFLRIAPLTPWWGALAAWVMILLIVEAIFAFFFPYRCCFNETEGDVQNNPHV
IRIS_313-11723 MELITACALLLIAVHIAIVISCPEPLAAVALVDVPLLYLCRYLLIIRRGVIIVSGAGAGGGRLRRFRLGAAMAIIYMAMSTLFFLRIAPLTPWWGALAAWVMILLIVEAIFAFFFPYRCCFNETEGDVQNNPHV
IRIS_313-11724 MELITACALLLIAVHIAIVISCPEPLAAVALVDVPLLYLCRYLLIIRRGVIIVSGAGGAAAAGCAASASGQPWRSSTWPCRPSSSSALLLSRHGGAPSPPGS*SCSSSRPSSPSSSPTAAASTKPKAMSRIIPM
IRIS_313-11725 MELITACALLLIAVHIAIVISCPEPLAAVALVDVPLLYLCRYLLIIRRGVIIVSGAGAGGGRLRRFRLGAAMAIIYMAMSTLFFLRIAPLTPWWGALAAWVMILLIVEAIFAFFFPYRCCFNETEGDVQNNPHV
IRIS_313-11726 MELITACALLLIAVHIAIVISCPEPLAAVALVDVPLLYLCRYLLIIRRGVIIVSGAGAGGGRLRRFRLGAAMAIIYMAMSTLFFLRIAPLTPWWGALAAWVMILLIVEAIFAFFFPYRCCFNETEGDVQNNPHV
IRIS_313-11727 MELITACALLLIAVHIAIVISCPEPLAAVALVDVPLLYLCRYLLIIRRGVIIVSGAGAGGGRLRRFRLGAAMAIIYMAMSTLFFLRIAPLTPWWGALAAWVMILLIVEAIFAFFFPYRCCFNETEGDVQNNPHV
IRIS_313-11728 MELITACALLLIAVHIAIVISCPEPLAAVALVDVPLLYLYRYLLIIRRGVIIVSGAGGGGGRLRRFRLGAAMAIIYMAMSTLFFLRIAPLTPWWGALAAWVMILLIVEAIFAFFFPYRCCFNETEGDVQNNPHV
IRIS_313-11729 MELITACALLLIAVHIAIVISCPEPLAAVALVDVPLLYLCRYLLIIRRGVIIVSGAGGGGGRLRRFRLGAAMAIIYMAMSTLFFLRIAPLTPWWGALAAWVMILLIVEAIFAFFFPYRCCFNETEGDVQNNPHV
IRIS_313-11730 MELITACALLLIAVHIAIVISCPEPLAAVALVDVPLLYLCRYLLIIRRGVIIVSGAGAGGGRLRRFRLGAAMAIIYMAMSTLFFLRIAPLTPWWGALAAWVMILLIVEAIFAFFFPYRCCFNETEGDVQNNPHV
IRIS_313-11731 MELITACALLLIAVHIAIVISCPEPLAAVALVDVPLLYLCRYLLIIRRGVIIVSGAGGGGGRLRRFRLGAAMAIIYMAMSTLFFLRIAPLTPWWGALAAWVMILLIVEAIFAFFFPYRCCFNETEGDVQNNPHV
IRIS_313-11732 MELITACALLLIAVHIAIVISCPEPLAAVALVDVPLLYLCRYLLIIRRGVIIVSGAGAGGGRLRRFRLGAAMAIIYMAMSTLFFLRIAPLTPWWGALAAWVMILLIVEAIFAFFFPYRCCFNETEGDVQNNPHV
IRIS_313-11733 MELITACALLLIAVHIAIVISCPEPLAAVALVDVPLLYLCRYLLIIRRGVIIVSGAGAGGGRLRRFRLGAAMAIIYMAMSTLFFLRIAPLTPWWGALAAWVMILLIVEAIFAFFFPYRCCFNETEGDVQNNPHV
IRIS_313-11734 MELITACALLLIAVHIAIVISCPEPLAAVALVDVPLLYLYRYLLIIRRGVIIVSGAGGGGGRLRRFRLGAAMAIIYMAMSTLFFLRIAPLTPWWGALAAWVMILLIVEAIFAFFFPYRCCFNETEGDVQNNPHV
IRIS_313-11735 MELITACALLLIAVHIAIVISCPEPLAAVALVDVPLLYLCRYLLIIRRGVIIVSGGGGGGRLRRFRLGAAMAIIYMAMSTLFFLRIAPLTPWWGALAAWVMILLIVEAIFAFFFPYRCCFNETEGDVQNNPHV
IRIS_313-11736 MELITACALLLIAVHIAIVISCPEPLAAVALVDVPLLYLCRYLLIIRRGVIIVSGAGGGGGRLRRFRLGAAMAIIYMAMSTLFFLRIAPLTPWWGALAAWVMILLIVEAIFAFFFPYRCCFNETEGDVQNNPHV
IRIS_313-11737 MELITACALLLIAVHIAIVISCPEPLAAVAFVNVPLLYLCRYLLIIRRGVIIVSGGGGGGGRLRRFRLGAAMAVIYMAMSTLFFLRIAPLTPWWGALAAWVMILLIVEAIFAFFFPYRCCFNETEGDVQNNPHV
IRIS_313-11738 MELITACALLLIAVHIAIVISCPEPLAAVALVDVPLLYLCRYLLIIRRGVIIVSGGGGGGRLRRFRLGAAMAIIYMAMSTLFFLRIAPLTPWWGALAAWVMILLIVEAIFAFFFPYRCCFNETEGDVQNNPHV
IRIS_313-11739 MELITACALLLIAVHIAIVISCPEPLAAVALVDVPLLYLCRYLLIIRRGVIIVSGAGAGGGRLRRFRLGAAMAIIYMAMSTLFFLRIAPLTPWWGALAAWVMILLIVEAIFAFFFPYRCCFNETEGDVQNNPHV
IRIS_313-11740 MELITACALLLIAVHIAIVISCPEPLAAVALVDVPLLYLCRYLLIIRRGVIIVSGAGGAAAAGCAASASGQPWRSSTWPCRPSSSSALLLSRHGGAPSPPGS*SCSSSRPSSPSSSPTAAASTKPKAMSRIIPM
IRIS_313-11741 MELITACALLLIAVHIAIVISCPEPLAAVALVDVPLLYLCRYLLIIRRGVIIVSGGGGGGRLRRFRLGAAMAIIYMAMSTLFFLRIAPLTPWWGALAAWVMILLIVEAIFAFFFPYRCCFNETEGDVQNNPHV
IRIS_313-11742 MELITACALLLIAVHIAIVISCPEPLAAVAFVNVPLLYLCRYLLIIRRGVIIVSGGGGGGGRLRRFRLGAAMAVIYMAMSTLFFLRIAPLTPWWGALAAWVMILLIVEAIFAFFFPYRCCFNETEGDVQNNPHV
IRIS_313-11743 MELITACALLLIAVHIAIVISCPEPLAAVALVDVPLLYLCRYLLIIRRGVIIVSGAGAGGGRLRRFRLGAAMAIIYMAMSTLFFLRIAPLTPWWGALAAWVMILLIVEAIFAFFFPYRCCFNETEGDVQNNPHV
IRIS_313-11744 MELITACALLLIAVHIAIVISCPEPLAAVALVDVPLLYLCRYLLIIRRGVIIVSGGGGGGRLRRFRLGAAMAIIYMAMSTLFFLRIAPLTPWWGALAAWVMILLIVEAIFAFFFPYRCCFNETEGDVQNNPHV
IRIS_313-11745 MELITACALLLIAVHIAIVISCPEPLAAVALVDVPLLYLYRYLLIIRRGVIIVSGAGGGGGRLRRFRLGAAMAIIYMAMSTLFFLRIAPLTPWWGALAAWVMILLIVEAIFAFFFPYRCCFNETEGDVQNNPHV
IRIS_313-11746 MELITACALLLIAVHIAIVISCPEPLAAVALVDVPLLYLCRYLLIIRRGVIIVSGAGAGGGRLRRFRLGAAMAIIYMAMSTLFFLRIAPLTPWWGALAAWVMILLIVEAIFAFFFPYRCCFNETEGDVQNNPHV
IRIS_313-11747 MELITACALLLIAVHIAIVISCPEPLAAVALVDVPLLYLCRYLLIIRRGVIIVSGGGGGGRLRRFRLGAAMAIIYMAMSTLFFLRIAPLTPWWGALAAWVMILLIVEAIFAFFFPYRCCFNETEGDVQNNPHV
IRIS_313-11748 MELITACALLLIAVHIAIVISCPEPLAAVALVDVPLLYLCRYLLIIRRGVIIVSGGGGGGRLRRFRLGAAMAIIYMAMSTLFFLRIAPLTPWWGALAAWVMILLIVEAIFAFFFPYRCCFNETEGDVQNNPHV
IRIS_313-11750 MELITACALLLIAVHIAIVISCPEPLAAVALVDVPLLYLCRYLLIIRRGVIIVSGGGGGGRLRRFRLGAAMAIIYMAMSTLFFLRIAPLTPWWGALAAWVMILLIVEAIFAFFFPYRCCFNETEGDVQNNPHV
IRIS_313-11751 MELITACALLLIAVHIAIVISCPEPLAAVALVDVPLLYLCRYLLIIRRGVIIVSGGGGGGRLRRFRLGAAMAIIYMAMSTLFFLRIAPLTPWWGALAAWVMILLIVEAIFAFFFPYRCCFNETEGDVQNNPHV
IRIS_313-11752 MELITACALLLIAVHIAIVISCPEPLAAVALVDVPLLYLYRYLLIIRRGVIIVSGAGGGGGRLRRFRLGAAMAIIYMAMSTLFFLRIAPLTPWWGALAAWVMILLIVEAIFAFFFPYRCCFNETEGDVQNNPHV
IRIS_313-11753 MELITACALLLIAVHIAIVISCPEPLAAVALVDVPLLYLCRYLLIIRRGVIIVSGGGAGGGRLRRFRLGAAMAIIYMAMSTLFFLRIAPLTPWWGALAAWVMILLIVEAIFAFFFPYRCCFNETEGDVQNNPHV
IRIS_313-11754 MELITACALLLIAVHIAIVISCPEPLAAVALVDVPLLYLCRYLLIIRRGVIIVSGAGAGGGRLRRFRLGAAMAIIYMAMSTLFFLRIAPLTPWWGALAAWVMILLIVEAIFAFFFPYRCCFNETEGDVQNNPHV
IRIS_313-11755 MELITACALLLIAVHIAIVISCPEPLAAVALVDVPLLYLCRYLLIIRRGVIIVSGAGAGGGRLRRFRLGAAMAIIYMAMSTLFFLRIAPLTPWWGALAAWVMILLIVEAIFAFFFPYRCCFNETEGDVQNNPHV
IRIS_313-11756 MELITACALLLIAVHIAIVISCPEPLAAVALVDVPLLYLYRYLLIIRRGVIIVSGAGGGGGRLRRFRLGAAMAIIYMAMSTLFFLRIAPLTPWWGALAAWVMILLIVEAIFAFFFPYRCCFNETEGDVQNNPHV
IRIS_313-11757 MELITACALLLIAVHIAIVISCPEPLAAVALVDVPLLYLCRYLLIIRRGVIIVSGGGGGGRLRRFRLGAAMAIIYMAMSTLFFLRIAPLTPWWGALAAWVMILLIVEAIFAFFFPYRCCFNETEGDVQNNPHV
IRIS_313-11758 MELITACALLLIAVHIAIVISCPEPLAAVALVDVPLLYLCRYLLIIRRGVIIVSGAGAGGGRLRRFRLGAAMAIIYMAMSTLFFLRIAPLTPWWGALAAWVMILLIVEAIFAFFFPYRCCFNETEGDVQNNPHV
IRIS_313-11759 MELITACALLLIAVHIAIVISCPEPLAAVALVDVPLLYLYRYLLIIRRGVIIVSGAGGGGGRLRRFRLGAAMAIIYMAMSTLFFLRIAPLTPWWGALAAWVMILLIVEAIFAFFFPYRCCFNETEGDVQNNPHV
IRIS_313-11760 MELITACALLLIAVHIAIVISCPEPLAAVALVDVPLLYLCRYLLIIRRGVIIVSGAGAGGGRLRRFRLGAAMAIIYMAMSTLFFLRIAPLTPWWGALAAWVMILLIVEAIFAFFFPYRCCFNETEGDVQNNPHV
IRIS_313-11761 MELITACALLLIAVHIAIVISCPEPLAAVALVDVPLLYLCRYLLIIRRGVIIVSGAGAGGGRLRRFRLGAAMAIIYMAMSTLFFLRIAPLTPWWGALAAWVMILLIVEAIFAFFFPYRCCFNETEGDVQNNPHV
IRIS_313-11762 MELITACALLLIAVHIAIVISCPEPLAAVALVDVPLLYLCRYLLIIRRGVIIVSGAGGGGGGRLRRFRLGAAMAIIYMAMSTLFFLRIAPLTPWWGALAAWVMILLIVEAIFAFFFPYRCCFNETEGDVQNNPHV
IRIS_313-11763 MELITACALLLIAVHIAIVISCPEPLAAVALVDVPLLYLCRYLLIIRRGVIIVSGAGAGGGRLRRFRLGAAMAIIYMAMSTLFFLRIAPLTPWWGALAAWVMILLIVEAIFAFFFPYRCCFNETEGDVQNNPHV
IRIS_313-11764 MELITACALLLIAVHIAIVISCPEPLAAVALVDVPLLYLYRYLLIIRRGVIIVSGAGGGGGRLRRFRLGAAMAIIYMAMSTLFFLRIAPLTPWWGALAAWVMILLIVEAIFAFFFPYRCCFNETEGDVQNNPHV
IRIS_313-11765 MELITACALLLIAVHIAIVISCPEPLAAVALVDVPLLYLCRYLLIIRRGVIIVSGAGAGGGRLRRFRLGAAMAIIYMAMSTLFFLRIAPLTPWWGALAAWVMILLIVEAIFAFFFPYRCCFNETEGDVQNNPHV
IRIS_313-11766 MELITACALLLIAVHIAIVISCPEPLAAVALVDVPLLYLCRYLLIIRRGVIIVSGAGGAAAAGCAASASGQPWRSSTWPCRPSSSSALLLSRHGGAPSPPGS*SCSSSRPSSPSSSPTAAASTKPKAMSRIIPM

IRIS_313-11767 MELITACALLLIAVHIAIVISCPEPLAAVALVDVPLLYLCRYLLIIRRGVIIVSGAGAGGGRLRRFRLGAAMAIIYMAMSTLFFLRIAPLTPWWGALAAWVMILLIVEAIFAFFFPYRCCFNETEGDVQNNPHV
IRIS_313-11772 MELITACALLLIAVHIAIVISCPEPLAAVALVDVPLLYLCRYLLIIRRGVIIVSGAGAGGGRLRRFRLGAAMAIIYMAMSTLFFLRIAPLTPWWGALAAWVMILLIVEAIFAFFFPYRCCFNETEGDVQNNPHV
IRIS_313-11773 MELITACALLLIAVHIAIVISCPEPLAAVALVDVPLLYLCRYLLIIRRGVIIVSGAGAGGGRLRRFRLGAAMAIIYMAMSTLFFLRIAPLTPWWGALAAWVMILLIVEAIFAFFFPYRCCFNETEGDVQNNPHV
IRIS_313-11779 MELITACALLLIAVHIAIVISCPEPLAAVALVDVPLLYLCRYLLIIRRGVIIVSGGGGGGRLRRFRLGAAMAIIYMAMSTLFFLRIAPLTPWWGALAAWVMILLIVEAIFAFFFPYRCCFNETEGDVQNNPHV
IRIS_313-11780 MELITACALLLIAVHIAIVISCPEPLAAVALVDVPLLYLCRYLLIIRRGVIIVSGGGGGGRLRRFRLGAAMAIIYMAMSTLFFLRIAPLTPWWGALAAWVMILLIVEAIFAFFFPYRCCFNETEGDVQNNPHV
IRIS_313-11782 MELITACALLLIAVHIAIVISCPEPLAAVALVDVPLLYLCRYLLIIRRGVIIVSGAGAGGGRLRRFRLGAAMAIIYMAMSTLFFLRIAPLTPWWGALAAWVMILLIVEAIFAFFFPYRCCFNETEGDVQNNPHV
IRIS_313-11783 MELITACALLLIAVHIAIVISCPEPLAAVALVDVPLLYLCRYLLIIRRGVIIVSGAGAGGGRLRRFRLGAAMAIIYMAMSTLFFLRIAPLTPWWGALAAWVMILLIVEAIFAFFFPYRCCFNETEGDVQNNPHV
IRIS_313-11784 MELITACALLLIAVHIAIVISCPEPLAAVALVDVPLLYLCRYLLIIRRGVIIVSGGGGGGRLRRFRLGAAMAIIYMAMSTLFFLRIAPLTPWWGALAAWVMILLIVEAIFAFFFPYRCCFNETEGDVQNNPHV
IRIS_313-11785 MELITACALLLIAVHIAIVISCPEPLAAVALVDVPLLYLCRYLLIIRRGVIIVSGGGGGGRLRRFRLGAAMAIIYMAMSTLFFLRIAPLTPWWGALAAWVMILLIVEAIFAFFFPYRCCFNETEGDVQNNPHV
IRIS_313-11786 MELITACALLLIAVHIAIVISCPEPLAAVALVDVPLLYLYRYLLIIRRGVIIVSGAGGGGGRLRRFRLGAAMAIIYMAMSTLFFLRIAPLTPWWGALAAWVMILLIVEAIFAFFFPYRCCFNETEGDVQNNPHV
IRIS_313-11787 MELITACALLLIAVHIAIVISCPEPLAAVALVDVPLLYLCRYLLIIRRGVIIVSGAGAGGGRLRRFRLGAAMAIIYMAMSTLFFLRIAPLTPWWGALAAWVMILLIVEAIFAFFFPYRCCFNETEGDVQNNPHV
IRIS_313-11788 MELITACALLLIAVHIAIVISCPEPLAAVALVDVPLLYLCRYLLIIRRGVIIVSGAGAGGGRLRRFRLGAAMAIIYMAMSTLFFLRIAPLTPWWGALAAWVMILLIVEAIFAFFFPYRCCFNETEGDVQNNPHV
IRIS_313-11789 MELITACALLLIAVHIAIVISCPEPLAAVALVDVPLLYLCRYLLIIRRGVIIVSGAGAGGGRLRRFRLGAAMAIIYMAMSTLFFLRIAPLTPWWGALAAWVMILLIVEAIFAFFFPYRCCFNETEGDVQNNPHV
IRIS_313-11790 MELITACALLLIAVHIAIVISCPEPLAAVALVDVPLLYLCRYLLIIRRGVIIVSGAGAGGGRLRRFRLGAAMAIIYMAMSTLFFLRIAPLTPWWGALAAWVMILLIVEAIFAFFFPYRCCFNETEGDVQNNPHV
IRIS_313-11791 MELITACALLLIAVHIAIVISCPEPLAAVALVDVPLLYLCRYLLIIRRGVIIVSGAGAGGGRLRRFRLGAAMAIIYMAMSTLFFLRIAPLTPWWGALAAWVMILLIVEAIFAFFFPYRCCFNETEGDVQNNPHV
IRIS_313-11792 MELITACALLLIAVHIAIVISCPEPLAAVALVDVPLLYLYRYLLIIRRGVIIVSGAGGGGGRLRRFRLGAAMAIIYMAMSTLFFLRIAPLTPWWGALAAWVMILLIVEAIFAFFFPYRCCFNETEGDVQNNPHV
IRIS_313-11793 MELITACALLLIAVHIAIVISCPEPLAAVALVDVPLLYLCRYLLIIRRGVIIVSGAGAGGGRLRRFRLGAAMAIIYMAMSTLFFLRIAPLTPWWGALAAWVMILLIVEAIFAFFFPYRCCFNETEGDVQNNPHV
IRIS_313-11794 MELITACALLLIAVHIAIVISCPEPLAAVALVDVPLLYLCRYLLIIRRGVIIVSGAGAGGGRLRRFRLGAAMAIIYMAMSTLFFLRIAPLTPWWGALAAWVMILLIVEAIFAFFFPYRCCFNETEGDVQNNPHV
IRIS_313-11795 MELITACALLLIAVHIAIVISCPEPLAAVALVDVPLLYLYRYLLIIRRGVIIVSGAGGGGGRLRRFRLGAAMAIIYMAMSTLFFLRIAPLTPWWGALAAWVMILLIVEAIFAFFFPYRCCFNETEGDVQNNPHV
IRIS_313-11796 MELITACALLLIAVHIAIVISCPEPLAAVALVDVPLLYLCRYLLIIRRGVIIVSGGGGGGRLRRFRLGAAMAIIYMAMSTLFFLRIAPLTPWWGALAAWVMILLIVEAIFAFFFPYRCCFNETEGDVQNNPHV
IRIS_313-11797 MELITACALLLIAVHIAIVISCPEPLAAVALVDVPLLYLYRYLLIIRRGVIIVSGAGGGGGRLRRFRLGAAMAIIYMAMSTLFFLRIAPLTPWWGALAAWVMILLIVEAIFAFFFPYRCCFNETEGDVQNNPHV
IRIS_313-11798 MELITACALLLIAVHIAIVISCPEPLAAVALVDVPLLYLCRYLLIIRRGVIIVSGGGGGGRLRRFRLGAAMAIIYMAMSTLFFLRIAPLTPWWGALAAWVMILLIVEAIFAFFFPYRCCFNETEGDVQNNPHV
IRIS_313-11799 MELITACALLLIAVHIAIVISCPEPLAAVALVDVPLLYLYRYLLIIRRGVIIVSGAGGGGGRLRRFRLGAAMAIIYMAMSTLFFLRIAPLTPWWGALAAWVMILLIVEAIFAFFFPYRCCFNETEGDVQNNPHV
IRIS_313-11800 MELITACALLLIAVHIAIVISCPEPLAAVALVDVPLLYLCRYLLIIRRGVIIVSGAGAGGGRLRRFRLGAAMAIIYMAMSTLFFLRIAPLTPWWGALAAWVMILLIVEAIFAFFFPYRCCFNETEGDVQNNPHV
IRIS_313-11801 MELITACALLLIAVHIAIVISCPEPLAAVALVDVPLLYLCRYLLIIRRGVIIVSGGGGGGRLRRFRLGAAMAIIYMAMSTLFFLRIAPLTPWWGALAAWVMILLIVEAIFAFFFPYRCCFNETEGDVQNNPHV
IRIS_313-11802 MELITACALLLIAVHIAIVISCPEPLAAVALVDVPLLYLYRYLLIIRRGVIIVSGAGGGGGRLRRFRLGAAMAIIYMAMSTLFFLRIAPLTPWWGALAAWVMILLIVEAIFAFFFPYRCCFNETEGDVQNNPHV
IRIS_313-11803 MELITACALLLIAVHIAIVISCPEPLAAVALVDVPLLYLCRYLLIIRRGVIIVSGAGGGGGRLRRFRLGAAMAIIYMAMSTLFFLRIAPLTPWWGALAAWVMILLIVEAIFAFFFPYRCCFNETEGDVQNNPHV
IRIS_313-11804 MELITACALLLIAVHIAIVISCPEPLAAVALVDVPLLYLCRYLLIIRRGVIIVSGGGGGGRLRRFRLGAAMAIIYMAMSTLFFLRIAPLTPWWGALAAWVMILLIVEAIFAFFFPYRCCFNETEGDVQNNPHV
IRIS_313-11805 MELITACALLLIAVHIAIVISCPEPLAAVALVDVPLLYLYRYLLIIRRGVIIVSGAGGGGGRLRRFRLGAAMAIIYMAMSTLFFLRIAPLTPWWGALAAWVMILLIVEAIFAFFFPYRCCFNETEGDVQNNPHV
IRIS_313-11806 MELITACALLLIAVHIAIVISCPEPLAAVALVDVPLLYLCRYLLIIRRGVIIVSGAGAGGGRLRRFRLGAAMAIIYMAMSTLFFLRIAPLTPWWGALAAWVMILLIVEAIFAFFFPYRCCFNETEGDVQNNPHV
IRIS_313-11807 MELITACALLLIAVHIAIVISCPEPLAAVALVDVPLLYLCRYLLIIRRGVIIVSGGGGGGRLRRFRLGAAMAIIYMAMSTLFFLRIAPLTPWWGALAAWVMILLIVEAIFAFFFPYRCCFNETEGDVQNNPHV
IRIS_313-11808 MELITACALLLIAVHIAIVISCPEPLAAVALVDVPLLYLCRYLLIIRRGVIIVSGGGGGGRLRRFRLGAAMAIIYMAMSTLFFLRIAPLTPWWGALAAWVMILLIVEAIFAFFFPYRCCFNETEGDVQNNPHV
IRIS_313-11809 MELITACALLLIAVHIAIVISCPEPLAAVALVDVPLLYLCRYLLIIRRGVIIVSGAGAGGGRLRRFRLGAAMAIIYMAMSTLFFLRIAPLTPWWGALAAWVMILLIVEAIFAFFFPYRCCFNETEGDVQNNPHV
IRIS_313-11810 MELITACALLLIAVHIAIVISCPEPLAAVALVDVPLLYLCRYLLIIRRGVIIVSGGGGGGRLRRFRLGAAMAIIYMAMSTLFFLRIAPLTPWWGALAAWVMILLIVEAIFAFFFPYRCCFNETEGDVQNNPHV
IRIS_313-11811 MELITACALLLIAVHIAIVISCPEPLAAVALVDVPLLYLCRYLLIIRRGVIIVSGAGAGGGRLRRFRLGAAMAIIYMAMSTLFFLRIAPLTPWWGALAAWVMILLIVEAIFAFFFPYRCCFNETEGDVQNNPHV
IRIS_313-11812 MELITACALLLIAVHIAIVISCPEPLAAVALVDVPLLYLYRYLLIIRRGVIIVSGAGGGGGRLRRFRLGAAMAIIYMAMSTLFFLRIAPLTPWWGALAAWVMILLIVEAIFAFFFPYRCCFNETEGDVQNNPHV
IRIS_313-11813 MELITACALLLIAVHIAIVISCPEPLAAVALVDVPLLYLCRYLLIIRRGVIIVSGGGGGGRLRRFRLGAAMAIIYMAMSTLFFLRIAPLTPWWGALAAWVMILLIVEAIFAFFFPYRCCFNETEGDVQNNPHV
IRIS_313-11814 MELITACALLLIAVHIAIVISCPEPLAAVALVDVPLLYLCRYLLIIRRGVIIVSGGGAGGGRLRRFRLGAAMAIIYMAMSTLFFLRIAPLTPWWGALAAWVMILLIVEAIFAFFFPYRCCFNETEGDVQNNPHV
IRIS_313-11815 MELITACALLLIAVHIAIVISCPEPLAAVALVDVPLLYLYRYLLIIRRGVIIVSGAGGGGGRLRRFRLGAAMAIIYMAMSTLFFLRIAPLTPWWGALAAWVMILLIVEAIFAFFFPYRCCFNETEGDVQNNPHV
IRIS_313-11816 MELITACALLLIAVHIAIVISCPEPLAAVALVDVPLLYLCRYLLIIRRGVIIVSGAGAGGGRLRRFRLGAAMAIIYMAMSTLFFLRIAPLTPWWGALAAWVMILLIVEAIFAFFFPYRCCFNETEGDVQNNPHV
IRIS_313-11817 MELITACALLLIAVHIAIVISCPEPLAAVALVDVPLLYLCRYLLIIRRGVIIVSGGGGGGRLRRFRLGAAMAIIYMAMSTLFFLRIAPLTPWWGALAAWVMILLIVEAIFAFFFPYRCCFNETEGDVQNNPHV
IRIS_313-11818 MELITACALLLIAVHIAIVISCPEPLAAVALVDVPLLYLCRYLLIIRRGVIIVSGAGGAAAAGCAASASGQPWRSSTWPCRPSSSSALLLSRHGGAPSPPGS*SCSSSRPSSPSSSPTAAASTKPKAMSRIIPM
IRIS_313-11819 MELITACALLLIAVHIAIVISCPEPLAAVALVDVPLLYLCRYLLIIRRGVIIVSGGGGGGRLRRFRLGAAMAIIYMAMSTLFFLRIAPLTPWWGALAAWVMILLIVEAIFAFFFPYRCCFNETEGDVQNNPHV
IRIS_313-11820 MELITACALLLIAVHIAIVISCPEPLAAVALVDVPLLYLCRYLLIIRRGVIIVSGGGAGGGRLRRFRLGAAMAIIYMAMSTLFFLRIAPLTPWWGALAAWVMILLIVEAIFAFFFPYRCCFNETEGDVQNNPHV
IRIS_313-11821 MELITACALLLIAVHIAIVISCPEPLAAVALVDVPLLYLCRYLLIIRRGVIIVSGGGGGGRLRRFRLGAAMAIIYMAMSTLFFLRIAPLTPWWGALAAWVMILLIVEAIFAFFFPYRCCFNETEGDVQNNPHV
IRIS_313-11822 MELITACALLLIAVHIAIVISCPEPLAAVALVDVPLLYLCRYLLIIRRGVIIVSGGGAGGGRLRRFRLGAAMAIIYMAMSTLFFLRIAPLTPWWGALAAWVMILLIVEAIFAFFFPYRCCFNETEGDVQNNPHV
IRIS_313-11823 MELITACALLLIAVHIAIVISCPEPLAAVALVDVPLLYLCRYLLIIRRGVIIVSGGGGGGRLRRFRLGAAMAIIYMAMSTLFFLRIAPLTPWWGALAAWVMILLIVEAIFAFFFPYRCCFNETEGDVQNNPHV
IRIS_313-11824 MELITACALLLIAVHIAIVISCPEPLAAVALVDVPLLYLCRYLLIIRRGVIIVSGGGGGGRLRRFRLGAAMAIIYMAMSTLFFLRIAPLTPWWGALAAWVMILLIVEAIFAFFFPYRCCFNETEGDVQNNPHV
IRIS_313-11825 MELITACALLLIAVHIAIVISCPEPLAAVALVDVPLLYLCRYLLIIRRGVIIVSGAGAGGGRLRRFRLGAAMAIIYMAMSTLFFLRIAPLTPWWGALAAWVMILLIVEAIFAFFFPYRCCFNETEGDVQNNPHV
IRIS_313-11826 MELITACALLLIAVHIAIVISCPEPLAAVALVDVPLLYLCRYLLIIRRGVIIVSGAGAGGGRLRRFRLGAAMAIIYMAMSTLFFLRIAPLTPWWGALAAWVMILLIVEAIFAFFFPYRCCFNETEGDVQNNPHV
IRIS_313-11827 MELITACALLLIAVHIAIVISCPEPLAAVALVDVPLLYLCRYLLIIRRGVIIVSGGGGGGRLRRFRLGAAMAIIYMAMSTLFFLRIAPLTPWWGALAAWVMILLIVEAIFAFFFPYRCCFNETEGDVQNNPHV
IRIS_313-11828 MELITACALLLIAVHIAIVISCPEPLAAVALVDVPLLYLCRYLLIIRRGVIIVSGGGGGGRLRRFRLGAAMAIIYMAMSTLFFLRIAPLTPWWGALAAWVMILLIVEAIFAFFFPYRCCFNETEGDVQNNPHV
IRIS_313-11829 MELITACALLLIAVHIAIVISCPEPLAAVALVDVPLLYLCRYLLIIRRGVIIVSGAGAGGGRLRRFRLGAAMAIIYMAMSTLFFLRIAPLTPWWGALAAWVMILLIVEAIFAFFFPYRCCFNETEGDVQNNPHV
IRIS_313-11830 MELITACALLLIAVHIAIVISCPEPLAAVALVDVPLLYLCRYLLIIRRGVIIVSGGGGGGRLRRFRLGAAMAIIYMAMSTLFFLRIAPLTPWWGALAAWVMILLIVEAIFAFFFPYRCCFNETEGDVQNNPHV
IRIS_313-11831 MELITACALLLIAVHIAIVISCPEPLAAVALVDVPLLYLCRYLLIIRRGVIIVSGAGGGGGRLRRFRLGAAMAIIYMAMSTLFFLRIAPLTPWWGALAAWVMILLIVEAIFAFFFPYRCCFNETEGDVQNNPHV
IRIS_313-11832 MELITACALLLIAVHIAIVISCPEPLAAVALVDVPLLYLYRYLLIIRRGVIIVSGAGGGGGRLRRFRLGAAMAIIYMAMSTLFFLRIAPLTPWWGALAAWVMILLIVEAIFAFFFPYRCCFNETEGDVQNNPHV
IRIS_313-11833 MELITACALLLIAVHIAIVISCPEPLAAVALVDVPLLYLCRYLLIIRRGVIIVSGGGGGGRLRRFRLGAAMAIIYMAMSTLFFLRIAPLTPWWGALAAWVMILLIVEAIFAFFFPYRCCFNETEGDVQNNPHV
IRIS_313-11834 MELITACALLLIAVHIAIVISCPEPLAAVALVDVPLLYLYRYLLIIRRGVIIVSGAGAGGGRLRRFRLGAAMAIIYMAMSTLFFLRIAPLTPWWGALAAWVMILLIVEAIFAFFFPYRCCFNETEGDVQNNPHV
IRIS_313-11835 MELITACALLLIAVHIAIVISCPEPLAAVALVDVPLLYLCRYLLIIRRGVIIVSGGGGGGRLRRFRLGAAMAIIYMAMSTLFFLRIAPLTPWWGALAAWVMILLIVEAIFAFFFPYRCCFNETEGDVQNNPHV
IRIS_313-11836 MELITACALLLIAVHIAIVISCPEPLAAVALVDVPLLYLCRYLLIIRRGVIIVSGGGGGGRLRRFRLGAAMAIIYMAMSTLFFLRIAPLTPWWGALAAWVMILLIVEAIFAFFFPYRCCFNETEGDVQNNPHV
IRIS_313-11837 MELITACALLLIAVHIAIVISCPEPLAAVALVDVPLLYLCRYLLIIRRGVIIVSGGGGGGRLRRFRLGAAMAIIYMAMSTLFFLRIAPLTPWWGALAAWVMILLIVEAIFAFFFPYRCCFNETEGDVQNNPHV
IRIS_313-11838 MELITACALLLIAVHIAIVISCPEPLAAVALVDVPLLYLCRYLLIIRRGVIIVSGGGGGGRLRRFRLGAAMAIIYMAMSTLFFLRIAPLTPWWGALAAWVMILLIVEAIFAFFFPYRCCFNETEGDVQNNPHV
IRIS_313-11839 MELITACALLLIAVHIAIVISCPEPLAAVALVDVPLLYLYRYLLIIRRGVIIVSGAGGGGGRLRRFRLGAAMAIIYMAMSTLFFLRIAPLTPWWGALAAWVMILLIVEAIFAFFFPYRCCFNETEGDVQNNPHV
IRIS_313-11840 MELITACALLLIAVHIAIVISCPEPLAAVALVDVPLLYLCRYLLIIRRGVIIVSGGGGGGRLRRFRLGAAMAIIYMAMSTLFFLRIAPLTPWWGALAAWVMILLIVEAIFAFFFPYRCCFNETEGDVQNNPHV
IRIS_313-11841 MELITACALLLIAVHIAIVISCPEPLAAVALVDVPLLYLYRYLLIIRRGVIIVSGAGGGGGRLRRFRLGAAMAIIYMAMSTLFFLRIAPLTPWWGALAAWVMILLIVEAIFAFFFPYRCCFNETEGDVQNNPHV
IRIS_313-11842 MELITACALLLIAVHIAIVISCPEPLAAVALVDVPLLYLCRYLLIIRRGVIIVSGGGGGGRLRRFRLGAAMAIIYMAMSTLFFLRIAPLTPWWGALAAWVMILLIVEAIFAFFFPYRCCFNETEGDVQNNPHV
IRIS_313-11843 MELITACALLLIAVHIAIVISCPEPLAAVALVDVPLLYLCRYLLIIRRGVIIVSGGGGGGRLRRFRLGAAMAIIYMAMSTLFFLRIAPLTPWWGALAAWVMILLIVEAIFAFFFPYRCCFNETEGDVQNNPHV
IRIS_313-11844 MELITACALLLIAVHIAIVISCPEPLAAVALVDVPLLYLYRYLLIIRRGVIIVSGAGGGGGRLRRFRLGAAMAIIYMAMSTLFFLRIAPLTPWWGALAAWVMILLIVEAIFAFFFPYRCCFNETEGDVQNNPHV
IRIS_313-11845 MELITACALLLIAVHIAIVISCPEPLAAVALVDVPLLYLYRYLLIIRRGVIIVSGAGGGGGRLRRFRLGAAMAIIYMAMSTLFFLRIAPLTPWWGALAAWVMILLIVEAIFAFFFPYRCCFNETEGDVQNNPHV
IRIS_313-11846 MELITACALLLIAVHIAIVISCPEPLAAVALVDVPLLYLCRYLLIIRRGVIIVSGAGAGGGRLRRFRLGAAMAIIYMAMSTLFFLRIAPLTPWWGALAAWVMILLIVEAIFAFFFPYRCCFNETEGDVQNNPHV
IRIS_313-11848 MELITACALLLIAVHIAIVISCPEPLAAVALVDVPLLYLCRYLLIIRRGVIIVSGGGGGGRLRRFRLGAAMAIIYMAMSTLFFLRIAPLTPWWGALAAWVMILLIVEAIFAFFFPYRCCFNETEGDVQNNPHV
IRIS_313-11849 MELITACALLLIAVHIAIVISCPEPLAAVALVDVPLLYLCRYLLIIRRGVIIVSGAGGAAAAGCAASASGQPWRSSTWPCRPSSSSALLLSRHGGAPSPPGS*SCSSSRPSSPSSSPTAAASTKPKAMSRIIPM
IRIS_313-11850 MELITACALLLIAVHIAIVISCPEPLAAVALVDVPLLYLCRYLLIIRRGVIIVSGAGAGGGRLRRFRLGAAMAIIYMAMSTLFFLRIAPLTPWWGALAAWVMILLIVEAIFAFFFPYRCCFNETEGDVQNNPHV
IRIS_313-11851 MELITACALLLIAVHIAIVISCPEPLAAVALVDVPLLYLCRYLLIIRRGVIIVSGAGAGGGRLRRFRLGAAMAIIYMAMSTLFFLRIAPLTPWWGALAAWVMILLIVEAIFAFFFPYRCCFNETEGDVQNNPHV
IRIS_313-11852 MELITACALLLIAVHIAIVISCPEPLAAVALVDVPLLYLCRYLLIIRRGVIIVSGGGGGGRLRRFRLGAAMAIIYMAMSTLFFLRIAPLTPWWGALAAWVMILLIVEAIFAFFFPYRCCFNETEGDVQNNPHV
IRIS_313-11853 MELITACALLLIAVHIAIVISCPEPLAAVALVDVPLLYLCRYLLIIRRGVIIVSGAGAGGGRLRRFRLGAAMAIIYMAMSTLFFLRIAPLTPWWGALAAWVMILLIVEAIFAFFFPYRCCFNETEGDVQNNPHV
IRIS_313-11854 MELITACALLLIAVHIAIVISCPEPLAAVALVDVPLLYLCRYLLIIRRGVIIVSGAGAGGGRLRRFRLGAAMAIIYMAMSTLFFLRIAPLTPWWGALAAWVMILLIVEAIFAFFFPYRCCFNETEGDVQNNPHV
IRIS_313-11855 MELITACALLLIAVHIAIVISCPEPLAAVALVDVPLLYLCRYLLIIRRGVIIVSGGGGGGRLRRFRLGAAMAIIYMAMSTLFFLRIAPLTPWWGALAAWVMILLIVEAIFAFFFPYRCCFNETEGDVQNNPHV
IRIS_313-11856 MELITACALLLIAVHIAIVISCPEPLAAVALVDVPLLYLCRYLLIIRRGVIIVSGGGGGGRLRRFRLGAAMAIIYMAMSTLFFLRIAPLTPWWGALAAWVMILLIVEAIFAFFFPYRCCFNETEGDVQNNPHV
IRIS_313-11857 MELITACALLLIAVHIAIVISCPEPLAAVALVDVPLLYLYRYLLIIRRGVIIVSGAGGGGGRLRRFRLGAAMAIIYMAMSTLFFLRIAPLTPWWGALAAWVMILLIVEAIFAFFFPYRCCFNETEGDVQNNPHV
IRIS_313-11858 MELITACALLLIAVHIAIVISCPEPLAAVALVDVPLLYLCRYLLIIRRGVIIVSGGGGGGRLRRFRLGAAMAIIYMAMSTLFFLRIAPLTPWWGALAAWVMILLIVEAIFAFFFPYRCCFNETEGDVQNNPHV
IRIS_313-11859 MELITACALLLIAVHIAIVISCPEPLAAVALVDVPLLYLCRYLLIIRRGVIIVSGGGAGGGRLRRFRLGAAMAIIYMAMSTLFFLRIAPLTPWWGALAAWVMILLIVEAIFAFFFPYRCCFNETEGDVQNNPHV
IRIS_313-11860 MELITACALLLIAVHIAIVISCPEPLAAVALVDVPLLYLCRYLLIIRRGVIIVSGAGGGGGRLRRFRLGAAMAIIYMAMSTLFFLRIAPLTPWWGALAAWVMILLIVEAIFAFFFPYRCCFNETEGDVQNNPHV
IRIS_313-11861 MELITACALLLIAVHIAIVISCPEPLAAVALVDVPLLYLCRYLLIIRRGVIIVSGAGGGGGRLRRFRLGAAMAIIYMAMSTLFFLRIAPLTPWWGALAAWVMILLIVEAIFAFFFPYRCCFNETEGDVQNNPHV
IRIS_313-11862 MELITACALLLIAVHIAIVISCPEPLAAVALVDVPLLYLCRYLLIIRRGVIIVSGGGGGGRLRRFRLGAAMAIIYMAMSTLFFLRIAPLTPWWGALAAWVMILLIVEAIFAFFFPYRCCFNETEGDVQNNPHV
IRIS_313-11863 MELITACALLLIAVHIAIVISCPEPLAAVALVDVPLLYLCRYLLIIRRGVIIVSGAGGGGGRLRRFRLGAAMAIIYMAMSTLFFLRIAPLTPWWGALAAWVMILLIVEAIFAFFFPYRCCFNETEGDVQNNPHV
IRIS_313-11865 MELITACALLLIAVHIAIVISCPEPLAAVALVDVPLLYLCRYLLIIRRGVIIVSGAGAGGGRLRRFRLGAAMAIIYMAMSTLFFLRIAPLTPWWGALAAWVMILLIVEAIFAFFFPYRCCFNETEGDVQNNPHV
IRIS_313-11866 MELITACALLLIAVHIAIVISCPEPLAAVALVDVPLLYLYRYLLIIRRGVIIVSGAGGGGGRLRRFRLGAAMAIIYMAMSTLFFLRIAPLTPWWGALAAWVMILLIVEAIFAFFFPYRCCFNETEGDVQNNPHV
IRIS_313-11867 MELITACALLLIAVHIAIVISCPEPLAAVALVDVPLLYLCRYLLIIRRGVIIVSGGGGGGRLRRFRLGAAMAIIYMAMSTLFFLRIAPLTPWWGALAAWVMILLIVEAIFAFFFPYRCCFNETEGDVQNNPHV
IRIS_313-11868 MELITACALLLIAVHIAIVISCPEPLAAVALVDVPLLYLCRYLLIIRRGVIIVSGGGGGGRLRRFRLGAAMAIIYMAMSTLFFLRIAPLTPWWGALAAWVMILLIVEAIFAFFFPYRCCFNETEGDVQNNPHV
IRIS_313-11869 MELITACALLLIAVHIAIVISCPEPLAAVALVDVPLLYLCRYLLIIRRGVIIVSGAGAGGGRLRRFRLGAAMAIIYMAMSTLFFLRIAPLTPWWGALAAWVMILLIVEAIFAFFFPYRCCFNETEGDVQNNPHV
IRIS_313-11870 MELITACALLLIAVHIAIVISCPEPLAAVALVDVPLLYLYRYLLIIRRGVIIVSGAGGGGGRLRRFRLGAAMAIIYMAMSTLFFLRIAPLTPWWGALAAWVMILLIVEAIFAFFFPYRCCFNETEGDVQNNPHV
IRIS_313-11871 MELITACALLLIAVHIAIVISCPEPLAAVALVDVPLLYLCRYLLIIRRGVIIVSGGGGGGRLRRFRLGAAMAIIYMAMSTLFFLRIAPLTPWWGALAAWVMILLIVEAIFAFFFPYRCCFNETEGDVQNNPHV
IRIS_313-11872 MELITACALLLIAVHIAIVISCPEPLAAVALVDVPLLYLCRYLLIIRRGVIIVSGGGGGGRLRRFRLGAAMAIIYMAMSTLFFLRIAPLTPWWGALAAWVMILLIVEAIFAFFFPYRCCFNETEGDVQNNPHV
IRIS_313-11875 MELITACALLLIAVHIAIVISCPEPLAAVALVDVPLLYLYRYLLIIRRGVIIVSGAGGGGGRLRRFRLGAAMAIIYMAMSTLFFLRIAPLTPWWGALAAWVMILLIVEAIFAFFFPYRCCFNETEGDVQNNPHV
IRIS_313-11876 MELITACALLLIAVHIAIVISCPEPLAAVALVDVPLLYLCRYLLIIRRGVIIVSGGGGGGRLRRFRLGAAMAIIYMAMSTLFFLRIAPLTPWWGALAAWVMILLIVEAIFAFFFPYRCCFNETEGDVQNNPHV
IRIS_313-11877 MELITACALLLIAVHIAIVISCPEPLAAVALVDVPLLYLYRYLLIIRRGVIIVSGAGGGGGRLRRFRLGAAMAIIYMAMSTLFFLRIAPLTPWWGALAAWVMILLIVEAIFAFFFPYRCCFNETEGDVQNNPHV
IRIS_313-11878 MELITACALLLIAVHIAIVISCPEPLAAVALVDVPLLYLCRYLLIIRRGVIIVSGAGGGGGRLRRFRLGAAMAIIYMAMSTLFFLRIAPLTPWWGALAAWVMILLIVEAIFAFFFPYRCCFNETEGDVQNNPHV
IRIS_313-11881 MELITACALLLIAVHIAIVISCPEPLAAVALVDVPLLYLYRYLLIIRRGVIIVSGAGGGGGRLRRFRLGAAMAIIYMAMSTLFFLRIAPLTPWWGALAAWVMILLIVEAIFAFFFPYRCCFNETEGDVQNNPHV
IRIS_313-11882 MELITACALLLIAVHIAIVISCPEPLAAVALVDVPLLYLYRYLLIIRRGVIIVSGAGGGGGRLRRFRLGAAMAIIYMAMSTLFFLRIAPLTPWWGALAAWVMILLIVEAIFAFFFPYRCCFNETEGDVQNNPHV
IRIS_313-11884 MELITACALLLIAVHIAIVISCPEPLAAVALVDVPLLYLYRYLLIIRRGVIIVSGAGGGGGRLRRFRLGAAMAIIYMAMSTLFFLRIAPLTPWWGALAAWVMILLIVEAIFAFFFPYRCCFNETEGDVQNNPHV
IRIS_313-11885 MELITACALLLIAVHIAIVISCPEPLAAVALVDVPLLYLYRYLLIIRRGVIIVSGAGGGGGRLRRFRLGAAMAIIYMAMSTLFFLRIAPLTPWWGALAAWVMILLIVEAIFAFFFPYRCCFNETEGDVQNNPHV
IRIS_313-11886 MELITACALLLIAVHIAIVISCPEPLAAVALVDVPLLYLCRYLLIIRRGVIIVSGGGGGGRLRRFRLGAAMAIIYMAMSTLFFLRIAPLTPWWGALAAWVMILLIVEAIFAFFFPYRCCFNETEGDVQNNPHV
IRIS_313-11887 MELITACALLLIAVHIAIVISCPEPLAAVALVDVPLLYLCRYLLIIRRGVIIVSGGGGGGRLRRFRLGAAMAIIYMAMSTLFFLRIAPLTPWWGALAAWVMILLIVEAIFAFFFPYRCCFNETEGDVQNNPHV
IRIS_313-11888 MELITACALLLIAVHIAIVISCPEPLAAVAFVNVPLLYLCRYLLIIRRGVIIVSGGGGGGGRLRRFRLGAAMAVIYMAMSTLFFLRIAPLTPWWGALAAWVMILLIVEAIFAFFFPYRCCFNETEGDVQNNPHV
IRIS_313-11889 MELITACALLLIAVHIAIVISCPEPLAAVALVDVPLLYLCRYLLIIRRGVIIVSGGGGGGRLRRFRLGAAMAIIYMAMSTLFFLRIAPLTPWWGALAAWVMILLIVEAIFAFFFPYRCCFNETEGDVQNNPHV
IRIS_313-11890 MELITACALLLIAVHIAIVISCPEPLAAVALVDVPLLYLCRYLLIIRRGVIIVSGAGAGGGRLRRFRLGAAMAIIYMAMSTLFFLRIAPLTPWWGALAAWVMILLIVEAIFAFFFPYRCCFNETEGDVQNNPHV
IRIS_313-11892 MELITACALLLIAVHIAIVISCPEPLAAVALVDVPLLYLYRYLLIIRRGVIIVSGAGGGGGRLRRFRLGAAMAIIYMAMSTLFFLRIAPLTPWWGALAAWVMILLIVEAIFAFFFPYRCCFNETEGDVQNNPHV
IRIS_313-11893 MELITACALLLIAVHIAIVISCPEPLAAVALVDVPLLYLCRYLLIIRRGVIIVSGGGGGGRLRRFRLGAAMAIIYMAMSTLFFLRIAPLTPWWGALAAWVMILLIVEAIFAFFFPYRCCFNETEGDVQNNPHV
IRIS_313-11894 MELITACALLLIAVHIAIVISCPEPLAAVALVDVPLLYLCRYLLIIRRGVIIVSGGGGGGRLRRFRLGAAMAIIYMAMSTLFFLRIAPLTPWWGALAAWVMILLIVEAIFAFFFPYRCCFNETEGDVQNNPHV
IRIS_313-11895 MELITACALLLIAVHIAIVISCPEPLAAVALVDVPLLYLCRYLLIIRRGVIIVSGAGGGGRLRRFRLGAAMAIIYMAMSTLFFLRIAPLTPWWGALAAWVMILLIVEAIFAFFFPYRCCFNETEGDVQNNPHV
IRIS_313-11896 MELITACALLLIAVHIAIVISCPEPLAAVALVDVPLLYLYRYLLIIRRGVIIVSGAGGGGGRLRRFRLGAAMAIIYMAMSTLFFLRIAPLTPWWGALAAWVMILLIVEAIFAFFFPYRCCFNETEGDVQNNPHV
IRIS_313-11897 MELITACALLLIAVHIAIVISCPEPLAAVALVDVPLLYLCRYLLIIRRGVIIVSGAGGGGGRLRRFRLGAAMAIIYMAMSTLFFLRIAPLTPWWGALAAWVMILLIVEAIFAFFFPYRCCFNETEGDVQNNPHV
IRIS_313-11898 MELITACALLLIAVHIAIVISCPEPLAAVALVDVPLLYLYRYLLIIRRGVIIVSGAGGGGGRLRRFRLGAAMAIIYMAMSTLFFLRIAPLTPWWGALAAWVMILLIVEAIFAFFFPYRCCFNETEGDVQNNPHV
IRIS_313-11899 MELITACALLLIAVHIAIVISCPEPLAAVALVDVPLLYLYRYLLIIRRGVIIVSGAGGGGGRLRRFRLGAAMAIIYMAMSTLFFLRIAPLTPWWGALAAWVMILLIVEAIFAFFFPYRCCFNETEGDVQNNPHV
IRIS_313-11900 MELITACALLLIAVHIAIVISCPEPLAAVALVDVPLLYLYRYLLIIRRGVIIVSGAGAGGGRLRRFRLGAAMAIIYMAMSTLFFLRIAPLTPWWGALAAWVMILLIVEAIFAFFFPYRCCFNETEGDVQNNPHV
IRIS_313-11901 MELITACALLLIAVHIAIVISCPEPLAAVALVDVPLLYLCRYLLIIRRGVIIVSGGGAGGGRLRRFRLGAAMAIIYMAMSTLFFLRIAPLTPWWGALAAWVMILLIVEAIFAFFFPYRCCFNETEGDVQNNPHV
IRIS_313-11902 MELITACALLLIAVHIAIVISCPEPLAAVALVDVPLLYLCRYLLIIRRGVIIVSGGGGGGRLRRFRLGAAMAIIYMAMSTLFFLRIAPLTPWWGALAAWVMILLIVEAIFAFFFPYRCCFNETEGDVQNNPHV
IRIS_313-11903 MELITACALLLIAVHIAIVISCPEPLAAVALVDVPLLYLCRYLLIIRRGVIIVSGAGGGGGGRLRRFRLGAAMAIIYMAMSTLFFLRIAPLTPWWGALAAWVMILLIVEAIFAFFFPYRCCFNETEGDVQNNPHV
IRIS_313-11904 MELITACALLLIAVHIAIVISCPEPLAAVALVDVPLLYLCRYLLIIRRGVIIVSGGGGGGRLRRFRLGAAMAIIYMAMSTLFFLRIAPLTPWWGALAAWVMILLIVEAIFAFFFPYRCCFNETEGDVQNNPHV
IRIS_313-11905 MELITACALLLIAVHIAIVISCPEPLAAVALVDVPLLYLCRYLLIIRRGVIIVSGGGGGGRLRRFRLGAAMAIIYMAMSTLFFLRIAPLTPWWGALAAWVMILLIVEAIFAFFFPYRCCFNETEGDVQNNPHV
IRIS_313-11906 MELITACALLLIAVHIAIVISCPEPLAAVALVDVPLLYLCRYLLIIRRGVIIVSGAGAGGGRLRRFRLGAAMAIIYMAMSTLFFLRIAPLTPWWGALAAWVMILLIVEAIFAFFFPYRCCFNETEGDVQNNPHV
IRIS_313-11907 MELITACALLLIAVHIAIVISCPEPLAAVALVDVPLLYLCRYLLIIRRGVIIVSGAGAGGGRLRRFRLGAAMAIIYMAMSTLFFLRIAPLTPWWGALAAWVMILLIVEAIFAFFFPYRCCFNETEGDVQNNPHV

IRIS_313-11908 MELITACALLLIAVHIAIVISCPEPLAAVALVDVPLLYLCRYLLIIRRGVIIVSGAGAGGGRLRRFRLGAAMAIIYMAMSTLFFLRIAPLTPWWGALAAWVMILLIVEAIFAFFFPYRCCFNETEGDVQNNPHV
IRIS_313-11909 MELITACALLLIAVHIAIVISCPEPLAAVALVDVPLLYLCRYLLIIRRGVIIVSGAGAGGGRLRRFRLGAAMAIIYMAMSTLFFLRIAPLTPWWGALAAWVMILLIVEAIFAFFFPYRCCFNETEGDVQNNPHV
IRIS_313-11910 MELITACALLLIAVHIAIVISCPEPLAAVALVDVPLLYLCRYLLIIRRGVIIVSGGGGGGRLRRFRLGAAMAIIYMAMSTLFFLRIAPLTPWWGALAAWVMILLIVEAIFAFFFPYRCCFNETEGDVQNNPHV
IRIS_313-11911 MELITACALLLIAVHIAIVISCPEPLAAVALVDVPLLYLCRYLLIIRRGVIIVSGAGGGGGRLRRFRLGAAMAIIYMAMSTLFFLRIAPLTPWWGALAAWVMILLIVEAIFAFFFPYRCCFNETEGDVQNNPHV
IRIS_313-11912 MELITACALLLIAVHIAIVISCPEPLAAVALVDVPLLYLCRYLLIIRRGVIIVSGGGGGGRLRRFRLGAAMAIIYMAMSTLFFLRIAPLTPWWGALAAWVMILLIVEAIFAFFFPYRCCFNETEGDVQNNPHV
IRIS_313-11913 MELITACALLLIAVHIAIVISCPEPLAAVALVDVPLLYLCRYLLIIRRGVIIVSGAGGGGGRLRRFRLGAAMAIIYMAMSTLFFLRIAPLTPWWGALAAWVMILLIVEAIFAFFFPYRCCFNETEGDVQNNPHV
IRIS_313-11914 MELITACALLLIAVHIAIVISCPEPLAAVALVDVPLLYLCRYLLIIRRGVIIVSGAGAGGGRLRRFRLGAAMAIIYMAMSTLFFLRIAPLTPWWGALAAWVMILLIVEAIFAFFFPYRCCFNETEGDVQNNPHV
IRIS_313-11915 MELITACALLLIAVHIAIVISCPEPLAAVALVDVPLLYLCRYLLIIRRGVIIVSGAGAGGGRLRRFRLGAAMAIIYMAMSTLFFLRIAPLTPWWGALAAWVMILLIVEAIFAFFFPYRCCFNETEGDVQNNPHV
IRIS_313-11916 MELITACALLLIAVHIAIVISCPEPLAAVALVDVPLLYLCRYLLIIRRGVIIVSGAGAGGGRLRRFRLGAAMAIIYMAMSTLFFLRIAPLTPWWGALAAWVMILLIVEAIFAFFFPYRCCFNETEGDVQNNPHV
IRIS_313-11917 MELITACALLLIAVHIAIVISCPEPLAAVAFVNVPLLYLCRYLLIIRRGVIIVSGGGGGGGRLRRFRLGAAMAVIYMAMSTLFFLRIAPLTPWWGALAAWVMILLIVEAIFAFFFPYRCCFNETEGDVQNNPHV
IRIS_313-11918 MELITACALLLIAVHIAIVISCPEPLAAVALVDVPLLYLCRYLLIIRRGVIIVSGGGGGGRLRRFRLGAAMAIIYMAMSTLFFLRIAPLTPWWGALAAWVMILLIVEAIFAFFFPYRCCFNETEGDVQNNPHV
IRIS_313-11919 MELITACALLLIAVHIAIVISCPEPLAAVALVDVPLLYLCRYLLIIRRGVIIVSGGGGGGRLRRFRLGAAMAIIYMAMSTLFFLRIAPLTPWWGALAAWVMILLIVEAIFAFFFPYRCCFNETEGDVQNNPHV
IRIS_313-11920 MELITACALLLIAVHIAIVISCPEPLAAVALVDVPLLYLCRYLLIIRRGVIIVSGGGGGGRLRRFRLGAAMAIIYMAMSTLFFLRIAPLTPWWGALAAWVMILLIVEAIFAFFFPYRCCFNETEGDVQNNPHV
IRIS_313-11921 MELITACALLLIAVHIAIVISCPEPLAAVALVDVPLLYLYRYLLIIRRGVIIVSGAGGGGGRLRRFRLGAAMAIIYMAMSTLFFLRIAPLTPWWGALAAWVMILLIVEAIFAFFFPYRCCFNETEGDVQNNPHV
IRIS_313-11922 MELITACALLLIAVHIAIVISCPEPLAAVALVDVPLLYLCRYLLIIRRGVIIVSGAGAGGGRLRRFRLGAAMAIIYMAMSTLFFLRIAPLTPWWGALAAWVMILLIVEAIFAFFFPYRCCFNETEGDVQNNPHV
IRIS_313-11923 MELITACALLLIAVHIAIVISCPEPLAAVALVDVPLLYLCRYLLIIRRGVIIVSGAGGGGGRLRRFRLGAAMAIIYMAMSTLFFLRIAPLTPWWGALAAWVMILLIVEAIFAFFFPYRCCFNETEGDVQNNPHV
IRIS_313-11924 MELITACALLLIAVHIAIVISCPEPLAAVALVDVPLLYLCRYLLIIRRGVIIVSGAGGGGGRLRRFRLGAAMAIIYMAMSTLFFLRIAPLTPWWGALAAWVMILLIVEAIFAFFFPYRCCFNETEGDVQNNPHV
IRIS_313-11925 MELITACALLLIAVHIAIVISCPEPLAAVALVDVPLLYLCRYLLIIRRGVIIVSGAGAGGGRLRRFRLGAAMAIIYMAMSTLFFLRIAPLTPWWGALAAWVMILLIVEAIFAFFFPYRCCFNETEGDVQNNPHV
IRIS_313-11926 MELITACALLLIAVHIAIVISCPEPLAAVALVDVPLLYLCRYLLIIRRGVIIVSGAGAGGGRLRRFRLGAAMAIIYMAMSTLFFLRIAPLTPWWGALAAWVMILLIVEAIFAFFFPYRCCFNETEGDVQNNPHV
IRIS_313-11927 MELITACALLLIAVHIAIVISCPEPLAAVALVDVPLLYLCRYLLIIRRGVIIVSGGGGGGRLRRFRLGAAMAIIYMAMSTLFFLRIAPLTPWWGALAAWVMILLIVEAIFAFFFPYRCCFNETEGDVQNNPHV
IRIS_313-11928 MELITACALLLIAVHIAIVISCPEPLAAVALVDVPLLYLCRYLLIIRRGVIIVSGAGGGGGRLRRFRLGAAMAIIYMAMSTLFFLRIAPLTPWWGALAAWVMILLIVEAIFAFFFPYRCCFNETEGDVQNNPHV
IRIS_313-11929 MELITACALLLIAVHIAIVISCPEPLAAVALVDVPLLYLCRYLLIIRRGVIIVSGAGAGGGRLRRFRLGAAMAIIYMAMSTLFFLRIAPLTPWWGALAAWVMILLIVEAIFAFFFPYRCCFNETEGDVQNNPHV
IRIS_313-11930 MELITACALLLIAVHIAIVISCPEPLAAVALVDVPLLYLCRYLLIIRRGVIIVSGGGGGGRLRRFRLGAAMAIIYMAMSTLFFLRIAPLTPWWGALAAWVMILLIVEAIFAFFFPYRCCFNETEGDVQNNPHV
IRIS_313-11931 MELITACALLLIAVHIAIVISCPEPLAAVALVDVPLLYLYRYLLIIRRGVIIVSGAGGGGGRLRRFRLGAAMAIIYMAMSTLFFLRIAPLTPWWGALAAWVMILLIVEAIFAFFFPYRCCFNETEGDVQNNPHV
IRIS_313-11932 MELITACALLLIAVHIAIVISCPEPLAAVALVDVPLLYLCRYLLIIRRGVIIVSGAGAGGGRLRRFRLGAAMAIIYMAMSTLFFLRIAPLTPWWGALAAWVMILLIVEAIFAFFFPYRCCFNETEGDVQNNPHV
IRIS_313-11934 MELITACALLLIAVHIAIVISCPEPLAAVALVDVPLLYLCRYLLIIRRGVIIVSGAGAGGGRLRRFRLGAAMAIIYMAMSTLFFLRIAPLTPWWGALAAWVMILLIVEAIFAFFFPYRCCFNETEGDVQNNPHV
IRIS_313-11935 MELITACALLLIAVHIAIVISCPEPLAAVALVDVPLLYLYRYLLIIRRGVIIVSGAGGGGGRLRRFRLGAAMAIIYMAMSTLFFLRIAPLTPWWGALAAWVMILLIVEAIFAFFFPYRCCFNETEGDVQNNPHV
IRIS_313-11936 MELITACALLLIAVHIAIVISCPEPLAAVALVDVPLLYLCRYLLIIRRGVIIVSGAGGGGGGRLRRFRLGAAMAIIYMAMSTLFFLRIAPLTPWWGALAAWVMILLIVEAIFAFFFPYRCCFNETEGDVQNNPHV
IRIS_313-11937 MELITACALLLIAVHIAIVISCPEPLAAVALVDVPLLYLCRYLLIIRRGVIIVSGGGGGGRLRRFRLGAAMAIIYMAMSTLFFLRIAPLTPWWGALAAWVMILLIVEAIFAFFFPYRCCFNETEGDVQNNPHV
IRIS_313-11938 MELITACALLLIAVHIAIVISCPEPLAAVALVDVPLLYLCRYLLIIRRGVIIVSGGGGGGRLRRFRLGAAMAIIYMAMSTLFFLRIAPLTPWWGALAAWVMILLIVEAIFAFFFPYRCCFNETEGDVQNNPHV
IRIS_313-11939 MELITACALLLIAVHIAIVISCPEPLAAVALVDVPLLYLYRYLLIIRRGVIIVSGAGGGGGRLRRFRLGAAMAIIYMAMSTLFFLRIAPLTPWWGALAAWVMILLIVEAIFAFFFPYRCCFNETEGDVQNNPHV
IRIS_313-11940 MELITACALLLIAVHIAIVISCPEPLAAVALVDVPLLYLYRYLLIIRRGVIIVSGAGGGGGRLRRFRLGAAMAIIYMAMSTLFFLRIAPLTPWWGALAAWVMILLIVEAIFAFFFPYRCCFNETEGDVQNNPHV
IRIS_313-11941 MELITACALLLIAVHIAIVISCPEPLAAVALVDVPLLYLCRYLLIIRRGVIIVSGAGAGGGRLRRFRLGAAMAIIYMAMSTLFFLRIAPLTPWWGALAAWVMILLIVEAIFAFFFPYRCCFNETEGDVQNNPHV
IRIS_313-11943 MELITACALLLIAVHIAIVISCPEPLAAVALVDVPLLYLCRYLLIIRRGVIIVSGGGGGGRLRRFRLGAAMAIIYMAMSTLFFLRIAPLTPWWGALAAWVMILLIVEAIFAFFFPYRCCFNETEGDVQNNPHV
IRIS_313-11944 MELITACALLLIAVHIAIVISCPEPLAAVALVDVPLLYLCRYLLIIRRGVIIVSGGGGGGRLRRFRLGAAMAIIYMAMSTLFFLRIAPLTPWWGALAAWVMILLIVEAIFAFFFPYRCCFNETEGDVQNNPHV
IRIS_313-11945 MELITACALLLIAVHIAIVISCPEPLAAVALVDVPLLYLCRYLLIIRRGVIIVSGAGAGGGRLRRFRLGAAMAIIYMAMSTLFFLRIAPLTPWWGALAAWVMILLIVEAIFAFFFPYRCCFNETEGDVQNNPHV
IRIS_313-11946 MELITACALLLIAVHIAIVISCPEPLAAVALVDVPLLYLCRYLLIIRRGVIIVSGAGGGGGRLRRFRLGAAMAIIYMAMSTLFFLRIAPLTPWWGALAAWVMILLIVEAIFAFFFPYRCCFNETEGDVQNNPHV
IRIS_313-11947 MELITACALLLIAVHIAIVISCPEPLAAVALVDVPLLYLYRYLLIIRRGVIIVSGAGGGGGRLRRFRLGAAMAIIYMAMSTLFFLRIAPLTPWWGALAAWVMILLIVEAIFAFFFPYRCCFNETEGDVQNNPHV
IRIS_313-11948 MELITACALLLIAVHIAIVISCPEPLAAVALVDVPLLYLCRYLLIIRRGVIIVSGGGGGGRLRRFRLGAAMAIIYMAMSTLFFLRIAPLTPWWGALAAWVMILLIVEAIFAFFFPYRCCFNETEGDVQNNPHV
IRIS_313-11949 MELITACALLLIAVHIAIVISCPEPLAAVALVDVPLLYLYRYLLIIRRGVIIVSGAGGGGGRLRRFRLGAAMAIIYMAMSTLFFLRIAPLTPWWGALAAWVMILLIVEAIFAFFFPYRCCFNETEGDVQNNPHV
IRIS_313-11950 MELITACALLLIAVHIAIVISCPEPLAAVALVDVPLLYLYRYLLIIRRGVIIVSGAGGGGGRLRRFRLGAAMAIIYMAMSTLFFLRIAPLTPWWGALAAWVMILLIVEAIFAFFFPYRCCFNETEGDVQNNPHV
IRIS_313-11951 MELITACALLLIAVHIAIVISCPEPLAAVALVDVPLLYLYRYLLIIRRGVIIVSGAGGGGGRLRRFRLGAAMAIIYMAMSTLFFLRIAPLTPWWGALAAWVMILLIVEAIFAFFFPYRCCFNETEGDVQNNPHV
IRIS_313-11952 MELITACALLLIAVHIAIVISCPEPLAAVALVDVPLLYLYRYLLIIRRGVIIVSGAGGGGGRLRRFRLGAAMAIIYMAMSTLFFLRIAPLTPWWGALAAWVMILLIVEAIFAFFFPYRCCFNETEGDVQNNPHV
IRIS_313-11953 MELITACALLLIAVHIAIVISCPEPLAAVALVDVPLLYLCRYLLIIRRGVIIVSGAGAGGGRLRRFRLGAAMAIIYMAMSTLFFLRIAPLTPWWGALAAWVMILLIVEAIFAFFFPYRCCFNETEGDVQNNPHV
IRIS_313-11954 MELITACALLLIAVHIAIVISCPEPLAAVALVDVPLLYLCRYLLIIRRGVIIVSGAGAGGGRLRRFRLGAAMAIIYMAMSTLFFLRIAPLTPWWGALAAWVMILLIVEAIFAFFFPYRCCFNETEGDVQNNPHV
IRIS_313-11955 MELITACALLLIAVHIAIVISCPEPLAAVALVDVPLLYLCRYLLIIRRGVIIVSGAGAGGGRLRRFRLGAAMAIIYMAMSTLFFLRIAPLTPWWGALAAWVMILLIVEAIFAFFFPYRCCFNETEGDVQNNPHV
IRIS_313-11956 MELITACALLLIAVHIAIVISCPEPLAAVAFVNVPLLYLCRYLLIIRRGVIIVSGGGGGGGRLRRFRLGAAMAVIYMAMSTLFFLRIAPLTPWWGALAAWVMILLIVEAIFAFFFPYRCCFNETEGDVQNNPHV
IRIS_313-11957 MELITACALLLIAVHIAIVISCPEPLAAVALVDVPLLYLCRYLLIIRRGVIIVSGGGGAAAAGCAASASGQPWRSSTWPCRPSSSSALLLSRHGGAPSPPGS*SCSSSRPSSPSSSPTAAASTKPKAMSRIIPM
IRIS_313-11958 MELITACALLLIAVHIAIVISCPEPLAAVALVDVPLLYLCRYLLIIRRGVIIVSGGGGGGRLRRFRLGAAMAIIYMAMSTLFFLRIAPLTPWWGALAAWVMILLIVEAIFAFFFPYRCCFNETEGDVQNNPHV
IRIS_313-11959 MELITACALLLIAVHIAIVISCPEPLAAVALVDVPLLYLCRYLLIIRRGVIIVSGGGGGGRLRRFRLGAAMAIIYMAMSTLFFLRIAPLTPWWGALAAWVMILLIVEAIFAFFFPYRCCFNETEGDVQNNPHV
IRIS_313-11960 MELITACALLLIAVHIAIVISCPEPLAAVALVDVPLLYLCRYLLIIRRGVIIVSGGGGGGRLRRFRLGAAMAIIYMAMSTLFFLRIAPLTPWWGALAAWVMILLIVEAIFAFFFPYRCCFNETEGDVQNNPHV
IRIS_313-11961 MELITACALLLIAVHIAIVISCPEPLAAVALVDVPLLYLCRYLLIIRRGVIIVSGGGGGGRLRRFRLGAAMAIIYMAMSTLFFLRIAPLTPWWGALAAWVMILLIVEAIFAFFFPYRCCFNETEGDVQNNPHV
IRIS_313-11962 MELITACALLLIAVHIAIVISCPEPLAAVALVDVPLLYLCRYLLIIRRGVIIVSGAGAGGGRLRRFRLGAAMAIIYMAMSTLFFLRIAPLTPWWGALAAWVMILLIVEAIFAFFFPYRCCFNETEGDVQNNPHV
IRIS_313-11963 MELITACALLLIAVHIAIVISCPEPLAAVALVDVPLLYLCRYLLIIRRGVIIVSGAGAGGGRLRRFRLGAAMAIIYMAMSTLFFLRIAPLTPWWGALAAWVMILLIVEAIFAFFFPYRCCFNETEGDVQNNPHV
IRIS_313-11964 MELITACALLLIAVHIAIVISCPEPLAAVALVDVPLLYLCRYLLIIRRGVIIVSGGGGGGRLRRFRLGAAMAIIYMAMSTLFFLRIAPLTPWWGALAAWVMILLIVEAIFAFFFPYRCCFNETEGDVQNNPHV
IRIS_313-11965 MELITACALLLIAVHIAIVISCPEPLAAVALVDVPLLYLCRYLLIIRRGVIIVSGGGGGGRLRRFRLGAAMAIIYMAMSTLFFLRIAPLTPWWGALAAWVMILLIVEAIFAFFFPYRCCFNETEGDVQNNPHV
IRIS_313-11966 MELITACALLLIAVHIAIVISCPEPLAAVALVDVPLLYLCRYLLIIRRGVIIVSGGGGGGRLRRFRLGAAMAIIYMAMSTLFFLRIAPLTPWWGALAAWVMILLIVEAIFAFFFPYRCCFNETEGDVQNNPHV
IRIS_313-11967 MELITACALLLIAVHIAIVISCPEPLAAVALVDVPLLYLCRYLLIIRRGVIIVSGGGGGGRLRRFRLGAAMAIIYMAMSTLFFLRIAPLTPWWGALAAWVMILLIVEAIFAFFFPYRCCFNETEGDVQNNPHV
IRIS_313-11968 MELITACALLLIAVHIAIVISCPEPLAAVALVDVPLLYLCRYLLIIRRGVIIVSGAGAGGGRLRRFRLGAAMAIIYMAMSTLFFLRIAPLTPWWGALAAWVMILLIVEAIFAFFFPYRCCFNETEGDVQNNPHV
IRIS_313-11969 MELITACALLLIAVHIAIVISCPEPLAAVALVDVPLLYLCRYLLIIRRGVIIVSGAGAGGGRLRRFRLGAAMAIIYMAMSTLFFLRIAPLTPWWGALAAWVMILLIVEAIFAFFFPYRCCFNETEGDVQNNPHV
IRIS_313-11970 MELITACALLLIAVHIAIVISCPEPLAAVALVDVPLLYLCRYLLIIRRGVIIVSGAGGAAAAGCAASASGQPWRSSTWPCRPSSSSALLLSRHGGAPSPPGS*SCSSSRPSSPSSSPTAAASTKPKAMSRIIPM
IRIS_313-11971 MELITACALLLIAVHIAIVISCPEPLAAVALVDVPLLYLCRYLLIIRRGVIIVSGAGAGGGRLRRFRLGAAMAIIYMAMSTLFFLRIAPLTPWWGALAAWVMILLIVEAIFAFFFPYRCCFNETEGDVQNNPHV
IRIS_313-11973 MELITACALLLIAVHIAIVISCPEPLAAVALVDVPLLYLCRYLLIIRRGVIIVSGAGAGGGRLRRFRLGAAMAIIYMAMSTLFFLRIAPLTPWWGALAAWVMILLIVEAIFAFFFPYRCCFNETEGDVQNNPHV
IRIS_313-11974 MELITACALLLIAVHIAIVISCPEPLAAVALVDVPLLYLCRYLLIIRRGVIIVSGAGAGGGRLRRFRLGAAMAIIYMAMSTLFFLRIAPLTPWWGALAAWVMILLIVEAIFAFFFPYRCCFNETEGDVQNNPHV
IRIS_313-11975 MELITACALLLIAVHIAIVISCPEPLAAVALVDVPLLYLYRYLLIIRRGVIIVSGAGGGGGRLRRFRLGAAMAIIYMAMSTLFFLRIAPLTPWWGALAAWVMILLIVEAIFAFFFPYRCCFNETEGDVQNNPHV
IRIS_313-11976 MELITACALLLIAVHIAIVISCPEPLAAVALVDVPLLYLCRYLLIIRRGVIIVSGAGGAAAAGCAASASGQPWRSSTWPCRPSSSSALLLSRHGGAPSPPGS*SCSSSRPSSPSSSPTAAASTKPKAMSRIIPM
IRIS_313-11977 MELITACALLLIAVHIAIVISCPEPLAAVALVDVPLLYLCRYLLIIRRGVIIVSGGGGGGRLRRFRLGAAMAIIYMAMSTLFFLRIAPLTPWWGALAAWVMILLIVEAIFAFFFPYRCCFNETEGDVQNNPHV
IRIS_313-11978 MELITACALLLIAVHIAIVISCPEPLAAVALVDVPLLYLCRYLLIIRRGVIIVSGGGGGGRLRRFRLGAAMAIIYMAMSTLFFLRIAPLTPWWGALAAWVMILLIVEAIFAFFFPYRCCFNETEGDVQNNPHV
IRIS_313-11979 MELITACALLLIAVHIAIVISCPEPLAAVALVDVPLLYLCRYLLIIRRGVIIVSGAGAGGGRLRRFRLGAAMAIIYMAMSTLFFLRIAPLTPWWGALAAWVMILLIVEAIFAFFFPYRCCFNETEGDVQNNPHV
IRIS_313-11980 MELITACALLLIAVHIAIVISCPEPLAAVALVDVPLLYLCRYLLIIRRGVIIVSGGGGGGRLRRFRLGAAMAIIYMAMSTLFFLRIAPLTPWWGALAAWVMILLIVEAIFAFFFPYRCCFNETEGDVQNNPHV
IRIS_313-11981 MELITACALLLIAVHIAIVISCPEPLAAVALVDVPLLYLCRYLLIIRRGVIIVSGAGAGGGRLRRFRLGAAMAIIYMAMSTLFFLRIAPLTPWWGALAAWVMILLIVEAIFAFFFPYRCCFNETEGDVQNNPHV
IRIS_313-11982 MELITACALLLIAVHIAIVISCPEPLAAVAFVNVPLLYLCRYLLIIRRGVIIVSGGGGGGGRLRRFRLGAAMAVIYMAMSTLFFLRIAPLTPWWGALAAWVMILLIVEAIFAFFFPYRCCFNETEGDVQNNPHV
IRIS_313-11983 MELITACALLLIAVHIAIVISCPEPLAAVALVDVPLLYLCRYLLIIRRGVIIVSGAGAGGGRLRRFRLGAAMAIIYMAMSTLFFLRIAPLTPWWGALAAWVMILLIVEAIFAFFFPYRCCFNETEGDVQNNPHV
IRIS_313-11984 MELITACALLLIAVHIAIVISCPEPLAAVALVDVPLLYLCRYLLIIRRGVIIVSGAGAGGGRLRRFRLGAAMAIIYMAMSTLFFLRIAPLTPWWGALAAWVMILLIVEAIFAFFFPYRCCFNETEGDVQNNPHV
IRIS_313-11985 MELITACALLLIAVHIAIVISCPEPLAAVALVDVPLLYLCRYLLIIRRGVIIVSGGGGGGRLRRFRLGAAMAIIYMAMSTLFFLRIAPLTPWWGALAAWVMILLIVEAIFAFFFPYRCCFNETEGDVQNNPHV
IRIS_313-11986 MELITACALLLIAVHIAIVISCPEPLAAVALVDVPLLYLCRYLLIIRRGVIIVSGAGAGGGRLRRFRLGAAMAIIYMAMSTLFFLRIAPLTPWWGALAAWVMILLIVEAIFAFFFPYRCCFNETEGDVQNNPHV
IRIS_313-11987 MELITACALLLIAVHIAIVISCPEPLAAVALVDVPLLYLCRYLLIIRRGVIIVSGGGGGGRLRRFRLGAAMAIIYMAMSTLFFLRIAPLTPWWGALAAWVMILLIVEAIFAFFFPYRCCFNETEGDVQNNPHV
IRIS_313-11988 MELITACALLLIAVHIAIVISCPEPLAAVALVDVPLLYLYRYLLIIRRGVIIVSGAGGGGGRLRRFRLGAAMAIIYMAMSTLFFLRIAPLTPWWGALAAWVMILLIVEAIFAFFFPYRCCFNETEGDVQNNPHV
IRIS_313-11989 MELITACALLLIAVHIAIVISCPEPLAAVALVDVPLLYLCRYLLIIRRGVIIVSGAGAGGGRLRRFRLGAAMAIIYMAMSTLFFLRIAPLTPWWGALAAWVMILLIVEAIFAFFFPYRCCFNETEGDVQNNPHV
IRIS_313-11990 MELITACALLLIAVHIAIVISCPEPLAAVALVDVPLLYLCRYLLIIRRGVIIVSGGGGGGRLRRFRLGAAMAIIYMAMSTLFFLRIAPLTPWWGALAAWVMILLIVEAIFAFFFPYRCCFNETEGDVQNNPHV
IRIS_313-11991 MELITACALLLIAVHIAIVISCPEPLAAVALVDVPLLYLYRYLLIIRRGVIIVSGAGGGGGRLRRFRLGAAMAIIYMAMSTLFFLRIAPLTPWWGALAAWVMILLIVEAIFAFFFPYRCCFNETEGDVQNNPHV
IRIS_313-11992 MELITACALLLIAVHIAIVISCPEPLAAVALVDVPLLYLYRYLLIIRRGVIIVSGAGGGGGRLRRFRLGAAMAIIYMAMSTLFFLRIAPLTPWWGALAAWVMILLIVEAIFAFFFPYRCCFNETEGDVQNNPHV
IRIS_313-11993 MELITACALLLIAVHIAIVISCPEPLAAVALVDVPLLYLCRYLLIIRRGVIIVSGAGGGGGGRLRRFRLGAAMAIIYMAMSTLFFLRIAPLTPWWGALAAWVMILLIVEAIFAFFFPYRCCFNETEGDVQNNPHV
IRIS_313-11994 MELITACALLLIAVHIAIVISCPEPLAAVALVDVPLLYLCRYLLIIRRGVIIVSGTGGGGGRLRRFRLGAAMAIIYMAMSTLFFLRIAPLTPWWGALAAWVMILLIVEAIFAFFFPYRCCFNETEGDVQNNPHV
IRIS_313-11995 MELITACALLLIAVHIAIVISCPEPLAAVALVDVPLLYLCRYLLIIRRGVIIVSGAGGGGGRLRRFRLGAAMAIIYMAMSTLFFLRIAPLTPWWGALAAWVMILLIVEAIFAFFFPYRCCFNETEGDVQNNPHV
IRIS_313-11996 MELITACALLLIAVHIAIVISCPEPLAAVALVDVPLLYLCRYLLIIRRGVIIVSGAGAGGGRLRRFRLGAAMAIIYMAMSTLFFLRIAPLTPWWGALAAWVMILLIVEAIFAFFFPYRCCFNETEGDVQNNPHV
IRIS_313-11997 MELITACALLLIAVHIAIVISCPEPLAAVALVDVPLLYLCRYLLIIRRGVIIVSGAGAGGGRLRRFRLGAAMAIIYMAMSTLFFLRIAPLTPWWGALAAWVMILLIVEAIFAFFFPYRCCFNETEGDVQNNPHV
IRIS_313-11998 MELITACALLLIAVHIAIVISCPEPLAAVALVDVPLLYLCRYLLIIRRGVIIVSGAGAGGGRLRRFRLGAAMAIIYMAMSTLFFLRIAPLTPWWGALAAWVMILLIVEAIFAFFFPYRCCFNETEGDVQNNPHV
IRIS_313-11999 MELITACALLLIAVHIAIVISCPEPLAAVALVDVPLLYLYRYLLIIRRGVIIVSGAGAGGGRLRRFRLGAAMAIIYMAMSTLFFLRIAPLTPWWGALAAWVMILLIVEAIFAFFFPYRCCFNETEGDVQNNPHV
IRIS_313-12000 MELITACALLLIAVHIAIVISCPEPLAAVALVDVPLLYLCRYLLIIRRGVIIVSGGGGGGRLRRFRLGAAMAIIYMAMSTLFFLRIAPLTPWWGALAAWVMILLIVEAIFAFFFPYRCCFNETEGDVQNNPHV
IRIS_313-12002 MELITACALLLIAVHIAIVISCPEPLAAVAFVNVPLLYLCRYLLIIRRGVIIVSGGGGGGGRLRRFRLGAAMAVIYMAMSTLFFLRIAPLTPWWGALAAWVMILLIVEAIFAFFFPYRCCFNETEGDVQNNPHV
IRIS_313-12003 MELITACALLLIAVHIAIVISCPEPLAAVALVDVPLLYLCRYLLIIRRGVIIVSGAGAGGGRLRRFRLGAAMAIIYMAMSTLFFLRIAPLTPWWGALAAWVMILLIVEAIFAFFFPYRCCFNETEGDVQNNPHV
IRIS_313-12004 MELITACALLLIAVHIAIVISCPEPLAAVALVDVPLLYLCRYLLIIRRGVIIVSGAGAGGGRLRRFRLGAAMAIIYMAMSTLFFLRIAPLTPWWGALAAWVMILLIVEAIFAFFFPYRCCFNETEGDVQNNPHV
IRIS_313-12005 MELITACALLLIAVHIAIVISCPEPLAAVALVDVPLLYLCRYLLIIRRGVIIVSGGGGGGRLRRFRLGAAMAIIYMAMSTLFFLRIAPLTPWWGALAAWVMILLIVEAIFAFFFPYRCCFNETEGDVQNNPHV
IRIS_313-12006 MELITACALLLIAVHIAIVISCPEPLAAVALVDVPLLYLYRYLLIIRRGVIIVSGAGGGGGRLRRFRLGAAMAIIYMAMSTLFFLRIAPLTPWWGALAAWVMILLIVEAIFAFFFPYRCCFNETEGDVQNNPHV
IRIS_313-12007 MELITACALLLIAVHIAIVISCPEPLAAVALVDVPLLYLCRYLLIIRRGVIIVSGGGGGGRLRRFRLGAAMAIIYMAMSTLFFLRIAPLTPWWGALAAWVMILLIVEAIFAFFFPYRCCFNETEGDVQNNPHV
IRIS_313-12009 MELITACALLLIAVHIAIVISCPEPLAAVALVDVPLLYLCRYLLIIRRGVIIVSGGGAGGGRLRRFRLGAAMAIIYMAMSTLFFLRIAPLTPWWGALAAWVMILLIVEAIFAFFFPYRCCFNETEGDVQNNPHV
IRIS_313-12010 MELITACALLLIAVHIAIVISCPEPLAAVALVDVPLLYLYRYLLIIRRGVIIVSGAGGGGGRLRRFRLGAAMAIIYMAMSTLFFLRIAPLTPWWGALAAWVMILLIVEAIFAFFFPYRCCFNETEGDVQNNPHV
IRIS_313-12011 MELITACALLLIAVHIAIVISCPEPLAAVALVDVPLLYLCRYLLIIRRGVIIVSGGGGGGRLRRFRLGAAMAIIYMAMSTLFFLRIAPLTPWWGALAAWVMILLIVEAIFAFFFPYRCCFNETEGDVQNNPHV
IRIS_313-12012 MELITACALLLIAVHIAIVISCPEPLAAVALVDVPLLYLCRYLLIIRRGVIIVSGAGAGGGRLRRFRLGAAMAIIYMAMSTLFFLRIAPLTPWWGALAAWVMILLIVEAIFAFFFPYRCCFNETEGDVQNNPHV
IRIS_313-12013 MELITACALLLIAVHIAIVISCPEPLAAVALVDVPLLYLCRYLLIIRRGVIIVSGAGAGGGRLRRFRLGAAMAIIYMAMSTLFFLRIAPLTPWWGALAAWVMILLIVEAIFAFFFPYRCCFNETEGDVQNNPHV
IRIS_313-12014 MELITACALLLIAVHIAIVISCPEPLAAVALVDVPLLYLCRYLLIIRRGVIIVSGAGAGGGRLRRFRLGAAMAIIYMAMSTLFFLRIAPLTPWWGALAAWVMILLIVEAIFAFFFPYRCCFNETEGDVQNNPHV
IRIS_313-12015 MELITACALLLIAVHIAIVISCPEPLAAVALVDVPLLYLCRYLLIIRRGVIIVSGAGAGGGRLRRFRLGAAMAIIYMAMSTLFFLRIAPLTPWWGALAAWVMILLIVEAIFAFFFPYRCCFNETEGDVQNNPHV
IRIS_313-12016 MELITACALLLIAVHIAIVISCPEPLAAVALVDVPLLYLCRYLLIIRRGVIIVSGGGGGGRLRRFRLGAAMAIIYMAMSTLFFLRIAPLTPWWGALAAWVMILLIVEAIFAFFFPYRCCFNETEGDVQNNPHV
IRIS_313-12017 MELITACALLLIAVHIAIVISCPEPLAAVALVDVPLLYLCRYLLIIRRGVIIVSGAGGAAAAGCAASASGQPWRSSTWPCRPSSSSALLLSRHGGAPSPPGS*SCSSSRPSSPSSSPTAAASTKPKAMSRIIPM
IRIS_313-12018 MELITACALLLIAVHIAIVISCPEPLAAVALVDVPLLYLCRYLLIIRRGVIIVSGAGAGGGRLRRFRLGAAMAIIYMAMSTLFFLRIAPLTPWWGALAAWVMILLIVEAIFAFFFPYRCCFNETEGDVQNNPHV
IRIS_313-12021 MELITACALLLIAVHIAIVISCPEPLAAVALVDVPLLYLCRYLLIIRRGVIIVSGAGAGGGRLRRFRLGAAMAIIYMAMSTLFFLRIAPLTPWWGALAAWVMILLIVEAIFAFFFPYRCCFNETEGDVQNNPHV
IRIS_313-12024 MELITACALLLIAVHIAIVISCPEPLAAVALVDVPLLYLCRYLLIIRRGVIIVSGGGGGGRLRRFRLGAAMAIIYMAMSTLFFLRIAPLTPWWGALAAWVMILLIVEAIFAFFFPYRCCFNETEGDVQNNPHV
IRIS_313-12027 MELITACALLLIAVHIAIVISCPEPLAAVALVDVPLLYLCRYLLIIRRGVIIVSGGGGGGRLRRFRLGAAMAIIYMAMSTLFFLRIAPLTPWWGALAAWVMILLIVEAIFAFFFPYRCCFNETEGDVQNNPHV
IRIS_313-12028 MELITACALLLIAVHIAIVISCPEPLAAVALVDVPLLYLCRYLLIIRRGVIIVSGGGGGGRLRRFRLGAAMAIIYMAMSTLFFLRIAPLTPWWGALAAWVMILLIVEAIFAFFFPYRCCFNETEGDVQNNPHV
IRIS_313-12029 MELITACALLLIAVHIAIVISCPEPLAAVALVDVPLLYLCRYLLIIRRGVIIVSGAGAGGGRLRRFRLGAAMAIIYMAMSTLFFLRIAPLTPWWGALAAWVMILLIVEAIFAFFFPYRCCFNETEGDVQNNPHV
IRIS_313-12030 MELITACALLLIAVHIAIVISCPEPLAAVALVDVPLLYLYRYLLIIRRGVIIVSGAGGGGGRLRRFRLGAAMAIIYMAMSTLFFLRIAPLTPWWGALAAWVMILLIVEAIFAFFFPYRCCFNETEGDVQNNPHV
IRIS_313-12031 MELITACALLLIAVHIAIVISCPEPLAAVALVDVPLLYLCRYLLIIRRGVIIVSGAGAGGGRLRRFRLGAAMAIIYMAMSTLFFLRIAPLTPWWGALAAWVMILLIVEAIFAFFFPYRCCFNETEGDVQNNPHV
IRIS_313-12033 MELITACALLLIAVHIAIVISCPEPLAAVALVDVPLLYLCRYLLIIRRGVIIVSGAGGGGGRLRRFRLGAAMAIIYMAMSTLFFLRIAPLTPWWGALAAWVMILLIVEAIFAFFFPYRCCFNETEGDVQNNPHV
IRIS_313-12036 MELITACALLLIAVHIAIVISCPEPLAAVALVDVPLLYLCRYLLIIRRGVIIVSGAGAGGGRLRRFRLGAAMAIIYMVMSTLFFLRIAPLTPWWGALAAWVMILLIVEAIFAFFFPYRCCFNETEGDVQNNPHV
IRIS_313-12037 MELITACALLLIAVHIAIVISCPEPLAAVALVDVPLLYLYRYLLIIRRGVIIVSGAGGGGGRLRRFRLGAAMAIIYMAMSTLFFLRIAPLTPWWGALAAWVMILLIVEAIFAFFFPYRCCFNETEGDVQNNPHV
IRIS_313-12038 MELITACALLLIAVHIAIVISCPEPLAAVALVDVPLLYLYRYLLIIRRGVIIVSGAGGGGGRLRRFRLGAAMAIIYMAMSTLFFLRIAPLTPWWGALAAWVMILLIVEAIFAFFFPYRCCFNETEGDVQNNPHV
IRIS_313-12039 MELITACALLLIAVHIAIVISCPEPLAAVALVDVPLLYLYRYLLIIRRGVIIVSGAGGGGGRLRRFRLGAAMAIIYMAMSTLFFLRIAPLTPWWGALAAWVMILLIVEAIFAFFFPYRCCFNETEGDVQNNPHV
IRIS_313-12040 MELITACALLLIAVHIAIVISCPEPLAAVALVDVPLLYLCRYLLIIRRGVIIVSGGGGGGRLRRFRLGAAMAIIYMAMSTLFFLRIAPLTPWWGALAAWVMILLIVEAIFAFFFPYRCCFNETEGDVQNNPHV
IRIS_313-12041 MELITACALLLIAVHIAIVISCPEPLAAVALVDVPLLYLCRYLLIIRRGVIIVSGGGGGGRLRRFRLGAAMAIIYMAMSTLFFLRIAPLTPWWGALAAWVMILLIVEAIFAFFFPYRCCFNETEGDVQNNPHV
IRIS_313-12042 MELITACALLLIAVHIAIVISCPEPLAAVAFVNVPLLYLCRYLLIIRRGVIIVSGGSGGGGRLRRFRLGAAMAVIYMAMSTLFFLRIAPLTPWWGALAAWVMILLIVEAIFAFFFPYRCCFNETEGDVQNNPHV
IRIS_313-12043 MELITACALLLIAVHIAIVISCPEPLAAVALVDVPLLYLCRYLLIIRRGVIIVSGGGGGGRLRRFRLGAAMAIIYMAMSTLFFLRIAPLTPWWGALAAWVMILLIVEAIFAFFFPYRCCFNETEGDVQNNPHV
IRIS_313-12044 MELITACALLLIAVHIAIVISCPEPLAAVALVDVPLLYLYRYLLIIRRGVIIVSGAGGGGGRLRRFRLGAAMAIIYMAMSTLFFLRIAPLTPWWGALAAWVMILLIVEAIFAFFFPYRCCFNETEGDVQNNPHV

```
IRIS_313-12045 MELITACALLLIAVHIAIVISCPEPLAAVALVDVPLLYLCRYLLIIRRGVIIVSGAGGGGGRLRRFRLGAAMAIIYMAMSTLFFLRIAPLTPWWGALAAWVMILLIVEAIFAFFFPYRCCFNETEGDVQNNPHV
IRIS_313-12046 MELITACALLLIAVHIAIVISCPEPLAAVALVDVPLLYLCRYLLIIRRGVIIVSGGGGGGRLRRFRLGAAMAIIYMAMSTLFFLRIAPLTPWWGALAAWVMILLIVEAIFAFFFPYRCCFNETEGDVQNNPHV
IRIS_313-12047 MELITACALLLIAVHIAIVISCPEPLAAVALVDVPLLYLCRYLLIIRRGVIIVSGGGGGGRLRRFRLGAAMAIIYMAMSTLFFLRIAPLTPWWGALAAWVMILLIVEAIFAFFFPYRCCFNETEGDVQNNPHV
IRIS_313-12048 MELITACALLLIAVHIAIVISCPEPLAAVALVDVPLLYLCRYLLIIRRGVIIVSGAGAGGGRLRRFRLGAAMAIIYMAMSTLFFLRIAPLTPWWGALAAWVMILLIVEAIFAFFFPYRCCFNETEGDVQNNPHV
IRIS_313-12049 MELITACALLLIAVHIAIVISCPEPLAAVALVDVPLLYLYRYLLIIRRGVIIVSGAGGGGGRLRRFRLGAAMAIIYMAMSTLFFLRIAPLTPWWGALAAWVMILLIVEAIFAFFFPYRCCFNETEGDVQNNPHV
IRIS_313-12050 MELITACALLLIAVHIAIVISCPEPLAAVALVDVPLLYLCRYLLIIRRGVIIVSGAGGAAAAGCAASASGQPWRSSTWPCRPSSSSALLLSRHGGAPSPPGS*SCSSSRPSSPSSSPTAAASTKPKAMSRIIPM
IRIS_313-12051 MELITACALLLIAVHIAIVISCPEPLAAVALVDVPLLYLYRYLLIIRRGVIIVSGAGGGGGRLRRFRLGAAMAIIYMAMSTLFFLRIAPLTPWWGALAAWVMILLIVEAIFAFFFPYRCCFNETEGDVQNNPHV
IRIS_313-12052 MELITACALLLIAVHIAIVISCPEPLAAVALVDVPLLYLCRYLLIIRRGVIIVSGAGGGGGRLRRFRLGAAMAIIYMAMSTLFFLRIAPLTPWWGALAAWVMILLIVEAIFAFFFPYRCCFNETEGDVQNNPHV
IRIS_313-12053 MELITACALLLIAVHIAIVISCPEPLAAVALVDVPLLYLCRYLLIIRRGVIIVSGGGGGGRLRRFRLGAAMAIIYMAMSTLFFLRIAPLTPWWGALAAWVMILLIVEAIFAFFFPYRCCFNETEGDVQNNPHV
IRIS_313-12054 MELITACALLLIAVHIAIVISCPEPLAAVALVDVPLLYLCRYLLIIRRGVIIVSGAGAGGGRLRRFRLGAAMAIIYMAMSTLFFLRIAPLTPWWGALAAWVMILLIVEAIFAFFFPYRCCFNETEGDVQNNPHV
IRIS_313-12055 MELITACALLLIAVHIAIVISCPEPLAAVAFVNVPLLYLCRYLLIIRRGVIIVSGGGGGGGRLRRFRLGAAMAVIYMAMSTLFFLRIAPLTPWWGALAAWVMILLIVEAIFAFFFPYRCCFNETEGDVQNNPHV
IRIS_313-12057 MELITACALLLIAVHIAIVISCPEPLAAVALVDVPLLYLCRYLLIIRRGVIIVSGAGGGGGGRLRRFRLGAAMAIIYMAMSTLFFLRIAPLTPWWGALAAWVMILLIVEAIFAFFFPYRCCFNETEGDVQNNPHV
IRIS_313-12058 MELITACALLLIAVHIAIVISCPEPLAAVALVDVPLLYLCRYLLIIRRGVIIVSGGGAGGGRLRRFRLGAAMAIIYMAMSTLFFLRIAPLTPWWGALAAWVMILLIVEAIFAFFFPYRCCFNETEGDVQNNPHV
IRIS_313-12059 MELITACALLLIAVHIAIVISCPEPLAAVALVDVPLLYLCRYLLIIRRGVIIVSGAGGGGGRLRRFRLGAAMAIIYMAMSTLFFLRIAPLTPWWGALAAWVMILLIVEAIFAFFFPYRCCFNETEGDVQNNPHV
IRIS_313-12060 MELITACALLLIAVHIAIVISCPEPLAAVALVDVPLLYLCRYLLIIRRGVIIVSGAGGGGGRLRRFRLGAAMAIIYMAMSTLFFLRIAPLTPWWGALAAWVMILLIVEAIFAFFFPYRCCFNETEGDVQNNPHV
IRIS_313-12061 MELITACALLLIAVHIAIVISCPEPLAAVALVDVPLLYLCRYLLIIRRGVIIVSGAGAGGGRLRRFRLGAAMAIIYMAMSTLFFLRIAPLTPWWGALAAWVMILLIVEAIFAFFFPYRCCFNETEGDVQNNPHV
IRIS_313-12063 MELITACALLLIAVHIAIVISCPEPLAAVALVDVPLLYLYRYLLIIRRGVIIVSGAGAGGGRLRRFRLGAAMAIIYMAMSTLFFLRIAPLTPWWGALAAWVMILLIVEAIFAFFFPYRCCFNETEGDVQNNPHV
IRIS_313-12065 MELITACALLLIAVHIAIVISCPEPLAAVALVDVPLLYLCRYLLIIRRGVIIVSGAGGAAAAGCAASASGQPWRSSTWPCRPSSSSALLLSRHGGAPSPPGS*SCSSSRPSSPSSSPTAAASTKPKAMSRIIPM
IRIS_313-12066 MELITACALLLIAVHIAIVISCPEPLAAVALVDVPLLYLCRYLLIIRRGVIIVSGGGGGGRLRRFRLGAAMAIIYMAMSTLFFLRIAPLTPWWGALAAWVMILLIVEAIFAFFFPYRCCFNETEGDVQNNPHV
IRIS_313-12067 MELITACALLLIAVHIAIVISCPEPLAAVALVDVPLLYLCRYLLIIRRGVIIVSGGGGGGRLRRFRLGAAMAIIYMAMSTLFFLRIAPLTPWWGALAAWVMILLIVEAIFAFFFPYRCCFNETEGDVQNNPHV
IRIS_313-12068 MELITACALLLIAVHIAIVISCPEPLAAVALVDVPLLYLCRYLLIIRRGVIIVSGAGAGGGRLRRFRLGAAMAIIYMAMSTLFFLRIAPLTPWWGALAAWVMILLIVEAIFAFFFPYRCCFNETEGDVQNNPHV
IRIS_313-12069 MELITACALLLIAVHIAIVISCPEPLAAVALVDVPLLYLCRYLLIIRRGVIIVSGAGAGGGRLRRFRLGAAMAIIYMAMSTLFFLRIAPLTPWWGALAAWVMILLIVEAIFAFFFPYRCCFNETEGDVQNNPHV
IRIS_313-12070 MELITACALLLIAVHIAIVISCPEPLAAVALVDVPLLYLYRYLLIIRRGVIIVSGAGGGGGRLRRFRLGAAMAIIYMAMSTLFFLRIAPLTPWWGALAAWVMILLIVEAIFAFFFPYRCCFNETEGDVQNNPHV
IRIS_313-12071 MELITACALLLIAVHIAIVISCPEPLAAVALVDVPLLYLCRYLLIIRRGVIIVSGAGGGGGRLRRFRLGAAMAIIYMAMSTLFFLRIAPLTPWWGALAAWVMILLIVEAIFAFFFPYRCCFNETEGDVQNNPHV
IRIS_313-12072 MELITACALLLIAVHIAIVISCPEPLAAVALVDVPLLYLCRYLLIIRRGVIIVSGAGGGGGGRLRRFRLGAAMAIIYMAMSTLFFLRIAPLTPWWGALAAWVMILLIVEAIFAFFFPYRCCFNETEGDVQNNPHV
IRIS_313-12073 MELITACALLLIAVHIAIVISCPEPLAAVALVDVPLLYLCRYLLIIRRGVIIVSGAGAGGGRLRRFRLGAAMAIIYMAMSTLFFLRIAPLTPWWGALAAWVMILLIVEAIFAFFFPYRCCFNETEGDVQNNPHV
IRIS_313-12074 MELITACALLLIAVHIAIVISCPEPLAAVALVDVPLLYLCRYLLIIRRGVIIVSGAGAGGGRLRRFRLGAAMAIIYMAMSTLFFLRIAPLTPWWGALAAWVMILLIVEAIFAFFFPYRCCFNETEGDVQNNPHV
IRIS_313-12076 MELITACALLLIAVHIAIVISCPEPLAAVALVDVPLLYLYRYLLIIRRGVIIVSGAGGGGGRLRRFRLGAAMAIIYMAMSTLFFLRIAPLTPWWGALAAWVMILLIVEAIFAFFFPYRCCFNETEGDVQNNPHV
IRIS_313-12077 MELITACALLLIAVHIAIVISCPEPLAAVALVDVPLLYLCRYLLIIRRGVIIVSGAGAGGGRLRRFRLGAAMAIIYMAMSTLFFLRIAPLTPWWGALAAWVMILLIVEAIFAFFFPYRCCFNETEGDVQNNPHV
IRIS_313-12078 MELITACALLLIAVHIAIVISCPEPLAAVALVDVPLLYLYRYLLIIRRGVIIVSGAGGGGGRLRRFRLGAAMAIIYMAMSTLFFLRIAPLTPWWGALAAWVMILLIVEAIFAFFFPYRCCFNETEGDVQNNPHV
IRIS_313-12079 MELITACALLLIAVHIAIVISCPEPLAAVALVDVPLLYLCRYLLIIRRGVIIVSGGGGGGRLRRFRLGAAMAIIYMAMSTLFFLRIAPLTPWWGALAAWVMILLIVEAIFAFFFPYRCCFNETEGDVQNNPHV
IRIS_313-12080 MELITACALLLIAVHIAIVISCPEPLAAVALVDVPLLYLCRYLLIIRRGVIIVSGGGGGGRLRRFRLGAAMAIIYMAMSTLFFLRIAPLTPWWGALAAWVMILLIVEAIFAFFFPYRCCFNETEGDVQNNPHV
IRIS_313-12081 MELITACALLLIAVHIAIVISCPEPLAAVALVDVPLLYLCRYLLIIRRGVIIVSGGGGGGRLRRFRLGAAMAIIYMAMSTLFFLRIAPLTPWWGALAAWVMILLIVEAIFAFFFPYRCCFNETEGDVQNNPHV
IRIS_313-12082 MELITACALLLIAVHIAIVISCPEPLAAVALVDVPLLYLCRYLLIIRRGVIIVSGAGGAAAAGCAASASGQPWRSSTWPCRPSSSSALLLSRHGGAPSPPGS*SCSSSRPSSPSSSPTAAASTKPKAMSRIIPM
IRIS_313-12083 MELITACALLLIAVHIAIVISCPEPLAAVALVDVPLLYLCRYLLIIRRGVIIVSGAGGAAAAGCAASASGQPWRSSTWPCRPSSSSALLLSRHGGAPSPPGS*SCSSSRPSSPSSSPTAAASTKPKAMSRIIPM
IRIS_313-12093 MELITACALLLIAVHIAIVISCPEPLAAVALVDVPLLYLYRYLLIIRRGVIIVSGAGGGGGRLRRFRLGAAMAIIYMAMSTLFFLRIAPLTPWWGALAAWVMILLIVEAIFAFFFPYRCCFNETEGDVQNNPHV
IRIS_313-12094 MELITACALLLIAVHIAIVISCPEPLAAVALVDVPLLYLCRYLLIIRRGVIIVSGAGAGGGRLRRFRLGAAMAIIYMAMSTLFFLRIAPLTPWWGALAAWVMILLIVEAIFAFFFPYRCCFNETEGDVQNNPHV
IRIS_313-12096 MELITACALLLIAVHIAIVISCPEPLAAVALVDVPLLYLCRYLLIIRRGVIIVSGAGAGGGRLRRFRLGAAMAIIYMAMSTLFFLRIAPLTPWWGALAAWVMILLIVEAIFAFFFPYRCCFNETEGDVQNNPHV
IRIS_313-12097 MELITACALLLIAVHIAIVISCPEPLAAVALVDVPLLYLCRYLLIIRRGVIIVSGGGGGGRLRRFRLGAAMAIIYMAMSTLFFLRIAPLTPWWGALAAWVMILLIVEAIFAFFFPYRCCFNETEGDVQNNPHV
IRIS_313-12101 MELITACALLLIAVHIAIVISCPEPLAAVALVDVPLLYLCRYLLIIRRGVIIVSGAGAGGGRLRRFRLGAAMAIIYMAMSTLFFLRIAPLTPWWGALAAWVMILLIVEAIFAFFFPYRCCFNETEGDVQNNPHV
IRIS_313-12102 MELITACALLLIAVHIAIVISCPEPLAAVALVDVPLLYLCRYLLIIRRGVIIVSGAGAGGGRLRRFRLGAAMAIIYMAMSTLFFLRIAPLTPWWGALAAWVMILLIVEAIFAFFFPYRCCFNETEGDVQNNPHV
IRIS_313-12108 MELITACALLLIAVHIAIVISCPEPLAAVALVDVPLLYLCRYLLIIRRGVIIVSGAGGAAAAGCAASASGQPWRSSTWPCRPSSSSALLLSRHGGAPSPPGS*SCSSSRPSSPSSSPTAAASTKPKAMSRIIPM
IRIS_313-12109 MELITACALLLIAVHIAIVISCPEPLAAVALVDVPLLYLYRYLLIIRRGVIIVSGAGGGGGRLRRFRLGAAMAIIYMAMSTLFFLRIAPLTPWWGALAAWVMILLIVEAIFAFFFPYRCCFNETEGDVQNNPHV
IRIS_313-12118 MELITACALLLIAVHIAIVISCPEPLAAVALVDVPLLYLCRYLLIIRRGVIIVSGAGAGGGRLRRFRLGAAMAIIYMAMSTLFFLRIAPLTPWWGALAAWVMILLIVEAIFAFFFPYRCCFNETEGDVQNNPHV
IRIS_313-12121 MELITACALLLIAVHIAIVISCPEPLAAVALVDVPLLYLCRYLLIIRRGVIIVSGGGGGGRLRRFRLGAAMAIIYMAMSTLFFLRIAPLTPWWGALAAWVMILLIVEAIFAFFFPYRCCFNETEGDVQNNPHV
IRIS_313-12127 MELITACALLLIAVHIAIVISCPEPLAAVALVDVPLLYLCRYLLIIRRGVIIVSGGGGGGRLRRFRLGAAMAIIYMAMSTLFFLRIAPLTPWWGALAAWVMILLIVEAIFAFFFPYRCCFNETEGDVQNNPHV
IRIS_313-12128 MELITACALLLIAVHIAIVISCPEPLAAVALVDVPLLYLCRYLLIIRRGVIIVSGGGGGGRLRRFRLGAAMAIIYMAMSTLFFLRIAPLTPWWGALAAWVMILLIVEAIFAFFFPYRCCFNETEGDVQNNPHV
IRIS_313-12129 MELITACALLLIAVHIAIVISCPEPLAAVALVDVPLLYLCRYLLIIRRGVIIVSGAGGGGGRLRRFRLGAAMAIIYMAMSTLFFLRIAPLTPWWGALAAWVMILLIVEAIFAFFFPYRCCFNETEGDVQNNPHV
IRIS_313-12130 MELITACALLLIAVHIAIVISCPEPLAAVALVDVPLLYLCRYLLIIRRGVIIVSGGGGGGRLRRFRLGAAMAIIYMAMSTLFFLRIAPLTPWWGALAAWVMILLIVEAIFAFFFPYRCCFNETEGDVQNNPHV
IRIS_313-12131 MELITACALLLIAVHIAIVISCPEPLAAVALVDVPLLYLCRYLLIIRRGVIIVSGGGGGGRLRRFRLGAAMAIIYMAMSTLFFLRIAPLTPWWGALAAWVMILLIVEAIFAFFFPYRCCFNETEGDVQNNPHV
IRIS_313-12133 MELITACALLLIAVHIAIVISCPEPLAAVALVDVPLLYLCRYLLIIRRGVIIVSGGGGGGRLRRFRLGAAMAIIYMAMSTLFFLRIAPLTPWWGALAAWVMILLIVEAIFAFFFPYRCCFNETEGDVQNNPHV
IRIS_313-12134 MELITACALLLIAVHIAIVISCPEPLAAVALVDVPLLYLYRYLLIIRRGVIIVSGAGAGGGRLRRFRLGAAMAIIYMAMSTLFFLRIAPLTPWWGALAAWVMILLIVEAIFAFFFPYRCCFNETEGDVQNNPHV
IRIS_313-12135 MELITACALLLIAVHIAIVISCPEPLAAVALVDVPLLYLCRYLLIIRRGVIIVSGGGGGGRLRRFRLGAAMAIIYMAMSTLFFLRIAPLTPWWGALAAWVMILLIVEAIFAFFFPYRCCFNETEGDVQNNPHV
IRIS_313-12138 MELITACALLLIAVHIAIVISCPEPLAAVALVDVPLLYLYRYLLIIRRGVIIVSGAGGGGGRLRRFRLGAAMAIIYMAMSTLFFLRIAPLTPWWGALAAWVMILLIVEAIFAFFFPYRCCFNETEGDVQNNPHV
IRIS_313-12139 MELITACALLLIAVHIAIVISCPEPLAAVAFVNVPLLYLCRYLLIIRRGVIIVSGGGGGGGRLRRFRLGAAMAVIYMAMSTLFFLRIAPLTPWWGALAAWVMILLIVEAIFAFFFPYRCCFNETEGDVQNNPHV
IRIS_313-12141 MELITACALLLIAVHIAIVISCPEPLAAVALVDVPLLYLCRYLLIIRRGVIIVSGAGAGGGRLRRFRLGAAMAIIYMAMSTLFFLRIAPLTPWWGALAAWVMILLIVEAIFAFFFPYRCCFNETEGDVQNNPHV
IRIS_313-12142 MELITACALLLIAVHIAIVISCPEPLAAVALVDVPLLYLYRYLLIIRRGVIIVSGAGGGGGRLRRFRLGAAMAIIYMAMSTLFFLRIAPLTPWWGALAAWVMILLIVEAIFAFFFPYRCCFNETEGDVQNNPHV
IRIS_313-12143 MELITACALLLIAVHIAIVISCPEPLAAVALVDVPLLYLCRYLLIIRRGVIIVSGGGGGGRLRRFRLGAAMAIIYMAMSTLFFLRIAPLTPWWGALAAWVMILLIVEAIFAFFFPYRCCFNETEGDVQNNPHV
IRIS_313-12144 MELITACALLLIAVHIAIVISCPEPLAAVALVDVPLLYLYRYLLIIRRGVIIVSGAGAGGGRLRRFRLGAAMAIIYMAMSTLFFLRIAPLTPWWGALAAWVMILLIVEAIFAFFFPYRCCFNETEGDVQNNPHV
IRIS_313-12146 MELITACALLLIAVHIAIVISCPEPLAAVALVDVPLLYLCRYLLIIRRGVIIVSGGGGGGRLRRFRLGAAMAIIYMAMSTLFFLRIAPLTPWWGALAAWVMILLIVEAIFAFFFPYRCCFNETEGDVQNNPHV
IRIS_313-12147 MELITACALLLIAVHIAIVISCPEPLAAVALVDVPLLYLYRYLLIIRRGVIIVSGAGGGGGRLRRFRLGAAMAIIYMAMSTLFFLRIAPLTPWWGALAAWVMILLIVEAIFAFFFPYRCCFNETEGDVQNNPHV
IRIS_313-12148 MELITACALLLIAVHIAIVISCPEPLAAVALVDVPLLYLYRYLLIIRRGVIIVSGAGGGGGRLRRFRLGAAMAIIYMAMSTLFFLRIAPLTPWWGALAAWVMILLIVEAIFAFFFPYRCCFNETEGDVQNNPHV
IRIS_313-12151 MELITACALLLIAVHIAIVISCPEPLAAVALVDVPLLYLYRYLLIIRRGVIIVSGAGGGGGRLRRFRLGAAMAIIYMAMSTLFFLRIAPLTPWWGALAAWVMILLIVEAIFAFFFPYRCCFNETEGDVQNNPHV
IRIS_313-12152 MELITACALLLIAVHIAIVISCPEPLAAVALVDVPLLYLCRYLLIIRRGVIIVSGGGGGGRLRRFRLGAAMAIIYMAMSTLFFLRIAPLTPWWGALAAWVMILLIVEAIFAFFFPYRCCFNETEGDVQNNPHV
IRIS_313-12161 MELITACALLLIAVHIAIVISCPEPLAAVALVDVPLLYLYRYLLIIRRGVIIVSGAGGGGGRLRRFRLGAAMAIIYMAMSTLFFLRIAPLTPWWGALAAWVMILLIVEAIFAFFFPYRCCFNETEGDVQNNPHV
IRIS_313-12164 MELITACALLLIAVHIAIVISCPEPLAAVALVDVPLLYLYRYLLIIRRGVIIVSGAGGGGGRLRRFRLGAAMAIIYMAMSTLFFLRIAPLTPWWGALAAWVMILLIVEAIFAFFFPYRCCFNETEGDVQNNPHV
IRIS_313-12180 MELITACALLLIAVHIAIVISCPEPLAAVALVDVPLLYLYRYLLIIRRGVIIVSGAGGGGGRLRRFRLGAAMAIIYMAMSTLFFLRIAPLTPWWGALAAWVMILLIVEAIFAFFFPYRCCFNETEGDVQNNPHV
IRIS_313-12182 MELITACALLLIAVHIAIVISCPEPLAAVALVDVPLLYLCRYLLIIRRGVIIVSGGGGGGRLRRFRLGAAMAIIYMAMSTLFFLRIAPLTPWWGALAAWVMILLIVEAIFAFFFPYRCCFNETEGDVQNNPHV
IRIS_313-12183 MELITACALLLIAVHIAIVISCPEPLAAVAFVNVPLLYLCRYLLIIRRGVIIVSGGGGGGGRLRRFRLGAAMAVIYMAMSTLFFLRIAPLTPWWGALAAWVMILLIVEAIFAFFFPYRCCFNETEGDVQNNPHV
IRIS_313-12185 MELITACALLLIAVHIAIVISCPEPLAAVALVDVPLLYLYRYLLIIRRGVIIVSGAGGGGGRLRRFRLGAAMAIIYMAMSTLFFLRIAPLTPWWGALAAWVMILLIVEAIFAFFFPYRCCFNETEGDVQNNPHV
IRIS_313-12186 MELITACALLLIAVHIAIVISCPEPLAAVALVDVPLLYLCRYLLIIRRGVIIVSGGGGGGRLRRFRLGAAMAIIYMAMSTLFFLRIAPLTPWWGALAAWVMILLIVEAIFAFFFPYRCCFNETEGDVQNNPHV
IRIS_313-12187 MELITACALLLIAVHIAIVISCPEPLAAVALVDVPLLYLCRYLLIIRRGVIIVSGAGGGGGGRLRRFRLGAAMAIIYMAMSTLFFLRIAPLTPWWGALAAWVMILLIVEAIFAFFFPYRCCFNETEGDVQNNPHV
IRIS_313-12188 MELITACALLLIAVHIAIVISCPEPLAAVALVDVPLLYLCRYLLIIRRGVIIVSGGGGGGRLRRFRLGAAMAIIYMAMSTLFFLRIAPLTPWWGALAAWVMILLIVEAIFAFFFPYRCCFNETEGDVQNNPHV
IRIS_313-12190 MELITACALLLIAVHIAIVISCPEPLAAVALVDVPLLYLCRYLLIIRRGVIIVSGGGGGGRLRRFRLGAAMAIIYMAMSTLFFLRIAPLTPWWGALAAWVMILLIVEAIFAFFFPYRCCFNETEGDVQNNPHV
IRIS_313-12193 MELITACALLLIAVHIAIVISCPEPLAAVALVDVPLLYLCRYLLIIRRGVIIVSGGGGGGRLRRFRLGAAMAIIYMAMSTLFFLRIAPLTPWWGALAAWVMILLIVEAIFAFFFPYRCCFNETEGDVQNNPHV
IRIS_313-12194 MELITACALLLIAVHIAIVISCPEPLAAVALVDVPLLYLCRYLLIIRRGVIIVSGGGGGGRLRRFRLGAAMAIIYMAMSTLFFLRIAPLTPWWGALAAWVMILLIVEAIFAFFFPYRCCFNETEGDVQNNPHV
IRIS_313-12200 MELITACALLLIAVHIAIVISCPEPLAAVALVDVPLLYLCRYLLIIRRGVIIVSGAGAGGGRLRRFRLGAAMAIIYMAMSTLFFLRIAPLTPWWGALAAWVMILLIVEAIFAFFFPYRCCFNETEGDVQNNPHV
IRIS_313-12207 MELITACALLLIAVHIAIVISCPEPLAAVALVDVPLLYLCRYLLIIRRGVIIVSGGGGGGRLRRFRLGAAMAIIYMAMSTLFFLRIAPLTPWWGALAAWVMILLIVEAIFAFFFPYRCCFNETEGDVQNNPHV
IRIS_313-12210 MELITACALLLIAVHIAIVISCPEPLAAVALVDVPLLYLCRYLLIIRRGVIIVSGAGGAAAAGCAASASGQPWRSSTWPCRPSSSSALLLSRHGGAPSPPGS*SCSSSRPSSPSSSPTAAASTKPKAMSRIIPM
IRIS_313-12217 MELITACALLLIAVHIAIVISCPEPLAAVALVDVPLLYLCRYLLIIRRGVIIVSGAGAGGGRLRRFRLGAAMAIIYMAMSTLFFLRIAPLTPWWGALAAWVMILLIVEAIFAFFFPYRCCFNETEGDVQNNPHV
IRIS_313-12220 MELITACALLLIAVHIAIVISCPEPLAAVALVDVPLLYLYRYLLIIRRGVIIVSGAGGGGGRLRRFRLGAAMAIIYMAMSTLFFLRIAPLTPWWGALAAWVMILLIVEAIFAFFFPYRCCFNETEGDVQNNPHV
IRIS_313-12221 MELITACALLLIAVHIAIVISCPEPLAAVALVDVPLLYLCRYLLIIRRGVIIVSGAGGAAAAGCAASASGQPWRSSTWPCRPSSSSALLLSRHGGAPSPPGS*SCSSSRPSSPSSSPTAAASTKPKAMSRIIPM
IRIS_313-12222 MELITACALLLIAVHIAIVISCPEPLAAVALVDVPLLYLYRYLLIIRRGVIIVSGAGGGGGRLRRFRLGAAMAIIYMAMSTLFFLRIAPLTPWWGALAAWVMILLIVEAIFAFFFPYRCCFNETEGDVQNNPHV
IRIS_313-12224 MELITACALLLIAVHIAIVISCPEPLAAVALVDVPLLYLCRYLLIIRRGVIIVSGGGGGGRLRRFRLGAAMAIIYMAMSTLFFLRIAPLTPWWGALAAWVMILLIVEAIFAFFFPYRCCFNETEGDVQNNPHV
IRIS_313-12225 MELITACALLLIAVHIAIVISCPEPLAAVALVDVPLLYLCRYLLIIRRGVIIVSGGGGGGRLRRFRLGAAMAIIYMAMSTLFFLRIAPLTPWWGALAAWVMILLIVEAIFAFFFPYRCCFNETEGDVQNNPHV
IRIS_313-12226 MELITACALLLIAVHIAIVISCPEPLAAVALVDVPLLYLCRYLLIIRRGVIIVSGAGGGGGRLRRFRLGAAMAIIYMAMSTLFFLRIAPLTPWWGALAAWVMILLIVEAIFAFFFPYRCCFNETEGDVQNNPHV
IRIS_313-12227 MELITACALLLIAVHIAIVISCPEPLAAVALVDVPLLYLYRYLLIIRRGVIIVSGAGGGGGRLRRFRLGAAMAIIYMAMSTLFFLRIAPLTPWWGALAAWVMILLIVEAIFAFFFPYRCCFNETEGDVQNNPHV
IRIS_313-12228 MELITACALLLIAVHIAIVISCPEPLAAVALVDVPLLYLCRYLLIIRRGVIIVSGAGGGGGRLRRFRLGAAMAIIYMAMSTLFFLRIAPLTPWWGALAAWVMILLIVEAIFAFFFPYRCCFNETEGDVQNNPHV
IRIS_313-12229 MELITACALLLIAVHIAIVISCPEPLAAVALVDVPLLYLYRYLLIIRRGVIIVSGAGGGGGRLRRFRLGAAMAIIYMAMSTLFFLRIAPLTPWWGALAAWVMILLIVEAIFAFFFPYRCCFNETEGDVQNNPHV
IRIS_313-12231 MELITACALLLIAVHIAIVISCPEPLAAVALVDVPLLYLYRYLLIIRRGVIIVSGAGGGGGRLRRFRLGAAMAIIYMAMSTLFFLRIAPLTPWWGALAAWVMILLIVEAIFAFFFPYRCCFNETEGDVQNNPHV
IRIS_313-12232 MELITACALLLIAVHIAIVISCPEPLAAVALVDVPLLYLCRYLLIIRRGVIIVSGGGGGGRLRRFRLGAAMAIIYMAMSTLFFLRIAPLTPWWGALAAWVMILLIVEAIFAFFFPYRCCFNETEGDVQNNPHV
IRIS_313-12234 MELITACALLLIAVHIAIVISCPEPLAAVALVDVPLLYLCRYLLIIRRGVIIVSGAGAGGGRLRRFRLGAAMAIIYMAMSTLFFLRIAPLTPWWGALAAWVMILLIVEAIFAFFFPYRCCFNETEGDVQNNPHV
IRIS_313-12236 MELITACALLLIAVHIAIVISCPEPLAAVALVDVPLLYLYRYLLIIRRGVIIVSGAGGGGGRLRRFRLGAAMAIIYMAMSTLFFLRIAPLTPWWGALAAWVMILLIVEAIFAFFFPYRCCFNETEGDVQNNPHV
IRIS_313-12237 MELITACALLLIAVHIAIVISCPEPLAAVALVDVPLLYLYRYLLIIRRGVIIVSGAGGGGGRLRRFRLGAAMAIIYMAMSTLFFLRIAPLTPWWGALAAWVMILLIVEAIFAFFFPYRCCFNETEGDVQNNPHV
IRIS_313-12240 MELITACALLLIAVHIAIVISCPEPLAAVALVDVPLLYLCRYLLIIRRGVIIVSGGGGGGRLRRFRLGAAMAIIYMAMSTLFFLRIAPLTPWWGALAAWVMILLIVEAIFAFFFPYRCCFNETEGDVQNNPHV
IRIS_313-12241 MELITACALLLIAVHIAIVISCPEPLAAVALVDVPLLYLCRYLLIIRRGVIIVSGAGGGGGGRLRRFRLGAAMAIIYMAMSTLFFLRIAPLTPWWGALAAWVMILLIVEAIFAFFFPYRCCFNETEGDVQNNPHV
IRIS_313-12242 MELITACALLLIAVHIAIVISCPEPLAAVALVDVPLLYLCRYLLIIRRGVIIVSGAGGGGGRLRRFRLGAAMAIIYMAMSTLFFLRIAPLTPWWGALAAWVMILLIVEAIFAFFFPYRCCFNETEGDVQNNPHV
IRIS_313-12244 MELITACALLLIAVHIAIVISCPEPLAAVALVDVPLLYLCRYLLIIRRGVIIVSGAGAGGGRLRRFRLGAAMAIIYMAMSTLFFLRIAPLTPWWGALAAWVMILLIVEAIFAFFFPYRCCFNETEGDVQNNPHV
IRIS_313-12246 MELITACALLLIAVHIAIVISCPEPLAAVALVDVPLLYLCRYLLIIRRGVIIVSGAGAGGGRLRRFRLGAAMAIIYMVMSTLFFLRIAPLTPWWGALAAWVMILLIVEAIFAFFFPYRCCFNETEGDVQNNPHV
IRIS_313-12247 MELITACALLLIAVHIAIVISCPEPLAAVALVDVPLLYLCRYLLIIRRGVIIVSGAGGGGRLRRFRLGAAMAIIYMAMSTLFFLRIAPLTPWWGALAAWVMILLIVEAIFAFFFPYRCCFNETEGDVQNNPHV
IRIS_313-12249 MELITACALLLIAVHIAIVISCPEPLAAVALVDVPLLYLYRYLLIIRRGVIIVSGAGGGGGRLRRFRLGAAMAIIYMAMSTLFFLRIAPLTPWWGALAAWVMILLIVEAIFAFFFPYRCCFNETEGDVQNNPHV
IRIS_313-12251 MELITACALLLIAVHIAIVISCPEPLAAVALVDVPLLYLYRYLLIIRRGVIIVSGAGGGGGRLRRFRLGAAMAIIYMAMSTLFFLRIAPLTPWWGALAAWVMILLIVEAIFAFFFPYRCCFNETEGDVQNNPHV
IRIS_313-12252 MELITACALLLIAVHIAIVISCPEPLAAVALVDVPLLYLYRYLLIIRRGVIIVSGAGAGGGRLRRFRLGAAMAIIYMAMSTLFFLRIAPLTPWWGALAAWVMILLIVEAIFAFFFPYRCCFNETEGDVQNNPHV
IRIS_313-12254 MELITACALLLIAVHIAIVISCPEPLAAVALVDVPLLYLCRYLLIIRRGVIIVSGAGGGGGRLRRFRLGAAMAIIYMAMSTLFFLRIAPLTPWWGALAAWVMILLIVEAIFAFFFPYRCCFNETEGDVQNNPHV
IRIS_313-12257 MELITACALLLIAVHIAIVISCPEPLAAVALVDVPLLYLCRYLLIIRRGVIIVSGAGGGGGRLRRFRLGAAMAIIYMAMSTLFFLRIAPLTPWWGALAAWVMILLIVEAIFAFFFPYRCCFNETEGDVQNNPHV
IRIS_313-12258 MELITACALLLIAVHIAIVISCPEPLAAVALVDVPLLYLYRYLLIIRRGVIIVSGAGGGGGRLRRFRLGAAMAIIYMAMSTLFFLRIAPLTPWWGALAAWVMILLIVEAIFAFFFPYRCCFNETEGDVQNNPHV
IRIS_313-12259 MELITACALLLIAVHIAIVISCPEPLAAVALVDVPLLYLCRYLLIIRRGVIIVSGAGAGGGRLRRFRLGAAMAIIYMVMSTLFFLRIAPLTPWWGALAAWVMILLIVEAIFAFFFPYRCCFNETEGDVQNNPHV
IRIS_313-12260 MELITACALLLIAVHIAIVISCPEPLAAVALVDVPLLYLCRYLLIIRRGVIIVSGGGGGGRLRRFRLGAAMAIIYMAMSTLFFLRIAPLTPWWGALAAWVMILLIVEAIFAFFFPYRCCFNETEGDVQNNPHV
IRIS_313-12261 MELITACALLLIAVHIAIVISCPEPLAAVALVDVPLLYLCRYLLIIRRGVIIVSGGGGGGRLRRFRLGAAMAIIYMAMSTLFFLRIAPLTPWWGALAAWVMILLIVEAIFAFFFPYRCCFNETEGDVQNNPHV
IRIS_313-12262 MELITACALLLIAVHIAIVISCPEPLAAVALVDVPLLYLYRYLLIIRRGVIIVSGAGAGGGRLRRFRLGAAMAIIYMAMSTLFFLRIAPLTPWWGALAAWVMILLIVEAIFAFFFPYRCCFNETEGDVQNNPHV
IRIS_313-12263 MELITACALLLIAVHIAIVISCPEPLAAVALVDVPLLYLYRYLLIIRRGVIIVSGAGGGGGRLRRFRLGAAMAIIYMAMSTLFFLRIAPLTPWWGALAAWVMILLIVEAIFAFFFPYRCCFNETEGDVQNNPHV
IRIS_313-12265 MELITACALLLIAVHIAIVISCPEPLAAVALVDVPLLYLYRYLLIIRRGVIIVSGAGGGGGRLRRFRLGAAMAIIYMAMSTLFFLRIAPLTPWWGALAAWVMILLIVEAIFAFFFPYRCCFNETEGDVQNNPHV
IRIS_313-12266 MELITACALLLIAVHIAIVISCPEPLAAVALVDVPLLYLCRYLLIIRRGVIIVSGAGAGGGRLRRFRLGAAMAIIYMAMSTLFFLRIAPLTPWWGALAAWVMILLIVEAIFAFFFPYRCCFNETEGDVQNNPHV
IRIS_313-12268 MELITACALLLIAVHIAIVISCPEPLAAVALVDVPLLYLCRYLLIIRRGVIIVSGGGGGGRLRRFRLGAAMAIIYMAMSTLFFLRIAPLTPWWGALAAWVMILLIVEAIFAFFFPYRCCFNETEGDVQNNPHV
IRIS_313-12269 MELITACALLLIAVHIAIVISCPEPLAAVALVDVPLLYLCRYLLIIRRGVIIVSGAGGGGGRLRRFRLGAAMAIIYMAMSTLFFLRIAPLTPWWGALAAWVMILLIVEAIFAFFFPYRCCFNETEGDVQNNPHV
IRIS_313-12270 MELITACALLLIAVHIAIVISCPEPLAAVALVDVPLYLCRYLLIIRRGVIIVSGGGGGGRLRRFRLGAAMAIIYMAMSTLFFLRIAPLTPWWGALAAWVMILLIVEAIFAFFFPYRCCFNETEGDVQNNPHV
IRIS_313-12271 MELITACALLLIAVHIAIVISCPEPLAAVALVDVPLLYLYRYLLIIRRGVIIVSGAGGGGGRLRRFRLGAAMAIIYMAMSTLFFLRIAPLTPWWGALAAWVMILLIVEAIFAFFFPYRCCFNETEGDVQNNPHV
IRIS_313-12272 MELITACALLLIAVHIAIVISCPEPLAAVALVDVPLLYLCRYLLIIRRGVIIVSGAGAGGGRLRRFRLGAAMAIIYMAMSTLFFLRIAPLTPWWGALAAWVMILLIVEAIFAFFFPYRCCFNETEGDVQNNPHV
IRIS_313-12273 MELITACALLLIAVHIAIVISCPEPLAAVALVDVPLLYLYRYLLIIRRGVIIVSGAGGGGGRLRRFRLGAAMAIIYMAMSTLFFLRIAPLTPWWGALAAWVMILLIVEAIFAFFFPYRCCFNETEGDVQNNPHV
IRIS_313-12275 MELITACALLLIAVHIAIVISCPEPLAAVALVDVPLLYLCRYLLIIRRGVIIVSGAGGAAAAGCAASASGQPWRSSTWPCRPSSSSALLLSRHGGAPSPPGS*SCSSSRPSSPSSSPTAAASTKPKAMSRIIPM
IRIS_313-12278 MELITACALLLIAVHIAIVISCPEPLAAVAFVNVPLLYLCRYLLIIRRGVIIVSGGGGGGGRLRRFRLGAAMAVIYMAMSTLFFLRIAPLTPWWGALAAWVMILLIVEAIFAFFFPYRCCFNETEGDVQNNPHV
IRIS_313-12279 MELITACALLLIAVHIAIVISCPEPLAAVALVDVPLLYLCRYLLIIRRGVIIVSGAGAGGGRLRRFRLGAAMAIIYMAMSTLFFLRIAPLTPWWGALAAWVMILLIVEAIFAFFFPYRCCFNETEGDVQNNPHV
IRIS_313-12280 MELITACALLLIAVHIAIVISCPEPLAAVALVDVPLLYLCRYLLIIRRGVIIVSGAGAGGGRLRRFRLGAAMAIIYMAMSTLFFLRIAPLTPWWGALAAWVMILLIVEAIFAFFFPYRCCFNETEGDVQNNPHV
```

IRIS_313-12281 MELITACALLLIAVHIAIVISCPEPLAAVALVDVPLLYLYRYLLIIRRGVIIVSGAGGGGGRLRRFRLGAAMAIIYMAMSTLFFLRIAPLTPWWGALAAWVMILLIVEAIFAFFFPYRCCFNETEGDVQNNPHV

IRIS_313-12282 MELITACALLLIAVHIAIVISCPEPLAAVALVDVPLLYLCRYLLIIRRGVIIVSGGGAGGGRLRRFRLGAAMAIIYMAMSTLFFLRIAPLTPWWGALAAWVMILLIVEAIFAFFFPYRCCFNETEGDVQNNPHV

IRIS_313-12283 MELITACALLLIAVHIAIVISCPEPLAAVALVDVPLLYLCRYLLIIRRGVIIVSGGGGGGRLRRFRLGAAMAIIYMAMSTLFFLRIAPLTPWWGALAAWVMILLIVEAIFAFFFPYRCCFNETEGDVQNNPHV

IRIS_313-12284 MELITACALLLIAVHIAIVISCPEPLAAVALVDVPLLYLCRYLLIIRRGVIIVSGAGGAAAAGCAASASGQPWRSSTWPCRPSSSSALLLSRHGGAPSPPGS*SCSSSRPSSPSSSPTAAASTKPKAMSRIIPM

IRIS_313-12285 MELITACALLLIAVHIAIVISCPEPLAAVALVDVPLLYLCRYLLIIRRGVIIVSGAGAGGGRLRRFRLGAAMAIIYMAMSTLFFLRIAPLTPWWGALAAWVMILLIVEAIFAFFFPYRCCFNETEGDVQNNPHV

IRIS_313-12286 MELITACALLLIAVHIAIVISCPEPLAAVALVDVPLLYLCRYLLIIRRGVIIVSGGGGGGRLRRFRLGAAMAIIYMAMSTLFFLRIAPLTPWWGALAAWVMILLIVEAIFAFFFPYRCCFNETEGDVQNNPHV

IRIS_313-12287 MELITACALLLIAVHIAIVISCPEPLAAVALVDVPLLYLCRYLLIIRRGVIIVSGGGGGGRLRRFRLGAAMAIIYMAMSTLFFLRIAPLTPWWGALAAWVMILLIVEAIFAFFFPYRCCFNETEGDVQNNPHV

IRIS_313-12288 MELITACALLLIAVHIAIVISCPEPLAAVALVDVPLLYLCRYLLIIRRGVIIVSGGGGGGRLRRFRLGAAMAIIYMAMSTLFFLRIAPLTPWWGALAAWVMILLIVEAIFAFFFPYRCCFNETEGDVQNNPHV

IRIS_313-12289 MELITACALLLIAVHIAIVISCPEPLAAVALVDVPLLYLYRYLLIIRRGVIIVSGAGGGGGRLRRFRLGAAMAIIYMAMSTLFFLRIAPLTPWWGALAAWVMILLIVEAIFAFFFPYRCCFNETEGDVQNNPHV

IRIS_313-12290 MELITACALLLIAVHIAIVISCPEPLAAVALVDVPLLYLCRYLLIIRRGVIIVSGAGGGGGGRLRRFRLGAAMAIIYMAMSTLFFLRIAPLTPWWGALAAWVMILLIVEAIFAFFFPYRCCFNETEGDVQNNPHV

IRIS_313-12291 MELITACALLLIAVHIAIVISCPEPLAAVALVDVPLLYLYRYLLIIRRGVIIVSGAGGGGGRLRRFRLGAAMAIIYMAMSTLFFLRIAPLTPWWGALAAWVMILLIVEAIFAFFFPYRCCFNETEGDVQNNPHV

IRIS_313-12292 MELITACALLLIAVHIAIVISCPEPLAAVALVDVPLLYLCRYLLIIRRGVIIVSGAGGAAAAGCAASASGQPWRSSTWPCRPSSSSALLLSRHGGAPSPPGS*SCSSSRPSSPSSSPTAAASTKPKAMSRIIPM

IRIS_313-12296 MELITACALLLIAVHIAIVISCPEPLAAVALVDVPLLYLCRYLLIIRRGVIIVSGGGGGGRLRRFRLGAAMAIIYMAMSTLFFLRIAPLTPWWGALAAWVMILLIVEAIFAFFFPYRCCFNETEGDVQNNPHV

IRIS_313-12297 MELITACALLLIAVHIAIVISCPEPLAAVALVDVPLLYLCRYLLIIRRGVIIVSGAGGGGGGRLRRFRLGAAMAIIYMAMSTLFFLRIAPLTPWWGALAAWVMILLIVEAIFAFFFPYRCCFNETEGDVQNNPHV

IRIS_313-12299 MELITACALLLIAVHIAIVISCPEPLAAVALVDVPLLYLYRYLLIIRRGVIIVSGAGGGGGRLRRFRLGAAMAIIYMAMSTLFFLRIAPLTPWWGALAAWVMILLIVEAIFAFFFPYRCCFNETEGDVQNNPHV

IRIS_313-12300 MELITACALLLIAVHIAIVISCPEPLAAVALVDVPLLYLCRYLLIIRRGVIIVSGAGGGGGGRLRRFRLGAAMAIIYMAMSTLFFLRIAPLTPWWGALAAWVMILLIVEAIFAFFFPYRCCFNETEGDVQNNPHV

IRIS_313-12301 MELITACALLLIAVHIAIVISCPEPLAAVALVDVPLLYLCRYLLIIRRGVIIVSGAGGGGGRLRRFRLGAAMAIIYMAMSTLFFLRIAPLTPWWGALAAWVMILLIVEAIFAFFFPYRCCFNETEGDVQNNPHV

IRIS_313-12302 MELITACALLLIAVHIAIVISCPEPLAAVALVDVPLLYLCRYLLIIRRGVIIVSGAGGAAAAGCAASASGQPWRSSTWPCRPSSSSALLLSRHGGAPSPPGS*SCSSSRPSSPSSSPTAAASTKPKAMSRIIPM

IRIS_313-12303 MELITACALLLIAVHIAIVISCPEPLAAVALVDVPLLYLYRYLLIIRRGVIIVSGAGGGGGRLRRFRLGAAMAIIYMAMSTLFFLRIAPLTPWWGALAAWVMILLIVEAIFAFFFPYRCCFNETEGDVQNNPHV

IRIS_313-12305 MELITACALLLIAVHIAIVISCPEPLAAVALVDVPLLYLYRYLLIIRRGVIIVSGAGGGGGRLRRFRLGAAMAIIYMAMSTLFFLRIAPLTPWWGALAAWVMILLIVEAIFAFFFPYRCCFNETEGDVQNNPHV

IRIS_313-12307 MELITACALLLIAVHIAIVISCPEPLAAVALVDVPLLYLCRYLLIIRRGVIIVSGAGAGGGRLRRFRLGAAMAIIYMAMSTLFFLRIAPLTPWWGALAAWVMILLIVEAIFAFFFPYRCCFNETEGDVQNNPHV

IRIS_313-12308 MELITACALLLIAVHIAIVISCPEPLAAVALVDVPLLYLCRYLLIIRRGVIIVSGGGGGGRLRRFRLGAAMAIIYMAMSTLFFLRIAPLTPWWGALAAWVMILLIVEAIFAFFFPYRCCFNETEGDVQNNPHV

IRIS_313-12309 MELITACALLLIAVHIAIVISCPEPLAAVALVDVPLLYLCRYLLIIRRGVIIVSGGGGGGRLRRFRLGAAMAIIYMAMSTLFFLRIAPLTPWWGALAAWVMILLIVEAIFAFFFPYRCCFNETEGDVQNNPHV

IRIS_313-12311 MELITACALLLIAVHIAIVISCPEPLAAVALVDVPLLYLYRYLLIIRRGVIIVSGAGGGGGRLRRFRLGAAMAIIYMAMSTLFFLRIAPLTPWWGALAAWVMILLIVEAIFAFFFPYRCCFNETEGDVQNNPHV

IRIS_313-12312 MELITACALLLIAVHIAIVISCPEPLAAVALVDVPLLYLCRYLLIIRRGVIIVSGAGAGGGRLRRFRLGAAMAIIYMAMSTLFFLRIAPLTPWWGALAAWVMILLIVEAIFAFFFPYRCCFNETEGDVQNNPHV

IRIS_313-12313 MELITACALLLIAVHIAIVISCPEPLAAVALVDVPLLYLYRYLLIIRRGVIIVSGAGGGGGRLRRFRLGAAMAIIYMAMSTLFFLRIAPLTPWWGALAAWVMILLIVEAIFAFFFPYRCCFNETEGDVQNNPHV

IRIS_313-12319 MELITACALLLIAVHIAIVISCPEPLAAVALVDVPLLYLCRYLLIIRRGVIIVSGGGGGGRLRRFRLGAAMAIIYMAMSTLFFLRIAPLTPWWGALAAWVMILLIVEAIFAFFFPYRCCFNETEGDVQNNPHV

IRIS_313-12321 MELITACALLLIAVHIAIVISCPEPLAAVALVDVPLLYLYRYLLIIRRGVIIVSGAGGGGGRLRRFRLGAAMAIIYMAMSTLFFLRIAPLTPWWGALAAWVMILLIVEAIFAFFFPYRCCFNETEGDVQNNPHV

IRIS_313-12322 MELITACALLLIAVHIAIVISCPEPLAAVALVDVPLLYLCRYLLIIRRGVIIVSGAGGGGGRLRRFRLGAAMAIIYMAMSTLFFLRIAPLTPWWGALAAWVMILLIVEAIFAFFFPYRCCFNETEGDVQNNPHV

IRIS_313-12323 MELITACALLLIAVHIAIVISCPEPLAAVALVDVPLLYLCRYLLIIRRGVIIVSGAGGGGGRLRRFRLGAAMAIIYMAMSTLFFLRIAPLTPWWGALAAWVMILLIVEAIFAFFFPYRCCFNETEGDVQNNPHV

IRIS_313-12324 MELITACALLLIAVHIAIVISCPEPLAAVALVDVPLLYLCRYLLIIRRGVIIVSGAGGGGGRLRRFRLGAAMAIIYMAMSTLFFLRIAPLTPWWGALAAWVMILLIVEAIFAFFFPYRCCFNETEGDVQNNPHV

IRIS_313-12325 MELITACALLLIAVHIAIVISCPEPLAAVALVDVPLLYLCRYLLIIRRGVIIVSGAGAGGGRLRRFRLGAAMAIIYMAMSTLFFLRIAPLTPWWGALAAWVMILLIVEAIFAFFFPYRCCFNETEGDVQNNPHV

IRIS_313-12329 MELITACALLLIAVHIAIVISCPEPLAAVALVDVPLLYLCRYLLIIRRGVIIVSGAGGAAAAGCAASASGQPWRSSTWPCRPSSSSALLLSRHGGAPSPPGS*SCSSSRPSSPSSSPTAAASTKPKAMSRIIPM

IRIS_313-12330 MELITACALLLIAVHIAIVISCPEPLAAVALVDVPLLYLCRYLLIIRRGVIIVSGAGAGGGRLRRFRLGAAMAIIYMAMSTLFFLRIAPLTPWWGALAAWVMILLIVEAIFAFFFPYRCCFNETEGDVQNNPHV

IRIS_313-12332 MELITACALLLIAVHIAIVISCPEPLAAVALVDVPLLYLCRYLLIIRRGVIIVSGAGGGGGRLRRFRLGAAMAIIYMAMSTLFFLRIAPLTPWWGALAAWVMILLIVEAIFAFFFPYRCCFNETEGDVQNNPHV

IRIS_313-12334 MELITACALLLIAVHIAIVISCPEPLAAVALVDVPLLYLCRYLLIIRRGVIIVSGGGGGGRLRRFRLGAAMAIIYMAMSTLFFLRIAPLTPWWGALAAWVMILLIVEAIFAFFFPYRCCFNETEGDVQNNPHV

IRIS_313-12336 MELITACALLLIAVHIAIVISCPEPLAAVALVDVPLLYLCRYLLIIRRGVIIVSGAGAGGGRLRRFRLGAAMAIIYMAMSTLFFLRIAPLTPWWGALAAWVMILLIVEAIFAFFFPYRCCFNETEGDVQNNPHV

IRIS_313-12337 MELITACALLLIAVHIAIVISCPEPLAAVALVDVPLLYLYRYLLIIRRGVIIVSGAGGGGGRLRRFRLGAAMAIIYMAMSTLFFLRIAPLTPWWGALAAWVMILLIVEAIFAFFFPYRCCFNETEGDVQNNPHV

IRIS_313-12340 MELITACALLLIAVHIAIVISCPEPLAAVALVDVPLLYLCRYLLIIRRGVIIVSGAGGGGGRLRRFRLGAAMAIIYMAMSTLFFLRIAPLTPWWGALAAWVMILLIVEAIFAFFFPYRCCFNETEGDVQNNPHV

IRIS_313-12341 MELITACALLLIAVHIAIVISCPEPLAAVALVDVPLLYLCRYLLIIRRGVIIVSGAGAGGGRLRRFRLGAAMAIIYMAMSTLFFLRIAPLTPWWGALAAWVMILLIVEAIFAFFFPYRCCFNETEGDVQNNPHV

IRIS_313-12342 MELITACALLLIAVHIAIVISCPEPLAAVALVDVPLLYLCRYLLIIRRGVIIVSGAGAGGGRLRRFRLGAAMAIIYMAMSTLFFLRIAPLTPWWGALAAWVMILLIVEAIFAFFFPYRCCFNETEGDVQNNPHV

IRIS_313-12344 MELITACALLLIAVHIAIVISCPEPLAAVALVDVPLLYLCRYLLIIRRGVIIVSGAGGGGGRLRRFRLGAAMAIIYMAMSTLFFLRIAPLTPWWGALAAWVMILLIVEAIFAFFFPYRCCFNETEGDVQNNPHV

IRIS_313-12345 MELITACALLLIAVHIAIVISCPEPLAAVALVDVPLLYLYRYLLIIRRGVIIVSGAGGGGGRLRRFRLGAAMAIIYMAMSTLFFLRIAPLTPWWGALAAWVMILLIVEAIFAFFFPYRCCFNETEGDVQNNPHV

IRIS_313-12346 MELITACALLLIAVHIAIVISCPEPLAAVALVDVPLLYLCRYLLIIRRGVIIVSGAGGGGGRLRRFRLGAAMAIIYMAMSTLFFLRIAPLTPWWGALAAWVMILLIVEAIFAFFFPYRCCFNETEGDVQNNPHV

IRIS_313-12347 MELITACALLLIAVHIAIVISCPEPLAAVALVDVPLLYLCRYLLIIRRGVIIVSGAGGGGGRLRRFRLGAAMAIIYMAMSTLFFLRIAPLTPWWGALAAWVMILLIVEAIFAFFFPYRCCFNETEGDVQNNPHV

IRIS_313-12348 MELITACALLLIAVHIAIVISCPEPLAAVALVDVPLLYLCRYLLIIRRGVIIVSGAGAGGGRLRRFRLGAAMAIIYMAMSTLFFLRIAPLTPWWGALAAWVMILLIVEAIFAFFFPYRCCFNETEGDVQNNPHV

IRIS_313-12349 MELITACALLLIAVHIAIVISCPEPLAAVALVDVPLLYLCRYLLIIRRGVIIVSGAGAGGGRLRRFRLGAAMAIIYMAMSTLFFLRIAPLTPWWGALAAWVMILLIVEAIFAFFFPYRCCFNETEGDVQNNPHV

IRIS_313-12350 MELITACALLLIAVHIAIVISCPEPLAAVALVDVPLLYLYRYLLIIRRGVIIVSGAGGGGGRLRRFRLGAAMAIIYMAMSTLFFLRIAPLTPWWGALAAWVMILLIVEAIFAFFFPYRCCFNETEGDVQNNPHV

IRIS_313-12351 MELITACALLLIAVHIAIVISCPEPLAAVALVDVPLLYLYRYLLIIRRGVIIVSGAGAGGGRLRRFRLGAAMAIIYMAMSTLFFLRIAPLTPWWGALAAWVMILLIVEAIFAFFFPYRCCFNETEGDVQNNPHV

IRIS_313-12352 MELITACALLLIAVHIAIVISCPEPLAAVALVDVPLLYLCRYLLIIRRGVIIVSGAGAGGGRLRRFRLGAAMAIIYMAMSTLFFLRIAPLTPWWGALAAWVMILLIVEAIFAFFFPYRCCFNETEGDVQNNPHV

IRIS_313-12353 MELITACALLLIAVHIAIVISCPEPLAAVALVDVPLLYLCRYLLIIRRGVIIVSGAGGGGGRLRRFRLGAAMAIIYMAMSTLFFLRIAPLTPWWGALAAWVMILLIVEAIFAFFFPYRCCFNETEGDVQNNPHV

IRIS_313-12354 MELITACALLLIAVHIAIVISCPEPLAAVALVDVPLLYLCRYLLIIRRGVIIVSGGGGGGRLRRFRLGAAMAIIYMAMSTLFFLRIAPLTPWWGALAAWVMILLIVEAIFAFFFPYRCCFNETEGDVQNNPHV

IRIS_313-12355 MELITACALLLIAVHIAIVISCPEPLAAVALVDVPLLYLCRYLLIIRRGVIIVSGAGAGGGRLRRFRLGAAMAIIYMAMSTLFFLRIAPLTPWWGALAAWVMILLIVEAIFAFFFPYRCCFNETEGDVQNNPHV

IRIS_313-15896 MELITACALLLIAVHIAIVISCPEPLAAVALVDVPLLYLCRYLLIIRRGVIIVSGAGAGGGRLRRFRLGAAMAIIYMAMSTLFFLRIAPLTPWWGALAAWVMILLIVEAIFAFFFPYRCCFNETEGDVQNNPHV

IRIS_313-15897 MELITACALLLIAVHIAIVISCPEPLAAVALVDVPLLYLCRYLLIIRRGVIIVSGAGAGGGRLRRFRLGAAMAIIYMAMSTLFFLRIAPLTPWWGALAAWVMILLIVEAIFAFFFPYRCCFNETEGDVQNNPHV

IRIS_313-15898 MELITACALLLIAVHIAIVISCPEPLAAVALVDVPLLYLCRYLLIIRRGVIIVSGGGAGGGRLRRFRLGAAMAIIYMAMSTLFFLRIAPLTPWWGALAAWVMILLIVEAIFAFFFPYRCCFNETEGDVQNNPHV

IRIS_313-15899 MELITACALLLIAVHIAIVISCPEPLAAVALVDVPLLYLCRYLLIIRRGVIIVSGGGAGGGRLRRFRLGAAMAIIYMAMSTLFFLRIAPLTPWWGALAAWVMILLIVEAIFAFFFPYRCCFNETEGDVQNNPHV

IRIS_313-15900 MELITACALLLIAVHIAIVISCPEPLAAVALVDVPLLYLCRYLLIIRRGVIIVSGGGGGGRLRRFRLGAAMAIIYMAMSTLFFLRIAPLTPWWGALAAWVMILLIVEAIFAFFFPYRCCFNETEGDVQNNPHV

IRIS_313-15901 MELITACALLLIAVHIAIVISCPEPLAAVALVDVPLLYLCRYLLIIRRGVIIVSGAGAGGGRLRRFRLGAAMAIIYMAMSTLFFLRIAPLTPWWGALAAWVMILLIVEAIFAFFFPYRCCFNETEGDVQNNPHV

IRIS_313-15902 MELITACALLLIAVHIAIVISCPEPLAAVALVDVPLLYLCRYLLIIRRGVIIVSGGGGGGRLRRFRLGAAMAIIYMAMSTLFFLRIAPLTPWWGALAAWVMILLIVEAIFAFFFPYRCCFNETEGDVQNNPHV

IRIS_313-15903 MELITACALLLIAVHIAIVISCPEPLAAVALVDVPLLYLCRYLLIIRRGVIIVSGAGAGGGRLRRFRLGAAMAIIYMAMSTLFFLRIAPLTPWWGALAAWVMILLIVEAIFAFFFPYRCCFNETEGDVQNNPHV

IRIS_313-15904 MELITACALLLIAVHIAIVISCPEPLAAVALVDVPLLYLCRYLLIIRRGVIIVSGAGAGGGRLRRFRLGAAMAIIYMAMSTLFFLRIAPLTPWWGALAAWVMILLIVEAIFAFFFPYRCCFNETEGDVQNNPHV

IRIS_313-15905 MELITACALLLIAVHIAIVISCPEPLAAVALVDVPLLYLCRYLLIIRRGVIIVSGAGAGGGRLRRFRLGAAMAIIYMAMSTLFFLRIAPLTPWWGALAAWVMILLIVEAIFAFFFPYRCCFNETEGDVQNNPHV

IRIS_313-15906 MELITACALLLIAVHIAIVISCPEPLAAVALVDVPLLYLCRYLLIIRRGVIIVSGGGAGGGRLRRFRLGAAMAIIYMAMSTLFFLRIAPLTPWWGALAAWVMILLIVEAIFAFFFPYRCCFNETEGDVQNNPHV

IRIS_313-15907 MELITACALLLIAVHIAIVISCPEPLAAVALVDVPLLYLYRYLLIIRRGVIIVSGAGGGGGRLRRFRLGAAMAIIYMAMSTLFFLRIAPLTPWWGALAAWVMILLIVEAIFAFFFPYRCCFNETEGDVQNNPHV

IRIS_313-15908 MELITACALLLIAVHIAIVISCPEPLAAVALVDVPLLYLCRYLLIIRRGVIIVSGAGAGGGRLRRFRLGAAMAIIYMAMSTLFFLRIAPLTPWWGALAAWVMILLIVEAIFAFFFPYRCCFNETEGDVQNNPHV

IRIS_313-15909 MELITACALLLIAVHIAIVISCPEPLAAVALVDVPLLYLCRYLLIIRRGVIIVSGAGAGGGRLRRFRLGAAMAIIYMAMSTLFFLRIAPLTPWWGALAAWVMILLIVEAIFAFFFPYRCCFNETEGDVQNNPHV

IRIS_313-15910 MELITACALLLIAVHIAIVISCPEPLAAVALVDVPLLYLYRYLLIIRRGVIIVSGAGGGGGRLRRFRLGAAMAIIYMAMSTLFFLRIAPLTPWWGALAAWVMILLIVEAIFAFFFPYRCCFNETEGDVQNNPHV

**Data S2**. **XA23 protein variants from 3,002 cultivated rice accessions.** Genomic sequences corresponding to the chr11:22,202,401-22,205,400 of the reference genome (*O. sativa* cv. Nipponbare) were extracted from the IRRI SNP-Seek Database (http://snp-seek.irri.org).[9] Start and stop codons of *Xa23* genes from 3,002 cultivated rice accessions were identified based on the genomic structure of *Xa23* from *O. sativa* cv. Nipponbare. None of the examined 3,002 accessions contained premature stop codons within the *Xa23* CDS.

```
B001        MLHHLKELAAVAGIHMILIYLCRFLLRRSRNVLFTVSNSLRFRLKVLTVLLYICLSVMLFYLFGSIMPLPPWGLVVGWVMALIAVELAYAFIFPYSFRYIADNDDDKMVILPV
B002        MLHHLKELAAVAGIHMILIYLCRFLLRRSRNVLFTVSNSLRFRLKVLTVLLYICLSVMLFYLFGSIMPLPPWGLVVGWVMALIAVELAYAFIFPYSFRYIADNDDDKMVILPV
B003        MLHHLKELAAVAGIHMILIYLCRFLLRRSRNVLFTVSNSLRFRLKVLTVLLYICLSVMLFYLFGSIMPLPPWGLVVGWVMALIAVELAYAFIFPYSFRYIADNDDDKMVILPV
B004        MLHHLKELAAVAGIHMILIYLCRFLLRRSRNVLFTVSNSLRFRLKVLTVLLYICLSVMLFYLFGSIMPLPPWGLVVGWVMALIAVELAYAFIFPYSFRYIADNDDDKMVILPV
B005        MLHHLKELAAVAGIHMILIYLCRFLLRRSRNVLFTVSNSLRFRLKVLTVLLYICLSVMLFYLFGSIMPLPPWGLVVGWVMALIAVELAYAFIFPYSFRYIADNDDDKMVILPV
B006        MLHHLKELAAVAGIHMILIYLCRFLLRRSRNVLFTVSNSLRFRLKVLTVLLYICLSVMLFYLFGSIMPLPPWGLVVGWVMALIAVELAYAFIFPYSFRYIADNDDDKMVILPV
B007        MLHHLKELAAVAGIHMILIYLCRFLLRRSRNVLFTVSNSLRFRLKVLTVLLYICLSVMLFYLFGSIMPLPPWGLVVGWVMALIAVELAYAFIFPYSFRYIADNDDDKMVILPV
B008        MLHHLKELAAVAGIHMILIYLCRFLLRRSRNVLFTVSNSLRFRLKVLTVLLYICLSVMLFYLFGSIMPLPPWGLVVGWVMALIAVELAYAFIFPYSFRYIADNDDDKMVILPV
B009        MLHHLKELAAVAGIHMILIYLCRFLLRLRRNVLFTVSNSLRFRLKVLTVLLYICLSVMLFYLFGSIMPLPPWGLVVGWVMALIAVELAYAFIFPYSFRYIADNDDDKMVILPV
B010        MLHHLKELAAVAGIHMILIYLCRFLLRRSRNVLFTVSNSLRFRLKVLTVLLYICLSVMLFYLFGSIMPLPPWGLVVGWVMALIAVELAYAFIFPYSFRYIADNDDDKMVILPV
B011        MLHHLKELAAVAGIHMILIYLCRFLLRRSRNVLFTVSNSLRFRLKVLTVLLYICLSVMLFYLFGSIMPLPPWGLVVGWVMALIAVELAYAFIFPYSFRYIADNDDDKMVILPV
B012        MLHHLKELAAVAGIHMILIYLCRFLLRRSRNVLFTVSNSLRFRLKVLTVLLYICLSVMLFYLFGSIMPLPPWGLVVGWVMALIAVELAYAFIFPYSFRYIADNDDDKMVILPV
B013        MLHHLKELAAVAGIHMILIYLCRFLLRRSRNVLFTVSNSLRFRLKVLTVLLYICLSVMLFYLFGSIMPLPPWGLVVGWVMALIAVELAYAFIFPYSFRYIADNDDDKMVILPV
B015        MLHHLKELAAVAGIHMILIYLCRFLLRRSRNVLFTVSNSLRFRLKVLTVLLYICLSVMLFYLFGSIMPLPPWGLVVGWVMALIAVELAYAFIFPYSFRYIADNDDDKMVILPV
B016        MLHHLKELAAVAGIHMILIYLCRFLLRRSRNVLFTVSNSLRFRLKVLTVLLYICLSVMLFYLFGSIMPLPPWGLVVGWVMALIAVELAYAFIFPYSFRYIADNDDDKMVILPV
B017        MLHHLKELAAVAGIHMILIYLCRFLLRRSRNVLFTVSNSLRFRLKVLTVLLYICLSVMLFYLFGSIMPLPPWGLVVGWVMALIAVELAYAFIFPYSFRYIADNDDDKMVILPV
B018        MLHHLKELAAVAGIHMILIYLCRFLLRRSRNVLFTVSNSLRFRLKVLTVLLYICLSVMLFYLFGSIMPLPPWGLVVGWVMALIAVELAYAFIFPYSFRYIADNDDDKMVILPV
B019        MLHHLKELAAVAGIHMILIYLCRFLLRRSRNVLFTVSNSLRFRLKVLTVLLYICLSVMLFYLFGSIMPLPPWGLVVGWVMALIAVELAYAFIFPYSFRYIADNDDDKMVILPV
B020        MLHHLKELAAVAGIHMILIYLCRFLLRRSRNVLFTVSNSLRFRLKVLTVLLYICLSVMLFYLFGSIMPLPPWGLVVGWVMALIAVELAYAFIFPYSFRYIADNDDDKMVILPV
B021        MLHHLKELAAVAGIHMILIYLCRFLLRRSRNVLFTVSNSLRFRLKVLTVLLYICLSVMLFYLFGSIMPLPPWGLVVGWVMALIAVELAYAFIFPYSFRYIADNDDDKMVILPV
B023        MLHHLKELAAVAGIHMILIYLCRFLLRRSRNVLFTVSNSLRFRLKVLTVLLYICLSVMLFYLFGSIMPLPPWGLVVGWVMALIAVELAYAFIFPYSFRYIADNDDDKMVILPV
B024        MLHHLKELAAVAGIHMILIYLCRFLLRRSRNVLFTVSNSLRFRLKVLTVLLYICLSVMLFYLFGSIMPLPPWGLVVGWVMALIAVELAYAFIFPYSFRYIADNDDDKMVILPV
B025        MLHHLKELAAVAGIHMILIYLCRFLLRRSRNVLFTVSNSLRFRLKVLTVLLYICLSVMLFYLFGSIMPLPPWGLVVGWVMALIAVELAYAFIFPYSFRYIADNDDDKMVILPV
B026        MLHHLKELAAVAGIHMILIYLCRFLLRRSRNVLFTVSNSLRFRLKVLTVLLYICLSVMLFYLFGSIMPLPPWGLVVGWVMALIAVELAYAFIFPYSFRYIADNDDDKMVILPV
B027        MLHHLKELAAVAGIHMILIYLCRFLLRRSRNVLFTVSNSLRFRLKVLTVLLYICLSVMLFYLFGSIMPLPPWGLVVGWVMALIAVELAYAFIFPYSFRYIADNDDDKMVILPV
B028        MLHHLKELAAVAGIHMILIYLCRFLLRRSRNVLFTVSNSLRFRLKVLTVLLYICLSVMLFYLFGSIMPLPPWGLVVGWVMALIAVELAYAFIFPYSFRYIADNDDDKMVILPV
B029        MLHHLKELAAVAGIHMILIYLCRFLLRRSRNVLFTVSNSLRFRLKVLTVLLYICLSVMLFYLFGSIMPLPPWGLVVGWVMALIAVELAYAFIFPYSFRYIADNDDDKMVILPV
B030        MLHHLKELAAVAGIHMILIYLCRFLLRRSRNVLFTVSNSLRFRLKVLTVLLYICLSVMLFYLFGSIMPLPPWGLVVGWVMALIAVELAYAFIFPYSFRYIADNDDDKMVILPV
B031        MLHHLKELAAVAGIHMILIYLCRFLLRRSRNVLFTVSNSLRFRLKVLTVLLYICLSVMLFYLFGSIMPLPPWGLVVGWVMALIAVELAYAFIFPYSFRYIADNDDDKMVILPV
B032        MLHHLKELAAVAGIHMILIYLCRFLLRRSRNVLFTVSNSLRFRLKVLTVLLYICLSVMLFYLFGSIMPLPPWGLVVGWVMALIAVELAYAFIFPYSFRYIADNDDDKMVILPV
B033        MLHHLKELAAVAGIHMILIYLCRFLLRRSRNVLFTVSNSLRFRLKVLTVLLYICLSVMLFYLFGSIMPLPPWGLVVGWVMALIAVELAYAFIFPYSFRYIADNDDDKMVILPV
B034        MLHHLKELAAVAGIHMILIYLCRFLLRRSRNVLFTVSNSLRFRLKVLTVLLYICLSVMLFYLFGSIMPLPPWGLVVGWVMALIAVELAYAFIFPYSFRYIADNDDDKMVILPV
B035        MLHHLKELAAVAGIHMILIYLCRFLLRRSRNVLFTVSNSLRFRLKVLTVLLYICLSVMLFYLFGSIMPLPPWGLVVGWVMALIAVELAYAFIFPYSFRYIADNDDDKMVILPV
B036        MLHHLKELAAVAGIHMILIYLCRFLLRRSRNVLFTVSNSLRFRLKVLTVLLYICLSVMLFYLFGSIMPLPPWGLVVGWVMALIAVELAYAFIFPYSFRYIADNDDDKMVILPV
B037        MLHHLKELAAVAGIHMILIYLCRFLLRRSRNVLFTVSNSLRFRLKVLTVLLYICLSVMLFYLFGSIMPLPPWGLVVGWVMALIAVELAYAFIFPYSFRYIADNDDDKMVILPV
B038        MLHQLKELAAVAGIHMILIYLCRFLLRRSRNVLFTVSNSLRFRLKVLTVLLYICLSVMLFYLFGSIMPLPPWGLVVGWVMALIAVELAYAFIFPYSFRYIADNDDDKMVILPV
B039        MLHHLKELAAVAGIHMILIYLCRFLLRRSRNVLFTVSNSLRFRLKVLTVLLYICLSVMLFYLFGSIMPLPPWGLVVGWVMALIAVELAYAFIFPYSFRYIADNDDDKMVILPV
B040        MLHHLKELAAVAGIHMILIYLCRFLLRRSRNVLFTVSNSLRFRLKVLTVLLYICLSVMLFYLFGSIMPLPPWGLVVGWVMALIAVELAYAFIFPYSFRYIADNDDDKMVILPV
B043        MLHHLKELAAVAGIHMILIYLCRFLLRRSRNVLFTVSNSLRFRLKVLTVLLYICLSVMLFYLFGSIMPLPPWGLVVGWVMALIAVELAYAFIFPYSFRYIADNDDDKMVILPV
B044        MLHHLKELAAVAGIHMILIYLCRFLLRRSRNVLFTVSNSLRFRLKVLTVLLYICLSVMLFYLFGSIMPLPPWGLVVGWVMALIAVELAYAFIFPYSFRYIADNDDDKMVILPV
B045        MLHHLKELAAVAGIHMILIYLCRFLLRRSRNVLFTVSNSLRFRLKVLTVLLYICLSVMLFYLFGSIMPLPPWGLVVGWVMALIAVELAYAFIFPYSFRYIADNDDDKMVILPV
B046        MLHHLKELAAVAGIHMILIYLCRFLLRRSRNVLFTVSNSLRFRLKVLTVLLYICLSVMLFYLFGSIMPLPPWGLVVGWVMALIAVELAYAFIFPYSFRYIADNDDDKMVILPV
B047        MLHHLKELAAVAGIHMILIYLCRFLLRRSRNVLFTVSNSLRFRLKVLTVLLYICLSVMLFYLFGSIMPLPPWGLVVGWVMALIAVELAYAFIFPYSFRYIADNDDDKMVILPV
B048        MLHHLKELAAVAGIHMILIYLCRFLLRRSRNVLFTVSNSLRFRLKVLTVLLYICLSVMLFYLFGSIMPLPPWGLVVGWVMALIAVELAYAFIFPYSFRYIADNDDDKMVILPV
B049        MLHHLKELAAVAGIHMILIYLCRFLLRRSRNVLFTVSNSLRFRLKVLTVLLYICLSVMLFYLFGSIMPLPPWGLVVGWVMALIAVELAYAFIFPYSFRYIADNDDDKMVILPV
B051        MLHHLKELAAVAGIHMILIYLCRFLLRRSRNVLFTVSNSLRFRLKVLTVLLYICLSVMLFYLFGSIMPLPPWGLVVGWVMALIAVELAYAFIFPYSFRYIADNDDDKMVILPV
B052        MLHHLKELAAVAGIHMILIYLCRFLLRRSRNVLFTVSNSLRFRLKVLTVLLYICLSVMLFYLFGSIMPLPPWGLVVGWVMALIAVELAYAFIFPYSFRYIADNDDDKMVILPV
B053        MLHHLKELAAVAGIHMILIYLCRFLLRRSRNVLFTVSNSLRFRLKVLTVLLYICLSVMLFYLFGSIMPLPPWGLVVGWVMALIAVELAYAFIFPYSFRYIADNDDDKMVILPV
B054        MLHHLKELAAVAGIHMILIYLCRFLLRRSRNVLFTVSNSLRFRLKVLTVLLYICLSVMLFYLFGSIMPLPPWGLVVGWVMALIAVELAYAFIFPYSFRYIADNDDDKMVILPV
B055        MLHHLKELAAVAGIHMILIYLCRFLLRRSRNVLFTVSNSLRFRLKVLTVLLYICLSVMLFYLFGSIMPLPPWGLVVGWVMALIAVELAYAFIFPYSFRYIADNDDDKMVILPV
B056        MLHHLKELAAVAGIHMILIYLCRFLLRRSRNVLFTVSNSLRFRLKVLTVLLYICLSVMLFYLFGSIMPLPPWGLVVGWVMALIAVELAYAFIFPYSFRYIADNDDDKMVILPV
B057        MLHHLKELAAVAGIHMILIYLCRFLLRRSRNVLFTVSNSLRFRLKVLTVLLYICLSVMLFYLFGSIMPLPPWGLVVGWVMALIAVELAYAFIFPYSFRYIADNDDDKMVILPV
B058        MLHHLKELAAVAGIHMILIYLCRFLLRRSRNVLFTVSNSLRFRLKVLTVLLYICLSVMLFYLFGSIMPLPPWGLVVGWVMALIAVELAYAFIFPYSFRYIADNDDDKMVILPV
B059        MLHHLKELAAVAGIHMILIYLCRFLLRRSRNVLFTVSNSLRFRLKVLTVLLYICLSVMLFYLFGSIMPLPPWGLVVGWVMALIAVELAYAFIFPYSFRYIADNDDDKMVILPV
B060        MLHHLKELAAVAGIHMILIYLCRFLLRRSRNVLFTVSNSLRFRLKVLTVLLYICLSVMLFYLFGSIMPLPPWGLVVGWVMALIAVELAYAFIFPYSFRYIADNDDDKMVILPV
B061        MLHHLKELAAVAGIHMILIYLCRFLLRRSRNVLFTVSNSLRFRLKVLTVLLYICLSVMLFYLFGSIMPLPPWGLVVGWVMALIAVELAYAFIFPYSFRYIADNDDDKMVILPV
B062        MLHHLKELAAVAGIHMILIYLCRFLLRRSRNVLFTVSNSLRFRLKVLTVLLYICLSVMLFYLFGSIMPLPPWGLVVGWVMALIAVELAYAFIFPYSFRYIADNDDDKMVILPV
B063        MLHHLKELAAVAGIHMILIYLCRFLLRRSRNVLFTVSNSLRFRLKVLTVLLYICLSVMLFYLFGSIMPLPPWGLVVGWVMALIAVELAYAFIFPYSFRYIADNDDDKMVILPV
B064        MLHHLKELAAVAGIHMILIYLCRFLLRRSRNVLFTVSNSLRFRLKVLTVLLYICLSVMLFYLFGSIMPLPPWGLVVGWVMALIAVELAYAFIFPYSFRYIADNDDDKMVILPV
B065        MLHHLKELAAVAGIHMILIYLCRFLLRRSRNVLFTVSNSLRFRLKVLTVLLYICLSVMLFYLFGSIMPLPPWGLVVGWVMALIAVELAYAFIFPYSFRYIADNDDDKMVILPV
B066        MLHQLKELAAVAGIHMILIYLCRFLLRRSRNVLFTVSNSLRFRLKVLTVLLYICLLVMLFYLFGSIMPLPPWGLVVGWVMALIAVELAYAFIFPYSFRYIADNDDDKMVILPV
B067        MLHHLKELAAVAGIHMILIYLCRFLLRRSRNVLFTVSNSLRFRLKVLTVLLYICLSVMLFYLFGSIMPLPPWGLVVGWVMALIAVELAYAFIFPYSFRYIADNDDDKMVILPV
B068        MLHHLKELAAVAGIHMILIYLCRFLLRRSRNVLFTVSNSLRFRLKVLTVLLYICLSVMLFYLFGSIMPLPPWGLVVGWVMALIAVELAYAFIFPYSFRYIADNDDDKMVILPV
B069        MLHHLKELAAVAGIHMILIYLCRFLLRRSRNVLFTVSNSLRFRLKVLTVLLYICLSVMLFYLFGSIMPLPPWGLVVGWVMALIAVELAYAFIFPYSFRYIADNDDDKMVILPV
B070        MLHHLKELAAVAGIHMILIYLCRFLLRRSRNVLFTVSNSLRFRLKVLTVLLYICLSVMLFYLFGSIMPLPPWGLVVGWVMALIAVELAYAFIFPYSFRYIADNDDDKMVILPV
B071        MLHHLKELAAVAGIHMILIYLCRFLLRRSRNVLFTVSNSLRFRLKVLTVLLYICLSVMLFYLFGSIMPLPPWGLVVGWVMALIAVELAYAFIFPYSFRYIADNDDDKMVILPV
B072        MLHHLKELAAVAGIHMILIYLCRFLLRRSRNVLFTVSNSLRFRLKVLTVLLYICLSVMLFYLFGSIMPLPPWGLVVGWVMALIAVELAYAFIFPYSFRYIADNDDDKMVILPV
B073        MLHHLKELAAVAGIHMILIYLCRFLLRRSRNVLFTVSNSLRFRLKVLTVLLYICLSVMLFYLFGSIMPLPPWGLVVGWVMALIAVELAYAFIFPYSFRYIADNDDDKMVILPV
B074        MLHHLKELAAVAGIHMILIYLCRFLLRRSRNVLFTVSNSLRFRLKVLTVLLYICLSVMLFYLFGSIMPLPPWGLVVGWVMALIAVELAYAFIFPYSFRYIADNDDDKMVILPV
B075        MLHHLKELAAVAGIHMILIYLCRFLLRRSRNVLFTVSNSLRFRLKVLTVLLYICLSVMLFYLFGSIMPLPPWGLVVGWVMALIAVELAYAFIFPYSFRYIADNDDDKMVILPV
B076        MLHHLKELAAVAGIHMILIYLCRFLLRRSRNVLFTVSNSLRFRLKVLTVLLYICLSVMLFYLFGSIMPLPPWGLVVGWVMALIAVELAYAFIFPYSFRYIADNDDDKMVILPV
B077        MLHHLKELAAVAGIHMILIYLCRFLLRRSRNVLFTVSNSLRFRLKVLTVLLYICLSVMLFYLFGSIMPLPPWGLVVGWVMALIAVELAYAFIFPYSFRYIADNDDDKMVILPV
B079        MLHHLKELAAVAGIHMILIYLCRFLLRRSRNVLFTVSNSLRFRLKVLTVLLYICLSVMLFYLFGSIMPLPPWGLVVGWVMALIAVELAYAFIFPYSFRYIADNDDDKMVILPV
B081        MLHHLKELAAVAGIHMILIYLCRFLLRRSRNVLFTVSNSLRFRLKVLTVLLYICLSVMLFYLFGSIMPLPPWGLVVGWVMALIAVELAYAFIFPYSFRYIADNDDDKMVILPV
B082        MLHHLKELAAVAGIHMILIYLCRFLLRRSRNVLFTVSNSLRFRLKVLTVLLYICLSVMLFYLFGSIMPLPPWGLVVGWVMALIAVELAYAFIFPYSFRYIADNDDDKMVILPV
B083        MLHHLKELAAVAGIHMILIYLCRFLLRRSRNVLFTVSNSLRFRLKVLTVLLYICLSVMLFYLFGSIMPLPPWGLVVGWVMALIAVELAYAFIFPYSFRYIADNDDDKMVILPV
B084        MLHHLKELAAVAGIHMILIYLCRFLLRRSRNVLFTVSNSLRFRLKVLTVLLYICLSVMLFYLFGSIMPLPPWGLVVGWVMALIAVELAYAFIFPYSFRYIADNDDDKMVILPV
B085        MLHHLKELAAVAGIHMILIYLCRFLLRRSRNVLFTVSNSLRFRLKVLTVLLYICLSVMLFYLFGSIMPLPPWGLVVGWVMALIAVELAYAFIFPYSFRYIADNDDDKMVILPV
B086        MLHHLKELAAVAGIHMILIYLCRFLLRRSRNVLFTVSNSLRFRLKVLTVLLYICLSVMLFYLFGSIMPLPPWGLVVGWVMALIAVELAYAFIFPYSFRYIADNDDDKMVILPV
B087        MLHHLKELAAVAGIHMILIYLCRFLLRRSRNVLFTVSNSLRFRLKVLTVLLYICLSVMLFYLFGSIMPLPPWGLVVGWVMALIAVELAYAFIFPYSFRYIADNDDDKMVILPV
B088        MLHHLKELAAVAGIHMILIYLCRFLLRRSRNVLFTVSNSLRFRLKVLTVLLYICLSVMLFYLFGSIMPLPPWGLVVGWVMALIAVELAYAFIFPYSFRYIADNDDDKMVILPV
B089        MLHHLKELAAVAGIHMILIYLCRFLLRRSRNVLFTVSNSLRFRLKVLTVLLYICLSVMLFYLFGSIMPLPPWGLVVGWVMALIAVELAYAFIFPYSFRYIADNDDDKMVILPV
B090        MLHHLKELAAVAGIHMILIYLCRFLLRRSRNVLFTVSNSLRFRLKVLTVLLYICLSVMLFYLFGSIMPLPPWGLVVGWVMALIAVELAYAFIFPYSFRYIADNDDDKMVILPV
B091        MLHHLKELAAVAGIHMILIYLCRFLLRRSRNVLFTVSNSLRFRLKVLTVLLYICLSVMLFYLFGSIMPLPPWGLVVGWVMALIAVELAYAFIFPYSFRYIADNDDDKMVILPV
B092        MLHHLKELAAVAGIHMILIYLCRFLLRRSRNVLFTVSNSLRFRLKVLTVLLYICLSVMLFYLFGSIMPLPPWGLVVGWVMALIAVELAYAFIFPYSFRYIADNDDDKMVILPV
B093        MLHHLKELAAVAGIHMILIYLCRFLLRRSRNVLFTVSNSLRFRLKVLTVLLYICLSVMLFYLFGSIMPLPPWGLVVGWVMALIAVELAYAFIFPYSFRYIADNDDDKMVILPV
B094        MLHHLKELAAVAGIHMILIYLCRFLLRRSRNVLFTVSNSLRFRLKVLTVLLYICLSVMLFYLFGSIMPLPPWGLVVGWVMALIAVELAYAFIFPYSFRYIADNDDDKMVILPV
B095        MLHHLKELAAVAGIHMILIYLCRFLLRRSRNVLFTVSNSLRFRLKVLTVLLYICLSVMLFYLFGSIMPLPPWGLVVGWVMALIAVELAYAFIFPYSFRYIADNDDDKMVILPV
B096        MLHHLKELAAVAGIHMILIYLCRFLLRRSRNVLFTVSNSLRFRLKVLTVLLYICLSVMLFYLFGSIMPLPPWGLVVGWVMALIAVELAYAFIFPYSFRYIADNDDDKMVILPV
B097        MLHHLKELAAVAGIHMILIYLCRFLLRRSRNVLFTVSNSLRFRLKVLTVLLYICLSVMLFYLFGSIMPLPPWGLVVGWVMALIAVELAYAFIFPYSFRYIADNDDDKMVILPV
B100        MLHQLKELAAVAGIHMILIYLCRFLLRRSRNVLFTVSNSLRFRLKVLTVLLYICLLVMLFYLFGSIMPLPPWGLVVGWVMALIAVELAYAFIFPYSFRYIADNDDDKMVILPV
B102        MLHHLKELAAVAGIHMILIYLCRFLLRRSRNVLFTVSNSLRFRLKVLTVLLYICLSVMLFYLFGSIMPLPPWGLVVGWVMALIAVELAYAFIFPYSFRYIADNDDDKMVILPV
B103        MLHQLKELAAVAGIHMILIYLCRFLLRRSRNVLFTVSNSLRFRLKVLTVLLYICLLVMLFYLFGSIMPLPPWGLVVGWVMALIAVELAYAFIFPYSFRYIADNDDDKMVILPV
B104        MLHHLKELAAVAGIHMILIYLCRFLLRRSRNVLFTVSNSLRFRLKVLTVLLYICLSVMLFYLFGSIMPLPPWGLVVGWVMALIAVELAYAFIFPYSFRYIADNDDDKMVILPV
B105        MLHHLKELAAVAGIHMILIYLCRFLLRRSRNVLFTVSNSLRFRLKVLTVLLYICLSVMLFYLFGSIMPLPPWGLVVGWVMALIAVELAYAFIFPYSFRYIADNDDDKMVILPV
B106        MLHHLKELAAVAGIHMILIYLCRFLLRRSRNVLFTVSNSLRFRLKVLTVLLYICLSVMLFYLFGSIMPLPPWGLVVGWVMALIAVELAYAFIFPYSFRYIADNDDDKMVILPV
B107        MLHHLKELAAVAGIHMILIYLCRFLLRRSRNVLFTVSNSLRFRLKVLTVLLYICLSVMLFYLFGSIMPLPPWGLVVGWVMALIAVELAYAFIFPYSFRYIADNDDDKMVILPV
B108        MLHHLKELAAVAGIHMILIYLCRFLLRRSRNVLFTVSNSLRFRLKVLTVLLYICLSVMLFYLFGSIMPLPPWGLVVGWVMALIAVELAYAFIFPYSFRYIADNDDDKMVILPV
B109        MLHHLKELAAVAGIHMILIYLCRFLLRRSRNVLFTVSNSLRFRLKVLTVLLYICLSVMLFYLFGSIMPLPPWGLVVGWVMALIAVELAYAFIFPYSFRYIADNDDDKMVILPV
B110        MLHHLKELAAVAGIHMILIYLCRFLLRRSRNVLFTVSNSLRFRLKVLTVLLYICLSVMLFYLFGSIMPLPPWGLVVGWVMALIAVELAYAFIFPYSFRYIADNDDDKMVILPV
B111        MLHHLKELAAVAGIHMILIYLCRFLLRRSRNVLFTVSNSLRFRLKVLTVLLYICLSVMLFYLFGSIMPLPPWGLVVGWVMALIAVELAYAFIFPYSFRYIADNDDDKMVILPV
B112        MLHHLKELAAVAGIHMILIYLCRFLLRLSRNVLFTVSNSLRFRLKVLTVLLYICLSVMLFYLFGSIMPLPPWGLVVGWVMALIAVELAYAFIFPYSFRYIADNDDDKMVILPV
B113        MLHHLKELAAVAGIHMILIYLCRFLLRRSRNVLFTVSNSLRFRLKVLTVLLYICLSVMLFYLFGSIMPLPPWGLVVGWVMALIAVELAYAFIFPYSFRYIADNDDDKMVILPV
B114        MLHHLKELAAVAGIHMILIYLCRFLLRRSRNVLFTVSNSLRFRLKVLTVLLYICLSVMLFYLFGSIMPLPPWGLVVGWVMALIAVELAYAFIFPYSFRYIADNDDDKMVILPV
B115        MLHHLKELAAVAGIHMILIYLCRFLLRRSRNVLFTVSNSLRFRLKVLTVLLYICLSVMLFYLFGSIMPLPPWGLVVGWVMALIAVELAYAFIFPYSFRYIADNDDDKMVILPV
B116        MLHHLKELAAVAGIHMILIYLCRFLLRRSRNVLFTVSNSLRFRLKVLTVLLYICLSVMLFYLFGSIMPLPPWGLVVGWVMALIAVELAYAFIFPYSFRYIADNDDDKMVILPV
B117        MLHHLKELAAVAGIHMILIYLCRFLLRRSRNVLFTVSNSLRFRLKVLTVLLYICLSVMLFYLFGSIMPLPPWGLVVGWVMALIAVELAYAFIFPYSFRYIADNDDDKMVILPV
```

```
B118        MLHHLKELAAVAGIHMILIYLCRFLLRRSRNVLFTVSNSLRFRLKVLTVLLYICLSVMLFYLFGSIMPLPPWGLVVGWVMALIAVELAYAFIFPYSFRYIADNDDDKMVILPV
B119        MLHHLKELAAVAGIHMILIYLCRFLLRRSRNVLFTVSNSLRFRLKVLTVLLYICLSVMLFYLFGSIMPLPPWGLVVGWVMALIAVELAYAFIFPYSFRYIADNDDDKMVILPV
B120        MLHHLKELAAVAGIHMILIYLCRFLLRRSRNVLFTVSNSLRFRLKVLTVLLYICLSVMLFYLFGSIMPLPPWGLVVGWVMALIAVELAYAFIFPYSFRYIADNDDDKMVILPV
B121        MLHHLKELAAVAGIHMILIYLCRFLLRRSRNVLFTVSNSLRFRLKVLTVLLYICLSVMLFYLFGSIMPLPPWGLVVGWVMALIAVELAYAFIFPYSFRYIADNDDDKMVILPV
B122        MLHHLKELAAVAGIHMILIYLCRFLLRRSRNVLFTVSNSLRFRLKVLTVLLYICLSVMLFYLFGSIMPLPPWGLVVGWVMALIAVELAYAFIFPYSFRYIADNDDDKMVILPV
B123        MLHHLKELAAVAGIHMILIYLCRFLLRRSRNVLFTVSNSLRFRLKVLTVLLYICLSVMLFYLFGSIMPLPPWGLVVGWVMALIAVELAYAFIFPYSFRYIADNDDDKMVILPV
B124        MLHHLKELAAVAGIHMILIYLCRFLLRRSRNVLFTVSNSLRFRLKVLTVLLYICLSVMLFYLFGSIMPLPPWGLVVGWVMALIAVELAYAFIFPYSFRYIADNDDDKMVILPV
B125        MLHHLKELAAVAGIHMILIYLCRFLLRRSRNVLFTVSNSLRFRLKVLTVLLYICLSVMLFYLFGSIMPLPPWGLVVGWVMALIAVELAYAFIFPYSFRYIADNDDDKMVILPV
B126        MLHHLKELAAVAGIHMILIYLCRFLLRRSRNVLFTVSNSLRFRLKVLTVLLYICLSVMLFYLFGSIMPLPPWGLVVGWVMALIAVELAYAFIFPYSFRYIADNDDDKMVILPV
B127        MLHHLKELAAVAGIHMILIYLCRFLLRRSRNVLFTVSNSLRFRLKVLTVLLYICLSVMLFYLFGSIMPLPPWGLVVGWVMALIAVELAYAFIFPYSFRYIADNDDDKMVILPV
B128        MLHHLKELAAVAGIHMILIYLCRFLLRRSRNVLFTVSNSLRFRLKVLTVLLYICLSVMLFYLFGSIMPLPPWGLVVGWVMALIAVELAYAFIFPYSFRYIADNDDDKMVILPV
B129        MLHHLKELAAVAGIHMILIYLCRFLLRRSRNVLFTVSNSLRFRLKVLTVLLYICLSVMLFYLFGSIMPLPPWGLVVGWVMALIAVELAYAFIFPYSFRYIADNDDDKMVILPV
B130        MLHHLKELAAVAGIHMILIYLCRFLLRRSRNVLFTVSNSLRFRLKVLTVLLYICLSVMLFYLFGSIMPLPPWGLVVGWVMALIAVELAYAFIFPYSFRYIADNDDDKMVILPV
B131        MLHHLKELAAVAGIHMILIYLCRFLLRRSRNVLFTVSNSLRFRLKVLTVLLYICLSVMLFYLFGSIMPLPPWGLVVGWVMALIAVELAYAFIFPYSFRYIADNDDDKMVILPV
B132        MLHHLKELAAVAGIHMILIYLCRFLLRRSRNVLFTVSNSLRFRLKVLTVLLYICLSVMLFYLFGSIMPLPPWGLVVGWVMALIAVELAYAFIFPYSFRYIADNDDDKMVILPV
B133        MLHHLKELAAVAGIHMILIYLCRFLLRRSRNVLFTVSNSLRFRLKVLTVLLYICLSVMLFYLFGSIMPLPPWGLVVGWVMALIAVELAYAFIFPYSFRYIADNDDDKMVILPV
B134        MLHQLKELAAVAGIHMILIYLCRFLLRRSRNVLFTVSNSLRFRLKVLTVLLYICLLVMLFYLFGSIMPLPPWGLVVGWVMALIAVELAYAFIFPYSFRYIADNDDDKMVILPV
B135        MLHHLKELAAVAGIHMILIYLCRFLLRRSRNVLFTVSNSLRFRLKVLTVLLYICLSVMLFYLFGSIMPLPPWGLVVGWVMALIAVELAYAFIFPYSFRYIADNDDDKMVILPV
B136        MLHHLKELAAVAGIHMILIYLCRFLLRRSRNVLFTVSNSLRFRLKVLTVLLYICLSVMLFYLFGSIMPLPPWGLVVGWVMALIAVELAYAFIFPYSFRYIADNDDDKMVILPV
B137        MLHHLKELAAVAGIHMILIYLCRFLLRRSRNVLFTVSNSLRFRLKVLTVLLYICLSVMLFYLFGSIMPLPPWGLVVGWVMALIAVELAYAFIFPYSFRYIADNDDDKMVILPV
B138        MLHHLKELAAVAGIHMILIYLCRFLLRRSRNVLFTVSNSLRFRLKVLTVLLYICLSVMLFYLFGSIMPLPPWGLVVGWVMALIAVELAYAFIFPYSFRYIADNDDDKMVILPV
B139        MLHHLKELAAVAGIHMILIYLCRFLLRRSRNVLFTVSNSLRFRLKVLTVLLYICLSVMLFYLFGSIMPLPPWGLVVGWVMALIAVELAYAFIFPYSFRYIADNDDDKMVILPV
B141        MLHHLKELAAVAGIHMILIYLCRFLLRRSRNVLFTVSNSLRFRLKVLTVLLYICLSVMLFYLFGSIMPLPPWGLVVGWVMALIAVELAYAFIFPYSFRYIADNDDDKMVILPV
B142        MLHHLKELAAVAGIHMILIYLCRFLLRRSRNVLFTVSNSLRFRLKVLTVLLYICLSVMLFYLFGSIMPLPPWGLVVGWVMALIAVELAYAFIFPYSFRYIADNDDDKMVILPV
B143        MLHHLKELAAVAGIHMILIYLCRFLLRRSRNVLFTVSNSLRFRLKVLTVLLYICLSVMLFYLFGSIMPLPPWGLVVGWVMALIAVELAYAFIFPYSFRYIADNDDDKMVILPV
B144        MLHQLKELAAVAGIHMILIYLCRFLLRRSRNVLFTVSNSLRFRLKVLTVLLYICLSVMLFYLFGSIMPLPPWGLVVGWVMALIAVELAYAFIFPYSFRYIADNDDDKMVILPV
B145        MLHHLKELAAVAGIHMILIYLCRFLLRRSRNVLFTVSNSLRFRLKVLTVLLYICLSVMLFYLFGSIMPLPPWGLVVGWVMALIAVELAYAFIFPYSFRYIADNDDDKMVILPV
B146        MLHHLKELAAVAGIHMILIYLCRFLLRRSRNVLFTVSNSLRFRLKVLTVLLYICLSVMLFYLFGSIMPLPPWGLVVGWVMALIAVELAYAFIFPYSFRYIADNDDDKMVILPV
B147        MLHHLKELAAVAGIHMILIYLCRFLLRRSRNVLFTVSNSLRFRLKVLTVLLYICLSVMLFYLFGSIMPLPPWGLVVGWVMALIAVELAYAFIFPYSFRYIADNDDDKMVILPV
B148        MLHHLKELAAVAGIHMILIYLCRFLLRRSRNVLFTVSNSLRFRLKVLTVLLYICLSVMLFYLFGSIMPLPPWGLVVGWVMALIAVELAYAFIFPYSFRYIADNDDDKMVILPV
B149        MLHHLKELAAVAGIHMILIYLCRFLLRRSRNVLFTVSNSLRFRLKVLTVLLYICLSVMLFYLFGSIMPLPPWGLVVGWVMALIAVELAYAFIFPYSFRYIADNDDDKMVILPV
B150        MLHHLKELAAVAGIHMILIYLCRFLLRRSRNVLFTVSNSLRFRLKVLTVLLYICLSVMLFYLFGSIMPLPPWGLVVGWVMALIAVELAYAFIFPYSFRYIADNDDDKMVILPV
B151        MLHHLKELAAVAGIHMILIYLCRFLLRRSRNVLFTVSNSLRFRLKVLTVLLYICLSVMLFYLFGSIMPLPPWGLVVGWVMALIAVELAYAFIFPYSFRYIADNDDDKMVILPV
B152        MLHHLKELAAVAGIHMILIYLCRFLLRRSRNVLFTVSNSLRFRLKVLTVLLYICLSVMLFYLFGSIMPLPPWGLVVGWVMALIAVELAYAFIFPYSFRYIADNDDDKMVILPV
B153        MLHHLKELAAVAGIHMILIYLCRFLLRRSRNVLFTVSNSLRFRLKVLTVLLYICLSVMLFYLFGSIMPLPPWGLVVGWVMALIAVELAYAFIFPYSFRYIADNDDDKMVILPV
B154        MLHHLKELAAVAGIHMILIYLCRFLLRRSRNVLFTVSNSLRFRLKVLTVLLYICLSVMLFYLFGSIMPLPPWGLVVGWVMALIAVELAYAFIFPYSFRYIADNDDDKMVILPV
B155        MLHHLKELAAVAGIHMILIYLCRFLLRRSRNVLFTVSNSLRFRLKVLTVLLYICLSVMLFYLFGSIMPLPPWGLVVGWVMALIAVELAYAFIFPYSFRYIADNDDDKMVILPV
B156        MLHHLKELAAVAGIHMILIYLCRFLLRRSRNVLFTVSNSLRFRLKVLTVLLYICLSVMLFYLFGSIMPLPPWGLVVGWVMALIAVELAYAFIFPYSFRYIADNDDDKMVILPV
B157        MLHHLKELAAVAGIHMILIYLCRFLLRRSRNVLFTVSNSLRFRLKVLTVLLYICLSVMLFYLFGSIMPLPPWGLVVGWVMALIAVELAYAFIFPYSFRYIADNDDDKMVILPV
B158        MLHHLKELAAVAGIHMILIYLCRFLLRRSRNVLFTVSNSLRFRLKVLTVLLYICLSVMLFYLFGSIMPLPPWGLVVGWVMALIAVELAYAFIFPYSFRYIADNDDDKMVILPV
B159        MLHHLKELAAVAGIHMILIYLCRFLLRRSRNVLFTVSNSLRFRLKVLTVLLYICLSVMLFYLFGSIMPLPPWGLVVGWVMALIAVELAYAFIFPYSFRYIADNDDDKMVILPV
B160        MLHHLKELAAVAGIHMILIYLCRFLLRRSRNVLFTVSNSLRFRLKVLTVLLYICLSVMLFYLFGSIMPLPPWGLVVGWVMALIAVELAYAFIFPYSFRYIADNDDDKMVILPV
B161        MLHHLKELAAVAGIHMILIYLCRFLLRRSRNVLFTVSNSLRFRLKVLTVLLYICLSVMLFYLFGSIMPLPPWGLVVGWVMALIAVELAYAFIFPYSFRYIADNDDDKMVILPV
B162        MLHHLKELAAVAGIHMILIYLCRFLLRRSRNVLFTVSNSLRFRLKVLTVLLYICLSVMLFYLFGSIMPLPPWGLVVGWVMALIAVELAYAFIFPYSFRYIADNDDDKMVILPV
B163        MLHHLKELAAVAGIHMILIYLCRFLLRRSRNVLFTVSNSLRFRLKVLTVLLYICLSVMLFYLFGSIMPLPPWGLVVGWVMALIAVELAYAFIFPYSFRYIADNDDDKMVILPV
B164        MLHHLKELAAVAGIHMILIYLCRFLLRRSRNVLFTVSNSLRFRLKVLTVLLYICLSVMLFYLFGSIMPLPPWGLVVGWVMALIAVELAYAFIFPYSFRYIADNDDDKMVILPV
B165        MLHHLKELAAVAGIHMILIYLCRFLLRRSRNVLFTVSNSLRFRLKVLTVLLYICLSVMLFYLFGSIMPLPPWGLVVGWVMALIAVELAYAFIFPYSFRYIADNDDDKMVILPV
B167        MLHHLKELAAVAGIHMILIYLCRFLLRRSRNVLFTVSNSLRFRLKVLTVLLYICLSVMLFYLFGSIMPLPPWGLVVGWVMALIAVELAYAFIFPYSFRYIADNDDDKMVILPV
B168        MLHHLKELAAVAGIHMILIYLCRFLLRRSRNVLFTVSNSLRFRLKVLTVLLYICLSVMLFYLFGSIMPLPPWGLVVGWVMALIAVELAYAFIFPYSFRYIADNDDDKMVILPV
B169        MLHHLKELAAVAGIHMILIYLCRFLLRRSRNVLFTVSNSLRFRLKVLTVLLYICLSVMLFYLFGSIMPLPPWGLVVGWVMALIAVELAYAFIFPYSFRYIADNDDDKMVILPV
B170        MLHHLKELAAVAGIHMILIYLCRFLLRRSRNVLFTVSNSLRFRLKVLTVLLYICLSVMLFYLFGSIMPLPPWGLVVGWVMALIAVELAYAFIFPYSFRYIADNDDDKMVILPV
B171        MLHHLKELAAVAGIHMILIYLCRFLLRRSRNVLFTVSNSLRFRLKVLTVLLYICLSVMLFYLFGSIMPLPPWGLVVGWVMALIAVELAYAFIFPYSFRYIADNDDDKMVILPV
B173        MLHHLKELAAVAGIHMILIYLCRFLLRRSRNVLFTVSNSLRFRLKVLTVLLYICLSVMLFYLFGSIMPLPPWGLVVGWVMALIAVELAYAFIFPYSFRYIADNDDDKMVILPV
B176        MLHHLKELAAVAGIHMILIYLCRFLLRRSRNVLFTVSNSLRFRLKVLTVLLYICLSVMLFYLFGSIMPLPPWGLVVGWVMALIAVELAYAFIFPYSFRYIADNDDDKMVILPV
B179        MLHHLKELAAVAGIHMILIYLCRFLLRRSRNVLFTVSNSLRFRLKVLTVLLYICLSVMLFYLFGSIMPLPPWGLVVGWVMALIAVELAYAFIFPYSFRYIADNDDDKMVILPV
B180        MLHHLKELAAVAGIHMILIYLCRFLLRLSRNVLFTVSNSLRFRLKVLTVLLYICLSVMLFYLFGSIMPLPPWGLVVGWVMALIAVELAYAFIFPYSFRYIADNDDDKMVILPV
B182        MLHHLKELAAVAGIHMILIYLCRFLLRRSRNVLFTVSNSLRFRLKVLTVLLYICLSVMLFYLFGSIMPLPPWGLVVGWVMALIAVELAYAFIFPYSFRYIADNDDDKMVILPV
B183        MLHHLKELAAVAGIHMILIYLCRFLLRRSRNVLFTVSNSLRFRLKVLTVLLYICLSVMLFYLFGSIMPLPPWGLVVGWVMALIAVELAYAFIFPYSFRYIADNDDDKMVILPV
B184        MLHHLKELAAVAGIHMILIYLCRFLLRRSRNVLFTVSNSLRFRLKVLTVLLYICLSVMLFYLFGSIMPLPPWGLVVGWVMALIAVELAYAFIFPYSFRYIADNDDDKMVILPV
B185        MLHHLKELAAVAGIHMILIYLCRFLLRRSRNVLFTVSNSLRFRLKVLTVLLYICLSVMLFYLFGSIMPLPPWGLVVGWVMALIAVELAYAFIFPYSFRYIADNDDDKMVILPV
B187        MLHHLKELAAVAGIHMILIYLCRFLLRRSRNVLFTVSNSLRFRLKVLTVLLYICLSVMLFYLFGSIMPLPPWGLVVGWVMALIAVELAYAFIFPYSFRYIADNDDDKMVILPV
B188        MLHQLKELAAVAGIHMILIYLCRFLLRRSRNVLFTVSNSLRFRLKVLTVLLYICLLVMLFYLFGSIMPLPPWGLVVGWVMALIAVELAYAFIFPYSFRYIADNDDDKMVILPV
B189        MLHHLKELAAVAGIHMILIYLCRFLLRRSRNVLFTVSNSLRFRLKVLTVLLYICLSVMLFYLFGSIMPLPPWGLVVGWVMALIAVELAYAFIFPYSFRYIADNDDDKMVILPV
B190        MLHHLKELAAVAGIHMILIYLCRFLLRRSRNVLFTVSNSLRFRLKVLTVLLYICLSVMLFYLFGSIMPLPPWGLVVGWVMALIAVELAYAFIFPYSFRYIADNDDDKMVILPV
B191        MLHHLKELAAVAGIHMILIYLCRFLLRRSRNVLFTVSNSLRFRLKVLTVLLYICLSVMLFYLFGSIMPLPPWGLVVGWVMALIAVELAYAFIFPYSFRYIADNDDDKMVILPV
B192        MLHHLKELAAVAGIHMILIYLCRFLLRRSRNVLFTVSNSLRFRLKVLTVLLYICLSVMLFYLFGSIMPLPPWGLVVGWVMALIAVELAYAFIFPYSFRYIADNDDDKMVILPV
B193        MLHHLKELAAVAGIHMILIYLCRFLLRRSRNVLFTVSNSLRFRLKVLTVLLYICLSVMLFYLFGSIMPLPPWGLVVGWVMALIAVELAYAFIFPYSFRYIADNDDDKMVILPV
B194        MLHHLKELAAVAGIHMILIYLCRFLLRRSRNVLFTVSNSLRFRLKVLTVLLYICLSVMLFYLFGSIMPLPPWGLVVGWVMALIAVELAYAFIFPYSFRYIADNDDDKMVILPV
B195        MLHHLKELAAVAGIHMILIYLCRFLLRRSRNVLFTVSNSLRFRLKVLTVLLYICLSVMLFYLFGSIMPLPPWGLVVGWVMALIAVELAYAFIFPYSFRYIADNDDDKMVILPV
B196        MLHHLKELAAVAGIHMILIYLCRFLLRRSRNVLFTVSNSLRFRLKVLTVLLYICLSVMLFYLFGSIMPLPPWGLVVGWVMALIAVELAYAFIFPYSFRYIADNDDDKMVILPV
B197        MLHHLKELAAVAGIHMILIYLCRFLLRRSRNVLFTVSNSLRFRLKVLTVLLYICLSVMLFYLFGSIMPLPPWGLVVGWVMALIAVELAYAFIFPYSFRYIADNDDDKMVILPV
B198        MLHHLKELAAVAGIHMILIYLCRFLLRRSRNVLFTVSNSLRFRLKVLTVLLYICLSVMLFYLFGSIMPLPPWGLVVGWVMALIAVELAYAFIFPYSFRYIADNDDDKMVILPV
B199        MLHHLKELAAVAGIHMILIYLCRFLLRLSRNVLFTVSNSLRFRLKVLTVLLYICLSVMLFYLFGSIMPLPPWGLVVGWVMALIAVELAYAFIFPYSFRYIADNDDDKMVILPV
B200        MLHHLKELAAVAGIHMILIYLCRFLLRRSRNVLFTVSNSLRFRLKVLTVLLYICLSVMLFYLFGSIMPLPPWGLVVGWVMALIAVELAYAFIFPYSFRYIADNDDDKMVILPV
B201        MLHHLKELAAVAGIHMILIYLCRFLLRRSRNVLFTVSNSLRFRLKVLTVLLYICLSVMLFYLFGSIMPLPPWGLVVGWVMALIAVELAYAFIFPYSFRYIADNDDDKMVILPV
B202        MLHHLKELAAVAGIHMILIYLCRFLLRRSRNVLFTVSNSLRFRLKVLTVLLYICLSVMLFYLFGSIMPLPPWGLVVGWVMALIAVELAYAFIFPYSFRYIADNDDDKMVILPV
B203        MLHHLKELAAVAGIHMILIYLCRFLLRRSRNVLFTVSNSLRFRLKVLTVLLYICLSVMLFYLFGSIMPLPPWGLVVGWVMALIAVELAYAFIFPYSFRYIADNDDDKMVILPV
B204        MLHHLKELAAVAGIHMILIYLCRFLLRRSRNVLFTVSNSLRFRLKVLTVLLYICLSVMLFYLFGSIMPLPPWGLVVGWVMALIAVELAYAFIFPYSFRYIADNDDDKMVILPV
B205        MLHQLKELAAVAGIHMILIYLCRFLLRRSRNVLFTVSNSLRFRLKVLTVLLYICLLVMLFYLFGSIMPLPPWGLVVGWVMALIAVELAYAFIFPYSFRYIADNDDDKMVILPV
B207        MLHHLKELAAVAGIHMILIYLCRFLLRRSRNVLFTVSNSLRFRLKVLTVLLYICLSVMLFYLFGSIMPLPPWGLVVGWVMALIAVELAYAFIFPYSFRYIADNDDDKMVILPV
B208        MLHHLKELAAVAGIHMILIYLCRFLLRRSRNVLFTVSNSLRFRLKVLTVLLYICLSVMLFYLFGSIMPLPPWGLVVGWVMALIAVELAYAFIFPYSFRYIADNDDDKMVILPV
B210        MLHHLKELAAVAGIHMILIYLCRFLLRRSRNVLFTVSNSLRFRLKVLTVLLYICLSVMLFYLFGSIMPLPPWGLVVGWVMALIAVELAYAFIFPYSFRYIADNDDDKMVILPV
B212        MLHHLKELAAVAGIHMILIYLCRFLLRRSRNVLFTVSNSLRFRLKVLTVLLYICLSVMLFYLFGSIMPLPPWGLVVGWVMALIAVELAYAFIFPYSFRYIADNDDDKMVILPV
B213        MLHHLKELAAVAGIHMILIYLCRFLLRRSRNVLFTVSNSLRFRLKVLTVLLYICLSVMLFYLFGSIMPLPPWGLVVGWVMALIAVELAYAFIFPYSFRYIADNDDDKMVILPV
B214        MLHHLKELAAVAGIHMILIYLCRFLLRRSRNVLFTVSNSLRFRLKVLTVLLYICLSVMLFYLFGSIMPLPPWGLVVGWVMALIAVELAYAFIFPYSFRYIADNDDDKMVILPV
B215        MLHHLKELAAVAGIHMILIYLCRFLLRRSRNVLFTVSNSLRFRLKVLTVLLYICLSVMLFYLFGSIMPLPPWGLVVGWVMALIAVELAYAFIFPYSFRYIADNDDDKMVILPV
B216        MLHHLKELAAVAGIHMILIYLCRFLLRRSRNVLFTVSNSLRFRLKVLTVLLYICLSVMLFYLFGSIMPLPPWGLVVGWVMALIAVELAYAFIFPYSFRYIADNDDDKMVILPV
B217        MLHHLKELAAVAGIHMILIYLCRFLLRRSRNVLFTVSNSLRFRLKVLTVLLYICLSVMLFYLFGSIMPLPPWGLVVGWVMALIAVELAYAFIFPYSFRYIADNDDDKMVILPV
B218        MLHQLKELAAVAGIHMILIYLCRFLLRRSRNVLFTVSNSLRFRLKVLTVLLYICLLVMLFYLFGSIMPLPPWGLVVGWVMALIAVELAYAFIFPYSFRYIADNDDDKMVILPV
B219        MLHHLKELAAVAGIHMILIYLCRFLLRRSRNVLFTVSNSLRFRLKVLTVLLYICLSVMLFYLFGSIMPLPPWGLVVGWVMALIAVELAYAFIFPYSFRYIADNDDDKMVILPV
B221        MLHHLKELAAVAGIHMILIYLCRFLLRRSRNVLFTVSNSLRFRLKVLTVLLYICLSVMLFYLFGSIMPLPPWGLVVGWVMALIAVELAYAFIFPYSFRYIADNDDDKMVILPV
B222        MLHHLKELAAVAGIHMILIYLCRFLLRRSRNVLFTVSNSLRFRLKVLTVLLYICLSVMLFYLFGSIMPLPPWGLVVGWVMALIAVELAYAFIFPYSFRYIADNDDDKMVILPV
B223        MLHHLKELAAVAGIHMILIYLCRFLLRRSRNVLFTVSNSLRFRLKVLTVLLYICLSVMLFYLFGSIMPLPPWGLVVGWVMALIAVELAYAFIFPYSFRYIADNDDDKMVILPV
B224        MLHHLKELAAVAGIHMILIYLCRFLLRRSRNVLFTVSNSLRFRLKVLTVLLYICLSVMLFYLFGSIMPLPPWGLVVGWVMALIAVELAYAFIFPYSFRYIADNDDDKMVILPV
B225        MLHHLKELAAVAGIHMILIYLCRFLLRRSRNVLFTVSNSLRFRLKVLTVLLYICLSVMLFYLFGSIMPLPPWGLVVGWVMALIAVELAYAFIFPYSFRYIADNDDDKMVILPV
B226        MLHHLKELAAVAGIHMILIYLCRFLLRRSRNVLFTVSNSLRFRLKVLTVLLYICLSVMLFYLFGSIMPLPPWGLVVGWVMALIAVELAYAFIFPYSFRYIADNDDDKMVILPV
B227        MLHHLKELAAVAGIHMILIYLCRFLLRRSRNVLFTVSNSLRFRLKVLTVLLYICLSVMLFYLFGSIMPLPPWGLVVGWVMALIAVELAYAFIFPYSFRYIADNDDDKMVILPV
B228        MLHHLKELAAVAGIHMILIYLCRFLLRRSRNVLFTVSNSLRFRLKVLTVLLYICLSVMLFYLFGSIMPLPPWGLVVGWVMALIAVELAYAFIFPYSFRYIADNDDDKMVILPV
B229        MLHHLKELAAVAGIHMILIYLCRFLLRRSRNVLFTVSNSLRFRLKVLTVLLYICLSVMLFYLFGSIMPLPPWGLVVGWVMALIAVELAYAFIFPYSFRYIADNDDDKMVILPV
B230        MLHHLKELAAVAGIHMILIYLCRFLLRRSRNVLFTVSNSLRFRLKVLTVLLYICLSVMLFYLFGSIMPLPPWGLVVGWVMALIAVELAYAFIFPYSFRYIADNDDDKMVILPV
B231        MLHHLKELAAVAGIHMILIYLCRFLLRRSRNVLFTVSNSLRFRLKVLTVLLYICLSVMLFYLFGSIMPLPPWGLVVGWVMALIAVELAYAFIFPYSFRYIADNDDDKMVILPV
B232        MLHHLKELAAVAGIHMILIYLCRFLLRRSRNVLFTVSNSLRFRLKVLTVLLYICLSVMLFYLFGSIMPLPPWGLVVGWVMALIAVELAYAFIFPYSFRYIADNDDDKMVILPV
B233        MLHHLKELAAVAGIHMILIYLCRFLLRRSRNVLFTVSNSLRFRLKVLTVLLYICLSVMLFYLFGSIMPLPPWGLVVGWVMALIAVELAYAFIFPYSFRYIADNDDDKMVILPV
B234        MLHHLKELAAVAGIHMILIYLCRFLLRRSRNVLFTVSNSLRFRLKVLTVLLYICLSVMLFYLFGSIMPLPPWGLVVGWVMALIAVELAYAFIFPYSFRYIADNDDDKMVILPV
B235        MLHHLKELAAVAGIHMILIYLCRFLLRRSRNVLFTVSNSLRFRLKVLTVLLYICLSVMLFYLFGSIMPLPPWGLVVGWVMALIAVELAYAFIFPYSFRYIADNDDDKMVILPV
B236        MLHHLKELAAVAGIHMILIYLCRFLLRRSRNVLFTVSNSLRFRLKVLTVLLYICLSVMLFYLFGSIMPLPPWGLVVGWVMALIAVELAYAFIFPYSFRYIADNDDDKMVILPV
B238        MLHHLKELAAVAGIHMILIYLCRFLLRRSRNVLFTVSNSLRFRLKVLTVLLYICLSVMLFYLFGSIMPLPPWGLVVGWVMALIAVELAYAFIFPYSFRYIADNDDDKMVILPV
B239        MLHHLKELAAVAGIHMILIYLCRFLLRRSRNVLFTVSNSLRFRLKVLTVLLYICLSVMLFYLFGSIMPLPPWGLVVGWVMALIAVELAYAFIFPYSFRYIADNDDDKMVILPV
B240        MLHHLKELAAVAGIHMILIYLCRFLLRRSRNVLFTVSNSLRFRLKVLTVLLYICLSVMLFYLFGSIMPLPPWGLVVGWVMALIAVELAYAFIFPYSFRYIADNDDDKMVILPV
B241        MLHHLKELAAVAGIHMILIYLCRFLLRRSRNVLFTVSNSLRFRLKVLTVLLYICLSVMLFYLFGSIMPLPPWGLVVGWVMALIAVELAYAFIFPYSFRYIADNDDDKMVILPV
B242        MLHHLKELAAVAGIHMILIYLCRFLLRRSRNVLFTVSNSLRFRLKVLTVLLYICLSVMLFYLFGSIMPLPPWGLVVGWVMALIAVELAYAFIFPYSFRYIADNDDDKMVILPV
B243        MLHHLKELAAVAGIHMILIYLCRFLLRRSRNVLFTVSNSLRFRLKVLTVLLYICLSVMLFYLFGSIMPLPPWGLVVGWVMALIAVELAYAFIFPYSFRYIADNDDDKMVILPV
B244        MLHHLKELAAVAGIHMILIYLCRFLLRRSRNVLFTVSNSLRFRLKVLTVLLYICLSVMLFYLFGSIMPLPPWGLVVGWVMALIAVELAYAFIFPYSFRYIADNDDDKMVILPV
B245        MLHQLKELAAVAGIHMILIYLCRFLLRRSRNVLFTVSNSLRFRLKVLTVLLYICLLVMLFYLFGSIMPLPPWGLVVGWVMALIAVELAYAFIFPYSFRYIADNDDDKMVILPV
B246        MLHHLKELAAVAGIHMILIYLCRFLLRRSRNVLFTVSNSLRFRLKVLTVLLYICLSVMLFYLFGSIMPLPPWGLVVGWVMALIAVELAYAFIFPYSFRYIADNDDDKMVILPV
B247        MLHHLKELAAVAGIHMILIYLCRFLLRRSRNVLFTVSNSLRFRLKVLTVLLYICLSVMLFYLFGSIMPLPPWGLVVGWVMALIAVELAYAFIFPYSFRYIADNDDDKMVILPV
B248        MLHHLKELAAVAGIHMILIYLCRFLLRRSRNVLFTVSNSLRFRLKVLTVLLYICLSVMLFYLFGSIMPLPPWGLVVGWVMALIAVELAYAFIFPYSFRYIADNDDDKMVILPV
B249        MLHHLKELAAVAGIHMILIYLCRFLLRLRRNVLFTVSNSLRFRLKVLTVLLYICLSVMLFYLFGSIMPLPPWGLVVGWVMALIAVELAYAFIFPYSFRYIADNDDDKMVILPV
B250        MLHHLKELAAVAGIHMILIYLCRFLLRRSRNVLFTVSNSLRFRLKVLTVLLYICLSVMLFYLFGSIMPLPPWGLVVGWVMALIAVELAYAFIFPYSFRYIADNDDDKMVILPV
B252        MLHHLKELAAVAGIHMILIYLCRFLLRLSRNVLFTVSNSLRFRLKVLTVLLYICLSVMLFYLFGSIMPLPPWGLVVGWVMALIAVELAYAFIFPYSFRYIADNDDDKMVILPV
B253        MLHHLKELAAVAGIHMILIYLCRFLLRRSRNVLFTVSNSLRFRLKVLTVLLYICLSVMLFYLFGSIMPLPPWGLVVGWVMALIAVELAYAFIFPYSFRYIADNDDDKMVILPV
B254        MLHHLKELAAVAGIHMILIYLCRFLLRLSRNVLFTVSNSLRFRLKVLTVLLYICLSVMLFYLFGSIMPLPPWGLVVGWVMALIAVELAYAFIFPYSFRYIADNDDDKMVILPV
B255        MLHHLKELAAVAGIHMILIYLCRFLLRRSRNVLFTVSNSLRFRLKVLTVLLYICLSVMLFYLFGSIMPLPPWGLVVGWVMALIAVELAYAFIFPYSFRYIADNDDDKMVILPV
```

```
B258    MLHHLKELAAVAGIHMILIYLCRFLLRRSRNVLFTVSNSLRFRLKVLTVLLYICLSVMLFYLFGSIMPLPPWGLVVGWVMALIAVELAYAFIFPYSFRYIADNDDDKMVILPV
B259    MLHHLKELAAVAGIHMILIYLCRFLLRRSRNVLFTVSNSLRFRLKVLTVLLYICLSVMLFYLFGSIMPLPPWGLVVGWVMALIAVELAYAFIFPYSFRYIADNDDDKMVILPV
B260    MLHHLKELAAVAGIHMILIYLCRFLLRRSRNVLFTVSNSLRFRLKVLTVLLYICLSVMLFYLFGSIMPLPPWGLVVGWVMALIAVELAYAFIFPYSFRYIADNDDDKMVILPV
B261    MLHHLKELAAVAGIHMILIYLCRFLLRRSRNVLFTVSNSLRFRLKVLTVLLYICLSVMLFYLFGSIMPLPPWGLVVGWVMALIAVELAYAFIFPYSFRYIADNDDDKMVILPV
B263    MLHHLKELAAVAGIHMILIYLCRFLLRRSRNVLFTVSNSLRFRLKVLTVLLYICLSVMLFYLFGSIMPLPPWGLVVGWVMALIAVELAYAFIFPYSFRYIADNDDDKMVILPV
B264    MLHHLKELAAVAGIHMILIYLCRFLLRRSRNVLFTVSNSLRFRLKVLTVLLYICLSVMLFYLFGSIMPLPPWGLVVGWVMALIAVELAYAFIFPYSFRYIADNDDDKMVILPV
B265    MLHHLKELAAVAGIHMILIYLCRFLLRRSRNVLFTVSNSLRFRLKVLTVLLYICLSVMLFYLFGSIMPLPPWGLVVGWVMALIAVELAYAFIFPYSFRYIADNDDDKMVILPV
B266    MLHHLKELAAVAGIHMILIYLCRFLLRRSRNVLFTVSNSLRFRLKVLTVLLYICLSVMLFYLFGSIMPLPPWGLVVGWVMALIAVELAYAFIFPYSFRYIADNDDDKMVILPV
B267    MLHHLKELAAVAGIHMILIYLCRFLLRRSRNVLFTVSNSLRFRLKVLTVLLYICLSVMLFYLFGSIMPLPPWGLVVGWVMALIAVELAYAFIFPYSFRYIADNDDDKMVILPV
B268    MLHHLKELAAVAGIHMILIYLCRFLLRRSRNVLFTVSNSLRFRLKVLTVLLYICLSVMLFYLFGSIMPLPPWGLVVGWVMALIAVELAYAFIFPYSFRYIADNDDDKMVILPV
B269    MLHHLKELAAVAGIHMILIYLCRFLLRRSRNVLFTVSNSLRFRLKVLTVLLYICLSVMLFYLFGSIMPLPPWGLVVGWVMALIAVELAYAFIFPYSFRYIADNDDDKMVILPV
CX2     MLHHLKELAAVAGIHMILIYLCRFLLRRSRNVLFTVSNSLRFRLKVLTVLLYICLSVMLFYLFGSIMPLPPWGLVVGWVMALIAVELAYAFIFPYSFRYIADNDDDKMVILPV
CX3     MLHHLKELAAVAGIHMILIYLCRFLLRRSRNVLFTVSNSLRFRLKVLTVLLYICLSVMLFYLFGSIMPLPPWGLVVGWVMALIAVELAYAFIFPYSFRYIADNDDDKMVILPV
CX4     MLHHLKELAAVAGIHMILIYLCRFLLRRSRNVLFTVSNSLRFRLKVLTVLLYICLSVMLFYLFGSIMPLPPWGLVVGWVMALIAVELAYAFIFPYSFRYIADNDDDKMVILPV
CX5     MLHHLKELAAVAGIHMILIYLCRFLLRRSRNVLFTVSNSLRFRLKVLTVLLYICLSVMLFYLFGSIMPLPPWGLVVGWVMALIAVELAYAFIFPYSFRYIADNDDDKMVILPV
CX6     MLHHLKELAAVAGIHMILIYLCRFLLRRSRNVLFTVSNSLRFRLKVLTVLLYICLSVMLFYLFGSIMPLPPWGLVVGWVMALIAVELAYAFIFPYSFRYIADNDDDKMVILPV
CX8     MLHHLKELAAVAGIHMILIYLCRFLLRRSRNVLFTVSNSLRFRLKVLTVLLYICLSVMLFYLFGSIMPLPPWGLVVGWVMALIAVELAYAFIFPYSFRYIADNDDDKMVILPV
CX9     MLHHLKELAAVAGIHMILIYLCRFLLRRSRNVLFTVSNSLRFRLKVLTVLLYICLSVMLFYLFGSIMPLPPWGLVVGWVMALIAVELAYAFIFPYSFRYIADNDDDKMVILPV
CX10    MLHHLKELAAVAGIHMILIYLCRFLLRRSRNVLFTVSNSLRFRLKVLTVLLYICLSVMLFYLFGSIMPLPPWGLVVGWVMALIAVELAYAFIFPYSFRYIADNDDDKMVILPV
CX11    MLHHLKELAAVAGIHMILIYLCRFLLRRSRNVLFTVSNSLRFRLKVLTVLLYICLSVMLFYLFGSIMPLPPWGLVVGWVMALIAVELAYAFIFPYSFRYIADNDDDKMVILPV
CX12    MLHHLKELAAVAGIHMILIYLCRFLLRRSRNVLFTVSNSLRFRLKVLTVLLYICLSVMLFYLFGSIMPLPPWGLVVGWVMALIAVELAYAFIFPYSFRYIADNDDDKMVILPV
CX13    MLHHLKELAAVAGIHMILIYLCRFLLRRSRNVLFTVSNSLRFRLKVLTVLLYICLSVMLFYLFGSIMPLPPWGLVVGWVMALIAVELAYAFIFPYSFRYIADNDDDKMVILPV
CX14    MLHHLKELAAVAGIHMILIYLCRFLLRRSRNVLFTVSNSLRFRLKVLTVLLYICLSVMLFYLFGSIMPLPPWGLVVGWVMALIAVELAYAFIFPYSFRYIADNDDDKMVILPV
CX15    MLHHLKELAAVAGIHMILIYLCRFLLRRSRNVLFTVSNSLRFRLKVLTVLLYICLSVMLFYLFGSIMPLPPWGLVVGWVMALIAVELAYAFIFPYSFRYIADNDDDKMVILPV
CX16    MLHHLKELAAVAGIHMILIYLCRFLLRRSRNVLFTVSNSLRFRLKVLTVLLYICLSVMLFYLFGSIMPLPPWGLVVGWVMALIAVELAYAFIFPYSFRYIADNDDDKMVILPV
CX17    MLHHLKELAAVAGIHMILIYLCRFLLRRSRNVLFTVSNSLRFRLKVLTVLLYICLSVMLFYLFGSIMPLPPWGLVVGWVMALIAVELAYAFIFPYSFRYIADNDDDKMVILPV
CX18    MLHHLKELAAVAGIHMILIYLCRFLLRRSRNVLFTVSNSLRFRLKVLTVLLYICLSVMLFYLFGSIMPLPPWGLVVGWVMALIAVELAYAFIFPYSFRYIADNDDDKMVILPV
CX19    MLHHLKELAAVAGIHMILIYLCRFLLRLSRNVLFTVSNSLRFRLKVLTVLLYICLSVMLFYLFGSIMPLPPWGLVVGWVMALIAVELAYAFIFPYSFRYIADNDDDKMVILPV
CX20    MLHHLKELAAVAGIHMILIYLCRFLLRRSRNVLFTVSNSLRFRLKVLTVLLYICLSVMLFYLFGSIMPLPPWGLVVGWVMALIAVELAYAFIFPYSFRYIADNDDDKMVILPV
CX21    MLHHLKELAAVAGIHMILIYLCRFLLRRSRNVLFTVSNSLRFRLKVLTVLLYICLSVMLFYLFGSIMPLPPWGLVVGWVMALIAVELAYAFIFPYSFRYIADNDDDKMVILPV
CX22    MLHHLKELAAVAGIHMILIYLCRFLLRRSRNVLFTVSNSLRFRLKVLTVLLYICLSVMLFYLFGSIMPLPPWGLVVGWVMALIAVELAYAFIFPYSFRYIADNDDDKMVILPV
CX23    MLHHLKELAAVAGIHMILIYLCRFLLRRSRNVLFTVSNSLRFRLKVLTVLLYICLSVMLFYLFGSIMPLPPWGLVVGWVMALIAVELAYAFIFPYSFRYIADNDDDKMVILPV
CX24    MLHHLKELAAVAGIHMILIYLCRFLLRRSRNVLFTVSNSLRFRLKVLTVLLYICLSVMLFYLFGSIMPLPPWGLVVGWVMALIAVELAYAFIFPYSFRYIADNDDDKMVILPV
CX25    MLHHLKELAAVAGIHMILIYLCRFLLRRSRNVLFTVSNSLRFRLKVLTVLLYICLSVMLFYLFGSIMPLPPWGLVVGWVMALIAVELAYAFIFPYSFRYIADNDDDKMVILPV
CX26    MLHHLKELAAVAGIHMILIYLCRFLLRRSRNVLFTVSNSLRFRLKVLTVLLYICLSVMLFYLFGSIMPLPPWGLVVGWVMALIAVELAYAFIFPYSFRYIADNDDDKMVILPV
CX27    MLHHLKELAAVAGIHMILIYLCRFLLRRSRNVLFTVSNSLRFRLKVLTVLLYICLSVMLFYLFGSIMPLPPWGLVVGWVMALIAVELAYAFIFPYSFRYIADNDDDKMVILPV
CX28    MLHHLKELAAVAGIHMILIYLCRFLLRRSRNVLFTVSNSLRFRLKVLTVLLYICLSVMLFYLFGSIMPLPPWGLVVGWVMALIAVELAYAFIFPYSFRYIADNDDDKMVILPV
CX29    MLHHLKELAAVAGIHMILIYLCRFLLRRSRNVLFTVSNSLRFRLKVLTVLLYICLSVMLFYLFGSIMPLPPWGLVVGWVMALIAVELAYAFIFPYSFRYIADNDDDKMVILPV
CX30    MLHHLKELAAVAGIHMILIYLCRFLLRRSRNVLFTVSNSLRFRLKVLTVLLYICLSVMLFYLFGSIMPLPPWGLVVGWVMALIAVELAYAFIFPYSFRYIADNDDDKMVILPV
CX31    MLHHLKELAAVAGIHMILIYLCRFLLRRSRNVLFTVSNSLRFRLKVLTVLLYICLSVMLFYLFGSIMPLPPWGLVVGWVMALIAVELAYAFIFPYSFRYIADNDDDKMVILPV
CX32    MLHHLKELAAVAGIHMILIYLCRFLLRRSRNVLFTVSNSLRFRLKVLTVLLYICLSVMLFYLFGSIMPLPPWGLVVGWVMALIAVELAYAFIFPYSFRYIADNDDDKMVILPV
CX33    MLHHLKELAAVAGIHMILIYLCRFLLRRSRNVLFTVSNSLRFRLKVLTVLLYICLSVMLFYLFGSIMPLPPWGLVVGWVMALIAVELAYAFIFPYSFRYIADNDDDKMVILPV
CX34    MLHHLKELAAVAGIHMILIYLCRFLLRRSRNVLFTVSNSLRFRLKVLTVLLYICLSVMLFYLFGSIMPLPPWGLVVGWVMALIAVELAYAFIFPYSFRYIADNDDDKMVILPV
CX35    MLHHLKELAAVAGIHMILIYLCRFLLRRSRNVLFTVSNSLRFRLKVLTVLLYICLSVMLFYLFGSIMPLPPWGLVVGWVMALIAVELAYAFIFPYSFRYIADNDDDKMVILPV
CX37    MLHHLKELAAVAGIHMILIYLCRFLLRRSRNVLFTVSNSLRFRLKVLTVLLYICLSVMLFYLFGSIMPLPPWGLVVGWVMALIAVELAYAFIFPYSFRYIADNDDDKMVILPV
CX42    MLHHLKELAAVAGIHMILIYLCRFLLRRSRNVLFTVSNSLRFRLKVLTVLLYICLSVMLFYLFGSIMPLPPWGLVVGWVMALIAVELAYAFIFPYSFRYIADNDDDKMVILPV
CX43    MLHHLKELAAVAGIHMILIYLCRFLLRRSRNVLFTVSNSLRFRLKVLTVLLYICLSVMLFYLFGSIMPLPPWGLVVGWVMALIAVELAYAFIFPYSFRYIADNDDDKMVILPV
CX44    MLHHLKELAAVAGIHMILIYLCRFLLRRSRNVLFTVSNSLRFRLKVLTVLLYICLSVMLFYLFGSIMPLPPWGLVVGWVMALIAVELAYAFIFPYSFRYIADNDDDKMVILPV
CX45    MLHHLKELAAVAGIHMILIYLCRFLLRRSRNVLFTVSNSLRFRLKVLTVLLYICLSVMLFYLFGSIMPLPPWGLVVGWVMALIAVELAYAFIFPYSFRYIADNDDDKMVILPV
CX46    MLHHLKELAAVAGIHMILIYLCRFLLRRSRNVLFTVSNSLRFRLKVLTVLLYICLSVMLFYLFGSIMPLPPWGLVVGWVMALIAVELAYAFIFPYSFRYIADNDDDKMVILPV
CX47    MLHHLKELAAVAGIHMILIYLCRFLLRRSRNVLFTVSNSLRFRLKVLTVLLYICLSVMLFYLFGSIMPLPPWGLVVGWVMALIAVELAYAFIFPYSFRYIADNDDDKMVILPV
CX48    MLHHLKELAAVAGIHMILIYLCRFLLRRSRNVLFTVSNSLRFRLKVLTVLLYICLSVMLFYLFGSIMPLPPWGLVVGWVMALIAVELAYAFIFPYSFRYIADNDDDKMVILPV
CX49    MLHHLKELAAVAGIHMILIYLCRFLLRRSRNVLFTVSNSLRFRLKVLTVLLYICLSVMLFYLFGSIMPLPPWGLVVGWVMALIAVELAYAFIFPYSFRYIADNDDDKMVILPV
CX50    MLHHLKELAAVAGIHMILIYLCRFLLRRSRNVLFTVSNSLRFRLKVLTVLLYICLSVMLFYLFGSIMPLPPWGLVVGWVMALIAVELAYAFIFPYSFRYIADNDDDKMVILPV
CX51    MLHHLKELAAVAGIHMILIYLCRFLLRRSRNVLFTVSNSLRFRLKVLTVLLYICLSVMLFYLFGSIMPLPPWGLVVGWVMALIAVELAYAFIFPYSFRYIADNDDDKMVILPV
CX52    MLHHLKELAAVAGIHMILIYLCRFLLRRSRNVLFTVSNSLRFRLKVLTVLLYICLSVMLFYLFGSIMPLPPWGLVVGWVMALIAVELAYAFIFPYSFRYIADNDDDKMVILPV
CX53    MLHHLKELAAVAGIHMILIYLCRFLLRRSRNVLFTVSNSLRFRLKVLTVLLYICLSVMLFYLFGSIMPLPPWGLVVGWVMALIAVELAYAFIFPYSFRYIADNDDDKMVILPV
CX54    MLHHLKELAAVAGIHMILIYLCRFLLRRSRNVLFTVSNSLRFRLKVLTVLLYICLSVMLFYLFGSIMPLPPWGLVVGWVMALIAVELAYAFIFPYSFRYIADNDDDKMVILPV
CX55    MLHHLKELAAVAGIHMILIYLCRFLLRRSRNVLFTVSNSLRFRLKVLTVLLYICLSVMLFYLFGSIMPLPPWGLVVGWVMALIAVELAYAFIFPYSFRYIADNDDDKMVILPV
CX56    MLHQLKELAAVAGIHMILIYLCRFLLRRSRNVLFTVSNSLRFRLKVLTVLLYICLLVMLFYLFGSIMPLPPWGLVVGWVMALIAVELAYAFIFPYSFRYIADNDDDKMVILPV
CX57    MLHHLKELAAVAGIHMILIYLCRFLLRRSRNVLFTVSNSLRFRLKVLTVLLYICLSVMLFYLFGSIMPLPPWGLVVGWVMALIAVELAYAFIFPYSFRYIADNDDDKMVILPV
CX58    MLHHLKELAAVAGIHMILIYLCRFLLRRSRNVLFTVSNSLRFRLKVLTVLLYICLSVMLFYLFGSIMPLPPWGLVVGWVMALIAVELAYAFIFPYSFRYIADNDDDKMVILPV
CX59    MLHHLKELAAVAGIHMILIYLCRFLLRRSRNVLFTVSNSLRFRLKVLTVLLYICLSVMLFYLFGSIMPLPPWGLVVGWVMALIAVELAYAFIFPYSFRYIADNDDDKMVILPV
CX60    MLHHLKELAAVAGIHMILIYLCRFLLRRSRNVLFTVSNSLRFRLKVLTVLLYICLSVMLFYLFGSIMPLPPWGLVVGWVMALIAVELAYAFIFPYSFRYIADNDDDKMVILPV
CX61    MLHHLKELAAVAGIHMILIYLCRFLLRRSRNVLFTVSNSLRFRLKVLTVLLYICLSVMLFYLFGSIMPLPPWGLVVGWVMALIAVELAYAFIFPYSFRYIADNDDDKMVILPV
CX63    MLHHLKELAAVAGIHMILIYLCRFLLRRSRNVLFTVSNSLRFRLKVLTVLLYICLSVMLFYLFGSIMPLPPWGLVVGWVMALIAVELAYAFIFPYSFRYIADNDDDKMVILPV
CX64    MLHHLKELAAVAGIHMILIYLCRFLLRRSRNVLFTVSNSLRFRLKVLTVLLYICLSVMLFYLFGSIMPLPPWGLVVGWVMALIAVELAYAFIFPYSFRYIADNDDDKMVILPV
CX65    MLHHLKELAAVAGIHMILIYLCRFLLRRSRNVLFTVSNSLRFRLKVLTVLLYICLSVMLFYLFGSIMPLPPWGLVVGWVMALIAVELAYAFIFPYSFRYIADNDDDKMVILPV
CX66    MLHHLKELAAVAGIHMILIYLCRFLLRRSRNVLFTVSNSLRFRLKVLTVLLYICLSVMLFYLFGSIMPLPPWGLVVGWVMALIAVELAYAFIFPYSFRYIADNDDDKMVILPV
CX67    MLHHLKELAAVAGIHMILIYLCRFLLRRSRNVLFTVSNSLRFRLKVLTVLLYICLSVMLFYLFGSIMPLPPWGLVVGWVMALIAVELAYAFIFPYSFRYIADNDDDKMVILPV
CX68    MLHHLKELAAVAGIHMILIYLCRFLLRRSRNVLFTVSNSLRFRLKVLTVLLYICLSVMLFYLFGSIMPLPPWGLVVGWVMALIAVELAYAFIFPYSFRYIADNDDDKMVILPV
CX69    MLHHLKELAAVAGIHMILIYLCRFLLRRSRNVLFTVSNSLRFRLKVLTVLLYICLSVMLFYLFGSIMPLPPWGLVVGWVMALIAVELAYAFIFPYSFRYIADNDDDKMVILPV
CX70    MLHHLKELAAVAGIHMILIYLCRFLLRRSRNVLFTVSNSLRFRLKVLTVLLYICLSVMLFYLFGSIMPLPPWGLVVGWVMALIAVELAYAFIFPYSFRYIADNDDDKMVILPV
CX71    MLHHLKELAAVAGIHMILIYLCRFLLRRSRNVLFTVSNSLRFRLKVLTVLLYICLSVMLFYLFGSIMPLPPWGLVVGWVMALIAVELAYAFIFPYSFRYIADNDDDKMVILPV
CX72    MLHHLKELAAVAGIHMILIYLCRFLLRRSRNVLFTVSNSLRFRLKVLTVLLYICLSVMLFYLFGSIMPLPPWGLVVGWVMALIAVELAYAFIFPYSFRYIADNDDDKMVILPV
CX73    MLHHLKELAAVAGIHMILIYLCRFLLRRSRNVLFTVSNSLRFRLKVLTVLLYICLSVMLFYLFGSIMPLPPWGLVVGWVMALIAVELAYAFIFPYSFRYIADNDDDKMVILPV
CX74    MLHHLKELAAVAGIHMILIYLCRFLLRRSRNVLFTVSNSLRFRLKVLTVLLYICLSVMLFYLFGSIMPLPPWGLVVGWVMALIAVELAYAFIFPYSFRYIADNDDDKMVILPV
CX75    MLHHLKELAAVAGIHMILIYLCRFLLRRSRNVLFTVSNSLRFRLKVLTVLLYICLSVMLFYLFGSIMPLPPWGLVVGWVMALIAVELAYAFIFPYSFRYIADNDDDKMVILPV
CX76    MLHHLKELAAVAGIHMILIYLCRFLLRRSRNVLFTVSNSLRFRLKVLTVLLYICLSVMLFYLFGSIMPLPPWGLVVGWVMALIAVELAYAFIFPYSFRYIADNDDDKMVILPV
CX77    MLHHLKELAAVAGIHMILIYLCRFLLRRSRNVLFTVSNSLRFRLKVLTVLLYICLSVMLFYLFGSIMPLPPWGLVVGWVMALIAVELAYAFIFPYSFRYIADNDDDKMVILPV
CX78    MLHHLKELAAVAGIHMILIYLCRFLLRRSRNVLFTVSNSLRFRLKVLTVLLYICLSVMLFYLFGSIMPLPPWGLVVGWVMALIAVELAYAFIFPYSFRYIADNDDDKMVILPV
CX79    MLHHLKELAAVAGIHMILIYLCRFLLRRSRNVLFTVSNSLRFRLKVLTVLLYICLSVMLFYLFGSIMPLPPWGLVVGWVMALIAVELAYAFIFPYSFRYIADNDDDKMVILPV
CX80    MLHHLKELAAVAGIHMILIYLCRFLLRLRRNVLFTVSNSLRFRLKVLTVLLYICLSVMLFYLFGSIMPLPPWGLVVGWVMALIAVELAYAFIFPYSFRYIADNDDDKMVILPV
CX82    MLHHLKELAAVAGIHMILIYLCRFLLRRSRNVLFTVSNSLRFRLKVLTVLLYICLSVMLFYLFGSIMPLPPWGLVVGWVMALIAVELAYAFIFPYSFRYIADNDDDKMVILPV
CX83    MLHHLKELAAVAGIHMILIYLCRFLLRRSRNVLFTVSNSLRFRLKVLTVLLYICLSVMLFYLFGSIMPLPPWGLVVGWVMALIAVELAYAFIFPYSFRYIADNDDDKMVILPV
CX84    MLHHLKELAAVAGIHMILIYLCRFLLRRSRNVLFTVSNSLRFRLKVLTVLLYICLSVMLFYLFGSIMPLPPWGLVVGWVMALIAVELAYAFIFPYSFRYIADNDDDKMVILPV
CX85    MLHHLKELAAVAGIHMILIYLCRFLLRRSRNVLFTVSNSLRFRLKVLTVLLYICLSVMLFYLFGSIMPLPPWGLVVGWVMALIAVELAYAFIFPYSFRYIADNDDDKMVILPV
CX86    MLHHLKELAAVAGIHMILIYLCRFLLRRSRNVLFTVSNSLRFRLKVLTVLLYICLSVMLFYLFGSIMPLPPWGLVVGWVMALIAVELAYAFIFPYSFRYIADNDDDKMVILPV
CX87    MLHHLKELAAVAGIHMILIYLCRFLLRRSRNVLFTVSNSLRFRLKVLTVLLYICLSVMLFYLFGSIMPLPPWGLVVGWVMALIAVELAYAFIFPYSFRYIADNDDDKMVILPV
CX88    MLHHLKELAAVAGIHMILIYLCRFLLRRSRNVLFTVSNSLRFRLKVLTVLLYICLSVMLFYLFGSIMPLPPWGLVVGWVMALIAVELAYAFIFPYSFRYIADNDDDKMVILPV
CX89    MLHHLKELAAVAGIHMILIYLCRFLLRRSRNVLFTVSNSLRFRLKVLTVLLYICLSVMLFYLFGSIMPLPPWGLVVGWVMALIAVELAYAFIFPYSFRYIADNDDDKMVILPV
CX90    MLHHLKELAAVAGIHMILIYLCRFLLRRSRNVLFTVSNSLRFRLKVLTVLLYICLSVMLFYLFGSIMPLPPWGLVVGWVMALIAVELAYAFIFPYSFRYIADNDDDKMVILPV
CX91    MLHHLKELAAVAGIHMILIYLCRFLLRRSRNVLFTVSNSLRFRLKVLTVLLYICLSVMLFYLFGSIMPLPPWGLVVGWVMALIAVELAYAFIFPYSFRYIADNDDDKMVILPV
CX92    MLHHLKELAAVAGIHMILIYLCRFLLRRSRNVLFTVSNSLRFRLKVLTVLLYICLSVMLFYLFGSIMPLPPWGLVVGWVMALIAVELAYAFIFPYSFRYIADNDDDKMVILPV
CX93    MLHHLKELAAVAGIHMILIYLCRFLLRRSRNVLFTVSNSLRFRLKVLTVLLYICLSVMLFYLFGSIMPLPPWGLVVGWVMALIAVELAYAFIFPYSFRYIADNDDDKMVILPV
CX94    MLHHLKELAAVAGIHMILIYLCRFLLRRSRNVLFTVSNSLRFRLKVLTVLLYICLSVMLFYLFGSIMPLPPWGLVVGWVMALIAVELAYAFIFPYSFRYIADNDDDKMVILPV
CX96    MLHHLKELAAVAGIHMILIYLCRFLLRRSRNVLFTVSNSLRFRLKVLTVLLYICLSVMLFYLFGSIMPLPPWGLVVGWVMALIAVELAYAFIFPYSFRYIADNDDDKMVILPV
CX97    MLHHLKELAAVAGIHMILIYLCRFLLRRSRNVLFTVSNSLRFRLKVLTVLLYICLSVMLFYLFGSIMPLPPWGLVVGWVMALIAVELAYAFIFPYSFRYIADNDDDKMVILPV
CX98    MLHHLKELAAVAGIHMILIYLCRFLLRRSRNVLFTVSNSLRFRLKVLTVLLYICLSVMLFYLFGSIMPLPPWGLVVGWVMALIAVELAYAFIFPYSFRYIADNDDDKMVILPV
CX99    MLHHLKELAAVAGIHMILIYLCRFLLRRSRNVLFTVSNSLRFRLKVLTVLLYICLSVMLFYLFGSIMPLPPWGLVVGWVMALIAVELAYAFIFPYSFRYIADNDDDKMVILPV
CX100   MLHHLKELAAVAGIHMILIYLCRFLLRRSRNVLFTVSNSLRFRLKVLTVLLYICLSVMLFYLFGSIMPLPPWGLVVGWVMALIAVELAYAFIFPYSFRYIADNDDDKMVILPV
CX101   MLHHLKELAAVAGIHMILIYLCRFLLRRSRNVLFTVSNSLRFRLKVLTVLLYICLSVMLFYLFGSIMPLPPWGLVVGWVMALIAVELAYAFIFPYSFRYIADNDDDKMVILPV
CX102   MLHHLKELAAVAGIHMILIYLCRFLLRRSRNVLFTVSNSLRFRLKVLTVLLYICLSVMLFYLFGSIMPLPPWGLVVGWVMALIAVELAYAFIFPYSFRYIADNDDDKMVILPV
CX103   MLHHLKELAAVAGIHMILIYLCRFLLRRSRNVLFTVSNSLRFRLKVLTVLLYICLSVMLFYLFGSIMPLPPWGLVVGWVMALIAVELAYAFIFPYSFRYIADNDDDKMVILPV
CX104   MLHHLKELAAVAGIHMILIYLCRFLLRRSRNVLFTVSNSLRFRLKVLTVLLYICLSVMLFYLFGSIMPLPPWGLVVGWVMALIAVELAYAFIFPYSFRYIADNDDDKMVILPV
CX106   MLHHLKELAAVAGIHMILIYLCRFLLRRSRNVLFTVSNSLRFRLKVLTVLLYICLSVMLFYLFGSIMPLPPWGLVVGWVMALIAVELAYAFIFPYSFRYIADNDDDKMVILPV
CX107   MLHHLKELAAVAGIHMILIYLCRFLLRRSRNVLFTVSNSLRFRLKVLTVLLYICLSVMLFYLFGSIMPLPPWGLVVGWVMALIAVELAYAFIFPYSFRYIADNDDDKMVILPV
CX108   MLHHLKELAAVAGIHMILIYLCRFLLRRSRNVLFTVSNSLRFRLKVLTVLLYISLSAMLFYLFGSIMPLPPWGLVVGWVMALIAVELAYAFIFPYSFRYIADNDDDKMVILPV
CX109   MLHHLKELAAVAGIHMILIYLCRFLLRRSRNVLFTVSNSLRFRLKVLTVLLYICLSVMLFYLFGSIMPLPPWGLVVGWVMALIAVELAYAFIFPYSFRYIADNDDDKMVILPV
CX110   MLHHLKELAAVAGIHMILIYLCRFLLRRSRNVLFTVSNSLRFRLKVLTVLLYICLSVMLFYLFGSIMPLPPWGLVVGWVMALIAVELAYAFIFPYSFRYIADNDDDKMVILPV
CX111   MLHHLKELAAVAGIHMILIYLCRFLLRRSRNVLFTVSNSLRFRLKVLTVLLYICLSVMLFYLFGSIMPLPPWGLVVGWVMALIAVELAYAFIFPYSFRYIADNDDDKMVILPV
CX112   MLHHLKELAAVAGIHMILIYLCRFLLRRSRNVLFTVSNSLRFRLKVLTVLLYICLSVMLFYLFGSIMPLPPWGLVVGWVMALIAVELAYAFIFPYSFRYIADNDDDKMVILPV
CX113   MLHHLKELAAVAGIHMILIYLCRFLLRRSRNVLFTVSNSLRFRLKVLTVLLYICLSVMLFYLFGSIMPLPPWGLVVGWVMALIAVELAYAFIFPYSFRYIADNDDDKMVILPV
CX114   MLHHLKELAAVAGIHMILIYLCRFLLRRSRNVLFTVSNSLRFRLKVLTVLLYICLSVMLFYLFGSIMPLPPWGLVVGWVMALIAVELAYAFIFPYSFRYIADNDDDKMVILPV
CX115   MLHHLKELAAVAGIHMILIYLCRFLLRRSRNVLFTVSNSLRFRLKVLTVLLYICLSVMLFYLFGSIMPLPPWGLVVGWVMALIAVELAYAFIFPYSFRYIADNDDDKMVILPV
CX116   MLHHLKELAAVAGIHMILIYLCRFLLRRSRNVLFTVSNSLRFRLKVLTVLLYICLSVMLFYLFGSIMPLPPWGLVVGWVMALIAVELAYAFIFPYSFRYIADNDDDKMVILPV
CX117   MLHHLKELAAVAGIHMILIYLCRFLLRRSRNVLFTVSNSLRFRLKVLTVLLYICLSVMLFYLFGSIMPLPPWGLVVGWVMALIAVELAYAFIFPYSFRYIADNDDDKMVILPV
CX118   MLHHLKELAAVAGIHMILIYLCRFLLRRSRNVLFTVSNSLRFRLKVLTVLLYICLSVMLFYLFGSIMPLPPWGLVVGWVMALIAVELAYAFIFPYSFRYIADNDDDKMVILPV
CX119   MLHHLKELAAVAGIHMILIYLCRFLLRRSRNVLFTVSNSLRFRLKVLTVLLYICLSVMLFYLFGSIMPLPPWGLVVGWVMALIAVELAYAFIFPYSFRYIADNDDDKMVILPV
CX120   MLHHLKELAAVAGIHMILIYLCRFLLRRSRNVLFTVSNSLRFRLKVLTVLLYICLSVMLFYLFGSIMPLPPWGLVVGWVMALIAVELAYAFIFPYSFRYIADNDDDKMVILPV
CX121   MLHHLKELAAVAGIHMILIYLCRFLLRRSRNVLFTVSNSLRFRLKVLTVLLYICLSVMLFYLFGSIMPLPPWGLVVGWVMALIAVELAYAFIFPYSFRYIADNDDDKMVILPV
CX122   MLHHLKELAAVAGIHMILIYLCRFLLRRSRNVLFTVSNSLRFRLKVLTVLLYICLSVMLFYLFGSIMPLPPWGLVVGWVMALIAVELAYAFIFPYSFRYIADNDDDKMVILPV
CX123   MLHHLKELAAVAGIHMILIYLCRFLLRRSRNVLFTVSNSLRFRLKVLTVLLYICLSVMLFYLFGSIMPLPPWGLVVGWVMALIAVELAYAFIFPYSFRYIADNDDDKMVILPV
```

| | |
|---|---|
| CX124 | MLHHLKELAAVAGIHMILIYLCRFLLRRSRNVLFTVSNSLRFRLKVLTVLLYICLSVMLFYLFGSIMPLPPWGLVVGWVMALIAVELAYAFIFPYSFRYIADNDDDKMVILPV |
| CX125 | MLHHLKELAAVAGIHMILIYLCRFLLRRSRNVLFTVSNSLRFRLKVLTVLLYICLSVMLFYLFGSIMPLPPWGLVVGWVMALIAVELAYAFIFPYSFRYIADNDDDKMVILPV |
| CX126 | MLHHLKELAAVAGIHMILIYLCRFLLRRSRNVLFTVSNSLRFRLKVLTVLLYICLSVMLFYLFGSIMPLPPWGLVVGWVMALIAVELAYAFIFPYSFRYIADNDDDKMVILPV |
| CX128 | MLHHLKELAAVAGIHMILIYLCRFLLRRSRNVLFTVSNSLRFRLKVLTVLLYICLSVMLFYLFGSIMPLPPWGLVVGWVMALIAVELAYAFIFPYSFRYIADNDDDKMVILPV |
| CX129 | MLHHLKELAAVAGIHMILIYLCRFLLRRSRNVLFTVSNSLRFRLKVLTVLLYICLSVMLFYLFGSIMPLPPWGLVVGWVMALIAVELAYAFIFPYSFRYIADNDDDKMVILPV |
| CX130 | MLHHLKELAAVAGIHMILIYLCRFLLRRSRNVLFTVSNSLRFRLKVLTVLLYICLSVMLFYLFGSIMPLPPWGLVVGWVMALIAVELAYAFIFPYSFRYIADNDDDKMVILPV |
| CX131 | MLHHLKELAAVAGIHMILIYLCRFLLRRSRNVLFTVSNSLRFRLKVLTVLLYICLSVMLFYLFGSIMPLPPWGLVVGWVMALIAVELAYAFIFPYSFRYIADNDDDKMVILPV |
| CX132 | MLHHLKELAAVAGIHMILIYLCRFLLRRSRNVLFTVSNSLRFRLKVLTVLLYICLSVMLFYLFGSIMPLPPWGLVVGWVMALIAVELAYAFIFPYSFRYIADNDDDKMVILPV |
| CX133 | MLHHLKELAAVAGIHMILIYLCRFLLRRSRNVLFTVSNSLRFRLKVLTVLLYICLSVMLFYLFGSIMPLPPWGLVVGWVMALIAVELAYAFIFPYSFRYIADNDDDKMVILPV |
| CX134 | MLHHLKELAAVAGIHMILIYLCRFLLRRSRNVLFTVSNSLRFRLKVLTVLLYICLSVMLFYLFGSIMPLPPWGLVVGWVMALIAVELAYAFIFPYSFRYIADNDDDKMVILPV |
| CX138 | MLHQLKELAAVAGIHMILIYLCRFLLRRSRNVLFTVSNSLRFRLKVLTVLLYICLLVMLFYLFGSIMPLPPWGLVVGWVMALIAVELAYAFIFPYSFRYIADNDDDKMVILPV |
| CX139 | MLHHLKELAAVAGIHMILIYLCRFLLRRSRNVLFTVSNSLRFRLKVLTVLLYICLSVMLFYLFGSIMPLPPWGLVVGWVMALIAVELAYAFIFPYSFRYIADNDDDKMVILPV |
| CX140 | MLHHLKELAAVAGIHMILIYLCRFLLRRSRNVLFTVSNSLRFRLKVLTVLLYICLSVMLFYLFGSIMPLPPWGLVVGWVMALIAVELAYAFIFPYSFRYIADNDDDKMVILPV |
| CX141 | MLHHLKELAAVAGIHMILIYLCRFLLRRSRNVLFTVSNSLRFRLKVLTVLLYICLSVMLFYLFGSIMPLPPWGLVVGWVMALIAVELAYAFIFPYSFRYIADNDDDKMVILPV |
| CX142 | MLHHLKELAAVAGIHMILIYLCRFLLRRSRNVLFTVSNSLRFRLKVLTVLLYICLSVMLFYLFGSIMPLPPWGLVVGWVMALIAVELAYAFIFPYSFRYIADNDDDKMVILPV |
| CX143 | MLHHLKELAAVAGIHMILIYLCRFLLRRSRNVLFTVSNSLRFRLKVLTVLLYICLSVMLFYLFGSIMPLPPWGLVVGWVMALIAVELAYAFIFPYSFRYIADNDDDKMVILPV |
| CX144 | MLHHLKELAAVAGIHMILIYLCRFLLRRSRNVLFTVSNSLRFRLKVLTVLLYICLSVMLFYLFGSIMPLPPWGLVVGWVMALIAVELAYAFIFPYSFRYIADNDDDKMVILPV |
| CX145 | MLHHLKELAAVAGIHMILIYLCRFLLRRSRNVLFTVSNSLRFRLKVLTVLLYICLSVMLFYLFGSIMPLPPWGLVVGWVMALIAVELAYAFIFPYSFRYIADNDDDKMVILPV |
| CX146 | MLHHLKELAAVAGIHMILIYLCRFLLRRSRNVLFTVSNSLRFRLKVLTVLLYICLSVMLFYLFGSIMPLPPWGLVVGWVMALIAVELAYAFIFPYSFRYIADNDDDKMVILPV |
| CX147 | MLHHLKELAAVAGIHMILIYLCRFLLRRSRNVLFTVSNSLRFRLKVLTVLLYICLSVMLFYLFGSIMPLPPWGLVVGWVMALIAVELAYAFIFPYSFRYIADNDDDKMVILPV |
| CX148 | MLHHLKELAAVAGIHMILIYLCRFLLRRSRNVLFTVSNSLRFRLKVLTVLLYICLSVMLFYLFGSIMPLPPWGLVVGWVMALIAVELAYAFIFPYSFRYIADNDDDKMVILPV |
| CX149 | MLHHLKELAAVAGIHMILIYLCRFLLRRSRNVLFTVSNSLRFRLKVLTVLLYICLSVMLFYLFGSIMPLPPWGLVVGWVMALIAVELAYAFIFPYSFRYIADNDDDKMVILPV |
| CX150 | MLHHLKELAAVAGIHMILIYLCRFLLRRSRNVLFTVSNSLRFRLKVLTVLLYICLSVMLFYLFGSIMPLPPWGLVVGWVMALIAVELAYAFIFPYSFRYIADNDDDKMVILPV |
| CX151 | MLHHLKELAAVAGIHMILIYLCRFLLRRSRNVLFTVSNSLRFRLKVLTVLLYICLSVMLFYLFGSIMPLPPWGLVVGWVMALIAVELAYAFIFPYSFRYIADNDDDKMVILPV |
| CX152 | MLHHLKELAAVAGIHMILIYLCRFLLRRSRNVLFTVSNSLRFRLKVLTVLLYICLSVMLFYLFGSIMPLPPWGLVVGWVMALIAVELAYAFIFPYSFRYIADNDDDKMVILPV |
| CX153 | MLHHLKELAAVAGIHMILIYLCRFLLRRSRNVLFTVSNSLRFRLKVLTVLLYICLSVMLFYLFGSIMPLPPWGLVVGWVMALIAVELAYAFIFPYSFRYIADNDDDKMVILPV |
| CX154 | MLHHLKELAAVAGIHMILIYLCRFLLRRSRNVLFTVSNSLRFRLKVLTVLLYICLSVMLFYLFGSIMPLPPWGLVVGWVMALIAVELAYAFIFPYSFRYIADNDDDKMVILPV |
| CX155 | MLHHLKELAAVAGIHMILIYLCRFLLRRSRNVLFTVSNSLRFRLKVLTVLLYICLSVMLFYLFGSIMPLPPWGLVVGWVMALIAVELAYAFIFPYSFRYIADNDDDKMVILPV |
| CX156 | MLHHLKELAAVAGIHMILIYLCRFLLRRSRNVLFTVSNSLRFRLKVLTVLLYICLSVMLFYLFGSIMPLPPWGLVVGWVMALIAVELAYAFIFPYSFRYIADNDDDKMVILPV |
| CX157 | MLHHLKELAAVAGIHMILIYLCRFLLRRSRNVLFTVSNSLRFRLKVLTVLLYICLSVMLFYLFGSIMPLPPWGLVVGWVMALIAVELAYAFIFPYSFRYIADNDDDKMVILPV |
| CX158 | MLHHLKELAAVAGIHMILIYLCRFLLRRSRNVLFTVSNSLRFRLKVLTVLLYICLSVMLFYLFGSIMPLPPWGLVVGWVMALIAVELAYAFIFPYSFRYIADNDDDKMVILPV |
| CX160 | MLHHLKELAAVAGIHMILIYLCRFLLRRSRNVLFTVSNSLRFRLKVLTVLLYICLSVMLFYLFGSIMPLPPWGLVVGWVMALIAVELAYAFIFPYSFRYIADNDDDKMVILPV |
| CX161 | MLHHLKELAAVAGIHMILIYLCRFLLRRSRNVLFTVSNSLRFRLKVLTVLLYICLSVMLFYLFGSIMPLPPWGLVVGWVMALIAVELAYAFIFPYSFRYIADNDDDKMVILPV |
| CX162 | MLHHLKELAAVAGIHMILIYLCRFLLRRSRNVLFTVSNSLRFRLKVLTVLLYICLSVMLFYLFGSIMPLPPWGLVVGWVMALIAVELAYAFIFPYSFRYIADNDDDKMVILPV |
| CX165 | MLHHLKELAAVAGIHMILIYLCRFLLRRSRNVLFTVSNSLRFRLKVLTVLLYICLSVMLFYLFGSIMPLPPWGLVVGWVMALIAVELAYAFIFPYSFRYIADNDDDKMVILPV |
| CX182 | MLHHLKELAAVAGIHMILIYLCRFLLRRSRNVLFTVSNSLRFRLKVLTVLLYICLSVMLFYLFGSIMPLPPWGLVVGWVMALIAVELAYAFIFPYSFRYIADNDDDKMVILPV |
| CX205 | MLHHLKELAAVAGIHMILIYLCRFLLRRSRNVLFTVSNSLRFRLKVLTVLLYICLSVMLFYLFGSIMPLPPWGLVVGWVMALIAVELAYAFIFPYSFRYIADNDDDKMVILPV |
| CX206 | MLHHLKELAAVAGIHMILIYLCRFLLRRSRNVLFTVSNSLRFRLKVLTVLLYICLSVMLFYLFGSIMPLPPWGLVVGWVMALIAVELAYAFIFPYSFRYIADNDDDKMVILPV |
| CX207 | MLHHLKELAAVAGIHMILIYLCRFLLRRSRNVLFTVSNSLRFRLKVLTVLLYICLSVMLFYLFGSIMPLPPWGLVVGWVMALIAVELAYAFIFPYSFRYIADNDDDKMVILPV |
| CX210 | MLHHLKELAAVAGIHMILIYLCRFLLRRSRNVLFTVSNSLRFRLKVLTVLLYICLSVMLFYLFGSIMPLPPWGLVVGWVMALIAVELAYAFIFPYSFRYIADNDDDKMVILPV |
| CX211 | MLHHLKELAAVAGIHMILIYLCRFLLRRSRNVLFTVSNSLRFRLKVLTVLLYICLSVMLFYLFGSIMPLPPWGLVVGWVMALIAVELAYAFIFPYSFRYIADNDDDKMVILPV |
| CX212 | MLHHLKELAAVAGIHMILIYLCRFLLRRSRNVLFTVSNSLRFRLKVLTVLLYICLSVMLFYLFGSIMPLPPWGLVVGWVMALIAVELAYAFIFPYSFRYIADNDDDKMVILPV |
| CX213 | MLHHLKELAAVAGIHMILIYLCRFLLRRSRNVLFTVSNSLRFRLKVLTVLLYICLSVMLFYLFGSIMPLPPWGLVVGWVMALIAVELAYAFIFPYSFRYIADNDDDKMVILPV |
| CX214 | MLHHLKELAAVAGIHMILIYLCRFLLRRSRNVLFTVSNSLRFRLKVLTVLLYICLSVMLFYLFGSIMPLPPWGLVVGWVMALIAVELAYAFIFPYSFRYIADNDDDKMVILPV |
| CX218 | MLHHLKELAAVAGIHMILIYLCRFLLRRSRNVLFTVSNSLRFRLKVLTVLLYICLSVMLFYLFGSIMPLPPWGLVVGWVMALIAVELAYAFIFPYSFRYIADNDDDKMVILPV |
| CX219 | MLHHLKELAAVAGIHMILIYLCRFLLRRSRNVLFTVSNSLRFRLKVLTVLLYICLSVMLFYLFGSIMPLPPWGLVVGWVMALIAVELAYAFIFPYSFRYIADNDDDKMVILPV |
| CX220 | MLHHLKELAAVAGIHMILIYLCRFLLRRSRNVLFTVSNSLRFRLKVLTVLLYICLSVMLFYLFGSIMPLPPWGLVVGWVMALIAVELAYAFIFPYSFRYIADNDDDKMVILPV |
| CX221 | MLHHLKELAAVAGIHMILIYLCRFLLRRSRNVLFTVSNSLRFRLKVLTVLLYICLSVMLFYLFGSIMPLPPWGLVVGWVMALIAVELAYAFIFPYSFRYIADNDDDKMVILPV |
| CX225 | MLHHLKELAAVAGIHMILIYLCRFLLRRSRNVLFTVSNSLRFRLKVLTVLLYICLSVMLFYLFGSIMPLPPWGLVVGWVMALIAVELAYAFIFPYSFRYIADNDDDKMVILPV |
| CX226 | MLHHLKELAAVAGIHMILIYLCRFLLRRSRNVLFTVSNSLRFRLKVLTVLLYICLSVMLFYLFGSIMPLPPWGLVVGWVMALIAVELAYAFIFPYSFRYIADNDDDKMVILPV |
| CX227 | MLHHLKELAAVAGIHMILIYLCRFLLRRSRNVLFTVSNSLRFRLKVLTVLLYICLSVMLFYLFGSIMPLPPWGLVVGWVMALIAVELAYAFIFPYSFRYIADNDDDKMVILPV |
| CX228 | MLHHLKELAAVAGIHMILIYLCRFLLRRSRNVLFTVSNSLRFRLKVLTVLLYICLSVMLFYLFGSIMPLPPWGLVVGWVMALIAVELAYAFIFPYSFRYIADNDDDKMVILPV |
| CX230 | MLHHLKELAAVAGIHMILIYLCRFLLRRSRNVLFTVSNSLRFRLKVLTVLLYICLSVMLFYLFGSIMPLPPWGLVVGWVMALIAVELAYAFIFPYSFRYIADNDDDKMVILPV |
| CX231 | MLHHLKELAAVAGIHMILIYLCRFLLRRSRNVLFTVSNSLRFRLKVLTVLLYICLSVMLFYLFGSIMPLPPWGLVVGWVMALIAVELAYAFIFPYSFRYIADNDDDKMVILPV |
| CX232 | MLHHLKELAAVAGIHMILIYLCRFLLRRSRNVLFTVSNSLRFRLKVLTVLLYICLSVMLFYLFGSIMPLPPWGLVVGWVMALIAVELAYAFIFPYSFRYIADNDDDKMVILPV |
| CX233 | MLHHLKELAAVAGIHMILIYLCRFLLRRSRNVLFTVSNSLRFRLKVLTVLLYICLSVMLFYLFGSIMPLPPWGLVVGWVMALIAVELAYAFIFPYSFRYIADNDDDKMVILPV |
| CX234 | MLHHLKELAAVAGIHMILIYLCRFLLRRSRNVLFTVSNSLRFRLKVLTVLLYICLSVMLFYLFGSIMPLPPWGLVVGWVMALIAVELAYAFIFPYSFRYIADNDDDKMVILPV |
| CX235 | MLHHLKELAAVAGIHMILIYLCRFLLRRSRNVLFTVSNSLRFRLKVLTVLLYICLSVMLFYLFGSIMPLPPWGLVVGWVMALIAVELAYAFIFPYSFRYIADNDDDKMVILPV |
| CX236 | MLHHLKELAAVAGIHMILIYLCRFLLRRSRNVLFTVSNSLRFRLKVLTVLLYICLSVMLFYLFGSIMPLPPWGLVVGWVMALIAVELAYAFIFPYSFRYIADNDDDKMVILPV |
| CX237 | MLHHLKELAAVAGIHMILIYLCRFLLRRSRNVLFTVSNSLRFRLKVLTVLLYICLSVMLFYLFGSIMPLPPWGLVVGWVMALIAVELAYAFIFPYSFRYIADNDDDKMVILPV |
| CX238 | MLHHLKELAAVAGIHMILIYLCRFLLRRSRNVLFTVSNSLRFRLKVLTVLLYICLSVMLFYLFGSIMPLPPWGLVVGWVMALIAVELAYAFIFPYSFRYIADNDDDKMVILPV |
| CX240 | MLHHLKELAAVAGIHMILIYLCRFLLRRSRNVLFTVSNSLRFRLKVLTVLLYICLSVMLFYLFGSIMPLPPWGLVVGWVMALIAVELAYAFIFPYSFRYIADNDDDKMVILPV |
| CX241 | MLHHLKELAAVAGIHMILIYLCRFLLRRSRNVLFTVSNSLRFRLKVLTVLLYICLSVMLFYLFGSIMPLPPWGLVVGWVMALIAVELAYAFIFPYSFRYIADNDDDKMVILPV |
| CX242 | MLHHLKELAAVAGIHMILIYLCRFLLRRSRNVLFTVSNSLRFRLKVLTVLLYICLSVMLFYLFGSIMPLPPWGLVVGWVMALIAVELAYAFIFPYSFRYIADNDDDKMVILPV |
| CX243 | MLHHLKELAAVAGIHMILIYLCRFLLRRSRNVLFTVSNSLRFRLKVLTVLLYICLSVMLFYLFGSIMPLPPWGLVVGWVMALIAVELAYAFIFPYSFRYIADNDDDKMVILPV |
| CX247 | MLHHLKELAAVAGIHMILIYLCRFLLRLSRNVLFTVSNSLRFRLKVLTVLLYICLSVMLFYLFGSIMPLPPWGLVVGWVMALIAVELAYAFIFPYSFRYIADNDDDKMVILPV |
| CX248 | MLHQLKELAAVAGIHMILIYLCRFLLRRSRNVLFTVSNSLRFRLKVLTVLLYICLLVMLFYLFGSIMPLPPWGLVVGWVMALIAVELAYAFIFPYSFRYIADNDDDKMVILPV |
| CX249 | MLHHLKELAAVAGIHMILIYLCRFLLRRSRNVLFTVSNSLRFRLKVLTVLLYICLSVMLFYLFGSIMPLPPWGLVVGWVMALIAVELAYAFIFPYSFRYIADNDDDKMVILPV |
| CX250 | MLHHLKELAAVAGIHMILIYLCRFLLRRSRNVLFTVSNSLRFRLKVLTVLLYICLSVMLFYLFGSIMPLPPWGLVVGWVMALIAVELAYAFIFPYSFRYIADNDDDKMVILPV |
| CX251 | MLHHLKELAAVAGIHMILIYLCRFLLRRSRNVLFTVSNSLRFRLKVLTVLLYISLSVMLLYLFGSIMPLPPWGLVVGWVMALIAVELAYAFIFPYSFRYIADNDDDKMVILPV |
| CX262 | MLHHLKELAAVAGIHMILIYLCRFLLRRSRNVLFTVSNSLRFRLKVLTVLLYICLSVMLFYLFGSIMPLPPWGLVVGWVMALIAVELAYAFIFPYSFRYIADNDDDKMVILPV |
| CX263 | MLHHLKELAAVAGIHMILIYLCRFLLRRSRNVLFTVSNSLRFRLKVLTVLLYICLSVMLFYLFGSIMPLPPWGLVVGWVMALIAVELAYAFIFPYSFRYIADNDDDKMVILPV |
| CX265 | MLHHLKELAAVAGIHMILIYLCRFLLRRSRNVLFTVSNSLRFRLKVLTVLLYICLSVMLFYLFGSIMPLPPWGLVVGWVMALIAVELAYAFIFPYSFRYIADNDDDKMVILPV |
| CX266 | MLHHLKELAAVAGIHMILIYLCRFLLRRSRNVLFTVSNSLRFRLKVLTVLLYICLSVMLFYLFGSIMPLPPWGLVVGWVMALIAVELAYAFIFPYSFRYIADNDDDKMVILPV |
| CX267 | MLHHLKELAAVAGIHMILIYLCRFLLRRSRNVLFTVSNSLRFRLKVLTVLLYICLSVMLFYLFGSIMPLPPWGLVVGWVMALIAVELAYAFIFPYSFRYIADNDDDKMVILPV |
| CX268 | MLHHLKELAAVAGIHMILIYLCRFLLRRSRNVLFTVSNSLRFRLKVLTVLLYICLSVMLFYLFGSIMPLPPWGLVVGWVMALIAVELAYAFIFPYSFRYIADNDDDKMVILPV |
| CX269 | MLHHLKELAAVAGIHMILIYLCRFLLRRSRNVLFTVSNSLRFRLKVLTVLLYICLSVMLFYLFGSIMPLPPWGLVVGWVMALIAVELAYAFIFPYSFRYIADNDDDKMVILPV |
| CX270 | MLHHLKELAAVAGIHMILIYLCRFLLRRSRNVLFTVSNSLRFRLKVLTVLLYICLSVMLFYLFGSIMPLPPWGLVVGWVMALIAVELAYAFIFPYSFRYIADNDDDKMVILPV |
| CX273 | MLHHLKELAAVAGIHMILIYLCRFLLRRSRNVLFTVSNSLRFRLKVLTVLLYICLSVMLFYLFGSIMPLPPWGLVVGWVMALIAVELAYAFIFPYSFRYIADNDDDKMVILPV |
| CX274 | MLHHLKELAAVAGIHMILIYLCRFLLRLRRNVLFTVSNSLRFRLKVLTVLLYICLSVMLFYLFGSIMPLPPWGLVVGWVMALIAVELAYAFIFPYSFRYIADNDDDKMVILPV |
| CX275 | MLHHLKELAAVAGIHMILIYLCRFLLRRSRNVLFTVSNSLRFRLKVLTVLLYICLSVMLFYLFGSIMPLPPWGLVVGWVMALIAVELAYAFIFPYSFRYIADNDDDKMVILPV |
| CX276 | MLHHLKELAAVAGIHMILIYLCRFLLRRSRNVLFTVSNSLRFRLKVLTVLLYICLSVMLFYLFGSIMPLPPWGLVVGWVMALIAVELAYAFIFPYSFRYIADNDDDKMVILPV |
| CX277 | MLHQLKELAAVAGIHMILIYLCRFLLRRSRNVLFTVSNSLRFRLKVLTVLLYICLLVMLFYLFGSIMPLPPWGLVVGWVMALIAVELAYAFIFPYSFRYIADNDDDKMVILPV |
| CX278 | MLHHLKELAAVAGIHMILIYLCRFLLRRSRNVLFTVSNSLRFRLKVLTVLLYICLSVMLFYLFGSIMPLPPWGLVVGWVMALIAVELAYAFIFPYSFRYIADNDDDKMVILPV |
| CX280 | MLHHLKELAAVAGIHMILIYLCRFLLRRSRNVLFTVSNSLRFRLKVLTVLLYICLSVMLFYLFGSIMPLPPWGLVVGWVMALIAVELAYAFIFPYSFRYIADNDDDKMVILPV |
| CX281 | MLHHLKELAAVAGIHMILIYLCRFLLRRSRNVLFTVSNSLRFRLKVLTVLLYICLSVMLFYLFGSIMPLPPWGLVVGWVMALIAVELAYAFIFPYSFRYIADNDDDKMVILPV |
| CX282 | MLHQLKELAAVAGIHMILIYLCRFLLRRSRNVLFTVSNSLRFRLKVLTVLLYICLLVMLFYLFGSIMPLPPWGLVVGWVMALIAVELAYAFIFPYSFRYIADNDDDKMVILPV |
| CX284 | MLHQLKELAAVAGIHMILIYLCRFLLRRSRNVLFTVSNSLRFRLKVLTVLLYICLLVMLFYLFGSIMPLPPWGLVVGWVMALIAVELAYAFIFPYSFRYIADNDDDKMVILPV |
| CX285 | MLHHLKELAAVAGIHMILIYLCRFLLRRSRNVLFTVSNSLRFRLKVLTVLLYICLSVMLFYLFGSIMPLPPWGLVVGWVMALIAVELAYAFIFPYSFRYIADNDDDKMVILPV |
| CX286 | MLHHLKELAAVAGIHMILIYLCRFLLRRSRNVLFTVSNSLRFRLKVLTVLLYICLSVMLFYLFGSIMPLPPWGLVVGWVMALIAVELAYAFIFPYSFRYIADNDDDKMVILPV |
| CX287 | MLHHLKELAAVAGIHMILIYLCRFLLRRSRNVLFTVSNSLRFRLKVLTVLLYICLSVMLFYLFGSIMPLPPWGLVVGWVMALIAVELAYAFIFPYSFRYIADNDDDKMVILPV |
| CX288 | MLHHLKELAAVAGIHMILIYLCRFLLRRSRNVLFTVSNSLRFRLKVLTVLLYICLSVMLFYLFGSIMPLPPWGLVVGWVMALIAVELAYAFIFPYSFRYIADNDDDKMVILPV |
| CX290 | MLHHLKELAAVAGIHMILIYLCRFLLRRSRNVLFTVSNSLRFRLKVLTVLLYICLSVMLFYLFGSIMPLPPWGLVVGWVMALIAVELAYAFIFPYSFRYIADNDDDKMVILPV |
| CX291 | MLHHLKELAAVAGIHMILIYLCRFLLRRSRNVLFTVSNSLRFRLKVLTVLLYICLSVMLFYLFGSIMPLPPWGLVVGWVMALIAVELAYAFIFPYSFRYIADNDDDKMVILPV |
| CX296 | MLHHLKELAAVAGIHMILIYLCRFLLRRSRNVLFTVSNSLRFRLKVLTVLLYICLSVMLFYLFGSIMPLPPWGLVVGWVMALIAVELAYAFIFPYSFRYIADNDDDKMVILPV |
| CX303 | MLHHLKELAAVAGIHMILIYLCRFLLRRSRNVLFTVSNSLRFRLKVLTVLLYICLSVMLFYLFGSIMPLPPWGLVVGWVMALIAVELAYAFIFPYSFRYIADNDDDKMVILPV |
| CX304 | MLHHLKELAAVAGIHMILIYLCRFLLRRSRNVLFTVSNSLRFRLKVLTVLLYICLSVMLFYLFGSIMPLPPWGLVVGWVMALIAVELAYAFIFPYSFRYIADNDDDKMVILPV |
| CX305 | MLHHLKELAAVAGIHMILIYLCRFLLRLRRNVLFTVSNSLRFRLKVLTVLLYICLSVMLFYLFGSIMPLPPWGLVVGWVMALIAVELAYAFIFPYSFRYIADNDDDKMVILPV |
| CX306 | MLHHLKELAAVAGIHMILIYLCRFLLRRSRNVLFTVSNSLRFRLKVLTVLLYICLSVMLFYLFGSIMPLPPWGLVVGWVMALIAVELAYAFIFPYSFRYIADNDDDKMVILPV |
| CX307 | MLHHLKELAAVAGIHMILIYLCRFLLRRSRNVLFTVSNSLRFRLKVLTVLLYICLSVMLFYLFGSIMPLPPWGLVVGWVMALIAVELAYAFIFPYSFRYIADNDDDKMVILPV |
| CX313 | MLHHLKELAAVAGIHMILIYLCRFLLRRSRNVLFTVSNSLRFRLKVLTVLLYICLSVMLFYLFGSIMPLPPWGLVVGWVMALIAVELAYAFIFPYSFRYIADNDDDKMVILPV |
| CX314 | MLHHLKELAAVAGIHMILIYLCRFLLRRSRNVLFTVSNSLRFRLKVLTVLLYICLSVMLFYLFGSIMPLPPWGLVVGWVMALIAVELAYAFIFPYSFRYIADNDDDKMVILPV |
| CX315 | MLHQLKELAAVAGIHMILIYLCRFLLRRSRNVLFTVSNSLRFRLKVLTVLLYICLLVMLFYLFGSIMPLPPWGLVVGWVMALIAVELAYAFIFPYSFRYIADNDDDKMVILPV |
| CX316 | MLHHLKELAAVAGIHMILIYLCRFLLRRSRNVLFTVSNSLRFRLKVLTVLLYICLSVMLFYLFGSIMPLPPWGLVVGWVMALIAVELAYAFIFPYSFRYIADNDDDKMVILPV |
| CX317 | MLHHLKELAAVAGIHMILIYLCRFLLRRSRNVLFTVSNSLRFRLKVLTVLLYICLSVMLFYLFGSIMPLPPWGLVVGWVMALIAVELAYAFIFPYSFRYIADNDDDKMVILPV |
| CX318 | MLHHLKELAAVAGIHMILIYLCRFLLRRSRNVLFTVSNSLRFRLKVLTVLLYICLSVMLFYLFGSIMPLPPWGLVVGWVMALIAVELAYAFIFPYSFRYIADNDDDKMVILPV |
| CX319 | MLHHLKELAAVAGIHMILIYLCRFLLRRSRNVLFTVSNSLRFRLKVLTVLLYICLSVMLFYLFGSIMPLPPWGLVVGWVMALIAVELAYAFIFPYSFRYIADNDDDKMVILPV |
| CX328 | MLHHLKELAAVAGIHMILIYLCRFLLRRSRNVLFTVSNSLRFRLKVLTVLLYICLSVMLFYLFGSIMPLPPWGLVVGWVMALIAVELAYAFIFPYSFRYIADNDDDKMVILPV |
| CX329 | MLHHLKELAAVAGIHMILIYLCRFLLRRSRNVLFTVSNSLRFRLKVLTVLLYICLSVMLFYLFGSIMPLPPWGLVVGWVMALIAVELAYAFIFPYSFRYIADNDDDKMVILPV |
| CX330 | MLHHLKELAAVAGIHMILIYLCRFLLRRSRNVLFTVSNSLRFRLKVLTVLLYICLSVMLFYLFGSIMPLPPWGLVVGWVMALIAVELAYAFIFPYSFRYIADNDDDKMVILPV |
| CX340 | MLHHLKELAAVAGIHMILIYLCRFLLRRSRNVLFTVSNSLRFRLKVLTVLLYICLSVMLFYLFGSIMPLPPWGLVVGWVMALIAVELAYAFIFPYSFRYIADNDDDKMVILPV |
| CX341 | MLHHLKELAAVAGIHMILIYLCRFLLRRSRNVLFTVSNSLRFRLKVLTVLLYICLSVMLFYLFGSIMPLPPWGLVVGWVMALIAVELAYAFIFPYSFRYIADNDDDKMVILPV |
| CX342 | MLHHLKELAAVAGIHMILIYLCRFLLRRSRNVLFTVSNSLRFRLKVLTVLLYICLSVMLFYLFGSIMPLPPWGLVVGWVMALIAVELAYAFIFPYSFRYIADNDDDKMVILPV |
| CX343 | MLHHLKELAAVAGIHMILIYLCRFLLRRSRNVLFTVSNSLRFRLKVLTVLLYICLSVMLFYLFGSIMPLPPWGLVVGWVMALIAVELAYAFIFPYSFRYIADNDDDKMVILPV |
| CX344 | MLHHLKELAAVAGIHMILIYLCRFLLRRSRNVLFTVSNSLRFRLKVLTVLLYICLSVMLFYLFGSIMPLPPWGLVVGWVMALIAVELAYAFIFPYSFRYIADNDDDKMVILPV |
| CX345 | MLHHLKELAAVAGIHMILIYLCRFLLRRSRNVLFTVSNSLRFRLKVLTVLLYISLSAMLLYLFGSIMPLPPWGLVVGWVMALIAVELAYAFIFPYSFRYIADNDDDKMVILPV |
| CX346 | MLHHLKELAAVAGIHMILIYLCRFLLRLRRNVLFTVSNSLRFRLKVLTVLLYICLSVMLFYLFGSIMPLPPWGLVVGWVMALIAVELAYAFIFPYSFRYIADNDDDKMVILPV |
| CX347 | MLHHLKELAAVAGIHMILIYLCRFLLRRSRNVLFTVSNSLRFRLKVLTVLLYICLSVMLFYLFGSIMPLPPWGLVVGWVMALIAVELAYAFIFPYSFRYIADNDDDKMVILPV |
| CX348 | MLHHLKELAAVAGIHMILIYLCRFLLRRSRNVLFTVSNSLRFRLKVLTVLLYICLSVMLFYLFGSIMPLPPWGLVVGWVMALIAVELAYAFIFPYSFRYIADNDDDKMVILPV |
| CX349 | MLHHLKELAAVAGIHMILIYLCRFLLRRSRNVLFTVSNSLRFRLKVLTVLLYICLSVMLFYLFGSIMPLPPWGLVVGWVMALIAVELAYAFIFPYSFRYIADNDDDKMVILPV |
| CX350 | MLHHLKELAAVAGIHMILIYLCRFLLRRSRNVLFTVSNSLRFRLKVLTVLLYICLSVMLFYLFGSIMPLPPWGLVVGWVMALIAVELAYAFIFPYSFRYIADNDDDKMVILPV |
| CX351 | MLHQLKELAAVAGIHMILIYLCRFLLRRSRNVLFTVSNSLRFRLKVLTVLLYICLLVMLFYLFGSIMPLPPWGLVVGWVMALIAVELAYAFIFPYSFRYIADNDDDKMVILPV |
| CX352 | MLHHLKELAAVAGIHMILIYLCRFLLRRSRNVLFTVSNSLRFRLKVLTVLLYICLSVMLFYLFGSIMPLPPWGLVVGWVMALIAVELAYAFIFPYSFRYIADNDDDKMVILPV |

CX353 MLHHLKELAAVAGIHMILIYLCRFLLRRSRNVLFTVSNSLRFRLKVLTVLLYICLSVMLFYLFGSIMPLPPWGLVVGWVMALIAVELAYAFIFPYSFRYIADNDDDKMVILPV
CX354 MLHHLKELAAVAGIHMILIYLCRFLLRRSRNVLFTVSNSLRFRLKVLTVLLYICLSVMLFYLFGSIMPLPPWGLVVGWVMALIAVELAYAFIFPYSFRYIADNDDDKMVILPV
CX355 MLHHLKELAAVAGIHMILIYLCRFLLRRSRNVLFTVSNSLRFRLKVLTVLLYICLSVMLFYLFGSIMPLPPWGLVVGWVMALIAVELAYAFIFPYSFRYIADNDDDKMVILPV
CX356 MLHHLKELAAVAGIHMILIYLCRFLLRRSRNVLFTVSNSLRFRLKVLTVLLYISLSAMLLYLFGSIMPLPPWGLVVGWVMALIAVELAYAFIFPYSFRYIADNDDDKMVILPV
CX357 MLHHLKELAAVAGIHMILIYLCRFLLRRSRNVLFTVSNSLRFRLKVLTVLLYICLSVMLFYLFGSIMPLPPWGLVVGWVMALIAVELAYAFIFPYSFRYIADNDDDKMVILPV
CX358 MLHHLKELAAVAGIHMILIYLCRFLLRRSRNVLFTVSNSLRFRLKVLTVLLYICLSVMLFYLFGSIMPLPPWGLVVGWVMALIAVELAYAFIFPYSFRYIADNDDDKMVILPV
CX359 MLHHLKELAAVAGIHMILIYLCRFLLRRSRNVLFTVSNSLRFRLKVLTVLLYICLSVMLFYLFGSIMPLPPWGLVVGWVMALIAVELAYAFLFPYSLRYIAHNDDDKMVILPV
CX360 MLHHLKELAAVAGIHMILIYLCRFLLRRSRNVLFTVSNSLRFRLKVLTVLLYICLSVMLFYLFGSIMPLPPWGLVVGWVMALIAVELAYAFIFPYSFRYIADNDDDKMVILPV
CX361 MLHHLKELAAVAGIHMILIYLCRFLLRRSRNVLFTVSNSLRFRLKVLTVLLYICLSVMLFYLFGSIMPLPPWGLVVGWVMALIAVELAYAFIFPYSFRYIADNDDDKMVILPV
CX362 MLHHLKELAAVAGIHMILIYLCRFLLRRSRNVLFTVSNSLRFRLKVLTVLLYICLSVMLFYLFGSIMPLPPWGLVVGWVMALIAVELAYAFIFPYSFRYIADNDDDKMVILPV
CX363 MLHHLKELAAVAGIHMILIYLCRFLLRRSRNVLFTVSNSLRFRLKVLTVLLYICLSVMLFYLFGSIMPLPPWGLVVGWVMALIAVELAYAFIFPYSFRYIADNDDDKMVILPV
CX364 MLHHLKELAAVAGIHMILIYLCRFLLRRSRNVLFTVSNSLRFRLKVLTVLLYICLSVMLFYLFGSIMPLPPWGLVVGWVMALIAVELAYAFIFPYSFRYIADNDDDKMVILPV
CX365 MLHHLKELAAVAGIHMILIYLCRFLLRRSRNVLFTVSNSLRFRLKVLTVLLYICLSVMLFYLFGSIMPLPPWGLVVGWVMALIAVELAYAFIFPYSFRYIADNDDDKMVILPV
CX366 MLHHLKELAAVAGIHMILIYLCRFLLRRSRNVLFTVSNSLRFRLKVLTVLLYICLSVMLFYLFGSIMPLPPWGLVVGWVMALIAVELAYAFIFPYSFRYIADNDDDKMVILPV
CX367 MLHHLKELAAVAGIHMILIYLCRFLLRRSRNVLFTVSNSLRFRLKVLTVLLYICLSVMLFYLFGSIMPLPPWGLVVGWVMALIAVELAYAFIFPYSFRYIADNDDDKMVILPV
CX368 MLHHLKELAAVAGIHMILIYLCRFLLRRSRNVLFTVSNSLRFRLKVLTVLLYICLSVMLFYLFGSIMPLPPWGLVVGWVMALIAVELAYAFIFPYSFRYIADNDDDKMVILPV
CX369 MLHHLKELAAVAGIHMILIYLCRFLLRRSRNVLFTVSNSLRFRLKVLTVLLYICLSVMLFYLFGSIMPLPPWGLVVGWVMALIAVELAYAFIFPYSFRYIADNDDDKMVILPV
CX370 MLHHLKELAAVAGIHMILIYLCRFLLRRSRNVLFTVSNSLRFRLKVLTVLLYICLSVMLFYLFGSIMPLPPWGLVVGWVMALIAVELAYAFIFPYSFRYIADNDDDKMVILPV
CX371 MLHHLKELAAVAGIHMILIYLCRFLLRRSRNVLFTVSNSLRFRLKVLTVLLYICLSVMLFYLFGSIMPLPPWGLVVGWVMALIAVELAYAFLFPYSLRYIAHNDDDKMVILPV
CX372 MLHHLKELAAVAGIHMILIYLCRFLLRRSRNVLFTVSNSLRFRLKVLTVLLYICLSVMLFYLFGSIMPLPPWGLVVGWVMALIAVELAYAFIFPYSFRYIADNDDDKMVILPV
CX373 MLHHLKELAAVAGIHMILIYLCRFLLRRSRNVLFTVSNSLRFRLKVLTVLLYICLSVMLFYLFGSIMPLPPWGLVVGWVMALIAVELAYAFIFPYSFRYIADNDDDKMVILPV
CX374 MLHHLKELAAVAGIHMILIYLCRFLLRRSRNVLFTVSNSLRFRLKVLTVLLYICLSVMLFYLFGSIMPLPPWGLVVGWVMALIAVELAYAFIFPYSFRYIADNDDDKMVILPV
CX375 MLHHLKELAAVAGIHMILIYLCRFLLRRSRNVLFTVSNSLRFRLKVLTVLLYICLSVMLFYLFGSIMPLPPWGLVVGWVMALIAVELAYAFIFPYSFRYIADNDDDKMVILPV
CX376 MLHHLKELAAVAGIHMILIYLCRFLLRRSRNVLFTVSNSLRFRLKVLTVLLYICLSVMLFYLFGSIMPLPPWGLVVGWVMALIAVELAYAFIFPYSFRYIADNDDDKMVILPV
CX377 MLHHLKELAAVAGIHMILIYLCRFLLRRSRNVLFTVSNSLRFRLKVLTVLLYICLSVMLFYLFGSIMPLPPWGLVVGWVMALIAVELAYAFIFPYSFRYIADNDDDKMVILPV
CX378 MLHHLKELAAVAGIHMILIYLCRFLLRRSRNVLFTVSNSLRFRLKVLTVLLYICLSVMLFYLFGSIMPLPPWGLVVGWVMALIAVELAYAFIFPYSFRYIADNDDDKMVILPV
CX379 MLHHLKELAAVAGIHMILIYLCRFLLRRSRNVLFTVSNSLRFRLKVLTVLLYICLSVMLFYLFGSIMPLPPWGLVVGWVMALIAVELAYAFIFPYSFRYIADNDDDKMVILPV
CX380 MLHHLKELAAVAGIHMILIYLCRFLLRRSRNVLFTVSNSLRFRLKVLTVLLYICLSVMLFYLFGSIMPLPPWGLVVGWVMALIAVELAYAFIFPYSFRYIADNDDDKMVILPV
CX381 MLHHLKELAAVAGIHMILIYLCRFLLRRSRNVLFTVSNSLRFRLKVLTVLLYICLSVMLFYLFGSIMPLPPWGLVVGWVMALIAVELAYAFIFPYSFRYIADNDDDKMVILPV
CX382 MLHHLKELAAVAGIHMILIYLCRFLLRRSRNVLFTVSNSLRFRLKVLTVLLYICLSVMLFYLFGSIMPLPPWGLVVGWVMALIAVELAYAFIFPYSFRYIADNDDDKMVILPV
CX383 MLHQLKELAAVAGIHMILIYLCRFLLRRSRNVLFTVSNSLRFRLKVLTVLLYICLLVMLFYLFGSIMPLPPWGLVVGWVMALIAVELAYAFIFPYSFRYIADNDDDKMVILPV
CX384 MLHQLKELAAVAGIHMILIYLCRFLLRRSRNVLFTVSNSLRFRLKVLTVLLYICLLVMLFYLFGSIMPLPPWGLVVGWVMALIAVELAYAFIFPYSFRYIADNDDDKMVILPV
CX385 MLHHLKELAAVAGIHMILIYLCRFLLRRSRNVLFTVSNSLRFRLKVLTVLLYICLSVMLFYLFGSIMPLPPWGLVVGWVMALIAVELAYAFIFPYSFRYIADNDDDKMVILPV
CX386 MLHHLKELAAVAGIHMILIYLCRFLLRRSRNVLFTVSNSLRFRLKVLTVLLYICLSVMLFYLFGSIMPLPPWGLVVGWVMALIAVELAYAFIFPYSFRYIADNDDDKMVILPV
CX387 MLHHLKELAAVAGIHMILIYLCRFLLRRSRNVLFTVSNSLRFRLKVLTVLLYICLSVMLFYLFGSIMPLPPWGLVVGWVMALIAVELAYAFIFPYSFRYIADNDDDKMVILPV
CX388 MLHHLKELAAVAGIHMILIYLCRFLLRRSRNVLFTVSNSLRFRLKVLTVLLYICLSVMLFYLFGSIMPLPPWGLVVGWVMALIAVELAYAFIFPYSFRYIADNDDDKMVILPV
CX389 MLHHLKELAAVAGIHMILIYLCRFLLRRSRNVLFTVSNSLRFRLKVLTVLLYICLSVMLFYLFGSIMPLPPWGLVVGWVMALIAVELAYAFIFPYSFRYIADNDDDKMVILPV
CX390 MLHHLKELAAVAGIHMILIYLCRFLLRRSRNVLFTVSNSLRFRLKVLTVLLYICLSVMLFYLFGSIMPLPPWGLVVGWVMALIAVELAYAFIFPYSFRYIADNDDDKMVILPV
CX391 MLHHLKELAAVAGIHMILIYLCRFLLRRSRNVLFTVSNSLRFRLKVLTVLLYICLSAMLLYLFGSIMPLPPWGLVVGWVMALIAVELAYAFIFPYSFRYIADNDDDKMVILPV
CX392 MLHHLKELAAVAGIHMILIYLCRFLLRRSRNVLFTVSNSLRFRLKVLTVLLYICLSVMLFYLFGSIMPLPPWGLVVGWVMALIAVELAYAFIFPYSFRYIADNDDDKMVILPV
CX393 MLHHLKELAAVAGIHMILIYLCRFLLRRSRNVLFTVSNSLRFRLKVLTVLLYICLSVMLFYLFGSIMPLPPWGLVVGWVMALIAVELAYAFIFPYSFRYIADNDDDKMVILPV
CX394 MLHHLKELAAVAGIHMILIYLCRFLLRRSRNVLFTVSNSLRFRLKVLTVLLYICLSVMLFYLFGSIMPLPPWGLVVGWVMALIAVELAYAFIFPYSFRYIADNDDDKMVILPV
CX395 MLHHLKELAAVAGIHMILIYLCRFLLRRSRNVLFTVSNSLRFRLKVLTVLLYICLSVMLFYLFGSIMPLPPWGLVVGWVMALIAVELAYAFIFPYSFRYIADNDDDKMVILPV
CX396 MLHHLKELAAVAGIHMILIYLCRFLLRRSRNVLFTVSNSLRFRLKVLTVLLYISLSAMLLYLFGSIMPLPPWGLVVGWVMALIAVELAYAFIFPYSFRYIADNDDDKMVILPV
CX397 MLHHLKELAAVAGIHMILIYLCRFLLRRSRNVLFTVSNSLRFRLKVLTVLLYICLSVMLFYLFGSIMPLPPWGLVVGWVMALIAVELAYAFIFPYSFRYIADNDDDKMVILPV
CX399 MLHHLKELAAVAGIHMILIYLCRFLLRRSRNVLFTVSNSLRFRLKVLTVLLYICLSVMLFYLFGSIMPLPPWGLVVGWVMALIAVELAYAFIFPYSFRYIADNDDDKMVILPV
CX403 MLHHLKELAAVAGIHMILIYLCRFLLRRSRNVLFTVSNSLRFRLKVLTVLLYICLSVMLFYLFGSIMPLPPWGLVVGWVMALIAVELAYAFIFPYSFRYIADNDDDKMVILPV
CX408 MLHHLKELAAVAGIHMILIYLCRFLLRRSRNVLFTVSNSLRFRLKVLTVLLYICLSVMLFYLFGSIMPLPPWGLVVGWVMALIAVELAYAFIFPYSFRYIADNDDDKMVILPV
CX409 MLHHLKELAAVAGIHMILIYLCRFLLRLSRNVLFTVSNSLRFRLKVLTVLLYICLSVMLFYLFGSIMPLPPWGLVVGWVMALIAVELAYAFIFPYSFRYIADNDDDKMVILPV
CX410 MLHHLKELAAVAGIHMILIYLCRFLLRRSRNVLFTVSNSLRFRLKVLTVLLYICLSVMLFYLFGSIMPLPPWGLVVGWVMALIAVELAYAFIFPYSFRYIADNDDDKMVILPV
CX411 MLHHLKELAAVAGIHMILIYLCRFLLRLSRNVLFTVSNSLRFRLKVLTVLLYICLSVMLFYLFGSIMPLPPWGLVVGWVMALIAVELAYAFIFPYSFRYIADNDDDKMVILPV
CX413 MLHHLKELAAVAGIHMILIYLCRFLLRRSRNVLFTVSNSLRFRLKVLTVLLYICLSVMLFYLFGSIMPLPPWGLVVGWVMALIAVELAYAFIFPYSFRYIADNDDDKMVILPV
CX414 MLHHLKELAAVAGIHMILIYLCRFLLRLSRNVLFTVSNSLRFRLKVLTVLLYICLSVMLFYLFGSIMPLPPWGLVVGWVMALIAVELAYAFIFPYSFRYIADNDDDKMVILPV
CX416 MLHHLKELAAVAGIHMILIYLCRFLLRRSRNVLFTVSNSLRFRLKVLTVLLYICLSVMLFYLFGSIMPLPPWGLVVGWVMALIAVELAYAFIFPYSFRYIADNDDDKMVILPV
CX417 MLHHLKELAAVAGIHMILIYLCRFLLRRSRNVLFTVSNSLRFRLKVLTVLLYICLSVMLFYLFGSIMPLPPWGLVVGWVMALIAVELAYAFIFPYSFRYIADNDDDKMVILPV
CX418 MLHHLKELAAVAGIHMILIYLCRFLLRRSRNVLFTVSNSLRFRLKVLTVLLYICLSVMLFYLFGSIMPLPPWGLVVGWVMALIAVELAYAFIFPYSFRYIADNDDDKMVILPV
CX419 MLHHLKELAAVAGIHMILIYLCRFLLRLSRNVLFTVSNSLRFRLKVLTVLLYICLSVMLFYLFGSIMPLPPWGLVVGWVMALIAVELAYAFIFPYSFRYIADNDDDKMVILPV
CX420 MLHHLKELAAVAGIHMILIYLCRFLLRRSRNVLFTVSNSLRFRLKVLTVLLYICLSVMLFYLFGSIMPLPPWGLVVGWVMALIAVELAYAFIFPYSFRYIADNDDDKMVILPV
CX421 MLHHLKELAAVAGIHMILIYLCRFLLRRSRNVLFTVSNSLRFRLKVLTVLLYICLSVMLFYLFGSIMPLPPWGLVVGWVMALIAVELAYAFIFPYSFRYIADNDDDKMVILPV
CX422 MLHHLKELAAVAGIHMILIYLCRFLLRRSRNVLFTVSNSLRFRLKVLTVLLYICLSVMLFYLFGSIMPLPPWGLVVGWVMALIAVELAYAFIFPYSFRYIADNDDDKMVILPV
CX423 MLHHLKELAAVAGIHMILIYLCRFLLRRSRNVLFTVSNSLRFRLKVLTVLLYICLSVMLFYLFGSIMPLPPWGLVVGWVMALIAVELAYAFIFPYSFRYIADNDDDKMVILPV
CX424 MLHHLKELAAVAGIHMILIYLCRFLLRRSRNVLFTVSNSLRFRLKVLTVLLYICLSVMLFYLFGSIMPLPPWGLVVGWVMALIAVELAYAFIFPYSFRYIADNDDDKMVILPV
CX425 MLHHLKELAAVAGIHMILIYLCRFLLRRSRNVLFTVSNSLRFRLKVLTVLLYICLSVMLFYLFGSIMPLPPWGLVVGWVMALIAVELAYAFIFPYSFRYIADNDDDKMVILPV
CX426 MLHHLKELAAVAGIHMILIYLCRFLLRRSRNVLFTVSNSLRFRLKVLTVLLYICLSVMLFYLFGSIMPLPPWGLVVGWVMALIAVELAYAFIFPYSFRYIADNDDDKMVILPV
CX427 MLHHLKELAAVAGIHMILIYLCRFLLRRSRNVLFTVSNSLRFRLKVLTVLLYICLSVMLFYLFGSIMPLPPWGLVVGWVMALIAVELAYAFIFPYSFRYIADNDDDKMVILPV
CX428 MLHHLKELAAVAGIHMILIYLCRFLLRRSRNVLFTVSNSLRFRLKVLTVLLYICLSVMLFYLFGSIMPLPPWGLVVGWVMALIAVELAYAFIFPYSFRYIADNDDDKMVILPV
CX429 MLHHLKELAAVAGIHMILIYLCRFLLRRSRNVLFTVSNSLRFRLKVLTVLLYICLSVMLFYLFGSIMPLPPWGLVVGWVMALIAVELAYAFIFPYSFRYIADNDDDKMVILPV
CX431 MLHHLKELAAVAGIHMILIYLCRFLLRRSRNVLFTVSNSLRFRLKVLTVLLYICLSVMLFYLFGSIMPLPPWGLVVGWVMALIAVELAYAFIFPYSFRYIADNDDDKMVILPV
CX534 MLHHLKELAAVAGIHMILIYLCRFLLRRSRNVLFTVSNSLRFRLKVLTVLLYICLSVMLFYLFGSIMPLPPWGLVVGWVMALIAVELAYAFIFPYSFRYIADNDDDKMVILPV
CX542 MLHHLKELAAVAGIHMILIYLCRFLLRRSRNVLFTVSNSLRFRLKVLTVLLYICLSVMLFYLFGSIMPLPPWGLVVGWVMALIAVELAYAFIFPYSFRYIADNDDDKMVILPV
CX548 MLHHLKELAAVAGIHMILIYLCRFLLRRSRNVLFTVSNSLRFRLKVLTVLLYICLSVMLFYLFGSIMPLPPWGLVVGWVMALIAVELAYAFIFPYSFRYIADNDDDKMVILPV
CX561 MLHHLKELAAVAGIHMILIYLCRFLLRRSRNVLFTVSNSLRFRLKVLTVLLYICLSVMLFYLFGSIMPLPPWGLVVGWVMALIAVELAYAFIFPYSFRYIADNDDDKMVILPV
CX578 MLHHLKELAAVAGIHMILIYLCRFLLRRSRNVLFTVSNSLRFRLKVLTVLLYICLSVMLFYLFGSIMPLPPWGLVVGWVMALIAVELAYAFIFPYSFRYIADNDDDKMVILPV
CX579 MLHHLKELAAVAGIHMILIYLCRFLLRRSRNVLFTVSNSLRFRLKVLTVLLYICLSVMLFYLFGSIMPLPPWGLVVGWVMALIAVELAYAFIFPYSFRYIADNDDDKMVILPV
IRIS_313-7620 MLHHLKELAAVAGIHMILIYLCRFLLRRSRNVLFTVSNSLRFRLKVLTVLLYICLSVMLFYLFGSIMPLPPWGLVVGWVMALIAVELAYAFIFPYSFRYIADNDDDKMVILPV
IRIS_313-7635 MLHHLKELAAVAGIHMILIYLCRFLLRRSRNVLFTVSNSLRFRLKVLTVLLYICLSVMLFYLFGSIMPLPPWGLVVGWVMALIAVELAYAFIFPYSFRYIADNDDDKMVILPV
IRIS_313-7636 MLHHLKELAAVAGIHMILIYLCRFLLRRSRNVLFTVSNSLRFRLKVLTVLLYICLSVMLFYLFGSIMPLPPWGLVVGWVMALIAVELAYAFIFPYSFRYIADNDDDKMVILPV
IRIS_313-7638 MLHHLKELAAVAGIHMILIYLCRFLLRRSRNVLFTVSNSLRFRLKVLTVLLYICLSVMLFYLFGSIMPLPPWGLVVGWVMALIAVELAYAFIFPYSFRYIADNDDDKMVILPV
IRIS_313-7641 MLHHLKELAAVAGIHMILIYLCRFLLRRSRNVLFTVSNSLRFRLKVLTVLLYICLSVMLFYLFGSIMPLPPWGLVVGWVMALIAVELAYAFIFPYSFRYIADNDDDKMVILPV
IRIS_313-7646 MLHHLKELAAVAGIHMILIYLCRFLLRRSRNVLFTVSNSLRFRLKVLTVLLYICLSVMLFYLFGSIMPLPPWGLVVGWVMALIAVELAYAFIFPYSFRYIADNDDDKMVILPV
IRIS_313-7650 MLHHLKELAAVAGIHMILIYLCRFLLRRSRNVLFTVSNSLRFRLKVLTVLLYICLSVMLFYLFGSIMPLPPWGLVVGWVMALIAVELAYAFIFPYSFRYIADNDDDKMVILPV
IRIS_313-7651 MLHHLKELAAVAGIHMILIYLCRFLLRRSRNVLFTVSNSLRFRLKVLTVLLYICLSVMLFYLFGSIMPLPPWGLVVGWVMALIAVELAYAFIFPYSFRYIADNDDDKMVILPV
IRIS_313-7654 MLHHLKELAAVAGIHMILIYLCRFLLRRSRNVLFTVSNSLRFRLKVLTVLLYICLSVMLFYLFGSIMPLPPWGLVVGWVMALIAVELAYAFIFPYSFRYIADNDDDKMVILPV
IRIS_313-7664 MLHHLKELAAVAGIHMILIYLCRFLLRRSRNVLFTVSNSLRFRLKVLTVLLYICLSVMLFYLFGSIMPLPPWGLVVGWVMALIAVELAYAFIFPYSFRYIADNDDDKMVILPV
IRIS_313-7665 MLHHLKELAAVAGIHMILIYLCRFLLRRSRNVLFTVSNSLRFRLKVLTVLLYICLSVMLFYLFGSIMPLPPWGLVVGWVMALIAVELAYAFIFPYSFRYIADNDDDKMVILPV
IRIS_313-7668 MLHHLKELAAVAGIHMILIYLCRFLLRRSRNVLFTVSNSLRFRLKVLTVLLYICLSVMLFYLFGSIMPLPPWGLVVGWVMALIAVELAYAFIFPYSFRYIADNDDDKMVILPV
IRIS_313-7681 MLHHLKELAAVAGIHMILIYLCRFLLRRSRNVLFTVSNSLRFRLKVLTVLLYICLSVMLFYLFGSIMPLPPWGLVVGWVMALIAVELAYAFIFPYSFRYIADNDDDKMVILPV
IRIS_313-7684 MLHHLKELAAVAGIHMILIYLCRFLLRRSRNVLFTVSNSLRFRLKVLTVLLYICLSVMLFYLFGSIMPLPPWGLVVGWVMALIAVELAYAFIFPYSFRYIADNDDDKMVILPV
IRIS_313-7685 MLHHLKELAAVAGIHMILIYLCRFLLRRSRNVLFTVSNSLRFRLKVLTVLLYICLSVMLFYLFGSIMPLPPWGLVVGWVMALIAVELAYAFIFPYSFRYIADNDDDKMVILPV
IRIS_313-7688 MLHHLKELAAVAGIHMILIYLCRFLLRRSRNVLFTVSNSLRFRLKVLTVLLYICLSVMLFYLFGSIMPLPPWGLVVGWVMALIAVELAYAFIFPYSFRYIADNDDDKMVILPV
IRIS_313-7689 MLHHLKELAAVAGIHMILIYLCRFLLRRSRNVLFTVSNSLRFRLKVLTVLLYICLSVMLFYLFGSIMPLPPWGLVVGWVMALIAVELAYAFIFPYSFRYIADNDDDKMVILPV
IRIS_313-7690 MLHHLKELAAVAGIHMILIYLCRFLLRRSRNVLFTVSNSLRFRLKVLTVLLYICLSVMLFYLFGSIMPLPPWGLVVGWVMALIAVELAYAFIFPYSFRYIADNDDDKMVILPV
IRIS_313-7691 MLHHLKELAAVAGIHMILIYLCRFLLRRSRNVLFTVSNSLRFRLKVLTVLLYICLSVMLFYLFGSIMPLPPWGLVVGWVMALIAVELAYAFIFPYSFRYIADNDDDKMVILPV
IRIS_313-7696 MLHHLKELAAVAGIHMILIYLCRFLLRRSRNVLFTVSNSLRFRLKVLTVLLYICLSVMLFYLFGSIMPLPPWGLVVGWVMALIAVELAYAFIFPYSFRYIADNDDDKMVILPV
IRIS_313-7698 MLHHLKELAAVAGIHMILIYLCRFLLRRSRNVLFTVSNSLRFRLKVLTVLLYICLSVMLFYLFGSIMPLPPWGLVVGWVMALIAVELAYAFIFPYSFRYIADNDDDKMVILPV
IRIS_313-7699 MLHHLKELAAVAGIHMILIYLCRFLLRRSRNVLFTVSNSLRFRLKVLTVLLYICLSVMLFYLFGSIMPLPPWGLVVGWVMALIAVELAYAFIFPYSFRYIADNDDDKMVILPV
IRIS_313-7719 MLHHLKELAAVAGIHMILIYLCRFLLRRSRNVLFTVSNSLRFRLKVLTVLLYICLSVMLFYLFGSIMPLPPWGLVVGWVMALIAVELAYAFIFPYSFRYIADNDDDKMVILPV
IRIS_313-7720 MLHHLKELAAVAGIHMILIYLCRFLLRRSRNVLFTVSNSLRFRLKVLTVLLYICLSVMLFYLFGSIMPLPPWGLVVGWVMALIAVELAYAFIFPYSFRYIADNDDDKMVILPV
IRIS_313-7722 MLHHLKELAAVAGIHMILIYLCRFLLRRSRNVLFTVSNSLRFRLKVLTVLLYICLSVMLFYLFGSIMPLPPWGLVVGWVMALIAVELAYAFIFPYSFRYIADNDDDKMVILPV
IRIS_313-7725 MLHHLKELAAVAGIHMILIYLCRFLLRRSRNVLFTVSNSLRFRLKVLTVLLYICLSVMLFYLFGSIMPLPPWGLVVGWVMALIAVELAYAFIFPYSFRYIADNDDDKMVILPV
IRIS_313-7728 MLHHLKELAAVAGIHMILIYLCRFLLRRSRNVLFTVSNSLRFRLKVLTVLLYICLSVMLFYLFGSIMPLPPWGLVVGWVMALIAVELAYAFIFPYSFRYIADNDDDKMVILPV
IRIS_313-7736 MLHHLKELAAVAGIHMILIYLCRFLLRRSRNVLFTVSNSLRFRLKVLTVLLYICLSVMLFYLFGSIMPLPPWGLVVGWVMALIAVELAYAFIFPYSFRYIADNDDDKMVILPV
IRIS_313-7758 MLHHLKELAAVAGIHMILIYLCRFLLRRSRNVLFTVSNSLRFRLKVLTVLLYICLSVMLFYLFGSIMPLPPWGLVVGWVMALIAVELAYAFIFPYSFRYIADNDDDKMVILPV
IRIS_313-7766 MLHHLKELAAVAGIHMILIYLCRFLLRRSRNVLFTVSNSLRFRLKVLTVLLYICLSVMLFYLFGSIMPLPPWGLVVGWVMALIAVELAYAFIFPYSFRYIADNDDDKMVILPV
IRIS_313-7769 MLHHLKELAAVAGIHMILIYLCRFLLRRSRNVLFTVSNSLRFRLKVLTVLLYICLSVMLFYLFGSIMPLPPWGLVVGWVMALIAVELAYAFIFPYSFRYIADNDDDKMVILPV
IRIS_313-7770 MLHHLKELAAVAGIHMILIYLCRFLLRRSRNVLFTVSNSLRFRLKVLTVLLYICLSVMLFYLFGSIMPLPPWGLVVGWVMALIAVELAYAFIFPYSFRYIADNDDDKMVILPV
IRIS_313-7773 MLHHLKELAAVAGIHMILIYLCRFLLRRSRNVLFTVSNSLRFRLKVLTVLLYICLSVMLFYLFGSIMPLPPWGLVVGWVMALIAVELAYAFIFPYSFRYIADNDDDKMVILPV
IRIS_313-7778 MLHHLKELAAVAGIHMILIYLCRFLLRRSRNVLFTVSNSLRFRLKVLTVLLYICLSVMLFYLFGSIMPLPPWGLVVGWVMALIAVELAYAFIFPYSFRYIADNDDDKMVILPV
IRIS_313-7780 MLHHLKELAAVAGIHMILIYLCRFLLRRSRNVLFTVSNSLRFRLKVLTVLLYICLSVMLFYLFGSIMPLPPWGLVVGWVMALIAVELAYAFIFPYSFRYIADNDDDKMVILPV
IRIS_313-7792 MLHHLKELAAVAGIHMILIYLCRFLLRRSRNVLFTVSNSLRFRLKVLTVLLYICLSVMLFYLFGSIMPLPPWGLVVGWVMALIAVELAYAFIFPYSFRYIADNDDDKMVILPV
IRIS_313-7793 MLHHLKELAAVAGIHMILIYLCRFLLRRSRNVLFTVSNSLRFRLKVLTVLLYICLSVMLFYLFGSIMPLPPWGLVVGWVMALIAVELAYAFIFPYSFRYIADNDDDKMVILPV
IRIS_313-7795 MLHHLKELAAVAGIHMILIYLCRFLLRRSRNVLFTVSNSLRFRLKVLTVLLYICLSVMLFYLFGSIMPLPPWGLVVGWVMALIAVELAYAFIFPYSFRYIADNDDDKMVILPV
IRIS_313-7797 MLHHLKELAAVAGIHMILIYLCRFLLRRSRNVLFTVSNSLRFRLKVLTVLLYICLSVMLFYLFGSIMPLPPWGLVVGWVMALIAVELAYAFIFPYSFRYIADNDDDKMVILPV
IRIS_313-7799 MLHHLKELAAVAGIHMILIYLCRFLLRRSRNVLFTVSNSLRFRLKVLTVLLYICLSVMLFYLFGSIMPLPPWGLVVGWVMALIAVELAYAFIFPYSFRYIADNDDDKMVILPV
IRIS_313-7807 MLHHLKELAAVAGIHMILIYLCRFLLRRSRNVLFTVSNSLRFRLKVLTVLLYICLSVMLFYLFGSIMPLPPWGLVVGWVMALIAVELAYAFIFPYSFRYIADNDDDKMVILPV
IRIS_313-7808 MLHHLKELAAVAGIHMILIYLCRFLLRRSRNVLFTVSNSLRFRLKVLTVLLYICLSVMLFYLFGSIMPLPPWGLVVGWVMALIAVELAYAFIFPYSFRYIADNDDDKMVILPV
IRIS_313-7809 MLHHLKELAAVAGIHMILIYLCRFLLRRSRNVLFTVSNSLRFRLKVLTVLLYICLSVMLFYLFGSIMPLPPWGLVVGWVMALIAVELAYAFIFPYSFRYIADNDDDKMVILPV
IRIS_313-7815 MLHHLKELAAVAGIHMILIYLCRFLLRRSRNVLFTVSNSLRFRLKVLTVLLYICLSVMLFYLFGSIMPLPPWGLVVGWVMALIAVELAYAFIFPYSFRYIADNDDDKMVILPV
IRIS_313-7816 MLHHLKELAAVAGIHMILIYLCRFLLRRSRNVLFTVSNSLRFRLKVLTVLLYICLSVMLFYLFGSIMPLPPWGLVVGWVMALIAVELAYAFIFPYSFRYIADNDDDKMVILPV
IRIS_313-7819 MLHHLKELAAVAGIHMILIYLCRFLLRRSRNVLFTVSNSLRFRLKVLTVLLYICLSVMLFYLFGSIMPLPPWGLVVGWVMALIAVELAYAFIFPYSFRYIADNDDDKMVILPV
IRIS_313-7820 MLHHLKELAAVAGIHMILIYLCRFLLRRSRNVLFTVSNSLRFRLKVLTVLLYICLSVMLFYLFGSIMPLPPWGLVVGWVMALIAVELAYAFIFPYSFRYIADNDDDKMVILPV
IRIS_313-7824 MLHHLKELAAVAGIHMILIYLCRFLLRRSRNVLFTVSNSLRFRLKVLTVLLYICLSVMLFYLFGSIMPLPPWGLVVGWVMALIAVELAYAFIFPYSFRYIADNDDDKMVILPV
IRIS_313-7826 MLHHLKELAAVAGIHMILIYLCRFLLRRSRNVLFTVSNSLRFRLKVLTVLLYICLSVMLFYLFGSIMPLPPWGLVVGWVMALIAVELAYAFIFPYSFRYIADNDDDKMVILPV

| | |
|---|---|
| IRIS_313-7832 | MLHHLKELAAVAGIHMILIYLCRFLLRRSRNVLFTVSNSLRFRLKVLTVLLYICLSVMLFYLFGSIMPLPPWGLVVGWVMALIAVELAYAFIFPYSFRYIADNDDDKMVILPV |
| IRIS_313-7838 | MLHHLKELAAVAGIHMILIYLCRFLLRRSRNVLFTVSNSLRFRLKVLTVLLYICLSVMLFYLFGSIMPLPPWGLVVGWVMALIAVELAYAFIFPYSFRYIADNDDDKMVILPV |
| IRIS_313-7850 | MLHHLKELAAVAGIHMILIYLCRFLLRRSRNVLFTVSNSLRFRLKVLTVLLYICLSVMLFYLFGSIMPLPPWGLVVGWVMALIAVELAYAFIFPYSFRYIADNDDDKMVILPV |
| IRIS_313-7856 | MLHHLKELAAVAGIHMILIYLCRFLLRRSRNVLFTVSNSLRFRLKVLTVLLYICLSVMLFYLFGSIMPLPPWGLVVGWVMALIAVELAYAFIFPYSFRYIADNDDDKMVILPV |
| IRIS_313-7859 | MLHQLKELAAVAGIHMILIYLCRFLLRRSRNVLFTVSNSLRFRLKVLTVLLYICLLVMLFYLFGSIMPLPPWGLVVGWVMALIAVELAYAFIFPYSFRYIADNDDDKMVILPV |
| IRIS_313-7863 | MLHHLKELAAVAGIHMILIYLCRFLLRRSRNVLFTVSNSLRFRLKVLTVLLYICLSVMLFYLFGSIMPLPPWGLVVGWVMALIAVELAYAFIFPYSFRYIADNDDDKMVILPV |
| IRIS_313-7866 | MLHHLKELAAVAGIHMILIYLCRFLLRRSRNVLFTVSNSLRFRLKVLTVLLYICLSVMLFYLFGSIMPLPPWGLVVGWVMALIAVELAYAFIFPYSFRYIADNDDDKMVILPV |
| IRIS_313-7868 | MLHHLKELAAVAGIHMILIYLCRFLLRRSRNVLFTVSNSLRFRLKVLTVLLYICLSVMLFYLFGSIMPLPPWGLVVGWVMALIAVELAYAFIFPYSFRYIADNDDDKMVILPV |
| IRIS_313-7870 | MLHQLKELAAVAGIHMILIYLCRFLLRRSRNVLFTVSNSLRFRLKVLTVLLYICLLVMLFYLFGSIMPLPPWGLVVGWVMALIAVELAYAFIFPYSFRYIADNDDDKMVILPV |
| IRIS_313-7876 | MLHHLKELAAVAGIHMILIYLCRFLLRRSRNVLFTVSNSLRFRLKVLTVLLYICLSVMLFYLFGSIMPLPPWGLVVGWVMALIAVELAYAFIFPYSFRYIADNDDDKMVILPV |
| IRIS_313-7883 | MLHHLKELAAVAGIHMILIYLCRFLLRRSRNVLFTVSNSLRFRLKVLTVLLYICLSVMLFYLFGSIMPLPPWGLVVGWVMALIAVELAYAFIFPYSFRYIADNDDDKMVILPV |
| IRIS_313-7885 | MLHHLKELAAVAGIHMILIYLCRFLLRRSRNVLFTVSNSLRFRLKVLTVLLYICLSVMLFYLFGSIMPLPPWGLVVGWVMALIAVELAYAFIFPYSFRYIADNDDDKMVILPV |
| IRIS_313-7902 | MLHHLKELAAVAGIHMILIYLCRFLLRRSRNVLFTVSNSLRFRLKVLTVLLYICLSVMLFYLFGSIMPLPPWGLVVGWVMALIAVELAYAFIFPYSFRYIADNDDDKMVILPV |
| IRIS_313-7907 | MLHHLKELAAVAGIHMILIYLCRFLLRRSRNVLFTVSNSLRFRLKVLTVLLYICLSVMLFYLFGSIMPLPPWGLVVGWVMALIAVELAYAFIFPYSFRYIADNDDDKMVILPV |
| IRIS_313-7909 | MLHHLKELAAVAGIHMILIYLCRFLLRRSRNVLFTVSNSLRFRLKVLTVLLYICLSVMLFYLFGSIMPLPPWGLVVGWVMALIAVELAYAFIFPYSFRYIADNDDDKMVILPV |
| IRIS_313-7911 | MLHHLKELAAVAGIHMILIYLCRFLLRRSRNVLFTVSNSLRFRLKVLTVLLYICLSVMLFYLFGSIMPLPPWGLVVGWVMALIAVELAYAFIFPYSFRYIADNDDDKMVILPV |
| IRIS_313-7912 | MLHHLKELAAVAGIHMILIYLCRFLLRRSRNVLFTVSNSLRFRLKVLTVLLYICLSVMLFYLFGSIMPLPPWGLVVGWVMALIAVELAYAFIFPYSFRYIADNDDDKMVILPV |
| IRIS_313-7914 | MLHHLKELAAVAGIHMILIYLCRFLLRRSRNVLFTVSNSLRFRLKVLTVLLYICLSVMLFYLFGSIMPLPPWGLVVGWVMALIAVELAYAFIFPYSFRYIADNDDDKMVILPV |
| IRIS_313-7922 | MLHHLKELAAVAGIHMILIYLCRFLLRRSRNVLFTVSNSLRFRLKVLTVLLYICLSVMLFYLFGSIMPLPPWGLVVGWVMALIAVELAYAFIFPYSFRYIADNDDDKMVILPV |
| IRIS_313-7924 | MLHHLKELAAVAGIHMILIYLCRFLLRRSRNVLFTVSNSLRFRLKVLTVLLYICLSVMLFYLFGSIMPLPPWGLVVGWVMALIAVELAYAFIFPYSFRYIADNDDDKMVILPV |
| IRIS_313-7933 | MLHHLKELAAVAGIHMILIYLCRFLLRRSRNVLFTVSNSLRFRLKVLTVLLYICLSVMLFYLFGSIMPLPPWGLVVGWVMALIAVELAYAFIFPYSFRYIADNDDDKMVILPV |
| IRIS_313-7959 | MLHHLKELAAVAGIHMILIYLCRFLLRRSRNVLFTVSNSLRFRLKVLTVLLYICLSVMLFYLFGSIMPLPPWGLVVGWVMALIAVELAYAFIFPYSFRYIADNDDDKMVILPV |
| IRIS_313-7992 | MLHHLKELAAVAGIHMILIYLCRFLLRRSRNVLFTVSNSLRFRLKVLTVLLYICLSVMLFYLFGSIMPLPPWGLVVGWVMALIAVELAYAFIFPYSFRYIADNDDDKMVILPV |
| IRIS_313-7993 | MLHHLKELAAVAGIHMILIYLCRFLLRRSRNVLFTVSNSLRFRLKVLTVLLYICLSVMLFYLFGSIMPLPPWGLVVGWVMALIAVELAYAFIFPYSFRYIADNDDDKMVILPV |
| IRIS_313-7994 | MLHHLKELAAVAGIHMILIYLCRFLLRRSRNVLFTVSNSLRFRLKVLTVLLYICLSVMLFYLFGSIMPLPPWGLVVGWVMALIAVELAYAFIFPYSFRYIADNDDDKMVILPV |
| IRIS_313-8003 | MLHHLKELAAVAGIHMILIYLCRFLLRRSRNVLFTVSNSLRFRLKVLTVLLYICLSVMLFYLFGSIMPLPPWGLVVGWVMALIAVELAYAFIFPYSFRYIADNDDDKMVILPV |
| IRIS_313-8010 | MLHHLKELAAVAGIHMILIYLCRFLLRRSRNVLFTVSNSLRFRLKVLTVLLYICLSVMLFYLFGSIMPLPPWGLVVGWVMALIAVELAYAFIFPYSFRYIADNDDDKMVILPV |
| IRIS_313-8011 | MLHQLKELAAVAGIHMILIYLCRFLLRRSRNVLFTVSNSLRFRLKVLTVLLYICLLVMLFYLFGSIMPLPPWGLVVGWVMALIAVELAYAFIFPYSFRYIADNDDDKMVILPV |
| IRIS_313-8023 | MLHHLKELAAVAGIHMILIYLCRFLLRRSRNVLFTVSNSLRFRLKVLTVLLYICLSVMLFYLFGSIMPLPPWGLVVGWVMALIAVELAYAFIFPYSFRYIADNDDDKMVILPV |
| IRIS_313-8024 | MLHHLKELAAVAGIHMILIYLCRFLLRRSRNVLFTVSNSLRFRLKVLTVLLYICLSVMLFYLFGSIMPLPPWGLVVGWVMALIAVELAYAFIFPYSFRYIADNDDDKMVILPV |
| IRIS_313-8025 | MLHHLKELAAVAGIHMILIYLCRFLLRRSRNVLFTVSNSLRFRLKVLTVLLYICLSVMLFYLFGSIMPLPPWGLVVGWVMALIAVELAYAFIFPYSFRYIADNDDDKMVILPV |
| IRIS_313-8026 | MLHHLKELAAVAGIHMILIYLCRFLLRRSRNVLFTVSNSLRFRLKVLTVLLYICLSVMLFYLFGSIMPLPPWGLVVGWVMALIAVELAYAFIFPYSFRYIADNDDDKMVILPV |
| IRIS_313-8027 | MLHHLKELAAVAGIHMILIYLCRFLLRRSRNVLFTVSNSLRFRLKVLTVLLYICLSVMLFYLFGSIMPLPPWGLVVGWVMALIAVELAYAFIFPYSFRYIADNDDDKMVILPV |
| IRIS_313-8029 | MLHHLKELAAVAGIHMILIYLCRFLLRRSRNVLFTVSNSLRFRLKVLTVLLYICLSVMLFYLFGSIMPLPPWGLVVGWVMALIAVELAYAFIFPYSFRYIADNDDDKMVILPV |
| IRIS_313-8031 | MLHHLKELAAVAGIHMILIYLCRFLLRRSRNVLFTVSNSLRFRLKVLTVLLYICLSVMLFYLFGSIMPLPPWGLVVGWVMALIAVELAYAFIFPYSFRYIADNDDDKMVILPV |
| IRIS_313-8032 | MLHHLKELAAVAGIHMILIYLCRFLLRRSRNVLFTVSNSLRFRLKVLTVLLYICLSVMLFYLFGSIMPLPPWGLVVGWVMALIAVELAYAFIFPYSFRYIADNDDDKMVILPV |
| IRIS_313-8033 | MLHHLKELAAVAGIHMILIYLCRFLLRRSRNVLFTVSNSLRFRLKVLTVLLYICLSVMLFYLFGSIMPLPPWGLVVGWVMALIAVELAYAFIFPYSFRYIADNDDDKMVILPV |
| IRIS_313-8037 | MLHHLKELAAVAGIHMILIYLCRFLLRRSRNVLFTVSNSLRFRLKVLTVLLYICLSVMLFYLFGSIMPLPPWGLVVGWVMALIAVELAYAFIFPYSFRYIADNDDDKMVILPV |
| IRIS_313-8039 | MLHHLKELAAVAGIHMILIYLCRFLLRRSRNVLFTVSNSLRFRLKVLTVLLYICLSVMLFYLFGSIMPLPPWGLVVGWVMALIAVELAYAFIFPYSFRYIADNDDDKMVILPV |
| IRIS_313-8041 | MLHHLKELAAVAGIHMILIYLCRFLLRRSRNVLFTVSNSLRFRLKVLTVLLYICLSVMLFYLFGSIMPLPPWGLVVGWVMALIAVELAYAFIFPYSFRYIADNDDDKMVILPV |
| IRIS_313-8044 | MLHHLKELAAVAGIHMILIYLCRFLLRRSRNVLFTVSNSLRFRLKVLTVLLYICLSVMLFYLFGSIMPLPPWGLVVGWVMALIAVELAYAFIFPYSFRYIADNDDDKMVILPV |
| IRIS_313-8046 | MLHHLKELAAVAGIHMILIYLCRFLLRRSRNVLFTVSNSLRFRLKVLTVLLYICLSVMLFYLFGSIMPLPPWGLVVGWVMALIAVELAYAFIFPYSFRYIADNDDDKMVILPV |
| IRIS_313-8048 | MLHHLKELAAVAGIHMILIYLCRFLLRRSRNVLFTVSNSLRFRLKVLTVLLYICLSVMLFYLFGSIMPLPPWGLVVGWVMALIAVELAYAFIFPYSFRYIADNDDDKMVILPV |
| IRIS_313-8049 | MLHHLKELAAVAGIHMILIYLCRFLLRRSRNVLFTVSNSLRFRLKVLTVLLYICLSVMLFYLFGSIMPLPPWGLVVGWVMALIAVELAYAFIFPYSFRYIADNDDDKMVILPV |
| IRIS_313-8050 | MLHHLKELAAVAGIHMILIYLCRFLLRRSRNVLFTVSNSLRFRLKVLTVLLYICLSVMLFYLFGSIMPLPPWGLVVGWVMALIAVELAYAFIFPYSFRYIADNDDDKMVILPV |
| IRIS_313-8052 | MLHHLKELAAVAGIHMILIYLCRFLLRRSRNVLFTVSNSLRFRLKVLTVLLYICLSVMLFYLFGSIMPLPPWGLVVGWVMALIAVELAYAFIFPYSFRYIADNDDDKMVILPV |
| IRIS_313-8053 | MLHHLKELAAVAGIHMILIYLCRFLLRRSRNVLFTVSNSLRFRLKVLTVLLYICLSVMLFYLFGSIMPLPPWGLVVGWVMALIAVELAYAFIFPYSFRYIADNDDDKMVILPV |
| IRIS_313-8057 | MLHHLKELAAVAGIHMILIYLCRFLLRRSRNVLFTVSNSLRFRLKVLTVLLYICLSVMLFYLFGSIMPLPPWGLVVGWVMALIAVELAYAFIFPYSFRYIADNDDDKMVILPV |
| IRIS_313-8058 | MLHHLKELAAVAGIHMILIYLCRFLLRRSRNVLFTVSNSLRFRLKVLTVLLYICLSVMLFYLFGSIMPLPPWGLVVGWVMALIAVELAYAFIFPYSFRYIADNDDDKMVILPV |
| IRIS_313-8060 | MLHHLKELAAVAGIHMILIYLCRFLLRRSRNVLFTVSNSLRFRLKVLTVLLYICLSVMLFYLFGSIMPLPPWGLVVGWVMALIAVELAYAFIFPYSFRYIADNDDDKMVILPV |
| IRIS_313-8061 | MLHHLKELAAVAGIHMILIYLCRFLLRRSRNVLFTVSNSLRFRLKVLTVLLYICLSVMLFYLFGSIMPLPPWGLVVGWVMALIAVELAYAFIFPYSFRYIADNDDDKMVILPV |
| IRIS_313-8062 | MLHHLKELAAVAGIHMILIYLCRFLLRRSRNVLFTVSNSLRFRLKVLTVLLYICLSVMLFYLFGSIMPLPPWGLVVGWVMALIAVELAYAFIFPYSFRYIADNDDDKMVILPV |
| IRIS_313-8063 | MLHHLKELAAVAGIHMILIYLCRFLLRRSRNVLFTVSNSLRFRLKVLTVLLYICLSVMLFYLFGSIMPLPPWGLVVGWVMALIAVELAYAFIFPYSFRYIADNDDDKMVILPV |
| IRIS_313-8064 | MLHHLKELAAVAGIHMILIYLCRFLLRRSRNVLFTVSNSLRFRLKVLTVLLYICLSVMLFYLFGSIMPLPPWGLVVGWVMALIAVELAYAFIFPYSFRYIADNDDDKMVILPV |
| IRIS_313-8065 | MLHHLKELAAVAGIHMILIYLCRFLLRRSRNVLFTVSNSLRFRLKVLTVLLYICLSVMLFYLFGSIMPLPPWGLVVGWVMALIAVELAYAFIFPYSFRYIADNDDDKMVILPV |
| IRIS_313-8066 | MLHHLKELAAVAGIHMILIYLCRFLLRRSRNVLFTVSNSLRFRLKVLTVLLYICLSVMLFYLFGSIMPLPPWGLVVGWVMALIAVELAYAFIFPYSFRYIADNDDDKMVILPV |
| IRIS_313-8068 | MLHHLKELAAVAGIHMILIYLCRFLLRRSRNVLFTVSNSLRFRLKVLTVLLYICLSVMLFYLFGSIMPLPPWGLVVGWVMALIAVELAYAFIFPYSFRYIADNDDDKMVILPV |
| IRIS_313-8069 | MLHHLKELAAVAGIHMILIYLCRFLLRRSRNVLFTVSNSLRFRLKVLTVLLYICLSVMLFYLFGSIMPLPPWGLVVGWVMALIAVELAYAFIFPYSFRYIADNDDDKMVILPV |
| IRIS_313-8072 | MLHHLKELAAVAGIHMILIYLCRFLLRRSRNVLFTVSNSLRFRLKVLTVLLYICLSVMLFYLFGSIMPLPPWGLVVGWVMALIAVELAYAFIFPYSFRYIADNDDDKMVILPV |
| IRIS_313-8073 | MLHHLKELAAVAGIHMILIYLCRFLLRRSRNVLFTVSNSLRFRLKVLTVLLYISLSVMLLYLFGSIMPLPPWGLVVGWVMALIAVELAYAFIFPYSFRYIADNDDDKMVILPV |
| IRIS_313-8074 | MLHHLKELAAVAGIHMILIYLCRFLLRRSRNVLFTVSNSLRFRLKVLTVLLYICLSVMLFYLFGSIMPLPPWGLVVGWVMALIAVELAYAFIFPYSFRYIADNDDDKMVILPV |
| IRIS_313-8075 | MLHHLKELAAVAGIHMILIYLCRFLLRRSRNVLFTVSNSLRFRLKVLTVLLYICLSVMLFYLFGSIMPLPPWGLVVGWVMALIAVELAYAFIFPYSFRYIADNDDDKMVILPV |
| IRIS_313-8076 | MLHHLKELAAVAGIHMILIYLCRFLLRRSRNVLFTVSNSLRFRLKVLTVLLYICLSVMLFYLFGSIMPLPPWGLVVGWVMALIAVELAYAFIFPYSFRYIADNDDDKMVILPV |
| IRIS_313-8084 | MLHHLKELAAVAGIHMILIYLCRFLLRRSRNVLFTVSNSLRFRLKVLTVLLYICLSVMLFYLFGSIMPLPPWGLVVGWVMALIAVELAYAFIFPYSFRYIADNDDDKMVILPV |
| IRIS_313-8085 | MLHHLKELAAVAGIHMILIYLCRFLLRRSRNVLFTVSNSLRFRLKVLTVLLYICLSVMLFYLFGSIMPLPPWGLVVGWVMALIAVELAYAFIFPYSFRYIADNDDDKMVILPV |
| IRIS_313-8087 | MLHHLKELAAVAGIHMILIYLCRFLLRRSRNVLFTVSNSLRFRLKVLTVLLYICLSVMLFYLFGSIMPLPPWGLVVGWVMALIAVELAYAFIFPYSFRYIADNDDDKMVILPV |
| IRIS_313-8090 | MLHHLKELAAVAGIHMILIYLCRFLLRRSRNVLFTVSNSLRFRLKVLTVLLYICLSVMLFYLFGSIMPLPPWGLVVGWVMALIAVELAYAFIFPYSFRYIADNDDDKMVILPV |
| IRIS_313-8095 | MLHHLKELAAVAGIHMILIYLCRFLLRRSRNVLFTVSNSLRFRLKVLTVLLYICLSVMLFYLFGSIMPLPPWGLVVGWVMALIAVELAYAFIFPYSFRYIADNDDDKMVILPV |
| IRIS_313-8096 | MLHHLKELAAVAGIHMILIYLCRFLLRRSRNVLFTVSNSLRFRLKVLTVLLYICLSVMLFYLFGSIMPLPPWGLVVGWVMALIAVELAYAFIFPYSFRYIADNDDDKMVILPV |
| IRIS_313-8097 | MLHHLKELAAVAGIHMILIYLCRFLLRRSRNVLFTVSNSLRFRLKVLTVLLYICLSVMLFYLFGSIMPLPPWGLVVGWVMALIAVELAYAFIFPYSFRYIADNDDDKMVILPV |
| IRIS_313-8099 | MLHHLKELAAVAGIHMILIYLCRFLLRRSRNVLFTVSNSLRFRLKVLTVLLYICLSVMLFYLFGSIMPLPPWGLVVGWVMALIAVELAYAFIFPYSFRYIADNDDDKMVILPV |
| IRIS_313-8102 | MLHHLKELAAVAGIHMILIYLCRFLLRRSRNVLFTVSNSLRFRLKVLTVLLYICLSVMLFYLFGSIMPLPPWGLVVGWVMALIAVELAYAFIFPYSFRYIADNDDDKMVILPV |
| IRIS_313-8105 | MLHHLKELAAVAGIHMILIYLCRFLLRRSRNVLFTVSNSLRFRLKVLTVLLYICLSVMLFYLFGSIMPLPPWGLVVGWVMALIAVELAYAFIFPYSFRYIADNDDDKMVILPV |
| IRIS_313-8109 | MLHHLKELAAVAGIHMILIYLCRFLLRRSRNVLFTVSNSLRFRLKVLTVLLYICLSVMLFYLFGSIMPLPPWGLVVGWVMALIAVELAYAFIFPYSFRYIADNDDDKMVILPV |
| IRIS_313-8111 | MLHHLKELAAVAGIHMILIYLCRFLLRRSRNVLFTVSNSLRFRLKVLTVLLYICLSVMLFYLFGSIMPLPPWGLVVGWVMALIAVELAYAFIFPYSFRYIADNDDDKMVILPV |
| IRIS_313-8112 | MLHHLKELAAVAGIHMILIYLCRFLLRRSRNVLFTVSNSLRFRLKVLTVLLYICLSVMLFYLFGSIMPLPPWGLVVGWVMALIAVELAYAFIFPYSFRYIADNDDDKMVILPV |
| IRIS_313-8113 | MLHHLKELAAVAGIHMILIYLCRFLLRRSRNVLFTVSNSLRFRLKVLTVLLYICLSVMLFYLFGSIMPLPPWGLVVGWVMALIAVELAYAFIFPYSFRYIADNDDDKMVILPV |
| IRIS_313-8114 | MLHHLKELAAVAGIHMILIYLCRFLLRRSRNVLFTVSNSLRFRLKVLTVLLYICLSVMLFYLFGSIMPLPPWGLVVGWVMALIAVELAYAFIFPYSFRYIADNDDDKMVILPV |
| IRIS_313-8115 | MLHHLKELAAVAGIHMILIYLCRFLLRRSRNVLFTVSNSLRFRLKVLTVLLYICLSVMLFYLFGSIMPLPPWGLVVGWVMALIAVELAYAFIFPYSFRYIADNDDDKMVILPV |
| IRIS_313-8116 | MLHHLKELAAVAGIHMILIYLCRFLLRRSRNVLFTVSNSLRFRLKVLTVLLYICLSVMLFYLFGSIMPLPPWGLVVGWVMALIAVELAYAFIFPYSFRYIADNDDDKMVILPV |
| IRIS_313-8118 | MLHHLKELAAVAGIHMILIYLCRFLLRRSRNVLFTVSNSLRFRLKVLTVLLYICLSVMLFYLFGSIMPLPPWGLVVGWVMALIAVELAYAFIFPYSFRYIADNDDDKMVILPV |
| IRIS_313-8119 | MLHHLKELAAVAGIHMILIYLCRFLLRRSRNVLFTVSNSLRFRLKVLTVLLYICLSVMLFYLFGSIMPLPPWGLVVGWVMALIAVELAYAFIFPYSFRYIADNDDDKMVILPV |
| IRIS_313-8121 | MLHHLKELAAVAGIHMILIYLCRFLLRRSRNVLFTVSNSLRFRLKVLTVLLYICLSVMLFYLFGSIMPLPPWGLVVGWVMALIAVELAYAFIFPYSFRYIADNDDDKMVILPV |
| IRIS_313-8123 | MLHHLKELAAVAGIHMILIYLCRFLLRRSRNVLFTVSNSLRFRLKVLTVLLYICLSVMLFYLFGSIMPLPPWGLVVGWVMALIAVELAYAFIFPYSFRYIADNDDDKMVILPV |
| IRIS_313-8124 | MLHHLKELAAVAGIHMILIYLCRFLLRRSRNVLFTVSNSLRFRLKVLTVLLYICLSVMLFYLFGSIMPLPPWGLVVGWVMALIAVELAYAFIFPYSFRYIADNDDDKMVILPV |
| IRIS_313-8125 | MLHHLKELAAVAGIHMILIYLCRFLLRRSRNVLFTVSNSLRFRLKVLTVLLYICLSVMLFYLFGSIMPLPPWGLVVGWVMALIAVELAYAFIFPYSFRYIADNDDDKMVILPV |
| IRIS_313-8126 | MLHHLKELAAVAGIHMILIYLCRFLLRRSRNVLFTVSNSLRFRLKVLTVLLYICLSVMLFYLFGSIMPLPPWGLVVGWVMALIAVELAYAFIFPYSFRYIADNDDDKMVILPV |
| IRIS_313-8127 | MLHHLKELAAVAGIHMILIYLCRFLLRRSRNVLFTVSNSLRFRLKVLTVLLYICLSVMLFYLFGSIMPLPPWGLVVGWVMALIAVELAYAFIFPYSFRYIADNDDDKMVILPV |
| IRIS_313-8128 | MLHHLKELAAVAGIHMILIYLCRFLLRRSRNVLFTVSNSLRFRLKVLTVLLYICLSVMLFYLFGSIMPLPPWGLVVGWVMALIAVELAYAFIFPYSFRYIADNDDDKMVILPV |
| IRIS_313-8129 | MLHHLKELAAVAGIHMILIYLCRFLLRRSRNVLFTVSNSLRFRLKVLTVLLYICLSVMLFYLFGSIMPLPPWGLVVGWVMALIAVELAYAFIFPYSFRYIADNDDDKMVILPV |
| IRIS_313-8132 | MLHHLKELAAVAGIHMILIYLCRFLLRRSRNVLFTVSNSLRFRLKVLTVLLYICLSVMLFYLFGSIMPLPPWGLVVGWVMALIAVELAYAFIFPYSFRYIADNDDDKMVILPV |
| IRIS_313-8134 | MLHHLKELAAVAGIHMILIYLCRFLLRRSRNVLFTVSNSLRFRLKVLTVLLYICLSVMLFYLFGSIMPLPPWGLVVGWVMALIAVELAYAFIFPYSFRYIADNDDDKMVILPV |
| IRIS_313-8135 | MLHHLKELAAVAGIHMILIYLCRFLLRRSRNVLFTVSNSLRFRLKVLTVLLYICLSVMLFYLFGSIMPLPPWGLVVGWVMALIAVELAYAFIFPYSFRYIADNDDDKMVILPV |
| IRIS_313-8136 | MLHHLKELAAVAGIHMILIYLCRFLLRRSRNVLFTVSNSLRFRLKVLTVLLYICLSVMLFYLFGSIMPLPPWGLVVGWVMALIAVELAYAFIFPYSFRYIADNDDDKMVILPV |
| IRIS_313-8137 | MLHHLKELAAVAGIHMILIYLCRFLLRRSRNVLFTVSNSLRFRLKVLTVLLYICLSVMLFYLFGSIMPLPPWGLVVGWVMALIAVELAYAFIFPYSFRYIADNDDDKMVILPV |
| IRIS_313-8138 | MLHHLKELAAVAGIHMILIYLCRFLLRRSRNVLFTVSNSLRFRLKVLTVLLYICLSVMLFYLFGSIMPLPPWGLVVGWVMALIAVELAYAFIFPYSFRYIADNDDDKMVILPV |
| IRIS_313-8139 | MLHHLKELAAVAGIHMILIYLCRFLLRRSRNVLFTVSNSLRFRLKVLTVLLYICLSVMLFYLFGSIMPLPPWGLVVGWVMALIAVELAYAFIFPYSFRYIADNDDDKMVILPV |
| IRIS_313-8140 | MLHHLKELAAVAGIHMILIYLCRFLLRRSRNVLFTVSNSLRFRLKVLTVLLYICLSVMLFYLFGSIMPLPPWGLVVGWVMALIAVELAYAFIFPYSFRYIADNDDDKMVILPV |
| IRIS_313-8141 | MLHHLKELAAVAGIHMILIYLCRFLLRRSRNVLFTVSNSLRFRLKVLTVLLYICLSVMLFYLFGSIMPLPPWGLVVGWVMALIAVELAYAFIFPYSFRYIADNDDDKMVILPV |
| IRIS_313-8142 | MLHHLKELAAVAGIHMILIYLCRFLLRRSRNVLFTVSNSLRFRLKVLTVLLYICLSVMLFYLFGSIMPLPPWGLVVGWVMALIAVELAYAFIFPYSFRYIADNDDDKMVILPV |
| IRIS_313-8143 | MLHHLKELAAVAGIHMILIYLCRFLLRRSRNVLFTVSNSLRFRLKVLTVLLYICLSVMLFYLFGSIMPLPPWGLVVGWVMALIAVELAYAFIFPYSFRYIADNDDDKMVILPV |
| IRIS_313-8145 | MLHHLKELAAVAGIHMILIYLCRFLLRRSRNVLFTVSNSLRFRLKVLTVLLYICLSVMLFYLFGSIMPLPPWGLVVGWVMALIAVELAYAFIFPYSFRYIADNDDDKMVILPV |
| IRIS_313-8147 | MLHHLKELAAVAGIHMILIYLCRFLLRRSRNVLFTVSNSLRFRLKVLTVLLYICLSVMLFYLFGSIMPLPPWGLVVGWVMALIAVELAYAFIFPYSFRYIADNDDDKMVILPV |
| IRIS_313-8148 | MLHHLKELAAVAGIHMILIYLCRFLLRRSRNVLFTVSNSLRFRLKVLTVLLYICLSVMLFYLFGSIMPLPPWGLVVGWVMALIAVELAYAFIFPYSFRYIADNDDDKMVILPV |
| IRIS_313-8149 | MLHHLKELAAVAGIHMILIYLCRFLLRRSRNVLFTVSNSLRFRLKVLTVLLYICLSVMLFYLFGSIMPLPPWGLVVGWVMALIAVELAYAFIFPYSFRYIADNDDDKMVILPV |
| IRIS_313-8151 | MLHHLKELAAVAGIHMILIYLCRFLLRRSRNVLFTVSNSLRFRLKVLTVLLYICLSVMLFYLFGSIMPLPPWGLVVGWVMALIAVELAYAFIFPYSFRYIADNDDDKMVILPV |
| IRIS_313-8154 | MLHHLKELAAVAGIHMILIYLCRFLLRRSRNVLFTVSNSLRFRLKVLTVLLYICLSVMLFYLFGSIMPLPPWGLVVGWVMALIAVELAYAFIFPYSFRYIADNDDDKMVILPV |
| IRIS_313-8155 | MLHHLKELAAVAGIHMILIYLCRFLLRRSRNVLFTVSNSLRFRLKVLTVLLYICLSVMLFYLFGSIMPLPPWGLVVGWVMALIAVELAYAFIFPYSFRYIADNDDDKMVILPV |
| IRIS_313-8158 | MLHHLKELAAVAGIHMILIYLCRFLLRRSRNVLFTVSNSLRFRLKVLTVLLYICLSVMLFYLFGSIMPLPPWGLVVGWVMALIAVELAYAFIFPYSFRYIADNDDDKMVILPV |
| IRIS_313-8159 | MLHHLKELAAVAGIHMILIYLCRFLLRRSRNVLFTVSNSLRFRLKVLTVLLYICLSVMLFYLFGSIMPLPPWGLVVGWVMALIAVELAYAFIFPYSFRYIADNDDDKMVILPV |
| IRIS_313-8160 | MLHHLKELAAVAGIHMILIYLCRFLLRRSRNVLFTVSNSLRFRLKVLTVLLYICLSVMLFYLFGSIMPLPPWGLVVGWVMALIAVELAYAFIFPYSFRYIADNDDDKMVILPV |
| IRIS_313-8161 | MLHHLKELAAVAGIHMILIYLCRFLLRRSRNVLFTVSNSLRFRLKVLTVLLYICLSVMLFYLFGSIMPLPPWGLVVGWVMALIAVELAYAFIFPYSFRYIADNDDDKMVILPV |
| IRIS_313-8162 | MLHHLKELAAVAGIHMILIYLCRFLLRRSRNVLFTVSNSLRFRLKVLTVLLYICLSVMLFYLFGSIMPLPPWGLVVGWVMALIAVELAYAFIFPYSFRYIADNDDDKMVILPV |
| IRIS_313-8164 | MLHHLKELAAVAGIHMILIYLCRFLLRRSRNVLFTVSNSLRFRLKVLTVLLYICLSVMLFYLFGSIMPLPPWGLVVGWVMALIAVELAYAFIFPYSFRYIADNDDDKMVILPV |
| IRIS_313-8165 | MLHHLKELAAVAGIHMILIYLCRFLLRRSRNVLFTVSNSLRFRLKVLTVLLYICLSVMLFYLFGSIMPLPPWGLVVGWVMALIAVELAYAFIFPYSFRYIADNDDDKMVILPV |
| IRIS_313-8166 | MLHHLKELAAVAGIHMILIYLCRFLLRRSRNVLFTVSNSLRFRLKVLTVLLYICLSVMLFYLFGSIMPLPPWGLVVGWVMALIAVELAYAFIFPYSFRYIADNDDDKMVILPV |
| IRIS_313-8167 | MLHHLKELAAVAGIHMILIYLCRFLPRRSRNVLFTVSNSLRFRLKVLTVLLYICLSVMLFYLFGSIMPLPPWGLVVGWVMALIAVELAYAFIFPYSFRYIADNDDDKMVILPV |
| IRIS_313-8168 | MLHHLKELAAVAGIHMILIYLCRFLLRRSRNVLFTVSNSLRFRLKVLTVLLYICLSVMLFYLFGSIMPLPPWGLVVGWVMALIAVELAYAFIFPYSFRYIADNDDDKMVILPV |
| IRIS_313-8170 | MLHHLKELAAVAGIHMILIYLCRFLLRRSRNVLFTVSNSLRFRLKVLTVLLYICLSVMLFYLFGSIMPLPPWGLVVGWVMALIAVELAYAFIFPYSFRYIADNDDDKMVILPV |
| IRIS_313-8171 | MLHHLKELAAVAGIHMILIYLCRFLLRRSRNVLFTVSNSLRFRLKVLTVLLYICLSVMLFYLFGSIMPLPPWGLVVGWVMALIAVELAYAFIFPYSFRYIADNDDDKMVILPV |
| IRIS_313-8172 | MLHHLKELAAVAGIHMILIYLCRFLLRRSRNVLFTVSNSLRFRLKVLTVLLYICLSVMLFYLFGSIMPLPPWGLVVGWVMALIAVELAYAFIFPYSFRYIADNDDDKMVILPV |
| IRIS_313-8173 | MLHHLKELAAVAGIHMILIYLCRFLLRRSRNVLFTVSNSLRFRLKVLTVLLYICLSVMLFYLFGSIMPLPPWGLVVGWVMALIAVELAYAFIFPYSFRYIADNDDDKMVILPV |
| IRIS_313-8177 | MLHHLKELAAVAGIHMILIYLCRFLLRRSRNVLFTVSNSLRFRLKVLTVLLYICLSVMLFYLFGSIMPLPPWGLVVGWVMALIAVELAYAFIFPYSFRYIADNDDDKMVILPV |

| | |
|---|---|
| IRIS_313-8180 | MLHHLKELAAVAGIHMILIYLCRFLLRRSRNVLFTVSNSLRFRLKVLTVLLYICLSVMLFYLFGSIMPLPPWGLVVGWVMALIAVELAYAFIFPYSFRYIADNDDDKMVILPV |
| IRIS_313-8183 | MLHHLKELAAVAGIHMILIYLCRFLLRRSRNVLFTVSNSLRFRLKVLTVLLYICLSVMLFYLFGSIMPLPPWGLVVGWVMALIAVELAYAFIFPYSFRYIADNDDDKMVILPV |
| IRIS_313-8184 | MLHHLKELAAVAGIHMILIYLCRFLLRRSRNVLFTVSNSLRFRLKVLTVLLYICLSVMLFYLFGSIMPLPPWGLVVGWVMALIAVELAYAFIFPYSFRYIADNDDDKMVILPV |
| IRIS_313-8185 | MLHHLKELAAVAGIHMILIYLCRFLLRRSRNVLFTVSNSLRFRLKVLTVLLYICLSVMLFYLFGSIMPLPPWGLVVGWVMALIAVELAYAFIFPYSFRYIADNDDDKMVILPV |
| IRIS_313-8186 | MLHHLKELAAVAGIHMILIYLCRFLLRRSRNVLFTVSNSLRFRLKVLTVLLYICLSVMLFYLFGSIMPLPPWGLVVGWVMALIAVELAYAFIFPYSFRYIADNDDDKMVILPV |
| IRIS_313-8192 | MLHHLKELAAVAGIHMILIYLCRFLLRRSRNVLFTVSNSLRFRLKVLTVLLYICLSVMLFYLFGSIMPLPPWGLVVGWVMALIAVELAYAFIFPYSFRYIADNDDDKMVILPV |
| IRIS_313-8193 | MLHHLKELAAVAGIHMILIYLCRFLLRRSRNVLFTVSNSLRFRLKVLTVLLYICLSVMLFYLFGSIMPLPPWGLVVGWVMALIAVELAYAFIFPYSFRYIADNDDDKMVILPV |
| IRIS_313-8195 | MLHHLKELAAVAGIHMILIYLCRFLLRRSRNVLFTVSNSLRFRLKVLTVLLYICLSVMLFYLFGSIMPLPPWGLVVGWVMALIAVELAYAFIFPYSFRYIADNDDDKMVILPV |
| IRIS_313-8200 | MLHHLKELAAVAGIHMILIYLCRFLLRRSRNVLFTVSNSLRFRLKVLTVLLYICLSVMLFYLFGSIMPLPPWGLVVGWVMALIAVELAYAFIFPYSFRYIADNDDDKMVILPV |
| IRIS_313-8202 | MLHHLKELAAVAGIHMILIYLCRFLLRRSRNVLFTVSNSLRFRLKVLTVLLYICLSVMLFYLFGSIMPLPPWGLVVGWVMALIAVELAYAFIFPYSFRYIADNDDDKMVILPV |
| IRIS_313-8204 | MLHHLKELAAVAGIHMILIYLCRFLLRRSRNVLFTVSNSLRFRLKVLTVLLYICLSVMLFYLFGSIMPLPPWGLVVGWVMALIAVELAYAFIFPYSFRYIADNDDDKMVILPV |
| IRIS_313-8205 | MLHHLKELAAVAGIHMILIYLCRFLLRRSRNVLFTVSNSLRFRLKVLTVLLYICLSVMLFYLFGSIMPLPPWGLVVGWVMALIAVELAYAFIFPYSFRYIADNDDDKMVILPV |
| IRIS_313-8208 | MLHHLKELAAVAGIHMILIYLCRFLLRRSRNVLFTVSNSLRFRLKVLTVLLYICLSVMLFYLFGSIMPLPPWGLVVGWVMALIAVELAYAFIFPYSFRYIADNDDDKMVILPV |
| IRIS_313-8209 | MLHHLKELAAVAGIHMILIYLCRFLLRRSRNVLFTVSNSLRFRLKVLTVLLYICLSVMLFYLFGSIMPLPPWGLVVGWVMALIAVELAYAFIFPYSFRYIADNDDDKMVILPV |
| IRIS_313-8212 | MLHHLKELAAVAGIHMILIYLCRFLLRRSRNVLFTVSNSLRFRLKVLTVLLYICLSVMLFYLFGSIMPLPPWGLVVGWVMALIAVELAYAFIFPYSFRYIADNDDDKMVILPV |
| IRIS_313-8213 | MLHHLKELAAVAGIHMILIYLCRFLLRRSRNVLFTVSNSLRFRLKVLTVLLYICLSVMLFYLFGSIMPLPPWGLVVGWVMALIAVELAYAFIFPYSFRYIADNDDDKMVILPV |
| IRIS_313-8214 | MLHHLKELAAVAGIHMILIYLCRFLLRRSRNVLFTVSNSLRFRLKVLTVLLYICLSVMLFYLFGSIMPLPPWGLVVGWVMALIAVELAYAFIFPYSFRYIADNDDDKMVILPV |
| IRIS_313-8215 | MLHHLKELAAVAGIHMILIYLCRFLLRRSRNVLFTVSNSLRFRLKVLTVLLYICLSVMLFYLFGSIMPLPPWGLVVGWVMALIAVELAYAFIFPYSFRYIADNDDDKMVILPV |
| IRIS_313-8216 | MLHHLKELAAVAGIHMILIYLCRFLLRRSRNVLFTVSNSLRFRLKVLTVLLYICLSVMLFYLFGSIMPLPPWGLVVGWVMALIAVELAYAFIFPYSFRYIADNDDDKMVILPV |
| IRIS_313-8217 | MLHHLKELAAVAGIHMILIYLCRFLLRRSRNVLFTVSNSLRFRLKVLTVLLYICLSVMLFYLFGSIMPLPPWGLVVGWVMALIAVELAYAFIFPYSFRYIADNDDDKMVILPV |
| IRIS_313-8218 | MLHHLKELAAVAGIHMILIYLCRFLLRRSRNVLFTVSNSLRFRLKVLTVLLYICLSVMLFYLFGSIMPLPPWGLVVGWVMALIAVELAYAFIFPYSFRYIADNDDDKMVILPV |
| IRIS_313-8232 | MLHHLKELAAVAGIHMILIYLCRFLLRRSRNVLFTVSNSLRFRLKVLTVLLYICLSVMLFYLFGSIMPLPPWGLVVGWVMALIAVELAYAFIFPYSFRYIADNDDDKMVILPV |
| IRIS_313-8244 | MLHHLKELAAVAGIHMILIYLCRFLLRRSRNVLFTVSNSLRFRLKVLTVLLYICLSVMLFYLFGSIMPLPPWGLVVGWVMALIAVELAYAFIFPYSFRYIADNDDDKMVILPV |
| IRIS_313-8252 | MLHHLKELAAVAGIHMILIYLCRFLLRRSRNVLFTVSNSLRFRLKVLTVLLYICLSVMLFYLFGSIMPLPPWGLVVGWVMALIAVELAYAFIFPYSFRYIADNDDDKMVILPV |
| IRIS_313-8253 | MLHHLKELAAVAGIHMILIYLCRFLLRRSRNVLFTVSNSLRFRLKVLTVLLYICLSVMLFYLFGSIMPLPPWGLVVGWVMALIAVELAYAFIFPYSFRYIADNDDDKMVILPV |
| IRIS_313-8256 | MLHHLKELAAVAGIHMILIYLCRFLLRRSRNVLFTVSNSLRFRLKVLTVLLYICLSVMLFYLFGSIMPLPPWGLVVGWVMALIAVELAYAFIFPYSFRYIADNDDDKMVILPV |
| IRIS_313-8265 | MLHHLKELAAVAGIHMILIYLCRFLLRRSRNVLFTVSNSLRFRLKVLTVLLYICLSVMLFYLFGSIMPLPPWGLVVGWVMALIAVELAYAFIFPYSFRYIADNDDDKMVILPV |
| IRIS_313-8268 | MLHHLKELAAVAGIHMILIYLCRFLLRRSRNVLFTVSNSLRFRLKVLTVLLYICLSVMLFYLFGSIMPLPPWGLVVGWVMALIAVELAYAFIFPYSFRYIADNDDDKMVILPV |
| IRIS_313-8277 | MLHHLKELAAVAGIHMILIYLCRFLLRRSRNVLFTVSNSLRFRLKVLTVLLYICLSVMLFYLFGSIMPLPPWGLVVGWVMALIAVELAYAFIFPYSFRYIADNDDDKMVILPV |
| IRIS_313-8279 | MLHHLKELAAVAGIHMILIYLCRFLLRRSRNVLFTVSNSLRFRLKVLTVLLYICLSVMLFYLFGSIMPLPPWGLVVGWVMALIAVELAYAFIFPYSFRYIADNDDDKMVILPV |
| IRIS_313-8283 | MLHHLKELAAVAGIHMILIYLCRFLLRRSRNVLFTVSNSLRFRLKVLTVLLYICLSVMLFYLFGSIMPLPPWGLVVGWVMALIAVELAYAFIFPYSFRYIADNDDDKMVILPV |
| IRIS_313-8285 | MLHHLKELAAVAGIHMILIYLCRFLLRRSRNVLFTVSNSLRFRLKVLTVLLYICLSVMLFYLFGSIMPLPPWGLVVGWVMALIAVELAYAFIFPYSFRYIADNDDDKMVILPV |
| IRIS_313-8288 | MLHHLKELAAVAGIHMILIYLCRFLLRRSRNVLFTVSNSLRFRLKVLTVLLYICLSVMLFYLFGSIMPLPPWGLVVGWVMALIAVELAYAFIFPYSFRYIADNDDDKMVILPV |
| IRIS_313-8291 | MLHHLKELAAVAGIHMILIYLCRFLLRRSRNVLFTVSNSLRFRLKVLTVLLYICLSVMLFYLFGSIMPLPPWGLVVGWVMALIAVELAYAFIFPYSFRYIADNDDDKMVILPV |
| IRIS_313-8292 | MLHHLKELAAVAGIHMILIYLCRFLLRRSRNVLFTVSNSLRFRLKVLTVLLYICLSVMLFYLFGSIMPLPPWGLVVGWVMALIAVELAYAFIFPYSFRYIADNDDDKMVILPV |
| IRIS_313-8293 | MLHHLKELAAVAGIHMILIYLCRFLLRRSRNVLFTVSNSLRFRLKVLTVLLYICLSVMLFYLFGSIMPLPPWGLVVGWVMALIAVELAYAFIFPYSFRYIADNDDDKMVILPV |
| IRIS_313-8302 | MLHHLKELAAVAGIHMILIYLCRFLLRRSRNVLFTVSNSLRFRLKVLTVLLYICLSVMLFYLFGSIMPLPPWGLVVGWVMALIAVELAYAFIFPYSFRYIADNDDDKMVILPV |
| IRIS_313-8303 | MLHHLKELAAVAGIHMILIYLCRFLLRRSRNVLFTVSNSLRFRLKVLTVLLYICLSVMLFYLFGSIMPLPPWGLVVGWVMALIAVELAYAFIFPYSFRYIADNDDDKMVILPV |
| IRIS_313-8305 | MLHHLKELAAVAGIHMILIYLCRFLLRRSRNVLFTVSNSLRFRLKVLTVLLYICLSVMLFYLFGSIMPLPPWGLVVGWVMALIAVELAYAFIFPYSFRYIADNDDDKMVILPV |
| IRIS_313-8306 | MLHHLKELAAVAGIHMILIYLCRFLLRRSRNVLFTVSNSLRFRLKVLTVLLYICLSVMLFYLFGSIMPLPPWGLVVGWVMALIAVELAYAFIFPYSFRYIADNDDDKMVILPV |
| IRIS_313-8312 | MLHHLKELAAVAGIHMILIYLCRFLLRRSRNVLFTVSNSLRFRLKVLTVLLYICLSVMLFYLFGSIMPLPPWGLVVGWVMALIAVELAYAFIFPYSFRYIADNDDDKMVILPV |
| IRIS_313-8314 | MLHHLKELAAVAGIHMILIYLCRFLLRRSRNVLFTVSNSLRFRLKVLTVLLYICLSVMLFYLFGSIMPLPPWGLVVGWVMALIAVELAYAFIFPYSFRYIADNDDDKMVILPV |
| IRIS_313-8315 | MLHHLKELAAVAGIHMILIYLCRFLLRRSRNVLFTVSNSLRFRLKVLTVLLYICLSVMLFYLFGSIMPLPPWGLVVGWVMALIAVELAYAFIFPYSFRYIADNDDDKMVILPV |
| IRIS_313-8316 | MLHHLKELAAVAGIHMILIYLCRFLLRRSRNVLFTVSNSLRFRLKVLTVLLYICLSVMLFYLFGSIMPLPPWGLVVGWVMALIAVELAYAFIFPYSFRYIADNDDDKMVILPV |
| IRIS_313-8321 | MLHHLKELAAVAGIHMILIYLCRFLLRRSRNVLFTVSNSLRFRLKVLTVLLYICLSVMLFYLFGSIMPLPPWGLVVGWVMALIAVELAYAFIFPYSFRYIADNDDDKMVILPV |
| IRIS_313-8323 | MLHHLKELAAVAGIHMILIYLCRFLLRRSRNVLFTVSNSLRFRLKVLTVLLYICLSVMLFYLFGSIMPLPPWGLVVGWVMALIAVELAYAFIFPYSFRYIADNDDDKMVILPV |
| IRIS_313-8324 | MLHHLKELAAVAGIHMILIYLCRFLLRRSRNVLFTVSNSLRFRLKVLTVLLYICLSVMLFYLFGSIMPLPPWGLVVGWVMALIAVELAYAFIFPYSFRYIADNDDDKMVILPV |
| IRIS_313-8326 | MLHHLKELAAVAGIHMILIYLCRFLLRRSRNVLFTVSNSLRFRLKVLTVLLYICLSVMLFYLFGSIMPLPPWGLVVGWVMALIAVELAYAFIFPYSFRYIADNDDDKMVILPV |
| IRIS_313-8332 | MLHHLKELAAVAGIHMILIYLCRFLLRRSRNVLFTVSNSLRFRLKVLTVLLYICLSVMLFYLFGSIMPLPPWGLVVGWVMALIAVELAYAFIFPYSFRYIADNDDDKMVILPV |
| IRIS_313-8339 | MLHHLKELAAVAGIHMILIYLCRFLLRRSRNVLFTVSNSLRFRLKVLTVLLYICLSVMLFYLFGSIMPLPPWGLVVGWVMALIAVELAYAFIFPYSFRYIADNDDDKMVILPV |
| IRIS_313-8341 | MLHHLKELAAVAGIHMILIYLCRFLLRRSRNVLFTVSNSLRFRLKVLTVLLYICLSVMLFYLFGSIMPLPPWGLVVGWVMALIAVELAYAFIFPYSFRYIADNDDDKMVILPV |
| IRIS_313-8342 | MLHHLKELAAVAGIHMILIYLCRFLLRRSRNVLFTVSNSLRFRLKVLTVLLYISLSAMLLYLFGSIMPLPPWGLVVGWVMALIAVELAYAFIFPYSFRYIADNDDDKMVILPV |
| IRIS_313-8349 | MLHHLKELAAVAGIHMILIYLCRFLLRRSRNVLFTVSNSLRFRLKVLTVLLYICLSVMLFYLFGSIMPLPPWGLVVGWVMALIAVELAYAFIFPYSFRYIADNDDDKMVILPV |
| IRIS_313-8356 | MLHHLKELAAVAGIHMILIYLCRFLLRRSRNVLFTVSNSLRFRLKVLTVLLYICLSVMLFYLFGSIMPLPPWGLVVGWVMALIAVELAYAFIFPYSFRYIADNDDDKMVILPV |
| IRIS_313-8368 | MLHHLKELAAVAGIHMILIYLCRFLLRRSRNVLFTVSNSLRFRLKVLTVLLYICLSVMLFYLFGSIMPLPPWGLVVGWVMALIAVELAYAFIFPYSFRYIADNDDDKMVILPV |
| IRIS_313-8380 | MLHHLKELAAVAGIHMILIYLCRFLLRRSRNVLFTVSNSLRFRLKVLTVLLYICLSVMLFYLFGSIMPLPPWGLVVGWVMALIAVELAYAFIFPYSFRYIADNDDDKMVILPV |
| IRIS_313-8381 | MLHHLKELAAVAGIHMILIYLCRFLLRRSRNVLFTVSNSLRFRLKVLTVLLYICLSVMLFYLFGSIMPLPPWGLVVGWVMALIAVELAYAFIFPYSFRYIADNDDDKMVILPV |
| IRIS_313-8382 | MLHHLKELAAVAGIHMILIYLCRFLLRRSRNVLFTVSNSLRFRLKVLTVLLYICLSVMLFYLFGSIMPLPPWGLVVGWVMALIAVELAYAFIFPYSFRYIADNDDDKMVILPV |
| IRIS_313-8383 | MLHHLKELAAVAGIHMILIYLCRFLLRRSRNVLFTVSNSLRFRLKVLTVLLYICLSVMLFYLFGSIMPLPPWGLVVGWVMALIAVELAYAFIFPYSFRYIADNDDDKMVILPV |
| IRIS_313-8385 | MLHHLKELAAVAGIHMILIYLCRFLLRRSRNVLFTVSNSLRFRLKVLTVLLYICLSVMLFYLFGSIMPLPPWGLVVGWVMALIAVELAYAFIFPYSFRYIADNDDDKMVILPV |
| IRIS_313-8386 | MLHHLKELAAVAGIHMILIYLCRFLLRRSRNVLFTVSNSLRFRLKVLTVLLYICLSVMLFYLFGSIMPLPPWGLVVGWVMALIAVELAYAFIFPYSFRYIADNDDDKMVILPV |
| IRIS_313-8387 | MLHHLKELAAVAGIHMILIYLCRFLLRRSRNVLFTVSNSLRFRLKVLTVLLYICLSVMLFYLFGSIMPLPPWGLVVGWVMALIAVELAYAFIFPYSFRYIADNDDDKMVILPV |
| IRIS_313-8390 | MLHHLKELAAVAGIHMILIYLCRFLLRRSRNVLFTVSNSLRFRLKVLTVLLYICLSVMLFYLFGSIMPLPPWGLVVGWVMALIAVELAYAFIFPYSFRYIADNDDDKMVILPV |
| IRIS_313-8391 | MLHHLKELAAVAGIHMILIYLCRFLLRRSRNVLFTVSNSLRFRLKVLTVLLYICLSVMLFYLFGSIMPLPPWGLVVGWVMALIAVELAYAFIFPYSFRYIADNDDDKMVILPV |
| IRIS_313-8392 | MLHHLKELAAVAGIHMILIYLCRFLLRRSRNVLFTVSNSLRFRLKVLTVLLYICLSVMLFYLFGSIMPLPPWGLVVGWVMALIAVELAYAFIFPYSFRYIADNDDDKMVILPV |
| IRIS_313-8398 | MLHHLKELAAVAGIHMILIYLCRFLLRRSRNVLFTVSNSLRFRLKVLTVLLYICLSVMLFYLFGSIMPLPPWGLVVGWVMALIAVELAYAFIFPYSFRYIADNDDDKMVILPV |
| IRIS_313-8399 | MLHHLKELAAVAGIHMILIYLCRFLLRRSRNVLFTVSNSLRFRLKVLTVLLYICLSVMLFYLFGSIMPLPPWGLVVGWVMALIAVELAYAFIFPYSFRYIADNDDDKMVILPV |
| IRIS_313-8400 | MLHHLKELAAVAGIHMILIYLCRFLLRRSRNVLFTVSNSLRFRLKVLTVLLYICLSVMLFYLFGSIMPLPPWGLVVGWVMALIAVELAYAFIFPYSFRYIADNDDDKMVILPV |
| IRIS_313-8401 | MLHHLKELAAVAGIHMILIYLCRFLLRRSRNVLFTVSNSLRFRLKVLTVLLYICLSVMLFYLFGSIMPLPPWGLVVGWVMALIAVELAYAFIFPYSFRYIADNDDDKMVILPV |
| IRIS_313-8405 | MLHHLKELAAVAGIHMILIYLCRFLLRRSRNVLFTVSNSLRFRLKVLTVLLYICLSVMLFYLFGSIMPLPPWGLVVGWVMALIAVELAYAFIFPYSFRYIADNDDDKMVILPV |
| IRIS_313-8407 | MLHHLKELAAVAGIHMILIYLCRFLLRRSRNVLFTVSNSLRFRLKVLTVLLYICLSVMLFYLFGSIMPLPPWGLVVGWVMALIAVELAYAFIFPYSFRYIADNDDDKMVILPV |
| IRIS_313-8409 | MLHHLKELAAVAGIHMILIYLCRFLLRRSRNVLFTVSNSLRFRLKVLTVLLYICLSVMLFYLFGSIMPLPPWGLVVGWVMALIAVELAYAFIFPYSFRYIADNDDDKMVILPV |
| IRIS_313-8410 | MLHHLKELAAVAGIHMILIYLCRFLLRRSRNVLFTVSNSLRFRLKVLTVLLYICLSVMLFYLFGSIMPLPPWGLVVGWVMALIAVELAYAFIFPYSFRYIADNDDDKMVILPV |
| IRIS_313-8412 | MLHHLKELAAVAGIHMILIYLCRFLLRRSRNVLFTVSNSLRFRLKVLTVLLYICLSVMLFYLFGSIMPLPPWGLVVGWVMALIAVELAYAFIFPYSFRYIADNDDDKMVILPV |
| IRIS_313-8414 | MLHHLKELAAVAGIHMILIYLCRFLLRRSRNVLFTVSNSLRFRLKVLTVLLYICLSVMLFYLFGSIMPLPPWGLVVGWVMALIAVELAYAFIFPYSFRYIADNDDDKMVILPV |
| IRIS_313-8431 | MLHHLKELAAVAGIHMILIYLCRFLLRRSRNVLFTVSNSLRFRLKVLTVLLYICLSVMLFYLFGSIMPLPPWGLVVGWVMALIAVELAYAFIFPYSFRYIADNDDDKMVILPV |
| IRIS_313-8433 | MLHHLKELAAVAGIHMILIYLCRFLLRLRRNVLFTVSNSLRFRLKVLTVLLYICLSVMLFYLFGSIMPLPPWGLVVGWVMALIAVELAYAFIFPYSFRYIADNDDDKMVILPV |
| IRIS_313-8434 | MLHHLKELAAVAGIHMILIYLCRFLLRRSRNVLFTVSNSLRFRLKVLTVLLYICLSVMLFYLFGSIMPLPPWGLVVGWVMALIAVELAYAFIFPYSFRYIADNDDDKMVILPV |
| IRIS_313-8435 | MLHHLKELAAVAGIHMILIYLCRFLLRRSRNVLFTVSNSLRFRLKVLTVLLYICLSVMLFYLFGSIMPLPPWGLVVGWVMALIAVELAYAFIFPYSFRYIADNDDDKMVILPV |
| IRIS_313-8436 | MLHQLKELAAVAGIHMILIYLCRFLLRRSRNVLFTVSNSLRFRLKVLTVLLYICLLVMLFYLFGSIMPLPPWGLVVGWVMALIAVELAYAFIFPYSFRYIADNDDDKMVILPV |
| IRIS_313-8437 | MLHHLKELAAVAGIHMILIYLCRFLLRRSRNVLFTVSNSLRFRLKVLTVLLYICLSVMLFYLFGSIMPLPPWGLVVGWVMALIAVELAYAFIFPYSFRYIADNDDDKMVILPV |
| IRIS_313-8439 | MLHHLKELAAVAGIHMILIYLCRFLLRRSRNVLFTVSNSLRFRLKVLTVLLYICLSVMLFYLFGSIMPLPPWGLVVGWVMALIAVELAYAFIFPYSFRYIADNDDDKMVILPV |
| IRIS_313-8450 | MLHHLKELAAVAGIHMILIYLCRFLLRRSRNVLFTVSNSLRFRLKVLTVLLYICLSVMLFYLFGSIMPLPPWGLVVGWVMALIAVELAYAFIFPYSFRYIADNDDDKMVILPV |
| IRIS_313-8453 | MLHHLKELAAVAGIHMILIYLCRFLLRRSRNVLFTVSNSLRFRLKVLTVLLYICLSVMLFYLFGSIMPLPPWGLVVGWVMALIAVELAYAFIFPYSFRYIADNDDDKMVILPV |
| IRIS_313-8454 | MLHHLKELAAVAGIHMILIYLCRFLLRRSRNVLFTVSNSLRFRLKVLTVLLYICLSVMLFYLFGSIMPLPPWGLVVGWVMALIAVELAYAFIFPYSFRYIADNDDDKMVILPV |
| IRIS_313-8457 | MLHHLKELAAVAGIHMILIYLCRFLLRRSRNVLFTVSNSLRFRLKVLTVLLYICLSVMLFYLFGSIMPLPPWGLVVGWVMALIAVELAYAFIFPYSFRYIADNDDDKMVILPV |
| IRIS_313-8458 | MLHHLKELAAVAGIHMILIYLCRFLLRRSRNVLFTVSNSLRFRLKVLTVLLYICLSVMLFYLFGSIMPLPPWGLVVGWVMALIAVELAYAFIFPYSFRYIADNDDDKMVILPV |
| IRIS_313-8466 | MLHHLKELAAVAGIHMILIYLCRFLLRRSRNVLFTVSNSLRFRLKVLTVLLYICLSVMLFYLFGSIMPLPPWGLVVGWVMALIAVELAYAFIFPYSFRYIADNDDDKMVILPV |
| IRIS_313-8468 | MLHHLKELAAVAGIHMILIYLCRFLLRRSRNVLFTVSNSLRFRLKVLTVLLYICLSVMLFYLFGSIMPLPPWGLVVGWVMALIAVELAYAFIFPYSFRYIADNDDDKMVILPV |
| IRIS_313-8474 | MLHHLKELAAVAGIHMILIYLCRFLLRRSRNVLFTVSNSLRFRLKVLTVLLYICLSVMLFYLFGSIMPLPPWGLVVGWVMALIAVELAYAFIFPYSFRYIADNDDDKMVILPV |
| IRIS_313-8481 | MLHHLKELAAVAGIHMILIYLCRFLLRRSRNVLFTVSNSLRFRLKVLTVLLYICLSVMLFYLFGSIMPLPPWGLVVGWVMALIAVELAYAFIFPYSFRYIADNDDDKMVILPV |
| IRIS_313-8485 | MLHHLKELAAVAGIHMILIYLCRFLLRRSRNVLFTVSNSLRFRLKVLTVLLYICLSVMLFYLFGSIMPLPPWGLVVGWVMALIAVELAYAFIFPYSFRYIADNDDDKMVILPV |
| IRIS_313-8486 | MLHHLKELAAVAGIHMILIYLCRFLLRRSRNVLFTVSNSLRFRLKVLTVLLYICLSVMLFYLFGSIMPLPPWGLVVGWVMALIAVELAYAFIFPYSFRYIADNDDDKMVILPV |
| IRIS_313-8492 | MLHHLKELAAVAGIHMILIYLCRFLLRRSRNVLFTVSNSLRFRLKVLTVLLYICLSVMLFYLFGSIMPLPPWGLVVGWVMALIAVELAYAFIFPYSFRYIADNDDDKMVILPV |
| IRIS_313-8493 | MLHHLKELAAVAGIHMILIYLCRFLLRRSRNVLFTVSNSLRFRLKVLTVLLYICLSVMLFYLFGSIMPLPPWGLVVGWVMALIAVELAYAFIFPYSFRYIADNDDDKMVILPV |
| IRIS_313-8498 | MLHHLKELAAVAGIHMILIYLCRFLLRRSRNVLFTVSNSLRFRLKVLTVLLYICLSVMLFYLFGSIMPLPPWGLVVGWVMALIAVELAYAFIFPYSFRYIADNDDDKMVILPV |
| IRIS_313-8509 | MLHHLKELAAVAGIHMILIYLCRFLLRRSRNVLFTVSNSLRFRLKVLTVLLYICLSVMLFYLFGSIMPLPPWGLVVGWVMALIAVELAYAFIFPYSFRYIADNDDDKMVILPV |
| IRIS_313-8514 | MLHHLKELAAVAGIHMILIYLCRFLLRRSRNVLFTVSNSLRFRLKVLTVLLYICLSVMLFYLFGSIMPLPPWGLVVGWVMALIAVELAYAFIFPYSFRYIADNDDDKMVILPV |
| IRIS_313-8519 | MLHHLKELAAVAGIHMILIYLCRFLLRRSRNVLFTVSNSLRFRLKVLTVLLYICLSVMLFYLFGSIMPLPPWGLVVGWVMALIAVELAYAFIFPYSFRYIADNDDDKMVILPV |
| IRIS_313-8523 | MLHQLKELAAVAGIHMILIYLCRFLLRRSRNVLFTVSNSLRFRLKVLTVLLYICLLVMLFYLFGSIMPLPPWGLVVGWVMALIAVELAYAFIFPYSFRYIADNDDDKMVILPV |
| IRIS_313-8530 | MLHHLKELAAVAGIHMILIYLCRFLLRRSRNVLFTVSNSLRFRLKVLTVLLYICLSVMLFYLFGSIMPLPPWGLVVGWVMALIAVELAYAFIFPYSFRYIADNDDDKMVILPV |
| IRIS_313-8536 | MLHHLKELAAVAGIHMILIYLCRFLLRRSRNVLFTVSNSLRFRLKVLTVLLYICLSVMLFYLFGSIMPLPPWGLVVGWVMALIAVELAYAFIFPYSFRYIADNDDDKMVILPV |
| IRIS_313-8554 | MLHHLKELAAVAGIHMILIYLCRFLLRRSRNVLFTVSNSLRFRLKVLTVLLYICLSVMLFYLFGSIMPLPPWGLVVGWVMALIAVELAYAFIFPYSFRYIADNDDDKMVILPV |
| IRIS_313-8557 | MLHHLKELAAVAGIHMILIYLCRFLLRRSRNVLFTVSNSLRFRLKVLTVLLYICLSVMLFYLFGSIMPLPPWGLVVGWVMALIAVELAYAFIFPYSFRYIADNDDDKMVILPV |
| IRIS_313-8559 | MLHHLKELAAVAGIHMILIYLCRFLLRRSRNVLFTVSNSLRFRLKVLTVLLYICLSVMLFYLFGSIMPLPPWGLVVGWVMALIAVELAYAFIFPYSFRYIADNDDDKMVILPV |
| IRIS_313-8565 | MLHQLKELAAVAGIHMILIYLCRFLLRRSRNVLFTVSNSLRFRLKVLTVLLYICLLVMLFYLFGSIMPLPPWGLVVGWVMALIAVELAYAFIFPYSFRYIADNDDDKMVILPV |
| IRIS_313-8567 | MLHHLKELAAVAGIHMILIYLCRFLLRRSRNVLFTVSNSLRFRLKVLTVLLYICLSVMLFYLFGSIMPLPPWGLVVGWVMALIAVELAYAFIFPYSFRYIADNDDDKMVILPV |
| IRIS_313-8568 | MLHHLKELAAVAGIHMILIYLCRFLLRRSRNVLFTVSNSLRFRLKVLTVLLYICLSVMLFYLFGSIMPLPPWGLVVGWVMALIAVELAYAFIFPYSFRYIADNDDDKMVILPV |
| IRIS_313-8571 | MLHHLKELAAVAGIHMILIYLCRFLLRRSRNVLFTVSNSLRFRLKVLTVLLYICLSVMLFYLFGSIMPLPPWGLVVGWVMALIAVELAYAFIFPYSFRYIADNDDDKMVILPV |
| IRIS_313-8572 | MLHHLKELAAVAGIHMILIYLCRFLLRRSRNVLFTVSNSLRFRLKVLTVLLYICLSVMLFYLFGSIMPLPPWGLVVGWVMALIAVELAYAFIFPYSFRYIADNDDDKMVILPV |
| IRIS_313-8578 | MLHHLKELAAVAGIHMILIYLCRFLLRRSRNVLFTVSNSLRFRLKVLTVLLYICLSVMLFYLFGSIMPLPPWGLVVGWVMALIAVELAYAFIFPYSFRYIADNDDDKMVILPV |
| IRIS_313-8580 | MLHHLKELAAVAGIHMILIYLCRFLLRRSRNVLFTVSNSLRFRLKVLTVLLYICLSVMLFYLFGSIMPLPPWGLVVGWVMALIAVELAYAFIFPYSFRYIADNDDDKMVILPV |
| IRIS_313-8585 | MLHHLKELAAVAGIHMILIYLCRFLLRRSRNVLFTVSNSLRFRLKVLTVLLYICLSVMLFYLFGSIMPLPPWGLVVGWVMALIAVELAYAFIFPYSFRYIADNDDDKMVILPV |
| IRIS_313-8586 | MLHHLKELAAVAGIHMILIYLCRFLLRRSRNVLFTVSNSLRFRLKVLTVLLYICLSVMLFYLFGSIMPLPPWGLVVGWVMALIAVELAYAFIFPYSFRYIADNDDDKMVILPV |
| IRIS_313-8591 | MLHHLKELAAVAGIHMILIYLCRFLLRRSRNVLFTVSNSLRFRLKVLTVLLYICLSVMLFYLFGSIMPLPPWGLVVGWVMALIAVELAYAFIFPYSFRYIADNDDDKMVILPV |
| IRIS_313-8594 | MLHHLKELAAVAGIHMILIYLCRFLLRRSRNVLFTVSNSLRFRLKVLTVLLYICLSVMLFYLFGSIMPLPPWGLVVGWVMALIAVELAYAFIFPYSFRYIADNDDDKMVILPV |
| IRIS_313-8595 | MLHHLKELAAVAGIHMILIYLCRFLLRRSRNVLFTVSNSLRFRLKVLTVLLYICLSVMLFYLFGSIMPLPPWGLVVGWVMALIAVELAYAFIFPYSFRYIADNDDDKMVILPV |
| IRIS_313-8599 | MLHHLKELAAVAGIHMILIYLCRFLLRRSRNVLFTVSNSLRFRLKVLTVLLYICLSVMLFYLFGSIMPLPPWGLVVGWVMALIAVELAYAFIFPYSFRYIADNDDDKMVILPV |
| IRIS_313-8603 | MLHHLKELAAVAGIHMILIYLCRFLLRRSRNVLFTVSNSLRFRLKVLTVLLYICLSVMLFYLFGSIMPLPPWGLVVGWVMALIAVELAYAFIFPYSFRYIADNDDDKMVILPV |
| IRIS_313-8606 | MLHHLKELAAVAGIHMILIYLCRFLLRRSRNVLFTVSNSLRFRLKVLTVLLYICLSVMLFYLFGSIMPLPPWGLVVGWVMALIAVELAYAFIFPYSFRYIADNDDDKMVILPV |
| IRIS_313-8608 | MLHHLKELAAVAGIHMILIYLCRFLLRRSRNVLFTVSNSLRFRLKVLTVLLYICLSVMLFYLFGSIMPLPPWGLVVGWVMALIAVELAYAFIFPYSFRYIADNDDDKMVILPV |
| IRIS_313-8614 | MLHHLKELAAVAGIHMILIYLCRFLLRRSRNVLFTVSNSLRFRLKVLTVLLYICLSVMLFYLFGSIMPLPPWGLVVGWVMALIAVELAYAFIFPYSFRYIADNDDDKMVILPV |
| IRIS_313-8616 | MLHHLKELAAVAGIHMILIYLCRFLLRRSRNVLFTVSNSLRFRLKVLTVLLYICLSVMLFYLFGSIMPLPPWGLVVGWVMALIAVELAYAFIFPYSFRYIADNDDDKMVILPV |

| | |
|---|---|
| IRIS_313-8621 | MLHHLKELAAVAGIHMILIYLCRFLLRRSRNVLFTVSNSLRFRLKVLTVLLYICLSVMLFYLFGSIMPLPPWGLVVGWVMALIAVELAYAFIFPYSFRYIADNDDDKMVILPV |
| IRIS_313-8622 | MLHHLKELAAVAGIHMILIYLCRFLLRRSRNVLFTVSNSLRFRLKVLTVLLYICLSVMLFYLFGSIMPLPPWGLVVGWVMALIAVELAYAFIFPYSFRYIADNDDDKMVILPV |
| IRIS_313-8626 | MLHQLKELAAVAGIHMILIYLCRFLLRRSRNVLFTVSNSLRFRLKVLTVLLYICLLVMLFYLFGSIMPLPPWGLVVGWVMALIAVELAYAFIFPYSFRYIADNDDDKMVILPV |
| IRIS_313-8627 | MLHHLKELAAVAGIHMILIYLCRFLLRRSRNVLFTVSNSLRFRLKVLTVLLYICLSVMLFYLFGSIMPLPPWGLVVGWVMALIAVELAYAFIFPYSFRYIADNDDDKMVILPV |
| IRIS_313-8631 | MLHHLKELAAVAGIHMILIYLCRFLLRRSRNVLFTVSNSLRFRLKVLTVLLYICLSVMLFYLFGSIMPLPPWGLVVGWVMALIAVELAYAFIFPYSFRYIADNDDDKMVILPV |
| IRIS_313-8632 | MLHHLKELAAVAGIHMILIYLCRFLLRRSRNVLFTVSNSLRFRLKVLTVLLYICLSVMLFYLFGSIMPLPPWGLVVGWVMALIAVELAYAFIFPYSFRYIADNDDDKMVILPV |
| IRIS_313-8637 | MLHHLKELAAVAGIHMILIYLCRFLLRRSRNVLFTVSNSLRFRLKVLTVLLYICLSVMLFYLFGSIMPLPPWGLVVGWVMALIAVELAYAFIFPYSFRYIADNDDDKMVILPV |
| IRIS_313-8638 | MLHHLKELAAVAGIHMILIYLCRFLLRRSRNVLFTVSNSLRFRLKVLTVLLYICLSVMLFYLFGSIMPLPPWGLVVGWVMALIAVELAYAFIFPYSFRYIADNDDDKMVILPV |
| IRIS_313-8641 | MLHHLKELAAVAGIHMILIYLCRFLLRRSRNVLFTVSNSLRFRLKVLTVLLYICLSVMLFYLFGSIMPLPPWGLVVGWVMALIAVELAYAFIFPYSFRYIADNDDDKMVILPV |
| IRIS_313-8643 | MLHHLKELAAVAGIHMILIYLCRFLLRRSRNVLFTVSNSLRFRLKVLTVLLYICLSVMLFYLFGSIMPLPPWGLVVGWVMALIAVELAYAFIFPYSFRYIADNDDDKMVILPV |
| IRIS_313-8645 | MLHHLKELAAVAGIHMILIYLCRFLLRRSRNVLFTVSNSLRFRLKVLTVLLYICLSVMLFYLFGSIMPLPPWGLVVGWVMALIAVELAYAFIFPYSFRYIADNDDDKMVILPV |
| IRIS_313-8647 | MLHHLKELAAVAGIHMILIYLCRFLLRRSRNVLFTVSNSLRFRLKVLTVLLYICLSVMLFYLFGSIMPLPPWGLVVGWVMALIAVELAYAFIFPYSFRYIADNDDDKMVILPV |
| IRIS_313-8655 | MLHHLKELAAVAGIHMILIYLCRFLLRRSRNVLFTVSNSLRFRLKVLTVLLYICLSVMLFYLFGSIMPLPPWGLVVGWVMALIAVELAYAFIFPYSFRYIADNDDDKMVILPV |
| IRIS_313-8656 | MLHHLKELAAVAGIHMILIYLCRFLLRRSRNVLFTVSNSLRFRLKVLTVLLYICLSVMLFYLFGSIMPLPPWGLVVGWVMALIAVELAYAFIFPYSFRYIADNDDDKMVILPV |
| IRIS_313-8657 | MLHHLKELAAVAGIHMILIYLCRFLLRRSRNVLFTVSNSLRFRLKVLTVLLYICLSVMLFYLFGSIMPLPPWGLVVGWVMALIAVELAYAFIFPYSFRYIADNDDDKMVILPV |
| IRIS_313-8658 | MLHHLKELAAVAGIHMILIYLCRFLLRRSRNVLFTVSNSLRFRLKVLTVLLYICLSVMLFYLFGSIMPLPPWGLVVGWVMALIAVELAYAFIFPYSFRYIADNDDDKMVILPV |
| IRIS_313-8659 | MLHHLKELAAVAGIHMILIYLCRFLLRRSRNVLFTVSNSLRFRLKVLTVLLYICLSVMLFYLFGSIMPLPPWGLVVGWVMALIAVELAYAFIFPYSFRYIADNDDDKMVILPV |
| IRIS_313-8660 | MLHHLKELAAVAGIHMILIYLCRFLLRRSRNVLFTVSNSLRFRLKVLTVLLYICLSVMLFYLFGSIMPLPPWGLVVGWVMALIAVELAYAFIFPYSFRYIADNDDDKMVILPV |
| IRIS_313-8664 | MLHHLKELAAVAGIHMILIYLCRFLLRRSRNVLFTVSNSLRFRLKVLTVLLYICLSVMLFYLFGSIMPLPPWGLVVGWVMALIAVELAYAFIFPYSFRYIADNDDDKMVILPV |
| IRIS_313-8665 | MLHHLKELAAVAGIHMILIYLCRFLLRRSRNVLFTVSNSLRFRLKVLTVLLYICLSVMLFYLFGSIMPLPPWGLVVGWVMALIAVELAYAFIFPYSFRYIADNDDDKMVILPV |
| IRIS_313-8669 | MLHHLKELAAVAGIHMILIYLCRFLLRRSRNVLFTVSNSLRFRLKVLTVLLYICLSVMLFYLFGSIMPLPPWGLVVGWVMALIAVELAYAFIFPYSFRYIADNDDDKMVILPV |
| IRIS_313-8674 | MLHHLKELAAVAGIHMILIYLCRFLLRRSRNVLFTVSNSLRFRLKVLTVLLYICLSVMLFYLFGSIMPLPPWGLVVGWVMALIAVELAYAFIFPYSFRYIADNDDDKMVILPV |
| IRIS_313-8679 | MLHHLKELAAVAGIHMILIYLCRFLLRRSRNVLFTVSNSLRFRLKVLTVLLYICLSVMLFYLFGSIMPLPPWGLVVGWVMALIAVELAYAFIFPYSFRYIADNDDDKMVILPV |
| IRIS_313-8681 | MLHHLKELAAVAGIHMILIYLCRFLLRRSRNVLFTVSNSLRFRLKVLTVLLYICLSVMLFYLFGSIMPLPPWGLVVGWVMALIAVELAYAFIFPYSFRYIADNDDDKMVILPV |
| IRIS_313-8683 | MLHHLKELAAVAGIHMILIYLCRFLLRRSRNVLFTVSNSLRFRLKVLTVLLYICLSVMLFYLFGSIMPLPPWGLVVGWVMALIAVELAYAFIFPYSFRYIADNDDDKMVILPV |
| IRIS_313-8687 | MLHHLKELAAVAGIHMILIYLCRFLLRRSRNVLFTVSNSLRFRLKVLTVLLYICLSVMLFYLFGSIMPLPPWGLVVGWVMALIAVELAYAFIFPYSFRYIADNDDDKMVILPV |
| IRIS_313-8690 | MLHHLKELAAVAGIHMILIYLCRFLLRRSRNVLFTVSNSLRFRLKVLTVLLYICLSVMLFYLFGSIMPLPPWGLVVGWVMALIAVELAYAFIFPYSFRYIADNDDDKMVILPV |
| IRIS_313-8694 | MLHHLKELAAVAGIHMILIYLCRFLLRRSRNVLFTVSNSLRFRLKVLTVLLYICLSVMLFYLFGSIMPLPPWGLVVGWVMALIAVELAYAFIFPYSFRYIADNDDDKMVILPV |
| IRIS_313-8697 | MLHHLKELAAVAGIHMILIYLCRFLLRRSRNVLFTVSNSLRFRLKVLTVLLYICLSVMLFYLFGSIMPLPPWGLVVGWVMALIAVELAYAFIFPYSFRYIADNDDDKMVILPV |
| IRIS_313-8699 | MLHHLKELAAVAGIHMILIYLCRFLLRRSRNVLFTVSNSLRFRLKVLTVLLYICLSVMLFYLFGSIMPLPPWGLVVGWVMALIAVELAYAFIFPYSFRYIADNDDDKMVILPV |
| IRIS_313-8702 | MLHHLKELAAVAGIHMILIYLCRFLLRRSRNVLFTVSNSLRFRLKVLTVLLYICLSVMLFYLFGSIMPLPPWGLVVGWVMALIAVELAYAFIFPYSFRYIADNDDDKMVILPV |
| IRIS_313-8703 | MLHHLKELAAVAGIHMILIYLCRFLLRRSRNVLFTVSNSLRFRLKVLTVLLYICLSVMLFYLFGSIMPLPPWGLVVGWVMALIAVELAYAFIFPYSFRYIADNDDDKMVILPV |
| IRIS_313-8704 | MLHHLKELAAVAGIHMILIYLCRFLLRRSRNVLFTVSNSLRFRLKVLTVLLYICLSVMLFYLFGSIMPLPPWGLVVGWVMALIAVELAYAFIFPYSFRYIADNDDDKMVILPV |
| IRIS_313-8712 | MLHHLKELAAVAGIHMILIYLCRFLLRRSRNVLFTVSNSLRFRLKVLTVLLYICLSVMLFYLFGSIMPLPPWGLVVGWVMALIAVELAYAFIFPYSFRYIADNDDDKMVILPV |
| IRIS_313-8713 | MLHHLKELAAVAGIHMILIYLCRFLLRRSRNVLFTVSNSLRFRLKVLTVLLYICLSVMLFYLFGSIMPLPPWGLVVGWVMALIAVELAYAFIFPYSFRYIADNDDDKMVILPV |
| IRIS_313-8717 | MLHHLKELAAVAGIHMILIYLCRFLLRRSRNVLFTVSNSLRFRLKVLTVLLYICLSVMLFYLFGSIMPLPPWGLVVGWVMALIAVELAYAFIFPYSFRYIADNDDDKMVILPV |
| IRIS_313-8721 | MLHHLKELAAVAGIHMILIYLCRFLLRRSRNVLFTVSNSLRFRLKVLTVLLYICLSVMLFYLFGSIMPLPPWGLVVGWVMALIAVELAYAFIFPYSFRYIADNDDDKMVILPV |
| IRIS_313-8722 | MLHHLKELAAVAGIHMILIYLCRFLLRRSRNVLFTVSNSLRFRLKVLTVLLYICLSVMLFYLFGSIMPLPPWGLVVGWVMALIAVELAYAFIFPYSFRYIADNDDDKMVILPV |
| IRIS_313-8723 | MLHHLKELAAVAGIHMILIYLCRFLLRRSRNVLFTVSNSLRFRLKVLTVLLYICLSVMLFYLFGSIMPLPPWGLVVGWVMALIAVELAYAFIFPYSFRYIADNDDDKMVILPV |
| IRIS_313-8725 | MLHHLKELAAVAGIHMILIYLCRFLLRRSRNVLFTVSNSLRFRLKVLTVLLYICLSVMLFYLFGSIMPLPPWGLVVGWVMALIAVELAYAFIFPYSFRYIADNDDDKMVILPV |
| IRIS_313-8727 | MLHHLKELAAVAGIHMILIYLCRFLLRRSRNVLFTVSNSLRFRLKVLTVLLYICLSVMLFYLFGSIMPLPPWGLVVGWVMALIAVELAYAFIFPYSFRYIADNDDDKMVILPV |
| IRIS_313-8731 | MLHHLKELAAVAGIHMILIYLCRFLLRRSRNVLFTVSNSLRFRLKVLTVLLYICLSVMLFYLFGSIMPLPPWGLVVGWVMALIAVELAYAFIFPYSFRYIADNDDDKMVILPV |
| IRIS_313-8732 | MLHHLKELAAVAGIHMILIYLCRFLLRRSRNVLFTVSNSLRFRLKVLTVLLYICLSVMLFYLFGSIMPLPPWGLVVGWVMALIAVELAYAFIFPYSFRYIADNDDDKMVILPV |
| IRIS_313-8733 | MLHHLKELAAVAGIHMILIYLCRFLLRRSRNVLFTVSNSLRFRLKVLTVLLYICLSVMLFYLFGSIMPLPPWGLVVGWVMALIAVELAYAFIFPYSFRYIADNDDDKMVILPV |
| IRIS_313-8735 | MLHHLKELAAVAGIHMILIYLCRFLLRRSRNVLFTVSNSLRFRLKVLTVLLYICLSVMLFYLFGSIMPLPPWGLVVGWVMALIAVELAYAFIFPYSFRYIADNDDDKMVILPV |
| IRIS_313-8737 | MLHHLKELAAVAGIHMILIYLCRFLLRRSRNVLFTVSNSLRFRLKVLTVLLYICLSVMLFYLFGSIMPLPPWGLVVGWVMALIAVELAYAFIFPYSFRYIADNDDDKMVILPV |
| IRIS_313-8739 | MLHHLKELAAVAGIHMILIYLCRFLLRRSRNVLFTVSNSLRFRLKVLTVLLYICLSVMLFYLFGSIMPLPPWGLVVGWVMALIAVELAYAFIFPYSFRYIADNDDDKMVILPV |
| IRIS_313-8743 | MLHHLKELAAVAGIHMILIYLCRFLLRRSRNVLFTVSNSLRFRLKVLTVLLYICLSVMLFYLFGSIMPLPPWGLVVGWVMALIAVELAYAFIFPYSFRYIADNDDDKMVILPV |
| IRIS_313-8744 | MLHHLKELAAVAGIHMILIYLCRFLLRRSRNVLFTVSNSLRFRLKVLTVLLYICLSVMLFYLFGSIMPLPPWGLVVGWVMALIAVELAYAFIFPYSFRYIADNDDDKMVILPV |
| IRIS_313-8745 | MLHHLKELAAVAGIHMILIYLCRFLLRRSRNVLFTVSNSLRFRLKVLTVLLYICLSVMLFYLFGSIMPLPPWGLVVGWVMALIAVELAYAFIFPYSFRYIADNDDDKMVILPV |
| IRIS_313-8747 | MLHHLKELAAVAGIHMILIYLCRFLLRRSRNVLFTVSNSLRFRLKVLTVLLYICLSVMLFYLFGSIMPLPPWGLVVGWVMALIAVELAYAFIFPYSFRYIADNDDDKMVILPV |
| IRIS_313-8751 | MLHHLKELAAVAGIHMILIYLCRFLLRRSRNVLFTVSNSLRFRLKVLTVLLYICLSVMLFYLFGSIMPLPPWGLVVGWVMALIAVELAYAFIFPYSFRYIADNDDDKMVILPV |
| IRIS_313-8754 | MLHHLKELAAVAGIHMILIYLCRFLLRRSRNVLFTVSNSLRFRLKVLTVLLYICLSVMLFYLFGSIMPLPPWGLVVGWVMALIAVELAYAFIFPYSFRYIADNDDDKMVILPV |
| IRIS_313-8755 | MLHHLKELAAVAGIHMILIYLCRFLLRRSRNVLFTVSNSLRFRLKVLTVLLYICLSVMLFYLFGSIMPLPPWGLVVGWVMALIAVELAYAFIFPYSFRYIADNDDDKMVILPV |
| IRIS_313-8757 | MLHHLKELAAVAGIHMILIYLCRFLLRRSRNVLFTVSNSLRFRLKVLTVLLYICLSVMLFYLFGSIMPLPPWGLVVGWVMALIAVELAYAFIFPYSFRYIADNDDDKMVILPV |
| IRIS_313-8765 | MLHHLKELAAVAGIHMILIYLCRFLLRRSRNVLFTVSNSLRFRLKVLTVLLYICLSVMLFYLFGSIMPLPPWGLVVGWVMALIAVELAYAFIFPYSFRYIADNDDDKMVILPV |
| IRIS_313-8767 | MLHHLKELAAVAGIHMILIYLCRFLLRRSRNVLFTVSNSLRFRLKVLTVLLYICLSVMLFYLFGSIMPLPPWGLVVGWVMALIAVELAYAFIFPYSFRYIADNDDDKMVILPV |
| IRIS_313-8768 | MLHHLKELAAVAGIHMILIYLCRFLLRRSRNVLFTVSNSLRFRLKVLTVLLYICLSVMLFYLFGSIMPLPPWGLVVGWVMALIAVELAYAFIFPYSFRYIADNDDDKMVILPV |
| IRIS_313-8769 | MLHHLKELAAVAGIHMILIYLCRFLLRRSRNVLFTVSNSLRFRLKVLTVLLYICLSVMLFYLFGSIMPLPPWGLVVGWVMALIAVELAYAFIFPYSFRYIADNDDDKMVILPV |
| IRIS_313-8771 | MLHHLKELAAVAGIHMILIYLCRFLLRRSRNVLFTVSNSLRFRLKVLTVLLYICLSVMLFYLFGSIMPLPPWGLVVGWVMALIAVELAYAFIFPYSFRYIADNDDDKMVILPV |
| IRIS_313-8778 | MLHHLKELAAVAGIHMILIYLCRFLLRRSRNVLFTVSNSLRFRLKVLTVLLYICLSVMLFYLFGSIMPLPPWGLVVGWVMALIAVELAYAFIFPYSFRYIADNDDDKMVILPV |
| IRIS_313-8781 | MLHHLKELAAVAGIHMILIYLCRFLLRRSRNVLFTVSNSLRFRLKVLTVLLYICLSVMLFYLFGSIMPLPPWGLVVGWVMALIAVELAYAFIFPYSFRYIADNDDDKMVILPV |
| IRIS_313-8783 | MLHHLKELAAVAGIHMILIYLCRFLLRRSRNVLFTVSNSLRFRLKVLTVLLYICLSVMLFYLFGSIMPLPPWGLVVGWVMALIAVELAYAFIFPYSFRYIADNDDDKMVILPV |
| IRIS_313-8785 | MLHHLKELAAVAGIHMILIYLCRFLLRRSRNVLFTVSNSLRFRLKVLTVLLYICLSVMLFYLFGSIMPLPPWGLVVGWVMALIAVELAYAFIFPYSFRYIADNDDDKMVILPV |
| IRIS_313-8789 | MLHHLKELAAVAGIHMILIYLCRFLLRRSRNVLFTVSNSLRFRLKVLTVLLYICLSVMLFYLFGSIMPLPPWGLVVGWVMALIAVELAYAFIFPYSFRYIADNDDDKMVILPV |
| IRIS_313-8791 | MLHHLKELAAVAGIHMILIYLCRFLLRRSRNVLFTVSNSLRFRLKVLTVLLYICLSVMLFYLFGSIMPLPPWGLVVGWVMALIAVELAYAFIFPYSFRYIADNDDDKMVILPV |
| IRIS_313-8793 | MLHHLKELAAVAGIHMILIYLCRFLLRRSRNVLFTVSNSLRFRLKVLTVLLYICLSVMLFYLFGSIMPLPPWGLVVGWVMALIAVELAYAFIFPYSFRYIADNDDDKMVILPV |
| IRIS_313-8796 | MLHHLKELAAVAGIHMILIYLCRFLLRRSRNVLFTVSNSLRFRLKVLTVLLYICLSVMLFYLFGSIMPLPPWGLVVGWVMALIAVELAYAFIFPYSFRYIADNDDDKMVILPV |
| IRIS_313-8803 | MLHHLKELAAVAGIHMILIYLCRFLLRRSRNVLFTVSNSLRFRLKVLTVLLYICLSVMLFYLFGSIMPLPPWGLVVGWVMALIAVELAYAFIFPYSFRYIADNDDDKMVILPV |
| IRIS_313-8811 | MLHHLKELAAVAGIHMILIYLCRFLLRRSRNVLFTVSNSLRFRLKVLTVLLYICLSVMLFYLFGSIMPLPPWGLVVGWVMALIAVELAYAFIFPYSFRYIADNDDDKMVILPV |
| IRIS_313-8812 | MLHHLKELAAVAGIHMILIYLCRFLLRRSRNVLFTVSNSLRFRLKVLTVLLYICLSVMLFYLFGSIMPLPPWGLVVGWVMALIAVELAYAFIFPYSFRYIADNDDDKMVILPV |
| IRIS_313-8813 | MLHHLKELAAVAGIHMILIYLCRFLLRRSRNVLFTVSNSLRFRLKVLTVLLYICLSVMLFYLFGSIMPLPPWGLVVGWVMALIAVELAYAFIFPYSFRYIADNDDDKMVILPV |
| IRIS_313-8814 | MLHHLKELAAVAGIHMILIYLCRFLLRRSRNVLFTVSNSLRFRLKVLTVLLYICLSVMLFYLFGSIMPLPPWGLVVGWVMALIAVELAYAFIFPYSFRYIADNDDDKMVILPV |
| IRIS_313-8815 | MLHHLKELAAVAGIHMILIYLCRFLLRRSRNVLFTVSNSLRFRLKVLTVLLYICLSVMLFYLFGSIMPLPPWGLVVGWVMALIAVELAYAFIFPYSFRYIADNDDDKMVILPV |
| IRIS_313-8822 | MLHHLKELAAVAGIHMILIYLCRFLLRRSRNVLFTVSNSLRFRLKVLTVLLYICLSVMLFYLFGSIMPLPPWGLVVGWVMALIAVELAYAFIFPYSFRYIADNDDDKMVILPV |
| IRIS_313-8831 | MLHHLKELAAVAGIHMILIYLCRFLLRRSRNVLFTVSNSLRFRLKVLTVLLYICLSVMLFYLFGSIMPLPPWGLVVGWVMALIAVELAYAFIFPYSFRYIADNDDDKMVILPV |
| IRIS_313-8833 | MLHHLKELAAVAGIHMILIYLCRFLLRRSRNVLFTVSNSLRFRLKVLTVLLYICLSVMLFYLFGSIMPLPPWGLVVGWVMALIAVELAYAFIFPYSFRYIADNDDDKMVILPV |
| IRIS_313-8844 | MLHHLKELAAVAGIHMILIYLCRFLLRRSRNVLFTVSNSLRFRLKVLTVLLYICLSVMLFYLFGSIMPLPPWGLVVGWVMALIAVELAYAFIFPYSFRYIADNDDDKMVILPV |
| IRIS_313-8846 | MLHHLKELAAVAGIHMILIYLCRFLLRRSRNVLFTVSNSLRFRLKVLTVLLYICLSVMLFYLFGSIMPLPPWGLVVGWVMALIAVELAYAFIFPYSFRYIADNDDDKMVILPV |
| IRIS_313-8850 | MLHHLKELAAVAGIHMILIYLCRFLLRRSRNVLFTVSNSLRFRLKVLTVLLYICLSVMLFYLFGSIMPLPPWGLVVGWVMALIAVELAYAFIFPYSFRYIADNDDDKMVILPV |
| IRIS_313-8854 | MLHHLKELAAVAGIHMILIYLCRFLLRRSRNVLFTVSNSLRFRLKVLTVLLYICLSVMLFYLFGSIMPLPPWGLVVGWVMALIAVELAYAFIFPYSFRYIADNDDDKMVILPV |
| IRIS_313-8856 | MLHHLKELAAVAGIHMILIYLCRFLLRRSRNVLFTVSNSLRFRLKVLTVLLYICLSVMLFYLFGSIMPLPPWGLVVGWVMALIAVELAYAFIFPYSFRYIADNDDDKMVILPV |
| IRIS_313-8857 | MLHHLKELAAVAGIHMILIYLCRFLLRRSRNVLFTVSNSLRFRLKVLTVLLYICLSVMLFYLFGSIMPLPPWGLVVGWVMALIAVELAYAFIFPYSFRYIADNDDDKMVILPV |
| IRIS_313-8859 | MLHHLKELAAVAGIHMILIYLCRFLLRRSRNVLFTVSNSLRFRLKVLTVLLYICLSVMLFYLFGSIMPLPPWGLVVGWVMALIAVELAYAFIFPYSFRYIADNDDDKMVILPV |
| IRIS_313-8864 | MLHHLKELAAVAGIHMILIYLCRFLLRRSRNVLFTVSNSLRFRLKVLTVLLYICLSVMLFYLFGSIMPLPPWGLVVGWVMALIAVELAYAFIFPYSFRYIADNDDDKMVILPV |
| IRIS_313-8865 | MLHQLKELAAVAGIHMILIYLCRFLLRRSRNVLFTVSNSLRFRLKVLTVLLYICLLVMLFYLFGSIMPLPPWGLVVGWVMALIAVELAYAFIFPYSFRYIADNDDDKMVILPV |
| IRIS_313-8870 | MLHHLKELAAVAGIHMILIYLCRFLLRRSRNVLFTVSNSLRFRLKVLTVLLYICLSVMLFYLFGSIMPLPPWGLVVGWVMALIAVELAYAFIFPYSFRYIADNDDDKMVILPV |
| IRIS_313-8872 | MLHHLKELAAVAGIHMILIYLCRFLLRRSRNVLFTVSNSLRFRLKVLTVLLYICLSVMLFYLFGSIMPLPPWGLVVGWVMALIAVELAYAFIFPYSFRYIADNDDDKMVILPV |
| IRIS_313-8873 | MLHHLKELAAVAGIHMILIYLCRFLLRRSRNVLFTVSNSLRFRLKVLTVLLYICLSVMLFYLFGSIMPLPPWGLVVGWVMALIAVELAYAFIFPYSFRYIADNDDDKMVILPV |
| IRIS_313-8876 | MLHHLKELAAVAGIHMILIYLCRFLLRRSRNVLFTVSNSLRFRLKVLTVLLYICLSVMLFYLFGSIMPLPPWGLVVGWVMALIAVELAYAFIFPYSFRYIADNDDDKMVILPV |
| IRIS_313-8880 | MLHQLKELAAVAGIHMILIYLCRFLLRRSRNVLFTVSNSLRFRLKVLTVLLYICLLVMLFYLFGSIMPLPPWGLVVGWVMALIAVELAYAFIFPYSFRYIADNDDDKMVILPV |
| IRIS_313-8883 | MLHQLKELAAVAGIHMILIYLCRFLLRRSRNVLFTVSNSLRFRLKVLTVLLYICLLVMLFYLFGSIMPLPPWGLVVGWVMALIAVELAYAFIFPYSFRYIADNDDDKMVILPV |
| IRIS_313-8884 | MLHHLKELAAVAGIHMILIYLCRFLLRRSRNVLFTVSNSLRFRLKVLTVLLYICLSVMLFYLFGSIMPLPPWGLVVGWVMALIAVELAYAFIFPYSFRYIADNDDDKMVILPV |
| IRIS_313-8889 | MLHHLKELAAVAGIHMILIYLCRFLLRRSRNVLFTVSNSLRFRLKVLTVLLYICLSVMLFYLFGSIMPLPPWGLVVGWVMALIAVELAYAFIFPYSFRYIADNDDDKMVILPV |
| IRIS_313-8890 | MLHHLKELAAVAGIHMILIYLCRFLLRRSRNVLFTVSNSLRFRLKVLTVLLYICLSVMLFYLFGSIMPLPPWGLVVGWVMALIAVELAYAFIFPYSFRYIADNDDDKMVILPV |
| IRIS_313-8894 | MLHHLKELAAVAGIHMILIYLCRFLLRRSRNVLFTVSNSLRFRLKVLTVLLYICLSVMLFYLFGSIMPLPPWGLVVGWVMALIAVELAYAFIFPYSFRYIADNDDDKMVILPV |
| IRIS_313-8895 | MLHHLKELAAVAGIHMILIYLCRFLLRRSRNVLFTVSNSLRFRLKVLTVLLYICLSVMLFYLFGSIMPLPPWGLVVGWVMALIAVELAYAFIFPYSFRYIADNDDDKMVILPV |
| IRIS_313-8900 | MLHHLKELAAVAGIHMILIYLCRFLLRRSRNVLFTVSNSLRFRLKVLTVLLYICLSVMLFYLFGSIMPLPPWGLVVGWVMALIAVELAYAFIFPYSFRYIADNDDDKMVILPV |
| IRIS_313-8903 | MLHHLKELAAVAGIHMILIYLCRFLLRRSRNVLFTVSNSLRFRLKVLTVLLYICLSVMLFYLFGSIMPLPPWGLVVGWVMALIAVELAYAFIFPYSFRYIADNDDDKMVILPV |
| IRIS_313-8909 | MLHHLKELAAVAGIHMILIYLCRFLLRRSRNVLFTVSNSLRFRLKVLTVLLYICLSVMLFYLFGSIMPLPPWGLVVGWVMALIAVELAYAFIFPYSFRYIADNDDDKMVILPV |
| IRIS_313-8911 | MLHHLKELAAVAGIHMILIYLCRFLLRRSRNVLFTVSNSLRFRLKVLTVLLYICLSVMLFYLFGSIMPLPPWGLVVGWVMALIAVELAYAFIFPYSFRYIADNDDDKMVILPV |
| IRIS_313-8914 | MLHHLKELAAVAGIHMILIYLCRFLLRRSRNVLFTVSNSLRFRLKVLTVLLYICLSVMLFYLFGSIMPLPPWGLVVGWVMALIAVELAYAFIFPYSFRYIADNDDDKMVILPV |
| IRIS_313-8920 | MLHHLKELAAVAGIHMILIYLCRFLLRRSRNVLFTVSNSLRFRLKVLTVLLYICLSVMLFYLFGSIMPLPPWGLVVGWVMALIAVELAYAFIFPYSFRYIADNDDDKMVILPV |
| IRIS_313-8922 | MLHHLKELAAVAGIHMILIYLCRFLLRRSRNVLFTVSNSLRFRLKVLTVLLYICLSVMLFYLFGSIMPLPPWGLVVGWVMALIAVELAYAFIFPYSFRYIADNDDDKMVILPV |
| IRIS_313-8923 | MLHHLKELAAVAGIHMILIYLCRFLLRRSRNVLFTVSNSLRFRLKVLTVLLYICLSVMLFYLFGSIMPLPPWGLVVGWVMALIAVELAYAFIFPYSFRYIADNDDDKMVILPV |
| IRIS_313-8924 | MLHHLKELAAVAGIHMILIYLCRFLLRRSRNVLFTVSNSLRFRLKVLTVLLYICLSVMLFYLFGSIMPLPPWGLVVGWVMALIAVELAYAFIFPYSFRYIADNDDDKMVILPV |
| IRIS_313-8925 | MLHHLKELAAVAGIHMILIYLCRFLLRRSRNVLFTVSNSLRFRLKVLTVLLYICLSVMLFYLFGSIMPLPPWGLVVGWVMALIAVELAYAFIFPYSFRYIADNDDDKMVILPV |
| IRIS_313-8927 | MLHQLKELAAVAGIHMILIYLCRFLLRRSRNVLFTVSNSLRFRLKVLTVLLYICLLVMLFYLFGSIMPLPPWGLVVGWVMALIAVELAYAFIFPYSFRYIADNDDDKMVILPV |
| IRIS_313-8930 | MLHHLKELAAVAGIHMILIYLCRFLLRRSRNVLFTVSNSLRFRLKVLTVLLYICLSVMLFYLFGSIMPLPPWGLVVGWVMALIAVELAYAFIFPYSFRYIADNDDDKMVILPV |
| IRIS_313-8932 | MLHHLKELAAVAGIHMILIYLCRFLLRRSRNVLFTVSNSLRFRLKVLTVLLYICLSVMLFYLFGSIMPLPPWGLVVGWVMALIAVELAYAFIFPYSFRYIADNDDDKMVILPV |
| IRIS_313-8935 | MLHHLKELAAVAGIHMILIYLCRFLLRRSRNVLFTVSNSLRFRLKVLTVLLYICLSVMLFYLFGSIMPLPPWGLVVGWVMALIAVELAYAFIFPYSFRYIADNDDDKMVILPV |
| IRIS_313-8940 | MLHHLKELAAVAGIHMILIYLCRFLLRLSRNVLFTVSNSLRFRLKVLTVLLYICLSVMLFYLFGSIMPLPPWGLVVGWVMALIAVELAYAFIFPYSFRYIADNDDDKMVILPV |
| IRIS_313-8946 | MLHHLKELAAVAGIHMILIYLCRFLLRRSRNVLFTVSNSLRFRLKVLTVLLYICLSVMLFYLFGSIMPLPPWGLVVGWVMALIAVELAYAFIFPYSFRYIADNDDDKMVILPV |
| IRIS_313-8948 | MLHHLKELAAVAGIHMILIYLCRFLLRRSRNVLFTVSNSLRFRLKVLTVLLYICLSVMLFYLFGSIMPLPPWGLVVGWVMALIAVELAYAFIFPYSFRYIADNDDDKMVILPV |
| IRIS_313-8955 | MLHQLKELAAVAGIHMILIYLCRFLLRRSRNVLFTVSNSLRFRLKVLTVLLYICLLVMLFYLFGSIMPLPPWGLVVGWVMALIAVELAYAFIFPYSFRYIADNDDDKMVILPV |
| IRIS_313-8956 | MLHHLKELAAVAGIHMILIYLCRFLLRRSRNVLFTVSNSLRFRLKVLTVLLYICLSVMLFYLFGSIMPLPPWGLVVGWVMALIAVELAYAFIFPYSFRYIADNDDDKMVILPV |
| IRIS_313-8957 | MLHHLKELAAVAGIHMILIYLCRFLLRRSRNVLFTVSNSLRFRLKVLTVLLYICLSVMLFYLFGSIMPLPPWGLVVGWVMALIAVELAYAFIFPYSFRYIADNDDDKMVILPV |
| IRIS_313-8960 | MLHHLKELAAVAGIHMILIYLCRFLLRRSRNVLFTVSNSLRFRLKVLTVLLYICLSVMLFYLFGSIMPLPPWGLVVGWVMALIAVELAYAFIFPYSFRYIADNDDDKMVILPV |
| IRIS_313-8963 | MLHHLKELAAVAGIHMILIYLCRFLLRRSRNVLFTVSNSLRFRLKVLTVLLYICLSVMLFYLFGSIMPLPPWGLVVGWVMALIAVELAYAFIFPYSFRYIADNDDDKMVILPV |
| IRIS_313-8967 | MLHHLKELAAVAGIHMILIYLCRFLLRRSRNVLFTVSNSLRFRLKVLTVLLYICLSVMLFYLFGSIMPLPPWGLVVGWVMALIAVELAYAFIFPYSFRYIADNDDDKMVILPV |
| IRIS_313-8968 | MLHHLKELAAVAGIHMILIYLCRFLLRRSRNVLFTVSNSLRFRLKVLTVLLYICLSVMLFYLFGSIMPLPPWGLVVGWVMALIAVELAYAFIFPYSFRYIADNDDDKMVILPV |
| IRIS_313-8976 | MLHHLKELAAVAGIHMILIYLCRFLLRRSRNVLFTVSNSLRFRLKVLTVLLYICLSVMLFYLFGSIMPLPPWGLVVGWVMALIAVELAYAFIFPYSFRYIADNDDDKMVILPV |
| IRIS_313-8978 | MLHHLKELAAVAGIHMILIYLCRFLLRRSRNVLFTVSNSLRFRLKVLTVLLYICLSVMLFYLFGSIMPLPPWGLVVGWVMALIAVELAYAFIFPYSFRYIADNDDDKMVILPV |

| | |
|---|---|
| IRIS_313-8980 | MLHHLKELAAVAGIHMILIYLCRFLLRRSRNVLFTVSNSLRFRLKVLTVLLYICLSVMLFYLFGSIMPLPPWGLVVGWVMALIAVELAYAFIFPYSFRYIADNDDDKMVILPV |
| IRIS_313-8982 | MLHHLKELAAVAGIHMILIYLCRFLLRRSRNVLFTVSNSLRFRLKVLTVLLYICLSVMLFYLFGSIMPLPPWGLVVGWVMALIAVELAYAFIFPYSFRYIADNDDDKMVILPV |
| IRIS_313-8985 | MLHHLKELAAVAGIHMILIYLCRFLLRRSRNVLFTVSNSLRFRLKVLTVLLYICLSVMLFYLFGSIMPLPPWGLVVGWVMALIAVELAYAFIFPYSFRYIADNDDDKMVILPV |
| IRIS_313-8986 | MLHHLKELAAVAGIHMILIYLCRFLLRLSRNVLFTVSNSLRFRLKVLTVLLYICLSVMLFYLFGSIMPLPPWGLVVGWVMALIAVELAYAFIFPYSFRYIADNDDDKMVILPV |
| IRIS_313-8987 | MLHHLKELAAVAGIHMILIYLCRFLLRRSRNVLFTVSNSLRFRLKVLTVLLYICLSVMLFYLFGSIMPLPPWGLVVGWVMALIAVELAYAFIFPYSFRYIADNDDDKMVILPV |
| IRIS_313-8988 | MLHHLKELAAVAGIHMILIYLCRFLLRRSRNVLFTVSNSLRFRLKVLTVLLYICLSVMLFYLFGSIMPLPPWGLVVGWVMALIAVELAYAFIFPYSFRYIADNDDDKMVILPV |
| IRIS_313-8994 | MLHHLKELAAVAGIHMILIYLCRFLLRRSRNVLFTVSNSLRFRLKVLTVLLYICLSVMLFYLFGSIMPLPPWGLVVGWVMALIAVELAYAFIFPYSFRYIADNDDDKMVILPV |
| IRIS_313-8996 | MLHHLKELAAVAGIHMILIYLCRFLLRRSRNVLFTVSNSLRFRLKVLTVLLYICLSVMLFYLFGSIMPLPPWGLVVGWVMALIAVELAYAFIFPYSFRYIADNDDDKMVILPV |
| IRIS_313-8998 | MLHHLKELAAVAGIHMILIYLCRFLLRRSRNVLFTVSNSLRFRLKVLTVLLYICLSVMLFYLFGSIMPLPPWGLVVGWVMALIAVELAYAFIFPYSFRYIADNDDDKMVILPV |
| IRIS_313-8999 | MLHHLKELAAVAGIHMILIYLCRFLLRRSRNVLFTVSNSLRFRLKVLTVLLYICLSVMLFYLFGSIMPLPPWGLVVGWVMALIAVELAYAFIFPYSFRYIADNDDDKMVILPV |
| IRIS_313-9002 | MLHHLKELAAVAGIHMILIYLCRFLLRRSRNVLFTVSNSLRFRLKVLTVLLYICLSVMLFYLFGSIMPLPPWGLVVGWVMALIAVELAYAFIFPYSFRYIADNDDDKMVILPV |
| IRIS_313-9005 | MLHHLKELAAVAGIHMILIYLCRFLLRRSRNVLFTVSNSLRFRLKVLTVLLYICLSVMLFYLFGSIMPLPPWGLVVGWVMALIAVELAYAFIFPYSFRYIADNDDDKMVILPV |
| IRIS_313-9006 | MLHHLKELAAVAGIHMILIYLCRFLLRRSRNVLFTVSNSLRFRLKVLTVLLYICLSVMLFYLFGSIMPLPPWGLVVGWVMALIAVELAYAFIFPYSFRYIADNDDDKMVILPV |
| IRIS_313-9017 | MLHHLKELAAVAGIHMILIYLCRFLLRRSRNVLFTVSNSLRFRLKVLTVLLYICLSVMLFYLFGSIMPLPPWGLVVGWVMALIAVELAYAFIFPYSFRYIADNDDDKMVILPV |
| IRIS_313-9019 | MLHHLKELAAVAGIHMILIYLCRFLLRRSRNVLFTVSNSLRFRLKVLTVLLYICLSVMLFYLFGSIMPLPPWGLVVGWVMALIAVELAYAFIFPYSFRYIADNDDDKMVILPV |
| IRIS_313-9020 | MLHHLKELAAVAGIHMILIYLCRFLLRRSRNVLFTVSNSLRFRLKVLTVLLYICLSVMLFYLFGSIMPLPPWGLVVGWVMALIAVELAYAFIFPYSFRYIADNDDDKMVILPV |
| IRIS_313-9023 | MLHHLKELAAVAGIHMILIYLCRFLLRRSRNVLFTVSNSLRFRLKVLTVLLYICLSVMLFYLFGSIMPLPPWGLVVGWVMALIAVELAYAFIFPYSFRYIADNDDDKMVILPV |
| IRIS_313-9032 | MLHHLKELAAVAGIHMILIYLCRFLLRRSRNVLFTVSNSLRFRLKVLTVLLYICLSVMLFYLFGSIMPLPPWGLVVGWVMALIAVELAYAFIFPYSFRYIADNDDDKMVILPV |
| IRIS_313-9039 | MLHHLKELAAVAGIHMILIYLCRFLLRRSRNVLFTVSNSLRFRLKVLTVLLYICLSVMLFYLFGSIMPLPPWGLVVGWVMALIAVELAYAFIFPYSFRYIADNDDDKMVILPV |
| IRIS_313-9044 | MLHHLKELAAVAGIHMILIYLCRFLLRRSRNVLFTVSNSLRFRLKVLTVLLYICLSVMLFYLFGSIMPLPPWGLVVGWVMALIAVELAYAFIFPYSFRYIADNDDDKMVILPV |
| IRIS_313-9048 | MLHHLKELAAVAGIHMILIYLCRFLLRRSRNVLFTVSNSLRFRLKVLTVLLYICLSVMLFYLFGSIMPLPPWGLVVGWVMALIAVELAYAFIFPYSFRYIADNDDDKMVILPV |
| IRIS_313-9049 | MLHHLKELAAVAGIHMILIYLCRFLLRRSRNVLFTVSNSLRFRLKVLTVLLYICLSVMLFYLFGSIMPLPPWGLVVGWVMALIAVELAYAFIFPYSFRYIADNDDDKMVILPV |
| IRIS_313-9050 | MLHHLKELAAVAGIHMILIYLCRFLLRRSRNVLFTVSNSLRFRLKVLTVLLYICLSVMLFYLFGSIMPLPPWGLVVGWVMALIAVELAYAFIFPYSFRYIADNDDDKMVILPV |
| IRIS_313-9053 | MLHHLKELAAVAGIHMILIYLCRFLLRRSRNVLFTVSNSLRFRLKVLTVLLYICLSVMLFYLFGSIMPLPPWGLVVGWVMALIAVELAYAFIFPYSFRYIADNDDDKMVILPV |
| IRIS_313-9054 | MLHHLKELAAVAGIHMILIYLCRFLLRRSRNVLFTVSNSLRFRLKVLTVLLYICLSVMLFYLFGSIMPLPPWGLVVGWVMALIAVELAYAFIFPYSFRYIADNDDDKMVILPV |
| IRIS_313-9062 | MLHHLKELAAVAGIHMILIYLCRFLLRRSRNVLFTVSNSLRFRLKVLTVLLYICLSVMLFYLFGSIMPLPPWGLVVGWVMALIAVELAYAFIFPYSFRYIADNDDDKMVILPV |
| IRIS_313-9065 | MLHHLKELAAVAGIHMILIYLCRFLLRRSRNVLFTVSNSLRFRLKVLTVLLYICLSVMLFYLFGSIMPLPPWGLVVGWVMALIAVELAYAFIFPYSFRYIADNDDDKMVILPV |
| IRIS_313-9066 | MLHHLKELAAVAGIHMILIYLCRFLLRRSRNVLFTVSNSLRFRLKVLTVLLYICLSVMLFYLFGSIMPLPPWGLVVGWVMALIAVELAYAFIFPYSFRYIADNDDDKMVILPV |
| IRIS_313-9067 | MLHHLKELAAVAGIHMILIYLCRFLLRRSRNVLFTVSNSLRFRLKVLTVLLYICLSVMLFYLFGSIMPLPPWGLVVGWVMALIAVELAYAFIFPYSFRYIADNDDDKMVILPV |
| IRIS_313-9070 | MLHHLKELAAVAGIHMILIYLCRFLLRRSRNVLFTVSNSLRFRLKVLTVLLYICLSVMLFYLFGSIMPLPPWGLVVGWVMALIAVELAYAFIFPYSFRYIADNDDDKMVILPV |
| IRIS_313-9072 | MLHHLKELAAVAGIHMILIYLCRFLLRRSRNVLFTVSNSLRFRLKVLTVLLYICLSVMLFYLFGSIMPLPPWGLVVGWVMALIAVELAYAFIFPYSFRYIADNDDDKMVILPV |
| IRIS_313-9081 | MLHHLKELAAVAGIHMILIYLCRFLLRRSRNVLFTVSNSLRFRLKVLTVLLYICLSVMLFYLFGSIMPLPPWGLVVGWVMALIAVELAYAFIFPYSFRYIADNDDDKMVILPV |
| IRIS_313-9083 | MLHHLKELAAVAGIHMILIYLCRFLLRRSRNVLFTVSNSLRFRLKVLTVLLYICLSVMLFYLFGSIMPLPPWGLVVGWVMALIAVELAYAFIFPYSFRYIADNDDDKMVILPV |
| IRIS_313-9091 | MLHHLKELAAVAGIHMILIYLCRFLLRRSRNVLFTVSNSLRFRLKVLTVLLYICLSVMLFYLFGSIMPLPPWGLVVGWVMALIAVELAYAFIFPYSFRYIADNDDDKMVILPV |
| IRIS_313-9097 | MLHHLKELAAVAGIHMILIYLCRFLLRRSRNVLFTVSNSLRFRLKVLTVLLYICLSVMLFYLFGSIMPLPPWGLVVGWVMALIAVELAYAFIFPYSFRYIADNDDDKMVILPV |
| IRIS_313-9098 | MLHHLKELAAVAGIHMILIYLCRFLLRRSRNVLFTVSNSLRFRLKVLTVLLYICLSVMLFYLFGSIMPLPPWGLVVGWVMALIAVELAYAFIFPYSFRYIADNDDDKMVILPV |
| IRIS_313-9101 | MLHQLKELAAVAGIHMILIYLCRFLLRRSRNVLFTVSNSLRFRLKVLTVLLYICLLVMLFYLFGSIMPLPPWGLVVGWVMALIAVELAYAFIFPYSFRYIADNDDDKMVILPV |
| IRIS_313-9102 | MLHHLKELAAVAGIHMILIYLCRFLLRRSRNVLFTVSNSLRFRLKVLTVLLYICLSVMLFYLFGSIMPLPPWGLVVGWVMALIAVELAYAFIFPYSFRYIADNDDDKMVILPV |
| IRIS_313-9108 | MLHHLKELAAVAGIHMILIYLCRFLLRRSRNVLFTVSNSLRFRLKVLTVLLYICLSVMLFYLFGSIMPLPPWGLVVGWVMALIAVELAYAFIFPYSFRYIADNDDDKMVILPV |
| IRIS_313-9111 | MLHHLKELAAVAGIHMILIYLCRFLLRRSRNVLFTVSNSLRFRLKVLTVLLYICLSVMLFYLFGSIMPLPPWGLVVGWVMALIAVELAYAFIFPYSFRYIADNDDDKMVILPV |
| IRIS_313-9112 | MLHHLKELAAVAGIHMILIYLCRFLLRRSRNVLFTVSNSLRFRLKVLTVLLYICLSVMLFYLFGSIMPLPPWGLVVGWVMALIAVELAYAFIFPYSFRYIADNDDDKMVILPV |
| IRIS_313-9114 | MLHHLKELAAVAGIHMILIYLCRFLLRRSRNVLFTVSNSLRFRLKVLTVLLYICLSVMLFYLFGSIMPLPPWGLVVGWVMALIAVELAYAFIFPYSFRYIADNDDDKMVILPV |
| IRIS_313-9115 | MLHHLKELAAVAGIHMILIYLCRFLLRRSRNVLFTVSNSLRFRLKVLTVLLYICLSVMLFYLFGSIMPLPPWGLVVGWVMALIAVELAYAFIFPYSFRYIADNDDDKMVILPV |
| IRIS_313-9116 | MLHHLKELAAVAGIHMILIYLCRFLLRRSRNVLFTVSNSLRFRLKVLTVLLYICLSVMLFYLFGSIMPLPPWGLVVGWVMALIAVELAYAFIFPYSFRYIADNDDDKMVILPV |
| IRIS_313-9117 | MLHHLKELAAVAGIHMILIYLCRFLLRRSRNVLFTVSNSLRFRLKVLTVLLYICLSVMLFYLFGSIMPLPPWGLVVGWVMALIAVELAYAFIFPYSFRYIADNDDDKMVILPV |
| IRIS_313-9119 | MLHHLKELAAVAGIHMILIYLCRFLLRRSRNVLFTVSNSLRFRLKVLTVLLYICLSVMLFYLFGSIMPLPPWGLVVGWVMALIAVELAYAFIFPYSFRYIADNDDDKMVILPV |
| IRIS_313-9120 | MLHHLKELAAVAGIHMILIYLCRFLLRRSRNVLFTVSNSLRFRLKVLTVLLYICLSVMLFYLFGSIMPLPPWGLVVGWVMALIAVELAYAFIFPYSFRYIADNDDDKMVILPV |
| IRIS_313-9121 | MLHHLKELAAVAGIHMILIYLCRFLLRRSRNVLFTVSNSLRFRLKVLTVLLYICLSVMLFYLFGSIMPLPPWGLVVGWVMALIAVELAYAFIFPYSFRYIADNDDDKMVILPV |
| IRIS_313-9123 | MLHHLKELAAVAGIHMILIYLCRFLLRRSRNVLFTVSNSLRFRLKVLTVLLYICLSVMLFYLFGSIMPLPPWGLVVGWVMALIAVELAYAFIFPYSFRYIADNDDDKMVILPV |
| IRIS_313-9129 | MLHHLKELAAVAGIHMILIYLCRFLLRRSRNVLFTVSNSLRFRLKVLTVLLYICLSVMLFYLFGSIMPLPPWGLVVGWVMALIAVELAYAFIFPYSFRYIADNDDDKMVILPV |
| IRIS_313-9131 | MLHHLKELAAVAGIHMILIYLCRFLLRRSRNVLFTVSNSLRFRLKVLTVLLYICLSVMLFYLFGSIMPLPPWGLVVGWVMALIAVELAYAFIFPYSFRYIADNDDDKMVILPV |
| IRIS_313-9137 | MLHHLKELAAVAGIHMILIYLCRFLLRRSRNVLFTVSNSLRFRLKVLTVLLYISLSAMLFYLFGSIMPLPPWGLVVGWVMALIAVELAYAFIFPYSFRYIADNDDDKMVILPV |
| IRIS_313-9139 | MLHHLKELAAVAGIHMILIYLCRFLLRRSRNVLFTVSNSLRFRLKVLTVLLYICLSVMLFYLFGSIMPLPPWGLVVGWVMALIAVELAYAFIFPYSFRYIADNDDDKMVILPV |
| IRIS_313-9140 | MLHHLKELAAVAGIHMILIYLCRFLLRRSRNVLFTVSNSLRFRLKVLTVLLYICLSVMLFYLFGSIMPLPPWGLVVGWVMALIAVELAYAFIFPYSFRYIADNDDDKMVILPV |
| IRIS_313-9148 | MLHHLKELAAVAGIHMILIYLCRFLLRRSRNVLFTVSNSLRFRLKVLTVLLYICLSVMLFYLFGSIMPLPPWGLVVGWVMALIAVELAYAFIFPYSFRYIADNDDDKMVILPV |
| IRIS_313-9156 | MLHHLKELAAVAGIHMILIYLCRFLLRRSRNVLFTVSNSLRFRLKVLTVLLYICLSVMLFYLFGSIMPLPPWGLVVGWVMALIAVELAYAFIFPYSFRYIADNDDDKMVILPV |
| IRIS_313-9160 | MLHHLKELAAVAGIHMILIYLCRFLLRRSRNVLFTVSNSLRFRLKVLTVLLYICLSVMLFYLFGSIMPLPPWGLVVGWVMALIAVELAYAFIFPYSFRYIADNDDDKMVILPV |
| IRIS_313-9170 | MLHHLKELAAVAGIHMILIYLCRFLLRRSRNVLFTVSNSLRFRLKVLTVLLYICLSVMLFYLFGSIMPLPPWGLVVGWVMALIAVELAYAFIFPYSFRYIADNDDDKMVILPV |
| IRIS_313-9172 | MLHHLKELAAVAGIHMILIYLCRFLLRRSRNVLFTVSNSLRFRLKVLTVLLYICLSVMLFYLFGSIMPLPPWGLVVGWVMALIAVELAYAFIFPYSFRYIADNDDDKMVILPV |
| IRIS_313-9174 | MLHHLKELAAVAGIHMILIYLCRFLLRRSRNVLFTVSNSLRFRLKVLTVLLYICLSVMLFYLFGSIMPLPPWGLVVGWVMALIAVELAYAFIFPYSFRYIADNDDDKMVILPV |
| IRIS_313-9176 | MLHHLKELAAVAGIHMILIYLCRFLLRRSRNVLFTVSNSLRFRLKVLTVLLYICLSVMLFYLFGSIMPLPPWGLVVGWVMALIAVELAYAFIFPYSFRYIADNDDDKMVILPV |
| IRIS_313-9182 | MLHHLKELAAVAGIHMILIYLCRFLLRRSRNVLFTVSNSLRFRLKVLTVLLYICLSVMLFYLFGSIMPLPPWGLVVGWVMALIAVELAYAFIFPYSFRYIADNDDDKMVILPV |
| IRIS_313-9187 | MLHHLKELAAVAGIHMILIYLCRFLLRRSRNVLFTVSNSLRFRLKVLTVLLYICLSVMLFYLFGSIMPLPPWGLVVGWVMALIAVELAYAFIFPYSFRYIADNDDDKMVILPV |
| IRIS_313-9188 | MLHHLKELAAVAGIHMILIYLCRFLLRRSRNVLFTVSNSLRFRLKVLTVLLYICLSVMLFYLFGSIMPLPPWGLVVGWVMALIAVELAYAFIFPYSFRYIADNDDDKMVILPV |
| IRIS_313-9190 | MLHHLKELAAVAGIHMILIYLCRFLLRRSRNVLFTVSNSLRFRLKVLTVLLYICLSVMLFYLFGSIMPLPPWGLVVGWVMALIAVELAYAFIFPYSFRYIADNDDDKMVILPV |
| IRIS_313-9193 | MLHHLKELAAVAGIHMILIYLCRFLLRRSRNVLFTVSNSLRFRLKVLTVLLYICLSVMLFYLFGSIMPLPPWGLVVGWVMALIAVELAYAFIFPYSFRYIADNDDDKMVILPV |
| IRIS_313-9197 | MLHHLKELAAVAGIHMILIYLCRFLLRRSRNVLFTVSNSLRFRLKVLTVLLYICLSVMLFYLFGSIMPLPPWGLVVGWVMALIAVELAYAFIFPYSFRYIADNDDDKMVILPV |
| IRIS_313-9198 | MLHHLKELAAVAGIHMILIYLCRFLLRRSRNVLFTVSNSLRFRLKVLTVLLYICLSVMLFYLFGSIMPLPPWGLVVGWVMALIAVELAYAFIFPYSFRYIADNDDDKMVILPV |
| IRIS_313-9201 | MLHHLKELAAVAGIHMILIYLCRFLLRRSRNVLFTVSNSLRFRLKVLTVLLYICLSVMLFYLFGSIMPLPPWGLVVGWVMALIAVELAYAFIFPYSFRYIADNDDDKMVILPV |
| IRIS_313-9204 | MLHHLKELAAVAGIHMILIYLCRFLLRRSRNVLFTVSNSLRFRLKVLTVLLYICLSVMLFYLFGSIMPLPPWGLVVGWVMALIAVELAYAFIFPYSFRYIADNDDDKMVILPV |
| IRIS_313-9208 | MLHHLKELAAVAGIHMILIYLCRFLLRRSRNVLFTVSNSLRFRLKVLTVLLYICLSVMLFYLFGSIMPLPPWGLVVGWVMALIAVELAYAFIFPYSFRYIADNDDDKMVILPV |
| IRIS_313-9209 | MLHHLKELAAVAGIHMILIYLCRFLLRRSRNVLFTVSNSLRFRLKVLTVLLYICLSVMLFYLFGSIMPLPPWGLVVGWVMALIAVELAYAFIFPYSFRYIADNDDDKMVILPV |
| IRIS_313-9210 | MLHHLKELAAVAGIHMILIYLCRFLLRRSRNVLFTVSNSLRFRLKVLTVLLYICLSVMLFYLFGSIMPLPPWGLVVGWVMALIAVELAYAFIFPYSFRYIADNDDDKMVILPV |
| IRIS_313-9218 | MLHHLKELAAVAGIHMILIYLCRFLLRRSRNVLFTVSNSLRFRLKVLTVLLYICLSVMLFYLFGSIMPLPPWGLVVGWVMALIAVELAYAFIFPYSFRYIADNDDDKMVILPV |
| IRIS_313-9227 | MLHHLKELAAVAGIHMILIYLCRFLLRRSRNVLFTVSNSLRFRLKVLTVLLYICLSVMLFYLFGSIMPLPPWGLVVGWVMALIAVELAYAFIFPYSFRYIADNDDDKMVILPV |
| IRIS_313-9228 | MLHHLKELAAVAGIHMILIYLCRFLLRRSRNVLFTVSNSLRFRLKVLTVLLYICLSVMLFYLFGSIMPLPPWGLVVGWVMALIAVELAYAFIFPYSFRYIADNDDDKMVILPV |
| IRIS_313-9239 | MLHHLKELAAVAGIHMILIYLCRFLLRRSRNVLFTVSNSLRFRLKVLTVLLYICLSVMLFYLFGSIMPLPPWGLVVGWVMALIAVELAYAFIFPYSFRYIADNDDDKMVILPV |
| IRIS_313-9242 | MLHHLKELAAVAGIHMILIYLCRFLLRRSRNVLFTVSNSLRFRLKVLTVLLYICLSVMLFYLFGSIMPLPPWGLVVGWVMALIAVELAYAFIFPYSFRYIADNDDDKMVILPV |
| IRIS_313-9243 | MLHHLKELAAVAGIHMILIYLCRFLLRRSRNVLFTVSNSLRFRLKVLTVLLYICLSVMLFYLFGSIMPLPPWGLVVGWVMALIAVELAYAFIFPYSFRYIADNDDDKMVILPV |
| IRIS_313-9249 | MLHHLKELAAVAGIHMILIYLCRFLLRRSRNVLFTVSNSLRFRLKVLTVLLYICLSVMLFYLFGSIMPLPPWGLVVGWVMALIAVELAYAFIFPYSFRYIADNDDDKMVILPV |
| IRIS_313-9251 | MLHHLKELAAVAGIHMILIYLCRFLLRRSRNVLFTVSNSLRFRLKVLTVLLYICLSVMLFYLFGSIMPLPPWGLVVGWVMALIAVELAYAFIFPYSFRYIADNDDDKMVILPV |
| IRIS_313-9253 | MLHHLKELAAVAGIHMILIYLCRFLLRRSRNVLFTVSNSLRFRLKVLTVLLYICLSVMLFYLFGSIMPLPPWGLVVGWVMALIAVELAYAFIFPYSFRYIADNDDDKMVILPV |
| IRIS_313-9256 | MLHHLKELAAVAGIHMILIYLCRFLLRRSRNVLFTVSNSLRFRLKVLTVLLYICLSVMLFYLFGSIMPLPPWGLVVGWVMALIAVELAYAFIFPYSFRYIADNDDDKMVILPV |
| IRIS_313-9258 | MLHHLKELAAVAGIHMILIYLCRFLLRRSRNVLFTVSNSLRFRLKVLTVLLYICLSVMLFYLFGSIMPLPPWGLVVGWVMALIAVELAYAFIFPYSFRYIADNDDDKMVILPV |
| IRIS_313-9259 | MLHHLKELAAVAGIHMILIYLCRFLLRRSRNVLFTVSNSLRFRLKVLTVLLYICLSVMLFYLFGSIMPLPPWGLVVGWVMALIAVELAYAFIFPYSFRYIADNDDDKMVILPV |
| IRIS_313-9262 | MLHHLKELAAVAGIHMILIYLCRFLLRRSRNVLFTVSNSLRFRLKVLTVLLYICLSVMLFYLFGSIMPLPPWGLVVGWVMALIAVELAYAFIFPYSFRYIADNDDDKMVILPV |
| IRIS_313-9267 | MLHHLKELAAVAGIHMILIYLCRFLLRRSRNVLFTVSNSLRFRLKVLTVLLYICLSVMLFYLFGSIMPLPPWGLVVGWVMALIAVELAYAFIFPYSFRYIADNDDDKMVILPV |
| IRIS_313-9271 | MLHHLKELAAVAGIHMILIYLCRFLLRRSRNVLFTVSNSLRFRLKVLTVLLYICLSVMLFYLFGSIMPLPPWGLVVGWVMALIAVELAYAFIFPYSFRYIADNDDDKMVILPV |
| IRIS_313-9273 | MLHHLKELAAVAGIHMILIYLCRFLLRRSRNVLFTVSNSLRFRLKVLTVLLYICLSVMLFYLFGSIMPLPPWGLVVGWVMALIAVELAYAFIFPYSFRYIADNDDDKMVILPV |
| IRIS_313-9281 | MLHHLKELAAVAGIHMILIYLCRFLLRRSRNVLFTVSNSLRFRLKVLTVLLYICLSVMLFYLFGSIMPLPPWGLVVGWVMALIAVELAYAFIFPYSFRYIADNDDDKMVILPV |
| IRIS_313-9283 | MLHHLKELAAVAGIHMILIYLCRFLLRRSRNVLFTVSNSLRFRLKVLTVLLYICLSVMLFYLFGSIMPLPPWGLVVGWVMALIAVELAYAFIFPYSFRYIADNDDDKMVILPV |
| IRIS_313-9285 | MLHHLKELAAVAGIHMILIYLCRFLLRRSRNVLFTVSNSLRFRLKVLTVLLYICLSVMLFYLFGSIMPLPPWGLVVGWVMALIAVELAYAFIFPYSFRYIADNDDDKMVILPV |
| IRIS_313-9286 | MLHHLKELAAVAGIHMILIYLCRFLLRRSRNVLFTVSNSLRFRLKVLTVLLYICLSVMLFYLFGSIMPLPPWGLVVGWVMALIAVELAYAFIFPYSFRYIADNDDDKMVILPV |
| IRIS_313-9287 | MLHHLKELAAVAGIHMILIYLCRFLLRRSRNVLFTVSNSLRFRLKVLTVLLYICLSVMLFYLFGSIMPLPPWGLVVGWVMALIAVELAYAFIFPYSFRYIADNDDDKMVILPV |
| IRIS_313-9288 | MLHHLKELAAVAGIHMILIYLCRFLLRRSRNVLFTVSNSLRFRLKVLTVLLYICLSVMLFYLFGSIMPLPPWGLVVGWVMALIAVELAYAFIFPYSFRYIADNDDDKMVILPV |
| IRIS_313-9294 | MLHHLKELAAVAGIHMILIYLCRFLLRRSRNVLFTVSNSLRFRLKVLTVLLYICLSVMLFYLFGSIMPLPPWGLVVGWVMALIAVELAYAFIFPYSFRYIADNDDDKMVILPV |
| IRIS_313-9297 | MLHHLKELAAVAGIHMILIYLCRFLLRRSRNVLFTVSNSLRFRLKVLTVLLYICLSVMLFYLFGSIMPLPPWGLVVGWVMALIAVELAYAFIFPYSFRYIADNDDDKMVILPV |
| IRIS_313-9301 | MLHHLKELAAVAGIHMILIYLCRFLLRRSRNVLFTVSNSLRFRLKVLTVLLYICLSVMLFYLFGSIMPLPPWGLVVGWVMALIAVELAYAFIFPYSFRYIADNDDDKMVILPV |
| IRIS_313-9302 | MLHHLKELAAVAGIHMILIYLCRFLLRRSRNVLFTVSNSLRFRLKVLTVLLYICLSVMLFYLFGSIMPLPPWGLVVGWVMALIAVELAYAFIFPYSFRYIADNDDDKMVILPV |
| IRIS_313-9310 | MLHHLKELAAVAGIHMILIYLCRFLLRRSRNVLFTVSNSLRFRLKVLTVLLYICLSVMLFYLFGSIMPLPPWGLVVGWVMALIAVELAYAFIFPYSFRYIADNDDDKMVILPV |
| IRIS_313-9313 | MLHHLKELAAVAGIHMILIYLCRFLLRRSRNVLFTVSNSLRFRLKVLTVLLYICLSVMLFYLFGSIMPLPPWGLVVGWVMALIAVELAYAFIFPYSFRYIADNDDDKMVILPV |
| IRIS_313-9314 | MLHHLKELAAVAGIHMILIYLCRFLLRRSRNVLFTVSNSLRFRLKVLTVLLYICLSVMLFYLFGSIMPLPPWGLVVGWVMALIAVELAYAFIFPYSFRYIADNDDDKMVILPV |
| IRIS_313-9317 | MLHHLKELAAVAGIHMILIYLCRFLLRRSRNVLFTVSNSLRFRLKVLTVLLYICLSVMLFYLFGSIMPLPPWGLVVGWVMALIAVELAYAFIFPYSFRYIADNDDDKMVILPV |
| IRIS_313-9320 | MLHHLKELAAVAGIHMILIYLCRFLLRRSRNVLFTVSNSLRFRLKVLTVLLYICLSVMLFYLFGSIMPLPPWGLVVGWVMALIAVELAYAFIFPYSFRYIADNDDDKMVILPV |
| IRIS_313-9324 | MLHHLKELAAVAGIHMILIYLCRFLLRLRRNVLFTVSNSLRFRLKVLTVLLYICLSVMLFYLFGSIMPLPPWGLVVGWVMALIAVELAYAFIFPYSFRYIADNDDDKMVILPV |
| IRIS_313-9325 | MLHHLKELAAVAGIHMILIYLCRFLLRRSRNVLFTVSNSLRFRLKVLTVLLYICLSVMLFYLFGSIMPLPPWGLVVGWVMALIAVELAYAFIFPYSFRYIADNDDDKMVILPV |
| IRIS_313-9329 | MLHHLKELAAVAGIHMILIYLCRFLLRRSRNVLFTVSNSLRFRLKVLTVLLYICLSVMLFYLFGSIMPLPPWGLVVGWVMALIAVELAYAFIFPYSFRYIADNDDDKMVILPV |
| IRIS_313-9342 | MLHHLKELAAVAGIHMILIYLCRFLLRRSRNVLFTVSNSLRFRLKVLTVLLYICLSVMLFYLFGSIMPLPPWGLVVGWVMALIAVELAYAFIFPYSFRYIADNDDDKMVILPV |
| IRIS_313-9346 | MLHHLKELAAVAGIHMILIYLCRFLLRRSRNVLFTVSNSLRFRLKVLTVLLYICLSVMLFYLFGSIMPLPPWGLVVGWVMALIAVELAYAFIFPYSFRYIADNDDDKMVILPV |
| IRIS_313-9347 | MLHHLKELAAVAGIHMILIYLCRFLLRRSRNVLFTVSNSLRFRLKVLTVLLYICLSVMLFYLFGSIMPLPPWGLVVGWVMALIAVELAYAFIFPYSFRYIADNDDDKMVILPV |
| IRIS_313-9348 | MLHHLKELAAVAGIHMILIYLCRFLLRRSRNVLFTVSNSLRFRLKVLTVLLYICLSVMLFYLFGSIMPLPPWGLVVGWVMALIAVELAYAFIFPYSFRYIADNDDDKMVILPV |
| IRIS_313-9351 | MLHHLKELAAVAGIHMILIYLCRFLLRRSRNVLFTVSNSLRFRLKVLTVLLYICLSVMLFYLFGSIMPLPPWGLVVGWVMALIAVELAYAFIFPYSFRYIADNDDDKMVILPV |
| IRIS_313-9357 | MLHHLKELAAVAGIHMILIYLCRFLLRRSRNVLFTVSNSLRFRLKVLTVLLYICLSVMLFYLFGSIMPLPPWGLVVGWVMALIAVELAYAFIFPYSFRYIADNDDDKMVILPV |
| IRIS_313-9360 | MLHHLKELAAVAGIHMILIYLCRFLLRRSRNVLFTVSNSLRFRLKVLTVLLYICLSVMLFYLFGSIMPLPPWGLVVGWVMALIAVELAYAFIFPYSFRYIADNDDDKMVILPV |
| IRIS_313-9363 | MLHHLKELAAVAGIHMILIYLCRFLLRRSRNVLFTVSNSLRFRLKVLTVLLYICLSVMLFYLFGSIMPLPPWGLVVGWVMALIAVELAYAFIFPYSFRYIADNDDDKMVILPV |
| IRIS_313-9366 | MLHHLKELAAVAGIHMILIYLCRFLLRRSRNVLFTVSNSLRFRLKVLTVLLYICLSVMLFYLFGSIMPLPPWGLVVGWVMALIAVELAYAFIFPYSFRYIADNDDDKMVILPV |
| IRIS_313-9368 | MLHHLKELAAVAGIHMILIYLCRFLLRRSRNVLFTVSNSLRFRLKVLTVLLYICLSVMLFYLFGSIMPLPPWGLVVGWVMALIAVELAYAFIFPYSFRYIADNDDDKMVILPV |
| IRIS_313-9372 | MLHHLKELAAVAGIHMILIYLCRFLLRRSRNVLFTVSNSLRFRLKVLTVLLYICLSVMLFYLFGSIMPLPPWGLVVGWVMALIAVELAYAFIFPYSFRYIADNDDDKMVILPV |
| IRIS_313-9375 | MLHHLKELAAVAGIHMILIYLCRFLLRRSRNVLFTVSNSLRFRLKVLTVLLYICLSVMLFYLFGSIMPLPPWGLVVGWVMALIAVELAYAFIFPYSFRYIADNDDDKMVILPV |
| IRIS_313-9379 | MLHHLKELAAVAGIHMILIYLCRFLLRRSRNVLFTVSNSLRFRLKVLTVLLYICLSVMLFYLFGSIMPLPPWGLVVGWVMALIAVELAYAFIFPYSFRYIADNDDDKMVILPV |
| IRIS_313-9384 | MLHHLKELAAVAGIHMILIYLCRFLLRRSRNVLFTVSNSLRFRLKVLTVLLYICLSVMLFYLFGSIMPLPPWGLVVGWVMALIAVELAYAFIFPYSFRYIADNDDDKMVILPV |
| IRIS_313-9388 | MLHHLKELAAVAGIHMILIYLCRFLLRRSRNVLFTVSNSLRFRLKVLTVLLYICLSVMLFYLFGSIMPLPPWGLVVGWVMALIAVELAYAFIFPYSFRYIADNDDDKMVILPV |
| IRIS_313-9389 | MLHHLKELAAVAGIHMILIYLCRFLLRRSRNVLFTVSNSLRFRLKVLTVLLYICLSVMLFYLFGSIMPLPPWGLVVGWVMALIAVELAYAFIFPYSFRYIADNDDDKMVILPV |

| | |
|---|---|
| IRIS_313-9391 | MLHHLKELAAVAGIHMILIYLCRFLLRRSRNVLFTVSNSLRFRLKVLTVLLYICLSVMLFYLFGSIMPLPPWGLVVGWVMALIAVELAYAFIFPYSFRYIADNDDDKMVILPV |
| IRIS_313-9392 | MLHHLKELAAVAGIHMILIYLCRFLLRRSRNVLFTVSNSLRFRLKVLTVLLYICLSVMLFYLFGSIMPLPPWGLVVGWVMALIAVELAYAFIFPYSFRYIADNDDDKMVILPV |
| IRIS_313-9397 | MLHHLKELAAVAGIHMILIYLCRFLLRRSRNVLFTVSNSLRFRLKVLTVLLYICLSVMLFYLFGSIMPLPPWGLVVGWVMALIAVELAYAFIFPYSFRYIADNDDDKMVILPV |
| IRIS_313-9400 | MLHHLKELAAVAGIHMILIYLCRFLLRRSRNVLFTVSNSLRFRLKVLTVLLYICLSVMLFYLFGSIMPLPPWGLVVGWVMALIAVELAYAFIFPYSFRYIADNDDDKMVILPV |
| IRIS_313-9402 | MLHHLKELAAVAGIHMILIYLCRFLLRRSRNVLFTVSNSLRFRLKVLTVLLYICLSVMLFYLFGSIMPLPPWGLVVGWVMALIAVELAYAFIFPYSFRYIADNDDDKMVILPV |
| IRIS_313-9403 | MLHHLKELAAVAGIHMILIYLCRFLLRRSRNVLFTVSNSLRFRLKVLTVLLYICLSVMLFYLFGSIMPLPPWGLVVGWVMALIAVELAYAFIFPYSFRYIADNDDDKMVILPV |
| IRIS_313-9405 | MLHHLKELAAVAGIHMILIYLCRFLLRRSRNVLFTVSNSLRFRLKVLTVLLYICLSVMLFYLFGSIMPLPPWGLVVGWVMALIAVELAYAFIFPYSFRYIADNDDDKMVILPV |
| IRIS_313-9406 | MLHHLKELAAVAGIHMILIYLCRFLLRRSRNVLFTVSNSLRFRLKVLTVLLYICLSVMLFYLFGSIMPLPPWGLVVGWVMALIAVELAYAFIFPYSFRYIADNDDDKMVILPV |
| IRIS_313-9409 | MLHHLKELAAVAGIHMILIYLCRFLLRRSRNVLFTVSNSLRFRLKVLTVLLYICLSVMLFYLFGSIMPLPPWGLVVGWVMALIAVELAYAFIFPYSFRYIADNDDDKMVILPV |
| IRIS_313-9410 | MLHHLKELAAVAGIHMILIYLCRFLLRRSRNVLFTVSNSLRFRLKVLTVLLYICLSVMLFYLFGSIMPLPPWGLVVGWVMALIAVELAYAFIFPYSFRYIADNDDDKMVILPV |
| IRIS_313-9415 | MLHHLKELAAVAGIHMILIYLCRFLLRRSRNVLFTVSNSLRFRLKVLTVLLYICLSVMLFYLFGSIMPLPPWGLVVGWVMALIAVELAYAFIFPYSFRYIADNDDDKMVILPV |
| IRIS_313-9422 | MLHHLKELAAVAGIHMILIYLCRFLLRRSRNVLFTVSNSLRFRLKVLTVLLYICLSVMLFYLFGSIMPLPPWGLVVGWVMALIAVELAYAFIFPYSFRYIADNDDDKMVILPV |
| IRIS_313-9423 | MLHHLKELAAVAGIHMILIYLCRFLLRRSRNVLFTVSNSLRFRLKVLTVLLYICLSVMLFYLFGSIMPLPPWGLVVGWVMALIAVELAYAFIFPYSFRYIADNDDDKMVILPV |
| IRIS_313-9424 | MLHHLKELAAVAGIHMILIYLCRFLLRRSRNVLFTVSNSLRFRLKVLTVLLYICLSVMLFYLFGSIMPLPPWGLVVGWVMALIAVELAYAFIFPYSFRYIADNDDDKMVILPV |
| IRIS_313-9427 | MLHHLKELAAVAGIHMILIYLCRFLLRRSRNVLFTVSNSLRFRLKVLTVLLYICLSVMLFYLFGSIMPLPPWGLVVGWVMALIAVELAYAFIFPYSFRYIADNDDDKMVILPV |
| IRIS_313-9429 | MLHHLKELAAVAGIHMILIYLCRFLLRRSRNVLFTVSNSLRFRLKVLTVLLYICLSVMLFYLFGSIMPLPPWGLVVGWVMALIAVELAYAFIFPYSFRYIADNDDDKMVILPV |
| IRIS_313-9433 | MLHHLKELAAVAGIHMILIYLCRFLLRRSRNVLFTVSNSLRFRLKVLTVLLYICLSVMLFYLFGSIMPLPPWGLVVGWVMALIAVELAYAFIFPYSFRYIADNDDDKMVILPV |
| IRIS_313-9438 | MLHHLKELAAVAGIHMILIYLCRFLLRRSRNVLFTVSNSLRFRLKVLTVLLYICLSVMLFYLFGSIMPLPPWGLVVGWVMALIAVELAYAFIFPYSFRYIADNDDDKMVILPV |
| IRIS_313-9445 | MLHHLKELAAVAGIHMILIYLCRFLLRRSRNVLFTVSNSLRFRLKVLTVLLYICLSVMLFYLFGSIMPLPPWGLVVGWVMALIAVELAYAFIFPYSFRYIADNDDDKMVILPV |
| IRIS_313-9449 | MLHHLKELAAVAGIHMILIYLCRFLLRRSRNVLFTVSNSLRFRLKVLTVLLYICLSVMLFYLFGSIMPLPPWGLVVGWVMALIAVELAYAFIFPYSFRYIADNDDDKMVILPV |
| IRIS_313-9451 | MLHHLKELAAVAGIHMILIYLCRFLLRRSRNVLFTVSNSLRFRLKVLTVLLYICLSVMLFYLFGSIMPLPPWGLVVGWVMALIAVELAYAFIFPYSFRYIADNDDDKMVILPV |
| IRIS_313-9452 | MLHHLKELAAVAGIHMILIYLCRFLLRRSRNVLFTVSNSLRFRLKVLTVLLYICLSVMLFYLFGSIMPLPPWGLVVGWVMALIAVELAYAFIFPYSFRYIADNDDDKMVILPV |
| IRIS_313-9461 | MLHHLKELAAVAGIHMILIYLCRFLLRRSRNVLFTVSNSLRFRLKVLTVLLYICLSVMLFYLFGSIMPLPPWGLVVGWVMALIAVELAYAFIFPYSFRYIADNDDDKMVILPV |
| IRIS_313-9463 | MLHHLKELAAVAGIHMILIYLCRFLLRRSRNVLFTVSNSLRFRLKVLTVLLYICLSVMLFYLFGSIMPLPPWGLVVGWVMALIAVELAYAFIFPYSFRYIADNDDDKMVILPV |
| IRIS_313-9464 | MLHHLKELAAVAGIHMILIYLCRFLLRRSRNVLFTVSNSLRFRLKVLTVLLYICLSVMLFYLFGSIMPLPPWGLVVGWVMALIAVELAYAFIFPYSFRYIADNDDDKMVILPV |
| IRIS_313-9468 | MLHHLKELAAVAGIHMILIYLCRFLLRRSRNVLFTVSNSLRFRLKVLTVLLYICLSVMLFYLFGSIMPLPPWGLVVGWVMALIAVELAYAFIFPYSFRYIADNDDDKMVILPV |
| IRIS_313-9469 | MLHHLKELAAVAGIHMILIYLCRFLLRRSRNVLFTVSNSLRFRLKVLTVLLYICLSVMLFYLFGSIMPLPPWGLVVGWVMALIAVELAYAFIFPYSFRYIADNDDDKMVILPV |
| IRIS_313-9470 | MLHQLKELAAVAGIHMILIYLCRFLLRRSRNVLFTVSNSLRFRLKVLTVLLYICLLVMLFYLFGSIMPLPPWGLVVGWVMALIAVELAYAFIFPYSFRYIADNDDDKMVILPV |
| IRIS_313-9472 | MLHHLKELAAVAGIHMILIYLCRFLLRRSRNVLFTVSNSLRFRLKVLTVLLYICLSVMLFYLFGSIMPLPPWGLVVGWVMALIAVELAYAFIFPYSFRYIADNDDDKMVILPV |
| IRIS_313-9482 | MLHHLKELAAVAGIHMILIYLCRFLLRRSRNVLFTVSNSLRFRLKVLTVLLYICLSVMLFYLFGSIMPLPPWGLVVGWVMALIAVELAYAFIFPYSFRYIADNDDDKMVILPV |
| IRIS_313-9484 | MLHHLKELAAVAGIHMILIYLCRFLLRRSRNVLFTVSNSLRFRLKVLTVLLYICLSVMLFYLFGSIMPLPPWGLVVGWVMALIAVELAYAFIFPYSFRYIADNDDDKMVILPV |
| IRIS_313-9491 | MLHHLKELAAVAGIHMILIYLCRFLLRRSRNVLFTVSNSLRFRLKVLTVLLYICLSVMLFYLFGSIMPLPPWGLVVGWVMALIAVELAYAFIFPYSFRYIADNDDDKMVILPV |
| IRIS_313-9492 | MLHHLKELAAVAGIHMILIYLCRFLLRRSRNVLFTVSNSLRFRLKVLTVLLYICLSVMLFYLFGSIMPLPPWGLVVGWVMALIAVELAYAFIFPYSFRYIADNDDDKMVILPV |
| IRIS_313-9503 | MLHHLKELAAVAGIHMILIYLCRFLLRRSRNVLFTVSNSLRFRLKVLTVLLYICLSVMLFYLFGSIMPLPPWGLVVGWVMALIAVELAYAFIFPYSFRYIADNDDDKMVILPV |
| IRIS_313-9505 | MLHHLKELAAVAGIHMILIYLCRFLLRRSRNVLFTVSNSLRFRLKVLTVLLYICLSVMLFYLFGSIMPLPPWGLVVGWVMALIAVELAYAFIFPYSFRYIADNDDDKMVILPV |
| IRIS_313-9506 | MLHHLKELAAVAGIHMILIYLCRFLLRRSRNVLFTVSNSLRFRLKVLTVLLYICLSVMLFYLFGSIMPLPPWGLVVGWVMALIAVELAYAFIFPYSFRYIADNDDDKMVILPV |
| IRIS_313-9516 | MLHHLKELAAVAGIHMILIYLCRFLLRRSRNVLFTVSNSLRFRLKVLTVLLYICLSVMLFYLFGSIMPLPPWGLVVGWVMALIAVELAYAFIFPYSFRYIADNDDDKMVILPV |
| IRIS_313-9519 | MLHHLKELAAVAGIHMILIYLCRFLLRRSRNVLFTVSNSLRFRLKVLTVLLYICLSVMLFYLFGSIMPLPPWGLVVGWVMALIAVELAYAFIFPYSFRYIADNDDDKMVILPV |
| IRIS_313-9522 | MLHHLKELAAVAGIHMILIYLCRFLLRRSRNVLFTVSNSLRFRLKVLTVLLYICLSVMLFYLFGSIMPLPPWGLVVGWVMALIAVELAYAFIFPYSFRYIADNDDDKMVILPV |
| IRIS_313-9523 | MLHHLKELAAVAGIHMILIYLCRFLLRRSRNVLFTVSNSLRFRLKVLTVLLYICLSVMLFYLFGSIMPLPPWGLVVGWVMALIAVELAYAFIFPYSFRYIADNDDDKMVILPV |
| IRIS_313-9529 | MLHHLKELAAVAGIHMILIYLCRFLLRRSRNVLFTVSNSLRFRLKVLTVLLYICLSVMLFYLFGSIMPLPPWGLVVGWVMALIAVELAYAFIFPYSFRYIADNDDDKMVILPV |
| IRIS_313-9533 | MLHHLKELAAVAGIHMILIYLCRFLLRRSRNVLFTVSNSLRFRLKVLTVLLYICLSVMLFYLFGSIMPLPPWGLVVGWVMALIAVELAYAFIFPYSFRYIADNDDDKMVILPV |
| IRIS_313-9539 | MLHHLKELAAVAGIHMILIYLCRFLLRRSRNVLFTVSNSLRFRLKVLTVLLYICLSVMLFYLFGSIMPLPPWGLVVGWVMALIAVELAYAFIFPYSFRYIADNDDDKMVILPV |
| IRIS_313-9547 | MLHHLKELAAVAGIHMILIYLCRFLLRRSRNVLFTVSNSLRFRLKVLTVLLYICLSVMLFYLFGSIMPLPPWGLVVGWVMALIAVELAYAFIFPYSFRYIADNDDDKMVILPV |
| IRIS_313-9550 | MLHHLKELAAVAGIHMILIYLCRFLLRRSRNVLFTVSNSLRFRLKVLTVLLYICLSVMLFYLFGSIMPLPPWGLVVGWVMALIAVELAYAFIFPYSFRYIADNDDDKMVILPV |
| IRIS_313-9551 | MLHHLKELAAVAGIHMILIYLCRFLLRRSRNVLFTVSNSLRFRLKVLTVLLYICLSVMLFYLFGSIMPLPPWGLVVGWVMALIAVELAYAFIFPYSFRYIADNDDDKMVILPV |
| IRIS_313-9555 | MLHHLKELAAVAGIHMILIYLCRFLLRRSRNVLFTVSNSLRFRLKVLTVLLYICLSVMLFYLFGSIMPLPPWGLVVGWVMALIAVELAYAFIFPYSFRYIADNDDDKMVILPV |
| IRIS_313-9557 | MLHHLKELAAVAGIHMILIYLCRFLLRRSRNVLFTVSNSLRFRLKVLTVLLYICLSVMLFYLFGSIMPLPPWGLVVGWVMALIAVELAYAFIFPYSFRYIADNDDDKMVILPV |
| IRIS_313-9558 | MLHHLKELAAVAGIHMILIYLCRFLLRRSRNVLFTVSNSLRFRLKVLTVLLYICLSVMLFYLFGSIMPLPPWGLVVGWVMALIAVELAYAFIFPYSFRYIADNDDDKMVILPV |
| IRIS_313-9560 | MLHHLKELAAVAGIHMILIYLCRFLLRRSRNVLFTVSNSLRFRLKVLTVLLYICLSVMLFYLFGSIMPLPPWGLVVGWVMALIAVELAYAFIFPYSFRYIADNDDDKMVILPV |
| IRIS_313-9566 | MLHHLKELAAVAGIHMILIYLCRFLLRRSRNVLFTVSNSLRFRLKVLTVLLYICLSVMLFYLFGSIMPLPPWGLVVGWVMALIAVELAYAFIFPYSFRYIADNDDDKMVILPV |
| IRIS_313-9568 | MLHHLKELAAVAGIHMILIYLCRFLLRRSRNVLFTVSNSLRFRLKVLTVLLYICLSVMLFYLFGSIMPLPPWGLVVGWVMALIAVELAYAFIFPYSFRYIADNDDDKMVILPV |
| IRIS_313-9570 | MLHHLKELAAVAGIHMILIYLCRFLLRRSRNVLFTVSNSLRFRLKVLTVLLYICLSVMLFYLFGSIMPLPPWGLVVGWVMALIAVELAYAFIFPYSFRYIADNDDDKMVILPV |
| IRIS_313-9572 | MLHHLKELAAVAGIHMILIYLCRFLLRRSRNVLFTVSNSLRFRLKVLTVLLYICLSVMLFYLFGSIMPLPPWGLVVGWVMALIAVELAYAFIFPYSFRYIADNDDDKMVILPV |
| IRIS_313-9574 | MLHHLKELAAVAGIHMILIYLCRFLLRRSRNVLFTVSNSLRFRLKVLTVLLYICLSVMLFYLFGSIMPLPPWGLVVGWVMALIAVELAYAFIFPYSFRYIADNDDDKMVILPV |
| IRIS_313-9575 | MLHHLKELAAVAGIHMILIYLCRFLLRRSRNVLFTVSNSLRFRLKVLTVLLYICLSVMLFYLFGSIMPLPPWGLVVGWVMALIAVELAYAFIFPYSFRYIADNDDDKMVILPV |
| IRIS_313-9582 | MLHHLKELAAVAGIHMILIYLCRFLLRRSRNVLFTVSNSLRFRLKVLTVLLYICLSVMLFYLFGSIMPLPPWGLVVGWVMALIAVELAYAFIFPYSFRYIADNDDDKMVILPV |
| IRIS_313-9590 | MLHHLKELAAVAGIHMILIYLCRFLLRRSRNVLFTVSNSLRFRLKVLTVLLYICLSVMLFYLFGSIMPLPPWGLVVGWVMALIAVELAYAFIFPYSFRYIADNDDDKMVILPV |
| IRIS_313-9593 | MLHHLKELAAVAGIHMILIYLCRFLLRRSRNVLFTVSNSLRFRLKVLTVLLYICLSVMLFYLFGSIMPLPPWGLVVGWVMALIAVELAYAFIFPYSFRYIADNDDDKMVILPV |
| IRIS_313-9594 | MLHHLKELAAVAGIHMILIYLCRFLLRRSRNVLFTVSNSLRFRLKVLTVLLYICLSVMLFYLFGSIMPLPPWGLVVGWVMALIAVELAYAFIFPYSFRYIADNDDDKMVILPV |
| IRIS_313-9600 | MLHHLKELAAVAGIHMILIYLCRFLLRRSRNVLFTVSNSLRFRLKVLTVLLYICLSVMLFYLFGSIMPLPPWGLVVGWVMALIAVELAYAFIFPYSFRYIADNDDDKMVILPV |
| IRIS_313-9601 | MLHHLKELAAVAGIHMILIYLCRFLLRRSRNVLFTVSNSLRFRLKVLTVLLYICLSVMLFYLFGSIMPLPPWGLVVGWVMALIAVELAYAFIFPYSFRYIADNDDDKMVILPV |
| IRIS_313-9602 | MLHHLKELAAVAGIHMILIYLCRFLLRRSRNVLFTVSNSLRFRLKVLTVLLYICLSVMLFYLFGSIMPLPPWGLVVGWVMALIAVELAYAFIFPYSFRYIADNDDDKMVILPV |
| IRIS_313-9604 | MLHHLKELAAVAGIHMILIYLCRFLLRRSRNVLFTVSNSLRFRLKVLTVLLYICLSVMLFYLFGSIMPLPPWGLVVGWVMALIAVELAYAFIFPYSFRYIADNDDDKMVILPV |
| IRIS_313-9605 | MLHHLKELAAVAGIHMILIYLCRFLLRRSRNVLFTVSNSLRFRLKVLTVLLYICLSVMLFYLFGSIMPLPPWGLVVGWVMALIAVELAYAFIFPYSFRYIADNDDDKMVILPV |
| IRIS_313-9606 | MLHHLKELAAVAGIHMILIYLCRFLLRRSRNVLFTVSNSLRFRLKVLTVLLYICLSVMLFYLFGSIMPLPPWGLVVGWVMALIAVELAYAFIFPYSFRYIADNDDDKMVILPV |
| IRIS_313-9609 | MLHHLKELAAVAGIHMILIYLCRFLLRRSRNVLFTVSNSLRFRLKVLTVLLYICLSVMLFYLFGSIMPLPPWGLVVGWVMALIAVELAYAFIFPYSFRYIADNDDDKMVILPV |
| IRIS_313-9610 | MLHHLKELAAVAGIHMILIYLCRFLLRRSRNVLFTVSNSLRFRLKVLTVLLYICLSVMLFYLFGSIMPLPPWGLVVGWVMALIAVELAYAFIFPYSFRYIADNDDDKMVILPV |
| IRIS_313-9611 | MLHHLKELAAVAGIHMILIYLCRFLLRRSRNVLFTVSNSLRFRLKVLTVLLYICLSVMLFYLFGSIMPLPPWGLVVGWVMALIAVELAYAFIFPYSFRYIADNDDDKMVILPV |
| IRIS_313-9616 | MLHHLKELAAVAGIHMILIYLCRFLLRRSRNVLFTVSNSLRFRLKVLTVLLYICLSVMLFYLFGSIMPLPPWGLVVGWVMALIAVELAYAFIFPYSFRYIADNDDDKMVILPV |
| IRIS_313-9617 | MLHHLKELAAVAGIHMILIYLCRFLLRRSRNVLFTVSNSLRFRLKVLTVLLYICLSVMLFYLFGSIMPLPPWGLVVGWVMALIAVELAYAFIFPYSFRYIADNDDDKMVILPV |
| IRIS_313-9619 | MLHHLKELAAVAGIHMILIYLCRFLLRRSRNVLFTVSNSLRFRLKVLTVLLYICLSVMLFYLFGSIMPLPPWGLVVGWVMALIAVELAYAFIFPYSFRYIADNDDDKMVILPV |
| IRIS_313-9623 | MLHHLKELAAVAGIHMILIYLCRFLLRRSRNVLFTVSNSLRFRLKVLTVLLYICLSVMLFYLFGSIMPLPPWGLVVGWVMALIAVELAYAFIFPYSFRYIADNDDDKMVILPV |
| IRIS_313-9626 | MLHHLKELAAVAGIHMILIYLCRFLLRRSRNVLFTVSNSLRFRLKVLTVLLYICLSVMLFYLFGSIMPLPPWGLVVGWVMALIAVELAYAFIFPYSFRYIADNDDDKMVILPV |
| IRIS_313-9629 | MLHHLKELAAVAGIHMILIYLCRFLLRRSRNVLFTVSNSLRFRLKVLTVLLYICLSVMLFYLFGSIMPLPPWGLVVGWVMALIAVELAYAFIFPYSFRYIADNDDDKMVILPV |
| IRIS_313-9634 | MLHHLKELAAVAGIHMILIYLCRFLLRRSRNVLFTVSNSLRFRLKVLTVLLYICLSVMLFYLFGSIMPLPPWGLVVGWVMALIAVELAYAFIFPYSFRYIADNDDDKMVILPV |
| IRIS_313-9636 | MLHHLKELAAVAGIHMILIYLCRFLLRRSRNVLFTVSNSLRFRLKVLTVLLYICLSVMLFYLFGSIMPLPPWGLVVGWVMALIAVELAYAFIFPYSFRYIADNDDDKMVILPV |
| IRIS_313-9637 | MLHHLKELAAVAGIHMILIYLCRFLLRRSRNVLFTVSNSLRFRLKVLTVLLYICLSVMLFYLFGSIMPLPPWGLVVGWVMALIAVELAYAFIFPYSFRYIADNDDDKMVILPV |
| IRIS_313-9641 | MLHHLKELAAVAGIHMILIYLCRFLLRRSRNVLFTVSNSLRFRLKVLTVLLYICLSVMLFYLFGSIMPLPPWGLVVGWVMALIAVELAYAFIFPYSFRYIADNDDDKMVILPV |
| IRIS_313-9648 | MLHHLKELAAVAGIHMILIYLCRFLLRRSRNVLFTVSNSLRFRLKVLTVLLYICLSVMLFYLFGSIMPLPPWGLVVGWVMALIAVELAYAFIFPYSFRYIADNDDDKMVILPV |
| IRIS_313-9661 | MLHHLKELAAVAGIHMILIYLCRFLLRRSRNVLFTVSNSLRFRLKVLTVLLYICLSVMLFYLFGSIMPLPPWGLVVGWVMALIAVELAYAFIFPYSFRYIADNDDDKMVILPV |
| IRIS_313-9669 | MLHHLKELAAVAGIHMILIYLCRFLLRRSRNVLFTVSNSLRFRLKVLTVLLYICLSVMLFYLFGSIMPLPPWGLVVGWVMALIAVELAYAFIFPYSFRYIADNDDDKMVILPV |
| IRIS_313-9682 | MLHHLKELAAVAGIHMILIYLCRFLLRRSRNVLFTVSNSLRFRLKVLTVLLYICLSVMLFYLFGSIMPLPPWGLVVGWVMALIAVELAYAFIFPYSFRYIADNDDDKMVILPV |
| IRIS_313-9685 | MLHHLKELAAVAGIHMILIYLCRFLLRRSRNVLFTVSNSLRFRLKVLTVLLYICLSVMLFYLFGSIMPLPPWGLVVGWVMALIAVELAYAFIFPYSFRYIADNDDDKMVILPV |
| IRIS_313-9687 | MLHHLKELAAVAGIHMILIYLCRFLLRRSRNVLFTVSNSLRFRLKVLTVLLYICLSVMLFYLFGSIMPLPPWGLVVGWVMALIAVELAYAFIFPYSFRYIADNDDDKMVILPV |
| IRIS_313-9691 | MLHHLKELAAVAGIHMILIYLCRFLLRRSRNVLFTVSNSLRFRLKVLTVLLYICLSVMLFYLFGSIMPLPPWGLVVGWVMALIAVELAYAFIFPYSFRYIADNDDDKMVILPV |
| IRIS_313-9694 | MLHHLKELAAVAGIHMILIYLCRFLLRRSRNVLFTVSNSLRFRLKVLTVLLYICLSVMLFYLFGSIMPLPPWGLVVGWVMALIAVELAYAFIFPYSFRYIADNDDDKMVILPV |
| IRIS_313-9695 | MLHHLKELAAVAGIHMILIYLCRFLLRRSRNVLFTVSNSLRFRLKVLTVLLYICLSVMLFYLFGSIMPLPPWGLVVGWVMALIAVELAYAFIFPYSFRYIADNDDDKMVILPV |
| IRIS_313-9696 | MLHHLKELAAVAGIHMILIYLCRFLLRRSRNVLFTVSNSLRFRLKVLTVLLYICLSVMLFYLFGSIMPLPPWGLVVGWVMALIAVELAYAFIFPYSFRYIADNDDDKMVILPV |
| IRIS_313-9697 | MLHHLKELAAVAGIHMILIYLCRFLLRRSRNVLFTVSNSLRFRLKVLTVLLYICLSVMLFYLFGSIMPLPPWGLVVGWVMALIAVELAYAFIFPYSFRYIADNDDDKMVILPV |
| IRIS_313-9698 | MLHHLKELAAVAGIHMILIYLCRFLLRRSRNVLFTVSNSLRFRLKVLTVLLYICLSVMLFYLFGSIMPLPPWGLVVGWVMALIAVELAYAFIFPYSFRYIADNDDDKMVILPV |
| IRIS_313-9699 | MLHHLKELAAVAGIHMILIYLCRFLLRRSRNVLFTVSNSLRFRLKVLTVLLYICLSVMLFYLFGSIMPLPPWGLVVGWVMALIAVELAYAFIFPYSFRYIADNDDDKMVILPV |
| IRIS_313-9701 | MLHHLKELAAVAGIHMILIYLCRFLLRRSRNVLFTVSNSLRFRLKVLTVLLYICLSVMLFYLFGSIMPLPPWGLVVGWVMALIAVELAYAFIFPYSFRYIADNDDDKMVILPV |
| IRIS_313-9702 | MLHHLKELAAVAGIHMILIYLCRFLLRRSRNVLFTVSNSLRFRLKVLTVLLYICLSVMLFYLFGSIMPLPPWGLVVGWVMALIAVELAYAFIFPYSFRYIADNDDDKMVILPV |
| IRIS_313-9703 | MLHHLKELAAVAGIHMILIYLCRFLLRRSRNVLFTVSNSLRFRLKVLTVLLYICLSVMLFYLFGSIMPLPPWGLVVGWVMALIAVELAYAFIFPYSFRYIADNDDDKMVILPV |
| IRIS_313-9705 | MLHHLKELAAVAGIHMILIYLCRFLLRRSRNVLFTVSNSLRFRLKVLTVLLYICLSVMLFYLFGSIMPLPPWGLVVGWVMALIAVELAYAFIFPYSFRYIADNDDDKMVILPV |
| IRIS_313-9706 | MLHHLKELAAVAGIHMILIYLCRFLLRRSRNVLFTVSNSLRFRLKVLTVLLYICLSVMLFYLFGSIMPLPPWGLVVGWVMALIAVELAYAFIFPYSFRYIADNDDDKMVILPV |
| IRIS_313-9708 | MLHHLKELAAVAGIHMILIYLCRFLLRRSRNVLFTVSNSLRFRLKVLTVLLYICLSVMLFYLFGSIMPLPPWGLVVGWVMALIAVELAYAFIFPYSFRYIADNDDDKMVILPV |
| IRIS_313-9709 | MLHHLKELAAVAGIHMILIYLCRFLLRRSRNVLFTVSNSLRFRLKVLTVLLYICLSVMLFYLFGSIMPLPPWGLVVGWVMALIAVELAYAFIFPYSFRYIADNDDDKMVILPV |
| IRIS_313-9723 | MLHHLKELAAVAGIHMILIYLCRFLLRRSRNVLFTVSNSLRFRLKVLTVLLYICLSVMLFYLFGSIMPLPPWGLVVGWVMALIAVELAYAFIFPYSFRYIADNDDDKMVILPV |
| IRIS_313-9724 | MLHHLKELAAVAGIHMILIYLCRFLLRRSRNVLFTVSNSLRFRLKVLTVLLYICLSVMLFYLFGSIMPLPPWGLVVGWVMALIAVELAYAFIFPYSFRYIADNDDDKMVILPV |
| IRIS_313-9725 | MLHHLKELAAVAGIHMILIYLCRFLLRRSRNVLFTVSNSLRFRLKVLTVLLYICLSVMLFYLFGSIMPLPPWGLVVGWVMALIAVELAYAFIFPYSFRYIADNDDDKMVILPV |
| IRIS_313-9727 | MLHHLKELAAVAGIHMILIYLCRFLLRRSRNVLFTVSNSLRFRLKVLTVLLYICLSVMLFYLFGSIMPLPPWGLVVGWVMALIAVELAYAFIFPYSFRYIADNDDDKMVILPV |
| IRIS_313-9730 | MLHHLKELAAVAGIHMILIYLCRFLLRRSRNVLFTVSNSLRFRLKVLTVLLYICLSVMLFYLFGSIMPLPPWGLVVGWVMALIAVELAYAFIFPYSFRYIADNDDDKMVILPV |
| IRIS_313-9732 | MLHHLKELAAVAGIHMILIYLCRFLLRRSRNVLFTVSNSLRFRLKVLTVLLYICLSVMLFYLFGSIMPLPPWGLVVGWVMALIAVELAYAFIFPYSFRYIADNDDDKMVILPV |
| IRIS_313-9740 | MLHHLKELAAVAGIHMILIYLCRFLLRRSRNVLFTVSNSLRFRLKVLTVLLYICLSVMLFYLFGSIMPLPPWGLVVGWVMALIAVELAYAFIFPYSFRYIADNDDDKMVILPV |
| IRIS_313-9742 | MLHHLKELAAVAGIHMILIYLCRFLLRRSRNVLFTVSNSLRFRLKVLTVLLYICLSVMLFYLFGSIMPLPPWGLVVGWVMALIAVELAYAFIFPYSFRYIADNDDDKMVILPV |
| IRIS_313-9745 | MLHHLKELAAVAGIHMILIYLCRFLLRRSRNVLFTVSNSLRFRLKVLTVLLYICLSVMLFYLFGSIMPLPPWGLVVGWVMALIAVELAYAFIFPYSFRYIADNDDDKMVILPV |
| IRIS_313-9758 | MLHHLKELAAVAGIHMILIYLCRFLLRRSRNVLFTVSNSLRFRLKVLTVLLYICLSVMLFYLFGSIMPLPPWGLVVGWVMALIAVELAYAFIFPYSFRYIADNDDDKMVILPV |
| IRIS_313-9759 | MLHHLKELAAVAGIHMILIYLCRFLLRRSRNVLFTVSNSLRFRLKVLTVLLYICLSVMLFYLFGSIMPLPPWGLVVGWVMALIAVELAYAFIFPYSFRYIADNDDDKMVILPV |
| IRIS_313-9767 | MLHHLKELAAVAGIHMILIYLCRFLLRRSRNVLFTVSNSLRFRLKVLTVLLYICLSVMLFYLFGSIMPLPPWGLVVGWVMALIAVELAYAFIFPYSFRYIADNDDDKMVILPV |
| IRIS_313-9769 | MLHHLKELAAVAGIHMILIYLCRFLLRRSRNVLFTVSNSLRFRLKVLTVLLYICLSVMLFYLFGSIMPLPPWGLVVGWVMALIAVELAYAFIFPYSFRYIADNDDDKMVILPV |
| IRIS_313-9770 | MLHHLKELAAVAGIHMILIYLCRFLLRRSRNVLFTVSNSLRFRLKVLTVLLYICLSVMLFYLFGSIMPLPPWGLVVGWVMALIAVELAYAFIFPYSFRYIADNDDDKMVILPV |
| IRIS_313-9771 | MLHHLKELAAVAGIHMILIYLCRFLLRRSRNVLFTVSNSLRFRLKVLTVLLYICLSVMLFYLFGSIMPLPPWGLVVGWVMALIAVELAYAFIFPYSFRYIADNDDDKMVILPV |
| IRIS_313-9774 | MLHQLKELAAVAGIHMILIYLCRFLLRRSRNVLFTVSNSLRFRLKVLTVLLYICLLVMLFYLFGSIMPLPPWGLVVGWVMALIAVELAYAFIFPYSFRYIADNDDDKMVILPV |
| IRIS_313-9778 | MLHHLKELAAVAGIHMILIYLCRFLLRRSRNVLFTVSNSLRFRLKVLTVLLYICLSVMLFYLFGSIMPLPPWGLVVGWVMALIAVELAYAFIFPYSFRYIADNDDDKMVILPV |
| IRIS_313-9782 | MLHHLKELAAVAGIHMILIYLCRFLLRRSRNVLFTVSNSLRFRLKVLTVLLYICLSVMLFYLFGSIMPLPPWGLVVGWVMALIAVELAYAFIFPYSFRYIADNDDDKMVILPV |
| IRIS_313-9783 | MLHHLKELAAVAGIHMILIYLCRFLLRRSRNVLFTVSNSLRFRLKVLTVLLYICLSVMLFYLFGSIMPLPPWGLVVGWVMALIAVELAYAFIFPYSFRYIADNDDDKMVILPV |
| IRIS_313-9789 | MLHHLKELAAVAGIHMILIYLCRFLLRRSRNVLFTVSNSLRFRLKVLTVLLYICLSVMLFYLFGSIMPLPPWGLVVGWVMALIAVELAYAFIFPYSFRYIADNDDDKMVILPV |
| IRIS_313-9790 | MLHHLKELAAVAGIHMILIYLCRFLLRRSRNVLFTVSNSLRFRLKVLTVLLYICLSVMLFYLFGSIMPLPPWGLVVGWVMALIAVELAYAFIFPYSFRYIADNDDDKMVILPV |
| IRIS_313-9791 | MLHHLKELAAVAGIHMILIYLCRFLLRRSRNVLFTVSNSLRFRLKVLTVLLYICLSVMLFYLFGSIMPLPPWGLVVGWVMALIAVELAYAFIFPYSFRYIADNDDDKMVILPV |
| IRIS_313-9795 | MLHHLKELAAVAGIHMILIYLCRFLLRRSRNVLFTVSNSLRFRLKVLTVLLYICLSVMLFYLFGSIMPLPPWGLVVGWVMALIAVELAYAFIFPYSFRYIADNDDDKMVILPV |
| IRIS_313-9800 | MLHHLKELAAVAGIHMILIYLCRFLLRRSRNVLFTVSNSLRFRLKVLTVLLYICLSVMLFYLFGSIMPLPPWGLVVGWVMALIAVELAYAFIFPYSFRYIADNDDDKMVILPV |

IRIS_313-9809 MLHHLKELAAVAGIHMILIYLCRFLLRRSRNVLFTVSNSLRFRLKVLTVLLYICLSVMLFYLFGSIMPLPPWGLVVGWVMALIAVELAYAFIFPYSFRYIADNDDDKMVILPV
IRIS_313-9811 MLHHLKELAAVAGIHMILIYLCRFLLRRSRNVLFTVSNSLRFRLKVLTVLLYICLSVMLFYLFGSIMPLPPWGLVVGWVMALIAVELAYAFIFPYSFRYIADNDDDKMVILPV
IRIS_313-9813 MLHHLKELAAVAGIHMILIYLCRFLLRRSRNVLFTVSNSLRFRLKVLTVLLYICLSVMLFYLFGSIMPLPPWGLVVGWVMALIAVELAYAFIFPYSFRYIADNDDDKMVILPV
IRIS_313-9814 MLHHLKELAAVAGIHMILIYLCRFLLRRSRNVLFTVSNSLRFRLKVLTVLLYICLSVMLFYLFGSIMPLPPWGLVVGWVMALIAVELAYAFIFPYSFRYIADNDDDKMVILPV
IRIS_313-9817 MLHHLKELAAVAGIHMILIYLCRFLLRRSRNVLFTVSNSLRFRLKVLTVLLYICLSVMLFYLFGSIMPLPPWGLVVGWVMALIAVELAYAFIFPYSFRYIADNDDDKMVILPV
IRIS_313-9818 MLHHLKELAAVAGIHMILIYLCRFLLRRSRNVLFTVSNSLRFRLKVLTVLLYICLSVMLFYLFGSIMPLPPWGLVVGWVMALIAVELAYAFIFPYSFRYIADNDDDKMVILPV
IRIS_313-9822 MLHHLKELAAVAGIHMILIYLCRFLLRRSRNVLFTVSNSLRFRLKVLTVLLYICLSVMLFYLFGSIMPLPPWGLVVGWVMALIAVELAYAFIFPYSFRYIADNDDDKMVILPV
IRIS_313-9825 MLHHLKELAAVAGIHMILIYLCRFLLRRSRNVLFTVSNSLRFRLKVLTVLLYICLSVMLFYLFGSIMPLPPWGLVVGWVMALIAVELAYAFIFPYSFRYIADNDDDKMVILPV
IRIS_313-9831 MLHHLKELAAVAGIHMILIYLCRFLLRRSRNVLFTVSNSLRFRLKVLTVLLYICLSVMLFYLFGSIMPLPPWGLVVGWVMALIAVELAYAFIFPYSFRYIADNDDDKMVILPV
IRIS_313-9832 MLHHLKELAAVAGIHMILIYLCRFLLRRSRNVLFTVSNSLRFRLKVLTVLLYICLSVMLFYLFGSIMPLPPWGLVVGWVMALIAVELAYAFIFPYSFRYIADNDDDKMVILPV
IRIS_313-9838 MLHHLKELAAVAGIHMILIYLCRFLLRRSRNVLFTVSNSLRFRLKVLTVLLYICLSVMLFYLFGSIMPLPPWGLVVGWVMALIAVELAYAFIFPYSFRYIADNDDDKMVILPV
IRIS_313-9839 MLHHLKELAAVAGIHMILIYLCRFLLRRSRNVLFTVSNSLRFRLKVLTVLLYICLSVMLFYLFGSIMPLPPWGLVVGWVMALIAVELAYAFIFPYSFRYIADNDDDKMVILPV
IRIS_313-9841 MLHHLKELAAVAGIHMILIYLCRFLLRRSRNVLFTVSNSLRFRLKVLTVLLYICLSVMLFYLFGSIMPLPPWGLVVGWVMALIAVELAYAFIFPYSFRYIADNDDDKMVILPV
IRIS_313-9851 MLHHLKELAAVAGIHMILIYLCRFLLRRSRNVLFTVSNSLRFRLKVLTVLLYICLSVMLFYLFGSIMPLPPWGLVVGWVMALIAVELAYAFIFPYSFRYIADNDDDKMVILPV
IRIS_313-9861 MLHHLKELAAVAGIHMILIYLCRFLLRRSRNVLFTVSNSLRFRLKVLTVLLYICLSVMLFYLFGSIMPLPPWGLVVGWVMALIAVELAYAFIFPYSFRYIADNDDDKMVILPV
IRIS_313-9862 MLHHLKELAAVAGIHMILIYLCRFLLRRSRNVLFTVSNSLRFRLKVLTVLLYICLSVMLFYLFGSIMPLPPWGLVVGWVMALIAVELAYAFIFPYSFRYIADNDDDKMVILPV
IRIS_313-9867 MLHHLKELAAVAGIHMILIYLCRFLLRRSRNVLFTVSNSLRFRLKVLTVLLYICLSVMLFYLFGSIMPLPPWGLVVGWVMALIAVELAYAFIFPYSFRYIADNDDDKMVILPV
IRIS_313-9880 MLHHLKELAAVAGIHMILIYLCRFLLRRSRNVLFTVSNSLRFRLKVLTVLLYICLSVMLFYLFGSIMPLPPWGLVVGWVMALIAVELAYAFIFPYSFRYIADNDDDKMVILPV
IRIS_313-9882 MLHHLKELAAVAGIHMILIYLCRFLLRRSRNVLFTVSNSLRFRLKVLTVLLYICLSVMLFYLFGSIMPLPPWGLVVGWVMALIAVELAYAFIFPYSFRYIADNDDDKMVILPV
IRIS_313-9884 MLHHLKELAAVAGIHMILIYLCRFLLRRSRNVLFTVSNSLRFRLKVLTVLLYICLSVMLFYLFGSIMPLPPWGLVVGWVMALIAVELAYAFIFPYSFRYIADNDDDKMVILPV
IRIS_313-9886 MLHHLKELAAVAGIHMILIYLCRFLLRRSRNVLFTVSNSLRFRLKVLTVLLYICLSVMLFYLFGSIMPLPPWGLVVGWVMALIAVELAYAFIFPYSFRYIADNDDDKMVILPV
IRIS_313-9887 MLHHLKELAAVAGIHMILIYLCRFLLRRSRNVLFTVSNSLRFRLKVLTVLLYICLSVMLFYLFGSIMPLPPWGLVVGWVMALIAVELAYAFIFPYSFRYIADNDDDKMVILPV
IRIS_313-9890 MLHHLKELAAVAGIHMILIYLCRFLLRRSRNVLFTVSNSLRFRLKVLTVLLYICLSVMLFYLFGSIMPLPPWGLVVGWVMALIAVELAYAFIFPYSFRYIADNDDDKMVILPV
IRIS_313-9891 MLHHLKELAAVAGIHMILIYLCRFLLRRSRNVLFTVSNSLRFRLKVLTVLLYICLSVMLFYLFGSIMPLPPWGLVVGWVMALIAVELAYAFIFPYSFRYIADNDDDKMVILPV
IRIS_313-9897 MLHHLKELAAVAGIHMILIYLCRFLLRRSRNVLFTVSNSLRFRLKVLTVLLYICLSVMLFYLFGSIMPLPPWGLVVGWVMALIAVELAYAFIFPYSFRYIADNDDDKMVILPV
IRIS_313-9898 MLHHLKELAAVAGIHMILIYLCRFLLRRSRNVLFTVSNSLRFRLKVLTVLLYICLSVMLFYLFGSIMPLPPWGLVVGWVMALIAVELAYAFIFPYSFRYIADNDDDKMVILPV
IRIS_313-9917 MLHHLKELAAVAGIHMILIYLCRFLLRRSRNVLFTVSNSLRFRLKVLTVLLYICLSVMLFYLFGSIMPLPPWGLVVGWVMALIAVELAYAFIFPYSFRYIADNDDDKMVILPV
IRIS_313-9922 MLHHLKELAAVAGIHMILIYLCRFLLRRSRNVLFTVSNSLRFRLKVLTVLLYICLSVMLFYLFGSIMPLPPWGLVVGWVMALIAVELAYAFIFPYSFRYIADNDDDKMVILPV
IRIS_313-9924 MLHHLKELAAVAGIHMILIYLCRFLLRRSRNVLFTVSNSLRFRLKVLTVLLYICLSVMLFYLFGSIMPLPPWGLVVGWVMALIAVELAYAFIFPYSFRYIADNDDDKMVILPV
IRIS_313-9925 MLHHLKELAAVAGIHMILIYLCRFLLRRSRNVLFTVSNSLRFRLKVLTVLLYICLSVMLFYLFGSIMPLPPWGLVVGWVMALIAVELAYAFIFPYSFRYIADNDDDKMVILPV
IRIS_313-9926 MLHHLKELAAVAGIHMILIYLCRFLLRRSRNVLFTVSNSLRFRLKVLTVLLYICLSVMLFYLFGSIMPLPPWGLVVGWVMALIAVELAYAFIFPYSFRYIADNDDDKMVILPV
IRIS_313-9928 MLHHLKELAAVAGIHMILIYLCRFLLRRSRNVLFTVSNSLRFRLKVLTVLLYICLSVMLFYLFGSIMPLPPWGLVVGWVMALIAVELAYAFIFPYSFRYIADNDDDKMVILPV
IRIS_313-9929 MLHHLKELAAVAGIHMILIYLCRFLLRRSRNVLFTVSNSLRFRLKVLTVLLYICLSVMLFYLFGSIMPLPPWGLVVGWVMALIAVELAYAFIFPYSFRYIADNDDDKMVILPV
IRIS_313-9935 MLHHLKELAAVAGIHMILIYLCRFLLRRSRNVLFTVSNSLRFRLKVLTVLLYICLSVMLFYLFGSIMPLPPWGLVVGWVMALIAVELAYAFIFPYSFRYIADNDDDKMVILPV
IRIS_313-9936 MLHHLKELAAVAGIHMILIYLCRFLLRRSRNVLFTVSNSLRFRLKVLTVLLYICLSVMLFYLFGSIMPLPPWGLVVGWVMALIAVELAYAFIFPYSFRYIADNDDDKMVILPV
IRIS_313-9937 MLHHLKELAAVAGIHMILIYLCRFLLRRSRNVLFTVSNSLRFRLKVLTVLLYICLSVMLFYLFGSIMPLPPWGLVVGWVMALIAVELAYAFIFPYSFRYIADNDDDKMVILPV
IRIS_313-9939 MLHHLKELAAVAGIHMILIYLCRFLLRRSRNVLFTVSNSLRFRLKVLTVLLYICLSVMLFYLFGSIMPLPPWGLVVGWVMALIAVELAYAFIFPYSFRYIADNDDDKMVILPV
IRIS_313-9940 MLHHLKELAAVAGIHMILIYLCRFLLRRSRNVLFTVSNSLRFRLKVLTVLLYICLSVMLFYLFGSIMPLPPWGLVVGWVMALIAVELAYAFIFPYSFRYIADNDDDKMVILPV
IRIS_313-9944 MLHHLKELAAVAGIHMILIYLCRFLLRRSRNVLFTVSNSLRFRLKVLTVLLYICLSVMLFYLFGSIMPLPPWGLVVGWVMALIAVELAYAFIFPYSFRYIADNDDDKMVILPV
IRIS_313-9949 MLHHLKELAAVAGIHMILIYLCRFLLRRSRNVLFTVSNSLRFRLKVLTVLLYICLSVMLFYLFGSIMPLPPWGLVVGWVMALIAVELAYAFIFPYSFRYIADNDDDKMVILPV
IRIS_313-9953 MLHHLKELAAVAGIHMILIYLCRFLLRRSRNVLFTVSNSLRFRLKVLTVLLYICLSVMLFYLFGSIMPLPPWGLVVGWVMALIAVELAYAFIFPYSFRYIADNDDDKMVILPV
IRIS_313-9961 MLHHLKELAAVAGIHMILIYLCRFLLRRSRNVLFTVSNSLRFRLKVLTVLLYICLSVMLFYLFGSIMPLPPWGLVVGWVMALIAVELAYAFIFPYSFRYIADNDDDKMVILPV
IRIS_313-9963 MLHHLKELAAVAGIHMILIYLCRFLLRRSRNVLFTVSNSLRFRLKVLTVLLYICLSVMLFYLFGSIMPLPPWGLVVGWVMALIAVELAYAFIFPYSFRYIADNDDDKMVILPV
IRIS_313-9964 MLHHLKELAAVAGIHMILIYLCRFLLRRSRNVLFTVSNSLRFRLKVLTVLLYICLSVMLFYLFGSIMPLPPWGLVVGWVMALIAVELAYAFIFPYSFRYIADNDDDKMVILPV
IRIS_313-9966 MLHHLKELAAVAGIHMILIYLCRFLLRRSRNVLFTVSNSLRFRLKVLTVLLYICLSVMLFYLFGSIMPLPPWGLVVGWVMALIAVELAYAFIFPYSFRYIADNDDDKMVILPV
IRIS_313-9968 MLHHLKELAAVAGIHMILIYLCRFLLRRSRNVLFTVSNSLRFRLKVLTVLLYICLSVMLFYLFGSIMPLPPWGLVVGWVMALIAVELAYAFIFPYSFRYIADNDDDKMVILPV
IRIS_313-9969 MLHHLKELAAVAGIHMILIYLCRFLLRRSRNVLFTVSNSLRFRLKVLTVLLYICLSVMLFYLFGSIMPLPPWGLVVGWVMALIAVELAYAFIFPYSFRYIADNDDDKMVILPV
IRIS_313-9970 MLHHLKELAAVAGIHMILIYLCRFLLRRSRNVLFTVSNSLRFRLKVLTVLLYICLSVMLFYLFGSIMPLPPWGLVVGWVMALIAVELAYAFIFPYSFRYIADNDDDKMVILPV
IRIS_313-9974 MLHHLKELAAVAGIHMILIYLCRFLLRRSRNVLFTVSNSLRFRLKVLTVLLYICLSVMLFYLFGSIMPLPPWGLVVGWVMALIAVELAYAFIFPYSFRYIADNDDDKMVILPV
IRIS_313-9976 MLHHLKELAAVAGIHMILIYLCRFLLRRSRNVLFTVSNSLRFRLKVLTVLLYICLSVMLFYLFGSIMPLPPWGLVVGWVMALIAVELAYAFIFPYSFRYIADNDDDKMVILPV
IRIS_313-9978 MLHHLKELAAVAGIHMILIYLCRFLLRRSRNVLFTVSNSLRFRLKVLTVLLYICLSVMLFYLFGSIMPLPPWGLVVGWVMALIAVELAYAFIFPYSFRYIADNDDDKMVILPV
IRIS_313-9980 MLHHLKELAAVAGIHMILIYLCRFLLRRSRNVLFTVSNSLRFRLKVLTVLLYICLSVMLFYLFGSIMPLPPWGLVVGWVMALIAVELAYAFIFPYSFRYIADNDDDKMVILPV
IRIS_313-9986 MLHHLKELAAVAGIHMILIYLCRFLLRRSRNVLFTVSNSLRFRLKVLTVLLYICLSVMLFYLFGSIMPLPPWGLVVGWVMALIAVELAYAFIFPYSFRYIADNDDDKMVILPV
IRIS_313-9989 MLHHLKELAAVAGIHMILIYLCRFLLRRSRNVLFTVSNSLRFRLKVLTVLLYICLSVMLFYLFGSIMPLPPWGLVVGWVMALIAVELAYAFIFPYSFRYIADNDDDKMVILPV
IRIS_313-9995 MLHHLKELAAVAGIHMILIYLCRFLLRRSRNVLFTVSNSLRFRLKVLTVLLYICLSVMLFYLFGSIMPLPPWGLVVGWVMALIAVELAYAFIFPYSFRYIADNDDDKMVILPV
IRIS_313-9996 MLHHLKELAAVAGIHMILIYLCRFLLRRSRNVLFTVSNSLRFRLKVLTVLLYICLSVMLFYLFGSIMPLPPWGLVVGWVMALIAVELAYAFIFPYSFRYIADNDDDKMVILPV
IRIS_313-10000 MLHHLKELAAVAGIHMILIYLCRFLLRRSRNVLFTVSNSLRFRLKVLTVLLYICLSVMLFYLFGSIMPLPPWGLVVGWVMALIAVELAYAFIFPYSFRYIADNDDDKMVILPV
IRIS_313-10001 MLHHLKELAAVAGIHMILIYLCRFLLRRSRNVLFTVSNSLRFRLKVLTVLLYICLSVMLFYLFGSIMPLPPWGLVVGWVMALIAVELAYAFIFPYSFRYIADNDDDKMVILPV
IRIS_313-10002 MLHHLKELAAVAGIHMILIYLCRFLLRRSRNVLFTVSNSLRFRLKVLTVLLYICLSVMLFYLFGSIMPLPPWGLVVGWVMALIAVELAYAFIFPYSFRYIADNDDDKMVILPV
IRIS_313-10007 MLHHLKELAAVAGIHMILIYLCRFLLRRSRNVLFTVSNSLRFRLKVLTVLLYICLSVMLFYLFGSIMPLPPWGLVVGWVMALIAVELAYAFIFPYSFRYIADNDDDKMVILPV
IRIS_313-10010 MLHHLKELAAVAGIHMILIYLCRFLLRRSRNVLFTVSNSLRFRLKVLTVLLYICLSVMLFYLFGSIMPLPPWGLVVGWVMALIAVELAYAFIFPYSFRYIADNDDDKMVILPV
IRIS_313-10014 MLHHLKELAAVAGIHMILIYLCRFLLRRSRNVLFTVSNSLRFRLKVLTVLLYICLSVMLFYLFGSIMPLPPWGLVVGWVMALIAVELAYAFIFPYSFRYIADNDDDKMVILPV
IRIS_313-10016 MLHHLKELAAVAGIHMILIYLCRFLLRRSRNVLFTVSNSLRFRLKVLTVLLYICLSVMLFYLFGSIMPLPPWGLVVGWVMALIAVELAYAFIFPYSFRYIADNDDDKMVILPV
IRIS_313-10020 MLHHLKELAAVAGIHMILIYLCRFLLRRSRNVLFTVSNSLRFRLKVLTVLLYICLSVMLFYLFGSIMPLPPWGLVVGWVMALIAVELAYAFIFPYSFRYIADNDDDKMVILPV
IRIS_313-10025 MLHHLKELAAVAGIHMILIYLCRFLLRRSRNVLFTVSNSLRFRLKVLTVLLYICLSVMLFYLFGSIMPLPPWGLVVGWVMALIAVELAYAFIFPYSFRYIADNDDDKMVILPV
IRIS_313-10026 MLHHLKELAAVAGIHMILIYLCRFLLRRSRNVLFTVSNSLRFRLKVLTVLLYICLSVMLFYLFGSIMPLPPWGLVVGWVMALIAVELAYAFIFPYSFRYIADNDDDKMVILPV
IRIS_313-10030 MLHHLKELAAVAGIHMILIYLCRFLLRRSRNVLFTVSNSLRFRLKVLTVLLYICLSVMLFYLFGSIMPLPPWGLVVGWVMALIAVELAYAFIFPYSFRYIADNDDDKMVILPV
IRIS_313-10032 MLHHLKELAAVAGIHMILIYLCRFLLRRSRNVLFTVSNSLRFRLKVLTVLLYICLSVMLFYLFGSIMPLPPWGLVVGWVMALIAVELAYAFIFPYSFRYIADNDDDKMVILPV
IRIS_313-10034 MLHHLKELAAVAGIHMILIYLCRFLLRRSRNVLFTVSNSLRFRLKVLTVLLYICLSVMLFYLFGSIMPLPPWGLVVGWVMALIAVELAYAFIFPYSFRYIADNDDDKMVILPV
IRIS_313-10035 MLHHLKELAAVAGIHMILIYLCRFLLRRSRNVLFTVSNSLRFRLKVLTVLLYICLSVMLFYLFGSIMPLPPWGLVVGWVMALIAVELAYAFIFPYSFRYIADNDDDKMVILPV
IRIS_313-10040 MLHHLKELAAVAGIHMILIYLCRFLLRRSRNVLFTVSNSLRFRLKVLTVLLYICLSVMLFYLFGSIMPLPPWGLVVGWVMALIAVELAYAFIFPYSFRYIADNDDDKMVILPV
IRIS_313-10041 MLHHLKELAAVAGIHMILIYLCRFLLRRSRNVLFTVSNSLRFRLKVLTVLLYICLSVMLFYLFGSIMPLPPWGLVVGWVMALIAVELAYAFIFPYSFRYIADNDDDKMVILPV
IRIS_313-10045 MLHHLKELAAVAGIHMILIYLCRFLLRRSRNVLFTVSNSLRFRLKVLTVLLYICLSVMLFYLFGSIMPLPPWGLVVGWVMALIAVELAYAFIFPYSFRYIADNDDDKMVILPV
IRIS_313-10046 MLHHLKELAAVAGIHMILIYLCRFLLRRSRNVLFTVSNSLRFRLKVLTVLLYICLSVMLFYLFGSIMPLPPWGLVVGWVMALIAVELAYAFIFPYSFRYIADNDDDKMVILPV
IRIS_313-10047 MLHHLKELAAVAGIHMILIYLCRFLLRRSRNVLFTVSNSLRFRLKVLTVLLYICLSVMLFYLFGSIMPLPPWGLVVGWVMALIAVELAYAFIFPYSFRYIADNDDDKMVILPV
IRIS_313-10048 MLHHLKELAAVAGIHMILIYLCRFLLRRSRNVLFTVSNSLRFRLKVLTVLLYICLSVMLFYLFGSIMPLPPWGLVVGWVMALIAVELAYAFIFPYSFRYIADNDDDKMVILPV
IRIS_313-10050 MLHHLKELAAVAGIHMILIYLCRFLLRRSRNVLFTVSNSLRFRLKVLTVLLYICLSVMLFYLFGSIMPLPPWGLVVGWVMALIAVELAYAFIFPYSFRYIADNDDDKMVILPV
IRIS_313-10051 MLHHLKELAAVAGIHMILIYLCRFLLRRSRNVLFTVSNSLRFRLKVLTVLLYICLSVMLFYLFGSIMPLPPWGLVVGWVMALIAVELAYAFIFPYSFRYIADNDDDKMVILPV
IRIS_313-10054 MLHHLKELAAVAGIHMILIYLCRFLLRRSRNVLFTVSNSLRFRLKVLTVLLYICLSVMLFYLFGSIMPLPPWGLVVGWVMALIAVELAYAFIFPYSFRYIADNDDDKMVILPV
IRIS_313-10056 MLHHLKELAAVAGIHMILIYLCRFLLRRSRNVLFTVSNSLRFRLKVLTVLLYICLSVMLFYLFGSIMPLPPWGLVVGWVMALIAVELAYAFIFPYSFRYIADNDDDKMVILPV
IRIS_313-10059 MLHHLKELAAVAGIHMILIYLCRFLLRRSRNVLFTVSNSLRFRLKVLTVLLYISLSVMLLYLFGSIMPLPPWGLVVGWVMALIAVELAYAFIFPYSFRYIADNDDDKMVILPV
IRIS_313-10061 MLHHLKELAAVAGIHMILIYLCRFLLRRSRNVLFTVSNSLRFRLKVLTVLLYICLSVMLFYLFGSIMPLPPWGLVVGWVMALIAVELAYAFIFPYSFRYIADNDDDKMVILPV
IRIS_313-10062 MLHHLKELAAVAGIHMILIYLCRFLLRRSRNVLFTVSNSLRFRLKVLTVLLYICLSVMLFYLFGSIMPLPPWGLVVGWVMALIAVELAYAFIFPYSFRYIADNDDDKMVILPV
IRIS_313-10065 MLHHLKELAAVAGIHMILIYLCRFLLRRSRNVLFTVSNSLRFRLKVLTVLLYICLSVMLFYLFGSIMPLPPWGLVVGWVMALIAVELAYAFIFPYSFRYIADNDDDKMVILPV
IRIS_313-10067 MLHHLKELAAVAGIHMILIYLCRFLLRRSRNVLFTVSNSLRFRLKVLTVLLYICLSVMLFYLFGSIMPLPPWGLVVGWVMALIAVELAYAFIFPYSFRYIADNDDDKMVILPV
IRIS_313-10071 MLHHLKELAAVAGIHMILIYLCRFLLRRSRNVLFTVSNSLRFRLKVLTVLLYICLSVMLFYLFGSIMPLPPWGLVVGWVMALIAVELAYAFIFPYSFRYIADNDDDKMVILPV
IRIS_313-10073 MLHHLKELAAVAGIHMILIYLCRFLLRRSRNVLFTVSNSLRFRLKVLTVLLYICLSVMLFYLFGSIMPLPPWGLVVGWVMALIAVELAYAFIFPYSFRYIADNDDDKMVILPV
IRIS_313-10074 MLHHLKELAAVAGIHMILIYLCRFLLRRSRNVLFTVSNSLRFRLKVLTVLLYICLSVMLFYLFGSIMPLPPWGLVVGWVMALIAVELAYAFIFPYSFRYIADNDDDKMVILPV
IRIS_313-10075 MLHHLKELAAVAGIHMILIYLCRFLLRRSRNVLFTVSNSLRFRLKVLTVLLYICLSVMLFYLFGSIMPLPPWGLVVGWVMALIAVELAYAFIFPYSFRYIADNDDDKMVILPV
IRIS_313-10076 MLHHLKELAAVAGIHMILIYLCRFLLRRSRNVLFTVSNSLRFRLKVLTVLLYICLSVMLFYLFGSIMPLPPWGLVVGWVMALIAVELAYAFIFPYSFRYIADNDDDKMVILPV
IRIS_313-10077 MLHHLKELAAVAGIHMILIYLCRFLLRRSRNVLFTVSNSLRFRLKVLTVLLYICLSVMLFYLFGSIMPLPPWGLVVGWVMALIAVELAYAFIFPYSFRYIADNDDDKMVILPV
IRIS_313-10078 MLHHLKELAAVAGIHMILIYLCRFLLRRSRNVLFTVSNSLRFRLKVLTVLLYICLSVMLFYLFGSIMPLPPWGLVVGWVMALIAVELAYAFIFPYSFRYIADNDDDKMVILPV
IRIS_313-10079 MLHHLKELAAVAGIHMILIYLCRFLLRRSRNVLFTVSNSLRFRLKVLTVLLYICLSVMLFYLFGSIMPLPPWGLVVGWVMALIAVELAYAFIFPYSFRYIADNDDDKMVILPV
IRIS_313-10080 MLHHLKELAAVAGIHMILIYLCRFLLRRSRNVLFTVSNSLRFRLKVLTVLLYICLSVMLFYLFGSIMPLPPWGLVVGWVMALIAVELAYAFIFPYSFRYIADNDDDKMVILPV
IRIS_313-10082 MLHHLKELAAVAGIHMILIYLCRFLLRRSRNVLFTVSNSLRFRLKVLTVLLYICLSVMLFYLFGSIMPLPPWGLVVGWVMALIAVELAYAFIFPYSFRYIADNDDDKMVILPV
IRIS_313-10083 MLHHLKELAAVAGIHMILIYLCRFLLRRSRNVLFTVSNSLRFRLKVLTVLLYICLSVMLFYLFGSIMPLPPWGLVVGWVMALIAVELAYAFIFPYSFRYIADNDDDKMVILPV
IRIS_313-10084 MLHHLKELAAVAGIHMILIYLCRFLLRRSRNVLFTVSNSLRFRLKVLTVLLYICLSVMLFYLFGSIMPLPPWGLVVGWVMALIAVELAYAFIFPYSFRYIADNDDDKMVILPV
IRIS_313-10089 MLHHLKELAAVAGIHMILIYLCRFLLRRSRNVLFTVSNSLRFRLKVLTVLLYICLSVMLFYLFGSIMPLPPWGLVVGWVMALIAVELAYAFIFPYSFRYIADNDDDKMVILPV
IRIS_313-10092 MLHHLKELAAVAGIHMILIYLCRFLLRRSRNVLFTVSNSLRFRLKVLTVLLYICLSVMLFYLFGSIMPLPPWGLVVGWVMALIAVELAYAFIFPYSFRYIADNDDDKMVILPV
IRIS_313-10093 MLHHLKELAAVAGIHMILIYLCRFLLRRSRNVLFTVSNSLRFRLKVLTVLLYICLSVMLFYLFGSIMPLPPWGLVVGWVMALIAVELAYAFIFPYSFRYIADNDDDKMVILPV
IRIS_313-10094 MLHHLKELAAVAGIHMILIYLCRFLLRRSRNVLFTVSNSLRFRLKVLTVLLYICLSVMLFYLFGSIMPLPPWGLVVGWVMALIAVELAYAFIFPYSFRYIADNDDDKMVILPV
IRIS_313-10096 MLHHLKELAAVAGIHMILIYLCRFLLRRSRNVLFTVSNSLRFRLKVLTVLLYICLSVMLFYLFGSIMPLPPWGLVVGWVMALIAVELAYAFIFPYSFRYIADNDDDKMVILPV
IRIS_313-10097 MLHHLKELAAVAGIHMILIYLCRFLLRRSRNVLFTVSNSLRFRLKVLTVLLYICLSVMLFYLFGSIMPLPPWGLVVGWVMALIAVELAYAFIFPYSFRYIADNDDDKMVILPV
IRIS_313-10099 MLHHLKELAAVAGIHMILIYLCRFLLRRSRNVLFTVSNSLRFRLKVLTVLLYICLSVMLFYLFGSIMPLPPWGLVVGWVMALIAVELAYAFIFPYSFRYIADNDDDKMVILPV
IRIS_313-10102 MLHHLKELAAVAGIHMILIYLCRFLLRRSRNVLFTVSNSLRFRLKVLTVLLYICLSVMLFYLFGSIMPLPPWGLVVGWVMALIAVELAYAFIFPYSFRYIADNDDDKMVILPV
IRIS_313-10103 MLHHLKELAAVAGIHMILIYLCRFLLRRSRNVLFTVSNSLRFRLKVLTVLLYICLSVMLFYLFGSIMPLPPWGLVVGWVMALIAVELAYAFIFPYSFRYIADNDDDKMVILPV
IRIS_313-10109 MLHHLKELAAVAGIHMILIYLCRFLLRRSRNVLFTVSNSLRFRLKVLTVLLYICLSVMLFYLFGSIMPLPPWGLVVGWVMALIAVELAYAFIFPYSFRYIADNDDDKMVILPV
IRIS_313-10111 MLHHLKELAAVAGIHMILIYLCRFLLRRSRNVLFTVSNSLRFRLKVLTVLLYICLSVMLFYLFGSIMPLPPWGLVVGWVMALIAVELAYAFIFPYSFRYIADNDDDKMVILPV
IRIS_313-10113 MLHHLKELAAVAGIHMILIYLCRFLLRRSRNVLFTVSNSLRFRLKVLTVLLYICLSVMLFYLFGSIMPLPPWGLVVGWVMALIAVELAYAFIFPYSFRYIADNDDDKMVILPV
IRIS_313-10114 MLHHLKELAAVAGIHMILIYLCRFLLRRSRNVLFTVSNSLRFRLKVLTVLLYICLSVMLFYLFGSIMPLPPWGLVVGWVMALIAVELAYAFIFPYSFRYIADNDDDKMVILPV
IRIS_313-10119 MLHHLKELAAVAGIHMILIYLCRFLLRRSRNVLFTVSNSLRFRLKVLTVLLYICLSVMLFYLFGSIMPLPPWGLVVGWVMALIAVELAYAFIFPYSFRYIADNDDDKMVILPV
IRIS_313-10124 MLHHLKELAAVAGIHMILIYLCRFLLRRSRNVLFTVSNSLRFRLKVLTVLLYICLSVMLFYLFGSIMPLPPWGLVVGWVMALIAVELAYAFIFPYSFRYIADNDDDKMVILPV
IRIS_313-10129 MLHHLKELAAVAGIHMILIYLCRFLLRRSRNVLFTVSNSLRFRLKVLTVLLYICLSVMLFYLFGSIMPLPPWGLVVGWVMALIAVELAYAFIFPYSFRYIADNDDDKMVILPV
IRIS_313-10134 MLHHLKELAAVAGIHMILIYLCRFLLRRSRNVLFTVSNSLRFRLKVLTVLLYICLSVMLFYLFGSIMPLPPWGLVVGWVMALIAVELAYAFIFPYSFRYIADNDDDKMVILPV
IRIS_313-10147 MLHHLKELAAVAGIHMILIYLCRFLLRRSRNVLFTVSNSLRFRLKVLTVLLYICLSVMLFYLFGSIMPLPPWGLVVGWVMALIAVELAYAFIFPYSFRYIADNDDDKMVILPV
IRIS_313-10148 MLHHLKELAAVAGIHMILIYLCRFLLRRSRNVLFTVSNSLRFRLKVLTVLLYICLSVMLFYLFGSIMPLPPWGLVVGWVMALIAVELAYAFIFPYSFRYIADNDDDKMVILPV
IRIS_313-10150 MLHHLKELAAVAGIHMILIYLCRFLLRRSRNVLFTVSNSLRFRLKVLTVLLYICLSVMLFYLFGSIMPLPPWGLVVGWVMALIAVELAYAFIFPYSFRYIADNDDDKMVILPV
IRIS_313-10151 MLHHLKELAAVAGIHMILIYLCRFLLRRSRNVLFTVSNSLRFRLKVLTVLLYICLSVMLFYLFGSIMPLPPWGLVVGWVMALIAVELAYAFIFPYSFRYIADNDDDKMVILPV
IRIS_313-10152 MLHHLKELAAVAGIHMILIYLCRFLLRRSRNVLFTVSNSLRFRLKVLTVLLYICLSVMLFYLFGSIMPLPPWGLVVGWVMALIAVELAYAFIFPYSFRYIADNDDDKMVILPV
IRIS_313-10154 MLHHLKELAAVAGIHMILIYLCRFLLRRSRNVLFTVSNSLRFRLKVLTVLLYICLSVMLFYLFGSIMPLPPWGLVVGWVMALIAVELAYAFIFPYSFRYIADNDDDKMVILPV
IRIS_313-10158 MLHHLKELAAVAGIHMILIYLCRFLLRRSRNVLFTVSNSLRFRLKVLTVLLYICLSVMLFYLFGSIMPLPPWGLVVGWVMALIAVELAYAFIFPYSFRYIADNDDDKMVILPV
IRIS_313-10161 MLHHLKELAAVAGIHMILIYLCRFLLRRSRNVLFTVSNSLRFRLKVLTVLLYICLSVMLFYLFGSIMPLPPWGLVVGWVMALIAVELAYAFIFPYSFRYIADNDDDKMVILPV
IRIS_313-10164 MLHHLKELAAVAGIHMILIYLCRFLLRRSRNVLFTVSNSLRFRLKVLTVLLYICLSVMLFYLFGSIMPLPPWGLVVGWVMALIAVELAYAFIFPYSFRYIADNDDDKMVILPV

4451 IRIS_313-10167 MLHHLKELAAVAGIHMILIYLCRFLLRRSRNVLFTVSNSLRFRLKVLTVLLYICLSVMLFYLFGSIMPLPPWGLVVGWVMALIAVELAYAFIFPYSFRYIADNDDDKMVILPV
4452 IRIS_313-10168 MLHHLKELAAVAGIHMILIYLCRFLLRRSRNVLFTVSNSLRFRLKVLTVLLYICLSVMLFYLFGSIMPLPPWGLVVGWVMALIAVELAYAFIFPYSFRYIADNDDDKMVILPV
4453 IRIS_313-10170 MLHHLKELAAVAGIHMILIYLCRFLLRLSRNVLFTVSNSLRFRLKVLTVLLYICLSVMLFYLFGSIMPLPPWGLVVGWVMALIAVELAYAFIFPYSFRYIADNDDDKMVILPV
4454 IRIS_313-10171 MLHHLKELAAVAGIHMILIYLCRFLLRRSRNVLFTVSNSLRFRLKVLTVLLYICLSVMLFYLFGSIMPLPPWGLVVGWVMALIAVELAYAFIFPYSFRYIADNDDDKMVILPV
4455 IRIS_313-10176 MLHHLKELAAVAGIHMILIYLCRFLLRRSRNVLFTVSNSLRFRLKVLTVLLYICLSVMLFYLFGSIMPLPPWGLVVGWVMALIAVELAYAFIFPYSFRYIADNDDDKMVILPV
4456 IRIS_313-10177 MLHHLKELAAVAGIHMILIYLCRFLLRRSRNVLFTVSNSLRFRLKVLTVLLYICLSVMLFYLFGSIMPLPPWGLVVGWVMALIAVELAYAFIFPYSFRYIADNDDDKMVILPV
4457 IRIS_313-10178 MLHHLKELAAVAGIHMILIYLCRFLLRRSRNVLFTVSNSLRFRLKVLTVLLYICLSVMLFYLFGSIMPLPPWGLVVGWVMALIAVELAYAFIFPYSFRYIADNDDDKMVILPV
4458 IRIS_313-10179 MLHHLKELAAVAGIHMILIYLCRFLLRRSRNVLFTVSNSLRFRLKVLTVLLYICLSVMLFYLFGSIMPLPPWGLVVGWVMALIAVELAYAFIFPYSFRYIADNDDDKMVILPV
4459 IRIS_313-10189 MLHHLKELAAVAGIHMILIYLCRFLLRRSRNVLFTVSNSLRFRLKVLTVLLYICLSVMLFYLFGSIMPLPPWGLVVGWVMALIAVELAYAFIFPYSFRYIADNDDDKMVILPV
4460 IRIS_313-10190 MLHHLKELAAVAGIHMILIYLCRFLLRRSRNVLFTVSNSLRFRLKVLTVLLYICLSVMLFYLFGSIMPLPPWGLVVGWVMALIAVELAYAFIFPYSFRYIADNDDDKMVILPV
4461 IRIS_313-10191 MLHHLKELAAVAGIHMILIYLCRFLLRRSRNVLFTVSNSLRFRLKVLTVLLYICLSVMLFYLFGSIMPLPPWGLVVGWVMALIAVELAYAFIFPYSFRYIADNDDDKMVILPV
4462 IRIS_313-10196 MLHHLKELAAVAGIHMILIYLCRFLLRRSRNVLFTVSNSLRFRLKVLTVLLYICLSVMLFYLFGSIMPLPPWGLVVGWVMALIAVELAYAFIFPYSFRYIADNDDDKMVILPV
4463 IRIS_313-10211 MLHHLKELAAVAGIHMILIYLCRFLLRRSRNVLFTVSNSLRFRLKVLTVLLYICLSVMLFYLFGSIMPLPPWGLVVGWVMALIAVELAYAFIFPYSFRYIADNDDDKMVILPV
4464 IRIS_313-10220 MLHHLKELAAVAGIHMILIYLCRFLLRLRRNVLFTVSNSLRFRLKVLTVLLYICLSVMLFYLFGSIMPLPPWGLVVGWVMALIAVELAYAFIFPYSFRYIADNDDDKMVILPV
4465 IRIS_313-10221 MLHHLKELAAVAGIHMILIYLCRFLLRRSRNVLFTVSNSLRFRLKVLTVLLYICLSVMLFYLFGSIMPLPPWGLVVGWVMALIAVELAYAFIFPYSFRYIADNDDDKMVILPV
4466 IRIS_313-10224 MLHHLKELAAVAGIHMILIYLCRFLLRRSRNVLFTVSNSLRFRLKVLTVLLYICLSVMLFYLFGSIMPLPPWGLVVGWVMALIAVELAYAFIFPYSFRYIADNDDDKMVILPV
4467 IRIS_313-10226 MLHHLKELAAVAGIHMILIYLCRFLLRRSRNVLFTVSNSLRFRLKVLTVLLYICLSVMLFYLFGSIMPLPPWGLVVGWVMALIAVELAYAFIFPYSFRYIADNDDDKMVILPV
4468 IRIS_313-10228 MLHHLKELAAVAGIHMILIYLCRFLLRRSRNVLFTVSNSLRFRLKVLTVLLYICLSVMLFYLFGSIMPLPPWGLVVGWVMALIAVELAYAFIFPYSFRYIADNDDDKMVILPV
4469 IRIS_313-10234 MLHHLKELAAVAGIHMILIYLCRFLLRRSRNVLFTVSNSLRFRLKVLTVLLYICLSVMLFYLFGSIMPLPPWGLVVGWVMALIAVELAYAFIFPYSFRYIADNDDDKMVILPV
4470 IRIS_313-10235 MLHHLKELAAVAGIHMILIYLCRFLLRRSRNVLFTVSNSLRFRLKVLTVLLYICLSVMLFYLFGSIMPLPPWGLVVGWVMALIAVELAYAFIFPYSFRYIADNDDDKMVILPV
4471 IRIS_313-10237 MLHHLKELAAVAGIHMILIYLCRFLLRRSRNVLFTVSNSLRFRLKVLTVLLYICLSVMLFYLFGSIMPLPPWGLVVGWVMALIAVELAYAFIFPYSFRYIADNDDDKMVILPV
4472 IRIS_313-10238 MLHHLKELAAVAGIHMILIYLCRFLLRRSRNVLFTVSNSLRFRLKVLTVLLYICLSVMLFYLFGSIMPLPPWGLVVGWVMALIAVELAYAFIFPYSFRYIADNDDDKMVILPV
4473 IRIS_313-10239 MLHHLKELAAVAGIHMILIYLCRFLLRRSRNVLFTVSNSLRFRLKVLTVLLYICLSVMLFYLFGSIMPLPPWGLVVGWVMALIAVELAYAFIFPYSFRYIADNDDDKMVILPV
4474 IRIS_313-10242 MLHHLKELAAVAGIHMILIYLCRFLLRRSRNVLFTVSNSLRFRLKVLTVLLYICLSVMLFYLFGSIMPLPPWGLVVGWVMALIAVELAYAFIFPYSFRYIADNDDDKMVILPV
4475 IRIS_313-10247 MLHHLKELAAVAGIHMILIYLCRFLLRRSRNVLFTVSNSLRFRLKVLTVLLYICLSVMLFYLFGSIMPLPPWGLVVGWVMALIAVELAYAFIFPYSFRYIADNDDDKMVILPV
4476 IRIS_313-10257 MLHHLKELAAVAGIHMILIYLCRFLLRRSRNVLFTVSNSLRFRLKVLTVLLYICLSVMLFYLFGSIMPLPPWGLVVGWVMALIAVELAYAFIFPYSFRYIADNDDDKMVILPV
4477 IRIS_313-10258 MLHHLKELAAVAGIHMILIYLCRFLLRRSRNVLFTVSNSLRFRLKVLTVLLYICLSVMLFYLFGSIMPLPPWGLVVGWVMALIAVELAYAFIFPYSFRYIADNDDDKMVILPV
4478 IRIS_313-10260 MLHHLKELAAVAGIHMILIYLCRFLLRRSRNVLFTVSNSLRFRLKVLTVLLYICLSVMLFYLFGSIMPLPPWGLVVGWVMALIAVELAYAFIFPYSFRYIADNDDDKMVILPV
4479 IRIS_313-10263 MLHHLKELAAVAGIHMILIYLCRFLLRRSRNVLFTVSNSLRFRLKVLTVLLYICLSVMLFYLFGSIMPLPPWGLVVGWVMALIAVELAYAFIFPYSFRYIADNDDDKMVILPV
4480 IRIS_313-10268 MLHHLKELAAVAGIHMILIYLCRFLLRRSRNVLFTVSNSLRFRLKVLTVLLYICLSVMLFYLFGSIMPLPPWGLVVGWVMALIAVELAYAFIFPYSFRYIADNDDDKMVILPV
4481 IRIS_313-10271 MLHHLKELAAVAGIHMILIYLCRFLLRRSRNVLFTVSNSLRFRLKVLTVLLYICLSVMLFYLFGSIMPLPPWGLVVGWVMALIAVELAYAFIFPYSFRYIADNDDDKMVILPV
4482 IRIS_313-10272 MLHHLKELAAVAGIHMILIYLCRFLLRRSRNVLFTVSNSLRFRLKVLTVLLYICLSVMLFYLFGSIMPLPPWGLVVGWVMALIAVELAYAFIFPYSFRYIADNDDDKMVILPV
4483 IRIS_313-10274 MLHHLKELAAVAGIHMILIYLCRFLLRRSRNVLFTVSNSLRFRLKVLTVLLYICLSVMLFYLFGSIMPLPPWGLVVGWVMALIAVELAYAFIFPYSFRYIADNDDDKMVILPV
4484 IRIS_313-10275 MLHHLKELAAVAGIHMILIYLCRFLLRRSRNVLFTVSNSLRFRLKVLTVLLYICLSVMLFYLFGSIMPLPPWGLVVGWVMALIAVELAYAFIFPYSFRYIADNDDDKMVILPV
4485 IRIS_313-10279 MLHHLKELAAVAGIHMILIYLCRFLLRRSRNVLFTVSNSLRFRLKVLTVLLYICLSVMLFYLFGSIMPLPPWGLVVGWVMALIAVELAYAFIFPYSFRYIADNDDDKMVILPV
4486 IRIS_313-10285 MLHHLKELAAVAGIHMILIYLCRFLLRRSRNVLFTVSNSLRFRLKVLTVLLYICLSVMLFYLFGSIMPLPPWGLVVGWVMALIAVELAYAFIFPYSFRYIADNDDDKMVILPV
4487 IRIS_313-10287 MLHHLKELAAVAGIHMILIYLCRFLLRRSRNVLFTVSNSLRFRLKVLTVLLYICLSVMLFYLFGSIMPLPPWGLVVGWVMALIAVELAYAFIFPYSFRYIADNDDDKMVILPV
4488 IRIS_313-10289 MLHHLKELAAVAGIHMILIYLCRFLLRRSRNVLFTVSNSLRFRLKVLTVLLYICLSVMLFYLFGSIMPLPPWGLVVGWVMALIAVELAYAFIFPYSFRYIADNDDDKMVILPV
4489 IRIS_313-10290 MLHHLKELAAVAGIHMILIYLCRFLLRRSRNVLFTVSNSLRFRLKVLTVLLYICLSVMLFYLFGSIMPLPPWGLVVGWVMALIAVELAYAFIFPYSFRYIADNDDDKMVILPV
4490 IRIS_313-10293 MLHHLKELAAVAGIHMILIYLCRFLLRRSRNVLFTVSNSLRFRLKVLTVLLYICLSVMLFYLFGSIMPLPPWGLVVGWVMALIAVELAYAFIFPYSFRYIADNDDDKMVILPV
4491 IRIS_313-10294 MLHHLKELAAVAGIHMILIYLCRFLLRRSRNVLFTVSNSLRFRLKVLTVLLYICLSVMLFYLFGSIMPLPPWGLVVGWVMALIAVELAYAFIFPYSFRYIADNDDDKMVILPV
4492 IRIS_313-10295 MLHHLKELAAVAGIHMILIYLCRFLLRRSRNVLFTVSNSLRFRLKVLTVLLYICLSVMLFYLFGSIMPLPPWGLVVGWVMALIAVELAYAFIFPYSFRYIADNDDDKMVILPV
4493 IRIS_313-10298 MLHHLKELAAVAGIHMILIYLCRFLLRRSRNVLFTVSNSLRFRLKVLTVLLYICLSVMLFYLFGSIMPLPPWGLVVGWVMALIAVELAYAFIFPYSFRYIADNDDDKMVILPV
4494 IRIS_313-10300 MLHHLKELAAVAGIHMILIYLCRFLLRRSRNVLFTVSNSLRFRLKVLTVLLYICLSVMLFYLFGSIMPLPPWGLVVGWVMALIAVELAYAFIFPYSFRYIADNDDDKMVILPV
4495 IRIS_313-10301 MLHHLKELAAVAGIHMILIYLCRFLLRRSRNVLFTVSNSLRFRLKVLTVLLYICLSVMLFYLFGSIMPLPPWGLVVGWVMALIAVELAYAFIFPYSFRYIADNDDDKMVILPV
4496 IRIS_313-10307 MLHHLKELAAVAGIHMILIYLCRFLLRRSRNVLFTVSNSLRFRLKVLTVLLYICLSVMLFYLFGSIMPLPPWGLVVGWVMALIAVELAYAFIFPYSFRYIADNDDDKMVILPV
4497 IRIS_313-10314 MLHHLKELAAVAGIHMILIYLCRFLLRRSRNVLFTVSNSLRFRLKVLTVLLYICLSVMLFYLFGSIMPLPPWGLVVGWVMALIAVELAYAFIFPYSFRYIADNDDDKMVILPV
4498 IRIS_313-10318 MLHHLKELAAVAGIHMILIYLCRFLLRRSRNVLFTVSNSLRFRLKVLTVLLYICLSVMLFYLFGSIMPLPPWGLVVGWVMALIAVELAYAFIFPYSFRYIADNDDDKMVILPV
4499 IRIS_313-10325 MLHHLKELAAVAGIHMILIYLCRFLLRRSRNVLFTVSNSLRFRLKVLTVLLYICLSVMLFYLFGSIMPLPPWGLVVGWVMALIAVELAYAFIFPYSFRYIADNDDDKMVILPV
4500 IRIS_313-10327 MLHHLKELAAVAGIHMILIYLCRFLLRRSRNVLFTVSNSLRFRLKVLTVLLYICLSVMLFYLFGSIMPLPPWGLVVGWVMALIAVELAYAFIFPYSFRYIADNDDDKMVILPV
4501 IRIS_313-10332 MLHHLKELAAVAGIHMILIYLCRFLLRRSRNVLFTVSNSLRFRLKVLTVLLYICLSVMLFYLFGSIMPLPPWGLVVGWVMALIAVELAYAFIFPYSFRYIADNDDDKMVILPV
4502 IRIS_313-10333 MLHHLKELAAVAGIHMILIYLCRFLLRRSRNVLFTVSNSLRFRLKVLTVLLYICLSVMLFYLFGSIMPLPPWGLVVGWVMALIAVELAYAFIFPYSFRYIADNDDDKMVILPV
4503 IRIS_313-10334 MLHHLKELAAVAGIHMILIYLCRFLLRRSRNVLFTVSNSLRFRLKVLTVLLYICLSVMLFYLFGSIMPLPPWGLVVGWVMALIAVELAYAFIFPYSFRYIADNDDDKMVILPV
4504 IRIS_313-10336 MLHHLKELAAVAGIHMILIYLCRFLLRRSRNVLFTVSNSLRFRLKVLTVLLYICLSVMLFYLFGSIMPLPPWGLVVGWVMALIAVELAYAFIFPYSFRYIADNDDDKMVILPV
4505 IRIS_313-10337 MLHHLKELAAVAGIHMILIYLCRFLLRRSRNVLFTVSNSLRFRLKVLTVLLYICLSVMLFYLFGSIMPLPPWGLVVGWVMALIAVELAYAFIFPYSFRYIADNDDDKMVILPV
4506 IRIS_313-10340 MLHHLKELAAVAGIHMILIYLCRFLLRRSRNVLFTVSNSLRFRLKVLTVLLYICLSVMLFYLFGSIMPLPPWGLVVGWVMALIAVELAYAFIFPYSFRYIADNDDDKMVILPV
4507 IRIS_313-10341 MLHHLKELAAVAGIHMILIYLCRFLLRRSRNVLFTVSNSLRFRLKVLTVLLYICLSVMLFYLFGSIMPLPPWGLVVGWVMALIAVELAYAFIFPYSFRYIADNDDDKMVILPV
4508 IRIS_313-10348 MLHHLKELAAVAGIHMILIYLCRFLLRRSRNVLFTVSNSLRFRLKVLTVLLYICLSVMLFYLFGSIMPLPPWGLVVGWVMALIAVELAYAFIFPYSFRYIADNDDDKMVILPV
4509 IRIS_313-10349 MLHHLKELAAVAGIHMILIYLCRFLLRRSRNVLFTVSNSLRFRLKVLTVLLYICLSVMLFYLFGSIMPLPPWGLVVGWVMALIAVELAYAFIFPYSFRYIADNDDDKMVILPV
4510 IRIS_313-10352 MLHHLKELAAVAGIHMILIYLCRFLLRRSRNVLFTVSNSLRFRLKVLTVLLYICLSVMLFYLFGSIMPLPPWGLVVGWVMALIAVELAYAFIFPYSFRYIADNDDDKMVILPV
4511 IRIS_313-10353 MLHHLKELAAVAGIHMILIYLCRFLLRRSRNVLFTVSNSLRFRLKVLTVLLYICLSVMLFYLFGSIMPLPPWGLVVGWVMALIAVELAYAFIFPYSFRYIADNDDDKMVILPV
4512 IRIS_313-10355 MLHHLKELAAVAGIHMILIYLCRFLLRRSRNVLFTVSNSLRFRLKVLTVLLYICLSVMLFYLFGSIMPLPPWGLVVGWVMALIAVELAYAFIFPYSFRYIADNDDDKMVILPV
4513 IRIS_313-10357 MLHHLKELAAVAGIHMILIYLCRFLLRRSRNVLFTVSNSLRFRLKVLTVLLYICLSVMLFYLFGSIMPLPPWGLVVGWVMALIAVELAYAFIFPYSFRYIADNDDDKMVILPV
4514 IRIS_313-10359 MLHHLKELAAVAGIHMILIYLCRFLLRRSRNVLFTVSNSLRFRLKVLTVLLYICLSVMLFYLFGSIMPLPPWGLVVGWVMALIAVELAYAFIFPYSFRYIADNDDDKMVILPV
4515 IRIS_313-10360 MLHHLKELAAVAGIHMILIYLCRFLLRRSRNVLFTVSNSLRFRLKVLTVLLYICLSVMLFYLFGSIMPLPPWGLVVGWVMALIAVELAYAFIFPYSFRYIADNDDDKMVILPV
4516 IRIS_313-10361 MLHHLKELAAVAGIHMILIYLCRFLLRRSRNVLFTVSNSLRFRLKVLTVLLYICLSVMLFYLFGSIMPLPPWGLVVGWVMALIAVELAYAFIFPYSFRYIADNDDDKMVILPV
4517 IRIS_313-10366 MLHHLKELAAVAGIHMILIYLCRFLLRRSRNVLFTVSNSLRFRLKVLTVLLYICLSVMLFYLFGSIMPLPPWGLVVGWVMALIAVELAYAFIFPYSFRYIADNDDDKMVILPV
4518 IRIS_313-10371 MLHHLKELAAVAGIHMILIYLCRFLLRRSRNVLFTVSNSLRFRLKVLTVLLYICLSVMLFYLFGSIMPLPPWGLVVGWVMALIAVELAYAFIFPYSFRYIADNDDDKMVILPV
4519 IRIS_313-10373 MLHHLKELAAVAGIHMILIYLCRFLLRRSRNVLFTVSNSLRFRLKVLTVLLYICLSVMLFYLFGSIMPLPPWGLVVGWVMALIAVELAYAFIFPYSFRYIADNDDDKMVILPV
4520 IRIS_313-10374 MLHHLKELAAVAGIHMILIYLCRFLLRRSRNVLFTVSNSLRFRLKVLTVLLYICLSVMLFYLFGSIMPLPPWGLVVGWVMALIAVELAYAFIFPYSFRYIADNDDDKMVILPV
4521 IRIS_313-10375 MLHHLKELAAVAGIHMILIYLCRFLLRRSRNVLFTVSNSLRFRLKVLTVLLYICLSVMLFYLFGSIMPLPPWGLVVGWVMALIAVELAYAFIFPYSFRYIADNDDDKMVILPV
4522 IRIS_313-10379 MLHHLKELAAVAGIHMILIYLCRFLLRRSRNVLFTVSNSLRFRLKVLTVLLYICLSVMLFYLFGSIMPLPPWGLVVGWVMALIAVELAYAFIFPYSFRYIADNDDDKMVILPV
4523 IRIS_313-10380 MLHHLKELAAVAGIHMILIYLCRFLLRRSRNVLFTVSNSLRFRLKVLTVLLYICLSVMLFYLFGSIMPLPPWGLVVGWVMALIAVELAYAFIFPYSFRYIADNDDDKMVILPV
4524 IRIS_313-10385 MLHHLKELAAVAGIHMILIYLCRFLLRLRRNVLFTVSNSLRFRLKVLTVLLYICLSVMLFYLFGSIMPLPPWGLVVGWVMALIAVELAYAFIFPYSFRYIADNDDDKMVILPV
4525 IRIS_313-10392 MLHHLKELAAVAGIHMILIYLCRFLLRRSRNVLFTVSNSLRFRLKVLTVLLYICLSVMLFYLFGSIMPLPPWGLVVGWVMALIAVELAYAFIFPYSFRYIADNDDDKMVILPV
4526 IRIS_313-10394 MLHHLKELAAVAGIHMILIYLCRFLLRRSRNVLFTVSNSLRFRLKVLTVLLYICLSVMLFYLFGSIMPLPPWGLVVGWVMALIAVELAYAFIFPYSFRYIADNDDDKMVILPV
4527 IRIS_313-10396 MLHHLKELAAVAGIHMILIYLCRFLLRRSRNVLFTVSNSLRFRLKVLTVLLYICLSVMLFYLFGSIMPLPPWGLVVGWVMALIAVELAYAFIFPYSFRYIADNDDDKMVILPV
4528 IRIS_313-10397 MLHHLKELAAVAGIHMILIYLCRFLLRRSRNVLFTVSNSLRFRLKVLTVLLYICLSVMLFYLFGSIMPLPPWGLVVGWVMALIAVELAYAFIFPYSFRYIADNDDDKMVILPV
4529 IRIS_313-10398 MLHHLKELAAVAGIHMILIYLCRFLLRRSRNVLFTVSNSLRFRLKVLTVLLYICLSVMLFYLFGSIMPLPPWGLVVGWVMALIAVELAYAFIFPYSFRYIADNDDDKMVILPV
4530 IRIS_313-10399 MLHHLKELAAVAGIHMILIYLCRFLLRRSRNVLFTVSNSLRFRLKVLTVLLYICLSVMLFYLFGSIMPLPPWGLVVGWVMALIAVELAYAFIFPYSFRYIADNDDDKMVILPV
4531 IRIS_313-10400 MLHHLKELAAVAGIHMILIYLCRFLLRRSRNVLFTVSNSLRFRLKVLTVLLYICLSVMLFYLFGSIMPLPPWGLVVGWVMALIAVELAYAFIFPYSFRYIADNDDDKMVILPV
4532 IRIS_313-10401 MLHHLKELAAVAGIHMILIYLCRFLLRRSRNVLFTVSNSLRFRLKVLTVLLYICLSVMLFYLFGSIMPLPPWGLVVGWVMALIAVELAYAFIFPYSFRYIADNDDDKMVILPV
4533 IRIS_313-10402 MLHHLKELAAVAGIHMILIYLCRFLLRRSRNVLFTVSNSLRFRLKVLTVLLYICLSVMLFYLFGSIMPLPPWGLVVGWVMALIAVELAYAFIFPYSFRYIADNDDDKMVILPV
4534 IRIS_313-10403 MLHHLKELAAVAGIHMILIYLCRFLLRRSRNVLFTVSNSLRFRLKVLTVLLYICLSVMLFYLFGSIMPLPPWGLVVGWVMALIAVELAYAFIFPYSFRYIADNDDDKMVILPV
4535 IRIS_313-10404 MLHHLKELAAVAGIHMILIYLCRFLLRRSRNVLFTVSNSLRFRLKVLTVLLYICLSVMLFYLFGSIMPLPPWGLVVGWVMALIAVELAYAFIFPYSFRYIADNDDDKMVILPV
4536 IRIS_313-10412 MLHHLKELAAVAGIHMILIYLCRFLLRRSRNVLFTVSNSLRFRLKVLTVLLYICLSVMLFYLFGSIMPLPPWGLVVGWVMALIAVELAYAFIFPYSFRYIADNDDDKMVILPV
4537 IRIS_313-10417 MLHHLKELAAVAGIHMILIYLCRFLLRRSRNVLFTVSNSLRFRLKVLTVLLYICLSVMLFYLFGSIMPLPPWGLVVGWVMALIAVELAYAFIFPYSFRYIADNDDDKMVILPV
4538 IRIS_313-10421 MLHHLKELAAVAGIHMILIYLCRFLLRRSRNVLFTVSNSLRFRLKVLTVLLYICLSVMLFYLFGSIMPLPPWGLVVGWVMALIAVELAYAFIFPYSFRYIADNDDDKMVILPV
4539 IRIS_313-10422 MLHHLKELAAVAGIHMILIYLCRFLLRRSRNVLFTVSNSLRFRLKVLTVLLYICLSVMLFYLFGSIMPLPPWGLVVGWVMALIAVELAYAFIFPYSFRYIADNDDDKMVILPV
4540 IRIS_313-10423 MLHHLKELAAVAGIHMILIYLCRFLLRRSRNVLFTVSNSLRFRLKVLTVLLYICLSVMLFYLFGSIMPLPPWGLVVGWVMALIAVELAYAFIFPYSFRYIADNDDDKMVILPV
4541 IRIS_313-10428 MLHHLKELAAVAGIHMILIYLCRFLLRRSRNVLFTVSNSLRFRLKVLTVLLYICLSVMLFYLFGSIMPLPPWGLVVGWVMALIAVELAYAFIFPYSFRYIADNDDDKMVILPV
4542 IRIS_313-10429 MLHHLKELAAVAGIHMILIYLCRFLLRRSRNVLFTVSNSLRFRLKVLTVLLYICLSVMLFYLFGSIMPLPPWGLVVGWVMALIAVELAYAFIFPYSFRYIADNDDDKMVILPV
4543 IRIS_313-10430 MLHHLKELAAVAGIHMILIYLCRFLLRRSRNVLFTVSNSLRFRLKVLTVLLYICLSVMLFYLFGSIMPLPPWGLVVGWVMALIAVELAYAFIFPYSFRYIADNDDDKMVILPV
4544 IRIS_313-10433 MLHHLKELAAVAGIHMILIYLCRFLLRRSRNVLFTVSNSLRFRLKVLTVLLYICLSVMLFYLFGSIMPLPPWGLVVGWVMALIAVELAYAFIFPYSFRYIADNDDDKMVILPV
4545 IRIS_313-10437 MLHHLKELAAVAGIHMILIYLCRFLLRRSRNVLFTVSNSLRFRLKVLTVLLYICLSVMLFYLFGSIMPLPPWGLVVGWVMALIAVELAYAFIFPYSFRYIADNDDDKMVILPV
4546 IRIS_313-10440 MLHHLKELAAVAGIHMILIYLCRFLLRRSRNVLFTVSNSLRFRLKVLTVLLYICLSVMLFYLFGSIMPLPPWGLVVGWVMALIAVELAYAFIFPYSFRYIADNDDDKMVILPV
4547 IRIS_313-10441 MLHHLKELAAVAGIHMILIYLCRFLLRRSRNVLFTVSNSLRFRLKVLTVLLYICLSVMLFYLFGSIMPLPPWGLVVGWVMALIAVELAYAFIFPYSFRYIADNDDDKMVILPV
4548 IRIS_313-10444 MLHHLKELAAVAGIHMILIYLCRFLLRRSRNVLFTVSNSLRFRLKVLTVLLYICLSVMLFYLFGSIMPLPPWGLVVGWVMALIAVELAYAFIFPYSFRYIADNDDDKMVILPV
4549 IRIS_313-10448 MLHHLKELAAVAGIHMILIYLCRFLLRRSRNVLFTVSNSLRFRLKVLTVLLYICLSVMLFYLFGSIMPLPPWGLVVGWVMALIAVELAYAFIFPYSFRYIADNDDDKMVILPV
4550 IRIS_313-10449 MLHHLKELAAVAGIHMILIYLCRFLLRRSRNVLFTVSNSLRFRLKVLTVLLYICLSVMLFYLFGSIMPLPPWGLVVGWVMALIAVELAYAFIFPYSFRYIADNDDDKMVILPV
4551 IRIS_313-10450 MLHHLKELAAVAGIHMILIYLCRFLLRRSRNVLFTVSNSLRFRLKVLTVLLYICLSVMLFYLFGSIMPLPPWGLVVGWVMALIAVELAYAFIFPYSFRYIADNDDDKMVILPV
4552 IRIS_313-10452 MLHHLKELAAVAGIHMILIYLCRFLLRRSRNVLFTVSNSLRFRLKVLTVLLYICLSVMLFYLFGSIMPLPPWGLVVGWVMALIAVELAYAFIFPYSFRYIADNDDDKMVILPV
4553 IRIS_313-10453 MLHHLKELAAVAGIHMILIYLCRFLLRRSRNVLFTVSNSLRFRLKVLTVLLYICLSVMLFYLFGSIMPLPPWGLVVGWVMALIAVELAYAFIFPYSFRYIADNDDDKMVILPV
4554 IRIS_313-10458 MLHHLKELAAVAGIHMILIYLCRFLLRRSRNVLFTVSNSLRFRLKVLTVLLYICLSVMLFYLFGSIMPLPPWGLVVGWVMALIAVELAYAFIFPYSFRYIADNDDDKMVILPV
4555 IRIS_313-10459 MLHHLKELAAVAGIHMILIYLCRFLLRRSRNVLFTVSNSLRFRLKVLTVLLYICLSVMLFYLFGSIMPLPPWGLVVGWVMALIAVELAYAFIFPYSFRYIADNDDDKMVILPV
4556 IRIS_313-10469 MLHHLKELAAVAGIHMILIYLCRFLLRRSRNVLFTVSNSLRFRLKVLTVLLYICLSVMLFYLFGSIMPLPPWGLVVGWVMALIAVELAYAFIFPYSFRYIADNDDDKMVILPV
4557 IRIS_313-10476 MLHHLKELAAVAGIHMILIYLCRFLLRRSRNVLFTVSNSLRFRLKVLTVLLYICLSVMLFYLFGSIMPLPPWGLVVGWVMALIAVELAYAFIFPYSFRYIADNDDDKMVILPV
4558 IRIS_313-10477 MLHHLKELAAVAGIHMILIYLCRFLLRRSRNVLFTVSNSLRFRLKVLTVLLYICLSVMLFYLFGSIMPLPPWGLVVGWVMALIAVELAYAFIFPYSFRYIADNDDDKMVILPV
4559 IRIS_313-10480 MLHHLKELAAVAGIHMILIYLCRFLLRRSRNVLFTVSNSLRFRLKVLTVLLYICLSVMLFYLFGSIMPLPPWGLVVGWVMALIAVELAYAFIFPYSFRYIADNDDDKMVILPV
4560 IRIS_313-10484 MLHHLKELAAVAGIHMILIYLCRFLLRRSRNVLFTVSNSLRFRLKVLTVLLYICLSVMLFYLFGSIMPLPPWGLVVGWVMALIAVELAYAFIFPYSFRYIADNDDDKMVILPV
4561 IRIS_313-10485 MLHHLKELAAVAGIHMILIYLCRFLLRRSRNVLFTVSNSLRFRLKVLTVLLYICLSVMLFYLFGSIMPLPPWGLVVGWVMALIAVELAYAFIFPYSFRYIADNDDDKMVILPV
4562 IRIS_313-10489 MLHHLKELAAVAGIHMILIYLCRFLLRRSRNVLFTVSNSLRFRLKVLTVLLYICLSVMLFYLFGSIMPLPPWGLVVGWVMALIAVELAYAFIFPYSFRYIADNDDDKMVILPV
4563 IRIS_313-10497 MLHHLKELAAVAGIHMILIYLCRFLLRRSRNVLFTVSNSLRFRLKVLTVLLYICLSVMLFYLFGSIMPLPPWGLVVGWVMALIAVELAYAFIFPYSFRYIADNDDDKMVILPV
4564 IRIS_313-10502 MLHHLKELAAVAGIHMILIYLCRFLLRRSRNVLFTVSNSLRFRLKVLTVLLYICLSVMLFYLFGSIMPLPPWGLVVGWVMALIAVELAYAFIFPYSFRYIADNDDDKMVILPV
4565 IRIS_313-10503 MLHHLKELAAVAGIHMILIYLCRFLLRRSRNVLFTVSNSLRFRLKVLTVLLYICLSVMLFYLFGSIMPLPPWGLVVGWVMALIAVELAYAFIFPYSFRYIADNDDDKMVILPV
4566 IRIS_313-10504 MLHHLKELAAVAGIHMILIYLCRFLLRRSRNVLFTVSNSLRFRLKVLTVLLYICLSVMLFYLFGSIMPLPPWGLVVGWVMALIAVELAYAFIFPYSFRYIADNDDDKMVILPV
4567 IRIS_313-10506 MLHHLKELAAVAGIHMILIYLCRFLLRRSRNVLFTVSNSLRFRLKVLTVLLYICLSVMLFYLFGSIMPLPPWGLVVGWVMALIAVELAYAFIFPYSFRYIADNDDDKMVILPV
4568 IRIS_313-10507 MLHHLKELAAVAGIHMILIYLCRFLLRRSRNVLFTVSNSLRFRLKVLTVLLYICLSVMLFYLFGSIMPLPPWGLVVGWVMALIAVELAYAFIFPYSFRYIADNDDDKMVILPV
4569 IRIS_313-10509 MLHHLKELAAVAGIHMILIYLCRFLLRRSRNVLFTVSNSLRFRLKVLTVLLYICLSVMLFYLFGSIMPLPPWGLVVGWVMALIAVELAYAFIFPYSFRYIADNDDDKMVILPV
4570 IRIS_313-10510 MLHHLKELAAVAGIHMILIYLCRFLLRRSRNVLFTVSNSLRFRLKVLTVLLYICLSVMLFYLFGSIMPLPPWGLVVGWVMALIAVELAYAFIFPYSFRYIADNDDDKMVILPV
4571 IRIS_313-10511 MLHHLKELAAVAGIHMILIYLCRFLLRRSRNVLFTVSNSLRFRLKVLTVLLYICLSVMLFYLFGSIMPLPPWGLVVGWVMALIAVELAYAFIFPYSFRYIADNDDDKMVILPV
4572 IRIS_313-10513 MLHHLKELAAVAGIHMILIYLCRFLLRRSRNVLFTVSNSLRFRLKVLTVLLYICLSVMLFYLFGSIMPLPPWGLVVGWVMALIAVELAYAFIFPYSFRYIADNDDDKMVILPV
4573 IRIS_313-10514 MLHHLKELAAVAGIHMILIYLCRFLLRRSRNVLFTVSNSLRFRLKVLTVLLYICLSVMLFYLFGSIMPLPPWGLVVGWVMALIAVELAYAFIFPYSFRYIADNDDDKMVILPV

| | |
|---|---|
| IRIS_313-10515 | MLHHLKELAAVAGIHMILIYLCRFLLRRSRNVLFTVSNSLRFRLKVLTVLLYICLSVMLFYLFGSIMPLPPWGLVVGWVMALIAVELAYAFIFPYSFRYIADNDDDKMVILPV |
| IRIS_313-10516 | MLHHLKELAAVAGIHMILIYLCRFLLRRSRNVLFTVSNSLRFRLKVLTVLLYICLSVMLFYLFGSIMPLPPWGLVVGWVMALIAVELAYAFIFPYSFRYIADNDDDKMVILPV |
| IRIS_313-10517 | MLHHLKELAAVAGIHMILIYLCRFLLRRSRNVLFTVSNSLRFRLKVLTVLLYICLSVMLFYLFGSIMPLPPWGLVVGWVMALIAVELAYAFIFPYSFRYIADNDDDKMVILPV |
| IRIS_313-10518 | MLHHLKELAAVAGIHMILIYLCRFLLRRSRNVLFTVSNSLRFRLKVLTVLLYICLSVMLFYLFGSIMPLPPWGLVVGWVMALIAVELAYAFIFPYSFRYIADNDDDKMVILPV |
| IRIS_313-10519 | MLHHLKELAAVAGIHMILIYLCRFLLRRSRNVLFTVSNSLRFRLKVLTVLLYICLSVMLFYLFGSIMPLPPWGLVVGWVMALIAVELAYAFIFPYSFRYIADNDDDKMVILPV |
| IRIS_313-10520 | MLHHLKELAAVAGIHMILIYLCRFLLRRSRNVLFTVSNSLRFRLKVLTVLLYICLSVMLFYLFGSIMPLPPWGLVVGWVMALIAVELAYAFIFPYSFRYIADNDDDKMVILPV |
| IRIS_313-10521 | MLHHLKELAAVAGIHMILIYLCRFLLRRSRNVLFTVSNSLRFRLKVLTVLLYICLSVMLFYLFGSIMPLPPWGLVVGWVMALIAVELAYAFIFPYSFRYIADNDDDKMVILPV |
| IRIS_313-10522 | MLHQLKELAAVAGIHMILIYLCRFLLRRSRNVLFTVSNSLRFRLKVLTVLLYICLLVMLFYLFGSIMPLPPWGLVVGWVMALIAVELAYAFIFPYSFRYIADNDDDKMVILPV |
| IRIS_313-10523 | MLHHLKELAAVAGIHMILIYLCRFLLRRSRNVLFTVSNSLRFRLKVLTVLLYICLSVMLFYLFGSIMPLPPWGLVVGWVMALIAVELAYAFIFPYSFRYIADNDDDKMVILPV |
| IRIS_313-10524 | MLHHLKELAAVAGIHMILIYLCRFLLRRSRNVLFTVSNSLRFRLKVLTVLLYICLSVMLFYLFGSIMPLPPWGLVVGWVMALIAVELAYAFIFPYSFRYIADNDDDKMVILPV |
| IRIS_313-10525 | MLHHLKELAAVAGIHMILIYLCRFLLRRSRNVLFTVSNSLRFRLKVLTVLLYICLSVMLFYLFGSIMPLPPWGLVVGWVMALIAVELAYAFIFPYSFRYIADNDDDKMVILPV |
| IRIS_313-10526 | MLHHLKELAAVAGIHMILIYLCRFLLRRSRNVLFTVSNSLRFRLKVLTVLLYICLSVMLFYLFGSIMPLPPWGLVVGWVMALIAVELAYAFIFPYSFRYIADNDDDKMVILPV |
| IRIS_313-10527 | MLHHLKELAAVAGIHMILIYLCRFLLRRSRNVLFTVSNSLRFRLKVLTVLLYICLSVMLFYLFGSIMPLPPWGLVVGWVMALIAVELAYAFIFPYSFRYIADNDDDKMVILPV |
| IRIS_313-10531 | MLHHLKELAAVAGIHMILIYLCRFLLRRSRNVLFTVSNSLRFRLKVLTVLLYICLSVMLFYLFGSIMPLPPWGLVVGWVMALIAVELAYAFIFPYSFRYIADNDDDKMVILPV |
| IRIS_313-10534 | MLHHLKELAAVAGIHMILIYLCRFLLRRSRNVLFTVSNSLRFRLKVLTVLLYICLSVMLFYLFGSIMPLPPWGLVVGWVMALIAVELAYAFIFPYSFRYIADNDDDKMVILPV |
| IRIS_313-10536 | MLHHLKELAAVAGIHMILIYLCRFLLRRSRNVLFTVSNSLRFRLKVLTVLLYICLSVMLFYLFGSIMPLPPWGLVVGWVMALIAVELAYAFIFPYSFRYIADNDDDKMVILPV |
| IRIS_313-10537 | MLHHLKELAAVAGIHMILIYLCRFLLRRSRNVLFTVSNSLRFRLKVLTVLLYICLSVMLFYLFGSIMPLPPWGLVVGWVMALIAVELAYAFIFPYSFRYIADNDDDKMVILPV |
| IRIS_313-10539 | MLHHLKELAAVAGIHMILIYLCRFLLRRSRNVLFTVSNSLRFRLKVLTVLLYICLSVMLFYLFGSIMPLPPWGLVVGWVMALIAVELAYAFIFPYSFRYIADNDDDKMVILPV |
| IRIS_313-10541 | MLHHLKELAAVAGIHMILIYLCRFLLRRSRNVLFTVSNSLRFRLKVLTVLLYICLSVMLFYLFGSIMPLPPWGLVVGWVMALIAVELAYAFIFPYSFRYIADNDDDKMVILPV |
| IRIS_313-10542 | MLHHLKELAAVAGIHMILIYLCRFLLRRSRNVLFTVSNSLRFRLKVLTVLLYICLSVMLFYLFGSIMPLPPWGLVVGWVMALIAVELAYAFIFPYSFRYIADNDDDKMVILPV |
| IRIS_313-10543 | MLHHLKELAAVAGIHMILIYLCRFLLRRSRNVLFTVSNSLRFRLKVLTVLLYICLSVMLFYLFGSIMPLPPWGLVVGWVMALIAVELAYAFIFPYSFRYIADNDDDKMVILPV |
| IRIS_313-10544 | MLHHLKELAAVAGIHMILIYLCRFLLRRSRNVLFTVSNSLRFRLKVLTVLLYICLSVMLFYLFGSIMPLPPWGLVVGWVMALIAVELAYAFIFPYSFRYIADNDDDKMVILPV |
| IRIS_313-10545 | MLHHLKELAAVAGIHMILIYLCRFLLRRSRNVLFTVSNSLRFRLKVLTVLLYICLSVMLFYLFGSIMPLPPWGLVVGWVMALIAVELAYAFIFPYSFRYIADNDDDKMVILPV |
| IRIS_313-10547 | MLHHLKELAAVAGIHMILIYLCRFLLRRSRNVLFTVSNSLRFRLKVLTVLLYICLSVMLFYLFGSIMPLPPWGLVVGWVMALIAVELAYAFIFPYSFRYIADNDDDKMVILPV |
| IRIS_313-10549 | MLHHLKELAAVAGIHMILIYLCRFLLRRSRNVLFTVSNSLRFRLKVLTVLLYICLSVMLFYLFGSIMPLPPWGLVVGWVMALIAVELAYAFIFPYSFRYIADNDDDKMVILPV |
| IRIS_313-10550 | MLHHLKELAAVAGIHMILIYLCRFLLRRSRNVLFTVSNSLRFRLKVLTVLLYICLSVMLFYLFGSIMPLPPWGLVVGWVMALIAVELAYAFIFPYSFRYIADNDDDKMVILPV |
| IRIS_313-10552 | MLHHLKELAAVAGIHMILIYLCRFLLRRSRNVLFTVSNSLRFRLKVLTVLLYICLSVMLFYLFGSIMPLPPWGLVVGWVMALIAVELAYAFIFPYSFRYIADNDDDKMVILPV |
| IRIS_313-10554 | MLHHLKELAAVAGIHMILIYLCRFLLRRSRNVLFTVSNSLRFRLKVLTVLLYICLSVMLFYLFGSIMPLPPWGLVVGWVMALIAVELAYAFIFPYSFRYIADNDDDKMVILPV |
| IRIS_313-10555 | MLHHLKELAAVAGIHMILIYLCRFLLRRSRNVLFTVSNSLRFRLKVLTVLLYICLSVMLFYLFGSIMPLPPWGLVVGWVMALIAVELAYAFIFPYSFRYIADNDDDKMVILPV |
| IRIS_313-10556 | MLHHLKELAAVAGIHMILIYLCRFLLRRSRNVLFTVSNSLRFRLKVLTVLLYICLSVMLFYLFGSIMPLPPWGLVVGWVMALIAVELAYAFIFPYSFRYIADNDDDKMVILPV |
| IRIS_313-10557 | MLHHLKELAAVAGIHMILIYLCRFLLRRSRNVLFTVSNSLRFRLKVLTVLLYICLSVMLFYLFGSIMPLPPWGLVVGWVMALIAVELAYAFIFPYSFRYIADNDDDKMVILPV |
| IRIS_313-10558 | MLHHLKELAAVAGIHMILIYLCRFLLRRSRNVLFTVSNSLRFRLKVLTVLLYICLSVMLFYLFGSIMPLPPWGLVVGWVMALIAVELAYAFIFPYSFRYIADNDDDKMVILPV |
| IRIS_313-10559 | MLHHLKELAAVAGIHMILIYLCRFLLRRSRNVLFTVSNSLRFRLKVLTVLLYICLSVMLFYLFGSIMPLPPWGLVVGWVMALIAVELAYAFIFPYSFRYIADNDDDKMVILPV |
| IRIS_313-10560 | MLHHLKELAAVAGIHMILIYLCRFLLRRSRNVLFTVSNSLRFRLKVLTVLLYICLSVMLFYLFGSIMPLPPWGLVVGWVMALIAVELAYAFIFPYSFRYIADNDDDKMVILPV |
| IRIS_313-10561 | MLHHLKELAAVAGIHMILIYLCRFLLRRSRNVLFTVSNSLRFRLKVLTVLLYICLSVMLFYLFGSIMPLPPWGLVVGWVMALIAVELAYAFIFPYSFRYIADNDDDKMVILPV |
| IRIS_313-10562 | MLHHLKELAAVAGIHMILIYLCRFLLRRSRNVLFTVSNSLRFRLKVLTVLLYICLSVMLFYLFGSIMPLPPWGLVVGWVMALIAVELAYAFIFPYSFRYIADNDDDKMVILPV |
| IRIS_313-10563 | MLHHLKELAAVAGIHMILIYLCRFLLRRSRNVLFTVSNSLRFRLKVLTVLLYICLSVMLFYLFGSIMPLPPWGLVVGWVMALIAVELAYAFIFPYSFRYIADNDDDKMVILPV |
| IRIS_313-10564 | MLHHLKELAAVAGIHMILIYLCRFLLRRSRNVLFTVSNSLRFRLKVLTVLLYICLSVMLFYLFGSIMPLPPWGLVVGWVMALIAVELAYAFIFPYSFRYIADNDDDKMVILPV |
| IRIS_313-10565 | MLHHLKELAAVAGIHMILIYLCRFLLRRSRNVLFTVSNSLRFRLKVLTVLLYICLSVMLFYLFGSIMPLPPWGLVVGWVMALIAVELAYAFIFPYSFRYIADNDDDKMVILPV |
| IRIS_313-10566 | MLHHLKELAAVAGIHMILIYLCRFLLRRSRNVLFTVSNSLRFRLKVLTVLLYICLSVMLFYLFGSIMPLPPWGLVVGWVMALIAVELAYAFIFPYSFRYIADNDDDKMVILPV |
| IRIS_313-10567 | MLHHLKELAAVAGIHMILIYLCRFLLRRSRNVLFTVSNSLRFRLKVLTVLLYICLSVMLFYLFGSIMPLPPWGLVVGWVMALIAVELAYAFIFPYSFRYIADNDDDKMVILPV |
| IRIS_313-10568 | MLHHLKELAAVAGIHMILIYLCRFLLRRSRNVLFTVSNSLRFRLKVLTVLLYICLSVMLFYLFGSIMPLPPWGLVVGWVMALIAVELAYAFIFPYSFRYIADNDDDKMVILPV |
| IRIS_313-10569 | MLHHLKELAAVAGIHMILIYLCRFLLRRSRNVLFTVSNSLRFRLKVLTVLLYICLSVMLFYLFGSIMPLPPWGLVVGWVMALIAVELAYAFIFPYSFRYIADNDDDKMVILPV |
| IRIS_313-10570 | MLHHLKELAAVAGIHMILIYLCRFLLRRSRNVLFTVSNSLRFRLKVLTVLLYICLSVMLFYLFGSIMPLPPWGLVVGWVMALIAVELAYAFIFPYSFRYIADNDDDKMVILPV |
| IRIS_313-10572 | MLHHLKELAAVAGIHMILIYLCRFLLRRSRNVLFTVSNSLRFRLKVLTVLLYICLSVMLFYLFGSIMPLPPWGLVVGWVMALIAVELAYAFIFPYSFRYIADNDDDKMVILPV |
| IRIS_313-10573 | MLHHLKELAAVAGIHMILIYLCRFLLRRSRNVLFTVSNSLRFRLKVLTVLLYICLSVMLFYLFGSIMPLPPWGLVVGWVMALIAVELAYAFIFPYSFRYIADNDDDKMVILPV |
| IRIS_313-10574 | MLHHLKELAAVAGIHMILIYLCRFLLRRSRNVLFTVSNSLRFRLKVLTVLLYICLSVMLFYLFGSIMPLPPWGLVVGWVMALIAVELAYAFIFPYSFRYIADNDDDKMVILPV |
| IRIS_313-10575 | MLHHLKELAAVAGIHMILIYLCRFLLRRSRNVLFTVSNSLRFRLKVLTVLLYICLSVMLFYLFGSIMPLPPWGLVVGWVMALIAVELAYAFIFPYSFRYIADNDDDKMVILPV |
| IRIS_313-10576 | MLHHLKELAAVAGIHMILIYLCRFLLRRSRNVLFTVSNSLRFRLKVLTVLLYICLSVMLFYLFGSIMPLPPWGLVVGWVMALIAVELAYAFIFPYSFRYIADNDDDKMVILPV |
| IRIS_313-10577 | MLHHLKELAAVAGIHMILIYLCRFLLRRSRNVLFTVSNSLRFRLKVLTVLLYICLSVMLFYLFGSIMPLPPWGLVVGWVMALIAVELAYAFIFPYSFRYIADNDDDKMVILPV |
| IRIS_313-10578 | MLHQLKELAAVAGIHMILIYLCRFLLRRSRNVLFTVSNSLRFRLKVLTVLLYICLLVMLFYLFGSIMPLPPWGLVVGWVMALIAVELAYAFIFPYSFRYIADNDDDKMVILPV |
| IRIS_313-10580 | MLHQLKELAAVAGIHMILIYLCRFLLRRSRNVLFTVSNSLRFRLKVLTVLLYICLLVMLFYLFGSIMPLPPWGLVVGWVMALIAVELAYAFIFPYSFRYIADNDDDKMVILPV |
| IRIS_313-10581 | MLHHLKELAAVAGIHMILIYLCRFLLRRSRNVLFTVSNSLRFRLKVLTVLLYICLSVMLFYLFGSIMPLPPWGLVVGWVMALIAVELAYAFIFPYSFRYIADNDDDKMVILPV |
| IRIS_313-10582 | MLHHLKELAAVAGIHMILIYLCRFLLRRSRNVLFTVSNSLRFRLKVLTVLLYICLSVMLFYLFGSIMPLPPWGLVVGWVMALIAVELAYAFIFPYSFRYIADNDDDKMVILPV |
| IRIS_313-10583 | MLHHLKELAAVAGIHMILIYLCRFLLRRSRNVLFTVSNSLRFRLKVLTVLLYICLSVMLFYLFGSIMPLPPWGLVVGWVMALIAVELAYAFIFPYSFRYIADNDDDKMVILPV |
| IRIS_313-10585 | MLHHLKELAAVAGIHMILIYLCRFLLRRSRNVLFTVSNSLRFRLKVLTVLLYICLSVMLFYLFGSIMPLPPWGLVVGWVMALIAVELAYAFIFPYSFRYIADNDDDKMVILPV |
| IRIS_313-10587 | MLHHLKELAAVAGIHMILIYLCRFLLRRSRNVLFTVSNSLRFRLKVLTVLLYICLSVMLFYLFGSIMPLPPWGLVVGWVMALIAVELAYAFIFPYSFRYIADNDDDKMVILPV |
| IRIS_313-10592 | MLHHLKELAAVAGIHMILIYLCRFLLRRSRNVLFTVSNSLRFRLKVLTVLLYISLSAMLLYLFGSIMPLPPWGLVVGWVMALIAVELAYAFIFPYSFRYIADNDDDKMVILPV |
| IRIS_313-10593 | MLHHLKELAAVAGIHMILIYLCRFLLRRSRNVLFTVSNSLRFRLKVLTVLLYICLSVMLFYLFGSIMPLPPWGLVVGWVMALIAVELAYAFIFPYSFRYIADNDDDKMVILPV |
| IRIS_313-10594 | MLHHLKELAAVAGIHMILIYLCRFLLRRSRNVLFTVSNSLRFRLKVLTVLLYICLSVMLFYLFGSIMPLPPWGLVVGWVMALIAVELAYAFIFPYSFRYIADNDDDKMVILPV |
| IRIS_313-10595 | MLHHLKELAAVAGIHMILIYLCRFLLRRSRNVLFTVSNSLRFRLKVLTVLLYICLSVMLFYLFGSIMPLPPWGLVVGWVMALIAVELAYAFIFPYSFRYIADNDDDKMVILPV |
| IRIS_313-10598 | MLHHLKELAAVAGIHMILIYLCRFLLRRSRNVLFTVSNSLRFRLKVLTVLLYICLSVMLFYLFGSIMPLPPWGLVVGWVMALIAVELAYAFIFPYSFRYIADNDDDKMVILPV |
| IRIS_313-10600 | MLHHLKELAAVAGIHMILIYLCRFLLRRSRNVLFTVSNSLRFRLKVLTVLLYICLSVMLFYLFGSIMPLPPWGLVVGWVMALIAVELAYAFIFPYSFRYIADNDDDKMVILPV |
| IRIS_313-10602 | MLHHLKELAAVAGIHMILIYLCRFLLRRSRNVLFTVSNSLRFRLKVLTVLLYICLSVMLFYLFGSIMPLPPWGLVVGWVMALIAVELAYAFIFPYSFRYIADNDDDKMVILPV |
| IRIS_313-10603 | MLHHLKELAAVAGIHMILIYLCRFLLRRSRNVLFTVSNSLRFRLKVLTVLLYICLSVMLFYLFGSIMPLPPWGLVVGWVMALIAVELAYAFIFPYSFRYIADNDDDKMVILPV |
| IRIS_313-10604 | MLHHLKELAAVAGIHMILIYLCRFLLRRSRNVLFTVSNSLRFRLKVLTVLLYICLSVMLFYLFGSIMPLPPWGLVVGWVMALIAVELAYAFIFPYSFRYIADNDDDKMVILPV |
| IRIS_313-10605 | MLHHLKELAAVAGIHMILIYLCRFLLRRSRNVLFTVSNSLRFRLKVLTVLLYICLSVMLFYLFGSIMPLPPWGLVVGWVMALIAVELAYAFIFPYSFRYIADNDDDKMVILPV |
| IRIS_313-10606 | MLHHLKELAAVAGIHMILIYLCRFLLRRSRNVLFTVSNSLRFRLKVLTVLLYICLSVMLFYLFGSIMPLPPWGLVVGWVMALIAVELAYAFIFPYSFRYIADNDDDKMVILPV |
| IRIS_313-10608 | MLHHLKELAAVAGIHMILIYLCRFLLRRSRNVLFTVSNSLRFRLKVLTVLLYICLSVMLFYLFGSIMPLPPWGLVVGWVMALIAVELAYAFIFPYSFRYIADNDDDKMVILPV |
| IRIS_313-10609 | MLHHLKELAAVAGIHMILIYLCRFLLRRSRNVLFTVSNSLRFRLKVLTVLLYICLSVMLFYLFGSIMPLPPWGLVVGWVMALIAVELAYAFIFPYSFRYIADNDDDKMVILPV |
| IRIS_313-10610 | MLHHLKELAAVAGIHMILIYLCRFLLRRSRNVLFTVSNSLRFRLKVLTVLLYICLSVMLFYLFGSIMPLPPWGLVVGWVMALIAVELAYAFIFPYSFRYIADNDDDKMVILPV |
| IRIS_313-10614 | MLHHLKELAAVAGIHMILIYLCRFFPSRRRNVLFTVSNSLRFRLKVLTVLLYICLSVMLFYLFGSIMPLPPWGLVVGWVMALIAVELAYAFIFPYSFRYIADNDDDKMVILPV |
| IRIS_313-10618 | MLHHLKELAAVAGIHMILIYLCRFLLRRSRNVLFTVSNSLRFRLKVLTVLLYICLSVMLFYLFGSIMPLPPWGLVVGWVMALIAVELAYAFIFPYSFRYIADNDDDKMVILPV |
| IRIS_313-10619 | MLHHLKELAAVAGIHMILIYLCRFLLRRSRNVLFTVSNSLRFRLKVLTVLLYICLSVMLFYLFGSIMPLPPWGLVVGWVMALIAVELAYAFIFPYSFRYIADNDDDKMVILPV |
| IRIS_313-10620 | MLHHLKELAAVAGIHMILIYLCRFLLRRSRNVLFTVSNSLRFRLKVLTVLLYICLSVMLFYLFGSIMPLPPWGLVVGWVMALIAVELAYAFIFPYSFRYIADNDDDKMVILPV |
| IRIS_313-10623 | MLHHLKELAAVAGIHMILIYLCRFLLRRSRNVLFTVSNSLRFRLKVLTVLLYICLSVMLFYLFGSIMPLPPWGLVVGWVMALIAVELAYAFIFPYSFRYIADNDDDKMVILPV |
| IRIS_313-10628 | MLHHLKELAAVAGIHMILIYLCRFLLRRSRNVLFTVSNSLRFRLKVLTVLLYICLSVMLFYLFGSIMPLPPWGLVVGWVMALIAVELAYAFIFPYSFRYIADNDDDKMVILPV |
| IRIS_313-10631 | MLHHLKELAAVAGIHMILIYLCRFLLRRSRNVLFTVSNSLRFRLKVLTVLLYICLSVMLFYLFGSIMPLPPWGLVVGWVMALIAVELAYAFIFPYSFRYIADNDDDKMVILPV |
| IRIS_313-10640 | MLHHLKELAAVAGIHMILIYLCRFLLRRSRNVLFTVSNSLRFRLKVLTVLLYICLSVMLFYLFGSIMPLPPWGLVVGWVMALIAVELAYAFIFPYSFRYIADNDDDKMVILPV |
| IRIS_313-10642 | MLHHLKELAAVAGIHMILIYLCRFLLRRSRNVLFTVSNSLRFRLKVLTVLLYICLSVMLFYLFGSIMPLPPWGLVVGWVMALIAVELAYAFIFPYSFRYIADNDDDKMVILPV |
| IRIS_313-10644 | MLHHLKELAAVAGIHMILIYLCRFLLRRSRNVLFTVSNSLRFRLKVLTVLLYICLSVMLFYLFGSIMPLPPWGLVVGWVMALIAVELAYAFIFPYSFRYIADNDDDKMVILPV |
| IRIS_313-10645 | MLHHLKELAAVAGIHMILIYLCRFLLRRSRNVLFTVSNSLRFRLKVLTVLLYICLSVMLFYLFGSIMPLPPWGLVVGWVMALIAVELAYAFIFPYSFRYIADNDDDKMVILPV |
| IRIS_313-10648 | MLHHLKELAAVAGIHMILIYLCRFLLRRSRNVLFTVSNSLRFRLKVLTVLLYICLSVMLFYLFGSIMPLPPWGLVVGWVMALIAVELAYAFIFPYSFRYIADNDDDKMVILPV |
| IRIS_313-10649 | MLHHLKELAAVAGIHMILIYLCRFLLRRSRNVLFTVSNSLRFRLKVLTVLLYICLSVMLFYLFGSIMPLPPWGLVVGWVMALIAVELAYAFIFPYSFRYIADNDDDKMVILPV |
| IRIS_313-10650 | MLHHLKELAAVAGIHMILIYLCRFLLRRSRNVLFTVSNSLRFRLKVLTVLLYICLSVMLFYLFGSIMPLPPWGLVVGWVMALIAVELAYAFIFPYSFRYIADNDDDKMVILPV |
| IRIS_313-10651 | MLHHLKELAAVAGIHMILIYLCRFLLRRSRNVLFTVSNSLRFRLKVLTVLLYICLSVMLFYLFGSIMPLPPWGLVVGWVMALIAVELAYAFIFPYSFRYIADNDDDKMVILPV |
| IRIS_313-10653 | MLHQLKELAAVAGIHMILIYLCRFLLRRSRNVLFTVSNSLRFRLKVLTVLLYICLLVMLFYLFGSIMPLPPWGLVVGWVMALIAVELAYAFIFPYSFRYIADNDDDKMVILPV |
| IRIS_313-10654 | MLHHLKELAAVAGIHMILIYLCRFLLRRSRNVLFTVSNSLRFRLKVLTVLLYICLSVMLFYLFGSIMPLPPWGLVVGWVMALIAVELAYAFIFPYSFRYIADNDDDKMVILPV |
| IRIS_313-10655 | MLHHLKELAAVAGIHMILIYLCRFLLRRSRNVLFTVSNSLRFRLKVLTVLLYICLSVMLFYLFGSIMPLPPWGLVVGWVMALIAVELAYAFIFPYSFRYIADNDDDKMVILPV |
| IRIS_313-10656 | MLHHLKELAAVAGIHMILIYLCRFLLRRSRNVLFTVSNSLRFRLKVLTVLLYICLSVMLFYLFGSIMPLPPWGLVVGWVMALIAVELAYAFIFPYSFRYIADNDDDKMVILPV |
| IRIS_313-10657 | MLHHLKELAAVAGIHMILIYLCRFLLRRSRNVLFTVSNSLRFRLKVLTVLLYICLSVMLFYLFGSIMPLPPWGLVVGWVMALIAVELAYAFIFPYSFRYIADNDDDKMVILPV |
| IRIS_313-10659 | MLHHLKELAAVAGIHMILIYLCRFLLRRSRNVLFTVSNSLRFRLKVLTVLLYICLSVMLFYLFGSIMPLPPWGLVVGWVMALIAVELAYAFIFPYSFRYIADNDDDKMVILPV |
| IRIS_313-10660 | MLHHLKELAAVAGIHMILIYLCRFLLRRSRNVLFTVSNSLRFRLKVLTVLLYICLSVMLFYLFGSIMPLPPWGLVVGWVMALIAVELAYAFIFPYSFRYIADNDDDKMVILPV |
| IRIS_313-10661 | MLHHLKELAAVAGIHMILIYLCRFLLRRSRNVLFTVSNSLRFRLKVLTVLLYICLSVMLFYLFGSIMPLPPWGLVVGWVMALIAVELAYAFIFPYSFRYIADNDDDKMVILPV |
| IRIS_313-10662 | MLHHLKELAAVAGIHMILIYLCRFLLRRSRNVLFTVSNSLRFRLKVLTVLLYICLSVMLFYLFGSIMPLPPWGLVVGWVMALIAVELAYAFIFPYSFRYIADNDDDKMVILPV |
| IRIS_313-10664 | MLHHLKELAAVAGIHMILIYLCRFLLRRSRNVLFTVSNSLRFRLKVLTVLLYICLSVMLFYLFGSIMPLPPWGLVVGWVMALIAVELAYAFIFPYSFRYIADNDDDKMVILPV |
| IRIS_313-10666 | MLHHLKELAAVAGIHMILIYLCRFLLRRSRNVLFTVSNSLRFRLKVLTVLLYICLSVMLFYLFGSIMPLPPWGLVVGWVMALIAVELAYAFIFPYSFRYIADNDDDKMVILPV |
| IRIS_313-10667 | MLHHLKELAAVAGIHMILIYLCRFLLRRSRNVLFTVSNSLRFRLKVLTVLLYICLSVMLFYLFGSIMPLPPWGLVVGWVMALIAVELAYAFIFPYSFRYIADNDDDKMVILPV |
| IRIS_313-10668 | MLHHLKELAAVAGIHMILIYLCRFLLRRSRNVLFTVSNSLRFRLKVLTVLLYICLSVMLFYLFGSIMPLPPWGLVVGWVMALIAVELAYAFIFPYSFRYIADNDDDKMVILPV |
| IRIS_313-10670 | MLHHLKELAAVAGIHMILIYLCRFLLRRSRNVLFTVSNSLRFRLKVLTVLLYICLSVMLFYLFGSIMPLPPWGLVVGWVMALIAVELAYAFIFPYSFRYIADNDDDKMVILPV |
| IRIS_313-10671 | MLHHLKELAAVAGIHMILIYLCRFLLRRSRNVLFTVSNSLRFRLKVLTVLLYICLSVMLFYLFGSIMPLPPWGLVVGWVMALIAVELAYAFIFPYSFRYIADNDDDKMVILPV |
| IRIS_313-10673 | MLHHLKELAAVAGIHMILIYLCRFLLRRSRNVLFTVSNSLRFRLKVLTVLLYICLSVMLFYLFGSIMPLPPWGLVVGWVMALIAVELAYAFIFPYSFRYIADNDDDKMVILPV |
| IRIS_313-10674 | MLHHLKELAAVAGIHMILIYLCRFLLRRSRNVLFTVSNSLRFRLKVLTVLLYICLSVMLFYLFGSIMPLPPWGLVVGWVMALIAVELAYAFIFPYSFRYIADNDDDKMVILPV |
| IRIS_313-10675 | MLHHLKELAAVAGIHMILIYLCRFLLRLRRNVLFTVSNSLRFRLKVLTVLLYICLSVMLFYLFGSIMPLPPWGLVVGWVMALIAVELAYAFIFPYSFRYIADNDDDKMVILPV |
| IRIS_313-10676 | MLHHLKELAAVAGIHMILIYLCRFLLRRSRNVLFTVSNSLRFRLKVLTVLLYICLSVMLFYLFGSIMPLPPWGLVVGWVMALIAVELAYAFIFPYSFRYIADNDDDKMVILPV |
| IRIS_313-10677 | MLHHLKELAAVAGIHMILIYLCRFLLRRSRNVLFTVSNSLRFRLKVLTVLLYICLSVMLFYLFGSIMPLPPWGLVVGWVMALIAVELAYAFIFPYSFRYIADNDDDKMVILPV |
| IRIS_313-10678 | MLHHLKELAAVAGIHMILIYLCRFLLRRSRNVLFTVSNSLRFRLKVLTVLLYICLSVMLFYLFGSIMPLPPWGLVVGWVMALIAVELAYAFIFPYSFRYIADNDDDKMVILPV |
| IRIS_313-10679 | MLHHLKELAAVAGIHMILIYLCRFLLRRSRNVLFTVSNSLRFRLKVLTVLLYICLSVMLFYLFGSIMPLPPWGLVVGWVMALIAVELAYAFIFPYSFRYIADNDDDKMVILPV |
| IRIS_313-10680 | MLHHLKELAAVAGIHMILIYLCRFLLRRSRNVLFTVSNSLRFRLKVLTVLLYICLSVMLFYLFGSIMPLPPWGLVVGWVMALIAVELAYAFIFPYSFRYIADNDDDKMVILPV |
| IRIS_313-10681 | MLHHLKELAAVAGIHMILIYLCRFLLRRSRNVLFTVSNSLRFRLKVLTVLLYICLSVMLFYLFGSIMPLPPWGLVVGWVMALIAVELAYAFIFPYSFRYIADNDDDKMVILPV |
| IRIS_313-10682 | MLHHLKELAAVAGIHMILIYLCRFLLRRSRNVLFTVSNSLRFRLKVLTVLLYICLSVMLFYLFGSIMPLPPWGLVVGWVMALIAVELAYAFIFPYSFRYIADNDDDKMVILPV |
| IRIS_313-10683 | MLHHLKELAAVAGIHMILIYLCRFLLRRSRNVLFTVSNSLRFRLKVLTVLLYICLSVMLFYLFGSIMPLPPWGLVVGWVMALIAVELAYAFIFPYSFRYIADNDDDKMVILPV |
| IRIS_313-10684 | MLHHLKELAAVAGIHMILIYLCRFLLRRSRNVLFTVSNSLRFRLKVLTVLLYICLSVMLFYLFGSIMPLPPWGLVVGWVMALIAVELAYAFIFPYSFRYIADNDDDKMVILPV |
| IRIS_313-10687 | MLHHLKELAAVAGIHMILIYLCRFLLRRSRNVLFTVSNSLRFRLKVLTVLLYICLSVMLFYLFGSIMPLPPWGLVVGWVMALIAVELAYAFIFPYSFRYIADNDDDKMVILPV |
| IRIS_313-10688 | MLHHLKELAAVAGIHMILIYLCRFLLRRSRNVLFTVSNSLRFRLKVLTVLLYICLSVMLFYLFGSIMPLPPWGLVVGWVMALIAVELAYAFIFPYSFRYIADNDDDKMVILPV |
| IRIS_313-10689 | MLHHLKELAAVAGIHMILIYLCRFLLRRSRNVLFTVSNSLRFRLKVLTVLLYICLSVMLFYLFGSIMPLPPWGLVVGWVMALIAVELAYAFIFPYSFRYIADNDDDKMVILPV |
| IRIS_313-10690 | MLHHLKELAAVAGIHMILIYLCRFLLRRSRNVLFTVSNSLRFRLKVLTVLLYICLSVMLFYLFGSIMPLPPWGLVVGWVMALIAVELAYAFIFPYSFRYIADNDDDKMVILPV |
| IRIS_313-10692 | MLHHLKELAAVAGIHMILIYLCRFLLRRSRNVLFTVSNSLRFRLKVLTVLLYICLSVMLFYLFGSIMPLPPWGLVVGWVMALIAVELAYAFIFPYSFRYIADNDDDKMVILPV |
| IRIS_313-10693 | MLHHLKELAAVAGIHMILIYLCRFLLRRSRNVLFTVSNSLRFRLKVLTVLLYICLSVMLFYLFGSIMPLPPWGLVVGWVMALIAVELAYAFIFPYSFRYIADNDDDKMVILPV |
| IRIS_313-10694 | MLHHLKELAAVAGIHMILIYLCRFLLRRSRNVLFTVSNSLRFRLKVLTVLLYICLSVMLFYLFGSIMPLPPWGLVVGWVMALIAVELAYAFIFPYSFRYIADNDDDKMVILPV |
| IRIS_313-10695 | MLHHLKELAAVAGIHMILIYLCRFLLRRSRNVLFTVSNSLRFRLKVLTVLLYICLSVMLFYLFGSIMPLPPWGLVVGWVMALIAVELAYAFIFPYSFRYIADNDDDKMVILPV |
| IRIS_313-10697 | MLHHLKELAAVAGIHMILIYLCRFLLRRSRNVLFTVSNSLRFRLKVLTVLLYICLSVMLFYLFGSIMPLPPWGLVVGWVMALIAVELAYAFIFPYSFRYIADNDDDKMVILPV |
| IRIS_313-10698 | MLHHLKELAAVAGIHMILIYLCRFLLRRSRNVLFTVSNSLRFRLKVLTVLLYICLSVMLFYLFGSIMPLPPWGLVVGWVMALIAVELAYAFIFPYSFRYIADNDDDKMVILPV |

| | |
|---|---|
| IRIS_313-10699 | MLHHLKELAAVAGIHMILIYLCRFLLRRSRNVLFTVSNSLRFRLKVLTVLLYICLSVMLFYLFGSIMPLPPWGLVVGWVMALIAVELAYAFIFPYSFRYIADNDDDKMVILPV |
| IRIS_313-10700 | MLHHLKELAAVAGIHMILIYLCRFLLRRSRNVLFTVSNSLRFRLKVLTVLLYICLSVMLFYLFGSIMPLPPWGLVVGWVMALIAVELAYAFIFPYSFRYIADNDDDKMVILPV |
| IRIS_313-10701 | MLHHLKELAAVAGIHMILIYLCRFLLRRSRNVLFTVSNSLRFRLKVLTVLLYICLSVMLFYLFGSIMPLPPWGLVVGWVMALIAVELAYAFIFPYSFRYIADNDDDKMVILPV |
| IRIS_313-10702 | MLHHLKELAAVAGIHMILIYLCRFLLRRSRNVLFTVSNSLRFRLKVLTVLLYICLSVMLFYLFGSIMPLPPWGLVVGWVMALIAVELAYAFIFPYSFRYIADNDDDKMVILPV |
| IRIS_313-10703 | MLHHLKELAAVAGIHMILIYLCRFLLRRSRNVLFTVSNSLRFRLKVLTVLLYICLSVMLFYLFGSIMPLPPWGLVVGWVMALIAVELAYAFIFPYSFRYIADNDDDKMVILPV |
| IRIS_313-10704 | MLHHLKELAAVAGIHMILIYLCRFLLRRSRNVLFTVSNSLRFRLKVLTVLLYICLSVMLFYLFGSIMPLPPWGLVVGWVMALIAVELAYAFIFPYSFRYIADNDDDKMVILPV |
| IRIS_313-10705 | MLHHLKELAAVAGIHMILIYLCRFLLRRSRNVLFTVSNSLRFRLKVLTVLLYICLSVMLFYLFGSIMPLPPWGLVVGWVMALIAVELAYAFIFPYSFRYIADNDDDKMVILPV |
| IRIS_313-10706 | MLHHLKELAAVAGIHMILIYLCRFLLRRSRNVLFTVSNSLRFRLKVLTVLLYICLSVMLFYLFGSIMPLPPWGLVVGWVMALIAVELAYAFIFPYSFRYIADNDDDKMVILPV |
| IRIS_313-10707 | MLHHLKELAAVAGIHMILIYLCRFLLRRSRNVLFTVSNSLRFRLKVLTVLLYICLSVMLFYLFGSIMPLPPWGLVVGWVMALIAVELAYAFIFPYSFRYIADNDDDKMVILPV |
| IRIS_313-10708 | MLHHLKELAAVAGIHMILIYLCRFLLRRSRNVLFTVSNSLRFRLKVLTVLLYICLSVMLFYLFGSIMPLPPWGLVVGWVMALIAVELAYAFIFPYSFRYIADNDDDKMVILPV |
| IRIS_313-10710 | MLHHLKELAAVAGIHMILIYLCRFLLRRSRNVLFTVSNSLRFRLKVLTVLLYICLSVMLFYLFGSIMPLPPWGLVVGWVMALIAVELAYAFIFPYSFRYIADNDDDKMVILPV |
| IRIS_313-10711 | MLHHLKELAAVAGIHMILIYLCRFLLRRSRNVLFTVSNSLRFRLKVLTVLLYICLSVMLFYLFGSIMPLPPWGLVVGWVMALIAVELAYAFIFPYSFRYIADNDDDKMVILPV |
| IRIS_313-10712 | MLHHLKELAAVAGIHMILIYLCRFLLRRSRNVLFTVSNSLRFRLKVLTVLLYICLSVMLFYLFGSIMPLPPWGLVVGWVMALIAVELAYAFIFPYSFRYIADNDDDKMVILPV |
| IRIS_313-10715 | MLHHLKELAAVAGIHMILIYLCRFLLRRSRNVLFTVSNSLRFRLKVLTVLLYICLSVMLFYLFGSIMPLPPWGLVVGWVMALIAVELAYAFIFPYSFRYIADNDDDKMVILPV |
| IRIS_313-10716 | MLHHLKELAAVAGIHMILIYLCRFLLRRSRNVLFTVSNSLRFRLKVLTVLLYICLSVMLFYLFGSIMPLPPWGLVVGWVMALIAVELAYAFIFPYSFRYIADNDDDKMVILPV |
| IRIS_313-10717 | MLHHLKELAAVAGIHMILIYLCRFLLRRSRNVLFTVSNSLRFRLKVLTVLLYICLSVMLFYLFGSIMPLPPWGLVVGWVMALIAVELAYAFIFPYSFRYIADNDDDKMVILPV |
| IRIS_313-10718 | MLHHLKELAAVAGIHMILIYLCRFLLRRSRNVLFTVSNSLRFRLKVLTVLLYICLSVMLFYLFGSIMPLPPWGLVVGWVMALIAVELAYAFIFPYSFRYIADNDDDKMVILPV |
| IRIS_313-10719 | MLHHLKELAAVAGIHMILIYLCRFLLRRSRNVLFTVSNSLRFRLKVLTVLLYICLSVMLFYLFGSIMPLPPWGLVVGWVMALIAVELAYAFIFPYSFRYIADNDDDKMVILPV |
| IRIS_313-10720 | MLHHLKELAAVAGIHMILIYLCRFLLRRSRNVLFTVSNSLRFRLKVLTVLLYICLSVMLFYLFGSIMPLPPWGLVVGWVMALIAVELAYAFIFPYSFRYIADNDDDKMVILPV |
| IRIS_313-10721 | MLHHLKELAAVAGIHMILIYLCRFLLRRSRNVLFTVSNSLRFRLKVLTVLLYICLSVMLFYLFGSIMPLPPWGLVVGWVMALIAVELAYAFIFPYSFRYIADNDDDKMVILPV |
| IRIS_313-10722 | MLHHLKELAAVAGIHMILIYLCRFLLRRSRNVLFTVSNSLRFRLKVLTVLLYICLSVMLFYLFGSIMPLPPWGLVVGWVMALIAVELAYAFIFPYSFRYIADNDDDKMVILPV |
| IRIS_313-10723 | MLHHLKELAAVAGIHMILIYLCRFLLRRSRNVLFTVSNSLRFRLKVLTVLLYICLSVMLFYLFGSIMPLPPWGLVVGWVMALIAVELAYAFIFPYSFRYIADNDDDKMVILPV |
| IRIS_313-10724 | MLHHLKELAAVAGIHMILIYLCRFLLRRSRNVLFTVSNSLRFRLKVLTVLLYICLSVMLFYLFGSIMPLPPWGLVVGWVMALIAVELAYAFIFPYSFRYIADNDDDKMVILPV |
| IRIS_313-10725 | MLHHLKELAAVAGIHMILIYLCRFLLRRSRNVLFTVSNSLRFRLKVLTVLLYICLSVMLFYLFGSIMPLPPWGLVVGWVMALIAVELAYAFIFPYSFRYIADNDDDKMVILPV |
| IRIS_313-10726 | MLHHLKELAAVAGIHMILIYLCRFLLRRSRNVLFTVSNSLRFRLKVLTVLLYICLSVMLFYLFGSIMPLPPWGLVVGWVMALIAVELAYAFIFPYSFRYIADNDDDKMVILPV |
| IRIS_313-10727 | MLHHLKELAAVAGIHMILIYLCRFLLRRSRNVLFTVSNSLRFRLKVLTVLLYICLSVMLFYLFGSIMPLPPWGLVVGWVMALIAVELAYAFIFPYSFRYIADNDDDKMVILPV |
| IRIS_313-10728 | MLHHLKELAAVAGIHMILIYLCRFLLRRSRNVLFTVSNSLRFRLKVLTVLLYICLSVMLFYLFGSIMPLPPWGLVVGWVMALIAVELAYAFIFPYSFRYIADNDDDKMVILPV |
| IRIS_313-10730 | MLHHLKELAAVAGIHMILIYLCRFLLRRSRNVLFTVSNSLRFRLKVLTVLLYICLSVMLFYLFGSIMPLPPWGLVVGWVMALIAVELAYAFIFPYSFRYIADNDDDKMVILPV |
| IRIS_313-10731 | MLHHLKELAAVAGIHMILIYLCRFLLRRSRNVLFTVSNSLRFRLKVLTVLLYICLSVMLFYLFGSIMPLPPWGLVVGWVMALIAVELAYAFIFPYSFRYIADNDDDKMVILPV |
| IRIS_313-10732 | MLHHLKELAAVAGIHMILIYLCRFLLRRSRNVLFTVSNSLRFRLKVLTVLLYICLSVMLFYLFGSIMPLPPWGLVVGWVMALIAVELAYAFIFPYSFRYIADNDDDKMVILPV |
| IRIS_313-10733 | MLHHLKELAAVAGIHMILIYLCRFLLRRSRNVLFTVSNSLRFRLKVLTVLLYICLSVMLFYLFGSIMPLPPWGLVVGWVMALIAVELAYAFIFPYSFRYIADNDDDKMVILPV |
| IRIS_313-10734 | MLHHLKELAAVAGIHMILIYLCRFLLRRSRNVLFTVSNSLRFRLKVLTVLLYICLSVMLFYLFGSIMPLPPWGLVVGWVMALIAVELAYAFIFPYSFRYIADNDDDKMVILPV |
| IRIS_313-10735 | MLHHLKELAAVAGIHMILIYLCRFLLRRSRNVLFTVSNSLRFRLKVLTVLLYICLSVMLFYLFGSIMPLPPWGLVVGWVMALIAVELAYAFIFPYSFRYIADNDDDKMVILPV |
| IRIS_313-10736 | MLHHLKELAAVAGIHMILIYLCRFLLRRSRNVLFTVSNSLRFRLKVLTVLLYISLSAMLLYLFGSIMPLPPWGLVVGWVMALIAVELAYAFIFPYSFRYIADNDDDKMVILPV |
| IRIS_313-10737 | MLHHLKELAAVAGIHMILIYLCRFLLRRSRNVLFTVSNSLRFRLKVLTVLLYICLSVMLFYLFGSIMPLPPWGLVVGWVMALIAVELAYAFIFPYSFRYIADNDDDKMVILPV |
| IRIS_313-10738 | MLHHLKELAAVAGIHMILIYLCRFLLRRSRNVLFTVSNSLRFRLKVLTVLLYICLSVMLFYLFGSIMPLPPWGLVVGWVMALIAVELAYAFIFPYSFRYIADNDDDKMVILPV |
| IRIS_313-10739 | MLHHLKELAAVAGIHMILIYLCRFLLRRSRNVLFTVSNSLRFRLKVLTVLLYICLSVMLFYLFGSIMPLPPWGLVVGWVMALIAVELAYAFIFPYSFRYIADNDDDKMVILPV |
| IRIS_313-10740 | MLHHLKELAAVAGIHMILIYLCRFLLRRSRNVLFTVSNSLRFRLKVLTVLLYICLSVMLFYLFGSIMPLPPWGLVVGWVMALIAVELAYAFIFPYSFRYIADNDDDKMVILPV |
| IRIS_313-10741 | MLHHLKELAAVAGIHMILIYLCRFLLRRSRNVLFTVSNSLRFRLKVLTVLLYICLSVMLFYLFGSIMPLPPWGLVVGWVMALIAVELAYAFIFPYSFRYIADNDDDKMVILPV |
| IRIS_313-10742 | MLHHLKELAAVAGIHMILIYLCRFLLRRSRNVLFTVSNSLRFRLKVLTVLLYICLSVMLFYLFGSIMPLPPWGLVVGWVMALIAVELAYAFIFPYSFRYIADNDDDKMVILPV |
| IRIS_313-10743 | MLHHLKELAAVAGIHMILIYLCRFLLRRSRNVLFTVSNSLRFRLKVLTVLLYICLSVMLFYLFGSIMPLPPWGLVVGWVMALIAVELAYAFIFPYSFRYIADNDDDKMVILPV |
| IRIS_313-10744 | MLHQLKELAAVAGIHMILIYLCRFLLRRSRNVLFTVSNSLRFRLKVLTVLLYICLLVMLFYLFGSIMPLPPWGLVVGWVMALIAVELAYAFIFPYSFRYIADNDDDKMVILPV |
| IRIS_313-10745 | MLHHLKELAAVAGIHMILIYLCRFLLRRSRNVLFTVSNSLRFRLKVLTVLLYICLSVMLFYLFGSIMPLPPWGLVVGWVMALIAVELAYAFIFPYSFRYIADNDDDKMVILPV |
| IRIS_313-10746 | MLHHLKELAAVAGIHMILIYLCRFLLRRSRNVLFTVSNSLRFRLKVLTVLLYICLSVMLFYLFGSIMPLPPWGLVVGWVMALIAVELAYAFIFPYSFRYIADNDDDKMVILPV |
| IRIS_313-10747 | MLHHLKELAAVAGIHMILIYLCRFLLRRSRNVLFTVSNSLRFRLKVLTVLLYICLSVMLFYLFGSIMPLPPWGLVVGWVMALIAVELAYAFIFPYSFRYIADNDDDKMVILPV |
| IRIS_313-10748 | MLHHLKELAAVAGIHMILIYLCRFLLRRSRNVLFTVSNSLRFRLKVLTVLLYICLSVMLFYLFGSIMPLPPWGLVVGWVMALIAVELAYAFIFPYSFRYIADNDDDKMVILPV |
| IRIS_313-10749 | MLHHLKELAAVAGIHMILIYLCRFLLRRSRNVLFTVSNSLRFRLKVLTVLLYICLSVMLFYLFGSIMPLPPWGLVVGWVMALIAVELAYAFIFPYSFRYIADNDDDKMVILPV |
| IRIS_313-10750 | MLHHLKELAAVAGIHMILIYLCRFLLRRSRNVLFTVSNSLRFRLKVLTVLLYICLSVMLFYLFGSIMPLPPWGLVVGWVMALIAVELAYAFIFPYSFRYIADNDDDKMVILPV |
| IRIS_313-10751 | MLHHLKELAAVAGIHMILIYLCRFLLRRSRNVLFTVSNSLRFRLKVLTVLLYICLSVMLFYLFGSIMPLPPWGLVVGWVMALIAVELAYAFIFPYSFRYIADNDDDKMVILPV |
| IRIS_313-10752 | MLHHLKELAAVAGIHMILIYLCRFLLRRSRNVLFTVSNSLRFRLKVLTVLLYICLSVMLFYLFGSIMPLPPWGLVVGWVMALIAVELAYAFIFPYSFRYIADNDDDKMVILPV |
| IRIS_313-10753 | MLHHLKELAAVAGIHMILIYLCRFLLRRSRNVLFTVSNSLRFRLKVLTVLLYICLSVMLFYLFGSIMPLPPWGLVVGWVMALIAVELAYAFIFPYSFRYIADNDDDKMVILPV |
| IRIS_313-10754 | MLHHLKELAAVAGIHMILIYLCRFLLRRSRNVLFTVSNSLRFRLKVLTVLLYICLSVMLFYLFGSIMPLPPWGLVVGWVMALIAVELAYAFIFPYSFRYIADNDDDKMVILPV |
| IRIS_313-10755 | MLHHLKELAAVAGIHMILIYLCRFLLRRSRNVLFTVSNSLRFRLKVLTVLLYICLSVMLFYLFGSIMPLPPWGLVVGWVMALIAVELAYAFIFPYSFRYIADNDDDKMVILPV |
| IRIS_313-10756 | MLHHLKELAAVAGIHMILIYLCRFLLRRSRNVLFTVSNSLRFRLKVLTVLLYICLSVMLFYLFGSIMPLPPWGLVVGWVMALIAVELAYAFIFPYSFRYIADNDDDKMVILPV |
| IRIS_313-10757 | MLHHLKELAAVAGIHMILIYLCRFLLRRSRNVLFTVSNSLRFRLKVLTVLLYICLSVMLFYLFGSIMPLPPWGLVVGWVMALIAVELAYAFIFPYSFRYIADNDDDKMVILPV |
| IRIS_313-10758 | MLHHLKELAAVAGIHMILIYLCRFLLRRSRNVLFTVSNSLRFRLKVLTVLLYICLSVMLFYLFGSIMPLPPWGLVVGWVMALIAVELAYAFIFPYSFRYIADNDDDKMVILPV |
| IRIS_313-10759 | MLHHLKELAAVAGIHMILIYLCRFLLRRSRNVLFTVSNSLRFRLKVLTVLLYICLSVMLFYLFGSIMPLPPWGLVVGWVMALIAVELAYAFIFPYSFRYIADNDDDKMVILPV |
| IRIS_313-10760 | MLHHLKELAAVAGIHMILIYLCRFLLRRSRNVLFTVSNSLRFRLKVLTVLLYICLSVMLFYLFGSIMPLPPWGLVVGWVMALIAVELAYAFIFPYSFRYIADNDDDKMVILPV |
| IRIS_313-10761 | MLHHLKELAAVAGIHMILIYLCRFLLRRSRNVLFTVSNSLRFRLKVLTVLLYICLSVMLFYLFGSIMPLPPWGLVVGWVMALIAVELAYAFIFPYSFRYIADNDDDKMVILPV |
| IRIS_313-10762 | MLHHLKELAAVAGIHMILIYLCRFLLRRSRNVLFTVSNSLRFRLKVLTVLLYICLSVMLFYLFGSIMPLPPWGLVVGWVMALIAVELAYAFIFPYSFRYIADNDDDKMVILPV |
| IRIS_313-10765 | MLHHLKELAAVAGIHMILIYLCRFLLRRSRNVLFTVSNSLRFRLKVLTVLLYICLSVMLFYLFGSIMPLPPWGLVVGWVMALIAVELAYAFIFPYSFRYIADNDDDKMVILPV |
| IRIS_313-10766 | MLHHLKELAAVAGIHMILIYLCRFLLRRSRNVLFTVSNSLRFRLKVLTVLLYICLSVMLFYLFGSIMPLPPWGLVVGWVMALIAVELAYAFIFPYSFRYIADNDDDKMVILPV |
| IRIS_313-10767 | MLHHLKELAAVAGIHMILIYLCRFLLRRSRNVLFTVSNSLRFRLKVLTVLLYICLSVMLFYLFGSIMPLPPWGLVVGWVMALIAVELAYAFIFPYSFRYIADNDDDKMVILPV |
| IRIS_313-10768 | MLHHLKELAAVAGIHMILIYLCRFLLRRSRNVLFTVSNSLRFRLKVLTVLLYICLSVMLFYLFGSIMPLPPWGLVVGWVMALIAVELAYAFIFPYSFRYIADNDDDKMVILPV |
| IRIS_313-10769 | MLHHLKELAAVAGIHMILIYLCRFLLRRSRNVLFTVSNSLRFRLKVLTVLLYICLSVMLFYLFGSIMPLPPWGLVVGWVMALIAVELAYAFIFPYSFRYIADNDDDKMVILPV |
| IRIS_313-10770 | MLHHLKELAAVAGIHMILIYLCRFLLRRSRNVLFTVSNSLRFRLKVLTVLLYICLSVMLFYLFGSIMPLPPWGLVVGWVMALIAVELAYAFIFPYSFRYIADNDDDKMVILPV |
| IRIS_313-10771 | MLHHLKELAAVAGIHMILIYLCRFLLRRSRNVLFTVSNSLRFRLKVLTVLLYICLSVMLFYLFGSIMPLPPWGLVVGWVMALIAVELAYAFIFPYSFRYIADNDDDKMVILPV |
| IRIS_313-10772 | MLHHLKELAAVAGIHMILIYLCRFLLRRSRNVLFTVSNSLRFRLKVLTVLLYICLSVMLFYLFGSIMPLPPWGLVVGWVMALIAVELAYAFIFPYSFRYIADNDDDKMVILPV |
| IRIS_313-10773 | MLHHLKELAAVAGIHMILIYLCRFLLRRSRNVLFTVSNSLRFRLKVLTVLLYICLSVMLFYLFGSIMPLPPWGLVVGWVMALIAVELAYAFIFPYSFRYIADNDDDKMVILPV |
| IRIS_313-10774 | MLHHLKELAAVAGIHMILIYLCRFLLRRSRNVLFTVSNSLRFRLKVLTVLLYICLSVMLFYLFGSIMPLPPWGLVVGWVMALIAVELAYAFIFPYSFRYIADNDDDKMVILPV |
| IRIS_313-10775 | MLHHLKELAAVAGIHMILIYLCRFLLRRSRNVLFTVSNSLRFRLKVLTVLLYICLSVMLFYLFGSIMPLPPWGLVVGWVMALIAVELAYAFIFPYSFRYIADNDDDKMVILPV |
| IRIS_313-10776 | MLHHLKELAAVAGIHMILIYLCRFLLRRSRNVLFTVSNSLRFRLKVLTVLLYICLSVMLFYLFGSIMPLPPWGLVVGWVMALIAVELAYAFIFPYSFRYIADNDDDKMVILPV |
| IRIS_313-10777 | MLHHLKELAAVAGIHMILIYLCRFLLRRSRNVLFTVSNSLRFRLKVLTVLLYICLSVMLFYLFGSIMPLPPWGLVVGWVMALIAVELAYAFIFPYSFRYIADNDDDKMVILPV |
| IRIS_313-10778 | MLHHLKELAAVAGIHMILIYLCRFLLRRSRNVLFTVSNSLRFRLKVLTVLLYICLSVMLFYLFGSIMPLPPWGLVVGWVMALIAVELAYAFIFPYSFRYIADNDDDKMVILPV |
| IRIS_313-10779 | MLHHLKELAAVAGIHMILIYLCRFLLRRSRNVLFTVSNSLRFRLKVLTVLLYICLSVMLFYLFGSIMPLPPWGLVVGWVMALIAVELAYAFIFPYSFRYIADNDDDKMVILPV |
| IRIS_313-10780 | MLHHLKELAAVAGIHMILIYLCRFLLRRSRNVLFTVSNSLRFRLKVLTVLLYICLSVMLFYLFGSIMPLPPWGLVVGWVMALIAVELAYAFIFPYSFRYIADNDDDKMVILPV |
| IRIS_313-10781 | MLHHLKELAAVAGIHMILIYLCRFLLRRSRNVLFTVSNSLRFRLKVLTVLLYICLSVMLFYLFGSIMPLPPWGLVVGWVMALIAVELAYAFIFPYSFRYIADNDDDKMVILPV |
| IRIS_313-10782 | MLHHLKELAAVAGIHMILIYLCRFLLRRSRNVLFTVSNSLRFRLKVLTVLLYICLSVMLFYLFGSIMPLPPWGLVVGWVMALIAVELAYAFIFPYSFRYIADNDDDKMVILPV |
| IRIS_313-10783 | MLHHLKELAAVAGIHMILIYLCRFLLRRSRNVLFTVSNSLRFRLKVLTVLLYICLSVMLFYLFGSIMPLPPWGLVVGWVMALIAVELAYAFIFPYSFRYIADNDDDKMVILPV |
| IRIS_313-10784 | MLHHLKELAAVAGIHMILIYLCRFLLRRSRNVLFTVSNSLRFRLKVLTVLLYICLSVMLFYLFGSIMPLPPWGLVVGWVMALIAVELAYAFIFPYSFRYIADNDDDKMVILPV |
| IRIS_313-10785 | MLHHLKELAAVAGIHMILIYLCRFLLRRSRNVLFTVSNSLRFRLKVLTVLLYICLSVMLFYLFGSIMPLPPWGLVVGWVMALIAVELAYAFIFPYSFRYIADNDDDKMVILPV |
| IRIS_313-10786 | MLHHLKELAAVAGIHMILIYLCRFLLRRSRNVLFTVSNSLRFRLKVLTVLLYICLSVMLFYLFGSIMPLPPWGLVVGWVMALIAVELAYAFIFPYSFRYIADNDDDKMVILPV |
| IRIS_313-10787 | MLHHLKELAAVAGIHMILIYLCRFLLRRSRNVLFTVSNSLRFRLKVLTVLLYICLSVMLFYLFGSIMPLPPWGLVVGWVMALIAVELAYAFIFPYSFRYIADNDDDKMVILPV |
| IRIS_313-10788 | MLHHLKELAAVAGIHMILIYLCRFLLRRSRNVLFTVSNSLRFRLKVLTVLLYICLSVMLFYLFGSIMPLPPWGLVVGWVMALIAVELAYAFIFPYSFRYIADNDDDKMVILPV |
| IRIS_313-10789 | MLHHLKELAAVAGIHMILIYLCRFLLRRSRNVLFTVSNSLRFRLKVLTVLLYICLSVMLFYLFGSIMPLPPWGLVVGWVMALIAVELAYAFIFPYSFRYIADNDDDKMVILPV |
| IRIS_313-10790 | MLHHLKELAAVAGIHMILIYLCRFLLRRSRNVLFTVSNSLRFRLKVLTVLLYICLSVMLFYLFGSIMPLPPWGLVVGWVMALIAVELAYAFIFPYSFRYIADNDDDKMVILPV |
| IRIS_313-10791 | MLHHLKELAAVAGIHMILIYLCRFLLRRSRNVLFTVSNSLRFRLKVLTVLLYICLSVMLFYLFGSIMPLPPWGLVVGWVMALIAVELAYAFIFPYSFRYIADNDDDKMVILPV |
| IRIS_313-10793 | MLHHLKELAAVAGIHMILIYLCRFLLRRSRNVLFTVSNSLRFRLKVLTVLLYICLSVMLFYLFGSIMPLPPWGLVVGWVMALIAVELAYAFIFPYSFRYIADNDDDKMVILPV |
| IRIS_313-10794 | MLHHLKELAAVAGIHMILIYLCRFLLRRSRNVLFTVSNSLRFRLKVLTVLLYICLSVMLFYLFGSIMPLPPWGLVVGWVMALIAVELAYAFIFPYSFRYIADNDDDKMVILPV |
| IRIS_313-10795 | MLHHLKELAAVAGIHMILIYLCRFLLRRSRNVLFTVSNSLRFRLKVLTVLLYICLSVMLFYLFGSIMPLPPWGLVVGWVMALIAVELAYAFIFPYSFRYIADNDDDKMVILPV |
| IRIS_313-10796 | MLHHLKELAAVAGIHMILIYLCRFLLRRSRNVLFTVSNSLRFRLKVLTVLLYICLSVMLFYLFGSIMPLPPWGLVVGWVMALIAVELAYAFIFPYSFRYIADNDDDKMVILPV |
| IRIS_313-10797 | MLHHLKELAAVAGIHMILIYLCRFLLRRSRNVLFTVSNSLRFRLKVLTVLLYICLSVMLFYLFGSIMPLPPWGLVVGWVMALIAVELAYAFIFPYSFRYIADNDDDKMVILPV |
| IRIS_313-10798 | MLHHLKELAAVAGIHMILIYLCRFLLRRSRNVLFTVSNSLRFRLKVLTVLLYICLSVMLFYLFGSIMPLPPWGLVVGWVMALIAVELAYAFIFPYSFRYIADNDDDKMVILPV |
| IRIS_313-10799 | MLHHLKELAAVAGIHMILIYLCRFLLRRSRNVLFTVSNSLRFRLKVLTVLLYICLSVMLFYLFGSIMPLPPWGLVVGWVMALIAVELAYAFIFPYSFRYIADNDDDKMVILPV |
| IRIS_313-10800 | MLHHLKELAAVAGIHMILIYLCRFLLRRSRNVLFTVSNSLRFRLKVLTVLLYICLSVMLFYLFGSIMPLPPWGLVVGWVMALIAVELAYAFIFPYSFRYIADNDDDKMVILPV |
| IRIS_313-10801 | MLHHLKELAAVAGIHMILIYLCRFLLRRSRNVLFTVSNSLRFRLKVLTVLLYICLSVMLFYLFGSIMPLPPWGLVVGWVMALIAVELAYAFIFPYSFRYIADNDDDKMVILPV |
| IRIS_313-10802 | MLHHLKELAAVAGIHMILIYLCRFLLRRSRNVLFTVSNSLRFRLKVLTVLLYICLSVMLFYLFGSIMPLPPWGLVVGWVMALIAVELAYAFIFPYSFRYIADNDDDKMVILPV |
| IRIS_313-10803 | MLHHLKELAAVAGIHMILIYLCRFLLRRSRNVLFTVSNSLRFRLKVLTVLLYICLSVMLFYLFGSIMPLPPWGLVVGWVMALIAVELAYAFIFPYSFRYIADNDDDKMVILPV |
| IRIS_313-10804 | MLHHLKELAAVAGIHMILIYLCRFLLRRSRNVLFTVSNSLRFRLKVLTVLLYICLSVMLFYLFGSIMPLPPWGLVVGWVMALIAVELAYAFIFPYSFRYIADNDDDKMVILPV |
| IRIS_313-10805 | MLHHLKELAAVAGIHMILIYLCRFLLRRSRNVLFTVSNSLRFRLKVLTVLLYICLSVMLFYLFGSIMPLPPWGLVVGWVMALIAVELAYAFIFPYSFRYIADNDDDKMVILPV |
| IRIS_313-10806 | MLHHLKELAAVAGIHMILIYLCRFLLRRSRNVLFTVSNSLRFRLKVLTVLLYICLSVMLFYLFGSIMPLPPWGLVVGWVMALIAVELAYAFIFPYSFRYIADNDDDKMVILPV |
| IRIS_313-10807 | MLHHLKELAAVAGIHMILIYLCRFLLRRSRNVLFTVSNSLRFRLKVLTVLLYICLSVMLFYLFGSIMPLPPWGLVVGWVMALIAVELAYAFIFPYSFRYIADNDDDKMVILPV |
| IRIS_313-10808 | MLHHLKELAAVAGIHMILIYLCRFLLRRSRNVLFTVSNSLRFRLKVLTVLLYICLSVMLFYLFGSIMPLPPWGLVVGWVMALIAVELAYAFIFPYSFRYIADNDDDKMVILPV |
| IRIS_313-10809 | MLHHLKELAAVAGIHMILIYLCRFLLRRSRNVLFTVSNSLRFRLKVLTVLLYICLSVMLFYLFGSIMPLPPWGLVVGWVMALIAVELAYAFIFPYSFRYIADNDDDKMVILPV |
| IRIS_313-10810 | MLHHLKELAAVAGIHMILIYLCRFLLRRSRNVLFTVSNSLRFRLKVLTVLLYICLSVMLFYLFGSIMPLPPWGLVVGWVMALIAVELAYAFIFPYSFRYIADNDDDKMVILPV |
| IRIS_313-10811 | MLHHLKELAAVAGIHMILIYLCRFLLRRSRNVLFTVSNSLRFRLKVLTVLLYICLSVMLFYLFGSIMPLPPWGLVVGWVMALIAVELAYAFIFPYSFRYIADNDDDKMVILPV |
| IRIS_313-10812 | MLHHLKELAAVAGIHMILIYLCRFLLRRSRNVLFTVSNSLRFRLKVLTVLLYICLSVMLFYLFGSIMPLPPWGLVVGWVMALIAVELAYAFIFPYSFRYIADNDDDKMVILPV |
| IRIS_313-10813 | MLHHLKELAAVAGIHMILIYLCRFLLRRSRNVLFTVSNSLRFRLKVLTVLLYICLSVMLFYLFGSIMPLPPWGLVVGWVMALIAVELAYAFIFPYSFRYIADNDDDKMVILPV |
| IRIS_313-10814 | MLHHLKELAAVAGIHMILIYLCRFLLRRSRNVLFTVSNSLRFRLKVLTVLLYICLSVMLFYLFGSIMPLPPWGLVVGWVMALIAVELAYAFIFPYSFRYIADNDDDKMVILPV |
| IRIS_313-10815 | MLHHLKELAAVAGIHMILIYLCRFLLRRSRNVLFTVSNSLRFRLKVLTVLLYICLSVMLFYLFGSIMPLPPWGLVVGWVMALIAVELAYAFIFPYSFRYIADNDDDKMVILPV |
| IRIS_313-10816 | MLHHLKELAAVAGIHMILIYLCRFLLRRSRNVLFTVSNSLRFRLKVLTVLLYICLSVMLFYLFGSIMPLPPWGLVVGWVMALIAVELAYAFIFPYSFRYIADNDDDKMVILPV |
| IRIS_313-10817 | MLHQLKELAAVAGIHMILIYLCRFLLRRSRNVLFTVSNSLRFRLKVLTVLLYICLLVMLFYLFGSIMPLPPWGLVVGWVMALIAVELAYAFIFPYSFRYIADNDDDKMVILPV |
| IRIS_313-10818 | MLHHLKELAAVAGIHMILIYLCRFLLRRSRNVLFTVSNSLRFRLKVLTVLLYICLSVMLFYLFGSIMPLPPWGLVVGWVMALIAVELAYAFIFPYSFRYIADNDDDKMVILPV |
| IRIS_313-10819 | MLHHLKELAAVAGIHMILIYLCRFLLRRSRNVLFTVSNSLRFRLKVLTVLLYICLSVMLFYLFGSIMPLPPWGLVVGWVMALIAVELAYAFIFPYSFRYIADNDDDKMVILPV |
| IRIS_313-10820 | MLHHLKELAAVAGIHMILIYLCRFLLRRSRNVLFTVSNSLRFRLKVLTVLLYICLSVMLFYLFGSIMPLPPWGLVVGWVMALIAVELAYAFIFPYSFRYIADNDDDKMVILPV |
| IRIS_313-10821 | MLHHLKELAAVAGIHMILIYLCRFLLRRSRNVLFTVSNSLRFRLKVLTVLLYICLSVMLFYLFGSIMPLPPWGLVVGWVMALIAVELAYAFIFPYSFRYIADNDDDKMVILPV |
| IRIS_313-10822 | MLHHLKELAAVAGIHMILIYLCRFLLRRSRNVLFTVSNSLRFRLKVLTVLLYICLSVMLFYLFGSIMPLPPWGLVVGWVMALIAVELAYAFIFPYSFRYIADNDDDKMVILPV |
| IRIS_313-10823 | MLHHLKELAAVAGIHMILIYLCRFLLRRSRNVLFTVSNSLRFRLKVLTVLLYICLSVMLFYLFGSIMPLPPWGLVVGWVMALIAVELAYAFIFPYSFRYIADNDDDKMVILPV |
| IRIS_313-10824 | MLHHLKELAAVAGIHMILIYLCRFLLRRSRNVLFTVSNSLRFRLKVLTVLLYICLSVMLFYLFGSIMPLPPWGLVVGWVMALIAVELAYAFIFPYSFRYIADNDDDKMVILPV |
| IRIS_313-10825 | MLHHLKELAAVAGIHMILIYLCRFLLRRSRNVLFTVSNSLRFRLKVLTVLLYICLSVMLFYLFGSIMPLPPWGLVVGWVMALIAVELAYAFIFPYSFRYIADNDDDKMVILPV |
| IRIS_313-10827 | MLHHLKELAAVAGIHMILIYLCRFLLRRSRNVLFTVSNSLRFRLKVLTVLLYICLSVMLFYLFGSIMPLPPWGLVVGWVMALIAVELAYAFIFPYSFRYIADNDDDKMVILPV |
| IRIS_313-10828 | MLHQLKELAAVAGIHMILIYLCRFLLRRSRNVLFTVSNSLRFRLKVLTVLLYICLLVMLFYLFGSIMPLPPWGLVVGWVMALIAVELAYAFIFPYSFRYIADNDDDKMVILPV |
| IRIS_313-10829 | MLHQLKELAAVAGIHMILIYLCRFLLRRSRNVLFTVSNSLRFRLKVLTVLLYICLLVMLFYLFGSIMPLPPWGLVVGWVMALIAVELAYAFIFPYSFRYIADNDDDKMVILPV |

| | |
|---|---|
| IRIS_313-10830 | MLHQLKELAAVAGIHMILIYLCRFLLRRSRNVLFTVSNSLRFRLKVLTVLLYICLLVMLFYLFGSIMPLPPWGLVVGWVMALIAVELAYAFIFPYSFRYIADNDDDKMVILPV |
| IRIS_313-10831 | MLHHLKELAAVAGIHMILIYLCRFLLRRSRNVLFTVSNSLRFRLKVLTVLLYICLSVMLFYLFGSIMPLPPWGLVVGWVMALIAVELAYAFIFPYSFRYIADNDDDKMVILPV |
| IRIS_313-10832 | MLHHLKELAAVAGIHMILIYLCRFLLRRSRNVLFTVSNSLRFRLKVLTVLLYICLSVMLFYLFGSIMPLPPWGLVVGWVMALIAVELAYAFIFPYSFRYIADNDDDKMVILPV |
| IRIS_313-10833 | MLHHLKELAAVAGIHMILIYLCRFLLRRSRNVLFTVSNSLRFRLKVLTVLLYICLSVMLFYLFGSIMPLPPWGLVVGWVMALIAVELAYAFIFPYSFRYIADNDDDKMVILPV |
| IRIS_313-10834 | MLHHLKELAAVAGIHMILIYLCRFLLRRSRNVLFTVSNSLRFRLKVLTVLLYICLSVMLFYLFGSIMPLPPWGLVVGWVMALIAVELAYAFIFPYSFRYIADNDDDKMVILPV |
| IRIS_313-10835 | MLHHLKELAAVAGIHMILIYLCRFLLRRSRNVLFTVSNSLRFRLKVLTVLLYICLSVMLFYLFGSIMPLPPWGLVVGWVMALIAVELAYAFIFPYSFRYIADNDDDKMVILPV |
| IRIS_313-10836 | MLHHLKELAAVAGIHMILIYLCRFLLRRSRNVLFTVSNSLRFRLKVLTVLLYICLSVMLFYLFGSIMPLPPWGLVVGWVMALIAVELAYAFIFPYSFRYIADNDDDKMVILPV |
| IRIS_313-10837 | MLHHLKELAAVAGIHMILIYLCRFLLRRSRNVLFTVSNSLRFRLKVLTVLLYICLSVMLFYLFGSIMPLPPWGLVVGWVMALIAVELAYAFIFPYSFRYIADNDDDKMVILPV |
| IRIS_313-10838 | MLHHLKELAAVAGIHMILIYLCRFLLRRSRNVLFTVSNSLRFRLKVLTVLLYICLSVMLFYLFGSIMPLPPWGLVVGWVMALIAVELAYAFIFPYSFRYIADNDDDKMVILPV |
| IRIS_313-10839 | MLHHLKELAAVAGIHMILIYLCRFLLRRSRNVLFTVSNSLRFRLKVLTVLLYICLSVMLFYLFGSIMPLPPWGLVVGWVMALIAVELAYAFIFPYSFRYIADNDDDKMVILPV |
| IRIS_313-10840 | MLHHLKELAAVAGIHMILIYLCRFLLRRSRNVLFTVSNSLRFRLKVLTVLLYICLSVMLFYLFGSIMPLPPWGLVVGWVMALIAVELAYAFIFPYSFRYIADNDDDKMVILPV |
| IRIS_313-10841 | MLHHLKELAAVAGIHMILIYLCRFLLRRSRNVLFTVSNSLRFRLKVLTVLLYICLSVMLFYLFGSIMPLPPWGLVVGWVMALIAVELAYAFIFPYSFRYIADNDDDKMVILPV |
| IRIS_313-10842 | MLHHLKELAAVAGIHMILIYLCRFLLRRSRNVLFTVSNSLRFRLKVLTVLLYICLSVMLFYLFGSIMPLPPWGLVVGWVMALIAVELAYAFIFPYSFRYIADNDDDKMVILPV |
| IRIS_313-10844 | MLHHLKELAAVAGIHMILIYLCRFLLRRSRNVLFTVSNSLRFRLKVLTVLLYICLSVMLFYLFGSIMPLPPWGLVVGWVMALIAVELAYAFIFPYSFRYIADNDDDKMVILPV |
| IRIS_313-10845 | MLHHLKELAAVAGIHMILIYLCRFLLRRSRNVLFTVSNSLRFRLKVLTVLLYICLSVMLFYLFGSIMPLPPWGLVVGWVMALIAVELAYAFIFPYSFRYIADNDDDKMVILPV |
| IRIS_313-10847 | MLHHLKELAAVAGIHMILIYLCRFLLRRSRNVLFTVSNSLRFRLKVLTVLLYICLSVMLFYLFGSIMPLPPWGLVVGWVMALIAVELAYAFIFPYSFRYIADNDDDKMVILPV |
| IRIS_313-10848 | MLHHLKELAAVAGIHMILIYLCRFLLRRSRNVLFTVSNSLRFRLKVLTVLLYICLSVMLFYLFGSIMPLPPWGLVVGWVMALIAVELAYAFIFPYSFRYIADNDDDKMVILPV |
| IRIS_313-10849 | MLHHLKELAAVAGIHMILIYLCRFLLRRSRNVLFTVSNSLRFRLKVLTVLLYICLSAMLLYLFGSIMPLPPWGLVVGWVMALIAVELAYAFIFPYSFRYIADNDDDKMVILPV |
| IRIS_313-10850 | MLHHLKELAAVAGIHMILIYLCRFLLRRSRNVLFTVSNSLRFRLKVLTVLLYICLSVMLFYLFGSIMPLPPWGLVVGWVMALIAVELAYAFIFPYSFRYIADNDDDKMVILPV |
| IRIS_313-10851 | MLHHLKELAAVAGIHMILIYLCRFLLRRSRNVLFTVSNSLRFRLKVLTVLLYICLSVMLFYLFGSIMPLPPWGLVVGWVMALIAVELAYAFIFPYSFRYIADNDDDKMVILPV |
| IRIS_313-10852 | MLHHLKELAAVAGIHMILIYLCRFLLRRSRNVLFTVSNSLRFRLKVLTVLLYICLSVMLFYLFGSIMPLPPWGLVVGWVMALIAVELAYAFIFPYSFRYIADNDDDKMVILPV |
| IRIS_313-10853 | MLHHLKELAAVAGIHMILIYLCRFLLRRSRNVLFTVSNSLRFRLKVLTVLLYICLSVMLFYLFGSIMPLPPWGLVVGWVMALIAVELAYAFIFPYSFRYIADNDDDKMVILPV |
| IRIS_313-10854 | MLHHLKELAAVAGIHMILIYLCRFLLRRSRNVLFTVSNSLRFRLKVLTVLLYICLSVMLFYLFGSIMPLPPWGLVVGWVMALIAVELAYAFIFPYSFRYIADNDDDKMVILPV |
| IRIS_313-10855 | MLHHLKELAAVAGIHMILIYLCRFLLRRSRNVLFTVSNSLRFRLKVLTVLLYICLSVMLFYLFGSIMPLPPWGLVVGWVMALIAVELAYAFIFPYSFRYIADNDDDKMVILPV |
| IRIS_313-10856 | MLHHLKELAAVAGIHMILIYLCRFLLRRSRNVLFTVSNSLRFRLKVLTVLLYICLSVMLFYLFGSIMPLPPWGLVVGWVMALIAVELAYAFIFPYSFRYIADNDDDKMVILPV |
| IRIS_313-10857 | MLHHLKELAAVAGIHMILIYLCRFLLRRSRNVLFTVSNSLRFRLKVLTVLLYICLSVMLFYLFGSIMPLPPWGLVVGWVMALIAVELAYAFIFPYSFRYIADNDDDKMVILPV |
| IRIS_313-10858 | MLHHLKELAAVAGIHMILIYLCRFLLRRSRNVLFTVSNSLRFRLKVLTVLLYICLSVMLFYLFGSIMPLPPWGLVVGWVMALIAVELAYAFIFPYSFRYIADNDDDKMVILPV |
| IRIS_313-10859 | MLHHLKELAAVAGIHMILIYLCRFLLRRSRNVLFTVSNSLRFRLKVLTVLLYICLSVMLFYLFGSIMPLPPWGLVVGWVMALIAVELAYAFIFPYSFRYIADNDDDKMVILPV |
| IRIS_313-10861 | MLHHLKELAAVAGIHMILIYLCRFLLRRSRNVLFTVSNSLRFRLKVLTVLLYISLSAMLLYLFGSIMPLPPWGLVVGWVMALIAVELAYAFIFPYSFRYIADNDDDKMVILPV |
| IRIS_313-10862 | MLHHLKELAAVAGIHMILIYLCRFLLRRSRNVLFTVSNSLRFRLKVLTVLLYICLSVMLFYLFGSIMPLPPWGLVVGWVMALIAVELAYAFIFPYSFRYIADNDDDKMVILPV |
| IRIS_313-10863 | MLHHLKELAAVAGIHMILIYLCRFLLRRSRNVLFTVSNSLRFRLKVLTVLLYICLSVMLFYLFGSIMPLPPWGLVVGWVMALIAVELAYAFIFPYSFRYIADNDDDKMVILPV |
| IRIS_313-10864 | MLHHLKELAAVAGIHMILIYLCRFLLRRSRNVLFTVSNSLRFRLKVLTVLLYICLSVMLFYLFGSIMPLPPWGLVVGWVMALIAVELAYAFIFPYSFRYIADNDDDKMVILPV |
| IRIS_313-10865 | MLHHLKELAAVAGIHMILIYLCRFLLRRSRNVLFTVSNSLRFRLKVLTVLLYICLSVMLFYLFGSIMPLPPWGLVVGWVMALIAVELAYAFIFPYSFRYIADNDDDKMVILPV |
| IRIS_313-10866 | MLHHLKELAAVAGIHMILIYLCRFLLRRSRNVLFTVSNSLRFRLKVLTVLLYICLSVMLFYLFGSIMPLPPWGLVVGWVMALIAVELAYAFIFPYSFRYIADNDDDKMVILPV |
| IRIS_313-10867 | MLHHLKELAAVAGIHMILIYLCRFLLRRSRNVLFTVSNSLRFRLKVLTVLLYICLSVMLFYLFGSIMPLPPWGLVVGWVMALIAVELAYAFIFPYSFRYIADNDDDKMVILPV |
| IRIS_313-10868 | MLHHLKELAAVAGIHMILIYLCRFLLRRSRNVLFTVSNSLRFRLKVLTVLLYICLSVMLFYLFGSIMPLPPWGLVVGWVMALIAVELAYAFIFPYSFRYIADNDDDKMVILPV |
| IRIS_313-10869 | MLHHLKELAAVAGIHMILIYLCRFLLRRSRNVLFTVSNSLRFRLKVLTVLLYICLSVMLFYLFGSIMPLPPWGLVVGWVMALIAVELAYAFIFPYSFRYIADNDDDKMVILPV |
| IRIS_313-10870 | MLHQLKELAAVAGIHMILIYLCRFLLRRSRNVLFTVSNSLRFRLKVLTVLLYICLLVMLFYLFGSIMPLPPWGLVVGWVMALIAVELAYAFIFPYSFRYIADNDDDKMVILPV |
| IRIS_313-10871 | MLHHLKELAAVAGIHMILIYLCRFLLRRSRNVLFTVSNSLRFRLKVLTVLLYICLSVMLFYLFGSIMPLPPWGLVVGWVMALIAVELAYAFIFPYSFRYIADNDDDKMVILPV |
| IRIS_313-10872 | MLHQLKELAAVAGIHMILIYLCRFLLRRSRNVLFTVSNSLRFRLKVLTVLLYICLLVMLFYLFGSIMPLPPWGLVVGWVMALIAVELAYAFIFPYSFRYIADNDDDKMVILPV |
| IRIS_313-10873 | MLHHLKELAAVAGIHMILIYLCRFLLRRSRNVLFTVSNSLRFRLKVLTVLLYICLSVMLFYLFGSIMPLPPWGLVVGWVMALIAVELAYAFIFPYSFRYIADNDDDKMVILPV |
| IRIS_313-10874 | MLHHLKELAAVAGIHMILIYLCRFLLRRSRNVLFTVSNSLRFRLKVLTVLLYICLSVMLFYLFGSIMPLPPWGLVVGWVMALIAVELAYAFIFPYSFRYIADNDDDKMVILPV |
| IRIS_313-10875 | MLHHLKELAAVAGIHMILIYLCRFLLRRSRNVLFTVSNSLRFRLKVLTVLLYICLSVMLFYLFGSIMPLPPWGLVVGWVMALIAVELAYAFIFPYSFRYIADNDDDKMVILPV |
| IRIS_313-10876 | MLHHLKELAAVAGIHMILIYLCRFLLRRSRNVLFTVSNSLRFRLKVLTVLLYICLSVMLFYLFGSIMPLPPWGLVVGWVMALIAVELAYAFIFPYSFRYIADNDDDKMVILPV |
| IRIS_313-10877 | MLHHLKELAAVAGIHMILIYLCRFLLRRSRNVLFTVSNSLRFRLKVLTVLLYICLSVMLFYLFGSIMPLPPWGLVVGWVMALIAVELAYAFIFPYSFRYIADNDDDKMVILPV |
| IRIS_313-10878 | MLHHLKELAAVAGIHMILIYLCRFLLRRSRNVLFTVSNSLRFRLKVLTVLLYICLSVMLFYLFGSIMPLPPWGLVVGWVMALIAVELAYAFIFPYSFRYIADNDDDKMVILPV |
| IRIS_313-10879 | MLHHLKELAAVAGIHMILIYLCRFLLRRSRNVLFTVSNSLRFRLKVLTVLLYICLSVMLFYLFGSIMPLPPWGLVVGWVMALIAVELAYAFIFPYSFRYIADNDDDKMVILPV |
| IRIS_313-10880 | MLHHLKELAAVAGIHMILIYLCRFLLRRSRNVLFTVSNSLRFRLKVLTVLLYICLSVMLFYLFGSIMPLPPWGLVVGWVMALIAVELAYAFIFPYSFRYIADNDDDKMVILPV |
| IRIS_313-10881 | MLHHLKELAAVAGIHMILIYLCRFLLRRSRNVLFTVSNSLRFRLKVLTVLLYICLSVMLFYLFGSIMPLPPWGLVVGWVMALIAVELAYAFIFPYSFRYIADNDDDKMVILPV |
| IRIS_313-10882 | MLHHLKELAAVAGIHMILIYLCRFLLRRSRNVLFTVSNSLRFRLKVLTVLLYICLSVMLFYLFGSIMPLPPWGLVVGWVMALIAVELAYAFIFPYSFRYIADNDDDKMVILPV |
| IRIS_313-10883 | MLHHLKELAAVAGIHMILIYLCRFLLRRSRNVLFTVSNSLRFRLKVLTVLLYICLSVMLFYLFGSIMPLPPWGLVVGWVMALIAVELAYAFIFPYSFRYIADNDDDKMVILPV |
| IRIS_313-10884 | MLHHLKELAAVAGIHMILIYLCRFLLRRSRNVLFTVSNSLRFRLKVLTVLLYICLSVMLFYLFGSIMPLPPWGLVVGWVMALIAVELAYAFIFPYSFRYIADNDDDKMVILPV |
| IRIS_313-10885 | MLHHLKELAAVAGIHMILIYLCRFLLRRSRNVLFTVSNSLRFRLKVLTVLLYICLSVMLFYLFGSIMPLPPWGLVVGWVMALIAVELAYAFIFPYSFRYIADNDDDKMVILPV |
| IRIS_313-10886 | MLHHLKELAAVAGIHMILIYLCRFLLRRSRNVLFTVSNSLRFRLKVLTVLLYICLSVMLFYLFGSIMPLPPWGLVVGWVMALIAVELAYAFIFPYSFRYIADNDDDKMVILPV |
| IRIS_313-10887 | MLHHLKELAAVAGIHMILIYLCRFLLRRSRNVLFTVSNSLRFRLKVLTVLLYICLSVMLFYLFGSIMPLPPWGLVVGWVMALIAVELAYAFIFPYSFRYIADNDDDKMVILPV |
| IRIS_313-10888 | MLHHLKELAAVAGIHMILIYLCRFLLRRSRNVLFTVSNSLRFRLKVLTVLLYICLSVMLFYLFGSIMPLPPWGLVVGWVMALIAVELAYAFIFPYSFRYIADNDDDKMVILPV |
| IRIS_313-10889 | MLHHLKELAAVAGIHMILIYLCRFLLRRSRNVLFTVSNSLRFRLKVLTVLLYICLSVMLFYLFGSIMPLPPWGLVVGWVMALIAVELAYAFIFPYSFRYIADNDDDKMVILPV |
| IRIS_313-10890 | MLHQLKELAAVAGIHMILIYLCRFLLRRSRNVLFTVSNSLRFRLKVLTVLLYICLLVMLFYLFGSIMPLPPWGLVVGWVMALIAVELAYAFIFPYSFRYIADNDDDKMVILPV |
| IRIS_313-10891 | MLHHLKELAAVAGIHMILIYLCRFLLRRSRNVLFTVSNSLRFRLKVLTVLLYICLSVMLFYLFGSIMPLPPWGLVVGWVMALIAVELAYAFIFPYSFRYIADNDDDKMVILPV |
| IRIS_313-10892 | MLHHLKELAAVAGIHMILIYLCRFLLRRSRNVLFTVSNSLRFRLKVLTVLLYICLSVMLFYLFGSIMPLPPWGLVVGWVMALIAVELAYAFIFPYSFRYIADNDDDKMVILPV |
| IRIS_313-10893 | MLHHLKELAAVAGIHMILIYLCRFLLRRSRNVLFTVSNSLRFRLKVLTVLLYICLSVMLFYLFGSIMPLPPWGLVVGWVMALIAVELAYAFIFPYSFRYIADNDDDKMVILPV |
| IRIS_313-10894 | MLHHLKELAAVAGIHMILIYLCRFLLRRSRNVLFTVSNSLRFRLKVLTVLLYICLSVMLFYLFGSIMPLPPWGLVVGWVMALIAVELAYAFIFPYSFRYIADNDDDKMVILPV |
| IRIS_313-10895 | MLHHLKELAAVAGIHMILIYLCRFLLRRSRNVLFTVSNSLRFRLKVLTVLLYICLSVMLFYLFGSIMPLPPWGLVVGWVMALIAVELAYAFIFPYSFRYIADNDDDKMVILPV |
| IRIS_313-10896 | MLHHLKELAAVAGIHMILIYLCRFLLRRSRNVLFTVSNSLRFRLKVLTVLLYICLSVMLFYLFGSIMPLPPWGLVVGWVMALIAVELAYAFIFPYSFRYIADNDDDKMVILPV |
| IRIS_313-10897 | MLHHLKELAAVAGIHMILIYLCRFLLRRSRNVLFTVSNSLRFRLKVLTVLLYICLSVMLFYLFGSIMPLPPWGLVVGWVMALIAVELAYAFIFPYSFRYIADNDDDKMVILPV |
| IRIS_313-10898 | MLHHLKELAAVAGIHMILIYLCRFLLRRSRNVLFTVSNSLRFRLKVLTVLLYICLSVMLFYLFGSIMPLPPWGLVVGWVMALIAVELAYAFIFPYSFRYIADNDDDKMVILPV |
| IRIS_313-10899 | MLHHLKELAAVAGIHMILIYLCRFLLRRSRNVLFTVSNSLRFRLKVLTVLLYICLSVMLFYLFGSIMPLPPWGLVVGWVMALIAVELAYAFIFPYSFRYIADNDDDKMVILPV |
| IRIS_313-10900 | MLHHLKELAAVAGIHMILIYLCRFLLRRSRNVLFTVSNSLRFRLKVLTVLLYICLSVMLFYLFGSIMPLPPWGLVVGWVMALIAVELAYAFIFPYSFRYIADNDDDKMVILPV |
| IRIS_313-10901 | MLHHLKELAAVAGIHMILIYLCRFLLRRSRNVLFTVSNSLRFRLKVLTVLLYICLSVMLFYLFGSIMPLPPWGLVVGWVMALIAVELAYAFIFPYSFRYIADNDDDKMVILPV |
| IRIS_313-10902 | MLHHLKELAAVAGIHMILIYLCRFLLRRSRNVLFTVSNSLRFRLKVLTVLLYICLSVMLFYLFGSIMPLPPWGLVVGWVMALIAVELAYAFIFPYSFRYIADNDDDKMVILPV |
| IRIS_313-10903 | MLHHLKELAAVAGIHMILIYLCRFLLRRSRNVLFTVSNSLRFRLKVLTVLLYICLSVMLFYLFGSIMPLPPWGLVVGWVMALIAVELAYAFIFPYSFRYIADNDDDKMVILPV |
| IRIS_313-10904 | MLHHLKELAAVAGIHMILIYLCRFLLRRSRNVLFTVSNSLRFRLKVLTVLLYICLSVMLFYLFGSIMPLPPWGLVVGWVMALIAVELAYAFIFPYSFRYIADNDDDKMVILPV |
| IRIS_313-10905 | MLHHLKELAAVAGIHMILIYLCRFLLRRSRNVLFTVSNSLRFRLKVLTVLLYICLSVMLFYLFGSIMPLPPWGLVVGWVMALIAVELAYAFIFPYSFRYIADNDDDKMVILPV |
| IRIS_313-10906 | MLHHLKELAAVAGIHMILIYLCRFLLRRSRNVLFTVSNSLRFRLKVLTVLLYICLSVMLFYLFGSIMPLPPWGLVVGWVMALIAVELAYAFIFPYSFRYIADNDDDKMVILPV |
| IRIS_313-10907 | MLHHLKELAAVAGIHMILIYLCRFLLRRSRNVLFTVSNSLRFRLKVLTVLLYICLSVMLFYLFGSIMPLPPWGLVVGWVMALIAVELAYAFIFPYSFRYIADNDDDKMVILPV |
| IRIS_313-10908 | MLHHLKELAAVAGIHMILIYLCRFLLRRSRNVLFTVSNSLRFRLKVLTVLLYICLSVMLFYLFGSIMPLPPWGLVVGWVMALIAVELAYAFIFPYSFRYIADNDDDKMVILPV |
| IRIS_313-10909 | MLHHLKELAAVAGIHMILIYLCRFLLRRSRNVLFTVSNSLRFRLKVLTVLLYICLSVMLFYLFGSIMPLPPWGLVVGWVMALIAVELAYAFIFPYSFRYIADNDDDKMVILPV |
| IRIS_313-10910 | MLHHLKELAAVAGIHMILIYLCRFLLRRSRNVLFTVSNSLRFRLKVLTVLLYICLSVMLFYLFGSIMPLPPWGLVVGWVMALIAVELAYAFIFPYSFRYIADNDDDKMVILPV |
| IRIS_313-10911 | MLHHLKELAAVAGIHMILIYLCRFLLRRSRNVLFTVSNSLRFRLKVLTVLLYICLSVMLFYLFGSIMPLPPWGLVVGWVMALIAVELAYAFIFPYSFRYIADNDDDKMVILPV |
| IRIS_313-10912 | MLHHLKELAAVAGIHMILIYLCRFLLRRSRNVLFTVSNSLRFRLKVLTVLLYICLSVMLFYLFGSIMPLPPWGLVVGWVMALIAVELAYAFIFPYSFRYIADNDDDKMVILPV |
| IRIS_313-10913 | MLHHLKELAAVAGIHMILIYLCRFLLRRSRNVLFTVSNSLRFRLKVLTVLLYICLSVMLFYLFGSIMPLPPWGLVVGWVMALIAVELAYAFIFPYSFRYIADNDDDKMVILPV |
| IRIS_313-10915 | MLHHLKELAAVAGIHMILIYLCRFLLRRSRNVLFTVSNSLRFRLKVLTVLLYICLSVMLFYLFGSIMPLPPWGLVVGWVMALIAVELAYAFIFPYSFRYIADNDDDKMVILPV |
| IRIS_313-10916 | MLHHLKELAAVAGIHMILIYLCRFLLRRSRNVLFTVSNSLRFRLKVLTVLLYICLSVMLFYLFGSIMPLPPWGLVVGWVMALIAVELAYAFIFPYSFRYIADNDDDKMVILPV |
| IRIS_313-10917 | MLHHLKELAAVAGIHMILIYLCRFLLRRSRNVLFTVSNSLRFRLKVLTVLLYICLSVMLFYLFGSIMPLPPWGLVVGWVMALIAVELAYAFIFPYSFRYIADNDDDKMVILPV |
| IRIS_313-10918 | MLHHLKELAAVAGIHMILIYLCRFLLRRSRNVLFTVSNSLRFRLKVLTVLLYICLSVMLFYLFGSIMPLPPWGLVVGWVMALIAVELAYAFIFPYSFRYIADNDDDKMVILPV |
| IRIS_313-10919 | MLHHLKELAAVAGIHMILIYLCRFLLRRSRNVLFTVSNSLRFRLKVLTVLLYICLSVMLFYLFGSIMPLPPWGLVVGWVMALIAVELAYAFIFPYSFRYIADNDDDKMVILPV |
| IRIS_313-10920 | MLHHLKELAAVAGIHMILIYLCRFLLRRSRNVLFTVSNSLRFRLKVLTVLLYICLSVMLFYLFGSIMPLPPWGLVVGWVMALIAVELAYAFIFPYSFRYIADNDDDKMVILPV |
| IRIS_313-10921 | MLHHLKELAAVAGIHMILIYLCRFLLRRSRNVLFTVSNSLRFRLKVLTVLLYICLSVMLFYLFGSIMPLPPWGLVVGWVMALIAVELAYAFIFPYSFRYIADNDDDKMVILPV |
| IRIS_313-10922 | MLHHLKELAAVAGIHMILIYLCRFLLRRSRNVLFTVSNSLRFRLKVLTVLLYICLSVMLFYLFGSIMPLPPWGLVVGWVMALIAVELAYAFIFPYSFRYIADNDDDKMVILPV |
| IRIS_313-10923 | MLHHLKELAAVAGIHMILIYLCRFLLRRSRNVLFTVSNSLRFRLKVLTVLLYICLSVMLFYLFGSIMPLPPWGLVVGWVMALIAVELAYAFIFPYSFRYIADNDDDKMVILPV |
| IRIS_313-10924 | MLHHLKELAAVAGIHMILIYLCRFLLRRSRNVLFTVSNSLRFRLKVLTVLLYICLSVMLFYLFGSIMPLPPWGLVVGWVMALIAVELAYAFIFPYSFRYIADNDDDKMVILPV |
| IRIS_313-10925 | MLHHLKELAAVAGIHMILIYLCRFLLRRSRNVLFTVSNSLRFRLKVLTVLLYICLSVMLFYLFGSIMPLPPWGLVVGWVMALIAVELAYAFIFPYSFRYIADNDDDKMVILPV |
| IRIS_313-10926 | MLHHLKELAAVAGIHMILIYLCRFLLRRSRNVLFTVSNSLRFRLKVLTVLLYICLSVMLFYLFGSIMPLPPWGLVVGWVMALIAVELAYAFIFPYSFRYIADNDDDKMVILPV |
| IRIS_313-10927 | MLHHLKELAAVAGIHMILIYLCRFLLRRSRNVLFTVSNSLRFRLKVLTVLLYICLSVMLFYLFGSIMPLPPWGLVVGWVMALIAVELAYAFIFPYSFRYIADNDDDKMVILPV |
| IRIS_313-10928 | MLHHLKELAAVAGIHMILIYLCRFLLRRSRNVLFTVSNSLRFRLKVLTVLLYICLSVMLFYLFGSIMPLPPWGLVVGWVMALIAVELAYAFIFPYSFRYIADNDDDKMVILPV |
| IRIS_313-10929 | MLHHLKELAAVAGIHMILIYLCRFLLRRSRNVLFTVSNSLRFRLKVLTVLLYICLSVMLFYLFGSIMPLPPWGLVVGWVMALIAVELAYAFIFPYSFRYIADNDDDKMVILPV |
| IRIS_313-10930 | MLHHLKELAAVAGIHMILIYLCRFLLRRSRNVLFTVSNSLRFRLKVLTVLLYICLSVMLFYLFGSIMPLPPWGLVVGWVMALIAVELAYAFIFPYSFRYIADNDDDKMVILPV |
| IRIS_313-10931 | MLHHLKELAAVAGIHMILIYLCRFLLRRSRNVLFTVSNSLRFRLKVLTVLLYICLSVMLFYLFGSIMPLPPWGLVVGWVMALIAVELAYAFIFPYSFRYIADNDDDKMVILPV |
| IRIS_313-10932 | MLHHLKELAAVAGIHMILIYLCRFLLRRSRNVLFTVSNSLRFRLKVLTVLLYICLSVMLFYLFGSIMPLPPWGLVVGWVMALIAVELAYAFIFPYSFRYIADNDDDKMVILPV |
| IRIS_313-10933 | MLHHLKELAAVAGIHMILIYLCRFLLRRSRNVLFTVSNSLRFRLKVLTVLLYICLSVMLFYLFGSIMPLPPWGLVVGWVMALIAVELAYAFIFPYSFRYIADNDDDKMVILPV |
| IRIS_313-10934 | MLHHLKELAAVAGIHMILIYLCRFLLRRSRNVLFTVSNSLRFRLKVLTVLLYICLSVMLFYLFGSIMPLPPWGLVVGWVMALIAVELAYAFIFPYSFRYIADNDDDKMVILPV |
| IRIS_313-10935 | MLHHLKELAAVAGIHMILIYLCRFLLRRSRNVLFTVSNSLRFRLKVLTVLLYICLSVMLFYLFGSIMPLPPWGLVVGWVMALIAVELAYAFIFPYSFRYIADNDDDKMVILPV |
| IRIS_313-10936 | MLHHLKELAAVAGIHMILIYLCRFLLRRSRNVLFTVSNSLRFRLKVLTVLLYICLSVMLFYLFGSIMPLPPWGLVVGWVMALIAVELAYAFIFPYSFRYIADNDDDKMVILPV |
| IRIS_313-10937 | MLHHLKELAAVAGIHMILIYLCRFLLRRSRNVLFTVSNSLRFRLKVLTVLLYICLSVMLFYLFGSIMPLPPWGLVVGWVMALIAVELAYAFIFPYSFRYIADNDDDKMVILPV |
| IRIS_313-10938 | MLHHLKELAAVAGIHMILIYLCRFLLRRSRNVLFTVSNSLRFRLKVLTVLLYICLSVMLFYLFGSIMPLPPWGLVVGWVMALIAVELAYAFIFPYSFRYIADNDDDKMVILPV |
| IRIS_313-10940 | MLHHLKELAAVAGIHMILIYLCRFLLRRSRNVLFTVSNSLRFRLKVLTVLLYICLSVMLFYLFGSIMPLPPWGLVVGWVMALIAVELAYAFIFPYSFRYIADNDDDKMVILPV |
| IRIS_313-10941 | MLHHLKELAAVAGIHMILIYLCRFLLRRSRNVLFTVSNSLRFRLKVLTVLLYICLSVMLFYLFGSIMPLPPWGLVVGWVMALIAVELAYAFIFPYSFRYIADNDDDKMVILPV |
| IRIS_313-10942 | MLHHLKELAAVAGIHMILIYLCRFLLRRSRNVLFTVSNSLRFRLKVLTVLLYICLSVMLFYLFGSIMPLPPWGLVVGWVMALIAVELAYAFIFPYSFRYIADNDDDKMVILPV |
| IRIS_313-10943 | MLHHLKELAAVAGIHMILIYLCRFLLRRSRNVLFTVSNSLRFRLKVLTVLLYICLSVMLFYLFGSIMPLPPWGLVVGWVMALIAVELAYAFIFPYSFRYIADNDDDKMVILPV |
| IRIS_313-10944 | MLHHLKELAAVAGIHMILIYLCRFLLRRSRNVLFTVSNSLRFRLKVLTVLLYICLSVMLFYLFGSIMPLPPWGLVVGWVMALIAVELAYAFIFPYSFRYIADNDDDKMVILPV |
| IRIS_313-10945 | MLHQLKELAAVAGIHMILIYLCRFLLRRSRNVLFTVSNSLRFRLKVLTVLLYICLLVMLFYLFGSIMPLPPWGLVVGWVMALIAVELAYAFIFPYSFRYIADNDDDKMVILPV |
| IRIS_313-10946 | MLHHLKELAAVAGIHMILIYLCRFLLRRSRNVLFTVSNSLRFRLKVLTVLLYICLSVMLFYLFGSIMPLPPWGLVVGWVMALIAVELAYAFIFPYSFRYIADNDDDKMVILPV |
| IRIS_313-10947 | MLHHLKELAAVAGIHMILIYLCRFLLRRSRNVLFTVSNSLRFRLKVLTVLLYICLSVMLFYLFGSIMPLPPWGLVVGWVMALIAVELAYAFIFPYSFRYIADNDDDKMVILPV |
| IRIS_313-10948 | MLHHLKELAAVAGIHMILIYLCRFLLRRSRNVLFTVSNSLRFRLKVLTVLLYICLSVMLFYLFGSIMPLPPWGLVVGWVMALIAVELAYAFIFPYSFRYIADNDDDKMVILPV |
| IRIS_313-10949 | MLHHLKELAAVAGIHMILIYLCRFLLRRSRNVLFTVSNSLRFRLKVLTVLLYICLSVMLFYLFGSIMPLPPWGLVVGWVMALIAVELAYAFIFPYSFRYIADNDDDKMVILPV |
| IRIS_313-10950 | MLHHLKELAAVAGIHMILIYLCRFLLRRSRNVLFTVSNSLRFRLKVLTVLLYICLSVMLFYLFGSIMPLPPWGLVVGWVMALIAVELAYAFIFPYSFRYIADNDDDKMVILPV |
| IRIS_313-10951 | MLHHLKELAAVAGIHMILIYLCRFLLRRSRNVLFTVSNSLRFRLKVLTVLLYICLSVMLFYLFGSIMPLPPWGLVVGWVMALIAVELAYAFIFPYSFRYIADNDDDKMVILPV |
| IRIS_313-10952 | MLHHLKELAAVAGIHMILIYLCRFLLRRSRNVLFTVSNSLRFRLKVLTVLLYICLSVMLFYLFGSIMPLPPWGLVVGWVMALIAVELAYAFIFPYSFRYIADNDDDKMVILPV |
| IRIS_313-10953 | MLHHLKELAAVAGIHMILIYLCRFLLRRSRNVLFTVSNSLRFRLKVLTVLLYICLSVMLFYLFGSIMPLPPWGLVVGWVMALIAVELAYAFIFPYSFRYIADNDDDKMVILPV |
| IRIS_313-10954 | MLHHLKELAAVAGIHMILIYLCRFLLRRSRNVLFTVSNSLRFRLKVLTVLLYICLSVMLFYLFGSIMPLPPWGLVVGWVMALIAVELAYAFIFPYSFRYIADNDDDKMVILPV |
| IRIS_313-10955 | MLHHLKELAAVAGIHMILIYLCRFLLRRSRNVLFTVSNSLRFRLKVLTVLLYICLSVMLFYLFGSIMPLPPWGLVVGWVMALIAVELAYAFIFPYSFRYIADNDDDKMVILPV |
| IRIS_313-10956 | MLHHLKELAAVAGIHMILIYLCRFLLRRSRNVLFTVSNSLRFRLKVLTVLLYICLSVMLFYLFGSIMPLPPWGLVVGWVMALIAVELAYAFIFPYSFRYIADNDDDKMVILPV |
| IRIS_313-10957 | MLHQLKELAAVAGIHMILIYLCRFLLRRSRNVLFTVSNSLRFRLKVLTVLLYICLLVMLFYLFGSIMPLPPWGLVVGWVMALIAVELAYAFIFPYSFRYIADNDDDKMVILPV |

| | |
|---|---|
| IRIS_313-10958 | MLHHLKELAAVAGIHMILIYLCRFLLRRSRNVLFTVSNSLRFRLKVLTVLLYICLSVMLFYLFGSIMPLPPWGLVVGWVMALIAVELAYAFIFPYSFRYIADNDDDKMVILPV |
| IRIS_313-10959 | MLHHLKELAAVAGIHMILIYLCRFLLRRSRNVLFTVSNSLRFRLKVLTVLLYICLSVMLFYLFGSIMPLPPWGLVVGWVMALIAVELAYAFIFPYSFRYIADNDDDKMVILPV |
| IRIS_313-10960 | MLHHLKELAAVAGIHMILIYLCRFLLRRSRNVLFTVSNSLRFRLKVLTVLLYICLSVMLFYLFGSIMPLPPWGLVVGWVMALIAVELAYAFIFPYSFRYIADNDDDKMVILPV |
| IRIS_313-10961 | MLHHLKELAAVAGIHMILIYLCRFLLRRSRNVLFTVSNSLRFRLKVLTVLLYICLSVMLFYLFGSIMPLPPWGLVVGWVMALIAVELAYAFIFPYSFRYIADNDDDKMVILPV |
| IRIS_313-10962 | MLHHLKELAAVAGIHMILIYLCRFLLRRSRNVLFTVSNSLRFRLKVLTVLLYICLSVMLFYLFGSIMPLPPWGLVVGWVMALIAVELAYAFIFPYSFRYIADNDDDKMVILPV |
| IRIS_313-10963 | MLHHLKELAAVAGIHMILIYLCRFLLRRSRNVLFTVSNSLRFRLKVLTVLLYICLSVMLFYLFGSIMPLPPWGLVVGWVMALIAVELAYAFIFPYSFRYIADNDDDKMVILPV |
| IRIS_313-10964 | MLHHLKELAAVAGIHMILIYLCRFLLRRSRNVLFTVSNSLRFRLKVLTVLLYICLSVMLFYLFGSIMPLPPWGLVVGWVMALIAVELAYAFIFPYSFRYIADNDDDKMVILPV |
| IRIS_313-10965 | MLHHLKELAAVAGIHMILIYLCRFLLRRSRNVLFTVSNSLRFRLKVLTVLLYICLSVMLFYLFGSIMPLPPWGLVVGWVMALIAVELAYAFIFPYSFRYIADNDDDKMVILPV |
| IRIS_313-10966 | MLHHLKELAAVAGIHMILIYLCRFLLRRSRNVLFTVSNSLRFRLKVLTVLLYICLSVMLFYLFGSIMPLPPWGLVVGWVMALIAVELAYAFIFPYSFRYIADNDDDKMVILPV |
| IRIS_313-10967 | MLHHLKELAAVAGIHMILIYLCRFLLRRSRNVLFTVSNSLRFRLKVLTVLLYICLSVMLFYLFGSIMPLPPWGLVVGWVMALIAVELAYAFIFPYSFRYIADNDDDKMVILPV |
| IRIS_313-10968 | MLHHLKELAAVAGIHMILIYLCRFLLRRSRNVLFTVSNSLRFRLKVLTVLLYICLSVMLFYLFGSIMPLPPWGLVVGWVMALIAVELAYAFIFPYSFRYIADNDDDKMVILPV |
| IRIS_313-10969 | MLHHLKELAAVAGIHMILIYLCRFLLRRSRNVLFTVSNSLRFRLKVLTVLLYICLSVMLFYLFGSIMPLPPWGLVVGWVMALIAVELAYAFIFPYSFRYIADNDDDKMVILPV |
| IRIS_313-10970 | MLHHLKELAAVAGIHMILIYLCRFLLRRSRNVLFTVSNSLRFRLKVLTVLLYICLSVMLFYLFGSIMPLPPWGLVVGWVMALIAVELAYAFIFPYSFRYIADNDDDKMVILPV |
| IRIS_313-10971 | MLHHLKELAAVAGIHMILIYLCRFLLRRSRNVLFTVSNSLRFRLKVLTVLLYICLSVMLFYLFGSIMPLPPWGLVVGWVMALIAVELAYAFIFPYSFRYIADNDDDKMVILPV |
| IRIS_313-10972 | MLHHLKELAAVAGIHMILIYLCRFLLRRSRNVLFTVSNSLRFRLKVLTVLLYICLSVMLFYLFGSIMPLPPWGLVVGWVMALIAVELAYAFIFPYSFRYIADNDDDKMVILPV |
| IRIS_313-10973 | MLHHLKELAAVAGIHMILIYLCRFLLRRSRNVLFTVSNSLRFRLKVLTVLLYICLSVMLFYLFGSIMPLPPWGLVVGWVMALIAVELAYAFIFPYSFRYIADNDDDKMVILPV |
| IRIS_313-10974 | MLHHLKELAAVAGIHMILIYLCRFLLRRSRNVLFTVSNSLRFRLKVLTVLLYICLSVMLFYLFGSIMPLPPWGLVVGWVMALIAVELAYAFIFPYSFRYIADNDDDKMVILPV |
| IRIS_313-10975 | MLHHLKELAAVAGIHMILIYLCRFLLRRSRNVLFTVSNSLRFRLKVLTVLLYICLSVMLFYLFGSIMPLPPWGLVVGWVMALIAVELAYAFIFPYSFRYIADNDDDKMVILPV |
| IRIS_313-10976 | MLHHLKELAAVAGIHMILIYLCRFLLRRSRNVLFTVSNSLRFRLKVLTVLLYICLSVMLFYLFGSIMPLPPWGLVVGWVMALIAVELAYAFIFPYSFRYIADNDDDKMVILPV |
| IRIS_313-10977 | MLHHLKELAAVAGIHMILIYLCRFLLRRSRNVLFTVSNSLRFRLKVLTVLLYICLSVMLFYLFGSIMPLPPWGLVVGWVMALIAVELAYAFIFPYSFRYIADNDDDKMVILPV |
| IRIS_313-10978 | MLHHLKELAAVAGIHMILIYLCRFLLRRSRNVLFTVSNSLRFRLKVLTVLLYICLSVMLFYLFGSIMPLPPWGLVVGWVMALIAVELAYAFIFPYSFRYIADNDDDKMVILPV |
| IRIS_313-10979 | MLHHLKELAAVAGIHMILIYLCRFLLRRSRNVLFTVSNSLRFRLKVLTVLLYICLSVMLFYLFGSIMPLPPWGLVVGWVMALIAVELAYAFIFPYSFRYIADNDDDKMVILPV |
| IRIS_313-10980 | MLHHLKELAAVAGIHMILIYLCRFLLRRSRNVLFTVSNSLRFRLKVLTVLLYICLSVMLFYLFGSIMPLPPWGLVVGWVMALIAVELAYAFIFPYSFRYIADNDDDKMVILPV |
| IRIS_313-10981 | MLHHLKELAAVAGIHMILIYLCRFFLRRSRNVLFTVSNSLRFRLKVLTVLLYICLSVMLFYLFGSIMPLPPWGLVVGWVMALIAVELAYAFIFPYSFRYIADNDDDKMVILPV |
| IRIS_313-10982 | MLHHLKELAAVAGIHMILIYLCRFLLRRSRNVLFTVSNSLRFRLKVLTVLLYICLSVMLFYLFGSIMPLPPWGLVVGWVMALIAVELAYAFIFPYSFRYIADNDDDKMVILPV |
| IRIS_313-10983 | MLHHLKELAAVAGIHMILIYLCRFLLRRSRNVLFTVSNSLRFRLKVLTVLLYICLSVMLFYLFGSIMPLPPWGLVVGWVMALIAVELAYAFIFPYSFRYIADNDDDKMVILPV |
| IRIS_313-10984 | MLHHLKELAAVAGIHMILIYLCRFLLRRSRNVLFTVSNSLRFRLKVLTVLLYICLSVMLFYLFGSIMPLPPWGLVVGWVMALIAVELAYAFIFPYSFRYIADNDDDKMVILPV |
| IRIS_313-10985 | MLHHLKELAAVAGIHMILIYLCRFLLRRSRNVLFTVSNSLRFRLKVLTVLLYICLSVMLFYLFGSIMPLPPWGLVVGWVMALIAVELAYAFIFPYSFRYIADNDDDKMVILPV |
| IRIS_313-10986 | MLHHLKELAAVAGIHMILIYLCRFLLRRSRNVLFTVSNSLRFRLKVLTVLLYICLSVMLFYLFGSIMPLPPWGLVVGWVMALIAVELAYAFIFPYSFRYIADNDDDKMVILPV |
| IRIS_313-10987 | MLHHLKELAAVAGIHMILIYLCRFLLRRSRNVLFTVSNSLRFRLKVLTVLLYICLSVMLFYLFGSIMPLPPWGLVVGWVMALIAVELAYAFIFPYSFRYIADNDDDKMVILPV |
| IRIS_313-10988 | MLHHLKELAAVAGIHMILIYLCRFLLRRSRNVLFTVSNSLRFRLKVLTVLLYICLSVMLFYLFGSIMPLPPWGLVVGWVMALIAVELAYAFIFPYSFRYIADNDDDKMVILPV |
| IRIS_313-10989 | MLHHLKELAAVAGIHMILIYLCRFLLRRSRNVLFTVSNSLRFRLKVLTVLLYICLSVMLFYLFGSIMPLPPWGLVVGWVMALIAVELAYAFIFPYSFRYIADNDDDKMVILPV |
| IRIS_313-10990 | MLHHLKELAAVAGIHMILIYLCRFLLRRSRNVLFTVSNSLRFRLKVLTVLLYICLSVMLFYLFGSIMPLPPWGLVVGWVMALIAVELAYAFIFPYSFRYIADNDDDKMVILPV |
| IRIS_313-10991 | MLHQLKELAAVAGIHMILIYLCRFLLRRSRNVLFTVSNSLRFRLKVLTVLLYICLLVMLFYLFGSIMPLPPWGLVVGWVMALIAVELAYAFIFPYSFRYIADNDDDKMVILPV |
| IRIS_313-10992 | MLHHLKELAAVAGIHMILIYLCRFLLRRSRNVLFTVSNSLRFRLKVLTVLLYICLSVMLFYLFGSIMPLPPWGLVVGWVMALIAVELAYAFIFPYSFRYIADNDDDKMVILPV |
| IRIS_313-10993 | MLHHLKELAAVAGIHMILIYLCRFLLRRSRNVLFTVSNSLRFRLKVLTVLLYICLSVMLFYLFGSIMPLPPWGLVVGWVMALIAVELAYAFIFPYSFRYIADNDDDKMVILPV |
| IRIS_313-10994 | MLHHLKELAAVAGIHMILIYLCRFLLRRSRNVLFTVSNSLRFRLKVLTVLLYICLSVMLFYLFGSIMPLPPWGLVVGWVMALIAVELAYAFIFPYSFRYIADNDDDKMVILPV |
| IRIS_313-10995 | MLHHLKELAAVAGIHMILIYLCRFLLRRSRNVLFTVSNSLRFRLKVLTVLLYICLSVMLFYLFGSIMPLPPWGLVVGWVMALIAVELAYAFIFPYSFRYIADNDDDKMVILPV |
| IRIS_313-10996 | MLHHLKELAAVAGIHMILIYLCRFLLRRSRNVLFTVSNSLRFRLKVLTVLLYICLSVMLFYLFGSIMPLPPWGLVVGWVMALIAVELAYAFIFPYSFRYIADNDDDKMVILPV |
| IRIS_313-10997 | MLHHLKELAAVAGIHMILIYLCRFLLRRSRNVLFTVSNSLRFRLKVLTVLLYICLSVMLFYLFGSIMPLPPWGLVVGWVMALIAVELAYAFIFPYSFRYIADNDDDKMVILPV |
| IRIS_313-10998 | MLHHLKELAAVAGIHMILIYLCRFLLRRSRNVLFTVSNSLRFRLKVLTVLLYICLSVMLFYLFGSIMPLPPWGLVVGWVMALIAVELAYAFIFPYSFRYIADNDDDKMVILPV |
| IRIS_313-10999 | MLHQLKELAAVAGIHMILIYLCRFLLRRSRNVLFTVSNSLRFRLKVLTVLLYICLLVMLFYLFGSIMPLPPWGLVVGWVMALIAVELAYAFIFPYSFRYIADNDDDKMVILPV |
| IRIS_313-11000 | MLHHLKELAAVAGIHMILIYLCRFLLRRSRNVLFTVSNSLRFRLKVLTVLLYICLSVMLFYLFGSIMPLPPWGLVVGWVMALIAVELAYAFIFPYSFRYIADNDDDKMVILPV |
| IRIS_313-11001 | MLHHLKELAAVAGIHMILIYLCRFLLRRSRNVLFTVSNSLRFRLKVLTVLLYICLSVMLFYLFGSIMPLPPWGLVVGWVMALIAVELAYAFIFPYSFRYIADNDDDKMVILPV |
| IRIS_313-11003 | MLHHLKELAAVAGIHMILIYLCRFLLRRSRNVLFTVSNSLRFRLKVLTVLLYICLSVMLFYLFGSIMPLPPWGLVVGWVMALIAVELAYAFIFPYSFRYIADNDDDKMVILPV |
| IRIS_313-11004 | MLHHLKELAAVAGIHMILIYLCRFFPSRRRNVLFTVSNSLRFRLKVLTVLLYICLSVMLFYLFGSIMPLPPWGLVVGWVMALIAVELAYAFIFPYSFRYIADNDDDKMVILPV |
| IRIS_313-11005 | MLHHLKELAAVAGIHMILIYLCRFLLRRSRNVLFTVSNSLRFRLKVLTVLLYICLSVMLFYLFGSIMPLPPWGLVVGWVMALIAVELAYAFIFPYSFRYIADNDDDKMVILPV |
| IRIS_313-11006 | MLHHLKELAAVAGIHMILIYLCRFLLRRSRNVLFTVSNSLRFRLKVLTVLLYICLSVMLFYLFGSIMPLPPWGLVVGWVMALIAVELAYAFIFPYSFRYIADNDDDKMVILPV |
| IRIS_313-11007 | MLHHLKELAAVAGIHMILIYLCRFLLRRSRNVLFTVSNSLRFRLKVLTVLLYICLSVMLFYLFGSIMPLPPWGLVVGWVMALIAVELAYAFIFPYSFRYIADNDDDKMVILPV |
| IRIS_313-11008 | MLHHLKELAAVAGIHMILIYLCRFLLRRSRNVLFTVSNSLRFRLKVLTVLLYICLSVMLFYLFGSIMPLPPWGLVVGWVMALIAVELAYAFIFPYSFRYIADNDDDKMVILPV |
| IRIS_313-11009 | MLHHLKELAAVAGIHMILIYLCRFLLRRSRNVLFTVSNSLRFRLKVLTVLLYICLSVMLFYLFGSIMPLPPWGLVVGWVMALIAVELAYAFIFPYSFRYIADNDDDKMVILPV |
| IRIS_313-11010 | MLHHLKELAAVAGIHMILIYLCRFLLRRSRNVLFTVSNSLRFRLKVLTVLLYICLSVMLFYLFGSIMPLPPWGLVVGWVMALIAVELAYAFIFPYSFRYIADNDDDKMVILPV |
| IRIS_313-11011 | MLHHLKELAAVAGIHMILIYLCRFLLRRSRNVLFTVSNSLRFRLKVLTVLLYICLSVMLFYLFGSIMPLPPWGLVVGWVMALIAVELAYAFIFPYSFRYIADNDDDKMVILPV |
| IRIS_313-11012 | MLHHLKELAAVAGIHMILIYLCRFLLRRSRNVLFTVSNSLRFRLKVLTVLLYICLSVMLFYLFGSIMPLPPWGLVVGWVMALIAVELAYAFIFPYSFRYIADNDDDKMVILPV |
| IRIS_313-11013 | MLHHLKELAAVAGIHMILIYLCRFLLRRSRNVLFTVSNSLRFRLKVLTVLLYICLSVMLFYLFGSIMPLPPWGLVVGWVMALIAVELAYAFIFPYSFRYIADNDDDKMVILPV |
| IRIS_313-11014 | MLHHLKELAAVAGIHMILIYLCRFLLRRSRNVLFTVSNSLRFRLKVLTVLLYICLSVMLFYLFGSIMPLPPWGLVVGWVMALIAVELAYAFIFPYSFRYIADNDDDKMVILPV |
| IRIS_313-11015 | MLHHLKELAAVAGIHMILIYLCRFLLRRSRNVLFTVSNSLRFRLKVLTVLLYICLSVMLFYLFGSIMPLPPWGLVVGWVMALIAVELAYAFIFPYSFRYIADNDDDKMVILPV |
| IRIS_313-11016 | MLHHLKELAAVAGIHMILIYLCRFLLRRSRNVLFTVSNSLRFRLKVLTVLLYICLSVMLFYLFGSIMPLPPWGLVVGWVMALIAVELAYAFIFPYSFRYIADNDDDKMVILPV |
| IRIS_313-11017 | MLHHLKELAAVAGIHMILIYLCRFLLRRSRNVLFTVSNSLRFRLKVLTVLLYISLSAMLLYLFGSIMPLPPWGLVVGWVMALIAVELAYAFIFPYSFRYIADNDDDKMVILPV |
| IRIS_313-11018 | MLHHLKELAAVAGIHMILIYLCRFLLRRSRNVLFTVSNSLRFRLKVLTVLLYICLSVMLFYLFGSIMPLPPWGLVVGWVMALIAVELAYAFIFPYSFRYIADNDDDKMVILPV |
| IRIS_313-11019 | MLHHLKELAAVAGIHMILIYLCRFLLRRSRNVLFTVSNSLRFRLKVLTVLLYICLSVMLFYLFGSIMPLPPWGLVVGWVMALIAVELAYAFIFPYSFRYIADNDDDKMVILPV |
| IRIS_313-11020 | MLHHLKELAAVAGIHMILIYLCRFLLRRSRNVLFTVSNSLRFRLKVLTVLLYISLSAMLLYLFGSIMPLPPWGLVVGWVMALIAVELAYAFIFPYSFRYIADNDDDKMVILPV |
| IRIS_313-11021 | MLHHLKELAAVAGIHMILIYLCRFLLRRSRNVLFTVSNSLRFRLKVLTVLLYICLSVMLFYLFGSIMPLPPWGLVVGWVMALIAVELAYAFIFPYSFRYIADNDDDKMVILPV |
| IRIS_313-11022 | MLHHLKELAAVAGIHMILIYLCRFLLRRSRNVLFTVSNSLRFRLKVLTVLLYICLSVMLFYLFGSIMPLPPWGLVVGWVMALIAVELAYAFIFPYSFRYIADNDDDKMVILPV |
| IRIS_313-11023 | MLHHLKELAAVAGIHMILIYLCRFLLRRSRNVLFTVSNSLRFRLKVLTVLLYICLSVMLFYLFGSIMPLPPWGLVVGWVMALIAVELAYAFIFPYSFRYIADNDDDKMVILPV |
| IRIS_313-11024 | MLHHLKELAAVAGIHMILIYLCRFLLRRSRNVLFTVSNSLRFRLKVLTVLLYICLSVMLFYLFGSIMPLPPWGLVVGWVMALIAVELAYAFIFPYSFRYIADNDDDKMVILPV |
| IRIS_313-11025 | MLHHLKELAAVAGIHMILIYLCRFLLRRSRNVLFTVSNSLRFRLKVLTVLLYICLSVMLFYLFGSIMPLPPWGLVVGWVMALIAVELAYAFIFPYSFRYIADNDDDKMVILPV |
| IRIS_313-11026 | MLHHLKELAAVAGIHMILIYLCRFLLRRSRNVLFTVSNSLRFRLKVLTVLLYICLSVMLFYLFGSIMPLPPWGLVVGWVMALIAVELAYAFIFPYSFRYIADNDDDKMVILPV |
| IRIS_313-11027 | MLHHLKELAAVAGIHMILIYLCRFLLRRSRNVLFTVSNSLRFRLKVLTVLLYICLSVMLFYLFGSIMPLPPWGLVVGWVMALIAVELAYAFIFPYSFRYIADNDDDKMVILPV |
| IRIS_313-11028 | MLHHLKELAAVAGIHMILIYLCRFLLRRSRNVLFTVSNSLRFRLKVLTVLLYICLSVMLFYLFGSIMPLPPWGLVVGWVMALIAVELAYAFIFPYSFRYIADNDDDKMVILPV |
| IRIS_313-11029 | MLHHLKELAAVAGIHMILIYLCRFLLRRSRNVLFTVSNSLRFRLKVLTVLLYICLSVMLFYLFGSIMPLPPWGLVVGWVMALIAVELAYAFIFPYSFRYIADNDDDKMVILPV |
| IRIS_313-11030 | MLHHLKELAAVAGIHMILIYLCRFLLRRSRNVLFTVSNSLRFRLKVLTVLLYICLSVMLFYLFGSIMPLPPWGLVVGWVMALIAVELAYAFIFPYSFRYIADNDDDKMVILPV |
| IRIS_313-11031 | MLHHLKELAAVAGIHMILIYLCRFLLRRSRNVLFTVSNSLRFRLKVLTVLLYICLSVMLFYLFGSIMPLPPWGLVVGWVMALIAVELAYAFIFPYSFRYIADNDDDKMVILPV |
| IRIS_313-11032 | MLHHLKELAAVAGIHMILIYLCRFLLRRSRNVLFTVSNSLRFRLKVLTVLLYICLSVMLFYLFGSIMPLPPWGLVVGWVMALIAVELAYAFIFPYSFRYIADNDDDKMVILPV |
| IRIS_313-11033 | MLHHLKELAAVAGIHMILIYLCRFLLRRSRNVLFTVSNSLRFRLKVLTVLLYICLSVMLFYLFGSIMPLPPWGLVVGWVMALIAVELAYAFIFPYSFRYIADNDDDKMVILPV |
| IRIS_313-11034 | MLHHLKELAAVAGIHMILIYLCRFLLRRSRNVLFTVSNSLRFRLKVLTVLLYISLSAMLLYLFGSIMPLPPWGLVVGWVMALIAVELAYAFIFPYSFRYIADNDDDKMVILPV |
| IRIS_313-11035 | MLHHLKELAAVAGIHMILIYLCRFLLRRSRNVLFTVSNSLRFRLKVLTVLLYICLSVMLFYLFGSIMPLPPWGLVVGWVMALIAVELAYAFIFPYSFRYIADNDDDKMVILPV |
| IRIS_313-11036 | MLHHLKELAAVAGIHMILIYLCRFLLRRSRNVLFTVSNSLRFRLKVLTVLLYICLSVMLFYLFGSIMPLPPWGLVVGWVMALIAVELAYAFIFPYSFRYIADNDDDKMVILPV |
| IRIS_313-11037 | MLHHLKELAAVAGIHMILIYLCRFLLRRSRNVLFTVSNSLRFRLKVLTVLLYISLSAMLLYLFGSIMPLPPWGLVVGWVMALIAVELAYAFIFPYSFRYIADNDDDKMVILPV |
| IRIS_313-11038 | MLHHLKELAAVAGIHMILIYLCRFLLRRSRNVLFTVSNSLRFRLKVLTVLLYICLSVMLFYLFGSIMPLPPWGLVVGWVMALIAVELAYAFIFPYSFRYIADNDDDKMVILPV |
| IRIS_313-11039 | MLHHLKELAAVAGIHMILIYLCRFLLRLRRNVLFTVSNSLRFRLKVLTVLLYICLSVMLFYLFGSIMPLPPWGLVVGWVMALIAVELAYAFIFPYSFRYIADNDDDKMVILPV |
| IRIS_313-11040 | MLHHLKELAAVAGIHMILIYLCRFLLRRSRNVLFTVSNSLRFRLKVLTVLLYICLSVMLFYLFGSIMPLPPWGLVVGWVMALIAVELAYAFIFPYSFRYIADNDDDKMVILPV |
| IRIS_313-11041 | MLHHLKELAAVAGIHMILIYLCRFLLRRSRNVLFTVSNSLRFRLKVLTVLLYICLSVMLFYLFGSIMPLPPWGLVVGWVMALIAVELAYAFIFPYSFRYIADNDDDKMVILPV |
| IRIS_313-11042 | MLHHLKELAAVAGIHMILIYLCRFLLRRSRNVLFTVSNSLRFRLKVLTVLLYICLSVMLFYLFGSIMPLPPWGLVVGWVMALIAVELAYAFIFPYSFRYIADNDDDKMVILPV |
| IRIS_313-11043 | MLHHLKELAAVAGIHMILIYLCRFLLRRSRNVLFTVSNSLRFRLKVLTVLLYICLSVMLFYLFGSIMPLPPWGLVVGWVMALIAVELAYAFIFPYSFRYIADNDDDKMVILPV |
| IRIS_313-11044 | MLHHLKELAAVAGIHMILIYLCRFLLRRSRNVLFTVSNSLRFRLKVLTVLLYICLSVMLFYLFGSIMPLPPWGLVVGWVMALIAVELAYAFIFPYSFRYIADNDDDKMVILPV |
| IRIS_313-11045 | MLHHLKELAAVAGIHMILIYLCRFLLRRSRNVLFTVSNSLRFRLKVLTVLLYICLSVMLFYLFGSIMPLPPWGLVVGWVMALIAVELAYAFIFPYSFRYIADNDDDKMVILPV |
| IRIS_313-11046 | MLHHLKELAAVAGIHMILIYLCRFLLRRSRNVLFTVSNSLRFRLKVLTVLLYICLSVMLFYLFGSIMPLPPWGLVVGWVMALIAVELAYAFIFPYSFRYIADNDDDKMVILPV |
| IRIS_313-11047 | MLHHLKELAAVAGIHMILIYLCRFLLRRSRNVLFTVSNSLRFRLKVLTVLLYICLSVMLFYLFGSIMPLPPWGLVVGWVMALIAVELAYAFIFPYSFRYIADNDDDKMVILPV |
| IRIS_313-11048 | MLHHLKELAAVAGIHMILIYLCRFLLRRSRNVLFTVSNSLRFRLKVLTVLLYICLSVMLFYLFGSIMPLPPWGLVVGWVMALIAVELAYAFIFPYSFRYIADNDDDKMVILPV |
| IRIS_313-11049 | MLHHLKELAAVAGIHMILIYLCRFLLRRSRNVLFTVSNSLRFRLKVLTVLLYICLSVMLFYLFGSIMPLPPWGLVVGWVMALIAVELAYAFIFPYSFRYIADNDDDKMVILPV |
| IRIS_313-11050 | MLHHLKELAAVAGIHMILIYLCRFLLRRSRNVLFTVSNSLRFRLKVLTVLLYICLSVMLFYLFGSIMPLPPWGLVVGWVMALIAVELAYAFIFPYSFRYIADNDDDKMVILPV |
| IRIS_313-11051 | MLHHLKELAAVAGIHMILIYLCRFLLRRSRNVLFTVSNSLRFRLKVLTVLLYISLSAMLLYLFGSIMPLPPWGLVVGWVMALIAVELAYAFIFPYSFRYIADNDDDKMVILPV |
| IRIS_313-11052 | MLHHLKELAAVAGIHMILIYLCRFLLRRSRNVLFTVSNSLRFRLKVLTVLLYICLSVMLFYLFGSIMPLPPWGLVVGWVMALIAVELAYAFIFPYSFRYIADNDDDKMVILPV |
| IRIS_313-11053 | MLHHLKELAAVAGIHMILIYLCRFLLRRSRNVLFTVSNSLRFRLKVLTVLLYICLSVMLFYLFGSIMPLPPWGLVVGWVMALIAVELAYAFIFPYSFRYIADNDDDKMVILPV |
| IRIS_313-11054 | MLHHLKELAAVAGIHMILIYLCRFLLRRSRNVLFTVSNSLRFRLKVLTVLLYICLSVMLFYLFGSIMPLPPWGLVVGWVMALIAVELAYAFIFPYSFRYIADNDDDKMVILPV |
| IRIS_313-11055 | MLHHLKELAAVAGIHMILIYLCRFLLRRSRNVLFTVSNSLRFRLKVLTVLLYICLSVMLFYLFGSIMPLPPWGLVVGWVMALIAVELAYAFIFPYSFRYIADNDDDKMVILPV |
| IRIS_313-11056 | MLHHLKELAAVAGIHMILIYLCRFLLRRSRNVLFTVSNSLRFRLKVLTVLLYICLSVMLFYLFGSIMPLPPWGLVVGWVMALIAVELAYAFIFPYSFRYIADNDDDKMVILPV |
| IRIS_313-11057 | MLHHLKELAAVAGIHMILIYLCRFLLRRSRNVLFTVSNSLRFRLKVLTVLLYICLSVMLFYLFGSIMPLPPWGLVVGWVMALIAVELAYAFIFPYSFRYIADNDDDKMVILPV |
| IRIS_313-11058 | MLHHLKELAAVAGIHMILIYLCRFLLRRSRNVLFTVSNSLRFRLKVLTVLLYICLSVMLFYLFGSIMPLPPWGLVVGWVMALIAVELAYAFIFPYSFRYIADNDDDKMVILPV |
| IRIS_313-11059 | MLHHLKELAAVAGIHMILIYLCRFLLRRSRNVLFTVSNSLRFRLKVLTVLLYICLSVMLFYLFGSIMPLPPWGLVVGWVMALIAVELAYAFIFPYSFRYIADNDDDKMVILPV |
| IRIS_313-11060 | MLHHLKELAAVAGIHMILIYLCRFLLRRSRNVLFTVSNSLRFRLKVLTVLLYICLSVMLFYLFGSIMPLPPWGLVVGWVMALIAVELAYAFIFPYSFRYIADNDDDKMVILPV |
| IRIS_313-11061 | MLHHLKELAAVAGIHMILIYLCRFLLRRSRNVLFTVSNSLRFRLKVLTVLLYICLSVMLFYLFGSIMPLPPWGLVVGWVMALIAVELAYAFIFPYSFRYIADNDDDKMVILPV |
| IRIS_313-11062 | MLHHLKELAAVAGIHMILIYLCRFLLRRSRNVLFTVSNSLRFRLKVLTVLLYISLSAMLLYLFGSIMPLPPWGLVVGWVMALIAVELAYAFIFPYSFRYIADNDDDKMVILPV |
| IRIS_313-11063 | MLHHLKELAAVAGIHMILIYLCRFLLRRSRNVLFTVSNSLRFRLKVLTVLLYISLSAMLLYLFGSIMPLPPWGLVVGWVMALIAVELAYAFIFPYSFRYIADNDDDKMVILPV |
| IRIS_313-11064 | MLHHLKELAAVAGIHMILIYLCRFLLRRSRNVLFTVSNSLRFRLKVLTVLLYISLSAMLLYLFGSIMPLPPWGLVVGWVMALIAVELAYAFIFPYSFRYIADNDDDKMVILPV |
| IRIS_313-11065 | MLHHLKELAAVAGIHMILIYLCRFLLRRSRNVLFTVSNSLRFRLKVLTVLLYICLSVMLFYLFGSIMPLPPWGLVVGWVMALIAVELAYAFIFPYSFRYIADNDDDKMVILPV |
| IRIS_313-11066 | MLHHLKELAAVAGIHMILIYLCRFLLRRSRNVLFTVSNSLRFRLKVLTVLLYISLSAMLLYLFGSIMPLPPWGLVVGWVMALIAVELAYAFIFPYSFRYIADNDDDKMVILPV |
| IRIS_313-11067 | MLHHLKELAAVAGIHMILIYLCRFLLRRSRNVLFTVSNSLRFRLKVLTVLLYICLSVMLFYLFGSIMPLPPWGLVVGWVMALIAVELAYAFIFPYSFRYIADNDDDKMVILPV |
| IRIS_313-11068 | MLHHLKELAAVAGIHMILIYLCRFLLRRSRNVLFTVSNSLRFRLKVLTVLLYICLSVMLFYLFGSIMPLPPWGLVVGWVMALIAVELAYAFIFPYSFRYIADNDDDKMVILPV |
| IRIS_313-11069 | MLHHLKELAAVAGIHMILIYLCRFLLRRSRNVLFTVSNSLRFRLKVLTVLLYICLSVMLFYLFGSIMPLPPWGLVVGWVMALIAVELAYAFIFPYSFRYIADNDDDKMVILPV |
| IRIS_313-11070 | MLHHLKELAAVAGIHMILIYLCRFLLRRSRNVLFTVSNSLRFRLKVLTVLLYICLSVMLFYLFGSIMPLPPWGLVVGWVMALIAVELAYAFIFPYSFRYIADNDDDKMVILPV |
| IRIS_313-11071 | MLHHLKELAAVAGIHMILIYLCRFLLRRSRNVLFTVSNSLRFRLKVLTVLLYICLSVMLFYLFGSIMPLPPWGLVVGWVMALIAVELAYAFIFPYSFRYIADNDDDKMVILPV |
| IRIS_313-11072 | MLHHLKELAAVAGIHMILIYLCRFLLRRSRNVLFTVSNSLRFRLKVLTVLLYICLSVMLFYLFGSIMPLPPWGLVVGWVMALIAVELAYAFIFPYSFRYIADNDDDKMVILPV |
| IRIS_313-11073 | MLHHLKELAAVAGIHMILIYLCRFLLRRSRNVLFTVSNSLRFRLKVLTVLLYICLSVMLFYLFGSIMPLPPWGLVVGWVMALIAVELAYAFIFPYSFRYIADNDDDKMVILPV |
| IRIS_313-11074 | MLHHLKELAAVAGIHMILIYLCRFLLRRSRNVLFTVSNSLRFRLKVLTVLLYICLSVMLFYLFGSIMPLPPWGLVVGWVMALIAVELAYAFIFPYSFRYIADNDDDKMVILPV |
| IRIS_313-11075 | MLHHLKELAAVAGIHMILIYLCRFLLRRSRNVLFTVSNSLRFRLKVLTVLLYICLSVMLFYLFGSIMPLPPWGLVVGWVMALIAVELAYAFIFPYSFRYIADNDDDKMVILPV |
| IRIS_313-11076 | MLHHLKELAAVAGIHMILIYLCRFLLRRSRNVLFTVSNSLRFRLKVLTVLLYICLSVMLFYLFGSIMPLPPWGLVVGWVMALIAVELAYAFIFPYSFRYIADNDDDKMVILPV |
| IRIS_313-11077 | MLHQLKELAAVAGIHMILIYLCRFLLRRSRNVLFTVSNSLRFRLKVLTVLLYICLLVMLFYLFGSIMPLPPWGLVVGWVMALIAVELAYAFIFPYSFRYIADNDDDKMVILPV |
| IRIS_313-11078 | MLHHLKELAAVAGIHMILIYLCRFLLRRSRNVLFTVSNSLRFRLKVLTVLLYICLSVMLFYLFGSIMPLPPWGLVVGWVMALIAVELAYAFIFPYSFRYIADNDDDKMVILPV |
| IRIS_313-11079 | MLHHLKELAAVAGIHMILIYLCRFLLRRSRNVLFTVSNSLRFRLKVLTVLLYICLSVMLFYLFGSIMPLPPWGLVVGWVMALIAVELAYAFIFPYSFRYIADNDDDKMVILPV |
| IRIS_313-11080 | MLHHLKELAAVAGIHMILIYLCRFLLRRSRNVLFTVSNSLRFRLKVLTVLLYICLSVMLFYLFGSIMPLPPWGLVVGWVMALIAVELAYAFIFPYSFRYIADNDDDKMVILPV |
| IRIS_313-11081 | MLHHLKELAAVAGIHMILIYLCRFLLRRSRNVLFTVSNSLRFRLKVLTVLLYICLSVMLFYLFGSIMPLPPWGLVVGWVMALIAVELAYAFIFPYSFRYIADNDDDKMVILPV |

| | |
|---|---|
| IRIS_313-11082 | MLHHLKELAAVAGIHMILIYLCRFLLRRSRNVLFTVSNSLRFRLKVLTVLLYICLSVMLFYLFGSIMPLPPWGLVVGWVMALIAVELAYAFIFPYSFRYIADNDDDKMVILPV |
| IRIS_313-11083 | MLHHLKELAAVAGIHMILIYLCRFLLRRSRNVLFTVSNSLRFRLKVLTVLLYICLSVMLFYLFGSIMPLPPWGLVVGWVMALIAVELAYAFIFPYSFRYIADNDDDKMVILPV |
| IRIS_313-11084 | MLHHLKELAAVAGIHMILIYLCRFLLRRSRNVLFTVSNSLRFRLKVLTVLLYICLSVMLFYLFGSIMPLPPWGLVVGWVMALIAVELAYAFIFPYSFRYIADNDDDKMVILPV |
| IRIS_313-11085 | MLHHLKELAAVAGIHMILIYLCRFLLRRSRNVLFTVSNSLRFRLKVLTVLLYICLSVMLFYLFGSIMPLPPWGLVVGWVMALIAVELAYAFIFPYSFRYIADNDDDKMVILPV |
| IRIS_313-11086 | MLHHLKELAAVAGIHMILIYLCRFLLRRSRNVLFTVSNSLRFRLKVLTVLLYICLSVMLFYLFGSIMPLPPWGLVVGWVMALIAVELAYAFIFPYSFRYIADNDDDKMVILPV |
| IRIS_313-11087 | MLHHLKELAAVAGIHMILIYLCRFLLRRSRNVLFTVSNSLRFRLKVLTVLLYICLSVMLFYLFGSIMPLPPWGLVVGWVMALIAVELAYAFIFPYSFRYIADNDDDKMVILPV |
| IRIS_313-11088 | MLHHLKELAAVAGIHMILIYLCRFLLRRSRNVLFTVSNSLRFRLKVLTVLLYICLSVMLFYLFGSIMPLPPWGLVVGWVMALIAVELAYAFIFPYSFRYIADNDDDKMVILPV |
| IRIS_313-11089 | MLHHLKELAAVAGIHMILIYLCRFLLRRSRNVLFTVSNSLRFRLKVLTVLLYICLSVMLFYLFGSIMPLPPWGLVVGWVMALIAVELAYAFIFPYSFRYIADNDDDKMVILPV |
| IRIS_313-11090 | MLHHLKELAAVAGIHMILIYLCRFLLRRSRNVLFTVSNSLRFRLKVLTVLLYICLSVMLFYLFGSIMPLPPWGLVVGWVMALIAVELAYAFIFPYSFRYIADNDDDKMVILPV |
| IRIS_313-11091 | MLHHLKELAAVAGIHMILIYLCRFLLRRSRNVLFTVSNSLRFRLKVLTVLLYICLSVMLFYLFGSIMPLPPWGLVVGWVMALIAVELAYAFIFPYSFRYIADNDDDKMVILPV |
| IRIS_313-11092 | MLHHLKELAAVAGIHMILIYLCRFLLRRSRNVLFTVSNSLRFRLKVLTVLLYICLSVMLFYLFGSIMPLPPWGLVVGWVMALIAVELAYAFIFPYSFRYIADNDDDKMVILPV |
| IRIS_313-11093 | MLHHLKELAAVAGIHMILIYLCRFLLRRSRNVLFTVSNSLRFRLKVLTVLLYICLSVMLFYLFGSIMPLPPWGLVVGWVMALIAVELAYAFIFPYSFRYIADNDDDKMVILPV |
| IRIS_313-11094 | MLHHLKELAAVAGIHMILIYLCRFLLRRSRNVLFTVSNSLRFRLKVLTVLLYICLSVMLFYLFGSIMPLPPWGLVVGWVMALIAVELAYAFIFPYSFRYIADNDDDKMVILPV |
| IRIS_313-11095 | MLHHLKELAAVAGIHMILIYLCRFLLRRSRNVLFTVSNSLRFRLKVLTVLLYICLSVMLFYLFGSIMPLPPWGLVVGWVMALIAVELAYAFIFPYSFRYIADNDDDKMVILPV |
| IRIS_313-11096 | MLHHLKELAAVAGIHMILIYLCRFLLRRSRNVLFTVSNSLRFRLKVLTVLLYICLSVMLFYLFGSIMPLPPWGLVVGWVMALIAVELAYAFIFPYSFRYIADNDDDKMVILPV |
| IRIS_313-11097 | MLHHLKELAAVAGIHMILIYLCRFLLRRSRNVLFTVSNSLRFRLKVLTVLLYICLSVMLFYLFGSIMPLPPWGLVVGWVMALIAVELAYAFIFPYSFRYIADNDDDKMVILPV |
| IRIS_313-11098 | MLHHLKELAAVAGIHMILIYLCRFLLRRSRNVLFTVSNSLRFRLKVLTVLLYICLSVMLFYLFGSIMPLPPWGLVVGWVMALIAVELAYAFIFPYSFRYIADNDDDKMVILPV |
| IRIS_313-11099 | MLHHLKELAAVAGIHMILIYLCRFLLRRSRNVLFTVSNSLRFRLKVLTVLLYICLSVMLFYLFGSIMPLPPWGLVVGWVMALIAVELAYAFIFPYSFRYIADNDDDKMVILPV |
| IRIS_313-11100 | MLHHLKELAAVAGIHMILIYLCRFLLRRSRNVLFTVSNSLRFRLKVLTVLLYICLSVMLFYLFGSIMPLPPWGLVVGWVMALIAVELAYAFIFPYSFRYIADNDDDKMVILPV |
| IRIS_313-11101 | MLHHLKELAAVAGIHMILIYLCRFLLRRSRNVLFTVSNSLRFRLKVLTVLLYICLSVMLFYLFGSIMPLPPWGLVVGWVMALIAVELAYAFIFPYSFRYIADNDDDKMVILPV |
| IRIS_313-11102 | MLHHLKELAAVAGIHMILIYLCRFLLRRSRNVLFTVSNSLRFRLKVLTVLLYICLSVMLFYLFGSIMPLPPWGLVVGWVMALIAVELAYAFIFPYSFRYIADNDDDKMVILPV |
| IRIS_313-11103 | MLHHLKELAAVAGIHMILIYLCRFLLRRSRNVLFTVSNSLRFRLKVLTVLLYICLSVMLFYLFGSIMPLPPWGLVVGWVMALIAVELAYAFIFPYSFRYIADNDDDKMVILPV |
| IRIS_313-11104 | MLHHLKELAAVAGIHMILIYLCRFLLRRSRNVLFTVSNSLRFRLKVLTVLLYICLSVMLFYLFGSIMPLPPWGLVVGWVMALIAVELAYAFIFPYSFRYIADNDDDKMVILPV |
| IRIS_313-11105 | MLHHLKELAAVAGIHMILIYLCRFLLRRSRNVLFTVSNSLRFRLKVLTVLLYICLSVMLFYLFGSIMPLPPWGLVVGWVMALIAVELAYAFIFPYSFRYIADNDDDKMVILPV |
| IRIS_313-11106 | MLHHLKELAAVAGIHMILIYLCRFLLRRSRNVLFTVSNSLRFRLKVLTVLLYICLSVMLFYLFGSIMPLPPWGLVVGWVMALIAVELAYAFIFPYSFRYIADNDDDKMVILPV |
| IRIS_313-11107 | MLHHLKELAAVAGIHMILIYLCRFFLRRSRNVLFTVSNSLRFRLKVLTVLLYICLSVMLFYLFGSIMPLPPWGLVVGWVMALIAVELAYAFIFPYSFRYIADNDDDKMVILPV |
| IRIS_313-11108 | MLHHLKELAAVAGIHMILIYLCRFLLRRSRNVLFTVSNSLRFRLKVLTVLLYICLSVMLFYLFGSIMPLPPWGLVVGWVMALIAVELAYAFIFPYSFRYIADNDDDKMVILPV |
| IRIS_313-11109 | MLHHLKELAAVAGIHMILIYLCRFLLRRSRNVLFTVSNSLRFRLKVLTVLLYICLSVMLFYLFGSIMPLPPWGLVVGWVMALIAVELAYAFIFPYSFRYIADNDDDKMVILPV |
| IRIS_313-11110 | MLHHLKELAAVAGIHMILIYLCRFLLRRSRNVLFTVSNSLRFRLKVLTVLLYICLSVMLFYLFGSIMPLPPWGLVVGWVMALIAVELAYAFIFPYSFRYIADNDDDKMVILPV |
| IRIS_313-11111 | MLHHLKELAAVAGIHMILIYLCRFLLRRSRNVLFTVSNSLRFRLKVLTVLLYICLSVMLFYLFGSIMPLPPWGLVVGWVMALIAVELAYAFIFPYSFRYIADNDDDKMVILPV |
| IRIS_313-11112 | MLHHLKELAAVAGIHMILIYLCRFLLRRSRNVLFTVSNSLRFRLKVLTVLLYICLSVMLFYLFGSIMPLPPWGLVVGWVMALIAVELAYAFIFPYSFRYIADNDDDKMVILPV |
| IRIS_313-11113 | MLHHLKELAAVAGIHMILIYLCRFLLRRSRNVLFTVSNSLRFRLKVLTVLLYICLSVMLFYLFGSIMPLPPWGLVVGWVMALIAVELAYAFIFPYSFRYIADNDDDKMVILPV |
| IRIS_313-11114 | MLHHLKELAAVAGIHMILIYLCRFLLRRSRNVLFTVSNSLRFRLKVLTVLLYICLSVMLFYLFGSIMPLPPWGLVVGWVMALIAVELAYAFIFPYSFRYIADNDDDKMVILPV |
| IRIS_313-11115 | MLHHLKELAAVAGIHMILIYLCRFLLRRSRNVLFTVSNSLRFRLKVLTVLLYICLSVMLFYLFGSIMPLPPWGLVVGWVMALIAVELAYAFIFPYSFRYIADNDDDKMVILPV |
| IRIS_313-11116 | MLHHLKELAAVAGIHMILIYLCRFLLRRSRNVLFTVSNSLRFRLKVLTVLLYICLSVMLFYLFGSIMPLPPWGLVVGWVMALIAVELAYAFIFPYSFRYIADNDDDKMVILPV |
| IRIS_313-11117 | MLHQLKELAAVAGIHMILIYLCRFLLRRSRNVLFTVSNSLRFRLKVLTVLLYICLSVMLFYLFGSIMPLPPWGLVVGWVMALIAVELAYAFIFPYSFRYIADNDDDKMVILPV |
| IRIS_313-11118 | MLHHLKELAAVAGIHMILIYLCRFLLRRSRNVLFTVSNSLRFRLKVLTVLLYICLSVMLFYLFGSIMPLPPWGLVVGWVMALIAVELAYAFIFPYSFRYIADNDDDKMVILPV |
| IRIS_313-11119 | MLHHLKELAAVAGIHMILIYLCRFLLRRSRNVLFTVSNSLRFRLKVLTVLLYICLSVMLFYLFGSIMPLPPWGLVVGWVMALIAVELAYAFIFPYSFRYIADNDDDKMVILPV |
| IRIS_313-11120 | MLHHLKELAAVAGIHMILIYLCRFLLRRSRNVLFTVSNSLRFRLKVLTVLLYICLSVMLFYLFGSIMPLPPWGLVVGWVMALIAVELAYAFIFPYSFRYIADNDDDKMVILPV |
| IRIS_313-11121 | MLHHLKELAAVAGIHMILIYLCRFLLRRSRNVLFTVSNSLRFRLKVLTVLLYICLSVMLFYLFGSIMPLPPWGLVVGWVMALIAVELAYAFIFPYSFRYIADNDDDKMVILPV |
| IRIS_313-11122 | MLHHLKELAAVAGIHMILIYLCRFLLRRSRNVLFTVSNSLRFRLKVLTVLLYICLSVMLFYLFGSIMPLPPWGLVVGWVMALIAVELAYAFIFPYSFRYIADNDDDKMVILPV |
| IRIS_313-11123 | MLHHLKELAAVAGIHMILIYLCRFLLRRSRNVLFTVSNSLRFRLKVLTVLLYICLSVMLFYLFGSIMPLPPWGLVVGWVMALIAVELAYAFIFPYSFRYIADNDDDKMVILPV |
| IRIS_313-11124 | MLHHLKELAAVAGIHMILIYLCRFLLRRSRNVLFTVSNSLRFRLKVLTVLLYICLSVMLFYLFGSIMPLPPWGLVVGWVMALIAVELAYAFIFPYSFRYIADNDDDKMVILPV |
| IRIS_313-11125 | MLHHLKELAAVAGIHMILIYLCRFLLRRSRNVLFTVSNSLRFRLKVLTVLLYICLSVMLFYLFGSIMPLPPWGLVVGWVMALIAVELAYAFIFPYSFRYIADNDDDKMVILPV |
| IRIS_313-11126 | MLHHLKELAAVAGIHMILIYLCRFLLRRSRNVLFTVSNSLRFRLKVLTVLLYICLSVMLFYLFGSIMPLPPWGLVVGWVMALIAVELAYAFIFPYSFRYIADNDDDKMVILPV |
| IRIS_313-11127 | MLHHLKELAAVAGIHMILIYLCRFLLRRSRNVLFTVSNSLRFRLKVLTVLLYICLSVMLFYLFGSIMPLPPWGLVVGWVMALIAVELAYAFIFPYSFRYIADNDDDKMVILPV |
| IRIS_313-11128 | MLHHLKELAAVAGIHMILIYLCRFLLRRSRNVLFTVSNSLRFRLKVLTVLLYICLSVMLFYLFGSIMPLPPWGLVVGWVMALIAVELAYAFIFPYSFRYIADNDDDKMVILPV |
| IRIS_313-11129 | MLHHLKELAAVAGIHMILIYLCRFLLRRSRNVLFTVSNSLRFRLKVLTVLLYICLSVMLFYLFGSIMPLPPWGLVVGWVMALIAVELAYAFIFPYSFRYIADNDDDKMVILPV |
| IRIS_313-11130 | MLHHLKELAAVAGIHMILIYLCRFLLRRSRNVLFTVSNSLRFRLKVLTVLLYICLSVMLFYLFGSIMPLPPWGLVVGWVMALIAVELAYAFIFPYSFRYIADNDDDKMVILPV |
| IRIS_313-11131 | MLHHLKELAAVAGIHMILIYLCRFLLRRSRNVLFTVSNSLRFRLKVLTVLLYICLSVMLFYLFGSIMPLPPWGLVVGWVMALIAVELAYAFIFPYSFRYIADNDDDKMVILPV |
| IRIS_313-11132 | MLHHLKELAAVAGIHMILIYLCRFLLRRSRNVLFTVSNSLRFRLKVLTVLLYICLSVMLFYLFGSIMPLPPWGLVVGWVMALIAVELAYAFIFPYSFRYIADNDDDKMVILPV |
| IRIS_313-11133 | MLHHLKELAAVAGIHMILIYLCRFLLRRSRNVLFTVSNSLRFRLKVLTVLLYICLSVMLFYLFGSIMPLPPWGLVVGWVMALIAVELAYAFIFPYSFRYIADNDDDKMVILPV |
| IRIS_313-11134 | MLHHLKELAAVAGIHMILIYLCRFLLRRSRNVLFTVSNSLRFRLKVLTVLLYICLSVMLFYLFGSIMPLPPWGLVVGWVMALIAVELAYAFIFPYSFRYIADNDDDKMVILPV |
| IRIS_313-11135 | MLHHLKELAAVAGIHMILIYLCRFLLRRSRNVLFTVSNSLRFRLKVLTVLLYICLSVMLFYLFGSIMPLPPWGLVVGWVMALIAVELAYAFIFPYSFRYIADNDDDKMVILPV |
| IRIS_313-11136 | MLHHLKELAAVAGIHMILIYLCRFLLRRSRNVLFTVSNSLRFRLKVLTVLLYICLSVMLFYLFGSIMPLPPWGLVVGWVMALIAVELAYAFIFPYSFRYIADNDDDKMVILPV |
| IRIS_313-11137 | MLHHLKELAAVAGIHMILIYLCRFLLRRSRNVLFTVSNSLRFRLKVLTVLLYICLSVMLFYLFGSIMPLPPWGLVVGWVMALIAVELAYAFIFPYSFRYIADNDDDKMVILPV |
| IRIS_313-11138 | MLHHLKELAAVAGIHMILIYLCRFLLRRSRNVLFTVSNSLRFRLKVLTVLLYICLSVMLFYLFGSIMPLPPWGLVVGWVMALIAVELAYAFIFPYSFRYIADNDDDKMVILPV |
| IRIS_313-11139 | MLHHLKELAAVAGIHMILIYLCRFLLRRSRNVLFTVSNSLRFRLKVLTVLLYICLSVMLFYLFGSIMPLPPWGLVVGWVMALIAVELAYAFIFPYSFRYIADNDDDKMVILPV |
| IRIS_313-11140 | MLHHLKELAAVAGIHMILIYLCRFLLRRSRNVLFTVSNSLRFRLKVLTVLLYICLSVMLFYLFGSIMPLPPWGLVVGWVMALIAVELAYAFIFPYSFRYIADNDDDKMVILPV |
| IRIS_313-11141 | MLHHLKELAAVAGIHMILIYLCRFLLRRSRNVLFTVSNSLRFRLKVLTVLLYICLSVMLFYLFGSIMPLPPWGLVVGWVMALIAVELAYAFIFPYSFRYIADNDDDKMVILPV |
| IRIS_313-11142 | MLHHLKELAAVAGIHMILIYLCRFLLRRSRNVLFTVSNSLRFRLKVLTVLLYICLSVMLFYLFGSIMPLPPWGLVVGWVMALIAVELAYAFIFPYSFRYIADNDDDKMVILPV |
| IRIS_313-11143 | MLHHLKELAAVAGIHMILIYLCRFLLRRSRNVLFTVSNSLRFRLKVLTVLLYICLSVMLFYLFGSIMPLPPWGLVVGWVMALIAVELAYAFIFPYSFRYIADNDDDKMVILPV |
| IRIS_313-11144 | MLHHLKELAAVAGIHMILIYLCRFLLRRSRNVLFTVSNSLRFRLKVLTVLLYICLSVMLFYLFGSIMPLPPWGLVVGWVMALIAVELAYAFIFPYSFRYIADNDDDKMVILPV |
| IRIS_313-11145 | MLHHLKELAAVAGIHMILIYLCRFLLRRSRNVLFTVSNSLRFRLKVLTVLLYICLSVMLFYLFGSIMPLPPWGLVVGWVMALIAVELAYAFIFPYSFRYIADNDDDKMVILPV |
| IRIS_313-11146 | MLHHLKELAAVAGIHMILIYLCRFLLRRSRNVLFTVSNSLRFRLKVLTVLLYICLSVMLFYLFGSIMPLPPWGLVVGWVMALIAVELAYAFIFPYSFRYIADNDDDKMVILPV |
| IRIS_313-11147 | MLHHLKELAAVAGIHMILIYLCRFLLRRSRNVLFTVSNSLRFRLKVLTVLLYICLSVMLFYLFGSIMPLPPWGLVVGWVMALIAVELAYAFIFPYSFRYIADNDDDKMVILPV |
| IRIS_313-11148 | MLHHLKELAAVAGIHMILIYLCRFLLRRSRNVLFTVSNSLRFRLKVLTVLLYICLSVMLFYLFGSIMPLPPWGLVVGWVMALIAVELAYAFIFPYSFRYIADNDDDKMVILPV |
| IRIS_313-11149 | MLHHLKELAAVAGIHMILIYLCRFLLRRSRNVLFTVSNSLRFRLKVLTVLLYICLSVMLFYLFGSIMPLPPWGLVVGWVMALIAVELAYAFIFPYSFRYIADNDDDKMVILPV |
| IRIS_313-11150 | MLHHLKELAAVAGIHMILIYLCRFLLRRSRNVLFTVSNSLRFRLKVLTVLLYICLSVMLFYLFGSIMPLPPWGLVVGWVMALIAVELAYAFIFPYSFRYIADNDDDKMVILPV |
| IRIS_313-11151 | MLHHLKELAAVAGIHMILIYLCRFLLRRSRNVLFTVSNSLRFRLKVLTVLLYICLSVMLFYLFGSIMPLPPWGLVVGWVMALIAVELAYAFIFPYSFRYIADNDDDKMVILPV |
| IRIS_313-11152 | MLHHLKELAAVAGIHMILIYLCRFLLRRSRNVLFTVSNSLRFRLKVLTVLLYICLSVMLFYLFGSIMPLPPWGLVVGWVMALIAVELAYAFIFPYSFRYIADNDDDKMVILPV |
| IRIS_313-11153 | MLHHLKELAAVAGIHMILIYLCRFLLRRSRNVLFTVSNSLRFRLKVLTVLLYICLSVMLFYLFGSIMPLPPWGLVVGWVMALIAVELAYAFIFPYSFRYIADNDDDKMVILPV |
| IRIS_313-11154 | MLHHLKELAAVAGIHMILIYLCRFLLRRSRNVLFTVSNSLRFRLKVLTVLLYICLSVMLFYLFGSIMPLPPWGLVVGWVMALIAVELAYAFIFPYSFRYIADNDDDKMVILPV |
| IRIS_313-11155 | MLHHLKELAAVAGIHMILIYLCRFLLRRSRNVLFTVSNSLRFRLKVLTVLLYICLSVMLFYLFGSIMPLPPWGLVVGWVMALIAVELAYAFIFPYSFRYIADNDDDKMVILPV |
| IRIS_313-11156 | MLHHLKELAAVAGIHMILIYLCRFLLRRSRNVLFTVSNSLRFRLKVLTVLLYICLSVMLFYLFGSIMPLPPWGLVVGWVMALIAVELAYAFIFPYSFRYIADNDDDKMVILPV |
| IRIS_313-11157 | MLHHLKELAAVAGIHMILIYLCRFLLRRSRNVLFTVSNSLRFRLKVLTVLLYICLSVMLFYLFGSIMPLPPWGLVVGWVMALIAVELAYAFIFPYSFRYIADNDDDKMVILPV |
| IRIS_313-11158 | MLHHLKELAAVAGIHMILIYLCRFLLRRSRNVLFTVSNSLRFRLKVLTVLLYICLSVMLFYLFGSIMPLPPWGLVVGWVMALIAVELAYAFIFPYSFRYIADNDDDKMVILPV |
| IRIS_313-11159 | MLHHLKELAAVAGIHMILIYLCRFLLRRSRNVLFTVSNSLRFRLKVLTVLLYICLSVMLFYLFGSIMPLPPWGLVVGWVMALIAVELAYAFIFPYSFRYIADNDDDKMVILPV |
| IRIS_313-11160 | MLHHLKELAAVAGIHMILIYLCRFLLRRSRNVLFTVSNSLRFRLKVLTVLLYICLSVMLFYLFGSIMPLPPWGLVVGWVMALIAVELAYAFIFPYSFRYIADNDDDKMVILPV |
| IRIS_313-11161 | MLHHLKELAAVAGIHMILIYLCRFLLRRSRNVLFTVSNSLRFRLKVLTVLLYICLSVMLFYLFGSIMPLPPWGLVVGWVMALIAVELAYAFIFPYSFRYIADNDDDKMVILPV |
| IRIS_313-11162 | MLHHLKELAAVAGIHMILIYLCRFLLRRSRNVLFTVSNSLRFRLKVLTVLLYICLSVMLFYLFGSIMPLPPWGLVVGWVMALIAVELAYAFIFPYSFRYIADNDDDKMVILPV |
| IRIS_313-11163 | MLHHLKELAAVAGIHMILIYLCRFLLRRSRNVLFTVSNSLRFRLKVLTVLLYICLSVMLFYLFGSIMPLPPWGLVVGWVMALIAVELAYAFIFPYSFRYIADNDDDKMVILPV |
| IRIS_313-11164 | MLHHLKELAAVAGIHMILIYLCRFLLRRSRNVLFTVSNSLRFRLKVLTVLLYICLSVMLFYLFGSIMPLPPWGLVVGWVMALIAVELAYAFIFPYSFRYIADNDDDKMVILPV |
| IRIS_313-11165 | MLHHLKELAAVAGIHMILIYLCRFLLRRSRNVLFTVSNSLRFRLKVLTVLLYICLSVMLFYLFGSIMPLPPWGLVVGWVMALIAVELAYAFIFPYSFRYIADNDDDKMVILPV |
| IRIS_313-11166 | MLHHLKELAAVAGIHMILIYLCRFLLRRSRNVLFTVSNSLRFRLKVLTVLLYICLSVMLFYLFGSIMPLPPWGLVVGWVMALIAVELAYAFIFPYSFRYIADNDDDKMVILPV |
| IRIS_313-11167 | MLHHLKELAAVAGIHMILIYLCRFLLRRSRNVLFTVSNSLRFRLKVLTVLLYICLSVMLFYLFGSIMPLPPWGLVVGWVMALIAVELAYAFIFPYSFRYIADNDDDKMVILPV |
| IRIS_313-11168 | MLHHLKELAAVAGIHMILIYLCRFLLRRSRNVLFTVSNSLRFRLKVLTVLLYICLSVMLFYLFGSIMPLPPWGLVVGWVMALIAVELAYAFIFPYSFRYIADNDDDKMVILPV |
| IRIS_313-11169 | MLHHLKELAAVAGIHMILIYLCRFLLRRSRNVLFTVSNSLRFRLKVLTVLLYICLSVMLFYLFGSIMPLPPWGLVVGWVMALIAVELAYAFIFPYSFRYIADNDDDKMVILPV |
| IRIS_313-11170 | MLHHLKELAAVAGIHMILIYLCRFLLRRSRNVLFTVSNSLRFRLKVLTVLLYICLSVMLFYLFGSIMPLPPWGLVVGWVMALIAVELAYAFIFPYSFRYIADNDDDKMVILPV |
| IRIS_313-11171 | MLHHLKELAAVAGIHMILIYLCRFLLRRSRNVLFTVSNSLRFRLKVLTVLLYICLSVMLFYLFGSIMPLPPWGLVVGWVMALIAVELAYAFIFPYSFRYIADNDDDKMVILPV |
| IRIS_313-11172 | MLHHLKELAAVAGIHMILIYLCRFLLRRSRNVLFTVSNSLRFRLKVLTVLLYICLSVMLFYLFGSIMPLPPWGLVVGWVMALIAVELAYAFIFPYSFRYIADNDDDKMVILPV |
| IRIS_313-11173 | MLHHLKELAAVAGIHMILIYLCRFLLRRSRNVLFTVSNSLRFRLKVLTVLLYICLSVMLFYLFGSIMPLPPWGLVVGWVMALIAVELAYAFIFPYSFRYIADNDDDKMVILPV |
| IRIS_313-11174 | MLHHLKELAAVAGIHMILIYLCRFLLRRSRNVLFTVSNSLRFRLKVLTVLLYICLSVMLFYLFGSIMPLPPWGLVVGWVMALIAVELAYAFIFPYSFRYIADNDDDKMVILPV |
| IRIS_313-11175 | MLHHLKELAAVAGIHMILIYLCRFLLRRSRNVLFTVSNSLRFRLKVLTVLLYICLSVMLFYLFGSIMPLPPWGLVVGWVMALIAVELAYAFIFPYSFRYIADNDDDKMVILPV |
| IRIS_313-11176 | MLHHLKELAAVAGIHMILIYLCRFLLRRSRNVLFTVSNSLRFRLKVLTVLLYICLSVMLFYLFGSIMPLPPWGLVVGWVMALIAVELAYAFIFPYSFRYIADNDDDKMVILPV |
| IRIS_313-11177 | MLHHLKELAAVAGIHMILIYLCRFLLRRSRNVLFTVSNSLRFRLKVLTVLLYICLSVMLFYLFGSIMPLPPWGLVVGWVMALIAVELAYAFIFPYSFRYIADNDDDKMVILPV |
| IRIS_313-11178 | MLHHLKELAAVAGIHMILIYLCRFLLRRSRNVLFTVSNSLRFRLKVLTVLLYICLSVMLFYLFGSIMPLPPWGLVVGWVMALIAVELAYAFIFPYSFRYIADNDDDKMVILPV |
| IRIS_313-11179 | MLHHLKELAAVAGIHMILIYLCRFLLRRSRNVLFTVSNSLRFRLKVLTVLLYICLSVMLFYLFGSIMPLPPWGLVVGWVMALIAVELAYAFIFPYSFRYIADNDDDKMVILPV |
| IRIS_313-11180 | MLHHLKELAAVAGIHMILIYLCRFLLRRSRNVLFTVSNSLRFRLKVLTVLLYICLSVMLFYLFGSIMPLPPWGLVVGWVMALIAVELAYAFIFPYSFRYIADNDDDKMVILPV |
| IRIS_313-11181 | MLHHLKELAAVAGIHMILIYLCRFLLRRSRNVLFTVSNSLRFRLKVLTVLLYICLSVMLFYLFGSIMPLPPWGLVVGWVMALIAVELAYAFIFPYSFRYIADNDDDKMVILPV |
| IRIS_313-11182 | MLHHLKELAAVAGIHMILIYLCRFLLRRSRNVLFTVSNSLRFRLKVLTVLLYICLSVMLFYLFGSIMPLPPWGLVVGWVMALIAVELAYAFIFPYSFRYIADNDDDKMVILPV |
| IRIS_313-11183 | MLHQLKELAAVAGIHMILIYLCRFLLRRSRNVLFTVSNSLRFRLKVLTVLLYICLLVMLFYLFGSIMPLPPWGLVVGWVMALIAVELAYAFIFPYSFRYIADNDDDKMVILPV |
| IRIS_313-11189 | MLHHLKELAAVAGIHMILIYLCRFLLRRSRNVLFTVSNSLRFRLKVLTVLLYICLSVMLFYLFGSIMPLPPWGLVVGWVMALIAVELAYAFIFPYSFRYIADNDDDKMVILPV |
| IRIS_313-11191 | MLHHLKELAAVAGIHMILIYLCRFLLRRSRNVLFTVSNSLRFRLKVLTVLLYICLSVMLFYLFGSIMPLPPWGLVVGWVMALIAVELAYAFIFPYSFRYIADNDDDKMVILPV |
| IRIS_313-11192 | MLHHLKELAAVAGIHMILIYLCRFLLRRSRNVLFTVSNSLRFRLKVLTVLLYICLSVMLFYLFGSIMPLPPWGLVVGWVMALIAVELAYAFIFPYSFRYIADNDDDKMVILPV |
| IRIS_313-11193 | MLHHLKELAAVAGIHMILIYLCRFLLRRSRNVLFTVSNSLRFRLKVLTVLLYICLSVMLFYLFGSIMPLPPWGLVVGWVMALIAVELAYAFIFPYSFRYIADNDDDKMVILPV |
| IRIS_313-11194 | MLHHLKELAAVAGIHMILIYLCRFLLRRSRNVLFTVSNSLRFRLKVLTVLLYICLSVMLFYLFGSIMPLPPWGLVVGWVMALIAVELAYAFIFPYSFRYIADNDDDKMVILPV |
| IRIS_313-11195 | MLHQLKELAAVAGIHMILIYLCRFLLRRSRNVLFTVSNSLRFRLKVLTVLLYICLLVMLFYLFGSIMPLPPWGLVVGWVMALIAVELAYAFIFPYSFRYIADNDDDKMVILPV |
| IRIS_313-11196 | MLHHLKELAAVAGIHMILIYLCRFLLRRSRNVLFTVSNSLRFRLKVLTVLLYICLSVMLFYLFGSIMPLPPWGLVVGWVMALIAVELAYAFIFPYSFRYIADNDDDKMVILPV |
| IRIS_313-11197 | MLHHLKELAAVAGIHMILIYLCRFLLRRSRNVLFTVSNSLRFRLKVLTVLLYICLSVMLFYLFGSIMPLPPWGLVVGWVMALIAVELAYAFIFPYSFRYIADNDDDKMVILPV |
| IRIS_313-11198 | MLHHLKELAAVAGIHMILIYLCRFLLRRSRNVLFTVSNSLRFRLKVLTVLLYICLSVMLFYLFGSIMPLPPWGLVVGWVMALIAVELAYAFIFPYSFRYIADNDDDKMVILPV |
| IRIS_313-11199 | MLHHLKELAAVAGIHMILIYLCRFLLRRSRNVLFTVSNSLRFRLKVLTVLLYICLSVMLFYLFGSIMPLPPWGLVVGWVMALIAVELAYAFIFPYSFRYIADNDDDKMVILPV |
| IRIS_313-11200 | MLHHLKELAAVAGIHMILIYLCRFLLRRSRNVLFTVSNSLRFRLKVLTVLLYICLSVMLFYLFGSIMPLPPWGLVVGWVMALIAVELAYAFIFPYSFRYIADNDDDKMVILPV |
| IRIS_313-11201 | MLHHLKELAAVAGIHMILIYLCRFLLRRSRNVLFTVSNSLRFRLKVLTVLLYICLSVMLFYLFGSIMPLPPWGLVVGWVMALIAVELAYAFIFPYSFRYIADNDDDKMVILPV |
| IRIS_313-11202 | MLHHLKELAAVAGIHMILIYLCRFLLRRSRNVLFTVSNSLRFRLKVLTVLLYICLSVMLFYLFGSIMPLPPWGLVVGWVMALIAVELAYAFIFPYSFRYIADNDDDKMVILPV |
| IRIS_313-11203 | MLHHLKELAAVAGIHMILIYLCRFLLRRSRNVLFTVSNSLRFRLKVLTVLLYICLSVMLFYLFGSIMPLPPWGLVVGWVMALIAVELAYAFIFPYSFRYIADNDDDKMVILPV |
| IRIS_313-11204 | MLHHLKELAAVAGIHMILIYLCRFLLRRSRNVLFTVSNSLRFRLKVLTVLLYICLSVMLFYLFGSIMPLPPWGLVVGWVMALIAVELAYAFIFPYSFRYIADNDDDKMVILPV |
| IRIS_313-11205 | MLHHLKELAAVAGIHMILIYLCRFLLRRSRNVLFTVSNSLRFRLKVLTVLLYICLSVMLFYLFGSIMPLPPWGLVVGWVMALIAVELAYAFIFPYSFRYIADNDDDKMVILPV |
| IRIS_313-11206 | MLHHLKELAAVAGIHMILIYLCRFLLRRSRNVLFTVSNSLRFRLKVLTVLLYICLSVMLFYLFGSIMPLPPWGLVVGWVMALIAVELAYAFIFPYSFRYIADNDDDKMVILPV |
| IRIS_313-11207 | MLHHLKELAAVAGIHMILIYLCRFLLRRSRNVLFTVSNSLRFRLKVLTVLLYICLSVMLFYLFGSIMPLPPWGLVVGWVMALIAVELAYAFIFPYSFRYIADNDDDKMVILPV |
| IRIS_313-11208 | MLHHLKELAAVAGIHMILIYLCRFLLRRSRNVLFTVSNSLRFRLKVLTVLLYICLSVMLFYLFGSIMPLPPWGLVVGWVMALIAVELAYAFIFPYSFRYIADNDDDKMVILPV |
| IRIS_313-11209 | MLHHLKELAAVAGIHMILIYLCRFLLRRSRNVLFTVSNSLRFRLKVLTVLLYICLSVMLFYLFGSIMPLPPWGLVVGWVMALIAVELAYAFIFPYSFRYIADNDDDKMVILPV |
| IRIS_313-11210 | MLHHLKELAAVAGIHMILIYLCRFLLRRSRNVLFTVSNSLRFRLKVLTVLLYICLSVMLFYLFGSIMPLPPWGLVVGWVMALIAVELAYAFIFPYSFRYIADNDDDKMVILPV |

| | |
|---|---|
| IRIS_313-11211 | MLHHLKELAAVAGIHMILIYLCRFLLRRSRNVLFTVSNSLRFRLKVLTVLLYICLSVMLFYLFGSIMPLPPWGLVVGWVMALIAVELAYAFIFPYSFRYIADNDDDKMVILPV |
| IRIS_313-11212 | MLHHLKELAAVAGIHMILIYLCRFLLRRSRNVLFTVSNSLRFRLKVLTVLLYICLSVMLFYLFGSIMPLPPWGLVVGWVMALIAVELAYAFIFPYSFRYIADNDDDKMVILPV |
| IRIS_313-11213 | MLHHLKELAAVAGIHMILIYLCRFLLRRSRNVLFTVSNSLRFRLKVLTVLLYICLSVMLFYLFGSIMPLPPWGLVVGWVMALIAVELAYAFIFPYSFRYIADNDDDKMVILPV |
| IRIS_313-11214 | MLHHLKELAAVAGIHMILIYLCRFLLRRSRNVLFTVSNSLRFRLKVLTVLLYICLSVMLFYLFGSIMPLPPWGLVVGWVMALIAVELAYAFIFPYSFRYIADNDDDKMVILPV |
| IRIS_313-11216 | MLHHLKELAAVAGIHMILIYLCRFLLRRSRNVLFTVSNSLRFRLKVLTVLLYICLSVMLFYLFGSIMPLPPWGLVVGWVMALIAVELAYAFIFPYSFRYIADNDDDKMVILPV |
| IRIS_313-11217 | MLHHLKELAAVAGIHMILIYLCRFLLRRSRNVLFTVSNSLRFRLKVLTVLLYICLSVMLFYLFGSIMPLPPWGLVVGWVMALIAVELAYAFIFPYSFRYIADNDDDKMVILPV |
| IRIS_313-11218 | MLHHLKELAAVAGIHMILIYLCRFLLRRSRNVLFTVSNSLRFRLKVLTVLLYICLSVMLFYLFGSIMPLPPWGLVVGWVMALIAVELAYAFIFPYSFRYIADNDDDKMVILPV |
| IRIS_313-11219 | MLHHLKELAAVAGIHMILIYLCRFLLRRSRNVLFTVSNSLRFRLKVLTVLLYICLSVMLFYLFGSIMPLPPWGLVVGWVMALIAVELAYAFIFPYSFRYIADNDDDKMVILPV |
| IRIS_313-11220 | MLHHLKELAAVAGIHMILIYLCRFLLRRSRNVLFTVSNSLRFRLKVLTVLLYICLSVMLFYLFGSIMPLPPWGLVVGWVMALIAVELAYAFIFPYSFRYIADNDDDKMVILPV |
| IRIS_313-11221 | MLHHLKELAAVAGIHMILIYLCRFLLRRSRNVLFTVSNSLRFRLKVLTVLLYICLSVMLFYLFGSIMPLPPWGLVVGWVMALIAVELAYAFIFPYSFRYIADNDDDKMVILPV |
| IRIS_313-11222 | MLHHLKELAAVAGIHMILIYLCRFLLRRSRNVLFTVSNSLRFRLKVLTVLLYICLSVMLFYLFGSIMPLPPWGLVVGWVMALIAVELAYAFIFPYSFRYIADNDDDKMVILPV |
| IRIS_313-11223 | MLHHLKELAAVAGIHMILIYLCRFLLRRSRNVLFTVSNSLRFRLKVLTVLLYICLSVMLFYLFGSIMPLPPWGLVVGWVMALIAVELAYAFIFPYSFRYIADNDDDKMVILPV |
| IRIS_313-11224 | MLHHLKELAAVAGIHMILIYLCRFLLRRSRNVLFTVSNSLRFRLKVLTVLLYICLSVMLFYLFGSIMPLPPWGLVVGWVMALIAVELAYAFIFPYSFRYIADNDDDKMVILPV |
| IRIS_313-11225 | MLHHLKELAAVAGIHMILIYLCRFLLRRSRNVLFTVSNSLRFRLKVLTVLLYICLSVMLFYLFGSIMPLPPWGLVVGWVMALIAVELAYAFIFPYSFRYIADNDDDKMVILPV |
| IRIS_313-11226 | MLHHLKELAAVAGIHMILIYLCRFLLRRSRNVLFTVSNSLRFRLKVLTVLLYICLSVMLFYLFGSIMPLPPWGLVVGWVMALIAVELAYAFIFPYSFRYIADNDDDKMVILPV |
| IRIS_313-11227 | MLHHLKELAAVAGIHMILIYLCRFLLRRSRNVLFTVSNSLRFRLKVLTVLLYICLSVMLFYLFGSIMPLPPWGLVVGWVMALIAVELAYAFIFPYSFRYIADNDDDKMVILPV |
| IRIS_313-11228 | MLHHLKELAAVAGIHMILIYLCRFLLRRSRNVLFTVSNSLRFRLKVLTVLLYICLSVMLFYLFGSIMPLPPWGLVVGWVMALIAVELAYAFIFPYSFRYIADNDDDKMVILPV |
| IRIS_313-11229 | MLHHLKELAAVAGIHMILIYLCRFLLRRSRNVLFTVSNSLRFRLKVLTVLLYICLSVMLFYLFGSIMPLPPWGLVVGWVMALIAVELAYAFIFPYSFRYIADNDDDKMVILPV |
| IRIS_313-11230 | MLHHLKELAAVAGIHMILIYLCRFLLRRSRNVLFTVSNSLRFRLKVLTVLLYICLSVMLFYLFGSIMPLPPWGLVVGWVMALIAVELAYAFIFPYSFRYIADNDDDKMVILPV |
| IRIS_313-11231 | MLHHLKELAAVAGIHMILIYLCRFLLRRSRNVLFTVSNSLRFRLKVLTVLLYICLSVMLFYLFGSIMPLPPWGLVVGWVMALIAVELAYAFIFPYSFRYIADNDDDKMVILPV |
| IRIS_313-11232 | MLHHLKELAAVAGIHMILIYLCRFLLRRSRNVLFTVSNSLRFRLKVLTVLLYICLSVMLFYLFGSIMPLPPWGLVVGWVMALIAVELAYAFIFPYSFRYIADNDDDKMVILPV |
| IRIS_313-11233 | MLHHLKELAAVAGIHMILIYLCRFLLRRSRNVLFTVSNSLRFRLKVLTVLLYICLSVMLFYLFGSIMPLPPWGLVVGWVMALIAVELAYAFIFPYSFRYIADNDDDKMVILPV |
| IRIS_313-11234 | MLHHLKELAAVAGIHMILIYLCRFLLRRSRNVLFTVSNSLRFRLKVLTVLLYICLSVMLFYLFGSIMPLPPWGLVVGWVMALIAVELAYAFIFPYSFRYIADNDDDKMVILPV |
| IRIS_313-11235 | MLHHLKELAAVAGIHMILIYLCRFLLRRSRNVLFTVSNSLRFRLKVLTVLLYICLSVMLFYLFGSIMPLPPWGLVVGWVMALIAVELAYAFIFPYSFRYIADNDDDKMVILPV |
| IRIS_313-11236 | MLHHLKELAAVAGIHMILIYLCRFLLRRSRNVLFTVSNSLRFRLKVLTVLLYICLSVMLFYLFGSIMPLPPWGLVVGWVMALIAVELAYAFIFPYSFRYIADNDDDKMVILPV |
| IRIS_313-11237 | MLHHLKELAAVAGIHMILIYLCRFLLRRSRNVLFTVSNSLRFRLKVLTVLLYICLSVMLFYLFGSIMPLPPWGLVVGWVMALIAVELAYAFIFPYSFRYIADNDDDKMVILPV |
| IRIS_313-11238 | MLHHLKELAAVAGIHMILIYLCRFLLRRSRNVLFTVSNSLRFRLKVLTVLLYICLSVMLFYLFGSIMPLPPWGLVVGWVMALIAVELAYAFIFPYSFRYIADNDDDKMVILPV |
| IRIS_313-11239 | MLHHLKELAAVAGIHMILIYLCRFLLRRSRNVLFTVSNSLRFRLKVLTVLLYICLSVMLFYLFGSIMPLPPWGLVVGWVMALIAVELAYAFIFPYSFRYIADNDDDKMVILPV |
| IRIS_313-11240 | MLHHLKELAAVAGIHMILIYLCRFLLRRSRNVLFTVSNSLRFRLKVLTVLLYICLSVMLFYLFGSIMPLPPWGLVVGWVMALIAVELAYAFIFPYSFRYIADNDDDKMVILPV |
| IRIS_313-11241 | MLHHLKELAAVAGIHMILIYLCRFLLRRSRNVLFTVSNSLRFRLKVLTVLLYICLSVMLFYLFGSIMPLPPWGLVVGWVMALIAVELAYAFIFPYSFRYIADNDDDKMVILPV |
| IRIS_313-11242 | MLHHLKELAAVAGIHMILIYLCRFLLRRSRNVLFTVSNSLRFRLKVLTVLLYICLSVMLFYLFGSIMPLPPWGLVVGWVMALIAVELAYAFIFPYSFRYIADNDDDKMVILPV |
| IRIS_313-11243 | MLHHLKELAAVAGIHMILIYLCRFLLRRSRNVLFTVSNSLRFRLKVLTVLLYICLSVMLFYLFGSIMPLPPWGLVVGWVMALIAVELAYAFIFPYSFRYIADNDDDKMVILPV |
| IRIS_313-11244 | MLHHLKELAAVAGIHMILIYLCRFLLRRSRNVLFTVSNSLRFRLKVLTVLLYICLSVMLFYLFGSIMPLPPWGLVVGWVMALIAVELAYAFIFPYSFRYIADNDDDKMVILPV |
| IRIS_313-11245 | MLHHLKELAAVAGIHMILIYLCRFLLRRSRNVLFTVSNSLRFRLKVLTVLLYICLSVMLFYLFGSIMPLPPWGLVVGWVMALIAVELAYAFIFPYSFRYIADNDDDKMVILPV |
| IRIS_313-11246 | MLHHLKELAAVAGIHMILIYLCRFLLRRSRNVLFTVSNSLRFRLKVLTVLLYICLSVMLFYLFGSIMPLPPWGLVVGWVMALIAVELAYAFIFPYSFRYIADNDDDKMVILPV |
| IRIS_313-11247 | MLHHLKELAAVAGIHMILIYLCRFLLRRSRNVLFTVSNSLRFRLKVLTVLLYICLSVMLFYLFGSIMPLPPWGLVVGWVMALIAVELAYAFIFPYSFRYIADNDDDKMVILPV |
| IRIS_313-11248 | MLHHLKELAAVAGIHMILIYLCRFLLRRSRNVLFTVSNSLRFRLKVLTVLLYICLSVMLFYLFGSIMPLPPWGLVVGWVMALIAVELAYAFIFPYSFRYIADNDDDKMVILPV |
| IRIS_313-11249 | MLHHLKELAAVAGIHMILIYLCRFLLRRSRNVLFTVSNSLRFRLKVLTVLLYICLSVMLFYLFGSIMPLPPWGLVVGWVMALIAVELAYAFIFPYSFRYIADNDDDKMVILPV |
| IRIS_313-11250 | MLHHLKELAAVAGIHMILIYLCRFLLRRSRNVLFTVSNSLRFRLKVLTVLLYICLSVMLFYLFGSIMPLPPWGLVVGWVMALIAVELAYAFIFPYSFRYIADNDDDKMVILPV |
| IRIS_313-11251 | MLHHLKELAAVAGIHMILIYLCRFLLRRSRNVLFTVSNSLRFRLKVLTVLLYICLSVMLFYLFGSIMPLPPWGLVVGWVMALIAVELAYAFIFPYSFRYIADNDDDKMVILPV |
| IRIS_313-11252 | MLHHLKELAAVAGIHMILIYLCRFLLRRSRNVLFTVSNSLRFRLKVLTVLLYICLSVMLFYLFGSIMPLPPWGLVVGWVMALIAVELAYAFIFPYSFRYIADNDDDKMVILPV |
| IRIS_313-11253 | MLHHLKELAAVAGIHMILIYLCRFLLRRSRNVLFTVSNSLRFRLKVLTVLLYICLSVMLFYLFGSIMPLPPWGLVVGWVMALIAVELAYAFIFPYSFRYIADNDDDKMVILPV |
| IRIS_313-11254 | MLHHLKELAAVAGIHMILIYLCRFLLRRSRNVLFTVSNSLRFRLKVLTVLLYICLSVMLFYLFGSIMPLPPWGLVVGWVMALIAVELAYAFIFPYSFRYIADNDDDKMVILPV |
| IRIS_313-11255 | MLHHLKELAAVAGIHMILIYLCRFLLRRSRNVLFTVSNSLRFRLKVLTVLLYICLSVMLFYLFGSIMPLPPWGLVVGWVMALIAVELAYAFIFPYSFRYIADNDDDKMVILPV |
| IRIS_313-11256 | MLHHLKELAAVAGIHMILIYLCRFLLRRSRNVLFTVSNSLRFRLKVLTVLLYICLSVMLFYLFGSIMPLPPWGLVVGWVMALIAVELAYAFIFPYSFRYIADNDDDKMVILPV |
| IRIS_313-11257 | MLHHLKELAAVAGIHMILIYLCRFLLRRSRNVLFTVSNSLRFRLKVLTVLLYICLSVMLFYLFGSIMPLPPWGLVVGWVMALIAVELAYAFIFPYSFRYIADNDDDKMVILPV |
| IRIS_313-11258 | MLHHLKELAAVAGIHMILIYLCRFLLRRSRNVLFTVSNSLRFRLKVLTVLLYICLSVMLFYLFGSIMPLPPWGLVVGWVMALIAVELAYAFIFPYSFRYIADNDDDKMVILPV |
| IRIS_313-11259 | MLHHLKELAAVAGIHMILIYLCRFLLRRSRNVLFTVSNSLRFRLKVLTVLLYICLSVMLFYLFGSIMPLPPWGLVVGWVMALIAVELAYAFIFPYSFRYIADNDDDKMVILPV |
| IRIS_313-11260 | MLHHLKELAAVAGIHMILIYLCRFLLRRSRNVLFTVSNSLRFRLKVLTVLLYICLSVMLFYLFGSIMPLPPWGLVVGWVMALIAVELAYAFIFPYSFRYIADNDDDKMVILPV |
| IRIS_313-11261 | MLHHLKELAAVAGIHMILIYLCRFLLRRSRNVLFTVSNSLRFRLKVLTVLLYICLSVMLFYLFGSIMPLPPWGLVVGWVMALIAVELAYAFIFPYSFRYIADNDDDKMVILPV |
| IRIS_313-11262 | MLHHLKELAAVAGIHMILIYLCRFLLRRSRNVLFTVSNSLRFRLKVLTVLLYICLSVMLFYLFGSIMPLPPWGLVVGWVMALIAVELAYAFIFPYSFRYIADNDDDKMVILPV |
| IRIS_313-11263 | MLHHLKELAAVAGIHMILIYLCRFLLRRSRNVLFTVSNSLRFRLKVLTVLLYICLSVMLFYLFGSIMPLPPWGLVVGWVMALIAVELAYAFIFPYSFRYIADNDDDKMVILPV |
| IRIS_313-11264 | MLHHLKELAAVAGIHMILIYLCRFLLRRSRNVLFTVSNSLRFRLKVLTVLLYICLSVMLFYLFGSIMPLPPWGLVVGWVMALIAVELAYAFIFPYSFRYIADNDDDKMVILPV |
| IRIS_313-11265 | MLHHLKELAAVAGIHMILIYLCRFLLRRSRNVLFTVSNSLRFRLKVLTVLLYISLSAMLLYLFGSIMPLPPWGLVVGWVMALIAVELAYAFIFPYSFRYIADNDDDKMVILPV |
| IRIS_313-11266 | MLHHLKELAAVAGIHMILIYLCRFLLRRSRNVLFTVSNSLRFRLKVLTVLLYICLSVMLFYLFGSIMPLPPWGLVVGWVMALIAVELAYAFIFPYSFRYIADNDDDKMVILPV |
| IRIS_313-11267 | MLHHLKELAAVAGIHMILIYLCRFLLRRSRNVLFTVSNSLRFRLKVLTVLLYICLSVMLFYLFGSIMPLPPWGLVVGWVMALIAVELAYAFIFPYSFRYIADNDDDKMVILPV |
| IRIS_313-11268 | MLHHLKELAAVAGIHMILIYLCRFLLRRSRNVLFTVSNSLRFRLKVLTVLLYICLSVMLFYLFGSIMPLPPWGLVVGWVMALIAVELAYAFIFPYSFRYIADNDDDKMVILPV |
| IRIS_313-11269 | MLHHLKELAAVAGIHMILIYLCRFLLRRSRNVLFTVSNSLRFRLKVLTVLLYICLSVMLFYLFGSIMPLPPWGLVVGWVMALIAVELAYAFIFPYSFRYIADNDDDKMVILPV |
| IRIS_313-11270 | MLHHLKELAAVAGIHMILIYLCRFLLRRSRNVLFTVSNSLRFRLKVLTVLLYICLSVMLFYLFGSIMPLPPWGLVVGWVMALIAVELAYAFIFPYSFRYIADNDDDKMVILPV |
| IRIS_313-11271 | MLHHLKELAAVAGIHMILIYLCRFLLRRSRNVLFTVSNSLRFRLKVLTVLLYICLSVMLFYLFGSIMPLPPWGLVVGWVMALIAVELAYAFIFPYSFRYIADNDDDKMVILPV |
| IRIS_313-11272 | MLHHLKELAAVAGIHMILIYLCRFLLRRSRNVLFTVSNSLRFRLKVLTVLLYICLSVMLFYLFGSIMPLPPWGLVVGWVMALIAVELAYAFIFPYSFRYIADNDDDKMVILPV |
| IRIS_313-11273 | MLHHLKELAAVAGIHMILIYLCRFLLRRSRNVLFTVSNSLRFRLKVLTVLLYICLSVMLFYLFGSIMPLPPWGLVVGWVMALIAVELAYAFIFPYSFRYIADNDDDKMVILPV |
| IRIS_313-11274 | MLHHLKELAAVAGIHMILIYLCRFLLRRSRNVLFTVSNSLRFRLKVLTVLLYICLSVMLFYLFGSIMPLPPWGLVVGWVMALIAVELAYAFIFPYSFRYIADNDDDKMVILPV |
| IRIS_313-11275 | MLHHLKELAAVAGIHMILIYLCRFLLRRSRNVLFTVSNSLRFRLKVLTVLLYICLSVMLFYLFGSIMPLPPWGLVVGWVMALIAVELAYAFIFPYSFRYIADNDDDKMVILPV |
| IRIS_313-11276 | MLHHLKELAAVAGIHMILIYLCRFLLRRSRNVLFTVSNSLRFRLKVLTVLLYICLSVMLFYLFGSIMPLPPWGLVVGWVMALIAVELAYAFIFPYSFRYIADNDDDKMVILPV |
| IRIS_313-11277 | MLHHLKELAAVAGIHMILIYLCRFLLRRSRNVLFTVSNSLRFRLKVLTVLLYICLSVMLFYLFGSIMPLPPWGLVVGWVMALIAVELAYAFIFPYSFRYIADNDDDKMVILPV |
| IRIS_313-11278 | MLHHLKELAAVAGIHMILIYLCRFLLRRSRNVLFTVSNSLRFRLKVLTVLLYICLSVMLFYLFGSIMPLPPWGLVVGWVMALIAVELAYAFIFPYSFRYIADNDDDKMVILPV |
| IRIS_313-11279 | MLHHLKELAAVAGIHMILIYLCRFLLRRSRNVLFTVSNSLRFRLKVLTVLLYICLSVMLFYLFGSIMPLPPWGLVVGWVMALIAVELAYAFIFPYSFRYIADNDDDKMVILPV |
| IRIS_313-11280 | MLHHLKELAAVAGIHMILIYLCRFLLRRSRNVLFTVSNSLRFRLKVLTVLLYICLSVMLFYLFGSIMPLPPWGLVVGWVMALIAVELAYAFIFPYSFRYIADNDDDKMVILPV |
| IRIS_313-11281 | MLHHLKELAAVAGIHMILIYLCRFLLRRSRNVLFTVSNSLRFRLKVLTVLLYICLSVMLFYLFGSIMPLPPWGLVVGWVMALIAVELAYAFIFPYSFRYIADNDDDKMVILPV |
| IRIS_313-11282 | MLHHLKELAAVAGIHMILIYLCRFLLRRSRNVLFTVSNSLRFRLKVLTVLLYICLSVMLFYLFGSIMPLPPWGLVVGWVMALIAVELAYAFIFPYSFRYIADNDDDKMVILPV |
| IRIS_313-11283 | MLHHLKELAAVAGIHMILIYLCRFLLRRSRNVLFTVSNSLRFRLKVLTVLLYICLSVMLFYLFGSIMPLPPWGLVVGWVMALIAVELAYAFIFPYSFRYIADNDDDKMVILPV |
| IRIS_313-11284 | MLHHLKELAAVAGIHMILIYLCRFLLRRSRNVLFTVSNSLRFRLKVLTVLLYICLSVMLFYLFGSIMPLPPWGLVVGWVMALIAVELAYAFIFPYSFRYIADNDDDKMVILPV |
| IRIS_313-11285 | MLHHLKELAAVAGIHMILIYLCRFLLRRSRNVLFTVSNSLRFRLKVLTVLLYICLSVMLFYLFGSIMPLPPWGLVVGWVMALIAVELAYAFIFPYSFRYIADNDDDKMVILPV |
| IRIS_313-11286 | MLHHLKELAAVAGIHMILIYLCRFLLRRSRNVLFTVSNSLRFRLKVLTVLLYICLSVMLFYLFGSIMPLPPWGLVVGWVMALIAVELAYAFIFPYSFRYIADNDDDKMVILPV |
| IRIS_313-11287 | MLHHLKELAAVAGIHMILIYLCRFLLRRSRNVLFTVSNSLRFRLKVLTVLLYICLSVMLFYLFGSIMPLPPWGLVVGWVMALIAVELAYAFIFPYSFRYIADNDDDKMVILPV |
| IRIS_313-11288 | MLHHLKELAAVAGIHMILIYLCRFLLRRSRNVLFTVSNSLRFRLKVLTVLLYICLSVMLFYLFGSIMPLPPWGLVVGWVMALIAVELAYAFIFPYSFRYIADNDDDKMVILPV |
| IRIS_313-11289 | MLHHLKELAAVAGIHMILIYLCRFLLRRSRNVLFTVSNSLRFRLKVLTVLLYICLSVMLFYLFGSIMPLPPWGLVVGWVMALIAVELAYAFIFPYSFRYIADNDDDKMVILPV |
| IRIS_313-11290 | MLHHLKELAAVAGIHMILIYLCRFLLRRSRNVLFTVSNSLRFRLKVLTVLLYICLSVMLFYLFGSIMPLPPWGLVVGWVMALIAVELAYAFIFPYSFRYIADNDDDKMVILPV |
| IRIS_313-11291 | MLHHLKELAAVAGIHMILIYLCRFLLRRSRNVLFTVSNSLRFRLKVLTVLLYICLSVMLFYLFGSIMPLPPWGLVVGWVMALIAVELAYAFIFPYSFRYIADNDDDKMVILPV |
| IRIS_313-11292 | MLHHLKELAAVAGIHMILIYLCRFLLRRSRNVLFTVSNSLRFRLKVLTVLLYICLSVMLFYLFGSIMPLPPWGLVVGWVMALIAVELAYAFIFPYSFRYIADNDDDKMVILPV |
| IRIS_313-11293 | MLHHLKELAAVAGIHMILIYLCRFLLRRSRNVLFTVSNSLRFRLKVLTVLLYICLSVMLFYLFGSIMPLPPWGLVVGWVMALIAVELAYAFIFPYSFRYIADNDDDKMVILPV |
| IRIS_313-11294 | MLHHLKELAAVAGIHMILIYLCRFLLRRSRNVLFTVSNSLRFRLKVLTVLLYICLSVMLFYLFGSIMPLPPWGLVVGWVMALIAVELAYAFIFPYSFRYIADNDDDKMVILPV |
| IRIS_313-11295 | MLHHLKELAAVAGIHMILIYLCRFLLRRSRNVLFTVSNSLRFRLKVLTVLLYICLSVMLFYLFGSIMPLPPWGLVVGWVMALIAVELAYAFIFPYSFRYIADNDDDKMVILPV |
| IRIS_313-11296 | MLHHLKELAAVAGIHMILIYLCRFLLRRSRNVLFTVSNSLRFRLKVLTVLLYICLSVMLFYLFGSIMPLPPWGLVVGWVMALIAVELAYAFIFPYSFRYIADNDDDKMVILPV |
| IRIS_313-11297 | MLHHLKELAAVAGIHMILIYLCRFLLRRSRNVLFTVSNSLRFRLKVLTVLLYICLSVMLFYLFGSIMPLPPWGLVVGWVMALIAVELAYAFIFPYSFRYIADNDDDKMVILPV |
| IRIS_313-11298 | MLHHLKELAAVAGIHMILIYLCRFLLRRSRNVLFTVSNSLRFRLKVLTVLLYICLSVMLFYLFGSIMPLPPWGLVVGWVMALIAVELAYAFIFPYSFRYIADNDDDKMVILPV |
| IRIS_313-11299 | MLHHLKELAAVAGIHMILIYLCRFLLRRSRNVLFTVSNSLRFRLKVLTVLLYICLSVMLFYLFGSIMPLPPWGLVVGWVMALIAVELAYAFIFPYSFRYIADNDDDKMVILPV |
| IRIS_313-11300 | MLHHLKELAAVAGIHMILIYLCRFLLRRSRNVLFTVSNSLRFRLKVLTVLLYICLSVMLFYLFGSIMPLPPWGLVVGWVMALIAVELAYAFIFPYSFRYIADNDDDKMVILPV |
| IRIS_313-11301 | MLHHLKELAAVAGIHMILIYLCRFLLRRSRNVLFTVSNSLRFRLKVLTVLLYICLSVMLFYLFGSIMPLPPWGLVVGWVMALIAVELAYAFIFPYSFRYIADNDDDKMVILPV |
| IRIS_313-11302 | MLHHLKELAAVAGIHMILIYLCRFLLRRSRNVLFTVSNSLRFRLKVLTVLLYICLSVMLFYLFGSIMPLPPWGLVVGWVMALIAVELAYAFIFPYSFRYIADNDDDKMVILPV |
| IRIS_313-11303 | MLHHLKELAAVAGIHMILIYLCRFLLRRSRNVLFTVSNSLRFRLKVLTVLLYICLSVMLFYLFGSIMPLPPWGLVVGWVMALIAVELAYAFIFPYSFRYIADNDDDKMVILPV |
| IRIS_313-11304 | MLHHLKELAAVAGIHMILIYLCRFLLRRSRNVLFTVSNSLRFRLKVLTVLLYICLSVMLFYLFGSIMPLPPWGLVVGWVMALIAVELAYAFIFPYSFRYIADNDDDKMVILPV |
| IRIS_313-11305 | MLHHLKELAAVAGIHMILIYLCRFLLRRSRNVLFTVSNSLRFRLKVLTVLLYICLSVMLFYLFGSIMPLPPWGLVVGWVMALIAVELAYAFIFPYSFRYIADNDDDKMVILPV |
| IRIS_313-11306 | MLHHLKELAAVAGIHMILIYLCRFLLRRSRNVLFTVSNSLRFRLKVLTVLLYICLSVMLFYLFGSIMPLPPWGLVVGWVMALIAVELAYAFIFPYSFRYIADNDDDKMVILPV |
| IRIS_313-11307 | MLHHLKELAAVAGIHMILIYLCRFLLRRSRNVLFTVSNSLRFRLKVLTVLLYICLSVMLFYLFGSIMPLPPWGLVVGWVMALIAVELAYAFIFPYSFRYIADNDDDKMVILPV |
| IRIS_313-11308 | MLHHLKELAAVAGIHMILIYLCRFLLRRSRNVLFTVSNSLRFRLKVLTVLLYICLSVMLFYLFGSIMPLPPWGLVVGWVMALIAVELAYAFIFPYSFRYIADNDDDKMVILPV |
| IRIS_313-11309 | MLHHLKELAAVAGIHMILIYLCRFLLRRSRNVLFTVSNSLRFRLKVLTVLLYICLSVMLFYLFGSIMPLPPWGLVVGWVMALIAVELAYAFIFPYSFRYIADNDDDKMVILPV |
| IRIS_313-11310 | MLHHLKELAAVAGIHMILIYLCRFLLRRSRNVLFTVSNSLRFRLKVLTVLLYICLSVMLFYLFGSIMPLPPWGLVVGWVMALIAVELAYAFIFPYSFRYIADNDDDKMVILPV |
| IRIS_313-11311 | MLHHLKELAAVAGIHMILIYLCRFLLRRSRNVLFTVSNSLRFRLKVLTVLLYICLSVMLFYLFGSIMPLPPWGLVVGWVMALIAVELAYAFIFPYSFRYIADNDDDKMVILPV |
| IRIS_313-11312 | MLHHLKELAAVAGIHMILIYLCRFLLRRSRNVLFTVSNSLRFRLKVLTVLLYICLSVMLFYLFGSIMPLPPWGLVVGWVMALIAVELAYAFIFPYSFRYIADNDDDKMVILPV |
| IRIS_313-11313 | MLHHLKELAAVAGIHMILIYLCRFLLRRSRNVLFTVSNSLRFRLKVLTVLLYICLSVMLFYLFGSIMPLPPWGLVVGWVMALIAVELAYAFIFPYSFRYIADNDDDKMVILPV |
| IRIS_313-11314 | MLHHLKELAAVAGIHMILIYLCRFLLRRSRNVLFTVSNSLRFRLKVLTVLLYICLSVMLFYLFGSIMPLPPWGLVVGWVMALIAVELAYAFIFPYSFRYIADNDDDKMVILPV |
| IRIS_313-11315 | MLHHLKELAAVAGIHMILIYLCRFLLRRSRNVLFTVSNSLRFRLKVLTVLLYICLSVMLFYLFGSIMPLPPWGLVVGWVMALIAVELAYAFIFPYSFRYIADNDDDKMVILPV |
| IRIS_313-11316 | MLHHLKELAAVAGIHMILIYLCRFLLRRSRNVLFTVSNSLRFRLKVLTVLLYICLSVMLFYLFGSIMPLPPWGLVVGWVMALIAVELAYAFIFPYSFRYIADNDDDKMVILPV |
| IRIS_313-11317 | MLHHLKELAAVAGIHMILIYLCRFLLRRSRNVLFTVSNSLRFRLKVLTVLLYICLSVMLFYLFGSIMPLPPWGLVVGWVMALIAVELAYAFIFPYSFRYIADNDDDKMVILPV |
| IRIS_313-11318 | MLHHLKELAAVAGIHMILIYLCRFLLRRSRNVLFTVSNSLRFRLKVLTVLLYICLSVMLFYLFGSIMPLPPWGLVVGWVMALIAVELAYAFIFPYSFRYIADNDDDKMVILPV |
| IRIS_313-11319 | MLHHLKELAAVAGIHMILIYLCRFLLRRSRNVLFTVSNSLRFRLKVLTVLLYICLSVMLFYLFGSIMPLPPWGLVVGWVMALIAVELAYAFIFPYSFRYIADNDDDKMVILPV |
| IRIS_313-11320 | MLHHLKELAAVAGIHMILIYLCRFLLRRSRNVLFTVSNSLRFRLKVLTVLLYICLSVMLFYLFGSIMPLPPWGLVVGWVMALIAVELAYAFIFPYSFRYIADNDDDKMVILPV |
| IRIS_313-11321 | MLHHLKELAAVAGIHMILIYLCRFLLRRSRNVLFTVSNSLRFRLKVLTVLLYICLSVMLFYLFGSIMPLPPWGLVVGWVMALIAVELAYAFIFPYSFRYIADNDDDKMVILPV |
| IRIS_313-11322 | MLHHLKELAAVAGIHMILIYLCRFLLRRSRNVLFTVSNSLRFRLKVLTVLLYICLSVMLFYLFGSIMPLPPWGLVVGWVMALIAVELAYAFIFPYSFRYIADNDDDKMVILPV |
| IRIS_313-11323 | MLHHLKELAAVAGIHMILIYLCRFLLRRSRNVLFTVSNSLRFRLKVLTVLLYICLSVMLFYLFGSIMPLPPWGLVVGWVMALIAVELAYAFIFPYSFRYIADNDDDKMVILPV |
| IRIS_313-11324 | MLHHLKELAAVAGIHMILIYLCRFLLRRSRNVLFTVSNSLRFRLKVLTVLLYICLSVMLFYLFGSIMPLPPWGLVVGWVMALIAVELAYAFIFPYSFRYIADNDDDKMVILPV |
| IRIS_313-11325 | MLHHLKELAAVAGIHMILIYLCRFLLRRSRNVLFTVSNSLRFRLKVLTVLLYICLSVMLFYLFGSIMPLPPWGLVVGWVMALIAVELAYAFIFPYSFRYIADNDDDKMVILPV |
| IRIS_313-11326 | MLHHLKELAAVAGIHMILIYLCRFLLRRSRNVLFTVSNSLRFRLKVLTVLLYICLSVMLFYLFGSIMPLPPWGLVVGWVMALIAVELAYAFIFPYSFRYIADNDDDKMVILPV |
| IRIS_313-11327 | MLHHLKELAAVAGIHMILIYLCRFLLRRSRNVLFTVSNSLRFRLKVLTVLLYICLSVMLFYLFGSIMPLPPWGLVVGWVMALIAVELAYAFIFPYSFRYIADNDDDKMVILPV |
| IRIS_313-11328 | MLHQLKELAAVAGIHMILIYLCRFLLRRSRNVLFTVSNSLRFRLKVLTVLLYICLSVMLFYLFGSIMPLPPWGLVVGWVMALIAVELAYAFIFPYSFRYIADNDDDKMVILPV |
| IRIS_313-11329 | MLHHLKELAAVAGIHMILIYLCRFLLRRSRNVLFTVSNSLRFRLKVLTVLLYICLSVMLFYLFGSIMPLPPWGLVVGWVMALIAVELAYAFIFPYSFRYIADNDDDKMVILPV |
| IRIS_313-11330 | MLHHLKELAAVAGIHMILIYLCRFLLRRSRNVLFTVSNSLRFRLKVLTVLLYICLSVMLFYLFGSIMPLPPWGLVVGWVMALIAVELAYAFIFPYSFRYIADNDDDKMVILPV |
| IRIS_313-11331 | MLHHLKELAAVAGIHMILIYLCRFLLRRSRNVLFTVSNSLRFRLKVLTVLLYICLSVMLFYLFGSIMPLPPWGLVVGWVMALIAVELAYAFIFPYSFRYIADNDDDKMVILPV |
| IRIS_313-11334 | MLHHLKELAAVAGIHMILIYLCRFLLRRSRNVLFTVSNSLRFRLKVLTVLLYICLSVMLFYLFGSIMPLPPWGLVVGWVMALIAVELAYAFIFPYSFRYIADNDDDKMVILPV |
| IRIS_313-11335 | MLHHLKELAAVAGIHMILIYLCRFLLRRSRNVLFTVSNSLRFRLKVLTVLLYICLSVMLFYLFGSIMPLPPWGLVVGWVMALIAVELAYAFIFPYSFRYIADNDDDKMVILPV |
| IRIS_313-11336 | MLHHLKELAAVAGIHMILIYLCRFLLRRSRNVLFTVSNSLRFRLKVLTVLLYICLSVMLFYLFGSIMPLPPWGLVVGWVMALIAVELAYAFIFPYSFRYIADNDDDKMVILPV |

| | | |
|---|---|---|
| 5312 | IRIS_313-11337 | MLHHLKELAAVAGIHMILIYLCRFLLRRSRNVLFTVSNSLRFRLKVLTVLLYICLSVMLFYLFGSIMPLPPWGLVVGWVMALIAVELAYAFIFPYSFRYIADNDDDKMVILPV |
| 5313 | IRIS_313-11338 | MLHHLKELAAVAGIHMILIYLCRFLLRRSRNVLFTVSNSLRFRLKVLTVLLYICLSVMLFYLFGSIMPLPPWGLVVGWVMALIAVELAYAFIFPYSFRYIADNDDDKMVILPV |
| 5314 | IRIS_313-11339 | MLHHLKELAAVAGIHMILIYLCRFLLRRSRNVLFTVSNSLRFRLKVLTVLLYICLSVMLFYLFGSIMPLPPWGLVVGWVMALIAVELAYAFIFPYSFRYIADNDDDKMVILPV |
| 5315 | IRIS_313-11340 | MLHQLKELAAVAGIHMILIYLCRFLLRRSRNVLFTVSNSLRFRLKVLTVLLYICLLVMLFYLFGSIMPLPPWGLVVGWVMALIAVELAYAFIFPYSFRYIADNDDDKMVILPV |
| 5316 | IRIS_313-11341 | MLHHLKELAAVAGIHMILIYLCRFLLRRSRNVLFTVSNSLRFRLKVLTVLLYICLSVMLFYLFGSIMPLPPWGLVVGWVMALIAVELAYAFIFPYSFRYIADNDDDKMVILPV |
| 5317 | IRIS_313-11342 | MLHHLKELAAVAGIHMILIYLCRFLLRRSRNVLFTVSNSLRFRLKVLTVLLYICLSVMLFYLFGSIMPLPPWGLVVGWVMALIAVELAYAFIFPYSFRYIADNDDDKMVILPV |
| 5318 | IRIS_313-11343 | MLHHLKELAAVAGIHMILIYLCRFLLRRSRNVLFTVSNSLRFRLKVLTVLLYICLSVMLFYLFGSIMPLPPWGLVVGWVMALIAVELAYAFIFPYSFRYIADNDDDKMVILPV |
| 5319 | IRIS_313-11344 | MLHHLKELAAVAGIHMILIYLCRFLLRRSRNVLFTVSNSLRFRLKVLTVLLYICLSVMLFYLFGSIMPLPPWGLVVGWVMALIAVELAYAFIFPYSFRYIADNDDDKMVILPV |
| 5320 | IRIS_313-11345 | MLHHLKELAAVAGIHMILIYLCRFLLRRSRNVLFTVSNSLRFRLKVLTVLLYICLSVMLFYLFGSIMPLPPWGLVVGWVMALIAVELAYAFIFPYSFRYIADNDDDKMVILPV |
| 5321 | IRIS_313-11346 | MLHHLKELAAVAGIHMILIYLCRFLLRRSRNVLFTVSNSLRFRLKVLTVLLYICLSVMLFYLFGSIMPLPPWGLVVGWVMALIAVELAYAFIFPYSFRYIADNDDDKMVILPV |
| 5322 | IRIS_313-11347 | MLHHLKELAAVAGIHMILIYLCRFLLRRSRNVLFTVSNSLRFRLKVLTVLLYICLSVMLFYLFGSIMPLPPWGLVVGWVMALIAVELAYAFIFPYSFRYIADNDDDKMVILPV |
| 5323 | IRIS_313-11348 | MLHHLKELAAVAGIHMILIYLCRFLLRRSRNVLFTVSNSLRFRLKVLTVLLYICLSVMLFYLFGSIMPLPPWGLVVGWVMALIAVELAYAFIFPYSFRYIADNDDDKMVILPV |
| 5324 | IRIS_313-11349 | MLHHLKELAAVAGIHMILIYLCRFLLRRSRNVLFTVSNSLRFRLKVLTVLLYICLSVMLFYLFGSIMPLPPWGLVVGWVMALIAVELAYAFIFPYSFRYIADNDDDKMVILPV |
| 5325 | IRIS_313-11350 | MLHHLKELAAVAGIHMILIYLCRFLLRRSRNVLFTVSNSLRFRLKVLTVLLYICLSVMLFYLFGSIMPLPPWGLVVGWVMALIAVELAYAFIFPYSFRYIADNDDDKMVILPV |
| 5326 | IRIS_313-11351 | MLHHLKELAAVAGIHMILIYLCRFLLRRSRNVLFTVSNSLRFRLKVLTVLLYICLSVMLFYLFGSIMPLPPWGLVVGWVMALIAVELAYAFIFPYSFRYIADNDDDKMVILPV |
| 5327 | IRIS_313-11352 | MLHHLKELAAVAGIHMILIYLCRFLLRRSRNVLFTVSNSLRFRLKVLTVLLYICLSVMLFYLFGSIMPLPPWGLVVGWVMALIAVELAYAFIFPYSFRYIADNDDDKMVILPV |
| 5328 | IRIS_313-11353 | MLHHLKELAAVAGIHMILIYLCRFLLRRSRNVLFTVSNSLRFRLKVLTVLLYICLSVMLFYLFGSIMPLPPWGLVVGWVMALIAVELAYAFIFPYSFRYIADNDDDKMVILPV |
| 5329 | IRIS_313-11354 | MLHHLKELAAVAGIHMILIYLCRFLLRRSRNVLFTVSNSLRFRLKVLTVLLYICLSVMLFYLFGSIMPLPPWGLVVGWVMALIAVELAYAFIFPYSFRYIADNDDDKMVILPV |
| 5330 | IRIS_313-11355 | MLHHLKELAAVAGIHMILIYLCRFLLRRSRNVLFTVSNSLRFRLKVLTVLLYICLSVMLFYLFGSIMPLPPWGLVVGWVMALIAVELAYAFIFPYSFRYIADNDDDKMVILPV |
| 5331 | IRIS_313-11356 | MLHHLKELAAVAGIHMILIYLCRFLLRRSRNVLFTVSNSLRFRLKVLTVLLYICLSVMLFYLFGSIMPLPPWGLVVGWVMALIAVELAYAFIFPYSFRYIADNDDDKMVILPV |
| 5332 | IRIS_313-11357 | MLHHLKELAAVAGIHMILIYLCRFLLRRSRNVLFTVSNSLRFRLKVLTVLLYICLSVMLFYLFGSIMPLPPWGLVVGWVMALIAVELAYAFIFPYSFRYIADNDDDKMVILPV |
| 5333 | IRIS_313-11358 | MLHHLKELAAVAGIHMILIYLCRFLLRRSRNVLFTVSNSLRFRLKVLTVLLYICLSVMLFYLFGSIMPLPPWGLVVGWVMALIAVELAYAFIFPYSFRYIADNDDDKMVILPV |
| 5334 | IRIS_313-11359 | MLHHLKELAAVAGIHMILIYLCRFLLRRSRNVLFTVSNSLRFRLKVLTVLLYICLSVMLFYLFGSIMPLPPWGLVVGWVMALIAVELAYAFIFPYSFRYIADNDDDKMVILPV |
| 5335 | IRIS_313-11360 | MLHHLKELAAVAGIHMILIYLCRFLLRRSRNVLFTVSNSLRFRLKVLTVLLYICLSVMLFYLFGSIMPLPPWGLVVGWVMALIAVELAYAFIFPYSFRYIADNDDDKMVILPV |
| 5336 | IRIS_313-11361 | MLHHLKELAAVAGIHMILIYLCRFLLRRSRNVLFTVSNSLRFRLKVLTVLLYICLSVMLFYLFGSIMPLPPWGLVVGWVMALIAVELAYAFIFPYSFRYIADNDDDKMVILPV |
| 5337 | IRIS_313-11362 | MLHHLKELAAVAGIHMILIYLCRFLLRRSRNVLFTVSNSLRFRLKVLTVLLYICLSVMLFYLFGSIMPLPPWGLVVGWVMALIAVELAYAFIFPYSFRYIADNDDDKMVILPV |
| 5338 | IRIS_313-11363 | MLHHLKELAAVAGIHMILIYLCRFLLRRSRNVLFTVSNSLRFRLKVLTVLLYICLSVMLFYLFGSIMPLPPWGLVVGWVMALIAVELAYAFIFPYSFRYIADNDDDKMVILPV |
| 5339 | IRIS_313-11364 | MLHHLKELAAVAGIHMILIYLCRFLLRRSRNVLFTVSNSLRFRLKVLTVLLYICLSVMLFYLFGSIMPLPPWGLVVGWVMALIAVELAYAFIFPYSFRYIADNDDDKMVILPV |
| 5340 | IRIS_313-11365 | MLHHLKELAAVAGIHMILIYLCRFLLRRSRNVLFTVSNSLRFRLKVLTVLLYICLSVMLFYLFGSIMPLPPWGLVVGWVMALIAVELAYAFIFPYSFRYIADNDDDKMVILPV |
| 5341 | IRIS_313-11367 | MLHHLKELAAVAGIHMILIYLCRFLLRRSRNVLFTVSNSLRFRLKVLTVLLYICLSVMLFYLFGSIMPLPPWGLVVGWVMALIAVELAYAFIFPYSFRYIADNDDDKMVILPV |
| 5342 | IRIS_313-11368 | MLHHLKELAAVAGIHMILIYLCRFLLRRSRNVLFTVSNSLRFRLKVLTVLLYICLSVMLFYLFGSIMPLPPWGLVVGWVMALIAVELAYAFIFPYSFRYIADNDDDKMVILPV |
| 5343 | IRIS_313-11369 | MLHHLKELAAVAGIHMILIYLCRFLLRRSRNVLFTVSNSLRFRLKVLTVLLYICLSVMLFYLFGSIMPLPPWGLVVGWVMALIAVELAYAFIFPYSFRYIADNDDDKMVILPV |
| 5344 | IRIS_313-11370 | MLHHLKELAAVAGIHMILIYLCRFLLRRSRNVLFTVSNSLRFRLKVLTVLLYICLSVMLFYLFGSIMPLPPWGLVVGWVMALIAVELAYAFIFPYSFRYIADNDDDKMVILPV |
| 5345 | IRIS_313-11371 | MLHHLKELAAVAGIHMILIYLCRFLLRRSRNVLFTVSNSLRFRLKVLTVLLYICLSVMLFYLFGSIMPLPPWGLVVGWVMALIAVELAYAFIFPYSFRYIADNDDDKMVILPV |
| 5346 | IRIS_313-11372 | MLHHLKELAAVAGIHMILIYLCRFLLRRSRNVLFTVSNSLRFRLKVLTVLLYICLSVMLFYLFGSIMPLPPWGLVVGWVMALIAVELAYAFIFPYSFRYIADNDDDKMVILPV |
| 5347 | IRIS_313-11373 | MLHHLKELAAVAGIHMILIYLCRFLLRRSRNVLFTVSNSLRFRLKVLTVLLYICLSVMLFYLFGSIMPLPPWGLVVGWVMALIAVELAYAFIFPYSFRYIADNDDDKMVILPV |
| 5348 | IRIS_313-11374 | MLHHLKELAAVAGIHMILIYLCRFLLRRSRNVLFTVSNSLRFRLKVLTVLLYICLSVMLFYLFGSIMPLPPWGLVVGWVMALIAVELAYAFIFPYSFRYIADNDDDKMVILPV |
| 5349 | IRIS_313-11375 | MLHHLKELAAVAGIHMILIYLCRFLLRRSRNVLFTVSNSLRFRLKVLTVLLYICLSVMLFYLFGSIMPLPPWGLVVGWVMALIAVELAYAFIFPYSFRYIADNDDDKMVILPV |
| 5350 | IRIS_313-11376 | MLHHLKELAAVAGIHMILIYLCRFLLRRSRNVLFTVSNSLRFRLKVLTVLLYICLSVMLFYLFGSIMPLPPWGLVVGWVMALIAVELAYAFIFPYSFRYIADNDDDKMVILPV |
| 5351 | IRIS_313-11377 | MLHHLKELAAVAGIHMILIYLCRFLLRRSRNVLFTVSNSLRFRLKVLTVLLYICLSVMLFYLFGSIMPLPPWGLVVGWVMALIAVELAYAFIFPYSFRYIADNDDDKMVILPV |
| 5352 | IRIS_313-11379 | MLHHLKELAAVAGIHMILIYLCRFLLRRSRNVLFTVSNSLRFRLKVLTVLLYICLSVMLFYLFGSIMPLPPWGLVVGWVMALIAVELAYAFIFPYSFRYIADNDDDKMVILPV |
| 5353 | IRIS_313-11380 | MLHQLKELAAVAGIHMILIYLCRFLLRRSRNVLFTVSNSLRFRLKVLTVLLYICLLVMLFYLFGSIMPLPPWGLVVGWVMALIAVELAYAFIFPYSFRYIADNDDDKMVILPV |
| 5354 | IRIS_313-11381 | MLHHLKELAAVAGIHMILIYLCRFLLRRSRNVLFTVSNSLRFRLKVLTVLLYICLSVMLFYLFGSIMPLPPWGLVVGWVMALIAVELAYAFIFPYSFRYIADNDDDKMVILPV |
| 5355 | IRIS_313-11382 | MLHHLKELAAVAGIHMILIYLCRFLLRRSRNVLFTVSNSLRFRLKVLTVLLYICLSVMLFYLFGSIMPLPPWGLVVGWVMALIAVELAYAFIFPYSFRYIADNDDDKMVILPV |
| 5356 | IRIS_313-11383 | MLHHLKELAAVAGIHMILIYLCRFLLRRSRNVLFTVSNSLRFRLKVLTVLLYICLSVMLFYLFGSIMPLPPWGLVVGWVMALIAVELAYAFIFPYSFRYIADNDDDKMVILPV |
| 5357 | IRIS_313-11384 | MLHHLKELAAVAGIHMILIYLCRFLLRRSRNVLFTVSNSLRFRLKVLTVLLYICLSVMLFYLFGSIMPLPPWGLVVGWVMALIAVELAYAFIFPYSFRYIADNDDDKMVILPV |
| 5358 | IRIS_313-11385 | MLHHLKELAAVAGIHMILIYLCRFLLRRSRNVLFTVSNSLRFRLKVLTVLLYICLSVMLFYLFGSIMPLPPWGLVVGWVMALIAVELAYAFIFPYSFRYIADNDDDKMVILPV |
| 5359 | IRIS_313-11386 | MLHHLKELAAVAGIHMILIYLCRFLLRRSRNVLFTVSNSLRFRLKVLTVLLYICLSVMLFYLFGSIMPLPPWGLVVGWVMALIAVELAYAFIFPYSFRYIADNDDDKMVILPV |
| 5360 | IRIS_313-11387 | MLHHLKELAAVAGIHMILIYLCRFLLRRSRNVLFTVSNSLRFRLKVLTVLLYICLSVMLFYLFGSIMPLPPWGLVVGWVMALIAVELAYAFIFPYSFRYIADNDDDKMVILPV |
| 5361 | IRIS_313-11388 | MLHHLKELAAVAGIHMILIYLCRFLLRRSRNVLFTVSNSLRFRLKVLTVLLYICLSVMLFYLFGSIMPLPPWGLVVGWVMALIAVELAYAFIFPYSFRYIADNDDDKMVILPV |
| 5362 | IRIS_313-11389 | MLHHLKELAAVAGIHMILIYLCRFLLRRSRNVLFTVSNSLRFRLKVLTVLLYICLSVMLFYLFGSIMPLPPWGLVVGWVMALIAVELAYAFIFPYSFRYIADNDDDKMVILPV |
| 5363 | IRIS_313-11390 | MLHHLKELAAVAGIHMILIYLCRFLLRRSRNVLFTVSNSLRFRLKVLTVLLYICLSVMLFYLFGSIMPLPPWGLVVGWVMALIAVELAYAFIFPYSFRYIADNDDDKMVILPV |
| 5364 | IRIS_313-11391 | MLHHLKELAAVAGIHMILIYLCRFLLRRSRNVLFTVSNSLRFRLKVLTVLLYICLSVMLFYLFGSIMPLPPWGLVVGWVMALIAVELAYAFIFPYSFRYIADNDDDKMVILPV |
| 5365 | IRIS_313-11392 | MLHHLKELAAVAGIHMILIYLCRFLLRRSRNVLFTVSNSLRFRLKVLTVLLYICLSVMLFYLFGSIMPLPPWGLVVGWVMALIAVELAYAFIFPYSFRYIADNDDDKMVILPV |
| 5366 | IRIS_313-11393 | MLHHLKELAAVAGIHMILIYLCRFLLRRSRNVLFTVSNSLRFRLKVLTVLLYICLSVMLFYLFGSIMPLPPWGLVVGWVMALIAVELAYAFIFPYSFRYIADNDDDKMVILPV |
| 5367 | IRIS_313-11394 | MLHHLKELAAVAGIHMILIYLCRFLLRRSRNVLFTVSNSLRFRLKVLTVLLYICLSVMLFYLFGSIMPLPPWGLVVGWVMALIAVELAYAFIFPYSFRYIADNDDDKMVILPV |
| 5368 | IRIS_313-11395 | MLHHLKELAAVAGIHMILIYLCRFLLRRSRNVLFTVSNSLRFRLKVLTVLLYICLSVMLFYLFGSIMPLPPWGLVVGWVMALIAVELAYAFIFPYSFRYIADNDDDKMVILPV |
| 5369 | IRIS_313-11396 | MLHHLKELAAVAGIHMILIYLCRFLLRRSRNVLFTVSNSLRFRLKVLTVLLYICLSVMLFYLFGSIMPLPPWGLVVGWVMALIAVELAYAFIFPYSFRYIADNDDDKMVILPV |
| 5370 | IRIS_313-11397 | MLHHLKELAAVAGIHMILIYLCRFLLRRSRNVLFTVSNSLRFRLKVLTVLLYICLSVMLFYLFGSIMPLPPWGLVVGWVMALIAVELAYAFIFPYSFRYIADNDDDKMVILPV |
| 5371 | IRIS_313-11398 | MLHHLKELAAVAGIHMILIYLCRFLLRRSRNVLFTVSNSLRFRLKVLTVLLYICLSVMLFYLFGSIMPLPPWGLVVGWVMALIAVELAYAFIFPYSFRYIADNDDDKMVILPV |
| 5372 | IRIS_313-11399 | MLHHLKELAAVAGIHMILIYLCRFLLRRSRNVLFTVSNSLRFRLKVLTVLLYICLSVMLFYLFGSIMPLPPWGLVVGWVMALIAVELAYAFIFPYSFRYIADNDDDKMVILPV |
| 5373 | IRIS_313-11400 | MLHHLKELAAVAGIHMILIYLCRFLLRRSRNVLFTVSNSLRFRLKVLTVLLYICLSVMLFYLFGSIMPLPPWGLVVGWVMALIAVELAYAFIFPYSFRYIADNDDDKMVILPV |
| 5374 | IRIS_313-11401 | MLHHLKELAAVAGIHMILIYLCRFLLRRSRNVLFTVSNSLRFRLKVLTVLLYICLSVMLFYLFGSIMPLPPWGLVVGWVMALIAVELAYAFIFPYSFRYIADNDDDKMVILPV |
| 5375 | IRIS_313-11402 | MLHHLKELAAVAGIHMILIYLCRFLLRRSRNVLFTVSNSLRFRLKVLTVLLYICLSVMLFYLFGSIMPLPPWGLVVGWVMALIAVELAYAFIFPYSFRYIADNDDDKMVILPV |
| 5376 | IRIS_313-11403 | MLHHLKELAAVAGIHMILIYLCRFLLRRSRNVLFTVSNSLRFRLKVLTVLLYICLSVMLFYLFGSIMPLPPWGLVVGWVMALIAVELAYAFIFPYSFRYIADNDDDKMVILPV |
| 5377 | IRIS_313-11404 | MLHHLKELAAVAGIHMILIYLCRFLLRRSRNVLFTVSNSLRFRLKVLTVLLYICLSVMLFYLFGSIMPLPPWGLVVGWVMALIAVELAYAFIFPYSFRYIADNDDDKMVILPV |
| 5378 | IRIS_313-11405 | MLHHLKELAAVAGIHMILIYLCRFLLRRSRNVLFTVSNSLRFRLKVLTVLLYICLSVMLFYLFGSIMPLPPWGLVVGWVMALIAVELAYAFIFPYSFRYIADNDDDKMVILPV |
| 5379 | IRIS_313-11406 | MLHHLKELAAVAGIHMILIYLCRFLLRRSRNVLFTVSNSLRFRLKVLTVLLYICLSVMLFYLFGSIMPLPPWGLVVGWVMALIAVELAYAFIFPYSFRYIADNDDDKMVILPV |
| 5380 | IRIS_313-11407 | MLHHLKELAAVAGIHMILIYLCRFLLRRSRNVLFTVSNSLRFRLKVLTVLLYICLSVMLFYLFGSIMPLPPWGLVVGWVMALIAVELAYAFIFPYSFRYIADNDDDKMVILPV |
| 5381 | IRIS_313-11408 | MLHHLKELAAVAGIHMILIYLCRFLLRRSRNVLFTVSNSLRFRLKVLTVLLYICLSVMLFYLFGSIMPLPPWGLVVGWVMALIAVELAYAFIFPYSFRYIADNDDDKMVILPV |
| 5382 | IRIS_313-11409 | MLHHLKELAAVAGIHMILIYLCRFLLRRSRNVLFTVSNSLRFRLKVLTVLLYICLSVMLFYLFGSIMPLPPWGLVVGWVMALIAVELAYAFIFPYSFRYIADNDDDKMVILPV |
| 5383 | IRIS_313-11410 | MLHHLKELAAVAGIHMILIYLCRFLLRRSRNVLFTVSNSLRFRLKVLTVLLYICLSVMLFYLFGSIMPLPPWGLVVGWVMALIAVELAYAFIFPYSFRYIADNDDDKMVILPV |
| 5384 | IRIS_313-11411 | MLHHLKELAAVAGIHMILIYLCRFLLRRSRNVLFTVSNSLRFRLKVLTVLLYICLSVMLFYLFGSIMPLPPWGLVVGWVMALIAVELAYAFIFPYSFRYIADNDDDKMVILPV |
| 5385 | IRIS_313-11413 | MLHHLKELAAVAGIHMILIYLCRFLLRRSRNVLFTVSNSLRFRLKVLTVLLYICLSVMLFYLFGSIMPLPPWGLVVGWVMALIAVELAYAFIFPYSFRYIADNDDDKMVILPV |
| 5386 | IRIS_313-11414 | MLHHLKELAAVAGIHMILIYLCRFLLRRSRNVLFTVSNSLRFRLKVLTVLLYICLSVMLFYLFGSIMPLPPWGLVVGWVMALIAVELAYAFIFPYSFRYIADNDDDKMVILPV |
| 5387 | IRIS_313-11416 | MLHHLKELAAVAGIHMILIYLCRFLLRRSRNVLFTVSNSLRFRLKVLTVLLYICLSVMLFYLFGSIMPLPPWGLVVGWVMALIAVELAYAFIFPYSFRYIADNDDDKMVILPV |
| 5388 | IRIS_313-11417 | MLHHLKELAAVAGIHMILIYLCRFLLRRSRNVLFTVSNSLRFRLKVLTVLLYICLSVMLFYLFGSIMPLPPWGLVVGWVMALIAVELAYAFIFPYSFRYIADNDDDKMVILPV |
| 5389 | IRIS_313-11418 | MLHHLKELAAVAGIHMILIYLCRFLLRRSRNVLFTVSNSLRFRLKVLTVLLYICLSVMLFYLFGSIMPLPPWGLVVGWVMALIAVELAYAFIFPYSFRYIADNDDDKMVILPV |
| 5390 | IRIS_313-11419 | MLHHLKELAAVAGIHMILIYLCRFLLRRSRNVLFTVSNSLRFRLKVLTVLLYICLSVMLFYLFGSIMPLPPWGLVVGWVMALIAVELAYAFIFPYSFRYIADNDDDKMVILPV |
| 5391 | IRIS_313-11420 | MLHHLKELAAVAGIHMILIYLCRFLLRRSRNVLFTVSNSLRFRLKVLTVLLYICLSVMLFYLFGSIMPLPPWGLVVGWVMALIAVELAYAFIFPYSFRYIADNDDDKMVILPV |
| 5392 | IRIS_313-11421 | MLHHLKELAAVAGIHMILIYLCRFLLRRSRNVLFTVSNSLRFRLKVLTVLLYICLSVMLFYLFGSIMPLPPWGLVVGWVMALIAVELAYAFIFPYSFRYIADNDDDKMVILPV |
| 5393 | IRIS_313-11422 | MLHHLKELAAVAGIHMILIYLCRFLLRRSRNVLFTVSNSLRFRLKVLTVLLYICLSVMLFYLFGSIMPLPPWGLVVGWVMALIAVELAYAFIFPYSFRYIADNDDDKMVILPV |
| 5394 | IRIS_313-11423 | MLHHLKELAAVAGIHMILIYLCRFLLRRSRNVLFTVSNSLRFRLKVLTVLLYICLSVMLFYLFGSIMPLPPWGLVVGWVMALIAVELAYAFIFPYSFRYIADNDDDKMVILPV |
| 5395 | IRIS_313-11424 | MLHHLKELAAVAGIHMILIYLCRFLLRRSRNVLFTVSNSLRFRLKVLTVLLYICLSVMLFYLFGSIMPLPPWGLVVGWVMALIAVELAYAFIFPYSFRYIADNDDDKMVILPV |
| 5396 | IRIS_313-11425 | MLHHLKELAAVAGIHMILIYLCRFLLRRSRNVLFTVSNSLRFRLKVLTVLLYICLSVMLFYLFGSIMPLPPWGLVVGWVMALIAVELAYAFIFPYSFRYIADNDDDKMVILPV |
| 5397 | IRIS_313-11426 | MLHQLKELAAVAGIHMILIYLCRFLLRRSRNVLFTVSNSLRFRLKVLTVLLYICLLVMLFYLFGSIMPLPPWGLVVGWVMALIAVELAYAFIFPYSFRYIADNDDDKMVILPV |
| 5398 | IRIS_313-11427 | MLHHLKELAAVAGIHMILIYLCRFLLRRSRNVLFTVSNSLRFRLKVLTVLLYICLSVMLFYLFGSIMPLPPWGLVVGWVMALIAVELAYAFIFPYSFRYIADNDDDKMVILPV |
| 5399 | IRIS_313-11428 | MLHHLKELAAVAGIHMILIYLCRFLLRRSRNVLFTVSNSLRFRLKVLTVLLYICLSVMLFYLFGSIMPLPPWGLVVGWVMALIAVELAYAFIFPYSFRYIADNDDDKMVILPV |
| 5400 | IRIS_313-11429 | MLHQLKELAAVAGIHMILIYLCRFLLRRSRNVLFTVSNSLRFRLKVLTVLLYICLLVMLFYLFGSIMPLPPWGLVVGWVMALIAVELAYAFIFPYSFRYIADNDDDKMVILPV |
| 5401 | IRIS_313-11431 | MLHHLKELAAVAGIHMILIYLCRFLLRRSRNVLFTVSNSLRFRLKVLTVLLYICLSVMLFYLFGSIMPLPPWGLVVGWVMALIAVELAYAFIFPYSFRYIADNDDDKMVILPV |
| 5402 | IRIS_313-11432 | MLHHLKELAAVAGIHMILIYLCRFLLRRSRNVLFTVSNSLRFRLKVLTVLLYICLSVMLFYLFGSIMPLPPWGLVVGWVMALIAVELAYAFIFPYSFRYIADNDDDKMVILPV |
| 5403 | IRIS_313-11433 | MLHHLKELAAVAGIHMILIYLCRFLLRRSRNVLFTVSNSLRFRLKVLTVLLYICLSVMLFYLFGSIMPLPPWGLVVGWVMALIAVELAYAFIFPYSFRYIADNDDDKMVILPV |
| 5404 | IRIS_313-11434 | MLHHLKELAAVAGIHMILIYLCRFLLRRSRNVLFTVSNSLRFRLKVLTVLLYICLSVMLFYLFGSIMPLPPWGLVVGWVMALIAVELAYAFIFPYSFRYIADNDDDKMVILPV |
| 5405 | IRIS_313-11435 | MLHHLKELAAVAGIHMILIYLCRFLLRRSRNVLFTVSNSLRFRLKVLTVLLYICLSVMLFYLFGSIMPLPPWGLVVGWVMALIAVELAYAFIFPYSFRYIADNDDDKMVILPV |
| 5406 | IRIS_313-11436 | MLHHLKELAAVAGIHMILIYLCRFLLRRSRNVLFTVSNSLRFRLKVLTVLLYICLSVMLFYLFGSIMPLPPWGLVVGWVMALIAVELAYAFIFPYSFRYIADNDDDKMVILPV |
| 5407 | IRIS_313-11437 | MLHHLKELAAVAGIHMILIYLCRFLLRRSRNVLFTVSNSLRFRLKVLTVLLYICLSVMLFYLFGSIMPLPPWGLVVGWVMALIAVELAYAFIFPYSFRYIADNDDDKMVILPV |
| 5408 | IRIS_313-11438 | MLHHLKELAAVAGIHMILIYLCRFLLRRSRNVLFTVSNSLRFRLKVLTVLLYICLSVMLFYLFGSIMPLPPWGLVVGWVMALIAVELAYAFIFPYSFRYIADNDDDKMVILPV |
| 5409 | IRIS_313-11439 | MLHHLKELAAVAGIHMILIYLCRFLLRRSRNVLFTVSNSLRFRLKVLTVLLYICLSVMLFYLFGSIMPLPPWGLVVGWVMALIAVELAYAFIFPYSFRYIADNDDDKMVILPV |
| 5410 | IRIS_313-11441 | MLHHLKELAAVAGIHMILIYLCRFLLRRSRNVLFTVSNSLRFRLKVLTVLLYICLSVMLFYLFGSIMPLPPWGLVVGWVMALIAVELAYAFIFPYSFRYIADNDDDKMVILPV |
| 5411 | IRIS_313-11442 | MLHHLKELAAVAGIHMILIYLCRFLLRRSRNVLFTVSNSLRFRLKVLTVLLYICLSVMLFYLFGSIMPLPPWGLVVGWVMALIAVELAYAFIFPYSFRYIADNDDDKMVILPV |
| 5412 | IRIS_313-11443 | MLHHLKELAAVAGIHMILIYLCRFLLRRSRNVLFTVSNSLRFRLKVLTVLLYICLSVMLFYLFGSIMPLPPWGLVVGWVMALIAVELAYAFIFPYSFRYIADNDDDKMVILPV |
| 5413 | IRIS_313-11445 | MLHHLKELAAVAGIHMILIYLCRFLLRRSRNVLFTVSNSLRFRLKVLTVLLYICLSVMLFYLFGSIMPLPPWGLVVGWVMALIAVELAYAFIFPYSFRYIADNDDDKMVILPV |
| 5414 | IRIS_313-11446 | MLHHLKELAAVAGIHMILIYLCRFLLRRSRNVLFTVSNSLRFRLKVLTVLLYICLSVMLFYLFGSIMPLPPWGLVVGWVMALIAVELAYAFIFPYSFRYIADNDDDKMVILPV |
| 5415 | IRIS_313-11447 | MLHHLKELAAVAGIHMILIYLCRFLLRRSRNVLFTVSNSLRFRLKVLTVLLYICLSVMLFYLFGSIMPLPPWGLVVGWVMALIAVELAYAFIFPYSFRYIADNDDDKMVILPV |
| 5416 | IRIS_313-11448 | MLHHLKELAAVAGIHMILIYLCRFLLRRSRNVLFTVSNSLRFRLKVLTVLLYICLSVMLFYLFGSIMPLPPWGLVVGWVMALIAVELAYAFIFPYSFRYIADNDDDKMVILPV |
| 5417 | IRIS_313-11449 | MLHHLKELAAVAGIHMILIYLCRFLLRRSRNVLFTVSNSLRFRLKVLTVLLYICLSVMLFYLFGSIMPLPPWGLVVGWVMALIAVELAYAFIFPYSFRYIADNDDDKMVILPV |
| 5418 | IRIS_313-11451 | MLHHLKELAAVAGIHMILIYLCRFLLRRSRNVLFTVSNSLRFRLKVLTVLLYICLSVMLFYLFGSIMPLPPWGLVVGWVMALIAVELAYAFIFPYSFRYIADNDDDKMVILPV |
| 5419 | IRIS_313-11452 | MLHHLKELAAVAGIHMILIYLCRFLLRRSRNVLFTVSNSLRFRLKVLTVLLYICLSVMLFYLFGSIMPLPPWGLVVGWVMALIAVELAYAFIFPYSFRYIADNDDDKMVILPV |
| 5420 | IRIS_313-11453 | MLHHLKELAAVAGIHMILIYLCRFLLRRSRNVLFTVSNSLRFRLKVLTVLLYICLSVMLFYLFGSIMPLPPWGLVVGWVMALIAVELAYAFIFPYSFRYIADNDDDKMVILPV |
| 5421 | IRIS_313-11454 | MLHHLKELAAVAGIHMILIYLCRFLLRRSRNVLFTVSNSLRFRLKVLTVLLYICLSVMLFYLFGSIMPLPPWGLVVGWVMALIAVELAYAFIFPYSFRYIADNDDDKMVILPV |
| 5422 | IRIS_313-11455 | MLHHLKELAAVAGIHMILIYLCRFLLRRSRNVLFTVSNSLRFRLKVLTVLLYICLSVMLFYLFGSIMPLPPWGLVVGWVMALIAVELAYAFIFPYSFRYIADNDDDKMVILPV |
| 5423 | IRIS_313-11456 | MLHHLKELAAVAGIHMILIYLCRFLLRRSRNVLFTVSNSLRFRLKVLTVLLYICLSVMLFYLFGSIMPLPPWGLVVGWVMALIAVELAYAFIFPYSFRYIADNDDDKMVILPV |
| 5424 | IRIS_313-11458 | MLHHLKELAAVAGIHMILIYLCRFLLRRSRNVLFTVSNSLRFRLKVLTVLLYICLSVMLFYLFGSIMPLPPWGLVVGWVMALIAVELAYAFIFPYSFRYIADNDDDKMVILPV |
| 5425 | IRIS_313-11460 | MLHHLKELAAVAGIHMILIYLCRFLLRRSRNVLFTVSNSLRFRLKVLTVLLYICLSVMLFYLFGSIMPLPPWGLVVGWVMALIAVELAYAFIFPYSFRYIADNDDDKMVILPV |
| 5426 | IRIS_313-11461 | MLHHLKELAAVAGIHMILIYLCRFLLRRSRNVLFTVSNSLRFRLKVLTVLLYICLSVMLFYLFGSIMPLPPWGLVVGWVMALIAVELAYAFIFPYSFRYIADNDDDKMVILPV |
| 5427 | IRIS_313-11462 | MLHHLKELAAVAGIHMILIYLCRFLLRRSRNVLFTVSNSLRFRLKVLTVLLYICLSVMLFYLFGSIMPLPPWGLVVGWVMALIAVELAYAFIFPYSFRYIADNDDDKMVILPV |
| 5428 | IRIS_313-11463 | MLHHLKELAAVAGIHMILIYLCRFLLRRSRNVLFTVSNSLRFRLKVLTVLLYICLSVMLFYLFGSIMPLPPWGLVVGWVMALIAVELAYAFIFPYSFRYIADNDDDKMVILPV |
| 5429 | IRIS_313-11464 | MLHHLKELAAVAGIHMILIYLCRFLLRRSRNVLFTVSNSLRFRLKVLTVLLYICLSVMLFYLFGSIMPLPPWGLVVGWVMALIAVELAYAFIFPYSFRYIADNDDDKMVILPV |
| 5430 | IRIS_313-11465 | MLHQLKELAAVAGIHMILIYLCRFLLRRSRNVLFTVSNSLRFRLKVLTVLLYICLLVMLFYLFGSIMPLPPWGLVVGWVMALIAVELAYAFIFPYSFRYIADNDDDKMVILPV |
| 5431 | IRIS_313-11467 | MLHHLKELAAVAGIHMILIYLCRFLLRRSRNVLFTVSNSLRFRLKVLTVLLYICLSVMLFYLFGSIMPLPPWGLVVGWVMALIAVELAYAFIFPYSFRYIADNDDDKMVILPV |
| 5432 | IRIS_313-11471 | MLHHLKELAAVAGIHMILIYLCRFLLRRSRNVLFTVSNSLRFRLKVLTVLLYICLSVMLFYLFGSIMPLPPWGLVVGWVMALIAVELAYAFIFPYSFRYIADNDDDKMVILPV |
| 5433 | IRIS_313-11472 | MLHHLKELAAVAGIHMILIYLCRFLLRRSRNVLFTVSNSLRFRLKVLTVLLYICLSVMLFYLFGSIMPLPPWGLVVGWVMALIAVELAYAFIFPYSFRYIADNDDDKMVILPV |
| 5434 | IRIS_313-11473 | MLHHLKELAAVAGIHMILIYLCRFLLRRSRNVLFTVSNSLRFRLKVLTVLLYICLSVMLFYLFGSIMPLPPWGLVVGWVMALIAVELAYAFIFPYSFRYIADNDDDKMVILPV |

| | |
|---|---|
| IRIS_313-11476 | MLHHLKELAAVAGIHMILIYLCRFLLRRSRNVLFTVSNSLRFRLKVLTVLLYICLSVMLFYLFGSIMPLPPWGLVVGWVMALIAVELAYAFIFPYSFRYIADNDDDKMVILPV |
| IRIS_313-11477 | MLHHLKELAAVAGIHMILIYLCRFLLRRSRNVLFTVSNSLRFRLKVLTVLLYICLSVMLFYLFGSIMPLPPWGLVVGWVMALIAVELAYAFIFPYSFRYIADNDDDKMVILPV |
| IRIS_313-11478 | MLHHLKELAAVAGIHMILIYLCRFLLRRSRNVLFTVSNSLRFRLKVLTVLLYICLSVMLFYLFGSIMPLPPWGLVVGWVMALIAVELAYAFIFPYSFRYIADNDDDKMVILPV |
| IRIS_313-11479 | MLHHLKELAAVAGIHMILIYLCRFLLRRSRNVLFTVSNSLRFRLKVLTVLLYICLSVMLFYLFGSIMPLPPWGLVVGWVMALIAVELAYAFIFPYSFRYIADNDDDKMVILPV |
| IRIS_313-11480 | MLHHLKELAAVAGIHMILIYLCRFLLRRSRNVLFTVSNSLRFRLKVLTVLLYICLSVMLFYLFGSIMPLPPWGLVVGWVMALIAVELAYAFIFPYSFRYIADNDDDKMVILPV |
| IRIS_313-11481 | MLHHLKELAAVAGIHMILIYLCRFLLRRSRNVLFTVSNSLRFRLKVLTVLLYICLSVMLFYLFGSIMPLPPWGLVVGWVMALIAVELAYAFIFPYSFRYIADNDDDKMVILPV |
| IRIS_313-11482 | MLHHLKELAAVAGIHMILIYLCRFLLRRSRNVLFTVSNSLRFRLKVLTVLLYICLSVMLFYLFGSIMPLPPWGLVVGWVMALIAVELAYAFIFPYSFRYIADNDDDKMVILPV |
| IRIS_313-11483 | MLHHLKELAAVAGIHMILIYLCRFLLRRSRNVLFTVSNSLRFRLKVLTVLLYISLSAMLLYLFGSIMPLPPWGLVVGWVMALIAVELAYAFIFPYSFRYIADNDDDKMVILPV |
| IRIS_313-11484 | MLHHLKELAAVAGIHMILIYLCRFLLRRSRNVLFTVSNSLRFRLKVLTVLLYICLSVMLFYLFGSIMPLPPWGLVVGWVMALIAVELAYAFIFPYSFRYIADNDDDKMVILPV |
| IRIS_313-11485 | MLHHLKELAAVAGIHMILIYLCRFLLRRSRNVLFTVSNSLRFRLKVLTVLLYICLSVMLFYLFGSIMPLPPWGLVVGWVMALIAVELAYAFIFPYSFRYIADNDDDKMVILPV |
| IRIS_313-11486 | MLHHLKELAAVAGIHMILIYLCRFLLRRSRNVLFTVSNSLRFRLKVLTVLLYICLSVMLFYLFGSIMPLPPWGLVVGWVMALIAVELAYAFIFPYSFRYIADNDDDKMVILPV |
| IRIS_313-11487 | MLHHLKELAAVAGIHMILIYLCRFLLRRSRNVLFTVSNSLRFRLKVLTVLLYICLSVMLFYLFGSIMPLPPWGLVVGWVMALIAVELAYAFIFPYSFRYIADNDDDKMVILPV |
| IRIS_313-11489 | MLHHLKELAAVAGIHMILIYLCRFLRRSRNVLFTVSNSLRFRLKVLTVLLYISLSAMLLYLFGSIMPLPPWGLVVGWVMALIAVELAYAFIFPYSFRYIADNDDDKMVILPV |
| IRIS_313-11490 | MLHHLKELAAVAGIHMILIYLCRFLLRRSRNVLFTVSNSLRFRLKVLTVLLYICLSVMLFYLFGSIMPLPPWGLVVGWVMALIAVELAYAFIFPYSFRYIADNDDDKMVILPV |
| IRIS_313-11491 | MLHHLKELAAVAGIHMILIYLCRFLLRRSRNVLFTVSNSLRFRLKVLTVLLYICLSVMLFYLFGSIMPLPPWGLVVGWVMALIAVELAYAFIFPYSFRYIADNDDDKMVILPV |
| IRIS_313-11492 | MLHHLKELAAVAGIHMILIYLCRFLLRRSRNVLFTVSNSLRFRLKVLTVLLYICLSVMLFYLFGSIMPLPPWGLVVGWVMALIAVELAYAFIFPYSFRYIADNDDDKMVILPV |
| IRIS_313-11493 | MLHHLKELAAVAGIHMILIYLCRFLLRRSRNVLFTVSNSLRFRLKVLTVLLYICLSVMLFYLFGSIMPLPPWGLVVGWVMALIAVELAYAFIFPYSFRYIADNDDDKMVILPV |
| IRIS_313-11494 | MLHHLKELAAVAGIHMILIYLCRFLLRRSRNVLFTVSNSLRFRLKVLTVLLYICLSVMLFYLFGSIMPLPPWGLVVGWVMALIAVELAYAFIFPYSFRYIADNDDDKMVILPV |
| IRIS_313-11495 | MLHHLKELAAVAGIHMILIYLCRFLLRRSRNVLFTVSNSLRFRLKVLTVLLYICLSVMLFYLFGSIMPLPPWGLVVGWVMALIAVELAYAFIFPYSFRYIADNDDDKMVILPV |
| IRIS_313-11496 | MLHHLKELAAVAGIHMILIYLCRFLLRRSRNVLFTVSNSLRFRLKVLTVLLYICLSVMLFYLFGSIMPLPPWGLVVGWVMALIAVELAYAFIFPYSFRYIADNDDDKMVILPV |
| IRIS_313-11497 | MLHHLKELAAVAGIHMILIYLCRFLLRRSRNVLFTVSNSLRFRLKVLTVLLYICLSVMLFYLFGSIMPLPPWGLVVGWVMALIAVELAYAFIFPYSFRYIADNDDDKMVILPV |
| IRIS_313-11498 | MLHHLKELAAVAGIHMILIYLCRFLLRRSRNVLFTVSNSLRFRLKVLTVLLYICLSVMLFYLFGSIMPLPPWGLVVGWVMALIAVELAYAFIFPYSFRYIADNDDDKMVILPV |
| IRIS_313-11499 | MLHHLKELAAVAGIHMILIYLCRFLLRRSRNVLFTVSNSLRFRLKVLTVLLYICLSVMLFYLFGSIMPLPPWGLVVGWVMALIAVELAYAFIFPYSFRYIADNDDDKMVILPV |
| IRIS_313-11500 | MLHHLKELAAVAGIHMILIYLCRFLLRRSRNVLFTVSNSLRFRLKVLTVLLYICLSVMLFYLFGSIMPLPPWGLVVGWVMALIAVELAYAFIFPYSFRYIADNDDDKMVILPV |
| IRIS_313-11505 | MLHHLKELAAVAGIHMILIYLCRFLLRRSRNVLFTVSNSLRFRLKVLTVLLYICLSVMLFYLFGSIMPLPPWGLVVGWVMALIAVELAYAFIFPYSFRYIADNDDDKMVILPV |
| IRIS_313-11506 | MLHHLKELAAVAGIHMILIYLCRFLLRRSRNVLFTVSNSLRFRLKVLTVLLYICLSVMLFYLFGSIMPLPPWGLVVGWVMALIAVELAYAFIFPYSFRYIADNDDDKMVILPV |
| IRIS_313-11507 | MLHHLKELAAVAGIHMILIYLCRFLLRRSRNVLFTVSNSLRFRLKVLTVLLYICLSVMLFYLFGSIMPLPPWGLVVGWVMALIAVELAYAFIFPYSFRYIADNDDDKMVILPV |
| IRIS_313-11508 | MLHHLKELAAVAGIHMILIYLCRFLLRRSRNVLFTVSNSLRFRLKVLTVLLYICLSVMLFYLFGSIMPLPPWGLVVGWVMALIAVELAYAFIFPYSFRYIADNDDDKMVILPV |
| IRIS_313-11509 | MLHHLKELAAVAGIHMILIYLCRFLLRRSRNVLFTVSNSLRFRLKVLTVLLYICLSVMLFYLFGSIMPLPPWGLVVGWVMALIAVELAYAFIFPYSFRYIADNDDDKMVILPV |
| IRIS_313-11510 | MLHHLKELAAVAGIHMILIYLCRFLLRRSRNVLFTVSNSLRFRLKVLTVLLYICLSVMLFYLFGSIMPLPPWGLVVGWVMALIAVELAYAFIFPYSFRYIADNDDDKMVILPV |
| IRIS_313-11511 | MLHHLKELAAVAGIHMILIYLCRFLLRRSRNVLFTVSNSLRFRLKVLTVLLYICLSVMLFYLFGSIMPLPPWGLVVGWVMALIAVELAYAFIFPYSFRYIADNDDDKMVILPV |
| IRIS_313-11512 | MLHHLKELAAVAGIHMILIYLCRFLLRRSRNVLFTVSNSLRFRLKVLTVLLYICLSVMLFYLFGSIMPLPPWGLVVGWVMALIAVELAYAFIFPYSFRYIADNDDDKMVILPV |
| IRIS_313-11513 | MLHHLKELAAVAGIHMILIYLCRFLLRRSRNVLFTVSNSLRFRLKVLTVLLYICLSVMLFYLFGSIMPLPPWGLVVGWVMALIAVELAYAFIFPYSFRYIADNDDDKMVILPV |
| IRIS_313-11515 | MLHHLKELAAVAGIHMILIYLCRFLLRRSRNVLFTVSNSLRFRLKVLTVLLYICLSVMLFYLFGSIMPLPPWGLVVGWVMALIAVELAYAFIFPYSFRYIADNDDDKMVILPV |
| IRIS_313-11516 | MLHHLKELAAVAGIHMILIYLCRFLLRRSRNVLFTVSNSLRFRLKVLTVLLYICLSVMLFYLFGSIMPLPPWGLVVGWVMALIAVELAYAFIFPYSFRYIADNDDDKMVILPV |
| IRIS_313-11517 | MLHHLKELAAVAGIHMILIYLCRFLLRRSRNVLFTVSNSLRFRLKVLTVLLYICLSVMLFYLFGSIMPLPPWGLVVGWVMALIAVELAYAFIFPYSFRYIADNDDDKMVILPV |
| IRIS_313-11521 | MLHHLKELAAVAGIHMILIYLCRFLLRRSRNVLFTVSNSLRFRLKVLTVLLYICLSVMLFYLFGSIMPLPPWGLVVGWVMALIAVELAYAFIFPYSFRYIADNDDDKMVILPV |
| IRIS_313-11522 | MLHQLKELAAVAGIHMILIYLCRFLLRRSRNVLFTVSNSLRFRLKVLTVLLYICLLVMLFYLFGSIMPLPPWGLVVGWVMALIAVELAYAFIFPYSFRYIADNDDDKMVILPV |
| IRIS_313-11523 | MLHHLKELAAVAGIHMILIYLCRFLLRRSRNVLFTVSNSLRFRLKVLTVLLYICLSVMLFYLFGSIMPLPPWGLVVGWVMALIAVELAYAFIFPYSFRYIADNDDDKMVILPV |
| IRIS_313-11524 | MLHHLKELAAVAGIHMILIYLCRFLLRRSRNVLFTVSNSLRFRLKVLTVLLYICLSVMLFYLFGSIMPLPPWGLVVGWVMALIAVELAYAFIFPYSFRYIADNDDDKMVILPV |
| IRIS_313-11525 | MLHHLKELAAVAGIHMILIYLCRFLLRRSRNVLFTVSNSLRFRLKVLTVLLYICLSVMLFYLFGSIMPLPPWGLVVGWVMALIAVELAYAFIFPYSFRYIADNDDDKMVILPV |
| IRIS_313-11526 | MLHHLKELAAVAGIHMILIYLCRFLLRRSRNVLFTVSNSLRFRLKVLTVLLYICLSVMLFYLFGSIMPLPPWGLVVGWVMALIAVELAYAFIFPYSFRYIADNDDDKMVILPV |
| IRIS_313-11527 | MLHHLKELAAVAGIHMILIYLCRFLLRRSRNVLFTVSNSLRFRLKVLTVLLYICLSVMLFYLFGSIMPLPPWGLVVGWVMALIAVELAYAFIFPYSFRYIADNDDDKMVILPV |
| IRIS_313-11528 | MLHHLKELAAVAGIHMILIYLCRFLLRRSRNVLFTVSNSLRFRLKVLTVLLYICLSVMLFYLFGSIMPLPPWGLVVGWVMALIAVELAYAFIFPYSFRYIADNDDDKMVILPV |
| IRIS_313-11530 | MLHHLKELAAVAGIHMILIYLCRFLLRRSRNVLFTVSNSLRFRLKVLTVLLYICLSVMLFYLFGSIMPLPPWGLVVGWVMALIAVELAYAFIFPYSFRYIADNDDDKMVILPV |
| IRIS_313-11532 | MLHHLKELAAVAGIHMILIYLCRFLLRRSRNVLFTVSNSLRFRLKVLTVLLYICLSVMLFYLFGSIMPLPPWGLVVGWVMALIAVELAYAFIFPYSFRYIADNDDDKMVILPV |
| IRIS_313-11534 | MLHHLKELAAVAGIHMILIYLCRFLLRRSRNVLFTVSNSLRFRLKVLTVLLYICLSVMLFYLFGSIMPLPPWGLVVGWVMALIAVELAYAFIFPYSFRYIADNDDDKMVILPV |
| IRIS_313-11535 | MLHHLKELAAVAGIHMILIYLCRFLLRRSRNVLFTVSNSLRFRLKVLTVLLYICLSVMLFYLFGSIMPLPPWGLVVGWVMALIAVELAYAFIFPYSFRYIADNDDDKMVILPV |
| IRIS_313-11536 | MLHHLKELAAVAGIHMILIYLCRFLLRRSRNVLFTVSNSLRFRLKVLTVLLYICLSVMLFYLFGSIMPLPPWGLVVGWVMALIAVELAYAFIFPYSFRYIADNDDDKMVILPV |
| IRIS_313-11537 | MLHHLKELAAVAGIHMILIYLCRFLLRRSRNVLFTVSNSLRFRLKVLTVLLYICLSVMLFYLFGSIMPLPPWGLVVGWVMALIAVELAYAFIFPYSFRYIADNDDDKMVILPV |
| IRIS_313-11538 | MLHHLKELAAVAGIHMILIYLCRFLLRRSRNVLFTVSNSLRFRLKVLTVLLYICLSVMLFYLFGSIMPLPPWGLVVGWVMALIAVELAYAFIFPYSFRYIADNDDDKMVILPV |
| IRIS_313-11539 | MLHHLKELAAVAGIHMILIYLCRFLLRRSRNVLFTVSNSLRFRLKVLTVLLYICLSVMLFYLFGSIMPLPPWGLVVGWVMALIAVELAYAFIFPYSFRYIADNDDDKMVILPV |
| IRIS_313-11540 | MLHHLKELAAVAGIHMILIYLCRFLLRRSRNVLFTVSNSLRFRLKVLTVLLYICLSVMLFYLFGSIMPLPPWGLVVGWVMALIAVELAYAFIFPYSFRYIADNDDDKMVILPV |
| IRIS_313-11541 | MLHHLKELAAVAGIHMILIYLCRFLLRRSRNVLFTVSNSLRFRLKVLTVLLYICLSVMLFYLFGSIMPLPPWGLVVGWVMALIAVELAYAFIFPYSFRYIADNDDDKMVILPV |
| IRIS_313-11542 | MLHHLKELAAVAGIHMILIYLCRFLLRRSRNVLFTVSNSLRFRLKVLTVLLYICLSVMLFYLFGSIMPLPPWGLVVGWVMALIAVELAYAFIFPYSFRYIADNDDDKMVILPV |
| IRIS_313-11543 | MLHHLKELAAVAGIHMILIYLCRFLLRRSRNVLFTVSNSLRFRLKVLTVLLYICLSVMLFYLFGSIMPLPPWGLVVGWVMALIAVELAYAFIFPYSFRYIADNDDDKMVILPV |
| IRIS_313-11544 | MLHHLKELAAVAGIHMILIYLCRFLLRRSRNVLFTVSNSLRFRLKVLTVLLYICLSVMLFYLFGSIMPLPPWGLVVGWVMALIAVELAYAFIFPYSFRYIADNDDDKMVILPV |
| IRIS_313-11545 | MLHHLKELAAVAGIHMILIYLCRFLLRRSRNVLFTVSNSLRFRLKVLTVLLYICLSVMLFYLFGSIMPLPPWGLVVGWVMALIAVELAYAFIFPYSFRYIADNDDDKMVILPV |
| IRIS_313-11546 | MLHHLKELAAVAGIHMILIYLCRFLLRRSRNVLFTVSNSLRFRLKVLTVLLYICLSVMLFYLFGSIMPLPPWGLVVGWVMALIAVELAYAFIFPYSFRYIADNDDDKMVILPV |
| IRIS_313-11547 | MLHHLKELAAVAGIHMILIYLCRFLLRRSRNVLFTVSNSLRFRLKVLTVLLYICLSVMLFYLFGSIMPLPPWGLVVGWVMALIAVELAYAFIFPYSFRYIADNDDDKMVILPV |
| IRIS_313-11548 | MLHHLKELAAVAGIHMILIYLCRFLLRRSRNVLFTVSNSLRFRLKVLTVLLYICLSVMLFYLFGSIMPLPPWGLVVGWVMALIAVELAYAFIFPYSFRYIADNDDDKMVILPV |
| IRIS_313-11549 | MLHHLKELAAVAGIHMILIYLCRFLLRRSRNVLFTVSNSLRFRLKVLTVLLYICLSVMLFYLFGSIMPLPPWGLVVGWVMALIAVELAYAFIFPYSFRYIADNDDDKMVILPV |
| IRIS_313-11551 | MLHHLKELAAVAGIHMILIYLCRFLLRRSRNVLFTVSNSLRFRLKVLTVLLYICLSVMLFYLFGSIMPLPPWGLVVGWVMALIAVELAYAFIFPYSFRYIADNDDDKMVILPV |
| IRIS_313-11554 | MLHHLKELAAVAGIHMILIYLCRFLLRRSRNVLFTVSNSLRFRLKVLTVLLYICLSVMLFYLFGSIMPLPPWGLVVGWVMALIAVELAYAFIFPYSFRYIADNDDDKMVILPV |
| IRIS_313-11555 | MLHHLKELAAVAGIHMILIYLCRFLLRRSRNVLFTVSNSLRFRLKVLTVLLYICLSVMLFYLFGSIMPLPPWGLVVGWVMALIAVELAYAFIFPYSFRYIADNDDDKMVILPV |
| IRIS_313-11556 | MLHHLKELAAVAGIHMILIYLCRFLLRRSRNVLFTVSNSLRFRLKVLTVLLYICLSVMLFYLFGSIMPLPPWGLVVGWVMALIAVELAYAFIFPYSFRYIADNDDDKMVILPV |
| IRIS_313-11557 | MLHHLKELAAVAGIHMILIYLCRFLLRRSRNVLFTVSNSLRFRLKVLTVLLYICLSVMLFYLFGSIMPLPPWGLVVGWVMALIAVELAYAFIFPYSFRYIADNDDDKMVILPV |
| IRIS_313-11558 | MLHHLKELAAVAGIHMILIYLCRFLLRRSRNVLFTVSNSLRFRLKVLTVLLYICLSVMLFYLFGSIMPLPPWGLVVGWVMALIAVELAYAFIFPYSFRYIADNDDDKMVILPV |
| IRIS_313-11561 | MLHHLKELAAVAGIHMILIYLCRFLLRRSRNVLFTVSNSLRFRLKVLTVLLYICLSVMLFYLFGSIMPLPPWGLVVGWVMALIAVELAYAFIFPYSFRYIADNDDDKMVILPV |
| IRIS_313-11563 | MLHHLKELAAVAGIHMILIYLCRFLLRRSRNVLFTVSNSLRFRLKVLTVLLYICLSVMLFYLFGSIMPLPPWGLVVGWVMALIAVELAYAFIFPYSFRYIADNDDDKMVILPV |
| IRIS_313-11564 | MLHHLKELAAVAGIHMILIYLCRFLLRRSRNVLFTVSNSLRFRLKVLTVLLYICLSVMLFYLFGSIMPLPPWGLVVGWVMALIAVELAYAFIFPYSFRYIADNDDDKMVILPV |
| IRIS_313-11565 | MLHHLKELAAVAGIHMILIYLCRFLLRRSRNVLFTVSNSLRFRLKVLTVLLYICLSVMLFYLFGSIMPLPPWGLVVGWVMALIAVELAYAFIFPYSFRYIADNDDDKMVILPV |
| IRIS_313-11566 | MLHHLKELAAVAGIHMILIYLCRFLLRRSRNVLFTVSNSLRFRLKVLTVLLYICLSVMLFYLFGSIMPLPPWGLVVGWVMALIAVELAYAFIFPYSFRYIADNDDDKMVILPV |
| IRIS_313-11567 | MLHHLKELAAVAGIHMILIYLCRFLLRRSRNVLFTVSNSLRFRLKVLTVLLYICLSVMLFYLFGSIMPLPPWGLVVGWVMALIAVELAYAFIFPYSFRYIADNDDDKMVILPV |
| IRIS_313-11568 | MLHHLKELAAVAGIHMILIYLCRFLLRRSRNVLFTVSNSLRFRLKVLTVLLYICLSVMLFYLFGSIMPLPPWGLVVGWVMALIAVELAYAFIFPYSFRYIADNDDDKMVILPV |
| IRIS_313-11570 | MLHQLKELAAVAGIHMILIYLCRFLLRRSRNVLFTVSNSLRFRLKVLTVLLYICLLVMLFYLFGSIMPLPPWGLVVGWVMALIAVELAYAFIFPYSFRYIADNDDDKMVILPV |
| IRIS_313-11571 | MLHHLKELAAVAGIHMILIYLCRFLLRRSRNVLFTVSNSLRFRLKVLTVLLYICLSVMLFYLFGSIMPLPPWGLVVGWVMALIAVELAYAFIFPYSFRYIADNDDDKMVILPV |
| IRIS_313-11572 | MLHHLKELAAVAGIHMILIYLCRFLLRRSRNVLFTVSNSLRFRLKVLTVLLYICLSVMLFYLFGSIMPLPPWGLVVGWVMALIAVELAYAFIFPYSFRYIADNDDDKMVILPV |
| IRIS_313-11573 | MLHHLKELAAVAGIHMILIYLCRFLLRRSRNVLFTVSNSLRFRLKVLTVLLYICLSVMLFYLFGSIMPLPPWGLVVGWVMALIAVELAYAFIFPYSFRYIADNDDDKMVILPV |
| IRIS_313-11574 | MLHHLKELAAVAGIHMILIYLCRFLLRRSRNVLFTVSNSLRFRLKVLTVLLYICLSVMLFYLFGSIMPLPPWGLVVGWVMALIAVELAYAFIFPYSFRYIADNDDDKMVILPV |
| IRIS_313-11575 | MLHHLKELAAVAGIHMILIYLCRFLLRRSRNVLFTVSNSLRFRLKVLTVLLYICLSVMLFYLFGSIMPLPPWGLVVGWVMALIAVELAYAFIFPYSFRYIADNDDDKMVILPV |
| IRIS_313-11576 | MLHHLKELAAVAGIHMILIYLCRFLLRRSRNVLFTVSNSLRFRLKVLTVLLYICLSVMLFYLFGSIMPLPPWGLVVGWVMALIAVELAYAFIFPYSFRYIADNDDDKMVILPV |
| IRIS_313-11577 | MLHHLKELAAVAGIHMILIYLCRFLLRRSRNVLFTVSNSLRFRLKVLTVLLYICLSVMLFYLFGSIMPLPPWGLVVGWVMALIAVELAYAFIFPYSFRYIADNDDDKMVILPV |
| IRIS_313-11578 | MLHHLKELAAVAGIHMILIYLCRFLLRRSRNVLFTVSNSLRFRLKVLTVLLYICLSVMLFYLFGSIMPLPPWGLVVGWVMALIAVELAYAFIFPYSFRYIADNDDDKMVILPV |
| IRIS_313-11579 | MLHHLKELAAVAGIHMILIYLCRFLLRRSRNVLFTVSNSLRFRLKVLTVLLYICLSVMLFYLFGSIMPLPPWGLVVGWVMALIAVELAYAFIFPYSFRYIADNDDDKMVILPV |
| IRIS_313-11580 | MLHHLKELAAVAGIHMILIYLCRFLLRRSRNVLFTVSNSLRFRLKVLTVLLYICLSVMLFYLFGSIMPLPPWGLVVGWVMALIAVELAYAFIFPYSFRYIADNDDDKMVILPV |
| IRIS_313-11582 | MLHHLKELAAVAGIHMILIYLCRFLLRRSRNVLFTVSNSLRFRLKVLTVLLYICLSVMLFYLFGSIMPLPPWGLVVGWVMALIAVELAYAFIFPYSFRYIADNDDDKMVILPV |
| IRIS_313-11584 | MLHHLKELAAVAGIHMILIYLCRFLLRRSRNVLFTVSNSLRFRLKVLTVLLYICLSVMLFYLFGSIMPLPPWGLVVGWVMALIAVELAYAFIFPYSFRYIADNDDDKMVILPV |
| IRIS_313-11585 | MLHHLKELAAVAGIHMILIYLCRFLLRRSRNVLFTVSNSLRFRLKVLTVLLYICLSVMLFYLFGSIMPLPPWGLVVGWVMALIAVELAYAFIFPYSFRYIADNDDDKMVILPV |
| IRIS_313-11586 | MLHHLKELAAVAGIHMILIYLCRFLLRRSRNVLFTVSNSLRFRLKVLTVLLYICLSVMLFYLFGSIMPLPPWGLVVGWVMALIAVELAYAFIFPYSFRYIADNDDDKMVILPV |
| IRIS_313-11588 | MLHHLKELAAVAGIHMILIYLCRFLLRRSRNVLFTVSNSLRFRLKVLTVLLYICLSVMLFYLFGSIMPLPPWGLVVGWVMALIAVELAYAFIFPYSFRYIADNDDDKMVILPV |
| IRIS_313-11591 | MLHHLKELAAVAGIHMILIYLCRFLLRRSRNVLFTVSNSLRFRLKVLTVLLYICLSVMLFYLFGSIMPLPPWGLVVGWVMALIAVELAYAFIFPYSFRYIADNDDDKMVILPV |
| IRIS_313-11594 | MLHHLKELAAVAGIHMILIYLCRFLLRRSRNVLFTVSNSLRFRLKVLTVLLYICLSVMLFYLFGSIMPLPPWGLVVGWVMALIAVELAYAFIFPYSFRYIADNDDDKMVILPV |
| IRIS_313-11595 | MLHHLKELAAVAGIHMILIYLCRFLLRRSRNVLFTVSNSLRFRLKVLTVLLYICLSVMLFYLFGSIMPLPPWGLVVGWVMALIAVELAYAFIFPYSFRYIADNDDDKMVILPV |
| IRIS_313-11596 | MLHHLKELAAVAGIHMILIYLCRFLLRRSRNVLFTVSNSLRFRLKVLTVLLYICLSVMLFYLFGSIMPLPPWGLVVGWVMALIAVELAYAFIFPYSFRYIADNDDDKMVILPV |
| IRIS_313-11597 | MLHHLKELAAVAGIHMILIYLCRFLLRRSRNVLFTVSNSLRFRLKVLTVLLYICLSVMLFYLFGSIMPLPPWGLVVGWVMALIAVELAYAFIFPYSFRYIADNDDDKMVILPV |
| IRIS_313-11598 | MLHHLKELAAVAGIHMILIYLCRFLLRRSRNVLFTVSNSLRFRLKVLTVLLYICLSVMLFYLFGSIMPLPPWGLVVGWVMALIAVELAYAFIFPYSFRYIADNDDDKMVILPV |
| IRIS_313-11599 | MLHHLKELAAVAGIHMILIYLCRFLLRRSRNVLFTVSNSLRFRLKVLTVLLYICLSVMLFYLFGSIMPLPPWGLVVGWVMALIAVELAYAFIFPYSFRYIADNDDDKMVILPV |
| IRIS_313-11600 | MLHHLKELAAVAGIHMILIYLCRFLLRRSRNVLFTVSNSLRFRLKVLTVLLYICLSVMLFYLFGSIMPLPPWGLVVGWVMALIAVELAYAFIFPYSFRYIADNDDDKMVILPV |
| IRIS_313-11602 | MLHHLKELAAVAGIHMILIYLCRFLLRRSRNVLFTVSNSLRFRLKVLTVLLYISLSAMLLYLFGSIMPLPPWGLVVGWVMALIAVELAYAFIFPYSFRYIADNDDDKMVILPV |
| IRIS_313-11603 | MLHHLKELAAVAGIHMILIYLCRFLLRRSRNVLFTVSNSLRFRLKVLTVLLYICLSVMLFYLFGSIMPLPPWGLVVGWVMALIAVELAYAFIFPYSFRYIADNDDDKMVILPV |
| IRIS_313-11604 | MLHHLKELAAVAGIHMILIYLCRFLLRLRRNVLFTVSNSLRFRLKVLTVLLYICLSVMLFYLFGSIMPLPPWGLVVGWVMALIAVELAYAFIFPYSFRYIADNDDDKMVILPV |
| IRIS_313-11606 | MLHHLKELAAVAGIHMILIYLCRFLLRRSRNVLFTVSNSLRFRLKVLTVLLYICLSVMLFYLFGSIMPLPPWGLVVGWVMALIAVELAYAFIFPYSFRYIADNDDDKMVILPV |
| IRIS_313-11607 | MLHHLKELAAVAGIHMILIYLCRFLLRRSRNVLFTVSNSLRFRLKVLTVLLYICLSVMLFYLFGSIMPLPPWGLVVGWVMALIAVELAYAFIFPYSFRYIADNDDDKMVILPV |
| IRIS_313-11608 | MLHHLKELAAVAGIHMILIYLCRFLLRRSRNVLFTVSNSLRFRLKVLTVLLYICLSVMLFYLFGSIMPLPPWGLVVGWVMALIAVELAYAFIFPYSFRYIADNDDDKMVILPV |
| IRIS_313-11609 | MLHHLKELAAVAGIHMILIYLCRFLLRRSRNVLFTVSNSLRFRLKVLTVLLYICLSVMLFYLFGSIMPLPPWGLVVGWVMALIAVELAYAFIFPYSFRYIADNDDDKMVILPV |
| IRIS_313-11610 | MLHHLKELAAVAGIHMILIYLCRFLLRRSRNVLFTVSNSLRFRLKVLTVLLYICLSVMLFYLFGSIMPLPPWGLVVGWVMALIAVELAYAFIFPYSFRYIADNDDDKMVILPV |
| IRIS_313-11611 | MLHHLKELAAVAGIHMILIYLCRFLLRRSRNVLFTVSNSLRFRLKVLTVLLYICLSVMLFYLFGSIMPLPPWGLVVGWVMALIAVELAYAFIFPYSFRYIADNDDDKMVILPV |
| IRIS_313-11615 | MLHHLKELAAVAGIHMILIYLCRFLLRRSRNVLFTVSNSLRFRLKVLTVLLYICLSVMLFYLFGSIMPLPPWGLVVGWVMALIAVELAYAFIFPYSFRYIADNDDDKMVILPV |
| IRIS_313-11616 | MLHHLKELAAVAGIHMILIYLCRFLLRRSRNVLFTVSNSLRFRLKVLTVLLYICLSVMLFYLFGSIMPLPPWGLVVGWVMALIAVELAYAFIFPYSFRYIADNDDDKMVILPV |
| IRIS_313-11617 | MLHHLKELAAVAGIHMILIYLCRFLLRRSRNVLFTVSNSLRFRLKVLTVLLYICLSVMLFYLFGSIMPLPPWGLVVGWVMALIAVELAYAFIFPYSFRYIADNDDDKMVILPV |
| IRIS_313-11618 | MLHHLKELAAVAGIHMILIYLCRFLLRRSRNVLFTVSNSLRFRLKVLTVLLYICLSVMLFYLFGSIMPLPPWGLVVGWVMALIAVELAYAFIFPYSFRYIADNDDDKMVILPV |
| IRIS_313-11619 | MLHHLKELAAVAGIHMILIYLCRFLLRRSRNVLFTVSNSLRFRLKVLTVLLYICLSVMLFYLFGSIMPLPPWGLVVGWVMALIAVELAYAFIFPYSFRYIADNDDDKMVILPV |
| IRIS_313-11620 | MLHHLKELAAVAGIHMILIYLCRFLLRRSRNVLFTVSNSLRFRLKVLTVLLYICLSVMLFYLFGSIMPLPPWGLVVGWVMALIAVELAYAFIFPYSFRYIADNDDDKMVILPV |
| IRIS_313-11621 | MLHHLKELAAVAGIHMILIYLCRFLLRRSRNVLFTVSNSLRFRLKVLTVLLYICLSVMLFYLFGSIMPLPPWGLVVGWVMALIAVELAYAFIFPYSFRYIADNDDDKMVILPV |
| IRIS_313-11622 | MLHHLKELAAVAGIHMILIYLCRFLLRRSRNVLFTVSNSLRFRLKVLTVLLYICLSVMLFYLFGSIMPLPPWGLVVGWVMALIAVELAYAFIFPYSFRYIADNDDDKMVILPV |
| IRIS_313-11623 | MLHQLKELAAVAGIHMILIYLCRFLLRRSRNVLFTVSNSLRFRLKVLTVLLYICLLVMLFYLFGSIMPLPPWGLVVGWVMALIAVELAYAFIFPYSFRYIADNDDDKMVILPV |
| IRIS_313-11624 | MLHHLKELAAVAGIHMILIYLCRFLLRRSRNVLFTVSNSLRFRLKVLTVLLYICLSVMLFYLFGSIMPLPPWGLVVGWVMALIAVELAYAFIFPYSFRYIADNDDDKMVILPV |
| IRIS_313-11625 | MLHHLKELAAVAGIHMILIYLCRFLLRRSRNVLFTVSNSLRFRLKVLTVLLYICLSVMLFYLFGSIMPLPPWGLVVGWVMALIAVELAYAFIFPYSFRYIADNDDDKMVILPV |
| IRIS_313-11626 | MLHHLKELAAVAGIHMILIYLCRFLLRRSRNVLFTVSNSLRFRLKVLTVLLYICLSVMLFYLFGSIMPLPPWGLVVGWVMALIAVELAYAFIFPYSFRYIADNDDDKMVILPV |
| IRIS_313-11627 | MLHHLKELAAVAGIHMILIYLCRFLLRRSRNVLFTVSNSLRFRLKVLTVLLYICLSVMLFYLFGSIMPLPPWGLVVGWVMALIAVELAYAFIFPYSFRYIADNDDDKMVILPV |
| IRIS_313-11628 | MLHHLKELAAVAGIHMILIYLCRFLLRRSRNVLFTVSNSLRFRLKVLTVLLYICLSVMLFYLFGSIMPLPPWGLVVGWVMALIAVELAYAFIFPYSFRYIADNDDDKMVILPV |
| IRIS_313-11629 | MLHHLKELAAVAGIHMILIYLCRFLLRRSRNVLFTVSNSLRFRLKVLTVLLYICLSVMLFYLFGSIMPLPPWGLVVGWVMALIAVELAYAFIFPYSFRYIADNDDDKMVILPV |

| | |
|---|---|
| IRIS_313-11630 | MLHHLKELAAVAGIHMILIYLCRFLLRRSRNVLFTVSNSLRFRLKVLTVLLYICLSVMLFYLFGSIMPLPPWGLVVGWVMALIAVELAYAFIFPYSFRYIADNDDDKMVILPV |
| IRIS_313-11632 | MLHHLKELAAVAGIHMILIYLCRFLLRRSRNVLFTVSNSLRFRLKVLTVLLYICLSVMLFYLFGSIMPLPPWGLVVGWVMALIAVELAYAFIFPYSFRYIADNDDDKMVILPV |
| IRIS_313-11635 | MLHHLKELAAVAGIHMILIYLCRFLLRRSRNVLFTVSNSLRFRLKVLTVLLYICLSVMLFYLFGSIMPLPPWGLVVGWVMALIAVELAYAFIFPYSFRYIADNDDDKMVILPV |
| IRIS_313-11636 | MLHHLKELAAVAGIHMILIYLCRFLLRRSRNVLFTVSNSLRFRLKVLTVLLYISLSVMLLYLFGSIMPLPPWGLVVGWVMALIAVELAYAFIFPYSFRYIADNDDDKMVILPV |
| IRIS_313-11638 | MLHHLKELAAVAGIHMILIYLCRFLLRRSRNVLFTVSNSLRFRLKVLTVLLYICLSVMLFYLFGSIMPLPPWGLVVGWVMALIAVELAYAFIFPYSFRYIADNDDDKMVILPV |
| IRIS_313-11639 | MLHHLKELAAVAGIHMILIYLCRFLLRRSRNVLFTVSNSLRFRLKVLTVLLYICLSVMLFYLFGSIMPLPPWGLVVGWVMALIAVELAYAFIFPYSFRYIADNDDDKMVILPV |
| IRIS_313-11640 | MLHHLKELAAVAGIHMILIYLCRFLLRRSRNVLFTVSNSLRFRLKVLTVLLYICLSVMLFYLFGSIMPLPPWGLVVGWVMALIAVELAYAFIFPYSFRYIADNDDDKMVILPV |
| IRIS_313-11641 | MLHHLKELAAVAGIHMILIYLCRFLLRRSRNVLFTVSNSLRFRLKVLTVLLYICLSVMLFYLFGSIMPLPPWGLVVGWVMALIAVELAYAFIFPYSFRYIADNDDDKMVILPV |
| IRIS_313-11642 | MLHHLKELAAVAGIHMILIYLCRFLLRRSRNVLFTVSNSLRFRLKVLTVLLYICLSVMLFYLFGSIMPLPPWGLVVGWVMALIAVELAYAFIFPYSFRYIADNDDDKMVILPV |
| IRIS_313-11643 | MLHHLKELAAVAGIHMILIYLCRFLLRRSRNVLFTVSNSLRFRLKVLTVLLYICLSVMLFYLFGSIMPLPPWGLVVGWVMALIAVELAYAFIFPYSFRYIADNDDDKMVILPV |
| IRIS_313-11644 | MLHHLKELAAVAGIHMILIYLCRFLLRRSRNVLFTVSNSLRFRLKVLTVLLYICLSVMLFYLFGSIMPLPPWGLVVGWVMALIAVELAYAFIFPYSFRYIADNDDDKMVILPV |
| IRIS_313-11645 | MLHHLKELAAVAGIHMILIYLCRFLLRRSRNVLFTVSNSLRFRLKVLTVLLYICLSVMLFYLFGSIMPLPPWGLVVGWVMALIAVELAYAFIFPYSFRYIADNDDDKMVILPV |
| IRIS_313-11646 | MLHHLKELAAVAGIHMILIYLCRFLLRRSRNVLFTVSNSLRFRLKVLTVLLYICLSVMLFYLFGSIMPLPPWGLVVGWVMALIAVELAYAFIFPYSFRYIADNDDDKMVILPV |
| IRIS_313-11647 | MLHHLKELAAVAGIHMILIYLCRFLLRRSRNVLFTVSNSLRFRLKVLTVLLYICLSVMLFYLFGSIMPLPPWGLVVGWVMALIAVELAYAFIFPYSFRYIADNDDDKMVILPV |
| IRIS_313-11648 | MLHHLKELAAVAGIHMILIYLCRFLLRRSRNVLFTVSNSLRFRLKVLTVLLYICLSVMLFYLFGSIMPLPPWGLVVGWVMALIAVELAYAFIFPYSFRYIADNDDDKMVILPV |
| IRIS_313-11650 | MLHHLKELAAVAGIHMILIYLCRFLLRRSRNVLFTVSNSLRFRLKVLTVLLYICLSVMLFYLFGSIMPLPPWGLVVGWVMALIAVELAYAFIFPYSFRYIADNDDDKMVILPV |
| IRIS_313-11651 | MLHQLKELAAVAGIHMILIYLCRFLLRRSRNVLFTVSNSLRFRLKVLTVLLYICLLVMLFYLFGSIMPLPPWGLVVGWVMALIAVELAYAFIFPYSFRYIADNDDDKMVILPV |
| IRIS_313-11652 | MLHHLKELAAVAGIHMILIYLCRFLLRRSRNVLFTVSNSLRFRLKVLTVLLYICLSVMLFYLFGSIMPLPPWGLVVGWVMALIAVELAYAFIFPYSFRYIADNDDDKMVILPV |
| IRIS_313-11653 | MLHQLKELAAVAGIHMILIYLCRFLLRRSRNVLFTVSNSLRFRLKVLTVLLYICLLVMLFYLFGSIMPLPPWGLVVGWVMALIAVELAYAFIFPYSFRYIADNDDDKMVILPV |
| IRIS_313-11654 | MLHHLKELAAVAGIHMILIYLCRFLLRRSRNVLFTVSNSLRFRLKVLTVLLYICLSVMLFYLFGSIMPLPPWGLVVGWVMALIAVELAYAFIFPYSFRYIADNDDDKMVILPV |
| IRIS_313-11655 | MLHHLKELAAVAGIHMILIYLCRFLLRRSRNVLFTVSNSLRFRLKVLTVLLYICLSVMLFYLFGSIMPLPPWGLVVGWVMALIAVELAYAFIFPYSFRYIADNDDDKMVILPV |
| IRIS_313-11656 | MLHHLKELAAVAGIHMILIYLCRFLLRRSRNVLFTVSNSLRFRLKVLTVLLYICLSVMLFYLFGSIMPLPPWGLVVGWVMALIAVELAYAFIFPYSFRYIADNDDDKMVILPV |
| IRIS_313-11657 | MLHHLKELAAVAGIHMILIYLCRFLLRRSRNVLFTVSNSLRFRLKVLTVLLYICLSVMLFYLFGSIMPLPPWGLVVGWVMALIAVELAYAFIFPYSFRYIADNDDDKMVILPV |
| IRIS_313-11658 | MLHHLKELAAVAGIHMILIYLCRFLLRRSRNVLFTVSNSLRFRLKVLTVLLYICLSVMLFYLFGSIMPLPPWGLVVGWVMALIAVELAYAFIFPYSFRYIADNDDDKMVILPV |
| IRIS_313-11659 | MLHHLKELAAVAGIHMILIYLCRFLLRRSRNVLFTVSNSLRFRLKVLTVLLYICLSVMLFYLFGSIMPLPPWGLVVGWVMALIAVELAYAFIFPYSFRYIADNDDDKMVILPV |
| IRIS_313-11660 | MLHHLKELAAVAGIHMILIYLCRFLLRRSRNVLFTVSNSLRFRLKVLTVLLYICLSVMLFYLFGSIMPLPPWGLVVGWVMALIAVELAYAFIFPYSFRYIADNDDDKMVILPV |
| IRIS_313-11661 | MLHHLKELAAVAGIHMILIYLCRFLLRRSRNVLFTVSNSLRFRLKVLTVLLYICLSVMLFYLFGSIMPLPPWGLVVGWVMALIAVELAYAFIFPYSFRYIADNDDDKMVILPV |
| IRIS_313-11663 | MLHHLKELAAVAGIHMILIYLCRFLLRRSRNVLFTVSNSLRFRLKVLTVLLYICLSVMLFYLFGSIMPLPPWGLVVGWVMALIAVELAYAFIFPYSFRYIADNDDDKMVILPV |
| IRIS_313-11664 | MLHHLKELAAVAGIHMILIYLCRFLLRRSRNVLFTVSNSLRFRLKVLTVLLYICLSVMLFYLFGSIMPLPPWGLVVGWVMALIAVELAYAFIFPYSFRYIADNDDDKMVILPV |
| IRIS_313-11665 | MLHHLKELAAVAGIHMILIYLCRFLLRRSRNVLFTVSNSLRFRLKVLTVLLYICLSVMLFYLFGSIMPLPPWGLVVGWVMALIAVELAYAFIFPYSFRYIADNDDDKMVILPV |
| IRIS_313-11666 | MLHHLKELAAVAGIHMILIYLCRFLLRRSRNVLFTVSNSLRFRLKVLTVLLYICLSVMLFYLFGSIMPLPPWGLVVGWVMALIAVELAYAFIFPYSFRYIADNDDDKMVILPV |
| IRIS_313-11667 | MLHHLKELAAVAGIHMILIYLCRFLLRRSRNVLFTVSNSLRFRLKVLTVLLYICLSVMLFYLFGSIMPLPPWGLVVGWVMALIAVELAYAFIFPYSFRYIADNDDDKMVILPV |
| IRIS_313-11668 | MLHHLKELAAVAGIHMILIYLCRFLLRRSRNVLFTVSNSLRFRLKVLTVLLYICLSVMLFYLFGSIMPLPPWGLVVGWVMALIAVELAYAFIFPYSFRYIADNDDDKMVILPV |
| IRIS_313-11669 | MLHHLKELAAVAGIHMILIYLCRFLLRRSRNVLFTVSNSLRFRLKVLTVLLYICLSVMLFYLFGSIMPLPPWGLVVGWVMALIAVELAYAFIFPYSFRYIADNDDDKMVILPV |
| IRIS_313-11671 | MLHHLKELAAVAGIHMILIYLCRFLLRRSRNVLFTVSNSLRFRLKVLTVLLYICLSVMLFYLFGSIMPLPPWGLVVGWVMALIAVELAYAFIFPYSFRYIADNDDDKMVILPV |
| IRIS_313-11672 | MLHHLKELAAVAGIHMILIYLCRFLLRRSRNVLFTVSNSLRFRLKVLTVLLYICLSVMLFYLFGSIMPLPPWGLVVGWVMALIAVELAYAFIFPYSFRYIADNDDDKMVILPV |
| IRIS_313-11673 | MLHQLKELAAVAGIHMILIYLCRFLLRRSRNVLFTVSNSLRFRLKVLTVLLYICLLVMLFYLFGSIMPLPPWGLVVGWVMALIAVELAYAFIFPYSFRYIADNDDDKMVILPV |
| IRIS_313-11674 | MLHHLKELAAVAGIHMILIYLCRFLLRRSRNVLFTVSNSLRFRLKVLTVLLYICLSVMLFYLFGSIMPLPPWGLVVGWVMALIAVELAYAFIFPYSFRYIADNDDDKMVILPV |
| IRIS_313-11676 | MLHHLKELAAVAGIHMILIYLCRFLLRRSRNVLFTVSNSLRFRLKVLTVLLYICLSVMLFYLFGSIMPLPPWGLVVGWVMALIAVELAYAFIFPYSFRYIADNDDDKMVILPV |
| IRIS_313-11677 | MLHHLKELAAVAGIHMILIYLCRFLLRRSRNVLFTVSNSLRFRLKVLTVLLYICLSVMLFYLFGSIMPLPPWGLVVGWVMALIAVELAYAFIFPYSFRYIADNDDDKMVILPV |
| IRIS_313-11678 | MLHHLKELAAVAGIHMILIYLCRFLLRRSRNVLFTVSNSLRFRLKVLTVLLYICLSVMLFYLFGSIMPLPPWGLVVGWVMALIAVELAYAFIFPYSFRYIADNDDDKMVILPV |
| IRIS_313-11679 | MLHHLKELAAVAGIHMILIYLCRFLLRRSRNVLFTVSNSLRFRLKVLTVLLYICLSVMLFYLFGSIMPLPPWGLVVGWVMALIAVELAYAFIFPYSFRYIADNDDDKMVILPV |
| IRIS_313-11680 | MLHHLKELAAVAGIHMILIYLCRFLLRRSRNVLFTVSNSLRFRLKVLTVLLYICLSVMLFYLFGSIMPLPPWGLVVGWVMALIAVELAYAFIFPYSFRYIADNDDDKMVILPV |
| IRIS_313-11681 | MLHHLKELAAVAGIHMILIYLCRFLLRRSRNVLFTVSNSLRFRLKVLTVLLYICLSVMLFYLFGSIMPLPPWGLVVGWVMALIAVELAYAFIFPYSFRYIADNDDDKMVILPV |
| IRIS_313-11682 | MLHHLKELAAVAGIHMILIYLCRFLLRRSRNVLFTVSNSLRFRLKVLTVLLYICLSVMLFYLFGSIMPLPPWGLVVGWVMALIAVELAYAFIFPYSFRYIADNDDDKMVILPV |
| IRIS_313-11683 | MLHHLKELAAVAGIHMILIYLCRFLLRRSRNVLFTVSNSLRFRLKVLTVLLYICLSVMLFYLFGSIMPLPPWGLVVGWVMALIAVELAYAFIFPYSFRYIADNDDDKMVILPV |
| IRIS_313-11684 | MLHHLKELAAVAGIHMILIYLCRFLLRRSRNVLFTVSNSLRFRLKVLTVLLYICLSVMLFYLFGSIMPLPPWGLVVGWVMALIAVELAYAFIFPYSFRYIADNDDDKMVILPV |
| IRIS_313-11685 | MLHHLKELAAVAGIHMILIYLCRFLLRRSRNVLFTVSNSLRFRLKVLTVLLYICLSVMLFYLFGSIMPLPPWGLVVGWVMALIAVELAYAFIFPYSFRYIADNDDDKMVILPV |
| IRIS_313-11686 | MLHHLKELAAVAGIHMILIYLCRFLLRRSRNVLFTVSNSLRFRLKVLTVLLYICLSVMLFYLFGSIMPLPPWGLVVGWVMALIAVELAYAFIFPYSFRYIADNDDDKMVILPV |
| IRIS_313-11687 | MLHHLKELAAVAGIHMILIYLCRFLLRRSRNVLFTVSNSLRFRLKVLTVLLYICLSVMLFYLFGSIMPLPPWGLVVGWVMALIAVELAYAFIFPYSFRYIADNDDDKMVILPV |
| IRIS_313-11688 | MLHHLKELAAVAGIHMILIYLCRFLLRRSRNVLFTVSNSLRFRLKVLTVLLYICLSVMLFYLFGSIMPLPPWGLVVGWVMALIAVELAYAFIFPYSFRYIADNDDDKMVILPV |
| IRIS_313-11689 | MLHHLKELAAVAGIHMILIYLCRFLLRRSRNVLFTVSNSLRFRLKVLTVLLYICLSVMLFYLFGSIMPLPPWGLVVGWVMALIAVELAYAFIFPYSFRYIADNDDDKMVILPV |
| IRIS_313-11690 | MLHHLKELAAVAGIHMILIYLCRFLLRRSRNVLFTVSNSLRFRLKVLTVLLYICLSVMLFYLFGSIMPLPPWGLVVGWVMALIAVELAYAFIFPYSFRYIADNDDDKMVILPV |
| IRIS_313-11691 | MLHHLKELAAVAGIHMILIYLCRFLLRRSRNVLFTVSNSLRFRLKVLTVLLYICLSVMLFYLFGSIMPLPPWGLVVGWVMALIAVELAYAFIFPYSFRYIADNDDDKMVILPV |
| IRIS_313-11692 | MLHHLKELAAVAGIHMILIYLCRFLLRRSRNVLFTVSNSLRFRLKVLTVLLYICLSVMLFYLFGSIMPLPPWGLVVGWVMALIAVELAYAFIFPYSFRYIADNDDDKMVILPV |
| IRIS_313-11693 | MLHHLKELAAVAGIHMILIYLCRFLLRRSRNVLFTVSNSLRFRLKVLTVLLYICLSVMLFYLFGSIMPLPPWGLVVGWVMALIAVELAYAFIFPYSFRYIADNDDDKMVILPV |
| IRIS_313-11694 | MLHHLKELAAVAGIHMILIYLCRFLLRRSRNVLFTVSNSLRFRLKVLTVLLYICLSVMLFYLFGSIMPLPPWGLVVGWVMALIAVELAYAFIFPYSFRYIADNDDDKMVILPV |
| IRIS_313-11695 | MLHHLKELAAVAGIHMILIYLCRFLLRRSRNVLFTVSNSLRFRLKVLTVLLYICLSVMLFYLFGSIMPLPPWGLVVGWVMALIAVELAYAFIFPYSFRYIADNDDDKMVILPV |
| IRIS_313-11698 | MLHHLKELAAVAGIHMILIYLCRFLLRRSRNVLFTVSNSLRFRLKVLTVLLYICLSVMLFYLFGSIMPLPPWGLVVGWVMALIAVELAYAFIFPYSFRYIADNDDDKMVILPV |
| IRIS_313-11700 | MLHHLKELAAVAGIHMILIYLCRFLLRRSRNVLFTVSNSLRFRLKVLTVLLYICLSVMLFYLFGSIMPLPPWGLVVGWVMALIAVELAYAFIFPYSFRYIADNDDDKMVILPV |
| IRIS_313-11702 | MLHHLKELAAVAGIHMILIYLCRFLLRRSRNVLFTVSNSLRFRLKVLTVLLYICLSVMLFYLFGSIMPLPPWGLVVGWVMALIAVELAYAFIFPYSFRYIADNDDDKMVILPV |
| IRIS_313-11704 | MLHHLKELAAVAGIHMILIYLCRFLLRRSRNVLFTVSNSLRFRLKVLTVLLYICLSVMLFYLFGSIMPLPPWGLVVGWVMALIAVELAYAFIFPYSFRYIADNDDDKMVILPV |
| IRIS_313-11705 | MLHHLKELAAVAGIHMILIYLCRFLLRRSRNVLFTVSNSLRFRLKVLTVLLYICLSVMLFYLFGSIMPLPPWGLVVGWVMALIAVELAYAFIFPYSFRYIADNDDDKMVILPV |
| IRIS_313-11706 | MLHHLKELAAVAGIHMILIYLCRFLLRRSRNVLFTVSNSLRFRLKVLTVLLYICLSVMLFYLFGSIMPLPPWGLVVGWVMALIAVELAYAFIFPYSFRYIADNDDDKMVILPV |
| IRIS_313-11707 | MLHHLKELAAVAGIHMILIYLCRFLLRRSRNVLFTVSNSLRFRLKVLTVLLYICLSVMLFYLFGSIMPLPPWGLVVGWVMALIAVELAYAFIFPYSFRYIADNDDDKMVILPV |
| IRIS_313-11708 | MLHHLKELAAVAGIHMILIYLCRFLLRRSRNVLFTVSNSLRFRLKVLTVLLYICLSVMLFYLFGSIMPLPPWGLVVGWVMALIAVELAYAFIFPYSFRYIADNDDDKMVILPV |
| IRIS_313-11709 | MLHHLKELAAVAGIHMILIYLCRFLLRRSRNVLFTVSNSLRFRLKVLTVLLYICLSVMLFYLFGSIMPLPPWGLVVGWVMALIAVELAYAFIFPYSFRYIADNDDDKMVILPV |
| IRIS_313-11710 | MLHHLKELAAVAGIHMILIYLCRFLLRRSRNVLFTVSNSLRFRLKVLTVLLYICLSVMLFYLFGSIMPLPPWGLVVGWVMALIAVELAYAFIFPYSFRYIADNDDDKMVILPV |
| IRIS_313-11711 | MLHHLKELAAVAGIHMILIYLCRFLLRRSRNVLFTVSNSLRFRLKVLTVLLYICLSVMLFYLFGSIMPLPPWGLVVGWVMALIAVELAYAFIFPYSFRYIADNDDDKMVILPV |
| IRIS_313-11712 | MLHHLKELAAVAGIHMILIYLCRFLLRRSRNVLFTVSNSLRFRLKVLTVLLYICLSVMLFYLFGSIMPLPPWGLVVGWVMALIAVELAYAFIFPYSFRYIADNDDDKMVILPV |
| IRIS_313-11713 | MLHHLKELAAVAGIHMILIYLCRFLLRRSRNVLFTVSNSLRFRLKVLTVLLYICLSVMLFYLFGSIMPLPPWGLVVGWVMALIAVELAYAFIFPYSFRYIADNDDDKMVILPV |
| IRIS_313-11714 | MLHHLKELAAVAGIHMILIYLCRFLLRRSRNVLFTVSNSLRFRLKVLTVLLYICLSVMLFYLFGSIMPLPPWGLVVGWVMALIAVELAYAFIFPYSFRYIADNDDDKMVILPV |
| IRIS_313-11716 | MLHHLKELAAVAGIHMILIYLCRFLLRRSRNVLFTVSNSLRFRLKVLTVLLYICLSVMLFYLFGSIMPLPPWGLVVGWVMALIAVELAYAFIFPYSFRYIADNDDDKMVILPV |
| IRIS_313-11717 | MLHHLKELAAVAGIHMILIYLCRFLLRRSRNVLFTVSNSLRFRLKVLTVLLYICLSVMLFYLFGSIMPLPPWGLVVGWVMALIAVELAYAFIFPYSFRYIADNDDDKMVILPV |
| IRIS_313-11719 | MLHHLKELAAVAGIHMILIYLCRFLLRRSRNVLFTVSNSLRFRLKVLTVLLYICLSVMLFYLFGSIMPLPPWGLVVGWVMALIAVELAYAFIFPYSFRYIADNDDDKMVILPV |
| IRIS_313-11720 | MLHHLKELAAVAGIHMILIYLCRFLLRRSRNVLFTVSNSLRFRLKVLTVLLYICLSVMLFYLFGSIMPLPPWGLVVGWVMALIAVELAYAFIFPYSFRYIADNDDDKMVILPV |
| IRIS_313-11721 | MLHHLKELAAVAGIHMILIYLCRFLLRRSRNVLFTVSNSLRFRLKVLTVLLYICLSVMLFYLFGSIMPLPPWGLVVGWVMALIAVELAYAFIFPYSFRYIADNDDDKMVILPV |
| IRIS_313-11722 | MLHHLKELAAVAGIHMILIYLCRFLLRRSRNVLFTVSNSLRFRLKVLTVLLYICLSVMLFYLFGSIMPLPPWGLVVGWVMALIAVELAYAFIFPYSFRYIADNDDDKMVILPV |
| IRIS_313-11723 | MLHHLKELAAVAGIHMILIYLCRFLLRRSRNVLFTVSNSLRFRLKVLTVLLYICLSVMLFYLFGSIMPLPPWGLVVGWVMALIAVELAYAFIFPYSFRYIADNDDDKMVILPV |
| IRIS_313-11724 | MLHHLKELAAVAGIHMILIYLCRFLLRRSRNVLFTVSNSLRFRLKVLTVLLYICLSVMLFYLFGSIMPLPPWGLVVGWVMALIAVELAYAFIFPYSFRYIADNDDDKMVILPV |
| IRIS_313-11725 | MLHHLKELAAVAGIHMILIYLCRFLLRRSRNVLFTVSNSLRFRLKVLTVLLYICLSVMLFYLFGSIMPLPPWGLVVGWVMALIAVELAYAFIFPYSFRYIADNDDDKMVILPV |
| IRIS_313-11726 | MLHHLKELAAVAGIHMILIYLCRFLLRRSRNVLFTVSNSLRFRLKVLTVLLYICLSVMLFYLFGSIMPLPPWGLVVGWVMALIAVELAYAFIFPYSFRYIADNDDDKMVILPV |
| IRIS_313-11727 | MLHHLKELAAVAGIHMILIYLCRFLLRRSRNVLFTVSNSLRFRLKVLTVLLYICLSVMLFYLFGSIMPLPPWGLVVGWVMALIAVELAYAFIFPYSFRYIADNDDDKMVILPV |
| IRIS_313-11728 | MLHHLKELAAVAGIHMILIYLCRFLLRRSRNVLFTVSNSLRFRLKVLTVLLYICLSVMLFYLFGSIMPLPPWGLVVGWVMALIAVELAYAFIFPYSFRYIADNDDDKMVILPV |
| IRIS_313-11729 | MLHHLKELAAVAGIHMILIYLCRFLLRRSRNVLFTVSNSLRFRLKVLTVLLYICLSVMLFYLFGSIMPLPPWGLVVGWVMALIAVELAYAFIFPYSFRYIADNDDDKMVILPV |
| IRIS_313-11730 | MLHHLKELAAVAGIHMILIYLCRFLLRRSRNVLFTVSNSLRFRLKVLTVLLYICLSVMLFYLFGSIMPLPPWGLVVGWVMALIAVELAYAFIFPYSFRYIADNDDDKMVILPV |
| IRIS_313-11731 | MLHHLKELAAVAGIHMILIYLCRFLLRRSRNVLFTVSNSLRFRLKVLTVLLYICLSVMLFYLFGSIMPLPPWGLVVGWVMALIAVELAYAFIFPYSFRYIADNDDDKMVILPV |
| IRIS_313-11732 | MLHHLKELAAVAGIHMILIYLCRFLLRRSRNVLFTVSNSLRFRLKVLTVLLYICLSVMLFYLFGSIMPLPPWGLVVGWVMALIAVELAYAFIFPYSFRYIADNDDDKMVILPV |
| IRIS_313-11733 | MLHHLKELAAVAGIHMILIYLCRFLLRRSRNVLFTVSNSLRFRLKVLTVLLYICLSVMLFYLFGSIMPLPPWGLVVGWVMALIAVELAYAFIFPYSFRYIADNDDDKMVILPV |
| IRIS_313-11734 | MLHHLKELAAVAGIHMILIYLCRFLLRRSRNVLFTVSNSLRFRLKVLTVLLYICLSVMLFYLFGSIMPLPPWGLVVGWVMALIAVELAYAFIFPYSFRYIADNDDDKMVILPV |
| IRIS_313-11735 | MLHHLKELAAVAGIHMILIYLCRFLLRRSRNVLFTVSNSLRFRLKVLTVLLYICLSVMLFYLFGSIMPLPPWGLVVGWVMALIAVELAYAFIFPYSFRYIADNDDDKMVILPV |
| IRIS_313-11736 | MLHQLKELAAVAGIHMILIYLCRFLLRRSRNVLFTVSNSLRFRLKVLTVLLYICLLVMLFYLFGSIMPLPPWGLVVGWVMALIAVELAYAFIFPYSFRYIADNDDDKMVILPV |
| IRIS_313-11737 | MLHHLKELAAVAGIHMILIYLCRFLLRRSRNVLFTVSNSLRFRLKVLTVLLYICLSVMLFYLFGSIMPLPPWGLVVGWVMALIAVELAYAFIFPYSFRYIADNDDDKMVILPV |
| IRIS_313-11738 | MLHHLKELAAVAGIHMILIYLCRFLLRRSRNVLFTVSNSLRFRLKVLTVLLYICLSVMLFYLFGSIMPLPPWGLVVGWVMALIAVELAYAFIFPYSFRYIADNDDDKMVILPV |
| IRIS_313-11739 | MLHHLKELAAVAGIHMILIYLCRFLLRRSRNVLFTVSNSLRFRLKVLTVLLYICLSVMLFYLFGSIMPLPPWGLVVGWVMALIAVELAYAFIFPYSFRYIADNDDDKMVILPV |
| IRIS_313-11740 | MLHHLKELAAVAGIHMILIYLCRFLLRRSRNVLFTVSNSLRFRLKVLTVLLYICLSVMLFYLFGSIMPLPPWGLVVGWVMALIAVELAYAFIFPYSFRYIADNDDDKMVILPV |
| IRIS_313-11741 | MLHHLKELAAVAGIHMILIYLCRFLLRRSRNVLFTVSNSLRFRLKVLTVLLYICLSVMLFYLFGSIMPLPPWGLVVGWVMALIAVELAYAFIFPYSFRYIADNDDDKMVILPV |
| IRIS_313-11742 | MLHHLKELAAVAGIHMILIYLCRFLLRRSRNVLFTVSNSLRFRLKVLTVLLYICLSVMLFYLFGSIMPLPPWGLVVGWVMALIAVELAYAFIFPYSFRYIADNDDDKMVILPV |
| IRIS_313-11743 | MLHHLKELAAVAGIHMILIYLCRFLLRRSRNVLFTVSNSLRFRLKVLTVLLYICLSVMLFYLFGSIMPLPPWGLVVGWVMALIAVELAYAFIFPYSFRYIADNDDDKMVILPV |
| IRIS_313-11744 | MLHHLKELAAVAGIHMILIYLCRFLLRRSRNVLFTVSNSLRFRLKVLTVLLYICLSVMLFYLFGSIMPLPPWGLVVGWVMALIAVELAYAFIFPYSFRYIADNDDDKMVILPV |
| IRIS_313-11745 | MLHHLKELAAVAGIHMILIYLCRFLLRRSRNVLFTVSNSLRFRLKVLTVLLYICLSVMLFYLFGSIMPLPPWGLVVGWVMALIAVELAYAFIFPYSFRYIADNDDDKMVILPV |
| IRIS_313-11746 | MLHHLKELAAVAGIHMILIYLCRFLLRRSRNVLFTVSNSLRFRLKVLTVLLYICLSVMLFYLFGSIMPLPPWGLVVGWVMALIAVELAYAFIFPYSFRYIADNDDDKMVILPV |
| IRIS_313-11747 | MLHHLKELAAVAGIHMILIYLCRFLLRRSRNVLFTVSNSLRFRLKVLTVLLYICLSVMLFYLFGSIMPLPPWGLVVGWVMALIAVELAYAFIFPYSFRYIADNDDDKMVILPV |
| IRIS_313-11748 | MLHHLKELAAVAGIHMILIYLCRFLLRRSRNVLFTVSNSLRFRLKVLTVLLYICLSVMLFYLFGSIMPLPPWGLVVGWVMALIAVELAYAFIFPYSFRYIADNDDDKMVILPV |
| IRIS_313-11750 | MLHHLKELAAVAGIHMILIYLCRFLLRRSRNVLFTVSNSLRFRLKVLTVLLYICLSVMLFYLFGSIMPLPPWGLVVGWVMALIAVELAYAFIFPYSFRYIADNDDDKMVILPV |
| IRIS_313-11751 | MLHHLKELAAVAGIHMILIYLCRFLLRRSRNVLFTVSNSLRFRLKVLTVLLYICLSVMLFYLFGSIMPLPPWGLVVGWVMALIAVELAYAFIFPYSFRYIADNDDDKMVILPV |
| IRIS_313-11752 | MLHHLKELAAVAGIHMILIYLCRFLLRRSRNVLFTVSNSLRFRLKVLTVLLYICLSVMLFYLFGSIMPLPPWGLVVGWVMALIAVELAYAFIFPYSFRYIADNDDDKMVILPV |
| IRIS_313-11753 | MLHHLKELAAVAGIHMILIYLCRFLLRRSRNVLFTVSNSLRFRLKVLTVLLYICLSVMLFYLFGSIMPLPPWGLVVGWVMALIAVELAYAFIFPYSFRYIADNDDDKMVILPV |
| IRIS_313-11754 | MLHHLKELAAVAGIHMILIYLCRFLLRRSRNVLFTVSNSLRFRLKVLTVLLYICLSVMLFYLFGSIMPLPPWGLVVGWVMALIAVELAYAFIFPYSFRYIADNDDDKMVILPV |
| IRIS_313-11755 | MLHHLKELAAVAGIHMILIYLCRFLLRRSRNVLFTVSNSLRFRLKVLTVLLYICLSVMLFYLFGSIMPLPPWGLVVGWVMALIAVELAYAFIFPYSFRYIADNDDDKMVILPV |
| IRIS_313-11756 | MLHHLKELAAVAGIHMILIYLCRFLLRRSRNVLFTVSNSLRFRLKVLTVLLYICLSVMLFYLFGSIMPLPPWGLVVGWVMALIAVELAYAFIFPYSFRYIADNDDDKMVILPV |
| IRIS_313-11757 | MLHHLKELAAVAGIHMILIYLCRFLLRRSRNVLFTVSNSLRFRLKVLTVLLYICLSVMLFYLFGSIMPLPPWGLVVGWVMALIAVELAYAFIFPYSFRYIADNDDDKMVILPV |
| IRIS_313-11758 | MLHHLKELAAVAGIHMILIYLCRFLLRRSRNVLFTVSNSLRFRLKVLTVLLYICLSVMLFYLFGSIMPLPPWGLVVGWVMALIAVELAYAFIFPYSFRYIADNDDDKMVILPV |
| IRIS_313-11759 | MLHHLKELAAVAGIHMILIYLCRFLLRRSRNVLFTVSNSLRFRLKVLTVLLYICLSVMLFYLFGSIMPLPPWGLVVGWVMALIAVELAYAFIFPYSFRYIADNDDDKMVILPV |
| IRIS_313-11760 | MLHHLKELAAVAGIHMILIYLCRFLLRRSRNVLFTVSNSLRFRLKVLTVLLYICLSVMLFYLFGSIMPLPPWGLVVGWVMALIAVELAYAFIFPYSFRYIADNDDDKMVILPV |
| IRIS_313-11761 | MLHHLKELAAVAGIHMILIYLCRFLLRRSRNVLFTVSNSLRFRLKVLTVLLYICLSVMLFYLFGSIMPLPPWGLVVGWVMALIAVELAYAFIFPYSFRYIADNDDDKMVILPV |
| IRIS_313-11762 | MLHHLKELAAVAGIHMILIYLCRFLLRRSRNVLFTVSNSLRFRLKVLTVLLYICLSVMLFYLFGSIMPLPPWGLVVGWVMALIAVELAYAFIFPYSFRYIADNDDDKMVILPV |
| IRIS_313-11763 | MLHHLKELAAVAGIHMILIYLCRFLLRRSRNVLFTVSNSLRFRLKVLTVLLYICLSVMLFYLFGSIMPLPPWGLVVGWVMALIAVELAYAFIFPYSFRYIADNDDDKMVILPV |
| IRIS_313-11764 | MLHHLKELAAVAGIHMILIYLCRFLLRRSRNVLFTVSNSLRFRLKVLTVLLYICLSVMLFYLFGSIMPLPPWGLVVGWVMALIAVELAYAFIFPYSFRYIADNDDDKMVILPV |
| IRIS_313-11765 | MLHHLKELAAVAGIHMILIYLCRFLLRRSRNVLFTVSNSLRFRLKVLTVLLYICLSVMLFYLFGSIMPLPPWGLVVGWVMALIAVELAYAFIFPYSFRYIADNDDDKMVILPV |
| IRIS_313-11766 | MLHHLKELAAVAGIHMILIYLCRFLLRRSRNVLFTVSNSLRFRLKVLTVLLYICLSVMLFYLFGSIMPLPPWGLVVGWVMALIAVELAYAFIFPYSFRYIADNDDDKMVILPV |
| IRIS_313-11767 | MLHHLKELAAVAGIHMILIYLCRFLLRRSRNVLFTVSNSLRFRLKVLTVLLYICLSVMLFYLFGSIMPLPPWGLVVGWVMALIAVELAYAFIFPYSFRYIADNDDDKMVILPV |
| IRIS_313-11772 | MLHHLKELAAVAGIHMILIYLCRFLLRRSRNVLFTVSNSLRFRLKVLTVLLYICLSVMLFYLFGSIMPLPPWGLVVGWVMALIAVELAYAFIFPYSFRYIADNDDDKMVILPV |

| | |
|---|---|
| IRIS_313-11773 | MLHHLKELAAVAGIHMILIYLCRFLLRRSRNVLFTVSNSLRFRLKVLTVLLYICLSVMLFYLFGSIMPLPPWGLVVGWVMALIAVELAYAFIFPYSFRYIADNDDDKMVILPV |
| IRIS_313-11779 | MLHHLKELAAVAGIHMILIYLCRFLLRRSRNVLFTVSNSLRFRLKVLTVLLYICLSVMLFYLFGSIMPLPPWGLVVGWVMALIAVELAYAFIFPYSFRYIADNDDDKMVILPV |
| IRIS_313-11780 | MLHHLKELAAVAGIHMILIYLCRFLLRRSRNVLFTVSNSLRFRLKVLTVLLYICLSVMLFYLFGSIMPLPPWGLVVGWVMALIAVELAYAFIFPYSFRYIADNDDDKMVILPV |
| IRIS_313-11782 | MLHHLKELAAVAGIHMILIYLCRFLLRRSRNVLFTVSNSLRFRLKVLTVLLYICLSVMLFYLFGSIMPLPPWGLVVGWVMALIAVELAYAFIFPYSFRYIADNDDDKMVILPV |
| IRIS_313-11783 | MLHHLKELAAVAGIHMILIYLCRFLLRRSRNVLFTVSNSLRFRLKVLTVLLYICLSVMLFYLFGSIMPLPPWGLVVGWVMALIAVELAYAFIFPYSFRYIADNDDDKMVILPV |
| IRIS_313-11784 | MLHHLKELAAVAGIHMILIYLCRFLLRRSRNVLFTVSNSLRFRLKVLTVLLYICLSVMLFYLFGSIMPLPPWGLVVGWVMALIAVELAYAFIFPYSFRYIADNDDDKMVILPV |
| IRIS_313-11785 | MLHHLKELAAVAGIHMILIYLCRFLLRRSRNVLFTVSNSLRFRLKVLTVLLYICLSVMLFYLFGSIMPLPPWGLVVGWVMALIAVELAYAFIFPYSFRYIADNDDDKMVILPV |
| IRIS_313-11786 | MLHHLKELAAVAGIHMILIYLCRFLLRRSRNVLFTVSNSLRFRLKVLTVLLYICLSVMLFYLFGSIMPLPPWGLVVGWVMALIAVELAYAFIFPYSFRYIADNDDDKMVILPV |
| IRIS_313-11787 | MLHHLKELAAVAGIHMILIYLCRFLLRRSRNVLFTVSNSLRFRLKVLTVLLYICLSVMLFYLFGSIMPLPPWGLVVGWVMALIAVELAYAFIFPYSFRYIADNDDDKMVILPV |
| IRIS_313-11788 | MLHHLKELAAVAGIHMILIYLCRFLLRRSRNVLFTVSNSLRFRLKVLTVLLYICLSVMLFYLFGSIMPLPPWGLVVGWVMALIAVELAYAFIFPYSFRYIADNDDDKMVILPV |
| IRIS_313-11789 | MLHHLKELAAVAGIHMILIYLCRFLLRRSRNVLFTVSNSLRFRLKVLTVLLYICLSVMLFYLFGSIMPLPPWGLVVGWVMALIAVELAYAFIFPYSFRYIADNDDDKMVILPV |
| IRIS_313-11790 | MLHHLKELAAVAGIHMILIYLCRFLLRRSRNVLFTVSNSLRFRLKVLTVLLYICLSVMLFYLFGSIMPLPPWGLVVGWVMALIAVELAYAFIFPYSFRYIADNDDDKMVILPV |
| IRIS_313-11791 | MLHHLKELAAVAGIHMILIYLCRFLLRRSRNVLFTVSNSLRFRLKVLTVLLYICLSVMLFYLFGSIMPLPPWGLVVGWVMALIAVELAYAFIFPYSFRYIADNDDDKMVILPV |
| IRIS_313-11792 | MLHHLKELAAVAGIHMILIYLCRFLLRRSRNVLFTVSNSLRFRLKVLTVLLYICLSVMLFYLFGSIMPLPPWGLVVGWVMALIAVELAYAFIFPYSFRYIADNDDDKMVILPV |
| IRIS_313-11793 | MLHHLKELAAVAGIHMILIYLCRFLLRRSRNVLFTVSNSLRFRLKVLTVLLYICLSVMLFYLFGSIMPLPPWGLVVGWVMALIAVELAYAFIFPYSFRYIADNDDDKMVILPV |
| IRIS_313-11794 | MLHHLKELAAVAGIHMILIYLCRFLLRRSRNVLFTVSNSLRFRLKVLTVLLYICLSVMLFYLFGSIMPLPPWGLVVGWVMALIAVELAYAFIFPYSFRYIADNDDDKMVILPV |
| IRIS_313-11795 | MLHHLKELAAVAGIHMILIYLCRFLLRRSRNVLFTVSNSLRFRLKVLTVLLYICLSVMLFYLFGSIMPLPPWGLVVGWVMALIAVELAYAFIFPYSFRYIADNDDDKMVILPV |
| IRIS_313-11796 | MLHHLKELAAVAGIHMILIYLCRFLLRRSRNVLFTVSNSLRFRLKVLTVLLYICLSVMLFYLFGSIMPLPPWGLVVGWVMALIAVELAYAFIFPYSFRYIADNDDDKMVILPV |
| IRIS_313-11797 | MLHHLKELAAVAGIHMILIYLCRFLLRRSRNVLFTVSNSLRFRLKVLTVLLYICLSVMLFYLFGSIMPLPPWGLVVGWVMALIAVELAYAFIFPYSFRYIADNDDDKMVILPV |
| IRIS_313-11798 | MLHHLKELAAVAGIHMILIYLCRFLLRRSRNVLFTVSNSLRFRLKVLTVLLYICLSVMLFYLFGSIMPLPPWGLVVGWVMALIAVELAYAFIFPYSFRYIADNDDDKMVILPV |
| IRIS_313-11799 | MLHHLKELAAVAGIHMILIYLCRFLLRRSRNVLFTVSNSLRFRLKVLTVLLYICLSVMLFYLFGSIMPLPPWGLVVGWVMALIAVELAYAFIFPYSFRYIADNDDDKMVILPV |
| IRIS_313-11800 | MLHHLKELAAVAGIHMILIYLCRFLLRRSRNVLFTVSNSLRFRLKVLTVLLYICLSVMLFYLFGSIMPLPPWGLVVGWVMALIAVELAYAFIFPYSFRYIADNDDDKMVILPV |
| IRIS_313-11801 | MLHHLKELAAVAGIHMILIYLCRFLLRRSRNVLFTVSNSLRFRLKVLTVLLYICLSVMLFYLFGSIMPLPPWGLVVGWVMALIAVELAYAFIFPYSFRYIADNDDDKMVILPV |
| IRIS_313-11802 | MLHHLKELAAVAGIHMILIYLCRFLLRRSRNVLFTVSNSLRFRLKVLTVLLYICLSVMLFYLFGSIMPLPPWGLVVGWVMALIAVELAYAFIFPYSFRYIADNDDDKMVILPV |
| IRIS_313-11803 | MLHHLKELAAVAGIHMILIYLCRFLLRRSRNVLFTVSNSLRFRLKVLTVLLYICLSVMLFYLFGSIMPLPPWGLVVGWVMALIAVELAYAFIFPYSFRYIADNDDDKMVILPV |
| IRIS_313-11804 | MLHHLKELAAVAGIHMILIYLCRFLLRRSRNVLFTVSNSLRFRLKVLTVLLYICLSVMLFYLFGSIMPLPPWGLVVGWVMALIAVELAYAFIFPYSFRYIADNDDDKMVILPV |
| IRIS_313-11805 | MLHHLKELAAVAGIHMILIYLCRFLLRRSRNVLFTVSNSLRFRLKVLTVLLYICLSVMLFYLFGSIMPLPPWGLVVGWVMALIAVELAYAFIFPYSFRYIADNDDDKMVILPV |
| IRIS_313-11806 | MLHHLKELAAVAGIHMILIYLCRFLLRRSRNVLFTVSNSLRFRLKVLTVLLYICLSVMLFYLFGSIMPLPPWGLVVGWVMALIAVELAYAFIFPYSFRYIADNDDDKMVILPV |
| IRIS_313-11807 | MLHHLKELAAVAGIHMILIYLCRFLLRRSRNVLFTVSNSLRFRLKVLTVLLYICLSVMLFYLFGSIMPLPPWGLVVGWVMALIAVELAYAFIFPYSFRYIADNDDDKMVILPV |
| IRIS_313-11808 | MLHHLKELAAVAGIHMILIYLCRFLLRRSRNVLFTVSNSLRFRLKVLTVLLYICLSVMLFYLFGSIMPLPPWGLVVGWVMALIAVELAYAFIFPYSFRYIADNDDDKMVILPV |
| IRIS_313-11809 | MLHHLKELAAVAGIHMILIYLCRFLLRRSRNVLFTVSNSLRFRLKVLTVLLYICLSVMLFYLFGSIMPLPPWGLVVGWVMALIAVELAYAFIFPYSFRYIADNDDDKMVILPV |
| IRIS_313-11810 | MLHHLKELAAVAGIHMILIYLCRFLLRRSRNVLFTVSNSLRFRLKVLTVLLYICLSVMLFYLFGSIMPLPPWGLVVGWVMALIAVELAYAFIFPYSFRYIADNDDDKMVILPV |
| IRIS_313-11811 | MLHHLKELAAVAGIHMILIYLCRFLLRRSRNVLFTVSNSLRFRLKVLTVLLYICLSVMLFYLFGSIMPLPPWGLVVGWVMALIAVELAYAFIFPYSFRYIADNDDDKMVILPV |
| IRIS_313-11812 | MLHHLKELAAVAGIHMILIYLCRFLLRRSRNVLFTVSNSLRFRLKVLTVLLYICLSVMLFYLFGSIMPLPPWGLVVGWVMALIAVELAYAFIFPYSFRYIADNDDDKMVILPV |
| IRIS_313-11813 | MLHHLKELAAVAGIHMILIYLCRFLLRRSRNVLFTVSNSLRFRLKVLTVLLYICLSVMLFYLFGSIMPLPPWGLVVGWVMALIAVELAYAFIFPYSFRYIADNDDDKMVILPV |
| IRIS_313-11814 | MLHHLKELAAVAGIHMILIYLCRFLLRRSRNVLFTVSNSLRFRLKVLTVLLYICLSVMLFYLFGSIMPLPPWGLVVGWVMALIAVELAYAFIFPYSFRYIADNDDDKMVILPV |
| IRIS_313-11815 | MLHHLKELAAVAGIHMILIYLCRFLLRRSRNVLFTVSNSLRFRLKVLTVLLYICLSVMLFYLFGSIMPLPPWGLVVGWVMALIAVELAYAFIFPYSFRYIADNDDDKMVILPV |
| IRIS_313-11816 | MLHHLKELAAVAGIHMILIYLCRFLLRRSRNVLFTVSNSLRFRLKVLTVLLYICLSVMLFYLFGSIMPLPPWGLVVGWVMALIAVELAYAFIFPYSFRYIADNDDDKMVILPV |
| IRIS_313-11817 | MLHHLKELAAVAGIHMILIYLCRFLLRRSRNVLFTVSNSLRFRLKVLTVLLYICLSVMLFYLFGSIMPLPPWGLVVGWVMALIAVELAYAFIFPYSFRYIADNDDDKMVILPV |
| IRIS_313-11818 | MLHHLKELAAVAGIHMILIYLCRFLLRRSRNVLFTVSNSLRFRLKVLTVLLYICLSVMLFYLFGSIMPLPPWGLVVGWVMALIAVELAYAFIFPYSFRYIADNDDDKMVILPV |
| IRIS_313-11819 | MLHHLKELAAVAGIHMILIYLCRFLLRRSRNVLFTVSNSLRFRLKVLTVLLYICLSVMLFYLFGSIMPLPPWGLVVGWVMALIAVELAYAFIFPYSFRYIADNDDDKMVILPV |
| IRIS_313-11820 | MLHHLKELAAVAGIHMILIYLCRFLLRRSRNVLFTVSNSLRFRLKVLTVLLYICLSVMLFYLFGSIMPLPPWGLVVGWVMALIAVELAYAFIFPYSFRYIADNDDDKMVILPV |
| IRIS_313-11821 | MLHHLKELAAVAGIHMILIYLCRFLLRRSRNVLFTVSNSLRFRLKVLTVLLYICLSVMLFYLFGSIMPLPPWGLVVGWVMALIAVELAYAFIFPYSFRYIADNDDDKMVILPV |
| IRIS_313-11822 | MLHHLKELAAVAGIHMILIYLCRFLLRRSRNVLFTVSNSLRFRLKVLTVLLYICLSVMLFYLFGSIMPLPPWGLVVGWVMALIAVELAYAFIFPYSFRYIADNDDDKMVILPV |
| IRIS_313-11823 | MLHHLKELAAVAGIHMILIYLCRFLLRRSRNVLFTVSNSLRFRLKVLTVLLYICLSVMLFYLFGSIMPLPPWGLVVGWVMALIAVELAYAFIFPYSFRYIADNDDDKMVILPV |
| IRIS_313-11824 | MLHHLKELAAVAGIHMILIYLCRFLLRRSRNVLFTVSNSLRFRLKVLTVLLYICLSVMLFYLFGSIMPLPPWGLVVGWVMALIAVELAYAFIFPYSFRYIADNDDDKMVILPV |
| IRIS_313-11825 | MLHHLKELAAVAGIHMILIYLCRFLLRRSRNVLFTVSNSLRFRLKVLTVLLYICLSVMLFYLFGSIMPLPPWGLVVGWVMALIAVELAYAFIFPYSFRYIADNDDDKMVILPV |
| IRIS_313-11826 | MLHHLKELAAVAGIHMILIYLCRFLLRRSRNVLFTVSNSLRFRLKVLTVLLYICLSVMLFYLFGSIMPLPPWGLVVGWVMALIAVELAYAFIFPYSFRYIADNDDDKMVILPV |
| IRIS_313-11827 | MLHHLKELAAVAGIHMILIYLCRFLLRRSRNVLFTVSNSLRFRLKVLTVLLYICLSVMLFYLFGSIMPLPPWGLVVGWVMALIAVELAYAFIFPYSFRYIADNDDDKMVILPV |
| IRIS_313-11828 | MLHHLKELAAVAGIHMILIYLCRFLLRRSRNVLFTVSNSLRFRLKVLTVLLYICLSVMLFYLFGSIMPLPPWGLVVGWVMALIAVELAYAFIFPYSFRYIADNDDDKMVILPV |
| IRIS_313-11829 | MLHHLKELAAVAGIHMILIYLCRFLLRRSRNVLFTVSNSLRFRLKVLTVLLYICLSVMLFYLFGSIMPLPPWGLVVGWVMALIAVELAYAFIFPYSFRYIADNDDDKMVILPV |
| IRIS_313-11830 | MLHHLKELAAVAGIHMILIYLCRFLLRRSRNVLFTVSNSLRFRLKVLTVLLYICLSVMLFYLFGSIMPLPPWGLVVGWVMALIAVELAYAFIFPYSFRYIADNDDDKMVILPV |
| IRIS_313-11831 | MLHQLKELAAVAGIHMILIYLCRFLLRRSRNVLFTVSNSLRFRLKVLTVLLYICLLVMLFYLFGSIMPLPPWGLVVGWVMALIAVELAYAFIFPYSFRYIADNDDDKMVILPV |
| IRIS_313-11832 | MLHHLKELAAVAGIHMILIYLCRFLLRRSRNVLFTVSNSLRFRLKVLTVLLYICLSVMLFYLFGSIMPLPPWGLVVGWVMALIAVELAYAFIFPYSFRYIADNDDDKMVILPV |
| IRIS_313-11833 | MLHHLKELAAVAGIHMILIYLCRFLLRRSRNVLFTVSNSLRFRLKVLTVLLYICLSVMLFYLFGSIMPLPPWGLVVGWVMALIAVELAYAFIFPYSFRYIADNDDDKMVILPV |
| IRIS_313-11834 | MLHHLKELAAVAGIHMILIYLCRFLLRRSRNVLFTVSNSLRFRLKVLTVLLYICLSVMLFYLFGSIMPLPPWGLVVGWVMALIAVELAYAFIFPYSFRYIADNDDDKMVILPV |
| IRIS_313-11835 | MLHHLKELAAVAGIHMILIYLCRFLLRRSRNVLFTVSNSLRFRLKVLTVLLYICLSVMLFYLFGSIMPLPPWGLVVGWVMALIAVELAYAFIFPYSFRYIADNDDDKMVILPV |
| IRIS_313-11836 | MLHHLKELAAVAGIHMILIYLCRFLLRRSRNVLFTVSNSLRFRLKVLTVLLYICLSVMLFYLFGSIMPLPPWGLVVGWVMALIAVELAYAFIFPYSFRYIADNDDDKMVILPV |
| IRIS_313-11837 | MLHHLKELAAVAGIHMILIYLCRFLLRRSRNVLFTVSNSLRFRLKVLTVLLYICLSVMLFYLFGSIMPLPPWGLVVGWVMALIAVELAYAFIFPYSFRYIADNDDDKMVILPV |
| IRIS_313-11838 | MLHHLKELAAVAGIHMILIYLCRFLLRRSRNVLFTVSNSLRFRLKVLTVLLYICLSVMLFYLFGSIMPLPPWGLVVGWVMALIAVELAYAFIFPYSFRYIADNDDDKMVILPV |
| IRIS_313-11839 | MLHHLKELAAVAGIHMILIYLCRFLLRRSRNVLFTVSNSLRFRLKVLTVLLYICLSVMLFYLFGSIMPLPPWGLVVGWVMALIAVELAYAFIFPYSFRYIADNDDDKMVILPV |
| IRIS_313-11840 | MLHHLKELAAVAGIHMILIYLCRFLLRRSRNVLFTVSNSLRFRLKVLTVLLYICLSVMLFYLFGSIMPLPPWGLVVGWVMALIAVELAYAFIFPYSFRYIADNDDDKMVILPV |
| IRIS_313-11841 | MLHHLKELAAVAGIHMILIYLCRFLLRRSRNVLFTVSNSLRFRLKVLTVLLYICLSVMLFYLFGSIMPLPPWGLVVGWVMALIAVELAYAFIFPYSFRYIADNDDDKMVILPV |
| IRIS_313-11842 | MLHHLKELAAVAGIHMILIYLCRFLLRRSRNVLFTVSNSLRFRLKVLTVLLYICLSVMLFYLFGSIMPLPPWGLVVGWVMALIAVELAYAFIFPYSFRYIADNDDDKMVILPV |
| IRIS_313-11843 | MLHHLKELAAVAGIHMILIYLCRFLLRRSRNVLFTVSNSLRFRLKVLTVLLYICLSVMLFYLFGSIMPLPPWGLVVGWVMALIAVELAYAFIFPYSFRYIADNDDDKMVILPV |
| IRIS_313-11844 | MLHHLKELAAVAGIHMILIYLCRFLLRRSRNVLFTVSNSLRFRLKVLTVLLYICLSVMLFYLFGSIMPLPPWGLVVGWVMALIAVELAYAFIFPYSFRYIADNDDDKMVILPV |
| IRIS_313-11845 | MLHHLKELAAVAGIHMILIYLCRFLLRRSRNVLFTVSNSLRFRLKVLTVLLYICLSVMLFYLFGSIMPLPPWGLVVGWVMALIAVELAYAFIFPYSFRYIADNDDDKMVILPV |
| IRIS_313-11846 | MLHHLKELAAVAGIHMILIYLCRFLLRRSRNVLFTVSNSLRFRLKVLTVLLYICLSVMLFYLFGSIMPLPPWGLVVGWVMALIAVELAYAFIFPYSFRYIADNDDDKMVILPV |
| IRIS_313-11848 | MLHHLKELAAVAGIHMILIYLCRFLLRRSRNVLFTVSNSLRFRLKVLTVLLYICLSVMLFYLFGSIMPLPPWGLVVGWVMALIAVELAYAFIFPYSFRYIADNDDDKMVILPV |
| IRIS_313-11849 | MLHHLKELAAVAGIHMILIYLCRFLLRRSRNVLFTVSNSLRFRLKVLTVLLYICLSVMLFYLFGSIMPLPPWGLVVGWVMALIAVELAYAFIFPYSFRYIADNDDDKMVILPV |
| IRIS_313-11850 | MLHHLKELAAVAGIHMILIYLCRFLLRRSRNVLFTVSNSLRFRLKVLTVLLYICLSVMLFYLFGSIMPLPPWGLVVGWVMALIAVELAYAFIFPYSFRYIADNDDDKMVILPV |
| IRIS_313-11851 | MLHHLKELAAVAGIHMILIYLCRFLLRRSRNVLFTVSNSLRFRLKVLTVLLYICLSVMLFYLFGSIMPLPPWGLVVGWVMALIAVELAYAFIFPYSFRYIADNDDDKMVILPV |
| IRIS_313-11852 | MLHHLKELAAVAGIHMILIYLCRFLLRRSRNVLFTVSNSLRFRLKVLTVLLYICLSVMLFYLFGSIMPLPPWGLVVGWVMALIAVELAYAFIFPYSFRYIADNDDDKMVILPV |
| IRIS_313-11853 | MLHHLKELAAVAGIHMILIYLCRFLLRRSRNVLFTVSNSLRFRLKVLTVLLYICLSVMLFYLFGSIMPLPPWGLVVGWVMALIAVELAYAFIFPYSFRYIADNDDDKMVILPV |
| IRIS_313-11854 | MLHHLKELAAVAGIHMILIYLCRFLLRRSRNVLFTVSNSLRFRLKVLTVLLYICLSVMLFYLFGSIMPLPPWGLVVGWVMALIAVELAYAFIFPYSFRYIADNDDDKMVILPV |
| IRIS_313-11855 | MLHHLKELAAVAGIHMILIYLCRFLLRRSRNVLFTVSNSLRFRLKVLTVLLYICLSVMLFYLFGSIMPLPPWGLVVGWVMALIAVELAYAFIFPYSFRYIADNDDDKMVILPV |
| IRIS_313-11856 | MLHHLKELAAVAGIHMILIYLCRFLLRRSRNVLFTVSNSLRFRLKVLTVLLYICLSVMLFYLFGSIMPLPPWGLVVGWVMALIAVELAYAFIFPYSFRYIADNDDDKMVILPV |
| IRIS_313-11857 | MLHHLKELAAVAGIHMILIYLCRFLLRRSRNVLFTVSNSLRFRLKVLTVLLYICLSVMLFYLFGSIMPLPPWGLVVGWVMALIAVELAYAFIFPYSFRYIADNDDDKMVILPV |
| IRIS_313-11858 | MLHHLKELAAVAGIHMILIYLCRFLLRRSRNVLFTVSNSLRFRLKVLTVLLYICLSVMLFYLFGSIMPLPPWGLVVGWVMALIAVELAYAFIFPYSFRYIADNDDDKMVILPV |
| IRIS_313-11859 | MLHHLKELAAVAGIHMILIYLCRFLLRRSRNVLFTVSNSLRFRLKVLTVLLYICLSVMLFYLFGSIMPLPPWGLVVGWVMALIAVELAYAFIFPYSFRYIADNDDDKMVILPV |
| IRIS_313-11860 | MLHHLKELAAVAGIHMILIYLCRFLLRRSRNVLFTVSNSLRFRLKVLTVLLYICLSVMLFYLFGSIMPLPPWGLVVGWVMALIAVELAYAFIFPYSFRYIADNDDDKMVILPV |
| IRIS_313-11861 | MLHHLKELAAVAGIHMILIYLCRFLLRRSRNVLFTVSNSLRFRLKVLTVLLYICLSVMLFYLFGSIMPLPPWGLVVGWVMALIAVELAYAFIFPYSFRYIADNDDDKMVILPV |
| IRIS_313-11862 | MLHHLKELAAVAGIHMILIYLCRFLLRRSRNVLFTVSNSLRFRLKVLTVLLYICLSVMLFYLFGSIMPLPPWGLVVGWVMALIAVELAYAFIFPYSFRYIADNDDDKMVILPV |
| IRIS_313-11863 | MLHHLKELAAVAGIHMILIYLCRFLLRRSRNVLFTVSNSLRFRLKVLTVLLYICLSVMLFYLFGSIMPLPPWGLVVGWVMALIAVELAYAFIFPYSFRYIADNDDDKMVILPV |
| IRIS_313-11865 | MLHHLKELAAVAGIHMILIYLCRFLLRRSRNVLFTVSNSLRFRLKVLTVLLYICLSVMLFYLFGSIMPLPPWGLVVGWVMALIAVELAYAFIFPYSFRYIADNDDDKMVILPV |
| IRIS_313-11866 | MLHHLKELAAVAGIHMILIYLCRFLLRRSRNVLFTVSNSLRFRLKVLTVLLYICLSVMLFYLFGSIMPLPPWGLVVGWVMALIAVELAYAFIFPYSFRYIADNDDDKMVILPV |
| IRIS_313-11867 | MLHHLKELAAVAGIHMILIYLCRFLLRRSRNVLFTVSNSLRFRLKVLTVLLYICLSVMLFYLFGSIMPLPPWGLVVGWVMALIAVELAYAFIFPYSFRYIADNDDDKMVILPV |
| IRIS_313-11868 | MLHHLKELAAVAGIHMILIYLCRFLLRRSRNVLFTVSNSLRFRLKVLTVLLYICLSVMLFYLFGSIMPLPPWGLVVGWVMALIAVELAYAFIFPYSFRYIADNDDDKMVILPV |
| IRIS_313-11869 | MLHHLKELAAVAGIHMILIYLCRFLLRRSRNVLFTVSNSLRFRLKVLTVLLYICLSVMLFYLFGSIMPLPPWGLVVGWVMALIAVELAYAFIFPYSFRYIADNDDDKMVILPV |
| IRIS_313-11870 | MLHHLKELAAVAGIHMILIYLCRFLLRRSRNVLFTVSNSLRFRLKVLTVLLYICLSVMLFYLFGSIMPLPPWGLVVGWVMALIAVELAYAFIFPYSFRYIADNDDDKMVILPV |
| IRIS_313-11871 | MLHHLKELAAVAGIHMILIYLCRFLLRRSRNVLFTVSNSLRFRLKVLTVLLYICLSVMLFYLFGSIMPLPPWGLVVGWVMALIAVELAYAFIFPYSFRYIADNDDDKMVILPV |
| IRIS_313-11872 | MLHHLKELAAVAGIHMILIYLCRFLLRRSRNVLFTVSNSLRFRLKVLTVLLYICLSVMLFYLFGSIMPLPPWGLVVGWVMALIAVELAYAFIFPYSFRYIADNDDDKMVILPV |
| IRIS_313-11875 | MLHHLKELAAVAGIHMILIYLCRFLLRRSRNVLFTVSNSLRFRLKVLTVLLYICLSVMLFYLFGSIMPLPPWGLVVGWVMALIAVELAYAFIFPYSFRYIADNDDDKMVILPV |
| IRIS_313-11876 | MLHHLKELAAVAGIHMILIYLCRFLLRRSRNVLFTVSNSLRFRLKVLTVLLYICLSVMLFYLFGSIMPLPPWGLVVGWVMALIAVELAYAFIFPYSFRYIADNDDDKMVILPV |
| IRIS_313-11877 | MLHHLKELAAVAGIHMILIYLCRFLLRRSRNVLFTVSNSLRFRLKVLTVLLYICLSVMLFYLFGSIMPLPPWGLVVGWVMALIAVELAYAFIFPYSFRYIADNDDDKMVILPV |
| IRIS_313-11878 | MLHHLKELAAVAGIHMILIYLCRFLLRRSRNVLFTVSNSLRFRLKVLTVLLYICLSVMLFYLFGSIMPLPPWGLVVGWVMALIAVELAYAFIFPYSFRYIADNDDDKMVILPV |
| IRIS_313-11881 | MLHHLKELAAVAGIHMILIYLCRFLLRRSRNVLFTVSNSLRFRLKVLTVLLYICLSVMLFYLFGSIMPLPPWGLVVGWVMALIAVELAYAFIFPYSFRYIADNDDDKMVILPV |
| IRIS_313-11882 | MLHHLKELAAVAGIHMILIYLCRFLLRRSRNVLFTVSNSLRFRLKVLTVLLYICLSVMLFYLFGSIMPLPPWGLVVGWVMALIAVELAYAFIFPYSFRYIADNDDDKMVILPV |
| IRIS_313-11884 | MLHHLKELAAVAGIHMILIYLCRFLLRRSRNVLFTVSNSLRFRLKVLTVLLYICLSVMLFYLFGSIMPLPPWGLVVGWVMALIAVELAYAFIFPYSFRYIADNDDDKMVILPV |
| IRIS_313-11885 | MLHHLKELAAVAGIHMILIYLCRFLLRRSRNVLFTVSNSLRFRLKVLTVLLYICLSVMLFYLFGSIMPLPPWGLVVGWVMALIAVELAYAFIFPYSFRYIADNDDDKMVILPV |
| IRIS_313-11886 | MLHHLKELAAVAGIHMILIYLCRFLLRRSRNVLFTVSNSLRFRLKVLTVLLYICLSVMLFYLFGSIMPLPPWGLVVGWVMALIAVELAYAFIFPYSFRYIADNDDDKMVILPV |
| IRIS_313-11887 | MLHHLKELAAVAGIHMILIYLCRFLLRRSRNVLFTVSNSLRFRLKVLTVLLYICLSVMLFYLFGSIMPLPPWGLVVGWVMALIAVELAYAFIFPYSFRYIADNDDDKMVILPV |
| IRIS_313-11888 | MLHHLKELAAVAGIHMILIYLCRFLLRRSRNVLFTVSNSLRFRLKVLTVLLYISLSVMLLYLFGSIMPLPPWGLVVGWVMALIAVELAYAFIFPYSFRYIADNDDDKMVILPV |
| IRIS_313-11889 | MLHHLKELAAVAGIHMILIYLCRFLLRRSRNVLFTVSNSLRFRLKVLTVLLYICLSVMLFYLFGSIMPLPPWGLVVGWVMALIAVELAYAFIFPYSFRYIADNDDDKMVILPV |
| IRIS_313-11890 | MLHHLKELAAVAGIHMILIYLCRFLLRRSRNVLFTVSNSLRFRLKVLTVLLYICLSVMLFYLFGSIMPLPPWGLVVGWVMALIAVELAYAFIFPYSFRYIADNDDDKMVILPV |
| IRIS_313-11892 | MLHHLKELAAVAGIHMILIYLCRFLLRRSRNVLFTVSNSLRFRLKVLTVLLYICLSVMLFYLFGSIMPLPPWGLVVGWVMALIAVELAYAFIFPYSFRYIADNDDDKMVILPV |
| IRIS_313-11893 | MLHHLKELAAVAGIHMILIYLCRFLLRRSRNVLFTVSNSLRFRLKVLTVLLYICLSVMLFYLFGSIMPLPPWGLVVGWVMALIAVELAYAFIFPYSFRYIADNDDDKMVILPV |
| IRIS_313-11894 | MLHHLKELAAVAGIHMILIYLCRFLLRRSRNVLFTVSNSLRFRLKVLTVLLYICLSVMLFYLFGSIMPLPPWGLVVGWVMALIAVELAYAFIFPYSFRYIADNDDDKMVILPV |
| IRIS_313-11895 | MLHHLKELAAVAGIHMILIYLCRFLLRRSRNVLFTVSNSLRFRLKVLTVLLYICLSVMLFYLFGSIMPLPPWGLVVGWVMALIAVELAYAFIFPYSFRYIADNDDDKMVILPV |
| IRIS_313-11896 | MLHHLKELAAVAGIHMILIYLCRFLLRRSRNVLFTVSNSLRFRLKVLTVLLYICLSVMLFYLFGSIMPLPPWGLVVGWVMALIAVELAYAFIFPYSFRYIADNDDDKMVILPV |
| IRIS_313-11897 | MLHQLKELAAVAGIHMILIYLCRFLLRRSRNVLFTVSNSLRFRLKVLTVLLYICLLVMLFYLFGSIMPLPPWGLVVGWVMALIAVELAYAFIFPYSFRYIADNDDDKMVILPV |
| IRIS_313-11898 | MLHHLKELAAVAGIHMILIYLCRFLLRRSRNVLFTVSNSLRFRLKVLTVLLYICLSVMLFYLFGSIMPLPPWGLVVGWVMALIAVELAYAFIFPYSFRYIADNDDDKMVILPV |
| IRIS_313-11899 | MLHHLKELAAVAGIHMILIYLCRFLLRRSRNVLFTVSNSLRFRLKVLTVLLYICLSVMLFYLFGSIMPLPPWGLVVGWVMALIAVELAYAFIFPYSFRYIADNDDDKMVILPV |
| IRIS_313-11900 | MLHHLKELAAVAGIHMILIYLCRFLLRRSRNVLFTVSNSLRFRLKVLTVLLYICLSVMLFYLFGSIMPLPPWGLVVGWVMALIAVELAYAFIFPYSFRYIADNDDDKMVILPV |
| IRIS_313-11901 | MLHHLKELAAVAGIHMILIYLCRFLLRRSRNVLFTVSNSLRFRLKVLTVLLYICLSVMLFYLFGSIMPLPPWGLVVGWVMALIAVELAYAFIFPYSFRYIADNDDDKMVILPV |
| IRIS_313-11902 | MLHHLKELAAVAGIHMILIYLCRFLLRRSRNVLFTVSNSLRFRLKVLTVLLYICLSVMLFYLFGSIMPLPPWGLVVGWVMALIAVELAYAFIFPYSFRYIADNDDDKMVILPV |
| IRIS_313-11903 | MLHHLKELAAVAGIHMILIYLCRFLLRRSRNVLFTVSNSLRFRLKVLTVLLYICLSVMLFYLFGSIMPLPPWGLVVGWVMALIAVELAYAFIFPYSFRYIADNDDDKMVILPV |
| IRIS_313-11904 | MLHHLKELAAVAGIHMILIYLCRFLLRRSRNVLFTVSNSLRFRLKVLTVLLYICLSVMLFYLFGSIMPLPPWGLVVGWVMALIAVELAYAFIFPYSFRYIADNDDDKMVILPV |
| IRIS_313-11905 | MLHHLKELAAVAGIHMILIYLCRFLLRRSRNVLFTVSNSLRFRLKVLTVLLYICLSVMLFYLFGSIMPLPPWGLVVGWVMALIAVELAYAFIFPYSFRYIADNDDDKMVILPV |
| IRIS_313-11906 | MLHHLKELAAVAGIHMILIYLCRFLLRRSRNVLFTVSNSLRFRLKVLTVLLYICLSVMLFYLFGSIMPLPPWGLVVGWVMALIAVELAYAFIFPYSFRYIADNDDDKMVILPV |
| IRIS_313-11907 | MLHHLKELAAVAGIHMILIYLCRFLLRRSRNVLFTVSNSLRFRLKVLTVLLYICLSVMLFYLFGSIMPLPPWGLVVGWVMALIAVELAYAFIFPYSFRYIADNDDDKMVILPV |
| IRIS_313-11908 | MLHHLKELAAVAGIHMILIYLCRFLLRRSRNVLFTVSNSLRFRLKVLTVLLYICLSVMLFYLFGSIMPLPPWGLVVGWVMALIAVELAYAFIFPYSFRYIADNDDDKMVILPV |
| IRIS_313-11909 | MLHHLKELAAVAGIHMILIYLCRFLLRRSRNVLFTVSNSLRFRLKVLTVLLYICLSVMLFYLFGSIMPLPPWGLVVGWVMALIAVELAYAFIFPYSFRYIADNDDDKMVILPV |

| | |
|---|---|
| IRIS_313-11910 | MLHHLKELAAVAGIHMILIYLCRFLLRRSRNVLFTVSNSLRFRLKVLTVLLYICLSVMLFYLFGSIMPLPPWGLVVGWVMALIAVELAYAFIFPYSFRYIADNDDDKMVILPV |
| IRIS_313-11911 | MLHHLKELAAVAGIHMILIYLCRFLLRRSRNVLFTVSNSLRFRLKVLTVLLYICLSVMLFYLFGSIMPLPPWGLVVGWVMALIAVELAYAFIFPYSFRYIADNDDDKMVILPV |
| IRIS_313-11912 | MLHHLKELAAVAGIHMILIYLCRFLLRRSRNVLFTVSNSLRFRLKVLTVLLYICLSVMLFYLFGSIMPLPPWGLVVGWVMALIAVELAYAFIFPYSFRYIADNDDDKMVILPV |
| IRIS_313-11913 | MLHQLKELAAVAGIHMILIYLCRFLLRRSRNVLFTVSNSLRFRLKVLTVLLYICLLVMLFYLFGSIMPLPPWGLVVGWVMALIAVELAYAFIFPYSFRYIADNDDDKMVILPV |
| IRIS_313-11914 | MLHHLKELAAVAGIHMILIYLCRFLLRRSRNVLFTVSNSLRFRLKVLTVLLYICLSVMLFYLFGSIMPLPPWGLVVGWVMALIAVELAYAFIFPYSFRYIADNDDDKMVILPV |
| IRIS_313-11915 | MLHHLKELAAVAGIHMILIYLCRFLLRRSRNVLFTVSNSLRFRLKVLTVLLYICLSVMLFYLFGSIMPLPPWGLVVGWVMALIAVELAYAFIFPYSFRYIADNDDDKMVILPV |
| IRIS_313-11916 | MLHHLKELAAVAGIHMILIYLCRFLLRRSRNVLFTVSNSLRFRLKVLTVLLYICLSVMLFYLFGSIMPLPPWGLVVGWVMALIAVELAYAFIFPYSFRYIADNDDDKMVILPV |
| IRIS_313-11917 | MLHHLKELAAVAGIHMILIYLCRFLLRRSRNVLFTVSNSLRFRLKVLTVLLYISLSAMLLYLFGSIMPLPPWGLVVGWVMALIAVELAYAFIFPYSFRYIADNDDDKMVILPV |
| IRIS_313-11918 | MLHHLKELAAVAGIHMILIYLCRFLLRRSRNVLFTVSNSLRFRLKVLTVLLYICLSVMLFYLFGSIMPLPPWGLVVGWVMALIAVELAYAFIFPYSFRYIADNDDDKMVILPV |
| IRIS_313-11919 | MLHHLKELAAVAGIHMILIYLCRFLLRRSRNVLFTVSNSLRFRLKVLTVLLYICLSVMLFYLFGSIMPLPPWGLVVGWVMALIAVELAYAFIFPYSFRYIADNDDDKMVILPV |
| IRIS_313-11920 | MLHHLKELAAVAGIHMILIYLCRFLLRRSRNVLFTVSNSLRFRLKVLTVLLYICLSVMLFYLFGSIMPLPPWGLVVGWVMALIAVELAYAFIFPYSFRYIADNDDDKMVILPV |
| IRIS_313-11921 | MLHHLKELAAVAGIHMILIYLCRFLLRRSRNVLFTVSNSLRFRLKVLTVLLYICLSVMLFYLFGSIMPLPPWGLVVGWVMALIAVELAYAFIFPYSFRYIADNDDDKMVILPV |
| IRIS_313-11922 | MLHHLKELAAVAGIHMILIYLCRFLLRRSRNVLFTVSNSLRFRLKVLTVLLYICLSVMLFYLFGSIMPLPPWGLVVGWVMALIAVELAYAFIFPYSFRYIADNDDDKMVILPV |
| IRIS_313-11923 | MLHQLKELAAVAGIHMILIYLCRFLLRRSRNVLFTVSNSLRFRLKVLTVLLYICLLVMLFYLFGSIMPLPPWGLVVGWVMALIAVELAYAFIFPYSFRYIADNDDDKMVILPV |
| IRIS_313-11924 | MLHQLKELAAVAGIHMILIYLCRFLLRRSRNVLFTVSNSLRFRLKVLTVLLYICLLVMLFYLFGSIMPLPPWGLVVGWVMALIAVELAYAFIFPYSFRYIADNDDDKMVILPV |
| IRIS_313-11925 | MLHHLKELAAVAGIHMILIYLCRFLLRRSRNVLFTVSNSLRFRLKVLTVLLYICLSVMLFYLFGSIMPLPPWGLVVGWVMALIAVELAYAFIFPYSFRYIADNDDDKMVILPV |
| IRIS_313-11926 | MLHQLKELAAVAGIHMILIYLCRFLLRRSRNVLFTVSNSLRFRLKVLTVLLYICLLVMLFYLFGSIMPLPPWGLVVGWVMALIAVELAYAFIFPYSFRYIADNDDDKMVILPV |
| IRIS_313-11927 | MLHHLKELAAVAGIHMILIYLCRFLLRRSRNVLFTVSNSLRFRLKVLTVLLYICLSVMLFYLFGSIMPLPPWGLVVGWVMALIAVELAYAFIFPYSFRYIADNDDDKMVILPV |
| IRIS_313-11928 | MLHQLKELAAVAGIHMILIYLCRFLLRRSRNVLFTVSNSLRFRLKVLTVLLYICLLVMLFYLFGSIMPLPPWGLVVGWVMALIAVELAYAFIFPYSFRYIADNDDDKMVILPV |
| IRIS_313-11929 | MLHHLKELAAVAGIHMILIYLCRFLLRRSRNVLFTVSNSLRFRLKVLTVLLYICLSVMLFYLFGSIMPLPPWGLVVGWVMALIAVELAYAFIFPYSFRYIADNDDDKMVILPV |
| IRIS_313-11930 | MLHHLKELAAVAGIHMILIYLCRFLLRRSRNVLFTVSNSLRFRLKVLTVLLYICLSVMLFYLFGSIMPLPPWGLVVGWVMALIAVELAYAFIFPYSFRYIADNDDDKMVILPV |
| IRIS_313-11931 | MLHHLKELAAVAGIHMILIYLCRFLLRRSRNVLFTVSNSLRFRLKVLTVLLYICLSVMLFYLFGSIMPLPPWGLVVGWVMALIAVELAYAFIFPYSFRYIADNDDDKMVILPV |
| IRIS_313-11932 | MLHHLKELAAVAGIHMILIYLCRFLLRRSRNVLFTVSNSLRFRLKVLTVLLYICLSVMLFYLFGSIMPLPPWGLVVGWVMALIAVELAYAFIFPYSFRYIADNDDDKMVILPV |
| IRIS_313-11934 | MLHHLKELAAVAGIHMILIYLCRFLLRRSRNVLFTVSNSLRFRLKVLTVLLYICLSVMLFYLFGSIMPLPPWGLVVGWVMALIAVELAYAFIFPYSFRYIADNDDDKMVILPV |
| IRIS_313-11935 | MLHHLKELAAVAGIHMILIYLCRFLLRRSRNVLFTVSNSLRFRLKVLTVLLYICLSVMLFYLFGSIMPLPPWGLVVGWVMALIAVELAYAFIFPYSFRYIADNDDDKMVILPV |
| IRIS_313-11936 | MLHHLKELAAVAGIHMILIYLCRFLLRRSRNVLFTVSNSLRFRLKVLTVLLYICLSVMLFYLFGSIMPLPPWGLVVGWVMALIAVELAYAFIFPYSFRYIADNDDDKMVILPV |
| IRIS_313-11937 | MLHHLKELAAVAGIHMILIYLCRFLLRRSRNVLFTVSNSLRFRLKVLTVLLYICLSVMLFYLFGSIMPLPPWGLVVGWVMALIAVELAYAFIFPYSFRYIADNDDDKMVILPV |
| IRIS_313-11938 | MLHHLKELAAVAGIHMILIYLCRFLLRRSRNVLFTVSNSLRFRLKVLTVLLYICLSVMLFYLFGSIMPLPPWGLVVGWVMALIAVELAYAFIFPYSFRYIADNDDDKMVILPV |
| IRIS_313-11939 | MLHHLKELAAVAGIHMILIYLCRFLLRRSRNVLFTVSNSLRFRLKVLTVLLYICLSVMLFYLFGSIMPLPPWGLVVGWVMALIAVELAYAFIFPYSFRYIADNDDDKMVILPV |
| IRIS_313-11940 | MLHHLKELAAVAGIHMILIYLCRFLLRRSRNVLFTVSNSLRFRLKVLTVLLYICLSVMLFYLFGSIMPLPPWGLVVGWVMALIAVELAYAFIFPYSFRYIADNDDDKMVILPV |
| IRIS_313-11941 | MLHHLKELAAVAGIHMILIYLCRFLLRRSRNVLFTVSNSLRFRLKVLTVLLYICLSVMLFYLFGSIMPLPPWGLVVGWVMALIAVELAYAFIFPYSFRYIADNDDDKMVILPV |
| IRIS_313-11943 | MLHHLKELAAVAGIHMILIYLCRFLLRRSRNVLFTVSNSLRFRLKVLTVLLYICLSVMLFYLFGSIMPLPPWGLVVGWVMALIAVELAYAFIFPYSFRYIADNDDDKMVILPV |
| IRIS_313-11944 | MLHHLKELAAVAGIHMILIYLCRFLLRRSRNVLFTVSNSLRFRLKVLTVLLYICLSVMLFYLFGSIMPLPPWGLVVGWVMALIAVELAYAFIFPYSFRYIADNDDDKMVILPV |
| IRIS_313-11945 | MLHHLKELAAVAGIHMILIYLCRFLLRRSRNVLFTVSNSLRFRLKVLTVLLYICLSVMLFYLFGSIMPLPPWGLVVGWVMALIAVELAYAFIFPYSFRYIADNDDDKMVILPV |
| IRIS_313-11946 | MLHQLKELAAVAGIHMILIYLCRFLLRRSRNVLFTVSNSLRFRLKVLTVLLYICLLVMLFYLFGSIMPLPPWGLVVGWVMALIAVELAYAFIFPYSFRYIADNDDDKMVILPV |
| IRIS_313-11947 | MLHHLKELAAVAGIHMILIYLCRFLLRRSRNVLFTVSNSLRFRLKVLTVLLYICLSVMLFYLFGSIMPLPPWGLVVGWVMALIAVELAYAFIFPYSFRYIADNDDDKMVILPV |
| IRIS_313-11948 | MLHHLKELAAVAGIHMILIYLCRFLLRRSRNVLFTVSNSLRFRLKVLTVLLYICLSVMLFYLFGSIMPLPPWGLVVGWVMALIAVELAYAFIFPYSFRYIADNDDDKMVILPV |
| IRIS_313-11949 | MLHHLKELAAVAGIHMILIYLCRFLLRRSRNVLFTVSNSLRFRLKVLTVLLYICLSVMLFYLFGSIMPLPPWGLVVGWVMALIAVELAYAFIFPYSFRYIADNDDDKMVILPV |
| IRIS_313-11950 | MLHHLKELAAVAGIHMILIYLCRFLLRRSRNVLFTVSNSLRFRLKVLTVLLYICLSVMLFYLFGSIMPLPPWGLVVGWVMALIAVELAYAFIFPYSFRYIADNDDDKMVILPV |
| IRIS_313-11951 | MLHHLKELAAVAGIHMILIYLCRFLLRRSRNVLFTVSNSLRFRLKVLTVLLYICLSVMLFYLFGSIMPLPPWGLVVGWVMALIAVELAYAFIFPYSFRYIADNDDDKMVILPV |
| IRIS_313-11952 | MLHHLKELAAVAGIHMILIYLCRFLLRRSRNVLFTVSNSLRFRLKVLTVLLYICLSVMLFYLFGSIMPLPPWGLVVGWVMALIAVELAYAFIFPYSFRYIADNDDDKMVILPV |
| IRIS_313-11953 | MLHHLKELAAVAGIHMILIYLCRFLLRRSRNVLFTVSNSLRFRLKVLTVLLYICLSVMLFYLFGSIMPLPPWGLVVGWVMALIAVELAYAFIFPYSFRYIADNDDDKMVILPV |
| IRIS_313-11954 | MLHHLKELAAVAGIHMILIYLCRFLLRRSRNVLFTVSNSLRFRLKVLTVLLYICLSVMLFYLFGSIMPLPPWGLVVGWVMALIAVELAYAFIFPYSFRYIADNDDDKMVILPV |
| IRIS_313-11955 | MLHHLKELAAVAGIHMILIYLCRFLLRRSRNVLFTVSNSLRFRLKVLTVLLYICLSVMLFYLFGSIMPLPPWGLVVGWVMALIAVELAYAFIFPYSFRYIADNDDDKMVILPV |
| IRIS_313-11956 | MLHHLKELAAVAGIHMILIYLCRFLLRRSRNVLFTVSNSLRFRLKVLTVLLYISLSAMLLYLFGSIMPLPPWGLVVGWVMALIAVELAYAFIFPYSFRYIADNDDDKMVILPV |
| IRIS_313-11957 | MLHHLKELAAVAGIHMILIYLCRFLLRRSRNVLFTVSNSLRFRLKVLTVLLYICLSVMLFYLFGSIMPLPPWGLVVGWVMALIAVELAYAFIFPYSFRYIADNDDDKMVILPV |
| IRIS_313-11958 | MLHHLKELAAVAGIHMILIYLCRFLLRRSRNVLFTVSNSLRFRLKVLTVLLYICLSVMLFYLFGSIMPLPPWGLVVGWVMALIAVELAYAFIFPYSFRYIADNDDDKMVILPV |
| IRIS_313-11959 | MLHHLKELAAVAGIHMILIYLCRFLLRRSRNVLFTVSNSLRFRLKVLTVLLYICLSVMLFYLFGSIMPLPPWGLVVGWVMALIAVELAYAFIFPYSFRYIADNDDDKMVILPV |
| IRIS_313-11960 | MLHHLKELAAVAGIHMILIYLCRFLLRRSRNVLFTVSNSLRFRLKVLTVLLYICLSVMLFYLFGSIMPLPPWGLVVGWVMALIAVELAYAFIFPYSFRYIADNDDDKMVILPV |
| IRIS_313-11961 | MLHHLKELAAVAGIHMILIYLCRFLLRRSRNVLFTVSNSLRFRLKVLTVLLYICLSVMLFYLFGSIMPLPPWGLVVGWVMALIAVELAYAFIFPYSFRYIADNDDDKMVILPV |
| IRIS_313-11962 | MLHHLKELAAVAGIHMILIYLCRFLLRRSRNVLFTVSNSLRFRLKVLTVLLYICLSVMLFYLFGSIMPLPPWGLVVGWVMALIAVELAYAFIFPYSFRYIADNDDDKMVILPV |
| IRIS_313-11963 | MLHHLKELAAVAGIHMILIYLCRFLLRRSRNVLFTVSNSLRFRLKVLTVLLYICLSVMLFYLFGSIMPLPPWGLVVGWVMALIAVELAYAFIFPYSFRYIADNDDDKMVILPV |
| IRIS_313-11964 | MLHHLKELAAVAGIHMILIYLCRFLLRRSRNVLFTVSNSLRFRLKVLTVLLYICLSVMLFYLFGSIMPLPPWGLVVGWVMALIAVELAYAFIFPYSFRYIADNDDDKMVILPV |
| IRIS_313-11965 | MLHHLKELAAVAGIHMILIYLCRFLLRRSRNVLFTVSNSLRFRLKVLTVLLYICLSVMLFYLFGSIMPLPPWGLVVGWVMALIAVELAYAFIFPYSFRYIADNDDDKMVILPV |
| IRIS_313-11966 | MLHHLKELAAVAGIHMILIYLCRFLLRRSRNVLFTVSNSLRFRLKVLTVLLYICLSVMLFYLFGSIMPLPPWGLVVGWVMALIAVELAYAFIFPYSFRYIADNDDDKMVILPV |
| IRIS_313-11967 | MLHHLKELAAVAGIHMILIYLCRFLLRRSRNVLFTVSNSLRFRLKVLTVLLYICLSVMLFYLFGSIMPLPPWGLVVGWVMALIAVELAYAFIFPYSFRYIADNDDDKMVILPV |
| IRIS_313-11968 | MLHHLKELAAVAGIHMILIYLCRFLLRRSRNVLFTVSNSLRFRLKVLTVLLYICLSVMLFYLFGSIMPLPPWGLVVGWVMALIAVELAYAFIFPYSFRYIADNDDDKMVILPV |
| IRIS_313-11969 | MLHHLKELAAVAGIHMILIYLCRFLLRRSRNVLFTVSNSLRFRLKVLTVLLYICLSVMLFYLFGSIMPLPPWGLVVGWVMALIAVELAYAFIFPYSFRYIADNDDDKMVILPV |
| IRIS_313-11970 | MLHHLKELAAVAGIHMILIYLCRFLLRRSRNVLFTVSNSLRFRLKVLTVLLYICLSVMLFYLFGSIMPLPPWGLVVGWVMALIAVELAYAFIFPYSFRYIADNDDDKMVILPV |
| IRIS_313-11971 | MLHHLKELAAVAGIHMILIYLCRFLLRRSRNVLFTVSNSLRFRLKVLTVLLYICLSVMLFYLFGSIMPLPPWGLVVGWVMALIAVELAYAFIFPYSFRYIADNDDDKMVILPV |
| IRIS_313-11973 | MLHHLKELAAVAGIHMILIYLCRFLLRRSRNVLFTVSNSLRFRLKVLTVLLYICLSVMLFYLFGSIMPLPPWGLVVGWVMALIAVELAYAFIFPYSFRYIADNDDDKMVILPV |
| IRIS_313-11974 | MLHHLKELAAVAGIHMILIYLCRFLLRRSRNVLFTVSNSLRFRLKVLTVLLYICLSVMLFYLFGSIMPLPPWGLVVGWVMALIAVELAYAFIFPYSFRYIADNDDDKMVILPV |
| IRIS_313-11975 | MLHHLKELAAVAGIHMILIYLCRFLLRRSRNVLFTVSNSLRFRLKVLTVLLYICLSVMLFYLFGSIMPLPPWGLVVGWVMALIAVELAYAFIFPYSFRYIADNDDDKMVILPV |
| IRIS_313-11976 | MLHHLKELAAVAGIHMILIYLCRFLLRRSRNVLFTVSNSLRFRLKVLTVLLYICLSVMLFYLFGSIMPLPPWGLVVGWVMALIAVELAYAFIFPYSFRYIADNDDDKMVILPV |
| IRIS_313-11977 | MLHHLKELAAVAGIHMILIYLCRFLLRRSRNVLFTVSNSLRFRLKVLTVLLYICLSVMLFYLFGSIMPLPPWGLVVGWVMALIAVELAYAFIFPYSFRYIADNDDDKMVILPV |
| IRIS_313-11978 | MLHHLKELAAVAGIHMILIYLCRFLLRRSRNVLFTVSNSLRFRLKVLTVLLYICLSVMLFYLFGSIMPLPPWGLVVGWVMALIAVELAYAFIFPYSFRYIADNDDDKMVILPV |
| IRIS_313-11979 | MLHHLKELAAVAGIHMILIYLCRFLLRRSRNVLFTVSNSLRFRLKVLTVLLYICLSVMLFYLFGSIMPLPPWGLVVGWVMALIAVELAYAFIFPYSFRYIADNDDDKMVILPV |
| IRIS_313-11980 | MLHHLKELAAVAGIHMILIYLCRFLLRRSRNVLFTVSNSLRFRLKVLTVLLYICLSVMLFYLFGSIMPLPPWGLVVGWVMALIAVELAYAFIFPYSFRYIADNDDDKMVILPV |
| IRIS_313-11981 | MLHHLKELAAVAGIHMILIYLCRFLLRRSRNVLFTVSNSLRFRLKVLTVLLYICLSVMLFYLFGSIMPLPPWGLVVGWVMALIAVELAYAFIFPYSFRYIADNDDDKMVILPV |
| IRIS_313-11982 | MLHHLKELAAVAGIHMILIYLCRFLLRRSRNVLFTVSNSLRFRLKVLTVLLYICLSVMLFYLFGSIMPLPPWGLVVGWVMALIAVELAYAFIFPYSFRYIADNDDDKMVILPV |
| IRIS_313-11983 | MLHHLKELAAVAGIHMILIYLCRFLLRRSRNVLFTVSNSLRFRLKVLTVLLYICLSVMLFYLFGSIMPLPPWGLVVGWVMALIAVELAYAFIFPYSFRYIADNDDDKMVILPV |
| IRIS_313-11984 | MLHHLKELAAVAGIHMILIYLCRFLLRRSRNVLFTVSNSLRFRLKVLTVLLYICLSVMLFYLFGSIMPLPPWGLVVGWVMALIAVELAYAFIFPYSFRYIADNDDDKMVILPV |
| IRIS_313-11985 | MLHHLKELAAVAGIHMILIYLCRFLLRRSRNVLFTVSNSLRFRLKVLTVLLYICLSVMLFYLFGSIMPLPPWGLVVGWVMALIAVELAYAFIFPYSFRYIADNDDDKMVILPV |
| IRIS_313-11986 | MLHHLKELAAVAGIHMILIYLCRFLLRRSRNVLFTVSNSLRFRLKVLTVLLYICLSVMLFYLFGSIMPLPPWGLVVGWVMALIAVELAYAFIFPYSFRYIADNDDDKMVILPV |
| IRIS_313-11987 | MLHHLKELAAVAGIHMILIYLCRFLLRRSRNVLFTVSNSLRFRLKVLTVLLYICLSVMLFYLFGSIMPLPPWGLVVGWVMALIAVELAYAFIFPYSFRYIADNDDDKMVILPV |
| IRIS_313-11988 | MLHHLKELAAVAGIHMILIYLCRFLLRRSRNVLFTVSNSLRFRLKVLTVLLYICLSVMLFYLFGSIMPLPPWGLVVGWVMALIAVELAYAFIFPYSFRYIADNDDDKMVILPV |
| IRIS_313-11989 | MLHHLKELAAVAGIHMILIYLCRFLLRRSRNVLFTVSNSLRFRLKVLTVLLYICLSVMLFYLFGSIMPLPPWGLVVGWVMALIAVELAYAFIFPYSFRYIADNDDDKMVILPV |
| IRIS_313-11990 | MLHHLKELAAVAGIHMILIYLCRFLLRRSRNVLFTVSNSLRFRLKVLTVLLYICLSVMLFYLFGSIMPLPPWGLVVGWVMALIAVELAYAFIFPYSFRYIADNDDDKMVILPV |
| IRIS_313-11991 | MLHHLKELAAVAGIHMILIYLCRFLLRRSRNVLFTVSNSLRFRLKVLTVLLYICLSVMLFYLFGSIMPLPPWGLVVGWVMALIAVELAYAFIFPYSFRYIADNDDDKMVILPV |
| IRIS_313-11992 | MLHHLKELAAVAGIHMILIYLCRFLLRRSRNVLFTVSNSLRFRLKVLTVLLYICLSVMLFYLFGSIMPLPPWGLVVGWVMALIAVELAYAFIFPYSFRYIADNDDDKMVILPV |
| IRIS_313-11993 | MLHHLKELAAVAGIHMILIYLCRFLLRRSRNVLFTVSNSLRFRLKVLTVLLYICLSVMLFYLFGSIMPLPPWGLVVGWVMALIAVELAYAFIFPYSFRYIADNDDDKMVILPV |
| IRIS_313-11994 | MLHQLKELAAVAGIHMILIYLCRFLLRRSRNVLFTVSNSLRFRLKVLTVLLYICLLVMLFYLFGSIMPLPPWGLVVGWVMALIAVELAYAFIFPYSFRYIADNDDDKMVILPV |
| IRIS_313-11995 | MLHQLKELAAVAGIHMILIYLCRFLLRRSRNVLFTVSNSLRFRLKVLTVLLYICLLVMLFYLFGSIMPLPPWGLVVGWVMALIAVELAYAFIFPYSFRYIADNDDDKMVILPV |
| IRIS_313-11996 | MLHHLKELAAVAGIHMILIYLCRFLLRRSRNVLFTVSNSLRFRLKVLTVLLYICLSVMLFYLFGSIMPLPPWGLVVGWVMALIAVELAYAFIFPYSFRYIADNDDDKMVILPV |
| IRIS_313-11997 | MLHHLKELAAVAGIHMILIYLCRFLLRRSRNVLFTVSNSLRFRLKVLTVLLYICLSVMLFYLFGSIMPLPPWGLVVGWVMALIAVELAYAFIFPYSFRYIADNDDDKMVILPV |
| IRIS_313-11998 | MLHHLKELAAVAGIHMILIYLCRFLLRRSRNVLFTVSNSLRFRLKVLTVLLYICLSVMLFYLFGSIMPLPPWGLVVGWVMALIAVELAYAFIFPYSFRYIADNDDDKMVILPV |
| IRIS_313-11999 | MLHHLKELAAVAGIHMILIYLCRFLLRRSRNVLFTVSNSLRFRLKVLTVLLYICLSVMLFYLFGSIMPLPPWGLVVGWVMALIAVELAYAFIFPYSFRYIADNDDDKMVILPV |
| IRIS_313-12000 | MLHHLKELAAVAGIHMILIYLCRFLLRRSRNVLFTVSNSLRFRLKVLTVLLYICLSVMLFYLFGSIMPLPPWGLVVGWVMALIAVELAYAFIFPYSFRYIADNDDDKMVILPV |
| IRIS_313-12002 | MLHHLKELAAVAGIHMILIYLCRFLLRRSRNVLFTVSNSLRFRLKVLTVLLYISLSAMLLYLFGSIMPLPPWGLVVGWVMALIAVELAYAFIFPYSFRYIADNDDDKMVILPV |
| IRIS_313-12003 | MLHHLKELAAVAGIHMILIYLCRFLLRRSRNVLFTVSNSLRFRLKVLTVLLYICLSVMLFYLFGSIMPLPPWGLVVGWVMALIAVELAYAFIFPYSFRYIADNDDDKMVILPV |
| IRIS_313-12004 | MLHHLKELAAVAGIHMILIYLCRFLLRRSRNVLFTVSNSLRFRLKVLTVLLYICLSVMLFYLFGSIMPLPPWGLVVGWVMALIAVELAYAFIFPYSFRYIADNDDDKMVILPV |
| IRIS_313-12005 | MLHHLKELAAVAGIHMILIYLCRFLLRRSRNVLFTVSNSLRFRLKVLTVLLYICLSVMLFYLFGSIMPLPPWGLVVGWVMALIAVELAYAFIFPYSFRYIADNDDDKMVILPV |
| IRIS_313-12006 | MLHHLKELAAVAGIHMILIYLCRFLLRRSRNVLFTVSNSLRFRLKVLTVLLYICLSVMLFYLFGSIMPLPPWGLVVGWVMALIAVELAYAFIFPYSFRYIADNDDDKMVILPV |
| IRIS_313-12007 | MLHHLKELAAVAGIHMILIYLCRFLLRRSRNVLFTVSNSLRFRLKVLTVLLYICLSVMLFYLFGSIMPLPPWGLVVGWVMALIAVELAYAFIFPYSFRYIADNDDDKMVILPV |
| IRIS_313-12009 | MLHHLKELAAVAGIHMILIYLCRFLLRRSRNVLFTVSNSLRFRLKVLTVLLYICLSVMLFYLFGSIMPLPPWGLVVGWVMALIAVELAYAFIFPYSFRYIADNDDDKMVILPV |
| IRIS_313-12010 | MLHHLKELAAVAGIHMILIYLCRFLLRRSRNVLFTVSNSLRFRLKVLTVLLYICLSVMLFYLFGSIMPLPPWGLVVGWVMALIAVELAYAFIFPYSFRYIADNDDDKMVILPV |
| IRIS_313-12011 | MLHHLKELAAVAGIHMILIYLCRFLLRRSRNVLFTVSNSLRFRLKVLTVLLYICLSVMLFYLFGSIMPLPPWGLVVGWVMALIAVELAYAFIFPYSFRYIADNDDDKMVILPV |
| IRIS_313-12012 | MLHHLKELAAVAGIHMILIYLCRFLLRRSRNVLFTVSNSLRFRLKVLTVLLYICLSVMLFYLFGSIMPLPPWGLVVGWVMALIAVELAYAFIFPYSFRYIADNDDDKMVILPV |
| IRIS_313-12013 | MLHHLKELAAVAGIHMILIYLCRFLLRRSRNVLFTVSNSLRFRLKVLTVLLYICLSVMLFYLFGSIMPLPPWGLVVGWVMALIAVELAYAFIFPYSFRYIADNDDDKMVILPV |
| IRIS_313-12014 | MLHHLKELAAVAGIHMILIYLCRFLLRRSRNVLFTVSNSLRFRLKVLTVLLYICLSVMLFYLFGSIMPLPPWGLVVGWVMALIAVELAYAFIFPYSFRYIADNDDDKMVILPV |
| IRIS_313-12015 | MLHHLKELAAVAGIHMILIYLCRFLLRRSRNVLFTVSNSLRFRLKVLTVLLYICLSVMLFYLFGSIMPLPPWGLVVGWVMALIAVELAYAFIFPYSFRYIADNDDDKMVILPV |
| IRIS_313-12016 | MLHHLKELAAVAGIHMILIYLCRFLLRRSRNVLFTVSNSLRFRLKVLTVLLYICLSVMLFYLFGSIMPLPPWGLVVGWVMALIAVELAYAFIFPYSFRYIADNDDDKMVILPV |
| IRIS_313-12017 | MLHHLKELAAVAGIHMILIYLCRFLLRRSRNVLFTVSNSLRFRLKVLTVLLYICLSVMLFYLFGSIMPLPPWGLVVGWVMALIAVELAYAFIFPYSFRYIADNDDDKMVILPV |
| IRIS_313-12018 | MLHHLKELAAVAGIHMILIYLCRFLLRRSRNVLFTVSNSLRFRLKVLTVLLYICLSVMLFYLFGSIMPLPPWGLVVGWVMALIAVELAYAFIFPYSFRYIADNDDDKMVILPV |
| IRIS_313-12021 | MLHHLKELAAVAGIHMILIYLCRFLLRRSRNVLFTVSNSLRFRLKVLTVLLYICLSVMLFYLFGSIMPLPPWGLVVGWVMALIAVELAYAFIFPYSFRYIADNDDDKMVILPV |
| IRIS_313-12024 | MLHHLKELAAVAGIHMILIYLCRFLLRRSRNVLFTVSNSLRFRLKVLTVLLYICLSVMLFYLFGSIMPLPPWGLVVGWVMALIAVELAYAFIFPYSFRYIADNDDDKMVILPV |
| IRIS_313-12027 | MLHHLKELAAVAGIHMILIYLCRFLLRRSRNVLFTVSNSLRFRLKVLTVLLYICLSVMLFYLFGSIMPLPPWGLVVGWVMALIAVELAYAFIFPYSFRYIADNDDDKMVILPV |
| IRIS_313-12028 | MLHHLKELAAVAGIHMILIYLCRFLLRRSRNVLFTVSNSLRFRLKVLTVLLYICLSVMLFYLFGSIMPLPPWGLVVGWVMALIAVELAYAFIFPYSFRYIADNDDDKMVILPV |
| IRIS_313-12029 | MLHQLKELAAVAGIHMILIYLCRFLLRRSRNVLFTVSNSLRFRLKVLTVLLYICLLVMLFYLFGSIMPLPPWGLVVGWVMALIAVELAYAFIFPYSFRYIADNDDDKMVILPV |
| IRIS_313-12030 | MLHHLKELAAVAGIHMILIYLCRFLLRRSRNVLFTVSNSLRFRLKVLTVLLYICLSVMLFYLFGSIMPLPPWGLVVGWVMALIAVELAYAFIFPYSFRYIADNDDDKMVILPV |
| IRIS_313-12031 | MLHHLKELAAVAGIHMILIYLCRFLLRRSRNVLFTVSNSLRFRLKVLTVLLYICLSVMLFYLFGSIMPLPPWGLVVGWVMALIAVELAYAFIFPYSFRYIADNDDDKMVILPV |
| IRIS_313-12033 | MLHHLKELAAVAGIHMILIYLCRFLLRRSRNVLFTVSNSLRFRLKVLTVLLYICLSVMLFYLFGSIMPLPPWGLVVGWVMALIAVELAYAFIFPYSFRYIADNDDDKMVILPV |
| IRIS_313-12036 | MLHHLKELAAVAGIHMILIYLCRFLLRRSRNVLFTVSNSLRFRLKVLTVLLYICLSVMLFYLFGSIMPLPPWGLVVGWVMALIAVELAYAFIFPYSFRYIADNDDDKMVILPV |
| IRIS_313-12037 | MLHHLKELAAVAGIHMILIYLCRFLLRRSRNVLFTVSNSLRFRLKVLTVLLYICLSVMLFYLFGSIMPLPPWGLVVGWVMALIAVELAYAFIFPYSFRYIADNDDDKMVILPV |
| IRIS_313-12038 | MLHHLKELAAVAGIHMILIYLCRFLLRRSRNVLFTVSNSLRFRLKVLTVLLYICLSVMLFYLFGSIMPLPPWGLVVGWVMALIAVELAYAFIFPYSFRYIADNDDDKMVILPV |
| IRIS_313-12039 | MLHHLKELAAVAGIHMILIYLCRFLLRRSRNVLFTVSNSLRFRLKVLTVLLYICLSVMLFYLFGSIMPLPPWGLVVGWVMALIAVELAYAFIFPYSFRYIADNDDDKMVILPV |
| IRIS_313-12040 | MLHHLKELAAVAGIHMILIYLCRFLLRRSRNVLFTVSNSLRFRLKVLTVLLYICLSVMLFYLFGSIMPLPPWGLVVGWVMALIAVELAYAFIFPYSFRYIADNDDDKMVILPV |
| IRIS_313-12041 | MLHHLKELAAVAGIHMILIYLCRFLLRRSRNVLFTVSNSLRFRLKVLTVLLYICLSVMLFYLFGSIMPLPPWGLVVGWVMALIAVELAYAFIFPYSFRYIADNDDDKMVILPV |
| IRIS_313-12042 | MLHHLKELAAVAGIHMILIYLCRFLLRRSRNVLFTVSNSLRFRLKVLTVLLYICLSVMLFYLFGSIMPLPPWGLVVGWVMALIAVELAYAFIFPYSFRYIADNDDDKMVILPV |
| IRIS_313-12043 | MLHHLKELAAVAGIHMILIYLCRFLLRRSRNVLFTVSNSLRFRLKVLTVLLYICLSVMLFYLFGSIMPLPPWGLVVGWVMALIAVELAYAFIFPYSFRYIADNDDDKMVILPV |
| IRIS_313-12044 | MLHHLKELAAVAGIHMILIYLCRFLLRRSRNVLFTVSNSLRFRLKVLTVLLYICLSVMLFYLFGSIMPLPPWGLVVGWVMALIAVELAYAFIFPYSFRYIADNDDDKMVILPV |
| IRIS_313-12045 | MLHQLKELAAVAGIHMILIYLCRFLLRRSRNVLFTVSNSLRFRLKVLTVLLYICLLVMLFYLFGSIMPLPPWGLVVGWVMALIAVELAYAFIFPYSFRYIADNDDDKMVILPV |
| IRIS_313-12046 | MLHHLKELAAVAGIHMILIYLCRFLLRRSRNVLFTVSNSLRFRLKVLTVLLYICLSVMLFYLFGSIMPLPPWGLVVGWVMALIAVELAYAFIFPYSFRYIADNDDDKMVILPV |

5927 IRIS_313-12047 MLHHLKELAAVAGIHMILIYLCRFLLRRSRNVLFTVSNSLRFRLKVLTVLLYICLSVMLFYLFGSIMPLPPWGLVVGWVMALIAVELAYAFIFPYSFRYIADNDDDKMVILPV
5928 IRIS_313-12048 MLHHLKELAAVAGIHMILIYLCRFLLRRSRNVLFTVSNSLRFRLKVLTVLLYICLSVMLFYLFGSIMPLPPWGLVVGWVMALIAVELAYAFIFPYSFRYIADNDDDKMVILPV
5929 IRIS_313-12049 MLHHLKELAAVAGIHMILIYLCRFLLRRSRNVLFTVSNSLRFRLKVLTVLLYICLSVMLFYLFGSIMPLPPWGLVVGWVMALIAVELAYAFIFPYSFRYIADNDDDKMVILPV
5930 IRIS_313-12050 MLHHLKELAAVAGIHMILIYLCRFLLRRSRNVLFTVSNSLRFRLKVLTVLLYICLSVMLFYLFGSIMPLPPWGLVVGWVMALIAVELAYAFIFPYSFRYIADNDDDKMVILPV
5931 IRIS_313-12051 MLHHLKELAAVAGIHMILIYLCRFLLRRSRNVLFTVSNSLRFRLKVLTVLLYICLSVMLFYLFGSIMPLPPWGLVVGWVMALIAVELAYAFIFPYSFRYIADNDDDKMVILPV
5932 IRIS_313-12052 MLHHLKELAAVAGIHMILIYLCRFLLRRSRNVLFTVSNSLRFRLKVLTVLLYICLSVMLFYLFGSIMPLPPWGLVVGWVMALIAVELAYAFIFPYSFRYIADNDDDKMVILPV
5933 IRIS_313-12053 MLHHLKELAAVAGIHMILIYLCRFLLRRSRNVLFTVSNSLRFRLKVLTVLLYICLSVMLFYLFGSIMPLPPWGLVVGWVMALIAVELAYAFIFPYSFRYIADNDDDKMVILPV
5934 IRIS_313-12054 MLHHLKELAAVAGIHMILIYLCRFLLRRSRNVLFTVSNSLRFRLKVLTVLLYICLSVMLFYLFGSIMPLPPWGLVVGWVMALIAVELAYAFIFPYSFRYIADNDDDKMVILPV
5935 IRIS_313-12055 MLHHLKELAAVAGIHMILIYLCRFLLRRSRNVLFTVSNSLRFRLKVLTVLLYISLSAMLLYLFGSIMPLPPWGLVVGWVMALIAVELAYAFIFPYSFRYIADNDDDKMVILPV
5936 IRIS_313-12057 MLHHLKELAAVAGIHMILIYLCRFLLRRSRNVLFTVSNSLRFRLKVLTVLLYICLSVMLFYLFGSIMPLPPWGLVVGWVMALIAVELAYAFIFPYSFRYIADNDDDKMVILPV
5937 IRIS_313-12058 MLHHLKELAAVAGIHMILIYLCRFLLRRSRNVLFTVSNSLRFRLKVLTVLLYICLSVMLFYLFGSIMPLPPWGLVVGWVMALIAVELAYAFIFPYSFRYIADNDDDKMVILPV
5938 IRIS_313-12059 MLHQLKELAAVAGIHMILIYLCRFLLRRSRNVLFTVSNSLRFRLKVLTVLLYICLLVMLFYLFGSIMPLPPWGLVVGWVMALIAVELAYAFIFPYSFRYIADNDDDKMVILPV
5939 IRIS_313-12060 MLHQLKELAAVAGIHMILIYLCRFLLRRSRNVLFTVSNSLRFRLKVLTVLLYICLLVMLFYLFGSIMPLPPWGLVVGWVMALIAVELAYAFIFPYSFRYIADNDDDKMVILPV
5940 IRIS_313-12061 MLHHLKELAAVAGIHMILIYLCRFLLRRSRNVLFTVSNSLRFRLKVLTVLLYICLSVMLFYLFGSIMPLPPWGLVVGWVMALIAVELAYAFIFPYSFRYIADNDDDKMVILPV
5941 IRIS_313-12063 MLHHLKELAAVAGIHMILIYLCRFLLRRSRNVLFTVSNSLRFRLKVLTVLLYICLSVMLFYLFGSIMPLPPWGLVVGWVMALIAVELAYAFIFPYSFRYIADNDDDKMVILPV
5942 IRIS_313-12065 MLHHLKELAAVAGIHMILIYLCRFLLRRSRNVLFTVSNSLRFRLKVLTVLLYICLSVMLFYLFGSIMPLPPWGLVVGWVMALIAVELAYAFIFPYSFRYIADNDDDKMVILPV
5943 IRIS_313-12066 MLHHLKELAAVAGIHMILIYLCRFLLRRSRNVLFTVSNSLRFRLKVLTVLLYICLSVMLFYLFGSIMPLPPWGLVVGWVMALIAVELAYAFIFPYSFRYIADNDDDKMVILPV
5944 IRIS_313-12067 MLHHLKELAAVAGIHMILIYLCRFLLRRSRNVLFTVSNSLRFRLKVLTVLLYICLSVMLFYLFGSIMPLPPWGLVVGWVMALIAVELAYAFIFPYSFRYIADNDDDKMVILPV
5945 IRIS_313-12068 MLHHLKELAAVAGIHMILIYLCRFLLRRSRNVLFTVSNSLRFRLKVLTVLLYICLSVMLFYLFGSIMPLPPWGLVVGWVMALIAVELAYAFIFPYSFRYIADNDDDKMVILPV
5946 IRIS_313-12069 MLHHLKELAAVAGIHMILIYLCRFLLRRSRNVLFTVSNSLRFRLKVLTVLLYICLSVMLFYLFGSIMPLPPWGLVVGWVMALIAVELAYAFIFPYSFRYIADNDDDKMVILPV
5947 IRIS_313-12070 MLHHLKELAAVAGIHMILIYLCRFLLRRSRNVLFTVSNSLRFRLKVLTVLLYICLSVMLFYLFGSIMPLPPWGLVVGWVMALIAVELAYAFIFPYSFRYIADNDDDKMVILPV
5948 IRIS_313-12071 MLHQLKELAAVAGIHMILIYLCRFLLRRSRNVLFTVSNSLRFRLKVLTVLLYICLLVMLFYLFGSIMPLPPWGLVVGWVMALIAVELAYAFIFPYSFRYIADNDDDKMVILPV
5949 IRIS_313-12072 MLHHLKELAAVAGIHMILIYLCRFLLRRSRNVLFTVSNSLRFRLKVLTVLLYICLSVMLFYLFGSIMPLPPWGLVVGWVMALIAVELAYAFIFPYSFRYIADNDDDKMVILPV
5950 IRIS_313-12073 MLHHLKELAAVAGIHMILIYLCRFLLRRSRNVLFTVSNSLRFRLKVLTVLLYICLSVMLFYLFGSIMPLPPWGLVVGWVMALIAVELAYAFIFPYSFRYIADNDDDKMVILPV
5951 IRIS_313-12074 MLHHLKELAAVAGIHMILIYLCRFLLRRSRNVLFTVSNSLRFRLKVLTVLLYICLSVMLFYLFGSIMPLPPWGLVVGWVMALIAVELAYAFIFPYSFRYIADNDDDKMVILPV
5952 IRIS_313-12076 MLHHLKELAAVAGIHMILIYLCRFLLRRSRNVLFTVSNSLRFRLKVLTVLLYICLSVMLFYLFGSIMPLPPWGLVVGWVMALIAVELAYAFIFPYSFRYIADNDDDKMVILPV
5953 IRIS_313-12077 MLHHLKELAAVAGIHMILIYLCRFLLRRSRNVLFTVSNSLRFRLKVLTVLLYICLSVMLFYLFGSIMPLPPWGLVVGWVMALIAVELAYAFIFPYSFRYIADNDDDKMVILPV
5954 IRIS_313-12078 MLHHLKELAAVAGIHMILIYLCRFLLRRSRNVLFTVSNSLRFRLKVLTVLLYICLSVMLFYLFGSIMPLPPWGLVVGWVMALIAVELAYAFIFPYSFRYIADNDDDKMVILPV
5955 IRIS_313-12079 MLHHLKELAAVAGIHMILIYLCRFLLRRSRNVLFTVSNSLRFRLKVLTVLLYICLSVMLFYLFGSIMPLPPWGLVVGWVMALIAVELAYAFIFPYSFRYIADNDDDKMVILPV
5956 IRIS_313-12080 MLHHLKELAAVAGIHMILIYLCRFLLRRSRNVLFTVSNSLRFRLKVLTVLLYICLSVMLFYLFGSIMPLPPWGLVVGWVMALIAVELAYAFIFPYSFRYIADNDDDKMVILPV
5957 IRIS_313-12081 MLHHLKELAAVAGIHMILIYLCRFLLRRSRNVLFTVSNSLRFRLKVLTVLLYICLSVMLFYLFGSIMPLPPWGLVVGWVMALIAVELAYAFIFPYSFRYIADNDDDKMVILPV
5958 IRIS_313-12082 MLHHLKELAAVAGIHMILIYLCRFLLRRSRNVLFTVSNSLRFRLKVLTVLLYICLSVMLFYLFGSIMPLPPWGLVVGWVMALIAVELAYAFIFPYSFRYIADNDDDKMVILPV
5959 IRIS_313-12083 MLHHLKELAAVAGIHMILIYLCRFLLRRSRNVLFTVSNSLRFRLKVLTVLLYICLSVMLFYLFGSIMPLPPWGLVVGWVMALIAVELAYAFIFPYSFRYIADNDDDKMVILPV
5960 IRIS_313-12093 MLHHLKELAAVAGIHMILIYLCRFLLRRSRNVLFTVSNSLRFRLKVLTVLLYICLSVMLFYLFGSIMPLPPWGLVVGWVMALIAVELAYAFIFPYSFRYIADNDDDKMVILPV
5961 IRIS_313-12094 MLHHLKELAAVAGIHMILIYLCRFLLRRSRNVLFTVSNSLRFRLKVLTVLLYICLSVMLFYLFGSIMPLPPWGLVVGWVMALIAVELAYAFIFPYSFRYIADNDDDKMVILPV
5962 IRIS_313-12096 MLHHLKELAAVAGIHMILIYLCRFLLRRSRNVLFTVSNSLRFRLKVLTVLLYICLSVMLFYLFGSIMPLPPWGLVVGWVMALIAVELAYAFIFPYSFRYIADNDDDKMVILPV
5963 IRIS_313-12097 MLHHLKELAAVAGIHMILIYLCRFLLRRSRNVLFTVSNSLRFRLKVLTVLLYICLSVMLFYLFGSIMPLPPWGLVVGWVMALIAVELAYAFIFPYSFRYIADNDDDKMVILPV
5964 IRIS_313-12101 MLHHLKELAAVAGIHMILIYLCRFLLRRSRNVLFTVSNSLRFRLKVLTVLLYICLSVMLFYLFGSIMPLPPWGLVVGWVMALIAVELAYAFIFPYSFRYIADNDDDKMVILPV
5965 IRIS_313-12102 MLHHLKELAAVAGIHMILIYLCRFLLRRSRNVLFTVSNSLRFRLKVLTVLLYICLSVMLFYLFGSIMPLPPWGLVVGWVMALIAVELAYAFIFPYSFRYIADNDDDKMVILPV
5966 IRIS_313-12108 MLHHLKELAAVAGIHMILIYLCRFLLRRSRNVLFTVSNSLRFRLKVLTVLLYICLSVMLFYLFGSIMPLPPWGLVVGWVMALIAVELAYAFIFPYSFRYIADNDDDKMVILPV
5967 IRIS_313-12109 MLHHLKELAAVAGIHMILIYLCRFLLRRSRNVLFTVSNSLRFRLKVLTVLLYICLSVMLFYLFGSIMPLPPWGLVVGWVMALIAVELAYAFIFPYSFRYIADNDDDKMVILPV
5968 IRIS_313-12118 MLHHLKELAAVAGIHMILIYLCRFLLRRSRNVLFTVSNSLRFRLKVLTVLLYICLSVMLFYLFGSIMPLPPWGLVVGWVMALIAVELAYAFIFPYSFRYIADNDDDKMVILPV
5969 IRIS_313-12121 MLHHLKELAAVAGIHMILIYLCRFLLRRSRNVLFTVSNSLRFRLKVLTVLLYICLSVMLFYLFGSIMPLPPWGLVVGWVMALIAVELAYAFIFPYSFRYIADNDDDKMVILPV
5970 IRIS_313-12127 MLHHLKELAAVAGIHMILIYLCRFLLRRSRNVLFTVSNSLRFRLKVLTVLLYICLSVMLFYLFGSIMPLPPWGLVVGWVMALIAVELAYAFIFPYSFRYIADNDDDKMVILPV
5971 IRIS_313-12128 MLHHLKELAAVAGIHMILIYLCRFLLRRSRNVLFTVSNSLRFRLKVLTVLLYICLSVMLFYLFGSIMPLPPWGLVVGWVMALIAVELAYAFIFPYSFRYIADNDDDKMVILPV
5972 IRIS_313-12129 MLHHLKELAAVAGIHMILIYLCRFLLRRSRNVLFTVSNSLRFRLKVLTVLLYICLSVMLFYLFGSIMPLPPWGLVVGWVMALIAVELAYAFIFPYSFRYIADNDDDKMVILPV
5973 IRIS_313-12130 MLHHLKELAAVAGIHMILIYLCRFLLRRSRNVLFTVSNSLRFRLKVLTVLLYICLSVMLFYLFGSIMPLPPWGLVVGWVMALIAVELAYAFIFPYSFRYIADNDDDKMVILPV
5974 IRIS_313-12131 MLHHLKELAAVAGIHMILIYLCRFLLRRSRNVLFTVSNSLRFRLKVLTVLLYICLSVMLFYLFGSIMPLPPWGLVVGWVMALIAVELAYAFIFPYSFRYIADNDDDKMVILPV
5975 IRIS_313-12133 MLHHLKELAAVAGIHMILIYLCRFLLRRSRNVLFTVSNSLRFRLKVLTVLLYICLSVMLFYLFGSIMPLPPWGLVVGWVMALIAVELAYAFIFPYSFRYIADNDDDKMVILPV
5976 IRIS_313-12134 MLHHLKELAAVAGIHMILIYLCRFLLRRSRNVLFTVSNSLRFRLKVLTVLLYICLSVMLFYLFGSIMPLPPWGLVVGWVMALIAVELAYAFIFPYSFRYIADNDDDKMVILPV
5977 IRIS_313-12135 MLHHLKELAAVAGIHMILIYLCRFLLRRSRNVLFTVSNSLRFRLKVLTVLLYICLSVMLFYLFGSIMPLPPWGLVVGWVMALIAVELAYAFIFPYSFRYIADNDDDKMVILPV
5978 IRIS_313-12138 MLHHLKELAAVAGIHMILIYLCRFLLRRSRNVLFTVSNSLRFRLKVLTVLLYICLSVMLFYLFGSIMPLPPWGLVVGWVMALIAVELAYAFIFPYSFRYIADNDDDKMVILPV
5979 IRIS_313-12139 MLHHLKELAAVAGIHMILIYLCRFLLRRSRNVLFTVSNSLRFRLKVLTVLLYICLSVMLFYLFGSIMPLPPWGLVVGWVMALIAVELAYAFIFPYSFRYIADNDDDKMVILPV
5980 IRIS_313-12141 MLHHLKELAAVAGIHMILIYLCRFLLRRSRNVLFTVSNSLRFRLKVLTVLLYICLSVMLFYLFGSIMPLPPWGLVVGWVMALIAVELAYAFIFPYSFRYIADNDDDKMVILPV
5981 IRIS_313-12142 MLHHLKELAAVAGIHMILIYLCRFLLRRSRNVLFTVSNSLRFRLKVLTVLLYICLSVMLFYLFGSIMPLPPWGLVVGWVMALIAVELAYAFIFPYSFRYIADNDDDKMVILPV
5982 IRIS_313-12143 MLHHLKELAAVAGIHMILIYLCRFLLRRSRNVLFTVSNSLRFRLKVLTVLLYICLSVMLFYLFGSIMPLPPWGLVVGWVMALIAVELAYAFIFPYSFRYIADNDDDKMVILPV
5983 IRIS_313-12144 MLHHLKELAAVAGIHMILIYLCRFLLRRSRNVLFTVSNSLRFRLKVLTVLLYICLSVMLFYLFGSIMPLPPWGLVVGWVMALIAVELAYAFIFPYSFRYIADNDDDKMVILPV
5984 IRIS_313-12146 MLHHLKELAAVAGIHMILIYLCRFLLRRSRNVLFTVSNSLRFRLKVLTVLLYICLSVMLFYLFGSIMPLPPWGLVVGWVMALIAVELAYAFIFPYSFRYIADNDDDKMVILPV
5985 IRIS_313-12147 MLHHLKELAAVAGIHMILIYLCRFLLRRSRNVLFTVSNSLRFRLKVLTVLLYICLSVMLFYLFGSIMPLPPWGLVVGWVMALIAVELAYAFIFPYSFRYIADNDDDKMVILPV
5986 IRIS_313-12148 MLHHLKELAAVAGIHMILIYLCRFLLRRSRNVLFTVSNSLRFRLKVLTVLLYICLSVMLFYLFGSIMPLPPWGLVVGWVMALIAVELAYAFIFPYSFRYIADNDDDKMVILPV
5987 IRIS_313-12151 MLHHLKELAAVAGIHMILIYLCRFLLRRSRNVLFTVSNSLRFRLKVLTVLLYICLSVMLFYLFGSIMPLPPWGLVVGWVMALIAVELAYAFIFPYSFRYIADNDDDKMVILPV
5988 IRIS_313-12152 MLHHLKELAAVAGIHMILIYLCRFLLRRSRNVLFTVSNSLRFRLKVLTVLLYICLSVMLFYLFGSIMPLPPWGLVVGWVMALIAVELAYAFIFPYSFRYIADNDDDKMVILPV
5989 IRIS_313-12161 MLHHLKELAAVAGIHMILIYLCRFLLRRSRNVLFTVSNSLRFRLKVLTVLLYICLSVMLFYLFGSIMPLPPWGLVVGWVMALIAVELAYAFIFPYSFRYIADNDDDKMVILPV
5990 IRIS_313-12164 MLHHLKELAAVAGIHMILIYLCRFLLRRSRNVLFTVSNSLRFRLKVLTVLLYICLSVMLFYLFGSIMPLPPWGLVVGWVMALIAVELAYAFIFPYSFRYIADNDDDKMVILPV
5991 IRIS_313-12180 MLHHLKELAAVAGIHMILIYLCRFLLRRSRNVLFTVSNSLRFRLKVLTVLLYICLSVMLFYLFGSIMPLPPWGLVVGWVMALIAVELAYAFIFPYSFRYIADNDDDKMVILPV
5992 IRIS_313-12182 MLHHLKELAAVAGIHMILIYLCRFLLRRSRNVLFTVSNSLRFRLKVLTVLLYICLSVMLFYLFGSIMPLPPWGLVVGWVMALIAVELAYAFIFPYSFRYIADNDDDKMVILPV
5993 IRIS_313-12183 MLHHLKELAAVAGIHMILIYLCRFLLRRSRNVLFTVSNSLRFRLKVLTVLLYICLSVMLFYLFGSIMPLPPWGLVVGWVMALIAVELAYAFIFPYSFRYIADNDDDKMVILPV
5994 IRIS_313-12185 MLHHLKELAAVAGIHMILIYLCRFLLRRSRNVLFTVSNSLRFRLKVLTVLLYICLSVMLFYLFGSIMPLPPWGLVVGWVMALIAVELAYAFIFPYSFRYIADNDDDKMVILPV
5995 IRIS_313-12186 MLHHLKELAAVAGIHMILIYLCRFLLRRSRNVLFTVSNSLRFRLKVLTVLLYICLSVMLFYLFGSIMPLPPWGLVVGWVMALIAVELAYAFIFPYSFRYIADNDDDKMVILPV
5996 IRIS_313-12187 MLHHLKELAAVAGIHMILIYLCRFLLRRSRNVLFTVSNSLRFRLKVLTVLLYICLSVMLFYLFGSIMPLPPWGLVVGWVMALIAVELAYAFIFPYSFRYIADNDDDKMVILPV
5997 IRIS_313-12188 MLHHLKELAAVAGIHMILIYLCRFLLRRSRNVLFTVSNSLRFRLKVLTVLLYICLSVMLFYLFGSIMPLPPWGLVVGWVMALIAVELAYAFIFPYSFRYIADNDDDKMVILPV
5998 IRIS_313-12190 MLHHLKELAAVAGIHMILIYLCRFLLRRSRNVLFTVSNSLRFRLKVLTVLLYICLSVMLFYLFGSIMPLPPWGLVVGWVMALIAVELAYAFIFPYSFRYIADNDDDKMVILPV
5999 IRIS_313-12193 MLHHLKELAAVAGIHMILIYLCRFLLRRSRNVLFTVSNSLRFRLKVLTVLLYICLSVMLFYLFGSIMPLPPWGLVVGWVMALIAVELAYAFIFPYSFRYIADNDDDKMVILPV
6000 IRIS_313-12194 MLHHLKELAAVAGIHMILIYLCRFLLRRSRNVLFTVSNSLRFRLKVLTVLLYICLSVMLFYLFGSIMPLPPWGLVVGWVMALIAVELAYAFIFPYSFRYIADNDDDKMVILPV
6001 IRIS_313-12200 MLHHLKELAAVAGIHMILIYLCRFLLRRSRNVLFTVSNSLRFRLKVLTVLLYICLSVMLFYLFGSIMPLPPWGLVVGWVMALIAVELAYAFIFPYSFRYIADNDDDKMVILPV
6002 IRIS_313-12207 MLHHLKELAAVAGIHMILIYLCRFLLRRSRNVLFTVSNSLRFRLKVLTVLLYICLSVMLFYLFGSIMPLPPWGLVVGWVMALIAVELAYAFIFPYSFRYIADNDDDKMVILPV
6003 IRIS_313-12210 MLHHLKELAAVAGIHMILIYLCRFLLRRSRNVLFTVSNSLRFRLKVLTVLLYICLSVMLFYLFGSIMPLPPWGLVVGWVMALIAVELAYAFIFPYSFRYIADNDDDKMVILPV
6004 IRIS_313-12217 MLHHLKELAAVAGIHMILIYLCRFLLRRSRNVLFTVSNSLRFRLKVLTVLLYICLSVMLFYLFGSIMPLPPWGLVVGWVMALIAVELAYAFIFPYSFRYIADNDDDKMVILPV
6005 IRIS_313-12220 MLHHLKELAAVAGIHMILIYLCRFLLRRSRNVLFTVSNSLRFRLKVLTVLLYICLSVMLFYLFGSIMPLPPWGLVVGWVMALIAVELAYAFIFPYSFRYIADNDDDKMVILPV
6006 IRIS_313-12221 MLHHLKELAAVAGIHMILIYLCRFLLRRSRNVLFTVSNSLRFRLKVLTVLLYICLSVMLFYLFGSIMPLPPWGLVVGWVMALIAVELAYAFIFPYSFRYIADNDDDKMVILPV
6007 IRIS_313-12222 MLHHLKELAAVAGIHMILIYLCRFLLRRSRNVLFTVSNSLRFRLKVLTVLLYICLSVMLFYLFGSIMPLPPWGLVVGWVMALIAVELAYAFIFPYSFRYIADNDDDKMVILPV
6008 IRIS_313-12224 MLHHLKELAAVAGIHMILIYLCRFLLRRSRNVLFTVSNSLRFRLKVLTVLLYICLSVMLFYLFGSIMPLPPWGLVVGWVMALIAVELAYAFIFPYSFRYIADNDDDKMVILPV
6009 IRIS_313-12225 MLHHLKELAAVAGIHMILIYLCRFLLRRSRNVLFTVSNSLRFRLKVLTVLLYICLSVMLFYLFGSIMPLPPWGLVVGWVMALIAVELAYAFIFPYSFRYIADNDDDKMVILPV
6010 IRIS_313-12226 MLHQLKELAAVAGIHMILIYLCRFLLRRSRNVLFTVSNSLRFRLKVLTVLLYICLLVMLFYLFGSIMPLPPWGLVVGWVMALIAVELAYAFIFPYSFRYIADNDDDKMVILPV
6011 IRIS_313-12227 MLHHLKELAAVAGIHMILIYLCRFLLRRSRNVLFTVSNSLRFRLKVLTVLLYICLSVMLFYLFGSIMPLPPWGLVVGWVMALIAVELAYAFIFPYSFRYIADNDDDKMVILPV
6012 IRIS_313-12228 MLHQLKELAAVAGIHMILIYLCRFLLRRSRNVLFTVSNSLRFRLKVLTVLLYICLLVMLFYLFGSIMPLPPWGLVVGWVMALIAVELAYAFIFPYSFRYIADNDDDKMVILPV
6013 IRIS_313-12229 MLHHLKELAAVAGIHMILIYLCRFLLRRSRNVLFTVSNSLRFRLKVLTVLLYICLSVMLFYLFGSIMPLPPWGLVVGWVMALIAVELAYAFIFPYSFRYIADNDDDKMVILPV
6014 IRIS_313-12231 MLHHLKELAAVAGIHMILIYLCRFLLRRSRNVLFTVSNSLRFRLKVLTVLLYICLSVMLFYLFGSIMPLPPWGLVVGWVMALIAVELAYAFIFPYSFRYIADNDDDKMVILPV
6015 IRIS_313-12232 MLHHLKELAAVAGIHMILIYLCRFLLRRSRNVLFTVSNSLRFRLKVLTVLLYICLSVMLFYLFGSIMPLPPWGLVVGWVMALIAVELAYAFIFPYSFRYIADNDDDKMVILPV
6016 IRIS_313-12234 MLHHLKELAAVAGIHMILIYLCRFLLRRSRNVLFTVSNSLRFRLKVLTVLLYICLSVMLFYLFGSIMPLPPWGLVVGWVMALIAVELAYAFIFPYSFRYIADNDDDKMVILPV
6017 IRIS_313-12236 MLHHLKELAAVAGIHMILIYLCRFLLRRSRNVLFTVSNSLRFRLKVLTVLLYICLSVMLFYLFGSIMPLPPWGLVVGWVMALIAVELAYAFIFPYSFRYIADNDDDKMVILPV
6018 IRIS_313-12237 MLHHLKELAAVAGIHMILIYLCRFLLRRSRNVLFTVSNSLRFRLKVLTVLLYICLSVMLFYLFGSIMPLPPWGLVVGWVMALIAVELAYAFIFPYSFRYIADNDDDKMVILPV
6019 IRIS_313-12240 MLHHLKELAAVAGIHMILIYLCRFLLRRSRNVLFTVSNSLRFRLKVLTVLLYICLSVMLFYLFGSIMPLPPWGLVVGWVMALIAVELAYAFIFPYSFRYIADNDDDKMVILPV
6020 IRIS_313-12241 MLHHLKELAAVAGIHMILIYLCRFLLRRSRNVLFTVSNSLRFRLKVLTVLLYICLSVMLFYLFGSIMPLPPWGLVVGWVMALIAVELAYAFIFPYSFRYIADNDDDKMVILPV
6021 IRIS_313-12242 MLHQLKELAAVAGIHMILIYLCRFLLRRSRNVLFTVSNSLRFRLKVLTVLLYICLLVMLFYLFGSIMPLPPWGLVVGWVMALIAVELAYAFIFPYSFRYIADNDDDKMVILPV
6022 IRIS_313-12244 MLHHLKELAAVAGIHMILIYLCRFLLRRSRNVLFTVSNSLRFRLKVLTVLLYICLSVMLFYLFGSIMPLPPWGLVVGWVMALIAVELAYAFIFPYSFRYIADNDDDKMVILPV
6023 IRIS_313-12246 MLHHLKELAAVAGIHMILIYLCRFLLRRSRNVLFTVSNSLRFRLKVLTVLLYICLSVMLFYLFGSIMPLPPWGLVVGWVMALIAVELAYAFIFPYSFRYIADNDDDKMVILPV
6024 IRIS_313-12247 MLHHLKELAAVAGIHMILIYLCRFLLRRSRNVLFTVSNSLRFRLKVLTVLLYICLSVMLFYLFGSIMPLPPWGLVVGWVMALIAVELAYAFIFPYSFRYIADNDDDKMVILPV
6025 IRIS_313-12249 MLHHLKELAAVAGIHMILIYLCRFLLRRSRNVLFTVSNSLRFRLKVLTVLLYICLSVMLFYLFGSIMPLPPWGLVVGWVMALIAVELAYAFIFPYSFRYIADNDDDKMVILPV
6026 IRIS_313-12251 MLHHLKELAAVAGIHMILIYLCRFLLRRSRNVLFTVSNSLRFRLKVLTVLLYICLSVMLFYLFGSIMPLPPWGLVVGWVMALIAVELAYAFIFPYSFRYIADNDDDKMVILPV
6027 IRIS_313-12252 MLHHLKELAAVAGIHMILIYLCRFLLRRSRNVLFTVSNSLRFRLKVLTVLLYICLSVMLFYLFGSIMPLPPWGLVVGWVMALIAVELAYAFIFPYSFRYIADNDDDKMVILPV
6028 IRIS_313-12254 MLHQLKELAAVAGIHMILIYLCRFLLRRSRNVLFTVSNSLRFRLKVLTVLLYICLLVMLFYLFGSIMPLPPWGLVVGWVMALIAVELAYAFIFPYSFRYIADNDDDKMVILPV
6029 IRIS_313-12257 MLHQLKELAAVAGIHMILIYLCRFLLRRSRNVLFTVSNSLRFRLKVLTVLLYICLLVMLFYLFGSIMPLPPWGLVVGWVMALIAVELAYAFIFPYSFRYIADNDDDKMVILPV
6030 IRIS_313-12258 MLHHLKELAAVAGIHMILIYLCRFLLRRSRNVLFTVSNSLRFRLKVLTVLLYICLSVMLFYLFGSIMPLPPWGLVVGWVMALIAVELAYAFIFPYSFRYIADNDDDKMVILPV
6031 IRIS_313-12259 MLHHLKELAAVAGIHMILIYLCRFLLRRSRNVLFTVSNSLRFRLKVLTVLLYICLSVMLFYLFGSIMPLPPWGLVVGWVMALIAVELAYAFIFPYSFRYIADNDDDKMVILPV
6032 IRIS_313-12260 MLHHLKELAAVAGIHMILIYLCRFLLRRSRNVLFTVSNSLRFRLKVLTVLLYICLSVMLFYLFGSIMPLPPWGLVVGWVMALIAVELAYAFIFPYSFRYIADNDDDKMVILPV
6033 IRIS_313-12261 MLHHLKELAAVAGIHMILIYLCRFLLRRSRNVLFTVSNSLRFRLKVLTVLLYICLSVMLFYLFGSIMPLPPWGLVVGWVMALIAVELAYAFIFPYSFRYIADNDDDKMVILPV
6034 IRIS_313-12262 MLHHLKELAAVAGIHMILIYLCRFLLRRSRNVLFTVSNSLRFRLKVLTVLLYICLSVMLFYLFGSIMPLPPWGLVVGWVMALIAVELAYAFIFPYSFRYIADNDDDKMVILPV
6035 IRIS_313-12263 MLHHLKELAAVAGIHMILIYLCRFLLRRSRNVLFTVSNSLRFRLKVLTVLLYICLSVMLFYLFGSIMPLPPWGLVVGWVMALIAVELAYAFIFPYSFRYIADNDDDKMVILPV
6036 IRIS_313-12265 MLHHLKELAAVAGIHMILIYLCRFFPSRSRNVLFTVSNSLRFRLKVLTVLLYICLSVMLFYLFGSIMPLPPWGLVVGWVMALIAVELAYAFIFPYSFRYIADNDDDKMVILPV
6037 IRIS_313-12266 MLHHLKELAAVAGIHMILIYLCRFLLRRSRNVLFTVSNSLRFRLKVLTVLLYICLSVMLFYLFGSIMPLPPWGLVVGWVMALIAVELAYAFIFPYSFRYIADNDDDKMVILPV
6038 IRIS_313-12268 MLHHLKELAAVAGIHMILIYLCRFLLRRSRNVLFTVSNSLRFRLKVLTVLLYICLSVMLFYLFGSIMPLPPWGLVVGWVMALIAVELAYAFIFPYSFRYIADNDDDKMVILPV
6039 IRIS_313-12269 MLHHLKELAAVAGIHMILIYLCRFLLRRSRNVLFTVSNSLRFRLKVLTVLLYICLSVMLFYLFGSIMPLPPWGLVVGWVMALIAVELAYAFIFPYSFRYIADNDDDKMVILPV
6040 IRIS_313-12270 MLHHLKELAAVAGIHMILIYLCRFLLRRSRNVLFTVSNSLRFRLKVLTVLLYICLSVMLFYLFGSIMPLPPWGLVVGWVMALIAVELAYAFIFPYSFRYIADNDDDKMVILPV
6041 IRIS_313-12271 MLHHLKELAAVAGIHMILIYLCRFLLRRSRNVLFTVSNSLRFRLKVLTVLLYICLSVMLFYLFGSIMPLPPWGLVVGWVMALIAVELAYAFIFPYSFRYIADNDDDKMVILPV
6042 IRIS_313-12272 MLHHLKELAAVAGIHMILIYLCRFLLRRSRNVLFTVSNSLRFRLKVLTVLLYICLSVMLFYLFGSIMPLPPWGLVVGWVMALIAVELAYAFIFPYSFRYIADNDDDKMVILPV
6043 IRIS_313-12273 MLHHLKELAAVAGIHMILIYLCRFLLRRSRNVLFTVSNSLRFRLKVLTVLLYICLSVMLFYLFGSIMPLPPWGLVVGWVMALIAVELAYAFIFPYSFRYIADNDDDKMVILPV
6044 IRIS_313-12275 MLHHLKELAAVAGIHMILIYLCRFLLRRSRNVLFTVSNSLRFRLKVLTVLLYICLSVMLFYLFGSIMPLPPWGLVVGWVMALIAVELAYAFIFPYSFRYIADNDDDKMVILPV
6045 IRIS_313-12278 MLHHLKELAAVAGIHMILIYLCRFLLRRSRNVLFTVSNSLRFRLKVLTVLLYICLSVMLFYLFGSIMPLPPWGLVVGWVMALIAVELAYAFIFPYSFRYIADNDDDKMVILPV
6046 IRIS_313-12279 MLHHLKELAAVAGIHMILIYLCRFLLRRSRNVLFTVSNSLRFRLKVLTVLLYICLSVMLFYLFGSIMPLPPWGLVVGWVMALIAVELAYAFIFPYSFRYIADNDDDKMVILPV
6047 IRIS_313-12280 MLHHLKELAAVAGIHMILIYLCRFLLRRSRNVLFTVSNSLRFRLKVLTVLLYICLSVMLFYLFGSIMPLPPWGLVVGWVMALIAVELAYAFIFPYSFRYIADNDDDKMVILPV
6048 IRIS_313-12281 MLHHLKELAAVAGIHMILIYLCRFLLRRSRNVLFTVSNSLRFRLKVLTVLLYICLSVMLFYLFGSIMPLPPWGLVVGWVMALIAVELAYAFIFPYSFRYIADNDDDKMVILPV
6049 IRIS_313-12282 MLHHLKELAAVAGIHMILIYLCRFLLRRSRNVLFTVSNSLRFRLKVLTVLLYICLSVMLFYLFGSIMPLPPWGLVVGWVMALIAVELAYAFIFPYSFRYIADNDDDKMVILPV

```
IRIS_313-12283          MLHHLKELAAVAGIHMILIYLCRFLLRRSRNVLFTVSNSLRFRLKVLTVLLYICLSVMLFYLFGSIMPLPPWGLVVGWVMALIAVELAYAFIFPYSFRYIADNDDDKMVILPV
IRIS_313-12284          MLHHLKELAAVAGIHMILIYLCRFLLRRSRNVLFTVSNSLRFRLKVLTVLLYICLSVMLFYLFGSIMPLPPWGLVVGWVMALIAVELAYAFIFPYSFRYIADNDDDKMVILPV
IRIS_313-12285          MLHHLKELAAVAGIHMILIYLCRFLLRRSRNVLFTVSNSLRFRLKVLTVLLYICLSVMLFYLFGSIMPLPPWGLVVGWVMALIAVELAYAFIFPYSFRYIADNDDDKMVILPV
IRIS_313-12286          MLHHLKELAAVAGIHMILIYLCRFLLRRSRNVLFTVSNSLRFRLKVLTVLLYICLSVMLFYLFGSIMPLPPWGLVVGWVMALIAVELAYAFIFPYSFRYIADNDDDKMVILPV
IRIS_313-12287          MLHHLKELAAVAGIHMILIYLCRFLLRRSRNVLFTVSNSLRFRLKVLTVLLYICLSVMLFYLFGSIMPLPPWGLVVGWVMALIAVELAYAFIFPYSFRYIADNDDDKMVILPV
IRIS_313-12288          MLHHLKELAAVAGIHMILIYLCRFLLRRSRNVLFTVSNSLRFRLKVLTVLLYICLSVMLFYLFGSIMPLPPWGLVVGWVMALIAVELAYAFIFPYSFRYIADNDDDKMVILPV
IRIS_313-12289          MLHHLKELAAVAGIHMILIYLCRFLLRRSRNVLFTVSNSLRFRLKVLTVLLYICLSVMLFYLFGSIMPLPPWGLVVGWVMALIAVELAYAFIFPYSFRYIADNDDDKMVILPV
IRIS_313-12290          MLHHLKELAAVAGIHMILIYLCRFLLRRSRNVLFTVSNSLRFRLKVLTVLLYICLSVMLFYLFGSIMPLPPWGLVVGWVMALIAVELAYAFIFPYSFRYIADNDDDKMVILPV
IRIS_313-12291          MLHHLKELAAVAGIHMILIYLCRFLLRRSRNVLFTVSNSLRFRLKVLTVLLYICLSVMLFYLFGSIMPLPPWGLVVGWVMALIAVELAYAFIFPYSFRYIADNDDDKMVILPV
IRIS_313-12292          MLHHLKELAAVAGIHMILIYLCRFLLRRSRNVLFTVSNSLRFRLKVLTVLLYICLSVMLFYLFGSIMPLPPWGLVVGWVMALIAVELAYAFIFPYSFRYIADNDDDKMVILPV
IRIS_313-12296          MLHHLKELAAVAGIHMILIYLCRFLLRRSRNVLFTVSNSLRFRLKVLTVLLYICLSVMLFYLFGSIMPLPPWGLVVGWVMALIAVELAYAFIFPYSFRYIADNDDDKMVILPV
IRIS_313-12297          MLHHLKELAAVAGIHMILIYLCRFLLRRSRNVLFTVSNSLRFRLKVLTVLLYICLSVMLFYLFGSIMPLPPWGLVVGWVMALIAVELAYAFIFPYSFRYIADNDDDKMVILPV
IRIS_313-12299          MLHHLKELAAVAGIHMILIYLCRFLLRRSRNVLFTVSNSLRFRLKVLTVLLYICLSVMLFYLFGSIMPLPPWGLVVGWVMALIAVELAYAFIFPYSFRYIADNDDDKMVILPV
IRIS_313-12300          MLHHLKELAAVAGIHMILIYLCRFLLRRSRNVLFTVSNSLRFRLKVLTVLLYICLSVMLFYLFGSIMPLPPWGLVVGWVMALIAVELAYAFIFPYSFRYIADNDDDKMVILPV
IRIS_313-12301          MLHHLKELAAVAGIHMILIYLCRFLLRRSRNVLFTVSNSLRFRLKVLTVLLYICLSVMLFYLFGSIMPLPPWGLVVGWVMALIAVELAYAFIFPYSFRYIADNDDDKMVILPV
IRIS_313-12302          MLHHLKELAAVAGIHMILIYLCRFLLRRSRNVLFTVSNSLRFRLKVLTVLLYICLSVMLFYLFGSIMPLPPWGLVVGWVMALIAVELAYAFIFPYSFRYIADNDDDKMVILPV
IRIS_313-12303          MLHHLKELAAVAGIHMILIYLCRFLLRRSRNVLFTVSNSLRFRLKVLTVLLYICLSVMLFYLFGSIMPLPPWGLVVGWVMALIAVELAYAFIFPYSFRYIADNDDDKMVILPV
IRIS_313-12305          MLHHLKELAAVAGIHMILIYLCRFLLRRSRNVLFTVSNSLRFRLKVLTVLLYICLSVMLFYLFGSIMPLPPWGLVVGWVMALIAVELAYAFIFPYSFRYIADNDDDKMVILPV
IRIS_313-12307          MLHHLKELAAVAGIHMILIYLCRFLLRRSRNVLFTVSNSLRFRLKVLTVLLYICLSVMLFYLFGSIMPLPPWGLVVGWVMALIAVELAYAFIFPYSFRYIADNDDDKMVILPV
IRIS_313-12308          MLHHLKELAAVAGIHMILIYLCRFLLRRSRNVLFTVSNSLRFRLKVLTVLLYICLSVMLFYLFGSIMPLPPWGLVVGWVMALIAVELAYAFIFPYSFRYIADNDDDKMVILPV
IRIS_313-12309          MLHHLKELAAVAGIHMILIYLCRFLLRRSRNVLFTVSNSLRFRLKVLTVLLYICLSVMLFYLFGSIMPLPPWGLVVGWVMALIAVELAYAFIFPYSFRYIADNDDDKMVILPV
IRIS_313-12311          MLHHLKELAAVAGIHMILIYLCRFLLRRSRNVLFTVSNSLRFRLKVLTVLLYICLSVMLFYLFGSIMPLPPWGLVVGWVMALIAVELAYAFIFPYSFRYIADNDDDKMVILPV
IRIS_313-12312          MLHQLKELAAVAGIHMILIYLCRFLLRRSRNVLFTVSNSLRFRLKVLTVLLYICLLVMLFYLFGSIMPLPPWGLVVGWVMALIAVELAYAFIFPYSFRYIADNDDDKMVILPV
IRIS_313-12313          MLHHLKELAAVAGIHMILIYLCRFLLRRSRNVLFTVSNSLRFRLKVLTVLLYICLSVMLFYLFGSIMPLPPWGLVVGWVMALIAVELAYAFIFPYSFRYIADNDDDKMVILPV
IRIS_313-12319          MLHHLKELAAVAGIHMILIYLCRFLLRRSRNVLFTVSNSLRFRLKVLTVLLYICLSVMLFYLFGSIMPLPPWGLVVGWVMALIAVELAYAFIFPYSFRYIADNDDDKMVILPV
IRIS_313-12321          MLHHLKELAAVAGIHMILIYLCRFLLRRSRNVLFTVSNSLRFRLKVLTVLLYICLSVMLFYLFGSIMPLPPWGLVVGWVMALIAVELAYAFIFPYSFRYIADNDDDKMVILPV
IRIS_313-12322          MLHQLKELAAVAGIHMILIYLCRFLLRRSRNVLFTVSNSLRFRLKVLTVLLYICLLVMLFYLFGSIMPLPPWGLVVGWVMALIAVELAYAFIFPYSFRYIADNDDDKMVILPV
IRIS_313-12323          MLHQLKELAAVAGIHMILIYLCRFLLRRSRNVLFTVSNSLRFRLKVLTVLLYICLLVMLFYLFGSIMPLPPWGLVVGWVMALIAVELAYAFIFPYSFRYIADNDDDKMVILPV
IRIS_313-12324          MLHQLKELAAVAGIHMILIYLCRFLLRRSRNVLFTVSNSLRFRLKVLTVLLYICLLVMLFYLFGSIMPLPPWGLVVGWVMALIAVELAYAFIFPYSFRYIADNDDDKMVILPV
IRIS_313-12325          MLHHLKELAAVAGIHMILIYLCRFLLRRSRNVLFTVSNSLRFRLKVLTVLLYICLSVMLFYLFGSIMPLPPWGLVVGWVMALIAVELAYAFIFPYSFRYIADNDDDKMVILPV
IRIS_313-12329          MLHHLKELAAVAGIHMILIYLCRFLLRRSRNVLFTVSNSLRFRLKVLTVLLYICLSVMLFYLFGSIMPLPPWGLVVGWVMALIAVELAYAFIFPYSFRYIADNDDDKMVILPV
IRIS_313-12330          MLHHLKELAAVAGIHMILIYLCRFLLRRSRNVLFTVSNSLRFRLKVLTVLLYICLSVMLFYLFGSIMPLPPWGLVVGWVMALIAVELAYAFIFPYSFRYIADNDDDKMVILPV
IRIS_313-12332          MLHQLKELAAVAGIHMILIYLCRFLLRRSRNVLFTVSNSLRFRLKVLTVLLYICLLVMLFYLFGSIMPLPPWGLVVGWVMALIAVELAYAFIFPYSFRYIADNDDDKMVILPV
IRIS_313-12334          MLHHLKELAAVAGIHMILIYLCRFLLRRSRNVLFTVSNSLRFRLKVLTVLLYICLSVMLFYLFGSIMPLPPWGLVVGWVMALIAVELAYAFIFPYSFRYIADNDDDKMVILPV
IRIS_313-12336          MLHHLKELAAVAGIHMILIYLCRFLLRRSRNVLFTVSNSLRFRLKVLTVLLYICLSVMLFYLFGSIMPLPPWGLVVGWVMALIAVELAYAFIFPYSFRYIADNDDDKMVILPV
IRIS_313-12337          MLHHLKELAAVAGIHMILIYLCRFLLRRSRNVLFTVSNSLRFRLKVLTVLLYICLSVMLFYLFGSIMPLPPWGLVVGWVMALIAVELAYAFIFPYSFRYIADNDDDKMVILPV
IRIS_313-12340          MLHQLKELAAVAGIHMILIYLCRFLLRRSRNVLFTVSNSLRFRLKVLTVLLYICLLVMLFYLFGSIMPLPPWGLVVGWVMALIAVELAYAFIFPYSFRYIADNDDDKMVILPV
IRIS_313-12341          MLHHLKELAAVAGIHMILIYLCRFLLRRSRNVLFTVSNSLRFRLKVLTVLLYICLSVMLFYLFGSIMPLPPWGLVVGWVMALIAVELAYAFIFPYSFRYIADNDDDKMVILPV
IRIS_313-12342          MLHHLKELAAVAGIHMILIYLCRFLLRRSRNVLFTVSNSLRFRLKVLTVLLYICLSVMLFYLFGSIMPLPPWGLVVGWVMALIAVELAYAFIFPYSFRYIADNDDDKMVILPV
IRIS_313-12344          MLHQLKELAAVAGIHMILIYLCRFLLRRSRNVLFTVSNSLRFRLKVLTVLLYICLLVMLFYLFGSIMPLPPWGLVVGWVMALIAVELAYAFIFPYSFRYIADNDDDKMVILPV
IRIS_313-12345          MLHHLKELAAVAGIHMILIYLCRFLLRRSRNVLFTVSNSLRFRLKVLTVLLYICLSVMLFYLFGSIMPLPPWGLVVGWVMALIAVELAYAFIFPYSFRYIADNDDDKMVILPV
IRIS_313-12346          MLHQLKELAAVAGIHMILIYLCRFLLRRSRNVLFTVSNSLRFRLKVLTVLLYICLLVMLFYLFGSIMPLPPWGLVVGWVMALIAVELAYAFIFPYSFRYIADNDDDKMVILPV
IRIS_313-12347          MLHQLKELAAVAGIHMILIYLCRFLLRRSRNVLFTVSNSLRFRLKVLTVLLYICLLVMLFYLFGSIMPLPPWGLVVGWVMALIAVELAYAFIFPYSFRYIADNDDDKMVILPV
IRIS_313-12348          MLHHLKELAAVAGIHMILIYLCRFLLRRSRNVLFTVSNSLRFRLKVLTVLLYICLSVMLFYLFGSIMPLPPWGLVVGWVMALIAVELAYAFIFPYSFRYIADNDDDKMVILPV
IRIS_313-12349          MLHHLKELAAVAGIHMILIYLCRFLLRRSRNVLFTVSNSLRFRLKVLTVLLYICLSVMLFYLFGSIMPLPPWGLVVGWVMALIAVELAYAFIFPYSFRYIADNDDDKMVILPV
IRIS_313-12350          MLHHLKELAAVAGIHMILIYLCRFLLRRSRNVLFTVSNSLRFRLKVLTVLLYICLSVMLFYLFGSIMPLPPWGLVVGWVMALIAVELAYAFIFPYSFRYIADNDDDKMVILPV
IRIS_313-12351          MLHHLKELAAVAGIHMILIYLCRFLLRRSRNVLFTVSNSLRFRLKVLTVLLYICLSVMLFYLFGSIMPLPPWGLVVGWVMALIAVELAYAFIFPYSFRYIADNDDDKMVILPV
IRIS_313-12352          MLHHLKELAAVAGIHMILIYLCRFLLRRSRNVLFTVSNSLRFRLKVLTVLLYICLSVMLFYLFGSIMPLPPWGLVVGWVMALIAVELAYAFIFPYSFRYIADNDDDKMVILPV
IRIS_313-12353          MLHQLKELAAVAGIHMILIYLCRFLLRRSRNVLFTVSNSLRFRLKVLTVLLYICLLVMLFYLFGSIMPLPPWGLVVGWVMALIAVELAYAFIFPYSFRYIADNDDDKMVILPV
IRIS_313-12354          MLHHLKELAAVAGIHMILIYLCRFLLRRSRNVLFTVSNSLRFRLKVLTVLLYICLSVMLFYLFGSIMPLPPWGLVVGWVMALIAVELAYAFIFPYSFRYIADNDDDKMVILPV
IRIS_313-12355          MLHHLKELAAVAGIHMILIYLCRFLLRRSRNVLFTVSNSLRFRLKVLTVLLYICLSVMLFYLFGSIMPLPPWGLVVGWVMALIAVELAYAFIFPYSFRYIADNDDDKMVILPV
IRIS_313-15896          MLHHLKELAAVAGIHMILIYLCRFLLRRSRNVLFTVSNSLRFRLKVLTVLLYICLSVMLFYLFGSIMPLPPWGLVVGWVMALIAVELAYAFIFPYSFRYIADNDDDKMVILPV
IRIS_313-15897          MLHHLKELAAVAGIHMILIYLCRFLLRRSRNVLFTVSNSLRFRLKVLTVLLYICLSVMLFYLFGSIMPLPPWGLVVGWVMALIAVELAYAFIFPYSFRYIADNDDDKMVILPV
IRIS_313-15898          MLHHLKELAAVAGIHMILIYLCRFLLRRSRNVLFTVSNSLRFRLKVLTVLLYICLSVMLFYLFGSIMPLPPWGLVVGWVMALIAVELAYAFIFPYSFRYIADNDDDKMVILPV
IRIS_313-15899          MLHHLKELAAVAGIHMILIYLCRFLLRRSRNVLFTVSNSLRFRLKVLTVLLYICLSVMLFYLFGSIMPLPPWGLVVGWVMALIAVELAYAFIFPYSFRYIADNDDDKMVILPV
IRIS_313-15900          MLHHLKELAAVAGIHMILIYLCRFLLRRSRNVLFTVSNSLRFRLKVLTVLLYICLSVMLFYLFGSIMPLPPWGLVVGWVMALIAVELAYAFIFPYSFRYIADNDDDKMVILPV
IRIS_313-15901          MLHHLKELAAVAGIHMILIYLCRFLLRLSRNVLFTVSNSLRFRLKVLTVLLYICLSVMLFYLFGSIMPLPPWGLVVGWVMALIAVELAYAFIFPYSFRYIADNDDDKMVILPV
IRIS_313-15902          MLHHLKELAAVAGIHMILIYLCRFLLRRSRNVLFTVSNSLRFRLKVLTVLLYICLSVMLFYLFGSIMPLPPWGLVVGWVMALIAVELAYAFIFPYSFRYIADNDDDKMVILPV
IRIS_313-15903          MLHHLKELAAVAGIHMILIYLCRFLLRRSRNVLFTVSNSLRFRLKVLTVLLYICLSVMLFYLFGSIMPLPPWGLVVGWVMALIAVELAYAFIFPYSFRYIADNDDDKMVILPV
IRIS_313-15904          MLHHLKELAAVAGIHMILIYLCRFLLRRSRNVLFTVSNSLRFRLKVLTVLLYICLSVMLFYLFGSIMPLPPWGLVVGWVMALIAVELAYAFIFPYSFRYIADNDDDKMVILPV
IRIS_313-15905          MLHHLKELAAVAGIHMILIYLCRFLLRRSRNVLFTVSNSLRFRLKVLTVLLYICLSVMLFYLFGSIMPLPPWGLVVGWVMALIAVELAYAFIFPYSFRYIADNDDDKMVILPV
IRIS_313-15906          MLHHLKELAAVAGIHMILIYLCRFLLRRSRNVLFTVSNSLRFRLKVLTVLLYICLSVMLFYLFGSIMPLPPWGLVVGWVMALIAVELAYAFIFPYSFRYIADNDDDKMVILPV
IRIS_313-15907          MLHHLKELAAVAGIHMILIYLCRFLLRRSRNVLFTVSNSLRFRLKVLTVLLYICLSVMLFYLFGSIMPLPPWGLVVGWVMALIAVELAYAFIFPYSFRYIADNDDDKMVILPV
IRIS_313-15908          MLHHLKELAAVAGIHMILIYLCRFLLRRSRNVLFTVSNSLRFRLKVLTVLLYICLSVMLFYLFGSIMPLPPWGLVVGWVMALIAVELAYAFIFPYSFRYIADNDDDKMVILPV
IRIS_313-15909          MLHHLKELAAVAGIHMILIYLCRFLLRRSRNVLFTVSNSLRFRLKVLTVLLYICLSVMLFYLFGSIMPLPPWGLVVGWVMALIAVELAYAFIFPYSFRYIADNDDDKMVILPV
IRIS_313-15910          MLHHLKELAAVAGIHMILIYLCRFLLRRSRNVLFTVSNSLRFRLKVLTVLLYICLSVMLFYLFGSIMPLPPWGLVVGWVMALIAVELAYAFIFPYSFRYIADNDDDKMVILPV
```

**Data S3. Alignment of the XA10 protein from the cultivated rice accession IRBB10A and XA10 homologs from wild rice and cutgrass species.** The alignment was constructed using the CLUSTAL OMEGA algorithm.[8] Dashes indicate the missing sequence or gaps generated by the alignment. Residues common to 80% of proteins are shaded in black using the pyBoxshade algorithm. Transmembrane topology was predicted with DeepTMHMM.[10] Predicted transmembrane helices of XA10 are colored in **red**.

```
L. perrieri IRGC 105164 XA10-1        -----MEMFFVWS-LLLV-AMILAILWLKQLAVAVAAFIHVLVIYLCRFTLLRGGD---D
L. perrieri IRGC 105164 XA10-2        MGRSTQLITF-CVGSILIAVL-AVMIFLNQLA--GAAFVYVLIIYLCRFVLLRRRI---F
O. brachyantha IRGC 101232 XA10-1     -----MQLTALSAGSLLLFASLLAMAYLKLLA--GAACVLVLTIYLCRFLLRRRIF---S
O. brachyantha IRGC 101232 XA10-2     -------------------------MLYLKQLA--GAACVLVLTIYLCRFLLLRRVF---S
O. coarctata TKM-2016 XA10            --MARLEIISACAAPLVIAML-VVMVWLNQLA--VAAYIHVGIIYLCRFVLLRRGG---S
O. longistaminata RD23 XA10           ---MQLIITF-FTGSLVFAVH-VLMVYLKQLA--GAACVDVLIIYLCRFLLLRGIF---F
O. meyeriana Menghai XA10             -----MEVITVCA-LLLLTL-VLVLVCLKQLA--VVACFHVLIIYLGRFVLLHGGD----
O. nivara IRGC 100897 XA10            ---MQLMLTF-CTGPLLFAVL-LLMVYLKQLA--AAACVDVLIIYLCRFLLLRGII---F
O. officinalis W0002 XA10-1           ----MQPITF-CTGPLLFAVL-LLMVYLKQLA--GAACVDVLIIYLCRFLLLRGIF---P
O. officinalis W0002 XA10-2           ----MQPITF-CTGPLLFAVL-LLMVYLKQLA--GAACVDVLIIYLCRFLLLRGIF----
O. punctata IRGC 105690 XA10-1        ---MQLMITFILTGALLFAVLVVLMVCIKQLA--VAACAHALIIYVCRFLLLRGIF----
O. punctata IRGC 105690 XA10-2        ----MELIITVCA-SILVAIIIVAMIFLKQLA--VAACVHVLVIYICRFVLLRGGG---G
O. sativa IRBB10A XA10                ---MQLMLTF-CTGPLLFAVL-LLMVYLKQLA--AAACVDVLIIYLCRFLLLRGII---F

L. perrieri IRGC 105164 XA10-1        G------DNLCKFRIKVAVGIMYISLSAILLCLFAPDM-LPPWGRVGHGPRCH-------
L. perrieri IRGC 105164 XA10-2        SDH-----SNILFRVKAGIGLMYIFLLAILLYISAAVMKLPPWGAVAMSVMAFAAVELGY
O. brachyantha IRGC 101232 XA10-1     DDH----HNHLRFRLKVGIGFMYISLSAILFYLSASVMALPPWGAAAMWGMALVAVELVY
O. brachyantha IRGC 101232 XA10-2     DDH----K--LRFRLKVGIGFLYISLSAVLFYLSAAVMALPPWGAVAMWGMVVVAVELAY
O. coarctata TKM-2016 XA10            GDK------LTRFRLKLAIGILYISLSGMLVYLFSPVMAAAPWGGVAVCATALVATELGF
O. longistaminata RD23 XA10           SDD-----DKLRFRVKVAIGFLYISLSAILFYLSAAVMALPPWGAVAIWGMALVAIELGY
O. meyeriana Menghai XA10             -------DRLRCFRIKVAVGILYLTLSAILFCLFGPV--MPPWGAVAAWAMSLAAIELGY
O. nivara IRGC 100897 XA10            SGD-----GKLRFRVKVAIGFLYISLSAILFYLSAAVMALPPWGAVAMWGMALVATELGN
O. officinalis W0002 XA10-1           DDD------KLRFRVKVAIGTTTERIFADG-----RIRSSHAVGRAAC------AKR---
O. officinalis W0002 XA10-2           SDD-----DKLRFRVKVAIGFLYISLSAILFYLSAAVLALPPWGAVAMWGMALVASELGY
O. punctata IRGC 105690 XA10-1        SDD-----GKLRLRVKVAIGFLYISLSAILFYLSAAVMALPPWGVVAMWGMALVATELGY
O. punctata IRGC 105690 XA10-2        SSSGDAERPLRCFRIRVAVGILYLSMSAILFGLFVSV--MPPWGAVAVWGMALAAVELGY
O. sativa IRBB10A XA10                SGD-----GKLRFRVKVAIGFLYISLSAILFYLSAAVMALPPWGAVAMWGMALVATELGY

L. perrieri IRGC 105164 XA10-1        ------------------------------
L. perrieri IRGC 105164 XA10-2        ALFFPYTCRCIDDLDEVNPPV---------
O. brachyantha IRGC 101232 XA10-1     AFFFPYSCICIGDDADETPPLSV-------
O. brachyantha IRGC 101232 XA10-2     AFFFPYSCICIGDDADKTPPLSV-------
O. coarctata TKM-2016 XA10            AFLSSYSC-CIDEDEESR-PL---------
O. longistaminata RD23 XA10           AFFFPYRCLCIRKDDDEETPSVSV------
O. meyeriana Menghai XA10             AFFFPYSCYHIDDEHEEMMINPTVGVSV--
O. nivara IRGC 100897 XA10            SFLCPYSCRCIGEDDEEISPV---------
O. officinalis W0002 XA10-1           -SVKITICACGRS-----------------
O. officinalis W0002 XA10-2           AFFFPYTCLCIGEDDDEEIPSVSV------
O. punctata IRGC 105690 XA10-1        AFFFPYRCLCIGDDDDEITPVDLRSITISA
O. punctata IRGC 105690 XA10-2        AFFFPYTCYHVGGDDDDDDDNIPITH-I--
O. sativa IRBB10A XA10                SFLCPYSCRCIGEDDEEISPV---------
```

**Data S4. Alignment of the XA10-Ni protein from the cultivated rice accession Nipponbare and XA10-Ni homologs from wild rice species.** The alignment was constructed using the CLUSTAL OMEGA algorithm.[8] Dashes indicate the missing sequence or gaps generated by the alignment. Residues common to 80% of proteins are shaded in black using the pyBoxshade algorithm. Transmembrane topology was predicted with DeepTMHMM.[10] Predicted transmembrane helices of XA10-Ni are colored in **red**.

```
O. australiensis KR-Keep River 1 XA10-Ni --MKLTGCALILIAVHVA-AAISCPEPLAAMAVVDVPLLFLCRYLLRRRDDD--------
O. glumipatula GEN 1233 XA10-Ni          -MELITACALLLIAVHIA-IVISCPEPLAAVALVDVPLLYLCRYLLIIRRGVIIVSGA--
O. longistaminata RD23 XA10-Ni           -MELITACALLLIAVHIA-IVISCPEPLAAVAFVDVPLLYLCRYLLIIRPGVIIVSGAG-
O. longistaminata W11 XA10-Ni            --MKLNLCALLVIAVHVA-AAISCPKPLAAMAIIDVPLLFLCRYLIRRRDDDFVVSS---
O. meridionalis OR44 W2112 XA10-Ni       -MELITACALLLIAVHIA-IVISCPEPLAAVALVDVPLLYLCRYLLIIRRGVIIVSGAGA
O. meyeriana Menghai XA10-Ni             ----------MLITVHIL-IVISCPELFTAIALVDVSILHLCRYLLHH--GVSVISG---
O. nivara WR04-6 XA10-Ni                 -MELITACALLLIAVHIA-IVISCPEPLAAVALVDVPLLYLCRYLLIIRRGVIIVSGAGA
O. officinalis W0002 XA10-Ni             --MVLTAYALLLILLHVA-IGISCPEPLAAMAFVDVPLLFLCRYLLLRGASVFFA-----
O. rufipogon Taxon A XA10-Ni             MEILITACALLLIAVNVVVTAISCPNPLAAIVLVDVPVLYLCRYLLRRGV--PVVT----
O. rufipogon W1943 XA10-Ni               -MELITACALLLIAVHIA-IVISCPEPLAAVALVDVPLLYLCRYLLIIRRGVIIVSGAGA
O. sativa Nipponbare XA10-Ni             -MELITACALLLIAVHIA-IVISCPEPLAAVALVDVPLLYLCRYLLIIRRGVIIVSGAGA

O. australiensis KR-Keep River 1 XA10-Ni --GGGRLKFRLKAAVLVLYMVMSALFFLRFAPI--TPWWGAVAAWAMMLLILELIFAFFF
O. glumipatula GEN 1233 XA10-Ni          GGGGRLRRFRLGAAMAIIYMAMSTLFFLRIAPL--TPWWGALAAWVMILLIVEAIF---A
O. longistaminata RD23 XA10-Ni           G-SGRMRRFRLGAAMAIIYMAMSTLFFLRIAPL--TPWWGALAAWVMILLIVEAIF---A
O. longistaminata W11 XA10-Ni            GGGGGRLQFQLKAAVLVLYMVMSALFFLRFAPI--TPWWGAVAAWAMMLLILEVIFAFFF
O. meridionalis OR44 W2112 XA10-Ni       GGGGRLRCFRLGAAMAIIYMAMSTLFFLRIAPL--TPWWGALAAWVMILLI---------
O. meyeriana Menghai XA10-Ni             --AGIRLRFRLRTADAIIYMALSVLLFLRFAPV--MSWWGALAAWVTILLIVEAVLAFFF
O. nivara WR04-6 XA10-Ni                 G-GGRLRRFRLGAAMAIIYMAMSTLFFLRIAPL--TPWWGALAAWVMILLIVEAIF---A
O. officinalis W0002 XA10-Ni             --GDGRLRFRLKAAVAILYMALSALFFLRFAPL--MPWWGVLVAWVTMVLIVEFVYAFFF
O. rufipogon Taxon A XA10-Ni             GAGVVRLRFRLVAAVATIYMAMPALFFLRFAPV--MPWWGGARRLGDHPAHRRGRLRLLL
O. rufipogon W1943 XA10-Ni               G-GGRLRRFRLGAAMAIIYMAMSTLFFLRIAPL--TPWWGALAAWVMILLIVEAIF---A
O. sativa Nipponbare XA10-Ni             G-GGRLRRFRLGAAMAIIYMAMSTLFFLRIAPL--TPWWGALAAWVMILLIVEAIF---A

O. australiensis KR-Keep River 1 XA10-Ni -----FPYSHCNKLGWDVENPGGILT
O. glumipatula GEN 1233 XA10-Ni          F--FFPYRCLLQRNRRRCPE------
O. longistaminata RD23 XA10-Ni           F--FFPYRCCFNETEGDVQNNPHV--
O. longistaminata W11 XA10-Ni            -----FPDGYSNKLRWDVENPGGILA
O. meridionalis OR44 W2112 XA10-Ni       ----LPYRCCFNETEGDVQNNPHV--
O. meyeriana Menghai XA10-Ni             F--FFPYGCCNERPEGDVHN-PHV--
O. nivara WR04-6 XA10-Ni                 F--FFPYRCCFNETEGDVQNNPHV--
O. officinalis W0002 XA10-Ni             -----HYSCYN-ELDGYVENPVVG--
O. rufipogon Taxon A XA10-Ni             LSLQLLQRSA----------------
O. rufipogon W1943 XA10-Ni               F--FFPYRCCFNETEGDVQNNPHV--
O. sativa Nipponbare XA10-Ni             F--FFPYRCCFNETEGDVQNNPHV--
```

**Data S5. Alignment of the XA23 protein from the cultivated rice accession Nipponbare and XA23 homologs from wild rice species.** The alignment was constructed using the CLUSTAL OMEGA algorithm.[8] Dashes indicate the missing sequence or gaps generated by the alignment. Residues common to 80% of proteins are shaded in black using the pyBoxshade algorithm. Transmembrane topology was predicted with DeepTMHMM.[10] Predicted transmembrane helices are colored in **red**. Amino acid substitutions within the putative transmembrane helices that might lead to the loss of cell death execution by XA23 proteins are shaded in yellow.

```
O. barthii IRGC 105608 XA23          -MLHLKEL-AAVAGIDMIVIYLCRFLLRRCRS-R-NASNSLRFRLKVLTVLLYISLSTML
O. glaberrima IRGC 96717 XA23        -MLHLKEL-AAVAGIDMIVIYLCRFLLRRCRS-R-NASNSLRFRLKVLTVLLYISLSTML
O. glumipatula GEN 1233 XA23         MLHHLKEL-AAVAGIHMILIYLCRFLLRRSRNVLFTVSNSLRFRLKVLTVLLYICLSVML
O. longistaminata RD23 XA23-1        MLHHLKEL-AAVAGTHMIVIFLCRFLLRLRRR---SASNSLRFRLKVVTVLLYISLSAML
O. longistaminata RD23 XA23-2        -MLHLKEL-AAVAGIHMILIYLCRFLLRLRRNASFTTSNSLRFRLKVLTALLYISLSAML
O. longistaminata RD23 XA23-3        MLHHLKEL-AAVAGIHMILIYLCRFLLRLRRR---SASTSLRFRLKVLTVLLYISLSAML
O. longistaminata RD23 XA23-4        -MLHLKEL-AAVAGIHMIVIYLCRFLLRLRHS-AFTTSNSLRFRLKVLTALLYISLSAIA
O. longistaminata RD23 XA23-5        -MLHLKELTAAIAGVHMIVIYLCRFLLRLRRS-AFTTSNSLRFRLKVLIVLLYVSLSTML
O. meridionalis OR44 W2112 XA23      MLHHLKEL-AAVAGIHMILIYLCRFLLRRSRNVLFTVSNSLRFRLKVLTVLLYICLSVMM
O. nivara WR04-6 XA23                MLHHLKEL-AAVAGIHMILIYLCRFLLRRSRNVLFTVSNSLRFRLKVLTVLLYICLSVML
O. officinalis W0002 XA23-1          -MLHLKKL-AAVTGIHMIVIYLCRFLLRRCRS-ASTTSNSLCFRLKILTALLYVSLSAML
O. officinalis W0002 XA23-2          -MLHLKKL-AAVAGIHMIVIYLCRFLLRRCRS-ASTTSNSLRFRLKILTALLYVSLSAIL
O. punctata IRGC 105690 XA23         -MLHMKEL-AAVAGIHMIVIYLCRFLLRRS---ASTTSNSLRFRLKVLTVLLYVSLSAML
O. rufipogon Taxon A XA23            MLHHLKEL-AAVAGIHMILIYLCRFLLRLRH----NASNSLRFRLKVLTALLYISLSAML
O. rufipogon W1943 XA23              ----------------MILIYLCRFLLRRSRNVLFTVSNSLRFRLKVLTVLLYICLSVML
O. sativa Nipponbare XA23            MLHHLKEL-AAVAGIHMILIYLCRFLLRRSRNVLFTVSNSLRFRLKVLTVLLYICLSVML

O. barthii IRGC 105608 XA23          LYLFGSIMPLPPWGALAG------------------------------------------
O. glaberrima IRGC 96717 XA23        LYLFGSIVPLPPWGALAGWVMALIAVELAYAFLFPYSLRYIAHNDDEDKMVIPPV-----
O. glumipatula GEN 1233 XA23         FYLFGSIMPLPPWGLVVGWVTALIAVELAYAFIFPYSFRYIADNDDD-KMVILPV-----
O. longistaminata RD23 XA23-1        FYLFASIMPLPPWGAVAGWVMALIAVELAYAFLFPYSLRYIDDD----NMVIDPPV----
O. longistaminata RD23 XA23-2        FCLFASIMPLQPWGAVAGWVMALIAVELAYAFLFPYCLCYFADNHDD-KMVIPPVYALKK
O. longistaminata RD23 XA23-3        FYLFASIMPLPPWGAVVGWVMALMAVELAYAFIFPYNLRYIADNGD-DKMVI-PPV----
O. longistaminata RD23 XA23-4        VLPVCLHHAAATVGRCGRLGHGP--HRRRARLRLPFSI----------------------
O. longistaminata RD23 XA23-5        FYLFASIMPLPPWGAVAGWVMALIAVELAYAFLFPYSLGYIPENND-DKMVIPPV-----
O. meridionalis OR44 W2112 XA23      FYLFGSIMPLPPWGLVVGWVMALIAVELAYAFIFPYSFRYIADNDDD-KMVILPV-----
O. nivara WR04-6 XA23                FYLFGSIMPLPPWGLVVGWVMALIAVELAYAFIFPYSFRYIADNDDD-KMVILPV-----
O. officinalis W0002 XA23-1          FYLFASIMPLPPWGAVAGWVMALIAVELAYASLFPYSLRYINENDD-EEMVTPPV-----
O. officinalis W0002 XA23-2          FYLFASIMPLPPWGAVAGWGVALIAVELAYAFLFPYSLRYIADNDD-EMMVIPPV-----
O. punctata IRGC 105690 XA23         FYLFASIMPLPPWGAVAGWVMALIAVELAYAFLFPYSLRYIADNDD-DMVV-PPV-----
O. rufipogon Taxon A XA23            FYLFASIMPLPPWGAVAGWVMALFAVELAYAFIFPYNLRYIADNHDEDNMVI-PPV----
O. rufipogon W1943 XA23              FYLFGSIMPLPPWGLVVGWVMALIAVELAYAFIFPYSFRYIADNDDD-KMVILPV-----
O. sativa Nipponbare XA23            FYLFGSIMPLPPWGLVVGWVMALIAVELAYAFIFPYSFRYIADNDDD-KMVILPV-----

O. barthii IRGC 105608 XA23          ---------
O. glaberrima IRGC 96717 XA23        ---------
O. glumipatula GEN 1233 XA23         ---------
O. longistaminata RD23 XA23-1        ---------
O. longistaminata RD23 XA23-2        LKPHILSLQ
O. longistaminata RD23 XA23-3        ---------
O. longistaminata RD23 XA23-4        ---------
O. longistaminata RD23 XA23-5        ---------
O. meridionalis OR44 W2112 XA23      ---------
O. nivara WR04-6 XA23                ---------
O. officinalis W0002 XA23-1          ---------
O. officinalis W0002 XA23-2          ---------
O. punctata IRGC 105690 XA23         ---------
O. rufipogon Taxon A XA23            ---------
O. rufipogon W1943 XA23              ---------
O. sativa Nipponbare XA23            ---------
```

**Data S6. Alignment of genomic sequences upstream of the *Xa10*-like loci from cultivated rice, wild rice, and cutgrass species.** 400 nucleotides upstream of start codons (**ATG**) of the *Xa10*-like loci were aligned using the MUSCLE algorithm.[11] Nucleotides common to 80% of sequences are shaded in black, while nucleotides common to 60% of sequences are shaded in grey using the pyBoxshade algorithm. The start codons (**ATG**) of each *Xa10*-like locus are colored in **red**. The AvrXa10 binding element for the activation of *Xa10* transcription in *O. sativa* cv. IRBB10A is shaded in magenta.[12]

```
                                     1 ........10........20........30........40........50........60........70........80........90
L. perrieri IRGC 105164 Xa10-1       1 ..................................................ACCATCATAACTGCTGACTAGTATTTTTTTTTCTGTCAC.
L. perrieri IRGC 105164 Xa10-2       1 TGCCGGGCCAGGGGAACTCGGCGCGGCGGCGGAGAGGAGCATGGCGCGGAAGCGCCTCCCCTTGAGGATGAATCTGTGCGTCTCGTGACG
O. brachyantha IRGC 101232 Xa10-1   1 .................GCCCATTTATTAATAGAAACCCAAGCATAGAGAGCCTGCAGAGGTTGAGAGCAATATATAAATCCCTGCGTCC
O. brachyantha IRGC 101232 Xa10-2   1 ...................................................................................CAATAGT
O. coarctata TKM-2016 Xa10           1 ......................ACTTTATCTATAATGT........TACTATTTTCATACCTGTTGATAAACGTAGAACA...CCTCTAA
O. longistaminata RD23 Xa10         1 ..............................CGTGTACATAAACGTTCCTCATCCCAGTTCCTCCTCATCGAACTTGTTAGCGGAACCTGA
O. meyeriana Menghai Xa10           1 .......................................................ACTACCTCCTGTGAAATATTAAATTATACGTGGTG
O. nivara IRGC 100897 Xa10          1 ........................TGAGGTCGCAATCTGACTTGGGTTCAAGCTTTCTCC.......................AGCCCCT
O. officinalis W0002 Xa10-1         1 .........................CTCACGGTGTAAATATTAGTACGTAGGATTAATCCCACATAGAGAGCTTACAAAGGTTGAGAGCA
O. officinalis W0002 Xa10-2         1 ....................ATTCCAAAACTAAAAGGAGCAAGAACTGCGATTCCAAAACTAAAAGG..AGCAAGAACAGTTCTACTAAA
O. punctata IRGC 105690 Xa10-1      1 ......................................................CATTTATTATATGCTACTGCAGTTCTTCCTGTAACA
O. punctata IRGC 105690 Xa10-2      1 ............................TTAACTATATCCATTTAATTAGTACCTAATTAACAATATATGATACAGTAATTACAGTAAGT
O. sativa IRBB10A Xa10              1 ....................................CACCAAAAGGTCCAACACCAAAAACTTATTGATGTCGCAATCACTTCAATTAC.

                                    91 .......100.......110.......120.......130.......140.......150.......160.......170.......180
L. perrieri IRGC 105164 Xa10-1      40 .............TACTTTCTTGACAGGACAGAAACTG.....AGTATTTTCGTGTACATTCTA....CAAAAGACAATC..........
L. perrieri IRGC 105164 Xa10-2      91 ACGTGGGGCCCATTCTTTGCCGAGC.TGAGATAATTGAGGAGGATGCTATATACGGTTGAAATA....CAGCCAGATGCC..........
O. brachyantha IRGC 101232 Xa10-1  74 TAACCGCAGTGCTCACGTTTAGCAA.TAGTCGAATAAACAGATGATGTCCCAAAGTTTAAATTA....AATTGGATTGGA..........
O. brachyantha IRGC 101232 Xa10-2   8 AGAATAAACAGATGATGTCCCAAAG..................ATTAAATTGGATTAAACTGAC....GTCAGAATCGTC..........
O. coarctata TKM-2016 Xa10          58 CGGATAAAAGATTATCATTTAAAAC.TG.......TTATGGATAAGACCACCCACTATACCATA....ATTACAGAT.............
O. longistaminata RD23 Xa10        61 ATAGCGAAGACGTCACTTTCCTATA..AACAACTTTCGAAGACACCACTTTCCTGTAAACAACT....TTCCTCCTCATC..........
O. meyeriana Menghai Xa10          36 AAACTAAAATATTAGTCATTGAAACATTTAAAAATTT.TGTGAAACACATTGAAATACTCGTTA..........................
O. nivara IRGC 100897 Xa10         44 TTGCATCAAAATTGATGTCGCAAA.............ATTTCAATAACTTCAATTGAGGTCGCA....ATCTGACTTGGG..........
O. officinalis W0002 Xa10-1        66 ATATATAAATCCTTGCGTTCTAGCC...ACAGTATATGGGGAAAAATATTTGTGGGGTGTTT....ATTAGATATGCA..........
O. officinalis W0002 Xa10-2        69 AGGAGCAAGAACTGCGATTCCAAAA.CTAAAAGGAGCAAGAACAGTTCTTCTGATGAAAGGCAA....ATTCCAAAAGGAGCAAGAACTG
O. punctata IRGC 105690 Xa10-1     37 CTGTTTCACACTGGATGTTCTATATGTTACAATATTTCTGTGTAACACAT...ACTATGTGCTA....CTGCCGCATTCC..........
O. punctata IRGC 105690 Xa10-2     63 TTAACCGACGAGTATTCCTCGG.........GCATATGTCAACAACCAATCAGACTATGCTTCAGGCTGCCTGACCAGTC..........
O. sativa IRBB10A Xa10             54 ...................................TTATTGATGTCGCAATCACGTTCACTCCT....CTTATATATACA..........

                                   181 .......190.......200.......210.......220.......230.......240.......250.......260.......270
L. perrieri IRGC 105164 Xa10-1      98 ...TCGGATAATTAATTTGGTAATTTCCGCGAGC....................GTGGTAAGCCTGTGGGCTGTGGCTAGTGGAGCATGA
L. perrieri IRGC 105164 Xa10-2     166 ...CAGATCCATCCACACGAATCATGGTA.........................AGATCTAACGCTTCGTAGGTCATGTGGAGGGATAA
O. brachyantha IRGC 101232 Xa10-1 149 ...TTAAACTGATGTCCCAAAGTTTGA...........................GAGCTAGGAATTTTGCAATTCCACATCCTTGTG...
O. brachyantha IRGC 101232 Xa10-2  66 ...AAAGAAAATCAGCCCGCGTAGCGATG.........................ACAGCGATAGCAACATATGGTCGTAGCCACAAGCGA
O. coarctata TKM-2016 Xa10         123 .....ATCATTATCTCTGGTATGTATTA..........GTTATGACCTGTCAATGTAGTGAAAATTTAAATTTTGAAATTTTGCAAA...
O. longistaminata RD23 Xa10       135 ...CCAGTTCCTTCTCATCCAACTTGTTAGCATAACCTGA..............ACAGCGAAGACTTTCCTTGTCAGAGCGACT......
O. meyeriana Menghai Xa10          99 ......AAACATTTCTATATCACATATTAA........................ATATTAAATTTCTATACCGTGAGAGGCCACGAAGAG
O. nivara IRGC 100897 Xa10        107 ...TTCAAGCTTTCTCCAGCCCCTTTGCATCA......................AAATTGAAGTGATTGCTTCTTTTGGTGTAACTGGAA
O. officinalis W0002 Xa10-1       139 ..............TGAAGCTATATATTACC.......................GCTTATAACGCGATTTCGCTGTCGAATCGCAACCAG
O. officinalis W0002 Xa10-2       154 CGGCAAATTTTTCCTCAGAGCGTGTGTTCTGAGCGTGTTTC.............AGAGTCAGAACGATGAAAGGCAAATTCCTCAGAGCG
O. punctata IRGC 105690 Xa10-1    110 ...CCTGTAACACGTTTAGTCATGTGTTGCAAGTTTGT................GCTGCAATAGCTTTGTTCTGTAGTAGTAGTGCTGTT
O. punctata IRGC 105690 Xa10-2    134 ...TAAATTCATTGGTTTGTCAAGTAGGTCTCAAGC..................ACATAAAAATCTCAAGCAGTTTACTGATCAGGG...
O. sativa IRBB10A Xa10              95 ...TCAATTACTTATTCTCTTATATAT............................ACATCAATTA..........................

                                   271 .......280.......290.......300.......310.......320.......330.......340.......350.......360
L. perrieri IRGC 105164 Xa10-1     165 TATACAGTTT............ATTTATATATTCGCCTAATCCCATCAAAGCCGTCTGAAAGAATGCATAAAGAAAGCCGGCCAAGCCCA
L. perrieri IRGC 105164 Xa10-2     228 AAG....ACT............TCTCGTCCACGACAACGATT..............................GAAGGATCGGTGA...AA
O. brachyantha IRGC 101232 Xa10-1 206 .......TTT............CGTTGCGTCGAAAGTCAATTCCCCT.........................GAAAGATCTCAAATTAAA
O. brachyantha IRGC 101232 Xa10-2 128 CGA....CCT............CAAGGGCAGCGATGTAAGTTAACTCATA.........................AGAGTGGCGATGTAA
O. coarctata TKM-2016 Xa10         195 ......................CGATTTTGGATGAAGATATGTCCTCCTATATTAAAGTTGTAAATCAAAGTAATAGATTTTTGA...CA
O. longistaminata RD23 Xa10       202 .......CTT............TACTTTCCTTTAAACAACTTTCCTCCTC......................ATCGAACTTGTTAGCATA
O. meyeriana Menghai Xa10         159 AAAAAGCATCTCCGTGACTCCGTCTCGCTGGCTAACTCTGTGATATCATA.........................CGTATGTGGAGAAAA
O. nivara IRGC 100897 Xa10        172 TAG....GCT............CCAGCCCCTTTGCATCAATGGACTCCTC.....................ATGTGCATGTTTGA.....
O. officinalis W0002 Xa10-1       192 GTTTCGCGTT............TCGCCTTAAACAGCCAGGTACTTTCTTT......................................CC
O. officinalis W0002 Xa10-2       231 GTG....TTT............CAATACTCGATACTCGATTTGCGGCCTA....................AATATGAACTAACGGCAAAT
O. punctata IRGC 105690 Xa10-1    181 GAAATGTCCC....CGAAGTCCAGTTACTTGAAAGCTACATGCATTCTTA.........ATTAACCGATGGTGACAAACTGTCAATCCCA
O. punctata IRGC 105690 Xa10-2    200 .............................CACCAATCAAATAGACCCTTA...........................ATTTGTTA...CG
O. sativa IRBB10A Xa10             129 .......CTT............ATTGATGTCGCAATCACGTTCACTCCTC........................................

                                   361 .......370.......380.......390.......400.......410.......420.......430.......440.......450
L. perrieri IRGC 105164 Xa10-1     243 CCATTGT.................CCATCATATGATATAATTTAAATTAAAATAAAACAAAATAAATAAATAAATAAAACAAACTAAAGG
L. perrieri IRGC 105164 Xa10-2     269 ACGATCCAACGATCCATGA.....CAACGACGACACTCGATT.................GACCAGCCTTCTTTG................
O. brachyantha IRGC 101232 Xa10-1 252 ...TTTGAGCTTTCTTATA.....TCCCGAACTGTTCCCCTTGAAACCGAGG......................................
O. brachyantha IRGC 101232 Xa10-2 177 TTTAATAGA...............TAGCGATGTAATTTAACTAGGTACCAGAACCCCATATATTTTTATATATT................
O. coarctata TKM-2016 Xa10         260 ACATCTAAAAATTTATAAC.....TAATAATAT..TTTTATTTAAAATTATTTACATCACCAAATATACATTAT................
O. longistaminata RD23 Xa10       251 ACCTGAA.................CAGCGAAGACTTTCCTCGTCAGAGCGACTCTATAAACACAGCGGAACAGA................
O. meyeriana Menghai Xa10         224 TTGGCCATGCATCATTGACGGCCGTGCAGATGTAATTTCATTAAAA........................CAAT................
O. nivara IRGC 100897 Xa10        220 ..................................GTTTTGTGAAAAGCATAGAAAAAAAAAGCACTTCATTCTT................
O. officinalis W0002 Xa10-1       232 TCCTCAGAACGGTGTAAACTTACTTTCTTTCTTTCCTCCTCCTCAGAACGGTGTAAAGTACTCCTCCTCAGAAC................
O. officinalis W0002 Xa10-2       285 TCCTCAGAGCGGTGTTTCG.....TCGCAGGTACTTTCCTCCTCAGAGCG........................................
O. punctata IRGC 105690 Xa10-1    258 TGTCCCGA................TTTCCGAAGACTTTCACCTAGGGTCTCTGGCAAATAGCCATGTATGC...................
O. punctata IRGC 105690 Xa10-2    231 TTCTCAAAGTATATATATAGCCTTCCTTCAGTATATTATATTAGGGAGGAGAGAGAGAGAGAGAGAGAGCGC................
O. sativa IRBB10A Xa10             160 ...............................................................TTATATATACA................

                                   451 .......460.......470.......480.......490.......500.......510.......520.......530.......540
L. perrieri IRGC 105164 Xa10-1     316 TGCAAGGATGCGAGAAATCTCGCTCTTTATATACACGTACATATGCATGC..........ATCGGAA.............GGGAAGGAGT
L. perrieri IRGC 105164 Xa10-2     321 ..........AGTGTTGTCCCGTGAATGACGAAT..CCAACGGCCTACGC..CTACGTAACTCTCA.............GTGCGAAGCT
O. brachyantha IRGC 101232 Xa10-1 297 ...................CCTTCTTATATATATCCC...GAACTGTGGC..CTTCTTGATCTTCA.............GAAAGCTTCT
O. brachyantha IRGC 101232 Xa10-2 236 ...............ATTCTATTATCAGGTATATAGCTAC...TCCTTGCACGTTCTTAATATGAA.............AAACATTATA
O. coarctata TKM-2016 Xa10         327 ..........AAC.........CGATAGATGTATTTTGAAAAACAAATGC..GAACC..ACTCGAA.............GAAAGAAGAA
O. longistaminata RD23 Xa10       308 ..........GACTCTATAAACACAGCGGAACATGCGAAAAAAAACTTGC..AGTCCTGCCAGACA.............GAGCGGTGTT
O. meyeriana Menghai Xa10         274 ..........GGCACCACCACATATTATATATATCAGCTGAGACAAAGAG..AAGCTAAGCTAAAG.............CAGCAATTAA
O. nivara IRGC 100897 Xa10        260 ..........CGTTTGCACTCCTCTTATATATACAGCCAGAAAGCACTTC..ATTCTTCATTAGCACAGCCGAAACTCGCGAGAGATCCT
O. officinalis W0002 Xa10-1       306 ..........GGTCTAAAGTACTCGGTACTCTTTCCTCCTCAGTACTCGGTACTCTTTCCTCCTCA.............GAACGGTGTT
O. officinalis W0002 Xa10-2       330 ...........GTGTTTCGTCCTCGTATATATATGCC........................TCCTCA.............GAACGGTGTT
O. punctata IRGC 105690 Xa10-1    313 ..........................CTATATATACCCGAAACATTCTCA..GCTCGAAGCTAAAA.............GAAACTCGAT
O. punctata IRGC 105690 Xa10-2    305 ..........ACTGCAGTGGCTCGATCAGTGAGCGCCGTGCAGACTAGAG..ATATCGACTTATCG............AGGAAGCAGTA
O. sativa IRBB10A Xa10             171 ..........CACGTTCACTCCTCTTATA[illegible]A[illegible]CAGCCAGAAAGCACTTC..ATTCTTCATTAGCACAGCCGAAACTCGCGAGAGATCCT
```

```
                                          541 .......550.......560.......570.......580.......590.......600.......610.......620.......630
L. perrieri IRGC 105164 Xa10-1            383 AGTAGC........................TTCCAT........................AAACAT........................
L. perrieri IRGC 105164 Xa10-2            383 CTCTCT........................CTCCCT........................CTCTCA........................
O. brachyantha IRGC 101232 Xa10-1         349 TGTATC........................CCAGTCGTGAAAAAAAGCTCAG....TTTCATTTCGTCGCAGGTAG..............
O. brachyantha IRGC 101232 Xa10-2         294 TATATCCGGTTCGATGCCCGTCCCCCAGGACTCCCC..CCACGAAGACGCAGCTGACACCATTTTG.CGCCGGATCTCTCCTCCTGTCTG
O. coarctata TKM-2016 Xa10                380 AGAATT........................TTCCTA........................CGATCGACA.....................
O. longistaminata RD23 Xa10               372 T.CATC........................CTCCTC........ATATTCAG........ATTTCT........................
O. meyeriana Menghai Xa10                 338 C.CATC......................ATACCCCTGATCACTTTGATTGAG........ATTCGGTCGGCGGCCGGCGAGCTAGG....
O. nivara IRGC 100897 Xa10                338 C.CTCC........................CTCCTC..........TTTCAGCCACTTCCAGATCT........CGAGAAAAGCACTTCA
O. officinalis W0002 Xa10-1               372 T.CATC........................CTCCTC........ATATTCAG........ATTTCT........................
O. officinalis W0002 Xa10-2               372 T.CATC........................CTCCTC........ATATTCAG........ATTTCT........................
O. punctata IRGC 105690 Xa10-1            361 TGAACC........................CACCAT..........ATTCAG........ATTTCTCCGGCGCTCGAT............
O. punctata IRGC 105690 Xa10-2            370 CAGATC........................GGGCTA...........GCTAGCTTACAACAATTAC........................
O. sativa IRBB10A Xa10                    249 C.CTCC........................CTCCTC....................TTTCATTCTT........................

                                          631 .......640.......650...
L. perrieri IRGC 105164 Xa10-1            401 ....................ATG
L. perrieri IRGC 105164 Xa10-2            401 ....................ATG
O. brachyantha IRGC 101232 Xa10-1         397 ............CGAG....ATG
O. brachyantha IRGC 101232 Xa10-2         381 CTTCTCTCTTAGCGAGCATCATG
O. coarctata TKM-2016 Xa10                401 ....................ATG
O. longistaminata RD23 Xa10               397 ............CGAG....ATG
O. meyeriana Menghai Xa10                 393 ............CTAGCTACATG
O. nivara IRGC 100897 Xa10                385 TTCTTCATTCTTCGAG....ATG
O. officinalis W0002 Xa10-1               397 ............CGAG....ATG
O. officinalis W0002 Xa10-2               397 ............CGAG....ATG
O. punctata IRGC 105690 Xa10-1            397 ............CGAG....ATG
O. punctata IRGC 105690 Xa10-2            401 ....................ATG
O. sativa IRBB10A Xa10                    270 ............CGAG....ATG
```

**Data S7. Alignment of genomic sequences upstream of the *Xa10-Ni*-like loci from cultivated and wild rice species.** 400 nucleotides upstream of start codons (**ATG**) of the *Xa10-Ni*-like loci were aligned using the MUSCLE algorithm.[11] Nucleotides common to 80% of sequences are shaded in black, while nucleotides common to 60% of sequences are shaded in grey using the pyBoxshade algorithm. The start codons (**ATG**) of each *Xa10-Ni*-like locus are colored in **red**. Note that the *Xa10-Ni* upstream sequence from *O. sativa* cv. Nipponbare does not contain binding elements for AvrXa10 and AvrXa23.[13]

```
                                                1 ........10........20........30........40........50........60........70........80........90
O. australiensis KR-Keep River 1 Xa10-Ni      1 .....................GCCCATCTAGCAACAATGAAATTAACAAAAGCAACAATGAAGTAACACAT.AAAGAATGAGGATCTTGT
O. glumipatula GEN 1233 Xa10-Ni               1 ............................................AGAAAACAATCCAAATATATATTGCT...AGCTGAAATTCGAATAA
O. longistaminata RD23 Xa10-Ni                1 .....................GTCAGCTCCAAGATACGTACCATGAGGCGCACTACAACAAAACAACCCC...AATGAAA..........
O. longistaminata W11 Xa10-Ni                 1 AAATTACTGAAACACATCCCGGCCAATCAAACTATACTACCTTGTTTAAGTCTATGGTTTTGTTTTTCT...AGCGATACATATTTGG
O. meridionalis OR44 W2112 Xa10-Ni            1 .............................CACACTTCCCTGCT...GCCTGAGCATTATGGGGCAT...ACTTTCAATATTCTTTA
O. meyeriana Menghai Xa10-Ni                  1 CGCCCCGTCGCCCACGCCGCCGGCCGCCCTCCCGCCTATGCCGCCGCCTACCCTGCCGCCACCTGCCTCCCCCACCACCGCCTGCCCTGC
O. nivara WR04-6 Xa10-Ni                      1 ...............GAGAGAGGGGAGAGGGGAGAGGGGATCTTGGAAGAGGGAGGAAGAAGATAGCTAG...GGGATGACATGTGGCCA
O. officinalis W0002 Xa10-Ni                  1 .......................CGTTGGCAATGCTCGCACGAAT..GCTGTCATATTTCCCCCCACC...ACTACGACTTCTCGTAT
O. rufipogon Taxon A Xa10-Ni                  1 ........................................TGAAGTAATCACATATTTATT.AACCAATAATGAGTTATC
O. rufipogon W1943 Xa10-Ni                    1 ...............AGAGAAAGAGAGAGGAGAGAGGGGATCTTGGAAGAGGGAGGAAGAAGATAGCTAG...GGGATGACATGTGGCCA
O. sativa Nipponbare Xa10-Ni                  1 ...............GAGAGAGGGGAGAGGGGAGAGGGGATCTTGGAAGAGGGAGGAAGAAGATAGCTAG...GGGATGACATGTGGCCA

                                               91 .......100.......110.......120.......130.......140.......150.......160.......170.......180
O. australiensis KR-Keep River 1 Xa10-Ni     69 CGCGTACTGCAAATCAGGTGTGGACCGTATATTATATGTCAGGAGAGAAGCTAACCTA.........................ACAAATTAAT
O. glumipatula GEN 1233 Xa10-Ni              44 TATATTTAACCGCCATAGTG...GCCTCCGACTACGT.CCATGTGGAGGAGATAGGGAGAGAAAGAGAGAGGAGAGAGGCATAAATT..C
O. longistaminata RD23 Xa10-Ni               57 .ACATCTTATTTGTAGGATAGAGACGTTTATTTCGTCTTCCACAGATAATAA..........................TTAAATT..C
O. longistaminata W11 Xa10-Ni                88 TACATATAA...GTAAGATG........AATTGGTGTAATATATAATTGGTTGTAGCTCGTTGA.................GCAAATG..C
O. meridionalis OR44 W2112 Xa10-Ni           53 CAGCTTTCGCCGCCATGTCGTCAGCTCCAAGGTACGTACCATGAGGTCACCTTTGAGATC.....................ATAAATT..C
O. meyeriana Menghai Xa10-Ni                 91 CACCGCCTACCTCCCTGGTAG...................AGGAAGAGAAGGAAGGG.......................GGAGAGTGAC
O. nivara WR04-6 Xa10-Ni                     73 TATTCGCAGTATCCATGGTGGATTCAATCTGTCTAGGTCCATG.......................................GTGGATTCAC
O. officinalis W0002 Xa10-Ni                 61 TAGTTATTA.CTGTAGGATG......CTGGATTCCATGCCACGTTT...................................GCAAGCCGGC
O. rufipogon Taxon A Xa10-Ni                 40 TATTTCAAGTAACAATGTCGG.................................................ATAAATATGT
O. rufipogon W1943 Xa10-Ni                   73 TATTCGCAGTATCCATGGTGGATTCAATCTGTCTAGGTCCATG.......................................GTGGATTCAC
O. sativa Nipponbare Xa10-Ni                 73 TATTCGCAGTATCCATGGTGGATTCAATCTGTCTAGGTCCATG.......................................GTGGATTCAC

                                              181 .......190.......200.......210.......220.......230.......240.......250.......260.......270
O. australiensis KR-Keep River 1 Xa10-Ni    137 CTTGTCGAAATGTAACA...AGTAATCTTGTCCAATG...........T..ATACGAAT.....ATACATGTA.............TAT
O. glumipatula GEN 1233 Xa10-Ni             128 CCTACCTTTATGTAACA...CAAATTCCAGACAATAGA..........T..CAAGAGGC.....ATGCACGCACAATACCCTGAAGCTAT
O. longistaminata RD23 Xa10-Ni              117 TCCACCTTTATATAACA...CAAATTCCAGACAATGGA..........T..CGAGAGGC.........ATGCACAATACCCTGACTCTAT
O. longistaminata W11 Xa10-Ni               149 CGGAATTATATTTCATAATAAAAAAACGTGAGCAAGGATGTAAGGATGT..AAGGAGAC.....AGGTGGTTGTTGAACTTTTATAGTAC
O. meridionalis OR44 W2112 Xa10-Ni          121 CCTACCTTTATATAACA...CAAATTCCAGACAATAGA..........T..CAAGAGGC.....ATGCACGCACAATACCCTGAAGCTAT
O. meyeriana Menghai Xa10-Ni                139 .........ATGTGGGGTTCTAAAAATTTGAAGAGTGG......GAGAT..AGAGAGACTATTGGAACATAAAAAAAATATAGAGAGGAA
O. nivara WR04-6 Xa10-Ni                    126 .........ATGCAGCGTCCTGGAGTGGAGAAAACAGA..............AAGGGAC.........ACTTTCAATATTCTTTACAGAT
O. officinalis W0002 Xa10-Ni                110 ATTTCAATTGTGCAAC....CGCGATGAGTACGACAG..........T..CAACGGGGAAATTGCGCATTCGC....CGCCCACCAGAC
O. rufipogon Taxon A Xa10-Ni                 71 AAACATAGTATATGATA...TAATATCATTATAATTGTCTCTAGGATATACCAATAGGC...........TACGCTTGACTTCTCTTAT
O. rufipogon W1943 Xa10-Ni                  126 .........ATGCAGCGTCCTGGAGTGGAGAAAACAGA..............AAGGGAC.........ACTTTCAATATTCTTTACAGAT
O. sativa Nipponbare Xa10-Ni                126 .........ATGCAGCGTCCTGGAGTGGAGAAAACAGA..............AAGGGAC.........ACTTTCAATATTCTTTACAGAT

                                              271 .......280.......290.......300.......310.......320.......330.......340.......350.......360
O. australiensis KR-Keep River 1 Xa10-Ni    192 TATATAGCCAGGAGAGAAGCTAACCTAACAA.................ATAA......TCTTGTCGAAA.....TGTAACA.........
O. glumipatula GEN 1233 Xa10-Ni             198 TCTG..........ATCAGCTTTACCAAAAACA...............ATTC......TCTCTTCGTGGCCCTTTACGTCG.........
O. longistaminata RD23 Xa10-Ni              183 TCTGTATTCAG...ATCGCCTTTACAAAAACA................ATTG......TCTCTTC......CTTTACGACC.........
O. longistaminata W11 Xa10-Ni               232 TATATTTA......ATCTGGTATCACGCAATAT.....GGGCG.....ACTC......TGTTGTGATAG..CAATGGGGCC.........
O. meridionalis OR44 W2112 Xa10-Ni          191 TCTG..........ATCACCTTTACCAAAAACA...............ATTC......TCTCTTCGTGGCCCTTTACGTCG.........
O. meyeriana Menghai Xa10-Ni                212 TCTTTTTTTG...GCTGGCTAAATAGGCAAATGGAGAGTGAG.....ATTTGATCAATCTGTTAGAGTTGCTCTTAGTCA.........
O. nivara WR04-6 Xa10-Ni                    184 TTCGTCGCCATGTCGTCAGCTCCAAT......................ATTC......TCTCTTCGTGGCCCTTTACGTCG.........
O. officinalis W0002 Xa10-Ni                179 CATG.....ATGATGCTAGGTTTCCCAGGGCGA.....AGGATGACGCATGC...TAGCTTTCCCATGGCGCAGGATGACGCATGCTAGG
O. rufipogon Taxon A Xa10-Ni                146 TAGCTTAGGATGCGATCGAGTTCCAAGAGAAAACTAACGCGGTTTGACATTTAGAGTTTGTGCTCGCGGCCTCCGCCATCG.........
O. rufipogon W1943 Xa10-Ni                  184 TTCGTCGCCATGTCGTCAGCTCCAAT......................ATTC......TCTCTTCGTGGCCCTTTACGTCG.........
O. sativa Nipponbare Xa10-Ni                184 TTCGTCGCCATGTCGTCAGCTCCAAT......................ATTC......TCTCTTCGTGGCCCTTTACGTCG.........

                                              361 .......370.......380.......390.......400.......410.......420.......430.......440.......450
O. australiensis KR-Keep River 1 Xa10-Ni    245 ..................AATAATCTTGTCC.............A.....ATGTATACGAATATACAAGTCAACGACAGTCGTGGTGGCTT
O. glumipatula GEN 1233 Xa10-Ni             248 ............GTTA.AACACCCTTTTCT.............TTCAGCGCTTTTA..GAAAGTGAAGCACATAG.......ATGCTTTT
O. longistaminata RD23 Xa10-Ni              233 ................CAACGGC.........................GCACCTTTACAAAAAGCGTAGCACATAGATGCTTCATGCTTTT
O. longistaminata W11 Xa10-Ni               289 ............ATTA..................................CTATATTAATATTGAA................AAACTGGT
O. meridionalis OR44 W2112 Xa10-Ni          241 ............GTCA.AACACCCTTTTCT.............TTCAGCGCCTTTAGAAAAAGTGAAGCACATAG.......ATGCTTTT
O. meyeriana Menghai Xa10-Ni                285 ..............................................CACGCACTCACACTAATA..........................TTC
O. nivara WR04-6 Xa10-Ni                    237 ............GTCA.AACACCCTTTTCT.............TTCAGCGCCTTTAGAAAAAGTGAAGCGCATAG.......ATGCTTTT
O. officinalis W0002 Xa10-Ni                256 TTTTTCTAACAAGTTACCATATCTTTGACC.............TTGCTGACAATTAGCAGAA..........................TTT
O. rufipogon Taxon A Xa10-Ni                227 .....TTGCAGAATTTTAACGAGACTGTCCGTATCGGCAGTCGTTGCAGACTTTTAACGAGCACATCGCACATCTCTACCGTCCAC.CGC
O. rufipogon W1943 Xa10-Ni                  237 ............GTCA.AACACCCTTTTCT.............TTCAGCGCCTTTAGAAAAAGTGAAGCGCATAG.......ATGCTTTT
O. sativa Nipponbare Xa10-Ni                237 ............GTCA.AACACCCTTTTCT.............TTCAGCGCCTTTAGAAAAAGTGAAGCGCATAG.......ATGCTTTT
                                              451 .......460.......470.......480.......490.......500.......510.......520.......530.......540
O. australiensis KR-Keep River 1 Xa10-Ni    300 CGGTATATA.............ACAGGAC...ATGGTTTCACGATTGATTCTCCCTAACAAC.CCCCCCAAA.CCACCACCAA...AAAAA
O. glumipatula GEN 1233 Xa10-Ni             303 ACGCCTATA.............T..GAAC.......TCACCAAATTGGGCT...TGAAGAAT.CATTCGGCA.TCGAGACCGGGCTAGGAA
O. longistaminata RD23 Xa10-Ni              283 ACGCCTATAGGCTATAAGAAGTAAGAAC.......TCACCAAATTAAGCT...TGAAGAAT.CATTCGGCA.TCGA...CCGGCTAGCTA
O. longistaminata W11 Xa10-Ni               317 GGGAGTATA................AAAC.........TACCAG..........CAGAAGAC.CATCCAAAA.AAAAAAACGA..GAGAAG
O. meridionalis OR44 W2112 Xa10-Ni          298 ACGCCTATA.............T..GAAC.......TCACCAAATTGGGCT...TGAAGAAT.CATTCGGCA.TCGAGACCGGGCTAG...
O. meyeriana Menghai Xa10-Ni                306 ACATATATT.............TATAAAC............CAAATTCTAT....TAAATAAA.TCTTTTGCA.AAGTCAATTGTTCGTTTT
O. nivara WR04-6 Xa10-Ni                    294 ACGCCTATA.............T..GAAC.......TCACCAAATTGGGCT...TGAAGAAT.CATTCGGCA.TCGAGACCGGGCTAGCTA
O. officinalis W0002 Xa10-Ni                308 AAGTCTATA.............T..AAAC.......TCACACGACT..................CATTTTAAACTTAAAAGTTAGTAACACA
O. rufipogon Taxon A Xa10-Ni                311 GCGCCTATA.............T..AAACTCATCAGTCATCAAATTAT......TGAAGAGTATTTTCGACA.TCAATTCC..........
O. rufipogon W1943 Xa10-Ni                  294 ACGCCTATA.............T..GAAC.......TCACCAAATTGGGCT...TGAAGAAT.CATTCGGCA.TCGAGACCGGGCTAGCTA
O. sativa Nipponbare Xa10-Ni                294 ACGCCTATA.............T..GAAC.......TCACCAAATTGGGCT...TGAAGAAT.CATTCGGCA.TCGAGACCGGGCTAGCTA

                                              541 .......550.......560.......570.......580......
O. australiensis KR-Keep River 1 Xa10-Ni    370 AAAACATCAGT............TTTCATCAGTTTGGCGAGGAATG
O. glumipatula GEN 1233 Xa10-Ni             367 CTAAATGGAGCTC.........ATTTAATTAGCTAGGAACTAAATG
O. longistaminata RD23 Xa10-Ni              358 GCTAGCTTAGTCTTGGCGGCGAGCTTAATTAGCTAGGAACTAAATG
O. longistaminata W11 Xa10-Ni               369 AAAACATCAGT...........GTTCATCAAATTGGGGGAGGAATG
O. meridionalis OR44 W2112 Xa10-Ni          359 .CTAGCTTAGTTTTGGCGGCGAGCTTAATTAGCTAGGAACTAAATG
O. meyeriana Menghai Xa10-Ni                367 AAAGTATTAATCATGATAATGATTTTACACGCTT.........ATG
O. nivara WR04-6 Xa10-Ni                    358 GCTAGCTTAGTTTTGGCGGCGAGCTTAATTAGCTAGGAACTAAATG
O. officinalis W0002 Xa10-Ni                360 AAAGCCTTAACCATTTTCAGTATGTCAA..ATTCAGCGAGAAAATG
O. rufipogon Taxon A Xa10-Ni                370 .CCGGCTTAGTTCTATCGGCGAGCT...........GAATGTAATG
O. rufipogon W1943 Xa10-Ni                  358 GCTAGCTTAGTTTTGGCGGCGAGCTTAATTAGCTAGGAACTAAATG
O. sativa Nipponbare Xa10-Ni                358 GCTAGCTTAGTTTTGGCGGCGAGCTTAATTAGCTAGGAACTAAATG
```

**Data S8. Alignment of genomic sequences upstream of the *Xa23*-like loci from cultivated and wild rice species.** 400 nucleotides upstream of start codons (**ATG**) of the *Xa23*-like loci were aligned using the MUSCLE algorithm.[11] Nucleotides common to 80% of sequences are shaded in black, while nucleotides common to 60% of sequences are shaded in grey using the pyBoxshade algorithm. The start codons (**ATG**) of each *Xa23* locus are colored in **red**. The AvrXa23 binding element for the activation of *Xa23* transcription in *O. sativa* cv. CBB23 is shaded in magenta.[12]

```
                                  1 ........10........20........30........40........50........60........70........80........90
O. barthii IRGC 105608 Xa23        1 CCCATATGGAGAATTTAGGTAAGTTGTGATAAGTATATAAACATATGCTTTCCTACTACAGTATAATAGCCGGTTAATCCTATGTTTGTC
O. glaberrima IRGC 96717 Xa23      1 ..........................................................................................
O. glumipatula GEN 1233 Xa23       1 ...............................GTATTGCATATTGTGGGGAGGGGAGTGCGGGATTATCGCTGGGCG.......CGCATTA
O. longistaminata RD23 Xa23-1      1 ..............................................................ATGTTCTTTAAGCA......TGACATAA
O. longistaminata RD23 Xa23-2      1 ..............................................................ATGTTCTTTAAGCA......TGACATAA
O. longistaminata RD23 Xa23-3      1 ............................................................AGCTCTGCACTGGACT.......AAAAGTT
O. longistaminata RD23 Xa23-4      1 ..............................................................ATAATTGCTTATAG..............
O. longistaminata RD23 Xa23-5      1 .........................................................AATATATATGTACTTAAAA..............
O. meridionalis OR44 W2112 Xa23    1 CCTATATTACTACTTT............................................GTATTATAGCTGGGTT.........CACTA
O. nivara WR04-6 Xa23              1 ............................................................GGGTGATCGCTTGGCG.......CGCATTA
O. officinalis W0002 Xa23-1        1 ...........................................TTTGGCTGTCTACAAAAATATTTTTTCTAGTGA..............
O. officinalis W0002 Xa23-2        1 ..........................................................CAATGTGATGCTACAGTA.............A
O. punctata IRGC 105690 Xa23       1 ..............................................TAAGTTTAACCGACGAGTATTCCTCGGGCA.......TATGTCA
O. rufipogon Taxon A Xa23          1 ......................................AAACATATGCTTTCCTACTACAGTATAATAGCCGGTTAATCCTATGTTTGTC
O. rufipogon W1943 Xa23            1 .....................................................................................CATTA
O. sativa Nipponbare Xa23          1 ............................................................GGGTGATCGCTTGGCG.......CGCATTA
O. sativa CBB23 Xa23               1 ............................................CCGCCACGGTGGCACAATGTCAGCGCCAGTCC.CGTTGGCGCTGAC

                                 91 .......100.......110.......120.......130.......140.......150.......160.......170.......180
O. barthii IRGC 105608 Xa23       91 TCGTCCGCCA...............TTCCCTAACTAGTATAAG.................................CGAAAGA.......
O. glaberrima IRGC 96717 Xa23      1 ...TCCGCCA...............TTCCCTAACTAGTATAAG.................................CGAAAGA.......
O. glumipatula GEN 1233 Xa23      53 GCAGTCCCCC...............CTTCCCTTCCGCTATA...................................TAAAAGT.......
O. longistaminata RD23 Xa23-1     23 GTA......................TTTTTATATTTGTATAAAAAATTTGAATAAAGCAAGTGGTCAAACGTTGGTTAAAAGT.......
O. longistaminata RD23 Xa23-2     23 GTA......................TTTTTATATTTGTATAAAAAATTTGAATAAAGCAAGTGGTCAAACGTTGGTTAAAAGT.......
O. longistaminata RD23 Xa23-3     24 ATTATCAGTC...............CTGGCCGAC..........................................TAAAAGT.......
O. longistaminata RD23 Xa23-4     15 .......ATA...............TTATTTAAATATGATATAACA..............................TAAAAGT.......
O. longistaminata RD23 Xa23-5     20 ..AGTCAACG...................GTAACAAATAT....................................TTAGGGT.......
O. meridionalis OR44 W2112 Xa23   38 TGTTGCATCA...............TCTCACAGCTAACCCGAACATCACTAACA......................TCGCAAT.......
O. nivara WR04-6 Xa23             24 GCAGTCCCCC...............CTTCCCTTCCGCTATT...................................TCAAAGT.......
O. officinalis W0002 Xa23-1       34 .TATAAGCTA...............CATCATGACACTTTCA...................................TGAAGTT.......
O. officinalis W0002 Xa23-2       20 ATATTTGCTA...............ATGACGGATTAATTAGGC.................................TCAAAAG.......
O. punctata IRGC 105690 Xa23      38 ACAACCAATCAGACTATGCTTCAGGCTGCCTGACCAGT......................................CTAAATT.......
O. rufipogon Taxon A Xa23         53 TCGTCCGCCA...............TTCCCTAACTAGTAGTATAAG..............................CGAAAGA.......
O. rufipogon W1943 Xa23            6 GCAGTCCCCC...............CTTCCCTTCCGCTATT...................................TCAAAGT.......
O. sativa Nipponbare Xa23         24 GCAGTCCCCC...............CTTCCCTTCCGCTATT...................................TCAAAGT.......
O. sativa CBB23 Xa23              46 ACGGCCAACG...............TCAGCGCCAATGTGTTTGGCGCTGAGGCGACGGCCTATTTTTGG......TTGAAGTTTTTGGC

                                181 .......190.......200.......210.......220.......230.......240.......250.......260.......270
O. barthii IRGC 105608 Xa23      126 .........AAACGGATAAATTCATCTCGTTTATATGCATAAACATCAAACGGAGCAACTCTCT..........................
O. glaberrima IRGC 96717 Xa23     33 .........AAACGGATAAATTCATCTCGTTTATATGCATAAACATCAAACGGAGCAACTCTCT..........................
O. glumipatula GEN 1233 Xa23      86 .........ATCCG.......TCTTCCTCCCTGTCCACATGAGCT..AAACGCTTCCTCTTCCCTGAGTCAAAGATCCAATATCTCCTTC
O. longistaminata RD23 Xa23-1     84 .........CAACG.......GCGTCATACATTAAAATACGGAGG..GAGTACTCCGTATTAATCAAATTAA.........GCAGCATC
O. longistaminata RD23 Xa23-2     84 .........CAACG.......GCGTCATACATTAAAATACGGAGG..GAGTACTCCGTATTAATCAAATTAA.........GCAGCATC
O. longistaminata RD23 Xa23-3     50 .........AATTA.......TCATATTACTAATCAAAATTGGCCGGGAAGATAGCTCTGCACTGGACTAAA..AGTTAT...TATCAGT
O. longistaminata RD23 Xa23-4     46 .........AATAGGTTAGAAAATT...........ATATAAGCAACTAA.........TACCAGTGAAAACAATTTATGAA.......
O. longistaminata RD23 Xa23-5     46 ...........CGGAGGGA............GCATATATAAATCCAACAGTCGATCAGTCGCTTAGC.....................
O. meridionalis OR44 W2112 Xa23   84 .........ATTCT.......CCTTCCGCGCCCTTCACATCAGCT..ACTTCCTTCCGCATCACTAACATTAGCTACTATAAAAGTCCCT
O. nivara WR04-6 Xa23             57 .........ATCCG.......TCTTCCTCCCTGTCCACATGAGCT..AAACGCTTCCTCTTCCCTGAGTCAA..AGTCTT...CCCTAT.
O. officinalis W0002 Xa23-1       66 .........ACATC.......TCTTAAATCTCCATAATATAAGCT..ACATCGTGACACTTTCACGAAATTA..CATCTCTTAAATCTCC
O. officinalis W0002 Xa23-2       55 .........ATTCG.......TCTCGCAATTTACACGCAAAATCT..GTAATTTGTTTTGTTATTAGACTAT................
O. punctata IRGC 105690 Xa23      83 .........CATTGGTTTGTCAAGTAGGTCTCAAGCACATAAAAATCTCAAGCAGTTTACTGATCAGGGCACCAATCAAATAG.......
O. rufipogon Taxon A Xa23         91 .........AAACGGATAAATTCATCTCGTTGTAAACTATGACGT..AAACA.......TCACTGAAGTAA..CATCACGAGACTTTTC
O. rufipogon W1943 Xa23           39 .........ATCCG.......TCTTCCTCCCTGTCCACATGAGCT..AAACGCTTCCTCTTCCCTGAGTCAA..AGTCTT...CCCTAT.
O. sativa Nipponbare Xa23         57 .........ATCCG.......TCTTCCTCCCTGTCCACATGAGCT..AAACGCTTCCTCTTCCCTGAGTCAA..AGTCTT...CCCTAT.
O. sativa CBB23 Xa23             114 AGGGGTTAGTTTCGAAATAAGTTTTCCAAAAGGGTCAATTTGTCA..AAAAAACGGCTCGTCCCTGAGTCAA..AGTCTT...CCCTAT.

                                271 .......280.......290.......300.......310.......320.......330.......340.......350.......360
O. barthii IRGC 105608 Xa23      181 ..................GCATTTGCGTCAAACAAGACTTGGACGAGAAGTGGTTACATC......GATA....GTAAACACATTGAAA
O. glaberrima IRGC 96717 Xa23     88 ..................GCATTTGCGTCAAACAAGACTTGGACGAGAAGTGGTTACATC......GATA....GTAAACACATTGAAA
O. glumipatula GEN 1233 Xa23     158 CGCG..............CCCTTCAGATCAGCTACTCCCTTCCGCTCACTAAGACCCTTCCGC...TATC....ATCAGCTACTATAAAA
O. longistaminata RD23 Xa23-1    146 CAGTCAACGCTCCTAGCTTCCAGTGCTTCTTCTACCACTCAAAT...............................TAAGCTAAAATAGA.
O. longistaminata RD23 Xa23-2    146 CAGTCAACGCTCCTAGCTTCCAGTGCTTCTTCTACCACTCAAAT...............................TAAGCTAAAATAGA.
O. longistaminata RD23 Xa23-3    119 CCTGGCCGACTAAAAGTAATTATCATATTGCTGATCGTAAAAGTAATTATCAGTCCTGGCCGGGAAGATA....CTTAATTAAAATTAAT
O. longistaminata RD23 Xa23-4     99 ..................GAAATCGTGGCAGTTGATCTCCGGTT.....................AATA....TTTCGCTCTTATAAA.
O. longistaminata RD23 Xa23-5     90 ..................TCTTTATCATCGTCCAGTTAGTGCTTAGCTCTATGCTGCATCTCAAGGAACT....CATAACTAGTATAAA.
O. meridionalis OR44 W2112 Xa23  156 TCCGCGTCACTAACATCTTCCTCCGCATCAC...................................TAAC....ATTAGCTACTATAAAA
O. nivara WR04-6 Xa23            123 ..................ATTACTACTTTGTATTATAGCTGGGT.TCACTATGTTGCATC......ATCT....CACAGCTAACCCGAA.
O. officinalis W0002 Xa23-1      136 ACATAA............TAAGCTACATCGTGCCAT..............CGTGCCTCCTCCCTACATC....TCCATGTAATATGAAG
O. officinalis W0002 Xa23-2      109 .............................GTTTAATACTTTAAATGTGTGTCCGTATATCTGATGTGATCTGATGTGACGTTTTTTGAAA
O. punctata IRGC 105690 Xa23     157 ..................ACCCTTAATTTGTTACGTTCTCAAAGTATATATATAGCCTTCCTTCAGTATA....TTATATTAGGGAGGAG
O. rufipogon Taxon A Xa23        160 CGTGCCTCAACATCAGGTTCCTCAACATCAGGCAAA.............................AACA....ATCAGGTAAGTCCAGG
O. rufipogon W1943 Xa23          105 ..................ATTACTACTTTGTATTATAGCTGGGT.TCACTATGTTGCATC......ATCT....CACAGCTAACCCGAA.
O. sativa Nipponbare Xa23        123 ..................ATTACTACTTTGTATTATAGCTGGGT.TCACTATGTTGCATC......ATCT....CACAGCTAACCCGAA.
O. sativa CBB23 Xa23             196 ..................TAAATTATGCGGCATCAC.............................TAAC....ATCAGCTACTATAAAA

                                361 .......370.......380.......390.......400.......410.......420.......430.......440.......450
O. barthii IRGC 105608 Xa23      243 CGGACAG...................................................................................
O. glaberrima IRGC 96717 Xa23    150 CGGACAGGAGAAGAGAAGAAGAAAAGAGAGTTGGAAAACGAGAAGTGGTTACATCGAGAAGATACATCGATAGTAAAACACATTG.AAAC
O. glumipatula GEN 1233 Xa23     227 AG...............................................TCCCTTTCGCGTTACTAATATCTT.................
O. longistaminata RD23 Xa23-1    204 ....................................ATCTCCTACTAGTAGCTTTCAGTGTTTCAGCATCCAGT............CCAG
O. longistaminata RD23 Xa23-2    204 ....................................ATCTCCTACTAGTAGCTTTCAGTGTTTCAGCATCCAGT............CCAG
O. longistaminata RD23 Xa23-3    205 CA..........................................................CTTCTCACATCTG.................
O. longistaminata RD23 Xa23-4    144 .....................................................TGTCGCATACTTAGCGTACGACGCATGCCCGCGCGAA
O. longistaminata RD23 Xa23-5    157 ..............................TCCAACGAATCCCATCTGTACCTCTGCACTTCTTAACTAGTAT.............AAAT
O. meridionalis OR44 W2112 Xa23  207 G................................................TCCCTTCCGCGTCACTAACATCTT.................
O. nivara WR04-6 Xa23            183 ............................CATCACTAACATCGCAATATTCTCCTTCCGCGCCCTTCACATCAG.............CTAC
O. officinalis W0002 Xa23-1      195 ATG.........................................ATTCCTATCTGTCTCGCCACTGACTTCCG.................
O. officinalis W0002 Xa23-2      170 AAAAAAG.......................TTTGGAACTAAACACGGCGGGGCTGTATCTGCTATCACATCAA.................
O. punctata IRGC 105690 Xa23     225 AGAG............................AGAGAGAGAGAGAGAGCGCACTGCAGTGGCTCGATCAGTGA.................
O. rufipogon Taxon A Xa23        216 TG...........................................................CCTCAACATCAG.............GTTA
O. rufipogon W1943 Xa23          165 ............................CATCACTAACATCGCAATATTCTCCTTCCGCGCCCTTCACATCAG.............CTAC
O. sativa Nipponbare Xa23        183 ............................CATCACTAACATCGCAATATTCTCCTTCCGCGCCCTTCACATCAG.............CTAC
O. sativa CBB23 Xa23             234 G................................................TCCCTTCCGCGTCACTAACATCAG.............CTAC
```

```
                                      451 .......460.......470.......480.......490.......500.......510.......520.......530.......540
O. barthii IRGC 105608 Xa23           250 ...........GAGAAGAAGAAAAGAGAGTTGGAAAACGAGAAG.......TGGTT.....ACATCGAG...AAGAT............
O. glaberrima IRGC 96717 Xa23         239 GGACGGGAGAAGAGAAGAAGAAAAGAGAGTTGGAAAACGAGAAG.......TGGTT.....ACATCGAG...AAGAT............
O. glumipatula GEN 1233 Xa23          253 ...........CCTCCGCATCACTGACATCAGCTATTATAAAAG..............................TCCCTTT.......
O. longistaminata RD23 Xa23-1         246 TGAGTCCACGCTCGCACAGCAGCACCTAGCTAAGCTTCTATAGC.......GCTCT.....ATATATAG...AGCATCCTTCT.......
O. longistaminata RD23 Xa23-2         246 TGAGTCCACGCTCGCACAGCAGCACCTAGCTAAGCTTCTATAGC.......GCTCT.....ATATATAG...AGCATCCTTCT.......
O. longistaminata RD23 Xa23-3         220 ...GGAAGATTCATAGCTTCTATAGCTAGC.......TTCTCAC.......TTCTC.....ACA..TCA...TCCTTCTATAT.......
O. longistaminata RD23 Xa23-4         181 TGCGCGGGCTACCTTCCTAGTTAATTTAATGTTGCATCTCAAAG.......AGATC.....TCAGCTAAGCTTGCATCATTTTAATTAAT
O. longistaminata RD23 Xa23-5         204 CCAACGAATCCCATATAAGTCACAATCTGCACCTCTGCATTTCT.......TAACTAGTATAAATCCAA...CGAATCCATATAAGTCAC
O. meridionalis OR44 W2112 Xa23       232 ...........CCTCTGCATCACTAACATCAGCTACTATAAAAG.......TCCCTTCCGCATCACTAA...CATCTTCCTCC.......
O. nivara WR04-6 Xa23                 232 TTCCACCACCTCCCATGTTTCAAAGGTAAA....TTCGTCGTGG.......TGGTG.....AGA..TTA...TGCCTTCCTTC.......
O. officinalis W0002 Xa23-1           227 ...............CGTACGAAAGTCATT....CGTCTCCCAA.......CATCT.....TCG..TAT...CACCTCACTTT.......
O. officinalis W0002 Xa23-2           220 CAATGGTATCTCCAAGCAGTCAAACAATGGTATCACTCTCAAGA.......AACTG.....GCAGCCAT...AGCCTCACTTT.......
O. punctata IRGC 105690 Xa23          270 .......GCGCCGTGCAGACTAGAGATATCGACTTATCGAGGAA.......GCAGT.....ACAGATCGGGCTAGCTAGCTT........
O. rufipogon Taxon A Xa23             234 GAACCTCAACATTTAGGGGGAGAAGTAACATCACTGTATAGAAGGATGCTCTATATATAGCATATATAG...AGCATCCTTCT.......
O. rufipogon W1943 Xa23               214 TTCCACCACCTCCCATGTTTCAAAGGTAAA....TTCGTCGTGG.......TGGTG.....AGA..TTA...TGCCTTCCTTC.......
O. sativa Nipponbare Xa23             232 TTCCACCACCTCCCATGTTTCAAAGGTAAA....TTCGTCGTGG.......TGGTG.....AGA..TTA...TGCCTTCCTTC.......
O. sativa CBB23 Xa23                  263 TATAAAAGTCCCTTCCG.........................................AAA...CATCTTCCTCC.......

                                      541 .......550.......560.......570.......580.......590.......600.......610.......620.......630
O. barthii IRGC 105608 Xa23           301 ................ACATCGATAGTAAAACAAAACGTC........CGGTCATGGTCGCGCGCCTTCCGCCTGCACGCCT......AC
O. glaberrima IRGC 96717 Xa23         301 ................ACATCGATAGTAAAACAAAACCTC........CGGTCATGGTCGCGCGCCTTCCGCCTGCACGCCT......AC
O. glumipatula GEN 1233 Xa23          293 ...CCGCATCACTA..ACATCAGC..TACTATAAAAGCC.........CTTCCTTGTTGCATCATCT....................CA
O. longistaminata RD23 Xa23-1         314 ................ATATCAAACGT.CTATCAAACACAGCAT....CCTTCTATATCAAACGTC.....................CA
O. longistaminata RD23 Xa23-2         314 ................ATATCAAACGT.CTATCAAACACAGCAT....CCTTCTATATCAAACGTC.....................CA
O. longistaminata RD23 Xa23-3         276 ....CAAATCTCCAGTTCTCCAGTC.CACAGCAGCAGCTAAGCTTTTCCACGCCTTATCAAACACATCATCCTTCTATATCAAACGTCCA
O. longistaminata RD23 Xa23-4         259 GTTGTATCTCAAAGAGATCTCAGCTAAAGAGCTCACAGCCGTATTTTATTTGCAACTTGAAACCTCT.............ATATATATG
O. longistaminata RD23 Xa23-5         284 AATCTGCACCTCT...ACACTAGTATAAATCCAATAGTCTATCACAACTTCTCCTAATATAAATCCAACAGTCTA.............TC
O. meridionalis OR44 W2112 Xa23       294 ....CGCGTCACTA..ACATCAGC..TACTATAAAAGCC.........CTTCCTTGTTGCATCATCT....................CA
O. nivara WR04-6 Xa23                 294 ....CGCCTCACTA..ACATCAGC..TACTATAAAAGCC.........CTTCCTTGTTGCATCATCT....................CA
O. officinalis W0002 Xa23-1           274 ....TGTTTTGCCAAAGCAGCACT..TTCTTCAAAGGTCAGCACTTTTCTTCAAAGGTCATTCGTCTCCCTACGTCTCCATATCTATATA
O. officinalis W0002 Xa23-2           288 .......................TGTTGTGCCAAAGCCAGCACTTT.CTTCAAATGTCATTCGTCTCCCTGCGTCTCCATATCTATATA
O. punctata IRGC 105690 Xa23          333 ................ACAACAAT.......................TACATGGAGTTGATCATC.......................
O. rufipogon Taxon A Xa23             314 ..........ATACAGCAGCAGCTTCTATATCAAACACAGCAT....CCTTCTATATCAAACGTCT......................
O. rufipogon W1943 Xa23               276 ....CGCCTCACTA..ACATCAGC..TACTATAAAAGCC.........CTTCCTTGTTGCATCATCT....................CA
O. sativa Nipponbare Xa23             294 ....CGCCTCACTA..ACATCAGC..TACTATAAAAGCC.........CTTCCTTGTTGCATCATCT....................CA
O. sativa CBB23 Xa23                  294 ....CGCATCACTA..ACATCAGC..TTCTATAAAAGCC.........CTTCCTTGTTGCATCATCT....................CA

                                      631 .......640.......650.......660.......670.......680.......690.......700.......710.......720
O. barthii IRGC 105608 Xa23           361 AAAA.GCCGCGCGC.......TCATTTT...TGCCA...............AACGGGCAGTGG...................CA.......
O. glaberrima IRGC 96717 Xa23         361 AAAA.GCCGCGCGC.......TCATTTT...TGCCA...............AACGGCCAATGG...................CA.......
O. glumipatula GEN 1233 Xa23          346 AGGA.GCTGCAAGCACT....TCCTCTC...TGGCAGCACTTCCTCATCTCAAGGA...GTTG...................CA.......
O. longistaminata RD23 Xa23-1         361 GTTC.GCCGCGCGC.......TCATTTT...TGCCGCCA............AACGG...GTGG...................CA.......
O. longistaminata RD23 Xa23-2         361 GTTC.GCCGCGCGC.......TCATTTT...TGCCGCCA............AACGG...GTGG...................CA.......
O. longistaminata RD23 Xa23-3         361 GTTC.GCCACGCGC.......TCATTTT...TGCCACCA............AACGG...GTGG...................CA.......
O. longistaminata RD23 Xa23-4         335 CATA.TAAGCGCGCAAT....TCATGTTGCATCTCA...............AAGAGCTCACAGCCGTAATCTCGCG.....CAATTAATT
O. longistaminata RD23 Xa23-5         358 AGTC.GCTTAGCTCTATATCATCGT........CCAGTTAGTGCTTACCTC.......................................
O. meridionalis OR44 W2112 Xa23       346 AGGA.GCTGCAAGCACT....TCCTCTC...TGGCAGCACTTCCTCATCTCAAGGA...GTTG...................CA.......
O. nivara WR04-6 Xa23                 346 AGGA.GCTGCAAGCACT....TCCTCTC...TGGCAGCACTTCCTCATCTCAAGGA...GTTG...................CA.......
O. officinalis W0002 Xa23-1           358 TGT.....ATGCGC.......TCATTTT...TGCCA...............AAGAG...GTAG...................CGCTTTGGT
O. officinalis W0002 Xa23-2           353 TGTATAGTATGCGC.......TCATTTT...TGCCA...............AAGAG...GTAG...................CGCTTTGGT
O. punctata IRGC 105690 Xa23          359 .....ACTGTTTGCGCTTCGATCCTCGT...CGCCATTATCATC...................GTAG...................CG.......
O. rufipogon Taxon A Xa23             366 ...A.GTTGCGC.........TCATTTT...TGCCGCCA............AACGG...GTGG...................CA.......
O. rufipogon W1943 Xa23               328 AGGA.GCTGCAAGCACT....TCCTCTC...TGGCAGCACTTCCTCATCTCAAGGA...GCTGGCAGCCGTAGCCGGTATACA.......
O. sativa Nipponbare Xa23             346 AGGA.GCTGCAAGCACT....TCCTCTC...TGGCAGCACTTCCTCATCTCAAGGA...GTTG...................CA.......
O. sativa CBB23 Xa23                  346 AGGA.GCTGCAAGCACT....TCCTCTC...TGGCAGCACTTCCTCATCTCAAGGA...GTTG...................CA.......

                                      721 .......
O. barthii IRGC 105608 Xa23           400 ...AATG
O. glaberrima IRGC 96717 Xa23         400 ...AATG
O. glumipatula GEN 1233 Xa23          400 ...AATG
O. longistaminata RD23 Xa23-1         400 ...AATG
O. longistaminata RD23 Xa23-2         400 ...AATG
O. longistaminata RD23 Xa23-3         400 ...AATG
O. longistaminata RD23 Xa23-4         400 ...AATG
O. longistaminata RD23 Xa23-5         400 ...TATG
O. meridionalis OR44 W2112 Xa23       400 ...AATG
O. nivara WR04-6 Xa23                 400 ...AATG
O. officinalis W0002 Xa23-1           397 GGCAATG
O. officinalis W0002 Xa23-2           397 GGCAATG
O. punctata IRGC 105690 Xa23          401 ....ATG
O. rufipogon Taxon A Xa23             400 ...AATG
O. rufipogon W1943 Xa23               400 ...CATG
O. sativa Nipponbare Xa23             400 ...AATG
O. sativa CBB23 Xa23                  400 ...AATG
```

**Data S9. Synthesized *E* gene homologs from wild rice and cutgrass species for transient overexpression in *N. benthamiana.*** Cloning adapters containing *Bsa*I sites are highlighted in yellow. The start (**ATG**) and stop (**TAA** / **TAG** / **TGA**) codons of *E* gene homologs are colored **RED**. The artificial intron in the *GFP* CDS is highlighted in magenta.

Code #1 *O. barthii* IRGC 105608 *Xa23*
GTATAGTGCACGGGTCCGTCCGACATGATGGATTGAAGGCAACATTAGAATTTGGTCTCAAATGTTGCATCTCAA
GGAACTCGCAGCCGTAGCCGGCATCGAC**ATG**ATCGTCATCTACCTCTGCCGCTTTCTCCTCCGTCGCTGCCGCAG
CCGCAACGCATCCAACAGCCTCCGTTTTCGCCTCAAGGTATTAACTGTATTGTTGTACATATCACTCTCAACCAT
GCTGTTGTACCTGTTTGGCTCCATCATGCCGCTGCCACCGTGGGGTGCTCTGGCCGGT**TGA**GCTTTGAGACCAAA

Code #2 *O. glaberrima* IRGC 96717 *Xa23*
TTTGGTCTCAA**ATG**TTGCATCTCAAGGAACTCGCAGCCGTAGCCGGCATCGACATGATTGTCATCTACCTCTGCC
GCTTTCTCCTCCGCCGCTGCCGCAGCCGCAACGCATCCAACAGCCTCCGTTTTCGCCTCAAGGTATTAACTGTAT
TGTTGTACATATCACTCTCAACCATGCTGTTGTACCTGTTTGGCTCCATCGTGCCGCTGCCACCGTGGGGTGCTC
TGGCCGGTTGGGTCATGGCCCTCATCGCCGTCGAGCTCGCCTACGCCTTCCTCTTTCCATATAGCTTGCGCTACA
TCGCTCACAACGACGACGAGGACAAGATGGTTATTCCCCCTGTT**TAA**GCTTTGAGACCAAA

Code #3 *O. glumipatula* GEN 1233 *Xa23*
TTTGGTCTCAA**ATG**TTGCATCATCTCAAGGAGCTGGCAGCCGTAGCCGGTATACACATGATCCTCATCTACCTCT
GCCGCTTTCTCCTCCGCCGCAGCCGCAACGTATTATTCACCGTTTCCAACAGCCTCCGTTTTCGCCTCAAGGTAT
TAACTGTATTGTTGTACATATGTCTCTCGGTCATGCTGTTCTACCTGTTTGGCTCCATCATGCCGCTGCCGCCGT
GGGGCCTCGTGGTCGGTTGGGTCACGGCCCTCATCGCCGTCGAGCTCGCCTACGCCTTCATCTTTCCATATAGCT
TTCGCTACATTGCTGACAACGACGACGACAAGATGGTTATTCTCCCTGTT**TAA**GCTTTGAGACCAAA

Code #4 *O. longistaminata* RD23 *Xa23-1*
TTTGGTCTCAA**ATG**TTGCACCATCTCAAGGAGCTCGCAGCCGTAGCCGGTACACACATGATCGTCATCTTCCTCT
GCCGCTTTCTCCTCCGCCTCCGGCGCCGCAGCGCATCCAACAGCCTCCGTTTTCGCCTCAAGGTAGTAACTGTAC
TGTTGTACATATCCCTCTCGGCCATGCTATTCTACCTGTTTGCCTCCATCATGCCGCTGCCGCCGTGGGGCGCTG
TGGCCGGTTGGGTCATGGCCCTCATCGCCGTCGAGCTCGCCTACGCCTTCCTCTTTCCATACAGCTTGCGCTACA
TCGACGACGACAATATGGTTATTGATCCCCCTGTT**TAA**GCTTTGAGACCAAA

Code #5 *O. longistaminata* RD23 *Xa23-2*
TTTGGTCTCAA**ATG**TTGCATCTCAAGGAACTCGCAGCCGTAGCCGGTATACACATGATCCTCATCTACCTCTGCC
GTTTTCTCCTCCGCCTTCGCCGCAACGCATCATTCACCACCTCCAACAGCCTCCGTTTTCGCCTCAAGGTATTAA
CTGCATTATTGTACATATCCCTCTCAGCCATGCTGTTCTGCCTGTTTGCCTCCATCATGCCGCTGCAACCGTGGG
GCGCTGTGGCCGGTTGGGTCATGGCCCTCATCGCCGTCGAGCTCGCCTACGCCTTCCTCTTTCCATATTGCTTGT
GCTACTTCGCTGACAACCACGACGACAAGATGGTTATTCCCCCTGTTTATGCTTTGAAGAAACTAAAGCCTCACA
TACTGTCTCTTCAA**TAA**GCTTTGAGACCAAA

Code #6 *O. longistaminata* RD23 *Xa23-3*
TTTGGTCTCAA**ATG**TTGCATCATCTCAAGGAGCTCGCAGCCGTAGCCGGTATACACATGATCCTCATCTACCTCT
GCCGTTTTCTCCTCCGCCTCCGCCGCCGCAGCGCATCCACCAGCCTCCGTTTTCGCCTCAAGGTATTAACTGTAT
TGTTGTACATATCCCTCTCAGCCATGCTGTTCTACCTGTTTGCCTCCATCATGCCGCTGCCGCCGTGGGGCGCTG
TGGTCGGTTGGGTCATGGCCCTTATGGCCGTCGAGCTCGCCTACGCTTTTATCTTTCCGTATAACTTGCGATACA
TAGCTGACAACGGCGACGACAAGATGGTTATCCCCCCTGTT**TAA**GCTTTGAGACCAAA

Code #7 *O. longistaminata* RD23 *Xa23-4*
TTTGGTCTCAA**ATG**TTGCATCTCAAGGAGCTCGCAGCCGTAGCCGGCATCCACATGATCGTCATCTACCTCTGCC
GCTTTCTCCTCCGCCTTCGCCACAGCGCATTCACCACCTCCAACAGCCTCCGTTTTCGCCTCAAGGTTTTAACTG
CATTGTTATACATATCCCTCTCAGCCATTGCTGTTCTACCTGTTTGCCTCCATCATGCCGCTGCAACCGTGGGGC
GCTGTGGCCGGTTGGGTCATGGCCCTCATCGCCGTCGAGCTCGCCTACGCCTTCCTTTTTCCATA**TAG**GCTTTGA
GACCAAA

Code #8 *O. longistaminata* RD23 *Xa23-5*
TTTGGTCTCAA**ATG**CTGCATCTCAAGGAACTCACCGCTGCTATAGCCGGTGTCCACATGATCGTCATCTACCTCT
GTCGCTTTCTCCTCCGCCTTCGCCGCAGTGCATTCACCACCTCCAACAGCCTCCGTTTTCGCCTCAAGGTATTAA
TTGTATTGTTGTACGTATCCCTCTCAACCATGCTGTTCTACCTATTTGCCTCCATCATGCCGCTGCCACCGTGGG
GCGCCGTGGCCGGTTGGGTCATGGCCCTCATCGCCGTCGAGCTCGCCTACGCCTTCCTCTTTCCATATAGCTTGG
GATACATCCCTGAGAACAACGACGACAAGATGGTTATCCCCCCTGTT**TAA**GCTTTGAGACCAAA

Code #9 *O. meridionalis* OR44 (W2112) *Xa23*
TTTGGTCTCAA**ATG**TTGCATCATCTCAAGGAGCTGGCAGCCGTAGCCGGTATACACATGATCCTCATCTACCTCT
GCCGCTTTCTCCTCCGCCGCAGCCGCAACGTATTATTCACCGTTTCCAACAGCCTCCGTTTTCGCCTCAAGGTAT
TAACTGTATTGTTGTACATATGTCTCTCGGTCATGATGTTCTACCTGTTTGGCTCCATCATGCCGCTGCCGCCGT
GGGGCCTCGTGGTCGGTTGGGTCATGGCCCTCATCGCCGTCGAGCTCGCCTACGCCTTCATCTTTCCATATAGCT
TTCGCTACATCGCTGACAACGACGACGACAAGATGGTTATTCTCCCTGTT**TAA**GCTTTGAGACCAAA

Code #10 *O. officinalis* W0002 *Xa23-1*
TTTGGTCTCAA**ATG**TTGCATCTCAAAAAGCTCGCAGCCGTAACCGGCATACACATGATCGTCATCTACCTCTGCC
GCTTTCTCCTCCGCCGCTGCCGCAGCGCATCCACCACCTCCAACAGCCTCTGTTTTCGCCTCAAGATATTAACTG
CATTGTTGTACGTATCTCTCTCAGCCATGCTGTTCTACCTGTTTGCCTCCATCATGCCGCTGCCACCGTGGGGCG
CTGTGGCCGGTTGGGTCATGGCCCTCATCGCCGTCGAGCTCGCCTACGCCTCCCTCTTTCCATATAGCTTGCGCT
ACATCAATGAGAACGACGACGAGGAGATGGTTACTCCCCCTGTT**TAA**GCTTTGAGACCAAA

Code #11 *O. officinalis* W0002 *Xa23-2*
TTTGGTCTCAA**ATG**TTGCATCTCAAGAAGCTCGCAGCCGTAGCCGGCATACACATGATCGTCATCTACCTCTGCC
GCTTTCTCCTCCGCCGCTGCCGCAGCGCATCCACCACCTCCAACAGCCTCCGTTTTCGCCTCAAGATATTAACTG
CATTGTTGTACGTATCTCTCTCGGCCATTCTGTTCTACCTGTTTGCCTCCATCATGCCGCTGCCGCCGTGGGGCG
CTGTGGCCGGTTGGGGCGTGGCCCTCATCGCCGTCGAGCTCGCCTACGCCTTCCTCTTTCCGTATAGCTTGCGCT
ACATCGCTGACAACGACGACGAGATGATGGTTATTCCCCCTGTT**TAA**GCTTTGAGACCAAA

Code #12 *O. punctata* IRGC 105690 *Xa23*
TTTGGTCTCAA**ATG**TTGCACATGAAGGAACTCGCAGCCGTAGCCGGCATCCACATGATCGTCATCTACCTCTGCC
GCTTTCTCCTCCGCCGCAGCGCATCCACCACCTCCAATAGCCTCCGTTTTCGCCTCAAGGTATTAACTGTATTGT
TGTACGTTTCCCTCTCAGCCATGCTGTTCTACCTGTTTGCCTCCATCATGCCGCTGCCGCCGTGGGGCGCTGTGG
CCGGTTGGGTCATGGCCCTCATCGCCGTCGAGCTCGCCTACGCCTTCCTCTTTCCATATAGCTTGCGCTACATCG
CTGACAACGACGACGACATGGTTGTCCCCCCTGTT**TAA**GCTTTGAGACCAAA

Code #13 *O. rufipogon* Taxon A *Xa23*
TTTGGTCTCAA**ATG**TTGCATCATCTCAAGGAGCTCGCAGCCGTAGCCGGTATACACATGATCCTCATCTACCTCT
GCCGCTTTCTCCTCCGCCTCCGGCACAACGCATCCAACAGCCTCCGTTTTCGCCTCAAGGTTTTAACTGCATTGT
TATACATATCCCTCTCAGCCATGCTGTTCTACCTGTTTGCCTCCATCATGCCGCTGCCGCCGTGGGGAGCTGTGG
CCGGTTGGGTCATGGCCCTCTTCGCCGTTGAGCTCGCCTACGCTTTTATCTTTCCGTATAACTTGCGATACATAG
CTGACAACCACGACGAGGACAATATGGTTATTCCCCCTGTT**TAA**GCTTTGAGACCAAA

Code #14 *O. rufipogon* W1943 *Xa23*
TTTGGTCTCAA**ATG**ATCCTCATCTACCTCTGCCGCTTTCTCCTCCGCCGCAGCCGCAACGTATTATTCACCGTTT
CCAACAGCCTCCGTTTTCGCCTCAAGGTATTAACTGTATTGTTGTACATATGTCTCTCGGTCATGCTGTTCTACC
TGTTTGGCTCCATCATGCCGCTGCCGCCGTGGGGCCTCGTGGTCGGTTGGGTCATGGCCCTCATCGCCGTCGAGC
TCGCCTACGCCTTCATCTTTCCATATAGCTTTCGCTACATCGCTGACAACGACGACGACAAGATGGTTATTCTCC
CTGTT**TAA**GCTTTGAGACCAAA

Code #15 *O. sativa* Nipponbare *Xa23*
TTTGGTCTCAA**ATG**TTGCATCATCTCAAGGAGCTGGCAGCCGTAGCCGGTATACACATGATCCTCATCTACCTCT
GCCGCTTTCTCCTCCGCCGCAGCCGCAACGTATTATTCACCGTTTCCAACAGCCTCCGTTTTCGCCTCAAGGTAT
TAACTGTATTGTTGTACATATGTCTCTCGGTCATGCTGTTCTACCTGTTTGGCTCCATCATGCCGCTGCCGCCGT
GGGGCCTCGTGGTCGGTTGGGTCATGGCCCTCATCGCCGTCGAGCTCGCCTACGCCTTCATCTTTCCATATAGCT
TTCGCTACATCGCTGACAACGACGACGACAAGATGGTTATTCTCCCTGTT**TAA**GCTTTGAGACCAAA

Code #16 *O. australiensis* KR-Keep River_1 *Xa10-Ni*
TTTGGTCTCAA**ATG**AAGCTCACTGGGTGCGCGCTTATCCTCATCGCTGTCCACGTAGCGGCGGCCATCTCCTGCC
CCGAGCCACTGGCGGCCATGGCCGTCGTGGACGTGCCGCTCCTCTTCCTCTGCCGCTACCTCCTACGCCGCCGCG
ACGACGACGGCGGTGGTAGGCTCAAGTTCCGGCTCAAAGCCGCCGTCTTGGTTCTCTACATGGTCATGTCGGCGC
TCTTCTTCCTCCGCTTTGCGCCAATCACGCCGTGGTGGGGCGCCGTTGCCGCCTGGGCCATGATGCTGCTCATCC
TTGAGCTCATCTTCGCCTTCTTCTTCTTCCCCTACAGCCACTGCAACAAGCTCGGTTGGGACGTGGAGAATCCGG
GAGGCATTTTAACT**TAA**GCTTTGAGACCAAA

Code #17 *O. glumipatula* GEN 1233 *Xa10-Ni*
TTTGGTCTCAA**ATG**GAGCTCATCACCGCGTGCGCGCTTCTCCTCATCGCCGTCCATATTGCGATTGTCATCTCCT
GCCCGGAGCCACTCGCGGCCGTTGCATTAGTCGACGTGCCCCTCCTCTACCTCTGCCGCTACCTCCTCATCATAC
GCCGCGGTGTGATCATCGTCTCCGGCGCCGGCGGCGGCGGCAGGTTGCGCCGCTTCCGCCTCGGAGCAGCCATGG
CGATCATCTACATGGCCATGTCGACCCTCTTCTTCCTCCGCATTGCTCCTCTCACGCCATGGTGGGGCGCCCCTCG
CCGCCTGGGTCATGATCCTGCTCATCGTCGAGGCCATCTTCGCCTTCTTCTTCCCCTACCGCTGCTTGCTTCAAC
GAAACCGAAGGCGATGTCCAGAA**TAA**GCTTTGAGACCAAA

Code #18 *O. longistaminata* RD23 *Xa10-Ni*
TTTGGTCTCAA**ATG**GAGCTCATCACCGCGTGCGCGCTTCTCCTCATCGCCGTCCACATTGCGATTGTCATCTCCT
GCCCGGAGCCACTCGCGGCCGTTGCATTCGTCGACGTGCCCCTCCTCTACCTCTGCCGCTACCTCCTCATCATAC
GCCCCGGCGTGATCATCGTCTCCGGCGCCGGCGGCAGCGGCAGGATGCGCCGCTTCCGCCTCGGAGCAGCCATGG
CGATCATCTACATGGCCATGTCGACCCTCTTCTTCCTCCGCATTGCTCCTCTCACGCCATGGTGGGGCGCCCCTCG
CCGCCTGGGTCATGATCCTGCTCATCGTCGAGGCCATCTTCGCCTTCTTCTTCCCCTACCGCTGCTGCTTCAACG
AAACCGAAGGCGATGTCCAGAATAATCCCCATGTC**TGA**GCTTTGAGACCAAA

Code #19 *O. longistaminata* W11 *Xa10-Ni*
TTTGGTCTCAA**ATG**AAGCTTAATTTGTGTGCGCTGCTCGTCATCGCTGTCCACGTAGCGGCGGCCATCTCCTGCC
CCAAGCCACTCGCGGCCATGGCCATCATCGACGTGCCGCTCCTCTTCCTCTGCCGCTACCTCATACGCCGCCGCG
ACGACGACTTCGTCGTCTCCAGCGGTGGCGGCGGTGGTAGGCTCCAGTTCCAGCTCAAAGCCGCCGTCTTGGTTC
TCTACATGGTCATGTCGGCGCTCTTCTTCCTCCGCTTTGCGCCAATCACGCCGTGGTGGGGCGCCGTTGCCGCCT
GGGCCATGATGCTGCTCATCCTTGAGGTCATCTTCGCCTTCTTCTTCTTCCCCGACGGCTACAGCAACAAGCTCC
GTTGGGACGTGGAGAATCCGGGAGGTATTTTAGCT**TAA**GCTTTGAGACCAAA

Code #20 O. *meridionalis* OR44 (W2112) *Xa10-Ni*
TTTGGTCTCAA**ATG**GAGCTCATCACCGCGTGCGCGCTTCTCCTCATCGCCGTCCATATTGCGATTGTCATCTCCT
GCCCGGAGCCACTCGCGGCCGTTGCATTAGTCGACGTGCCCCTCCTCTACCTCTGCCGCTACCTCCTCATCATAC
GCCGCGGTGTGATCATCGTCTCCGGCGCCGGCGCCGGCGGCGGCGGCAGGTTGCGCTGCTTCCGCCTCGGAGCAG
CCATGGCGATCATCTACATGGCCATGTCGACCCTCTTCTTCCTCCGCATTGCTCCTCTCACGCCATGGTGGGGCG
CCCTCGCCGCCTGGGTCATGATCCTGCTCATCCTCCCCTACCGCTGCTGCTTCAACGAAACCGAAGGCGATGTCC
AGAATAATCCCCATGTC**TGA**GCTTTGAGACCAAA

Code #21 *O. meyeriana* Menghai *Xa10-Ni*
TTTGGTCTCAA**ATG**CTCATCACCGTCCACATTTTGATTGTCATATCCTGCCCGGAGCTGTTCACAGCCATTGCCC
TCGTCGATGTGTCCATCCTCCACCTCTGCCGCTACCTCCTGCACCACGGCGTCTCCGTCATCTCCGGCGCCGGCA
TCAGGCTCCGGTTTCGCCTTCGAACAGCTGACGCGATCATCTACATGGCCCTGTCCGTCCTCCTCTTCCTCCGCT
TTGCTCCGGTCATGTCGTGGTGGGGCGCGCTCGCCGCCTGGGTCACCATCCTGCTCATCGTCGAGGCCGTCTTGG
CCTTCTTCTTCTTCTTCTTCCCCTACGGCTGCTGCAACGAGCGGCCTGAAGGCGACGTCCACAATCCCCATGTC**T**
**GA**GCTTTGAGACCAAA

Code #22 *O. officinalis* W0002 *Xa10-Ni*
TTTGGTCTCAA**ATG**GTACTCACCGCGTACGCGCTTCTCCTCATCCTCCTTCACGTAGCCATCGGCATCTCCTGCC
CCGAGCCACTCGCTGCCATGGCCTTCGTCGACGTGCCCCTCCTCTTCCTCTGCCGCTACCTCCTGCTCCGTGGCG
CCTCCGTCTTCTTCGCCGGCGACGGCAGGCTCCGCTTCCGTCTCAAAGCCGCCGTGGCGATACTTTACATGGCCC
TGTCGGCCCTCTTCTTCCTCCGATTTGCTCCCCTCATGCCGTGGTGGGGTGTCCTCGTAGCCTGGGTCACGATGG
TGCTCATCGTTGAATTCGTGTACGCCTTCTTCTTCCACTACAGCTGCTACAACGAACTCGACGGTTATGTGGAGA
ATCCCGTTGTCGGA**TGA**GCTTTGAGACCAAA

Code #23 *O. rufipogon* Taxon A *Xa10-Ni*
TTTGGTCTCAA**ATG**GAAATCCTCATCACCGCCTGTGCGCTTCTCCTCATTGCCGTCAACGTCGTAGTGACGGCCA
TATCCTGCCCCAATCCACTCGCAGCCATCGTCCTCGTCGACGTGCCCGTGCTCTACCTCTGCCGCTACCTCCTGC
GCCGCGGCGTCCCCGTCGTCACCGGCGCCGGCGTCGTCAGGCTCCGCTTCCGCCTGGTCGCAGCGGTGGCGACCA
TCTACATGGCCATGCCAGCGCTCTTCTTCCTCCGCTTTGCTCCGGTCATGCCGTGGTGGGGGGGCGCTCGCCGCC
TGGGTGACCATCCTGCTCATCGTCGAGGCCGTCTTCGCCTTCTTCTTCTTTCCCTACAGCTGCTGCAACGATCAG
CC**TGA**GCTTTGAGACCAAA

Code #24 *O. sativa* Nipponbare *Xa10-Ni*
TTTGGTCTCAA**ATG**GAGCTCATCACCGCGTGCGCGCTTCTCCTCATCGCCGTCCATATTGCGATTGTCATCTCCT
GCCCGGAGCCACTCGCGGCCGTTGCATTAGTCGACGTGCCCCTCCTCTACCTCTGCCGCTACCTCCTCATCATAC
GCCGCGGTGTGATCATCGTCTCCGGCGCCGGCGCCGGCGGCGGCAGGTTGCGCCGCTTCCGCCTCGGGGCAGCCA
TGGCGATCATCTACATGGCCATGTCGACCCTCTTCTTCCTCCGCATTGCTCCTCTCACGCCATGGTGGGGCGCCC
TCGCCGCCTGGGTCATGATCCTGCTCATCGTCGAGGCCATCTTCGCCTTCTTCTTCCCCTACCGCTGCTGCTTCA
ACGAAACCGAAGGCGATGTCCAGAATAATCCCCATGTC**TGA**GCTTTGAGACCAAA

Code #25 *L. perrieri* IRGC 105164 *Xa10-1*
TTTGGTCTCAA**ATG**GAGATGTTCTTTGTTTGGTCGCTGCTTCTCGTCGCCATGATCCTGGCGATCTTGTGGCTGA
AGCAGCTCGCAGTCGCAGTCGCAGCCTTCATCCACGTGCTCGTCATCTACCTCTGCCGCTTCACCCTCCTCCGCG
GCGGCGACGATGGAGACAACCTCTGCAAATTCCGTATTAAGGTAGCTGTCGGTATCATGTACATCTCTCTCTCAG
CTATACTGCTGTGCCTGTTTGCACCTGATATGCTGCCGCCATGGGGCCGCGTTGGGCATGGCCCTCGTTGTCAT**T**
**GA**GCTTTGAGACCAAA

Code #26 *L. perrieri* IRGC 105164 *Xa10-2*
TTTGGTCTCAA**ATG**GGTCGATCGACGCAGCTGATCACATTCTGCGTGGGATCAATCCTGATTGCCGTTCTCGCAG
TGATGATCTTCCTCAACCAACTCGCCGGCGCAGCCTTCGTCTACGTGCTCATAATCTACCTCTGCCGCTTCGTCC
TCCTCCGGCGACGCATCTTCTCCGATCACAGCAACATACTATTTCGTGTTAAGGCAGGTATCGGGCTCATGTACA
TATTTCTCTTAGCCATACTGTTGTACATCTCAGCCGCAGTCATGAAGTTACCGCCGTGGGGCGCCGTCGCCATGT
CGGTTATGGCGTTTGCCGCTGTTGAGCTTGGCTACGCATTATTCTTCCCGTACACATGCCGCTGCATTGATGACC
TGGACGAGGTGAATCCCCCTGTT**TAA**GCTTTGAGACCAAA

Code #27 *O. brachyantha* IRGC 101232 *Xa10-1*
TTTGGTCTCAA**ATG**CAGCTGACAGCATTATCCGCGGGATCCCTCCTCCTGTTTGCTTCTCTCTTAGCGATGGCAT
ACCTGAAGCTACTCGCCGGCGCAGCCTGCGTCCTCGTGCTCACCATCTACCTCTGCCGCTTCCTCCTCCGCCGCC
GCATCTTCTCCGACGACCACCACAACCACCTACGGTTCCGTCTCAAGGTAGGGATCGGATTCATGTACATCTCCC
TCTCGGCCATACTCTTCTACCTCTCTGCCTCTGTCATGGCCTTGCCGCCATGGGGTGCCGCCGCCATGTGGGGTA
TGGCGCTCGTCGCCGTTGAGCTTGTCTACGCGTTCTTCTTCCCCTATAGCTGCATCTGCATTGGTGACGATGCCG
ACGAGACTCCCCCTCTTTCTGTT**TGA**GCTTTGAGACCAAA

Code #28 *O. brachyantha* IRGC 101232 *Xa10-2*
TTTGGTCTCAA**ATG**CTATACCTGAAGCAACTCGCCGGCGCAGCCTGCGTCCTCGTGCTCACCATCTACCTCTGCC
GCTTCCTCCTCCTCCGCCGCGTCTTCTCCGACGACCACAAGCTACGGTTCCGTCTCAAGGTAGGGATCGGATTCC
TGTACATCTCCCTCTCAGCCGTACTCTTCTACCTCTCCGCCGCCGTCATGGCGTTGCCTCCATGGGGTGCCGTCG
CCATGTGGGGTATGGTGGTCGTCGCCGTTGAGCTTGCCTACGCGTTCTTCTTCCCGTACAGCTGCATCTGCATCG
GCGACGACGCCGACAAGACTCCCCCTCTTTCTGTT**TGA**GCTTTGAGACCAAA

Code #29 *O. coarctata* TKM-2016 *Xa10*
TTTGGTCTCAA**ATG**GCTAGGTTAGAGATCATCTCCGCTTGCGCGGCGCCGCTCGTCATCGCCATGCTTGTGGTGA
TGGTGTGGCTCAACCAGCTCGCTGTGGCGGCCTACATCCACGTGGGCATCATCTACCTCTGCCGCTTCGTCCTCC
TCCGCCGCGGCGGCTCCGGCGACAAACTGACCCGCTTCCGCCTCAAGCTAGCCATCGGAATCCTGTACATATCTC
TCTCAGGCATGTTGGTGTACCTCTTCTCCCCCGTCATGGCGGCAGCACCCTGGGGCGGCGTCGCCGTGTGCGCGA
CGGCGCTCGTCGCCACTGAGCTCGGGTTTGCATTTCTCTCTTCCTATAGCTGCTGCATTGATGAAGACGAGGAGA
GTCGGCCTCTG**TAG**GCTTTGAGACCAAA

Code #30 *O. longistaminata* RD23 *Xa10*
TTTGGTCTCAA**ATG**CAGCTGATTATCACATTCTTCACGGGGTCCCTCGTGTTCGCGGTTCACGTACTGATGGTAT
ACCTCAAGCAACTCGCCGGCGCGGCCTGCGTCGACGTGCTCATCATCTACCTCTGCCGCTTCCTCCTCCTCCGCG
GCATCTTCTTCTCCGACGACGACAAGCTACGATTCCGCGTCAAGGTAGCGATCGGGTTCCTCTACATCTCCCTCT
CGGCCATACTCTTCTACCTCTCTGCCGCTGTCATGGCGTTGCCGCCGTGGGGTGCCGTCGCCATATGGGGGATGG
CGCTCGTCGCCATTGAGCTTGGCTACGCGTTCTTCTTCCCGTATCGCTGCCTCTGCATTCGTAAGGATGACGACG
AGGAGACTCCCTCTGTTTCTGTT**TGA**GCTTTGAGACCAAA

Code #31 *O. meyeriana* Menghai *Xa10*
TTTGGTCTCAA**ATG**GAGGTGATCACCGTTTGCGCTTTGCTTCTCCTCACCCTTGTCCTAGTGCTGGTCTGCCTCA
AGCAACTTGCAGTCGTAGCCTGCTTCCACGTGCTCATCATCTACCTCGGCCGCTTCGTCCTCCTCCACGGCGGCG
ACGACAGGCTCCGGTGCTTTCGCATTAAGGTAGCCGTGGGAATCCTCTACCTCACTCTGTCAGCCATATTGTTCT
GCCTCTTTGGCCCCGTCATGCCGCCGTGGGGCGCCGTCGCCGCCTGGGCGATGAGCCTCGCTGCCATTGAGCTCG
GATACGCCTTCTTCTTCCCCTACAGCTGCTACCACATTGACGACGAACACGAGGAGATGATGATCAACCCCACTG
TTGGAGTGTCTGTT**TAA**GCTTTGAGACCAAA

Code #32 *O. nivara* IRGC 100897 *Xa10*
TTTGGTCTCAA**ATG**CAGCTGATGCTCACATTCTGCACGGGCCCCCTCCTGTTTGCCGTCCTCCTACTGATGGTAT
ACCTCAAGCAACTCGCCGCCGCGGCCTGCGTCGACGTGCTCATCATCTACCTCTGCCGCTTCCTCCTCCTCCGCG
GCATCATCTTCTCAGGCGACGGCAAGCTACGATTCCGCGTCAAGGTAGCGATCGGGTTCCTCTACATCTCCCTCT
CGGCCATACTCTTCTACCTCTCTGCCGCTGTCATGGCGTTGCCGCCGTGGGGTGCGGTGGCCATGTGGGGAATGG
CGCTCGTCGCCACTGAGCTTGGCAACTCCTTCTTATGCCCGTATAGCTGCCGCTGCATTGGTGAAGACGACGAGG
AGATTTCCCCCGTC**TGA**GCTTTGAGACCAAA

Code #33 *O. officinalis* W0002 *Xa10-1*
TTTGGTCTCAA**ATG**CAGCCGATCACATTCTGCACGGGCCCCCTCCTGTTTGCCGTCCTCCTACTGATGGTATACC
TCAAGCAACTCGCCGGCGCGGCCTGCGTCGACGTGCTCATCATCTACCTCTGCCGCTTCCTCCTCCTTCGCGGCA
TCTTCCCCGACGACGACAAGCTAAGATTCCGTGTCAAGGTAGCGATCGGCACTACTACAGAAAGGATTTTTGCAG
ACGGGCGGATTCGATCTTCGCATGCGGTTGGCCGGGCCGCATGCGCGAAAAGGTCTGTGAAGATCACCATTTGCG
CATGCGGCCGGAGC**TAG**GCTTTGAGACCAAA

Code #34 *O. officinalis* W0002 *Xa10-2*
TTTGGTCTCAA**ATG**CAGCCGATCACATTCTGCACGGGCCCCCTCCTGTTTGCCGTCCTCCTACTGATGGTATACC
TCAAGCAACTCGCCGGCGCGGCCTGCGTCGACGTGCTCATCATCTACCTCTGCCGCTTCCTCCTCCTTCGCGGCA
TCTTCTCCGACGACGACAAGCTAAGATTCCGTGTCAAGGTAGCGATCGGGTTCCTGTACATCTCCCTCTCCGCCA
TACTGTTTTACCTCTCCGCCGCTGTCTTGGCGTTGCCGCCGTGGGGTGCCGTCGCCATGTGGGGTATGGCGCTCG
TCGCCAGTGAGCTTGGCTACGCGTTCTTCTTCCCGTATACCTGCCTCTGCATCGGTGAGGATGACGACGAGGAGA
TTCCCTCTGTTTCTGTT**TGA**GCTTTGAGACCAAA

Code #35 *O. punctata* IRGC 105690 *Xa10-1*
TTTGGTCTCAA**ATG**CAGCTGATGATCACATTTATCTTGACGGGGGCCCTCCTGTTTGCCGTCCTCGTAGTACTGA
TGGTATGCATCAAGCAACTCGCCGTCGCGGCCTGCGCCCACGCGCTCATCATCTACGTCTGCCGCTTCCTCCTCC
TCCGCGGCATCTTCTCCGACGACGGCAAGCTACGACTCCGTGTCAAGGTAGCGATCGGGTTCCTGTACATCTCCC
TCTCAGCCATACTGTTCTACCTCTCTGCCGCTGTCATGGCGTTGCCGCCGTGGGGTGTCGTCGCCATGTGGGGTA
TGGCGCTCGTCGCCACTGAGCTTGGCTACGCATTCTTCTTCCCGTACAGATGCCTCTGCATTGGTGACGACGACG
ACGAGATTACCCCTGTCGATCTGAGGTCCATAACTATTTCAGCA**TAG**GCTTTGAGACCAAA

Code #36 *O. punctata* IRGC 105690 *Xa10-2*
TTTGGTCTCAA**ATG**GAGTTGATCATCACTGTTTGCGCTTCGATCCTCGTCGCCATTATCATCGTAGCGATGATCT
TCCTCAAGCAACTCGCAGTCGCGGCCTGCGTCCACGTGCTCGTCATCTACATCTGCCGCTTCGTCCTCCTCCGCG
GCGGTGGCGGCTCTTCCTCCGGCGACGCCGAGAGGCCGCTCCGCTGCTTTCGCATTAGGGTAGCCGTTGGAATCC
TATACCTCTCTATGTCAGCGATATTGTTCGGCCTCTTTGTCTCAGTCATGCCGCCGTGGGGCGCCGTCGCCGTGT
GGGGCATGGCCCTCGCCGCCGTTGAGCTCGGCTACGCCTTCTTCTTCCCCTATACCTGCTACCACGTCGGTGGAG
ACGACGACGACGACGACGACAATATTCCTATTACCCATATT**TAA**GCTTTGAGACCAAA

Code #37 *O. sativa* IRBB10A *Xa10* (JX025645)
TTTGGTCTCAA**ATG**CAGCTGATGCTCACATTCTGCACGGGCCCCCTCCTGTTTGCCGTCCTCCTACTGATGGTAT
ACCTCAAGCAACTCGCCGCCGCGGCCTGCGTCGACGTGCTCATCATCTACCTCTGCCGCTTCCTCCTCCTCCGCG
GCATCATCTTCTCCGGCGACGGCAAGCTACGATTCCGCGTCAAGGTAGCGATCGGGTTCCTCTACATCTCCCTCT
CGGCCATACTCTTCTACCTCTCTGCCGCTGTCATGGCGTTGCCGCCGTGGGGTGCGGTGGCCATGTGGGGAATGG
CGCTCGTCGCCACTGAGCTTGGCTACTCCTTCTTATGCCCGTATAGCTGCCGCTGCATTGGTGAAGACGACGAGG
AGATTTCCCCCGTC**TGA**GCTTTGAGACCAAA

Code #40 *A. victoria GFPi*
TTTGGTCTCAA**ATG**TCTAAGGGCGAAGAACTCTTCACCGGCGTGGTGCCGATCCTCGTTGAACTCGATGGTGACG
TGAACGGCCACAAGTTCTCCGTTTCTGGCGAAGGTGAAGGCGACGCCACATACGGCAAGCTCACCCTCAAGTTCA
TCTGCACCACAGGCAAGCTCCCGGTGCCATGGCCAACACTCGTGACCACCTTCACCTACGGCGTGCAATGCTTCT
CTCGCTACCCGGACCATATGAAGCAGCACGACTTCTTCAAGTCCGCCATGCCAGAGGGCTACGTGCAAGAGAGGA
CCATTTTCTTCAAGGACGATGGCAACTACAAGACACGCGCCGAGGTAAGTTGTTACTTATGATTGTTTTCCTCTC
TGCTACATGTATTTTGTTGTTCATTTCTGTAAGATATAAGAATTGAGTTTTCCTCTGATGATATTATTAGGTCAA
GTTCGAGGGCGATACCCTCGTGAACCGCATCGAGCTGAAGGGCATCGACTTTAAAGAGGACGGCAACATCCTGGG
CCACAAGCTCGAGTACAACTACAACTCCCACAACGTGTACATCATGGCCGACAAGCAAAAGAACGGCATCAAGGT
GAACTTCAAGATCAGGCACAACATCGAGGACGGCTCCGTGCAGCTCGCCGATCATTACCAGCAGAACACCCCAAT
TGGCGACGGCCCAGTGCTCCTGCCAGATAACCATTACCTCAGCACCCAGAGCGCCCTCAGCAAGGATCCAAACGA
GAAGCGCGACCACATGGTGCTCCTCGAGTTCGTTACAGCCGCTGGCATCACCCACGGCATGGACGAGTTGTACAA
A**TAA**GCTTTGAGACCAAA

**Data S10. Publicly available RNA-seq datasets from the RAP-DB database screened for RNA-seq reads mapping to *O. sativa Xa10-Ni* and *Xa23* loci.**[2]

1. Strand-specific RNA-seq from 8 tissues, such as flower, leaf, root, seed, etc. published by Wang *et al*. (2015). *The Plant Journal*. DOI: 10.1111/tpj.13018
2. Total RNA-seq from 8 tissues, such as flower, leaf, root, seed, etc. published by Wang *et al*. (2015). *The Plant Journal*. DOI: 10.1111/tpj.13018
3. Strand-specific RNA-seq from shoot, seed, anther, and pistil published by Zhang YC *et al.* (2014). *Genome Biology*. DOI: 10.1186/s13059-014-0512-1
4. Strand-specific mRNA-seq (leaf) published by Lu *et al.* (2019). *Nature Plants*. DOI: 10.1038/s41477-019-0548-z
5. Strand-specific mRNA-seq from seedlings under salinity, cold, and drought stress conditions published by Lu *et al.* (2012). *BMC Genomics*. DOI: 10.1186/1471-2164-13-721
6. Strand-specific mRNA-seq from shoot under dehydration and rehydration stress conditions published by Park *et al.* (2023). *Int J Mol Sci*. DOI: 10.3390/ijms24098439
7. Strand-specific mRNA-seq from shoot and root under nitrogen starvation published by Shin *et al.* (2018). *BMC Genomics*. DOI: 10.1186/s12864-018-4897-1
8. Strand-specific mRNA-seq from lamina joint, *OsDCL3a* RNAi mutant published by Wei *et al.* (2014). *Proc Natl Acad Sci*. DOI: 10.1073/pnas.1318131111
9. Strand-specific total RNA-seq from root under phosphate starvation and recovery conditions, *dcl3a* RNAi mutant published by Secco *et al.* (2015). *eLife*. DOI: 10.7554/eLife.09343
10. mRNA-seq from root under phosphate starvation and recovery conditions published by Secco *et al.* (2015). *eLife*. DOI: 10.7554/eLife.09343
11. Strand-specific RNA-seq from root and shoot under iron-deficient conditions, *osbhlh156-1* mutant published by Wang *et al.* (2019). *New Phytologist*. DOI: 10.1111/nph.16232
12. Strand-specific RNA-seq from root after inoculation of arbuscular mycorrhizal fungus, *phr2* mutant published by Das *et al.* (2022). *Nat Commun*. DOI: 10.1038/s41467-022-27976-8
13. mRNA-seq from 7 tissues, such as leaf, root, panicle, callus, etc. published by Sakai *et al.* (2011). *Genome Biol Evol*. DOI: 10.1093/gbe/evr111
14. mRNA-seq from 9 tissues, such as leaf, flower, anther, still, etc published by Davidson RM *et al.* (2012). *The Plant Journal*. DOI: 10.1111/j.1365-313X.2012.05005.x
15. mRNA-seq from leaf published by Yang *et al.* (2019). *BMC Plant Biology*. DOI: 10.1186/s12870-019-1872-1
16. mRNA-seq from leaf and callus published by Wu *et al.* 2011 Plant Cell. DOI: 10.1105/tpc.111.090043
17. mRNA-seq from anther published by Komiya *et al.* 2014 Plant J. DOI: 10.1111/tpj.12483
18. mRNA-seq from egg, sperm, zygote published by Rahman *et al.* 2019 Plant Cell Physiol. DOI: 10.1093/pcp/pcz030
19. mRNA-seq from aleurone published by Watanabe *et al.* (2014). *Genomics*. DOI: 10.1016/j.ygeno.2013.10.007
20. mRNA-seq from root meristematic, elongation, and differentiation zones published by Huang *et al.* (2015). *Plant Cell*. DOI: 10.1104/pp.114.254623
21. mRNA-seq from culm tissue containing shoot apical meristem published by Song *et al.* (2015). *Proc Natl Acad Sci*. DOI: 10.1073/pnas.1421127112
22. mRNA-seq from root and root tips in early development published by Kyndt *et al.* (2012). *J Exp Bot*. DOI: 10.1093/jxb/err435
23. mRNA-seq under salinity stress conditions published by Mizuno *et al.* (2010). *BMC Genomics*. DOI: 10.1186/1471-2164-11-683
24. mRNA-seq under phosphate stress conditions published by Oono *et al.* (2011). *Rice*. DOI: 10.1007/s12284-011-9064-0
25. mRNA-seq under phosphate stress conditions published by Oono *et al.* 2013. *Plant Mol Biol*. DOI: 10.1007/s11103-013-0106-4

26. mRNA-seq under cadmium stress conditions published by Oono *et al.* (2014). *PLoS One*. DOI: 10.1371/journal.pone.0096946
27. mRNA-seq under cadmium stress conditions published by Kawahara *et al.* (2016). *Plant Cell Physiol*. DOI: 10.1093/pcp/pcv179
28. mRNA-seq under cold, flood, and osmotic stress conditions published by Kawahara *et al.* (2016). *Plant Cell Physiol*. DOI: 10.1093/pcp/pcv179
29. mRNA-seq under drought stress conditions published by Kawahara *et al.* (2016). *Plant Cell Physiol*. DOI: 10.1093/pcp/pcv179
30. mRNA-seq under JA treatment published by Kawahara *et al.* (2016). *Plant Cell Physiol*. DOI: 10.1093/pcp/pcv179
31. mRNA-seq under ABA treatment published by Kawahara *et al.* (2016). *Plant Cell Physiol*. DOI: 10.1093/pcp/pcv179
32. mRNA-seq under no treatment/control data published by Kawahara *et al.* (2016). *Plant Cell Physiol*. DOI: 10.1093/pcp/pcv179
33. mRNA-seq from root and shoot under phosphate starvation and recovery conditions published by Secco *et al.* (2013). *Plant Cell*. DOI: 10.1105/tpc.113.117325
34. mRNA-seq from shoot and root under arsenic stress published by Yu *et al.* (2012). *New Phytol*. DOI: 10.1111/j.1469-8137.2012.04154.x
35. mRNA-seq from coleoptile and the first leaf under shade published by Liu *et al.* (2016). *J Integr Plant Biol*. DOI: 10.1111/jipb.12468
36. mRNA-seq from leaf under drought stress condition in long day and short day published by Galbiati *et al.* (2016). *Plant Cell Environ*. DOI: 10.1111/pce.12760
37. mRNA-seq from shoot, *MET1-2* mutant published by Hu *et al.* (2014). *Proc Natl Acad Sci*. DOI: 10.1073/pnas.141076111
38. mRNA-seq from shoot, *osino80* RNAi mutant published by Li *et al.* (2018). *J Integr Plant Biol*. DOI: 10.1111/jipb.12603
39. mRNA-seq from whole tissue of above-ground seedling, *oscmt3a* mutant published by Cheng *et al.* (2015). *The Plant Journal*. DOI: 10.1111/tpj.12952
40. mRNA-seq from leaf after inoculation of *Magnaporthe oryzae* strains published by Kawahara *et al.* (2012). *PLoS One*. DOI: 10.1371/journal.pone.0049423
41. mRNA-seq from leaf after inoculation of *Xanthomonas oryzae* pv. *oryzicola* (*Xoc*) strains published by Wilkins *et al.* (2015). *Front Plant Sci*. DOI: 10.3389/fpls.2015.00536
42. mRNA-seq from leaf after inoculation of *Xanthomonas oryzae* pv. *oryzicola* (*Xoo*) strains published by Hummel *et al.* (2017). *Mol Plant Pathol*. DOI: 10.1111/mpp.12377
43. mRNA-seq from root after inoculation of arbuscular mycorrhizal fungus published by Wang *et al.* (2020). *Proc Natl Acad Sci*. DOI: 10.1073/pnas.2000926117
44. mRNA-seq from root tip and root after inoculation of root knot and root rot nematodes published by Kyndt *et al.* (2012). *New Phytologist*. DOI: 10.1093/jxb/err435
45. mRNA-seq from root vascular cells after inoculation of root knot nematode published by Ji *et al.* (2013). *J Exp Bot*. DOI: 10.1093/jxb/ert219
46. mRNA-seq from root under AM fungus-secreted substances treatment, *oscerk1* mutant published by Miyata *et al.* (2014). *Plant Cell Physiol*. DOI: 10.1093/pcp/pcu129

**Data S11. Alignment of the genomic *Xa23* sequence and putative full-length native *Xa23* transcripts reconstructed *in silico* using the published RNA-seq datasets**. Transcripts MSTRG.23300.1 and MSTRG.23300.2 derived from the dataset published by Rahman *et al.* [1] and transcripts MSTRG.28072.1 – MSTRG.28072.4 derived from the dataset published by Toda *et al.* [3]. The alignment was generated using the MUSCLE algorithm.[7] The dTALE binding element for transcriptional activation of Xa23 is underlined and highlighted in magenta.[13] The published 5'UTR, CDS, and 3'UTR of the dTALE-induced *Xa23* mRNA transcripts are highlighted in blue, yellow, and grey, respectively.[13] The Xa23 start (**ATG**) and stop (**TAA**) codons are colored in **red**. The first nucleotide of the *Xa23* start codon is designated as "+1".

```
              -500......-490......-480......-470......-460......-450......-440......-430......-420......-410.........
Xa23 locus    GACTAATTTTTTTTGGTGTACTGTAGTAGGAAAAAAAGAAGGGGAGGAGGAAGGGGGAGAAGAGGGAGGAGGTATTGCATATTGGGGGGAGGGGAATGCG
MSTRG.23300.1 ....................................................................................................
MSTRG.23300.2 ....................................................................................................
MSTRG.28072.1 .......................................................................GTATTGCATATTGGGGGGAGGGGAATGCG
MSTRG.28072.2 .......................................................................GTATTGCATATTGGGGGGAGGGGAATGCG
MSTRG.28072.3 .......................................................................GTATTGCATATTGGGGGGAGGGGAATGCG
MSTRG.28072.4 .......................................................................GTATTGCATATTGGGGGGAGGGGAATGCG

              -400......-390......-380......-370......-360......-350......-340......-330......-320......-310.........
Xa23 locus    GGGTGATCGCTTGGCGCGCATTAGCAGTCCCCCCTTCCCTTCCGCTATTTCAAAGTATCCGTCTTCCTCCCTGTCCACATGAGCTAAACGCTTCCTCTTC
MSTRG.23300.1 ....................................................................................................
MSTRG.23300.2 ....................................................................................................
MSTRG.28072.1 GGGTGATCGCTTGGCGCGCATTAGCAGTCCCCCCTTCCCTTCCGCTATTTCAAAGTATCCGTCTTCCTCCCTGTCCACATGAGCTAAACGCTTCCTCTTC
MSTRG.28072.2 GGGTGATCGCTTGGCGCGCATTAGCAGTCCCCCCTTCCCTTCCGCTATTTCAAAGTATCCGTCTTCCTCCCTGTCCACATGAGCTAAACGCTTCCTCTTC
MSTRG.28072.3 GGGTGATCGCTTGGCGCGCATTAGCAGTCCCCCCTTCCCTTCCGCTATTTCAAAGTATCCGTCTTCCTCCCTGTCCACATGAGCTAAACGCTTCCTCTTC
MSTRG.28072.4 GGGTGATCGCTTGGCGCGCATTAGCAGTCCCCCCTTCCCTTCCGCTATTTCAAAGTATCCGTCTTCCTCCCTGTCCACATGAGCTAAACGCTTCCTCTTC

              -300......-290......-280......-270......-260......-250......-240......-230......-220......-210.........
Xa23 locus    CCTGAGTCAAAGTCTTCCCTATATTACTACTTTGTATTATAGCTGGGTTCACTATGTTGCATCATCTCACAGCTAACCCGAACATCACTAACATCGCAAT
MSTRG.23300.1 ....................................................................................................
MSTRG.23300.2 ....................................................................................................
MSTRG.28072.1 CCTGAGTCAAAGTCTTCCCTATATTACTACTTTGTATTATAGCTGGGTTCACTATGTTGCATCATCTCACAGCTAACCCGAACATCACTAACATCGCAAT
MSTRG.28072.2 CCTGAGTCAAAGTCTTCCCTATATTACTACTTTGTATTATAGCTGGGTTCACTATGTTGCATCATCTCACAGCTAACCCGAACATCACTAACATCGCAAT
MSTRG.28072.3 CCTGAGTCAAAGTCTTCCCTATATTACTACTTTGTATTATAGCTGGGTTCACTATGTTGCATCATCTCACAGCTAACCCGAACATCACTAACATCGCAAT
MSTRG.28072.4 CCTGAGTCAAAGTCTTCCCTATATTACTACTTTGTATTATAGCTGGGTTCACTATGTTGCATCATCTCACAGCTAACCCGAACATCACTAACATCGCAAT

              -200......-190......-180......-170......-160......-150......-140......-130......-120......-110.........
Xa23 locus    ATTCTCCTTCCGCGCCCTTCACATCAGCTACTTCCACCACCTCCCATGTTTCAAAGGTAAATTCGTCGTGGTGGTGAGATTATGCCTTCCTTCCGCCTCA
MSTRG.23300.1 ......CTTCCGCGCCCTTCACATCAGCTACTTCCACCACCTCCCATGTTTCAAAGGTAAATTCGTCGTGGTGGTGAGATTATGCCTTCCTTCCGCCTCA
MSTRG.23300.2 ......CTTCCGCGCCCTTCACATCAGCTACTTCCACCACCTCCCATGTTTCAAAGGTAAATTCGTCGTGGTGGTGAGATTATGCCTTCCTTCCGCCTCA
MSTRG.28072.1 ATTCTCCTTCCGCGCCCTTCACATCAGCTACTTCCACCACCTCCCATGTTTC................................................
MSTRG.28072.2 ATTCTCCTTCCGCGCCCTTCACATCAGCTACTTCCACCACCTCCCATGTTTCAAAGGTAAATTCGTCGTGGTGGTGAGATTATGCCTTCCTTCCGCCTCA
MSTRG.28072.3 ATTCTCCTTCCGCGCCCTTCACATCAGCTACTTCCACCACCTCCCATGTTTCAAAGGTAAATTCGTCGTGGTGGTGAGATTATGCCTTCCTTCCGCCTCA
MSTRG.28072.4 ATTCTCCTTCCGCGCCCTTCACATCAGCTACTTCCACCACCTCCCATGTTTCAAAGGTAAATTCGTCGTGGTGGTGAGATTATGCCTTCCTTCCGCCTCA

              -100.......-90.......-80.......-70.......-60.......-50.......-40.......-30.......-20.......-10.......-1
Xa23 locus    CTAACATCAGCTACTATAAAAGCCCTTCCTTGTTGCATCATCTCAAGGAGCTGCAAGCACTTCCTCTCTGGCAGCACTTCCTCATCTCAAGGAGTTGCAA
MSTRG.23300.1 CTAACATCAGCTACTATAAAAGCCCTTCCTTGTTGCATCATCTCAAGGAGCTGCAAGCACTTCCTCTCTGGCAGCACTTCCTCATCTCAAGGAGTTGCAA
MSTRG.23300.2 CTAACATCAGCTACTATAAAAGCCCTTCCTTGTTGCATCATCTCAAGGAGCTGCAAGCACTTCCTCTCTGGCAGCACTTCCTCATCTCAAGGAGTTGCAA
MSTRG.28072.1 ..................AAAGCCCTTCCTTGTTGCATCATCTCAAGGAGCTGCAAGCACTTCCTCTCTGGCAGCACTTCCTCATCTCAAGGAGTTGCAA
MSTRG.28072.2 CTAACATCAGCTACTATAAAAGCCCTTCCTTGTTGCATCATCTCAAGGAGCTGCAAGCACTTCCTCTCTGGCAGCACTTCCTCATCTCAAGGAGTTGCAA
MSTRG.28072.3 CTAACATCAGCTACTATAAAAGCCCTTCCTTGTTGCATCATCTCAAGGAGCTGCAAGCACTTCCTCTCTGGCAGCACTTCCTCATCTCAAGGAGTTGCAA
MSTRG.28072.4 CTAACATCAGCTACTATAAAAGCCCTTCCTTGTTGCATCATCTCAAGGAGCTGCAAGCACTTCCTCTCTGGCAGCACTTCCTCATCTCAAGGAGTTGCAA

                +1......+10.......+20.......+30.......+40.......+50.......+60.......+70.......+80.......+90......+100
Xa23 locus    ATGTTGCATCATCTCAAGGAGCTGGCAGCCGTAGCCGGTATACACATGATCCTCATCTACCTCTGCCGCTTTCTCCTCCGCCGCAGCCGCAACGTATTAT
MSTRG.23300.1 ATGTTGCATCATCTCAAGGAGCTGGCAGCCGTAGCCGGTATACACATGATCCTCATCTACCTCTGCCGCTTTCTCCTCCGCCGCAGCCGCAACGTATTAT
MSTRG.23300.2 ATGTTGCATCATCTCAAGGAGCTGGCAGCCGTAGCCGGTATACACATGATCCTCATCTACCTCTGCCGCTTTCTCCTCCGCCGCAGCCGCAACGTATTAT
MSTRG.28072.1 ATGTTGCATCATCTCAAGGAGCTGGCAGCCGTAGCCGGTATACACATGATCCTCATCTACCTCTGCCGCTTTCTCCTCCGCCGCAGCCGCAACGTATTAT
MSTRG.28072.2 ATGTTGCATCATCTCAAGGAGCTGGCAGCCGTAGCCGGTATACACATGATCCTCATCTACCTCTGCCGCTTTCTCCTCCGCCGCAGCCGCAACGTATTAT
MSTRG.28072.3 ATGTTGCATCATCTCAAGGAGCTGGCAGCCGTAGCCGGTATACACATGATCCTCATCTACCTCTGCCGCTTTCTCCTCCGCCGCAGCCGCAACGTATTAT
MSTRG.28072.4 ATGTTGCATCATCTCAAGGAGCTGGCAGCCGTAGCCGGTATACACATGATCCTCATCTACCTCTGCCGCTTTCTCCTCCGCCGCAGCCGCAACGTATTAT

              ......+110......+120......+130......+140......+150......+160......+170......+180......+190......+200
Xa23 locus    TCACCGTTTCCAACAGCCTCCGTTTTCGCCTCAAGGTATTAACTGTATTGTTGTACATATGTCTCTCGGTCATGCTGTTCTACCTGTTTGGCTCCATCAT
MSTRG.23300.1 TCACCGTTTCCAACAGCCTCCGTTTTCGCCTCAAGGTATTAACTGTATTGTTGTACATATGTCTCTCGGTCATGCTGTTCTACCTGTTTGGCTCCATCAT
MSTRG.23300.2 TCACCGTTTCCAACAGCCTCCGTTTTCGCCTCAAG.................................................................
MSTRG.28072.1 TCACCGTTTCCAACAGCCTCCGTTTTCGCCTCAAGGTATTAACTGTATTGTTGTACATATGTCTCTCGGTCATGCTGTTCTACCTGTTTGGCTCCATCAT
MSTRG.28072.2 TCACCGTTTCCAACAGCCTCCGTTTTCGCCTCAAGGTATTAACTGTATTGTTGTACATATGTCTCTCGGTCATGCTGTTCTACCTGTTTGGCTCCATCAT
MSTRG.28072.3 TCACCGTTTCCAACAGCCTCCGTTTTCGCCTCAAGGTATTAACTGTATTGTTGTACATATGTCTCTCGGTCATGCTGTTCTACCTGTTTGGCTCCATCAT
MSTRG.28072.4 TCACCGTTTCCAACAGCCTCCGTTTTCGCCTCAAG.................................................................

              ......+210......+220......+230......+240......+250......+260......+270......+280......+290......+300
Xa23 locus    GCCGCTGCCGCCGTGGGGCCTCGTGGTCGGTTGGGTCATGGCCCTCATCGCCGTCGAGCTCGCCTACGCCTTCATCTTTCCATATAGCTTTCGCTACATC
MSTRG.23300.1 GCCGCTGCCGCCGTGGGGCCTCGTGGTCGGTTGGGTCATGGCCCTCATCGCCGTCGAGCTCGCCTACGCCTTCATCTTTCCATATAGCTTTCGCTACATC
MSTRG.23300.2 ....................................................................................................
MSTRG.28072.1 GCCGCTGCCGCCGTGGGGCCTCGTGGTCGGTTGGGTCATGGCCCTCATCGCCGTCGAGCTCGCCTACGCCTTCATCTTTCCATATAGCTTTCGCTACATC
MSTRG.28072.2 GCCGCTGCCGCCGTGGGGCCTCGTGGTCGGTTGGGTCATGGCCCTCATCGCCGTCGAGCTCGCCTACGCCTTCATCTTTCCATATAGCTTTCGCTACATC
MSTRG.28072.3 GCCGCTGCCGCCGTGGGGCCTCGTGGTCGGTTGGGTCATGGCCCTCATCGCCGTCGAGCTCGCCTACGCCTTCATCTTTCCATATAGCTTTCGCTACATC
MSTRG.28072.4 ....................................................................................................
```

```
             ......+310......+320......+330......+340......+350......+360......+370......+380......+390......+400
Xa23         GCTGACAACGACGACGACAAGATGGTTATTCTCCCTGTTTAAGCCTTCAGGGCCTATATATATAGTATATATATAAAGCCTTCCATACTGTCTCTTCAAT
MSTRG.23300.1 GCTGACAACGACGACGACAAGATGGTTATTCTCCCTGTTTAAGCCTTCAGGGCCTATATATATAGTATATATATAAAGCCTTCCATACTGTCTCTTCAAT
MSTRG.23300.2 ....................................................................................................
MSTRG.28072.1 GCTGACAACGACGACGACAAGATGGTTATTCTCCCTGTTTAAGCCTTCAGGGCCTATATATATAGTATATATATAAAGCCTTCCATACTGTCTCTTCAAT
MSTRG.28072.2 GCTGACAACGACGACGACAAGATGGTTATTCTCCCTGTTTAAGCCTTCAGGGCCTATATATATAGTATATATATAAAGCCTTCCATACTGTCTCTTCAAT
MSTRG.28072.3 GCTGACAACGACGACGACAAGATGGTTATTCTCCCTGTTTAAGCCTTCAGGGCCTATATATATAGTATATATATAAAGCCTTCCATACTGTCTCTTCAAT
MSTRG.28072.4 ....................................................................................................

             ......+410......+420......+430......+440......+450......+460......+470......+480......+490......+500
Xa23 locus    AAAGGCTAGCTTGTGTTGTGAGTTGTATCTGTGTACGTATTTTGTTTGGTTGGTTATATATTGTCACGTAGGTATGCCATATATATATGTATTGCTGTAT
MSTRG.23300.1 AAAGGCTAGCTTGTGTTGTGAGTTGTATCTGTGTACGTATTTTGTTTGGTTGGTTATATATTGTCACGTAGGTATGCCATATATATATGTATTGCTGTAT
MSTRG.23300.2 ....................................................................................................
MSTRG.28072.1 AAAGGCTAGCTTGTGTTGTGAGTTGTATCTGTGTACGTATTTTGTTTGGTTGGTTATATATTGTCACGTAGGTATGCCATATATATATGTATTGCTGTAT
MSTRG.28072.2 AAAGGCTAGCTTGTGTTGTGAGTTGTATCTGT....................................................................
MSTRG.28072.3 AAAGGCTAGCTTGTGTTGTGAGTTGTATCTGTGTACGTATTTTGTTTGGTTGGTTATATATTGTCACGTAGGTATGCCATATATATATGTATTGCTGTAT
MSTRG.28072.4 ....................................................................................................

             ......+510......+520......+530......+540......+550......+560......+570......+580......+590......+600
Xa23 locus    TTATATTTGTTACTATCTTTTGTTTTTCAGATAATAAAATTCAGCCAGCTTTGCTTGCTTCGTCGTACGTGTATGCTCATCATATCCTCATCCATCAGCT
MSTRG.23300.1 TTATATTTGTTACTATCTTTTGTTTTTCAGATAATAAAATTCAGCCAGCTTTGCTTGCTTCGTCGTACGTGTATGCTCATCATATCCTCATCCATCAGCT
MSTRG.23300.2 ..............................ATAATAAAATTCAGCCAGCTTTGCTTGCTTCGTCGTACGTGTATGCTCATCATATCCTCATCCATCAGCT
MSTRG.28072.1 TTATATTTGTTACTATCTTTTGTTTTTCAGATAATAAAATTCAGCCAGCTTTGCTTGCTTCGTCGTACGTGTATGCTCATCATATCCTCATCCATCAGCT
MSTRG.28072.2 ..............................ATAATAAAATTCAGCCAGCTTTGCTTGCTTCGTCGTACGTGTATGCTCATCATATCCTCATCCATCAGCT
MSTRG.28072.3 TTATATTTGTTACTATCTTTTGTTTTTCAGATAATAAAATTCAGCCAGCTTTGCTTGCTTCGTCGTACGTGTATGCTCATCATATCCTCATCCATCAGCT
MSTRG.28072.4 ..............................ATAATAAAATTCAGCCAGCTTTGCTTGCTTCGTCGTACGTGTATGCTCATCATATCCTCATCCATCAGCT

             ......+610......+620......+630......+640......+650......+660......+670......+680......+690......+700
Xa23 locus    GCTCATAGCTAGCTGGGCCGTGTTATATGTGTGTAGTGACCAGTCACAACCATGTATTTCATCCCATGTATGTTAGTTTGTTTTTCATTTTTGAAAGAAA
MSTRG.23300.1 GCTCATAGCTAGCTGGGCCGTGTTATATGTGTGTAGTGACCAGTCACAACCATGTATTTCATCCCATGTATGTTAGTTTGTTTTTCATTTTTGAAAGAAA
MSTRG.23300.2 GCTCATAGCTAGCTGGGCCGTGTTATATGTGTGTAGTGACCAGTCAC.....................................................
MSTRG.28072.1 GCTCATAGCTAGCTGGGCCGTGTTATATGTGTGTAGTGACCAGTCACAACCATGTATTTCATCCCATGTATGTTAGTTTGTTTTTCATTTTTGAAAGAAA
MSTRG.28072.2 GCTCATAGCTAGCTGGGCCGTGTTATATGTGTGTAGTGACCAGTCACAACCATGTATTTCATCCCATGTATGTTAGTTTGTTTTTCATTTTTGAAAGAAA
MSTRG.28072.3 GCTCATAGCTAGCTGGGCCGTGTTATATGTGTGTAGTGACCAGTCACAACCATGTATTTCATCCCATGTATGTTAGTTTGTTTTTCATTTTTGAAAGAAA
MSTRG.28072.4 GCTCATAGCTAGCTGGGCCGTGTTATATGTGTGTAGTGACCAGTCACAACCATGTATTTCATCCCATGTATGTTAGTTTGTTTTTCATTTTTGAAAGAAA

             ......+710......+720......+730......+740......+750......+760......+770......+780......+790......+800
Xa23 locus    ACATGCATATATGTTATAACTCAAAGTTTTGAGATGAATTTGATCATCTAATTTATTTCTGTTTTGAATTATTCTGCTCTGTTGTTGCAAACTTTTTTTT
MSTRG.23300.1 ACATGCATATATGTTATAACTCAAAGTTTTGAGATGAATTTGATCAT.....................................................
MSTRG.23300.2 ....................................................................................................
MSTRG.28072.1 ACATGCATATATGTTATAACTCAAAGT.........................................................................
MSTRG.28072.2 ACATGCATATATGTTATAACT...............................................................................
MSTRG.28072.3 ACATGCATATATGTTATAAC................................................................................
MSTRG.28072.4 ACATGCATATATGTTATAAC................................................................................

             ......+810......+820......+830......+840..
Xa23 locus    TTATGTTGGGACACTTTCTCCTCGGTTTGATTGTTATATTGA
MSTRG.23300.1 ..........................................
MSTRG.23300.2 ..........................................
MSTRG.28072.1 ..........................................
MSTRG.28072.2 ..........................................
MSTRG.28072.3 ..........................................
MSTRG.28072.4 ..........................................
```

**Data S12. Alignment of the genomic *Xa23* sequences from two cultivated rice accessions (Nipponbare and CBB23) and 5'RACE-derived parts of the TSA-induced *Xa23* transcripts**. The alignment was constructed using the MUSCLE algorithm.[11] Binding sites of the primers KS535, KS536, KS537, KS538, and KS539 that were used for nested 5'RACE PCRs are annotated and underlined. The binding elements for the dTALE and AvrXa23 activating *Xa23* transcription are annotated and highlighted in magenta.[13,14] dTALE and AvrXa23 bind to different elements. The published 5'UTR, CDS, and 3'UTR of the dTALE-induced *Xa23* mRNA transcripts are highlighted in blue, yellow, and grey, respectively.[13] The *Xa23* start (**ATG**) and stop (**TAA**) codons are colored in **red**. TATA boxes and TATA box-like sequences detected upstream of the *Xa23* start codon are colored in **green**. Additional potential CREs within the 5' upstream sequences identified using PlantCARE[15] (GCN4 motifs, G boxes, and **CARE motifs**) are highlighted and annotated in the alignment.

```
Xa23Nipponbare locus  .................................AGCAGTCCCCCCTTCCCTTCCGCTATT
KS535+KS536_Col1      ............................................................
KS535+KS536_Col3      ............................................................
KS535+KS536_Col6      ............................................................
KS535+KS536_Col7      ............................................................
KS535+KS536_Col9      ............................................................
KS535+KS536_Col10     ............................................................
KS536+KS537_Col13     ............................................................
KS536+KS537_Col15     ............................................................
KS536+KS537_Col17     ............................................................
KS536+KS537_Col20     ............................................................
KS536+KS537_Col21     ............................................................
KS536+KS537_Col23     ............................................................
KS538+KS539_Col5      ............................................................
KS538+KS539_Col8      ............................................................
KS538+KS539_Col113    ............................................................
KS538+KS539_Col114    ............................................................
KS538+KS539_Col115    ............................................................
Xa23CBB23 locus       CAACGTCAGCGCCAATGTGTTTGGCGCTGAGGCGACGGCCTATTTTTGGTTGAAGTTTTT

                                 dTALEXa23Ni-1                           GCN4 motif
                      ........70........80........90.......100.......110.......120
Xa23Nipponbare locus  TCAAAGTATCCGTCTTCCTCCCTGTCCACATGAGCTAAACGCTTCCTCTTCCCTGAGTCA
KS535+KS536_Col1      ............................................................
KS535+KS536_Col3      ............................................................
KS535+KS536_Col6      ............................................................
KS535+KS536_Col7      ............................................................
KS535+KS536_Col9      ............................................................
KS535+KS536_Col10     ............................................................
KS536+KS537_Col13     ............................................................
KS536+KS537_Col15     ............................................................
KS536+KS537_Col17     ............................................................
KS536+KS537_Col20     ............................................................
KS536+KS537_Col21     ............................................................
KS536+KS537_Col23     ............................................................
KS538+KS539_Col5      ............................................................
KS538+KS539_Col8      ............................................................
KS538+KS539_Col113    ............................................................
KS538+KS539_Col114    ............................................................
KS538+KS539_Col115    ............................................................
Xa23CBB23 locus       GGCAGGGGTTAGTTTCGAAATAAGTTTTCCAAAAGGGTCAATTTGTCAAAAAAACGGCTC
```

```
                            TATA box-like1  TATA box-like2
                        .......130.......140.......150.......160.......170.......180
Xa23^Nipponbare locus   AAGTCTTCCCTATATTACTACTTTGTATTATAGCTGGGTTCACTATGTTGCATCATCTCA
KS535+KS536_Col1        ............................................................
KS535+KS536_Col3        ............................................................
KS535+KS536_Col6        ............................................................
KS535+KS536_Col7        ............................................................
KS535+KS536_Col9        ............................................................
KS535+KS536_Col10       ............................................................
KS536+KS537_Col13       ............................................................
KS536+KS537_Col15       ............................................................
KS536+KS537_Col17       ............................................................
KS536+KS537_Col20       ............................................................
KS536+KS537_Col21       ............................................................
KS536+KS537_Col23       ............................................................
KS538+KS539_Col5        ............................................................
KS538+KS539_Col8        ............................................................
KS538+KS539_Col113      ............................................................
KS538+KS539_Col114      ............................................................
KS538+KS539_Col115      ............................................................
Xa23^CBB23 locus        GTCCCTGAGTCAAAGTCTTCCCTATTAAATTATGCGGCATCACTAACATCAGCTACTATA
                           GCN4 motif        TATA box-like3                       TATA box

                        .......190.......200.......210.......220.......230.......240
Xa23^Nipponbare locus   CAGCTAACCCGAACATCACTAACAT......................CGCAATATTCTCC
KS535+KS536_Col1        ............................................................
KS535+KS536_Col3        ............................................................
KS535+KS536_Col6        ............................................................
KS535+KS536_Col7        ............................................................
KS535+KS536_Col9        ............................................................
KS535+KS536_Col10       ............................................................
KS536+KS537_Col13       ............................................................
KS536+KS537_Col15       ............................................................
KS536+KS537_Col17       ............................................................
KS536+KS537_Col20       ............................................................
KS536+KS537_Col21       ............................................................
KS536+KS537_Col23       ............................................................
KS538+KS539_Col5        ....................AACAT......................CGCAATATTCTCC
KS538+KS539_Col8        ....................AACAT......................CGCAATATTCTCC
KS538+KS539_Col113      ....................AACAT......................CGCAATATTCTCC
KS538+KS539_Col114      ....................AACAT......................CGCAATATTCTCC
KS538+KS539_Col115      ....................AACAT......................CGCAATATTCTCC
Xa23^CBB23 locus        AAAGTCCCTTCCGCGTCACTAACATCAGCTACTATAAAAGTCCCTTCCGAAACATCTTCC
                      TATA box                           TATA box        AvrXa23

                        .......250.......260.......270.......280.......290.......300
Xa23^Nipponbare locus   TTCCGCGCCCTTCACATCAGCTACT.TCCACCACCTCCCATGTTTCAAAGGTAAATTCGT
KS535+KS536_Col1        ............................................................
KS535+KS536_Col3        ............................................................
KS535+KS536_Col6        ............................................................
KS535+KS536_Col7        ............................................................
KS535+KS536_Col9        ............................................................
KS535+KS536_Col10       ............................................................
KS536+KS537_Col13       ............................................................
KS536+KS537_Col15       ............................................................
KS536+KS537_Col17       ............................................................
KS536+KS537_Col20       ............................................................
KS536+KS537_Col21       ............................................................
KS536+KS537_Col23       ............................................................
KS538+KS539_Col5        TTCCGCGCCCTTCACATCAGCTACT.TCCACCACCTCCCATGTTTCA.............
KS538+KS539_Col8        TTCCGCGCCCTTCACATCAGCTACT.TCCACCACCTCCCATGTTTCA.............
KS538+KS539_Col113      TTCCGCGCCCTTCACATCAGCTACT.TCCACCACCTCCCATGTTTCA.............
KS538+KS539_Col114      TTCCGCGCCCTTCACATCAGCTACT.TCCACCACCTCCCATGTTTCA.............
KS538+KS539_Col115      TTCCGCGCCCTTCACATCAGCTACT.TCCACCACCTCCCATGTTTCA.............
Xa23^CBB23 locus        TCCCGCATCACTAACATCAGCTTCTATAAAAGCCCTTCCTTGTTGCA.............
                           AvrXa23            TATA box
```

```
                                                                   TATA box
                         .......310.......320.......330.......340.......350.......360
Xa23Nipponbare locus     CGTGGTGGTGAGATTATGCCTTCCTTCCGCCTCACTAACATCAGCTACTATAAAAGCCCT
KS535+KS536_Col1         ............................................................
KS535+KS536_Col3         ............................................................
KS535+KS536_Col6         ............................................................
KS535+KS536_Col7         ............................................................
KS535+KS536_Col9         ............................................................
KS535+KS536_Col10        ............................................................
KS536+KS537_Col13        ............................................................
KS536+KS537_Col15        ............................................................
KS536+KS537_Col17        ............................................................
KS536+KS537_Col20        ............................................................
KS536+KS537_Col21        ............................................................
KS536+KS537_Col23        ............................................................
KS538+KS539_Col5         ............................................................
KS538+KS539_Col8         ............................................................
KS538+KS539_Col113       ............................................................
KS538+KS539_Col114       ............................................................
KS538+KS539_Col115       ............................................................
Xa23CBB23 locus          ............................................................

                                                                   G box
                         .......370.......380.......390.......400.......410.......420
Xa23Nipponbare locus     TCCTTGTTGCATCATCTCAAGGAGCTGCAAGCACTTCCTCTCTGGCAGCACTTCCTCATC
KS535+KS536_Col1         ............................AAGCACTTCCTCTCTGGCAGCACTTCCTCATC
KS535+KS536_Col3         ............................AAGCACTTCCTCTCTGGCAGCACTTCCTCATC
KS535+KS536_Col6         ............................AAGCACTTCCTCTCTGGCAGCACTTCCTCATC
KS535+KS536_Col7         ............................AAGCACTTCCTCTCTGGCAGCACTTCCTCATC
KS535+KS536_Col9         ............................AAGCACTTCCTCTCTGGCAGCACTTCCTCATC
KS535+KS536_Col10        ............................AAGCACTTCCTCTCTGGCAGCACTTCCTCATC
KS536+KS537_Col13        ..................AAGGAGCTGCAAGCACTTCCTCTCTGGCAGCACTTCCTCATC
KS536+KS537_Col15        ..................AAGGAGCTGCAAGCACTTCCTCTCTGGTAGCACTTCCTCATC
KS536+KS537_Col17        ..................AAGGAGCTGCAAGCACTTCCTCTCTGGCAGCACTTCCTCATC
KS536+KS537_Col20        .......................................CTCTGGCAGCACTTCCTCATC
KS536+KS537_Col21        ........................CTGCAAGCACTTCCTCTCTGGCAGCACTTCCTCATC
KS536+KS537_Col23        ........................CTGCAAGCACTTCCTCTCTGGCAGCACTTCCTCATC
KS538+KS539_Col5         ..................AAGGAGCTGCAAGCACTTCCTCTCTGGCAGCACTTCCTCATC
KS538+KS539_Col8         ..................AAGGAGCTGCAAGCACTTCCTCTCTGGCAGCACTTCCTCATC
KS538+KS539_Col113       ..................AAGGAGCTGCAAGCACTTCCTCTCTGGCAGCACTTCCTCATC
KS538+KS539_Col114       ..................AAGGAGCTGCAAGCACTTCCTCTCTGGCAGCACTTCCTCATC
KS538+KS539_Col115       ..................AAGGAGCTGCAAGCACTTCCTCTCTGGCAGCACTTCCTCATC
Xa23CBB23 locus          ...........TCATCTCAAGGAGCTGCAAGCACTTCCTCTCTGGCAGCACTTCCTCATC
                                                                   G box

                                                                   primer KS539
                             CARE motif                         <------------------
                         .......430.......440.......450.......460.......470.......480
Xa23Nipponbare locus     TCAAGGAGTTGCAAATGTTGCATCATCTCAAGGAGCTGGCAGCCGTAGCCGGTATACACA
KS535+KS536_Col1         TCAAGGAGTTGCAAATGTTGCATCATCTCAAGGAGCTGGCAGCCGTAGCCGGTATACACA
KS535+KS536_Col3         TCAAGGAGTTGCAAATGTTGCATCATCTCAAGGAGCTGGCAGCCGTAGCCGGTATACACA
KS535+KS536_Col6         TCAAGGAGTTGCAAATGTTGCATCATCTCAAGGAGCTGGCAGCCGTAGCCGGTATACACA
KS535+KS536_Col7         TCAAGGAGTTGCAAATGTTGCATCATCTCAAGGAGCTGGCAGCCGTAGCCGGTATACACA
KS535+KS536_Col9         TCAAGGAGTTGCAAATGTTGCATCATCTCAAGGAGCTGGCAGCCGTAGCCGGTATACACA
KS535+KS536_Col10        TCAAGGAGTTGCAAATGTTGCATCATCTCAAGGAGCTGGCAGCCGTAGCCGGTATACACA
KS536+KS537_Col13        TCAAGGAGTTGCAAATGTTGCATCATCTCAAGGAGCTGGCAGCCGTAGCCGGTATACACA
KS536+KS537_Col15        TCAAGGAGTTGCAAATGTTGCATCATCTCAAGGAGCTGGCAGCCGTAGCCGGTATACACA
KS536+KS537_Col17        TCAAGGAGTTGCAAATGTTGCATCATCTCAAGGAGCTGGCAGCCGTAGCCGGTATACACA
KS536+KS537_Col20        TCAAGGAGTTGCAAATGTTGCATCATCTCAAGGAGCTGGCAGCCGTAGCCGGTATACACA
KS536+KS537_Col21        TCAAGGAGTTGCAAATGTTGCATCATCTCAAGGAGCTGGCAGCCGTAGCCGGTATACACA
KS536+KS537_Col23        TCAAGGAGTTGCAAATGTTGCATCATCTCAAGGAGCTGGCAGCCGTAGCCGGTATACACA
KS538+KS539_Col5         TCAAGGAGTTGCAAATGTTGCATCATCTCAAGGAGCTGGCAGCCGTAGCCGGTATACACA
KS538+KS539_Col8         TCAAGGAGTTGCAAATGTTGCATCATCTCAAGAAGCTGGCAGCCGTAGCCGGTATACACA
KS538+KS539_Col113       TCAAGGAGTTGCAAATGTTGCATCATCTCAAGGAGCTGGCAGCCGTAGCCGGTATACACA
KS538+KS539_Col114       TCAAGGAGTTGCAAATGTTGCATCATCTCAAGGAGCTGGCAGCCGTAGCCGGTATACACA
KS538+KS539_Col115       TCAAGGAGTTGCAAATGTTGCATCATCTCAAGGAGCTGGCAGCCGTAGCCGGTATACACA
Xa23CBB23 locus          TCAAGGAGTTGCAAATGTTGCATCATCTCAAGGAGCTGGCAGCCGTAGCCGGTATACACA
                             CARE motif
```

```
                          primer KS539
                          ------------
                          .......490.......500.......510.......520.......530.......540
Xa23^Nipponbare locus     TGATCCTCATCTACCTCTGCCGCTTTCTCCTCCGCCGCAGCCGCAACGTATTATTCACCG
KS535+KS536_Col1          TGATCCTCATCTACCTCTGCCGCTTTCTCCTCCGCCGCAGCCGCAACGTATTATTCACCG
KS535+KS536_Col3          TGATCCTCATCTACCTCTGCCGCTTTCTCCTCCGCCGCAGCCGCAACGTATTATTCACCG
KS535+KS536_Col6          TGATCCTCATCTACCTCTGCCGCTTTCTCCTCCGCCGCAGCCGCAACGTATTATTCACCG
KS535+KS536_Col7          TGATCCTCATCTACCTCTGCCGCTTTCTCCTCCGCCGCAGCCGCAACGTATTATTCACCG
KS535+KS536_Col9          TGATCCTCATCTACCTCTGCCGCTTTCTCCTCCGCCGCAGCCGCAACGTATTATTCACCG
KS535+KS536_Col10         TGATCCTCATCTACCTCTGCCGCTTTCTCCTCCGCCGCAGCCGCAACGTATTATTCACCG
KS536+KS537_Col13         TGATCCTCATCTACCTCTGCCGCTTTCTCCTCCGCCGCAGCCGCAACGTATTATTCACCG
KS536+KS537_Col15         TGATCCTCATCTACCTCTGCCGCTTTCTCCTCCGCCGCAGCCGCAACGTATTATTCACCG
KS536+KS537_Col17         TGATCCTCATCTACCTCTGCCGCTTTCTCCTCCGCCGCAGCCGCAACGTATTATTCACCG
KS536+KS537_Col20         TGATCCTCATCTACCTCTGCCGCTTTCTCCTCCGCCGCAGCCGCAACGTATTATTCACCG
KS536+KS537_Col21         TGATCCTCATCTACCTCTGCCGCTTTCTCCTCCGCCGCAGCCGCAACGTATTATTCACCG
KS536+KS537_Col23         TGATCCTCATCTACCCCTGCCGCTTTCTCCTCCGCCGCAGCCGCAACGTATTATTCACCG
KS538+KS539_Col5          TGATCCT.....................................................
KS538+KS539_Col8          TGATCCTCATC.................................................
KS538+KS539_Col113        TGATCCTCATC.................................................
KS538+KS539_Col114        TGATCCTCATC.................................................
KS538+KS539_Col115        TGATCCTCATC.................................................
Xa23^CBB23 locus          TGATCCTCATCTACCTCTGCCGCTTTCTCCTCCGCCGCAGCCGCAACGTATTATTCACCG

                          .......550.......560.......570.......580.......590.......600
Xa23^Nipponbare locus     TTTCCAACAGCCTCCGTTTTCGCCTCAAGGTATTAACTGTATTGTTGTACATATGTCTCT
KS535+KS536_Col1          TTTCCAACAGCCTCCGTTTTCGCCTCAAGGTATTAACTGTATTGTTGTACATATGTCTCT
KS535+KS536_Col3          TTTCCAACAGCCTCCGTTTTCGCCTCAAGGTATTAACTGTATTGTTGTACATATGTCTCT
KS535+KS536_Col6          TTTCCAACAGCCTCCGTTTTCGCCTCAAGGTATTAACTGTATTGTTGTACATATGTCTCT
KS535+KS536_Col7          TTTCCAACAGCCTCCGTTTTCGCCTCAAGGTATTAACTGTATTGTTGTACATATGTCTCT
KS535+KS536_Col9          TTTCCAACAGCCTCCGTTTTCGCCTCAAGGTATTAACTGTATTGTTGTACATATGTCTCT
KS535+KS536_Col10         TTTCCAACAGCCTCCGTTTTCGCCTCAAGGTATTAACTGTATTGTTGTACATATGTCTCT
KS536+KS537_Col13         TTTCCAACAGCCTCCGTTTTCGCCTCAAGGTATTAACTGTATTGTTGTACATATGTCTCT
KS536+KS537_Col15         TTTCCAACAGCCTCCGTTTTCGCCTCAAGGTATTAACTGTATTGTTGTACATATGTCTCT
KS536+KS537_Col17         TTTCCAACAGCCTCCGTTTTCGCCTCAAGGTATTAACTGTATTGTTGTACATATGTCTCT
KS536+KS537_Col20         TTTCCAACAGCCTCCGTTTTCGCCTCAAGGTATTAACTGTATTGTTGTACATATGTCTCT
KS536+KS537_Col21         TTTCCAACAGCCTCCGTTTTCGCCTCAAGGTATTAACTGTATTGTTGTACATATGTCTCT
KS536+KS537_Col23         TTTCCAACAGCCTCCGTTTTCGCCTCAAGGTATTAACTGTATTGTTGTACATATGTCTCT
KS538+KS539_Col5          ............................................................
KS538+KS539_Col8          ............................................................
KS538+KS539_Col113        ............................................................
KS538+KS539_Col114        ............................................................
KS538+KS539_Col115        ............................................................
Xa23^CBB23 locus          TTTCCAACAGCCTCCGTTTTCGCCTCAAGGTATTAACTGTATTGTTGTACATATGTCTCT

                          .......610.......620.......630.......640.......650.......660
Xa23^Nipponbare locus     CGGTCATGCTGTTCTACCTGTTTGGCTCCATCATGCCGCTGCCGCCGTGGGGCCTCGTGG
KS535+KS536_Col1          CGGTCATGCTGTTCTACCTGTTTGGCTCCATCATGCCGCTGCCGCCGTGGGGCCTCGTGG
KS535+KS536_Col3          CGGTCATGCTGTTCTACCTGTTTGGCTCCATCATGCCGCTGCCGCCGTGGGGCCTCGTGG
KS535+KS536_Col6          CGGTCATGCTGTTCTACCTGTTTGGCTCCATCATGCCGCTGCCGCCGTGGGGCCTCGTGG
KS535+KS536_Col7          CGGTCATGCTGTTCTACCTGTTTGGCTCCATCATGCCGCTGCCGCCGTGGGGCCTCGTGG
KS535+KS536_Col9          CGGTCATGCTGTTCTACCTGTTTGGCTCCATCATGCCGCTGCCGCCGTGGGGCCTCGTGG
KS535+KS536_Col10         CGGTCATGCTGTTCTACCTGTTTGGCTCCATCATGCCGCTGCCGCCGTGGGGCCTCGTGG
KS536+KS537_Col13         CGGTCATGCTGTTCTACCTGTTTGGCTCCATCATGCCGCTGCCGCCGTGGGGCCTCGTGG
KS536+KS537_Col15         CGGTCATGCTGTTCTACCTGTTTGGCTCCATCATGCCGCTGCCGCCGTGGGGCCTCGTGG
KS536+KS537_Col17         CGGTCATGCTGTTCTACCTGTTTGGCTCCATCATGCCGCTGCCGCCGTGGGGCCTCGTGG
KS536+KS537_Col20         CGGTCATGCTGTTCTACCTGTTTGGCTCCATCATGCCGCTGCCGCCGTGGGGCCTCGTGG
KS536+KS537_Col21         CGGTCATGCTGTTCTACCTGTTTGGCTCCATCATGCCGCTGCCGCCGTGGGGCCTCGTGG
KS536+KS537_Col23         CGGTCATGCTGTTCTACCTGTTTGGCTCCATCATGCCGCTGCCGCCGTGGGGCCTCGTGG
KS538+KS539_Col5          ............................................................
KS538+KS539_Col8          ............................................................
KS538+KS539_Col113        ............................................................
KS538+KS539_Col114        ............................................................
KS538+KS539_Col115        ............................................................
Xa23^CBB23 locus          CGGTCATGCTGTTCTACCTGTTTGGCTCCATCATGCCGCTGCCGCCGTGGGGCCTCGTGG
```

```
                                                        primer KS538
                                              ←----------------------------
                      .......670.......680.......690.......700.......710.......720
Xa23^Nipponbare locus  TCGGTTGGGTCATGGCCCTCATCGCCGTCGAGCTCGCCTACGCCTTCATCTTTCCATATA
KS535+KS536_Col1       TCGGTTGGGTCATGGCCCTCATCGCCGTCGAGCTCGCCTACGCCTTCATCTTTCCATATA
KS535+KS536_Col3       TCGGTTGGGTCATGGCCCTCATCGCCGTCGAGCTCGCCTACGCCTTCATCTTTCCATATA
KS535+KS536_Col6       TCGGTTGGGTCATGGCCCTCATCGCCGTCGAGCTCGCCTACGCCTTCATCTTTCCATATA
KS535+KS536_Col7       TCGGTTGGGTCATGGCCCTCATCGCCGTCGAGCTCGCCTACGCCTTCATCTTTCCATATA
KS535+KS536_Col9       TCGGTTGGGTCATGGCCCTCATCGCCGTCGAGCTCGCCTACGCCTTCATCTTTCCATATA
KS535+KS536_Col10      TCGGTTGGGTCATGGCCCTCATCGCCGTCGAGCTCGCCTACGCCTTCATCTTTCCATATA
KS536+KS537_Col13      TCGGTTGGGTCATGGCCCTCATCGCCGTCGAGCTCGCCTACGCCTTCATCTTTCCATATA
KS536+KS537_Col15      TCGGTTGGGTCATGGCCCTCATCGCCGTCGAGCTCGCCTACGCCTTCATCTTTCCATATA
KS536+KS537_Col17      TCGGTTGGGTCATGGCCCTCATCGCCGTCGAGCTCGCCTACGCCTTCATCTTTCCATATA
KS536+KS537_Col20      TCGGTTGGGTCATGGCCCTCATCGCCGTCGAGCTCGCCTACGCCTTCATCTTTCCATATA
KS536+KS537_Col21      TCGGTTGGGTCATGGCCCTCATCGCCGTCGAGCTCGCCTACGCCTTCATCTTTCCATATA
KS536+KS537_Col23      TCGGTTGGGTCATGGCCCTCATCGCCGTCGAGCTCGCCTACGCCTTCATCTTTCCATATA
KS538+KS539_Col5       ............................................................
KS538+KS539_Col8       ............................................................
KS538+KS539_Col113     ............................................................
KS538+KS539_Col114     ............................................................
KS538+KS539_Col115     ............................................................
Xa23^CBB23 locus       TCGGTTGGGTCATGGCCCTCATCGCCGTCGAGCTCGCCTACGCCTTCATCTTTCCATATA

                                 primer KS537                      primer KS536
                       ←----------------------------          ←----------------
                      .......730.......740.......750.......760.......770.......780
Xa23^Nipponbare locus  GCTTTCGCTACATCGCTGACAACGACGACGACAAGATGGTTATTCTCCCTGTTTAAGCCT
KS535+KS536_Col1       GCTTTCGCTACATCGCTGACAACGACGACGACAAGATGGTTATTCTCCCTGTTTAAGCCT
KS535+KS536_Col3       GCTTTCGCTACATCGCTGACAACGACGACGACAAGATGGTTATTCTCCCTGTTTAAGCCT
KS535+KS536_Col6       GCTTTCGCTACATCGCTGACAACGACGACGACAAGATGGTTATTCTCCCTGTTTAAGCCT
KS535+KS536_Col7       GCTTTCGCTACATCGCTGACAACGACGACGACAAGATGGTTATTCTCCCTGTTTAAGCCT
KS535+KS536_Col9       GCTTTCGCTACATCGCTGACAACGACGACGACAAGATGGTTATTCTCCCTGTTTAAGCCT
KS535+KS536_Col10      GCTTTCGCTGCATCGCTGACAACGACGACGACAAGATGGTTATTCTCCCTGTTTAAGCCT
KS536+KS537_Col13      GCTTTCGCTACATCGCTGACAACGACG.................................
KS536+KS537_Col15      GCTTTCGCTACATCGCTGACAACGACG.................................
KS536+KS537_Col17      GCTTTCGCTACATCGCTGACAACGACG.................................
KS536+KS537_Col20      GCTTTCGCTACATCGCTGACAACGACG.................................
KS536+KS537_Col21      GCTTTCGCTACATCGCTGACAACGACG.................................
KS536+KS537_Col23      GCTTTCGCTACATCGCTGACAACGACG.................................
KS538+KS539_Col5       ............................................................
KS538+KS539_Col8       ............................................................
KS538+KS539_Col113     ............................................................
KS538+KS539_Col114     ............................................................
KS538+KS539_Col115     ............................................................
Xa23^CBB23 locus       GCTTTCGCTACATCGCTGACAACGACGACGACAAGATGGTTATTCTCCCTGTTTAAGCCT

                     primer KS536                                         primer KS535
                       ----------                                          ←--
                      .......790.......800.......810.......820.......830.......840
Xa23^Nipponbare locus  TCAGGGCCTATATATATAGTATATATATAAAGCCTTCCATACTGTCTCTTCAATAAAGGC
KS535+KS536_Col1       TCAGGGCCT...................................................
KS535+KS536_Col3       TCAGGGCCT...................................................
KS535+KS536_Col6       TCAGGGCCT...................................................
KS535+KS536_Col7       TCAGGGCCT...................................................
KS535+KS536_Col9       TCAGGGCCT...................................................
KS535+KS536_Col10      TCAGGGCCT...................................................
KS536+KS537_Col13      ............................................................
KS536+KS537_Col15      ............................................................
KS536+KS537_Col17      ............................................................
KS536+KS537_Col20      ............................................................
KS536+KS537_Col21      ............................................................
KS536+KS537_Col23      ............................................................
KS538+KS539_Col5       ............................................................
KS538+KS539_Col8       ............................................................
KS538+KS539_Col113     ............................................................
KS538+KS539_Col114     ............................................................
KS538+KS539_Col115     ............................................................
Xa23^CBB23 locus       TCAGGGCCTATATATATAGTATATATATAAAGCCTTCCATACTGTCTCTTCAATAAAGGC
```

```
                            primer KS535
                          -----------------
                          .......850.......860.......870.......880.......890.......900
Xa23^Nipponbare locus     TAGCTTGTGTTGTGAGTTGTATCTGTGTACGTATTTTGTTTGGTTGGTTATATATTGTCA
KS535+KS536_Col1          ............................................................
KS535+KS536_Col3          ............................................................
KS535+KS536_Col6          ............................................................
KS535+KS536_Col7          ............................................................
KS535+KS536_Col9          ............................................................
KS535+KS536_Col10         ............................................................
KS536+KS537_Col13         ............................................................
KS536+KS537_Col15         ............................................................
KS536+KS537_Col17         ............................................................
KS536+KS537_Col20         ............................................................
KS536+KS537_Col21         ............................................................
KS536+KS537_Col23         ............................................................
KS538+KS539_Col5          ............................................................
KS538+KS539_Col8          ............................................................
KS538+KS539_Col113        ............................................................
KS538+KS539_Col114        ............................................................
KS538+KS539_Col115        ............................................................
Xa23^CBB23 locus          TAGCTTGTGTTGTGAGTTGTATCTGTGTACGTATTTTGTTTGGTTGGTTATATATTGTCA

                          .......910.......920.......930.......940.......950.......960
Xa23^Nipponbare locus     CGTAGGTATGCCATATATATATGTATTGCTGTATTTATATTTGTTACTATCTTTTGTTTT
KS535+KS536_Col1          ............................................................
KS535+KS536_Col3          ............................................................
KS535+KS536_Col6          ............................................................
KS535+KS536_Col7          ............................................................
KS535+KS536_Col9          ............................................................
KS535+KS536_Col10         ............................................................
KS536+KS537_Col13         ............................................................
KS536+KS537_Col15         ............................................................
KS536+KS537_Col17         ............................................................
KS536+KS537_Col20         ............................................................
KS536+KS537_Col21         ............................................................
KS536+KS537_Col23         ............................................................
KS538+KS539_Col5          ............................................................
KS538+KS539_Col8          ............................................................
KS538+KS539_Col113        ............................................................
KS538+KS539_Col114        ............................................................
KS538+KS539_Col115        ............................................................
Xa23^CBB23 locus          CGTAGGTATGCCATATATATATGTATTGCTGTATTTATATTTGTTACTATCTTTTGTTTT

                          .......970.......980.......990......1000......1010......1020
Xa23^Nipponbare locus     TCAGATAATAAAATTCAGCCAGCTTTGCTTGCTTCGTCGTACGTGTATGCTCATCATATC
KS535+KS536_Col1          ............................................................
KS535+KS536_Col3          ............................................................
KS535+KS536_Col6          ............................................................
KS535+KS536_Col7          ............................................................
KS535+KS536_Col9          ............................................................
KS535+KS536_Col10         ............................................................
KS536+KS537_Col13         ............................................................
KS536+KS537_Col15         ............................................................
KS536+KS537_Col17         ............................................................
KS536+KS537_Col20         ............................................................
KS536+KS537_Col21         ............................................................
KS536+KS537_Col23         ............................................................
KS538+KS539_Col5          ............................................................
KS538+KS539_Col8          ............................................................
KS538+KS539_Col113        ............................................................
KS538+KS539_Col114        ............................................................
KS538+KS539_Col115        ............................................................
Xa23^CBB23 locus          TCAGATAATAAAATTCAGCCAGCTTTGCTTGCTTCGTCGTACGTGTATGCTCATCATATC
```

......1030......1040......1050......
*Xa23*$^{Nipponbare}$ locus CTCATCCATCAGCTGCTCATAGCTAGCTGGGCCGTG
KS535+KS536_Col1 ....................................
KS535+KS536_Col3 ....................................
KS535+KS536_Col6 ....................................
KS535+KS536_Col7 ....................................
KS535+KS536_Col9 ....................................
KS535+KS536_Col10 ....................................
KS536+KS537_Col13 ....................................
KS536+KS537_Col15 ....................................
KS536+KS537_Col17 ....................................
KS536+KS537_Col20 ....................................
KS536+KS537_Col21 ....................................
KS536+KS537_Col23 ....................................
KS538+KS539_Col5 ....................................
KS538+KS539_Col8 ....................................
KS538+KS539_Col113 ....................................
KS538+KS539_Col114 ....................................
KS538+KS539_Col115 ....................................
*Xa23*$^{CBB23}$ locus CTCATCCATCAGCTGCTCATAGCTAGCTGGGCCGTG

**Data S13**. **Alignment of the genomic *Xa23* sequences from two cultivated rice accessions (Nipponbare and CBB23) and 3'RACE-derived parts of the TSA-induced *Xa23* transcripts.** The alignment was constructed using the MUSCLE algorithm.[11] Binding sites of the primers KS505, KS507, and KS529 that were used for nested 3'RACE PCRs are annotated and underlined. The binding elements for the dTALE and AvrXa23 activating *Xa23* transcription are annotated and highlighted in magenta.[13,14] dTALE and AvrXa23 bind to different elements. The published 5'UTR, CDS, and 3'UTR of the dTALE-induced *Xa23* mRNA transcripts are highlighted in blue, yellow, and grey, respectively.[13] The *Xa23* start (**ATG**) and stop (**TAA**) codons are colored in **red**. TATA boxes and TATA box-like sequences detected upstream of the *Xa23* start codon are colored in **green**. Additional potential CREs within the 5' upstream sequences identified using PlantCARE[15] (GCN4 motifs, **G boxes**, and **CARE motifs**) are highlighted and annotated in the alignment. Poly(A) tails at the 3' ends of the cloned TSA-induced *Xa23* transcripts are colored in **blue**.

```
                           ........10........20........30........40........50........60
Xa23^Nipponbare locus      .............................................AGCAGTCCCCCCTTC
KS505+KS507_Col27          ............................................................
KS505+KS507_Col30          ............................................................
KS505+KS507_Col71          ............................................................
KS505+KS507_Col72          ............................................................
KS505+KS507_Col91          ............................................................
KS507+KS529_Col52          ............................................................
KS507+KS529_Col54          ............................................................
KS507+KS529_Col55          ............................................................
KS507+KS529_Col56          ............................................................
KS507+KS529_Col57          ............................................................
KS507+KS529_Col58          ............................................................
KS507+KS529_Col59          ............................................................
Xa23^CBB23 locus           CAACGTCAGCGCCAATGTGTTTGGCGCTGAGGCGACGGCCTATTTTTGGTTGAAGTTTTT

                                                      dTALE^Xa23Ni-1
                           ........70........80........90.......100.......110.......120
Xa23^Nipponbare locus      CCTTCCGCTATTTCAAAGTATCCGTCTTCCTCCCTGTCCACATGAGCTAAACGCTTCCTC
KS505+KS507_Col27          ............................................................
KS505+KS507_Col30          ............................................................
KS505+KS507_Col71          ............................................................
KS505+KS507_Col72          ............................................................
KS505+KS507_Col91          ............................................................
KS507+KS529_Col52          ............................................................
KS507+KS529_Col54          ............................................................
KS507+KS529_Col55          ............................................................
KS507+KS529_Col56          ............................................................
KS507+KS529_Col57          ............................................................
KS507+KS529_Col58          ............................................................
KS507+KS529_Col59          ............................................................
Xa23^CBB23 locus           GGCAGGGGTTAGTTTCGAAATAAGTTTTCCAAAAGGGTCAATTTGTCAAAAAAACGGCTC

                               GCN4 motif          TATA box-like1    TATA box-like2
                           .......130.......140.......150.......160.......170.......180
Xa23^Nipponbare locus      TTCCCTGAGTCAAAGTCTTCCCTATATTACTACTTTGTATTATAGCTGGGTTCACTATGT
KS505+KS507_Col27          ............................................................
KS505+KS507_Col30          ............................................................
KS505+KS507_Col71          ............................................................
KS505+KS507_Col72          ............................................................
KS505+KS507_Col91          ............................................................
KS507+KS529_Col52          ............................................................
KS507+KS529_Col54          ............................................................
KS507+KS529_Col55          ............................................................
KS507+KS529_Col56          ............................................................
KS507+KS529_Col57          ............................................................
KS507+KS529_Col58          ............................................................
KS507+KS529_Col59          ............................................................
Xa23^CBB23 locus           GTCCCTGAGTCAAAGTCTTCCCTATTAAATTATGCGGCATCAC.................
                                GCN4 motif          TATA box-like3
```

```
                         .......190.......200.......210.......220.......230.......240
Xa23^Nipponbare locus    TGCATCATCTCACAGCTAACCCGAACATCACTAACATCGCAATATTCTCCTTCCGCGCCC
KS505+KS507_Col27        ............................................................
KS505+KS507_Col30        ............................................................
KS505+KS507_Col71        ............................................................
KS505+KS507_Col72        ............................................................
KS505+KS507_Col91        ............................................................
KS507+KS529_Col52        ............................................................
KS507+KS529_Col54        ............................................................
KS507+KS529_Col55        ............................................................
KS507+KS529_Col56        ............................................................
KS507+KS529_Col57        ............................................................
KS507+KS529_Col58        ............................................................
KS507+KS529_Col59        ............................................................
Xa23^CBB23 locus         .........................TAACATCAGCTACTATAAAAGTCCCTTCCGCGTCA
                                                              TATA box

                         .......250.......260.......270.......280.......290.......300
Xa23^Nipponbare locus    TTCACATCAGCTACTTCCACCACCTCCCATGTTTCAAAGGTAAATTCGTCGTGGTGGTGA
KS505+KS507_Col27        ............................................................
KS505+KS507_Col30        ............................................................
KS505+KS507_Col71        ............................................................
KS505+KS507_Col72        ............................................................
KS505+KS507_Col91        ............................................................
KS507+KS529_Col52        ............................................................
KS507+KS529_Col54        ............................................................
KS507+KS529_Col55        ............................................................
KS507+KS529_Col56        ............................................................
KS507+KS529_Col57        ............................................................
KS507+KS529_Col58        ............................................................
KS507+KS529_Col59        ............................................................
Xa23^CBB23 locus         CTAACATCAGCTACTATAAAAGTCCCTTCCGAAACA........................
                                       TATA box    AvrXa23

                                                             TATA box
                         .......310.......320.......330.......340.......350.......360
Xa23^Nipponbare locus    GATTATGCCTTCCTTCCGCCTCACTAACATCAGCTACTATAAAAGCCCTTCCTTGTTGCA
KS505+KS507_Col27        ............................................................
KS505+KS507_Col30        ............................................................
KS505+KS507_Col71        ............................................................
KS505+KS507_Col72        ............................................................
KS505+KS507_Col91        ............................................................
KS507+KS529_Col52        ............................................................
KS507+KS529_Col54        ............................................................
KS507+KS529_Col55        ............................................................
KS507+KS529_Col56        ............................................................
KS507+KS529_Col57        ............................................................
KS507+KS529_Col58        ............................................................
KS507+KS529_Col59        ............................................................
Xa23^CBB23 locus         .......TCTTCCTCCCGCATCACTAACATCAGCTTCTATAAAAGCCCTTCCTTGTTGCA
                                     AvrXa23                TATA box

                                                          G box            CARE motif
                         .......370.......380.......390.......400.......410.......420
Xa23^Nipponbare locus    TCATCTCAAGGAGCTGCAAGCACTTCCTCTCTGGCAGCACTTCCTCATCTCAAGGAGTTG
KS505+KS507_Col27        ............................................................
KS505+KS507_Col30        ............................................................
KS505+KS507_Col71        ............................................................
KS505+KS507_Col72        ............................................................
KS505+KS507_Col91        ............................................................
KS507+KS529_Col52        ............................................................
KS507+KS529_Col54        ............................................................
KS507+KS529_Col55        ............................................................
KS507+KS529_Col56        ............................................................
KS507+KS529_Col57        ............................................................
KS507+KS529_Col58        ............................................................
KS507+KS529_Col59        ............................................................
Xa23^CBB23 locus         TCATCTCAAGGAGCTGCAAGCACTTCCTCTCTGGCAGCACTTCCTCATCTCAAGGAGTTG
                                                           G box           CARE motif
```

```
                                      primer KS505            primer KS507
                                ----------------------------->   ----------------
                    .......430.......440.......450.......460.......470.......480
Xa23Nipponbare locus CAAATGTTGCATCATCTCAAGGAGCTGGCAGCCGTAGCCGGTATACACATGATCCTCATC
KS505+KS507_Col27   .............................................CACATGATCCTCATC
KS505+KS507_Col30   .............................................CACATGATCCTCATC
KS505+KS507_Col71   .......................................NNNNNNNNNNTGATCCTCATC
KS505+KS507_Col72   .....................................NNNNNNNNNNNNTGATCCTCATC
KS505+KS507_Col91   .....................................NNNNNNNNNNNNTGATCCTCATC
KS507+KS529_Col52   ............................................................
KS507+KS529_Col54   ............................................................
KS507+KS529_Col55   ............................................................
KS507+KS529_Col56   ............................................................
KS507+KS529_Col57   ............................................................
KS507+KS529_Col58   ............................................................
KS507+KS529_Col59   ............................................................
Xa23CBB23 locus      CAAATGTTGCATCATCTCAAGGAGCTGGCAGCCGTAGCCGGTATACACATGATCCTCATC

                    primer KS507                                   primer KS529
                    ------------->                             --------------------
                    .......490.......500.......510.......520.......530.......540
Xa23Nipponbare locus TACCTCTGCCGCTTTCTCCTCCGCCGCAGCCGCAACGTATTATTCACCGTTTCCAACAGC
KS505+KS507_Col27   TACCTCTGCCGCTTTCTCCTCCGCCGCAGCCGCAACGTATTATTCACCGTTTCCAACAGC
KS505+KS507_Col30   TACCTCTGCCGCTTTCTCCTCCGCCGCAGCCGCAACGTATTATTCACCGTTTCCAACAGC
KS505+KS507_Col71   TACCTCTGCCGCTTTCTCCTCCGCCGCAGCCGCAACGTATTATTCACCGTTTCCAACAGC
KS505+KS507_Col72   TACCTCTGCCGCTTTCTCCTCCGCCGCAGCCGCAACGTATTATTCACCGTTTCCAACAGC
KS505+KS507_Col91   TACCTCTGCCGCTTTCTCCTCCGCCGCAGCCGCAACGTATTATTCACCGTTTCCAACAGC
KS507+KS529_Col52   ...........................................TCACCGTTTCCAACAGC
KS507+KS529_Col54   ...........................................TCACCGTTTCCAACAGC
KS507+KS529_Col55   ...........................................TCACCGTTTCCAACAGC
KS507+KS529_Col56   ...........................................TCACCGTTTCCAACAGC
KS507+KS529_Col57   ...........................................TCACCGTTTCCAACAGC
KS507+KS529_Col58   ...........................................TCACCGTTTCCAACAGC
KS507+KS529_Col59   ...........................................TCACCGTTTCCAACAGC
Xa23CBB23 locus      TACCTCTGCCGCTTTCTCCTCCGCCGCAGCCGCAACGTATTATTCACCGTTTCCAACAGC

                 primer KS529
                    ---->
                    .......550.......560.......570.......580.......590.......600
Xa23Nipponbare locus CTCCGTTTTCGCCTCAAGGTATTAACTGTATTGTTGTACATATGTCTCTCGGTCATGCTG
KS505+KS507_Col27   CTCCGTTTTCGCCTCAAGGTATTAACTGTATTGTTGTACATATGTCTCTCGGTCATGCTG
KS505+KS507_Col30   CTCCGTTTTCGCCTCAAGGTATTAACTGTATTGTTGTACATATGTCTCTCGGTCATGCTG
KS505+KS507_Col71   CTCCGTTTTCGCCTCAAGGTATTAACTGTATTGTTGTACATATGTCTCTCGGTCATGCTG
KS505+KS507_Col72   CTCCGTTTTCGCCTCAAGGTATTAACTGTATTGTTGTACATATGTCTCTCGGTCATGCTG
KS505+KS507_Col91   CTCCGTTTTCGCCTCAAGGTATTAACTGTATTGTTGTACATATGTCTCTCGGTCATGCTG
KS507+KS529_Col52   CTCCGTTTTCGCCTCAAGGTATTAACTGTATTGTTGTACATATGTCTCTCGGTCATGCTG
KS507+KS529_Col54   CTCCGTTTTCGCCTCAAGGTATTAACTGTATTGTTGTACATATGTCTCTCGGTCATGCTG
KS507+KS529_Col55   CTCCGTTTTCGCCTCAAGGTATTAACTGTATTGTTGTACATATGTCTTTCGGTCATGCTG
KS507+KS529_Col56   CTCCGTTTTCGCCTCAAGGTATTAACTGTATTGTTGTACATATGTCTCTCGGTCATGCTG
KS507+KS529_Col57   CTCCGTTTTCGCCTCAAGGTATTAACTGTATTGTTGTACATATGTCTCTCGGTCATGCTG
KS507+KS529_Col58   CTCCGTTTTCGCCTCAAGGTATTAACTGTATTGTTGTACATATGTCTCTCGGTCATGCTG
KS507+KS529_Col59   CTCCGTTTTCGCCTCAAGGTATTAACTGTATTGTTGTACATATGTCTCTCGGTCATGCTG
Xa23CBB23 locus      CTCCGTTTTCGCCTCAAGGTATTAACTGTATTGTTGTACATATGTCTCTCGGTCATGCTG

                    .......610.......620.......630.......640.......650.......660
Xa23Nipponbare locus TTCTACCTGTTTGGCTCCATCATGCCGCTGCCGCCGTGGGGCCTCGTGGTCGGTTGGGTC
KS505+KS507_Col27   TTCTACCTGTTTGGCTCCATCATGCCGCTGCCGCCGTGGGGCCTCGTGGTCGGTTGGGTC
KS505+KS507_Col30   TTCTACCTGTTTGGCTCCATCATGCCGCTGCCGCCGTGGGGCCTCGTGGTCGGTTGGGTC
KS505+KS507_Col71   TTCTACCTGTTTGGCTCCATCATGCCGCTGCCGCCGTGGGGCCTCGTGGTCGGTTGGGTC
KS505+KS507_Col72   TTCTACCTGTTTGGCTCCATCATGCCGCTGCCGCCGTGGGGCCTCGTGGTCGGTTGGGTC
KS505+KS507_Col91   TTCTACCTGTTTGGCTCCATCATGCCGCTGCCGCCGTGGGGCCTCGTGGTCGGTTGGGTC
KS507+KS529_Col52   TTCTACCTGTTTGGCTCCATCATGCCGCTGCCGCCGTGGGGCCTCGTGGTCGGTTGGGTC
KS507+KS529_Col54   TTCTACCTGTTTGGCTCCATCATGCCGCTGCCGCCGTGGGGCCTCGTGGTCGGTTGGGTC
KS507+KS529_Col55   TTCTACCTGTTTGGCTCCATCATGCCGCTGCCGCCGTGGGGCCTCGTGGTCGGTTGGGTC
KS507+KS529_Col56   TTCTACCTGTTTGGCTCCATCATGCCGCTGCCGCCGTGGGGCCTCGTGGTCGGTTGGGTC
KS507+KS529_Col57   TTCTACCTGTTTGGCTCCATCATGCCGCTGCCGCCGTGGGGCCTCGTGGTCGGTTGGGTC
KS507+KS529_Col58   TTNTACCTGTTTGGCTCCATCATGCCGCTGCCGCCGTGGGGCCTCGTGGTCGGTTGGGTC
KS507+KS529_Col59   TTCTACCTGTTTGGCTCCATCATGCCGCTGCCGCCGTGGGGCCTCGTGGTCGGTTGGGTC
Xa23CBB23 locus      TTCTACCTGTTTGGCTCCATCATGCCGCTGCCGCCGTGGGGCCTCGTGGTCGGTTGGGTC
```

.......670.......680.......690.......700.......710.......720
*Xa23*$^{Nipponbare}$ locus ATGGCCCTCATCGCCGTCGAGCTCGCCTACGCCTTCATCTTTCCATATAGCTTTCGCTAC
KS505+KS507_Col27 ATGGCCCTCATCGCCGTCGAGCTCGCCTACGCCTTCATCTTTCCATATAGCTTTCGCTAC
KS505+KS507_Col30 ATGGCCCTCATCGCCGTCGAGCTCGCCTACGCCTTCATCTTTCCATATAGCTTTCGCTAC
KS505+KS507_Col71 ATGGCCCTCATCGCCGTCGAGCTCGCCTACGCCTTCATCTTTCCATATAGCTTTCGCTAC
KS505+KS507_Col72 ATGGCCCTCATCGCCGTCGAGCTCGCCTACGCCTTCATCTTTCCATATAGCTTTCGCTAC
KS505+KS507_Col91 ATGGCCCTCATCGCCGTCGAGCTCGCCTACGCCTTCATCTTTCCATATAGCTTTCGCTAC
KS507+KS529_Col52 ATGGCCCTCATCGCCGTCGAGCTCGCCTACGCCTTCATCTTTCCATATAGCTTTCGCTAC
KS507+KS529_Col54 ATGGCCCTCATCGCCGTCGAGCTCGCCTACGCCTTCATCTTTCCATATAGCTTTCGCTAC
KS507+KS529_Col55 ATGGCCCTCATCGCCGTTGAGCTCGCCTACGCCTTCATCTTTCCATATAGCTTTCGCTAC
KS507+KS529_Col56 ATGGCCCTCATCGCCGTCGAGCTCGCCTACGCCTTCATCTTTCCATATAGCTTTCGCTAC
KS507+KS529_Col57 ATGGCCCTCATCGCCGTCGAGCTCGCCTACGCCTTCATCTTTCCATATAGCTTTCGCTAC
KS507+KS529_Col58 ATGGCCCTCATCGCCGTCGAGCTCGCNTACGCCTTCATCTTTCCATATAGCTTTCGCTAC
KS507+KS529_Col59 ATGGCCCTCATCGCCGTCGAGCTCGCCTACGCCTTCATCTTTCCATATAGCTTTCGCTAC
*Xa23*$^{CBB23}$ locus ATGGCCCTCATCGCCGTCGAGCTCGCCTACGCCTTCATCTTTCCATATAGCTTTCGCTAC

.......730.......740.......750.......760.......770.......780
*Xa23*$^{Nipponbare}$ locus ATCGCTGACAACGACGACGACAAGATGGTTATTCTCCCTGTT**TAA**GCCTTCAGGGCCTAT
KS505+KS507_Col27 ATCGCTGACAACGACGACGACAAGATGGTTATTCTCCCTGTTTAAGCCTTCAGGGCCTAT
KS505+KS507_Col30 ATCGCTGACAACGACGACGACAAGATGGTTATTCTCCCTGTTTAAGCCTTCAGGGCCTAT
KS505+KS507_Col71 ATCGCTGACAACGACGACGACAAGATGGTTATTCTCCCTGTTTAAGCCTTCAGGGCCTAT
KS505+KS507_Col72 ATCGCTGACAACGACGACGACAAGATGGTTATTCTCCCTGTTTAAGCCTTCAGGGCCTAT
KS505+KS507_Col91 ATCGCTGACAACGACGACGACAAGATGGTTATTCTCCCTGTTTAAGCCTTCAGGGCCTAT
KS507+KS529_Col52 ATCGCTGACAACGACGACGACAAGATGGTTATTCTCCCTGTTTAAGCCTTCAGGGCCTAT
KS507+KS529_Col54 ATCGCTGACAACGACGACGACAAGATGGTTATTCTCCCTGTTTAAGCCTTCAGGGCCTAT
KS507+KS529_Col55 ATCGCTGACAACGAGGACGACAAGATGGTTATTCTCCCTGTTTAAGCCTTCAGGGCCTAT
KS507+KS529_Col56 ATCGCTGACAACGACGACGACAAGATGGTTATTCTCCCTGTTTAAGCCTTCAGGGCCTAT
KS507+KS529_Col57 ATCGCTGACAACGACGACGACAAGATGGTTATTCTCCCTGTTTAAGCCTTCAGGGCCTAT
KS507+KS529_Col58 ATCGCTGACAACGAGGACGACAAGATGGTTATTCTCCCTGTTTAAGCCTTCAGGGCCTAT
KS507+KS529_Col59 ATCGCTGACAACGACGACGACAAGATGGTTATTCTCCCTGTTTAAGCCTTCAGGGCCTAT
*Xa23*$^{CBB23}$ locus ATCGCTGACAACGACGACGACAAGATGGTTATTCTCCCTGTT**TAA**GCCTTCAGGGCCTAT

.......790.......800.......810.......820.......830.......840
*Xa23*$^{Nipponbare}$ locus ATATATAGTATATATATAAAGCCTTCCATACTGTCTCTTCAATAAAGGCTAGCTTGTGTT
KS505+KS507_Col27 ATATATAGTATATATATAAAGCCTTCCATACTGTCTCTTCAATAAAGGCTAGCTTGTGTT
KS505+KS507_Col30 ATATATAGTATATATATAAAGCCTTCCATACTGTCTCTTCAATAAAGGCTAGCTTGTGTT
KS505+KS507_Col71 ATATATAGTATATATATAAAGCCTTCCATACTGTCTCTTCAATAAAGGCTAGCTTGTGTT
KS505+KS507_Col72 ATATATAGTATATATATAAAGCCTTCCATACTGTCTCTTCAATAAAGGCTAGCTTGTGTT
KS505+KS507_Col91 ATATATAGTATATATATAAAGCCTTCCATACTGTCTCTTCAATAAAGGCTAGCTTGTGTT
KS507+KS529_Col52 ATATATAGTATATATATAAAGCCTTCCATACTGTCTCTTCAATAAAGGCTAGCTTGTGTT
KS507+KS529_Col54 ATATATAGTATATATATAAAGCCTTCCATACTGTCTCTTCAATAAAGGCTAGCTTGTGTT
KS507+KS529_Col55 ATATATAGTATATATATAAACCCTTCCATACTGTCTCTTCAATAAAGGCTAGCTTGTGTT
KS507+KS529_Col56 ATATATAGTATATATATAAAGCCTTCCATACTGTCTCTTCAATAAAGGCTAGCTTGTGTT
KS507+KS529_Col57 ATATATAGTATATATATAAAGCCTTCCATACTGTCTCTTCAATAAAGGCTAGCTTGTGTT
KS507+KS529_Col58 ATATATAGTATATATATAAAGCCTTCCATACTGTCTCTTCAATAAAGGCTAGCTTGTGTT
KS507+KS529_Col59 ATATATAGTATATATATAAAGCCTTCCATACTGTCTCTTCAATAAAGGCTAGCTTGTGTT
*Xa23*$^{CBB23}$ locus ATATATAGTATATATATAAAGCCTTCCATACTGTCTCTTCAATAAAGGCTAGCTTGTGTT

.......850.......860.......870.......880.......890.......900
*Xa23*$^{Nipponbare}$ locus GTGAGTTGTATCTGTGTACGTATTTTGTTTGGTTGGTTATATATTGTCACGTAGGTATGC
KS505+KS507_Col27 GTGAGTTGTATCTGTGTACGTATTTTGTTTGGTTGGTTATATATTGTCACGTAGGTATGC
KS505+KS507_Col30 GTGAGTTGTATCTGTGTACGTATTTTGTTTGGTTGGTTATATATTGTCACGTAGGTATGC
KS505+KS507_Col71 GTGAGTTGTATCTGTGTACGTATTTTGTTTGGTTGGTTATATATTGTCACGTAGGTATGC
KS505+KS507_Col72 GTGAGTTGTATCTGTGTACGTATTTTGTTTGGTTGGTTATATATTGTCACGTAGGTATGC
KS505+KS507_Col91 GTGAGTTGTATCTGTGTACGTATTTTGTTTGGTTGGTTATATATTGTCACGTAGGTATGC
KS507+KS529_Col52 GTGAGTTGTATCTGTGTACGTATTTTGTTTGGTTGGTTATATATTGTCACGTAGGTATGC
KS507+KS529_Col54 GTGAGTTGTATCTGTGTACGTATTTTGTTTGGTTGGTTATATATTGTCACGTAGGTATGC
KS507+KS529_Col55 GTGAGTTGTATCTGTGTACGTATTTTGTTTGGTTGGTTATATATTGTCACGTAGGTATCC
KS507+KS529_Col56 GTGAGTTGTATCTGTGTACGTATTTTGTTTGGTTGGTTATATATTGTCACGTAGGTATGC
KS507+KS529_Col57 GTGAGTTGTATCTGTGTACGTATTTTGTTTGGTTGGTTATATATTGTCACGTAGGTATGC
KS507+KS529_Col58 GTGAGTTGTATCTGTGTACGTATTTTGTTTGGTTGGTTATATATTGTCACGTAGGTATGC
KS507+KS529_Col59 GTGAGTTGTATCTGTGTACGTATTTTGTTTGGTTGGTTATATATTGTCACGTAGGTATGC
*Xa23*$^{CBB23}$ locus GTGAGTTGTATCTGTGTACGTATTTTGTTTGGTTGGTTATATATTGTCACGTAGGTATGC

.......910.......920.......930.......940.......950.......960
*Xa23*$^{Nipponbare}$ locus CATATATATATGTATTGCTGTATTTATATTTGTTACTATCTTTTGTTTTTCAGATAATAA
KS505+KS507_Col27 CATATATATATGTATTGCTGTATTTATATTTGTTACTATCTTTTGTTTTTCAGATAATAA
KS505+KS507_Col30 CATATATATATGTATTGCTGTATTTATATTTGTTACTATCTTTTGTTTTTCAGATAATAA
KS505+KS507_Col71 CATATATATATGTATTGCTGTATTTATATTTGTTACTATCTTTTGTTTTTCAGATAATAA
KS505+KS507_Col72 CATATATATATGTATTGCTGTATTTATATTTGTTACTATCTTTTGTTTTTCAGATAATAA
KS505+KS507_Col91 CATATATATATGTATTGCTGTATTTATATTTGTTACTATCTTTTGTTTTTCAGATAATAA
KS507+KS529_Col52 CATATATATATGTATTGCTGTATTTATATTTGTTACTATCTTTTGTTTTTCAGATAATAA
KS507+KS529_Col54 CATATATATATGTATTGCTGTATTTATATTTGTTACTATCTTTTGTTTTTCAGATAATAA
KS507+KS529_Col55 CATATATATATGTATTGCTGTATTTATATTTGTTACTATCTTTTGTTTTTCAGATAATAA
KS507+KS529_Col56 CATATATATATGTATTGCTGTATTTATATTTGTTACTATCTTTTGTTTTTCAGATAATAA
KS507+KS529_Col57 CATATATATATGTATTGCTGTATTTATATTTGTTACTATCTTTTGTTTTTCAGATAATAA
KS507+KS529_Col58 CATATATATATGTATTGCTGTATTTATATTTGTTACTATCTTTTGTTTTTCAGATAATAA
KS507+KS529_Col59 CATATATATATGTATTGCTGTATTTATATTTGTTACTATCTTTTGTTTTTCAGATAATAA
*Xa23*$^{CBB23}$ locus CATATATATATGTATTGCTGTATTTATATTTGTTACTATCTTTTGTTTTTCAGATAATAA

.......970.......980.......990......1000......1010......1020
*Xa23*$^{Nipponbare}$ locus AATTCAGCCAGCTTTGCTTGCTTCGTCGTACGTGTATGCTCATCATATCCTCATCCATCA
KS505+KS507_Col27 AATTCAGCCAGCTTTGCTTGCTTC**AAAAAAAAAAAAAAAAAAAAAAAAAAA**.........
KS505+KS507_Col30 AATTCAGCCAGCTTTGCTTGCTTC**AAAAAAAAAAAAAAAAAAAAAA**..............
KS505+KS507_Col71 AATTCAGCCAGCTTTGCTTGCTTC**AAAAAAAAAAAAAAAAAAAAAAA**.............
KS505+KS507_Col72 AATTCAGCCAGCTTTGCTTGCTTC**AAAAAAAAAAAAAAAAAAAAAAAAAAAAA**.......
KS505+KS507_Col91 AATTCAGCCAGCTTTGCTTGCTTC**AAAAAAAAAAAAAAAAAAAAAAAAAAA**.........
KS507+KS529_Col52 AATTCAGCCAGCTTTGCTTGCGT**AAAAAAAAAAAAAAAAAAAAAA**...............
KS507+KS529_Col54 AATTCAGCCAGCTTTGCTTGCGT**AAAAAAAAAAAAAAAAAAAAAAAAAAAAA**........
KS507+KS529_Col55 AATTCANCCAGCTTTGCTTGCGC**AAAAAAAAAAAAAAAAAAAAAAAAAAAAAA**.......
KS507+KS529_Col56 AATTCAGCCAGCTTTGCTTGCGT**AAAAAAAAAAAAAAAAAAAAA**................
KS507+KS529_Col57 AATTCAGCCAGCTTTGCTTGCGT**AAAAAAAAAAAAAAAAAAAAAAAA**.............
KS507+KS529_Col58 AATTCAGCCAGCTTTGCTTGCGC**AAAAAAAAAAAAAAAAAAAAAAAAAA**...........
KS507+KS529_Col59 AATTCAGCCAGCTTTGCTTGCGT**AAAAAAAAAAAAAAAAAAAAA**................
*Xa23*$^{CBB23}$ locus AATTCAGCCAGCTTTGCTTGCTTCGTCGTACGTGTATGCTCATCATATCCTCATCCATCA

......1030......1040.....
*Xa23*$^{Nipponbare}$ locus GCTGCTCATAGCTAGCTGGGCCGTG
KS505+KS507_Col27 .........................
KS505+KS507_Col30 .........................
KS505+KS507_Col71 .........................
KS505+KS507_Col72 .........................
KS505+KS507_Col91 .........................
KS507+KS529_Col52 .........................
KS507+KS529_Col54 .........................
KS507+KS529_Col55 .........................
KS507+KS529_Col56 .........................
KS507+KS529_Col57 .........................
KS507+KS529_Col58 .........................
KS507+KS529_Col59 .........................
*Xa23*$^{CBB23}$ locus GCTGCTCATAGCTAGCTGGGCCGTG

### References


1. Rahman, M.H., Toda, E., Kobayashi, M., Kudo, T., Koshimizu, S., Takahara, M., Iwami, M., Watanabe, Y., Sekimoto, H., Yano, K., and Okamoto, T. (2019). Expression of genes from paternal alleles in rice zygotes and involvement of *OsASGR-BBML1* in initiation of zygotic development. Plant and Cell Physiology 60, 725-737. 10.1093/pcp/pcz030.
2. Sakai, H., Lee, S.S., Tanaka, T., Numa, H., Kim, J., Kawahara, Y., Wakimoto, H., Yang, C.-c., Iwamoto, M., Abe, T., *et al*. (2013). Rice annotation project database (RAP-DB): an integrative and interactive database for rice genomics. Plant and Cell Physiology 54, e6-e6. 10.1093/pcp/pcs183.
3. Toda, E., Koshimizu, S., Kinoshita, A., Higashiyama, T., Izawa, T., Yano, K., and Okamoto, T. (2025). Transcriptional dynamics during karyogamy in rice zygotes. Development. 10.1242/dev.204497.
4. Langmead, B., and Salzberg, S.L. (2012). Fast gapped-read alignment with Bowtie 2. Nat. Methods 9, 357-359. 10.1038/nmeth.1923.
5. Li, H., Handsaker, B., Wysoker, A., Fennell, T., Ruan, J., Homer, N., Marth, G., Abecasis, G., Durbin, R., and Subgroup, G.P.D.P. (2009). The sequence alignment / map format and SAMtools. Bioinformatics 25, 2078-2079. 10.1093/bioinformatics/btp352.
6. Robinson, J.T., Thorvaldsdóttir, H., Winckler, W., Guttman, M., Lander, E.S., Getz, G., and Mesirov, J.P. (2011). Integrative genomics viewer. Nat. Biotechnol. 29, 24-26. 10.1038/nbt.1754.
7. Edgar, R.C. (2022). Muscle5: high-accuracy alignment ensembles enable unbiased assessments of sequence homology and phylogeny. Nat. Commun. 13, 6968. 10.1038/s41467-022-34630-w.
8. Sievers, F., Wilm, A., Dineen, D., Gibson, T.J., Karplus, K., Li, W., Lopez, R., McWilliam, H., Remmert, M., Söding, J., *et al*. (2011). Fast, scalable generation of high-quality protein multiple sequence alignments using Clustal Omega. Mol. Syst. Biol. 7, 539. 10.1038/msb.2011.75.
9. Mansueto, L., Fuentes, R.R., Borja, F.N., Detras, J., Abriol-Santos, J.M., Chebotarov, D., Sanciangco, M., Palis, K., Copetti, D., Poliakov, A., *et al*. (2017). Rice SNP-seek database update: new SNPs, indels, and queries. Nucleic Acids Res. 45, D1075-D1081. 10.1093/nar/gkw1135.
10. Hallgren, J., Tsirigos, K.D., Pedersen, M.D., Almagro Armenteros, J.J., Marcatili, P., Nielsen, H., Krogh, A., and Winther, O. (2022). DeepTMHMM predicts alpha and beta transmembrane proteins using deep neural networks. bioRxiv, 2022.2004.2008.487609. 10.1101/2022.04.08.487609.
11. Madeira, F., Madhusoodanan, N., Lee, J., Eusebi, A., Niewielska, A., Tivey, A.R.N., Lopez, R., and Butcher, S. (2024). The EMBL-EBI job dispatcher sequence analysis tools framework in 2024. Nucleic Acids Res. 52, W521-W525. 10.1093/nar/gkae241.
12. Tian, D., Wang, J., Zeng, X., Gu, K., Qiu, C., Yang, X., Zhou, Z., Goh, M., Luo, Y., Murata-Hori, M., *et al*. (2014). The rice TAL effector–dependent resistance protein XA10 triggers cell death and calcium depletion in the endoplasmic reticulum. The Plant Cell 26, 497-515. 10.1105/tpc.113.119255.
13. Wang, J., Tian, D., Gu, K., Yang, X., Wang, L., Zeng, X., and Yin, Z. (2017). Induction of *Xa10*-like genes in rice cultivar Nipponbare confers disease resistance to rice bacterial blight. Mol. Plant-Microbe Interact. 30, 466-477. 10.1094/MPMI-11-16-0229-R.

14. Wang, C., Zhang, X., Fan, Y., Gao, Y., Zhu, Q., Zheng, C., Qin, T., Li, Y., Che, J., Zhang, M., *et al*. (2015). XA23 is an executor R protein and confers broad-spectrum disease resistance in rice. Molecular Plant 8, 290-302. 10.1016/j.molp.2014.10.010.

15. Lescot, M., Déhais, P., Thijs, G., Marchal, K., Moreau, Y., Van de Peer, Y., Rouzé, P., and Rombauts, S. (2002). PlantCARE, a database of plant *cis*-acting regulatory elements and a portal to tools for *in silico* analysis of promoter sequences. Nucleic Acids Res. 30, 325-327. 10.1093/nar/30.1.325.
